# Interferons are the key cytokines acting on pancreatic islets in type 1 diabetes

**DOI:** 10.1101/2023.06.29.547000

**Authors:** Alexandra Coomans de Brachène, Maria Ines Alvelos, Florian Szymczak, Priscila Laiz Zimath, Angela Castela, Bianca Marmontel de Souza, Arturo Roca Rivada, Sandra Marín-Cañas, Xiaoyan Yi, Anne Op de Beeck, Noel G. Morgan, Sebastian Sonntag, Sayro Jawurek, Alexandra C. Title, Burcak Yesildag, François Pattou, Julie Kerr-Conte, Eduard Montanya, Montserrat Nacher, Lorella Marselli, Piero Marchetti, Sarah J. Richardson, Decio L. Eizirik

**Author notes:** These authors contributed equally to the study (Alexandra Coomans de Brachène, Maria Ines Alvelos, Florian Szymczak). Correspondence should be addressed to: Alexandra Coomans de Brachène or Decio L. Eizirik ULB Center for Diabetes Research Campus Erasme, Université Libre de Bruxelles Route de Lennik, 808-CP618, 1070 Brussels, Belgium (ACdB) (DLE).

## Abstract

The pro-inflammatory cytokines IFNα, IFNγ, IL-1β and TNFα may contribute to innate and adaptive immune responses during islet inflammation (insulitis) in type 1 diabetes (T1D). We used deep RNA-sequencing analysis to characterize the response of human pancreatic beta cells to each cytokine individually and compared the signatures obtained with those present in islets of individuals affected by T1D. IFNα and IFNγ had a much greater impact on the beta cell transcriptome when compared to IL-1β and TNFα. The IFN-induced gene signatures have a strong correlation with those observed in beta cells from T1D patients, and the level of expression of specific IFN-stimulated genes is positively correlated with proteins present in islets of these individuals, regulating beta cell responses to “danger signals” such as viral infections. These data suggest that IFNα and IFNγ are the central cytokines at the islet level in T1D, contributing to the triggering and amplification of autoimmunity.

## Introduction

Pancreatic beta cells are unique in their ability to maintain glucose homeostasis by producing, storing, and releasing the hormone insulin. Type 1 diabetes (T1D) is caused by severe insulin deficiency due to an immune-mediated assault on the pancreatic beta cells, leading to islet inflammation (insulitis) and loss of beta cells in genetically susceptible individuals submitted to environmental triggers^1, 2^. There is still no cure for T1D and the standard treatment relies on lifelong exogenous insulin administration^3^.

Cytokines are signaling molecules that play a central role in T1D by controlling innate and adaptative immune responses and thus impacting on the initiation and amplification of beta cell autoimmunity^1, 2, 4^. Specifically, the pro-inflammatory cytokines interferon α (IFNα), interferon γ (IFNγ), interleukin 1β (IL-1β), and tumor necrosis factor α (TNFα) are produced mainly by islet-infiltrating immune cells and affect beta cell function and survival, contributing to T1D development^4–7^.

IFNα, a type I interferon (IFN) cytokine secreted by microbial infected cells, initiates the innate immune response. IFNα is expressed in islets from T1D individuals and its signaling is a key component of the early stages of T1D, as there is evidence for a type I IFN signature before the onset of the disease^1, 8^. This cytokine induces endoplasmic reticulum (ER) stress, insulitis and overexpression of the human leukocyte antigen (HLA) class I in human beta cells – three hallmarks of T1D^4, 9, 10^. IFNα signals via JAK1/TYK2 and STAT1/STAT2/IRF9 (or IRF7), triggering the expression of interferon-stimulated genes (ISG), including HLAs and chemokines, among others^11^. Interestingly, *TYK2* is a susceptibility gene for T1D and single nucleotide polymorphisms that decrease *TYK2* expression have a protective effect against the disease^12^. IFNγ is secreted by adaptive immune cells, such as CD4^+^ and CD8^+^ T cells, and signals via JAK1/JAK2/STAT1 to regulate the expression of many genes, including the transcription factors of the interferon regulatory factor (IRF) family^13, 14^. An IFNγ signature was found in young islet autoantibody-positive individuals^15^. Blocking IFNγ in NOD mice, either by using specific anti-IFNγ antibodies or soluble IFNγ receptors^16^, decreases the incidence of diabetes and prevents disease transfer by splenocytes from diabetic NOD donor mice^17^. IFNγ alone does not induce beta cell apoptosis, but its combination with either IL-1β or TNFα causes beta cell death^18^.

IL-1β is a powerful pro-inflammatory cytokine secreted by monocytes, macrophages and dendritic cells during inflammation and host response to infections, and serum levels of IL-1β are higher in T1D patients when compared to non-diabetic individuals^19^. IL-1β, together with IFNγ, induces ER stress and the unfolded protein response in both rodent and human islets^20^. IL-1β also acts synergistically with IFNα to induce human beta cell apoptosis^9^. TNFα is also produced by immune cells, including macrophages and dendritic cells, and elevated levels of TNFα are found in the plasma and the islets of patients with T1D. A significant association between the plasma levels of TNFα and glycemic control in T1D individuals has been observed^19^.

Cytokines participate to an effective immune response to different pathogens, but their dysregulation contributes to the development and progression of T1D and other autoimmune diseases, making them attractive immunotherapeutic targets^12, 21^. Until now, most studies evaluating the impact of pro-inflammatory cytokines on human pancreatic beta cells in the context of T1D used combinations of 2-4 pro-inflammatory cytokines to simulate the inflammatory environment in T1D. To identify key targets for therapy, however, it is crucial to clarify the role and relevance of each individual cytokine. Having this in mind, we performed deep RNA-sequencing (RNA-seq) analysis on human insulin-producing EndoC-βH1 cells treated with each individual cytokine (IFNα, IFNγ, IL-1β and TNFα) alone and investigated the main molecular pathways activated by these four cytokines. Our data indicate a predominant role for the IFNs, both IFNα and IFNγ, in the pathogenic response observed at the beta cell level in T1D as compared to a milder impact of IL-1β and TNFα. Therefore, we suggest that future studies and clinical trials dealing with the role of beta cells in the pathogenesis of T1D should primarily focus on IFNs in preference to the other cytokines studied.

## Results

### IFNα, IFNγ, IL-1β and TNFα have distinct effects on human pancreatic beta cells

Exposure to high concentrations of each individual cytokine for 24h did not induce apoptosis in either EndoC-βH1 cells or dispersed human islets (Fig. 1A, C). This aligns with our previous findings indicating that even longer exposure of primary human islets to individual cytokines does not induce apoptosis^1, 7, 22^. However, a 48h exposure of EndoC-βH1 cells to IFNα or IFNγ induced a mild increase (16-35%) in cell death (Fig. 1B).

**Figure 1:**
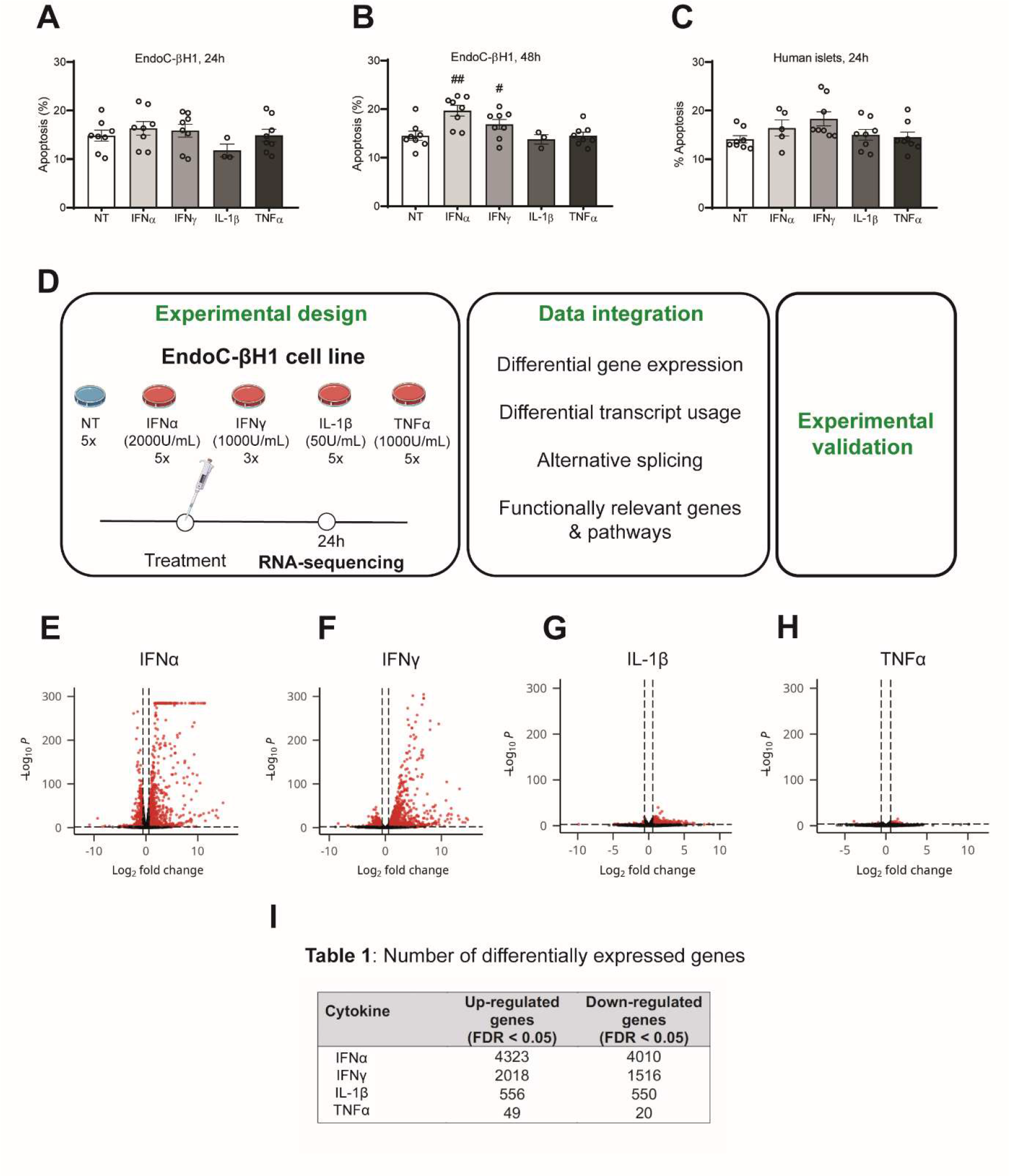
IFNα, IFNγ, IL-1β, and TNFα have distinct effects on human pancreatic beta cells. EndoC-βH1 cells and dispersed human islets were treated with IFNα (2000 U/mL), IFNγ (1000 U/mL), IL-1β (50 U/mL) or TNFα (1000 U/mL). (**A**) Apoptosis was evaluated by Hoechst 33342 + propidium iodide after 24h (**A**) and 48h (**B**) in EndoC-βH1 cells (n = 3-8), and after 24h in dispersed human islets (n = 6-9) (**C**). Results are expressed as mean ± SEM. #*p*<0.05, ##*p*<0.01 vs NT, (ANOVA). (**D**) Experimental design for the RNA-sequencing experiments. EndoC-βH1 cells were treated with IFNα, IFNγ, IL-1β or TNFα and RNA-sequencing performed after 24h of treatment (n = 5 to all conditions, except for IFNγ: n = 3). The data analysis and integration were performed as described in the Methods section. (**E-H**) Volcano plots for the differentially expressed genes in EndoC-βH1 cells after the respective treatments. The horizontal dashed line delimits the genes with an adjusted p-value < 0.05. The vertical dashed lines delimit the genes with |Log_2_ Fold Change| > 0.58. (**I**) Summary table with the number of genes up- and down-regulated after treatment with the respective pro-inflammatory cytokines.

We then treated EndoC-βH1 cells for 24h with IFNγ, IL-1β or TNFα individually to perform deep RNA-seq analysis (> 200 million reads) (Fig 1D). Our group has previously published an IFNα RNA-seq dataset^6^ which was re-analyzed here using the pipeline described in Methods. We have previously shown that most IFNα-stimulated genes are modified after 24h of stimulation^23^, thus performed the present experiments at 24h to enable data comparison and also to avoid possible changes associated with the apoptosis seen at later time points.

RNA-seq analysis showed that IFNα, IFNγ, IL-1β and TNFα have a distinct impact on gene expression in EndoC-βH1 cells (Fig. 1E-I), with both IFNα and IFNγ (Fig. 1E-F, I) modifying a larger number of genes than IL-1β or TNFα (Figs. 1G-I). IFNα and IFNγ induced the upregulation of 4323 and 2018 genes, respectively, and the downregulation of 4010 and 1516 genes (Fig. 1I). By contrast, treatment with IL-1β or TNFα induced the upregulation of only 556 and 49 genes, and the downregulation of 550 and 20 genes, respectively (Fig. 1I). All genes differentially expressed after treatment with the respective pro-inflammatory cytokines are listed in Supplementary Table 1.

The analysis of the expression of the different cytokine receptors under basal condition, based on previous and present RNA-seq of EndoC-βH1 cells, dispersed human islets and FACS-purified beta cells from donors with and without T1D, indicates that all receptors are expressed in human beta cells but with a clear predominance of the receptors for IFNs (*IFNAR1*, *IFNAR2*, *IFNGR1* and *IFNGR2*) as compared to IL-1β (*IL1R1* and *IL1R2*) and TNFα (*TNFRSF1A* and *TNFRSF1B*) receptors (Supplementary fig. 1). This is in marked contrast from observations in rat FACS-purified beta cells, where expression of the IL-1 receptor is nearly 15-fold higher than IFN receptors^24^. These data indicate that all of these four cytokines can transduce signal to human beta cells and induce changes in gene expression.

### IFNα and IFN**γ** induce a more marked immune-related response in human beta cells than IL-1β or TNFα

Using gene set enrichment analysis (GSEA) based on the Kyoto Encyclopaedia of Genes and Genomes (KEGG) and REACTOME databases (Fig. 2A-D), we re-analyzed the IFNα dataset and, in line with our previous findings^6^, observed an enrichment in IFN signaling–related pathways, antigen processing and presentation, and antiviral response among the up-regulated pathways, while there was depletion in oxidative phosphorylation and mitochondrial function among the downregulated pathways (Fig. 2A). GSEA analysis of the IFNγ up-regulated genes showed enrichment in IFN-signaling related pathways, antigen processing and presentation, assembly and peptide loading of HLA class I molecules. Analysis of the downregulated genes revealed a depletion in chromosome maintenance, DNA replication and RNA processing-related pathways (Fig. 2B). IL-1β up-regulated genes were involved in translation regulation, nonsense-mediated decay, interleukin 1 signaling and NFκB signaling, while the downregulated genes were involved in adherent junctions, cilium structure and assembly, and global DNA packaging (Fig. 2C). GSEA analysis performed on the TNFα dataset showed an over-representation of nonsense-mediated decay, eukaryotic translation and RNA processing related pathways among the up-regulated genes. The down-regulated genes were enriched in pathways related to beta cell development and different signaling pathways (Fig. 2D).

**Figure 2:**
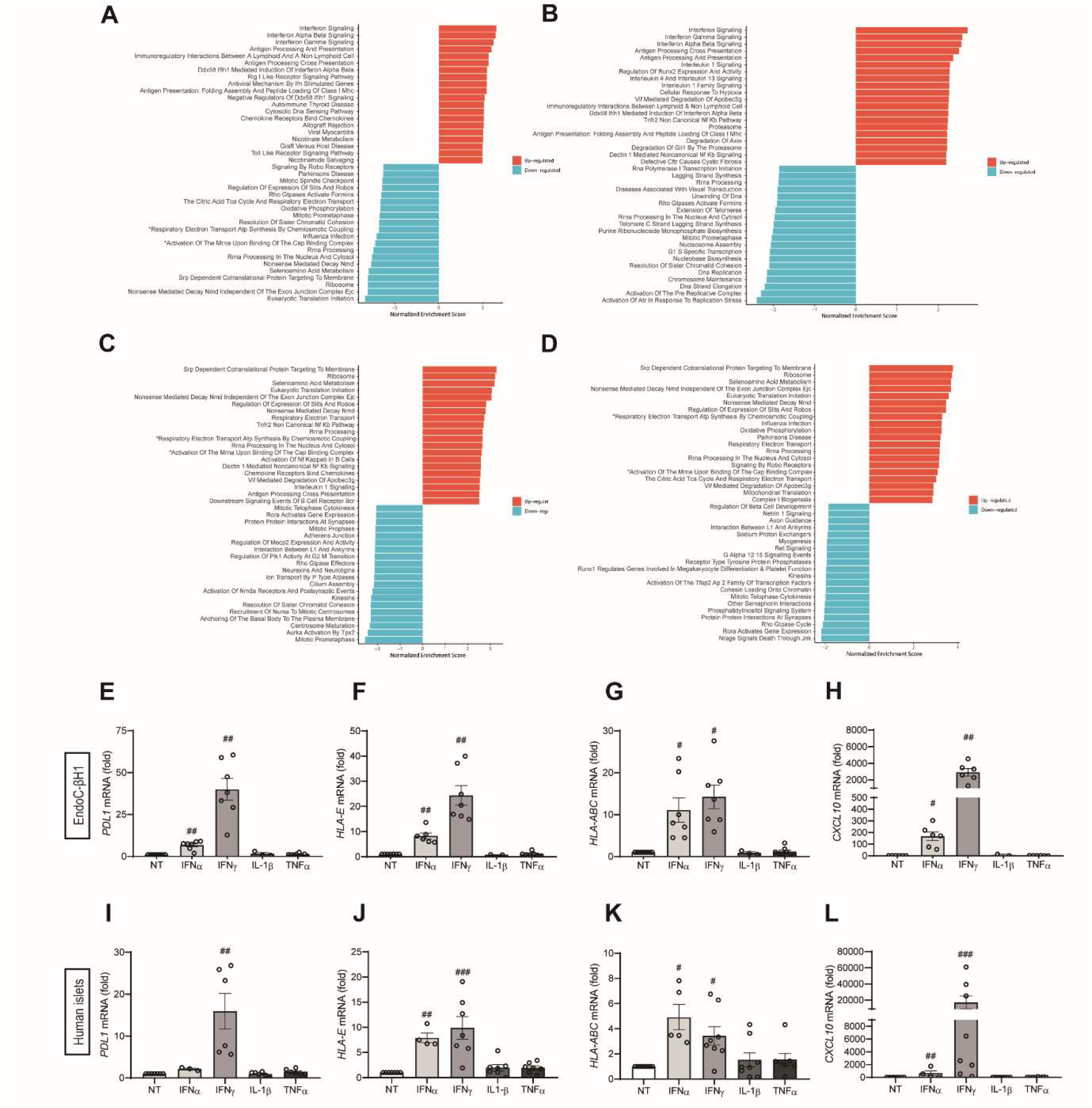
IFNα and IFNγ induce a more marked immune-related response in human beta cells than IL-1β and TNFα. (**A-D**) Results from the differential gene expression analysis were input in fast pre-ranked gene set enrichment analysis (fGSEA). The horizontal bar plots show the significantly enriched pathways for IFN⍺ (**A**), IFNγ (**B**), IL-1β (**C**) and TNF⍺ (**D**). The up-regulated pathways are represented in red and the down-regulated pathways in blue. EndoC-βH1 cells (**E-H**) and dispersed human islets (**I-L**) were left untreated (non-treated: NT) or exposed to IFNα (2000 U/mL), IFNγ (1000 U/mL), IL-1β (50 U/mL) or TNFα (1000 U/mL) for 24h. mRNA expression of *PDL1* (**E, I**), *HLA-E* (**F, J**), *HLA-ABC* (**G, K**) and *CXCL10* (**H, L**) was measured by RT-qPCR and normalized by the geometric mean of *ACTB* and *VAPA* and then by the NT value considered as 1. Results are expressed as mean ± SEM of 2-7 independent experiments. #*p*<0.05, ##*p*<0.01 vs. NT (ANOVA). (**A**) The name of some pathways indicated by an * has been shortened due to space restrictions. Full names are: “Respiratory Electron Transport ATP Synthesis By Chemiosmotic Coupling and Heat Production By Uncoupling Proteins” and “Activation of the mRNA upon binding of the cap binding complex and Eifs and subsequent binding to 43s”.

Together, these data indicate that IFNα and IFNγ are the main drivers of the beta cell responses to pro-inflammatory conditions. To confirm these findings we analyzed the impact of the cytokines on the gene expression of selected relevant immunomodulators, namely *PDL1, HLA-E*, *HLA-ABC* and *CXCL10*, known to be more expressed in pancreatic beta cells or serum of individuals with T1D as compared to non-diabetic individuals^6, 25–27^. IFNα induced the expression of these four genes in EndoC-βH1 cells (Fig. 2E-H), as previously described^6, 9, 23, 28^, and *HLA-E*, *HLA-ABC* and *CXCL10* in dispersed human islets (Fig. 2J-K)^6^. Interestingly, IFNγ also upregulated these four genes in both EndoC-βH1 cells and dispersed human islets (Fig. 2E-L). By contrast, neither IL-1β nor TNFα had any effect on the expression of these genes (Fig. 2E-L). This effect was confirmed in human islet microtissues where only IFNγ at different concentrations (alone or mixed with IL-1β and TNFα), but not IL-1β or TNFα alone, induced HLA class I upregulation after 1, 4 and 7 days of treatment (Supplementary fig. 2A-C). Of note, only the mixture of these three cytokines induced cell death and even a 7-day exposure to individual cytokines failed to induce beta cell death in human islet microtissues (Supplementary fig. 2D-F).

We next evaluated the expression of selected transcription factors known to regulate gene expression mediated by the different cytokines (Supplementary fig. 3A). Apart from NF-κB-related transcriptions factors induced by the four cytokines, the expression of the other transcription factors was induced more strongly by IFNα and IFNγ than by IL-1β and TNFα, with a marked induction of *STAT1*^6^. IFNα and IFNγ up-regulated most IRF family members, with *IRF1*^6^, *IRF7* and *IRF9* being most induced (Supplementary fig. 3A). We validated *IRF7* and *IRF9* induction by IFNα in EndoC-βH1 cells and showed that the silencing of these transcription factors repressed IFNα-induced *HLA-ABC* (only for *IRF7*), *CXCL10*, *PDL1*, *HLA-E* and *MX1* expression (Supplementary fig. 3B-M).

### IFN**α** and IFN**γ** induce a gene signature comparable to that observed in beta cells from T1D patients

RRHO analysis between the different pairs of pro-inflammatory cytokines (Supplementary fig. 4A) showed that IFNα and IFNγ share many differentially expressed genes, with commonly up-regulated genes involved in pathways related mainly to IFN signaling and antigen processing and presentation (Supplementary fig. 4B). However, there was no enrichment for the commonly down-regulated genes. We also observed a close correlation between genes modulated by IL-1β and IFNα (Supplementary figs. 4A and C), which is in line with the central role for NF-kB in signaling of both cytokines (Supplementary fig. 3A). We did not find any significantly enriched pathway for the other pairs of cytokines.

Using the same RRHO approach, we evaluated the similarities between the gene signatures induced by individual cytokines and those observed in FACS-purified beta cells from T1D patients compared to control individuals (Figs. 3A-D). There was a predominance of commonly up-regulated genes between the different pro-inflammatory cytokines and T1D samples, with a closer correlation between IFNs (IFNα and IFNγ) and T1D samples (Fig. 3A-B). For these two cytokines, the commonly up-regulated genes were related to IFN signaling, antigen processing and presentation, chemokine-related signaling and interleukin signaling (Fig. 3E-F). For the commonly up-regulated genes between T1D and IL-1β, there was an enrichment for IFN-related pathways, and pathways related to extracellular matrix organization, histone acetylation and DNA methylation (Fig. 3G). Regarding the commonly up-regulated genes between TNFα and T1D samples, there were pathways related to chromatin modifications and DNA methylation (Fig. 3H).

**Figure 3:**
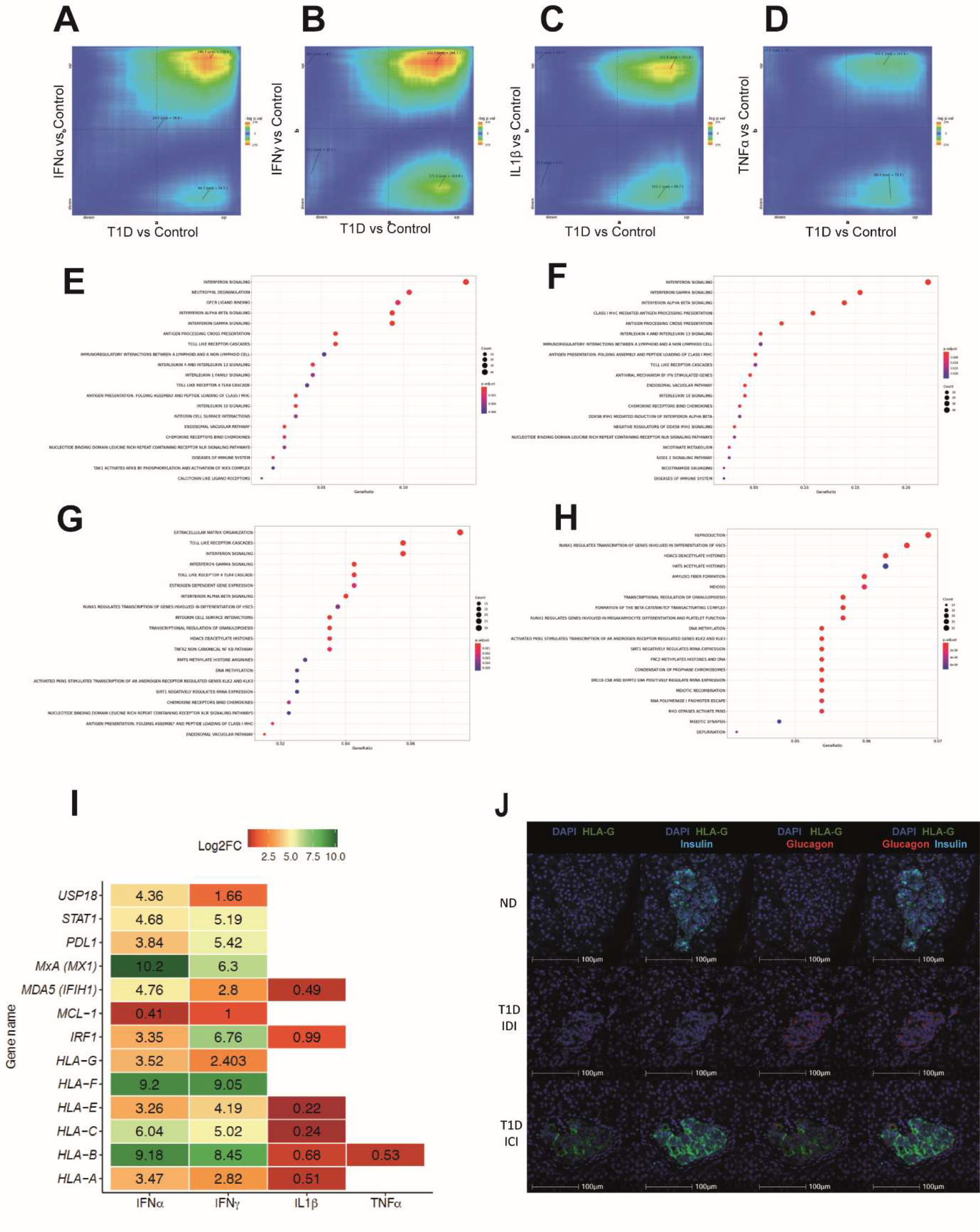
IFNα and IFNγ induce a gene signature similar to the one observed in beta cells from T1D patients. (**A-D**) RRHO algorithm was conducted for each pairwise analysis. The genes were ranked according to their fold change. The significance of overlap between genes commonly up-regulated (top right quadrant), commonly down-regulated (bottom left quadrant), up-regulated by the pro-inflammatory cytokine and down-regulated in FACS-purified beta cells (top left quadrant), and down-regulated by the pro-inflammatory cytokine and up-regulated in FACS-purified beta cells (bottom right quadrant) is represented by the level map. The colors represent the –log(adjusted p values). For comparison purposes, all the level maps are plotted using the same scale that corresponds to the highest scale number when no scale restriction is applied. (**E-H**) Pathway enrichment analysis for the commonly up-regulated genes between IFN⍺ (**E**), IFNγ (**F**), IL-1β (**G**) and TNF⍺ (**H**), and T1D samples was done using gProfiler. (**I**) Heatmap showing statistically significant (adjusted p-value < 0.05) gene expression changes (in Log_2_ Fold Change) after the treatment of EndoC-βH1 cells with IFN⍺, IFNγ, IL-1β or TNFα for 24h of genes known to be altered in T1D pancreatic samples. (**J**) Immunohistochemistry of pancreatic sections stained for HLA-G (green), glucagon (red), and insulin (cyan) from non-diabetic (ND), TD1 insulin-deficient islets (IDI) and T1D insulin-containing islets (ICI) donors.

We selected 12 genes related to the immune and antiviral response that are highly expressed at the protein level in pancreatic islets from T1D donors when compared to controls^25, 26, 29–31^, and investigated the impact of the different cytokines on their expression (Fig. 3I). Reinforcing the close correlation between IFNα- and IFNγ-induced genes as compared to T1D donors, both IFNs induced a significant upregulation of all 12 genes, while IL-1β and TNFα had only a mild impact (6/12 and 1/12 genes induced by these two cytokines, respectively, with a low fold change compared to IFNs). We validated the expression of HLA-G, not previously studied, in human pancreatic islets observing that it was more expressed in insulin-containing islets (ICI) of T1D individuals compared to insulin-deficient islets (IDI) of T1D donors or islets from control subjects (Fig. 3J). Further examples of elevated expression of these proteins in T1D islets are shown in Supplementary Figure 5.

Since IFNs have the most marked impact on the beta cell transcriptome, we evaluated the presence of an ISG signature (defined as described in Methods) in beta cells obtained from non-diabetic controls, individuals presenting 1 or 2 islet-specific autoantibodies (AAB1+ and AAB2+, respectively) or T1D donors, and analysed by single-cell RNA-seq. There was an increase in the ISG score that positively correlated with the natural evolution of the disease, *i.e.* the lowest ISG score was observed in beta cells from non-diabetic individuals and individuals with one autoantibody, but ISG score increased in individuals with two autoantibodies and reached the highest level in beta cells from individuals affected by T1D (Fig. 4A). Furthermore, we analyzed bulk RNA-seq data of FACS-purified beta cells from non-diabetic and T1D donors and again observed a significantly higher ISG score in beta cells isolated from T1D individuals as compared to non-diabetic individuals (nearly 3-fold increase; Fig. 4B). This confirms the relevance of our findings supporting the key role for IFNs on human T1D.

**Figure 4:**
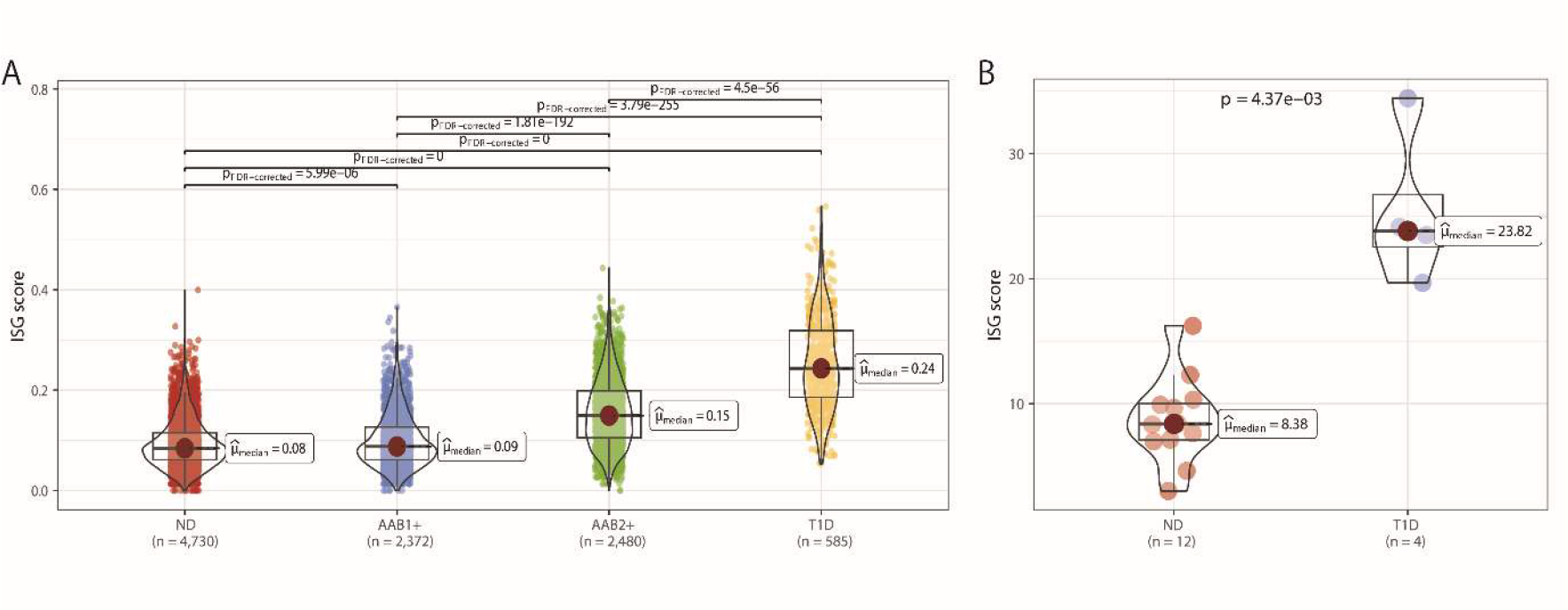
Beta cells from autoantibody-positive and T1D individuals present an elevated interferon-stimulated gene (ISG) score. (**A**) An ISG score (described in Methods) was calculated for beta cells from single-cell RNA-seq data of non-diabetic donors (ND, n = 15), one autoantibody-positive donors (AAB1+, n = 8), 2 or more autoantibody-positive donors (AAB2+, n = 2) and T1D donors (n = 9) obtained from HPAP (https://hpap.pmacs.upenn.edu). (**B**) An ISG score (see Methods) was calculated from bulk RNA-seq data of FACS-purified beta cells of ND (n = 12) and T1D (n = 4) donors^58^. The ISG score was calculated as the average expression of the defined ISGs in each cell for (**A**) and each donor for (**B**). Box limits correspond to the 25^th^ and 75^th^ percentiles, bold lines represent the median score; p-values were determined by the Dunn test for multiple groups comparison in (**A**) and the two-tailed Mann-Whitney U test for (**B**); p value was adjusted for multiple comparisons by “FDR” for (**A**).

### The expression of T1D candidate genes is preferentially affected by IFNα and IFNγ

Genome-wide association studies (GWAS) have identified more than 60 loci that influence T1D risk and many susceptibility genes for T1D – called candidate genes – have been identified in these regions^32^. Since more than 60% of the T1D candidate genes are expressed in pancreatic beta cells^33^, we investigated their regulation by the different individual cytokines and observed that they were mainly regulated by IFNα and IFNγ (51/66 and 41/66, respectively) as compared to IL-1β and TNFα (18/66 and 1/66 respectively) (Fig. 5A). In a previous study, we found that the expression of *BACH2*, one of the candidate genes for T1D, was significantly induced by IFNγ + IL-1β in human beta cells, and *BACH2* silencing enhanced IFNγ + IL-1β-induced beta cell apoptosis^34^. In the present study we observed that both IFNγ and IL-1β, studied separately, as well as IFNα, induced the expression of *BACH2* in EndoC-βH1 cells, while TNFα had no effect (Fig. 5A-B). By contrast, none of these cytokines induced the expression of the *BACH2* paralog, *BACH1* (Fig. 5C). *BACH2* silencing exacerbated all individual cytokine-induced beta cell apoptosis, supporting a key role for this candidate gene in the survival of human beta cells exposed to a pro-inflammatory environment (Fig. 5D-E).

**Figure 5:**
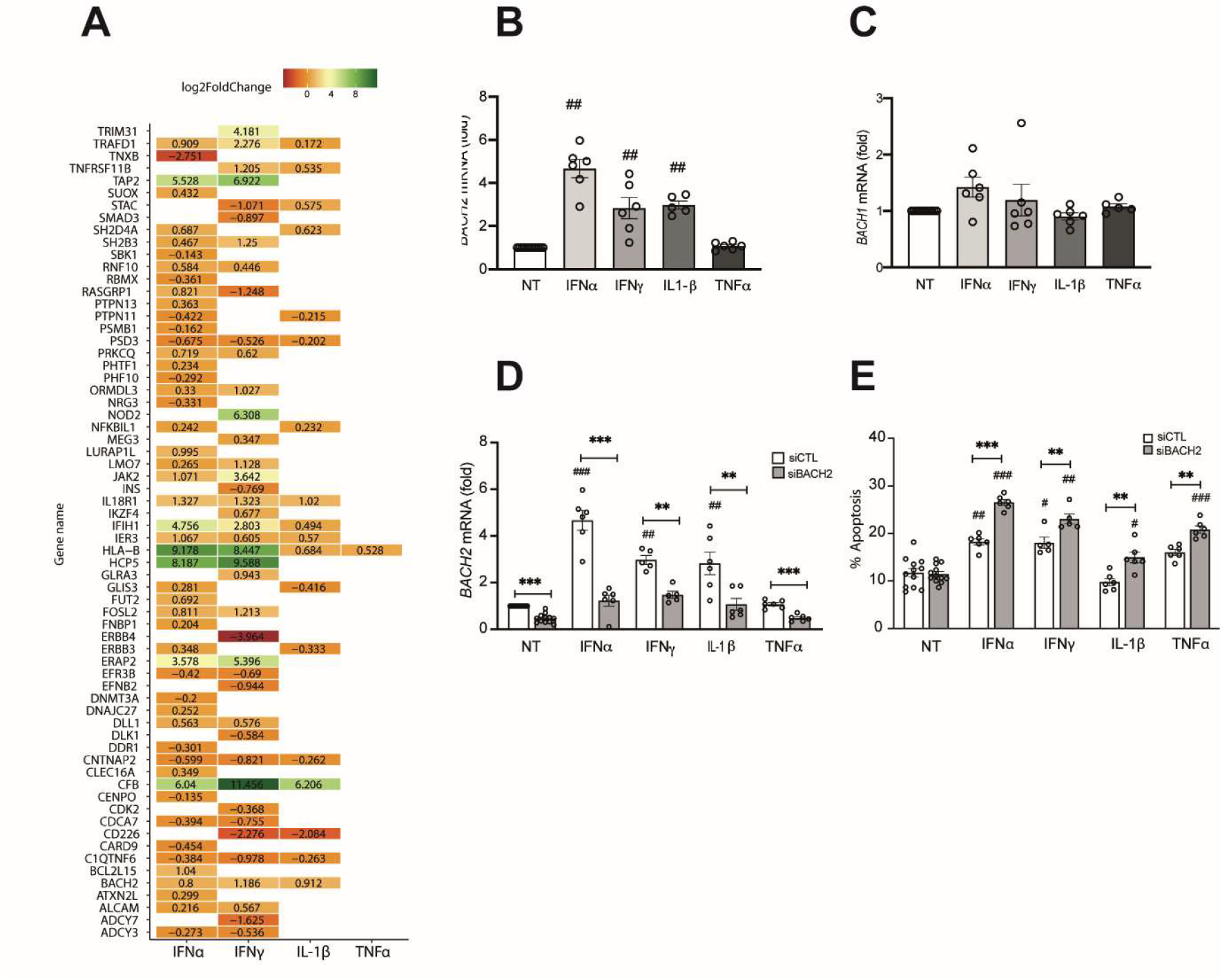
The expression of T1D candidate genes is more affected by IFNα and IFNγ than IL-1β and TNFα in human beta cells. (**A**) Heatmap showing statistically significant (adjusted p-value < 0.05) gene expression changes (Log_2_ Fold Change) of T1D candidate genes after the treatment with the indicated pro-inflammatory cytokines. (**B-C**) EndoC-βH1 cells were left untreated (non-treated: NT) or treated with IFNα (2000 U/mL), IFNγ (1000 U/mL), IL-1β (50 U/mL) and TNFα (1000 U/mL) for 48h. (**D-E**) EndoC-βH1 cells were transfected with a siRNA control (siCTL, white bars) or a siRNA targeting *BACH2* (siBACH2, grey bars). The cells were left untreated (NT) or treated with the respective pro-inflammatory cytokines (at same concentrations than B and C) for 48h. *BACH2* (**B, D**) and *BACH1* (**C**) mRNA expression was evaluated by RT-qPCR and normalized by the geometric mean of *ACTB* and *VAPA* and then by the value of NT considered as 1. (**E**) The impact of BACH2 silencing on EndoC-βH1 cell apoptosis was evaluated by direct cell counting after Hoechst 33342 + propidium iodide staining. Results are expressed as mean ± SEM of 5-6 independent experiments. \**p*<0.05, \*\**p*<0.01 and \*\*\**p*<0.001 vs siCTL treated at the same time point with IFNα, IFNγ or TNFα and #*p*<0.05, ##*p*<0.01 and ###*p*<0.001 vs NT (ANOVA).

### The expression of antiviral and immune-related genes in beta cells is mainly regulated by IFN**α** and IFN**γ.**

Pancreatic beta cells of T1D individuals have a strong antiviral and inflammatory gene signature^10^ and we next investigated the impact of the different cytokines on the expression of 17 genes related to these pathways. Most of these genes were modified by IFNα (15/17) and IFNγ (12/17), while IL-1β had a limited effect (4/17 genes modified), and TNFα did not affect the expression of these genes (Fig. 6A). Recent findings in other cell types suggest that the Zinc Finger NFX1-Type Containing 1 gene (*ZNFX1)*, one of the most interferon-induced antiviral gene in our analysis, is of particular interest for the early antiviral response^35^. We analyzed some of our previous RNA-seq datasets and confirmed *ZNFX1* upregulation in EndoC-βH1 cells (Fig. 6B) and dispersed human islets (Fig. 6C) exposed to IFNα for different time points, starting at 2h. *ZNFX1* expression also tended to be up-regulated in FACS-purified beta cells from T1D individuals as compared to non-diabetic subjects (Fig. 6D). For comparison, we also evaluated the expression of two other well-known double-stranded RNA (dsRNA) sensors, namely *MDA5* (Melanoma Differentiation-Associated Gene 5, also known as *IFIH1*) and *RIG-I* (Retinoic Acid-Inducible Gene I, also known as *DDX58*), involved in the antiviral response of human beta cells. Both genes followed the same pattern of induction as *ZNFX1* in EndoC-βH1 cells (Supplementary fig. 6A, E) and dispersed human islets (Supplementary fig. 6B, F) exposed to IFNα. There was a significant induction as early as 2h post-stimulation, reaching a peak after 8h of treatment. We validated *ZNFX1*, *MDA5* and *RIG-I* induction by IFNα in EndoC-βH1 cells (Fig. 6E-G and Supplementary fig. 6C, G) and showed that *ZNFX1* upregulation was repressed by the JAK1/2 inhibitor Baricitinib (Fig. 6H) or by the silencing of STAT2, but not of STAT1 (Fig. 6I). IFNα also significantly induced *ZNFX1* expression in dispersed human islets in a TYK2-dependent manner as observed with the use of the TYK2 inhibitor BMS-986165 (Fig. 6J). We next analyzed *ZNFX1* expression in beta cells obtained from single-cell RNA-seq data of non-diabetic, AAB1+, AAB2+ and T1D donors (Supplementary fig. 7). There was a gradual increase in the percentage of cells expressing *ZNFX1* and in the average expression in beta cells, from the non-diabetic to the AAB2+ and T1D donors. There was also an increase in *RIG-I* and *MDA5* expression in beta cells from T1D donors, as observed in human beta cells exposed to IFNα (Fig. 6A and Supplementary fig. 6 and 7).

**Figure 6:**
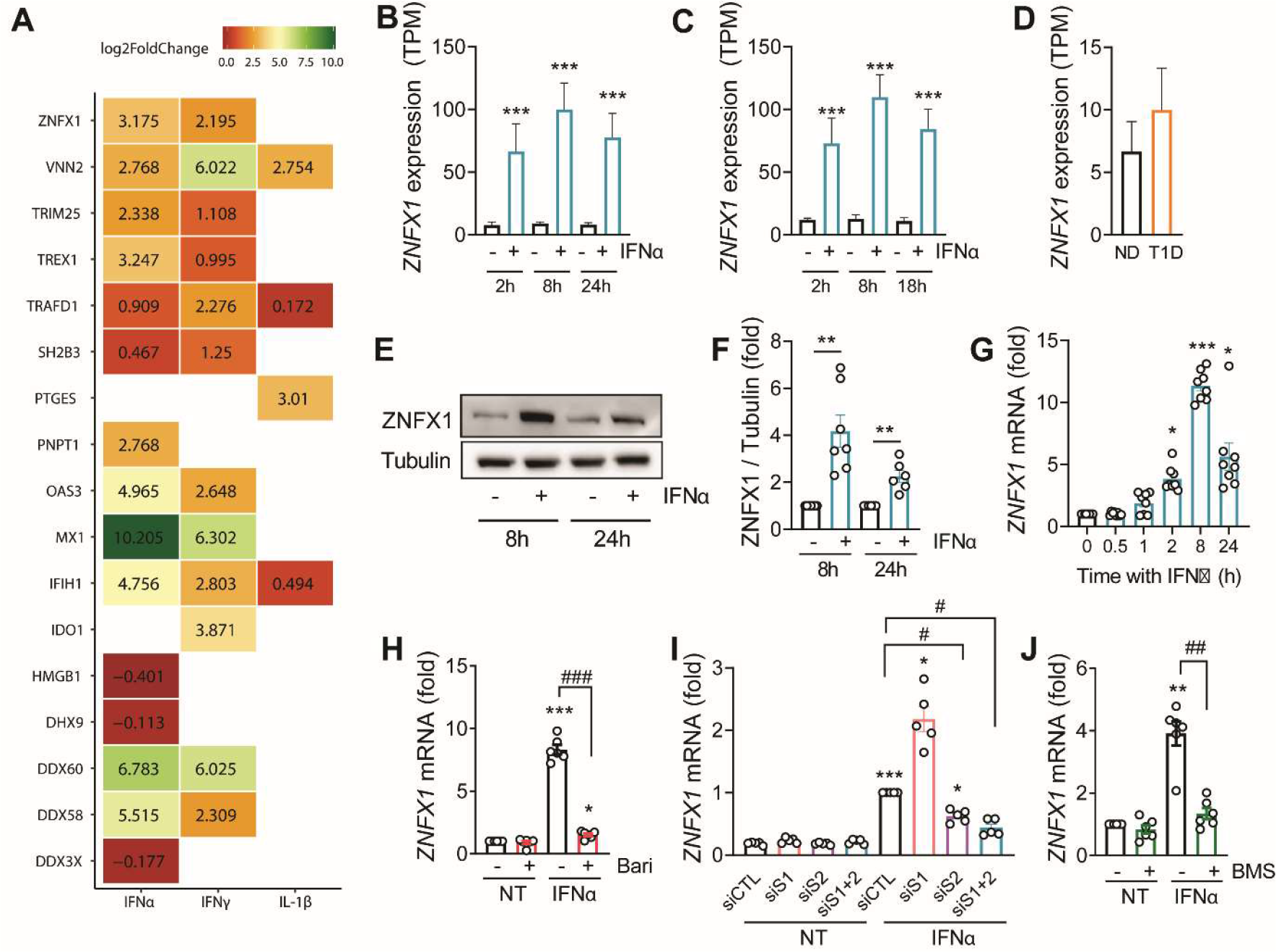
The expression of antiviral and immune-related genes in beta cells is mainly regulated by IFNα and IFNγ. (**A**) Heatmap showing statistically significant (adjusted p-value < 0.05) gene expression changes (in Log_2_ Fold Change) of immune-related genes and antiviral response-related genes after the treatment of EndoC-βH1 cells with IFN⍺, IFNγ or IL-1β for 24h (TNFα did not induce any of these genes and is thus not shown here). *ZNFX1* expression from RNA-seq data of EndoC-βH1 cells (N = 5) (**B**) and human islets (N = 5) (**C**) exposed to IFNα for different time points, and FACS-purified beta cells from non-diabetic (ND) or T1D individuals^58^ (**D**). EndoC-βH1 cells (**E-I**) or dispersed human islets (**J**) were exposed to IFNα (2000 U/mL) alone (**E-G**) or in combination with siRNAs (**I**) or inhibitors (**H, J**). EndoC-βH1 cells (**H**) and dispersed human islets (**J**) were pre-treated for 2h with baricitinib (4 µM) or BMS-986165 (0.3 µM) respectively, and then IFNα was added for an additional 24h. (**I**) EndoC-βH1 cells were transfected with a siRNA control (siCTL: black outline) or with a siRNA targeting STAT1 (siS1: pink outline), STAT2 (siS2: purple outline), or both (siS1+2: blue outline) and then treated with IFNα for 24h. (**E**) Protein expression was measured by western blot and representative images of 6 (24h) or 7 (8h) independent experiments are shown. Densitometry results are shown for ZNFX1 (**F**). mRNA expression of *ZNFX1* (**G-J**) was analyzed by RT-qPCR and normalized by the geomean of *ACTB* and *VAPA* or *ACTB* and *GAPDH* and then by the control value considered as 1. Results are mean ± SEM of 5-8 independent experiments. \**p*<0.05, \*\**p*<0.01 and \*\*\**p*<0.001 vs non-treated condition and #*p*<0.05, ##*p*<0.01 and ###*p*<0.001 as indicated (ANOVA).

**Figure 7:**
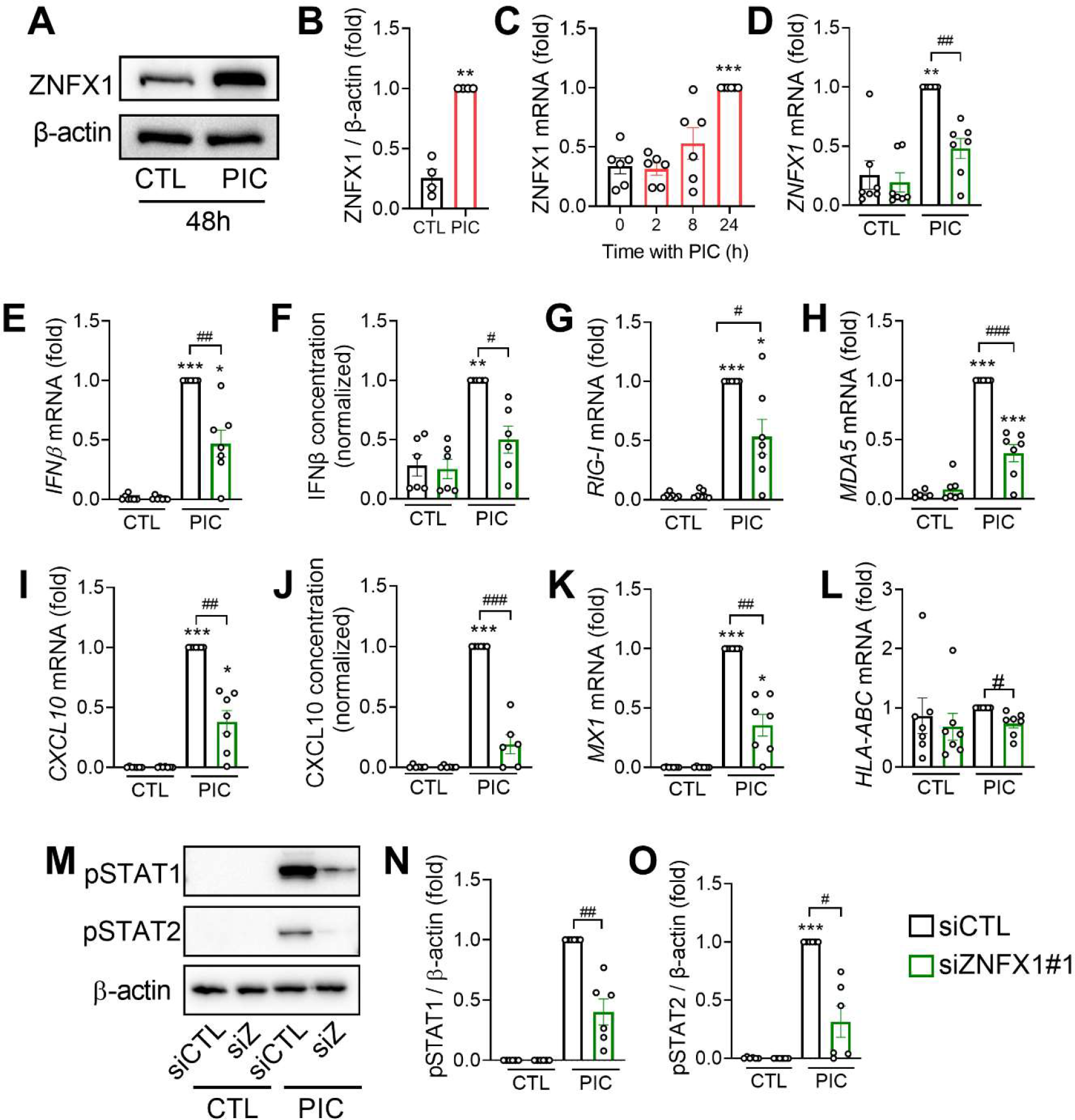
ZNFX1 knock-down prevents the human beta cell response to dsRNA (poly-IC). EndoC-βH1 cells were transfected with poly-IC (PIC, 1 µg/mL) for 48h (**A-B**) or for different periods of time (**C**); or they were transfected with a siRNA control (siCTL: black outline) or with the siRNA#1 targeting ZNFX1 (green outline) for 72h prior to transfection with poly-IC for an additional 24h (**D-N**). Protein expression was measured by western blot and representative images of four (**A**) or six (**M**) independent experiments are shown. Densitometry results are shown for ZNFX1 (**B**), pSTAT1 (**N**) and pSTAT2 (**O**) with the values normalized by β-actin. The mRNA expression of *ZNFX1* (**D**), *IFNβ* (**E**), *RIG-I* (**G**), *MDA5* (**H**), *CXCL10* (**I**), *MX1* (**K**) and *HLA-ABC* (**L**) was analyzed by RT-qPCR and the values normalized by *ACTB*. The release of IFNβ (**F**) and CXCL10 (**J**) to the culture medium (by 35.000 cells/200 µL) was determined by ELISA. In all experiments the values were normalized by the condition PIC (**B, C**) or siZNFX1 + PIC (**D-O**) considered as 1. Results are mean ± SEM of 6-7 (RT-qPCR) or six (ELISA) independent experiments. \**p*<0.05, \*\**p*<0.01 and \*\*\**p*<0.001 vs control (CTL or siCTL + CTL), ##*p*<0.05, ##*p*<0.01 and ###*p*<0.001 vs siCTL + PIC (ANOVA).

### *ZNFX1* is involved in the antiviral response of human pancreatic beta cells

To evaluate the potential antiviral role of ZNFX1 in EndoC-βH1 cells, we first tested its regulation by the dsRNA analog polyinosinic-polycytidylic acid (poly-IC or PIC), used to mimic a viral infection. Treatment with PIC significantly increased ZNFX1 expression both at the protein level after 48h (Fig. 7A-B) and at the mRNA level after 24h (Fig. 7C). *MDA5* and *RIG-I* were also significantly induced after 24h of exposure to PIC (Supplementary fig. 6D, H). To assess the functional role of ZNFX1 in the antiviral response against PIC, we first validated two different siRNAs targeting *ZNFX1* that partially reduced its expression (Supplementary fig. 8A-C). Partial *ZNFX1* silencing reduced PIC-induced *IFNβ* (a central player of the innate immune and antiviral response) expression (Supplementary fig. 8D). Furthermore, in subsequent experiments we observed that *ZNFX1* silencing (Fig. 7D) reduced not only PIC-induced *IFNβ* expression (Fig. 7E) but also IFNβ secretion in the cell supernatant (Fig. 7F). In addition, PIC-induced *RIG-I*, *MDA5*, *CXCL10*, *MX1* and *HLA-ABC* expression, as well as CXCL10 secretion, was repressed by *ZNFX1* silencing (Fig. 7G-L). The phosphorylation of STAT1 and STAT2, the two key transcription factors activated by type I IFN, was also significantly reduced following *ZNFX1* knock-down (Fig. 7M-O). Altogether, these data identified *ZNFX1* as a novel important player of the antiviral response in human beta cells.

### The pro-inflammatory cytokines IFN**α**, IFN**γ**, IL-1**β** and TNF**α** change the alternative splicing landscape of human beta cells

The data described above point to IFNα and IFNγ, but not IL-1β and TNFα, as the main cytokines modulating beta cell transcriptome (Fig. 1E-I). However, modifications that occur at the transcript level may be overlooked when analyzing global differential gene expression only. To address this potential gap, we used transcript-level analysis to identify isoform switches^36^, *i.e.* preferentially expressed isoforms in the non-treated condition that switch to another preferentially expressed isoform following cytokine treatment. Differential transcript usage (DTU) analysis showed that IFNα and IFNγ modified more transcripts than IL-1β and TNFα (Fig. 8A-D), with each dot representing a different transcript plotted according to the difference in inclusion fraction (*i.e.* the ratio between the isoform expression and the gene expression) between the non-treated and the cytokine-treated conditions. Interestingly, some genes that were not differentially expressed in the global analysis had differences in terms of DTU (represented in yellow in the volcano plot in Fig. 8A-D). This observation was particularly relevant for TNFα. Indeed, while this cytokine modified the expression of only 69 genes (Fig. 1I), it induced a switch in isoform usage in 40 additional genes that were not differentially expressed, adding another level of gene regulation (Fig. 8D). We validated the DTU analysis for *CD47* (Cluster of differentiation 47, also known as Integrin-associated protein *IAP*) and *LARP1* (La ribonucleoprotein 1) by RT-PCR in EndoC-βH1 cells and dispersed human islets. Visualization plots reporting structural and functional information about the transcript isoforms produced by these genes, different gene expression (DGE) and DTU are shown in Fig. 8E-F. The DTU analysis indicated that the four *CD47* isoforms expressed in EndoC-βH1 cells were affected differently by TNFα treatment. The canonical isoform ENST00000355354.13 (CD47-201) and the non-coding transcript ENST0000000644850.1 (CD47-206) were unchanged, but there was a switch of isoform usage between the two others: the coding isoform ENST00000361309.6 (CD47-202, mentioned as 09.6) was decreased by TNFα, while the non-coding isoform ENST0000000517766.5 (CD47-205) was increased, resulting in no change in the total gene expression (Fig. 8E). Using primers that allow to discriminate between two different isoforms we observed a significantly reduced proportion of the isoform 09.6 relative to the canonical one in EndoC-βH1 cells treated with TNFα (Fig. 8G), but not in dispersed human islets (Fig. 8H). The DTU analysis of *LARP1* isoforms (Fig. 8F) indicated that TNFα increased the usage of the canonical isoform (ENST00000336314.9, LARP1-201) while decreasing the usage of the coding isoform ENST00000518297.6 (LARP1-204, mentioned as 97.6). These findings were validated in EndoC-βH1 cells and isolated human islets after TNFα treatment (Fig. 8I-J). Altogether, these data indicate that cytokines can also affect beta cell gene expression via a switch in isoform usage, possibly leading to a change in function.

**Figure 8:**
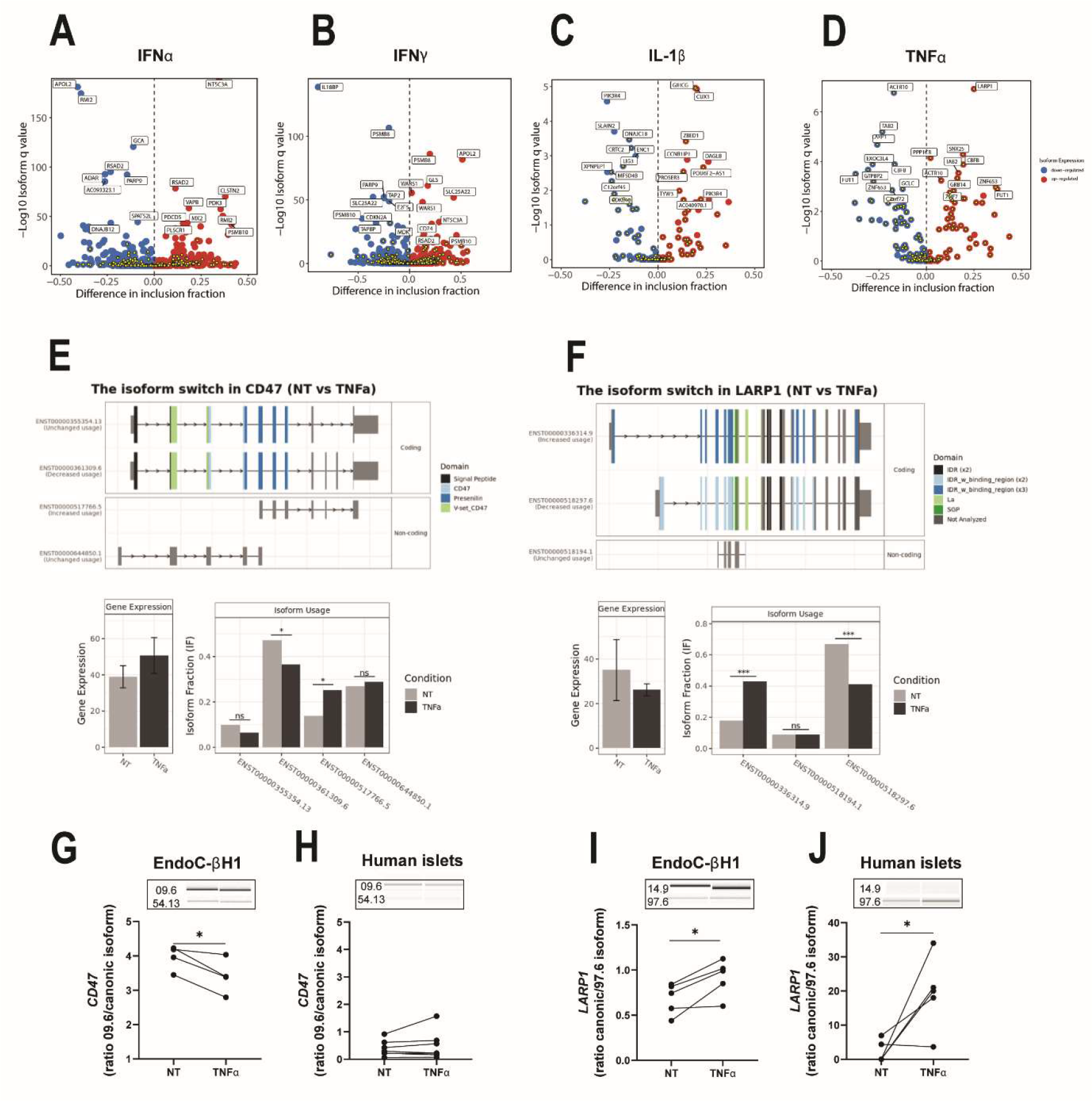
IFNα, IFNγ, IL-1β and TNFα induce differential transcript usage independently of total gene expression. **(A-D)** Volcano plots showing the genes with differential transcript usage after exposure to IFN⍺ (**A**), IFNγ (**B**), IL-1β (**C**) or TNF⍺ (**D**) for 24h. Each dot corresponds to a different transcript, the X axis represents the difference in inclusion fraction measured as the ratio between the isoform expression and the gene expression while the Y axis represents the isoform q value (-log_10_). Blue circles correspond to the down-regulated transcripts and red circles to the up-regulated ones. The yellow dots indicate differentially used transcripts from which the corresponding gene is not differentially expressed under the same experimental conditions. Representation of *CD47* isoform switch comparing NT and TNFα treatment (24h), and validation of the isoform usage in EndoC-βH1 cells (**G**) and dispersed human islets (**H**); results were expressed as ratio between ENST00000361309.6 and ENST00000355354.13 isoforms. Representation of *LARP1* isoform switch comparing NT and TNFα treatment (24h), and validation of the isoform usage in EndoC-βH1 cells (**I**) and dispersed human islets (**J**); results were expressed as ratio between ENST00000336324.9 and ENST00000518297.6 isoforms. cDNA was amplified by RT-PCR using primers located in the upstream and downstream exons of the splicing event and the product evaluated using a Bioanalyzer 2100. (**F-G, I-J**) n = 4-5 (EndoC-βH1) and n = 5-6 (human islets), two-sided paired t-test, \**p*<0.05 vs NT.

## Discussion

We presently identified IFNα and IFNγ as the two main pro-inflammatory cytokines responsible for the transcriptomic modulation of beta cells in the context of T1D, showing that they induce a gene signature that closely correlates with the gene signature observed in beta cells from individuals affected by T1D.

The understanding of the dynamics and natural history of the autoimmune process in T1D has improved through years of intensive research, opening a window of opportunity for the introduction of beta cell protective therapies^12^. Therefore, a good comprehension of the action of the main pro-inflammatory cytokines present in inflamed islets is crucial to identify the most relevant therapeutic targets to preserve beta cell mass and also to define experimental conditions to better model the human disease using human tissues.

IFNα is found in beta cells of individuals with T1D^37^ and a type I IFN gene signature is present in individuals at risk of developing T1D, even before the development of autoantibodies^8^, supporting the role of the innate immune response and IFNα as an early event in T1D pathogenesis. We presently observed that IFNα induces the expression of all the genes encoding proteins previously shown to be overexpressed in pancreatic islets of individuals affected by T1D. IFNα also modulates the expression of most of the candidate genes for T1D and triggers a strong antiviral and inflammatory response in human beta cells. As the disease progresses, there is a transition from innate to adaptive immunity with the local release of pro-inflammatory cytokines – such as IFNγ - and chemokines – such as CXCL10-mostly by the immune cells attracted to the islets in response to the initial inflammatory response^1^. Our GSEA analysis showed that exposure of human beta cells to IFNα and IFNγ, but not IL-1β and TNFα, induced an enrichment in genes involved in antigen presentation and attraction of immune cells, particularly HLA class I and related mRNAs. By doing so, they initiate (IFNα) and then amplify (IFNγ) a dialogue between beta cells and the immune cells that contributes to the transition and amplification of the adaptive immune response against the beta cells. The role of IFNα and IFNγ in the progression of the disease is further reinforced by the gradual increase in the ISG score that we observed from non-diabetic subjects to T1D donors, passing through individuals with AAB2+.

Considering the present findings pointing to IFNα and IFNγ as major players in the deleterious dialogue between the immune system and beta cells, targeting these cytokines for therapy may be of particular interest. The candidate gene *TYK2* is an important mediator of IFNα signaling, acting directly downstream of its receptor to transduce cell signaling in response to type I IFN stimulation. Polymorphisms in the *TYK2* gene confer protection against several autoimmune diseases, including T1D, and it has been suggested that *TYK2* might be a good therapeutic target for T1D^12^. Our group and others previously validated *in vitro* the use of different TYK2 inhibitors in EndoC-βH1 cells, dispersed human islets and human stem-cell derived beta-like cells to block IFNα signaling and also reduce T-cell-mediated cytotoxicity^38, 39^. JAK inhibitors can block the signaling of cytokines that function via the JAK-STAT pathway, including IFNα and IFNγ. Therefore, many JAK inhibitors have been developed (with different selectivity for the four JAK molecules: JAK1, JAK2, JAK3 and TYK2) and studied in the context of T1D. Baricitinib, a JAK1/2 inhibitor, protects human beta cells against cytokine-induced apoptosis and HLA class I upregulation^6^ and is currently under investigation in a phase 2 clinical trial involving patients with recent-onset T1D^40^.

Many studies in rodents have reported an important role for IL-1β in the destruction of beta cells, suggesting the possibility that disease progression might be slowed in humans by blocking its action. Unfortunately, clinical trials with monoclonal anti-IL-1β antibody (canakinumab) or IL1R antagonist (anakinra) failed to preserve beta cell function in T1D patients, suggesting that targeting IL-1β is not effective as single immunomodulatory therapy for T1D^41^. Regarding the present study, this absence of effect is not surprising since human beta cells, but not rat beta cells, have a much higher expression of IFN receptors than IL-1 receptors. On the other hand, golimumab, a monoclonal anti-TNFα antibody already approved to treat several autoimmune diseases such as ulcerative colitis and rheumatoid arthritis, has been tested with more success in a phase 2 clinical trial in children and young adults with recent onset T1D. Golimumab responders had better preservation of C-peptide production and glycemic control than individuals who received a placebo, and this positive result was still present after one year off-therapy^42, 43^. This beneficial effect of golimumab contrasts with the limited impact of TNFα on human beta cell gene expression (present data). Possible explanations for this dichotomy are: 1) The main role of TNFα is on the immune system, and not on the beta cells; 2) TNFα has a more prominent effect in alternative splicing, rather than on the regulation of gene expression; 3) TNFα signaling is associated with deleterious polymorphisms, as recently shown for the T1D candidate gene PTPN2^7^, that may contribute to beta cell death only in a specific cohort of individuals carrying these polymorphisms. Additional studies are now required to distinguish between these possibilities.

The present findings indicate that IFNα also induces a strong antiviral gene signature in human beta cells. Several epidemiological and clinical studies, and experimental data, support the involvement of viral infection, particularly by the enterovirus coxsackievirus B (CVB), in the pathogenesis of T1D^44, 45^. The cytosolic viral dsRNA sensor *MDA5* is a candidate gene for T1D that is expressed in islet cells of T1D individuals^31, 46^ and, together with *RIG-I*, another viral dsRNA sensor, plays an important role in viral detection in beta cells and subsequent induction of type I IFN^45, 47^. The present data show that these two genes are mainly induced by IFNα and IFNγ in human beta cells, with also a mild effect of IL-1β on *MDA5* expression. Besides these two genes, we identified *ZNFX1*^35^ as a new beta cell viral dsRNA sensor, potentially acting in the early stages of viral infections (or exposure to other danger signals) to induce a type I IFN response. *ZNFX1* is induced by IFNα, IFNγ and by the mimic of viral infection poly-IC in human beta cells. Gene silencing of *ZNFX1* indicates that it acts as a viral dsRNA sensor in beta cells exposed to poly-IC, leading to the production of IFNβ and the induction of downstream ISGs, including the two other viral sensors *MDA5* and *RIG-I*. ZNFX1 seems thus to be a critical regulator of the IFN-mediated defense responses in human beta cells and could be an important player in the initiation of the innate immune response in the early stages of T1D.

The primary focus of our present study was on the variation of gene expression induced by different pro-inflammatory cytokines. However, other regulatory mechanisms can significantly affect beta cells. Alternative splicing plays an important regulatory role for beta cell development, function and survival and might contribute to pancreatic beta cell failure and their recognition by the immune system in the context of T1D^4, 6, 48^. The present DTU analysis highlights changes in transcripts that are preferentially used following cytokine exposure, even in the absence of changes in total gene expression. We validated the DTU analysis of *CD47*, a gene that results in a membrane protein expressed in several cell types and is well known for providing a “don’t eat me” signal via binding to SIRPα on macrophages, preventing phagocytosis^49^. Interestingly, CD47 is strongly expressed in human beta cells and regulates their viability; CD47 expression is altered in T1D with differences observed between putative T1D endotypes^50^. In the present study, we observed that although TNFα did not alter *CD47* total gene expression there was a reduced expression of a protein-coding transcript counterbalanced by the increase in a non-coding transcript.

In conclusion, we have applied a deep RNA-sequencing approach to study the individual impact of the four major pro-inflammatory cytokines involved in T1D – IFNα, IFNγ, IL-1β and TNFα - on the human beta cell transcriptome. Our data identified IFNα and IFNγ as the main drivers of the beta cell response, inducing a gene signature that strongly correlates with the transcriptomic profile of beta cells from T1D individuals. Therefore, therapies that aim to protect beta cells from an immune attack should focus on these cytokines to stop the dialogue between beta cells and immune cells and break the infernal circle of immune amplification.

## Methods

### Culture of EndoC-βH1 cells and human islets

The human insulin-secreting EndoC-βH1 cells were provided by Dr. R. Scharfmann (Institut Cochin, Université Paris Descartes, Paris, France)^51^. EndoC-βH1 cells were cultured in DMEM containing 5.6 mmol/L glucose (Gibco, Thermo Fisher Scientific, Waltham, MA, USA), 2% BSA fraction V, fatty acid free (Roche, Basel, Switzerland), 50 μmol/L 2-mercaptoethanol (Sigma-Aldrich, St Louis, MO, USA), 10 mmol/L nicotinamide (Calbiochem, Darmstadt, Germany), 5.5 μg/mL transferrin (Sigma-Aldrich), 6.7 ng/mL sodium selenite (Sigma-Aldrich), 100 U/mL penicillin + 100 μg/mL streptomycin (Lonza, Leusden, Netherlands), in matrigel-fibronectin-coated plates, as previously described^20^.

Human islets were isolated from non-diabetic organ donors by collagenase digestion and density gradient purification^52, 53^ and cultured as previously described^6^. Beta cell purity was determined by insulin staining using immunofluorescence technique^54^. All the experimental replicates correspond to different donors, and the donor characteristics are detailed in Supplementary Table S2. The use of pancreatic human islets for the present project was approved by the Comité d’Ethique Hospitalo-Facultaire Erasme-ULB.

### RNA-sequencing of EndoC-**β**H1 cells

After treatment with the respective cytokines for 24h, total RNA was isolated from EndoC-βH1 cells using the RNeasy Plus Micro Kit (Qiagen, Venlo, Netherlands), according to the recommended protocol. For all the samples, a minimum of 500 ng of high-quality RNA with RNA integrity number (RIN) > 9 (determined using Agilent Bioanalyzer) was used for the library preparation. The RNA-seq was performed on NovaSeq 6000 (Eurofins Genomics Europe Sequencing GmbH, Konstanz, Germany).

### RNA-sequencing analysis

The raw RNA-seq data of EndoC-βH1 cells treated with IFNγ, IL-1β, and TNFα [2 x 150-bp paired-end with a sequencing depth > 200 million reads] were first subjected to a quality check using fastp^55^ to remove low quality reads and adaptor trimming. The passing reads were then subjected to quality control (QC) using FastQC (Babraham Bioinformatics, Cambridge, UK). The five samples of RNA-seq data from EndoC-βH1 cells treated with IFNα used in the present study were previously published by our group^6^ (available at NCBI Gene Expression Omnibus (GEO) under accession code GSE148058). For comparison purposes, these five paired RNA-seq replicates were re-analyzed as described below. Gene expression was quantified using Salmon v1.4^56^ with additional parameters “-seqBias-gcBias - validateMappings” and expressed in TPM (transcripts per million). GENCODE v36 was used as the reference genome and indexed with default k-mer values. The genes differentially expressed after exposure to cytokines were identified by DESeq2 v1.30.1^57^. For each gene, a Log_2_ Fold Change value was obtained based on the comparison performed (cytokine-treated condition *versus* non-treated condition). Each gene comparison has an associated Wald test statistic, p-value, and an adjusted p-value (Benjamini-Hochberg correction). Genes presenting an adjusted p-value < 0.05 were considered significantly differentially expressed. Bulk RNA-seq data of FACS-sorted beta cells from non-diabetic (n = 12) and T1D (n = 4) donors from the GEO portal (GSE121863)^58^ were analyzed using the same pipeline to enable data comparison. From our RNA-seq data, we generated a “beta cell ISG-signature” (ISG score) based on the selection of mRNAs that had a >3-fold upregulation following treatment of EndoC-βH1 cells with IFNα or IFNγ but were not induced by IL-1β or TNFα. The 53 selected ISGs were: *IRF1*, *IRF2*, *IRF7*, *IRF8*, *IRF9*, *STAT1*, *STAT2*, *STAT3*, *C1QB*, *C4A*, *CCL14*, *CIITA*, *CSF1*, *CXCL10*, *CXCL11*, *CXCL14*, *CXCL17*, *CXCL8*, *CXCL9*, *HCP5*, *HLA-A*, *HLA-B*, *HLA-C*, *HLA-DMB*, *HLA-DQB1*, *HLA-DRA*, *HLA-E*, *HLA-F*, *IL12A*, *IL15RA*, *IL18BP*, *IL6*, *NLRC5*, *PSMB10*, *PSMB8*, *PSMB9*, *TNFRSF1B*, *CASP1*, *RIG-I*, *DDX60*, *IDO1*, *IFI16*, *MDA5*, *MX1, OAS1*, *OAS3*, *PNPT1*, *TREX1*, *ZBP1*, *ZNFX1*, *SOCS1*, *SOCS3*, *USP18*. The ISG score was calculated as the average expression of these selected 53 ISGs21,59.

### Gene set enrichment analysis

Gene set enrichment analysis (GSEA) was done using the previously generated differentially gene expression (DGE) results. The enrichment or depletion in metabolic pathways was assessed with fGSEA^60^ using KEGG^61^ and REACTOME^62^ databases as references.

### RRHO analysis

Rank-rank hypergeometric (RRHO) test is used to evaluate the level of agreement between two ranked lists^63, 64^. Briefly, RRHO was used to evaluate the concordance between gene expression signatures of the different cytokines (IFNα, IFNγ, IL-1β, and TNFα), and also between each cytokine with the gene expression signature observed by RNA-seq on FACS-purified beta cells from T1D patients^58^. The RRHO map interpretation is detailed in Supplementary fig. 4 and the respective figure legend.

### Cell treatment and small RNA interference

EndoC-βH1 cells and dispersed human islets were exposed or not to IFNα (2000 U/mL, PeproTech Inc., Rocky Hill, NJ), IFNγ (1000 U/mL, PeproTech Inc.), IL-1β (50 U/mL, R&D Systems, Minneapolis, MN, USA), and TNFα (1000 U/mL, PeproTech Inc.) for 24h. After this period, the cells were washed with PBS and harvested for RNA and protein extraction. In some experiments, the cells were pre-treated for 2h with 4 µM of the JAK1/2 inhibitor Baricitinib (Selleckchem, Planegg, Germany)^6^ or 0.3 µM of the TYK2 inhibitor BMS-986165 (MedChemExpress, Monmouth Junction, NJ, USA)^39^ or their respective vehicle before treatment with IFNα.

EndoC-βH1 cells were transfected with siRNAs targeting *IRF7*, *IRF9*^9^, *ZNFX1* and *BACH2* at a final concentration of 30 nM, using previously described transfection conditions^9^. The Allstars unspecific control siRNA (Qiagen) was used as a negative control (siCTL). The sequence of the siRNAs used in the present study is detailed in Supplementary Table S3.

EndoC-βH1 cells were transfected for 24h with the synthetic dsRNA analog polyinosinic-polycytidylic acid (PIC, P0913, Sigma-Aldrich) at a final concentration of 1 µg/mL using Lipofectamine 2000 (Invitrogen, Carlsbad, CA, USA).

### Cell viability assessment

The percentage of apoptotic EndoC-βH1 cells and dispersed human islets was determined using fluorescent microscopy after 15 minutes of incubation with the DNA binding dyes Hoechst 33342 (10 µg/mL, Sigma-Aldrich) and propidium iodide (10 µg/mL, Sigma-Aldrich)^29, 30^. The cell counting was performed by two observers, one of them unaware of the identity of the samples, and the agreement between them was >90%. The results are expressed as percentage of apoptosis (number of apoptotic cells / total number of cells).

### mRNA extraction, quantitative real-time PCR and validation of the differential transcript usage

After the respective treatments, Poly(A)+ mRNA was isolated using the Dynabeads mRNA DIRECT Kit (Invitrogen), according to the manufacturer’s recommendations. The mRNA molecules were retrieved in Tris-HCl elution buffer. The mRNA was reverse transcribed using the Reverse Transcriptase Core Kit (Eurogentec, Liège, Belgium) following the manufacturer’s instructions. The quantitative RT-PCR (RT-qPCR) amplification reactions were conducted using IQ SYBR Green Supermix (Bio-Rad, Hercules, CA, USA) and ran in the CFX Connect Real-Time PCR Detection System (Bio-Rad). The PCR product concentration was determined using the standard curve method^65^, and it was expressed as number of copies per μL. The target gene expression level was expressed as a ratio of the geometric mean of the two reference genes *ACTB* and *VAPA* mRNAs, or only *ACTB* for experiments performed before our identification of *VAPA* as a useful reference gene^66^.

For the validation of the differential transcript usage (DTU) analysis the cDNA sequences of statistically significant over- and down-represented transcripts of a specific gene were aligned using the pairwise sequence alignment tool EMBOSS Needle to identify isoform-specific sequence variations. Whenever possible, the different isoforms were detected by RT-PCR using manually designed primers annealing to the flanking common exons of the distinct or mismatched genomic region between the affected isoforms. This approach allowed the simultaneous amplification of different regions that correspond to the different isoforms of a particular gene, based on their different amplicon size.

RT-PCR was done using the RedTaq DNA Polymerase (Bioline, Memphis, TN, USA) following the recommended protocol. The analysis of the PCR products was performed using the LabChip electrophoretic Agilent 2100 Bioanalyzer microchip electrophoresis system (Agilent Technologies, Wokingham, UK). Briefly, after the PCR reaction, 1 μL of each PCR product was loaded into a DNA 1000 LabChip kit, prepared according to the supplied protocol. The PCR products were then separated using the LabChip electrophoretic Agilent 2100 Bioanalyzer system, and the molarity of each PCR product corresponding to a specific transcript isoform was quantified using the 2100 Expert Software, and it was used to calculate the ratio between the isoforms. The primers used in the present study are listed in Supplementary Table S4.

### Protein extraction, western blotting analysis and ELISA

After treatment, EndoC-βH1 cells were washed with PBS, lysed using 1x Laemmli Sample Buffer (60 mM tris(hydroxymethyl)aminomethane pH 6.8, 10% Glycerol, 2% Sodium dodecyl sulfate, 1.5% 2-mercaptoethanol, 1.5% Dithiothreitol and 0.005% bromophenol blue) and proteins were analysed by western blot. To evaluate ZNFX1 expression we used a monoclonal rabbit anti-ZNFX1 antibody. To evaluate the type I IFN signaling activation we measured STAT1 and STAT2 phosphorylation using a rabbit anti-phospho-STAT1 and a rabbit anti-phospho-STAT2 antibody. The data were normalized for the expression of the housekeeping genes β-actin or α-tubulin. Secondary donkey anti-rabbit and anti-mouse antibodies coupled with the horseradish peroxidase (HRP) were used and the signal was detected using the SuperSignal West Femto chemiluminescent substrate (Thermo Fisher Scientific). Immunoreactive bands were visualized using ChemiDoc XRS+ (Bio-Rad) and quantified with the Image Studio Lite v5.2 software (LI-COR Biosciences, Lincoln, NE, USA). The list of antibodies and conditions of use is provided as Supplementary Table S5.

IFNβ and CXCL10 release in the culture medium of EndoC-βH1 cells was measured using enzyme-linked immunosorbent assays (Quantikine ELISA kit, R&D Systems). Each condition was measured from the supernatant of 35.000 cells cultured in 200 µL medium and culture conditions are indicated in the legend of the figures.

### HPAP single-cell RNA sequencing data analyses

Raw FASTQ files of single cell RNA-sequencing (scRNA-seq) data from human pancreatic islets were downloaded from the Human Pancreas Analysis Program (HPAP) (https://hpap.pmacs.upenn.edu), which includes samples from non-diabetic donors, individuals presenting 1 islet-specific autoantibody (AAB1+), individuals presenting 2 islet-specific autoantibodies (AAB2+), and T1D donors. The analysis was performed as previously described by our group^48^. Briefly, the raw scRNA-seq data files (FASTQ) were aligned on the human genome GRCh38 using Cell Ranger v6.1.2, following the recommendations provided by 10X Genomics. We analyzed samples from 15 non-diabetic donors (HPAP-044, HPAP-039, HPAP-104, HPAP-047, HPAP-034, HPAP-036, HPAP-026, HPAP-082, HPAP-099, HPAP-027, HPAP-035, HPAP-056, HPAP-037, HPAP-040, HPAP-022), 8 AAB1+ (HPAP-024, HPAP-029, HPAP-038, HPAP-045, HPAP-049, HPAP-50, HPAP-072, HPAP-092), 2 AAB2+ (HPAP-043, HPAP-107) and 9 T1D donors (HPAP-020, HPAP-021, HPAP-023, HPAP-032, HPAP-055, HPAP-064, HPAP-071, HPAP-084, HPAP-087). Raw FASTQ data (10x Genomics) was processed with Cell Ranger (V6.1.2)^67^ regarding quality control, read alignment to a reference genome (hg38), barcode processing and molecule counting. For each sample we performed an initial clustering using the Seurat software (V4.1.1)^68^, followed by decontamination of the background mRNA with SoupX (V1.6.1)^69^ using genes (INS, GCG, SST, TTR, IAPP, PYY, KRT19 and TPH1) representing the major cell types pre-identified in the initial clustering. The adjusted gene expression count was imported into Seurat software for further filtering. Potential doublet cells were evaluated and removed using scDblFinder (V3.16)^70^. We further filtered the cells using the criteria of nFeature_RNA >200 and <9000, percent of mitochondrial genes <15% and nCount_RNA <10000. We obtained 79,545 single cells for downstream analyses. Next, we used Seurat’s SCTransform function to measure the differences in sequencing depth per cell and normalize the counts by removing the variation due to sequencing depth. Since mitochondrial genes are indicators of cell state, we also regressed the variation from the percentage of mitochondrial reads. The top 3000 variable genes were selected to perform a Principal Component Analysis (PCA). Harmony (V0.1.1) software was used to integrate the data using the top 50 PCA components, considering the sample ID and batches of Reagent Kits as confounding factors. The integrated components were subsequently used to construct a Uniform Manifold Approximation and Project (UMAP) embedding of the 79,545 cells. Lastly, scSorter (V0.0.2)^71^ was used to annotate cell types based on the known marker genes. We obtained 10,167 beta cells and calculated the ISG score (see method above) for each of these cells.

### Differential transcript usage

Differential transcript usage (DTU) analysis measures the proportional differences in the expressed transcript composition of a gene, and the relative contribution of these transcripts for the expression of that specific gene. DTU analysis was performed using the R library IsoformSwitchAnalyzeR^36^ following the pipeline instructions, and DEXSEq to test differential isoform usage (dIF). The files used as input in the *isoformSwitchAnalysisPart2* function were generated externally by running Coding Potential Calculator 2 (CPC2), Pfam, Signal peptide (SignalIP 6.0) and IUPred2A, as recommended in the pipeline. We selected all the isoforms with overall false discovery rate (OFDR) < 0.05, and |dIF > 0.05|.

### Human islet for microtissue production at InSphero, culture and treatment

Human islets were acquired from Prodo Laboratories Inc (Irvine, CA, USA). All islets were procured from deceased donors with consent obtained from their next of kin. To generate 3D InSight™ Islet Microtissues (MTs) (InSphero AG, Schlieren, Switzerland), human islets were dispersed using dissociation solution (1X TrypLE™ Express solution - Thermo Fisher Scientific, with 40 µg/mL DNase I - Sigma-Aldrich) through gentle pipetting for 10 minutes at 37°C, as previously described^72–74^. Approximately 1700 dispersed live islet cells were reaggregated in each well of the Akura™ PLUS Spheroid Hanging Drop System for 5 days (InSphero AG, CS-06-001-02) to achieve a volume equivalent to approximately 1 islet (IEQ) per MT, corresponding to an islet with a diameter of 150 µm. The reaggregated islets were then transferred and cultured in Akura™ 96 Spheroid Microplates (InSphero AG) using 3D InSight™ Human Islet Maintenance Medium (InSphero AG). The islet MT cultures were maintained at 37°C in a humidified atmosphere containing 5% CO_2_, with the cell culture medium being replaced every 2-3 days.

Islet MTs were treated with IL-1β (2, 5 or 10 ng/mL, Sigma-Aldrich), IFNγ (10, 25 or 50 ng/mL, Sigma-Aldrich), or TNFα (10, 25 or 50 ng/mL, Sigma-Aldrich), or a cocktail of these cytokines for a duration of 1, 4 or 7 days. The culture medium was replenished with fresh cytokines every 2-3 days.

### Immunofluorescence staining of islet microtissues, confocal imaging and image analysis

The islet MTs were fixed using a paraformaldehyde (PFA) solution 2%, permeabilized with 0.5% Triton X-100 in PBS, and blocked with 10% fetal calf serum (FCS). Overnight incubation with mouse anti-HLA class I-ABC and 4h incubation with goat anti-Mouse Alexa Fluor 488 (Supplementary Table S5) and DAPI (1 mg/mL, Sigma-Aldrich) were performed in 10% FCS and 0.2% Triton X-100 in PBS. After each antibody incubation, non-specific binding was eliminated by rinsing the MTs with 0.2% Triton X-100 in PBS through multiple washing steps. The stained MTs were then transferred to Akura™ 384 Spheroid Microplates (InSphero AG) and treated with ScaleS4 solution (consisting of 40% w/v D-(-) sorbitol, 10% w/v glycerol, 4M Urea, 0.2% w/v Triton X-100, and 15% v/v DMSO in MilliQ water) for clearing. Imaging was conducted with a Yokogawa CQ1 confocal benchtop high-content analysis system (Yokogawa Electric Corp., Tokyo, Japan) equipped with a 40x dry objective lens (Olympus, Tokyo, Japan). Image stacks were acquired with a Z-step size of 3 µm to cover the entire MT structure. The analysis procedure with the Yokogawa CellPathfinder software encompassed three distinct steps. To obtain the nuclear count in islet MTs, the entire spheroid region was initially identified by applying Otsu thresholding and a Gaussian function to the DAPI channel. Subsequently, the nuclei were individually segmented and labelled using a dynamic thresholding approach in the DAPI channel. The analysis was performed in 3D to prevent double counting. Finally, mean HLA signal intensity was calculated for the whole spheroid region.

### Pancreatic samples

Pancreatic tissue from the Exeter Archival Diabetes Biobank (https://pancreatlas.org/datasets/960/overview) was used in this study. This biobank contains paraffin-embedded human pancreas samples recovered from people who died between 1940 and 1995 with recent-onset T1D, and individuals without diabetes. Ethical approval for research with the Biobank was granted by the West of Scotland Research Ethics Service (REC reference 15-WS-0258). Immunostaining was performed on 4-µm-thick sections of tissue.

### Immunofluorescence of pancreatic samples

After dewaxing and rehydration, sections were subjected to heat–induced epitope retrieval (HIER) in 10 mM citrate pH 6 buffer for 20 minutes. After blocking, the sections were probed in a sequential manner with relevant primary antibodies (Supplementary Table S5), followed by detection with either goat anti-rabbit Alexa Fluor™ 488 or TSA Alexa Fluor™ 488-conjugated goat anti mouse/ rabbit versions. Following washes, and if using TSA-conjugates a further HIER, sections were then probed with mouse monoclonal anti-glucagon followed by detection with goat anti-mouse Alexa Fluor™ 555. Finally sections were probed with guinea pig anti-insulin antibody, followed by detection with goat anti-guinea pig Alexa Fluor™ 647 in the presence of DAPI (1 µg/mL) to identify cell nuclei. After mounting, the sections were imaged via a Leica DMi8 confocal microscope (Leica Microsystems UK, Milton Keynes, UK) or sections were scanned at 40x magnification using an Akoya Biosciences Vectra® Polaris™ Automated Quantitative Pathology Imaging System. The distribution of proteins of interest within beta cells (insulin+) or alpha cells (glucagon+) was examined in multiple islets. The list of antibodies and conditions of use is provided in Supplementary Table S5.

### Statistical analysis

The data are shown as means ± SEM. A normality test was performed to assess Gaussian distribution and outliers were detected and eliminated using the outliers removal test Grubbs (alpha = 0.05). When the distribution was normal, significant differences between experimental conditions were determined by paired t-test or one-way analysis of variance (ANOVA) followed by Bonferroni post-hoc. If the distribution was not normal, the non-parametric test equivalent to one-way ANOVA was used (Friedman or Kruskal-Wallis if values were missing). P value < 0.05 was considered statistically significant. For the experiments using InSphero microtissues, outliers were detected and eliminated using the robust regression and outlier removal test (ROUT) with a significance level (Q) of 5%. A conventional one-way ANOVA with Dunnett post-hoc test was conducted to compare each condition with the control. Data analysis and visualization were performed using Prism software version 8 and ggplot2 package with R. The RNA-seq was analyzed as described above.

## Data availability

All newly generated RNA-seq data that support the findings of the present study have been deposited on GEO under accession code GSE235683.

## Acknowledgements

The authors are grateful to A.M. Musuaya, L. Sauvage, Y. Cai and I. Millard of the ULB Center for Diabetes Research, Université Libre de Bruxelles, Belgium, for excellent technical support, and to J. Hopkinson, C. Flaxman, J. Hill, R. Wyatt, M. Russell and K. Murrall from EXCEED for imaging assistance. D.L.E. acknowledges the support of grants from the Juvenile Diabetes Foundation International (3-SRA-2022-1201-S-B(1) and 3-SRA-2022-1201-S-B (2)), Welbio-FNRS (Fonds National de la Recherche Scientifique) (WELBIO-CR-2019C-04), Belgium; the Dutch Diabetes Research Foundation (Innovate2CureType1), Holland; the JDRF (3-SRA-2022-1201-S-B); the National Institutes of Health Human Islet Research Network Consortium on Beta Cell Death & Survival from Pancreatic β-Cell Gene Networks to Therapy [HIRN-CBDS]) (grant U01 DK127786) and the National Institutes of Health Grants, NIDDK, grants RO1DK126444 and RO1DK133881-01. D.L.E. P.M., S.J.R. and N.G.M. acknowledge the support from the Innovative Medicines Initiative 2 Joint Undertaking under grant agreements 115797 (INNODIA) and 945268 (INNODIA HARVEST), supported by the European Union’s Horizon 2020 research and innovation programme. These joint undertakings receive support from the European Union’s Horizon 2020 research and innovation programme and European Federation of Pharmaceutical Industries and Associations (EFPIA), JDRF, and the Leona M. and Harry B. Helmsley Charitable Trust. F.S. is supported by a Research Fellow (Aspirant) fellowship from the FNRS (Belgium). X.Y. is supported by the Fondation ULB, Wallonie-Bruxelles International (WBI) and the China Scholarship Council. A.O.d.B. is supported by a grant (CDR) from the FNRS (35248676). E.M. acknowledges the support of grants PI19/00246 and PI22/00334 from Instituto de Salud Carlos III, co-financed by the European Regional Development Fund (ERCF). Parts of this study were supported by the National Institute for Health and Care Research Exeter Biomedical Research Centre. The views expressed are those of the author(s) and not necessarily those of the NIHR or the Department of Health and Social Care.

## Author contributions

A.C.d.B., M.I.A. and F.S. contributed to the original idea, to the design, performance and interpretation of the experiments, investigation and formal analysis, and wrote, revised and edited the manuscript. P.L.Z. performed and analysed experiments, and wrote, revised and edited the manuscript. A.C., B.M.d.S., S.J.R., N.G.M., A.R.R., S.M.C., X.Y., A.O.d.B., S.S., S.J., A.C.T. and B.Y. performed and analysed experiments. F.P., J.K.C, E.M., M.N., L.M. and P.M. contributed with material. D.L.E. contributed to the original idea and the design, supervision and interpretation of the experiments, and wrote and revised the manuscript. All authors have read and approved the final version of the manuscript. A.C.d.B., M.I.A. and D.L.E. are the guarantors of this work and, as such, have full access to all the data in the study and take responsibility for the integrity of the data and the accuracy of the data analysis.

## Material and correspondence

All the data that support the findings of the present study are available from the corresponding authors upon reasonable request.

