## Supplementary figures and tables for "Interferons are the key cytokines acting on pancreatic islets in type 1 diabetes"

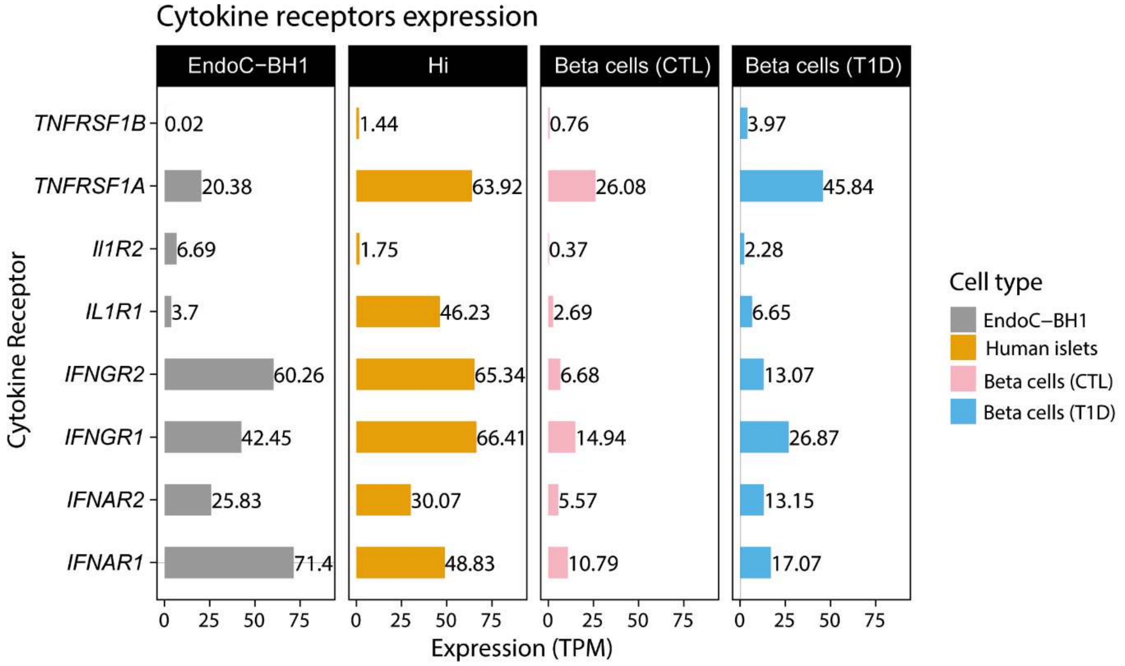


**Supplementary figure 1: Gene expression of the different cytokine receptors on different models of human pancreatic beta cells.** (**A**) Gene expression of IFN$\alpha$ (*IFNAR1, IFNAR2*), IFN$\gamma$ (*IFNGR1, IFNGR2*), IL-1$\beta$ (*IL1R1, IL1R2*), and TNF$\alpha$ (*TNFRSF1A, TNFRS1B*) receptors in EndoC-$\beta$H1 cells, dispersed human islets (Hi) and FACS-purified beta cells from non-diabetic (CTL) and type 1 diabetes (T1D) donors, under basal conditions. The quantification of RNA-sequencing was conducted using Salmon with GENCODE v36 as the genome reference and the expression level is expressed in transcript per million (TPM). Data were obtained from the following sources: EndoC-$\beta$H1 cells, present study; human islets[1]; T1D_Ctl and T1D[2].

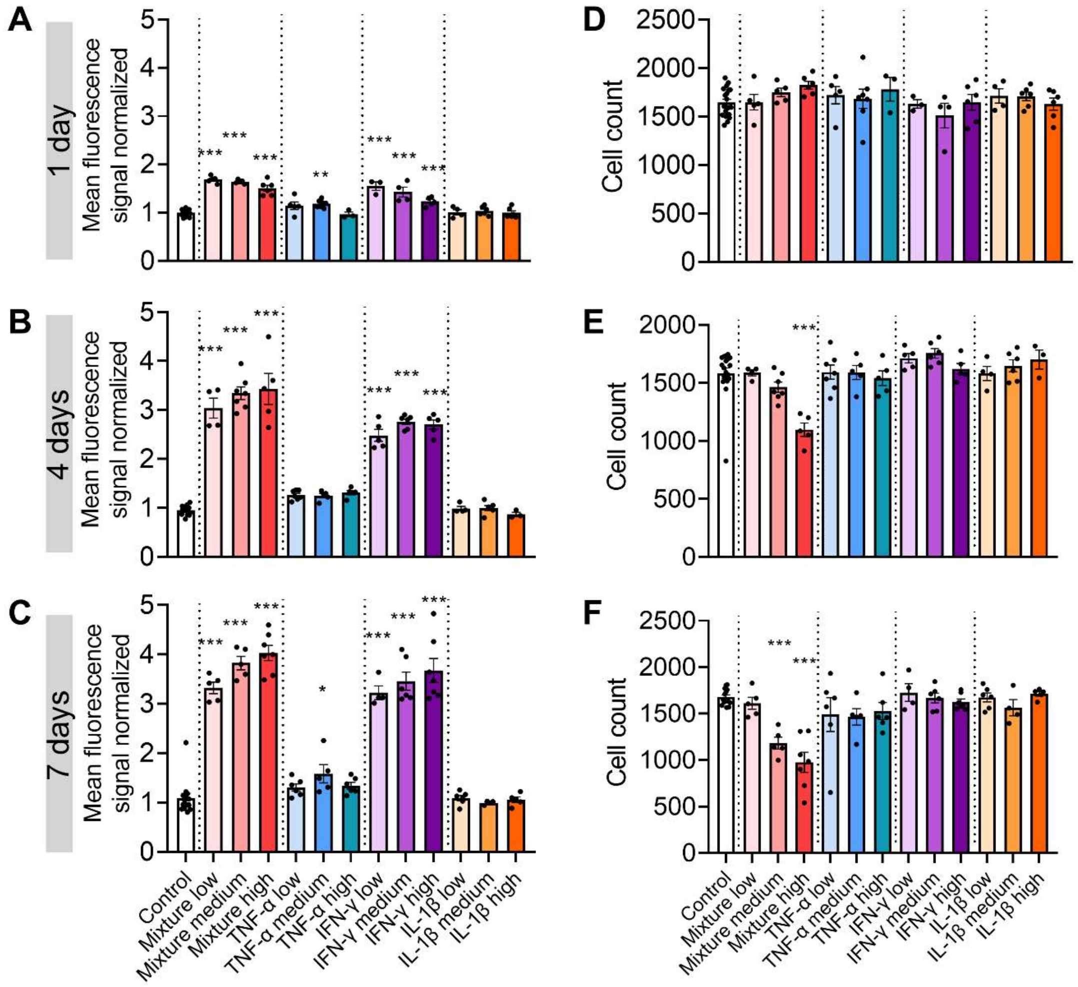


**Supplementary figure 2: IFNγ, but not IL-1β or TNFα, induces HLA class I expression in human islet microtissues.**

Islet microtissues were treated with TNFα (low= 10, medium = 25 and high = 50ng/ml), IFNγ (low= 10, medium = 25 and high = 50ng/ml), IL-1β (low = 2, medium = 5 and high = 10ng/ml), or a cocktail of these cytokines for a duration of 1, 4 or 7 days. (**A-C**) HLA class I expression was analysed by confocal microscopy and the mean HLA signal intensity (Mean fluorescence signal normalized) was calculated for the entire spheroid region. (**D-F**) Cell count was done based on DAPI staining. *$p<0.05$, **$p<0.01$, ***$p<0.001$ vs Control, ANOVA.

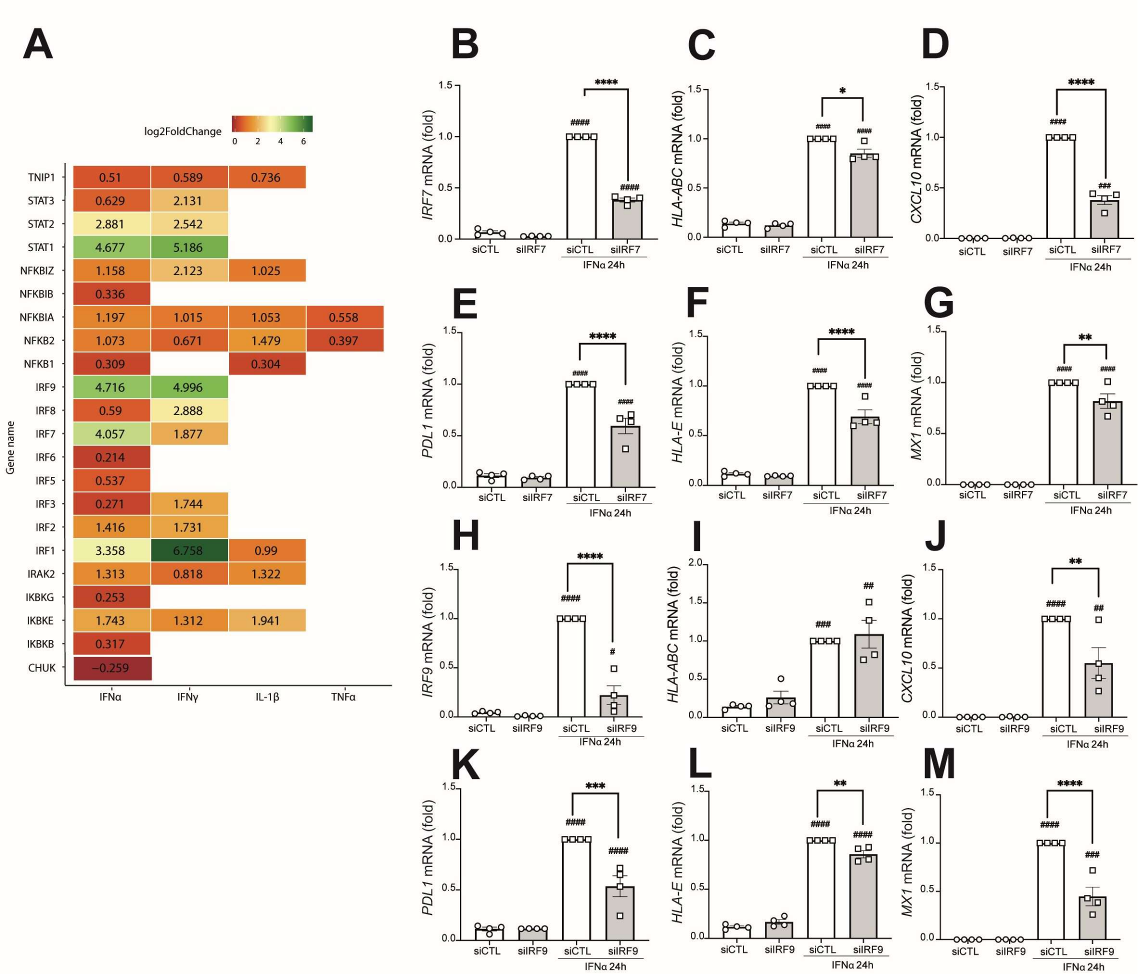


**Supplementary figure 3: The impact of the different pro-inflammatory cytokines on the expression of selected immune-related transcription factors.** (**A**) Heatmap showing the changes at the mRNA expression level (Log$_2$ Fold Change) induced by the four pro-inflammatory cytokines - IFN$\alpha$, IFN$\gamma$, IL1$\beta$, and TNF$\alpha$ - on a panel of immune-related transcription factors. Only the Log2 Fold Change of statistically significant changes are shown (Adjusted P-value < 0.05). (**B-M**) EndoC-$\beta$H1 cells were transfected with siRNA control (siCTL), siRNA against IRF7 (**B-G**) or siRNA against IRF9 (**H-M**) for 48h, and then treated or not with IFN$\alpha$ (2000 U/ml) for 24h. The mRNA expression of *IRF7* (**B**), *HLA-ABC* (**C, I**), *CXCL10* (**D, J**), *MX1* (**E, K**), *PDL1* (**F, L**), *HLA-E* (**G, M**) and *IRF9* (**H**) was measured by RT qPCR and the values were normalized by the geometric mean of *VAPA* and *ACTB* and then by the condition siCTL + IFNα considered as 1. Results are the mean ± SEM of four independent experiments. #$p$<0.05, ##$p$<0.01, ###$p$<0.001 vs siCTL non treated; and *$p$<0.05, **$p$<0.01, ***$p$<0.001 vs siCTL + IFNα as indicated; one-way ANOVA.

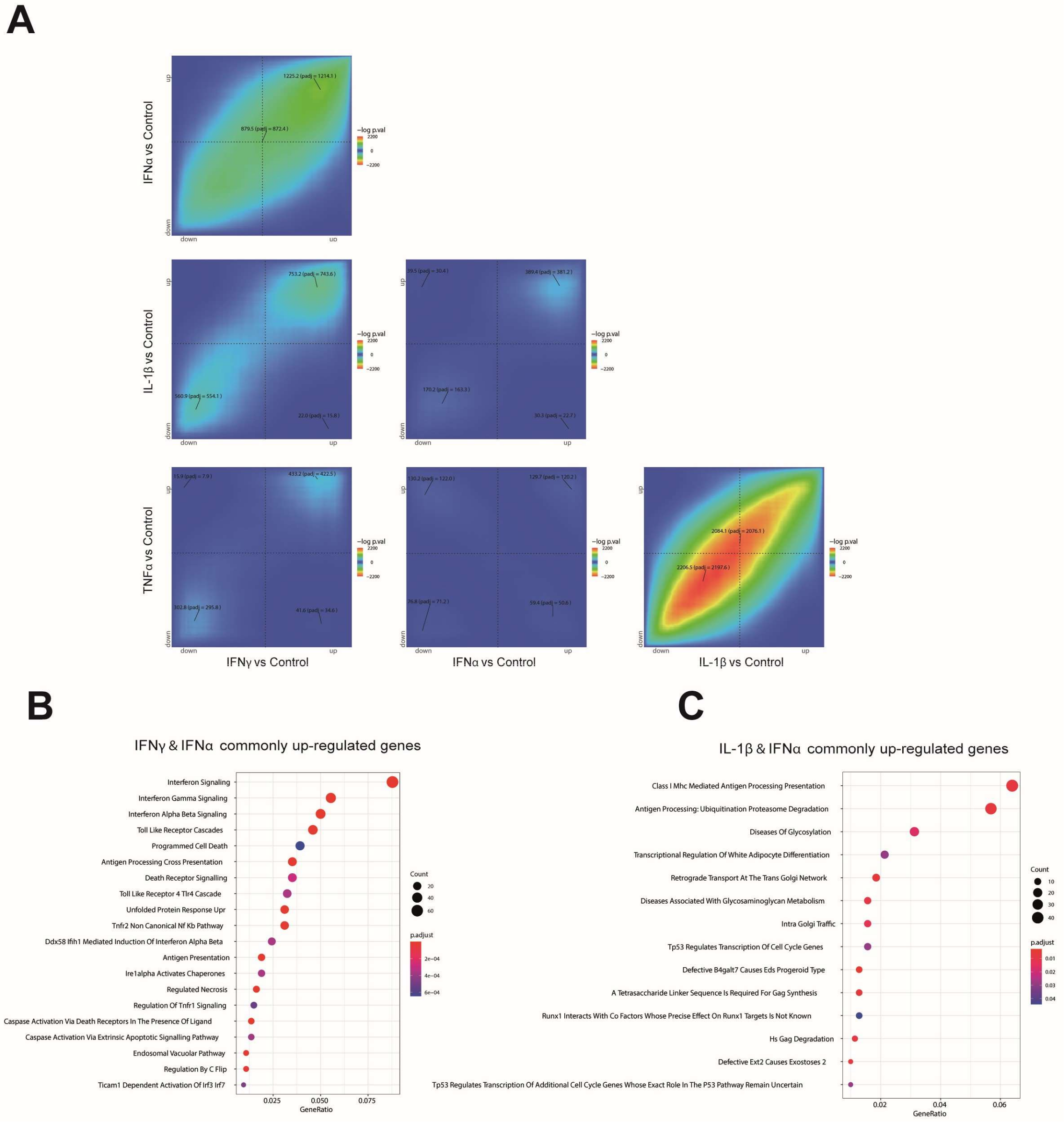


**Supplementary figure 4: Pairwise rank-rank hypergeometric (RRHO) analysis of the gene expression signatures of EndoC-BH1 cells exposed to the proinflammatory cytokines IFNα, IFNγ, IL-1β, and TNFα, and respective pathway enrichment analysis.** (**A**) RRHO algorithm was conducted for each pairwise analysis. The genes were ranked according to their fold change. The significance of overlap between genes up-regulated in both cytokines (top right quadrant), down-regulated in bot (bottom left quadrant), up-regulated by one and down by the other cytokine (top left quadrant), and down-regulated by one cytokine and down-regulated by the other (bottom right quadrant) is represented by the level map. The colors represent the –log(adjusted p-values). For comparison purposes, all the level maps are plotted using the same scale that corresponds to the highest scale number when no scale restriction was applied. Pathway enrichment analysis of the commonly up-regulated genes between IFNα and IFN$\gamma$ (**B**), and IL-1β and IFNα (**C**) performed using gProfiler.

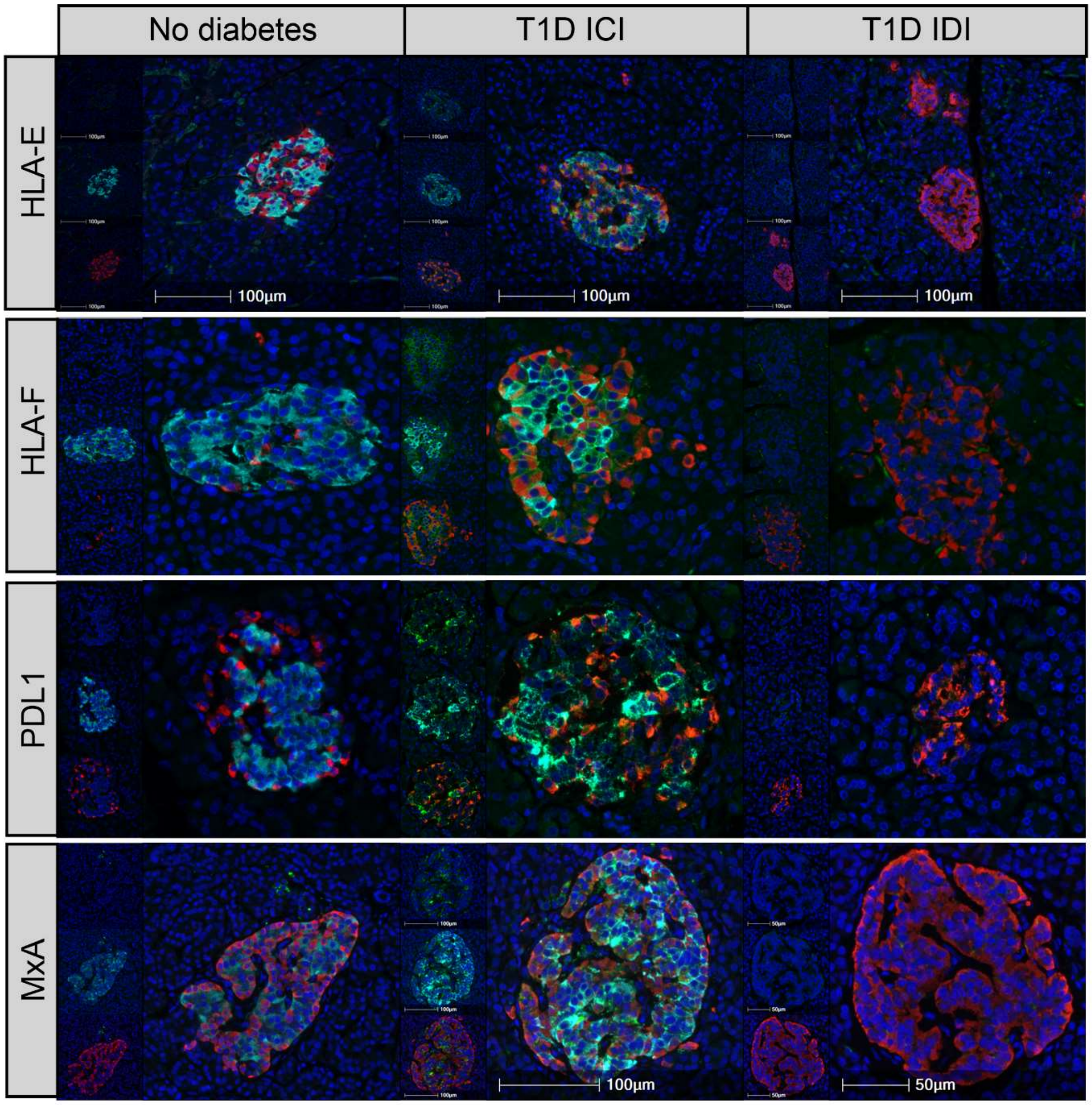


**Supplementary figure 5: HLA-E, HLA-F, PDL1 and MxA are all elevated in insulin-containing islets (ICIs) of donors with T1D, but not in T1D insulin-deficient islets (IDIs) or islets from donors without diabetes.**

Representative immunostaining of HLA-E, HLA-F, PDL1 and MxA in donors without diabetes and in insulin-containing islets (ICI) or insulin-deficient islets (IDI) of donors with type 1 diabetes (T1D). Each panel contains an inset where upper panel is protein of interest (green) and DAPI (dark blue); middle panel is protein of interest (green), insulin (cyan) and DAPI; lower panel contains protein of interest (green), glucagon (red) and DAPI. The larger panels contain the overlay image of protein of interest (green), insulin (cyan), glucagon (red) and DAPI (dark blue).

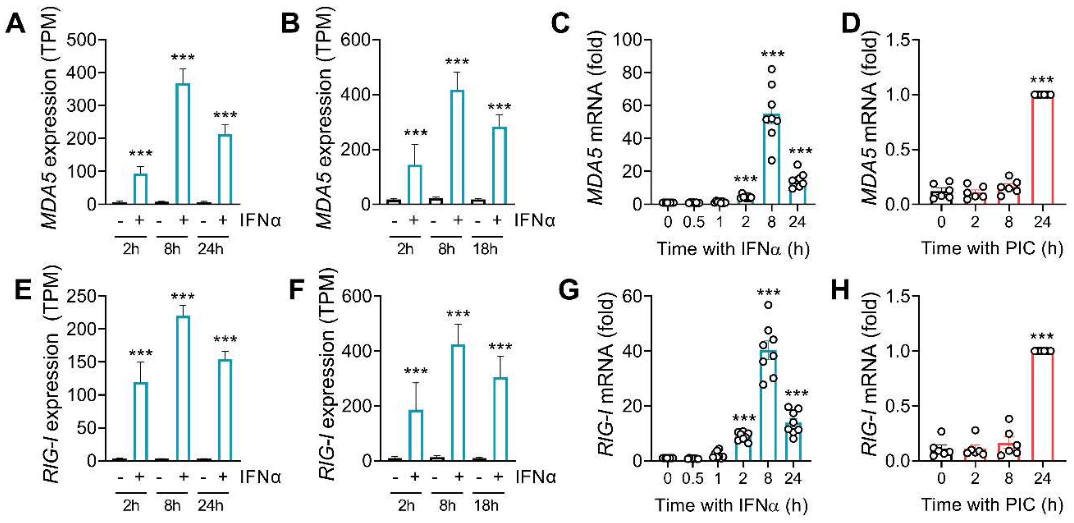


**Supplementary figure 6: Expression of the viral sensors *MDA5* and *RIG-I* follow the same temporal pattern of expression as *ZNFX1* in EndoC-βH1 cells.**

*MDA5* (**A-D**) and *RIG-I* (**E-H**) expression from RNAseq data of EndoC-βH1 cells (N = 5) (**A, E**) and human islets (N = 5) (**B, F**) exposed to IFNα for different time points. EndoC-βH1 cells were either treated with IFNα (2000 U/ml) for different periods of time (**C, G**) or transfected with poly-IC (PIC, 1 µg/ml) for different periods of time as indicated (**D, H**). The mRNA expression of *MDA5* (**C, D**) and *RIG-I* (**G, H**) was analyzed by RT-qPCR and the values were normalized by the geometric mean of *VAPA* and *ACTB*. Results are mean ± SEM of eight (IFNα) or six (PIC) independent experiments. ***$p$<0.001 vs control (time 0); one-way ANOVA.

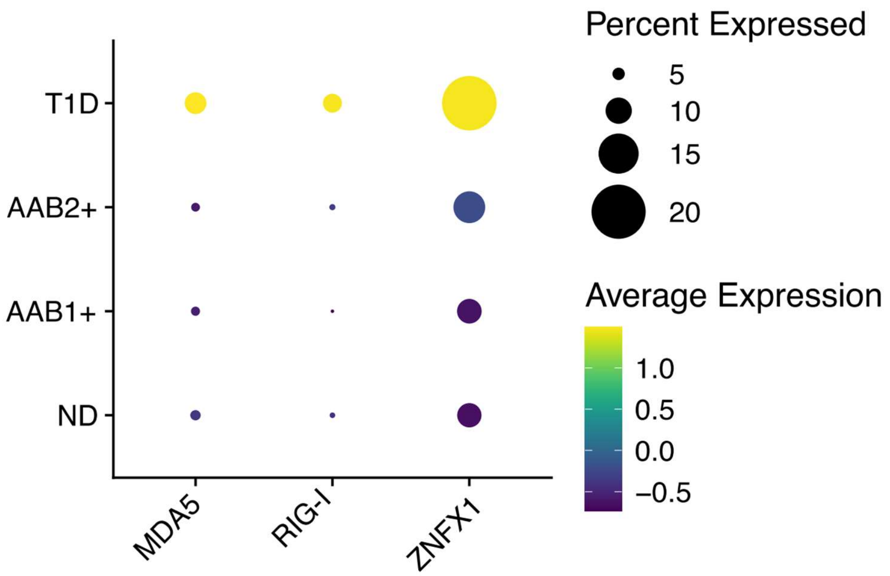


**Supplementary figure 7: Expression of genes involved in viral response in beta cells from non-diabetic, autoantibody-positive and T1D individuals.**

Single-cell RNAseq data of non-diabetic donors (ND, n = 15), one autoantibody-positive donors (AAB1+, n = 8), 2 or more autoantibody-positive donors (AAB2+, n = 2) and T1D donors (n = 9). The size of the circles represents the proportion of cells expressing each gene, while the color scale indicates the normalized average gene expression. Data were downloaded from Human Pancreas Analysis Program (HPAP: https://hpap.pmacs.upenn.edu) data portal and re-analyzed at our own pipeline (see Methods). MDA5: Melanoma Differentiation-Associated Gene 5; RIG-I: Retinoic Acid-Inducible Gene I; ZNFX1: Zinc Finger NFX1-Type Containing 1.

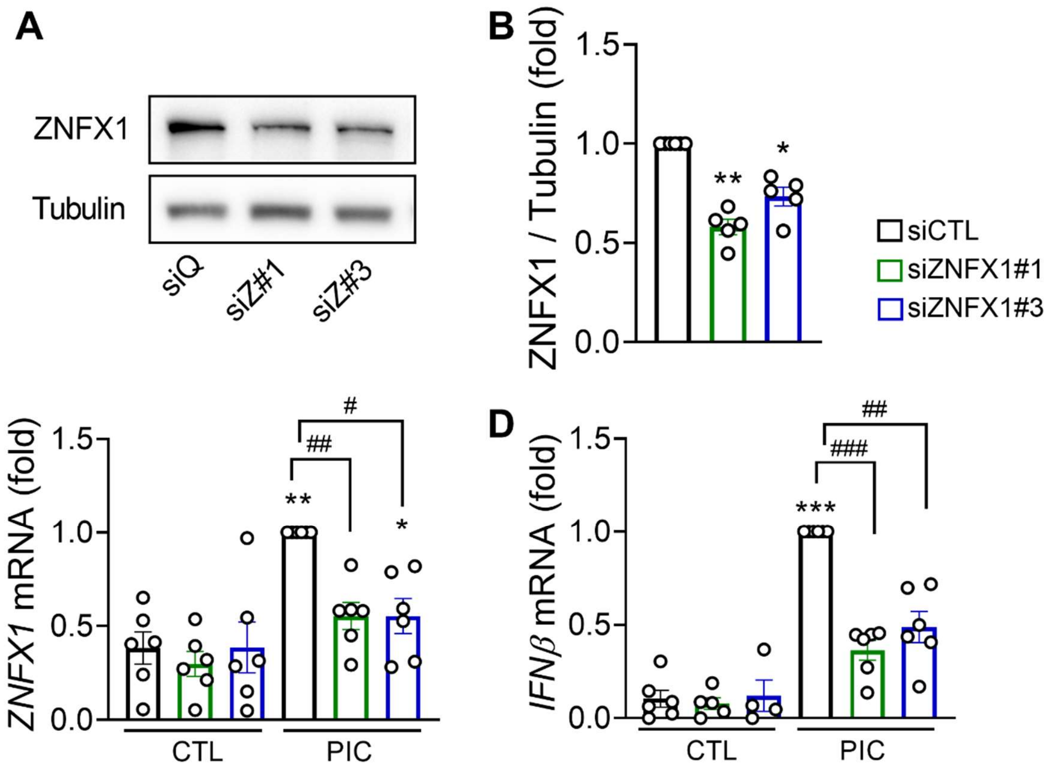


**Supplementary figure 8: *ZNFX1* silencing using two different siRNAs decreases poly-IC-induced IFNβ expression.**

EndoC-βH1 cells were transfected with a siRNA control (siQ: black outline) or with two different siRNAs targeting ZNFX1 (siZNFX1#1: green outline; siZNFX1#3: blue outline) for 72h. (**A, B**) Protein expression was measured by western blot and representative images of five independent experiments are shown. Densitometry results are shown for ZNFX1 (**B**), and the values were normalized by tubulin and then by the value of siCTL considered as 1. (**C, D**) Cells already transfected with siZNFX1#1 or #3 were transfected with poly-IC (PIC, 1 µg/ml) for an additional 24h. The mRNA expression of *ZNFX1* (**C**) and *IFNβ* (**D**) was analysed by RT-qPCR and the values were normalized by the geometric mean of *ACTB* and *VAPA* and then by the value of siCTL + PIC considered as 1. Results are mean ± SEM of six independent experiments. **p*<0.05, ***p*<0.01 and ****p*<0.001 vs siQ CTL, ##*p*<0.05, ##*p*<0.01 and ###*p*<0.001 vs siQ PIC; one-way ANOVA.

**SUPPLEMENTARY TABLES**

**Supplementary table S2:** Human islet donors

| | Age (years) | Gender | BMI (kg/m$^2$) | Cause of death | Beta cell purity (%) |
|---|---|---|---|---|---|
| Donor 1 | 84 | F | 23.7 | Cardiovascular disease | 62% |
| Donor 2 | 63 | M | 21.2 | Cardiovascular disease | 37% |
| Donor 3 | 91 | F | 22.2 | Cardiovascular disease | 65% |
| Donor 4 | 39 | F | 36 | Brain aneurysm | 56% |
| Donor 5 | 34 | | 21.5 | Anoxia | 51% |
| Donor 6 | 66 | M | 25.6 | Brain aneurysm | 56% |
| Donor 7 | 48 | F | 22 | NDD - neurological | 33% |
| Donor 8 | 57 | F | 33.4 | Brain aneurysm | 54% |
| Donor 9 | 69 | F | 26.7 | Respiratory failure | 54% |
| Donor 10 | 86 | F | 27.1 | Cardiovascular disease | 33% |
| Donor 11 | 81 | M | 27.1 | Cardiovascular disease | 11% |
| Donor 12 | 62 | F | 23.5 | Cardiovascular disease | 55% |
| Donor 13 | 36 | M | 29.4 | Cardiovascular disease | 35% |
| Donor 14 | 76 | F | 23.9 | Cardiovascular disease | 37% |
| Donor 15 | 89 | M | 26.1 | Cardiovascular disease | 25% |

Beta cell purity was determined by immunofluorescence for insulin, as described in Methods.

**Supplementary table S3**: List of siRNAs used in the present study.

| siRNA | Sequence (5'-3') | Reference | Supplier |
|---|---|---|---|
| Allstars negative control siRNA | Sequence not provided | 1027281 | Qiagen |
| siZNFX1#1 | CCCAUGCUAUGUGCCUUGUACUAAG | HSS126101 | TFS |
| siZNFX1#3 | AAGAAGCAACCAGCUUGCUUCUGAA | HSS183742 | TFS |
| siBACH2 | GAUAUUCUCUGUGACGUGATT | Silencer Select S34070 | TFS |
| siIRF7 | CAGCGCCAACAGCCTCTATGA | SI03171763 Hs | Qiagen |
| siIRF9 | CCACCGAAGTTCCAGGTAACACTG | HSS115879 | TFS |

Qiagen (Venlo, Netherlands), TFS : Thermo Fisher Scientific (Waltham, MA, USA)

**Supplementary table S4:** List of the primers used for quantitative RT-PCR and alternative splicing analysis in the present study.

| Gene | Sequence (5' to 3') | Primer | Application | Supplier |
|---|---|---|---|---|
| *ACTB* | CTGTACGCCAACACAGTGCT | Forward | RT-qPCR | Eurogentec |
| | GCTCAGGAGGAGCAATGATC | Reverse | | |
| *BACH2* | TGGTGGTCAGCTTGCC | Forward | RT-qPCR | Eurogentec |
| | CGGATGACCTCGCGGATGTT | Reverse | | |
| *BACH1* | CAGAAGAGGTGACAGTTAAAGGA | Forward | RT-qPCR | Eurogentec |
| | AAACTCCACACATTTGCACAC | Reverse | | |
| *CXCL10* | GTGGCATTCAAGGAGTACCTC | Forward | RT-qPCR | Eurogentec |
| | GCCTTCGATTCTGGATTCAG | Reverse | | |
| *HLA-ABC* | CAGGAGACACGGAATGTGAA | Forward | RT-qPCR | Eurogentec |
| | TTATCTGGATGGTGTGAGAACC | Reverse | | |
| *HLA-E* | TGGTTGCTGCTGTGATATGGA | Forward | RT-qPCR | Eurogentec |
| | GCTCCACTCAGCCTTAGAGT | Reverse | | |
| *PDL1* | CCAGTCACCTCTGAACATGAA | Forward | RT-qPCR | Eurogentec |
| | ACTTGATGGTCACTGCTTGT | Reverse | | |
| *VAPA* | TACCGAAACAAGGAAACTAATGGAA | Forward | RT-qPCR | Eurogentec |
| | GCCTTAAACCTTCATCTCTCAGGT | Reverse | | |
| *ZNFX1* | CAGTGCAGGATAGTGAATGG | Forward | RT-qPCR | Eurogentec |
| | CTTCTGCCTGGGCTGTATTC | Reverse | | |
| *CD47* | GCGGCGTGTATACCAATGCATG | Forward | Splicing (PCR) | Eurogentec |
| | CACGTAAGGGTCTCATAGGTGA | Reverse | | |
| *LARP1* | GGGCCTCGGATTTACGGC | Forward | Splicing (PCR) | Eurogentec |
| | GGAAAAACAGCGTGGCCTTGGC | Forward | | |
| | CAAAGTCACCAACCTTGCTGC | Reverse | | |
| *IRF7* | AGGCAGAGCCGTACCTGTC | Forward | RT-qPCR | Eurogentec |
| | GGCCCTTGTACATGATGGTC | Reverse | | |
| *IFNβ* | GTTGAGAACCTCCTGGCTAATG | Forward | RT-qPCR | Eurogentec |
| | GGTAATGCAGAATCCTCCCATAAT | Reverse | | |
| *MX1* | AGACAGGACCATCGGAATCT | Forward | RT-qPCR | Eurogentec |
| | GTAACCCTTCTTCAGGTGGAAC | Reverse | | |
| *MDA5* | GAGGAATCAGCACGAGGAATAA | Forward | RT-qPCR | Eurogentec |
| | TCAGATGGTGGGCTTTGAC | Reverse | | |
| *RIG-I* | AGCACTTGTGGACGCTTTA | Forward | RT-qPCR | Eurogentec |
| | GGTCATTCCTGTGTTCTGATTTG | Reverse | | |
| *IRF9* | CTCTTCAGAACCGCCTACTTC | Forward | RT-qPCR | Eurogentec |
| | GGCTCTCTTCCCAGAAATTCA | Reverse | | |
| *GAPDH* | CAGCCTCAAGATCATCAGCA | Forward | RT-qPCR | Eurogentec |
| | TGTGGTCATGAGTCCTTCCA | Reverse | | |

Eurogentec, Liège, Belgium

**Supplementary Table S5:** List of antibodies used in the present study and conditions of use.

| Antibody | Source | Company | Cat. # | Dilution | Application |
|---|---|---|---|---|---|
| β-actin | Rabbit | CST | 4967 | 1:5000 | WB |
| α-tubulin | Mouse | SA | T5168 | 1:5000 | WB |
| Phospho-STAT1 | Rabbit | CST | 9167 | 1:4000 | WB |
| Phospho-STAT2 | Rabbit | CST | 88410 | 1:4000 | WB |
| ZNFX1 | Rabbit | Abcam | ab179452 | 1:1000 | WB |
| HRP anti-rabbit IgG | Donkey | JIRL | 711-036-152 | 1:5000 | WB |
| HRP anti-mouse IgG | Donkey | JIRL | 715-036-150 | 1/5000 | WB |
| HLA class I ABC | Mouse | Abcam | ab70328 | 1:10000 (O/N) | IF islet microtissues |
| anti-Mouse Alexa Fluor 488 | Goat | JIRL | 115-545-166 | 1:50 | IF islet microtissues |
| HLA-G [4H84] | Mouse | Abcam | ab52455 | 1:100 (O/N) | IF (FFPE) TSA |
| HLA-E [MEM-E/02] | Mouse | Abcam | ab2216 | 1:150 (O/N) | IF (FFPE) TSA |
| HLA-F | Rabbit | Abcam | ab126624 | 1:400 (O/N) | IF (FFPE) |
| STAT1 | Rabbit | Abcam | ab2415 | 1:100 (1h) | IF (FFPE) |
| PDL1 | Rabbit | Abcam | ab205921 | 1:100 (1h) | IF (FFPE) TSA |
| Insulin | Guinea-pig | Agilent | IR000261-2 | 1:4 (1h) | IF (FFPE) |
| Glucagon | Rabbit | Abcam | ab92517 | 1:4000 (1h) | IF (FFPE) |
| Glucagon | Mouse | Abcam | ab10988 | 1:2000 (1h) | IF (FFPE) |
| anti-mouse Alexa Fluor™ 488 | Goat | Invitrogen | A11001 | 1:400 | IF (FFPE) |
| anti-rabbit Alexa Fluor™ 488 | Goat | Invitrogen | A11008 | 1:400 | IF (FFPE) |
| anti-rabbit Alexa Fluor™ 555 | Goat | Invitrogen | A21428 | 1:400 | IF (FFPE) |
| anti-mouse Alexa Fluor™ 555 | Goat | Invitrogen | A31572 | 1:400 | IF (FFPE) |
| anti-guinea pig Alexa Fluor™ 647 | Goat | Invitrogen | A21450 | 1:400 | IF (FFPE) |

| | | | | | |
|---|---|---|---|---|---|
| anti-mouse/ rabbit Alexa Fluor™ 488 (Tyramide SuperBoost) | Goat | Invitrogen | B40941 /B40943 | 1:100 | IF (FFPE) |
| Anti-goat Alexa Fluor™ 488 | Donkey | Invitrogen | A11055 | 1:400 | IF (FFPE) |

HRP: horseradish peroxidase; O/N: overnight; FFPE: formalin-fixed paraffin-embedded; CST: Cell Signaling Technologies (Danvers, MA, USA); SA: Sigma-Aldrich (Bornem, Belgium); JIRL: Jackson ImmunoResearch Laboratories (Wes Grove, PA, USA); Abcam (Cambridge, UK).
