## Supplementary material for "Interferons are the key cytokines acting on pancreatic islets in type 1 diabetes": Supplmentary table S1

**DEG_significative_IFNα vs NT 24h**

| Ensembl_gene_ID | gene_name | gene_type | baseMean | log2FoldChange | lfcSE | stat | pvalue | padj |
|---|---|---|---|---|---|---|---|---|
| ENSG00000000457.14 | SCYL3 | protein_coding | 2458,510098 | 0,263607536 | 0,057905419 | 4,552381126 | 5,30E-06 | 3,10E-05 |
| ENSG00000000460.17 | C1orf112 | protein_coding | 3102,625333 | -0,156197769 | 0,057054303 | -2,737703571 | 0,006186982 | 0,019697704 |
| ENSG00000001036.13 | FUCA2 | protein_coding | 22048,40905 | 0,346565417 | 0,044234492 | 7,834732688 | 4,70E-15 | 7,10E-14 |
| ENSG00000001167.14 | NFYA | protein_coding | 9110,828192 | -0,22325705 | 0,046018604 | -4,851452057 | 1,23E-06 | 7,87E-06 |
| ENSG00000001460.18 | STPG1 | protein_coding | 1253,401792 | 0,231666011 | 0,076604129 | 3,024197445 | 0,002492937 | 0,008781458 |
| ENSG00000001629.10 | ANKIB1 | protein_coding | 15280,38923 | 0,784675395 | 0,048342124 | 16,23171123 | 3,01E-59 | 2,10E-57 |
| ENSG00000002330.13 | BAD | protein_coding | 2363,516328 | -0,21281065 | 0,071601528 | -2,972152366 | 0,002957199 | 0,010221722 |
| ENSG00000002549.12 | LAP3 | protein_coding | 32485,29271 | 3,527801519 | 0,051773392 | 68,1392776 | 0 | 0 |
| ENSG00000002586.20 | CD99 | protein_coding | 19157,74485 | 0,493906782 | 0,04643645 | 10,63618739 | 2,02E-26 | 5,57E-25 |
| ENSG00000002726.20 | AOC1 | protein_coding | 497,6934256 | 0,370748648 | 0,107828778 | 3,438308908 | 0,00058536 | 0,002391697 |
| ENSG00000002822.15 | MAD1L1 | protein_coding | 623,7861169 | -0,654987484 | 0,133944278 | -4,889999748 | 1,01E-06 | 6,54E-06 |
| ENSG00000002933.9 | TMEM176A | protein_coding | 291,719529 | 0,456335834 | 0,152328368 | 2,995737688 | 0,002737818 | 0,009549908 |
| ENSG00000003056.8 | M6PR | protein_coding | 16082,9607 | 0,182097111 | 0,041754117 | 4,361177381 | 1,29E-05 | 7,13E-05 |
| ENSG00000003147.19 | ICA1 | protein_coding | 2349,91117 | 0,180566858 | 0,06330894 | 2,852154203 | 0,004342403 | 0,014401525 |
| ENSG00000003249.13 | DBNDD1 | protein_coding | 1798,162007 | -0,178670412 | 0,067340948 | -2,653220934 | 0,007972767 | 0,024603648 |
| ENSG00000003393.15 | ALS2 | protein_coding | 10922,12592 | 0,594560049 | 0,04551708 | 13,06235048 | 5,40E-39 | 2,35E-37 |
| ENSG00000003402.20 | CFLAR | protein_coding | 6991,716935 | 0,690001362 | 0,064888119 | 10,63370877 | 2,08E-26 | 5,71E-25 |
| ENSG00000004059.11 | ARF5 | protein_coding | 18848,77838 | -0,226722385 | 0,046572439 | -4,868166473 | 1,13E-06 | 7,27E-06 |
| ENSG00000004399.12 | PLXND1 | protein_coding | 1693,709733 | -0,789001426 | 0,071048507 | -11,1051092 | 1,18E-28 | 3,55E-27 |
| ENSG00000004660.14 | CAMKK1 | protein_coding | 2998,964996 | -0,262011256 | 0,067646109 | -3,873264235 | 0,000107387 | 0,000507918 |
| ENSG00000004700.16 | RECQL | protein_coding | 7934,79793 | -0,163391068 | 0,050241398 | -3,25212026 | 0,001145475 | 0,004378249 |
| ENSG00000004776.13 | HSPB6 | protein_coding | 64,61497189 | -1,468797356 | 0,309839343 | -4,740512748 | 2,13E-06 | 1,32E-05 |
| ENSG00000004777.18 | ARHGAP33 | protein_coding | 2612,893308 | -0,670752087 | 0,077621902 | -8,641273472 | 5,56E-18 | 1,01E-16 |
| ENSG00000004779.10 | NDUFAB1 | protein_coding | 7298,631958 | -0,330156128 | 0,055895659 | -5,906650595 | 3,49E-09 | 3,07E-08 |
| ENSG00000004838.14 | ZMYND10 | protein_coding | 99,85452127 | -1,577512411 | 0,263837198 | -5,979112976 | 2,24E-09 | 2,01E-08 |
| ENSG00000004864.13 | SLC25A13 | protein_coding | 14901,20029 | -0,245604063 | 0,052360339 | -4,690650748 | 2,72E-06 | 1,66E-05 |
| ENSG00000004866.20 | ST7 | protein_coding | 3353,6643 | 0,212633199 | 0,054412248 | 3,907818665 | 9,31E-05 | 0,000445839 |
| ENSG00000004961.15 | HCCS | protein_coding | 5628,149946 | 0,192398506 | 0,054244923 | 3,546848179 | 0,000389869 | 0,001653895 |
| ENSG00000004975.12 | DVL2 | protein_coding | 5817,576573 | -0,150431835 | 0,055130944 | -2,728627929 | 0,006359842 | 0,020173855 |
| ENSG00000005001.10 | PRSS22 | protein_coding | 602,3459143 | 0,322987056 | 0,119965695 | 2,692328477 | 0,007095502 | 0,02222553 |
| ENSG00000005007.12 | UPF1 | protein_coding | 18371,67916 | 0,190153717 | 0,045365611 | 4,191582804 | 2,77E-05 | 0,000144403 |
| ENSG00000005020.13 | SKAP2 | protein_coding | 4130,265416 | -0,280856894 | 0,054606166 | -5,143318334 | 2,70E-07 | 1,89E-06 |
| ENSG00000005022.6 | SLC25A5 | protein_coding | 32243,49341 | -0,423581857 | 0,048445321 | -8,743504033 | 2,26E-18 | 4,22E-17 |
| ENSG00000005059.16 | MCUB | protein_coding | 2283,941416 | -0,248459094 | 0,070750185 | -3,511780138 | 0,000445116 | 0,001869587 |

| | | | | | | | | |
|---|---|---|---|---|---|---|---|---|
| ENSG00000005075.15 | POLR2J | protein_coding | 5194,911214 | -0,371367359 | 0,05171596 | -7,180904289 | 6,93E-13 | 8,88E-12 |
| ENSG00000005108.16 | THSD7A | protein_coding | 331,7742786 | 1,439312159 | 0,139642322 | 10,30713425 | 6,54E-25 | 1,69E-23 |
| ENSG00000005187.12 | ACSM3 | protein_coding | 1706,789164 | -0,286159211 | 0,088338895 | -3,239334285 | 0,001198091 | 0,004556915 |
| ENSG00000005189.19 | REXO5 | protein_coding | 2800,954307 | 0,192095357 | 0,072031372 | 2,666829069 | 0,00765706 | 0,023746593 |
| ENSG00000005249.13 | PRKAR2B | protein_coding | 3445,093824 | -0,537637173 | 0,061187424 | -8,786726703 | 1,54E-18 | 2,90E-17 |
| ENSG00000005339.14 | CREBBP | protein_coding | 12237,14735 | 0,402227047 | 0,05781133 | 6,957581671 | 3,46E-12 | 4,16E-11 |
| ENSG00000005379.17 | TSPOAP1 | protein_coding | 1783,823609 | -0,280549841 | 0,107584175 | -2,607724061 | 0,00911464 | 0,027657242 |
| ENSG00000005448.16 | WDR54 | protein_coding | 2307,447318 | -0,217514765 | 0,077103085 | -2,821090291 | 0,004786073 | 0,015690317 |
| ENSG00000005469.11 | CROT | protein_coding | 5626,474062 | 0,275285051 | 0,049016628 | 5,61615639 | 1,95E-08 | 1,58E-07 |
| ENSG00000005471.17 | ABCB4 | protein_coding | 2370,425625 | -0,250073484 | 0,064676673 | -3,866517459 | 0,000110401 | 0,000521075 |
| ENSG00000005483.21 | KMT2E | protein_coding | 11570,84929 | 0,308455378 | 0,050312877 | 6,130744205 | 8,75E-10 | 8,24E-09 |
| ENSG00000005513.10 | SOX8 | protein_coding | 110,4549884 | -1,731669744 | 0,282079819 | -6,13893525 | 8,31E-10 | 7,84E-09 |
| ENSG00000005700.15 | IBTK | protein_coding | 14681,60302 | 0,395245356 | 0,042482909 | 9,303632016 | 1,36E-20 | 2,87E-19 |
| ENSG00000005801.18 | ZNF195 | protein_coding | 5107,501182 | 0,2898992 | 0,052483367 | 5,523639518 | 3,32E-08 | 2,61E-07 |
| ENSG00000005810.18 | MYCBP2 | protein_coding | 17964,17758 | 0,449179303 | 0,061074351 | 7,354630761 | 1,91E-13 | 2,56E-12 |
| ENSG00000005882.11 | PDK2 | protein_coding | 1549,228608 | -0,24287612 | 0,084558421 | -2,872287802 | 0,004075117 | 0,01362549 |
| ENSG00000005884.18 | ITGA3 | protein_coding | 3069,391346 | 1,23938237 | 0,081908075 | 15,13138198 | 1,01E-51 | 6,04E-50 |
| ENSG00000005889.15 | ZFX | protein_coding | 8578,251005 | 0,258452621 | 0,045030471 | 5,739505171 | 9,50E-09 | 7,91E-08 |
| ENSG00000005893.15 | LAMP2 | protein_coding | 18133,20308 | 0,329145951 | 0,040963985 | 8,035008095 | 9,36E-16 | 1,48E-14 |
| ENSG00000005961.18 | ITGA2B | protein_coding | 89,4758523 | -0,995760958 | 0,340493151 | -2,924466924 | 0,003450467 | 0,011758754 |
| ENSG00000006007.12 | GDE1 | protein_coding | 16004,04151 | 0,196009302 | 0,04774794 | 4,105083982 | 4,04E-05 | 0,000205638 |
| ENSG00000006015.17 | REX1BD | protein_coding | 667,5429689 | -0,340004808 | 0,140150793 | -2,425992761 | 0,015266575 | 0,043062072 |
| ENSG00000006042.12 | TMEM98 | protein_coding | 4730,619504 | -0,283806481 | 0,052695219 | -5,385810781 | 7,21E-08 | 5,40E-07 |
| ENSG00000006062.17 | MAP3K14 | protein_coding | 942,1575328 | 0,367795911 | 0,104880542 | 3,506807894 | 0,000453516 | 0,001902031 |
| ENSG00000006071.14 | ABCC8 | protein_coding | 18931,39833 | -0,239299829 | 0,048921008 | -4,891555591 | 1,00E-06 | 6,49E-06 |
| ENSG00000006118.14 | TMEM132A | protein_coding | 15024,46708 | 0,206260971 | 0,047133841 | 4,376069632 | 1,21E-05 | 6,69E-05 |
| ENSG00000006125.18 | AP2B1 | protein_coding | 27601,2487 | -0,207774381 | 0,043904449 | -4,73242203 | 2,22E-06 | 1,37E-05 |
| ENSG00000006194.10 | ZNF263 | protein_coding | 7095,087608 | 0,222046064 | 0,058422669 | 3,800683315 | 0,000144298 | 0,000666939 |
| ENSG00000006210.7 | CX3CL1 | protein_coding | 2276,10149 | 1,198290414 | 0,083935489 | 14,27632615 | 3,07E-46 | 1,62E-44 |
| ENSG00000006283.18 | CACNA1G | protein_coding | 623,584856 | 0,309283914 | 0,121103983 | 2,553870692 | 0,010653281 | 0,031593505 |
| ENSG00000006327.13 | TNFRSF12A | protein_coding | 1373,006401 | 0,357999265 | 0,101444668 | 3,529010179 | 0,000417117 | 0,001759207 |
| ENSG00000006377.11 | DLX6 | protein_coding | 752,5475347 | -0,274721661 | 0,108153399 | -2,540111208 | 0,011081723 | 0,032674678 |
| ENSG00000006432.15 | MAP3K9 | protein_coding | 6239,48421 | -0,137582698 | 0,047749679 | -2,881332399 | 0,003959978 | 0,013286777 |
| ENSG00000006451.8 | RALA | protein_coding | 11087,72453 | -0,125753602 | 0,046211755 | -2,721247049 | 0,006503614 | 0,020583498 |
| ENSG00000006453.14 | BAIAP2L1 | protein_coding | 7652,524074 | 0,209152906 | 0,04746729 | 4,406253317 | 1,05E-05 | 5,87E-05 |
| ENSG00000006459.11 | KDM7A | protein_coding | 9490,353788 | 0,125355366 | 0,049492345 | 2,532823362 | 0,011314795 | 0,033253 |

| | | | | | | | | |
|---|---|---|---|---|---|---|---|---|
| ENSG00000006468.14 | ETV1 | protein_coding | 6373,900262 | -0,218532363 | 0,054146223 | -4,03596691 | 5,44E-05 | 0,000270974 |
| ENSG00000006625.18 | GGCT | protein_coding | 6561,04134 | -0,431021689 | 0,051962321 | -8,294889077 | 1,09E-16 | 1,82E-15 |
| ENSG00000006634.8 | DBF4 | protein_coding | 5143,818645 | -0,250654248 | 0,048911497 | -5,124648893 | 2,98E-07 | 2,08E-06 |
| ENSG00000006652.14 | IFRD1 | protein_coding | 5488,473501 | 0,351802567 | 0,054813073 | 6,418223741 | 1,38E-10 | 1,42E-09 |
| ENSG00000006695.12 | COX10 | protein_coding | 2621,155421 | 0,26229732 | 0,068536128 | 3,827139447 | 0,000129641 | 0,000605129 |
| ENSG00000006712.15 | PAF1 | protein_coding | 6824,059952 | -0,132399773 | 0,05309924 | -2,493440065 | 0,012651194 | 0,036635241 |
| ENSG00000006744.19 | ELAC2 | protein_coding | 5852,607331 | 0,236185533 | 0,049623455 | 4,7595544 | 1,94E-06 | 1,21E-05 |
| ENSG00000006747.15 | SCIN | protein_coding | 6107,886726 | -0,156455096 | 0,054417415 | -2,875092418 | 0,004039093 | 0,013519827 |
| ENSG00000006756.16 | ARSD | protein_coding | 4286,735602 | 0,160901904 | 0,062169433 | 2,588119234 | 0,009650157 | 0,029031727 |
| ENSG00000006757.12 | PNPLA4 | protein_coding | 3521,029248 | -0,43568902 | 0,060348592 | -7,219539089 | 5,22E-13 | 6,74E-12 |
| ENSG00000007080.11 | CCDC124 | protein_coding | 2299,428406 | -0,307460488 | 0,062052464 | -4,954847398 | 7,24E-07 | 4,80E-06 |
| ENSG00000007168.13 | PAFAH1B1 | protein_coding | 33022,69393 | -0,108333802 | 0,039492132 | -2,743174338 | 0,006084837 | 0,019402691 |
| ENSG00000007202.15 | KIAA0100 | protein_coding | 21628,55874 | 0,108439443 | 0,042637978 | 2,543259505 | 0,010982363 | 0,032428441 |
| ENSG00000007216.14 | SLC13A2 | protein_coding | 41,37385197 | -1,276333072 | 0,388022863 | -3,2893244 | 0,001004282 | 0,003890102 |
| ENSG00000007255.10 | TRAPPC6A | protein_coding | 426,0202107 | -0,391036121 | 0,13221607 | -2,957553659 | 0,003100908 | 0,010669265 |
| ENSG00000007341.19 | ST7L | protein_coding | 2697,225193 | 0,189931236 | 0,075145415 | 2,527515974 | 0,01148726 | 0,033645638 |
| ENSG00000007392.16 | LUC7L | protein_coding | 8780,814744 | -0,12062137 | 0,043962533 | -2,743731108 | 0,006074527 | 0,019372561 |
| ENSG00000007402.11 | CACNA2D2 | protein_coding | 8753,153563 | -0,186933601 | 0,045230136 | -4,132943595 | 3,58E-05 | 0,000183467 |
| ENSG00000007866.21 | TEAD3 | protein_coding | 626,002172 | 0,376514107 | 0,104185502 | 3,613881989 | 0,000301646 | 0,0013095 |
| ENSG00000007944.15 | MYLIP | protein_coding | 1832,410367 | 0,583889101 | 0,069765979 | 8,369252601 | 5,80E-17 | 9,87E-16 |
| ENSG00000008018.9 | PSMB1 | protein_coding | 15359,88939 | -0,161866083 | 0,043124708 | -3,753441821 | 0,000174423 | 0,00079171 |
| ENSG00000008056.14 | SYN1 | protein_coding | 7484,20627 | 0,112335254 | 0,044056698 | 2,549788338 | 0,010778834 | 0,031911189 |
| ENSG00000008086.11 | CDKL5 | protein_coding | 5361,177637 | 0,476764583 | 0,067308788 | 7,083244201 | 1,41E-12 | 1,76E-11 |
| ENSG00000008282.8 | SYPL1 | protein_coding | 9869,55817 | 0,238864658 | 0,046307924 | 5,158181056 | 2,49E-07 | 1,76E-06 |
| ENSG00000008300.17 | CELSR3 | protein_coding | 16650,79085 | 0,389943653 | 0,051343148 | 7,594852892 | 3,08E-14 | 4,38E-13 |
| ENSG00000008311.15 | AASS | protein_coding | 1883,539399 | 0,423075685 | 0,084506007 | 5,006456923 | 5,54E-07 | 3,74E-06 |
| ENSG00000008324.11 | SS18L2 | protein_coding | 2597,263546 | 0,206571628 | 0,067243814 | 3,071979661 | 0,002126442 | 0,007628959 |
| ENSG00000008382.15 | MPND | protein_coding | 1260,695046 | -0,474500407 | 0,079238842 | -5,988229967 | 2,12E-09 | 1,91E-08 |
| ENSG00000008441.16 | NFIX | protein_coding | 808,7788379 | 0,855628669 | 0,107775777 | 7,938970052 | 2,04E-15 | 3,15E-14 |
| ENSG00000008517.16 | IL32 | protein_coding | 20,70780033 | 4,277983675 | 0,743879103 | 5,750912555 | 8,88E-09 | 7,42E-08 |
| ENSG00000008735.14 | MAPK8IP2 | protein_coding | 7259,306849 | 0,59638693 | 0,054872552 | 10,86858388 | 1,63E-27 | 4,66E-26 |
| ENSG00000008869.12 | HEATR5B | protein_coding | 6863,314496 | -0,171756441 | 0,058612319 | -2,930381238 | 0,003385464 | 0,011558229 |
| ENSG00000008988.9 | RPS20 | protein_coding | 35485,28926 | -0,587438482 | 0,042153287 | -13,93576922 | 3,84E-44 | 1,93E-42 |
| ENSG00000009307.16 | CSDE1 | protein_coding | 70218,33616 | -0,361182317 | 0,046953142 | -7,692399385 | 1,44E-14 | 2,10E-13 |
| ENSG00000009413.15 | REV3L | protein_coding | 14116,57357 | 0,29765355 | 0,056253782 | 5,291262884 | 1,21E-07 | 8,89E-07 |
| ENSG00000009780.15 | FAM76A | protein_coding | 1560,920848 | 0,829239596 | 0,082249712 | 10,08197568 | 6,64E-24 | 1,64E-22 |

| | | | | | | | | |
|---|---|---|---|---|---|---|---|---|
| ENSG00000009954.11 | BAZ1B | protein_coding | 30724,97029 | 0,12223299 | 0,038637919 | 3,163550041 | 0,001558576 | 0,005791972 |
| ENSG00000010030.14 | ETV7 | protein_coding | 3283,104517 | 7,503336996 | 0,294297563 | 25,49574971 | 2,20E-143 | 3,48E-141 |
| ENSG00000010256.11 | UQCRC1 | protein_coding | 11871,9564 | -0,134617994 | 0,043716583 | -3,079334761 | 0,002074634 | 0,007463312 |
| ENSG00000010292.13 | NCAPD2 | protein_coding | 14342,8389 | -0,686130727 | 0,050869767 | -13,48798639 | 1,84E-41 | 8,56E-40 |
| ENSG00000010310.9 | GIPR | protein_coding | 6413,85747 | -0,522161741 | 0,065970057 | -7,915132417 | 2,47E-15 | 3,80E-14 |
| ENSG00000010322.16 | NISCH | protein_coding | 11473,71377 | 0,138059544 | 0,048493222 | 2,846986431 | 0,004413524 | 0,01459868 |
| ENSG00000010361.13 | FUZ | protein_coding | 2895,637278 | -0,240116312 | 0,06191424 | -3,878208198 | 0,000105229 | 0,000498651 |
| ENSG00000010404.18 | IDS | protein_coding | 22894,14562 | -0,621451821 | 0,050022036 | -12,42356109 | 1,95E-35 | 7,62E-34 |
| ENSG00000010704.18 | HFE | protein_coding | 426,8478716 | 1,653763198 | 0,150421848 | 10,99416887 | 4,08E-28 | 1,19E-26 |
| ENSG00000010803.16 | SCMH1 | protein_coding | 4504,590852 | 0,265558864 | 0,059882497 | 4,434665808 | 9,22E-06 | 5,20E-05 |
| ENSG00000010818.10 | HIVEP2 | protein_coding | 1780,424356 | 1,268776711 | 0,08643341 | 14,67923932 | 8,76E-49 | 4,82E-47 |
| ENSG00000011007.12 | ELOA | protein_coding | 11084,89321 | 0,21617731 | 0,045082794 | 4,79511782 | 1,63E-06 | 1,03E-05 |
| ENSG00000011009.11 | LYPLA2 | protein_coding | 6350,894836 | -0,13460129 | 0,047120838 | -2,856513076 | 0,004283224 | 0,014240928 |
| ENSG00000011021.23 | CLCN6 | protein_coding | 3115,884704 | 0,213932533 | 0,061066336 | 3,503280954 | 0,000459564 | 0,001923812 |
| ENSG00000011105.14 | TSPAN9 | protein_coding | 2990,691797 | 0,177486133 | 0,072760161 | 2,439331218 | 0,014714475 | 0,041734893 |
| ENSG00000011201.12 | ANOS1 | protein_coding | 458,7518853 | 0,320523963 | 0,129424776 | 2,476527088 | 0,013266756 | 0,03818688 |
| ENSG00000011243.18 | AKAP8L | protein_coding | 9031,246486 | 0,148855335 | 0,052400129 | 2,840743666 | 0,004500847 | 0,014859134 |
| ENSG00000011258.16 | MBTD1 | protein_coding | 7129,590108 | -0,334175156 | 0,054725122 | -6,106430483 | 1,02E-09 | 9,51E-09 |
| ENSG00000011275.18 | RNF216 | protein_coding | 3714,516764 | 0,315687037 | 0,064238831 | 4,914271195 | 8,91E-07 | 5,83E-06 |
| ENSG00000011295.16 | TTC19 | protein_coding | 7092,544065 | -0,190534913 | 0,047754399 | -3,989892374 | 6,61E-05 | 0,00032466 |
| ENSG00000011304.20 | PTBP1 | protein_coding | 30004,48034 | -0,133814224 | 0,039832827 | -3,35939558 | 0,000781132 | 0,00310261 |
| ENSG00000011347.9 | SYT7 | protein_coding | 11848,51407 | -0,279368024 | 0,054243602 | -5,150248405 | 2,60E-07 | 1,83E-06 |
| ENSG00000011376.12 | LARS2 | protein_coding | 7763,469752 | 0,235877641 | 0,049916966 | 4,72540019 | 2,30E-06 | 1,42E-05 |
| ENSG00000011426.11 | ANLN | protein_coding | 13295,77803 | -0,620587066 | 0,047460006 | -13,07600068 | 4,52E-39 | 1,98E-37 |
| ENSG00000011454.17 | RABGAP1 | protein_coding | 10738,7483 | 0,167831644 | 0,047059446 | 3,566375234 | 0,000361953 | 0,001546545 |
| ENSG00000011485.14 | PPP5C | protein_coding | 11304,95257 | -0,19003025 | 0,04301247 | -4,418026859 | 9,96E-06 | 5,59E-05 |
| ENSG00000011523.14 | CEP68 | protein_coding | 3643,444388 | -0,193446423 | 0,062426836 | -3,098770237 | 0,001943257 | 0,007043601 |
| ENSG00000011638.10 | TMEM159 | protein_coding | 1215,2675 | 0,605847272 | 0,089209479 | 6,791288088 | 1,11E-11 | 1,27E-10 |
| ENSG00000011677.13 | GABRA3 | protein_coding | 1520,018685 | -0,392299948 | 0,075271443 | -5,21180317 | 1,87E-07 | 1,34E-06 |
| ENSG00000012048.22 | BRCA1 | protein_coding | 8590,844884 | -0,167099229 | 0,057635724 | -2,899230166 | 0,003740802 | 0,012622052 |
| ENSG00000012211.13 | PRICKLE3 | protein_coding | 674,5128932 | 0,753367967 | 0,11439165 | 6,585865002 | 4,52E-11 | 4,87E-10 |
| ENSG00000012232.8 | EXTL3 | protein_coding | 5392,567098 | 0,581196039 | 0,068327373 | 8,506049794 | 1,80E-17 | 3,16E-16 |
| ENSG00000012504.15 | NR1H4 | protein_coding | 569,1273363 | 0,934404079 | 0,130612507 | 7,154016857 | 8,43E-13 | 1,07E-11 |
| ENSG00000012660.14 | ELOVL5 | protein_coding | 34142,51283 | -0,142392273 | 0,04284962 | -3,323069669 | 0,000890327 | 0,003490137 |
| ENSG00000012822.16 | CALCOCO1 | protein_coding | 3772,484201 | -0,169807111 | 0,061146088 | -2,777072382 | 0,005485096 | 0,017711093 |
| ENSG00000012963.15 | UBR7 | protein_coding | 13492,72026 | -0,403821125 | 0,057258219 | -7,052631632 | 1,76E-12 | 2,17E-11 |

| | | | | | | | | |
|---|---|---|---|---|---|---|---|---|
| ENSG00000013016.16 | EHD3 | protein_coding | 702,9092227 | 0,429752549 | 0,0985707 | 4,359840668 | 1,30E-05 | 7,17E-05 |
| ENSG00000013288.8 | MAN2B2 | protein_coding | 4271,54744 | 0,563986961 | 0,054546751 | 10,33951513 | 4,67E-25 | 1,22E-23 |
| ENSG00000013297.11 | CLDN11 | protein_coding | 40,19168595 | -0,937888561 | 0,361396105 | -2,595181708 | 0,009454092 | 0,028544891 |
| ENSG00000013306.16 | SLC25A39 | protein_coding | 6188,438293 | -0,33119097 | 0,063106985 | -5,248087391 | 1,54E-07 | 1,11E-06 |
| ENSG00000013364.19 | MVP | protein_coding | 474,6648224 | 2,637398811 | 0,165855819 | 15,90175629 | 6,16E-57 | 4,12E-55 |
| ENSG00000013374.16 | NUB1 | protein_coding | 18164,05041 | 1,971231952 | 0,049319723 | 39,96843076 | 0 | 0 |
| ENSG00000013523.9 | ANGEL1 | protein_coding | 8535,196634 | 0,212239365 | 0,044530612 | 4,766145286 | 1,88E-06 | 1,17E-05 |
| ENSG00000013563.14 | DNASE1L1 | protein_coding | 1827,000696 | -0,19801757 | 0,080420399 | -2,462280361 | 0,01380567 | 0,039561022 |
| ENSG00000013573.17 | DDX11 | protein_coding | 6510,587371 | -0,254500128 | 0,06050236 | -4,206449623 | 2,59E-05 | 0,000135984 |
| ENSG00000013619.14 | MAMLD1 | protein_coding | 1437,111246 | 0,785668566 | 0,08532575 | 9,20787181 | 3,33E-20 | 6,82E-19 |
| ENSG00000013810.19 | TACC3 | protein_coding | 7001,580613 | -0,665551322 | 0,048394737 | -13,75255594 | 4,92E-43 | 2,42E-41 |
| ENSG00000014216.15 | CAPN1 | protein_coding | 6849,223959 | 0,173876659 | 0,060808111 | 2,859432021 | 0,004244004 | 0,014123045 |
| ENSG00000014641.18 | MDH1 | protein_coding | 21555,20235 | -0,293570523 | 0,040653326 | -7,221316214 | 5,15E-13 | 6,65E-12 |
| ENSG00000014824.14 | SLC30A9 | protein_coding | 28095,66929 | -0,247193854 | 0,053472242 | -4,622844358 | 3,79E-06 | 2,26E-05 |
| ENSG00000014914.21 | MTMR11 | protein_coding | 1472,442344 | 0,844696804 | 0,084128755 | 10,04052428 | 1,01E-23 | 2,47E-22 |
| ENSG00000014919.12 | COX15 | protein_coding | 5314,902832 | -0,173584539 | 0,056812881 | -3,055372952 | 0,002247808 | 0,008010345 |
| ENSG00000015133.19 | CCDC88C | protein_coding | 3413,406547 | 0,361285871 | 0,073046814 | 4,945949731 | 7,58E-07 | 5,01E-06 |
| ENSG00000015171.19 | ZMYND11 | protein_coding | 8443,996443 | -0,384290722 | 0,051414364 | -7,474384434 | 7,76E-14 | 1,07E-12 |
| ENSG00000015285.10 | WAS | protein_coding | 199,054147 | -0,706494529 | 0,169272713 | -4,173705934 | 3,00E-05 | 0,000155464 |
| ENSG00000015413.9 | DPEP1 | protein_coding | 2273,747598 | 0,318254669 | 0,085050341 | 3,741956434 | 0,000182593 | 0,000824306 |
| ENSG00000015520.14 | NPC1L1 | protein_coding | 575,197363 | 0,501526597 | 0,104324927 | 4,807351525 | 1,53E-06 | 9,66E-06 |
| ENSG00000015676.18 | NUDCD3 | protein_coding | 6433,014024 | 0,112344898 | 0,046641205 | 2,408704894 | 0,016009237 | 0,044892478 |
| ENSG00000016082.15 | ISL1 | protein_coding | 19745,5355 | 0,219927895 | 0,040148197 | 5,477902195 | 4,30E-08 | 3,32E-07 |
| ENSG00000016391.11 | CHDH | protein_coding | 1742,906451 | 0,173105899 | 0,068512655 | 2,526626644 | 0,011516386 | 0,0337134 |
| ENSG00000016402.13 | IL20RA | protein_coding | 744,3364237 | 0,773243425 | 0,112589729 | 6,867797208 | 6,52E-12 | 7,64E-11 |
| ENSG00000016864.18 | GLT8D1 | protein_coding | 6319,59681 | -0,1494906 | 0,054360149 | -2,750003478 | 0,005959463 | 0,019051492 |
| ENSG00000018236.15 | CNTN1 | protein_coding | 38682,65625 | -0,657047007 | 0,050016504 | -13,13660403 | 2,03E-39 | 9,02E-38 |
| ENSG00000018510.15 | AGPS | protein_coding | 12799,69013 | -0,227273542 | 0,046047023 | -4,935683733 | 7,99E-07 | 5,26E-06 |
| ENSG00000019505.8 | SYT13 | protein_coding | 24850,79998 | -0,301581681 | 0,047055581 | -6,409052384 | 1,46E-10 | 1,50E-09 |
| ENSG00000019582.15 | CD74 | protein_coding | 11676,66948 | 1,90394173 | 0,057951528 | 32,85403855 | 9,97E-237 | 2,24E-234 |
| ENSG00000020129.16 | NCDN | protein_coding | 1900,455669 | 0,185650081 | 0,071071024 | 2,612176797 | 0,008996772 | 0,02734382 |
| ENSG00000020181.17 | ADGRA2 | protein_coding | 420,6529484 | -0,437208105 | 0,135140126 | -3,235220491 | 0,001215488 | 0,004616845 |
| ENSG00000020577.13 | SAMD4A | protein_coding | 624,8017078 | 0,891286068 | 0,122573775 | 7,271425452 | 3,56E-13 | 4,66E-12 |
| ENSG00000020922.12 | MRE11 | protein_coding | 6084,288 | -0,251786221 | 0,056280267 | -4,473792226 | 7,68E-06 | 4,37E-05 |
| ENSG00000021300.14 | PLEKHB1 | protein_coding | 14968,56101 | -0,166531458 | 0,046370073 | -3,591356382 | 0,000328961 | 0,001417961 |
| ENSG00000021355.13 | SERPINB1 | protein_coding | 10802,86353 | 0,346761098 | 0,044969811 | 7,71097527 | 1,25E-14 | 1,82E-13 |

| | | | | | | | | |
|---|---|---|---|---|---|---|---|---|
| ENSG00000021488.13 | SLC7A9 | protein_coding | 3531,593777 | 0,138808969 | 0,058460361 | 2,374411753 | 0,017576937 | 0,048604334 |
| ENSG00000021776.11 | AQR | protein_coding | 10534,03554 | -0,126218516 | 0,047039304 | -2,683256459 | 0,007290906 | 0,022742655 |
| ENSG00000022840.16 | RNF10 | protein_coding | 11723,13238 | 0,583680574 | 0,050670179 | 11,51921286 | 1,06E-30 | 3,47E-29 |
| ENSG00000023171.18 | GRAMD1B | protein_coding | 5351,381543 | 0,276227731 | 0,058480062 | 4,723451414 | 2,32E-06 | 1,43E-05 |
| ENSG00000023228.14 | NDUFS1 | protein_coding | 21570,73177 | -0,236767738 | 0,054307859 | -4,359732467 | 1,30E-05 | 7,17E-05 |
| ENSG00000023318.8 | ERP44 | protein_coding | 12235,80586 | 0,405374782 | 0,043432416 | 9,333461465 | 1,02E-20 | 2,17E-19 |
| ENSG00000023445.14 | BIRC3 | protein_coding | 1036,351153 | 3,737616415 | 0,30543559 | 12,23700359 | 1,97E-34 | 7,40E-33 |
| ENSG00000023697.13 | DERA | protein_coding | 5582,743506 | -0,280734441 | 0,051693894 | -5,430707898 | 5,61E-08 | 4,28E-07 |
| ENSG00000023734.11 | STRAP | protein_coding | 19512,19239 | -0,368570973 | 0,041233253 | -8,938682984 | 3,94E-19 | 7,66E-18 |
| ENSG00000023839.12 | ABCC2 | protein_coding | 401,8963876 | 0,469076013 | 0,176709896 | 2,654497708 | 0,007942658 | 0,024527547 |
| ENSG00000023909.10 | GCLM | protein_coding | 7529,423095 | 0,409359478 | 0,047820872 | 8,560267914 | 1,13E-17 | 2,01E-16 |
| ENSG00000024048.10 | UBR2 | protein_coding | 9614,939068 | 0,627258867 | 0,047825697 | 13,11551965 | 2,68E-39 | 1,18E-37 |
| ENSG00000024526.17 | DEPDC1 | protein_coding | 7082,612212 | -0,808595763 | 0,064815303 | -12,47538357 | 1,02E-35 | 4,03E-34 |
| ENSG00000025293.17 | PHF20 | protein_coding | 9426,135964 | -0,170298942 | 0,047869023 | -3,557602196 | 0,000374256 | 0,001595175 |
| ENSG00000025708.14 | TYMP | protein_coding | 581,0231077 | 3,880957108 | 0,399101995 | 9,724223763 | 2,38E-22 | 5,46E-21 |
| ENSG00000026036.22 | RTEL1-TNFRSF6E | protein_coding | 1638,935875 | -0,223273708 | 0,082692776 | -2,700038858 | 0,006933138 | 0,021789723 |
| ENSG00000026103.22 | FAS | protein_coding | 160,4253203 | 3,31205136 | 0,240172807 | 13,79028461 | 2,92E-43 | 1,45E-41 |
| ENSG00000026297.15 | RNASET2 | protein_coding | 2546,944245 | 0,588807041 | 0,065725266 | 8,958610237 | 3,29E-19 | 6,41E-18 |
| ENSG00000026508.18 | CD44 | protein_coding | 12903,37192 | 1,582432131 | 0,065577791 | 24,13061051 | 1,19E-128 | 1,73E-126 |
| ENSG00000026652.14 | AGPAT4 | protein_coding | 2102,949466 | 0,398686139 | 0,074634598 | 5,34184081 | 9,20E-08 | 6,81E-07 |
| ENSG00000026950.17 | BTN3A1 | protein_coding | 5792,183876 | 3,963472995 | 0,064086574 | 61,84560551 | 0 | 0 |
| ENSG00000027075.15 | PRKCH | protein_coding | 4395,540097 | 0,462904988 | 0,053366785 | 8,674028022 | 4,17E-18 | 7,66E-17 |
| ENSG00000027697.14 | IFNGR1 | protein_coding | 5470,932132 | 0,166455832 | 0,051615238 | 3,224935885 | 0,001260009 | 0,004771457 |
| ENSG00000027847.14 | B4GALT7 | protein_coding | 1219,271714 | 0,214892703 | 0,082660209 | 2,599711589 | 0,009330214 | 0,028223866 |
| ENSG00000028116.17 | VRK2 | protein_coding | 2065,850625 | 0,282483726 | 0,065637705 | 4,303680754 | 1,68E-05 | 9,08E-05 |
| ENSG00000028137.19 | TNFRSF1B | protein_coding | 23,12950303 | 2,033104911 | 0,601238684 | 3,381527111 | 0,000720841 | 0,002887142 |
| ENSG00000028277.21 | POU2F2 | protein_coding | 129,6049685 | -1,233338419 | 0,273521799 | -4,509104666 | 6,51E-06 | 3,75E-05 |
| ENSG00000028839.10 | TBPL1 | protein_coding | 3232,970607 | 0,253205981 | 0,065830906 | 3,846308617 | 0,000119911 | 0,00056348 |
| ENSG00000029153.14 | ARNTL2 | protein_coding | 5625,14053 | -0,248760469 | 0,070835729 | -3,511793732 | 0,000445093 | 0,001869587 |
| ENSG00000029364.12 | SLC39A9 | protein_coding | 21525,07591 | -0,244078384 | 0,044761004 | -5,452924634 | 4,95E-08 | 3,80E-07 |
| ENSG00000029725.17 | RABEP1 | protein_coding | 19535,70611 | -0,113220189 | 0,046701767 | -2,424323461 | 0,015336938 | 0,04323887 |
| ENSG00000029993.15 | HMGB3 | protein_coding | 12408,09265 | -0,597822844 | 0,043420696 | -13,7681543 | 3,96E-43 | 1,96E-41 |
| ENSG00000030066.13 | NUP160 | protein_coding | 18795,90586 | 0,177988915 | 0,041684387 | 4,269918042 | 1,96E-05 | 0,000104499 |
| ENSG00000030110.13 | BAK1 | protein_coding | 3382,825769 | 0,326927829 | 0,053904633 | 6,064930068 | 1,32E-09 | 1,22E-08 |
| ENSG00000030419.16 | IKZF2 | protein_coding | 1197,114125 | 0,493028039 | 0,093197528 | 5,290140736 | 1,22E-07 | 8,94E-07 |
| ENSG00000030582.18 | GRN | protein_coding | 17796,04982 | 0,558947282 | 0,04654679 | 12,00828855 | 3,21E-33 | 1,17E-31 |

| | | | | | | | | |
|---|---|---|---|---|---|---|---|---|
| ENSG00000031081.10 | ARHGAP31 | protein_coding | 174,3275156 | 0,848266566 | 0,203950972 | 4,159169037 | 3,19E-05 | 0,000165125 |
| ENSG00000031698.13 | SARS | protein_coding | 12435,91842 | 0,167517092 | 0,043741751 | 3,829684188 | 0,000128308 | 0,000599187 |
| ENSG00000032219.18 | ARID4A | protein_coding | 3492,812776 | 0,173517948 | 0,057182565 | 3,034455483 | 0,002409703 | 0,008518927 |
| ENSG00000032444.16 | PNPLA6 | protein_coding | 4186,226641 | -0,246689236 | 0,078060026 | -3,160250507 | 0,001576335 | 0,005849723 |
| ENSG00000032742.17 | IFT88 | protein_coding | 2228,167388 | -0,223244744 | 0,066264229 | -3,369008422 | 0,000754391 | 0,003005131 |
| ENSG00000033030.13 | ZCCHC8 | protein_coding | 4399,60417 | 0,230069099 | 0,055947404 | 4,112239083 | 3,92E-05 | 0,000199908 |
| ENSG00000033100.16 | CHPF2 | protein_coding | 6569,713928 | 0,267106486 | 0,050029716 | 5,338956641 | 9,35E-08 | 6,90E-07 |
| ENSG00000033178.13 | UBA6 | protein_coding | 14498,54662 | 0,652008628 | 0,045419466 | 14,35526837 | 9,87E-47 | 5,24E-45 |
| ENSG00000033627.16 | ATP6V0A1 | protein_coding | 21966,35854 | 0,254693315 | 0,044038158 | 5,783468854 | 7,32E-09 | 6,19E-08 |
| ENSG00000033800.13 | PIAS1 | protein_coding | 8717,626965 | 0,191401781 | 0,050054643 | 3,823856693 | 0,00013138 | 0,000612138 |
| ENSG00000034053.14 | APBA2 | protein_coding | 197,9756279 | 0,976260486 | 0,203648531 | 4,79384988 | 1,64E-06 | 1,03E-05 |
| ENSG00000034510.6 | TMSB10 | protein_coding | 50376,7178 | 0,541450543 | 0,052304877 | 10,35181747 | 4,11E-25 | 1,07E-23 |
| ENSG00000034677.13 | RNF19A | protein_coding | 5227,319744 | 0,370098828 | 0,0569351 | 6,500363142 | 8,01E-11 | 8,41E-10 |
| ENSG00000034713.8 | GABARAPL2 | protein_coding | 11885,52085 | -0,139249919 | 0,048081304 | -2,896134398 | 0,003777906 | 0,012735791 |
| ENSG00000035141.8 | FAM136A | protein_coding | 9647,659232 | 0,134966161 | 0,048005195 | 2,811490751 | 0,004931251 | 0,016123987 |
| ENSG00000035499.13 | DEPDC1B | protein_coding | 2997,941497 | -0,774061205 | 0,058176585 | -13,30537373 | 2,15E-40 | 9,78E-39 |
| ENSG00000035664.11 | DAPK2 | protein_coding | 605,0117189 | -0,460792677 | 0,153389164 | -3,004075809 | 0,002663889 | 0,009319473 |
| ENSG00000036530.8 | CYP46A1 | protein_coding | 986,5018193 | -0,528873705 | 0,103495416 | -5,110117202 | 3,22E-07 | 2,24E-06 |
| ENSG00000037241.7 | RPL26L1 | protein_coding | 3682,694086 | -0,305603262 | 0,060727365 | -5,032381429 | 4,84E-07 | 3,29E-06 |
| ENSG00000037474.15 | NSUN2 | protein_coding | 8246,614974 | 0,137413583 | 0,045130626 | 3,0447967 | 0,002328376 | 0,008265167 |
| ENSG00000037637.11 | FBXO42 | protein_coding | 4987,345865 | 0,170411885 | 0,055221711 | 3,085958088 | 0,002028974 | 0,007316594 |
| ENSG00000037757.14 | MRI1 | protein_coding | 2019,230646 | -0,177847298 | 0,067562516 | -2,632336794 | 0,008479975 | 0,025969492 |
| ENSG00000037897.17 | METTL1 | protein_coding | 539,7800669 | 0,429407891 | 0,108080981 | 3,973019932 | 7,10E-05 | 0,000346884 |
| ENSG00000038002.9 | AGA | protein_coding | 3867,558408 | 0,350362453 | 0,057814818 | 6,060080531 | 1,36E-09 | 1,25E-08 |
| ENSG00000038210.14 | PI4K2B | protein_coding | 10140,91876 | 0,438001522 | 0,04852812 | 9,02572619 | 1,79E-19 | 3,51E-18 |
| ENSG00000038219.13 | BOD1L1 | protein_coding | 9235,144885 | 0,367500842 | 0,050015746 | 7,347702895 | 2,02E-13 | 2,69E-12 |
| ENSG00000038274.17 | MAT2B | protein_coding | 3411,639964 | 0,472556698 | 0,056969182 | 8,294953218 | 1,09E-16 | 1,82E-15 |
| ENSG00000038382.20 | TRIO | protein_coding | 12679,99519 | 0,305069065 | 0,047716452 | 6,393372755 | 1,62E-10 | 1,66E-09 |
| ENSG00000038427.16 | VCAN | protein_coding | 432,243292 | 1,43308514 | 0,16507192 | 8,681580345 | 3,90E-18 | 7,18E-17 |
| ENSG00000038532.16 | CLEC16A | protein_coding | 4281,799556 | 0,348944376 | 0,056920757 | 6,130353767 | 8,77E-10 | 8,25E-09 |
| ENSG00000039068.19 | CDH1 | protein_coding | 24732,35567 | 0,28893591 | 0,046023627 | 6,277990909 | 3,43E-10 | 3,39E-09 |
| ENSG00000039319.17 | ZFYVE16 | protein_coding | 9727,244253 | 0,348966621 | 0,058137461 | 6,002440032 | 1,94E-09 | 1,76E-08 |
| ENSG00000039523.20 | RIPOR1 | protein_coding | 1366,776588 | 0,302517526 | 0,098508558 | 3,07097709 | 0,002133595 | 0,007649744 |
| ENSG00000039560.14 | RAI14 | protein_coding | 1942,184747 | 0,18663161 | 0,068258732 | 2,734179277 | 0,006253599 | 0,019890102 |
| ENSG00000040199.18 | PHLPP2 | protein_coding | 5604,522126 | 0,53453509 | 0,051959374 | 10,28755824 | 8,02E-25 | 2,07E-23 |
| ENSG00000040275.17 | SPDL1 | protein_coding | 6459,617002 | -0,41505606 | 0,057599049 | -7,205953281 | 5,76E-13 | 7,43E-12 |

| | | | | | | | | |
|---|---|---|---|---|---|---|---|---|
| ENSG00000040487.13 | PQLC2 | protein_coding | 3224,227257 | 0,34196591 | 0,054419933 | 6,283835551 | 3,30E-10 | 3,27E-09 |
| ENSG00000040933.15 | INPP4A | protein_coding | 6064,208496 | 0,260319405 | 0,048088587 | 5,413330249 | 6,19E-08 | 4,69E-07 |
| ENSG00000041353.10 | RAB27B | protein_coding | 1444,524363 | -0,294451608 | 0,080764658 | -3,645797758 | 0,000266564 | 0,00116915 |
| ENSG00000041357.16 | PSMA4 | protein_coding | 20539,456 | 0,137499433 | 0,042351996 | 3,24658686 | 0,001167978 | 0,004453671 |
| ENSG00000041880.14 | PARP3 | protein_coding | 464,2941054 | 0,787448958 | 0,141063905 | 5,582214363 | 2,37E-08 | 1,90E-07 |
| ENSG00000042088.13 | TDP1 | protein_coding | 3112,676453 | -0,322070695 | 0,057681641 | -5,583591108 | 2,36E-08 | 1,88E-07 |
| ENSG00000042429.12 | MED17 | protein_coding | 5396,307774 | 0,219338975 | 0,051464149 | 4,261976137 | 2,03E-05 | 0,000108079 |
| ENSG00000042445.14 | RETSAT | protein_coding | 4357,112591 | 0,147754618 | 0,053759319 | 2,748446597 | 0,005987839 | 0,019131336 |
| ENSG00000042493.16 | CAPG | protein_coding | 845,0618421 | 0,278854908 | 0,103157447 | 2,703197055 | 0,006867602 | 0,021604871 |
| ENSG00000042753.11 | AP2S1 | protein_coding | 5498,399215 | -0,178558033 | 0,047672639 | -3,74550343 | 0,000180032 | 0,000813725 |
| ENSG00000043143.20 | JADE2 | protein_coding | 2074,646818 | 0,961425226 | 0,083598481 | 11,50051065 | 1,31E-30 | 4,28E-29 |
| ENSG00000044090.8 | CUL7 | protein_coding | 6620,119658 | -0,166891511 | 0,051301823 | -3,253130221 | 0,001141412 | 0,00436494 |
| ENSG00000044574.8 | HSPA5 | protein_coding | 124466,9485 | 0,613275998 | 0,044856705 | 13,67189124 | 1,49E-42 | 7,27E-41 |
| ENSG00000046651.15 | OFD1 | protein_coding | 8185,342548 | 0,162709093 | 0,049958835 | 3,25686324 | 0,001126507 | 0,004314539 |
| ENSG00000047188.16 | YTHDC2 | protein_coding | 5561,741818 | -0,153512754 | 0,053890184 | -2,848621828 | 0,004390904 | 0,014530264 |
| ENSG00000047315.16 | POLR2B | protein_coding | 29721,30477 | -0,210103811 | 0,039929366 | -5,261886998 | 1,43E-07 | 1,03E-06 |
| ENSG00000047410.14 | TPR | protein_coding | 33216,53568 | -0,149223751 | 0,041097658 | -3,630955079 | 0,000282374 | 0,001231776 |
| ENSG00000047457.14 | CP | protein_coding | 33,23482622 | 1,635480972 | 0,550549653 | 2,970633011 | 0,002971867 | 0,010266681 |
| ENSG00000047579.19 | DTNBP1 | protein_coding | 1867,929529 | 0,306993404 | 0,069359765 | 4,426102153 | 9,60E-06 | 5,39E-05 |
| ENSG00000047597.7 | XK | protein_coding | 4874,432417 | -0,181994646 | 0,05232686 | -3,478034915 | 0,000505104 | 0,002091499 |
| ENSG00000048028.11 | USP28 | protein_coding | 4072,443179 | 0,367536928 | 0,058357798 | 6,297991722 | 3,02E-10 | 3,00E-09 |
| ENSG00000048392.11 | RRM2B | protein_coding | 6277,370369 | -0,170554106 | 0,048340468 | -3,528184841 | 0,00041842 | 0,001764041 |
| ENSG00000048405.10 | ZNF800 | protein_coding | 4282,660763 | 0,34497557 | 0,057834669 | 5,964857708 | 2,45E-09 | 2,19E-08 |
| ENSG00000048471.14 | SNX29 | protein_coding | 7852,29688 | 0,277807713 | 0,059171524 | 4,694956223 | 2,67E-06 | 1,63E-05 |
| ENSG00000048540.15 | LMO3 | protein_coding | 345,4285988 | -0,435193818 | 0,137896294 | -3,155950063 | 0,001599762 | 0,005926403 |
| ENSG00000048544.6 | MRPS10 | protein_coding | 5838,068616 | -0,166034384 | 0,05734615 | -2,895301335 | 0,003787947 | 0,012763908 |
| ENSG00000048649.13 | RSF1 | protein_coding | 12657,85494 | 0,281727097 | 0,051222584 | 5,500056286 | 3,80E-08 | 2,95E-07 |
| ENSG00000048707.15 | VPS13D | protein_coding | 6344,483466 | 0,190137826 | 0,069720058 | 2,727161018 | 0,006388186 | 0,020252348 |
| ENSG00000048828.17 | FAM120A | protein_coding | 15537,92646 | -0,270469569 | 0,045409106 | -5,95628477 | 2,58E-09 | 2,30E-08 |
| ENSG00000049130.16 | KITLG | protein_coding | 1243,401109 | -0,49246264 | 0,090097626 | -5,4658781 | 4,61E-08 | 3,54E-07 |
| ENSG00000049167.14 | ERCC8 | protein_coding | 1929,267729 | -0,295089623 | 0,078768596 | -3,746285163 | 0,000179473 | 0,000811846 |
| ENSG00000049192.15 | ADAMTS6 | protein_coding | 1303,705152 | 1,07845 | 0,079011656 | 13,64925198 | 2,04E-42 | 9,83E-41 |
| ENSG00000049239.12 | H6PD | protein_coding | 2949,527102 | 0,142388872 | 0,059250629 | 2,403162225 | 0,016253975 | 0,045465481 |
| ENSG00000049245.13 | VAMP3 | protein_coding | 10026,68212 | -0,267023164 | 0,050820721 | -5,254218314 | 1,49E-07 | 1,08E-06 |
| ENSG00000049249.8 | TNFRSF9 | protein_coding | 30,71018354 | 1,245596437 | 0,474379724 | 2,62573709 | 0,008646157 | 0,026428076 |
| ENSG00000049323.16 | LTBP1 | protein_coding | 486,1653189 | -0,81874392 | 0,115677247 | -7,077830265 | 1,46E-12 | 1,83E-11 |

| | | | | | | | | |
|---|---|---|---|---|---|---|---|---|
| ENSG00000049449.9 | RCN1 | protein_coding | 29393,69052 | 0,662544075 | 0,045084963 | 14,69545576 | 6,89E-49 | 3,83E-47 |
| ENSG00000049759.18 | NEDD4L | protein_coding | 8139,09669 | 0,134635249 | 0,05040878 | 2,670869001 | 0,007565517 | 0,023475629 |
| ENSG00000049768.15 | FOXP3 | protein_coding | 243,8132083 | 0,6160071 | 0,158759719 | 3,880122142 | 0,000104404 | 0,000495265 |
| ENSG00000049860.14 | HEXB | protein_coding | 14692,97447 | 0,534116257 | 0,042438262 | 12,58572412 | 2,53E-36 | 1,02E-34 |
| ENSG00000050130.18 | JKAMP | protein_coding | 7520,204764 | -0,197730286 | 0,050498272 | -3,915585159 | 9,02E-05 | 0,000432739 |
| ENSG00000050165.17 | DKK3 | protein_coding | 2111,64423 | 0,220584095 | 0,067356823 | 3,274858964 | 0,001057147 | 0,004076637 |
| ENSG00000050327.15 | ARHGEF5 | protein_coding | 4730,293945 | 0,385176923 | 0,070902927 | 5,432454435 | 5,56E-08 | 4,24E-07 |
| ENSG00000050344.9 | NFE2L3 | protein_coding | 3644,684744 | 0,37138824 | 0,061621383 | 6,026937746 | 1,67E-09 | 1,52E-08 |
| ENSG00000050393.11 | MCUR1 | protein_coding | 7610,228274 | -0,28642952 | 0,05527147 | -5,182230862 | 2,19E-07 | 1,55E-06 |
| ENSG00000050405.13 | LIMA1 | protein_coding | 15071,58584 | 0,434143014 | 0,054709188 | 7,935468028 | 2,10E-15 | 3,24E-14 |
| ENSG00000050438.17 | SLC4A8 | protein_coding | 16787,03245 | -0,353546519 | 0,054577642 | -6,477863589 | 9,30E-11 | 9,72E-10 |
| ENSG00000050628.20 | PTGER3 | protein_coding | 1517,744672 | 0,25067207 | 0,072105678 | 3,476453941 | 0,000508091 | 0,002100885 |
| ENSG00000050748.17 | MAPK9 | protein_coding | 6177,610507 | -0,35986613 | 0,054886648 | -6,556533198 | 5,51E-11 | 5,89E-10 |
| ENSG00000050767.17 | COL23A1 | protein_coding | 290,8178073 | -0,729123313 | 0,160853455 | -4,532842107 | 5,82E-06 | 3,38E-05 |
| ENSG00000051108.15 | HERPUD1 | protein_coding | 10988,51419 | 0,642513843 | 0,054349266 | 11,82194144 | 3,01E-32 | 1,07E-30 |
| ENSG00000051341.14 | POLQ | protein_coding | 4548,268892 | 0,577582645 | 0,057816804 | 9,989875024 | 1,69E-23 | 4,08E-22 |
| ENSG00000051596.9 | THOC3 | protein_coding | 3923,266911 | -0,273996501 | 0,07613138 | -3,598995569 | 0,000319449 | 0,001380128 |
| ENSG00000051620.11 | HEBP2 | protein_coding | 3332,449823 | -0,78814514 | 0,071411877 | -11,03661143 | 2,54E-28 | 7,53E-27 |
| ENSG00000052795.13 | FNIP2 | protein_coding | 27786,96227 | 0,139125639 | 0,040075896 | 3,471554012 | 0,000517455 | 0,002135179 |
| ENSG00000052802.13 | MSMO1 | protein_coding | 28820,04185 | 0,231980055 | 0,049334443 | 4,702192691 | 2,57E-06 | 1,58E-05 |
| ENSG00000052841.15 | TTC17 | protein_coding | 9295,774968 | 0,164110233 | 0,050627765 | 3,241506562 | 0,001188997 | 0,004526151 |
| ENSG00000053108.17 | FSTL4 | protein_coding | 154,8548444 | 0,621968812 | 0,245475189 | 2,53373392 | 0,011285439 | 0,033175388 |
| ENSG00000053371.12 | AKR7A2 | protein_coding | 2957,063475 | -0,35819945 | 0,060959107 | -5,876061316 | 4,20E-09 | 3,66E-08 |
| ENSG00000053372.5 | MRTO4 | protein_coding | 3009,266131 | -0,220072315 | 0,061961821 | -3,551740613 | 0,000382692 | 0,001627127 |
| ENSG00000053438.11 | NNAT | protein_coding | 3628,309376 | -0,497660184 | 0,063368256 | -7,853461952 | 4,05E-15 | 6,15E-14 |
| ENSG00000053524.12 | MCF2L2 | protein_coding | 163,9296086 | 1,052634734 | 0,293761329 | 3,5832992 | 0,000339282 | 0,001458257 |
| ENSG00000054148.17 | PHPT1 | protein_coding | 6592,819865 | -0,621005793 | 0,051212978 | -12,12594584 | 7,70E-34 | 2,82E-32 |
| ENSG00000054219.11 | LY75 | protein_coding | 2907,325968 | 0,205254591 | 0,056310487 | 3,645050874 | 0,000267339 | 0,001172323 |
| ENSG00000054267.22 | ARID4B | protein_coding | 11353,18447 | 0,369831678 | 0,046909981 | 7,883858994 | 3,17E-15 | 4,85E-14 |
| ENSG00000054356.14 | PTPRN | protein_coding | 36480,43776 | 0,235620281 | 0,040495291 | 5,818461294 | 5,94E-09 | 5,08E-08 |
| ENSG00000054392.13 | HHAT | protein_coding | 958,4593866 | 0,525323196 | 0,086242878 | 6,091206698 | 1,12E-09 | 1,04E-08 |
| ENSG00000054523.17 | KIF1B | protein_coding | 14370,892 | -0,167379393 | 0,055111062 | -3,037128775 | 0,002388434 | 0,008454358 |
| ENSG00000054793.13 | ATP9A | protein_coding | 38191,42431 | -0,171612057 | 0,050759587 | -3,380879699 | 0,000722542 | 0,002892351 |
| ENSG00000054965.10 | FAM168A | protein_coding | 13552,40372 | -0,139791601 | 0,050378971 | -2,774800672 | 0,005523555 | 0,017817383 |
| ENSG00000055070.17 | SZRD1 | protein_coding | 23387,21552 | -0,32146068 | 0,050775825 | -6,330978919 | 2,44E-10 | 2,44E-09 |
| ENSG00000055118.14 | KCNH2 | protein_coding | 6762,660135 | -0,45959206 | 0,054164647 | -8,48509282 | 2,16E-17 | 3,77E-16 |

| | | | | | | | | |
|---|---|---|---|---|---|---|---|---|
| ENSG00000055130.17 | CUL1 | protein_coding | 7166,02315 | 0,351648097 | 0,047599669 | 7,387616371 | 1,49E-13 | 2,03E-12 |
| ENSG00000055163.20 | CYFIP2 | protein_coding | 10573,69371 | -0,16884569 | 0,047524267 | -3,552831016 | 0,000381109 | 0,00162193 |
| ENSG00000055208.19 | TAB2 | protein_coding | 13485,84379 | 0,228542733 | 0,042962224 | 5,31962067 | 1,04E-07 | 7,65E-07 |
| ENSG00000055211.14 | GINM1 | protein_coding | 5608,289786 | 0,30351826 | 0,049801021 | 6,094619275 | 1,10E-09 | 1,02E-08 |
| ENSG00000055332.18 | EIF2AK2 | protein_coding | 45835,31826 | 2,978564878 | 0,063677969 | 46,77543815 | 0 | 0 |
| ENSG00000055609.18 | KMT2C | protein_coding | 21685,94313 | 0,276332462 | 0,045775877 | 6,036639448 | 1,57E-09 | 1,44E-08 |
| ENSG00000055732.13 | MCOLN3 | protein_coding | 5560,872693 | 0,157541844 | 0,057113456 | 2,758401505 | 0,005808481 | 0,018621719 |
| ENSG00000055950.16 | MRPL43 | protein_coding | 2631,679437 | -0,281226904 | 0,063798443 | -4,4080528 | 1,04E-05 | 5,83E-05 |
| ENSG00000056277.16 | ZNF280C | protein_coding | 1280,918861 | 0,417536302 | 0,080002227 | 5,219058477 | 1,80E-07 | 1,29E-06 |
| ENSG00000056291.17 | NPFFR2 | protein_coding | 153,8445969 | 0,827389452 | 0,198985448 | 4,158039987 | 3,21E-05 | 0,000165867 |
| ENSG00000056736.10 | IL17RB | protein_coding | 1558,370477 | 0,242151709 | 0,074035788 | 3,270738597 | 0,00107267 | 0,004130129 |
| ENSG00000056998.20 | GYG2 | protein_coding | 4551,395575 | -0,231576283 | 0,05618104 | -4,121965077 | 3,76E-05 | 0,000191911 |
| ENSG00000057019.16 | DCBLD2 | protein_coding | 3203,111948 | -0,17038966 | 0,057561778 | -2,960118077 | 0,003075212 | 0,010592186 |
| ENSG00000057608.17 | GDI2 | protein_coding | 19925,21132 | 0,173510776 | 0,050894164 | 3,409246965 | 0,000651425 | 0,0026339 |
| ENSG00000057704.13 | TMCC3 | protein_coding | 15010,3278 | 0,448358579 | 0,048328735 | 9,277266928 | 1,74E-20 | 3,63E-19 |
| ENSG00000058056.9 | USP13 | protein_coding | 5620,390046 | -0,279130948 | 0,055726246 | -5,008967345 | 5,47E-07 | 3,70E-06 |
| ENSG00000058063.16 | ATP11B | protein_coding | 7105,027633 | -0,17474475 | 0,052240712 | -3,344991754 | 0,00082285 | 0,003252245 |
| ENSG00000058085.14 | LAMC2 | protein_coding | 504,5056256 | 0,495080634 | 0,106876991 | 4,632247112 | 3,62E-06 | 2,16E-05 |
| ENSG00000058091.17 | CDK14 | protein_coding | 36840,3577 | -0,343774705 | 0,054650983 | -6,29036631 | 3,17E-10 | 3,14E-09 |
| ENSG00000058262.10 | SEC61A1 | protein_coding | 31790,40108 | 0,15194027 | 0,04573536 | 3,322161926 | 0,000893228 | 0,003499683 |
| ENSG00000058335.15 | RASGRF1 | protein_coding | 603,9070341 | -0,366959467 | 0,129823846 | -2,82659526 | 0,004704575 | 0,015456908 |
| ENSG00000058404.20 | CAMK2B | protein_coding | 2093,335904 | -0,202077197 | 0,078315841 | -2,5802851 | 0,009871878 | 0,029599853 |
| ENSG00000058453.17 | CROCC | protein_coding | 1062,791707 | -0,404865699 | 0,114480818 | -3,53653745 | 0,000405409 | 0,001713359 |
| ENSG00000058673.16 | ZC3H11A | protein_coding | 23302,98908 | 0,388403343 | 0,042161929 | 9,21218151 | 3,20E-20 | 6,57E-19 |
| ENSG00000058729.11 | RIOK2 | protein_coding | 6365,66621 | 0,161121718 | 0,051555518 | 3,125208036 | 0,001776794 | 0,006509415 |
| ENSG00000058804.12 | NDC1 | protein_coding | 6548,96396 | -0,54589509 | 0,062778878 | -8,695521627 | 3,45E-18 | 6,37E-17 |
| ENSG00000058866.15 | DGKG | protein_coding | 1788,126759 | 2,07829804 | 0,070527452 | 29,46793049 | 7,42E-191 | 1,45E-188 |
| ENSG00000059145.18 | UNKL | protein_coding | 465,4232601 | 0,699983926 | 0,142443747 | 4,91410778 | 8,92E-07 | 5,83E-06 |
| ENSG00000059378.12 | PARP12 | protein_coding | 13995,60783 | 4,797729221 | 0,054939524 | 87,32746198 | 0 | 0 |
| ENSG00000059588.10 | TARBP1 | protein_coding | 4600,229401 | 0,371400359 | 0,066111402 | 5,617795845 | 1,93E-08 | 1,56E-07 |
| ENSG00000059728.11 | MXD1 | protein_coding | 5804,731069 | 0,141324727 | 0,058410313 | 2,419516711 | 0,015541147 | 0,043710816 |
| ENSG00000059758.8 | CDK17 | protein_coding | 3639,728166 | 0,828013266 | 0,066863431 | 12,38364908 | 3,20E-35 | 1,24E-33 |
| ENSG00000059769.20 | DNAJC25 | protein_coding | 1744,001762 | -0,212823411 | 0,071060094 | -2,994977903 | 0,002744647 | 0,009570761 |
| ENSG00000059915.17 | PSD | protein_coding | 3158,455336 | -0,429411808 | 0,082068353 | -5,232367782 | 1,67E-07 | 1,20E-06 |
| ENSG00000060339.14 | CCAR1 | protein_coding | 15967,82176 | -0,276185227 | 0,054933359 | -5,027641381 | 4,97E-07 | 3,37E-06 |
| ENSG00000060491.16 | OGFR | protein_coding | 9384,193791 | 1,952649881 | 0,0564227 | 34,60752263 | 1,95E-262 | 4,87E-260 |

| | | | | | | | | |
|---|---|---|---|---|---|---|---|---|
| ENSG00000060558.3 | GNA15 | protein_coding | 574,2520914 | 0,279843001 | 0,107169498 | 2,61121873 | 0,009022017 | 0,027409445 |
| ENSG00000060688.13 | SNRNP40 | protein_coding | 5755,081114 | -0,143646087 | 0,04844726 | -2,9649992 | 0,003026837 | 0,010441544 |
| ENSG00000060749.15 | QSER1 | protein_coding | 8114,053555 | -0,383068602 | 0,046230464 | -8,286064379 | 1,17E-16 | 1,96E-15 |
| ENSG00000060762.19 | MPC1 | protein_coding | 3530,438241 | -0,222523271 | 0,054790847 | -4,061321996 | 4,88E-05 | 0,000244641 |
| ENSG00000060971.18 | ACAA1 | protein_coding | 2541,510566 | 0,455649142 | 0,062492225 | 7,291293302 | 3,07E-13 | 4,05E-12 |
| ENSG00000060982.15 | BCAT1 | protein_coding | 760,9140168 | -0,711737996 | 0,119897437 | -5,93622362 | 2,92E-09 | 2,58E-08 |
| ENSG00000061273.18 | HDAC7 | protein_coding | 207,5055293 | 0,820247088 | 0,200629245 | 4,088372495 | 4,34E-05 | 0,00022008 |
| ENSG00000061337.15 | LZTS1 | protein_coding | 906,9844517 | -0,47522442 | 0,090108193 | -5,273931333 | 1,34E-07 | 9,72E-07 |
| ENSG00000061794.13 | MRPS35 | protein_coding | 10195,24594 | -0,135998354 | 0,047627933 | -2,855432615 | 0,004297824 | 0,014283143 |
| ENSG00000061938.18 | TNK2 | protein_coding | 3115,807348 | 0,33674185 | 0,068490848 | 4,916596294 | 8,81E-07 | 5,76E-06 |
| ENSG00000062194.16 | GPBP1 | protein_coding | 16278,04012 | 0,444082348 | 0,048216323 | 9,210207669 | 3,25E-20 | 6,68E-19 |
| ENSG00000062282.15 | DGAT2 | protein_coding | 2179,849355 | -0,301414309 | 0,065318119 | -4,614558911 | 3,94E-06 | 2,34E-05 |
| ENSG00000062370.16 | ZNF112 | protein_coding | 2733,76912 | 0,267609873 | 0,061894109 | 4,32367273 | 1,53E-05 | 8,34E-05 |
| ENSG00000062485.19 | CS | protein_coding | 19384,79084 | -0,283695419 | 0,039692122 | -7,147398719 | 8,84E-13 | 1,12E-11 |
| ENSG00000062582.14 | MRPS24 | protein_coding | 2401,936428 | -0,435469571 | 0,075708131 | -5,751952489 | 8,82E-09 | 7,38E-08 |
| ENSG00000062650.19 | WAPL | protein_coding | 12213,4978 | 0,16652716 | 0,042722538 | 3,897876101 | 9,70E-05 | 0,000462672 |
| ENSG00000062725.10 | APPBP2 | protein_coding | 8808,876765 | 0,207882567 | 0,057260113 | 3,630495219 | 0,000282878 | 0,001233734 |
| ENSG00000063046.17 | EIF4B | protein_coding | 44302,02109 | -0,612310213 | 0,043750284 | -13,99557104 | 1,66E-44 | 8,45E-43 |
| ENSG00000063177.13 | RPL18 | protein_coding | 23692,68796 | -0,367710138 | 0,051314291 | -7,165842705 | 7,73E-13 | 9,87E-12 |
| ENSG00000063244.12 | U2AF2 | protein_coding | 22250,375 | -0,134754254 | 0,045571843 | -2,956962985 | 0,003106854 | 0,010683193 |
| ENSG00000063587.14 | ZNF275 | protein_coding | 1666,190332 | 0,278683613 | 0,0672586 | 4,143464377 | 3,42E-05 | 0,000175926 |
| ENSG00000063660.9 | GPC1 | protein_coding | 3126,172005 | -0,577972889 | 0,063942135 | -9,038998931 | 1,58E-19 | 3,12E-18 |
| ENSG00000063761.16 | ADCK1 | protein_coding | 752,3753392 | 0,351314969 | 0,10135983 | 3,466017752 | 0,000528228 | 0,00217644 |
| ENSG00000063978.16 | RNF4 | protein_coding | 6479,611048 | 0,147514218 | 0,059711973 | 2,470429459 | 0,013495093 | 0,038740091 |
| ENSG00000064012.21 | CASP8 | protein_coding | 1528,472759 | 2,464376708 | 0,096954607 | 25,41784009 | 1,60E-142 | 2,52E-140 |
| ENSG00000064042.18 | LIMCH1 | protein_coding | 4701,581379 | 0,142783146 | 0,051360196 | 2,780035077 | 0,005435303 | 0,017567954 |
| ENSG00000064102.15 | INTS13 | protein_coding | 4735,883885 | 0,127297769 | 0,049057965 | 2,594844072 | 0,009463384 | 0,028565283 |
| ENSG00000064115.11 | TM7SF3 | protein_coding | 10893,4121 | -0,315591647 | 0,047231351 | -6,681825483 | 2,36E-11 | 2,61E-10 |
| ENSG00000064199.6 | SPA17 | protein_coding | 1840,228977 | -0,435368961 | 0,075199641 | -5,789508499 | 7,06E-09 | 5,98E-08 |
| ENSG00000064201.15 | TSPAN32 | protein_coding | 735,4223438 | -0,57513518 | 0,120947071 | -4,755263406 | 1,98E-06 | 1,23E-05 |
| ENSG00000064225.12 | ST3GAL6 | protein_coding | 8441,42429 | -0,250675563 | 0,058856934 | -4,259065977 | 2,05E-05 | 0,000109392 |
| ENSG00000064309.14 | CDON | protein_coding | 5870,008962 | 0,324743881 | 0,055013219 | 5,9030154 | 3,57E-09 | 3,13E-08 |
| ENSG00000064393.16 | HIPK2 | protein_coding | 21382,05765 | -0,78303231 | 0,055541092 | -14,09825186 | 3,89E-45 | 2,01E-43 |
| ENSG00000064490.14 | RFXANK | protein_coding | 2066,881917 | 0,194787337 | 0,06315637 | 3,084207278 | 0,002040953 | 0,007353901 |
| ENSG00000064601.18 | CTSA | protein_coding | 36351,15469 | 0,255980352 | 0,046659968 | 5,486080735 | 4,11E-08 | 3,18E-07 |
| ENSG00000064607.17 | SUGP2 | protein_coding | 19625,83377 | -0,110315565 | 0,044157795 | -2,498212694 | 0,012482128 | 0,036220216 |

| | | | | | | | | |
|---|---|---|---|---|---|---|---|---|
| ENSG00000064651.14 | SLC12A2 | protein_coding | 2976,777385 | -0,280225068 | 0,064071924 | -4,373601561 | 1,22E-05 | 6,76E-05 |
| ENSG00000064726.9 | BTBD1 | protein_coding | 13684,35574 | -0,120245404 | 0,048900081 | -2,459002115 | 0,01393238 | 0,039868286 |
| ENSG00000064932.16 | SBNO2 | protein_coding | 1174,459281 | 0,31351955 | 0,08441621 | 3,713973309 | 0,00020403 | 0,000912481 |
| ENSG00000064995.17 | TAF11 | protein_coding | 4750,199676 | -0,138311747 | 0,056842762 | -2,433234092 | 0,014964623 | 0,042332373 |
| ENSG00000064999.15 | ANKS1A | protein_coding | 4655,925791 | 0,153629744 | 0,05398408 | 2,845834248 | 0,004429524 | 0,014642997 |
| ENSG00000065057.9 | NTHL1 | protein_coding | 870,9176019 | -0,319282907 | 0,087245797 | -3,659579241 | 0,00025263 | 0,001113242 |
| ENSG00000065060.17 | UHRF1BP1 | protein_coding | 8490,592798 | 0,732400611 | 0,046044898 | 15,90622715 | 5,74E-57 | 3,84E-55 |
| ENSG00000065150.20 | IPO5 | protein_coding | 22024,30651 | -0,428983873 | 0,050317952 | -8,525463698 | 1,52E-17 | 2,70E-16 |
| ENSG00000065243.20 | PKN2 | protein_coding | 9692,276671 | 0,242268552 | 0,046856945 | 5,170387328 | 2,34E-07 | 1,65E-06 |
| ENSG00000065308.5 | TRAM2 | protein_coding | 3663,066583 | 0,325394541 | 0,052246725 | 6,228037111 | 4,72E-10 | 4,60E-09 |
| ENSG00000065361.15 | ERBB3 | protein_coding | 15918,5886 | 0,348100636 | 0,053248267 | 6,537313901 | 6,26E-11 | 6,66E-10 |
| ENSG00000065427.14 | KARS | protein_coding | 11693,05133 | 0,227988332 | 0,050013179 | 4,558565082 | 5,15E-06 | 3,01E-05 |
| ENSG00000065485.20 | PDIA5 | protein_coding | 908,2385798 | 0,629933281 | 0,132577533 | 4,75143311 | 2,02E-06 | 1,25E-05 |
| ENSG00000065518.8 | NDUFB4 | protein_coding | 14586,88435 | -0,165319096 | 0,046003568 | -3,593614651 | 0,000326122 | 0,001407339 |
| ENSG00000065526.11 | SPEN | protein_coding | 15474,22029 | 0,353778658 | 0,06170011 | 5,733841647 | 9,82E-09 | 8,16E-08 |
| ENSG00000065534.18 | MYLK | protein_coding | 602,5294294 | -0,972769378 | 0,121177959 | -8,027609831 | 9,94E-16 | 1,57E-14 |
| ENSG00000065609.14 | SNAP91 | protein_coding | 2443,355411 | -0,235869008 | 0,067613781 | -3,488475334 | 0,000485784 | 0,002019299 |
| ENSG00000065621.15 | GSTO2 | protein_coding | 2188,260284 | -0,519553474 | 0,075303584 | -6,899452122 | 5,22E-12 | 6,16E-11 |
| ENSG00000065675.14 | PRKCQ | protein_coding | 844,8366039 | 0,71895026 | 0,102222986 | 7,03315655 | 2,02E-12 | 2,49E-11 |
| ENSG00000065802.12 | ASB1 | protein_coding | 5956,340245 | -0,818465754 | 0,058325605 | -14,03270055 | 9,83E-45 | 5,03E-43 |
| ENSG00000065809.13 | FAM107B | protein_coding | 10319,80099 | -0,285230643 | 0,05126902 | -5,563411264 | 2,65E-08 | 2,10E-07 |
| ENSG00000065883.16 | CDK13 | protein_coding | 6020,530711 | 0,198324803 | 0,051117178 | 3,879807334 | 0,000104539 | 0,000495488 |
| ENSG00000065911.12 | MTHFD2 | protein_coding | 6771,066682 | 0,196602687 | 0,054843094 | 3,584821246 | 0,000337309 | 0,001450333 |
| ENSG00000065970.9 | FOXJ2 | protein_coding | 2819,099043 | 0,177217398 | 0,056381379 | 3,143190223 | 0,001671172 | 0,00616318 |
| ENSG00000065978.19 | YBX1 | protein_coding | 38492,53013 | -0,491549247 | 0,044765762 | -10,98047317 | 4,74E-28 | 1,39E-26 |
| ENSG00000066032.18 | CTNNA2 | protein_coding | 2587,443107 | -0,458213363 | 0,065719195 | -6,972291179 | 3,12E-12 | 3,76E-11 |
| ENSG00000066044.15 | ELAVL1 | protein_coding | 8140,05644 | -0,290475046 | 0,04855715 | -5,982127121 | 2,20E-09 | 1,98E-08 |
| ENSG00000066117.14 | SMARCD1 | protein_coding | 8295,55704 | 0,296434619 | 0,048633316 | 6,095299359 | 1,09E-09 | 1,02E-08 |
| ENSG00000066135.13 | KDM4A | protein_coding | 8228,848988 | 0,199676719 | 0,047438767 | 4,209146431 | 2,56E-05 | 0,000134685 |
| ENSG00000066279.18 | ASPM | protein_coding | 11589,42318 | -0,716145804 | 0,055460274 | -12,9127707 | 3,81E-38 | 1,61E-36 |
| ENSG00000066379.15 | ZNRD1 | protein_coding | 1302,642765 | 0,191203669 | 0,079484519 | 2,405546042 | 0,016148317 | 0,045227693 |
| ENSG00000066468.22 | FGFR2 | protein_coding | 200,9084227 | -0,50304386 | 0,202005897 | -2,490243438 | 0,012765562 | 0,036910224 |
| ENSG00000066557.6 | LRRC40 | protein_coding | 4739,199242 | -0,144654715 | 0,056213148 | -2,573325288 | 0,010072649 | 0,030121596 |
| ENSG00000066583.12 | ISOC1 | protein_coding | 4691,205277 | -0,188668509 | 0,050819909 | -3,712492041 | 0,000205229 | 0,000917658 |
| ENSG00000066654.14 | THUMPD1 | protein_coding | 15578,85025 | 0,259466954 | 0,040610511 | 6,389157628 | 1,67E-10 | 1,70E-09 |
| ENSG00000066697.14 | MSANTD3 | protein_coding | 2190,51678 | -0,237193787 | 0,068520382 | -3,46165302 | 0,000536869 | 0,002210018 |

| | | | | | | | | |
|---|---|---|---|---|---|---|---|---|
| ENSG00000066777.9 | ARFGEF1 | protein_coding | 10576,05232 | 0,137222665 | 0,053095443 | 2,584452777 | 0,009753366 | 0,029283515 |
| ENSG00000066926.11 | FECH | protein_coding | 2918,284234 | -0,203204741 | 0,059137521 | -3,436138993 | 0,000590068 | 0,002408313 |
| ENSG00000066933.16 | MYO9A | protein_coding | 3159,205875 | 0,417783919 | 0,073546089 | 5,68057287 | 1,34E-08 | 1,10E-07 |
| ENSG00000067048.17 | DDX3Y | protein_coding | 12525,5394 | 0,16795127 | 0,045203664 | 3,715434894 | 0,000202854 | 0,00090806 |
| ENSG00000067057.17 | PFKP | protein_coding | 4118,334246 | 0,328199378 | 0,070500868 | 4,655252982 | 3,24E-06 | 1,95E-05 |
| ENSG00000067066.17 | SP100 | protein_coding | 5795,857227 | 9,722493456 | 0,291744495 | 33,32537075 | 1,66E-243 | 3,85E-241 |
| ENSG00000067082.15 | KLF6 | protein_coding | 2705,611781 | 0,473025846 | 0,062036725 | 7,62493257 | 2,44E-14 | 3,48E-13 |
| ENSG00000067113.17 | PLPP1 | protein_coding | 688,4241093 | 0,225057175 | 0,094900137 | 2,371515812 | 0,017715289 | 0,048932806 |
| ENSG00000067141.17 | NEO1 | protein_coding | 36794,83059 | -0,138752753 | 0,042257976 | -3,283468993 | 0,001025379 | 0,003962957 |
| ENSG00000067167.8 | TRAM1 | protein_coding | 16021,92851 | 0,296879766 | 0,042263466 | 7,024501232 | 2,15E-12 | 2,64E-11 |
| ENSG00000067182.7 | TNFRSF1A | protein_coding | 440,2534502 | 0,508961838 | 0,147760078 | 3,444515221 | 0,000572084 | 0,002343412 |
| ENSG00000067191.16 | CACNB1 | protein_coding | 400,3279909 | 0,397874432 | 0,144767701 | 2,748364641 | 0,005989336 | 0,019133404 |
| ENSG00000067221.14 | STOML1 | protein_coding | 2095,116454 | 0,177200413 | 0,069730704 | 2,541210739 | 0,011046931 | 0,032580631 |
| ENSG00000067365.14 | METTL22 | protein_coding | 2146,859601 | 0,201697306 | 0,070153745 | 2,875075404 | 0,00403931 | 0,013519827 |
| ENSG00000067445.20 | TRO | protein_coding | 8764,796944 | 0,24277189 | 0,053443112 | 4,542622668 | 5,56E-06 | 3,23E-05 |
| ENSG00000067533.6 | RRP15 | protein_coding | 3240,905598 | -0,239401122 | 0,083770752 | -2,85781273 | 0,004265721 | 0,014191118 |
| ENSG00000067560.12 | RHOA | protein_coding | 36951,11208 | -0,296356264 | 0,052357093 | -5,660288693 | 1,51E-08 | 1,23E-07 |
| ENSG00000067646.12 | ZFY | protein_coding | 3434,086201 | 0,187073447 | 0,055690998 | 3,35913259 | 0,000781875 | 0,003105017 |
| ENSG00000067704.10 | IARS2 | protein_coding | 18096,05928 | -0,279630075 | 0,045317544 | -6,170459604 | 6,81E-10 | 6,48E-09 |
| ENSG00000067840.12 | PDZD4 | protein_coding | 2448,117987 | -0,209901845 | 0,080373474 | -2,611581104 | 0,009012461 | 0,027387806 |
| ENSG00000067955.15 | CBFB | protein_coding | 11712,68585 | -0,157214022 | 0,049286169 | -3,189820259 | 0,001423613 | 0,005330057 |
| ENSG00000067992.15 | PDK3 | protein_coding | 9362,559338 | -0,683105885 | 0,052036064 | -13,12754718 | 2,29E-39 | 1,01E-37 |
| ENSG00000068024.16 | HDAC4 | protein_coding | 2921,686304 | -0,281488769 | 0,099317555 | -2,834229753 | 0,004593629 | 0,015127725 |
| ENSG00000068078.18 | FGFR3 | protein_coding | 10293,32409 | -0,312346658 | 0,05227788 | -5,974738444 | 2,30E-09 | 2,06E-08 |
| ENSG00000068079.7 | IFI35 | protein_coding | 14702,90044 | 5,704798065 | 0,07382253 | 77,27719561 | 0 | 0 |
| ENSG00000068120.15 | COASY | protein_coding | 6908,566761 | 0,160827206 | 0,055134929 | 2,916974935 | 0,003534442 | 0,012004942 |
| ENSG00000068305.17 | MEF2A | protein_coding | 8144,281218 | 0,177305901 | 0,058951914 | 3,007636034 | 0,002632883 | 0,009225335 |
| ENSG00000068308.13 | OTUD5 | protein_coding | 5945,096357 | 0,16528024 | 0,052069364 | 3,174231939 | 0,001502336 | 0,005607043 |
| ENSG00000068323.17 | TFE3 | protein_coding | 6984,603214 | 0,232916337 | 0,050477282 | 4,614280462 | 3,94E-06 | 2,35E-05 |
| ENSG00000068366.19 | ACSL4 | protein_coding | 6801,203497 | 0,136463061 | 0,04813972 | 2,834729046 | 0,004586457 | 0,015106315 |
| ENSG00000068383.19 | INPP5A | protein_coding | 856,8812255 | -0,321655753 | 0,088754639 | -3,624100746 | 0,000289969 | 0,001263434 |
| ENSG00000068394.11 | GPKOW | protein_coding | 3593,369854 | 0,265960014 | 0,057768668 | 4,603879991 | 4,15E-06 | 2,46E-05 |
| ENSG00000068400.13 | GRIPAP1 | protein_coding | 15185,79208 | 0,48623811 | 0,056114043 | 8,665176894 | 4,51E-18 | 8,25E-17 |
| ENSG00000068489.12 | PRR11 | protein_coding | 3133,8975 | -1,712243284 | 0,074262212 | -23,05672349 | 1,26E-117 | 1,69E-115 |
| ENSG00000068615.18 | REEP1 | protein_coding | 10762,98504 | -0,881272838 | 0,055465789 | -15,88858369 | 7,60E-57 | 5,03E-55 |
| ENSG00000068650.18 | ATP11A | protein_coding | 12032,56543 | -0,119132774 | 0,046962704 | -2,536752877 | 0,01118859 | 0,032942313 |

| | | | | | | | | |
|---|---|---|---|---|---|---|---|---|
| ENSG00000068654.16 | POLR1A | protein_coding | 6022,252976 | -0,318572153 | 0,081848219 | -3,892230713 | 9,93E-05 | 0,000472673 |
| ENSG00000068697.7 | LAPTM4A | protein_coding | 30988,28638 | 0,339821281 | 0,042614129 | 7,974380452 | 1,53E-15 | 2,38E-14 |
| ENSG00000068796.16 | KIF2A | protein_coding | 11009,40381 | -0,106810621 | 0,044569655 | -2,396487472 | 0,016553059 | 0,046203981 |
| ENSG00000068878.15 | PSME4 | protein_coding | 14414,34022 | 0,299164993 | 0,044911877 | 6,661155453 | 2,72E-11 | 2,99E-10 |
| ENSG00000068885.15 | IFT80 | protein_coding | 7261,676836 | -0,454305814 | 0,051053959 | -8,898542259 | 5,66E-19 | 1,09E-17 |
| ENSG00000068971.14 | PPP2R5B | protein_coding | 2397,575724 | 0,645367074 | 0,070461247 | 9,159177631 | 5,23E-20 | 1,06E-18 |
| ENSG00000069020.18 | MAST4 | protein_coding | 5530,163022 | 0,501031676 | 0,058061267 | 8,629361672 | 6,17E-18 | 1,12E-16 |
| ENSG00000069275.13 | NUCKS1 | protein_coding | 61053,61544 | -0,54079089 | 0,047563215 | -11,36993976 | 5,90E-30 | 1,85E-28 |
| ENSG00000069399.15 | BCL3 | protein_coding | 154,9443621 | 0,579710779 | 0,200193141 | 2,895757452 | 0,003782446 | 0,012749189 |
| ENSG00000069509.6 | FUNDC1 | protein_coding | 4738,884845 | 0,143184864 | 0,04979119 | 2,875706789 | 0,00403124 | 0,013498837 |
| ENSG00000069535.14 | MAOB | protein_coding | 6744,5358 | 0,483982544 | 0,059064096 | 8,194192024 | 2,52E-16 | 4,13E-15 |
| ENSG00000069667.16 | RORA | protein_coding | 5724,249952 | 0,277763719 | 0,0672411 | 4,130862213 | 3,61E-05 | 0,000185052 |
| ENSG00000069943.10 | PIGB | protein_coding | 881,1027968 | 0,314441149 | 0,104171023 | 3,018508805 | 0,00254022 | 0,008934954 |
| ENSG00000069974.16 | RAB27A | protein_coding | 5111,582133 | -0,184266526 | 0,050181606 | -3,671993418 | 0,000240666 | 0,001064691 |
| ENSG00000070010.19 | UFD1 | protein_coding | 5243,791897 | -0,120253012 | 0,048288422 | -2,49030736 | 0,012763267 | 0,036910224 |
| ENSG00000070018.9 | LRP6 | protein_coding | 7982,56619 | 0,140210392 | 0,048071196 | 2,916723588 | 0,003537291 | 0,012009183 |
| ENSG00000070019.5 | GUCY2C | protein_coding | 28920,06 | 0,123505066 | 0,043346882 | 2,849226072 | 0,004382573 | 0,01450696 |
| ENSG00000070061.15 | ELP1 | protein_coding | 11220,78422 | -0,125325137 | 0,04439906 | -2,822698013 | 0,00476214 | 0,015625513 |
| ENSG00000070081.16 | NUCB2 | protein_coding | 31681,49607 | 0,369406779 | 0,047899834 | 7,712068021 | 1,24E-14 | 1,81E-13 |
| ENSG00000070182.21 | SPTB | protein_coding | 3217,923833 | 0,300370924 | 0,055841134 | 5,379026238 | 7,49E-08 | 5,60E-07 |
| ENSG00000070269.14 | TMEM260 | protein_coding | 4910,936793 | 0,256166552 | 0,050751316 | 5,047485872 | 4,48E-07 | 3,06E-06 |
| ENSG00000070366.14 | SMG6 | protein_coding | 4292,470573 | 0,294897182 | 0,057754465 | 5,106049938 | 3,29E-07 | 2,29E-06 |
| ENSG00000070367.16 | EXOC5 | protein_coding | 18143,02527 | -0,151126311 | 0,041539281 | -3,638154239 | 0,000274599 | 0,001199951 |
| ENSG00000070413.20 | DGCR2 | protein_coding | 12715,31716 | 0,223853134 | 0,055744696 | 4,015684902 | 5,93E-05 | 0,000293486 |
| ENSG00000070444.15 | MNT | protein_coding | 1856,732121 | -0,367747733 | 0,085488342 | -4,301729606 | 1,69E-05 | 9,15E-05 |
| ENSG00000070540.13 | WIPI1 | protein_coding | 782,4418319 | 0,552937632 | 0,100851427 | 5,482695176 | 4,19E-08 | 3,23E-07 |
| ENSG00000070756.16 | PABPC1 | protein_coding | 45065,61254 | -0,554749689 | 0,043506178 | -12,75105553 | 3,08E-37 | 1,26E-35 |
| ENSG00000070778.13 | AL162171.1 | protein_coding | 2161,617352 | 0,461695329 | 0,075898154 | 6,083090324 | 1,18E-09 | 1,09E-08 |
| ENSG00000070831.17 | CDC42 | protein_coding | 45661,33035 | -0,2532364 | 0,045201159 | -5,602431611 | 2,11E-08 | 1,70E-07 |
| ENSG00000070882.13 | OSBPL3 | protein_coding | 338,4500241 | -0,644758475 | 0,17905855 | -3,60082484 | 0,000317209 | 0,001372034 |
| ENSG00000070886.12 | EPHA8 | protein_coding | 241,4724365 | -0,401678168 | 0,165184487 | -2,431694256 | 0,015028387 | 0,042486055 |
| ENSG00000070985.13 | TRPM5 | protein_coding | 237,8071827 | -0,479879539 | 0,163797644 | -2,929709652 | 0,003392789 | 0,011579724 |
| ENSG00000071073.13 | MGAT4A | protein_coding | 34813,12046 | -0,254596801 | 0,042492924 | -5,991510446 | 2,08E-09 | 1,87E-08 |
| ENSG00000071082.11 | RPL31 | protein_coding | 34964,89165 | -0,467547629 | 0,039353275 | -11,88078053 | 1,49E-32 | 5,34E-31 |
| ENSG00000071242.12 | RPS6KA2 | protein_coding | 2669,981999 | -0,257873025 | 0,08051306 | -3,202871982 | 0,001360645 | 0,005118962 |
| ENSG00000071246.11 | VASH1 | protein_coding | 5299,958448 | -0,146803502 | 0,054368995 | -2,700132741 | 0,006931182 | 0,021786617 |

| | | | | | | | | |
|---|---|---|---|---|---|---|---|---|
| ENSG00000071539.14 | TRIP13 | protein_coding | 1971,40658 | -0,490755204 | 0,072569048 | -6,76259666 | 1,36E-11 | 1,53E-10 |
| ENSG00000071564.16 | TCF3 | protein_coding | 3025,709857 | -0,601573979 | 0,056999997 | -10,55393004 | 4,87E-26 | 1,32E-24 |
| ENSG00000071575.11 | TRIB2 | protein_coding | 1391,668037 | 0,28008652 | 0,092863743 | 3,016101971 | 0,002560471 | 0,008999633 |
| ENSG00000071626.16 | DAZAP1 | protein_coding | 4335,6198 | -0,2171269 | 0,052403581 | -4,143360011 | 3,42E-05 | 0,000175926 |
| ENSG00000071655.18 | MBD3 | protein_coding | 6279,731665 | -0,166426081 | 0,061239555 | -2,717623922 | 0,006575253 | 0,020786849 |
| ENSG00000071794.16 | HLTF | protein_coding | 23608,691 | -0,21264131 | 0,046825566 | -4,541136976 | 5,60E-06 | 3,26E-05 |
| ENSG00000071859.15 | FAM50A | protein_coding | 6639,798087 | -0,120358244 | 0,050140545 | -2,400417552 | 0,016376379 | 0,045745337 |
| ENSG00000071894.17 | CPSF1 | protein_coding | 7529,205183 | -0,1795682 | 0,066630793 | -2,694973191 | 0,007039429 | 0,022083696 |
| ENSG00000072062.13 | PRKACA | protein_coding | 8941,171762 | -0,312940607 | 0,048061505 | -6,511252716 | 7,45E-11 | 7,85E-10 |
| ENSG00000072071.16 | ADGRL1 | protein_coding | 10840,66345 | 0,228300746 | 0,046159863 | 4,945871426 | 7,58E-07 | 5,01E-06 |
| ENSG00000072110.13 | ACTN1 | protein_coding | 6330,529657 | 0,138771145 | 0,055692135 | 2,491754793 | 0,012711376 | 0,036776396 |
| ENSG00000072121.16 | ZFYVE26 | protein_coding | 8794,111552 | 0,981401489 | 0,269319948 | 3,643998505 | 0,000268435 | 0,001176441 |
| ENSG00000072133.11 | RPS6KA6 | protein_coding | 5551,469087 | -0,358491146 | 0,0526385 | -6,810436207 | 9,73E-12 | 1,12E-10 |
| ENSG00000072163.19 | LIMS2 | protein_coding | 70,95169939 | -1,384643091 | 0,295183204 | -4,690792263 | 2,72E-06 | 1,66E-05 |
| ENSG00000072195.15 | SPEG | protein_coding | 6876,22255 | -0,130509084 | 0,049952677 | -2,612654464 | 0,008984208 | 0,027309324 |
| ENSG00000072274.13 | TFRC | protein_coding | 21791,99613 | -0,865885767 | 0,156370298 | -5,537405626 | 3,07E-08 | 2,42E-07 |
| ENSG00000072364.13 | AFF4 | protein_coding | 25300,2515 | 0,154947072 | 0,042562128 | 3,640491618 | 0,000272118 | 0,001190959 |
| ENSG00000072501.17 | SMC1A | protein_coding | 42137,87498 | -0,347939847 | 0,054415297 | -6,394155065 | 1,61E-10 | 1,65E-09 |
| ENSG00000072518.20 | MARK2 | protein_coding | 8647,202994 | 0,400760115 | 0,059901247 | 6,690346764 | 2,23E-11 | 2,47E-10 |
| ENSG00000072571.20 | HMMR | protein_coding | 6752,035028 | -1,133335189 | 0,056748427 | -19,97121762 | 9,80E-89 | 1,02E-86 |
| ENSG00000072609.17 | CHFR | protein_coding | 3652,907625 | 0,222327715 | 0,057890066 | 3,840515826 | 0,000122776 | 0,000575503 |
| ENSG00000072682.18 | P4HA2 | protein_coding | 7370,796857 | 0,435583326 | 0,048136339 | 9,048950059 | 1,44E-19 | 2,85E-18 |
| ENSG00000072736.19 | NFATC3 | protein_coding | 10476,03534 | 0,162381987 | 0,044819238 | 3,623042111 | 0,000291158 | 0,001267925 |
| ENSG00000072756.17 | TRNT1 | protein_coding | 3038,763453 | 0,237946147 | 0,060308704 | 3,94546943 | 7,96E-05 | 0,000386025 |
| ENSG00000072786.13 | STK10 | protein_coding | 1255,676205 | 0,287884441 | 0,095961044 | 3,000013639 | 0,002699675 | 0,009431474 |
| ENSG00000072803.17 | FBXW11 | protein_coding | 13206,14129 | -0,104370714 | 0,04402246 | -2,370851466 | 0,017747162 | 0,048996794 |
| ENSG00000072832.14 | CRMP1 | protein_coding | 29028,65597 | -0,163649029 | 0,043243193 | -3,784388179 | 0,000154087 | 0,000707971 |
| ENSG00000072849.11 | DERL2 | protein_coding | 4933,103983 | -0,115195845 | 0,04863831 | -2,368417914 | 0,017864344 | 0,049284045 |
| ENSG00000072858.10 | SIDT1 | protein_coding | 256,1584989 | 3,720383298 | 0,221625336 | 16,78681401 | 3,05E-63 | 2,22E-61 |
| ENSG00000072864.15 | NDE1 | protein_coding | 1858,20039 | -0,18321202 | 0,066569737 | -2,752181801 | 0,005919965 | 0,018933288 |
| ENSG00000072954.7 | TMEM38A | protein_coding | 1604,243257 | -0,304551272 | 0,084978034 | -3,583882292 | 0,000338525 | 0,001455281 |
| ENSG00000072958.8 | AP1M1 | protein_coding | 5230,88403 | -0,185925361 | 0,064408737 | -2,886648165 | 0,003893694 | 0,013079115 |
| ENSG00000073008.15 | PVR | protein_coding | 10318,71693 | 0,134731745 | 0,0507065 | 2,657090229 | 0,007881834 | 0,024353083 |
| ENSG00000073067.14 | CYP2W1 | protein_coding | 14,05634498 | -1,687994051 | 0,640328802 | -2,636136381 | 0,008385602 | 0,025708461 |
| ENSG00000073150.13 | PANX2 | protein_coding | 75,16699011 | -0,847911534 | 0,273296117 | -3,102537805 | 0,00191869 | 0,006963528 |
| ENSG00000073331.18 | ALPK1 | protein_coding | 129,4248772 | 0,915927851 | 0,268579372 | 3,410268796 | 0,000648989 | 0,002625937 |

| | | | | | | | | |
|---|---|---|---|---|---|---|---|---|
| ENSG00000073417.15 | PDE8A | protein_coding | 4975,209374 | 0,167268849 | 0,053649114 | 3,117830567 | 0,001821875 | 0,006655077 |
| ENSG00000073464.12 | CLCN4 | protein_coding | 11525,94393 | -0,455231415 | 0,053718114 | -8,474448888 | 2,36E-17 | 4,12E-16 |
| ENSG00000073584.20 | SMARCE1 | protein_coding | 28006,18993 | -0,20489967 | 0,041544046 | -4,932106795 | 8,13E-07 | 5,35E-06 |
| ENSG00000073605.18 | GSDMB | protein_coding | 589,8875406 | 1,573273018 | 0,283580417 | 5,547890201 | 2,89E-08 | 2,29E-07 |
| ENSG00000073614.12 | KDM5A | protein_coding | 12240,89779 | 0,282355201 | 0,041817063 | 6,75215286 | 1,46E-11 | 1,64E-10 |
| ENSG00000073712.15 | FERMT2 | protein_coding | 6082,422774 | 0,258414871 | 0,047919519 | 5,392684973 | 6,94E-08 | 5,21E-07 |
| ENSG00000073756.12 | PTGS2 | protein_coding | 68082,7098 | 0,533582373 | 0,043143402 | 12,36764709 | 3,91E-35 | 1,50E-33 |
| ENSG00000073803.14 | MAP3K13 | protein_coding | 4844,267928 | -0,290511014 | 0,053226317 | -5,458033354 | 4,81E-08 | 3,70E-07 |
| ENSG00000073910.21 | FRY | protein_coding | 3634,415308 | 0,716866321 | 0,069436176 | 10,32410434 | 5,48E-25 | 1,43E-23 |
| ENSG00000073921.18 | PICALM | protein_coding | 19680,63801 | -0,135821726 | 0,045938731 | -2,956584217 | 0,003110673 | 0,010693056 |
| ENSG00000074071.14 | MRPS34 | protein_coding | 4839,280483 | -0,287330504 | 0,064403275 | -4,461426903 | 8,14E-06 | 4,62E-05 |
| ENSG00000074201.8 | CLNS1A | protein_coding | 8203,085339 | -0,229375176 | 0,049985324 | -4,588850408 | 4,46E-06 | 2,63E-05 |
| ENSG00000074211.14 | PPP2R2C | protein_coding | 1001,576532 | -1,481540014 | 0,114861794 | -12,89845794 | 4,59E-38 | 1,93E-36 |
| ENSG00000074266.20 | EED | protein_coding | 4214,006401 | 0,164763406 | 0,055867196 | 2,94919768 | 0,003186001 | 0,010931966 |
| ENSG00000074276.10 | CDHR2 | protein_coding | 13573,29954 | 0,469750824 | 0,056937918 | 8,250228366 | 1,58E-16 | 2,63E-15 |
| ENSG00000074317.11 | SNCB | protein_coding | 285,2666314 | -0,525808466 | 0,165643817 | -3,174331983 | 0,001501818 | 0,00560604 |
| ENSG00000074356.17 | NCBP3 | protein_coding | 9221,879522 | -0,155541786 | 0,055463845 | -2,804381583 | 0,00504132 | 0,016433755 |
| ENSG00000074370.18 | ATP2A3 | protein_coding | 69118,77506 | -0,21240738 | 0,050469381 | -4,208638474 | 2,57E-05 | 0,000134925 |
| ENSG00000074582.14 | BCS1L | protein_coding | 2948,541412 | -0,213844609 | 0,056610768 | -3,777454653 | 0,000158439 | 0,00072545 |
| ENSG00000074590.13 | NUAK1 | protein_coding | 2412,802238 | 0,536373084 | 0,07912166 | 6,779092918 | 1,21E-11 | 1,38E-10 |
| ENSG00000074696.13 | HACD3 | protein_coding | 44804,20352 | -0,27414872 | 0,048642889 | -5,635946521 | 1,74E-08 | 1,41E-07 |
| ENSG00000074755.15 | ZZEF1 | protein_coding | 9308,18556 | 0,529381164 | 0,057040013 | 9,280873913 | 1,68E-20 | 3,52E-19 |
| ENSG00000074800.16 | ENO1 | protein_coding | 82699,8715 | -0,145568689 | 0,056605234 | -2,571647133 | 0,0101216 | 0,030235499 |
| ENSG00000074803.19 | SLC12A1 | protein_coding | 568,5183263 | -0,918729832 | 0,142319224 | -6,455416235 | 1,08E-10 | 1,12E-09 |
| ENSG00000074842.7 | MYDGF | protein_coding | 7849,840271 | 0,58460393 | 0,056403717 | 10,3646349 | 3,59E-25 | 9,39E-24 |
| ENSG00000074935.14 | TUBE1 | protein_coding | 2148,615285 | 0,702063101 | 0,080048874 | 8,77043069 | 1,78E-18 | 3,34E-17 |
| ENSG00000075035.10 | WSCD2 | protein_coding | 392,2807989 | 0,805191495 | 0,176494027 | 4,562145877 | 5,06E-06 | 2,96E-05 |
| ENSG00000075043.18 | KCNQ2 | protein_coding | 4503,458023 | -0,241959888 | 0,080453883 | -3,007435789 | 0,002634618 | 0,009228542 |
| ENSG00000075188.9 | NUP37 | protein_coding | 2344,92652 | -0,418791392 | 0,064090565 | -6,534368903 | 6,39E-11 | 6,78E-10 |
| ENSG00000075213.11 | SEMA3A | protein_coding | 10591,78269 | 0,364932541 | 0,046861785 | 7,787422971 | 6,84E-15 | 1,02E-13 |
| ENSG00000075218.19 | GTSE1 | protein_coding | 3452,164475 | -0,471453821 | 0,057174024 | -8,245944388 | 1,64E-16 | 2,72E-15 |
| ENSG00000075234.17 | TTC38 | protein_coding | 5555,646503 | 0,356464102 | 0,059881802 | 5,952795167 | 2,64E-09 | 2,35E-08 |
| ENSG00000075239.13 | ACAT1 | protein_coding | 6771,636785 | -0,183557792 | 0,053516184 | -3,429949174 | 0,000603694 | 0,002457847 |
| ENSG00000075303.13 | SLC25A40 | protein_coding | 4953,884789 | -0,151476109 | 0,057113206 | -2,652208136 | 0,007996723 | 0,024670811 |
| ENSG00000075336.12 | TIMM21 | protein_coding | 3199,394209 | -0,418766594 | 0,063865721 | -6,556985337 | 5,49E-11 | 5,87E-10 |
| ENSG00000075391.16 | RASAL2 | protein_coding | 4416,663782 | 0,275772661 | 0,057107584 | 4,829002439 | 1,37E-06 | 8,73E-06 |

| | | | | | | | | |
|---|---|---|---|---|---|---|---|---|
| ENSG00000075407.18 | ZNF37A | protein_coding | 8431,505747 | 0,135467978 | 0,045207203 | 2,996601642 | 0,002730072 | 0,00952584 |
| ENSG00000075413.18 | MARK3 | protein_coding | 11473,70679 | 0,147140649 | 0,050415806 | 2,918542025 | 0,003516725 | 0,011951979 |
| ENSG00000075415.12 | SLC25A3 | protein_coding | 24088,34166 | -0,549947389 | 0,044145363 | -12,45764781 | 1,27E-35 | 5,01E-34 |
| ENSG00000075420.13 | FNDC3B | protein_coding | 6825,31201 | 0,517730565 | 0,04986025 | 10,38363364 | 2,94E-25 | 7,71E-24 |
| ENSG00000075426.12 | FOSL2 | protein_coding | 1822,447009 | 0,811203829 | 0,072025627 | 11,2627111 | 2,00E-29 | 6,15E-28 |
| ENSG00000075429.8 | CACNG5 | protein_coding | 208,9082697 | -0,404495714 | 0,163146251 | -2,47934421 | 0,013162422 | 0,037925364 |
| ENSG00000075539.14 | FRYL | protein_coding | 9731,468569 | 0,405838652 | 0,052747074 | 7,694050494 | 1,43E-14 | 2,07E-13 |
| ENSG00000075618.18 | FSCN1 | protein_coding | 101,4133242 | -0,581055172 | 0,240885312 | -2,412165218 | 0,015858094 | 0,044501911 |
| ENSG00000075624.16 | ACTB | protein_coding | 59608,07972 | -0,280282838 | 0,047689824 | -5,877204289 | 4,17E-09 | 3,64E-08 |
| ENSG00000075643.6 | MOCOS | protein_coding | 196,60207 | 1,530363574 | 0,192448316 | 7,952075685 | 1,83E-15 | 2,85E-14 |
| ENSG00000075651.16 | PLD1 | protein_coding | 3011,142351 | 0,359642136 | 0,064322205 | 5,591259431 | 2,25E-08 | 1,81E-07 |
| ENSG00000075702.18 | WDR62 | protein_coding | 3215,44599 | -0,173325915 | 0,058488093 | -2,963439324 | 0,00304222 | 0,010490076 |
| ENSG00000075711.21 | DLG1 | protein_coding | 14468,34891 | -0,329716724 | 0,045718086 | -7,211953846 | 5,52E-13 | 7,12E-12 |
| ENSG00000076003.5 | MCM6 | protein_coding | 33417,32388 | -0,296890821 | 0,042601007 | -6,969103398 | 3,19E-12 | 3,85E-11 |
| ENSG00000076043.10 | REXO2 | protein_coding | 5170,068748 | -0,209664664 | 0,052728707 | -3,976290655 | 7,00E-05 | 0,000342521 |
| ENSG00000076053.10 | RBM7 | protein_coding | 6166,211514 | 0,551157299 | 0,058993682 | 9,342649544 | 9,40E-21 | 2,00E-19 |
| ENSG00000076108.11 | BAZ2A | protein_coding | 19879,65666 | 0,485515632 | 0,049250634 | 9,858058437 | 6,33E-23 | 1,49E-21 |
| ENSG00000076248.10 | UNG | protein_coding | 6706,6892 | -0,386605482 | 0,052784649 | -7,324202966 | 2,40E-13 | 3,20E-12 |
| ENSG00000076321.11 | KLHL20 | protein_coding | 2646,421741 | 0,242666736 | 0,066045119 | 3,674256958 | 0,000238543 | 0,001056128 |
| ENSG00000076344.16 | RGS11 | protein_coding | 82,40442004 | -1,635829919 | 0,294469699 | -5,555172323 | 2,77E-08 | 2,20E-07 |
| ENSG00000076382.17 | SPAG5 | protein_coding | 12439,9881 | -0,599026984 | 0,048174623 | -12,43449248 | 1,70E-35 | 6,66E-34 |
| ENSG00000076706.17 | AP002956.1 | protein_coding | 112,0115163 | 1,248491983 | 0,287783963 | 4,338295888 | 1,44E-05 | 7,84E-05 |
| ENSG00000076770.14 | MBNL3 | protein_coding | 933,969168 | -0,746135711 | 0,11753603 | -6,34814456 | 2,18E-10 | 2,20E-09 |
| ENSG00000076864.19 | RAP1GAP | protein_coding | 9818,409275 | -0,396785961 | 0,049999386 | -7,935816619 | 2,09E-15 | 3,23E-14 |
| ENSG00000076924.12 | XAB2 | protein_coding | 3654,019946 | 0,465277415 | 0,057364726 | 8,110862625 | 5,03E-16 | 8,09E-15 |
| ENSG00000077063.11 | CTTNBP2 | protein_coding | 3262,839073 | 0,720494168 | 0,074827189 | 9,628775014 | 6,04E-22 | 1,36E-20 |
| ENSG00000077080.9 | ACTL6B | protein_coding | 3607,539062 | -0,389156298 | 0,065775704 | -5,916413995 | 3,29E-09 | 2,90E-08 |
| ENSG00000077097.15 | TOP2B | protein_coding | 23548,04452 | -0,116609573 | 0,046424938 | -2,51178738 | 0,012012144 | 0,035018954 |
| ENSG00000077147.16 | TM9SF3 | protein_coding | 35547,91989 | 0,21667773 | 0,054081042 | 4,006537672 | 6,16E-05 | 0,000304545 |
| ENSG00000077150.20 | NFKB2 | protein_coding | 1700,895512 | 1,072520077 | 0,098681642 | 10,86848636 | 1,63E-27 | 4,66E-26 |
| ENSG00000077152.11 | UBE2T | protein_coding | 7611,517236 | -0,393191172 | 0,046539298 | -8,448584015 | 2,95E-17 | 5,10E-16 |
| ENSG00000077157.22 | PPP1R12B | protein_coding | 3873,458345 | 0,837357356 | 0,192578121 | 4,348143755 | 1,37E-05 | 7,53E-05 |
| ENSG00000077232.18 | DNAJC10 | protein_coding | 17114,42603 | 0,329340826 | 0,050637643 | 6,503873518 | 7,83E-11 | 8,23E-10 |
| ENSG00000077254.14 | USP33 | protein_coding | 18410,69224 | 0,304211467 | 0,043128826 | 7,053553219 | 1,74E-12 | 2,16E-11 |
| ENSG00000077264.15 | PAK3 | protein_coding | 3728,559202 | 0,154452874 | 0,058298283 | 2,649355467 | 0,008064546 | 0,024859609 |
| ENSG00000077279.18 | DCX | protein_coding | 20184,12902 | -0,563800309 | 0,056824483 | -9,921785066 | 3,35E-23 | 7,96E-22 |

| | | | | | | | | |
|---|---|---|---|---|---|---|---|---|
| ENSG00000077312.9 | SNRPA | protein_coding | 4565,284688 | -0,128581767 | 0,051670619 | -2,488488993 | 0,012828721 | 0,037066592 |
| ENSG00000077348.9 | EXOSC5 | protein_coding | 763,9681497 | -0,600033569 | 0,099387822 | -6,037294664 | 1,57E-09 | 1,43E-08 |
| ENSG00000077380.15 | DYNC1I2 | protein_coding | 18357,94449 | -0,128636412 | 0,051616838 | -2,492140469 | 0,012697581 | 0,036745931 |
| ENSG00000077458.13 | FAM76B | protein_coding | 2950,664229 | 0,159731211 | 0,062431224 | 2,55851483 | 0,010512034 | 0,031228107 |
| ENSG00000077514.8 | POLD3 | protein_coding | 5412,130192 | -0,247111164 | 0,049862397 | -4,95586215 | 7,20E-07 | 4,77E-06 |
| ENSG00000077522.13 | ACTN2 | protein_coding | 832,184772 | -0,395796655 | 0,099613868 | -3,973308774 | 7,09E-05 | 0,000346613 |
| ENSG00000077549.18 | CAPZB | protein_coding | 16124,55859 | -0,196255971 | 0,047409033 | -4,139632409 | 3,48E-05 | 0,000178605 |
| ENSG00000078018.19 | MAP2 | protein_coding | 35224,8272 | 0,254693583 | 0,042521964 | 5,989694778 | 2,10E-09 | 1,89E-08 |
| ENSG00000078043.16 | PIAS2 | protein_coding | 4000,123944 | -0,15618356 | 0,05838702 | -2,674970573 | 0,007473581 | 0,023228781 |
| ENSG00000078061.13 | ARAF | protein_coding | 5036,642875 | -0,155000091 | 0,050170486 | -3,089467599 | 0,002005156 | 0,007243468 |
| ENSG00000078081.8 | LAMP3 | protein_coding | 13002,81549 | 7,618670935 | 0,082659204 | 92,16966293 | 0 | 0 |
| ENSG00000078114.18 | NEBL | protein_coding | 8374,193375 | -0,306926375 | 0,055910592 | -5,489592628 | 4,03E-08 | 3,12E-07 |
| ENSG00000078124.12 | ACER3 | protein_coding | 4511,189997 | -0,209857961 | 0,072882429 | -2,879404047 | 0,003984275 | 0,01335548 |
| ENSG00000078142.13 | PIK3C3 | protein_coding | 10987,27669 | 0,180456882 | 0,04650982 | 3,879973748 | 0,000104468 | 0,000495328 |
| ENSG00000078177.14 | N4BP2 | protein_coding | 11613,38805 | -0,28620392 | 0,04614242 | -6,202620514 | 5,55E-10 | 5,37E-09 |
| ENSG00000078237.7 | TIGAR | protein_coding | 1721,837039 | 0,46047018 | 0,104577084 | 4,403165224 | 1,07E-05 | 5,95E-05 |
| ENSG00000078295.16 | ADCY2 | protein_coding | 1134,685694 | 0,770057287 | 0,242862182 | 3,170758335 | 0,001520416 | 0,005665128 |
| ENSG00000078304.19 | PPP2R5C | protein_coding | 11211,40268 | -0,227554188 | 0,043810208 | -5,194090547 | 2,06E-07 | 1,46E-06 |
| ENSG00000078319.9 | PMS2P1 | unprocessed_pseud | 963,2614964 | -0,295323425 | 0,085789578 | -3,442416113 | 0,000576543 | 0,002360385 |
| ENSG00000078328.20 | RBFOX1 | protein_coding | 306,7790877 | 1,227410026 | 0,172953076 | 7,096780538 | 1,28E-12 | 1,60E-11 |
| ENSG00000078369.18 | GNB1 | protein_coding | 57349,83893 | -0,352044967 | 0,043698261 | -8,056269397 | 7,87E-16 | 1,25E-14 |
| ENSG00000078487.17 | ZCWPW1 | protein_coding | 537,9815983 | 0,45329732 | 0,121627462 | 3,726932335 | 0,000193824 | 0,000872036 |
| ENSG00000078618.21 | NRDC | protein_coding | 22110,32248 | 0,344123673 | 0,039449002 | 8,72325421 | 2,70E-18 | 5,02E-17 |
| ENSG00000078668.14 | VDAC3 | protein_coding | 24460,82807 | -0,205097169 | 0,039809868 | -5,151917836 | 2,58E-07 | 1,81E-06 |
| ENSG00000078674.17 | PCM1 | protein_coding | 28716,92074 | 0,199548388 | 0,039267674 | 5,081747028 | 3,74E-07 | 2,58E-06 |
| ENSG00000078687.17 | TNRC6C | protein_coding | 2867,262318 | 0,170641102 | 0,06713813 | 2,541642157 | 0,011033307 | 0,032548978 |
| ENSG00000078699.21 | CBFA2T2 | protein_coding | 27138,07314 | 0,274133718 | 0,040648881 | 6,743942595 | 1,54E-11 | 1,73E-10 |
| ENSG00000078804.13 | TP53INP2 | protein_coding | 7039,664438 | -0,350927616 | 0,050917743 | -6,892049707 | 5,50E-12 | 6,47E-11 |
| ENSG00000078808.17 | SDF4 | protein_coding | 9913,387721 | 0,254617381 | 0,052830748 | 4,819492228 | 1,44E-06 | 9,13E-06 |
| ENSG00000078900.15 | TP73 | protein_coding | 79,36698374 | -0,935785805 | 0,279497585 | -3,34809979 | 0,000813677 | 0,003220513 |
| ENSG00000078902.16 | TOLLIP | protein_coding | 12683,01439 | 0,27748644 | 0,050431085 | 5,502289706 | 3,75E-08 | 2,92E-07 |
| ENSG00000079101.16 | CLUL1 | protein_coding | 285,049806 | -0,597247107 | 0,1791942 | -3,332960037 | 0,000859273 | 0,003380468 |
| ENSG00000079102.16 | RUNX1T1 | protein_coding | 5980,076024 | 0,159397556 | 0,055672414 | 2,863133563 | 0,004194737 | 0,013979765 |
| ENSG00000079112.10 | CDH17 | protein_coding | 54456,71391 | -0,209481224 | 0,04425031 | -4,734005818 | 2,20E-06 | 1,36E-05 |
| ENSG00000079134.12 | THOC1 | protein_coding | 4105,646329 | -0,248314818 | 0,057323625 | -4,331805941 | 1,48E-05 | 8,05E-05 |
| ENSG00000079156.17 | OSBPL6 | protein_coding | 4427,109726 | 0,425722837 | 0,054262526 | 7,845614072 | 4,31E-15 | 6,53E-14 |

| | | | | | | | | |
|---|---|---|---|---|---|---|---|---|
| ENSG00000079246.16 | XRCC5 | protein_coding | 80926,22143 | -0,36048145 | 0,043415916 | -8,302979306 | 1,02E-16 | 1,71E-15 |
| ENSG00000079257.8 | LXN | protein_coding | 1437,112456 | 0,363841323 | 0,075605399 | 4,812372229 | 1,49E-06 | 9,45E-06 |
| ENSG00000079277.21 | MKNK1 | protein_coding | 2925,155604 | 0,156855509 | 0,058204681 | 2,694895113 | 0,007041079 | 0,02208426 |
| ENSG00000079335.20 | CDC14A | protein_coding | 1439,886105 | 0,468979921 | 0,070057432 | 6,69422081 | 2,17E-11 | 2,40E-10 |
| ENSG00000079385.22 | CEACAM1 | protein_coding | 17969,27964 | 1,650035859 | 0,048675734 | 33,89853042 | 7,00E-252 | 1,73E-249 |
| ENSG00000079432.7 | CIC | protein_coding | 5955,139228 | 0,138304402 | 0,057162044 | 2,41951464 | 0,015541235 | 0,043710816 |
| ENSG00000079459.13 | FDFT1 | protein_coding | 22217,54555 | 0,191648611 | 0,043469318 | 4,408824898 | 1,04E-05 | 5,82E-05 |
| ENSG00000079462.7 | PAFAH1B3 | protein_coding | 5082,604586 | -0,145066319 | 0,052175643 | -2,780345569 | 0,005430108 | 0,017556205 |
| ENSG00000079616.12 | KIF22 | protein_coding | 5807,816562 | -0,521922454 | 0,051750156 | -10,0854277 | 6,41E-24 | 1,59E-22 |
| ENSG00000079689.14 | SCGN | protein_coding | 27723,32376 | 0,179034416 | 0,043209421 | 4,143411645 | 3,42E-05 | 0,000175926 |
| ENSG00000079691.18 | CARMIL1 | protein_coding | 2934,33282 | 0,50633159 | 0,058962921 | 8,587288094 | 8,90E-18 | 1,60E-16 |
| ENSG00000079819.19 | EPB41L2 | protein_coding | 4282,267207 | 0,685565462 | 0,059495359 | 11,52300749 | 1,01E-30 | 3,32E-29 |
| ENSG00000079950.14 | STX7 | protein_coding | 9338,620213 | 0,14480354 | 0,044550031 | 3,250357795 | 0,001152599 | 0,004403233 |
| ENSG00000080031.10 | AC010327.1 | protein_coding | 1063,367355 | 0,490283174 | 0,08109625 | 6,045694756 | 1,49E-09 | 1,36E-08 |
| ENSG00000080200.10 | CRYBG3 | protein_coding | 4513,649825 | 0,265836204 | 0,059512703 | 4,46688169 | 7,94E-06 | 4,51E-05 |
| ENSG00000080298.15 | RFX3 | protein_coding | 6577,210497 | -0,179671672 | 0,052606085 | -3,415416142 | 0,000636846 | 0,002580982 |
| ENSG00000080345.17 | RIF1 | protein_coding | 19988,06517 | 0,255461912 | 0,041785349 | 6,113671829 | 9,74E-10 | 9,10E-09 |
| ENSG00000080493.17 | SLC4A4 | protein_coding | 103,2048222 | -0,662301763 | 0,270848852 | -2,445281779 | 0,014473901 | 0,041145875 |
| ENSG00000080561.13 | MID2 | protein_coding | 848,1626628 | 0,443414009 | 0,101249154 | 4,379434238 | 1,19E-05 | 6,60E-05 |
| ENSG00000080603.17 | SRCAP | protein_coding | 12126,87586 | 0,35506836 | 0,047640151 | 7,453132489 | 9,11E-14 | 1,25E-12 |
| ENSG00000080644.16 | CHRNA3 | protein_coding | 219,4951356 | -1,215694746 | 0,182075985 | -6,676853857 | 2,44E-11 | 2,70E-10 |
| ENSG00000080709.15 | KCNN2 | protein_coding | 23,15144017 | 1,28787508 | 0,508906966 | 2,530668992 | 0,011384523 | 0,03340735 |
| ENSG00000080802.18 | CNOT4 | protein_coding | 4344,249051 | 0,172127752 | 0,05157449 | 3,337459095 | 0,000845482 | 0,003333499 |
| ENSG00000080815.19 | PSEN1 | protein_coding | 10607,22654 | 0,256408788 | 0,046631064 | 5,498669076 | 3,83E-08 | 2,98E-07 |
| ENSG00000080819.9 | CPOX | protein_coding | 4471,66799 | -0,224495533 | 0,052231904 | -4,298053775 | 1,72E-05 | 9,29E-05 |
| ENSG00000080824.18 | HSP90AA1 | protein_coding | 281198,5021 | -0,219259987 | 0,043967261 | -4,986892064 | 6,14E-07 | 4,11E-06 |
| ENSG00000080845.17 | DLGAP4 | protein_coding | 10082,04564 | -0,17247415 | 0,047999764 | -3,593229153 | 0,000326605 | 0,001409154 |
| ENSG00000080854.14 | IGSF9B | protein_coding | 1207,6715 | -0,438884621 | 0,094058758 | -4,666068645 | 3,07E-06 | 1,86E-05 |
| ENSG00000080986.13 | NDC80 | protein_coding | 6108,392192 | -0,344110417 | 0,055797318 | -6,167149763 | 6,95E-10 | 6,61E-09 |
| ENSG00000081019.13 | RSBN1 | protein_coding | 4232,016642 | 0,182664229 | 0,057331719 | 3,186093704 | 0,001442079 | 0,005395571 |
| ENSG00000081041.9 | CXCL2 | protein_coding | 175,6872081 | 4,212526738 | 0,271360824 | 15,52371 | 2,40E-54 | 1,49E-52 |
| ENSG00000081051.7 | AFP | protein_coding | 114,6909571 | 0,730600945 | 0,253567883 | 2,88128345 | 0,003960593 | 0,013286777 |
| ENSG00000081059.20 | TCF7 | protein_coding | 465,0492247 | -0,700616485 | 0,113956553 | -6,148101786 | 7,84E-10 | 7,43E-09 |
| ENSG00000081087.15 | OSTM1 | protein_coding | 2892,288074 | 0,560900115 | 0,061477851 | 9,123612951 | 7,27E-20 | 1,46E-18 |
| ENSG00000081138.14 | CDH7 | protein_coding | 77,40975725 | -1,214421078 | 0,397001859 | -3,058980833 | 0,002220914 | 0,007924944 |
| ENSG00000081154.11 | PCNP | protein_coding | 21968,03342 | -0,23230712 | 0,046022091 | -5,047730718 | 4,47E-07 | 3,06E-06 |

| | | | | | | | | |
|---|---|---|---|---|---|---|---|---|
| ENSG00000081177.18 | EXD2 | protein_coding | 5825,281889 | 0,119506793 | 0,048940956 | 2,441856539 | 0,014611952 | 0,041480694 |
| ENSG00000081320.11 | STK17B | protein_coding | 1412,654873 | -0,236958524 | 0,085808939 | -2,761466652 | 0,005754239 | 0,018479408 |
| ENSG00000081386.12 | ZNF510 | protein_coding | 5093,753296 | 0,172446861 | 0,055569204 | 3,103281114 | 0,001913878 | 0,006948301 |
| ENSG00000081479.15 | LRP2 | protein_coding | 29,31668973 | 2,029068446 | 0,674236052 | 3,009433326 | 0,002617355 | 0,009183795 |
| ENSG00000081665.14 | ZNF506 | protein_coding | 4667,726934 | 0,211712854 | 0,051194073 | 4,135495415 | 3,54E-05 | 0,000181688 |
| ENSG00000081803.16 | CADPS2 | protein_coding | 263,4636225 | 2,579640027 | 0,205364441 | 12,56127894 | 3,45E-36 | 1,38E-34 |
| ENSG00000082014.16 | SMARCD3 | protein_coding | 122,9683912 | 0,87226654 | 0,238844658 | 3,652024484 | 0,000260181 | 0,001142938 |
| ENSG00000082146.13 | STRADB | protein_coding | 2555,291613 | 0,318386432 | 0,087231986 | 3,64988173 | 0,000262361 | 0,001151616 |
| ENSG00000082153.18 | BZW1 | protein_coding | 32748,99326 | -0,294250533 | 0,040344951 | -7,293366996 | 3,02E-13 | 3,99E-12 |
| ENSG00000082175.14 | PGR | protein_coding | 1184,082657 | 0,495300014 | 0,103213932 | 4,798770883 | 1,60E-06 | 1,01E-05 |
| ENSG00000082212.13 | ME2 | protein_coding | 10576,83554 | 0,26526014 | 0,043906482 | 6,041480152 | 1,53E-09 | 1,40E-08 |
| ENSG00000082213.18 | C5orf22 | protein_coding | 3455,143682 | 0,342269875 | 0,054595711 | 6,269171483 | 3,63E-10 | 3,57E-09 |
| ENSG00000082269.16 | FAM135A | protein_coding | 9948,918482 | 0,174054159 | 0,061319535 | 2,838478143 | 0,004532922 | 0,01495406 |
| ENSG00000082397.17 | EPB41L3 | protein_coding | 18721,32684 | 0,155620092 | 0,042952982 | 3,623033458 | 0,000291168 | 0,001267925 |
| ENSG00000082438.16 | COBLL1 | protein_coding | 4748,740943 | 0,248789948 | 0,057359063 | 4,337413021 | 1,44E-05 | 7,87E-05 |
| ENSG00000082497.12 | SERTAD4 | protein_coding | 226,3031601 | -0,986249315 | 0,189445393 | -5,205982058 | 1,93E-07 | 1,38E-06 |
| ENSG00000082512.15 | TRAF5 | protein_coding | 451,7573802 | 0,634130802 | 0,130875967 | 4,845280729 | 1,26E-06 | 8,10E-06 |
| ENSG00000082641.16 | NFE2L1 | protein_coding | 33240,98017 | 0,2099487 | 0,040682903 | 5,160612532 | 2,46E-07 | 1,74E-06 |
| ENSG00000082701.15 | GSK3B | protein_coding | 18914,32868 | -0,269316821 | 0,045575274 | -5,909274901 | 3,44E-09 | 3,02E-08 |
| ENSG00000082805.20 | ERC1 | protein_coding | 10490,5021 | -0,14820018 | 0,057521311 | -2,576439552 | 0,009982366 | 0,029883372 |
| ENSG00000083067.22 | TRPM3 | protein_coding | 8190,961922 | -0,222888622 | 0,054833186 | -4,064849039 | 4,81E-05 | 0,000241272 |
| ENSG00000083093.9 | PALB2 | protein_coding | 4786,539119 | 0,131387704 | 0,050189912 | 2,617811019 | 0,00884958 | 0,026961992 |
| ENSG00000083097.14 | DOP1A | protein_coding | 9123,817227 | 0,420734848 | 0,05349567 | 7,864839291 | 3,70E-15 | 5,63E-14 |
| ENSG00000083099.11 | LYRM2 | protein_coding | 11697,39671 | -0,145226048 | 0,048568748 | -2,990113081 | 0,002788742 | 0,00970948 |
| ENSG00000083123.15 | BCKDHB | protein_coding | 4568,609631 | 0,254622818 | 0,056266721 | 4,525282673 | 6,03E-06 | 3,49E-05 |
| ENSG00000083168.11 | KAT6A | protein_coding | 15984,12506 | 0,306404783 | 0,044887172 | 6,826110206 | 8,72E-12 | 1,01E-10 |
| ENSG00000083223.18 | TUT7 | protein_coding | 4197,292293 | 0,305268731 | 0,066865394 | 4,565421879 | 4,98E-06 | 2,92E-05 |
| ENSG00000083312.17 | TNPO1 | protein_coding | 44225,94715 | -0,366481395 | 0,052537612 | -6,975600555 | 3,05E-12 | 3,68E-11 |
| ENSG00000083444.17 | PLOD1 | protein_coding | 7869,436429 | 0,185367386 | 0,049175642 | 3,769495985 | 0,000163578 | 0,000746244 |
| ENSG00000083454.22 | P2RX5 | protein_coding | 95,9901387 | 0,61026184 | 0,230672818 | 2,645573262 | 0,008155263 | 0,025087714 |
| ENSG00000083457.12 | ITGAE | protein_coding | 2189,163413 | -1,232104975 | 0,063912877 | -19,27788318 | 8,24E-83 | 7,82E-81 |
| ENSG00000083720.13 | OXCT1 | protein_coding | 6968,136994 | -0,608774485 | 0,056058401 | -10,85964769 | 1,79E-27 | 5,12E-26 |
| ENSG00000083799.17 | CYLD | protein_coding | 9617,1305 | 0,262226324 | 0,047487358 | 5,522023914 | 3,35E-08 | 2,63E-07 |
| ENSG00000083812.12 | ZNF324 | protein_coding | 2573,193383 | 0,258503135 | 0,082010061 | 3,152090503 | 0,00162106 | 0,005996458 |
| ENSG00000083814.13 | ZNF671 | protein_coding | 517,9219876 | 0,49746673 | 0,105056595 | 4,735226096 | 2,19E-06 | 1,35E-05 |
| ENSG00000083828.16 | ZNF586 | protein_coding | 2014,07015 | 0,226998602 | 0,064382871 | 3,525760772 | 0,000422268 | 0,001778599 |

| | | | | | | | | |
|---|---|---|---|---|---|---|---|---|
| ENSG00000083844.10 | ZNF264 | protein_coding | 6404,337523 | 0,320866244 | 0,060617391 | 5,293303391 | 1,20E-07 | 8,81E-07 |
| ENSG00000083845.9 | RPS5 | protein_coding | 29713,81312 | -0,370148398 | 0,04851971 | -7,62882548 | 2,37E-14 | 3,39E-13 |
| ENSG00000083896.12 | YTHDC1 | protein_coding | 28099,53 | 0,562142094 | 0,049097192 | 11,44957732 | 2,36E-30 | 7,61E-29 |
| ENSG00000083937.9 | CHMP2B | protein_coding | 6855,770936 | 0,220765256 | 0,049393429 | 4,469526815 | 7,84E-06 | 4,46E-05 |
| ENSG00000084090.13 | STARD7 | protein_coding | 20456,27426 | -0,163620992 | 0,045159906 | -3,623147266 | 0,00029104 | 0,001267857 |
| ENSG00000084234.17 | APLP2 | protein_coding | 56921,9537 | 0,1707857 | 0,042938896 | 3,9774125 | 6,97E-05 | 0,000340984 |
| ENSG00000084444.14 | FAM234B | protein_coding | 3399,38126 | -0,452561691 | 0,068295643 | -6,626508971 | 3,44E-11 | 3,74E-10 |
| ENSG00000084453.16 | SLCO1A2 | protein_coding | 51,03894464 | -1,186236846 | 0,381519122 | -3,109246109 | 0,001875654 | 0,00682715 |
| ENSG00000084623.11 | EIF3I | protein_coding | 14759,56562 | -0,269305291 | 0,044908862 | -5,996707038 | 2,01E-09 | 1,82E-08 |
| ENSG00000084628.10 | NKAIN1 | protein_coding | 374,463243 | -0,35011018 | 0,134235909 | -2,608170817 | 0,009102752 | 0,027628619 |
| ENSG00000084674.14 | APOB | protein_coding | 29390,69863 | 0,34122657 | 0,048879865 | 6,980922943 | 2,93E-12 | 3,55E-11 |
| ENSG00000084676.15 | NCOA1 | protein_coding | 9001,317482 | 0,310627162 | 0,04332429 | 7,169815366 | 7,51E-13 | 9,60E-12 |
| ENSG00000084693.16 | AGBL5 | protein_coding | 2324,844987 | -0,20632426 | 0,063068532 | -3,271429547 | 0,001070052 | 0,00412146 |
| ENSG00000084710.14 | EFR3B | protein_coding | 1823,03542 | -0,419688076 | 0,078642565 | -5,336652925 | 9,47E-08 | 6,98E-07 |
| ENSG00000084731.15 | KIF3C | protein_coding | 6564,540273 | -0,145914465 | 0,051526119 | -2,831854362 | 0,004627892 | 0,015227193 |
| ENSG00000084754.12 | HADHA | protein_coding | 18314,64745 | -0,262271212 | 0,041101256 | -6,381099638 | 1,76E-10 | 1,79E-09 |
| ENSG00000084774.14 | CAD | protein_coding | 6496,807964 | -0,282197328 | 0,059318928 | -4,757289761 | 1,96E-06 | 1,22E-05 |
| ENSG00000085063.16 | CD59 | protein_coding | 25052,2183 | 0,237451836 | 0,043206958 | 5,4956851 | 3,89E-08 | 3,02E-07 |
| ENSG00000085117.12 | CD82 | protein_coding | 5043,448881 | 0,371774201 | 0,051119825 | 7,272603237 | 3,53E-13 | 4,62E-12 |
| ENSG00000085185.15 | BCORL1 | protein_coding | 1287,994075 | 0,231866456 | 0,090067185 | 2,5743722 | 0,010042218 | 0,030034584 |
| ENSG00000085276.18 | MECOM | protein_coding | 10733,06077 | 0,360960548 | 0,04715798 | 7,654283497 | 1,94E-14 | 2,80E-13 |
| ENSG00000085415.16 | SEH1L | protein_coding | 4806,894043 | -0,316832782 | 0,054490106 | -5,814501087 | 6,08E-09 | 5,20E-08 |
| ENSG00000085433.15 | WDR47 | protein_coding | 2795,90335 | 0,256920896 | 0,05807602 | 4,423872321 | 9,69E-06 | 5,44E-05 |
| ENSG00000085449.15 | WDFY1 | protein_coding | 14880,97076 | 0,300379316 | 0,047198972 | 6,364107223 | 1,96E-10 | 1,99E-09 |
| ENSG00000085491.17 | SLC25A24 | protein_coding | 8236,698003 | -0,319847275 | 0,046762433 | -6,839833851 | 7,93E-12 | 9,18E-11 |
| ENSG00000085552.17 | IGSF9 | protein_coding | 5731,708828 | -0,468986128 | 0,058388716 | -8,032136397 | 9,58E-16 | 1,52E-14 |
| ENSG00000085563.14 | ABCB1 | protein_coding | 18086,96905 | 0,265882937 | 0,043696123 | 6,084817526 | 1,17E-09 | 1,08E-08 |
| ENSG00000085719.13 | CPNE3 | protein_coding | 14616,1175 | -0,327792107 | 0,051024676 | -6,424187894 | 1,33E-10 | 1,37E-09 |
| ENSG00000085788.13 | DDHD2 | protein_coding | 7270,749291 | -0,181386517 | 0,071344635 | -2,542398835 | 0,011009447 | 0,032487105 |
| ENSG00000085832.17 | EPS15 | protein_coding | 10363,19037 | 0,171883584 | 0,047900423 | 3,588352109 | 0,000332775 | 0,001432752 |
| ENSG00000085982.13 | USP40 | protein_coding | 5734,654802 | 0,186079536 | 0,05478695 | 3,396420769 | 0,000682733 | 0,002748148 |
| ENSG00000085998.14 | POMGNT1 | protein_coding | 5905,706613 | -0,223520271 | 0,055075889 | -4,058405147 | 4,94E-05 | 0,000247584 |
| ENSG00000086061.16 | DNAJA1 | protein_coding | 65919,41415 | 0,604751286 | 0,039740718 | 15,21742207 | 2,71E-52 | 1,64E-50 |
| ENSG00000086065.14 | CHMP5 | protein_coding | 19576,99052 | 1,09294986 | 0,048643803 | 22,46842965 | 8,45E-112 | 1,09E-109 |
| ENSG00000086102.19 | NFX1 | protein_coding | 7178,177553 | 0,274407001 | 0,047185785 | 5,815459026 | 6,05E-09 | 5,17E-08 |
| ENSG00000086200.16 | IPO11 | protein_coding | 3591,056337 | -0,148330737 | 0,055654072 | -2,665227043 | 0,007693636 | 0,023846882 |

| | | | | | | | | |
|---|---|---|---|---|---|---|---|---|
| ENSG00000086289.12 | EPDR1 | protein_coding | 2487,35772 | 0,461603662 | 0,070245971 | 6,571247531 | 4,99E-11 | 5,35E-10 |
| ENSG00000086300.15 | SNX10 | protein_coding | 1687,722643 | -0,196445389 | 0,069653385 | -2,820327948 | 0,004797459 | 0,015720771 |
| ENSG00000086475.15 | SEPHS1 | protein_coding | 4582,158869 | -0,400189713 | 0,062888675 | -6,363462298 | 1,97E-10 | 1,99E-09 |
| ENSG00000086827.9 | ZW10 | protein_coding | 4224,238926 | -0,209955578 | 0,059038443 | -3,556251975 | 0,000376183 | 0,001602481 |
| ENSG00000086967.10 | MYBPC2 | protein_coding | 94,1556362 | -1,78323895 | 0,315611805 | -5,650102181 | 1,60E-08 | 1,30E-07 |
| ENSG00000087053.18 | MTMR2 | protein_coding | 6178,96617 | -0,285718239 | 0,06760479 | -4,226301668 | 2,38E-05 | 0,000125583 |
| ENSG00000087074.8 | PPP1R15A | protein_coding | 2206,740823 | 0,464173982 | 0,07740393 | 5,996775346 | 2,01E-09 | 1,82E-08 |
| ENSG00000087076.9 | HSD17B14 | protein_coding | 253,2307501 | -0,695353167 | 0,152993257 | -4,544992232 | 5,49E-06 | 3,20E-05 |
| ENSG00000087086.15 | FTL | protein_coding | 161395,4061 | -0,3580476 | 0,045247682 | -7,913059453 | 2,51E-15 | 3,86E-14 |
| ENSG00000087088.20 | BAX | protein_coding | 4905,679746 | -0,19608383 | 0,052740597 | -3,717891764 | 0,000200892 | 0,000900594 |
| ENSG00000087095.13 | NLK | protein_coding | 6223,671757 | 0,137379419 | 0,048172613 | 2,851815804 | 0,004347028 | 0,01441474 |
| ENSG00000087116.16 | ADAMTS2 | protein_coding | 4160,870551 | -0,32053785 | 0,079083911 | -4,05313606 | 5,05E-05 | 0,000252948 |
| ENSG00000087152.15 | ATXN7L3 | protein_coding | 6089,625086 | 0,168250451 | 0,050100949 | 3,358228791 | 0,000784437 | 0,003112991 |
| ENSG00000087206.17 | UIMC1 | protein_coding | 4137,070541 | 0,247233972 | 0,065155618 | 3,794515049 | 0,000147932 | 0,000682758 |
| ENSG00000087250.8 | MT3 | protein_coding | 92,05259572 | 0,927381675 | 0,246100951 | 3,768297813 | 0,000164365 | 0,00074953 |
| ENSG00000087266.17 | SH3BP2 | protein_coding | 2425,758918 | -0,160757816 | 0,065943841 | -2,437798771 | 0,014776997 | 0,041881363 |
| ENSG00000087301.9 | TXNDC16 | protein_coding | 13352,87659 | -0,32864531 | 0,04341969 | -7,56903853 | 3,76E-14 | 5,32E-13 |
| ENSG00000087303.18 | NID2 | protein_coding | 12,10961227 | -2,562353384 | 0,715677983 | -3,580316072 | 0,000343179 | 0,001473881 |
| ENSG00000087338.5 | GMCL1 | protein_coding | 10014,65083 | -0,117992637 | 0,044961162 | -2,624323589 | 0,008682126 | 0,026523611 |
| ENSG00000087586.17 | AURKA | protein_coding | 9937,361563 | -0,854051823 | 0,047716898 | -17,89830994 | 1,22E-71 | 1,00E-69 |
| ENSG00000087842.11 | PIR | protein_coding | 895,1531651 | 0,211709778 | 0,085135295 | 2,486745118 | 0,012891773 | 0,037218104 |
| ENSG00000087903.13 | RFX2 | protein_coding | 1238,939506 | -0,321633369 | 0,094603123 | -3,399817656 | 0,000674308 | 0,002717636 |
| ENSG00000088035.18 | ALG6 | protein_coding | 3079,380018 | -0,333119625 | 0,055799692 | -5,969918711 | 2,37E-09 | 2,12E-08 |
| ENSG00000088179.9 | PTPN4 | protein_coding | 4081,222285 | 0,551174268 | 0,168327717 | 3,274411823 | 0,001058822 | 0,004082395 |
| ENSG00000088256.9 | GNA11 | protein_coding | 15612,51471 | -0,225909919 | 0,051882896 | -4,35422725 | 1,34E-05 | 7,34E-05 |
| ENSG00000088298.13 | EDEM2 | protein_coding | 2957,767068 | 0,711731732 | 0,06255707 | 11,37731881 | 5,42E-30 | 1,70E-28 |
| ENSG00000088305.18 | DNMT3B | protein_coding | 603,1601929 | -0,36288121 | 0,101441448 | -3,577247932 | 0,000347231 | 0,001489295 |
| ENSG00000088325.16 | TPX2 | protein_coding | 19765,9159 | -0,704970992 | 0,045318542 | -15,55590615 | 1,45E-54 | 9,12E-53 |
| ENSG00000088386.17 | SLC15A1 | protein_coding | 1267,767276 | 0,238668034 | 0,07785036 | 3,065728078 | 0,002171407 | 0,007765523 |
| ENSG00000088387.19 | DOCK9 | protein_coding | 4658,040574 | 0,396982691 | 0,082062191 | 4,837583398 | 1,31E-06 | 8,39E-06 |
| ENSG00000088451.11 | TGDS | protein_coding | 3664,053129 | 0,24136799 | 0,054836515 | 4,401592413 | 1,07E-05 | 5,99E-05 |
| ENSG00000088538.13 | DOCK3 | protein_coding | 4193,28577 | 0,276774143 | 0,062872088 | 4,402178318 | 1,07E-05 | 5,98E-05 |
| ENSG00000088756.12 | ARHGAP28 | protein_coding | 1198,13347 | -0,603430067 | 0,075292828 | -8,014442868 | 1,11E-15 | 1,74E-14 |
| ENSG00000088808.17 | PPP1R13B | protein_coding | 2363,764085 | 0,266294164 | 0,062501121 | 4,260630189 | 2,04E-05 | 0,00010868 |
| ENSG00000088812.18 | ATRN | protein_coding | 17004,69435 | -0,357192957 | 0,045978515 | -7,768692718 | 7,93E-15 | 1,18E-13 |
| ENSG00000088832.17 | FKBP1A | protein_coding | 22406,61391 | -0,270507877 | 0,046381091 | -5,832287896 | 5,47E-09 | 4,71E-08 |

| | | | | | | | | |
|---|---|---|---|---|---|---|---|---|
| ENSG00000088854.12 | C20orf194 | protein_coding | 3004,693974 | 0,316462622 | 0,057710583 | 5,483615094 | 4,17E-08 | 3,22E-07 |
| ENSG00000088876.11 | ZNF343 | protein_coding | 2835,244803 | 0,2184859 | 0,058035908 | 3,764667583 | 0,000166771 | 0,000759734 |
| ENSG00000088881.20 | EBF4 | protein_coding | 1669,378935 | -0,174218141 | 0,072804146 | -2,392970045 | 0,016712606 | 0,046568957 |
| ENSG00000088888.18 | MAVS | protein_coding | 18566,72389 | -0,577224273 | 0,064374265 | -8,966693051 | 3,06E-19 | 5,96E-18 |
| ENSG00000088899.15 | LZTS3 | protein_coding | 3440,327614 | 0,298215491 | 0,062573979 | 4,765806758 | 1,88E-06 | 1,17E-05 |
| ENSG00000088970.16 | KIZ | protein_coding | 2300,927979 | 0,356630959 | 0,077765578 | 4,585974508 | 4,52E-06 | 2,67E-05 |
| ENSG00000088986.11 | DYNLL1 | protein_coding | 17220,26775 | -0,143390286 | 0,044709819 | -3,207131919 | 0,001340655 | 0,005051357 |
| ENSG00000088992.18 | TESC | protein_coding | 1542,959935 | -0,502356304 | 0,094526351 | -5,314457793 | 1,07E-07 | 7,86E-07 |
| ENSG00000089006.16 | SNX5 | protein_coding | 13592,51701 | -0,241027245 | 0,044474122 | -5,419494174 | 5,98E-08 | 4,54E-07 |
| ENSG00000089009.15 | AC004086.1 | protein_coding | 58923,19351 | -0,743497799 | 0,039106164 | -19,01229151 | 1,35E-80 | 1,25E-78 |
| ENSG00000089050.16 | RBBP9 | protein_coding | 7158,094409 | -0,410423069 | 0,055843632 | -7,349505362 | 1,99E-13 | 2,66E-12 |
| ENSG00000089053.13 | ANAPC5 | protein_coding | 15578,03686 | -0,198201118 | 0,048984011 | -4,046241122 | 5,20E-05 | 0,000260164 |
| ENSG00000089057.15 | SLC23A2 | protein_coding | 7521,766735 | -0,175747133 | 0,051747002 | -3,396276658 | 0,000683093 | 0,002748612 |
| ENSG00000089063.15 | TMEM230 | protein_coding | 22234,39531 | -0,297236209 | 0,047797531 | -6,218651888 | 5,01E-10 | 4,87E-09 |
| ENSG00000089094.19 | KDM2B | protein_coding | 4070,31688 | -0,394901829 | 0,059380742 | -6,650335084 | 2,92E-11 | 3,20E-10 |
| ENSG00000089123.16 | TASP1 | protein_coding | 2309,494346 | 0,214296877 | 0,064575935 | 3,318525337 | 0,000904941 | 0,003540028 |
| ENSG00000089127.13 | AC004551.1 | protein_coding | 61396,2198 | 7,10644263 | 0,346331513 | 20,51919148 | 1,45E-93 | 1,58E-91 |
| ENSG00000089157.16 | RPLP0 | protein_coding | 89287,17605 | -0,803717469 | 0,045147039 | -17,80221898 | 6,79E-71 | 5,52E-69 |
| ENSG00000089169.15 | RPH3A | protein_coding | 107,8235743 | 4,628257715 | 0,384025073 | 12,05196753 | 1,89E-33 | 6,91E-32 |
| ENSG00000089177.18 | KIF16B | protein_coding | 6422,551287 | 0,226644264 | 0,048642267 | 4,65940996 | 3,17E-06 | 1,91E-05 |
| ENSG00000089199.10 | CHGB | protein_coding | 89895,16092 | 0,18335824 | 0,038613179 | 4,748592181 | 2,05E-06 | 1,27E-05 |
| ENSG00000089234.16 | BRAP | protein_coding | 4186,590963 | 0,276634689 | 0,050800435 | 5,445518142 | 5,17E-08 | 3,96E-07 |
| ENSG00000089248.7 | ERP29 | protein_coding | 51020,86222 | 0,38065935 | 0,054905363 | 6,933008486 | 4,12E-12 | 4,92E-11 |
| ENSG00000089280.18 | FUS | protein_coding | 53577,86553 | -0,115388929 | 0,042889778 | -2,690359687 | 0,007137504 | 0,022335338 |
| ENSG00000089289.16 | IGBP1 | protein_coding | 3577,142206 | -0,162047811 | 0,053249715 | -3,043167636 | 0,002341019 | 0,008307424 |
| ENSG00000089335.21 | ZNF302 | protein_coding | 8794,91827 | 0,199989845 | 0,046770061 | 4,276022799 | 1,90E-05 | 0,000101918 |
| ENSG00000089356.18 | FXYD3 | protein_coding | 12663,33238 | 0,376623598 | 0,071670603 | 5,254924367 | 1,48E-07 | 1,07E-06 |
| ENSG00000089558.9 | KCNH4 | protein_coding | 52,26686202 | 1,177097587 | 0,339028953 | 3,471967737 | 0,000516658 | 0,002132673 |
| ENSG00000089639.11 | GMIP | protein_coding | 621,1460997 | 0,703704434 | 0,113993352 | 6,173205921 | 6,69E-10 | 6,39E-09 |
| ENSG00000089685.15 | BIRC5 | protein_coding | 6900,00545 | -0,8339956 | 0,066985793 | -12,45033551 | 1,39E-35 | 5,47E-34 |
| ENSG00000089737.17 | DDX24 | protein_coding | 20763,13214 | 0,42376061 | 0,04227676 | 10,02348839 | 1,20E-23 | 2,92E-22 |
| ENSG00000089818.18 | NECAP1 | protein_coding | 11370,60314 | 0,305401604 | 0,042141406 | 7,247067241 | 4,26E-13 | 5,54E-12 |
| ENSG00000089916.17 | GPATCH2L | protein_coding | 8313,462957 | -0,337492744 | 0,053873499 | -6,264541041 | 3,74E-10 | 3,67E-09 |
| ENSG00000090013.10 | BLVRB | protein_coding | 671,9883954 | -0,276604151 | 0,107176745 | -2,58082247 | 0,009856525 | 0,029565636 |
| ENSG00000090060.18 | PAPOLA | protein_coding | 40558,70816 | -0,176408701 | 0,042830669 | -4,118747306 | 3,81E-05 | 0,000194522 |
| ENSG00000090061.17 | CCNK | protein_coding | 4781,710657 | 0,184393968 | 0,056651711 | 3,254870231 | 0,001134442 | 0,004342713 |

| | | | | | | | | |
|---|---|---|---|---|---|---|---|---|
| ENSG00000090104.12 | RGS1 | protein_coding | 273,6199296 | 1,172380217 | 0,151642652 | 7,731203588 | 1,07E-14 | 1,56E-13 |
| ENSG00000090238.11 | YPEL3 | protein_coding | 990,0827646 | -0,298957697 | 0,10230735 | -2,92215267 | 0,003476211 | 0,011839315 |
| ENSG00000090263.16 | MRPS33 | protein_coding | 8787,23233 | -0,224219275 | 0,043702451 | -5,130588106 | 2,89E-07 | 2,02E-06 |
| ENSG00000090266.13 | NDUFB2 | protein_coding | 7072,998763 | -0,362454945 | 0,0546288 | -6,634869212 | 3,25E-11 | 3,55E-10 |
| ENSG00000090339.9 | ICAM1 | protein_coding | 41,16106661 | 2,865527266 | 0,45679726 | 6,27308331 | 3,54E-10 | 3,49E-09 |
| ENSG00000090432.7 | MUL1 | protein_coding | 3906,713336 | 0,137821181 | 0,051330992 | 2,68495064 | 0,007254052 | 0,022646526 |
| ENSG00000090487.11 | SPG21 | protein_coding | 9943,882351 | -0,175902915 | 0,044985056 | -3,910252253 | 9,22E-05 | 0,000441652 |
| ENSG00000090520.11 | DNAJB11 | protein_coding | 13237,24853 | 0,419019464 | 0,047312546 | 8,856413386 | 8,26E-19 | 1,57E-17 |
| ENSG00000090554.13 | FLT3LG | protein_coding | 106,2064177 | 1,583805885 | 0,243226016 | 6,511663147 | 7,43E-11 | 7,83E-10 |
| ENSG00000090565.16 | RAB11FIP3 | protein_coding | 8975,71931 | -0,178576695 | 0,060489379 | -2,952199184 | 0,003155193 | 0,010832862 |
| ENSG00000090621.14 | PABPC4 | protein_coding | 11286,921 | -0,14330346 | 0,042684824 | -3,357246128 | 0,00078723 | 0,003123526 |
| ENSG00000090776.6 | EFNB1 | protein_coding | 6049,910706 | -0,270454573 | 0,052383046 | -5,163017321 | 2,43E-07 | 1,71E-06 |
| ENSG00000090861.15 | AARS | protein_coding | 25810,75852 | 0,4688499 | 0,044446278 | 10,54868755 | 5,15E-26 | 1,40E-24 |
| ENSG00000090863.11 | GLG1 | protein_coding | 30544,14328 | 0,269699293 | 0,042796516 | 6,301898364 | 2,94E-10 | 2,93E-09 |
| ENSG00000090889.12 | KIF4A | protein_coding | 6068,571091 | -0,627632771 | 0,052297448 | -12,00121224 | 3,50E-33 | 1,27E-31 |
| ENSG00000090905.19 | TNRC6A | protein_coding | 13084,79897 | 0,147712761 | 0,047505977 | 3,109351102 | 0,001874988 | 0,006825828 |
| ENSG00000090975.12 | PITPNM2 | protein_coding | 1871,440138 | 0,744771411 | 0,077466738 | 9,614079971 | 6,97E-22 | 1,57E-20 |
| ENSG00000090989.18 | EXOC1 | protein_coding | 4233,871907 | 0,293183783 | 0,054404077 | 5,389003908 | 7,08E-08 | 5,32E-07 |
| ENSG00000091039.17 | OSBPL8 | protein_coding | 8640,177683 | -0,203723129 | 0,046861173 | -4,347375744 | 1,38E-05 | 7,55E-05 |
| ENSG00000091073.19 | DTX2 | protein_coding | 426,2768507 | 0,452124334 | 0,159047509 | 2,842699878 | 0,004473317 | 0,014776915 |
| ENSG00000091127.14 | PUS7 | protein_coding | 4123,087238 | 0,167196912 | 0,059662729 | 2,80236784 | 0,0050729 | 0,016527124 |
| ENSG00000091128.13 | LAMB4 | protein_coding | 94,37219076 | -0,98427151 | 0,284736219 | -3,456783668 | 0,000546664 | 0,002247461 |
| ENSG00000091136.14 | LAMB1 | protein_coding | 31401,17734 | 0,515856619 | 0,045452739 | 11,3492966 | 7,48E-30 | 2,33E-28 |
| ENSG00000091137.13 | SLC26A4 | protein_coding | 194,840133 | -1,737697473 | 0,225970018 | -7,689947053 | 1,47E-14 | 2,13E-13 |
| ENSG00000091262.15 | ABCC6 | protein_coding | 6145,089483 | 0,234658211 | 0,048679701 | 4,820453026 | 1,43E-06 | 9,09E-06 |
| ENSG00000091317.8 | CMTM6 | protein_coding | 22555,00004 | -0,437800333 | 0,047049133 | -9,305173143 | 1,34E-20 | 2,83E-19 |
| ENSG00000091409.15 | ITGA6 | protein_coding | 3664,873386 | 0,361696906 | 0,054249005 | 6,667346366 | 2,60E-11 | 2,87E-10 |
| ENSG00000091428.18 | RAPGEF4 | protein_coding | 3174,579183 | -0,208047984 | 0,077266478 | -2,692603425 | 0,007089654 | 0,022210303 |
| ENSG00000091436.17 | MAP3K20 | protein_coding | 1495,020411 | -0,542916531 | 0,076839115 | -7,065627083 | 1,60E-12 | 1,99E-11 |
| ENSG00000091483.6 | FH | protein_coding | 10227,76949 | -0,206133265 | 0,046113908 | -4,470088859 | 7,82E-06 | 4,45E-05 |
| ENSG00000091490.11 | SEL1L3 | protein_coding | 4541,31013 | 0,373113729 | 0,062004517 | 6,017524942 | 1,77E-09 | 1,61E-08 |
| ENSG00000091527.15 | CDV3 | protein_coding | 19742,39739 | -0,193747735 | 0,044281613 | -4,375354061 | 1,21E-05 | 6,71E-05 |
| ENSG00000091592.16 | NLRP1 | protein_coding | 174,2714128 | -0,632215602 | 0,192571905 | -3,283010577 | 0,001027048 | 0,003968726 |
| ENSG00000091704.10 | CPA1 | protein_coding | 1773,100917 | -0,29987271 | 0,064537998 | -4,646451998 | 3,38E-06 | 2,03E-05 |
| ENSG00000091732.16 | ZC3HC1 | protein_coding | 2585,433056 | 0,170825583 | 0,068781644 | 2,48359263 | 0,01300645 | 0,037514376 |
| ENSG00000091879.13 | ANGPT2 | protein_coding | 93,93671244 | 1,419381805 | 0,303812938 | 4,671893876 | 2,98E-06 | 1,81E-05 |

| | | | | | | | | |
|---|---|---|---|---|---|---|---|---|
| ENSG00000092010.15 | PSME1 | protein_coding | 16564,70487 | 1,726865229 | 0,043596466 | 39,61021161 | 0 | 0 |
| ENSG00000092054.13 | MYH7 | protein_coding | 32,26737297 | 2,121593578 | 0,456680092 | 4,645688773 | 3,39E-06 | 2,04E-05 |
| ENSG00000092096.16 | SLC22A17 | protein_coding | 9089,013463 | 0,291822738 | 0,065995221 | 4,421876842 | 9,78E-06 | 5,49E-05 |
| ENSG00000092098.17 | RNF31 | protein_coding | 2866,979922 | 1,224490003 | 0,062845879 | 19,48401429 | 1,50E-84 | 1,47E-82 |
| ENSG00000092140.16 | G2E3 | protein_coding | 5369,488682 | -0,351656085 | 0,054934138 | -6,401412603 | 1,54E-10 | 1,58E-09 |
| ENSG00000092201.10 | SUPT16H | protein_coding | 37313,96331 | -0,177439369 | 0,050618984 | -3,505391739 | 0,000455936 | 0,00191111 |
| ENSG00000092203.14 | TOX4 | protein_coding | 8426,542419 | 0,146309817 | 0,046738666 | 3,130380706 | 0,001745799 | 0,006410471 |
| ENSG00000092330.18 | AL096870.1 | protein_coding | 5667,125916 | 0,192283352 | 0,052546236 | 3,659317337 | 0,000252888 | 0,001114163 |
| ENSG00000092439.16 | TRPM7 | protein_coding | 12068,5519 | 0,201779195 | 0,047231019 | 4,272175398 | 1,94E-05 | 0,00010357 |
| ENSG00000092445.12 | TYRO3 | protein_coding | 387,6342815 | -0,481527741 | 0,164884791 | -2,920389067 | 0,003495946 | 0,011897529 |
| ENSG00000092470.12 | WDR76 | protein_coding | 10975,91935 | -0,321450336 | 0,047518149 | -6,764790791 | 1,34E-11 | 1,51E-10 |
| ENSG00000092607.15 | TBX15 | protein_coding | 429,5936977 | 0,830832191 | 0,126254302 | 6,580624827 | 4,68E-11 | 5,04E-10 |
| ENSG00000092621.12 | PHGDH | protein_coding | 4035,158054 | 0,18007056 | 0,062885664 | 2,863459639 | 0,004190422 | 0,013967453 |
| ENSG00000092820.18 | EZR | protein_coding | 16033,34816 | 0,260657661 | 0,042520161 | 6,130213345 | 8,78E-10 | 8,25E-09 |
| ENSG00000092847.12 | AGO1 | protein_coding | 10224,507 | -0,373456566 | 0,048256183 | -7,739040708 | 1,00E-14 | 1,48E-13 |
| ENSG00000092850.12 | TEKT2 | protein_coding | 626,5054913 | -0,268238438 | 0,105268145 | -2,548144448 | 0,010829762 | 0,032037893 |
| ENSG00000092929.11 | UNC13D | protein_coding | 70,03319715 | -0,886751593 | 0,304687464 | -2,910364545 | 0,003610074 | 0,012237825 |
| ENSG00000092964.18 | DPYSL2 | protein_coding | 28259,96237 | -0,431455831 | 0,048472012 | -8,901133151 | 5,53E-19 | 1,07E-17 |
| ENSG00000092978.11 | GPATCH2 | protein_coding | 4065,360663 | 0,140085251 | 0,057329198 | 2,443523654 | 0,014544616 | 0,04131038 |
| ENSG00000093000.19 | NUP50 | protein_coding | 13155,32235 | -0,138748703 | 0,041640483 | -3,332062734 | 0,000862048 | 0,003390502 |
| ENSG00000093009.9 | CDC45 | protein_coding | 3203,369382 | -0,345089958 | 0,062664778 | -5,506920634 | 3,65E-08 | 2,85E-07 |
| ENSG00000093010.13 | COMT | protein_coding | 3338,754882 | -0,187001448 | 0,059109824 | -3,163627226 | 0,001558162 | 0,005791972 |
| ENSG00000093134.14 | VNN3 | protein_coding | 241,4258498 | 3,5996842 | 0,214997152 | 16,74293898 | 6,38E-63 | 4,61E-61 |
| ENSG00000093183.14 | SEC22C | protein_coding | 8306,430043 | -0,204180861 | 0,05532296 | -3,690707485 | 0,000223631 | 0,000994805 |
| ENSG00000093217.10 | XYLB | protein_coding | 946,8871623 | -0,283229482 | 0,092810011 | -3,051712622 | 0,002275398 | 0,00809754 |
| ENSG00000094631.20 | HDAC6 | protein_coding | 13167,65619 | 0,165720039 | 0,054252187 | 3,054624138 | 0,002253427 | 0,008026966 |
| ENSG00000094914.13 | AAAS | protein_coding | 3507,417258 | -0,131921565 | 0,055088079 | -2,394738876 | 0,016632206 | 0,04637363 |
| ENSG00000094916.16 | CBX5 | protein_coding | 38689,00007 | -1,163659833 | 0,059084833 | -19,6947299 | 2,39E-86 | 2,45E-84 |
| ENSG00000094975.14 | SUCO | protein_coding | 14349,89616 | 0,344292653 | 0,044472389 | 7,741717086 | 9,81E-15 | 1,45E-13 |
| ENSG00000095002.14 | MSH2 | protein_coding | 17045,21603 | -0,17128076 | 0,040611647 | -4,217528072 | 2,47E-05 | 0,000130171 |
| ENSG00000095015.6 | MAP3K1 | protein_coding | 2119,853302 | 0,391953438 | 0,074381639 | 5,269491812 | 1,37E-07 | 9,96E-07 |
| ENSG00000095059.16 | DHPS | protein_coding | 4071,119482 | -0,167219643 | 0,057167877 | -2,925063009 | 0,003443865 | 0,011739808 |
| ENSG00000095209.12 | TMEM38B | protein_coding | 4943,214414 | -0,227891151 | 0,056649258 | -4,022844455 | 5,75E-05 | 0,000285141 |
| ENSG00000095321.17 | CRAT | protein_coding | 3489,003538 | -0,199095694 | 0,062138004 | -3,204088963 | 0,001354906 | 0,005100783 |
| ENSG00000095380.11 | NANS | protein_coding | 2018,113147 | 0,301949784 | 0,072350854 | 4,173410081 | 3,00E-05 | 0,000155631 |
| ENSG00000095397.14 | WHRN | protein_coding | 179,4297366 | 0,597779273 | 0,188600037 | 3,169560738 | 0,001526695 | 0,005683822 |

| | | | | | | | | |
|---|---|---|---|---|---|---|---|---|
| ENSG00000095485.18 | CWF19L1 | protein_coding | 3468,818922 | 0,266656144 | 0,061186522 | 4,358086327 | 1,31E-05 | 7,22E-05 |
| ENSG00000095564.14 | BTAF1 | protein_coding | 12833,76144 | 0,230226578 | 0,043748637 | 5,262485756 | 1,42E-07 | 1,03E-06 |
| ENSG00000095585.16 | BLNK | protein_coding | 42,28347125 | 2,248242856 | 0,499343431 | 4,502397983 | 6,72E-06 | 3,86E-05 |
| ENSG00000095587.9 | TLL2 | protein_coding | 307,8585765 | 1,559787576 | 0,185930614 | 8,389084218 | 4,90E-17 | 8,38E-16 |
| ENSG00000095906.17 | NUBP2 | protein_coding | 3852,272979 | -0,351165595 | 0,059315324 | -5,920318252 | 3,21E-09 | 2,84E-08 |
| ENSG00000095932.6 | SMIM24 | protein_coding | 7171,230912 | -0,225287473 | 0,050526145 | -4,458829656 | 8,24E-06 | 4,67E-05 |
| ENSG00000095951.17 | HIVEP1 | protein_coding | 2223,656177 | 0,534999964 | 0,083861707 | 6,379550151 | 1,78E-10 | 1,80E-09 |
| ENSG00000095981.10 | KCNK16 | protein_coding | 17568,45272 | -0,150937952 | 0,055562156 | -2,716560392 | 0,006596416 | 0,020833272 |
| ENSG00000096070.19 | BRPF3 | protein_coding | 12177,41752 | 0,506374012 | 0,050345688 | 10,057942 | 8,48E-24 | 2,08E-22 |
| ENSG00000096080.11 | MRPS18A | protein_coding | 2525,372743 | 0,219077861 | 0,068507581 | 3,197863008 | 0,0013845 | 0,005200888 |
| ENSG00000096092.6 | TMEM14A | protein_coding | 5882,496177 | -0,261721973 | 0,057410651 | -4,558770308 | 5,15E-06 | 3,01E-05 |
| ENSG00000096093.16 | EFHC1 | protein_coding | 5101,7911 | -0,173442277 | 0,062017662 | -2,796659402 | 0,005163393 | 0,016800061 |
| ENSG00000096717.12 | SIRT1 | protein_coding | 6932,347901 | 0,201552234 | 0,049316799 | 4,086887972 | 4,37E-05 | 0,000221392 |
| ENSG00000096872.16 | IFT74 | protein_coding | 2565,374287 | 0,186787692 | 0,064477875 | 2,896926919 | 0,003768375 | 0,012708705 |
| ENSG00000096968.13 | JAK2 | protein_coding | 1793,236103 | 1,071024564 | 0,092839999 | 11,53624057 | 8,66E-31 | 2,86E-29 |
| ENSG00000097007.18 | ABL1 | protein_coding | 8617,034332 | 0,286907669 | 0,046417579 | 6,181013223 | 6,37E-10 | 6,10E-09 |
| ENSG00000097021.19 | ACOT7 | protein_coding | 2520,723869 | 0,305276395 | 0,064434679 | 4,737765453 | 2,16E-06 | 1,34E-05 |
| ENSG00000097033.14 | SH3GLB1 | protein_coding | 18368,40469 | 0,243308104 | 0,044910536 | 5,417617425 | 6,04E-08 | 4,58E-07 |
| ENSG00000097046.13 | CDC7 | protein_coding | 9491,239625 | -0,168257346 | 0,048554235 | -3,465348514 | 0,000529545 | 0,002180666 |
| ENSG00000099139.13 | PCSK5 | protein_coding | 1099,976531 | 0,299906893 | 0,091699739 | 3,270531591 | 0,001073456 | 0,00413192 |
| ENSG00000099194.6 | SCD | protein_coding | 297872,0535 | -0,225390011 | 0,057466242 | -3,922128954 | 8,78E-05 | 0,00042232 |
| ENSG00000099203.7 | TMED1 | protein_coding | 2634,497888 | 0,306578869 | 0,057139354 | 5,365459098 | 8,07E-08 | 6,00E-07 |
| ENSG00000099204.20 | ABLIM1 | protein_coding | 4303,714446 | -0,236254126 | 0,058296079 | -4,052658951 | 5,06E-05 | 0,000253408 |
| ENSG00000099219.14 | ERMP1 | protein_coding | 2167,783061 | -0,248768699 | 0,061600911 | -4,038393154 | 5,38E-05 | 0,000268425 |
| ENSG00000099250.18 | NRP1 | protein_coding | 1336,266554 | 1,51620829 | 0,236861733 | 6,401237842 | 1,54E-10 | 1,58E-09 |
| ENSG00000099256.19 | PRTFDC1 | protein_coding | 2332,310359 | -0,321511922 | 0,07470735 | -4,303618351 | 1,68E-05 | 9,08E-05 |
| ENSG00000099260.11 | PALMD | protein_coding | 81,54414883 | -0,854244413 | 0,260891638 | -3,274326539 | 0,001059141 | 0,004082928 |
| ENSG00000099282.10 | TSPAN15 | protein_coding | 2134,46119 | -0,198850775 | 0,061640989 | -3,225950435 | 0,001255551 | 0,004757777 |
| ENSG00000099284.14 | H2AFY2 | protein_coding | 5386,906996 | -0,146249232 | 0,059292036 | -2,46659151 | 0,013640587 | 0,039132803 |
| ENSG00000099326.8 | MZF1 | protein_coding | 2773,78871 | -0,192902105 | 0,063394028 | -3,04290659 | 0,002343051 | 0,008312013 |
| ENSG00000099338.23 | CATSPERG | protein_coding | 125,3437784 | 1,370043859 | 0,266157549 | 5,147492013 | 2,64E-07 | 1,85E-06 |
| ENSG00000099377.14 | HSD3B7 | protein_coding | 304,2775271 | 0,442769523 | 0,147485331 | 3,002125837 | 0,002681013 | 0,009372096 |
| ENSG00000099381.18 | SETD1A | protein_coding | 3307,336211 | -0,274582063 | 0,082127153 | -3,343377326 | 0,000827653 | 0,003267784 |
| ENSG00000099385.11 | BCL7C | protein_coding | 875,3792646 | -0,418268934 | 0,092114518 | -4,540749332 | 5,61E-06 | 3,26E-05 |
| ENSG00000099617.4 | EFNA2 | protein_coding | 249,5934231 | -0,681662509 | 0,164876639 | -4,134378974 | 3,56E-05 | 0,000182448 |
| ENSG00000099622.14 | CIRBP | protein_coding | 19085,71162 | -0,202672281 | 0,052729288 | -3,843637749 | 0,000121224 | 0,000569295 |

| | | | | | | | | |
|---|---|---|---|---|---|---|---|---|
| ENSG00000099769.5 | IGFALS | protein_coding | 991,1173252 | -0,694361149 | 0,101582117 | -6,835466404 | 8,17E-12 | 9,45E-11 |
| ENSG00000099783.12 | HNRNPM | protein_coding | 28089,09626 | -0,160894935 | 0,041301103 | -3,89565707 | 9,79E-05 | 0,000466336 |
| ENSG00000099795.7 | NDUFB7 | protein_coding | 3314,839233 | -0,231595346 | 0,090037335 | -2,572214588 | 0,010105024 | 0,030202361 |
| ENSG00000099797.15 | TECR | protein_coding | 13894,5759 | -0,197960383 | 0,043820096 | -4,517570784 | 6,26E-06 | 3,61E-05 |
| ENSG00000099800.8 | TIMM13 | protein_coding | 3227,730229 | -0,274000382 | 0,083836315 | -3,26827796 | 0,00108204 | 0,004160514 |
| ENSG00000099804.9 | CDC34 | protein_coding | 3778,64347 | -0,388584651 | 0,059147447 | -6,569761963 | 5,04E-11 | 5,40E-10 |
| ENSG00000099812.9 | MISP | protein_coding | 4575,808386 | 0,585358726 | 0,052083211 | 11,238914 | 2,63E-29 | 8,01E-28 |
| ENSG00000099814.16 | CEP170B | protein_coding | 3945,667951 | 0,306589312 | 0,053649037 | 5,714721643 | 1,10E-08 | 9,09E-08 |
| ENSG00000099817.12 | POLR2E | protein_coding | 7346,850856 | -0,220101318 | 0,053390311 | -4,122495508 | 3,75E-05 | 0,000191513 |
| ENSG00000099821.14 | POLRMT | protein_coding | 2802,236067 | -0,208557465 | 0,062944484 | -3,313355688 | 0,000921837 | 0,003601739 |
| ENSG00000099822.3 | HCN2 | protein_coding | 964,3132826 | -0,677158538 | 0,102684396 | -6,594561213 | 4,27E-11 | 4,61E-10 |
| ENSG00000099834.18 | CDHR5 | protein_coding | 13254,87871 | 0,624323255 | 0,054839027 | 11,38465228 | 4,99E-30 | 1,57E-28 |
| ENSG00000099860.9 | GADD45B | protein_coding | 226,5516142 | 0,832187767 | 0,193245746 | 4,306370441 | 1,66E-05 | 8,97E-05 |
| ENSG00000099864.18 | PALM | protein_coding | 2020,985736 | -0,35223339 | 0,074201217 | -4,747002845 | 2,06E-06 | 1,28E-05 |
| ENSG00000099889.14 | ARVCF | protein_coding | 3577,692368 | -0,306947484 | 0,071428841 | -4,297248583 | 1,73E-05 | 9,32E-05 |
| ENSG00000099901.17 | RANBP1 | protein_coding | 7408,469915 | -0,286135341 | 0,061785509 | -4,631107585 | 3,64E-06 | 2,17E-05 |
| ENSG00000099904.15 | ZDHHC8 | protein_coding | 1951,30036 | -0,310011862 | 0,070633104 | -4,389044849 | 1,14E-05 | 6,32E-05 |
| ENSG00000099910.17 | KLHL22 | protein_coding | 2663,312963 | 0,220336337 | 0,062726095 | 3,512674223 | 0,000443621 | 0,001864351 |
| ENSG00000099917.17 | MED15 | protein_coding | 2641,466265 | 0,247322779 | 0,062995921 | 3,926012558 | 8,64E-05 | 0,000416543 |
| ENSG00000099949.21 | LZTR1 | protein_coding | 4318,517688 | -0,215536901 | 0,079143574 | -2,723365768 | 0,006462047 | 0,020466331 |
| ENSG00000099953.10 | MMP11 | protein_coding | 562,6838099 | -0,398931311 | 0,10630655 | -3,75265034 | 0,000174975 | 0,000793734 |
| ENSG00000099956.19 | SMARCB1 | protein_coding | 7454,585929 | -0,21331861 | 0,047274862 | -4,512305324 | 6,41E-06 | 3,70E-05 |
| ENSG00000099960.13 | SLC7A4 | protein_coding | 775,0350116 | -0,450652208 | 0,098827658 | -4,559980637 | 5,12E-06 | 2,99E-05 |
| ENSG00000099968.17 | BCL2L13 | protein_coding | 6888,664579 | 0,370986303 | 0,058534822 | 6,337873543 | 2,33E-10 | 2,34E-09 |
| ENSG00000099977.14 | DDT | protein_coding | 2942,188234 | -0,382220079 | 0,08342262 | -4,581731877 | 4,61E-06 | 2,72E-05 |
| ENSG00000100027.15 | YPEL1 | protein_coding | 1130,593404 | -0,441778908 | 0,081979679 | -5,3888831 | 7,09E-08 | 5,32E-07 |
| ENSG00000100028.12 | SNRPD3 | protein_coding | 14791,38297 | -0,46082174 | 0,052160672 | -8,834658873 | 1,00E-18 | 1,91E-17 |
| ENSG00000100030.14 | MAPK1 | protein_coding | 23086,73456 | -0,551926206 | 0,042640171 | -12,94380847 | 2,55E-38 | 1,08E-36 |
| ENSG00000100033.16 | PRODH | protein_coding | 8920,708207 | -0,34166021 | 0,063971033 | -5,340858077 | 9,25E-08 | 6,84E-07 |
| ENSG00000100038.19 | TOP3B | protein_coding | 2193,164503 | 0,201832814 | 0,069119013 | 2,920076641 | 0,003499453 | 0,011904763 |
| ENSG00000100056.12 | ESS2 | protein_coding | 2332,909894 | 0,198182377 | 0,064129826 | 3,090330814 | 0,001999337 | 0,007224766 |
| ENSG00000100058.12 | CRYBB2P1 | transcribed_unproc | 2179,025434 | -0,347536643 | 0,079743657 | -4,358172891 | 1,31E-05 | 7,22E-05 |
| ENSG00000100060.17 | MFNG | protein_coding | 33,71561583 | -1,107294291 | 0,42808598 | -2,586616576 | 0,009692338 | 0,029143045 |
| ENSG00000100068.13 | LRP5L | protein_coding | 1055,234077 | 0,488585983 | 0,084391339 | 5,789527545 | 7,06E-09 | 5,98E-08 |
| ENSG00000100079.7 | LGALS2 | protein_coding | 914,4864778 | -0,777314511 | 0,110194273 | -7,054037268 | 1,74E-12 | 2,16E-11 |
| ENSG00000100084.14 | HIRA | protein_coding | 13207,26828 | 1,06074086 | 0,047562763 | 22,30191849 | 3,54E-110 | 4,50E-108 |

| | | | | | | | | |
|---|---|---|---|---|---|---|---|---|
| ENSG00000100105.18 | PATZ1 | protein_coding | 4375,276232 | 0,209764746 | 0,052005125 | 4,033539874 | 5,49E-05 | 0,000273608 |
| ENSG00000100106.22 | TRIOBP | protein_coding | 2156,580283 | -0,163859248 | 0,069316696 | -2,363921792 | 0,018082629 | 0,049831281 |
| ENSG00000100109.17 | TFIP11 | protein_coding | 4698,000309 | 0,190148175 | 0,057369372 | 3,314454505 | 0,000918221 | 0,003589483 |
| ENSG00000100129.18 | EIF3L | protein_coding | 19787,08356 | -0,845883727 | 0,04603873 | -18,37330723 | 2,15E-75 | 1,90E-73 |
| ENSG00000100139.13 | MICALL1 | protein_coding | 786,3345079 | 0,550250303 | 0,092167613 | 5,970104719 | 2,37E-09 | 2,12E-08 |
| ENSG00000100147.13 | CCDC134 | protein_coding | 665,1160395 | 0,265334021 | 0,101176332 | 2,622491014 | 0,008728957 | 0,026641368 |
| ENSG00000100162.15 | CENPM | protein_coding | 2016,340178 | -0,41085259 | 0,070526225 | -5,825529312 | 5,69E-09 | 4,89E-08 |
| ENSG00000100167.20 | sept-03 | protein_coding | 10655,02072 | -0,475803364 | 0,053531819 | -8,888234545 | 6,21E-19 | 1,19E-17 |
| ENSG00000100196.11 | KDELR3 | protein_coding | 215,4214763 | -0,665729523 | 0,172989674 | -3,848377229 | 0,000118903 | 0,000558978 |
| ENSG00000100206.10 | DMC1 | protein_coding | 2772,225432 | -0,574416047 | 0,06738503 | -8,524386562 | 1,54E-17 | 2,72E-16 |
| ENSG00000100209.10 | HSCB | protein_coding | 1140,788893 | 0,185841275 | 0,077368961 | 2,402013326 | 0,016305114 | 0,045580206 |
| ENSG00000100211.10 | CBY1 | protein_coding | 2973,242313 | -0,224890408 | 0,059630779 | -3,771381328 | 0,000162346 | 0,000741078 |
| ENSG00000100216.6 | TOMM22 | protein_coding | 5986,553775 | -0,368684913 | 0,054305683 | -6,789066807 | 1,13E-11 | 1,29E-10 |
| ENSG00000100219.16 | XBP1 | protein_coding | 20433,96377 | 0,49257459 | 0,041326273 | 11,91916313 | 9,40E-33 | 3,38E-31 |
| ENSG00000100220.12 | RTCB | protein_coding | 12145,24467 | 0,178107851 | 0,049374335 | 3,607296169 | 0,000309404 | 0,00134111 |
| ENSG00000100226.16 | GTPBP1 | protein_coding | 7559,635655 | 1,181572844 | 0,052385543 | 22,55532304 | 1,19E-112 | 1,54E-110 |
| ENSG00000100234.11 | TIMP3 | protein_coding | 2558,325678 | -0,607383308 | 0,067402984 | -9,011222782 | 2,04E-19 | 4,00E-18 |
| ENSG00000100242.15 | SUN2 | protein_coding | 5148,524635 | -0,579905633 | 0,050733995 | -11,43031676 | 2,95E-30 | 9,43E-29 |
| ENSG00000100243.21 | CYB5R3 | protein_coding | 10284,92664 | -0,261341323 | 0,04266551 | -6,125353294 | 9,05E-10 | 8,48E-09 |
| ENSG00000100263.14 | RHBDD3 | protein_coding | 2400,780121 | 1,244567148 | 0,064166401 | 19,39593201 | 8,35E-84 | 8,10E-82 |
| ENSG00000100281.14 | HMGXB4 | protein_coding | 6260,032815 | -0,156842595 | 0,054809894 | -2,861574478 | 0,004215424 | 0,014036241 |
| ENSG00000100284.21 | TOM1 | protein_coding | 2563,2929 | 0,277320297 | 0,066992948 | 4,139544598 | 3,48E-05 | 0,000178633 |
| ENSG00000100290.3 | BIK | protein_coding | 468,3250643 | -0,754489073 | 0,127740242 | -5,906432174 | 3,50E-09 | 3,07E-08 |
| ENSG00000100292.17 | HMOX1 | protein_coding | 75,23339257 | 2,006347134 | 0,390698189 | 5,135286496 | 2,82E-07 | 1,97E-06 |
| ENSG00000100294.12 | MCAT | protein_coding | 1388,631695 | -0,280206837 | 0,074013857 | -3,785869932 | 0,000153172 | 0,00070434 |
| ENSG00000100297.16 | MCM5 | protein_coding | 10574,65699 | -0,203648502 | 0,057346013 | -3,551223352 | 0,000383445 | 0,001629405 |
| ENSG00000100299.18 | ARSA | protein_coding | 2892,781223 | 0,447771251 | 0,064575816 | 6,934039357 | 4,09E-12 | 4,89E-11 |
| ENSG00000100302.7 | RASD2 | protein_coding | 16,47850155 | -1,506832464 | 0,601398609 | -2,50554697 | 0,012226216 | 0,035573926 |
| ENSG00000100304.13 | TTLL12 | protein_coding | 3698,768859 | -0,294958187 | 0,061786386 | -4,773837804 | 1,81E-06 | 1,13E-05 |
| ENSG00000100307.13 | CBX7 | protein_coding | 1124,839601 | -0,377712146 | 0,079104121 | -4,774873169 | 1,80E-06 | 1,13E-05 |
| ENSG00000100314.4 | CABP7 | protein_coding | 2583,886074 | -0,539322287 | 0,093117582 | -5,791841631 | 6,96E-09 | 5,91E-08 |
| ENSG00000100316.16 | RPL3 | protein_coding | 94270,333 | -0,769959413 | 0,047845313 | -16,09268208 | 2,87E-58 | 1,95E-56 |
| ENSG00000100324.14 | TAB1 | protein_coding | 3998,003558 | 0,19772368 | 0,059990714 | 3,29590476 | 0,000981053 | 0,003811273 |
| ENSG00000100330.15 | MTMR3 | protein_coding | 10483,9743 | 0,363757801 | 0,049055793 | 7,415185477 | 1,21E-13 | 1,66E-12 |
| ENSG00000100336.17 | APOL4 | protein_coding | 56,2581413 | 1,187041019 | 0,37544352 | 3,161703307 | 0,001568493 | 0,005825857 |
| ENSG00000100341.11 | PNPLA5 | protein_coding | 59,66932469 | -0,961433921 | 0,344606094 | -2,789950435 | 0,005271611 | 0,017087946 |

| | | | | | | | | |
|---|---|---|---|---|---|---|---|---|
| ENSG00000100342.20 | APOL1 | protein_coding | 8252,959592 | 5,802487216 | 0,082850588 | 70,03555874 | 0 | 0 |
| ENSG00000100347.15 | SAMM50 | protein_coding | 2742,257805 | -0,199391 | 0,060434361 | -3,299298555 | 0,000969268 | 0,003770696 |
| ENSG00000100348.10 | TXN2 | protein_coding | 6483,326677 | -0,229494267 | 0,053902253 | -4,257600677 | 2,07E-05 | 0,000110007 |
| ENSG00000100350.14 | FOXRED2 | protein_coding | 8899,082718 | -0,335362363 | 0,05742091 | -5,840422332 | 5,21E-09 | 4,49E-08 |
| ENSG00000100354.21 | TNRC6B | protein_coding | 4772,392395 | 0,277100716 | 0,082946012 | 3,340735847 | 0,000835567 | 0,003296717 |
| ENSG00000100372.15 | SLC25A17 | protein_coding | 3909,045999 | -0,208994264 | 0,057818796 | -3,614642307 | 0,000300763 | 0,001306167 |
| ENSG00000100376.12 | FAM118A | protein_coding | 1176,874904 | 0,262804362 | 0,108637081 | 2,419103667 | 0,015558805 | 0,043741874 |
| ENSG00000100379.17 | KCTD17 | protein_coding | 1977,69526 | -0,204516028 | 0,069145805 | -2,957750319 | 0,00309893 | 0,010664091 |
| ENSG00000100380.14 | ST13 | protein_coding | 33334,60239 | -0,12921783 | 0,041268798 | -3,131126572 | 0,001741371 | 0,006395255 |
| ENSG00000100393.13 | EP300 | protein_coding | 11920,62158 | 0,380703795 | 0,058274211 | 6,53297212 | 6,45E-11 | 6,84E-10 |
| ENSG00000100412.16 | ACO2 | protein_coding | 6627,648169 | -0,184916173 | 0,048465702 | -3,81540279 | 0,000135961 | 0,000630739 |
| ENSG00000100418.8 | DESI1 | protein_coding | 5376,257166 | 0,220049212 | 0,057390116 | 3,834270236 | 0,000125938 | 0,000588974 |
| ENSG00000100422.14 | CERK | protein_coding | 6548,942935 | -1,855192484 | 0,0600778 | -30,87983368 | 2,23E-209 | 4,60E-207 |
| ENSG00000100425.18 | BRD1 | protein_coding | 5145,293072 | 0,151258907 | 0,054428586 | 2,779034304 | 0,005452077 | 0,017613975 |
| ENSG00000100462.16 | PRMT5 | protein_coding | 5983,455809 | 0,274732202 | 0,065534837 | 4,192155107 | 2,76E-05 | 0,000144073 |
| ENSG00000100479.13 | POLE2 | protein_coding | 2174,267822 | -0,335455619 | 0,065312994 | -5,136123731 | 2,80E-07 | 1,96E-06 |
| ENSG00000100485.12 | SOS2 | protein_coding | 7974,12316 | -0,161183203 | 0,047471928 | -3,395337229 | 0,000685441 | 0,002757076 |
| ENSG00000100490.9 | CDKL1 | protein_coding | 133,1770779 | 0,885626138 | 0,268071646 | 3,303691951 | 0,000954206 | 0,003719174 |
| ENSG00000100503.23 | NIN | protein_coding | 5988,731882 | 0,23668465 | 0,051266515 | 4,616749333 | 3,90E-06 | 2,32E-05 |
| ENSG00000100505.13 | TRIM9 | protein_coding | 1513,659499 | -0,431245544 | 0,077868798 | -5,538104553 | 3,06E-08 | 2,41E-07 |
| ENSG00000100522.10 | GNPNAT1 | protein_coding | 9396,842638 | -0,295963288 | 0,053644192 | -5,517154355 | 3,45E-08 | 2,70E-07 |
| ENSG00000100526.20 | CDKN3 | protein_coding | 3331,453324 | -0,748792263 | 0,067063382 | -11,16544134 | 6,02E-29 | 1,82E-27 |
| ENSG00000100528.12 | CNIH1 | protein_coding | 15921,22429 | -0,211177396 | 0,051872645 | -4,0710744 | 4,68E-05 | 0,000235649 |
| ENSG00000100564.9 | PIGH | protein_coding | 2109,156323 | 0,191119877 | 0,062214949 | 3,071928526 | 0,002126806 | 0,00762905 |
| ENSG00000100567.13 | PSMA3 | protein_coding | 12913,92229 | 0,358549444 | 0,044645158 | 8,031093608 | 9,66E-16 | 1,53E-14 |
| ENSG00000100568.11 | VTI1B | protein_coding | 7093,181873 | -0,161966082 | 0,058587549 | -2,764513701 | 0,005700771 | 0,018326001 |
| ENSG00000100578.16 | KIAA0586 | protein_coding | 5457,859799 | -0,175623997 | 0,054286221 | -3,235148717 | 0,001215794 | 0,004617227 |
| ENSG00000100580.8 | TMED8 | protein_coding | 17083,60126 | -0,31248005 | 0,049077199 | -6,367112589 | 1,93E-10 | 1,95E-09 |
| ENSG00000100596.6 | SPTLC2 | protein_coding | 9929,153507 | -0,343975096 | 0,049223453 | -6,988032662 | 2,79E-12 | 3,38E-11 |
| ENSG00000100600.15 | LGMN | protein_coding | 8873,253131 | 1,178796429 | 0,071286126 | 16,53612698 | 2,02E-61 | 1,43E-59 |
| ENSG00000100601.10 | ALKBH1 | protein_coding | 2232,072271 | 0,23386193 | 0,064014649 | 3,65325643 | 0,000258935 | 0,00113791 |
| ENSG00000100603.13 | SNW1 | protein_coding | 9407,29145 | 0,107688714 | 0,045159754 | 2,384616911 | 0,017096915 | 0,047510426 |
| ENSG00000100604.13 | CHGA | protein_coding | 1341239,439 | 0,268130348 | 0,041395624 | 6,477263113 | 9,34E-11 | 9,76E-10 |
| ENSG00000100605.17 | ITPK1 | protein_coding | 6502,395574 | -0,149055686 | 0,053858178 | -2,767558994 | 0,005647782 | 0,018176427 |
| ENSG00000100625.9 | SIX4 | protein_coding | 1419,357159 | 0,691651814 | 0,076217303 | 9,074734858 | 1,14E-19 | 2,26E-18 |
| ENSG00000100628.11 | ASB2 | protein_coding | 47,03933494 | 1,080136091 | 0,368030681 | 2,934907728 | 0,003336469 | 0,011404795 |

| | | | | | | | | |
|---|---|---|---|---|---|---|---|---|
| ENSG00000100697.14 | DICER1 | protein_coding | 23936,66416 | -0,336511221 | 0,046322426 | -7,264542231 | 3,74E-13 | 4,90E-12 |
| ENSG00000100731.15 | PCNX1 | protein_coding | 12891,32953 | 0,484940553 | 0,047912314 | 10,12141787 | 4,44E-24 | 1,11E-22 |
| ENSG00000100744.15 | GSKIP | protein_coding | 8670,498791 | 0,126573042 | 0,052566165 | 2,40788047 | 0,016045433 | 0,04498277 |
| ENSG00000100767.16 | PAPLN | protein_coding | 936,1149446 | -0,345492602 | 0,088467806 | -3,90529185 | 9,41E-05 | 0,000449949 |
| ENSG00000100796.17 | PPP4R3A | protein_coding | 19490,61351 | 0,233781121 | 0,043257376 | 5,4044222 | 6,50E-08 | 4,91E-07 |
| ENSG00000100804.18 | PSMB5 | protein_coding | 15455,74548 | -0,327201172 | 0,051972589 | -6,295648851 | 3,06E-10 | 3,04E-09 |
| ENSG00000100813.14 | ACIN1 | protein_coding | 24399,01799 | 0,139294208 | 0,043486405 | 3,203166796 | 0,001359252 | 0,005114579 |
| ENSG00000100814.18 | CCNB1IP1 | protein_coding | 5456,09595 | -0,976953614 | 0,050125267 | -19,49024262 | 1,33E-84 | 1,31E-82 |
| ENSG00000100815.12 | TRIP11 | protein_coding | 9026,95161 | 0,130416185 | 0,048230843 | 2,703999716 | 0,006851035 | 0,021558778 |
| ENSG00000100823.12 | APEX1 | protein_coding | 17161,33674 | -0,185667178 | 0,040734398 | -4,557994876 | 5,16E-06 | 3,02E-05 |
| ENSG00000100852.13 | ARHGAP5 | protein_coding | 16986,71124 | 0,330665771 | 0,042156859 | 7,843700324 | 4,37E-15 | 6,62E-14 |
| ENSG00000100867.14 | DHRS2 | protein_coding | 2073,764966 | -0,190834926 | 0,063757437 | -2,993139845 | 0,002761232 | 0,009621141 |
| ENSG00000100883.12 | SRP54 | protein_coding | 14089,02292 | 0,141692577 | 0,044578644 | 3,178485564 | 0,001480466 | 0,005533678 |
| ENSG00000100888.13 | CHD8 | protein_coding | 16925,60548 | 0,4834293 | 0,041982292 | 11,51507636 | 1,11E-30 | 3,62E-29 |
| ENSG00000100889.11 | PCK2 | protein_coding | 1083,15249 | 0,506293334 | 0,087699378 | 5,773055043 | 7,78E-09 | 6,55E-08 |
| ENSG00000100897.17 | DCAF11 | protein_coding | 6624,634186 | 0,188470473 | 0,058156464 | 3,240748508 | 0,001192163 | 0,004536669 |
| ENSG00000100902.10 | PSMA6 | protein_coding | 15702,34978 | 0,527168731 | 0,052296337 | 10,08041413 | 6,74E-24 | 1,67E-22 |
| ENSG00000100906.10 | NFKBIA | protein_coding | 3621,709642 | 1,197097027 | 0,058156664 | 20,58400438 | 3,82E-94 | 4,21E-92 |
| ENSG00000100908.14 | EMC9 | protein_coding | 1061,817946 | 0,441327787 | 0,083806961 | 5,266003945 | 1,39E-07 | 1,01E-06 |
| ENSG00000100911.16 | PSME2 | protein_coding | 18495,51565 | 2,359556998 | 0,042521478 | 55,49094445 | 0 | 0 |
| ENSG00000100916.14 | BRMS1L | protein_coding | 2713,389073 | 0,239290202 | 0,060285199 | 3,969302685 | 7,21E-05 | 0,000352033 |
| ENSG00000100918.13 | REC8 | protein_coding | 14212,73179 | 0,443717954 | 0,054051843 | 8,209117873 | 2,23E-16 | 3,66E-15 |
| ENSG00000100926.15 | TM9SF1 | protein_coding | 7017,227357 | 0,397637544 | 0,046326228 | 8,58342155 | 9,21E-18 | 1,65E-16 |
| ENSG00000100934.15 | SEC23A | protein_coding | 5653,998642 | 0,42502608 | 0,052061346 | 8,163947141 | 3,24E-16 | 5,28E-15 |
| ENSG00000100941.9 | PNN | protein_coding | 19408,32598 | -0,16984866 | 0,04488748 | -3,78387602 | 0,000154405 | 0,000709285 |
| ENSG00000100968.14 | NFATC4 | protein_coding | 720,9236628 | -0,452449216 | 0,100340421 | -4,509142106 | 6,51E-06 | 3,75E-05 |
| ENSG00000100979.15 | PLTP | protein_coding | 126,3051947 | 0,500851562 | 0,21026003 | 2,382057888 | 0,017216191 | 0,047788783 |
| ENSG00000100991.12 | TRPC4AP | protein_coding | 22796,98915 | 0,135970869 | 0,042152991 | 3,225651772 | 0,001256862 | 0,004761943 |
| ENSG00000100994.12 | PYGB | protein_coding | 5485,647231 | 0,259586671 | 0,051449906 | 5,045425559 | 4,53E-07 | 3,09E-06 |
| ENSG00000100997.19 | ABHD12 | protein_coding | 9464,679371 | 0,113441842 | 0,045296364 | 2,504435919 | 0,012264682 | 0,035664613 |
| ENSG00000101000.6 | PROCR | protein_coding | 568,7208228 | 1,510027248 | 0,120016535 | 12,58182675 | 2,66E-36 | 1,07E-34 |
| ENSG00000101003.10 | GINS1 | protein_coding | 4592,915541 | -0,211427015 | 0,051877754 | -4,075485153 | 4,59E-05 | 0,00023164 |
| ENSG00000101004.15 | NINL | protein_coding | 4482,302982 | 0,28725295 | 0,05452434 | 5,268343464 | 1,38E-07 | 1,00E-06 |
| ENSG00000101052.12 | IFT52 | protein_coding | 6013,20365 | -0,143652123 | 0,049618317 | -2,895142973 | 0,003789859 | 0,012768437 |
| ENSG00000101076.16 | HNF4A | protein_coding | 17168,42452 | 0,143519034 | 0,043167485 | 3,324702207 | 0,00088513 | 0,00347158 |
| ENSG00000101079.21 | NDRG3 | protein_coding | 10345,90664 | -0,327462673 | 0,047411661 | -6,906796117 | 4,96E-12 | 5,87E-11 |

| | | | | | | | | |
|---|---|---|---|---|---|---|---|---|
| ENSG00000101084.18 | RAB5IF | protein_coding | 8017,573272 | -0,153463699 | 0,050492366 | -3,03934459 | 0,002370935 | 0,008396377 |
| ENSG00000101096.20 | NFATC2 | protein_coding | 182,9949439 | 0,992362908 | 0,199439909 | 4,975748903 | 6,50E-07 | 4,34E-06 |
| ENSG00000101098.12 | RIMS4 | protein_coding | 283,3159176 | -0,744637635 | 0,153254807 | -4,858820756 | 1,18E-06 | 7,59E-06 |
| ENSG00000101126.17 | ADNP | protein_coding | 35252,74116 | 0,138598201 | 0,042279423 | 3,278147902 | 0,001044906 | 0,004032887 |
| ENSG00000101158.14 | NELFCD | protein_coding | 25560,93036 | -0,178826766 | 0,043023145 | -4,156524737 | 3,23E-05 | 0,000166894 |
| ENSG00000101161.8 | PRPF6 | protein_coding | 12618,06546 | 0,124029399 | 0,043510932 | 2,850534181 | 0,004364586 | 0,014458052 |
| ENSG00000101162.3 | TUBB1 | protein_coding | 4087,201196 | -0,450860822 | 0,078907919 | -5,713758838 | 1,11E-08 | 9,13E-08 |
| ENSG00000101166.16 | PRELID3B | protein_coding | 64068,42593 | -0,228187953 | 0,047136225 | -4,841031558 | 1,29E-06 | 8,27E-06 |
| ENSG00000101180.16 | HRH3 | protein_coding | 55,44251216 | -1,353016612 | 0,332899082 | -4,064344684 | 4,82E-05 | 0,00024174 |
| ENSG00000101181.17 | MTG2 | protein_coding | 4068,279962 | 0,486292034 | 0,058665057 | 8,289296155 | 1,14E-16 | 1,91E-15 |
| ENSG00000101182.15 | PSMA7 | protein_coding | 15902,18535 | -0,182339082 | 0,045323057 | -4,023097604 | 5,74E-05 | 0,00028496 |
| ENSG00000101189.7 | MRGBP | protein_coding | 2333,077764 | 0,326023608 | 0,063131423 | 5,164204979 | 2,41E-07 | 1,70E-06 |
| ENSG00000101193.8 | GID8 | protein_coding | 12183,09103 | -0,310169451 | 0,051454133 | -6,028076535 | 1,66E-09 | 1,51E-08 |
| ENSG00000101204.17 | CHRNA4 | protein_coding | 154,1542363 | -1,07134453 | 0,250830973 | -4,271181177 | 1,94E-05 | 0,000103959 |
| ENSG00000101210.12 | EEF1A2 | protein_coding | 66522,68093 | -0,117286905 | 0,046753536 | -2,508621074 | 0,012120342 | 0,035293245 |
| ENSG00000101220.17 | C20orf27 | protein_coding | 1111,978185 | -0,504133946 | 0,086263444 | -5,844120307 | 5,09E-09 | 4,40E-08 |
| ENSG00000101224.17 | CDC25B | protein_coding | 10288,80215 | -0,458240983 | 0,047972826 | -9,552094804 | 1,27E-21 | 2,81E-20 |
| ENSG00000101255.11 | TRIB3 | protein_coding | 4698,928812 | 0,309625902 | 0,057474673 | 5,387171204 | 7,16E-08 | 5,37E-07 |
| ENSG00000101280.8 | ANGPT4 | protein_coding | 29,16997116 | -1,187208105 | 0,457768349 | -2,593469184 | 0,009501305 | 0,028668215 |
| ENSG00000101294.17 | HM13 | protein_coding | 19732,88064 | 0,175708177 | 0,046910435 | 3,745609655 | 0,000179956 | 0,000813544 |
| ENSG00000101298.15 | SNPH | protein_coding | 1762,129738 | 0,588825866 | 0,074350902 | 7,919552412 | 2,38E-15 | 3,67E-14 |
| ENSG00000101331.16 | CCM2L | protein_coding | 204,8250491 | 0,444362644 | 0,171823317 | 2,586160314 | 0,009705178 | 0,029173858 |
| ENSG00000101333.16 | PLCB4 | protein_coding | 6076,411477 | -0,195816709 | 0,067133759 | -2,916814288 | 0,003536263 | 0,012009183 |
| ENSG00000101342.10 | TLDC2 | protein_coding | 1280,249511 | -0,406839413 | 0,087725443 | -4,637644413 | 3,52E-06 | 2,11E-05 |
| ENSG00000101346.14 | POFUT1 | protein_coding | 17478,28424 | -0,132882771 | 0,053259708 | -2,494996249 | 0,012595846 | 0,036498443 |
| ENSG00000101347.9 | SAMHD1 | protein_coding | 33306,17352 | 2,470556317 | 0,047008894 | 52,55508336 | 0 | 0 |
| ENSG00000101350.8 | KIF3B | protein_coding | 30332,6722 | -0,235387441 | 0,047938857 | -4,91015968 | 9,10E-07 | 5,94E-06 |
| ENSG00000101367.9 | MAPRE1 | protein_coding | 73179,36375 | -0,143698945 | 0,045362574 | -3,16778641 | 0,001536043 | 0,005716732 |
| ENSG00000101384.12 | JAG1 | protein_coding | 894,1582006 | 0,924693623 | 0,107755385 | 8,581414486 | 9,37E-18 | 1,68E-16 |
| ENSG00000101391.21 | CDK5RAP1 | protein_coding | 4220,026889 | 0,152765466 | 0,062846214 | 2,430782332 | 0,015066263 | 0,042577088 |
| ENSG00000101400.5 | SNTA1 | protein_coding | 1532,311625 | -0,237992676 | 0,081935443 | -2,904636451 | 0,003676799 | 0,012435923 |
| ENSG00000101412.13 | E2F1 | protein_coding | 10882,79139 | -0,43291128 | 0,047239277 | -9,164223312 | 4,99E-20 | 1,01E-18 |
| ENSG00000101417.12 | PXMP4 | protein_coding | 4453,589142 | -0,927861959 | 0,067844931 | -13,67621639 | 1,41E-42 | 6,86E-41 |
| ENSG00000101438.4 | SLC32A1 | protein_coding | 330,5189384 | 0,494283131 | 0,129271327 | 3,823609954 | 0,000131512 | 0,00061237 |
| ENSG00000101444.13 | AHCY | protein_coding | 33627,75044 | -0,593686066 | 0,044965054 | -13,20327696 | 8,40E-40 | 3,78E-38 |
| ENSG00000101445.10 | PPP1R16B | protein_coding | 234,3810219 | -0,741767052 | 0,187194807 | -3,962540752 | 7,42E-05 | 0,00036153 |

| | | | | | | | | |
|---|---|---|---|---|---|---|---|---|
| ENSG00000101447.15 | FAM83D | protein_coding | 1883,769802 | -0,952388984 | 0,077260965 | -12,3269103 | 6,49E-35 | 2,47E-33 |
| ENSG00000101460.13 | MAP1LC3A | protein_coding | 2573,909886 | -0,292279574 | 0,065467681 | -4,464486457 | 8,03E-06 | 4,55E-05 |
| ENSG00000101464.10 | PIGU | protein_coding | 5548,665743 | -0,319032948 | 0,054036132 | -5,904067096 | 3,55E-09 | 3,11E-08 |
| ENSG00000101470.10 | TNNC2 | protein_coding | 95,53381679 | -0,739363633 | 0,262732059 | -2,814135568 | 0,00489086 | 0,016005868 |
| ENSG00000101474.12 | APMAP | protein_coding | 20755,42041 | -0,243008162 | 0,041401107 | -5,869605441 | 4,37E-09 | 3,80E-08 |
| ENSG00000101542.10 | CDH20 | protein_coding | 117,8563991 | 0,792359234 | 0,226060572 | 3,505074891 | 0,000456479 | 0,00191303 |
| ENSG00000101544.9 | ADNP2 | protein_coding | 4323,052383 | 0,255679252 | 0,059171449 | 4,320990183 | 1,55E-05 | 8,43E-05 |
| ENSG00000101557.15 | USP14 | protein_coding | 15742,63636 | -0,12094602 | 0,041362868 | -2,924024022 | 0,003455381 | 0,011773716 |
| ENSG00000101577.9 | LPIN2 | protein_coding | 15267,59589 | 0,199091577 | 0,046680226 | 4,265008807 | 2,00E-05 | 0,000106723 |
| ENSG00000101596.15 | SMCHD1 | protein_coding | 22676,01815 | 0,674540133 | 0,05547054 | 12,16033103 | 5,06E-34 | 1,87E-32 |
| ENSG00000101608.12 | MYL12A | protein_coding | 11672,67628 | 0,411238797 | 0,0440465 | 9,336469378 | 9,96E-21 | 2,11E-19 |
| ENSG00000101624.10 | CEP76 | protein_coding | 2587,489854 | -0,206011867 | 0,063321757 | -3,253413649 | 0,001140274 | 0,00436207 |
| ENSG00000101654.17 | RNMT | protein_coding | 8991,202577 | -0,254288786 | 0,049254262 | -5,162777302 | 2,43E-07 | 1,72E-06 |
| ENSG00000101670.12 | LIPG | protein_coding | 4446,20405 | 0,362730506 | 0,062181821 | 5,833385072 | 5,43E-09 | 4,68E-08 |
| ENSG00000101680.15 | LAMA1 | protein_coding | 973,2808846 | 0,353274905 | 0,120296953 | 2,936690389 | 0,003317351 | 0,011346339 |
| ENSG00000101745.17 | ANKRD12 | protein_coding | 9405,864941 | 0,383339058 | 0,048582784 | 7,890429963 | 3,01E-15 | 4,61E-14 |
| ENSG00000101746.15 | NOL4 | protein_coding | 7633,166616 | 0,135504943 | 0,052139054 | 2,598914477 | 0,009351907 | 0,028281885 |
| ENSG00000101752.11 | MIB1 | protein_coding | 20268,4887 | -0,403485515 | 0,043557824 | -9,263215644 | 1,98E-20 | 4,14E-19 |
| ENSG00000101773.19 | RBBP8 | protein_coding | 12122,34541 | -0,186348116 | 0,051134625 | -3,644264864 | 0,000268157 | 0,001175453 |
| ENSG00000101782.15 | RIOK3 | protein_coding | 13409,67633 | -0,273196589 | 0,050885298 | -5,368870745 | 7,92E-08 | 5,91E-07 |
| ENSG00000101843.19 | PSMD10 | protein_coding | 5665,816263 | -0,250807775 | 0,056610879 | -4,430381195 | 9,41E-06 | 5,30E-05 |
| ENSG00000101846.7 | STS | protein_coding | 4020,632326 | -0,287095991 | 0,058667961 | -4,893573667 | 9,90E-07 | 6,43E-06 |
| ENSG00000101850.13 | GPR143 | protein_coding | 248,562462 | 0,494671709 | 0,162301278 | 3,047860833 | 0,002304766 | 0,008191692 |
| ENSG00000101868.10 | POLA1 | protein_coding | 18145,93099 | -0,430926361 | 0,04586486 | -9,395566836 | 5,69E-21 | 1,22E-19 |
| ENSG00000101888.11 | NXT2 | protein_coding | 2123,225693 | 0,216967781 | 0,0683289 | 3,175344244 | 0,001496588 | 0,0055893 |
| ENSG00000101898.6 | MCTS2P | protein_coding | 592,6349697 | -0,303375009 | 0,113802763 | -2,66579651 | 0,007680617 | 0,023813086 |
| ENSG00000101901.11 | ALG13 | protein_coding | 3898,270132 | -0,262778127 | 0,053686089 | -4,894715411 | 9,84E-07 | 6,40E-06 |
| ENSG00000101928.13 | MOSPD1 | protein_coding | 1650,042318 | 0,18395178 | 0,06974724 | 2,63740588 | 0,008354281 | 0,025622905 |
| ENSG00000101935.10 | AMMECR1 | protein_coding | 2527,059288 | -0,184807298 | 0,060002209 | -3,080008256 | 0,002069949 | 0,007447646 |
| ENSG00000101938.14 | CHRDL1 | protein_coding | 52,1069577 | -1,679932895 | 0,398777962 | -4,21270244 | 2,52E-05 | 0,000132705 |
| ENSG00000101945.16 | SUV39H1 | protein_coding | 3407,65919 | -0,162345955 | 0,057841079 | -2,806758775 | 0,00500427 | 0,016334268 |
| ENSG00000101955.15 | SRPX | protein_coding | 227,5294279 | -0,542881757 | 0,193200624 | -2,809937908 | 0,004955106 | 0,016192578 |
| ENSG00000101966.12 | XIAP | protein_coding | 10724,72726 | 0,296400859 | 0,054009627 | 5,487926464 | 4,07E-08 | 3,15E-07 |
| ENSG00000101972.18 | STAG2 | protein_coding | 18186,44141 | -0,189542004 | 0,042260562 | -4,485080075 | 7,29E-06 | 4,16E-05 |
| ENSG00000101974.14 | ATP11C | protein_coding | 111,6944297 | 2,123596446 | 0,228598222 | 9,289645499 | 1,55E-20 | 3,24E-19 |
| ENSG00000102007.11 | PLP2 | protein_coding | 20182,49301 | -0,150560033 | 0,041150853 | -3,658734212 | 0,000253464 | 0,001116263 |

| | | | | | | | | |
|---|---|---|---|---|---|---|---|---|
| ENSG00000102024.18 | PLS3 | protein_coding | 9875,915719 | -0,189312447 | 0,05073823 | -3,731159875 | 0,0001906 | 0,000858216 |
| ENSG00000102043.16 | MTMR8 | protein_coding | 140,2206099 | 0,612529825 | 0,250650631 | 2,443759358 | 0,014535118 | 0,041288613 |
| ENSG00000102048.15 | ASB9 | protein_coding | 4301,671304 | -0,450801691 | 0,060261816 | -7,480718673 | 7,39E-14 | 1,03E-12 |
| ENSG00000102054.17 | RBBP7 | protein_coding | 49753,30796 | -0,243243991 | 0,039117671 | -6,218263638 | 5,03E-10 | 4,88E-09 |
| ENSG00000102057.10 | KCND1 | protein_coding | 4062,845733 | -0,18338004 | 0,060285093 | -3,04188033 | 0,002351053 | 0,008335149 |
| ENSG00000102078.16 | SLC25A14 | protein_coding | 2587,029711 | -0,200706109 | 0,061611627 | -3,257601196 | 0,001123582 | 0,004305954 |
| ENSG00000102081.14 | FMR1 | protein_coding | 9282,305596 | 0,920398535 | 0,058069115 | 15,85005267 | 1,40E-56 | 9,16E-55 |
| ENSG00000102096.9 | PIM2 | protein_coding | 6911,864562 | 0,373488513 | 0,059065868 | 6,323254468 | 2,56E-10 | 2,56E-09 |
| ENSG00000102100.16 | SLC35A2 | protein_coding | 7212,710484 | 0,169044887 | 0,050457671 | 3,35023169 | 0,00080744 | 0,003199202 |
| ENSG00000102103.16 | PQBP1 | protein_coding | 10957,68837 | 0,13038442 | 0,045992106 | 2,834930419 | 0,004583567 | 0,015099005 |
| ENSG00000102119.10 | EMD | protein_coding | 2387,679448 | -0,158334197 | 0,058970357 | -2,684979443 | 0,007253427 | 0,022646526 |
| ENSG00000102125.16 | TAZ | protein_coding | 1530,978265 | -0,20653576 | 0,083527622 | -2,472664195 | 0,013411009 | 0,038528195 |
| ENSG00000102144.15 | PGK1 | protein_coding | 40289,50846 | -0,117758764 | 0,049016213 | -2,402445182 | 0,016285875 | 0,045532079 |
| ENSG00000102158.19 | MAGT1 | protein_coding | 13196,5099 | 0,33754812 | 0,052486888 | 6,431094176 | 1,27E-10 | 1,31E-09 |
| ENSG00000102181.21 | CD99L2 | protein_coding | 5412,117278 | -0,12958537 | 0,048408012 | -2,67694056 | 0,007429781 | 0,02311499 |
| ENSG00000102218.6 | RP2 | protein_coding | 6450,16506 | -0,209981717 | 0,046737608 | -4,492778406 | 7,03E-06 | 4,03E-05 |
| ENSG00000102225.16 | CDK16 | protein_coding | 12997,76779 | -0,170813592 | 0,046578644 | -3,667208327 | 0,000245213 | 0,001082043 |
| ENSG00000102226.9 | USP11 | protein_coding | 24952,1102 | 0,152811911 | 0,040121854 | 3,808695184 | 0,000139702 | 0,000646894 |
| ENSG00000102230.13 | PCYT1B | protein_coding | 1619,669928 | -0,262421172 | 0,077688084 | -3,377881908 | 0,000730464 | 0,002921988 |
| ENSG00000102241.12 | HTATSF1 | protein_coding | 15672,58132 | -0,147487397 | 0,041017973 | -3,595677351 | 0,000323549 | 0,001396769 |
| ENSG00000102265.12 | TIMP1 | protein_coding | 1433,667553 | 0,295939169 | 0,071752443 | 4,124447264 | 3,72E-05 | 0,000190026 |
| ENSG00000102309.14 | PIN4 | protein_coding | 3200,784481 | -0,236385371 | 0,069896595 | -3,381929691 | 0,000719786 | 0,002883883 |
| ENSG00000102316.17 | MAGED2 | protein_coding | 26593,74688 | 0,175411357 | 0,044390799 | 3,951525158 | 7,77E-05 | 0,00037679 |
| ENSG00000102317.18 | RBM3 | protein_coding | 17380,1629 | -0,453764311 | 0,0472624 | -9,600957846 | 7,92E-22 | 1,77E-20 |
| ENSG00000102383.14 | ZDHHC15 | protein_coding | 1544,126192 | -0,299275514 | 0,069950933 | -4,278363445 | 1,88E-05 | 0,000100925 |
| ENSG00000102384.13 | CENPI | protein_coding | 2786,007625 | -0,821390351 | 0,072125524 | -11,38834494 | 4,78E-30 | 1,51E-28 |
| ENSG00000102390.10 | PBDC1 | protein_coding | 2102,672222 | 0,321777227 | 0,065749562 | 4,893982832 | 9,88E-07 | 6,42E-06 |
| ENSG00000102393.10 | GLA | protein_coding | 2533,765636 | 0,170683929 | 0,061048742 | 2,795863149 | 0,005176131 | 0,016836639 |
| ENSG00000102401.20 | ARMCX3 | protein_coding | 17429,89036 | 0,205060114 | 0,042626585 | 4,810615564 | 1,50E-06 | 9,52E-06 |
| ENSG00000102452.17 | NALCN | protein_coding | 278,1454836 | -1,231766468 | 0,201713966 | -6,106500662 | 1,02E-09 | 9,51E-09 |
| ENSG00000102466.15 | FGF14 | protein_coding | 1861,147695 | -0,545641577 | 0,08112219 | -6,726169227 | 1,74E-11 | 1,95E-10 |
| ENSG00000102524.11 | TNFSF13B | protein_coding | 314,9233703 | 1,033850728 | 0,168502135 | 6,135534886 | 8,49E-10 | 8,00E-09 |
| ENSG00000102547.19 | CAB39L | protein_coding | 1863,047322 | -0,486532159 | 0,076295112 | -6,376976798 | 1,81E-10 | 1,83E-09 |
| ENSG00000102554.14 | KLF5 | protein_coding | 2275,496281 | 0,532548121 | 0,077659682 | 6,857459427 | 7,01E-12 | 8,18E-11 |
| ENSG00000102572.14 | STK24 | protein_coding | 14680,81021 | 0,333333071 | 0,041763368 | 7,981470033 | 1,45E-15 | 2,25E-14 |
| ENSG00000102575.12 | ACP5 | protein_coding | 536,1626741 | 0,624246128 | 0,123179668 | 5,067769194 | 4,03E-07 | 2,76E-06 |

| | | | | | | | | |
|---|---|---|---|---|---|---|---|---|
| ENSG00000102580.15 | DNAJC3 | protein_coding | 9264,759816 | 0,231511414 | 0,045358741 | 5,104008827 | 3,33E-07 | 2,31E-06 |
| ENSG00000102595.20 | UGGT2 | protein_coding | 8267,79541 | 0,22008841 | 0,05312279 | 4,143013043 | 3,43E-05 | 0,000176152 |
| ENSG00000102678.6 | FGF9 | protein_coding | 144,4717792 | -1,173670247 | 0,226276362 | -5,186888448 | 2,14E-07 | 1,52E-06 |
| ENSG00000102699.6 | PARP4 | protein_coding | 16110,94666 | 0,67124909 | 0,043158187 | 15,55322738 | 1,51E-54 | 9,49E-53 |
| ENSG00000102710.20 | SUPT20H | protein_coding | 9521,944342 | 0,517647634 | 0,057955455 | 8,931819009 | 4,19E-19 | 8,11E-18 |
| ENSG00000102738.7 | MRPS31 | protein_coding | 2471,36526 | 0,196865261 | 0,059931614 | 3,284831591 | 0,001020433 | 0,003946553 |
| ENSG00000102781.14 | KATNAL1 | protein_coding | 7802,720742 | -0,441844354 | 0,047897757 | -9,22474002 | 2,84E-20 | 5,85E-19 |
| ENSG00000102786.15 | INTS6 | protein_coding | 8251,309892 | 0,292699153 | 0,047614243 | 6,147302406 | 7,88E-10 | 7,47E-09 |
| ENSG00000102804.15 | TSC22D1 | protein_coding | 18985,53189 | 0,356140899 | 0,043385325 | 8,208787197 | 2,23E-16 | 3,67E-15 |
| ENSG00000102854.16 | MSLN | protein_coding | 493,3755467 | -0,306880234 | 0,113376895 | -2,706726384 | 0,006795024 | 0,021400474 |
| ENSG00000102858.13 | MGRN1 | protein_coding | 6078,887095 | 0,205870137 | 0,05242301 | 3,927094912 | 8,60E-05 | 0,00041494 |
| ENSG00000102870.6 | ZNF629 | protein_coding | 4876,656346 | 0,245767241 | 0,054716004 | 4,491688448 | 7,07E-06 | 4,05E-05 |
| ENSG00000102871.16 | TRADD | protein_coding | 383,6331531 | 0,723767016 | 0,128943811 | 5,613041914 | 1,99E-08 | 1,60E-07 |
| ENSG00000102879.15 | CORO1A | protein_coding | 1447,084007 | -0,319166633 | 0,092022175 | -3,468366544 | 0,000523633 | 0,002158294 |
| ENSG00000102882.12 | MAPK3 | protein_coding | 2899,420664 | -0,245330606 | 0,055815034 | -4,395421623 | 1,11E-05 | 6,15E-05 |
| ENSG00000102890.15 | ELMO3 | protein_coding | 1414,318689 | 0,207182459 | 0,081839419 | 2,531572934 | 0,01135522 | 0,033345681 |
| ENSG00000102893.16 | PHKB | protein_coding | 11001,03332 | -0,522043266 | 0,055663386 | -9,378575465 | 6,69E-21 | 1,43E-19 |
| ENSG00000102897.10 | LYRM1 | protein_coding | 7359,20015 | -0,172014495 | 0,049719025 | -3,459731831 | 0,000540714 | 0,002224659 |
| ENSG00000102898.12 | NUTF2 | protein_coding | 6564,840599 | -0,224068967 | 0,053248423 | -4,207992568 | 2,58E-05 | 0,000135279 |
| ENSG00000102901.13 | CENPT | protein_coding | 4056,814986 | -0,319370585 | 0,061185466 | -5,219713187 | 1,79E-07 | 1,28E-06 |
| ENSG00000102908.22 | NFAT5 | protein_coding | 6314,741895 | 0,511163827 | 0,058175034 | 8,786652816 | 1,54E-18 | 2,90E-17 |
| ENSG00000102921.8 | AC026470.1 | protein_coding | 8337,479004 | 0,770367088 | 0,056962576 | 13,52409148 | 1,13E-41 | 5,31E-40 |
| ENSG00000102924.12 | CBLN1 | protein_coding | 5507,536401 | -0,379802449 | 0,056715488 | -6,696626707 | 2,13E-11 | 2,37E-10 |
| ENSG00000102974.16 | CTCF | protein_coding | 10941,10465 | -0,116071107 | 0,042035827 | -2,761242393 | 0,005758192 | 0,018489465 |
| ENSG00000102977.16 | ACD | protein_coding | 2245,779326 | -0,203824464 | 0,072873824 | -2,796950306 | 0,005158747 | 0,016787369 |
| ENSG00000102984.15 | ZNF821 | protein_coding | 1596,483074 | 0,18960096 | 0,074756136 | 2,536259496 | 0,011204367 | 0,032975825 |
| ENSG00000102996.5 | MMP15 | protein_coding | 6662,960225 | 0,14436724 | 0,047803525 | 3,020012451 | 0,002527643 | 0,008898141 |
| ENSG00000103024.7 | NME3 | protein_coding | 1865,557199 | -0,20830665 | 0,083045972 | -2,50832936 | 0,012130354 | 0,03531783 |
| ENSG00000103037.11 | SETD6 | protein_coding | 2916,342977 | -0,185734619 | 0,063008205 | -2,947784648 | 0,003200599 | 0,010980383 |
| ENSG00000103042.9 | SLC38A7 | protein_coding | 2723,257238 | 0,435313687 | 0,067174212 | 6,480369089 | 9,15E-11 | 9,57E-10 |
| ENSG00000103056.12 | SMPD3 | protein_coding | 1061,428837 | 0,566976945 | 0,114479764 | 4,952638994 | 7,32E-07 | 4,85E-06 |
| ENSG00000103061.13 | SLC7A6OS | protein_coding | 2871,038827 | 0,329568898 | 0,068741986 | 4,794288276 | 1,63E-06 | 1,03E-05 |
| ENSG00000103126.15 | AXIN1 | protein_coding | 2923,90741 | 0,224614089 | 0,055074308 | 4,078382415 | 4,54E-05 | 0,000228927 |
| ENSG00000103145.10 | HCFC1R1 | protein_coding | 1295,843368 | -0,36825319 | 0,080394488 | -4,580577572 | 4,64E-06 | 2,73E-05 |
| ENSG00000103154.9 | NECAB2 | protein_coding | 525,6887784 | -0,669729426 | 0,11417565 | -5,86578158 | 4,47E-09 | 3,88E-08 |
| ENSG00000103160.12 | HSDL1 | protein_coding | 7128,527747 | -0,155870283 | 0,045471236 | -3,427887555 | 0,000608297 | 0,002475084 |

| | | | | | | | | |
|---|---|---|---|---|---|---|---|---|
| ENSG00000103174.13 | NAGPA | protein_coding | 2190,158869 | -0,155974712 | 0,06429082 | -2,426080622 | 0,015262879 | 0,043062072 |
| ENSG00000103187.8 | COTL1 | protein_coding | 2946,093164 | -0,206449646 | 0,055358327 | -3,729333205 | 0,000191987 | 0,000864115 |
| ENSG00000103194.15 | USP10 | protein_coding | 10671,52344 | 0,125328471 | 0,045898479 | 2,730558289 | 0,006322715 | 0,020075891 |
| ENSG00000103202.13 | NME4 | protein_coding | 2797,488346 | -0,447550598 | 0,070996753 | -6,303817818 | 2,90E-10 | 2,89E-09 |
| ENSG00000103222.19 | ABCC1 | protein_coding | 5833,030694 | 0,265296308 | 0,05323316 | 4,983666324 | 6,24E-07 | 4,18E-06 |
| ENSG00000103226.18 | NOMO3 | protein_coding | 13698,08694 | 0,404455846 | 0,049375294 | 8,191462101 | 2,58E-16 | 4,22E-15 |
| ENSG00000103257.9 | SLC7A5 | protein_coding | 799,1516629 | 0,874251047 | 0,117744466 | 7,424986323 | 1,13E-13 | 1,54E-12 |
| ENSG00000103260.9 | METRN | protein_coding | 654,4522706 | -0,452970234 | 0,104771381 | -4,323415725 | 1,54E-05 | 8,34E-05 |
| ENSG00000103264.18 | FBXO31 | protein_coding | 2585,445588 | 0,253923839 | 0,064854163 | 3,915305134 | 9,03E-05 | 0,000433149 |
| ENSG00000103266.11 | STUB1 | protein_coding | 4500,422333 | -0,146557931 | 0,059144607 | -2,477959316 | 0,013213621 | 0,038063143 |
| ENSG00000103275.20 | UBE2I | protein_coding | 13796,63618 | -0,198921936 | 0,050948727 | -3,904355348 | 9,45E-05 | 0,000451599 |
| ENSG00000103316.11 | CRYM | protein_coding | 525,1066387 | 0,366289574 | 0,104353456 | 3,510085719 | 0,000447962 | 0,001880489 |
| ENSG00000103319.12 | EEF2K | protein_coding | 3662,028951 | -0,232020483 | 0,055092 | -4,211509539 | 2,54E-05 | 0,000133346 |
| ENSG00000103351.13 | CLUAP1 | protein_coding | 3891,699446 | 0,213152951 | 0,081522766 | 2,614643273 | 0,008932069 | 0,027176526 |
| ENSG00000103363.14 | ELOB | protein_coding | 13649,4838 | -0,191353535 | 0,043919555 | -4,356909665 | 1,32E-05 | 7,26E-05 |
| ENSG00000103423.14 | DNAJA3 | protein_coding | 10498,63575 | 0,142544984 | 0,051438865 | 2,771153386 | 0,005585811 | 0,018005301 |
| ENSG00000103426.12 | CORO7-PAM16 | protein_coding | 14,28457061 | -3,580193787 | 0,976619494 | -3,66590449 | 0,000246466 | 0,001087145 |
| ENSG00000103449.11 | SALL1 | protein_coding | 1476,162204 | 0,211377126 | 0,082246917 | 2,570031004 | 0,010168941 | 0,030353169 |
| ENSG00000103472.10 | RRN3P2 | lncRNA | 14,40918169 | 1,737231962 | 0,650658061 | 2,669961485 | 0,007585995 | 0,023535928 |
| ENSG00000103479.16 | RBL2 | protein_coding | 14037,91601 | -0,181685073 | 0,047276377 | -3,843041366 | 0,000121519 | 0,000570442 |
| ENSG00000103490.14 | PYCARD | protein_coding | 22,50154564 | 1,278450454 | 0,505961418 | 2,526774591 | 0,011511536 | 0,03370797 |
| ENSG00000103494.14 | RPGRIP1L | protein_coding | 2508,280366 | -0,501765858 | 0,071338402 | -7,033600983 | 2,01E-12 | 2,48E-11 |
| ENSG00000103495.14 | MAZ | protein_coding | 22861,94906 | -0,305604106 | 0,045186422 | -6,763184432 | 1,35E-11 | 1,53E-10 |
| ENSG00000103496.15 | STX4 | protein_coding | 1822,242635 | 0,384348826 | 0,069575715 | 5,524180803 | 3,31E-08 | 2,60E-07 |
| ENSG00000103510.20 | KAT8 | protein_coding | 2381,607174 | -0,363981987 | 0,070419173 | -5,168791002 | 2,36E-07 | 1,66E-06 |
| ENSG00000103512.15 | NOMO1 | protein_coding | 50979,11087 | 0,42071492 | 0,038784405 | 10,84752811 | 2,05E-27 | 5,83E-26 |
| ENSG00000103528.17 | SYT17 | protein_coding | 3111,348119 | -0,178008491 | 0,06091068 | -2,922451201 | 0,00347288 | 0,011829762 |
| ENSG00000103540.16 | CCP110 | protein_coding | 13119,06785 | -0,217942734 | 0,043508276 | -5,00922477 | 5,46E-07 | 3,70E-06 |
| ENSG00000103544.14 | VPS35L | protein_coding | 7810,957911 | -0,125836845 | 0,050102514 | -2,511587437 | 0,012018951 | 0,035034261 |
| ENSG00000103642.12 | LACTB | protein_coding | 1848,0739 | 0,404037991 | 0,069749268 | 5,792720167 | 6,93E-09 | 5,88E-08 |
| ENSG00000103647.12 | CORO2B | protein_coding | 123,4174474 | -0,885289224 | 0,240326197 | -3,683698392 | 0,000229874 | 0,001020761 |
| ENSG00000103723.14 | AP3B2 | protein_coding | 6140,513551 | 0,137291333 | 0,054078702 | 2,538732044 | 0,0111255 | 0,032782285 |
| ENSG00000103740.10 | ACSBG1 | protein_coding | 35,33005417 | 1,721712464 | 0,589639527 | 2,919940721 | 0,00350098 | 0,011907457 |
| ENSG00000103742.12 | IGDCC4 | protein_coding | 1419,868483 | -1,907056599 | 0,109355306 | -17,43908604 | 4,17E-68 | 3,29E-66 |
| ENSG00000103966.11 | EHD4 | protein_coding | 5838,000885 | 2,081845337 | 0,058371742 | 35,66529386 | 1,37E-278 | 3,53E-276 |
| ENSG00000103978.15 | TMEM87A | protein_coding | 10248,07973 | 0,186387708 | 0,045843783 | 4,065713949 | 4,79E-05 | 0,000240432 |

| | | | | | | | | |
|---|---|---|---|---|---|---|---|---|
| ENSG00000103994.17 | ZNF106 | protein_coding | 10617,54556 | -0,152647073 | 0,05956296 | -2,562785203 | 0,010383627 | 0,030887934 |
| ENSG00000104044.16 | OCA2 | protein_coding | 1703,429995 | -0,175962892 | 0,06883334 | -2,556361394 | 0,01057732 | 0,031394354 |
| ENSG00000104067.16 | TJP1 | protein_coding | 10348,61401 | 0,43374144 | 0,05086665 | 8,527029848 | 1,50E-17 | 2,66E-16 |
| ENSG00000104081.14 | BMF | protein_coding | 3246,568135 | 0,234917147 | 0,06930246 | 3,389737503 | 0,000699596 | 0,002810495 |
| ENSG00000104112.9 | SCG3 | protein_coding | 95903,40586 | 0,368022873 | 0,040644726 | 9,05462807 | 1,37E-19 | 2,71E-18 |
| ENSG00000104129.10 | DNAJC17 | protein_coding | 755,8528299 | -0,292477328 | 0,111720704 | -2,617933095 | 0,008846415 | 0,026955997 |
| ENSG00000104133.15 | SPG11 | protein_coding | 11222,40613 | 0,623792749 | 0,04984511 | 12,5146228 | 6,21E-36 | 2,47E-34 |
| ENSG00000104142.11 | VPS18 | protein_coding | 3615,248193 | 0,407678219 | 0,062758192 | 6,496016011 | 8,25E-11 | 8,65E-10 |
| ENSG00000104147.9 | OIP5 | protein_coding | 1818,62454 | -0,600465529 | 0,072079268 | -8,330627425 | 8,04E-17 | 1,36E-15 |
| ENSG00000104164.10 | BLOC1S6 | protein_coding | 12785,74323 | -0,270193244 | 0,044018108 | -6,138229393 | 8,34E-10 | 7,87E-09 |
| ENSG00000104177.18 | MYEF2 | protein_coding | 16585,29293 | -0,238672385 | 0,04686753 | -5,092489103 | 3,53E-07 | 2,44E-06 |
| ENSG00000104205.15 | SGK3 | protein_coding | 6016,808792 | 0,24038745 | 0,051879824 | 4,633544083 | 3,59E-06 | 2,15E-05 |
| ENSG00000104228.13 | TRIM35 | protein_coding | 2703,039743 | 0,30510004 | 0,058589973 | 5,207376323 | 1,92E-07 | 1,37E-06 |
| ENSG00000104231.11 | ZFAND1 | protein_coding | 3422,898897 | 0,572451739 | 0,056130905 | 10,1985126 | 2,01E-24 | 5,09E-23 |
| ENSG00000104290.11 | FZD3 | protein_coding | 7689,327935 | 0,195811291 | 0,060059087 | 3,260310796 | 0,001112902 | 0,004268969 |
| ENSG00000104312.8 | RIPK2 | protein_coding | 587,5199786 | 1,350260538 | 0,117295624 | 11,51160202 | 1,15E-30 | 3,77E-29 |
| ENSG00000104320.13 | NBN | protein_coding | 9036,560099 | 0,702418918 | 0,04666473 | 15,05245852 | 3,33E-51 | 1,97E-49 |
| ENSG00000104321.11 | TRPA1 | protein_coding | 60,17998032 | -1,146816338 | 0,362048704 | -3,167574763 | 0,001537162 | 0,00571995 |
| ENSG00000104324.16 | CPQ | protein_coding | 6121,002906 | 0,484699746 | 0,050809376 | 9,539572861 | 1,43E-21 | 3,16E-20 |
| ENSG00000104325.7 | DECR1 | protein_coding | 4384,243331 | 0,411651198 | 0,055017166 | 7,482231937 | 7,31E-14 | 1,01E-12 |
| ENSG00000104331.9 | IMPAD1 | protein_coding | 16164,70277 | -0,207768577 | 0,047485616 | -4,375400236 | 1,21E-05 | 6,71E-05 |
| ENSG00000104332.12 | SFRP1 | protein_coding | 11771,72547 | 0,129784976 | 0,049188411 | 2,638527525 | 0,008326694 | 0,025555707 |
| ENSG00000104365.15 | IKBKB | protein_coding | 3014,178754 | 0,316837683 | 0,064277774 | 4,929195039 | 8,26E-07 | 5,42E-06 |
| ENSG00000104369.5 | JPH1 | protein_coding | 1096,679129 | -0,295681689 | 0,08053368 | -3,671528349 | 0,000241104 | 0,001066002 |
| ENSG00000104381.12 | GDAP1 | protein_coding | 8836,795457 | -0,186211459 | 0,048504151 | -3,839082922 | 0,000123495 | 0,00057851 |
| ENSG00000104388.15 | RAB2A | protein_coding | 17981,05948 | -0,140215063 | 0,057253629 | -2,449016153 | 0,014324702 | 0,040835198 |
| ENSG00000104408.9 | EIF3E | protein_coding | 21317,52572 | -0,495328545 | 0,042984828 | -11,52333446 | 1,01E-30 | 3,32E-29 |
| ENSG00000104419.14 | NDRG1 | protein_coding | 13006,92065 | -0,173905642 | 0,067988743 | -2,557859332 | 0,010531869 | 0,031278774 |
| ENSG00000104432.14 | IL7 | protein_coding | 949,2241764 | 1,551092477 | 0,097582253 | 15,89523122 | 6,84E-57 | 4,54E-55 |
| ENSG00000104435.13 | STMN2 | protein_coding | 29304,30349 | -0,103355441 | 0,040840034 | -2,530738358 | 0,011382272 | 0,03340735 |
| ENSG00000104442.10 | ARMC1 | protein_coding | 8330,605623 | -0,279498664 | 0,052469546 | -5,326874085 | 9,99E-08 | 7,36E-07 |
| ENSG00000104450.12 | SPAG1 | protein_coding | 1845,943626 | 0,45239492 | 0,06679253 | 6,773136492 | 1,26E-11 | 1,43E-10 |
| ENSG00000104472.10 | CHRAC1 | protein_coding | 3932,850115 | -0,247804503 | 0,057395013 | -4,31752671 | 1,58E-05 | 8,55E-05 |
| ENSG00000104490.18 | NCALD | protein_coding | 4177,227492 | 0,654673492 | 0,068437854 | 9,565955827 | 1,11E-21 | 2,47E-20 |
| ENSG00000104517.13 | UBR5 | protein_coding | 18546,27019 | 0,315440431 | 0,042494261 | 7,423130215 | 1,14E-13 | 1,56E-12 |
| ENSG00000104518.11 | GSDMD | protein_coding | 2273,522954 | 3,554709895 | 0,079399941 | 44,76967952 | 0 | 0 |

| | | | | | | | | |
|---|---|---|---|---|---|---|---|---|
| ENSG00000104522.16 | TSTA3 | protein_coding | 1952,162319 | -0,318416733 | 0,069509314 | -4,580921832 | 4,63E-06 | 2,73E-05 |
| ENSG00000104537.17 | ANXA13 | protein_coding | 1074,791571 | -0,448427128 | 0,085171545 | -5,264987583 | 1,40E-07 | 1,02E-06 |
| ENSG00000104549.12 | SQLE | protein_coding | 25575,92702 | 0,371216971 | 0,042288916 | 8,778114994 | 1,66E-18 | 3,12E-17 |
| ENSG00000104611.12 | SH2D4A | protein_coding | 225,3520654 | 0,687019511 | 0,163789585 | 4,194525025 | 2,73E-05 | 0,00014274 |
| ENSG00000104635.14 | SLC39A14 | protein_coding | 28043,64804 | 0,259052327 | 0,040847638 | 6,341916922 | 2,27E-10 | 2,28E-09 |
| ENSG00000104689.9 | TNFRSF10A | protein_coding | 36,43100285 | 3,834172396 | 0,587003591 | 6,531769917 | 6,50E-11 | 6,88E-10 |
| ENSG00000104691.15 | UBXN8 | protein_coding | 591,4481358 | -0,49793996 | 0,106785721 | -4,662982603 | 3,12E-06 | 1,88E-05 |
| ENSG00000104714.14 | ERICH1 | protein_coding | 1147,991643 | 0,210741555 | 0,078682079 | 2,678393318 | 0,007397629 | 0,023034063 |
| ENSG00000104723.20 | TUSC3 | protein_coding | 44279,68768 | 0,13148371 | 0,038291958 | 3,433715991 | 0,000595368 | 0,002426862 |
| ENSG00000104728.16 | ARHGEF10 | protein_coding | 586,737934 | -0,4630628 | 0,119215348 | -3,884254902 | 0,000102644 | 0,000487636 |
| ENSG00000104731.14 | KLHDC4 | protein_coding | 2172,992644 | 0,303274466 | 0,072258263 | 4,197090432 | 2,70E-05 | 0,000141166 |
| ENSG00000104738.18 | MCM4 | protein_coding | 29783,02435 | -0,208301627 | 0,049422235 | -4,2147351 | 2,50E-05 | 0,000131638 |
| ENSG00000104756.16 | KCTD9 | protein_coding | 3753,877187 | -0,299879835 | 0,0606324 | -4,945867761 | 7,58E-07 | 5,01E-06 |
| ENSG00000104760.17 | FGL1 | protein_coding | 127,4230931 | 1,609837117 | 0,240114781 | 6,704448237 | 2,02E-11 | 2,25E-10 |
| ENSG00000104765.16 | BNIP3L | protein_coding | 14829,4844 | -0,368755741 | 0,070399177 | -5,238068898 | 1,62E-07 | 1,17E-06 |
| ENSG00000104774.13 | MAN2B1 | protein_coding | 8314,855004 | -0,233961851 | 0,073315032 | -3,191185272 | 0,001416904 | 0,005310205 |
| ENSG00000104805.16 | NUCB1 | protein_coding | 21905,56094 | 0,548054324 | 0,051521945 | 10,63729876 | 2,00E-26 | 5,51E-25 |
| ENSG00000104825.17 | NFKBIB | protein_coding | 1753,522934 | 0,336031016 | 0,071381433 | 4,707540905 | 2,51E-06 | 1,54E-05 |
| ENSG00000104852.15 | SNRNP70 | protein_coding | 18501,93726 | -0,21971772 | 0,04842415 | -4,537358357 | 5,70E-06 | 3,31E-05 |
| ENSG00000104856.14 | RELB | protein_coding | 376,97675 | 1,754985711 | 0,14884516 | 11,79068037 | 4,36E-32 | 1,54E-30 |
| ENSG00000104870.12 | FCGRT | protein_coding | 17471,04325 | 0,184197733 | 0,054238679 | 3,396058605 | 0,000683637 | 0,002750311 |
| ENSG00000104872.11 | PIH1D1 | protein_coding | 5190,313722 | -0,227654285 | 0,052270573 | -4,355304924 | 1,33E-05 | 7,30E-05 |
| ENSG00000104880.18 | ARHGEF18 | protein_coding | 3691,931932 | 0,450132162 | 0,066736635 | 6,744903444 | 1,53E-11 | 1,72E-10 |
| ENSG00000104883.8 | PEX11G | protein_coding | 99,88610411 | -0,579959963 | 0,236469325 | -2,452580113 | 0,014183579 | 0,040489307 |
| ENSG00000104889.6 | RNASEH2A | protein_coding | 4026,713564 | -0,400451391 | 0,058066272 | -6,896454263 | 5,33E-12 | 6,29E-11 |
| ENSG00000104904.12 | OAZ1 | protein_coding | 26056,32925 | -0,175645209 | 0,068949482 | -2,547447834 | 0,010851408 | 0,032079634 |
| ENSG00000104915.15 | STX10 | protein_coding | 1620,228309 | -0,256291134 | 0,075414822 | -3,398418588 | 0,000677766 | 0,002730596 |
| ENSG00000104936.17 | DMPK | protein_coding | 2560,355074 | -0,177381194 | 0,073869048 | -2,40129255 | 0,016337269 | 0,045647418 |
| ENSG00000104946.13 | TBC1D17 | protein_coding | 3257,997984 | 0,229938887 | 0,057510863 | 3,998181851 | 6,38E-05 | 0,000314667 |
| ENSG00000104960.15 | PTOV1 | protein_coding | 17315,38865 | -0,2021386 | 0,043222761 | -4,676670242 | 2,92E-06 | 1,77E-05 |
| ENSG00000104967.7 | NOVA2 | protein_coding | 931,645145 | 0,379107356 | 0,102838209 | 3,686444546 | 0,000227409 | 0,001010412 |
| ENSG00000104973.18 | MED25 | protein_coding | 25777,94079 | 0,107883508 | 0,04322623 | 2,495788011 | 0,012567768 | 0,036426461 |
| ENSG00000104979.9 | C19orf53 | protein_coding | 3427,702797 | -0,264262729 | 0,058912072 | -4,485714429 | 7,27E-06 | 4,15E-05 |
| ENSG00000104980.8 | TIMM44 | protein_coding | 1978,66997 | 0,19520136 | 0,067756844 | 2,880909852 | 0,00396529 | 0,013299765 |
| ENSG00000105053.11 | VRK3 | protein_coding | 5608,870667 | 0,167117349 | 0,051941674 | 3,217403985 | 0,001293563 | 0,004887832 |
| ENSG00000105058.12 | FAM32A | protein_coding | 9284,678492 | 0,137788074 | 0,04852003 | 2,839818417 | 0,004513922 | 0,014895744 |

| | | | | | | | | |
|---|---|---|---|---|---|---|---|---|
| ENSG00000105063.19 | PPP6R1 | protein_coding | 19025,64491 | -0,258656137 | 0,046775851 | -5,529693943 | 3,21E-08 | 2,52E-07 |
| ENSG00000105137.13 | SYDE1 | protein_coding | 282,4125875 | -0,686773405 | 0,150639896 | -4,559040611 | 5,14E-06 | 3,01E-05 |
| ENSG00000105173.14 | CCNE1 | protein_coding | 1829,239938 | -0,79011825 | 0,064373665 | -12,27393603 | 1,25E-34 | 4,73E-33 |
| ENSG00000105176.18 | URI1 | protein_coding | 6425,812232 | -0,300165175 | 0,05304898 | -5,658264815 | 1,53E-08 | 1,25E-07 |
| ENSG00000105185.12 | PDCD5 | protein_coding | 10610,74206 | -0,540140486 | 0,048479749 | -11,1415693 | 7,87E-29 | 2,37E-27 |
| ENSG00000105193.9 | RPS16 | protein_coding | 23497,83401 | -0,674721609 | 0,044519635 | -15,15559627 | 6,96E-52 | 4,20E-50 |
| ENSG00000105202.9 | FBL | protein_coding | 12263,75722 | -0,305945772 | 0,054861634 | -5,576679956 | 2,45E-08 | 1,96E-07 |
| ENSG00000105245.9 | NUMBL | protein_coding | 1238,407648 | -0,687342485 | 0,104959024 | -6,548674536 | 5,80E-11 | 6,19E-10 |
| ENSG00000105248.16 | YJU2 | protein_coding | 1252,964786 | 0,186049839 | 0,074703624 | 2,490506206 | 0,012756127 | 0,036901126 |
| ENSG00000105254.12 | TBCB | protein_coding | 14880,30002 | -0,283849451 | 0,045275389 | -6,269398335 | 3,62E-10 | 3,57E-09 |
| ENSG00000105255.11 | FSD1 | protein_coding | 962,1751594 | -0,252441142 | 0,084852326 | -2,975064484 | 0,00292927 | 0,010143835 |
| ENSG00000105258.9 | POLR2I | protein_coding | 2763,429119 | -0,357347489 | 0,063895485 | -5,592687617 | 2,24E-08 | 1,79E-07 |
| ENSG00000105270.15 | CLIP3 | protein_coding | 4574,75338 | 0,570475818 | 0,055316896 | 10,31286748 | 6,16E-25 | 1,60E-23 |
| ENSG00000105287.12 | PRKD2 | protein_coding | 5818,775427 | 2,555895039 | 0,20307804 | 12,58577754 | 2,53E-36 | 1,02E-34 |
| ENSG00000105289.15 | TJP3 | protein_coding | 443,0574129 | 0,547291167 | 0,135380585 | 4,042611921 | 5,29E-05 | 0,000264017 |
| ENSG00000105290.12 | APLP1 | protein_coding | 29233,20802 | 0,113893746 | 0,045828867 | 2,485196639 | 0,012947989 | 0,037350543 |
| ENSG00000105321.14 | CCDC9 | protein_coding | 951,1095197 | 0,532522647 | 0,088534224 | 6,014879052 | 1,80E-09 | 1,64E-08 |
| ENSG00000105323.16 | HNRNPUL1 | protein_coding | 26269,19773 | -0,236360896 | 0,042633903 | -5,543965666 | 2,96E-08 | 2,34E-07 |
| ENSG00000105339.10 | DENND3 | protein_coding | 3230,21508 | 0,939609619 | 0,069909987 | 13,44027743 | 3,51E-41 | 1,63E-39 |
| ENSG00000105341.18 | DMAC2 | protein_coding | 4176,612328 | -0,17597294 | 0,063851464 | -2,755973455 | 0,005851775 | 0,018744499 |
| ENSG00000105372.7 | RPS19 | protein_coding | 30988,916 | -0,385815325 | 0,042853135 | -9,003199591 | 2,19E-19 | 4,29E-18 |
| ENSG00000105373.19 | NOP53 | protein_coding | 4886,59567 | -0,332055338 | 0,051543322 | -6,442257227 | 1,18E-10 | 1,22E-09 |
| ENSG00000105379.9 | ETFB | protein_coding | 1031,515799 | -0,25129894 | 0,081141329 | -3,097052298 | 0,001954554 | 0,00707885 |
| ENSG00000105393.16 | BABAM1 | protein_coding | 4729,476798 | -0,204789939 | 0,05182883 | -3,951274558 | 7,77E-05 | 0,000377104 |
| ENSG00000105402.8 | NAPA | protein_coding | 15043,5634 | 1,377938185 | 0,050942404 | 27,04894292 | 3,93E-161 | 6,81E-159 |
| ENSG00000105419.17 | MEIS3 | protein_coding | 590,0391093 | 0,51826456 | 0,119391708 | 4,340875665 | 1,42E-05 | 7,75E-05 |
| ENSG00000105426.16 | PTPRS | protein_coding | 1645,341404 | -0,364081019 | 0,082724449 | -4,401129571 | 1,08E-05 | 6,00E-05 |
| ENSG00000105427.10 | CNFN | protein_coding | 441,2727346 | -0,679592348 | 0,121069318 | -5,613250001 | 1,99E-08 | 1,60E-07 |
| ENSG00000105429.13 | MEGF8 | protein_coding | 6813,168378 | 0,161780194 | 0,057650449 | 2,806226086 | 0,005012551 | 0,016349607 |
| ENSG00000105443.15 | CYTH2 | protein_coding | 6500,112109 | -0,339588508 | 0,063793779 | -5,323222917 | 1,02E-07 | 7,51E-07 |
| ENSG00000105472.13 | CLEC11A | protein_coding | 127,1854336 | -1,052360368 | 0,223752678 | -4,703230274 | 2,56E-06 | 1,57E-05 |
| ENSG00000105483.18 | CARD8 | protein_coding | 5599,814961 | 0,27485023 | 0,049166419 | 5,59020229 | 2,27E-08 | 1,82E-07 |
| ENSG00000105486.14 | LIG1 | protein_coding | 6302,429024 | -0,140118059 | 0,057817706 | -2,423445474 | 0,015374061 | 0,043310981 |
| ENSG00000105499.14 | PLA2G4C | protein_coding | 353,7126148 | 1,423516069 | 0,147620732 | 9,643063321 | 5,26E-22 | 1,19E-20 |
| ENSG00000105514.8 | RAB3D | protein_coding | 13583,69939 | -0,396593761 | 0,041425942 | -9,573560424 | 1,03E-21 | 2,30E-20 |
| ENSG00000105552.15 | BCAT2 | protein_coding | 4436,808727 | -0,188875825 | 0,060644739 | -3,114463493 | 0,001842798 | 0,006721687 |

| | | | | | | | | |
|---|---|---|---|---|---|---|---|---|
| ENSG00000105556.11 | MIER2 | protein_coding | 1170,715339 | 0,220310327 | 0,092738914 | 2,375597453 | 0,017520565 | 0,048494444 |
| ENSG00000105559.12 | PLEKHA4 | protein_coding | 1728,953126 | 1,406288091 | 0,109410106 | 12,85336563 | 8,23E-38 | 3,44E-36 |
| ENSG00000105568.18 | PPP2R1A | protein_coding | 20413,5082 | -0,131662581 | 0,043487153 | -3,027620177 | 0,002464877 | 0,008692137 |
| ENSG00000105583.11 | WDR83OS | protein_coding | 5427,093867 | -0,2498029 | 0,071035478 | -3,516593521 | 0,000437123 | 0,001837728 |
| ENSG00000105612.9 | DNASE2 | protein_coding | 1516,0581 | 0,872569493 | 0,088163067 | 9,897222514 | 4,28E-23 | 1,01E-21 |
| ENSG00000105613.10 | MAST1 | protein_coding | 1402,52427 | -0,500523959 | 0,089314224 | -5,604078897 | 2,09E-08 | 1,68E-07 |
| ENSG00000105618.14 | PRPF31 | protein_coding | 5794,791209 | -0,240396038 | 0,049065908 | -4,899451534 | 9,61E-07 | 6,26E-06 |
| ENSG00000105640.13 | RPL18A | protein_coding | 18662,53132 | -0,48175657 | 0,046145623 | -10,43991908 | 1,63E-25 | 4,32E-24 |
| ENSG00000105643.10 | ARRDC2 | protein_coding | 321,9369813 | 0,641345232 | 0,153606824 | 4,1752392 | 2,98E-05 | 0,000154563 |
| ENSG00000105647.17 | PIK3R2 | protein_coding | 7413,981305 | -0,704726692 | 0,05388448 | -13,07847254 | 4,37E-39 | 1,92E-37 |
| ENSG00000105650.22 | AC005759.1 | protein_coding | 377,7527479 | -0,699451281 | 0,192871807 | -3,626508674 | 0,000287279 | 0,001252444 |
| ENSG00000105655.18 | ISYNA1 | protein_coding | 5214,583778 | -0,360950613 | 0,076494377 | -4,718655485 | 2,37E-06 | 1,46E-05 |
| ENSG00000105656.13 | ELL | protein_coding | 1078,339048 | 0,311095016 | 0,079851677 | 3,895910845 | 9,78E-05 | 0,000466072 |
| ENSG00000105662.16 | CRTC1 | protein_coding | 2888,300606 | -0,442900157 | 0,066534901 | -6,65665914 | 2,80E-11 | 3,07E-10 |
| ENSG00000105672.14 | ETV2 | protein_coding | 93,18206904 | -1,178576477 | 0,25144211 | -4,687267694 | 2,77E-06 | 1,69E-05 |
| ENSG00000105677.12 | TMEM147 | protein_coding | 3098,032348 | -0,168321369 | 0,060234408 | -2,794438845 | 0,005198987 | 0,016901214 |
| ENSG00000105696.9 | AC003112.1 | protein_coding | 808,8583933 | 0,297446984 | 0,088861402 | 3,34731364 | 0,000815988 | 0,003227391 |
| ENSG00000105698.16 | USF2 | protein_coding | 5137,544342 | -0,35005237 | 0,056381088 | -6,208684221 | 5,34E-10 | 5,17E-09 |
| ENSG00000105699.16 | LSR | protein_coding | 9483,755215 | 0,363245542 | 0,049160363 | 7,388992213 | 1,48E-13 | 2,01E-12 |
| ENSG00000105708.9 | ZNF14 | protein_coding | 2156,604849 | 0,215037781 | 0,061564498 | 3,492886159 | 0,00047783 | 0,001988809 |
| ENSG00000105722.10 | ERF | protein_coding | 1462,458951 | 0,575367407 | 0,078164048 | 7,361023692 | 1,83E-13 | 2,45E-12 |
| ENSG00000105737.9 | GRIK5 | protein_coding | 76,33713425 | 0,691212047 | 0,260534868 | 2,653050061 | 0,007976804 | 0,024612732 |
| ENSG00000105784.15 | RUNDC3B | protein_coding | 688,5393389 | -0,51873723 | 0,094252919 | -5,50367281 | 3,72E-08 | 2,90E-07 |
| ENSG00000105793.15 | GTPBP10 | protein_coding | 3568,587336 | 0,230871476 | 0,075140276 | 3,072539628 | 0,002122456 | 0,007617088 |
| ENSG00000105808.17 | RASA4 | protein_coding | 20672,61654 | -0,275066242 | 0,050972767 | -5,396337264 | 6,80E-08 | 5,12E-07 |
| ENSG00000105810.9 | CDK6 | protein_coding | 645,441978 | 0,683421832 | 0,105370966 | 6,485864698 | 8,82E-11 | 9,24E-10 |
| ENSG00000105819.14 | PMPCB | protein_coding | 12632,16716 | -0,130999428 | 0,04565508 | -2,869328623 | 0,004113442 | 0,013743426 |
| ENSG00000105821.15 | DNAJC2 | protein_coding | 3222,439728 | 0,170889986 | 0,060003976 | 2,847977684 | 0,004399801 | 0,014555426 |
| ENSG00000105835.12 | NAMPT | protein_coding | 18528,02306 | 1,814649737 | 0,061637494 | 29,44067987 | 1,66E-190 | 3,22E-188 |
| ENSG00000105852.11 | PON3 | protein_coding | 11244,98412 | 0,176030069 | 0,048657253 | 3,617755983 | 0,000297168 | 0,001292054 |
| ENSG00000105855.10 | ITGB8 | protein_coding | 1923,574755 | 0,249470864 | 0,070740976 | 3,526539778 | 0,000421028 | 0,001774371 |
| ENSG00000105856.14 | HBP1 | protein_coding | 5563,010896 | 0,211047807 | 0,065310943 | 3,231431029 | 0,00123172 | 0,004672193 |
| ENSG00000105879.12 | CBLL1 | protein_coding | 5555,161961 | 0,192581631 | 0,050773169 | 3,79298036 | 0,00014885 | 0,000686149 |
| ENSG00000105887.11 | MTPN | protein_coding | 37888,08812 | -0,275400255 | 0,043163755 | -6,380358987 | 1,77E-10 | 1,80E-09 |
| ENSG00000105894.11 | PTN | protein_coding | 18,5326252 | 2,537649871 | 0,633718806 | 4,004378356 | 6,22E-05 | 0,00030707 |
| ENSG00000105939.13 | ZC3HAV1 | protein_coding | 16059,74606 | 1,280741664 | 0,053716386 | 23,84266237 | 1,21E-125 | 1,70E-123 |

| | | | | | | | | |
|---|---|---|---|---|---|---|---|---|
| ENSG00000105948.13 | TTC26 | protein_coding | 2608,907744 | -0,591012215 | 0,0641818 | -9,2084082 | 3,31E-20 | 6,79E-19 |
| ENSG00000105963.15 | ADAP1 | protein_coding | 940,6942235 | 0,263432622 | 0,100043614 | 2,633177784 | 0,008459006 | 0,025908798 |
| ENSG00000105968.18 | H2AFV | protein_coding | 23045,18619 | -0,684872669 | 0,048789819 | -14,03720442 | 9,23E-45 | 4,73E-43 |
| ENSG00000105971.15 | CAV2 | protein_coding | 345,4886887 | 1,917568999 | 0,146930118 | 13,05089127 | 6,28E-39 | 2,72E-37 |
| ENSG00000105976.15 | MET | protein_coding | 10621,42946 | 0,624985103 | 0,050614686 | 12,34790054 | 5,00E-35 | 1,91E-33 |
| ENSG00000106003.13 | LFNG | protein_coding | 1095,491482 | -0,933320122 | 0,082028487 | -11,37799993 | 5,38E-30 | 1,69E-28 |
| ENSG00000106009.16 | BRAT1 | protein_coding | 1372,326509 | -0,253298481 | 0,08271469 | -3,06231555 | 0,002196318 | 0,007843395 |
| ENSG00000106012.18 | IQCE | protein_coding | 1636,248663 | -0,183238405 | 0,066352828 | -2,761576431 | 0,005752305 | 0,018475833 |
| ENSG00000106013.15 | ANKRD7 | protein_coding | 368,9006054 | -0,463317053 | 0,140685153 | -3,293290324 | 0,000990222 | 0,003839604 |
| ENSG00000106018.14 | VIPR2 | protein_coding | 57,11166012 | -1,962258516 | 0,373178648 | -5,258228266 | 1,45E-07 | 1,05E-06 |
| ENSG00000106025.8 | TSPAN12 | protein_coding | 3196,998598 | -0,188631335 | 0,063283752 | -2,980723018 | 0,002875688 | 0,009981319 |
| ENSG00000106028.11 | SSBP1 | protein_coding | 11331,95324 | -0,141293446 | 0,043472712 | -3,250164029 | 0,001153385 | 0,004405487 |
| ENSG00000106069.22 | CHN2 | protein_coding | 1323,677624 | 0,384743888 | 0,081680313 | 4,710362539 | 2,47E-06 | 1,52E-05 |
| ENSG00000106070.19 | GRB10 | protein_coding | 191,8984067 | -0,752853273 | 0,207003976 | -3,636902475 | 0,000275936 | 0,001205561 |
| ENSG00000106080.10 | FKBP14 | protein_coding | 937,8273771 | 0,70552698 | 0,10685226 | 6,602826937 | 4,03E-11 | 4,37E-10 |
| ENSG00000106089.12 | STX1A | protein_coding | 15397,85362 | -0,225869561 | 0,049903826 | -4,526097056 | 6,01E-06 | 3,48E-05 |
| ENSG00000106105.14 | GARS | protein_coding | 9849,727655 | 0,18550136 | 0,048684058 | 3,810310134 | 0,000138793 | 0,000642815 |
| ENSG00000106133.18 | NSUN5P2 | transcribed_unproc | 540,6214478 | -0,319793667 | 0,105089563 | -3,043058299 | 0,00234187 | 0,008309133 |
| ENSG00000106153.13 | CHCHD2 | protein_coding | 20971,3023 | -0,317105896 | 0,054446497 | -5,824174488 | 5,74E-09 | 4,93E-08 |
| ENSG00000106244.13 | PDAP1 | protein_coding | 12546,27475 | -0,133564999 | 0,044035558 | -3,033116979 | 0,002420418 | 0,008554118 |
| ENSG00000106261.17 | ZKSCAN1 | protein_coding | 49294,97402 | -0,113569506 | 0,043389126 | -2,617464725 | 0,008858564 | 0,026985712 |
| ENSG00000106268.15 | NUDT1 | protein_coding | 409,4453774 | -0,456982437 | 0,121871928 | -3,749693988 | 0,00017705 | 0,00080218 |
| ENSG00000106302.9 | HYAL4 | protein_coding | 48,10041137 | 1,137141552 | 0,382645696 | 2,971787121 | 0,002960719 | 0,010232277 |
| ENSG00000106330.12 | MOSPD3 | protein_coding | 588,7001339 | -0,439389163 | 0,109808581 | -4,001410068 | 6,30E-05 | 0,000310744 |
| ENSG00000106336.13 | FBXO24 | protein_coding | 84,79948165 | -0,649067641 | 0,2642909 | -2,455883428 | 0,014053874 | 0,040164886 |
| ENSG00000106346.11 | USP42 | protein_coding | 1696,68976 | 0,41196619 | 0,078414136 | 5,253723498 | 1,49E-07 | 1,08E-06 |
| ENSG00000106348.18 | IMPDH1 | protein_coding | 2679,203541 | -0,41796453 | 0,074133743 | -5,637979622 | 1,72E-08 | 1,40E-07 |
| ENSG00000106351.13 | AGFG2 | protein_coding | 2388,383617 | 0,302424103 | 0,070053143 | 4,317066887 | 1,58E-05 | 8,57E-05 |
| ENSG00000106355.10 | LSM5 | protein_coding | 6522,90567 | -0,424578687 | 0,055527681 | -7,646252825 | 2,07E-14 | 2,97E-13 |
| ENSG00000106392.10 | C1GALT1 | protein_coding | 5392,41692 | 0,290781134 | 0,060855975 | 4,778185462 | 1,77E-06 | 1,11E-05 |
| ENSG00000106397.12 | PLOD3 | protein_coding | 5354,772147 | 0,35119244 | 0,051894232 | 6,767465768 | 1,31E-11 | 1,49E-10 |
| ENSG00000106399.11 | RPA3 | protein_coding | 3637,760178 | -0,195436626 | 0,058186586 | -3,358791764 | 0,00078284 | 0,003107752 |
| ENSG00000106400.12 | ZNHIT1 | protein_coding | 5160,38878 | -0,329322691 | 0,053241821 | -6,185413764 | 6,19E-10 | 5,94E-09 |
| ENSG00000106459.15 | NRF1 | protein_coding | 1884,100882 | -0,236830214 | 0,062871861 | -3,766871388 | 0,000165306 | 0,000753491 |
| ENSG00000106462.10 | EZH2 | protein_coding | 8460,703157 | -0,187536335 | 0,04438988 | -4,224754294 | 2,39E-05 | 0,00012636 |
| ENSG00000106477.19 | CEP41 | protein_coding | 3739,972911 | -0,277386452 | 0,063187535 | -4,389891967 | 1,13E-05 | 6,30E-05 |

| | | | | | | | | |
|---|---|---|---|---|---|---|---|---|
| ENSG00000106484.15 | MEST | protein_coding | 18703,98778 | -0,304982779 | 0,050526955 | -6,036041182 | 1,58E-09 | 1,44E-08 |
| ENSG00000106524.9 | ANKMY2 | protein_coding | 1578,806417 | 0,49066121 | 0,074916568 | 6,549435232 | 5,78E-11 | 6,16E-10 |
| ENSG00000106537.8 | TSPAN13 | protein_coding | 5077,652223 | -0,206846425 | 0,054688149 | -3,782289778 | 0,000155392 | 0,000712661 |
| ENSG00000106541.12 | AGR2 | protein_coding | 2468,309808 | 0,556423789 | 0,066761489 | 8,33450243 | 7,78E-17 | 1,32E-15 |
| ENSG00000106546.14 | AHR | protein_coding | 2211,398007 | 0,404944343 | 0,088449078 | 4,578276595 | 4,69E-06 | 2,76E-05 |
| ENSG00000106560.11 | GIMAP2 | protein_coding | 534,0440535 | 4,618087897 | 0,191219898 | 24,15066599 | 7,35E-129 | 1,07E-126 |
| ENSG00000106603.18 | COA1 | protein_coding | 5491,054464 | -0,21656113 | 0,062045908 | -3,490337028 | 0,000482412 | 0,002007136 |
| ENSG00000106617.14 | PRKAG2 | protein_coding | 4591,541807 | -0,261227675 | 0,055050099 | -4,74527167 | 2,08E-06 | 1,29E-05 |
| ENSG00000106624.11 | AEBP1 | protein_coding | 423,6168049 | 0,582042173 | 0,135664094 | 4,290318498 | 1,78E-05 | 9,59E-05 |
| ENSG00000106628.10 | POLD2 | protein_coding | 6941,990929 | -0,445844386 | 0,052081213 | -8,560560726 | 1,12E-17 | 2,01E-16 |
| ENSG00000106631.8 | MYL7 | protein_coding | 348,6423135 | -0,616294786 | 0,181882816 | -3,388416778 | 0,000702974 | 0,00282205 |
| ENSG00000106633.16 | GCK | protein_coding | 48777,07408 | -0,190184247 | 0,046589012 | -4,082169557 | 4,46E-05 | 0,000225531 |
| ENSG00000106638.16 | TBL2 | protein_coding | 3366,708855 | 0,232412675 | 0,061545022 | 3,776303403 | 0,000159173 | 0,000728365 |
| ENSG00000106665.15 | CLIP2 | protein_coding | 8137,375468 | 0,306664766 | 0,061963163 | 4,949146425 | 7,45E-07 | 4,93E-06 |
| ENSG00000106682.14 | EIF4H | protein_coding | 39316,47712 | -0,120955202 | 0,040317659 | -3,000055186 | 0,002699307 | 0,009431474 |
| ENSG00000106686.16 | SPATA6L | protein_coding | 346,700549 | 0,479357221 | 0,135179117 | 3,546089296 | 0,000390994 | 0,00165804 |
| ENSG00000106692.14 | FKTN | protein_coding | 5535,117824 | -0,165099092 | 0,053560673 | -3,082468606 | 0,002052914 | 0,007393446 |
| ENSG00000106701.12 | FSD1L | protein_coding | 7807,803276 | -0,269179533 | 0,051036838 | -5,274220397 | 1,33E-07 | 9,71E-07 |
| ENSG00000106723.17 | SPIN1 | protein_coding | 19734,66977 | -0,195541287 | 0,046508568 | -4,204414256 | 2,62E-05 | 0,000137022 |
| ENSG00000106733.21 | NMRK1 | protein_coding | 3669,412646 | 0,242725717 | 0,061450138 | 3,949962104 | 7,82E-05 | 0,000379014 |
| ENSG00000106772.18 | PRUNE2 | protein_coding | 5999,197361 | 0,234624461 | 0,074220076 | 3,161199418 | 0,001571209 | 0,005834104 |
| ENSG00000106780.9 | MEGF9 | protein_coding | 11184,61713 | -0,207951299 | 0,046043739 | -4,516385979 | 6,29E-06 | 3,63E-05 |
| ENSG00000106785.15 | TRIM14 | protein_coding | 9114,698625 | 3,508937473 | 0,080211075 | 43,74629668 | 0 | 0 |
| ENSG00000106789.13 | CORO2A | protein_coding | 741,2790918 | 0,807788124 | 0,093122621 | 8,674456493 | 4,16E-18 | 7,64E-17 |
| ENSG00000106799.13 | TGFBR1 | protein_coding | 8042,386677 | -0,556501567 | 0,055375778 | -10,04954852 | 9,23E-24 | 2,26E-22 |
| ENSG00000106804.7 | C5 | protein_coding | 2456,811223 | 0,402225848 | 0,074709789 | 5,383843969 | 7,29E-08 | 5,46E-07 |
| ENSG00000106823.12 | ECM2 | protein_coding | 136,1059886 | 1,033950542 | 0,246298375 | 4,197959242 | 2,69E-05 | 0,000140658 |
| ENSG00000106829.18 | TLE4 | protein_coding | 9108,103906 | 0,511311989 | 0,051641512 | 9,90118163 | 4,11E-23 | 9,76E-22 |
| ENSG00000106853.20 | PTGR1 | protein_coding | 7458,116683 | -0,473174327 | 0,054712343 | -8,648401822 | 5,22E-18 | 9,53E-17 |
| ENSG00000106868.16 | SUSD1 | protein_coding | 5560,546865 | 0,383551767 | 0,051084478 | 7,50818615 | 6,00E-14 | 8,37E-13 |
| ENSG00000106948.16 | AKNA | protein_coding | 434,1251855 | 0,417676509 | 0,161106416 | 2,592550434 | 0,009526721 | 0,028737197 |
| ENSG00000106992.19 | AK1 | protein_coding | 1948,805003 | -0,221947828 | 0,082021156 | -2,705982683 | 0,00681026 | 0,02143946 |
| ENSG00000106993.12 | CDC37L1 | protein_coding | 2973,018543 | 0,144086329 | 0,059227642 | 2,432754788 | 0,014984445 | 0,04238312 |
| ENSG00000107036.12 | RIC1 | protein_coding | 4362,834756 | 0,536865932 | 0,066768423 | 8,04071604 | 8,93E-16 | 1,41E-14 |
| ENSG00000107099.15 | DOCK8 | protein_coding | 5024,538906 | -0,275871887 | 0,06128982 | -4,501104547 | 6,76E-06 | 3,88E-05 |
| ENSG00000107130.10 | NCS1 | protein_coding | 2487,23027 | -0,500532878 | 0,073170655 | -6,840623217 | 7,88E-12 | 9,13E-11 |

| | | | | | | | | |
|---|---|---|---|---|---|---|---|---|
| ENSG00000107164.16 | FUBP3 | protein_coding | 10553,55463 | -0,221833771 | 0,045931522 | -4,829663002 | 1,37E-06 | 8,71E-06 |
| ENSG00000107175.12 | CREB3 | protein_coding | 3447,154157 | 0,30089437 | 0,053716227 | 5,601554415 | 2,12E-08 | 1,71E-07 |
| ENSG00000107185.9 | RGP1 | protein_coding | 13319,0204 | -0,153611077 | 0,043862454 | -3,502108555 | 0,000461592 | 0,001930861 |
| ENSG00000107186.16 | MPDZ | protein_coding | 2741,23094 | -0,230538217 | 0,072208651 | -3,192667546 | 0,001409651 | 0,005283904 |
| ENSG00000107201.10 | DDX58 | protein_coding | 25674,61424 | 5,514912085 | 0,053934041 | 102,2529004 | 0 | 0 |
| ENSG00000107242.19 | PIP5K1B | protein_coding | 624,2657542 | 0,380487046 | 0,108437344 | 3,508819291 | 0,000450101 | 0,001888761 |
| ENSG00000107249.23 | GLIS3 | protein_coding | 7050,341839 | 0,280540476 | 0,056679749 | 4,949571589 | 7,44E-07 | 4,92E-06 |
| ENSG00000107282.8 | APBA1 | protein_coding | 796,2404773 | -0,496614307 | 0,100672108 | -4,93298806 | 8,10E-07 | 5,33E-06 |
| ENSG00000107290.14 | SETX | protein_coding | 20534,59714 | 0,688643172 | 0,045724483 | 15,06070982 | 2,94E-51 | 1,75E-49 |
| ENSG00000107295.10 | SH3GL2 | protein_coding | 8007,818114 | 0,200426058 | 0,045926691 | 4,364043087 | 1,28E-05 | 7,04E-05 |
| ENSG00000107331.17 | ABCA2 | protein_coding | 14908,49247 | -0,48937229 | 0,05634049 | -8,685978634 | 3,75E-18 | 6,92E-17 |
| ENSG00000107338.10 | SHB | protein_coding | 1847,279058 | 0,327280573 | 0,070337783 | 4,652983915 | 3,27E-06 | 1,97E-05 |
| ENSG00000107372.13 | ZFAND5 | protein_coding | 17434,28401 | 0,241439196 | 0,046033251 | 5,24488689 | 1,56E-07 | 1,13E-06 |
| ENSG00000107518.18 | ATRNL1 | protein_coding | 4450,204061 | -0,233032141 | 0,059317401 | -3,928562922 | 8,55E-05 | 0,000412593 |
| ENSG00000107537.14 | PHYH | protein_coding | 1810,456704 | -0,417907009 | 0,07361772 | -5,676717608 | 1,37E-08 | 1,13E-07 |
| ENSG00000107551.21 | RASSF4 | protein_coding | 4738,673842 | 0,219413889 | 0,053109393 | 4,131357486 | 3,61E-05 | 0,000184695 |
| ENSG00000107554.17 | DNMBP | protein_coding | 1962,228612 | 0,267221365 | 0,0706367 | 3,783038645 | 0,000154925 | 0,000711096 |
| ENSG00000107560.12 | RAB11FIP2 | protein_coding | 1846,071603 | 0,189986356 | 0,074395156 | 2,553746304 | 0,010657087 | 0,03160063 |
| ENSG00000107562.16 | CXCL12 | protein_coding | 5397,115864 | 0,441870939 | 0,054841887 | 8,057179774 | 7,81E-16 | 1,24E-14 |
| ENSG00000107566.14 | ERLIN1 | protein_coding | 10948,29037 | 0,133013682 | 0,054118622 | 2,457817207 | 0,01397843 | 0,039988739 |
| ENSG00000107643.16 | MAPK8 | protein_coding | 11645,3518 | -0,252156451 | 0,04665634 | -5,404548438 | 6,50E-08 | 4,91E-07 |
| ENSG00000107672.15 | NSMCE4A | protein_coding | 1820,837352 | 0,247355417 | 0,072124262 | 3,429572934 | 0,000604532 | 0,002460653 |
| ENSG00000107719.9 | PALD1 | protein_coding | 90,86069445 | -1,504450269 | 0,35579855 | -4,22837662 | 2,35E-05 | 0,000124518 |
| ENSG00000107736.20 | CDH23 | protein_coding | 1311,921126 | -0,692436956 | 0,074402013 | -9,306696534 | 1,32E-20 | 2,79E-19 |
| ENSG00000107738.20 | VSIR | protein_coding | 284,9304616 | 0,54072745 | 0,209614347 | 2,579629963 | 0,009890623 | 0,029640267 |
| ENSG00000107742.13 | SPOCK2 | protein_coding | 1087,60328 | -1,279881046 | 0,115288411 | -11,10155858 | 1,23E-28 | 3,68E-27 |
| ENSG00000107758.15 | PPP3CB | protein_coding | 11562,89349 | -0,411929609 | 0,046406842 | -8,876484463 | 6,90E-19 | 1,32E-17 |
| ENSG00000107771.17 | CCSER2 | protein_coding | 7792,213613 | 0,44093867 | 0,043778067 | 10,0721365 | 7,34E-24 | 1,81E-22 |
| ENSG00000107789.16 | MINPP1 | protein_coding | 7151,063558 | 0,206694547 | 0,048122804 | 4,295147611 | 1,75E-05 | 9,40E-05 |
| ENSG00000107798.18 | LIPA | protein_coding | 27436,6882 | 2,198562524 | 0,044327336 | 49,5983455 | 0 | 0 |
| ENSG00000107815.10 | TWNK | protein_coding | 1568,121664 | 0,234258499 | 0,070285926 | 3,332936068 | 0,000859347 | 0,003380468 |
| ENSG00000107829.15 | FBXW4 | protein_coding | 3696,646792 | 0,239307328 | 0,072252319 | 3,31210583 | 0,000925965 | 0,003617242 |
| ENSG00000107854.6 | TNKS2 | protein_coding | 15973,00342 | 0,30542652 | 0,044530227 | 6,8588584 | 6,94E-12 | 8,11E-11 |
| ENSG00000107862.4 | GBF1 | protein_coding | 14271,56136 | 0,24490482 | 0,043602017 | 5,616823167 | 1,95E-08 | 1,57E-07 |
| ENSG00000107863.18 | ARHGAP21 | protein_coding | 3248,76081 | 0,525504751 | 0,060145339 | 8,737248177 | 2,39E-18 | 4,45E-17 |
| ENSG00000107864.15 | CPEB3 | protein_coding | 1439,596743 | 0,23167533 | 0,078977185 | 2,933446291 | 0,003352216 | 0,011455145 |

| | | | | | | | | |
|---|---|---|---|---|---|---|---|---|
| ENSG00000107874.11 | CUEDC2 | protein_coding | 3207,989184 | -0,249611795 | 0,055015687 | -4,537102196 | 5,70E-06 | 3,31E-05 |
| ENSG00000107890.16 | ANKRD26 | protein_coding | 2326,932241 | 0,160764568 | 0,067470026 | 2,382755381 | 0,017183609 | 0,047714722 |
| ENSG00000107897.19 | ACBD5 | protein_coding | 3702,734616 | 0,262521636 | 0,054401835 | 4,825602606 | 1,40E-06 | 8,88E-06 |
| ENSG00000107954.10 | NEURL1 | protein_coding | 5447,465511 | -0,184982674 | 0,053789287 | -3,439024465 | 0,000583814 | 0,002388419 |
| ENSG00000107957.16 | SH3PXD2A | protein_coding | 5668,315369 | -0,291594079 | 0,053620044 | -5,438154444 | 5,38E-08 | 4,11E-07 |
| ENSG00000107959.16 | PITRM1 | protein_coding | 6275,220725 | 0,148348373 | 0,051068838 | 2,904870727 | 0,003674048 | 0,012430355 |
| ENSG00000107968.10 | MAP3K8 | protein_coding | 217,3877122 | 1,738395082 | 0,183931083 | 9,451339374 | 3,35E-21 | 7,25E-20 |
| ENSG00000108021.20 | TASOR2 | protein_coding | 6536,416908 | 0,309410649 | 0,066438565 | 4,65709408 | 3,21E-06 | 1,94E-05 |
| ENSG00000108064.11 | TFAM | protein_coding | 11225,47348 | -0,132175919 | 0,049365949 | -2,677471435 | 0,007418018 | 0,023084773 |
| ENSG00000108100.18 | CCNY | protein_coding | 13861,92289 | -0,415320514 | 0,041608552 | -9,981614093 | 1,83E-23 | 4,43E-22 |
| ENSG00000108106.14 | UBE2S | protein_coding | 5301,798095 | -0,60193241 | 0,184431381 | -3,263720133 | 0,001099597 | 0,004221531 |
| ENSG00000108176.15 | DNAJC12 | protein_coding | 5702,632977 | 0,456566594 | 0,05536651 | 8,246259169 | 1,63E-16 | 2,71E-15 |
| ENSG00000108187.16 | PBLD | protein_coding | 6592,291657 | 0,315338196 | 0,05282932 | 5,968999703 | 2,39E-09 | 2,13E-08 |
| ENSG00000108239.8 | TBC1D12 | protein_coding | 2193,921188 | 0,215884297 | 0,062651564 | 3,445792621 | 0,000569387 | 0,002334487 |
| ENSG00000108256.9 | NUFIP2 | protein_coding | 28685,20705 | 0,418348178 | 0,039760256 | 10,52176759 | 6,86E-26 | 1,85E-24 |
| ENSG00000108298.11 | RPL19 | protein_coding | 49173,45745 | -0,377955376 | 0,042495998 | -8,89390518 | 5,90E-19 | 1,13E-17 |
| ENSG00000108309.14 | RUNDC3A | protein_coding | 8960,624744 | -0,339364142 | 0,049228201 | -6,893693752 | 5,44E-12 | 6,40E-11 |
| ENSG00000108375.12 | RNF43 | protein_coding | 5228,46269 | 1,032559387 | 0,078827812 | 13,0989224 | 3,34E-39 | 1,47E-37 |
| ENSG00000108384.15 | RAD51C | protein_coding | 4582,906577 | -0,304719479 | 0,052266278 | -5,830135461 | 5,54E-09 | 4,77E-08 |
| ENSG00000108387.14 | sept-04 | protein_coding | 158,687214 | 0,486002047 | 0,189405217 | 2,56593802 | 0,010289721 | 0,030664813 |
| ENSG00000108389.9 | MTMR4 | protein_coding | 13748,82154 | 0,29719393 | 0,04964691 | 5,986151616 | 2,15E-09 | 1,93E-08 |
| ENSG00000108406.10 | DHX40 | protein_coding | 10840,18484 | -0,22689954 | 0,049607374 | -4,573907525 | 4,79E-06 | 2,81E-05 |
| ENSG00000108424.10 | KPNB1 | protein_coding | 67221,48208 | 0,214371138 | 0,03849117 | 5,569358857 | 2,56E-08 | 2,04E-07 |
| ENSG00000108433.16 | GOSR2 | protein_coding | 6712,802158 | 0,359611715 | 0,047548437 | 7,563060695 | 3,94E-14 | 5,57E-13 |
| ENSG00000108439.10 | PNPO | protein_coding | 2335,819473 | 0,259968183 | 0,061636089 | 4,217791679 | 2,47E-05 | 0,00013005 |
| ENSG00000108448.21 | TRIM16L | protein_coding | 368,3241745 | -0,335376119 | 0,125008838 | -2,68281927 | 0,007300443 | 0,022762941 |
| ENSG00000108468.15 | CBX1 | protein_coding | 26624,46068 | -0,222029286 | 0,04196984 | -5,290210453 | 1,22E-07 | 8,94E-07 |
| ENSG00000108474.16 | PIGL | protein_coding | 1423,159673 | 0,276451912 | 0,073703067 | 3,750887497 | 0,00017621 | 0,000798853 |
| ENSG00000108506.12 | INTS2 | protein_coding | 6282,820478 | -0,16897507 | 0,046039315 | -3,67023423 | 0,000242328 | 0,001070153 |
| ENSG00000108510.10 | MED13 | protein_coding | 21950,58481 | 0,473572589 | 0,041114558 | 11,51836764 | 1,07E-30 | 3,50E-29 |
| ENSG00000108551.5 | RASD1 | protein_coding | 281,2620633 | -0,6369321 | 0,170626352 | -3,73290581 | 0,000189283 | 0,000852457 |
| ENSG00000108557.19 | RAI1 | protein_coding | 3701,386545 | -0,29699606 | 0,079828062 | -3,720446842 | 0,000198871 | 0,000892598 |
| ENSG00000108578.15 | BLMH | protein_coding | 15088,52874 | -0,171850897 | 0,044460499 | -3,865248937 | 0,000110976 | 0,000523352 |
| ENSG00000108582.12 | CPD | protein_coding | 30298,62688 | -0,23696919 | 0,04545879 | -5,212835414 | 1,86E-07 | 1,33E-06 |
| ENSG00000108587.15 | GOSR1 | protein_coding | 13199,69351 | 0,131040452 | 0,046092445 | 2,842991989 | 0,004469219 | 0,014767713 |
| ENSG00000108588.14 | CCDC47 | protein_coding | 25174,10654 | -0,166740989 | 0,046055445 | -3,620440277 | 0,000294102 | 0,001279464 |

| | | | | | | | | |
|---|---|---|---|---|---|---|---|---|
| ENSG00000108590.11 | MED31 | protein_coding | 2089,397848 | -0,202323116 | 0,064947329 | -3,115187618 | 0,001838279 | 0,006707381 |
| ENSG00000108592.17 | FTSJ3 | protein_coding | 6130,949472 | 0,231261326 | 0,055778304 | 4,146080311 | 3,38E-05 | 0,000174088 |
| ENSG00000108599.15 | AKAP10 | protein_coding | 5042,696572 | 0,232220913 | 0,058664479 | 3,958458649 | 7,54E-05 | 0,000366811 |
| ENSG00000108622.11 | ICAM2 | protein_coding | 72,48624641 | 1,349347253 | 0,280653745 | 4,807871903 | 1,53E-06 | 9,65E-06 |
| ENSG00000108651.10 | UTP6 | protein_coding | 7791,932852 | 0,541878982 | 0,054967437 | 9,85818163 | 6,32E-23 | 1,49E-21 |
| ENSG00000108666.10 | C17orf75 | protein_coding | 3462,332656 | -0,18941765 | 0,059124833 | -3,203690216 | 0,001356784 | 0,005106998 |
| ENSG00000108669.16 | CYTH1 | protein_coding | 4791,009571 | 1,049489292 | 0,053385067 | 19,65885489 | 4,86E-86 | 4,90E-84 |
| ENSG00000108671.11 | PSMD11 | protein_coding | 12824,78625 | 0,178377342 | 0,046738998 | 3,816456281 | 0,000135382 | 0,000628571 |
| ENSG00000108679.13 | LGALS3BP | protein_coding | 90976,75462 | 2,82466971 | 0,064422616 | 43,845933 | 0 | 0 |
| ENSG00000108684.14 | ASIC2 | protein_coding | 181,1532894 | -0,476690892 | 0,200664236 | -2,375564783 | 0,017522116 | 0,048494444 |
| ENSG00000108733.10 | PEX12 | protein_coding | 1405,991201 | 0,264433677 | 0,07446986 | 3,550881907 | 0,000383943 | 0,001631213 |
| ENSG00000108771.13 | DHX58 | protein_coding | 1886,16554 | 5,757828827 | 0,114292709 | 50,37791872 | 0 | 0 |
| ENSG00000108773.11 | KAT2A | protein_coding | 5240,317391 | 0,709114286 | 0,054536971 | 13,00245076 | 1,18E-38 | 5,10E-37 |
| ENSG00000108774.14 | RAB5C | protein_coding | 9435,243855 | 0,118310954 | 0,046458122 | 2,54661507 | 0,010877334 | 0,032143615 |
| ENSG00000108784.10 | NAGLU | protein_coding | 3352,987209 | 0,399343068 | 0,066540823 | 6,001474722 | 1,96E-09 | 1,77E-08 |
| ENSG00000108799.13 | EZH1 | protein_coding | 3419,138318 | -0,217309621 | 0,076136403 | -2,854214441 | 0,00431434 | 0,014325339 |
| ENSG00000108819.11 | PPP1R9B | protein_coding | 3202,381464 | 0,166260088 | 0,057263437 | 2,90342487 | 0,003691055 | 0,012478516 |
| ENSG00000108821.13 | COL1A1 | protein_coding | 400,0759438 | 1,010588042 | 0,146403225 | 6,902771709 | 5,10E-12 | 6,03E-11 |
| ENSG00000108828.16 | VAT1 | protein_coding | 8601,714629 | -0,62470319 | 0,056222798 | -11,11120782 | 1,11E-28 | 3,32E-27 |
| ENSG00000108830.10 | RND2 | protein_coding | 199,9400115 | -0,47488312 | 0,177412334 | -2,676719871 | 0,007434677 | 0,023123827 |
| ENSG00000108924.14 | HLF | protein_coding | 1984,543419 | 0,571717135 | 0,065032006 | 8,791319426 | 1,48E-18 | 2,79E-17 |
| ENSG00000108946.14 | PRKAR1A | protein_coding | 50896,05786 | -0,144271533 | 0,038758158 | -3,722352676 | 0,000197375 | 0,000886416 |
| ENSG00000108947.5 | EFNB3 | protein_coding | 1026,522264 | 0,452060721 | 0,088627548 | 5,100679511 | 3,38E-07 | 2,35E-06 |
| ENSG00000108953.17 | YWHAE | protein_coding | 78657,13055 | -0,232892389 | 0,042148266 | -5,525550878 | 3,28E-08 | 2,58E-07 |
| ENSG00000108963.17 | DPH1 | protein_coding | 2204,904439 | -0,236447354 | 0,063225278 | -3,739759824 | 0,000184196 | 0,000831209 |
| ENSG00000108984.15 | MAP2K6 | protein_coding | 11802,39468 | 1,006369453 | 0,048137609 | 20,90609565 | 4,71E-97 | 5,36E-95 |
| ENSG00000109016.17 | DHRS7B | protein_coding | 1394,101925 | 0,225898269 | 0,077979408 | 2,896896424 | 0,003768742 | 0,012708705 |
| ENSG00000109072.14 | VTN | protein_coding | 8387,406437 | 0,151402977 | 0,049956379 | 3,030703563 | 0,002439847 | 0,008610616 |
| ENSG00000109111.15 | SUPT6H | protein_coding | 24428,7798 | 0,209131756 | 0,042870837 | 4,878182258 | 1,07E-06 | 6,93E-06 |
| ENSG00000109113.20 | RAB34 | protein_coding | 895,082459 | 0,252944214 | 0,089148816 | 2,837325566 | 0,00454932 | 0,014999361 |
| ENSG00000109118.14 | PHF12 | protein_coding | 3863,987697 | 0,270835736 | 0,054217577 | 4,995349274 | 5,87E-07 | 3,95E-06 |
| ENSG00000109265.14 | KIAA1211 | protein_coding | 3198,800576 | -0,219354729 | 0,061871438 | -3,545331042 | 0,00039212 | 0,001662191 |
| ENSG00000109320.13 | NFKB1 | protein_coding | 3316,997158 | 0,309020894 | 0,062703107 | 4,928318702 | 8,29E-07 | 5,44E-06 |
| ENSG00000109321.11 | AREG | protein_coding | 23,35723003 | 1,485044343 | 0,498321966 | 2,980090073 | 0,002881636 | 0,009998883 |
| ENSG00000109323.10 | MANBA | protein_coding | 12511,17169 | 0,452570029 | 0,04720869 | 9,586583076 | 9,11E-22 | 2,03E-20 |
| ENSG00000109339.22 | MAPK10 | protein_coding | 5526,577379 | 0,733292189 | 0,08208909 | 8,9328824 | 4,15E-19 | 8,05E-18 |

| | | | | | | | | |
|---|---|---|---|---|---|---|---|---|
| ENSG00000109381.19 | ELF2 | protein_coding | 8295,32718 | 0,1209581 | 0,046727367 | 2,588592265 | 0,009636913 | 0,029018041 |
| ENSG00000109390.12 | NDUFC1 | protein_coding | 3322,258637 | -0,162247352 | 0,0611146 | -2,654805087 | 0,007935425 | 0,024508572 |
| ENSG00000109436.8 | TBC1D9 | protein_coding | 6772,184524 | 0,336115892 | 0,047648283 | 7,054102889 | 1,74E-12 | 2,16E-11 |
| ENSG00000109452.12 | INPP4B | protein_coding | 5916,148021 | 1,397093676 | 0,067125745 | 20,81308268 | 3,29E-96 | 3,73E-94 |
| ENSG00000109458.8 | GAB1 | protein_coding | 1444,452456 | 0,437189613 | 0,101784629 | 4,295241975 | 1,75E-05 | 9,40E-05 |
| ENSG00000109472.14 | CPE | protein_coding | 187958,9684 | -0,140936432 | 0,038291085 | -3,680659117 | 0,000232632 | 0,001031785 |
| ENSG00000109475.16 | RPL34 | protein_coding | 22326,21018 | -0,533583535 | 0,040790771 | -13,08098684 | 4,23E-39 | 1,86E-37 |
| ENSG00000109511.11 | ANXA10 | protein_coding | 67002,63002 | -0,220864799 | 0,039638335 | -5,571999903 | 2,52E-08 | 2,01E-07 |
| ENSG00000109534.17 | GAR1 | protein_coding | 2639,69489 | -0,208583222 | 0,059867612 | -3,484074494 | 0,000493842 | 0,002049012 |
| ENSG00000109572.13 | CLCN3 | protein_coding | 16046,51938 | -0,20775469 | 0,043924556 | -4,729807372 | 2,25E-06 | 1,39E-05 |
| ENSG00000109576.14 | AADAT | protein_coding | 980,8290304 | -0,428019919 | 0,097637499 | -4,383765681 | 1,17E-05 | 6,47E-05 |
| ENSG00000109586.12 | GALNT7 | protein_coding | 1957,788979 | 0,544587674 | 0,068686733 | 7,928571499 | 2,22E-15 | 3,42E-14 |
| ENSG00000109618.12 | SEPSECS | protein_coding | 3398,469245 | 0,147961948 | 0,058661744 | 2,522290328 | 0,011659343 | 0,034105284 |
| ENSG00000109654.15 | TRIM2 | protein_coding | 5805,36062 | -0,340782057 | 0,049498554 | -6,884687114 | 5,79E-12 | 6,81E-11 |
| ENSG00000109670.16 | FBXW7 | protein_coding | 3509,001575 | 0,23963787 | 0,065161927 | 3,67757495 | 0,000235462 | 0,001043515 |
| ENSG00000109674.4 | NEIL3 | protein_coding | 2542,831773 | -0,155965407 | 0,062571989 | -2,492575518 | 0,012682036 | 0,036719828 |
| ENSG00000109680.11 | TBC1D19 | protein_coding | 1563,554179 | -0,186962581 | 0,072880483 | -2,565331256 | 0,010307735 | 0,030710364 |
| ENSG00000109685.18 | NSD2 | protein_coding | 24394,66692 | -0,297041981 | 0,048084511 | -6,177498251 | 6,51E-10 | 6,22E-09 |
| ENSG00000109705.7 | NKX3-2 | protein_coding | 213,9035493 | 0,508035568 | 0,165018281 | 3,078662349 | 0,002079322 | 0,007477785 |
| ENSG00000109738.11 | GLRB | protein_coding | 2162,370673 | -0,206363126 | 0,064460802 | -3,201373844 | 0,00136774 | 0,005143935 |
| ENSG00000109756.9 | RAPGEF2 | protein_coding | 4208,802444 | 0,416080443 | 0,067887008 | 6,129014316 | 8,84E-10 | 8,31E-09 |
| ENSG00000109762.16 | SNX25 | protein_coding | 4357,470266 | 0,151279528 | 0,05348059 | 2,828680977 | 0,004674027 | 0,015363269 |
| ENSG00000109775.11 | UFSP2 | protein_coding | 1665,325801 | 0,251129293 | 0,081295591 | 3,089088701 | 0,002007715 | 0,007250386 |
| ENSG00000109787.13 | KLF3 | protein_coding | 7454,947937 | 0,442273122 | 0,046008833 | 9,612787185 | 7,06E-22 | 1,58E-20 |
| ENSG00000109805.10 | NCAPG | protein_coding | 16130,90094 | -0,303898829 | 0,040226177 | -7,554752859 | 4,20E-14 | 5,92E-13 |
| ENSG00000109814.12 | UGDH | protein_coding | 5724,348521 | 0,266247284 | 0,055718434 | 4,778441635 | 1,77E-06 | 1,11E-05 |
| ENSG00000109854.13 | HTATIP2 | protein_coding | 5410,773765 | 0,569836099 | 0,055721517 | 10,22650003 | 1,51E-24 | 3,84E-23 |
| ENSG00000109861.16 | CTSC | protein_coding | 289,5938607 | 0,579774238 | 0,167138854 | 3,468817827 | 0,000522754 | 0,002155067 |
| ENSG00000109881.17 | CCDC34 | protein_coding | 2675,025187 | -0,577175812 | 0,06817157 | -8,466517815 | 2,53E-17 | 4,40E-16 |
| ENSG00000109911.19 | ELP4 | protein_coding | 8138,618119 | -0,243585265 | 0,056335251 | -4,323851617 | 1,53E-05 | 8,33E-05 |
| ENSG00000109919.10 | MTCH2 | protein_coding | 17583,24628 | -0,363837697 | 0,046241348 | -7,868232897 | 3,60E-15 | 5,49E-14 |
| ENSG00000109920.12 | FNBP4 | protein_coding | 6393,243261 | 0,227802587 | 0,071252712 | 3,197107608 | 0,001388131 | 0,005211049 |
| ENSG00000109944.10 | JHY | protein_coding | 609,4300876 | 0,323224984 | 0,103275674 | 3,129730085 | 0,00174967 | 0,006423637 |
| ENSG00000109971.14 | HSPA8 | protein_coding | 198509,8205 | -0,178236534 | 0,042895301 | -4,155152845 | 3,25E-05 | 0,000167706 |
| ENSG00000110002.15 | VWA5A | protein_coding | 15624,89913 | 0,242507807 | 0,044615867 | 5,435460996 | 5,47E-08 | 4,17E-07 |
| ENSG00000110031.12 | LPXN | protein_coding | 376,4089316 | 0,65354277 | 0,131158765 | 4,982837186 | 6,27E-07 | 4,19E-06 |

| | | | | | | | | |
|---|---|---|---|---|---|---|---|---|
| ENSG00000110048.12 | OSBP | protein_coding | 12402,56159 | 0,223090099 | 0,04291198 | 5,198783582 | 2,01E-07 | 1,43E-06 |
| ENSG00000110057.8 | UNC93B1 | protein_coding | 1973,223148 | 4,630824397 | 0,4734334 | 9,781363955 | 1,35E-22 | 3,15E-21 |
| ENSG00000110075.14 | PPP6R3 | protein_coding | 21558,9064 | -0,118025511 | 0,042981955 | -2,74593169 | 0,006033933 | 0,019259471 |
| ENSG00000110080.18 | ST3GAL4 | protein_coding | 1586,160705 | -0,448941813 | 0,087279339 | -5,14373525 | 2,69E-07 | 1,89E-06 |
| ENSG00000110090.13 | CPT1A | protein_coding | 3174,289306 | -0,537017972 | 0,056480861 | -9,50796362 | 1,94E-21 | 4,24E-20 |
| ENSG00000110107.9 | PRPF19 | protein_coding | 14425,22165 | -0,136455079 | 0,045666989 | -2,988046353 | 0,00280767 | 0,009767826 |
| ENSG00000110172.12 | CHORDC1 | protein_coding | 10999,89377 | 0,156312114 | 0,047587418 | 3,284736166 | 0,001020779 | 0,003947211 |
| ENSG00000110200.8 | ANAPC15 | protein_coding | 1834,115149 | -0,772377892 | 0,070227577 | -10,99821357 | 3,90E-28 | 1,14E-26 |
| ENSG00000110218.9 | PANX1 | protein_coding | 3578,340593 | 0,288552836 | 0,061084016 | 4,723868114 | 2,31E-06 | 1,43E-05 |
| ENSG00000110237.4 | ARHGEF17 | protein_coding | 86,55501897 | -0,901241565 | 0,322112066 | -2,797913084 | 0,005143395 | 0,016744675 |
| ENSG00000110244.7 | APOA4 | protein_coding | 262,5911295 | -0,581588861 | 0,158863644 | -3,660931139 | 0,0002513 | 0,001107601 |
| ENSG00000110274.16 | CEP164 | protein_coding | 4916,091439 | 0,469437926 | 0,050212613 | 9,349004044 | 8,85E-21 | 1,89E-19 |
| ENSG00000110315.7 | RNF141 | protein_coding | 4059,436885 | -0,145382938 | 0,057532269 | -2,526980765 | 0,01150478 | 0,033692571 |
| ENSG00000110321.17 | EIF4G2 | protein_coding | 180970,9736 | -0,185249712 | 0,042360425 | -4,373178791 | 1,22E-05 | 6,77E-05 |
| ENSG00000110330.8 | BIRC2 | protein_coding | 9036,965904 | 0,578042633 | 0,044993669 | 12,84719931 | 8,92E-38 | 3,72E-36 |
| ENSG00000110344.9 | UBE4A | protein_coding | 12312,9975 | -0,198293078 | 0,047275763 | -4,194391905 | 2,74E-05 | 0,000142791 |
| ENSG00000110367.13 | DDX6 | protein_coding | 25720,93369 | -0,274558028 | 0,041405387 | -6,630973613 | 3,33E-11 | 3,64E-10 |
| ENSG00000110395.6 | CBL | protein_coding | 4215,958732 | 0,148042541 | 0,055551481 | 2,664961196 | 0,007699721 | 0,023861296 |
| ENSG00000110429.14 | FBXO3 | protein_coding | 8906,419317 | -0,169553426 | 0,051455469 | -3,295148807 | 0,000983696 | 0,003818904 |
| ENSG00000110435.11 | PDHX | protein_coding | 5532,767773 | 0,220684001 | 0,052254968 | 4,2232157 | 2,41E-05 | 0,000127107 |
| ENSG00000110442.12 | COMMD9 | protein_coding | 6962,507857 | -0,200070186 | 0,056375368 | -3,548893668 | 0,000386853 | 0,001642339 |
| ENSG00000110455.13 | ACCS | protein_coding | 570,9064104 | 0,437275374 | 0,125295857 | 3,489942796 | 0,000483124 | 0,002009356 |
| ENSG00000110492.15 | MDK | protein_coding | 12496,81212 | 1,877350319 | 0,056869891 | 33,01132288 | 5,59E-239 | 1,28E-236 |
| ENSG00000110497.14 | AMBRA1 | protein_coding | 4707,58242 | 0,923038606 | 0,080574638 | 11,45569659 | 2,20E-30 | 7,10E-29 |
| ENSG00000110514.19 | MADD | protein_coding | 14452,41147 | 0,190354431 | 0,051412362 | 3,702503115 | 0,000213483 | 0,000952295 |
| ENSG00000110619.17 | CARS | protein_coding | 3718,920096 | 0,729106973 | 0,075427541 | 9,666323006 | 4,19E-22 | 9,49E-21 |
| ENSG00000110628.15 | SLC22A18 | protein_coding | 149,8279788 | 0,649951139 | 0,195814829 | 3,319213061 | 0,000902715 | 0,003531934 |
| ENSG00000110660.15 | SLC35F2 | protein_coding | 133,0235123 | 0,573186214 | 0,220154629 | 2,603561944 | 0,009226059 | 0,02795388 |
| ENSG00000110665.11 | C11orf21 | protein_coding | 714,6589984 | -0,637238325 | 0,117466016 | -5,424873897 | 5,80E-08 | 4,41E-07 |
| ENSG00000110693.18 | SOX6 | protein_coding | 4791,762888 | 0,331779499 | 0,05290084 | 6,271724624 | 3,57E-10 | 3,52E-09 |
| ENSG00000110697.13 | PITPNM1 | protein_coding | 7581,610543 | -0,292075943 | 0,075142946 | -3,886937587 | 0,000101517 | 0,000482586 |
| ENSG00000110700.7 | RPS13 | protein_coding | 37786,11263 | -0,554877264 | 0,044233586 | -12,54425234 | 4,27E-36 | 1,71E-34 |
| ENSG00000110711.10 | AIP | protein_coding | 4012,141773 | -0,198537101 | 0,061215437 | -3,243252221 | 0,001181735 | 0,004501554 |
| ENSG00000110717.13 | NDUFS8 | protein_coding | 4057,940862 | -0,163266578 | 0,058413785 | -2,795000821 | 0,005189958 | 0,016879175 |
| ENSG00000110756.17 | HPS5 | protein_coding | 4729,836885 | 0,21725712 | 0,048316745 | 4,496518151 | 6,91E-06 | 3,96E-05 |
| ENSG00000110841.14 | PPFIBP1 | protein_coding | 2178,719888 | -0,422582152 | 0,086937557 | -4,860754857 | 1,17E-06 | 7,53E-06 |

| | | | | | | | | |
|---|---|---|---|---|---|---|---|---|
| ENSG00000110851.12 | PRDM4 | protein_coding | 9307,300956 | 0,146992083 | 0,045123349 | 3,257561507 | 0,001123739 | 0,004305954 |
| ENSG00000110881.11 | ASIC1 | protein_coding | 1066,633953 | -0,307112911 | 0,091034784 | -3,373577632 | 0,000741981 | 0,002962791 |
| ENSG00000110917.8 | MLEC | protein_coding | 29451,01273 | -0,385709536 | 0,053253835 | -7,24284994 | 4,39E-13 | 5,71E-12 |
| ENSG00000110925.6 | CSRNP2 | protein_coding | 5109,505341 | 0,431848644 | 0,065631941 | 6,579854854 | 4,71E-11 | 5,06E-10 |
| ENSG00000110931.19 | CAMKK2 | protein_coding | 3364,796017 | 0,175373948 | 0,057579438 | 3,045773869 | 0,002320823 | 0,008244856 |
| ENSG00000110955.9 | ATP5F1B | protein_coding | 50661,20116 | -0,209145134 | 0,039888873 | -5,243194905 | 1,58E-07 | 1,14E-06 |
| ENSG00000110958.16 | PTGES3 | protein_coding | 57480,71381 | -0,158553911 | 0,044715669 | -3,545824413 | 0,000391387 | 0,001659395 |
| ENSG00000111011.18 | RSRC2 | protein_coding | 10851,67309 | 0,104364836 | 0,042400608 | 2,461399505 | 0,013839616 | 0,039633069 |
| ENSG00000111057.11 | KRT18 | protein_coding | 8030,174936 | 0,385724212 | 0,054767594 | 7,042927788 | 1,88E-12 | 2,33E-11 |
| ENSG00000111144.10 | LTA4H | protein_coding | 9635,617684 | -0,246069475 | 0,044051458 | -5,58595532 | 2,32E-08 | 1,86E-07 |
| ENSG00000111186.13 | WNT5B | protein_coding | 498,0721974 | -1,136958633 | 0,122770695 | -9,260830755 | 2,03E-20 | 4,23E-19 |
| ENSG00000111206.12 | FOXM1 | protein_coding | 8739,038262 | -0,19911375 | 0,056363222 | -3,532689286 | 0,000411356 | 0,001736535 |
| ENSG00000111229.16 | ARPC3 | protein_coding | 22728,21735 | -0,120580399 | 0,048005113 | -2,511824102 | 0,012010894 | 0,035018954 |
| ENSG00000111231.9 | GPN3 | protein_coding | 1699,512233 | -0,343904269 | 0,082974725 | -4,144687038 | 3,40E-05 | 0,00017503 |
| ENSG00000111237.18 | VPS29 | protein_coding | 12372,29382 | -0,177711379 | 0,04775518 | -3,721300587 | 0,000198199 | 0,000889762 |
| ENSG00000111252.10 | SH2B3 | protein_coding | 2765,853886 | 0,466858471 | 0,07101148 | 6,574408395 | 4,88E-11 | 5,24E-10 |
| ENSG00000111254.8 | AKAP3 | protein_coding | 109,5049927 | 0,819419243 | 0,262750969 | 3,118615498 | 0,001817029 | 0,006639531 |
| ENSG00000111262.6 | KCNA1 | protein_coding | 21,22951213 | 3,640932426 | 0,73939006 | 4,924237722 | 8,47E-07 | 5,55E-06 |
| ENSG00000111266.8 | DUSP16 | protein_coding | 1859,126543 | 1,021617173 | 0,089691487 | 11,39034721 | 4,67E-30 | 1,48E-28 |
| ENSG00000111271.14 | ACAD10 | protein_coding | 4215,272918 | -0,315700492 | 0,059157922 | -5,336571684 | 9,47E-08 | 6,99E-07 |
| ENSG00000111275.13 | ALDH2 | protein_coding | 3215,96467 | -0,259365169 | 0,089330626 | -2,90342942 | 0,003691002 | 0,012478516 |
| ENSG00000111276.11 | CDKN1B | protein_coding | 4152,140274 | 0,165922333 | 0,052891951 | 3,137005372 | 0,00170683 | 0,006276595 |
| ENSG00000111305.19 | GSG1 | protein_coding | 226,9461934 | -0,840518201 | 0,164836363 | -5,099106686 | 3,41E-07 | 2,36E-06 |
| ENSG00000111321.11 | AC005840.1 | protein_coding | 2316,25528 | 0,335464334 | 0,067616254 | 4,961297238 | 7,00E-07 | 4,65E-06 |
| ENSG00000111328.7 | CDK2AP1 | protein_coding | 4710,603859 | -0,360190474 | 0,054024484 | -6,667171047 | 2,61E-11 | 2,87E-10 |
| ENSG00000111331.13 | OAS3 | protein_coding | 79298,05492 | 4,964871575 | 0,207030995 | 23,98129597 | 4,36E-127 | 6,25E-125 |
| ENSG00000111344.11 | RASAL1 | protein_coding | 233,7992479 | -1,314777425 | 0,16620162 | -7,9107377 | 2,56E-15 | 3,93E-14 |
| ENSG00000111358.14 | GTF2H3 | protein_coding | 4634,096591 | -0,152192412 | 0,058631428 | -2,595748024 | 0,009438525 | 0,028509363 |
| ENSG00000111361.12 | EIF2B1 | protein_coding | 5306,755793 | -0,122042362 | 0,050981929 | -2,393835688 | 0,016673216 | 0,046470706 |
| ENSG00000111371.15 | SLC38A1 | protein_coding | 61,99713295 | 1,397501384 | 0,394563662 | 3,541890751 | 0,00039727 | 0,001680539 |
| ENSG00000111412.6 | C12orf49 | protein_coding | 4492,414884 | -0,380819546 | 0,058274265 | -6,534952388 | 6,36E-11 | 6,75E-10 |
| ENSG00000111445.14 | RFC5 | protein_coding | 6520,803871 | -0,147240959 | 0,047183045 | -3,120632839 | 0,001804629 | 0,006599634 |
| ENSG00000111450.14 | STX2 | protein_coding | 6351,950778 | -0,180234828 | 0,04771593 | -3,777246477 | 0,000158572 | 0,000725909 |
| ENSG00000111481.10 | COPZ1 | protein_coding | 20749,97024 | -0,178020537 | 0,055436702 | -3,211239693 | 0,001321636 | 0,004988048 |
| ENSG00000111530.13 | CAND1 | protein_coding | 41861,77989 | 0,2137501 | 0,048014253 | 4,451805221 | 8,52E-06 | 4,82E-05 |
| ENSG00000111581.10 | NUP107 | protein_coding | 7342,646345 | -0,204201119 | 0,051197099 | -3,988529087 | 6,65E-05 | 0,000326389 |

| | | | | | | | | |
|---|---|---|---|---|---|---|---|---|
| ENSG00000111639.8 | MRPL51 | protein_coding | 11253,12464 | -0,521224091 | 0,043069574 | -12,10190951 | 1,03E-33 | 3,78E-32 |
| ENSG00000111640.15 | GAPDH | protein_coding | 199845,7861 | -0,108776089 | 0,043523397 | -2,4992555 | 0,012445455 | 0,036137092 |
| ENSG00000111641.11 | NOP2 | protein_coding | 3693,863799 | 0,178683653 | 0,062612955 | 2,853780845 | 0,004320232 | 0,014336551 |
| ENSG00000111642.15 | CHD4 | protein_coding | 61504,187 | 0,09140005 | 0,038538818 | 2,371636037 | 0,017709526 | 0,048922892 |
| ENSG00000111665.11 | CDCA3 | protein_coding | 2512,461237 | -1,112353347 | 0,082259119 | -13,52255367 | 1,15E-41 | 5,41E-40 |
| ENSG00000111669.15 | TPI1 | protein_coding | 75330,14414 | -0,146493883 | 0,05207324 | -2,813227744 | 0,00490469 | 0,016048796 |
| ENSG00000111671.9 | SPSB2 | protein_coding | 1519,33154 | 0,22957844 | 0,071843287 | 3,195544754 | 0,001395671 | 0,005235861 |
| ENSG00000111674.9 | ENO2 | protein_coding | 16126,28158 | -0,147268322 | 0,043335803 | -3,398306048 | 0,000678045 | 0,002731231 |
| ENSG00000111676.15 | ATN1 | protein_coding | 7406,34537 | 0,218289667 | 0,050396267 | 4,331465014 | 1,48E-05 | 8,06E-05 |
| ENSG00000111678.11 | C12orf57 | protein_coding | 4095,79475 | -0,174759915 | 0,061472005 | -2,842918724 | 0,004470247 | 0,01476894 |
| ENSG00000111679.17 | PTPN6 | protein_coding | 2617,229885 | 0,27546385 | 0,068445904 | 4,024548371 | 5,71E-05 | 0,000283459 |
| ENSG00000111696.12 | NT5DC3 | protein_coding | 3715,991468 | 0,223181453 | 0,05546917 | 4,023522466 | 5,73E-05 | 0,000284572 |
| ENSG00000111707.12 | SUDS3 | protein_coding | 7783,538879 | -0,169253929 | 0,048939424 | -3,458437291 | 0,000543319 | 0,002234934 |
| ENSG00000111711.10 | GOLT1B | protein_coding | 7194,42062 | -0,239069121 | 0,058157193 | -4,110740373 | 3,94E-05 | 0,000201028 |
| ENSG00000111716.12 | LDHB | protein_coding | 19005,97612 | -0,317722659 | 0,044312392 | -7,170063385 | 7,50E-13 | 9,59E-12 |
| ENSG00000111725.11 | PRKAB1 | protein_coding | 2459,416838 | 0,179720389 | 0,06180756 | 2,907741219 | 0,003640495 | 0,012331876 |
| ENSG00000111726.13 | CMAS | protein_coding | 7567,130185 | -0,169075577 | 0,047257894 | -3,577721368 | 0,000346603 | 0,001487167 |
| ENSG00000111731.12 | C2CD5 | protein_coding | 28503,01163 | -0,101177491 | 0,042311974 | -2,391225995 | 0,016792213 | 0,046773409 |
| ENSG00000111752.11 | PHC1 | protein_coding | 5400,60259 | -0,14598247 | 0,054443939 | -2,681335565 | 0,007332894 | 0,022857792 |
| ENSG00000111775.3 | COX6A1 | protein_coding | 21864,02547 | -0,290987731 | 0,042386053 | -6,865176424 | 6,64E-12 | 7,77E-11 |
| ENSG00000111786.9 | SRSF9 | protein_coding | 12580,98455 | -0,13963907 | 0,046738712 | -2,987653354 | 0,002811282 | 0,00977786 |
| ENSG00000111790.14 | FGFR1OP2 | protein_coding | 4497,022919 | 0,16922718 | 0,06079193 | 2,783711282 | 0,005374085 | 0,017395064 |
| ENSG00000111801.16 | BTN3A3 | protein_coding | 1825,93132 | 4,86560684 | 0,120164896 | 40,49108354 | 0 | 0 |
| ENSG00000111802.14 | TDP2 | protein_coding | 13071,64201 | -0,149134906 | 0,044544866 | -3,347970661 | 0,000814056 | 0,003221447 |
| ENSG00000111834.12 | RSPH4A | protein_coding | 306,6242805 | 0,560068188 | 0,140683276 | 3,981057341 | 6,86E-05 | 0,000336309 |
| ENSG00000111837.11 | MAK | protein_coding | 519,4556932 | 1,285997163 | 0,123494622 | 10,41338594 | 2,15E-25 | 5,68E-24 |
| ENSG00000111846.17 | GCNT2 | protein_coding | 3073,998614 | 1,055798991 | 0,064366795 | 16,40285167 | 1,82E-60 | 1,28E-58 |
| ENSG00000111850.11 | SMIM8 | protein_coding | 2987,852011 | -0,183487748 | 0,065776833 | -2,789549747 | 0,005278139 | 0,017104179 |
| ENSG00000111859.17 | NEDD9 | protein_coding | 4893,76852 | 0,331324302 | 0,053270994 | 6,219600553 | 4,98E-10 | 4,85E-09 |
| ENSG00000111875.8 | ASF1A | protein_coding | 3569,113232 | -0,213164005 | 0,054975908 | -3,877407612 | 0,000105575 | 0,000499978 |
| ENSG00000111885.7 | MAN1A1 | protein_coding | 91911,26807 | -0,29633723 | 0,041348028 | -7,166901251 | 7,67E-13 | 9,80E-12 |
| ENSG00000111897.7 | SERINC1 | protein_coding | 59744,30963 | -0,173334112 | 0,042463196 | -4,081984588 | 4,47E-05 | 0,00022561 |
| ENSG00000111906.17 | HDDC2 | protein_coding | 2497,668697 | -0,417776771 | 0,060141559 | -6,946556992 | 3,74E-12 | 4,49E-11 |
| ENSG00000111911.7 | HINT3 | protein_coding | 5038,224037 | -0,36824191 | 0,052291334 | -7,042121159 | 1,89E-12 | 2,34E-11 |
| ENSG00000111913.19 | RIPOR2 | protein_coding | 211,0458719 | 3,165216035 | 0,253726593 | 12,47490851 | 1,02E-35 | 4,05E-34 |
| ENSG00000111981.5 | ULBP1 | protein_coding | 121,6680202 | 2,204865676 | 0,248226503 | 8,882474891 | 6,54E-19 | 1,25E-17 |

| | | | | | | | | |
|---|---|---|---|---|---|---|---|---|
| ENSG00000112029.10 | FBXO5 | protein_coding | 6006,249746 | 0,118792989 | 0,049055037 | 2,421626738 | 0,015451212 | 0,043517434 |
| ENSG00000112078.14 | KCTD20 | protein_coding | 10117,75723 | -0,257337553 | 0,050526887 | -5,093081488 | 3,52E-07 | 2,44E-06 |
| ENSG00000112096.18 | SOD2 | protein_coding | 11332,24921 | -0,747641291 | 0,046979842 | -15,91408691 | 5,06E-57 | 3,40E-55 |
| ENSG00000112110.10 | MRPL18 | protein_coding | 5010,521725 | -0,274794644 | 0,053557845 | -5,130800946 | 2,89E-07 | 2,01E-06 |
| ENSG00000112118.19 | MCM3 | protein_coding | 30132,38011 | -0,136601571 | 0,045259826 | -3,018163829 | 0,002543114 | 0,008941417 |
| ENSG00000112137.17 | PHACTR1 | protein_coding | 1068,865004 | -0,362610552 | 0,095420325 | -3,800139552 | 0,000144615 | 0,000668267 |
| ENSG00000112146.16 | FBXO9 | protein_coding | 8177,497291 | -0,223704905 | 0,05070322 | -4,412045363 | 1,02E-05 | 5,73E-05 |
| ENSG00000112149.10 | CD83 | protein_coding | 1976,584362 | 0,326656196 | 0,066294255 | 4,927368111 | 8,33E-07 | 5,46E-06 |
| ENSG00000112175.8 | BMP5 | protein_coding | 8298,013376 | -0,158541895 | 0,052259748 | -3,033728647 | 0,002415516 | 0,008538136 |
| ENSG00000112182.15 | BACH2 | protein_coding | 967,7576863 | 0,800482366 | 0,096370132 | 8,306332593 | 9,87E-17 | 1,66E-15 |
| ENSG00000112200.17 | ZNF451 | protein_coding | 9409,144596 | 0,282524903 | 0,045553751 | 6,202011795 | 5,57E-10 | 5,39E-09 |
| ENSG00000112218.9 | GPR63 | protein_coding | 1045,702158 | 0,311226499 | 0,082177033 | 3,787268621 | 0,000152312 | 0,000700961 |
| ENSG00000112234.9 | FBXL4 | protein_coding | 7291,283694 | 0,143500384 | 0,046626991 | 3,077624786 | 0,002086575 | 0,007501471 |
| ENSG00000112246.9 | SIM1 | protein_coding | 23776,55614 | 0,152605748 | 0,042864458 | 3,56019314 | 0,000370582 | 0,001580715 |
| ENSG00000112249.14 | ASCC3 | protein_coding | 19963,42823 | 0,46300952 | 0,053086375 | 8,721814586 | 2,74E-18 | 5,08E-17 |
| ENSG00000112276.14 | BVES | protein_coding | 1865,034827 | 0,212515677 | 0,067674814 | 3,140247653 | 0,001688051 | 0,006212612 |
| ENSG00000112280.16 | COL9A1 | protein_coding | 76,32690059 | 0,810311932 | 0,261629527 | 3,097173097 | 0,001953758 | 0,007077105 |
| ENSG00000112293.15 | GPLD1 | protein_coding | 354,7207793 | -0,45855179 | 0,144388227 | -3,17582534 | 0,001494109 | 0,005581891 |
| ENSG00000112294.12 | ALDH5A1 | protein_coding | 5116,189737 | -0,168638455 | 0,051819041 | -3,254372345 | 0,001136432 | 0,004349592 |
| ENSG00000112297.15 | CRYBG1 | protein_coding | 4161,904363 | 1,844953573 | 0,075737756 | 24,35976006 | 4,57E-131 | 6,86E-129 |
| ENSG00000112299.8 | VNN1 | protein_coding | 107,1386074 | 0,61012448 | 0,235669377 | 2,58890013 | 0,009628301 | 0,029004743 |
| ENSG00000112303.14 | VNN2 | protein_coding | 58,88769043 | 2,768281301 | 0,445064603 | 6,21995387 | 4,97E-10 | 4,84E-09 |
| ENSG00000112306.8 | RPS12 | protein_coding | 34871,92835 | -0,352152647 | 0,041459436 | -8,493908346 | 2,00E-17 | 3,50E-16 |
| ENSG00000112308.13 | C6orf62 | protein_coding | 17920,42787 | 1,106501523 | 0,051562882 | 21,45926454 | 3,74E-102 | 4,43E-100 |
| ENSG00000112309.10 | B3GAT2 | protein_coding | 59,5466968 | -1,231201183 | 0,314982517 | -3,908792133 | 9,28E-05 | 0,000444141 |
| ENSG00000112339.14 | HBS1L | protein_coding | 8834,419603 | -0,134910351 | 0,051335905 | -2,627992055 | 0,008589052 | 0,026264227 |
| ENSG00000112343.11 | TRIM38 | protein_coding | 185,1019285 | 3,239759775 | 0,227277014 | 14,25467414 | 4,19E-46 | 2,20E-44 |
| ENSG00000112365.4 | ZBTB24 | protein_coding | 2040,006405 | 0,238057993 | 0,066257854 | 3,592902248 | 0,000327015 | 0,001410344 |
| ENSG00000112378.12 | PERP | protein_coding | 14630,71744 | -0,396961872 | 0,050809776 | -7,812706617 | 5,60E-15 | 8,43E-14 |
| ENSG00000112379.9 | ARFGEF3 | protein_coding | 20615,38175 | -0,238463135 | 0,071064044 | -3,355608843 | 0,000791905 | 0,00314152 |
| ENSG00000112511.17 | PHF1 | protein_coding | 3072,66837 | 0,296049912 | 0,067026521 | 4,416907024 | 1,00E-05 | 5,62E-05 |
| ENSG00000112530.11 | PACRG | protein_coding | 230,2318001 | -0,653294833 | 0,200567631 | -3,257229637 | 0,001125054 | 0,004309706 |
| ENSG00000112531.17 | QKI | protein_coding | 2224,613523 | 1,0859599 | 0,07259225 | 14,95972238 | 1,35E-50 | 7,85E-49 |
| ENSG00000112539.15 | C6orf118 | protein_coding | 13,78242537 | 2,075376126 | 0,658669699 | 3,150860179 | 0,001627904 | 0,006017818 |
| ENSG00000112561.18 | TFEB | protein_coding | 374,8374012 | 0,762999475 | 0,140321935 | 5,437492534 | 5,40E-08 | 4,13E-07 |
| ENSG00000112562.18 | SMOC2 | protein_coding | 22,78763388 | 1,547466494 | 0,504729648 | 3,065931435 | 0,002169931 | 0,007761476 |

| | | | | | | | | |
|---|---|---|---|---|---|---|---|---|
| ENSG00000112619.8 | PRPH2 | protein_coding | 1079,018265 | 0,50506107 | 0,080676036 | 6,260360523 | 3,84E-10 | 3,77E-09 |
| ENSG00000112640.15 | PPP2R5D | protein_coding | 4445,357579 | -0,165700207 | 0,059969112 | -2,763092555 | 0,005725653 | 0,018398104 |
| ENSG00000112658.8 | SRF | protein_coding | 2592,381013 | 0,183096913 | 0,062406074 | 2,933959826 | 0,003346675 | 0,011437946 |
| ENSG00000112667.12 | DNPH1 | protein_coding | 822,4104181 | -0,284003165 | 0,092836308 | -3,059182017 | 0,002219423 | 0,007920879 |
| ENSG00000112679.14 | DUSP22 | protein_coding | 5049,517875 | -0,198336113 | 0,054224582 | -3,657678961 | 0,00025451 | 0,001120649 |
| ENSG00000112695.11 | COX7A2 | protein_coding | 16782,29946 | -0,289286406 | 0,044819689 | -6,454449175 | 1,09E-10 | 1,13E-09 |
| ENSG00000112697.16 | TMEM30A | protein_coding | 24927,84631 | -0,142336573 | 0,042242897 | -3,369479402 | 0,000753103 | 0,003000814 |
| ENSG00000112699.11 | GMDS | protein_coding | 1017,042043 | -0,280040034 | 0,087308063 | -3,207493367 | 0,001338971 | 0,005046703 |
| ENSG00000112701.18 | SENP6 | protein_coding | 13093,41097 | 0,137962378 | 0,046341505 | 2,977080205 | 0,002910079 | 0,010083527 |
| ENSG00000112742.10 | TTK | protein_coding | 5624,64078 | -0,534044649 | 0,078388492 | -6,812794036 | 9,57E-12 | 1,10E-10 |
| ENSG00000112759.18 | SLC29A1 | protein_coding | 35672,00694 | -0,175479185 | 0,044610091 | -3,933620849 | 8,37E-05 | 0,000404349 |
| ENSG00000112763.17 | BTN2A1 | protein_coding | 11373,15305 | 0,748155008 | 0,056838563 | 13,16280652 | 1,44E-39 | 6,40E-38 |
| ENSG00000112773.16 | TENT5A | protein_coding | 32066,98013 | 2,175400849 | 0,04886759 | 44,51622924 | 0 | 0 |
| ENSG00000112782.17 | CLIC5 | protein_coding | 491,2589567 | 1,934252756 | 0,152382105 | 12,69343769 | 6,43E-37 | 2,63E-35 |
| ENSG00000112787.13 | FBRSL1 | protein_coding | 3664,713412 | -0,328580909 | 0,074557117 | -4,407103192 | 1,05E-05 | 5,85E-05 |
| ENSG00000112796.10 | ENPP5 | protein_coding | 2123,851624 | 0,409360779 | 0,062188187 | 6,582613158 | 4,62E-11 | 4,97E-10 |
| ENSG00000112812.16 | PRSS16 | protein_coding | 1468,217484 | 0,398020929 | 0,081818167 | 4,86470109 | 1,15E-06 | 7,38E-06 |
| ENSG00000112855.17 | HARS2 | protein_coding | 4481,19576 | 0,239181798 | 0,055795462 | 4,286760783 | 1,81E-05 | 9,74E-05 |
| ENSG00000112874.10 | NUDT12 | protein_coding | 4089,519974 | -0,245622596 | 0,052619 | -4,667944953 | 3,04E-06 | 1,84E-05 |
| ENSG00000112877.8 | CEP72 | protein_coding | 1138,805326 | 0,509851376 | 0,081949303 | 6,221546241 | 4,92E-10 | 4,79E-09 |
| ENSG00000112964.14 | GHR | protein_coding | 581,2141584 | -0,554977213 | 0,121883486 | -4,553342152 | 5,28E-06 | 3,08E-05 |
| ENSG00000112983.17 | BRD8 | protein_coding | 12970,9918 | -0,278168947 | 0,052379505 | -5,31064478 | 1,09E-07 | 8,02E-07 |
| ENSG00000112984.12 | KIF20A | protein_coding | 4470,577657 | -1,12217996 | 0,064563359 | -17,38106537 | 1,15E-67 | 8,97E-66 |
| ENSG00000112992.18 | NNT | protein_coding | 14296,24333 | -0,585615789 | 0,065494421 | -8,941460687 | 3,84E-19 | 7,48E-18 |
| ENSG00000113048.16 | MRPS27 | protein_coding | 7306,10098 | -0,231537686 | 0,053882349 | -4,297097139 | 1,73E-05 | 9,33E-05 |
| ENSG00000113070.8 | HBEGF | protein_coding | 203,615232 | 0,957516684 | 0,165867312 | 5,772787149 | 7,80E-09 | 6,56E-08 |
| ENSG00000113161.16 | HMGCR | protein_coding | 55184,08932 | 0,134055936 | 0,044955838 | 2,981947255 | 0,002864213 | 0,009947588 |
| ENSG00000113205.5 | PCDHB3 | protein_coding | 371,7735627 | -0,554840769 | 0,141680159 | -3,91615011 | 9,00E-05 | 0,000431819 |
| ENSG00000113209.8 | PCDHB5 | protein_coding | 1276,7664 | -0,266592913 | 0,082183048 | -3,243891764 | 0,001179085 | 0,00449222 |
| ENSG00000113231.13 | PDE8B | protein_coding | 1300,457243 | 0,302837566 | 0,105515214 | 2,870084352 | 0,004103623 | 0,013714692 |
| ENSG00000113249.12 | HAVCR1 | protein_coding | 434,7061634 | -0,473736699 | 0,12689353 | -3,733340076 | 0,000188957 | 0,000851329 |
| ENSG00000113269.14 | RNF130 | protein_coding | 8805,402929 | -0,126808106 | 0,048742856 | -2,601573141 | 0,009279727 | 0,028105138 |
| ENSG00000113300.12 | CNOT6 | protein_coding | 12947,91016 | -0,255896248 | 0,044254902 | -5,782325474 | 7,37E-09 | 6,22E-08 |
| ENSG00000113303.11 | BTNL8 | protein_coding | 551,9103437 | -0,359311952 | 0,136114631 | -2,639774654 | 0,008296117 | 0,025475757 |
| ENSG00000113356.12 | POLR3G | protein_coding | 229,1570381 | 0,74651028 | 0,19088766 | 3,910730957 | 9,20E-05 | 0,000440966 |
| ENSG00000113360.16 | DROSHA | protein_coding | 8247,646265 | 0,156271582 | 0,062494674 | 2,5005584 | 0,012399769 | 0,036013728 |

| | | | | | | | | |
|---|---|---|---|---|---|---|---|---|
| ENSG00000113361.13 | CDH6 | protein_coding | 2760,479722 | 0,620224768 | 0,064058315 | 9,682189925 | 3,59E-22 | 8,15E-21 |
| ENSG00000113368.12 | LMNB1 | protein_coding | 21601,46386 | -0,448173418 | 0,039505033 | -11,34471693 | 7,88E-30 | 2,46E-28 |
| ENSG00000113369.9 | ARRDC3 | protein_coding | 3710,408037 | 0,671495006 | 0,057419549 | 11,69453646 | 1,36E-31 | 4,66E-30 |
| ENSG00000113391.19 | FAM172A | protein_coding | 4960,355461 | -0,960220782 | 0,063043383 | -15,23111136 | 2,20E-52 | 1,34E-50 |
| ENSG00000113407.13 | TARS | protein_coding | 22876,45575 | 0,580725027 | 0,047504316 | 12,2246792 | 2,29E-34 | 8,57E-33 |
| ENSG00000113441.16 | LNPEP | protein_coding | 14600,76614 | 0,587596775 | 0,043337871 | 13,55850583 | 7,06E-42 | 3,35E-40 |
| ENSG00000113448.19 | PDE4D | protein_coding | 3936,360598 | 0,298314975 | 0,051918676 | 5,745812456 | 9,15E-09 | 7,63E-08 |
| ENSG00000113456.19 | RAD1 | protein_coding | 6201,974823 | -0,169184285 | 0,054282513 | -3,116736444 | 0,00182865 | 0,006675491 |
| ENSG00000113504.21 | SLC12A7 | protein_coding | 10766,39474 | 0,588850088 | 0,050591376 | 11,63933731 | 2,60E-31 | 8,79E-30 |
| ENSG00000113522.14 | RAD50 | protein_coding | 13924,62596 | 0,157033391 | 0,047198269 | 3,32710062 | 0,000877547 | 0,003444237 |
| ENSG00000113532.13 | ST8SIA4 | protein_coding | 3978,293495 | 0,131120653 | 0,052872468 | 2,479941972 | 0,013140377 | 0,037876391 |
| ENSG00000113569.16 | NUP155 | protein_coding | 11008,75099 | -0,237279826 | 0,053328264 | -4,449419646 | 8,61E-06 | 4,87E-05 |
| ENSG00000113580.15 | NR3C1 | protein_coding | 21447,98277 | 0,379438319 | 0,04106499 | 9,239946715 | 2,47E-20 | 5,11E-19 |
| ENSG00000113583.8 | C5orf15 | protein_coding | 13260,98582 | 0,20622694 | 0,048171326 | 4,281114006 | 1,86E-05 | 9,98E-05 |
| ENSG00000113594.10 | LIFR | protein_coding | 1902,295161 | -0,317750808 | 0,066278467 | -4,794178589 | 1,63E-06 | 1,03E-05 |
| ENSG00000113597.18 | TRAPPC13 | protein_coding | 5186,885095 | 0,137265034 | 0,050965404 | 2,693298259 | 0,007074895 | 0,022173323 |
| ENSG00000113615.13 | SEC24A | protein_coding | 7665,60165 | 0,233091883 | 0,055933724 | 4,167287006 | 3,08E-05 | 0,000159685 |
| ENSG00000113621.14 | TXNDC15 | protein_coding | 6029,448841 | 0,196757796 | 0,050883895 | 3,866799058 | 0,000110273 | 0,000520692 |
| ENSG00000113638.14 | TTC33 | protein_coding | 3708,790104 | -0,172802404 | 0,064291652 | -2,687789131 | 0,00719268 | 0,022476751 |
| ENSG00000113648.16 | H2AFY | protein_coding | 36567,96395 | -0,119031814 | 0,042527223 | -2,798955705 | 0,005126817 | 0,016697949 |
| ENSG00000113657.13 | DPYSL3 | protein_coding | 297,4980353 | -0,505892104 | 0,135895908 | -3,722644119 | 0,000197147 | 0,000885746 |
| ENSG00000113712.18 | CSNK1A1 | protein_coding | 22786,85708 | 0,16440665 | 0,043859046 | 3,748523166 | 0,000177879 | 0,000805609 |
| ENSG00000113716.13 | HMGXB3 | protein_coding | 6467,82147 | 0,287387757 | 0,057601132 | 4,989272693 | 6,06E-07 | 4,06E-06 |
| ENSG00000113719.16 | ERGIC1 | protein_coding | 6519,723401 | -0,17320504 | 0,06371307 | -2,718516634 | 0,006557536 | 0,020739578 |
| ENSG00000113742.14 | CPEB4 | protein_coding | 9218,395274 | 0,169345749 | 0,046322988 | 3,655760508 | 0,000256421 | 0,00112774 |
| ENSG00000113749.7 | HRH2 | protein_coding | 44,7116611 | 1,257883496 | 0,361286883 | 3,481674964 | 0,000498288 | 0,002066317 |
| ENSG00000113758.13 | DBN1 | protein_coding | 12631,18977 | -0,244168013 | 0,055445529 | -4,403745717 | 1,06E-05 | 5,94E-05 |
| ENSG00000113761.12 | ZNF346 | protein_coding | 2355,348168 | 0,246521989 | 0,078431209 | 3,143161918 | 0,001671334 | 0,00616318 |
| ENSG00000113763.12 | UNC5A | protein_coding | 3935,923643 | -0,609284625 | 0,064153032 | -9,49736292 | 2,15E-21 | 4,69E-20 |
| ENSG00000113790.11 | EHHADH | protein_coding | 1631,478802 | -0,231748721 | 0,083661584 | -2,770073311 | 0,005604368 | 0,018054775 |
| ENSG00000113805.8 | CNTN3 | protein_coding | 501,7302778 | 0,290175008 | 0,114798774 | 2,527683855 | 0,011481769 | 0,033633933 |
| ENSG00000113810.16 | SMC4 | protein_coding | 35275,33321 | -0,344438864 | 0,04100817 | -8,399274114 | 4,49E-17 | 7,70E-16 |
| ENSG00000113811.11 | SELENOK | protein_coding | 11367,05421 | 0,372335273 | 0,052169775 | 7,136992154 | 9,54E-13 | 1,21E-11 |
| ENSG00000113916.18 | BCL6 | protein_coding | 1624,17562 | 0,346049668 | 0,068365555 | 5,061754722 | 4,15E-07 | 2,85E-06 |
| ENSG00000113971.20 | NPHP3 | protein_coding | 1886,866747 | 0,482763436 | 0,075362304 | 6,405900722 | 1,49E-10 | 1,53E-09 |
| ENSG00000114030.13 | KPNA1 | protein_coding | 12244,0375 | 0,183907176 | 0,046493962 | 3,955506692 | 7,64E-05 | 0,000370889 |

| | | | | | | | | |
|---|---|---|---|---|---|---|---|---|
| ENSG00000114062.20 | UBE3A | protein_coding | 17482,05418 | -0,117136512 | 0,045206263 | -2,591156727 | 0,009565392 | 0,028834526 |
| ENSG00000114098.17 | ARMC8 | protein_coding | 7268,488546 | 0,127726131 | 0,053115568 | 2,404683506 | 0,016186477 | 0,045310461 |
| ENSG00000114115.9 | RBP1 | protein_coding | 1761,839769 | -0,567313695 | 0,076903091 | -7,376994719 | 1,62E-13 | 2,18E-12 |
| ENSG00000114125.14 | RNF7 | protein_coding | 5535,32338 | 0,169534381 | 0,053727217 | 3,155465523 | 0,001602422 | 0,00593434 |
| ENSG00000114126.17 | TFDP2 | protein_coding | 4882,612331 | 0,232275578 | 0,080854319 | 2,872766478 | 0,004068948 | 0,013608907 |
| ENSG00000114127.10 | XRN1 | protein_coding | 13173,45238 | 0,869255876 | 0,076452069 | 11,36994567 | 5,90E-30 | 1,85E-28 |
| ENSG00000114166.8 | KAT2B | protein_coding | 7859,716405 | 0,246537345 | 0,047578672 | 5,181677756 | 2,20E-07 | 1,56E-06 |
| ENSG00000114204.14 | SERPINI2 | protein_coding | 62,45327925 | 1,511806204 | 0,333252567 | 4,536517802 | 5,72E-06 | 3,32E-05 |
| ENSG00000114248.9 | LRRC31 | protein_coding | 23,78005427 | 1,954636125 | 0,542519123 | 3,602888897 | 0,0003147 | 0,001363014 |
| ENSG00000114251.14 | WNT5A | protein_coding | 3922,460246 | 0,156856651 | 0,05707687 | 2,748164881 | 0,005992986 | 0,019136917 |
| ENSG00000114268.12 | PFKFB4 | protein_coding | 16125,53653 | 0,564790142 | 0,047901825 | 11,79057661 | 4,37E-32 | 1,54E-30 |
| ENSG00000114270.17 | COL7A1 | protein_coding | 1978,635968 | 0,331774792 | 0,088816499 | 3,735508537 | 0,000187336 | 0,000844702 |
| ENSG00000114302.16 | PRKAR2A | protein_coding | 10763,29376 | -0,885299324 | 0,067143986 | -13,18508746 | 1,07E-39 | 4,79E-38 |
| ENSG00000114315.4 | HES1 | protein_coding | 27,67900915 | 1,261518786 | 0,443142217 | 2,846758305 | 0,004416688 | 0,014606999 |
| ENSG00000114316.12 | USP4 | protein_coding | 5544,376336 | 0,343851121 | 0,062582887 | 5,49433142 | 3,92E-08 | 3,04E-07 |
| ENSG00000114346.13 | ECT2 | protein_coding | 15646,04029 | -0,667324259 | 0,050571103 | -13,19576247 | 9,28E-40 | 4,17E-38 |
| ENSG00000114353.17 | GNAI2 | protein_coding | 28606,14056 | -0,113238119 | 0,042980726 | -2,634625535 | 0,008423015 | 0,025802858 |
| ENSG00000114354.14 | TFG | protein_coding | 11575,218 | 0,222617842 | 0,044315307 | 5,023497737 | 5,07E-07 | 3,44E-06 |
| ENSG00000114391.13 | RPL24 | protein_coding | 30039,59099 | -0,39700308 | 0,039881528 | -9,954560444 | 2,41E-23 | 5,77E-22 |
| ENSG00000114439.19 | BBX | protein_coding | 6372,921761 | 0,369958853 | 0,05313924 | 6,962065132 | 3,35E-12 | 4,04E-11 |
| ENSG00000114446.5 | IFT57 | protein_coding | 3702,917509 | -0,199402469 | 0,076491156 | -2,606869588 | 0,009137415 | 0,027716101 |
| ENSG00000114450.10 | GNB4 | protein_coding | 6591,543629 | 1,433741657 | 0,053821739 | 26,63870938 | 2,42E-156 | 4,06E-154 |
| ENSG00000114473.14 | IQCG | protein_coding | 672,3257061 | -0,424391114 | 0,111212805 | -3,816027433 | 0,000135617 | 0,000629275 |
| ENSG00000114480.13 | GBE1 | protein_coding | 6282,994738 | -0,380950212 | 0,056781613 | -6,709041797 | 1,96E-11 | 2,19E-10 |
| ENSG00000114520.11 | SNX4 | protein_coding | 11966,14685 | -0,172890831 | 0,045366669 | -3,810965933 | 0,000138425 | 0,000641244 |
| ENSG00000114529.12 | C3orf52 | protein_coding | 1425,368544 | 0,882914237 | 0,071554593 | 12,33902963 | 5,58E-35 | 2,13E-33 |
| ENSG00000114541.15 | FRMD4B | protein_coding | 7360,745432 | -0,218130024 | 0,047355152 | -4,606257511 | 4,10E-06 | 2,43E-05 |
| ENSG00000114573.10 | ATP6V1A | protein_coding | 28363,90238 | -0,271427752 | 0,044729868 | -6,068154527 | 1,29E-09 | 1,20E-08 |
| ENSG00000114631.11 | PODXL2 | protein_coding | 9453,374038 | 0,31511184 | 0,046033685 | 6,845244728 | 7,63E-12 | 8,86E-11 |
| ENSG00000114646.10 | CSPG5 | protein_coding | 599,4515922 | -0,824808845 | 0,108695795 | -7,588231401 | 3,24E-14 | 4,60E-13 |
| ENSG00000114656.12 | KIAA1257 | protein_coding | 126,3256489 | 0,732804336 | 0,25204836 | 2,90739577 | 0,003644518 | 0,012341575 |
| ENSG00000114686.9 | MRPL3 | protein_coding | 11010,98814 | -0,304850829 | 0,052117646 | -5,849282404 | 4,94E-09 | 4,27E-08 |
| ENSG00000114698.15 | PLSCR4 | protein_coding | 2959,286229 | 0,167087035 | 0,067709279 | 2,4677125 | 0,013597948 | 0,039015452 |
| ENSG00000114737.15 | CISH | protein_coding | 314,397649 | -0,944552957 | 0,135977025 | -6,946415788 | 3,75E-12 | 4,49E-11 |
| ENSG00000114738.11 | MAPKAPK3 | protein_coding | 4375,440327 | -0,182153694 | 0,059134268 | -3,080340703 | 0,002067639 | 0,007441716 |
| ENSG00000114739.14 | ACVR2B | protein_coding | 3670,30067 | -0,583004452 | 0,060187346 | -9,686495402 | 3,44E-22 | 7,82E-21 |

| | | | | | | | | |
|---|---|---|---|---|---|---|---|---|
| ENSG00000114757.19 | PEX5L | protein_coding | 336,3329185 | 1,462755336 | 0,207346956 | 7,054626533 | 1,73E-12 | 2,15E-11 |
| ENSG00000114770.17 | ABCC5 | protein_coding | 4172,985478 | -0,189405448 | 0,059835464 | -3,165437917 | 0,001548497 | 0,005760225 |
| ENSG00000114790.13 | ARHGEF26 | protein_coding | 3021,512671 | -0,305578199 | 0,058654787 | -5,209774213 | 1,89E-07 | 1,35E-06 |
| ENSG00000114805.17 | PLCH1 | protein_coding | 1580,010926 | 1,067483083 | 0,087265052 | 12,23265285 | 2,08E-34 | 7,78E-33 |
| ENSG00000114853.14 | ZBTB47 | protein_coding | 348,4374586 | -0,353856806 | 0,129703059 | -2,728207094 | 0,006367962 | 0,02019392 |
| ENSG00000114854.7 | TNNC1 | protein_coding | 7,439742529 | 2,493428045 | 0,89264503 | 2,793303006 | 0,00521728 | 0,016940252 |
| ENSG00000114861.20 | FOXP1 | protein_coding | 6364,28209 | 0,155210615 | 0,049623266 | 3,127779092 | 0,001761325 | 0,006461158 |
| ENSG00000114923.17 | SLC4A3 | protein_coding | 1115,985653 | -0,247916673 | 0,090808544 | -2,730102958 | 0,006331455 | 0,020099933 |
| ENSG00000114933.16 | INO80D | protein_coding | 3948,994702 | 0,233787882 | 0,05942785 | 3,933978467 | 8,36E-05 | 0,000403834 |
| ENSG00000114942.14 | EEF1B2 | protein_coding | 22516,43864 | -0,786634886 | 0,043612914 | -18,03674215 | 1,00E-72 | 8,55E-71 |
| ENSG00000114948.13 | ADAM23 | protein_coding | 619,0508123 | 0,327507858 | 0,119681871 | 2,736486789 | 0,006209909 | 0,019762313 |
| ENSG00000114978.18 | MOB1A | protein_coding | 11155,24039 | 0,343286948 | 0,050985468 | 6,733035097 | 1,66E-11 | 1,86E-10 |
| ENSG00000114982.19 | KANSL3 | protein_coding | 13102,53851 | 0,217537257 | 0,045577888 | 4,77286834 | 1,82E-06 | 1,14E-05 |
| ENSG00000114993.16 | RTKN | protein_coding | 4118,246348 | 0,320741074 | 0,055297483 | 5,800283463 | 6,62E-09 | 5,64E-08 |
| ENSG00000114999.8 | TTL | protein_coding | 6310,525018 | -0,402291467 | 0,066779569 | -6,024169859 | 1,70E-09 | 1,55E-08 |
| ENSG00000115020.17 | PIKFYVE | protein_coding | 9802,743391 | 0,160713542 | 0,045933043 | 3,498865523 | 0,000467242 | 0,001951235 |
| ENSG00000115073.7 | ACTR1B | protein_coding | 5854,348645 | -0,113207307 | 0,047012264 | -2,408037758 | 0,016038522 | 0,044968995 |
| ENSG00000115112.8 | TFCP2L1 | protein_coding | 830,0842155 | -0,797169013 | 0,110336125 | -7,224914004 | 5,01E-13 | 6,49E-12 |
| ENSG00000115128.7 | SF3B6 | protein_coding | 6835,920778 | -0,157115582 | 0,050780744 | -3,0939992 | 0,00197478 | 0,007144058 |
| ENSG00000115129.14 | TP53I3 | protein_coding | 2068,725691 | 0,173938783 | 0,068966877 | 2,522062639 | 0,011666893 | 0,034122934 |
| ENSG00000115137.11 | DNAJC27 | protein_coding | 986,0447008 | 0,25151072 | 0,105438979 | 2,385367562 | 0,017062065 | 0,047425292 |
| ENSG00000115159.16 | GPD2 | protein_coding | 13050,86201 | 0,351377083 | 0,062923869 | 5,584162098 | 2,35E-08 | 1,88E-07 |
| ENSG00000115163.15 | CENPA | protein_coding | 1306,813328 | -1,050843766 | 0,089238049 | -11,77573662 | 5,21E-32 | 1,83E-30 |
| ENSG00000115165.10 | CYTIP | protein_coding | 314,8947012 | 0,640762991 | 0,14748681 | 4,344544373 | 1,40E-05 | 7,64E-05 |
| ENSG00000115170.13 | ACVR1 | protein_coding | 4269,775504 | 0,314218434 | 0,055556909 | 5,655794016 | 1,55E-08 | 1,27E-07 |
| ENSG00000115221.12 | ITGB6 | protein_coding | 110,3393138 | 1,480350941 | 0,23805699 | 6,218472905 | 5,02E-10 | 4,87E-09 |
| ENSG00000115232.14 | ITGA4 | protein_coding | 34,47302627 | 0,991820093 | 0,395756068 | 2,50613995 | 0,01220573 | 0,035523503 |
| ENSG00000115252.18 | PDE1A | protein_coding | 862,2810291 | -1,192034977 | 0,306506353 | -3,889103647 | 0,000100615 | 0,000478501 |
| ENSG00000115255.11 | REEP6 | protein_coding | 202,2693863 | -0,724573728 | 0,165412181 | -4,380413358 | 1,18E-05 | 6,57E-05 |
| ENSG00000115267.8 | IFIH1 | protein_coding | 21980,08453 | 4,755730476 | 0,061070713 | 77,87252296 | 0 | 0 |
| ENSG00000115268.9 | RPS15 | protein_coding | 24020,97712 | -0,226274469 | 0,054587168 | -4,145195225 | 3,40E-05 | 0,000174682 |
| ENSG00000115271.11 | GCA | protein_coding | 6024,486174 | 0,68852204 | 0,064520989 | 10,6712877 | 1,39E-26 | 3,85E-25 |
| ENSG00000115275.13 | MOGS | protein_coding | 6195,859219 | 0,180926767 | 0,054928273 | 3,293873221 | 0,000988171 | 0,003833632 |
| ENSG00000115295.20 | CLIP4 | protein_coding | 1534,187609 | 0,736182201 | 0,074911346 | 9,82737914 | 8,58E-23 | 2,01E-21 |
| ENSG00000115310.18 | RTN4 | protein_coding | 36249,54975 | -0,107755952 | 0,043534709 | -2,475173366 | 0,013317151 | 0,038312342 |
| ENSG00000115317.11 | HTRA2 | protein_coding | 2643,985886 | -0,235158443 | 0,068188281 | -3,448663592 | 0,000563368 | 0,002311493 |

| | | | | | | | | |
|---|---|---|---|---|---|---|---|---|
| ENSG00000115318.12 | LOXL3 | protein_coding | 4622,397133 | -0,171081218 | 0,053062607 | -3,224138967 | 0,001263521 | 0,004783146 |
| ENSG00000115339.13 | GALNT3 | protein_coding | 8845,441797 | 0,381449505 | 0,056035304 | 6,807306748 | 9,94E-12 | 1,14E-10 |
| ENSG00000115380.20 | EFEMP1 | protein_coding | 169,3798759 | 0,623782289 | 0,187368806 | 3,32916829 | 0,000871058 | 0,003419364 |
| ENSG00000115414.19 | FN1 | protein_coding | 4986,221057 | 0,399252395 | 0,067537105 | 5,9116007 | 3,39E-09 | 2,98E-08 |
| ENSG00000115415.18 | STAT1 | protein_coding | 141933,5654 | 4,676622977 | 0,045460004 | 102,8733519 | 0 | 0 |
| ENSG00000115421.13 | PAPOLG | protein_coding | 5178,676284 | 0,190793858 | 0,054615808 | 3,4933816 | 0,000476944 | 0,00198549 |
| ENSG00000115423.18 | DNAH6 | protein_coding | 712,5803299 | 0,893208861 | 0,12253161 | 7,289619861 | 3,11E-13 | 4,09E-12 |
| ENSG00000115425.14 | PECR | protein_coding | 1400,448599 | -0,249450916 | 0,078832837 | -3,164302141 | 0,001554553 | 0,005780842 |
| ENSG00000115459.18 | ELMOD3 | protein_coding | 1395,06807 | 0,314719124 | 0,083198953 | 3,782729383 | 0,000155118 | 0,000711835 |
| ENSG00000115461.5 | IGFBP5 | protein_coding | 38770,03654 | -0,33388115 | 0,050071115 | -6,668138944 | 2,59E-11 | 2,86E-10 |
| ENSG00000115464.14 | USP34 | protein_coding | 23200,18046 | -0,147052014 | 0,045834064 | -3,208356393 | 0,00133496 | 0,005032424 |
| ENSG00000115484.15 | CCT4 | protein_coding | 21521,86178 | -0,160362819 | 0,044618248 | -3,594108395 | 0,000325504 | 0,001404943 |
| ENSG00000115504.14 | EHBP1 | protein_coding | 7603,664358 | -0,196230343 | 0,062160569 | -3,156829923 | 0,001594943 | 0,005910534 |
| ENSG00000115507.10 | OTX1 | protein_coding | 49,41875471 | 1,30216472 | 0,333661479 | 3,902652246 | 9,51E-05 | 0,000454404 |
| ENSG00000115520.8 | COQ10B | protein_coding | 2276,771318 | 0,391343549 | 0,0712382 | 5,493450849 | 3,94E-08 | 3,06E-07 |
| ENSG00000115548.17 | KDM3A | protein_coding | 17167,36613 | 0,223665093 | 0,064321149 | 3,47731807 | 0,000506457 | 0,002095172 |
| ENSG00000115561.16 | CHMP3 | protein_coding | 13158,79855 | -0,489712709 | 0,046570987 | -10,5154033 | 7,34E-26 | 1,97E-24 |
| ENSG00000115594.12 | IL1R1 | protein_coding | 807,4065756 | 0,514153339 | 0,100028539 | 5,140066464 | 2,75E-07 | 1,92E-06 |
| ENSG00000115604.10 | IL18R1 | protein_coding | 1702,575079 | 1,327297645 | 0,087039081 | 15,24944469 | 1,66E-52 | 1,02E-50 |
| ENSG00000115657.14 | ABCB6 | protein_coding | 3533,280627 | 0,150993573 | 0,060160851 | 2,509831084 | 0,012078893 | 0,035186203 |
| ENSG00000115661.14 | STK16 | protein_coding | 3843,736149 | -0,1821555 | 0,064195363 | -2,83751803 | 0,004546578 | 0,014992517 |
| ENSG00000115677.17 | HDLBP | protein_coding | 48773,26648 | -0,14048977 | 0,040098683 | -3,503600575 | 0,000459013 | 0,001921862 |
| ENSG00000115687.13 | PASK | protein_coding | 4325,346847 | -0,502728053 | 0,054728484 | -9,185857397 | 4,08E-20 | 8,32E-19 |
| ENSG00000115694.15 | STK25 | protein_coding | 6401,146521 | -0,186036932 | 0,061345336 | -3,032617373 | 0,002424428 | 0,008565602 |
| ENSG00000115705.21 | TPO | protein_coding | 20,81150503 | -1,435041478 | 0,600913858 | -2,388098491 | 0,016935802 | 0,047117873 |
| ENSG00000115738.10 | ID2 | protein_coding | 16731,4678 | 0,218279535 | 0,043697037 | 4,995293693 | 5,87E-07 | 3,95E-06 |
| ENSG00000115750.17 | TAF1B | protein_coding | 2638,211789 | -0,227927506 | 0,068905327 | -3,307835762 | 0,000940199 | 0,003667755 |
| ENSG00000115758.13 | ODC1 | protein_coding | 22363,93841 | -0,480993726 | 0,043807692 | -10,97966363 | 4,79E-28 | 1,40E-26 |
| ENSG00000115760.14 | BIRC6 | protein_coding | 23322,70839 | 0,42016478 | 0,048277733 | 8,703076046 | 3,23E-18 | 5,97E-17 |
| ENSG00000115761.16 | NOL10 | protein_coding | 5572,605244 | -0,162531326 | 0,057866807 | -2,808714257 | 0,004973977 | 0,016244813 |
| ENSG00000115762.16 | PLEKHB2 | protein_coding | 12290,84907 | -0,37211516 | 0,055534949 | -6,700558254 | 2,08E-11 | 2,31E-10 |
| ENSG00000115816.15 | CEBPZ | protein_coding | 12777,22648 | 0,166231923 | 0,046544408 | 3,57146928 | 0,000354984 | 0,001518499 |
| ENSG00000115828.17 | QPCT | protein_coding | 16532,12764 | 0,253761428 | 0,041976281 | 6,045352765 | 1,49E-09 | 1,37E-08 |
| ENSG00000115839.17 | RAB3GAP1 | protein_coding | 18826,41037 | 0,222989443 | 0,044464341 | 5,015017326 | 5,30E-07 | 3,59E-06 |
| ENSG00000115840.14 | SLC25A12 | protein_coding | 4152,574789 | 0,18266705 | 0,06626392 | 2,75665928 | 0,005839516 | 0,018710559 |
| ENSG00000115875.19 | SRSF7 | protein_coding | 19108,27151 | -0,222421804 | 0,041658899 | -5,339118646 | 9,34E-08 | 6,90E-07 |

| | | | | | | | | |
|---|---|---|---|---|---|---|---|---|
| ENSG00000115904.12 | SOS1 | protein_coding | 13555,68633 | 0,330796173 | 0,076161394 | 4,343357648 | 1,40E-05 | 7,67E-05 |
| ENSG00000115947.13 | ORC4 | protein_coding | 8815,854576 | 0,196366059 | 0,044885419 | 4,37482955 | 1,22E-05 | 6,72E-05 |
| ENSG00000115963.13 | RND3 | protein_coding | 3646,050509 | 1,007788101 | 0,061669817 | 16,34167488 | 4,99E-60 | 3,50E-58 |
| ENSG00000115977.19 | AAK1 | protein_coding | 5601,936825 | -0,189794869 | 0,05201227 | -3,649040274 | 0,000263222 | 0,001155169 |
| ENSG00000115993.13 | TRAK2 | protein_coding | 5093,871427 | -0,278307893 | 0,054165151 | -5,138135596 | 2,77E-07 | 1,94E-06 |
| ENSG00000115998.7 | C2orf42 | protein_coding | 1099,885102 | 0,348202645 | 0,100128428 | 3,477560295 | 0,000505999 | 0,00209405 |
| ENSG00000116001.16 | TIA1 | protein_coding | 16470,78728 | -0,110588871 | 0,045172136 | -2,448165636 | 0,014358563 | 0,040905822 |
| ENSG00000116005.12 | PCYOX1 | protein_coding | 24691,63005 | -0,5582413 | 0,05031374 | -11,09520578 | 1,32E-28 | 3,95E-27 |
| ENSG00000116014.10 | KISS1R | protein_coding | 88,99178841 | -1,065760471 | 0,267240923 | -3,988013732 | 6,66E-05 | 0,000326956 |
| ENSG00000116016.14 | EPAS1 | protein_coding | 16453,9169 | -0,158258328 | 0,062938363 | -2,514497026 | 0,011920231 | 0,034764508 |
| ENSG00000116017.11 | ARID3A | protein_coding | 12651,82434 | -0,641852203 | 0,058829251 | -10,91042615 | 1,03E-27 | 2,96E-26 |
| ENSG00000116030.16 | SUMO1 | protein_coding | 21936,34542 | -0,142610108 | 0,043513279 | -3,277392844 | 0,001047705 | 0,004042995 |
| ENSG00000116039.13 | ATP6V1B1 | protein_coding | 1580,731606 | -0,646534879 | 0,075573266 | -8,555073914 | 1,18E-17 | 2,10E-16 |
| ENSG00000116044.16 | NFE2L2 | protein_coding | 10653,77256 | 0,121426664 | 0,042818313 | 2,835858193 | 0,004570273 | 0,01506624 |
| ENSG00000116095.11 | PLEKHA3 | protein_coding | 2001,23856 | 0,194788107 | 0,063079644 | 3,087970926 | 0,002015282 | 0,007271879 |
| ENSG00000116106.12 | EPHA4 | protein_coding | 2783,116246 | 0,18402526 | 0,064143435 | 2,868964855 | 0,004118176 | 0,0137572 |
| ENSG00000116120.10 | FARSB | protein_coding | 9313,245582 | -0,210823873 | 0,056606663 | -3,724364969 | 0,000195807 | 0,000880254 |
| ENSG00000116127.18 | ALMS1 | protein_coding | 5804,092014 | 0,368239559 | 0,067147998 | 5,483999055 | 4,16E-08 | 3,21E-07 |
| ENSG00000116138.13 | DNAJC16 | protein_coding | 2494,514316 | 0,414974952 | 0,068134818 | 6,090497673 | 1,13E-09 | 1,04E-08 |
| ENSG00000116147.17 | TNR | protein_coding | 14740,719 | -0,24198766 | 0,063961637 | -3,783324986 | 0,000154747 | 0,000710568 |
| ENSG00000116157.6 | GPX7 | protein_coding | 4764,815712 | 0,184689529 | 0,051708966 | 3,57171191 | 0,000354655 | 0,001517669 |
| ENSG00000116161.17 | CACYBP | protein_coding | 16959,80155 | -0,18019941 | 0,050400722 | -3,575333874 | 0,000349781 | 0,001498806 |
| ENSG00000116199.12 | FAM20B | protein_coding | 9848,585846 | -0,67640795 | 0,059167486 | -11,43208868 | 2,89E-30 | 9,26E-29 |
| ENSG00000116209.11 | TMEM59 | protein_coding | 48899,37218 | 0,361522512 | 0,037812342 | 9,560965812 | 1,17E-21 | 2,58E-20 |
| ENSG00000116212.15 | LRRC42 | protein_coding | 9989,811493 | 0,128182777 | 0,04243758 | 3,020501531 | 0,002523564 | 0,008886563 |
| ENSG00000116221.15 | MRPL37 | protein_coding | 6002,845952 | -0,336673844 | 0,056721494 | -5,935560205 | 2,93E-09 | 2,59E-08 |
| ENSG00000116237.16 | ICMT | protein_coding | 25805,00485 | -0,326768085 | 0,046857628 | -6,973636925 | 3,09E-12 | 3,73E-11 |
| ENSG00000116251.11 | RPL22 | protein_coding | 49959,35443 | -0,778711796 | 0,045911858 | -16,96101677 | 1,60E-64 | 1,19E-62 |
| ENSG00000116260.17 | QSOX1 | protein_coding | 13297,61763 | 0,457938376 | 0,065394635 | 7,002690271 | 2,51E-12 | 3,07E-11 |
| ENSG00000116266.11 | STXBP3 | protein_coding | 5094,484204 | 0,192475424 | 0,049589671 | 3,881361151 | 0,000103873 | 0,000493163 |
| ENSG00000116273.6 | PHF13 | protein_coding | 4685,989451 | -0,254139757 | 0,052409827 | -4,849085932 | 1,24E-06 | 7,96E-06 |
| ENSG00000116288.13 | PARK7 | protein_coding | 22113,03981 | -0,359798236 | 0,049137476 | -7,322277541 | 2,44E-13 | 3,24E-12 |
| ENSG00000116299.17 | KIAA1324 | protein_coding | 11532,39724 | 0,221255599 | 0,043568645 | 5,078321792 | 3,81E-07 | 2,62E-06 |
| ENSG00000116337.17 | AMPD2 | protein_coding | 9436,709146 | -0,197085211 | 0,05105345 | -3,860370102 | 0,000113215 | 0,000533354 |
| ENSG00000116459.11 | ATP5PB | protein_coding | 19503,54287 | -0,653971115 | 0,057135355 | -11,44599727 | 2,46E-30 | 7,92E-29 |
| ENSG00000116473.14 | RAP1A | protein_coding | 13237,80669 | -0,11150181 | 0,04441213 | -2,510616154 | 0,012052067 | 0,035117148 |

| | | | | | | | | |
|---|---|---|---|---|---|---|---|---|
| ENSG00000116478.12 | HDAC1 | protein_coding | 25352,77411 | -0,455255495 | 0,043709641 | -10,41544795 | 2,11E-25 | 5,56E-24 |
| ENSG00000116514.16 | RNF19B | protein_coding | 3441,408781 | 0,811005422 | 0,062647268 | 12,94558322 | 2,49E-38 | 1,06E-36 |
| ENSG00000116525.14 | TRIM62 | protein_coding | 895,3384767 | 0,343627212 | 0,091090222 | 3,772383097 | 0,000161696 | 0,000738257 |
| ENSG00000116539.13 | ASH1L | protein_coding | 16079,40037 | 0,12396457 | 0,046828067 | 2,647228015 | 0,008115461 | 0,024985764 |
| ENSG00000116560.11 | SFPQ | protein_coding | 41452,18167 | -0,152315063 | 0,042476018 | -3,585907301 | 0,000335908 | 0,001444861 |
| ENSG00000116574.6 | RHOU | protein_coding | 6425,953165 | 0,124914015 | 0,05226811 | 2,38987053 | 0,016854313 | 0,046923158 |
| ENSG00000116580.18 | GON4L | protein_coding | 7000,613932 | 0,245720017 | 0,061703205 | 3,98228937 | 6,83E-05 | 0,000334642 |
| ENSG00000116584.18 | ARHGEF2 | protein_coding | 912,026077 | 0,617766944 | 0,244607325 | 2,525545562 | 0,01155188 | 0,033808513 |
| ENSG00000116586.11 | LAMTOR2 | protein_coding | 3098,418273 | -0,164732344 | 0,061798579 | -2,665633193 | 0,007684349 | 0,023821375 |
| ENSG00000116663.11 | FBXO6 | protein_coding | 914,8837769 | 3,844120175 | 0,107719356 | 35,68643852 | 6,42E-279 | 1,68E-276 |
| ENSG00000116678.19 | LEPR | protein_coding | 834,1626534 | -0,551368548 | 0,109190525 | -5,049600686 | 4,43E-07 | 3,03E-06 |
| ENSG00000116698.21 | SMG7 | protein_coding | 16359,52357 | 0,618278213 | 0,047610258 | 12,98623954 | 1,46E-38 | 6,28E-37 |
| ENSG00000116704.8 | SLC35D1 | protein_coding | 3679,584206 | -0,149217555 | 0,056932294 | -2,620965068 | 0,008768125 | 0,026746404 |
| ENSG00000116717.13 | GADD45A | protein_coding | 2669,478394 | 0,312788001 | 0,062938844 | 4,969713111 | 6,71E-07 | 4,47E-06 |
| ENSG00000116741.8 | RGS2 | protein_coding | 1933,767176 | 0,235308871 | 0,063737804 | 3,691825832 | 0,00022265 | 0,000990832 |
| ENSG00000116747.12 | RO60 | protein_coding | 5068,12142 | 0,233901383 | 0,049504922 | 4,724810609 | 2,30E-06 | 1,42E-05 |
| ENSG00000116754.13 | SRSF11 | protein_coding | 27972,1965 | -0,220182153 | 0,050348244 | -4,373184381 | 1,22E-05 | 6,77E-05 |
| ENSG00000116761.11 | CTH | protein_coding | 386,2698797 | 1,518070114 | 0,144181082 | 10,52891329 | 6,36E-26 | 1,72E-24 |
| ENSG00000116771.6 | AGMAT | protein_coding | 1714,533892 | -0,452382432 | 0,067645619 | -6,687534849 | 2,27E-11 | 2,51E-10 |
| ENSG00000116774.12 | OLFML3 | protein_coding | 107,7394574 | 1,615085668 | 0,262370609 | 6,155741571 | 7,47E-10 | 7,09E-09 |
| ENSG00000116785.13 | CFHR3 | protein_coding | 159,3436818 | -0,509104143 | 0,211884835 | -2,402739886 | 0,016272758 | 0,045506709 |
| ENSG00000116793.16 | PHTF1 | protein_coding | 8021,313545 | 0,234404206 | 0,04866327 | 4,816860967 | 1,46E-06 | 9,24E-06 |
| ENSG00000116815.16 | CD58 | protein_coding | 436,1482856 | 0,440023324 | 0,125875988 | 3,495689143 | 0,000472839 | 0,001971682 |
| ENSG00000116833.14 | NR5A2 | protein_coding | 117,8247383 | 0,847552125 | 0,239087176 | 3,544950168 | 0,000392687 | 0,001663968 |
| ENSG00000116852.14 | KIF21B | protein_coding | 651,5928433 | -0,624974465 | 0,100263651 | -6,233310468 | 4,57E-10 | 4,46E-09 |
| ENSG00000116857.17 | TMEM9 | protein_coding | 10333,89046 | 0,221964855 | 0,04590299 | 4,835520608 | 1,33E-06 | 8,48E-06 |
| ENSG00000116863.11 | ADPRHL2 | protein_coding | 2926,681163 | 0,315509061 | 0,063474029 | 4,970679628 | 6,67E-07 | 4,45E-06 |
| ENSG00000116871.15 | MAP7D1 | protein_coding | 5183,677053 | 0,160003477 | 0,050623518 | 3,160655046 | 0,001574148 | 0,00584309 |
| ENSG00000116874.11 | WARS2 | protein_coding | 1829,455141 | -0,244146502 | 0,074133659 | -3,29332863 | 0,000990087 | 0,003839604 |
| ENSG00000116883.8 | AL591845.1 | lncRNA | 115,6045334 | -0,559180271 | 0,224658959 | -2,489018349 | 0,012809635 | 0,037017856 |
| ENSG00000116903.7 | EXOC8 | protein_coding | 3549,335513 | 0,185831878 | 0,053856612 | 3,450493307 | 0,000559563 | 0,002296719 |
| ENSG00000116906.13 | GNPAT | protein_coding | 8797,744581 | -0,237448757 | 0,048041468 | -4,942579078 | 7,71E-07 | 5,09E-06 |
| ENSG00000116954.8 | RRAGC | protein_coding | 5581,97797 | 0,533437306 | 0,050901797 | 10,47973428 | 1,07E-25 | 2,86E-24 |
| ENSG00000116977.18 | LGALS8 | protein_coding | 5200,938219 | 0,749059863 | 0,058521239 | 12,79979498 | 1,64E-37 | 6,79E-36 |
| ENSG00000116984.13 | MTR | protein_coding | 5966,396596 | -0,256685285 | 0,065703217 | -3,906738439 | 9,36E-05 | 0,000447645 |
| ENSG00000116991.10 | SIPA1L2 | protein_coding | 5164,962287 | 0,192371854 | 0,055844661 | 3,444767166 | 0,000571551 | 0,002341655 |

| | | | | | | | | |
|---|---|---|---|---|---|---|---|---|
| ENSG00000117000.9 | RLF | protein_coding | 5855,243626 | 0,308536016 | 0,052187726 | 5,912041748 | 3,38E-09 | 2,98E-08 |
| ENSG00000117009.12 | KMO | protein_coding | 117,1126633 | 2,441803319 | 0,690475524 | 3,536408223 | 0,000405607 | 0,001713876 |
| ENSG00000117010.17 | ZNF684 | protein_coding | 398,1632236 | 0,372532963 | 0,138624926 | 2,687344724 | 0,007202258 | 0,022503557 |
| ENSG00000117020.17 | AKT3 | protein_coding | 3956,459859 | -0,370048818 | 0,077291034 | -4,787732822 | 1,69E-06 | 1,06E-05 |
| ENSG00000117036.12 | ETV3 | protein_coding | 3694,486565 | 0,220464347 | 0,055958041 | 3,93981535 | 8,15E-05 | 0,000394558 |
| ENSG00000117069.15 | ST6GALNAC5 | protein_coding | 5815,566494 | -0,305005317 | 0,054571749 | -5,5890698 | 2,28E-08 | 1,83E-07 |
| ENSG00000117115.13 | PADI2 | protein_coding | 287,9275697 | -1,598402426 | 0,160454441 | -9,961721316 | 2,24E-23 | 5,38E-22 |
| ENSG00000117122.13 | MFAP2 | protein_coding | 1212,323001 | -0,482750854 | 0,074903647 | -6,44495794 | 1,16E-10 | 1,20E-09 |
| ENSG00000117139.17 | KDM5B | protein_coding | 14562,74535 | 0,153559312 | 0,048286449 | 3,180174074 | 0,001471866 | 0,005503361 |
| ENSG00000117143.13 | UAP1 | protein_coding | 10545,78739 | 0,21264611 | 0,051496237 | 4,12935244 | 3,64E-05 | 0,000186186 |
| ENSG00000117151.13 | CTBS | protein_coding | 5216,557893 | 0,249589038 | 0,050251248 | 4,966822693 | 6,81E-07 | 4,53E-06 |
| ENSG00000117152.13 | RGS4 | protein_coding | 14887,65297 | 0,610931356 | 0,056755983 | 10,76417545 | 5,08E-27 | 1,42E-25 |
| ENSG00000117153.16 | KLHL12 | protein_coding | 7356,923201 | 0,236897421 | 0,048771522 | 4,857289856 | 1,19E-06 | 7,65E-06 |
| ENSG00000117155.16 | SSX2IP | protein_coding | 29192,37852 | 0,182544755 | 0,047165352 | 3,870314698 | 0,000108695 | 0,000513671 |
| ENSG00000117226.12 | GBP3 | protein_coding | 5528,507508 | 6,827085931 | 0,09201313 | 74,19686626 | 0 | 0 |
| ENSG00000117228.10 | GBP1 | protein_coding | 5247,845763 | 8,895083963 | 0,36493968 | 24,37412114 | 3,22E-131 | 4,86E-129 |
| ENSG00000117242.7 | PINK1-AS | lncRNA | 509,325935 | -0,644067889 | 0,1122455 | -5,738028575 | 9,58E-09 | 7,98E-08 |
| ENSG00000117262.19 | GPR89A | protein_coding | 3826,136485 | 0,299821604 | 0,059928161 | 5,003016883 | 5,64E-07 | 3,80E-06 |
| ENSG00000117266.15 | CDK18 | protein_coding | 229,5508565 | 1,647288749 | 0,180872026 | 9,107482156 | 8,43E-20 | 1,69E-18 |
| ENSG00000117298.16 | ECE1 | protein_coding | 21227,61217 | 0,572046806 | 0,043076497 | 13,27978937 | 3,03E-40 | 1,37E-38 |
| ENSG00000117305.15 | HMGCL | protein_coding | 4662,030599 | 0,146655621 | 0,056393912 | 2,600557685 | 0,009307237 | 0,028169504 |
| ENSG00000117318.9 | ID3 | protein_coding | 2861,239574 | 0,792309221 | 0,078055408 | 10,15059989 | 3,29E-24 | 8,25E-23 |
| ENSG00000117322.17 | CR2 | protein_coding | 79,65372783 | 2,424776683 | 0,31879717 | 7,606016954 | 2,83E-14 | 4,02E-13 |
| ENSG00000117335.19 | CD46 | protein_coding | 61620,11539 | -0,118541185 | 0,043503944 | -2,724837683 | 0,006433311 | 0,020381054 |
| ENSG00000117360.13 | PRPF3 | protein_coding | 6584,171959 | 0,222218132 | 0,048863763 | 4,54770808 | 5,42E-06 | 3,16E-05 |
| ENSG00000117362.13 | APH1A | protein_coding | 12859,15668 | 0,149630511 | 0,045061538 | 3,320581533 | 0,000898301 | 0,003515886 |
| ENSG00000117395.12 | EBNA1BP2 | protein_coding | 4025,18319 | -0,169233483 | 0,058798745 | -2,878181885 | 0,003999745 | 0,013398166 |
| ENSG00000117399.14 | CDC20 | protein_coding | 3542,528146 | -1,013597172 | 0,071599477 | -14,15648841 | 1,70E-45 | 8,87E-44 |
| ENSG00000117408.11 | IPO13 | protein_coding | 4556,269212 | 0,144914546 | 0,056665237 | 2,557380056 | 0,010546392 | 0,031317776 |
| ENSG00000117419.15 | ERI3 | protein_coding | 5717,101042 | -0,240611618 | 0,059541385 | -4,041081958 | 5,32E-05 | 0,000265592 |
| ENSG00000117425.14 | PTCH2 | protein_coding | 238,1825835 | -0,549524533 | 0,157031588 | -3,499452176 | 0,000466215 | 0,001947307 |
| ENSG00000117450.14 | PRDX1 | protein_coding | 14135,62846 | -0,186592085 | 0,049275114 | -3,786740839 | 0,000152636 | 0,000702307 |
| ENSG00000117461.15 | PIK3R3 | protein_coding | 13293,04655 | 0,276837695 | 0,053704766 | 5,154806829 | 2,54E-07 | 1,79E-06 |
| ENSG00000117475.14 | BLZF1 | protein_coding | 4121,805099 | 1,104560341 | 0,065703264 | 16,81134667 | 2,02E-63 | 1,47E-61 |
| ENSG00000117481.10 | NSUN4 | protein_coding | 2758,124067 | 0,398585935 | 0,057477588 | 6,934632232 | 4,07E-12 | 4,87E-11 |
| ENSG00000117523.16 | PRRC2C | protein_coding | 35221,02105 | 0,26617828 | 0,03890386 | 6,841950462 | 7,81E-12 | 9,06E-11 |

| | | | | | | | | |
|---|---|---|---|---|---|---|---|---|
| ENSG00000117525.14 | F3 | protein_coding | 2274,422169 | 0,639731565 | 0,063885783 | 10,01367648 | 1,33E-23 | 3,23E-22 |
| ENSG00000117528.13 | ABCD3 | protein_coding | 27284,90895 | -0,217237429 | 0,042487864 | -5,112928896 | 3,17E-07 | 2,21E-06 |
| ENSG00000117543.21 | DPH5 | protein_coding | 2885,422882 | -0,238824798 | 0,063300495 | -3,772874076 | 0,000161378 | 0,000737105 |
| ENSG00000117569.18 | PTBP2 | protein_coding | 3956,130661 | -0,224411466 | 0,052569684 | -4,268837987 | 1,96E-05 | 0,000104957 |
| ENSG00000117592.9 | PRDX6 | protein_coding | 7459,685815 | -0,179448189 | 0,057695562 | -3,11025984 | 0,001869228 | 0,006808165 |
| ENSG00000117593.12 | DARS2 | protein_coding | 7034,24428 | -0,642027203 | 0,055280631 | -11,61396292 | 3,50E-31 | 1,17E-29 |
| ENSG00000117595.12 | IRF6 | protein_coding | 9641,447322 | 0,213584506 | 0,044913755 | 4,755436453 | 1,98E-06 | 1,23E-05 |
| ENSG00000117597.18 | UTP25 | protein_coding | 2227,290347 | 0,157726851 | 0,059867129 | 2,634615235 | 0,008423271 | 0,025802858 |
| ENSG00000117600.12 | PLPPR4 | protein_coding | 666,6200269 | -0,376139141 | 0,102474691 | -3,670556484 | 0,000242023 | 0,001069434 |
| ENSG00000117602.12 | RCAN3 | protein_coding | 20350,53979 | -0,289288414 | 0,041752131 | -6,928710074 | 4,25E-12 | 5,07E-11 |
| ENSG00000117625.13 | RCOR3 | protein_coding | 5629,82234 | 0,188436057 | 0,04693541 | 4,014795178 | 5,95E-05 | 0,000294465 |
| ENSG00000117632.23 | STMN1 | protein_coding | 96387,46963 | -0,344337399 | 0,042547639 | -8,092984799 | 5,82E-16 | 9,33E-15 |
| ENSG00000117650.13 | NEK2 | protein_coding | 3993,318895 | -0,823201994 | 0,059138384 | -13,91992705 | 4,79E-44 | 2,40E-42 |
| ENSG00000117682.17 | DHDDS | protein_coding | 2725,80356 | 0,259126519 | 0,060144545 | 4,308396011 | 1,64E-05 | 8,90E-05 |
| ENSG00000117697.15 | NSL1 | protein_coding | 7704,663375 | -0,173056808 | 0,051357304 | -3,369663052 | 0,000752602 | 0,002999346 |
| ENSG00000117707.15 | PROX1 | protein_coding | 12901,89384 | 0,483129908 | 0,046149359 | 10,46883236 | 1,20E-25 | 3,20E-24 |
| ENSG00000117724.13 | CENPF | protein_coding | 22117,19631 | -0,739343277 | 0,06391966 | -11,56675855 | 6,07E-31 | 2,02E-29 |
| ENSG00000117748.10 | RPA2 | protein_coding | 11337,49527 | -0,166549114 | 0,043593847 | -3,820472985 | 0,000133196 | 0,000619316 |
| ENSG00000117758.14 | STX12 | protein_coding | 6369,495893 | 0,285489541 | 0,046606766 | 6,125495605 | 9,04E-10 | 8,48E-09 |
| ENSG00000117791.16 | mars-02 | protein_coding | 1066,554682 | 0,270010588 | 0,090643304 | 2,978825531 | 0,002893555 | 0,010033245 |
| ENSG00000117868.16 | ESYT2 | protein_coding | 11162,50294 | -0,346963415 | 0,050814307 | -6,828065494 | 8,61E-12 | 9,94E-11 |
| ENSG00000117899.11 | MESD | protein_coding | 14085,68553 | 0,288476073 | 0,050085113 | 5,759716879 | 8,43E-09 | 7,06E-08 |
| ENSG00000117906.14 | RCN2 | protein_coding | 11413,04247 | -0,122959562 | 0,043368194 | -2,835247462 | 0,00457902 | 0,015088445 |
| ENSG00000117971.12 | CHRNB4 | protein_coding | 699,7552597 | -1,085837792 | 0,113502872 | -9,566610683 | 1,10E-21 | 2,45E-20 |
| ENSG00000117984.14 | CTSD | protein_coding | 19506,65507 | 0,243033904 | 0,056040549 | 4,336750985 | 1,45E-05 | 7,89E-05 |
| ENSG00000118007.13 | STAG1 | protein_coding | 8773,726819 | -0,324270167 | 0,050747389 | -6,389888639 | 1,66E-10 | 1,69E-09 |
| ENSG00000118058.21 | KMT2A | protein_coding | 14564,54854 | 0,285244713 | 0,05669071 | 5,031595317 | 4,86E-07 | 3,30E-06 |
| ENSG00000118137.9 | APOA1 | protein_coding | 2521,924021 | -0,204648659 | 0,079859989 | -2,56259314 | 0,010389372 | 0,030900419 |
| ENSG00000118160.14 | SLC8A2 | protein_coding | 467,7852909 | 0,484972276 | 0,122704156 | 3,952370426 | 7,74E-05 | 0,000375623 |
| ENSG00000118181.11 | RPS25 | protein_coding | 33227,58185 | -0,534868579 | 0,043092548 | -12,41208989 | 2,25E-35 | 8,75E-34 |
| ENSG00000118193.12 | KIF14 | protein_coding | 4517,72141 | -0,917875212 | 0,056919394 | -16,12587804 | 1,68E-58 | 1,14E-56 |
| ENSG00000118200.14 | CAMSAP2 | protein_coding | 8427,895382 | 0,314603192 | 0,050868308 | 6,184660005 | 6,22E-10 | 5,97E-09 |
| ENSG00000118217.6 | ATF6 | protein_coding | 8815,568783 | -0,27972291 | 0,049564247 | -5,6436429 | 1,66E-08 | 1,35E-07 |
| ENSG00000118231.5 | CRYGD | protein_coding | 342,7004239 | -0,604490735 | 0,167096581 | -3,617612825 | 0,000297333 | 0,001292519 |
| ENSG00000118263.15 | KLF7 | protein_coding | 4795,72126 | -0,241961343 | 0,053396562 | -4,53140306 | 5,86E-06 | 3,40E-05 |
| ENSG00000118271.10 | TTR | protein_coding | 197661,5213 | 0,253619763 | 0,044708859 | 5,672695971 | 1,41E-08 | 1,15E-07 |

| | | | | | | | | |
|---|---|---|---|---|---|---|---|---|
| ENSG00000118322.14 | ATP10B | protein_coding | 36,47749724 | 1,740418115 | 0,456076039 | 3,816070055 | 0,000135594 | 0,000629275 |
| ENSG00000118369.13 | USP35 | protein_coding | 863,0390698 | 0,314919445 | 0,085067567 | 3,701991921 | 0,000213913 | 0,000954028 |
| ENSG00000118402.6 | ELOVL4 | protein_coding | 10857,83076 | 0,285837809 | 0,0431654 | 6,621919662 | 3,55E-11 | 3,85E-10 |
| ENSG00000118418.14 | HMGN3 | protein_coding | 8918,194524 | -0,274077659 | 0,050757382 | -5,399759587 | 6,67E-08 | 5,03E-07 |
| ENSG00000118454.13 | ANKRD13C | protein_coding | 5359,408658 | 0,148075652 | 0,051882071 | 2,854081344 | 0,004316148 | 0,014329229 |
| ENSG00000118495.19 | PLAGL1 | protein_coding | 90399,25835 | -0,488639603 | 0,041584021 | -11,75065787 | 7,01E-32 | 2,44E-30 |
| ENSG00000118503.15 | TNFAIP3 | protein_coding | 1730,909109 | 1,346287163 | 0,095313588 | 14,12481879 | 2,67E-45 | 1,39E-43 |
| ENSG00000118507.17 | AKAP7 | protein_coding | 486,9991997 | 2,015281753 | 0,123970914 | 16,25608535 | 2,02E-59 | 1,41E-57 |
| ENSG00000118508.5 | RAB32 | protein_coding | 2332,964113 | 0,211048968 | 0,062479563 | 3,37788803 | 0,000730448 | 0,002921988 |
| ENSG00000118513.19 | MYB | protein_coding | 748,2153673 | -0,484874262 | 0,106921277 | -4,53487161 | 5,76E-06 | 3,34E-05 |
| ENSG00000118523.6 | CCN2 | protein_coding | 201,4498387 | 1,267746028 | 0,176957225 | 7,164138265 | 7,83E-13 | 9,98E-12 |
| ENSG00000118596.12 | SLC16A7 | protein_coding | 1804,795682 | -0,465719225 | 0,08510514 | -5,47228085 | 4,44E-08 | 3,42E-07 |
| ENSG00000118600.11 | RXYLT1 | protein_coding | 5045,205471 | -0,253239888 | 0,055468579 | -4,565465601 | 4,98E-06 | 2,92E-05 |
| ENSG00000118620.13 | ZNF430 | protein_coding | 3398,106845 | 0,514101528 | 0,06018478 | 8,542052162 | 1,32E-17 | 2,34E-16 |
| ENSG00000118689.15 | FOXO3 | protein_coding | 6523,927097 | 0,151403934 | 0,050328755 | 3,008298786 | 0,002627147 | 0,009212409 |
| ENSG00000118690.13 | ARMC2 | protein_coding | 506,5993506 | -0,358119338 | 0,13232559 | -2,706349828 | 0,006802734 | 0,021421761 |
| ENSG00000118705.17 | RPN2 | protein_coding | 88108,06343 | 0,196562489 | 0,039093443 | 5,028016804 | 4,96E-07 | 3,36E-06 |
| ENSG00000118707.10 | TGIF2 | protein_coding | 2437,042555 | 0,849939335 | 0,084435738 | 10,06610893 | 7,80E-24 | 1,92E-22 |
| ENSG00000118762.8 | PKD2 | protein_coding | 13522,28131 | 0,260570895 | 0,04320512 | 6,031018886 | 1,63E-09 | 1,49E-08 |
| ENSG00000118777.12 | ABCG2 | protein_coding | 2759,573814 | -0,152525072 | 0,059314759 | -2,571452289 | 0,010127297 | 0,030241002 |
| ENSG00000118804.8 | STBD1 | protein_coding | 1271,224972 | 0,607431695 | 0,089968568 | 6,751599023 | 1,46E-11 | 1,65E-10 |
| ENSG00000118816.10 | CCNI | protein_coding | 33172,96136 | -0,158651939 | 0,045617593 | -3,477867403 | 0,00050542 | 0,002092224 |
| ENSG00000118849.10 | RARRES1 | protein_coding | 1726,606542 | 0,263765498 | 0,06987117 | 3,775026217 | 0,000159991 | 0,000731661 |
| ENSG00000118855.19 | MFSD1 | protein_coding | 4964,491659 | -0,143813314 | 0,052600532 | -2,734065779 | 0,006255755 | 0,019894148 |
| ENSG00000118894.14 | EEF2KMT | protein_coding | 1248,902178 | 0,212474705 | 0,085006837 | 2,499501372 | 0,012436822 | 0,036116684 |
| ENSG00000118900.14 | UBN1 | protein_coding | 14926,3387 | 0,261959927 | 0,043899848 | 5,967217131 | 2,41E-09 | 2,16E-08 |
| ENSG00000118922.17 | KLF12 | protein_coding | 5680,886681 | -0,42553988 | 0,052701043 | -8,074600718 | 6,77E-16 | 1,08E-14 |
| ENSG00000118946.12 | PCDH17 | protein_coding | 5419,541806 | 1,435825047 | 0,069238628 | 20,73734124 | 1,60E-95 | 1,80E-93 |
| ENSG00000118961.15 | LDAH | protein_coding | 8321,592786 | -0,240561933 | 0,065947476 | -3,647780742 | 0,000264515 | 0,001160619 |
| ENSG00000118965.14 | WDR35 | protein_coding | 4637,554099 | -0,345152844 | 0,060541926 | -5,701054901 | 1,19E-08 | 9,81E-08 |
| ENSG00000118985.16 | ELL2 | protein_coding | 25069,73438 | 0,230426382 | 0,039633307 | 5,813957934 | 6,10E-09 | 5,21E-08 |
| ENSG00000118997.14 | DNAH7 | protein_coding | 340,0571345 | 0,860944209 | 0,169349964 | 5,083816894 | 3,70E-07 | 2,55E-06 |
| ENSG00000119013.9 | NDUFB3 | protein_coding | 7902,166409 | -0,191865567 | 0,055136859 | -3,479805916 | 0,000501777 | 0,002079189 |
| ENSG00000119042.17 | SATB2 | protein_coding | 533,7217219 | 0,319291557 | 0,107615696 | 2,966960847 | 0,003007593 | 0,010383114 |
| ENSG00000119121.22 | TRPM6 | protein_coding | 173,5582817 | 0,914770435 | 0,223019188 | 4,101756646 | 4,10E-05 | 0,000208477 |
| ENSG00000119139.20 | TJP2 | protein_coding | 25393,2064 | 0,183911876 | 0,039792299 | 4,621795718 | 3,80E-06 | 2,27E-05 |

| | | | | | | | | |
|---|---|---|---|---|---|---|---|---|
| ENSG00000119185.12 | ITGB1BP1 | protein_coding | 7314,108178 | -0,426099702 | 0,064480285 | -6,608216739 | 3,89E-11 | 4,22E-10 |
| ENSG00000119203.14 | CPSF3 | protein_coding | 12035,29454 | -0,154792386 | 0,044402242 | -3,486138978 | 0,000490046 | 0,002034764 |
| ENSG00000119227.8 | PIGZ | protein_coding | 220,8161497 | -0,539313588 | 0,202836475 | -2,658859005 | 0,007840577 | 0,024235587 |
| ENSG00000119283.15 | TRIM67 | protein_coding | 57,69139627 | 0,856883512 | 0,309606286 | 2,767655405 | 0,005646112 | 0,018176249 |
| ENSG00000119318.13 | RAD23B | protein_coding | 50228,39473 | -0,132359875 | 0,047423853 | -2,790997888 | 0,005254582 | 0,017047474 |
| ENSG00000119321.9 | FKBP15 | protein_coding | 4080,806187 | 0,44924585 | 0,168831473 | 2,660912936 | 0,00779291 | 0,024108572 |
| ENSG00000119326.15 | CTNNAL1 | protein_coding | 12672,62367 | -0,137487035 | 0,047932381 | -2,868353981 | 0,004126136 | 0,013781748 |
| ENSG00000119333.11 | WDR34 | protein_coding | 6203,416104 | -0,509542267 | 0,059772679 | -8,524668379 | 1,53E-17 | 2,71E-16 |
| ENSG00000119335.16 | SET | protein_coding | 55252,71521 | -0,241515356 | 0,03897859 | -6,196102894 | 5,79E-10 | 5,57E-09 |
| ENSG00000119396.11 | RAB14 | protein_coding | 21542,57867 | -0,131938136 | 0,044914385 | -2,937547392 | 0,003308196 | 0,011320186 |
| ENSG00000119397.16 | CNTRL | protein_coding | 4488,39528 | -0,21755743 | 0,074527546 | -2,919154625 | 0,003509821 | 0,011933921 |
| ENSG00000119402.17 | FBXW2 | protein_coding | 14875,87063 | -0,182272754 | 0,066155338 | -2,755223704 | 0,005865202 | 0,018774151 |
| ENSG00000119403.15 | PHF19 | protein_coding | 5531,245147 | -0,601417171 | 0,053078606 | -11,33068897 | 9,25E-30 | 2,87E-28 |
| ENSG00000119431.9 | HDHD3 | protein_coding | 1696,643884 | 0,218629808 | 0,07022037 | 3,113481293 | 0,001848942 | 0,006739732 |
| ENSG00000119446.14 | RBM18 | protein_coding | 4618,828668 | 0,227571319 | 0,052002891 | 4,376128202 | 1,21E-05 | 6,69E-05 |
| ENSG00000119471.15 | HSDL2 | protein_coding | 7910,266476 | -0,563820564 | 0,054218892 | -10,39896879 | 2,51E-25 | 6,59E-24 |
| ENSG00000119487.17 | MAPKAP1 | protein_coding | 4538,880249 | -0,298089803 | 0,055479187 | -5,37300232 | 7,74E-08 | 5,78E-07 |
| ENSG00000119509.13 | INVS | protein_coding | 3411,380832 | 0,196534686 | 0,063667741 | 3,086880155 | 0,002022691 | 0,00729563 |
| ENSG00000119537.18 | KDSR | protein_coding | 3657,702733 | 0,435428305 | 0,065570054 | 6,640657998 | 3,12E-11 | 3,41E-10 |
| ENSG00000119599.16 | DCAF4 | protein_coding | 730,9705215 | 0,466670113 | 0,120518022 | 3,872201899 | 0,000107857 | 0,00051003 |
| ENSG00000119616.11 | FCF1 | protein_coding | 7048,007464 | -0,181496601 | 0,049646664 | -3,65576628 | 0,000256415 | 0,00112774 |
| ENSG00000119640.9 | ACYP1 | protein_coding | 3355,4054 | -0,145767962 | 0,05477412 | -2,661256121 | 0,007784971 | 0,024093486 |
| ENSG00000119655.10 | NPC2 | protein_coding | 13878,83819 | 0,109572134 | 0,044426662 | 2,466359839 | 0,013649414 | 0,039153135 |
| ENSG00000119673.14 | ACOT2 | protein_coding | 1016,460295 | 0,539445656 | 0,088239353 | 6,11343623 | 9,75E-10 | 9,11E-09 |
| ENSG00000119681.12 | LTBP2 | protein_coding | 126,6383584 | -0,67955707 | 0,241724119 | -2,811291951 | 0,004934299 | 0,01613161 |
| ENSG00000119684.15 | MLH3 | protein_coding | 4233,92632 | 0,394881649 | 0,053544804 | 7,374789368 | 1,65E-13 | 2,21E-12 |
| ENSG00000119688.21 | ABCD4 | protein_coding | 4763,985657 | 0,219361375 | 0,056999018 | 3,84851148 | 0,000118838 | 0,000558788 |
| ENSG00000119698.12 | PPP4R4 | protein_coding | 2120,712353 | 0,223551612 | 0,066916099 | 3,340774742 | 0,00083545 | 0,003296717 |
| ENSG00000119703.14 | ZC2HC1C | protein_coding | 1004,210363 | 0,596309013 | 0,087790722 | 6,792392167 | 1,10E-11 | 1,26E-10 |
| ENSG00000119705.9 | SLIRP | protein_coding | 4527,974527 | -0,150090447 | 0,056841811 | -2,640493744 | 0,008278532 | 0,025425226 |
| ENSG00000119711.12 | ALDH6A1 | protein_coding | 1731,100681 | -0,225071918 | 0,078656633 | -2,861448688 | 0,004217098 | 0,014039735 |
| ENSG00000119718.11 | EIF2B2 | protein_coding | 3492,863742 | -0,150108852 | 0,055608218 | -2,699400493 | 0,006946453 | 0,021828522 |
| ENSG00000119725.18 | ZNF410 | protein_coding | 7703,29936 | 0,232555723 | 0,049562049 | 4,692213682 | 2,70E-06 | 1,65E-05 |
| ENSG00000119729.12 | RHOQ | protein_coding | 1757,301586 | 0,228239723 | 0,076168498 | 2,996510746 | 0,002730886 | 0,009527203 |
| ENSG00000119772.16 | DNMT3A | protein_coding | 8882,611939 | -0,200009 | 0,056684614 | -3,528453064 | 0,000417996 | 0,001762584 |
| ENSG00000119778.15 | ATAD2B | protein_coding | 5420,589922 | 0,385016846 | 0,0512101 | 7,518377153 | 5,55E-14 | 7,77E-13 |

| | | | | | | | | |
|---|---|---|---|---|---|---|---|---|
| ENSG00000119787.14 | ATL2 | protein_coding | 6977,880115 | -0,123367593 | 0,049497817 | -2,492384539 | 0,012688858 | 0,036730344 |
| ENSG00000119801.13 | YPEL5 | protein_coding | 10231,1675 | 0,195128686 | 0,04307928 | 4,52952525 | 5,91E-06 | 3,42E-05 |
| ENSG00000119812.19 | FAM98A | protein_coding | 6524,449004 | 0,317084724 | 0,067644631 | 4,687507645 | 2,77E-06 | 1,69E-05 |
| ENSG00000119820.11 | YIPF4 | protein_coding | 9619,059282 | -0,210283222 | 0,067907565 | -3,096609676 | 0,001957474 | 0,007088287 |
| ENSG00000119844.15 | AFTPH | protein_coding | 10208,36468 | 0,203170596 | 0,043529586 | 4,667413896 | 3,05E-06 | 1,85E-05 |
| ENSG00000119862.13 | LGALSL | protein_coding | 4804,421042 | 0,203409921 | 0,055908292 | 3,638278187 | 0,000274467 | 0,001199607 |
| ENSG00000119878.5 | CRIPT | protein_coding | 2721,223522 | -0,213133781 | 0,05919725 | -3,600400012 | 0,000317728 | 0,001373749 |
| ENSG00000119888.10 | EPCAM | protein_coding | 74121,27835 | 0,340954311 | 0,040010181 | 8,521688838 | 1,57E-17 | 2,78E-16 |
| ENSG00000119900.9 | OGFRL1 | protein_coding | 2902,956639 | -0,465880421 | 0,060004917 | -7,764037399 | 8,23E-15 | 1,22E-13 |
| ENSG00000119912.16 | IDE | protein_coding | 4604,665645 | 0,140119624 | 0,056078363 | 2,498639706 | 0,012467099 | 0,03618127 |
| ENSG00000119915.5 | ELOVL3 | protein_coding | 70,47785028 | 0,979085099 | 0,281806054 | 3,47432245 | 0,000512145 | 0,002115595 |
| ENSG00000119917.14 | IFIT3 | protein_coding | 60248,19773 | 11,33818476 | 0,214862753 | 52,76942883 | 0 | 0 |
| ENSG00000119922.10 | IFIT2 | protein_coding | 15405,56994 | 10,05426499 | 0,130368328 | 77,1219906 | 0 | 0 |
| ENSG00000119927.14 | GPAM | protein_coding | 1954,353375 | -0,784572987 | 0,069798436 | -11,24055253 | 2,58E-29 | 7,87E-28 |
| ENSG00000119946.11 | CNNM1 | protein_coding | 5649,810048 | 0,837880289 | 0,049684335 | 16,86407367 | 8,27E-64 | 6,08E-62 |
| ENSG00000119950.21 | MXI1 | protein_coding | 715,5760456 | 0,284805273 | 0,093855428 | 3,034510405 | 0,002409264 | 0,008518714 |
| ENSG00000119953.12 | SMNDC1 | protein_coding | 2675,989551 | 0,181183738 | 0,058479084 | 3,098265664 | 0,001946568 | 0,007052199 |
| ENSG00000120053.12 | GOT1 | protein_coding | 5330,89714 | -0,233945977 | 0,055389157 | -4,223678202 | 2,40E-05 | 0,000126905 |
| ENSG00000120071.14 | KANSL1 | protein_coding | 7073,240003 | 0,380458305 | 0,056276697 | 6,760494616 | 1,38E-11 | 1,56E-10 |
| ENSG00000120129.6 | DUSP1 | protein_coding | 919,8454289 | 0,303687133 | 0,08396377 | 3,616883024 | 0,000298172 | 0,001295667 |
| ENSG00000120137.7 | PANK3 | protein_coding | 26127,63852 | -0,403676771 | 0,058892265 | -6,854495582 | 7,16E-12 | 8,33E-11 |
| ENSG00000120217.14 | CD274 | protein_coding | 1909,217756 | 3,837432942 | 0,114184601 | 33,60727202 | 1,31E-247 | 3,18E-245 |
| ENSG00000120251.20 | GRIA2 | protein_coding | 1089,09194 | 0,378073587 | 0,096397463 | 3,922028382 | 8,78E-05 | 0,000422406 |
| ENSG00000120253.14 | NUP43 | protein_coding | 7403,838642 | -0,397039691 | 0,05131391 | -7,737467072 | 1,01E-14 | 1,49E-13 |
| ENSG00000120278.16 | PLEKHG1 | protein_coding | 316,3473383 | 1,43080793 | 0,150642377 | 9,498044054 | 2,14E-21 | 4,66E-20 |
| ENSG00000120306.11 | CYSTM1 | protein_coding | 12954,02436 | 0,394703923 | 0,057882503 | 6,819054204 | 9,16E-12 | 1,05E-10 |
| ENSG00000120334.15 | CENPL | protein_coding | 1576,06568 | -0,378433202 | 0,086647906 | -4,367482352 | 1,26E-05 | 6,93E-05 |
| ENSG00000120341.18 | SEC16B | protein_coding | 1366,709172 | 3,648423081 | 0,11689879 | 31,21010145 | 7,77E-214 | 1,65E-211 |
| ENSG00000120438.12 | TCP1 | protein_coding | 26116,24287 | -0,346800746 | 0,04679923 | -7,410394231 | 1,26E-13 | 1,72E-12 |
| ENSG00000120458.11 | MSANTD2 | protein_coding | 1327,324038 | -0,27104984 | 0,094342979 | -2,873026084 | 0,004065605 | 0,01360274 |
| ENSG00000120509.10 | PDZD11 | protein_coding | 4364,473206 | -0,158569306 | 0,055518321 | -2,856161775 | 0,004287966 | 0,014254589 |
| ENSG00000120519.14 | SLC10A7 | protein_coding | 1277,156898 | 0,258815804 | 0,075455957 | 3,430024824 | 0,000603526 | 0,002457847 |
| ENSG00000120539.14 | MASTL | protein_coding | 3227,117047 | 1,333502287 | 0,064367394 | 20,7170463 | 2,43E-95 | 2,72E-93 |
| ENSG00000120549.18 | KIAA1217 | protein_coding | 1703,05881 | 0,564813604 | 0,113708353 | 4,967212968 | 6,79E-07 | 4,52E-06 |
| ENSG00000120594.17 | PLXDC2 | protein_coding | 2813,755696 | -0,280289167 | 0,077808046 | -3,602315988 | 0,000315395 | 0,001364972 |
| ENSG00000120616.16 | EPC1 | protein_coding | 3134,571888 | 0,215540485 | 0,058126342 | 3,70813781 | 0,000208789 | 0,000932837 |

| | | | | | | | | |
|---|---|---|---|---|---|---|---|---|
| ENSG00000120645.11 | IQSEC3 | protein_coding | 5414,664974 | -0,130405998 | 0,051524076 | -2,530972066 | 0,011374691 | 0,033394145 |
| ENSG00000120675.6 | DNAJC15 | protein_coding | 5637,127049 | 0,179343491 | 0,051902472 | 3,455393973 | 0,000549489 | 0,002257429 |
| ENSG00000120688.9 | WBP4 | protein_coding | 3393,215782 | 0,65969942 | 0,055477447 | 11,89130825 | 1,31E-32 | 4,71E-31 |
| ENSG00000120690.16 | ELF1 | protein_coding | 8778,413221 | 1,187795524 | 0,053864727 | 22,05145371 | 9,25E-108 | 1,13E-105 |
| ENSG00000120693.13 | SMAD9 | protein_coding | 1844,972026 | 0,254870471 | 0,072427658 | 3,518966045 | 0,000433232 | 0,001821712 |
| ENSG00000120699.13 | EXOSC8 | protein_coding | 4391,386544 | -0,210459243 | 0,057316789 | -3,671860309 | 0,000240791 | 0,001065017 |
| ENSG00000120709.11 | FAM53C | protein_coding | 7219,713592 | 0,343365426 | 0,047448837 | 7,236540419 | 4,60E-13 | 5,97E-12 |
| ENSG00000120725.13 | SIL1 | protein_coding | 4875,190913 | 0,470477389 | 0,048848962 | 9,631266952 | 5,90E-22 | 1,33E-20 |
| ENSG00000120733.14 | KDM3B | protein_coding | 12387,91831 | 0,248322967 | 0,043184737 | 5,750248407 | 8,91E-09 | 7,44E-08 |
| ENSG00000120742.11 | SERP1 | protein_coding | 15586,09337 | 0,229921723 | 0,042839542 | 5,367044419 | 8,00E-08 | 5,96E-07 |
| ENSG00000120756.13 | PLS1 | protein_coding | 8940,570202 | -0,167768747 | 0,053612584 | -3,129279248 | 0,001752357 | 0,006432453 |
| ENSG00000120802.13 | TMPO | protein_coding | 25347,0464 | -0,295663605 | 0,042833315 | -6,902655248 | 5,10E-12 | 6,03E-11 |
| ENSG00000120875.9 | DUSP4 | protein_coding | 2160,83966 | -0,220746803 | 0,073129506 | -3,018573678 | 0,002539676 | 0,008934913 |
| ENSG00000120885.21 | CLU | protein_coding | 98098,17383 | 0,277559832 | 0,042628765 | 6,511092477 | 7,46E-11 | 7,85E-10 |
| ENSG00000120889.13 | TNFRSF10B | protein_coding | 107,9025209 | 1,702805629 | 0,27503054 | 6,191332886 | 5,97E-10 | 5,73E-09 |
| ENSG00000120899.18 | PTK2B | protein_coding | 1674,680616 | 0,38097274 | 0,084351635 | 4,51648315 | 6,29E-06 | 3,63E-05 |
| ENSG00000120910.14 | PPP3CC | protein_coding | 1150,273525 | 0,253541235 | 0,084967225 | 2,983988665 | 0,002845173 | 0,009886076 |
| ENSG00000120913.23 | PDLIM2 | protein_coding | 335,2181223 | -0,701957742 | 0,184172063 | -3,811423566 | 0,000138169 | 0,000640321 |
| ENSG00000120915.13 | EPHX2 | protein_coding | 5254,742619 | 0,231240024 | 0,070573499 | 3,276584364 | 0,001050709 | 0,004053198 |
| ENSG00000120992.18 | LYPLA1 | protein_coding | 7918,827945 | -0,284614937 | 0,049350987 | -5,767157999 | 8,06E-09 | 6,77E-08 |
| ENSG00000121022.14 | COPS5 | protein_coding | 8668,06617 | -0,126735687 | 0,049920017 | -2,538774915 | 0,011124137 | 0,032782285 |
| ENSG00000121039.10 | RDH10 | protein_coding | 5108,330619 | 0,25732901 | 0,062121202 | 4,142370096 | 3,44E-05 | 0,000176607 |
| ENSG00000121058.5 | COIL | protein_coding | 6341,076696 | -0,160494218 | 0,051794843 | -3,098652472 | 0,001944029 | 0,007045267 |
| ENSG00000121060.18 | TRIM25 | protein_coding | 28591,65024 | 2,337699929 | 0,060090172 | 38,9031991 | 0 | 0 |
| ENSG00000121064.12 | SCPEP1 | protein_coding | 6272,316982 | 0,280386449 | 0,048880231 | 5,736193219 | 9,68E-09 | 8,06E-08 |
| ENSG00000121067.19 | SPOP | protein_coding | 8587,714876 | 0,128212588 | 0,050114757 | 2,558379903 | 0,010516114 | 0,031236105 |
| ENSG00000121104.8 | FAM117A | protein_coding | 1495,414506 | 0,239430186 | 0,078178856 | 3,062595142 | 0,002194267 | 0,007838559 |
| ENSG00000121152.10 | NCAPH | protein_coding | 6180,703125 | -0,150153031 | 0,057953535 | -2,590920982 | 0,009571947 | 0,028846559 |
| ENSG00000121210.16 | TMEM131L | protein_coding | 3237,217445 | -0,146508604 | 0,059082274 | -2,479738754 | 0,013147867 | 0,03789313 |
| ENSG00000121211.7 | MND1 | protein_coding | 783,6716779 | -0,472806085 | 0,105475611 | -4,482610548 | 7,37E-06 | 4,20E-05 |
| ENSG00000121274.12 | TENT4B | protein_coding | 4551,605721 | 0,214148878 | 0,055869216 | 3,833038921 | 0,00012657 | 0,000591562 |
| ENSG00000121297.7 | TSHZ3 | protein_coding | 898,248502 | 0,410328964 | 0,097138704 | 4,224155233 | 2,40E-05 | 0,000126667 |
| ENSG00000121351.7 | IAPP | protein_coding | 1973,565546 | 0,466530469 | 0,07435017 | 6,274773394 | 3,50E-10 | 3,46E-09 |
| ENSG00000121380.12 | BCL2L14 | protein_coding | 29,66630286 | 2,093677617 | 0,513258843 | 4,079184692 | 4,52E-05 | 0,00022819 |
| ENSG00000121406.9 | ZNF549 | protein_coding | 1937,205844 | 0,25316084 | 0,066294767 | 3,818715259 | 0,000134148 | 0,000623101 |
| ENSG00000121410.11 | A1BG | protein_coding | 416,3161157 | -0,930462047 | 0,163343757 | -5,696342886 | 1,22E-08 | 1,01E-07 |

| | | | | | | | | |
|---|---|---|---|---|---|---|---|---|
| ENSG00000121440.15 | PDZRN3 | protein_coding | 3985,524595 | -0,28871721 | 0,06066194 | -4,759445692 | 1,94E-06 | 1,21E-05 |
| ENSG00000121454.6 | LHX4 | protein_coding | 464,2835739 | -0,869854687 | 0,150900214 | -5,764436414 | 8,19E-09 | 6,88E-08 |
| ENSG00000121578.13 | B4GALT4 | protein_coding | 12625,43017 | 0,34261351 | 0,050009024 | 6,851033787 | 7,33E-12 | 8,53E-11 |
| ENSG00000121579.13 | NAA50 | protein_coding | 21175,68954 | -0,2377368 | 0,045646911 | -5,20816835 | 1,91E-07 | 1,36E-06 |
| ENSG00000121621.7 | KIF18A | protein_coding | 3332,631958 | -0,257246715 | 0,055622554 | -4,6248634 | 3,75E-06 | 2,24E-05 |
| ENSG00000121644.19 | DESI2 | protein_coding | 9108,293809 | -0,407918524 | 0,048987681 | -8,326961216 | 8,29E-17 | 1,40E-15 |
| ENSG00000121653.11 | MAPK8IP1 | protein_coding | 9470,351629 | -0,384064725 | 0,051268852 | -7,491190216 | 6,83E-14 | 9,50E-13 |
| ENSG00000121671.11 | CRY2 | protein_coding | 3016,251862 | 0,291244589 | 0,06660634 | 4,372625599 | 1,23E-05 | 6,78E-05 |
| ENSG00000121691.7 | CAT | protein_coding | 6552,043092 | -0,391541952 | 0,04924442 | -7,950991193 | 1,85E-15 | 2,87E-14 |
| ENSG00000121741.16 | ZMYM2 | protein_coding | 22626,03773 | 0,18933988 | 0,041940152 | 4,5145254 | 6,35E-06 | 3,66E-05 |
| ENSG00000121753.12 | ADGRB2 | protein_coding | 2131,731896 | 1,167940825 | 0,0705763 | 16,54862637 | 1,64E-61 | 1,17E-59 |
| ENSG00000121774.18 | KHDRBS1 | protein_coding | 25172,493 | -0,139001228 | 0,045174294 | -3,076998349 | 0,002090965 | 0,007511255 |
| ENSG00000121858.11 | TNFSF10 | protein_coding | 16445,54435 | 3,186415972 | 0,067576221 | 47,15291754 | 0 | 0 |
| ENSG00000121864.10 | ZNF639 | protein_coding | 5889,242228 | 0,199062133 | 0,049258866 | 4,041143255 | 5,32E-05 | 0,00026559 |
| ENSG00000121879.5 | PIK3CA | protein_coding | 3695,261104 | 0,413631995 | 0,078367582 | 5,278100752 | 1,31E-07 | 9,51E-07 |
| ENSG00000121892.15 | PDS5A | protein_coding | 21173,25443 | -0,143048377 | 0,045915473 | -3,115472138 | 0,001836507 | 0,006702 |
| ENSG00000121897.14 | LIAS | protein_coding | 1208,342862 | 0,284447174 | 0,085092222 | 3,3428105 | 0,000829345 | 0,003273892 |
| ENSG00000121903.14 | ZSCAN20 | protein_coding | 912,5905779 | 0,43997028 | 0,086893143 | 5,063348654 | 4,12E-07 | 2,82E-06 |
| ENSG00000121905.10 | HPCA | protein_coding | 6123,443503 | -0,329405829 | 0,04876753 | -6,754613699 | 1,43E-11 | 1,62E-10 |
| ENSG00000121931.16 | LRIF1 | protein_coding | 4679,025823 | 0,301655643 | 0,054406116 | 5,544517148 | 2,95E-08 | 2,33E-07 |
| ENSG00000121940.15 | CLCC1 | protein_coding | 7246,043866 | 0,120161542 | 0,045494452 | 2,641235093 | 0,008260438 | 0,025376579 |
| ENSG00000121957.14 | GPSM2 | protein_coding | 4798,709464 | -0,558161831 | 0,067736896 | -8,240144793 | 1,72E-16 | 2,85E-15 |
| ENSG00000121964.14 | GTDC1 | protein_coding | 2787,080955 | -0,646377853 | 0,070876203 | -9,119814869 | 7,53E-20 | 1,51E-18 |
| ENSG00000121988.18 | ZRANB3 | protein_coding | 2569,773584 | -0,191812301 | 0,065079932 | -2,94733407 | 0,003205267 | 0,010994721 |
| ENSG00000121989.15 | ACVR2A | protein_coding | 1634,198351 | 0,3193315 | 0,073106715 | 4,368018724 | 1,25E-05 | 6,92E-05 |
| ENSG00000122008.15 | POLK | protein_coding | 6203,975765 | 0,35941452 | 0,048108231 | 7,470956828 | 7,96E-14 | 1,10E-12 |
| ENSG00000122026.10 | RPL21 | protein_coding | 62876,53687 | -0,795698548 | 0,041948393 | -18,96851094 | 3,11E-80 | 2,85E-78 |
| ENSG00000122035.6 | RASL11A | protein_coding | 289,9804556 | 0,508222646 | 0,14985331 | 3,391467592 | 0,000695194 | 0,002793309 |
| ENSG00000122042.9 | UBL3 | protein_coding | 15474,56345 | -0,269046979 | 0,05273372 | -5,101991283 | 3,36E-07 | 2,33E-06 |
| ENSG00000122068.13 | FYTTD1 | protein_coding | 15534,63199 | 0,11417333 | 0,044793339 | 2,548890819 | 0,010806613 | 0,031989222 |
| ENSG00000122203.15 | KIAA1191 | protein_coding | 8066,710465 | 0,258518882 | 0,055352329 | 4,670424607 | 3,01E-06 | 1,82E-05 |
| ENSG00000122257.20 | RBBP6 | protein_coding | 11360,84265 | 0,465223155 | 0,067460181 | 6,896263021 | 5,34E-12 | 6,30E-11 |
| ENSG00000122299.12 | ZC3H7A | protein_coding | 14438,06649 | 0,132376606 | 0,043786365 | 3,023238102 | 0,002500854 | 0,008807968 |
| ENSG00000122335.17 | SERAC1 | protein_coding | 4397,683718 | 0,236052717 | 0,058092368 | 4,06340325 | 4,84E-05 | 0,000242609 |
| ENSG00000122378.14 | PRXL2A | protein_coding | 49804,94029 | -0,231662006 | 0,048158944 | -4,810363032 | 1,51E-06 | 9,53E-06 |
| ENSG00000122386.10 | ZNF205 | protein_coding | 1017,668652 | 0,257340098 | 0,085555202 | 3,007883716 | 0,002630738 | 0,009222376 |

| | | | | | | | | |
|---|---|---|---|---|---|---|---|---|
| ENSG00000122390.19 | NAA60 | protein_coding | 8988,308324 | 0,285764961 | 0,048544038 | 5,886715853 | 3,94E-09 | 3,45E-08 |
| ENSG00000122406.14 | RPL5 | protein_coding | 52788,40306 | -0,910388417 | 0,04652394 | -19,56817113 | 2,89E-85 | 2,89E-83 |
| ENSG00000122432.17 | SPATA1 | protein_coding | 34,0881286 | 1,470141435 | 0,471721428 | 3,116545797 | 0,001829833 | 0,006678726 |
| ENSG00000122483.17 | CCDC18 | protein_coding | 1892,846706 | -0,196895482 | 0,082876341 | -2,375774303 | 0,017512171 | 0,04847884 |
| ENSG00000122484.9 | RPAP2 | protein_coding | 3931,708293 | -0,183642371 | 0,071974565 | -2,551489827 | 0,010726346 | 0,031776693 |
| ENSG00000122507.20 | BBS9 | protein_coding | 1517,912264 | -0,351877096 | 0,083649671 | -4,206556863 | 2,59E-05 | 0,000135951 |
| ENSG00000122512.16 | PMS2 | protein_coding | 1529,199554 | -0,222494721 | 0,075539679 | -2,945402005 | 0,003225353 | 0,011056386 |
| ENSG00000122515.15 | ZMIZ2 | protein_coding | 5701,517806 | 0,274828131 | 0,06935006 | 3,962911231 | 7,40E-05 | 0,000361048 |
| ENSG00000122545.19 | sept-07 | protein_coding | 22419,4971 | -0,125170323 | 0,050303952 | -2,488280087 | 0,01283626 | 0,037078378 |
| ENSG00000122547.11 | EEPD1 | protein_coding | 200,8423282 | 0,582209891 | 0,172994756 | 3,365477108 | 0,000764114 | 0,003041671 |
| ENSG00000122557.10 | HERPUD2 | protein_coding | 3405,342163 | 0,402335455 | 0,062212222 | 6,467144912 | 9,99E-11 | 1,04E-09 |
| ENSG00000122565.19 | CBX3 | protein_coding | 28604,45484 | -0,212207783 | 0,044642899 | -4,753449831 | 2,00E-06 | 1,24E-05 |
| ENSG00000122566.21 | HNRNPA2B1 | protein_coding | 170316,541 | -0,131604794 | 0,039556967 | -3,326968803 | 0,000877962 | 0,003445266 |
| ENSG00000122584.13 | NXPH1 | protein_coding | 18,76437238 | 1,905363584 | 0,633245982 | 3,008883813 | 0,002622094 | 0,009197555 |
| ENSG00000122643.21 | NT5C3A | protein_coding | 22344,33194 | 2,649016285 | 0,069139007 | 38,31435243 | 0 | 0 |
| ENSG00000122678.17 | POLM | protein_coding | 1118,146548 | 0,240010775 | 0,08961871 | 2,678132438 | 0,007403394 | 0,023048823 |
| ENSG00000122679.8 | RAMP3 | protein_coding | 87,74026435 | 0,780826306 | 0,262127547 | 2,978802938 | 0,002893768 | 0,010033245 |
| ENSG00000122696.14 | SLC25A51 | protein_coding | 1535,069998 | -0,172644221 | 0,071400984 | -2,417952975 | 0,015608094 | 0,043855022 |
| ENSG00000122705.17 | CLTA | protein_coding | 7574,842861 | -0,133956963 | 0,054267322 | -2,468464602 | 0,013569407 | 0,038938525 |
| ENSG00000122729.19 | ACO1 | protein_coding | 3947,771574 | -1,632702078 | 0,064618249 | -25,26688835 | 7,39E-141 | 1,15E-138 |
| ENSG00000122741.16 | DCAF10 | protein_coding | 7995,188003 | 0,191732607 | 0,048856802 | 3,924378995 | 8,70E-05 | 0,000418841 |
| ENSG00000122783.16 | CYREN | protein_coding | 4128,517386 | -0,410060477 | 0,080365421 | -5,102449178 | 3,35E-07 | 2,32E-06 |
| ENSG00000122786.20 | CALD1 | protein_coding | 28,60658074 | 1,508592414 | 0,468844007 | 3,217685181 | 0,001292296 | 0,004884683 |
| ENSG00000122870.11 | BICC1 | protein_coding | 786,7997135 | -0,365770476 | 0,093383562 | -3,916861465 | 8,97E-05 | 0,000430639 |
| ENSG00000122877.16 | EGR2 | protein_coding | 44,80287335 | 2,09747257 | 0,485295337 | 4,322053828 | 1,55E-05 | 8,39E-05 |
| ENSG00000122912.15 | SLC25A16 | protein_coding | 1799,255004 | -0,526266475 | 0,067538546 | -7,792090649 | 6,59E-15 | 9,87E-14 |
| ENSG00000122952.17 | ZWINT | protein_coding | 12659,40663 | -0,526874307 | 0,041324692 | -12,74962467 | 3,13E-37 | 1,28E-35 |
| ENSG00000122966.16 | CIT | protein_coding | 5155,235183 | -0,772276863 | 0,07160301 | -10,78553631 | 4,03E-27 | 1,13E-25 |
| ENSG00000122970.16 | IFT81 | protein_coding | 7329,210632 | -0,205191708 | 0,047533056 | -4,31682128 | 1,58E-05 | 8,58E-05 |
| ENSG00000122971.9 | ACADS | protein_coding | 1861,74725 | 0,239951606 | 0,076739729 | 3,126823751 | 0,001767058 | 0,006476912 |
| ENSG00000123066.8 | MED13L | protein_coding | 10364,81626 | 0,28962718 | 0,055979196 | 5,173835976 | 2,29E-07 | 1,62E-06 |
| ENSG00000123094.15 | RASSF8 | protein_coding | 2075,907883 | 0,381184051 | 0,078999217 | 4,825162422 | 1,40E-06 | 8,89E-06 |
| ENSG00000123095.6 | BHLHE41 | protein_coding | 164,1160481 | 0,955552443 | 0,222185076 | 4,300704895 | 1,70E-05 | 9,19E-05 |
| ENSG00000123119.12 | NECAB1 | protein_coding | 144,1310889 | -1,119451055 | 0,208333018 | -5,373373207 | 7,73E-08 | 5,77E-07 |
| ENSG00000123124.13 | WWP1 | protein_coding | 2570,739577 | 0,196120716 | 0,059996026 | 3,268895121 | 0,001079683 | 0,004152159 |
| ENSG00000123130.17 | ACOT9 | protein_coding | 668,0723567 | 0,522639942 | 0,128252071 | 4,075099425 | 4,60E-05 | 0,000231973 |

| | | | | | | | | |
|---|---|---|---|---|---|---|---|---|
| ENSG00000123131.13 | PRDX4 | protein_coding | 20654,44831 | 0,213845211 | 0,041980692 | 5,093894359 | 3,51E-07 | 2,43E-06 |
| ENSG00000123136.14 | DDX39A | protein_coding | 6661,671138 | -0,284302847 | 0,058427883 | -4,86587621 | 1,14E-06 | 7,35E-06 |
| ENSG00000123144.11 | TRIR | protein_coding | 4549,625398 | -0,207008091 | 0,076544561 | -2,704412804 | 0,006842523 | 0,021535003 |
| ENSG00000123213.23 | NLN | protein_coding | 2882,460066 | -0,377939629 | 0,070291691 | -5,376732651 | 7,58E-08 | 5,67E-07 |
| ENSG00000123219.13 | CENPK | protein_coding | 5190,104188 | -0,537111675 | 0,049367153 | -10,87994017 | 1,44E-27 | 4,12E-26 |
| ENSG00000123240.17 | OPTN | protein_coding | 11309,39035 | 1,406493165 | 0,05179231 | 27,15640933 | 2,13E-162 | 3,71E-160 |
| ENSG00000123338.13 | NCKAP1L | protein_coding | 216,706249 | 2,72168727 | 0,179471416 | 15,16501807 | 6,03E-52 | 3,65E-50 |
| ENSG00000123349.14 | PFDN5 | protein_coding | 16738,30061 | -0,299759058 | 0,045237838 | -6,626290562 | 3,44E-11 | 3,74E-10 |
| ENSG00000123416.15 | TUBA1B | protein_coding | 128367,5925 | -0,290787269 | 0,040248393 | -7,22481694 | 5,02E-13 | 6,49E-12 |
| ENSG00000123427.17 | EEF1AKMT3 | protein_coding | 395,6336392 | 0,484086578 | 0,12826866 | 3,774005124 | 0,000160647 | 0,000734067 |
| ENSG00000123485.12 | HJURP | protein_coding | 3801,50775 | -0,422483413 | 0,05420047 | -7,794829318 | 6,45E-15 | 9,68E-14 |
| ENSG00000123505.16 | AMD1 | protein_coding | 13305,45133 | -0,119394655 | 0,045373463 | -2,63137629 | 0,008503982 | 0,026034437 |
| ENSG00000123570.4 | RAB9B | protein_coding | 4705,450683 | -0,338226932 | 0,055501103 | -6,094057787 | 1,10E-09 | 1,02E-08 |
| ENSG00000123575.9 | FAM199X | protein_coding | 10018,9423 | -0,143191572 | 0,046920561 | -3,05178727 | 0,002274833 | 0,008096807 |
| ENSG00000123600.19 | METTL8 | protein_coding | 1131,464846 | 0,525672668 | 0,112768413 | 4,661524015 | 3,14E-06 | 1,90E-05 |
| ENSG00000123607.15 | TTC21B | protein_coding | 5874,614402 | 0,273277553 | 0,054930664 | 4,974954479 | 6,53E-07 | 4,36E-06 |
| ENSG00000123609.10 | NMI | protein_coding | 5387,458975 | 3,562638813 | 0,057052914 | 62,44446748 | 0 | 0 |
| ENSG00000123612.16 | ACVR1C | protein_coding | 141978,3795 | -0,408080192 | 0,042077104 | -9,698390612 | 3,06E-22 | 7,00E-21 |
| ENSG00000123636.18 | BAZ2B | protein_coding | 12472,48765 | 0,220069207 | 0,047561402 | 4,627054683 | 3,71E-06 | 2,21E-05 |
| ENSG00000123684.13 | LPGAT1 | protein_coding | 20902,64651 | -0,400633565 | 0,055964923 | -7,15865477 | 8,15E-13 | 1,04E-11 |
| ENSG00000123700.4 | KCNJ2 | protein_coding | 3790,362474 | 0,704183145 | 0,057029284 | 12,34774656 | 5,01E-35 | 1,91E-33 |
| ENSG00000123728.10 | RAP2C | protein_coding | 3232,387571 | 0,219561626 | 0,063202935 | 3,473915023 | 0,000512923 | 0,002118032 |
| ENSG00000123737.12 | EXOSC9 | protein_coding | 5755,398798 | 0,308100217 | 0,052856059 | 5,829042606 | 5,57E-09 | 4,79E-08 |
| ENSG00000123739.11 | PLA2G12A | protein_coding | 7168,107042 | -0,464823832 | 0,072264603 | -6,432247753 | 1,26E-10 | 1,30E-09 |
| ENSG00000123810.8 | B9D2 | protein_coding | 399,8422689 | -0,385212636 | 0,121524287 | -3,169840747 | 0,001525225 | 0,005679287 |
| ENSG00000123815.12 | COQ8B | protein_coding | 3874,767837 | -0,240995392 | 0,070087691 | -3,438483848 | 0,000584981 | 0,002390933 |
| ENSG00000123836.15 | PFKFB2 | protein_coding | 15388,62375 | -0,192023062 | 0,042076746 | -4,563638654 | 5,03E-06 | 2,94E-05 |
| ENSG00000123843.13 | C4BPB | protein_coding | 38,10836485 | 1,703528029 | 0,389917898 | 4,368940325 | 1,25E-05 | 6,89E-05 |
| ENSG00000123870.10 | ZNF137P | transcribed_unproc | 407,23368 | 1,062453221 | 0,158934581 | 6,684846159 | 2,31E-11 | 2,56E-10 |
| ENSG00000123908.12 | AGO2 | protein_coding | 4776,572243 | -0,64991078 | 0,190285211 | -3,415456069 | 0,000636753 | 0,002580982 |
| ENSG00000123933.17 | MXD4 | protein_coding | 4798,281315 | -0,296579262 | 0,066422422 | -4,465047393 | 8,01E-06 | 4,54E-05 |
| ENSG00000123975.5 | CKS2 | protein_coding | 9058,267131 | -0,397982556 | 0,049217217 | -8,086246732 | 6,15E-16 | 9,84E-15 |
| ENSG00000123983.14 | ACSL3 | protein_coding | 17296,83251 | 0,411070173 | 0,046874195 | 8,769647617 | 1,79E-18 | 3,36E-17 |
| ENSG00000123992.19 | DNPEP | protein_coding | 4561,537849 | 1,219849515 | 0,067706875 | 18,016627 | 1,44E-72 | 1,23E-70 |
| ENSG00000123999.5 | INHA | protein_coding | 146,4179799 | -0,602645981 | 0,20858728 | -2,889178963 | 0,003862492 | 0,012993697 |
| ENSG00000124006.15 | OBSL1 | protein_coding | 12094,85735 | -0,265676766 | 0,054100867 | -4,910767217 | 9,07E-07 | 5,93E-06 |

| | | | | | | | | |
|---|---|---|---|---|---|---|---|---|
| ENSG00000124074.12 | ENKD1 | protein_coding | 1220,467832 | -0,242288545 | 0,07869756 | -3,078730096 | 0,002078849 | 0,00747728 |
| ENSG00000124098.10 | FAM210B | protein_coding | 8527,584953 | -0,348124099 | 0,048317123 | -7,204983968 | 5,81E-13 | 7,48E-12 |
| ENSG00000124104.19 | SNX21 | protein_coding | 1265,084895 | 0,291503835 | 0,083371977 | 3,496424647 | 0,000471538 | 0,001967468 |
| ENSG00000124120.11 | TTPAL | protein_coding | 8254,568189 | 0,217055728 | 0,047115859 | 4,606850696 | 4,09E-06 | 2,43E-05 |
| ENSG00000124126.14 | PREX1 | protein_coding | 271,9887192 | -0,890970216 | 0,1498397 | -5,946155919 | 2,75E-09 | 2,44E-08 |
| ENSG00000124140.13 | SLC12A5 | protein_coding | 1423,421348 | -0,285316634 | 0,092535272 | -3,083328427 | 0,002046991 | 0,007374475 |
| ENSG00000124145.6 | SDC4 | protein_coding | 3491,31284 | 0,366198587 | 0,054852776 | 6,676026567 | 2,46E-11 | 2,71E-10 |
| ENSG00000124151.19 | NCOA3 | protein_coding | 5697,616766 | 0,446447779 | 0,061198626 | 7,295062124 | 2,99E-13 | 3,95E-12 |
| ENSG00000124155.18 | PIGT | protein_coding | 15165,31919 | 0,266819292 | 0,046692218 | 5,714427418 | 1,10E-08 | 9,10E-08 |
| ENSG00000124164.15 | VAPB | protein_coding | 32300,29759 | -0,177522561 | 0,051131856 | -3,471858379 | 0,000516869 | 0,00213315 |
| ENSG00000124171.9 | PARD6B | protein_coding | 1093,701548 | 0,323413067 | 0,081121828 | 3,986757624 | 6,70E-05 | 0,000328548 |
| ENSG00000124172.10 | ATP5F1E | protein_coding | 115904,0838 | -0,370703426 | 0,053628929 | -6,91237792 | 4,77E-12 | 5,65E-11 |
| ENSG00000124177.15 | CHD6 | protein_coding | 21794,11686 | 0,530032552 | 0,045196089 | 11,72739873 | 9,23E-32 | 3,20E-30 |
| ENSG00000124191.18 | TOX2 | protein_coding | 76,30755636 | -0,923609059 | 0,277416817 | -3,32931893 | 0,000870587 | 0,00341864 |
| ENSG00000124193.16 | SRSF6 | protein_coding | 41369,1048 | -0,187445891 | 0,040808241 | -4,593334244 | 4,36E-06 | 2,58E-05 |
| ENSG00000124198.9 | ARFGEF2 | protein_coding | 25661,38755 | -0,195832662 | 0,050463343 | -3,880691398 | 0,00010416 | 0,000494216 |
| ENSG00000124201.15 | ZNFX1 | protein_coding | 25540,35208 | 3,174766693 | 0,054255803 | 58,51478572 | 0 | 0 |
| ENSG00000124203.6 | ZNF831 | protein_coding | 376,4198739 | 0,377497071 | 0,130078682 | 2,902067166 | 0,00370709 | 0,012525203 |
| ENSG00000124207.17 | CSE1L | protein_coding | 39014,51208 | -0,160786561 | 0,04259849 | -3,774466232 | 0,000160351 | 0,000732859 |
| ENSG00000124209.4 | RAB22A | protein_coding | 8631,467646 | -0,301839366 | 0,049083746 | -6,149476932 | 7,77E-10 | 7,37E-09 |
| ENSG00000124214.19 | STAU1 | protein_coding | 28386,11225 | 0,148184162 | 0,043347218 | 3,418539132 | 0,000629583 | 0,002553843 |
| ENSG00000124222.22 | STX16 | protein_coding | 18404,23448 | -0,147064472 | 0,04785127 | -3,073366109 | 0,002116586 | 0,007598444 |
| ENSG00000124225.16 | PMEPA1 | protein_coding | 83,81868554 | 1,223767181 | 0,326872888 | 3,743862599 | 0,000181213 | 0,000818403 |
| ENSG00000124226.11 | RNF114 | protein_coding | 16068,28178 | 0,362441544 | 0,046474241 | 7,798761911 | 6,25E-15 | 9,39E-14 |
| ENSG00000124243.17 | BCAS4 | protein_coding | 433,8269107 | 0,809244961 | 0,189479419 | 4,270885814 | 1,95E-05 | 0,000104072 |
| ENSG00000124299.15 | PEPD | protein_coding | 12532,72696 | -0,218556512 | 0,047816887 | -4,570697186 | 4,86E-06 | 2,85E-05 |
| ENSG00000124302.13 | CHST8 | protein_coding | 315,9876124 | -0,560579053 | 0,203430173 | -2,755633767 | 0,005857855 | 0,018755967 |
| ENSG00000124333.16 | VAMP7 | protein_coding | 8428,168212 | -0,118445613 | 0,047053688 | -2,517243964 | 0,01182769 | 0,034543853 |
| ENSG00000124343.13 | XG | protein_coding | 75,14443507 | 1,048620314 | 0,29525503 | 3,551574766 | 0,000382933 | 0,001627846 |
| ENSG00000124374.9 | PAIP2B | protein_coding | 2778,87141 | -0,735442024 | 0,062677561 | -11,7337371 | 8,56E-32 | 2,98E-30 |
| ENSG00000124406.16 | ATP8A1 | protein_coding | 31492,29228 | -0,235666919 | 0,051958775 | -4,535651949 | 5,74E-06 | 3,33E-05 |
| ENSG00000124422.12 | USP22 | protein_coding | 32841,67585 | -0,744004438 | 0,050409459 | -14,75922289 | 2,68E-49 | 1,51E-47 |
| ENSG00000124459.12 | ZNF45 | protein_coding | 4880,710974 | 0,283802028 | 0,052509614 | 5,404763132 | 6,49E-08 | 4,90E-07 |
| ENSG00000124493.13 | GRM4 | protein_coding | 720,4119978 | -0,503144042 | 0,105993211 | -4,746945951 | 2,07E-06 | 1,28E-05 |
| ENSG00000124496.12 | TRERF1 | protein_coding | 1848,301448 | 0,480402721 | 0,063849957 | 7,52393175 | 5,32E-14 | 7,46E-13 |
| ENSG00000124508.16 | BTN2A2 | protein_coding | 845,4025462 | 2,133057792 | 0,117021809 | 18,22786544 | 3,10E-74 | 2,69E-72 |

| | | | | | | | | |
|---|---|---|---|---|---|---|---|---|
| ENSG00000124523.15 | SIRT5 | protein_coding | 1335,788049 | 0,246887138 | 0,083775907 | 2,946994505 | 0,003208789 | 0,011005125 |
| ENSG00000124532.14 | MRS2 | protein_coding | 6161,807724 | -0,138362886 | 0,05071491 | -2,728248681 | 0,006367159 | 0,02019392 |
| ENSG00000124549.14 | BTN2A3P | transcribed_unproc | 213,789369 | 1,713231795 | 0,174265395 | 9,83116469 | 8,27E-23 | 1,94E-21 |
| ENSG00000124562.10 | SNRPC | protein_coding | 8585,789029 | -0,267581137 | 0,049205015 | -5,438086679 | 5,39E-08 | 4,11E-07 |
| ENSG00000124568.10 | SLC17A1 | protein_coding | 310,5323089 | 0,543550506 | 0,141907916 | 3,830304333 | 0,000127985 | 0,000597803 |
| ENSG00000124571.18 | XPO5 | protein_coding | 10783,58131 | 0,115207338 | 0,042494462 | 2,711114132 | 0,006705754 | 0,021142847 |
| ENSG00000124574.15 | ABCC10 | protein_coding | 2603,695948 | 0,240062055 | 0,068423764 | 3,50846023 | 0,000450709 | 0,001890608 |
| ENSG00000124588.20 | NQO2 | protein_coding | 919,336566 | 0,310371635 | 0,084430955 | 3,676040814 | 0,000236882 | 0,001049187 |
| ENSG00000124596.17 | OARD1 | protein_coding | 6506,537708 | -0,114782926 | 0,047451426 | -2,418956299 | 0,01556511 | 0,043745169 |
| ENSG00000124613.8 | ZNF391 | protein_coding | 507,7591848 | 1,044218094 | 0,132688077 | 7,8697206 | 3,55E-15 | 5,43E-14 |
| ENSG00000124614.16 | RPS10 | protein_coding | 32564,93181 | -0,561438755 | 0,045084856 | -12,45293444 | 1,35E-35 | 5,30E-34 |
| ENSG00000124615.20 | MOCS1 | protein_coding | 1306,697705 | 0,309632691 | 0,095704133 | 3,235311583 | 0,001215101 | 0,004616152 |
| ENSG00000124635.8 | HIST1H2BJ | protein_coding | 234,1727614 | 0,6751419 | 0,178139657 | 3,789958452 | 0,000150672 | 0,000693839 |
| ENSG00000124659.6 | TBCC | protein_coding | 5774,424642 | 0,237545622 | 0,049730273 | 4,776680453 | 1,78E-06 | 1,12E-05 |
| ENSG00000124688.13 | MAD2L1BP | protein_coding | 3253,595374 | 0,186187619 | 0,056889023 | 3,272821544 | 0,001064797 | 0,004102621 |
| ENSG00000124702.18 | KLHDC3 | protein_coding | 9020,556475 | -0,464077285 | 0,054856504 | -8,459840685 | 2,68E-17 | 4,65E-16 |
| ENSG00000124733.4 | MEA1 | protein_coding | 6751,649292 | -0,166961049 | 0,053114402 | -3,143423294 | 0,001669842 | 0,006160706 |
| ENSG00000124762.13 | CDKN1A | protein_coding | 1335,51888 | 0,667032912 | 0,091113601 | 7,320892908 | 2,46E-13 | 3,27E-12 |
| ENSG00000124766.6 | SOX4 | protein_coding | 16394,79128 | -0,131195086 | 0,0512167 | -2,561568537 | 0,010420068 | 0,030979436 |
| ENSG00000124767.7 | GLO1 | protein_coding | 17360,82653 | -0,359790829 | 0,043510672 | -8,26902481 | 1,35E-16 | 2,25E-15 |
| ENSG00000124772.12 | CPNE5 | protein_coding | 80,79926761 | -0,754940399 | 0,292339555 | -2,582409341 | 0,009811314 | 0,029449641 |
| ENSG00000124782.20 | RREB1 | protein_coding | 3228,659688 | 0,198700029 | 0,066149426 | 3,003805772 | 0,002666255 | 0,009326298 |
| ENSG00000124783.14 | SSR1 | protein_coding | 31683,69084 | 0,204399073 | 0,046141545 | 4,429827205 | 9,43E-06 | 5,31E-05 |
| ENSG00000124785.9 | NRN1 | protein_coding | 179,7079537 | -0,869374792 | 0,298532144 | -2,912164764 | 0,003589333 | 0,012171179 |
| ENSG00000124788.18 | ATXN1 | protein_coding | 87,88861576 | 1,12688942 | 0,338256136 | 3,331467788 | 0,000863893 | 0,003396571 |
| ENSG00000124789.11 | NUP153 | protein_coding | 13000,5005 | 0,253556196 | 0,045925437 | 5,521040507 | 3,37E-08 | 2,64E-07 |
| ENSG00000124795.17 | DEK | protein_coding | 43612,6404 | -0,209110423 | 0,039585246 | -5,282534441 | 1,27E-07 | 9,30E-07 |
| ENSG00000124802.12 | EEF1E1 | protein_coding | 5069,761924 | -0,206113343 | 0,067998047 | -3,031165624 | 0,002436116 | 0,008600146 |
| ENSG00000124831.19 | LRRFIP1 | protein_coding | 6856,853992 | -0,201622262 | 0,054572259 | -3,694592591 | 0,00022024 | 0,000980493 |
| ENSG00000124875.10 | CXCL6 | protein_coding | 230,6583962 | 2,465231444 | 0,17889821 | 13,78007888 | 3,36E-43 | 1,66E-41 |
| ENSG00000124920.13 | MYRF | protein_coding | 10825,1897 | 0,158427869 | 0,047663695 | 3,323868832 | 0,000887779 | 0,003481364 |
| ENSG00000124942.14 | AHNAK | protein_coding | 139,6292717 | 0,673810721 | 0,244187063 | 2,759403843 | 0,005790693 | 0,018572627 |
| ENSG00000125144.13 | MT1G | protein_coding | 50,07879253 | 2,480907211 | 0,380547111 | 6,519316902 | 7,06E-11 | 7,45E-10 |
| ENSG00000125148.7 | MT2A | protein_coding | 6768,142002 | 4,675599694 | 0,075592596 | 61,85261456 | 0 | 0 |
| ENSG00000125149.11 | C16orf70 | protein_coding | 2645,659707 | 0,191198469 | 0,062380802 | 3,065020997 | 0,002176547 | 0,007782669 |
| ENSG00000125247.15 | TMTC4 | protein_coding | 3523,792441 | -0,286019886 | 0,063766591 | -4,485419149 | 7,28E-06 | 4,15E-05 |

| | | | | | | | | |
|---|---|---|---|---|---|---|---|---|
| ENSG00000125249.7 | RAP2A | protein_coding | 9629,822787 | -0,208508179 | 0,048645845 | -4,286248499 | 1,82E-05 | 9,76E-05 |
| ENSG00000125304.9 | TM9SF2 | protein_coding | 45442,16156 | 0,270893278 | 0,040837952 | 6,633370864 | 3,28E-11 | 3,58E-10 |
| ENSG00000125347.14 | IRF1 | protein_coding | 4687,620497 | 3,358041389 | 0,077066751 | 43,57315346 | 0 | 0 |
| ENSG00000125352.5 | RNF113A | protein_coding | 926,9427874 | 0,237305529 | 0,088046931 | 2,695216355 | 0,007034294 | 0,022070661 |
| ENSG00000125354.23 | sept-06 | protein_coding | 2207,528163 | 0,607041009 | 0,081529218 | 7,44568665 | 9,64E-14 | 1,32E-12 |
| ENSG00000125355.15 | TMEM255A | protein_coding | 388,5038915 | 0,405150525 | 0,163316547 | 2,480768376 | 0,013109953 | 0,037793537 |
| ENSG00000125356.7 | NDUFA1 | protein_coding | 8883,880814 | -0,244259931 | 0,046605145 | -5,241050736 | 1,60E-07 | 1,15E-06 |
| ENSG00000125384.7 | PTGER2 | protein_coding | 2130,548321 | 0,164811433 | 0,064194596 | 2,567372396 | 0,01024725 | 0,030562521 |
| ENSG00000125386.15 | FAM193A | protein_coding | 3432,648366 | 0,157919784 | 0,055999497 | 2,820021492 | 0,004802043 | 0,015731217 |
| ENSG00000125388.20 | GRK4 | protein_coding | 1449,016082 | -0,4372532 | 0,078805364 | -5,548520804 | 2,88E-08 | 2,29E-07 |
| ENSG00000125398.8 | SOX9 | protein_coding | 55,56403268 | 2,17079941 | 0,358678596 | 6,05221341 | 1,43E-09 | 1,31E-08 |
| ENSG00000125449.7 | ARMC7 | protein_coding | 783,2828231 | 0,253912194 | 0,099624196 | 2,548700068 | 0,010812525 | 0,032002514 |
| ENSG00000125450.11 | NUP85 | protein_coding | 6046,019049 | -0,262762631 | 0,05144915 | -5,107229741 | 3,27E-07 | 2,27E-06 |
| ENSG00000125482.13 | TTF1 | protein_coding | 2755,424111 | 0,200289609 | 0,0570071 | 3,513415182 | 0,000442386 | 0,001859507 |
| ENSG00000125485.17 | DDX31 | protein_coding | 1819,469779 | 0,261488965 | 0,066037215 | 3,959721264 | 7,50E-05 | 0,000365113 |
| ENSG00000125505.17 | MBOAT7 | protein_coding | 5638,35009 | 0,173416801 | 0,05379423 | 3,223706343 | 0,001265431 | 0,004788767 |
| ENSG00000125520.14 | SLC2A4RG | protein_coding | 5941,974294 | -0,487613767 | 0,080139246 | -6,084581429 | 1,17E-09 | 1,08E-08 |
| ENSG00000125531.7 | FNDC11 | protein_coding | 130,3442136 | -1,276888175 | 0,237968794 | -5,365779906 | 8,06E-08 | 5,99E-07 |
| ENSG00000125534.10 | PPDPF | protein_coding | 9952,27936 | -0,290149541 | 0,068927239 | -4,209504769 | 2,56E-05 | 0,000134503 |
| ENSG00000125551.18 | PLGLB2 | protein_coding | 164,5304142 | 0,690356694 | 0,207700699 | 3,323805343 | 0,000887982 | 0,00348155 |
| ENSG00000125618.17 | PAX8 | protein_coding | 371,0217384 | 0,46898458 | 0,15335357 | 3,058191477 | 0,002226772 | 0,007943331 |
| ENSG00000125637.15 | PSD4 | protein_coding | 1590,445482 | 0,205844578 | 0,085197975 | 2,416073602 | 0,01568889 | 0,044065537 |
| ENSG00000125651.14 | GTF2F1 | protein_coding | 7117,624474 | 0,444510158 | 0,063488129 | 7,00146883 | 2,53E-12 | 3,09E-11 |
| ENSG00000125652.8 | ALKBH7 | protein_coding | 588,205402 | -0,417230879 | 0,163709806 | -2,548600408 | 0,010815615 | 0,032007451 |
| ENSG00000125656.10 | CLPP | protein_coding | 1148,899703 | -0,233851513 | 0,083684566 | -2,794440155 | 0,005198966 | 0,016901214 |
| ENSG00000125675.18 | GRIA3 | protein_coding | 756,4885135 | 0,338778093 | 0,102020556 | 3,320684638 | 0,000897969 | 0,003515199 |
| ENSG00000125691.13 | RPL23 | protein_coding | 38402,37219 | -0,640266091 | 0,042337555 | -15,12288784 | 1,14E-51 | 6,85E-50 |
| ENSG00000125743.10 | SNRPD2 | protein_coding | 13795,42299 | -0,214086476 | 0,048765311 | -4,390138664 | 1,13E-05 | 6,30E-05 |
| ENSG00000125744.12 | RTN2 | protein_coding | 2554,016821 | -0,395405871 | 0,080081532 | -4,937541289 | 7,91E-07 | 5,21E-06 |
| ENSG00000125753.14 | VASP | protein_coding | 3534,146324 | -0,26333555 | 0,05986115 | -4,399106084 | 1,09E-05 | 6,05E-05 |
| ENSG00000125755.18 | SYMPK | protein_coding | 7478,702672 | 0,267356626 | 0,057316348 | 4,664578879 | 3,09E-06 | 1,87E-05 |
| ENSG00000125772.12 | GPCPD1 | protein_coding | 1488,70294 | 0,249527454 | 0,09674705 | 2,579173769 | 0,009903695 | 0,02967549 |
| ENSG00000125812.16 | GZF1 | protein_coding | 6006,119207 | 0,133047548 | 0,052389832 | 2,539568127 | 0,011098943 | 0,032716881 |
| ENSG00000125814.17 | NAPB | protein_coding | 10518,78583 | -0,293506588 | 0,053105517 | -5,526856767 | 3,26E-08 | 2,56E-07 |
| ENSG00000125820.5 | NKX2-2 | protein_coding | 3842,823593 | -0,189795574 | 0,065547476 | -2,895543583 | 0,003785025 | 0,012755969 |
| ENSG00000125821.12 | DTD1 | protein_coding | 3561,996622 | -0,170099376 | 0,054132115 | -3,142300559 | 0,001676259 | 0,006178305 |

| | | | | | | | | |
|---|---|---|---|---|---|---|---|---|
| ENSG00000125826.20 | RBCK1 | protein_coding | 10081,69922 | 1,491413161 | 0,059359481 | 25,12510436 | 2,64E-139 | 4,05E-137 |
| ENSG00000125827.9 | TMX4 | protein_coding | 39505,33394 | -0,340625999 | 0,044119535 | -7,720525677 | 1,16E-14 | 1,70E-13 |
| ENSG00000125841.13 | NRSN2 | protein_coding | 9425,359004 | 0,333686399 | 0,048307728 | 6,907515828 | 4,93E-12 | 5,84E-11 |
| ENSG00000125844.15 | RRBP1 | protein_coding | 52900,72789 | 0,271762852 | 0,048435384 | 5,610833042 | 2,01E-08 | 1,62E-07 |
| ENSG00000125845.7 | BMP2 | protein_coding | 1218,371457 | 0,890091468 | 0,087498968 | 10,17259393 | 2,63E-24 | 6,62E-23 |
| ENSG00000125846.15 | ZNF133 | protein_coding | 2602,644959 | 0,169438793 | 0,067916664 | 2,494804411 | 0,012602658 | 0,036513479 |
| ENSG00000125848.10 | FLRT3 | protein_coding | 2756,702183 | 0,454473349 | 0,062668455 | 7,252027292 | 4,11E-13 | 5,35E-12 |
| ENSG00000125863.20 | MKKS | protein_coding | 9948,6983 | -0,225645576 | 0,048125026 | -4,688736689 | 2,75E-06 | 1,68E-05 |
| ENSG00000125864.14 | BFSP1 | protein_coding | 54,34633071 | -1,369550746 | 0,385995748 | -3,548098016 | 0,000388024 | 0,001646997 |
| ENSG00000125868.16 | DSTN | protein_coding | 27501,10126 | -0,44580723 | 0,04126749 | -10,80286764 | 3,34E-27 | 9,41E-26 |
| ENSG00000125875.14 | TBC1D20 | protein_coding | 14703,38203 | 0,162636913 | 0,041238299 | 3,943831769 | 8,02E-05 | 0,000388422 |
| ENSG00000125877.13 | ITPA | protein_coding | 5593,553096 | -0,385397094 | 0,058499994 | -6,587985154 | 4,46E-11 | 4,80E-10 |
| ENSG00000125885.13 | MCM8 | protein_coding | 9373,843201 | -0,143174731 | 0,043154362 | -3,31773487 | 0,000907506 | 0,003549198 |
| ENSG00000125944.20 | HNRNPR | protein_coding | 35178,46708 | -0,17416455 | 0,043358686 | -4,016831812 | 5,90E-05 | 0,000292126 |
| ENSG00000125945.15 | ZNF436 | protein_coding | 3125,560677 | 0,197832106 | 0,059321553 | 3,334911095 | 0,000853267 | 0,003360662 |
| ENSG00000125952.19 | MAX | protein_coding | 6592,323422 | 0,801969725 | 0,079504522 | 10,08709571 | 6,30E-24 | 1,56E-22 |
| ENSG00000125962.14 | ARMCX5 | protein_coding | 2887,252971 | 0,172643867 | 0,067094298 | 2,573152591 | 0,010077677 | 0,030132628 |
| ENSG00000125966.10 | MMP24 | protein_coding | 4465,342813 | -0,151575848 | 0,05662614 | -2,676782271 | 0,007433292 | 0,023122716 |
| ENSG00000125967.16 | NECAB3 | protein_coding | 2896,256668 | -0,392273946 | 0,064206333 | -6,109583428 | 9,99E-10 | 9,33E-09 |
| ENSG00000125968.9 | ID1 | protein_coding | 2442,131535 | 0,664577151 | 0,087515289 | 7,593840542 | 3,11E-14 | 4,41E-13 |
| ENSG00000125971.16 | DYNLRB1 | protein_coding | 16488,82763 | -0,136616359 | 0,045557657 | -2,998757382 | 0,002710831 | 0,009467508 |
| ENSG00000125977.7 | EIF2S2 | protein_coding | 28149,79281 | 0,191283532 | 0,042115992 | 4,54182655 | 5,58E-06 | 3,25E-05 |
| ENSG00000125995.16 | ROMO1 | protein_coding | 9019,184606 | -0,162465625 | 0,066303498 | -2,450332632 | 0,01427243 | 0,040706807 |
| ENSG00000126001.16 | CEP250 | protein_coding | 8620,848494 | 0,329376954 | 0,044046575 | 7,477924258 | 7,55E-14 | 1,05E-12 |
| ENSG00000126012.11 | KDM5C | protein_coding | 13623,9214 | 0,13608291 | 0,043690606 | 3,114694923 | 0,001841352 | 0,006717505 |
| ENSG00000126016.15 | AMOT | protein_coding | 244,3388293 | -0,79031335 | 0,15542667 | -5,08479884 | 3,68E-07 | 2,54E-06 |
| ENSG00000126088.14 | UROD | protein_coding | 2573,140438 | -0,414040036 | 0,06496059 | -6,373711193 | 1,85E-10 | 1,87E-09 |
| ENSG00000126106.14 | TMEM53 | protein_coding | 1119,970851 | -0,62470002 | 0,085341892 | -7,319969189 | 2,48E-13 | 3,29E-12 |
| ENSG00000126214.21 | KLC1 | protein_coding | 7400,383442 | -0,238238189 | 0,056301114 | -4,231500423 | 2,32E-05 | 0,000122888 |
| ENSG00000126215.14 | XRCC3 | protein_coding | 2002,239294 | 0,344757823 | 0,076481029 | 4,507756053 | 6,55E-06 | 3,77E-05 |
| ENSG00000126216.15 | TUBGCP3 | protein_coding | 4767,083837 | -0,14247473 | 0,056387539 | -2,526705958 | 0,011513786 | 0,033710172 |
| ENSG00000126217.21 | MCF2L | protein_coding | 10569,94585 | -0,297290905 | 0,04358324 | -6,821220846 | 9,03E-12 | 1,04E-10 |
| ENSG00000126218.11 | F10 | protein_coding | 14748,55617 | 0,322047895 | 0,045724258 | 7,043261242 | 1,88E-12 | 2,32E-11 |
| ENSG00000126247.10 | CAPNS1 | protein_coding | 13184,86738 | -0,209051079 | 0,055998763 | -3,733137421 | 0,000189109 | 0,000851844 |
| ENSG00000126249.8 | PDCD2L | protein_coding | 879,1703289 | 0,328394404 | 0,087864546 | 3,73750754 | 0,000185854 | 0,00083852 |
| ENSG00000126251.6 | GPR42 | protein_coding | 23,81224033 | -1,475758041 | 0,565508561 | -2,609612203 | 0,009064491 | 0,027519913 |

| | | | | | | | | |
|---|---|---|---|---|---|---|---|---|
| ENSG00000126259.19 | KIRREL2 | protein_coding | 5360,897568 | 0,164931293 | 0,050808857 | 3,246113022 | 0,001169924 | 0,004460335 |
| ENSG00000126262.4 | FFAR2 | protein_coding | 1134,555079 | 0,394276429 | 0,087940528 | 4,483443945 | 7,34E-06 | 4,19E-05 |
| ENSG00000126368.6 | NR1D1 | protein_coding | 482,0697958 | 0,508011763 | 0,11928345 | 4,258862087 | 2,05E-05 | 0,000109466 |
| ENSG00000126391.14 | FRMD8 | protein_coding | 2521,494749 | 0,190363555 | 0,079640334 | 2,39029077 | 0,016835038 | 0,046881094 |
| ENSG00000126432.14 | PRDX5 | protein_coding | 9977,613179 | -0,274376471 | 0,053517241 | -5,126879972 | 2,95E-07 | 2,06E-06 |
| ENSG00000126456.16 | IRF3 | protein_coding | 3045,596414 | 0,270652896 | 0,066485225 | 4,070872843 | 4,68E-05 | 0,000235756 |
| ENSG00000126457.21 | PRMT1 | protein_coding | 9434,513949 | -0,280752767 | 0,044966572 | -6,243588417 | 4,28E-10 | 4,18E-09 |
| ENSG00000126460.11 | PRRG2 | protein_coding | 1636,68344 | 0,231441513 | 0,076652936 | 3,019343101 | 0,002533235 | 0,008915037 |
| ENSG00000126461.15 | SCAF1 | protein_coding | 3371,509348 | 0,165016845 | 0,06042002 | 2,731161691 | 0,00631115 | 0,020050483 |
| ENSG00000126464.14 | PRR12 | protein_coding | 2635,952398 | 0,226225058 | 0,065865124 | 3,434671407 | 0,000593273 | 0,002419199 |
| ENSG00000126500.3 | FLRT1 | protein_coding | 16,22461419 | -2,091198356 | 0,65264906 | -3,20416972 | 0,001354526 | 0,005100205 |
| ENSG00000126561.16 | STAT5A | protein_coding | 928,0252277 | -0,469459055 | 0,139089738 | -3,375224235 | 0,000737556 | 0,002947211 |
| ENSG00000126653.18 | NSRP1 | protein_coding | 5726,993258 | 0,185244385 | 0,052827912 | 3,506562703 | 0,000453934 | 0,00190343 |
| ENSG00000126705.14 | AHDC1 | protein_coding | 3661,604679 | -0,259191124 | 0,057715172 | -4,490866345 | 7,09E-06 | 4,06E-05 |
| ENSG00000126709.15 | IFI6 | protein_coding | 123611,1857 | 6,516177896 | 0,204976788 | 31,78983319 | 8,95E-222 | 1,96E-219 |
| ENSG00000126733.21 | DACH2 | protein_coding | 4137,452305 | -0,400512545 | 0,051683224 | -7,749372316 | 9,23E-15 | 1,36E-13 |
| ENSG00000126756.12 | UXT | protein_coding | 7093,207711 | -0,164686702 | 0,050371487 | -3,269442961 | 0,001077595 | 0,004146961 |
| ENSG00000126768.12 | TIMM17B | protein_coding | 10628,35629 | -0,284068432 | 0,056066678 | -5,066617859 | 4,05E-07 | 2,78E-06 |
| ENSG00000126778.11 | SIX1 | protein_coding | 609,3769982 | 1,716396364 | 0,146741698 | 11,69671872 | 1,32E-31 | 4,55E-30 |
| ENSG00000126787.13 | DLGAP5 | protein_coding | 7347,741268 | -0,845541831 | 0,057034887 | -14,82499358 | 1,01E-49 | 5,77E-48 |
| ENSG00000126803.9 | HSPA2 | protein_coding | 10518,54634 | -0,321620024 | 0,04890937 | -6,575836633 | 4,84E-11 | 5,20E-10 |
| ENSG00000126822.17 | PLEKHG3 | protein_coding | 21025,80715 | 0,280008766 | 0,042105246 | 6,650210931 | 2,93E-11 | 3,20E-10 |
| ENSG00000126878.13 | AIF1L | protein_coding | 172,0918465 | -0,490735575 | 0,20549407 | -2,388076576 | 0,016936812 | 0,047117873 |
| ENSG00000126883.17 | NUP214 | protein_coding | 8114,427174 | 0,186970617 | 0,047229599 | 3,958759369 | 7,53E-05 | 0,000366428 |
| ENSG00000126945.9 | HNRNPH2 | protein_coding | 20646,31777 | -0,191068285 | 0,039314185 | -4,860034205 | 1,17E-06 | 7,55E-06 |
| ENSG00000126947.12 | ARMCX1 | protein_coding | 4702,665015 | 0,982080932 | 0,05924348 | 16,5770298 | 1,02E-61 | 7,32E-60 |
| ENSG00000126953.8 | TIMM8A | protein_coding | 927,9452999 | -0,393734728 | 0,088728936 | -4,437500813 | 9,10E-06 | 5,13E-05 |
| ENSG00000127022.15 | CANX | protein_coding | 138911,6666 | 0,106899979 | 0,039660666 | 2,69536523 | 0,007031151 | 0,022063877 |
| ENSG00000127080.10 | IPPK | protein_coding | 2588,876361 | 0,288429139 | 0,060485569 | 4,768561241 | 1,86E-06 | 1,16E-05 |
| ENSG00000127081.14 | ZNF484 | protein_coding | 2262,28602 | 0,350628913 | 0,066078227 | 5,306269986 | 1,12E-07 | 8,21E-07 |
| ENSG00000127124.16 | HIVEP3 | protein_coding | 1071,582304 | -0,292675285 | 0,090169987 | -3,245817085 | 0,001171141 | 0,004464217 |
| ENSG00000127125.9 | PPCS | protein_coding | 4077,815416 | 0,188958851 | 0,06600179 | 2,862935249 | 0,004197363 | 0,013986447 |
| ENSG00000127184.13 | COX7C | protein_coding | 23376,32314 | -0,434343438 | 0,062143771 | -6,989331866 | 2,76E-12 | 3,36E-11 |
| ENSG00000127311.9 | HELB | protein_coding | 543,6389949 | 1,122476584 | 0,122426439 | 9,168579851 | 4,79E-20 | 9,75E-19 |
| ENSG00000127314.18 | RAP1B | protein_coding | 24806,00732 | 0,192483079 | 0,041492277 | 4,63900982 | 3,50E-06 | 2,10E-05 |
| ENSG00000127328.21 | RAB3IP | protein_coding | 14552,67269 | -0,113206017 | 0,0462342 | -2,448534119 | 0,014343884 | 0,040884702 |

| | | | | | | | | |
|---|---|---|---|---|---|---|---|---|
| ENSG00000127334.10 | DYRK2 | protein_coding | 14774,37813 | 0,178289445 | 0,042704369 | 4,174969639 | 2,98E-05 | 0,000154711 |
| ENSG00000127399.15 | LRRC61 | protein_coding | 2490,667291 | 0,270965868 | 0,064445603 | 4,204567203 | 2,62E-05 | 0,000136961 |
| ENSG00000127419.17 | TMEM175 | protein_coding | 2407,216573 | -0,173193279 | 0,062740372 | -2,760475817 | 0,005771723 | 0,018524984 |
| ENSG00000127445.13 | PIN1 | protein_coding | 6440,809144 | -0,253569768 | 0,054288958 | -4,67074298 | 3,00E-06 | 1,82E-05 |
| ENSG00000127481.15 | UBR4 | protein_coding | 25336,4657 | 0,390454003 | 0,046598833 | 8,379051042 | 5,34E-17 | 9,11E-16 |
| ENSG00000127483.18 | HP1BP3 | protein_coding | 28910,90766 | -0,400656048 | 0,053361228 | -7,508373871 | 5,99E-14 | 8,36E-13 |
| ENSG00000127526.15 | SLC35E1 | protein_coding | 5520,373607 | -0,353882383 | 0,060425065 | -5,856549495 | 4,73E-09 | 4,09E-08 |
| ENSG00000127527.14 | EPS15L1 | protein_coding | 3222,408691 | 0,203409519 | 0,063117465 | 3,222713719 | 0,001269824 | 0,004802969 |
| ENSG00000127540.12 | UQCR11 | protein_coding | 7157,392473 | -0,368820969 | 0,05220985 | -7,06420276 | 1,62E-12 | 2,01E-11 |
| ENSG00000127554.13 | GFER | protein_coding | 2862,245775 | 0,219460138 | 0,061703077 | 3,556713018 | 0,000375524 | 0,001599975 |
| ENSG00000127561.15 | SYNGR3 | protein_coding | 265,0401802 | 0,808441022 | 0,149803527 | 5,396675498 | 6,79E-08 | 5,11E-07 |
| ENSG00000127586.17 | CHTF18 | protein_coding | 3740,802352 | -0,42235934 | 0,065651096 | -6,433393586 | 1,25E-10 | 1,29E-09 |
| ENSG00000127666.9 | TICAM1 | protein_coding | 2202,923498 | 1,126322242 | 0,062705306 | 17,96215198 | 3,86E-72 | 3,21E-70 |
| ENSG00000127774.7 | EMC6 | protein_coding | 1292,230925 | -0,234091542 | 0,096240555 | -2,432358606 | 0,015000847 | 0,042424182 |
| ENSG00000127804.13 | METTL16 | protein_coding | 4683,170192 | -0,199397401 | 0,07713473 | -2,585053473 | 0,00973639 | 0,029252055 |
| ENSG00000127824.14 | TUBA4A | protein_coding | 14745,81467 | -0,167347057 | 0,042234671 | -3,962314691 | 7,42E-05 | 0,000361716 |
| ENSG00000127831.10 | VIL1 | protein_coding | 98184,58053 | -0,297770878 | 0,047055056 | -6,328137805 | 2,48E-10 | 2,49E-09 |
| ENSG00000127863.15 | TNFRSF19 | protein_coding | 5520,716718 | -0,160614791 | 0,04940649 | -3,250884465 | 0,001150466 | 0,004396577 |
| ENSG00000127884.5 | ECHS1 | protein_coding | 4432,446883 | -0,161288527 | 0,050108185 | -3,218806031 | 0,001287255 | 0,004866448 |
| ENSG00000127947.16 | PTPN12 | protein_coding | 5923,299924 | 0,362738129 | 0,064779201 | 5,599607985 | 2,15E-08 | 1,73E-07 |
| ENSG00000127989.13 | MTERF1 | protein_coding | 3138,905687 | 0,21994305 | 0,054942168 | 4,00317387 | 6,25E-05 | 0,000308503 |
| ENSG00000127990.18 | SGCE | protein_coding | 6707,657857 | 0,163380594 | 0,050886639 | 3,210677675 | 0,001324224 | 0,004996137 |
| ENSG00000127993.16 | RBM48 | protein_coding | 2228,350647 | 0,218364715 | 0,072401045 | 3,016043692 | 0,002560964 | 0,008999958 |
| ENSG00000128016.6 | ZFP36 | protein_coding | 1219,139969 | 0,474923461 | 0,08577032 | 5,53715388 | 3,07E-08 | 2,42E-07 |
| ENSG00000128039.11 | SRD5A3 | protein_coding | 912,8568398 | -0,719421387 | 0,097766483 | -7,358568784 | 1,86E-13 | 2,49E-12 |
| ENSG00000128045.7 | RASL11B | protein_coding | 202,8731796 | -0,83042281 | 0,17862745 | -4,648909272 | 3,34E-06 | 2,01E-05 |
| ENSG00000128050.8 | PAICS | protein_coding | 27054,72884 | -0,426604107 | 0,047901347 | -8,905889675 | 5,30E-19 | 1,02E-17 |
| ENSG00000128165.9 | ADM2 | protein_coding | 379,8088003 | 0,857281364 | 0,121395994 | 7,061858789 | 1,64E-12 | 2,04E-11 |
| ENSG00000128185.9 | DGCR6L | protein_coding | 3792,371144 | -0,260693636 | 0,071997166 | -3,620887441 | 0,000293594 | 0,001277501 |
| ENSG00000128203.7 | ASPHD2 | protein_coding | 726,6816634 | 0,614343603 | 0,099072816 | 6,200930062 | 5,61E-10 | 5,42E-09 |
| ENSG00000128228.5 | SDF2L1 | protein_coding | 2034,283455 | 0,622133494 | 0,077016431 | 8,077932069 | 6,59E-16 | 1,05E-14 |
| ENSG00000128245.15 | YWHAH | protein_coding | 10959,11719 | -0,157742071 | 0,04290287 | -3,676725399 | 0,000236247 | 0,001046583 |
| ENSG00000128268.12 | MGAT3 | protein_coding | 6228,565106 | 0,156796557 | 0,064581063 | 2,427903024 | 0,015186403 | 0,042872543 |
| ENSG00000128272.14 | ATF4 | protein_coding | 23635,67172 | 0,731829506 | 0,056974217 | 12,84492434 | 9,18E-38 | 3,82E-36 |
| ENSG00000128274.17 | A4GALT | protein_coding | 145,3787997 | -0,633664266 | 0,216167776 | -2,931353961 | 0,00337488 | 0,011527341 |
| ENSG00000128309.16 | MPST | protein_coding | 1734,322122 | -0,254907162 | 0,076358686 | -3,338286398 | 0,000842968 | 0,003324754 |

| | | | | | | | | |
|---|---|---|---|---|---|---|---|---|
| ENSG00000128335.14 | APOL2 | protein_coding | 8424,988274 | 2,872263878 | 0,054074229 | 53,11705638 | 0 | 0 |
| ENSG00000128394.17 | APOBEC3F | protein_coding | 1207,458292 | 2,7342915 | 0,086588553 | 31,57797886 | 7,41E-219 | 1,60E-216 |
| ENSG00000128463.13 | EMC4 | protein_coding | 6524,928504 | -0,115795983 | 0,045749238 | -2,531101877 | 0,011370482 | 0,033386143 |
| ENSG00000128487.16 | SPECC1 | protein_coding | 2794,592241 | -0,264087601 | 0,077119794 | -3,424381586 | 0,0006162 | 0,002504072 |
| ENSG00000128512.22 | DOCK4 | protein_coding | 837,1578925 | 0,636448928 | 0,086964289 | 7,318508953 | 2,51E-13 | 3,33E-12 |
| ENSG00000128563.13 | PRKRIP1 | protein_coding | 3114,904194 | 0,227072508 | 0,061999749 | 3,662474604 | 0,000249791 | 0,001101162 |
| ENSG00000128564.7 | VGF | protein_coding | 1788,581355 | 0,78466281 | 0,070032984 | 11,20418929 | 3,89E-29 | 1,18E-27 |
| ENSG00000128567.17 | PODXL | protein_coding | 1429,846562 | 0,303643812 | 0,101195627 | 3,000562582 | 0,002694814 | 0,009418876 |
| ENSG00000128573.25 | FOXP2 | protein_coding | 2714,948566 | -0,373215122 | 0,063062296 | -5,918197502 | 3,25E-09 | 2,87E-08 |
| ENSG00000128581.16 | IFT22 | protein_coding | 3771,031874 | -0,177607672 | 0,060723593 | -2,924854484 | 0,003446173 | 0,011745899 |
| ENSG00000128585.18 | MKLN1 | protein_coding | 13194,00428 | 0,195618674 | 0,050931309 | 3,840833428 | 0,000122617 | 0,000574879 |
| ENSG00000128590.5 | DNAJB9 | protein_coding | 10258,93402 | 0,51234562 | 0,059517976 | 8,608250096 | 7,42E-18 | 1,34E-16 |
| ENSG00000128596.17 | CCDC136 | protein_coding | 365,2865929 | 0,4419776 | 0,170181335 | 2,59709797 | 0,00940151 | 0,02842044 |
| ENSG00000128602.11 | SMO | protein_coding | 235,8859067 | -0,729514645 | 0,16094639 | -4,532656156 | 5,82E-06 | 3,38E-05 |
| ENSG00000128604.20 | IRF5 | protein_coding | 317,3340217 | 0,536956813 | 0,144394605 | 3,718676416 | 0,000200269 | 0,00089816 |
| ENSG00000128607.14 | KLHDC10 | protein_coding | 10389,75212 | -0,227454052 | 0,064249792 | -3,540152371 | 0,000399896 | 0,00169133 |
| ENSG00000128609.15 | NDUFA5 | protein_coding | 18719,61079 | -0,218493357 | 0,044931029 | -4,862861201 | 1,16E-06 | 7,45E-06 |
| ENSG00000128656.14 | CHN1 | protein_coding | 4270,727714 | -0,331066055 | 0,06725356 | -4,92265471 | 8,54E-07 | 5,59E-06 |
| ENSG00000128694.12 | OSGEPL1 | protein_coding | 1546,51001 | -0,265829812 | 0,073179859 | -3,6325543 | 0,000280629 | 0,001224639 |
| ENSG00000128699.14 | ORMDL1 | protein_coding | 7838,45762 | -0,131215353 | 0,049545917 | -2,64835855 | 0,008088369 | 0,024915984 |
| ENSG00000128833.13 | MYO5C | protein_coding | 3991,129499 | -0,194125469 | 0,05169983 | -3,754856985 | 0,00017344 | 0,000788044 |
| ENSG00000128849.10 | CGNL1 | protein_coding | 5531,038547 | -0,502340715 | 0,061535997 | -8,163363582 | 3,26E-16 | 5,30E-15 |
| ENSG00000128872.10 | TMOD2 | protein_coding | 9085,844072 | -0,186661978 | 0,05368691 | -3,476862014 | 0,000507319 | 0,002098353 |
| ENSG00000128881.17 | TTBK2 | protein_coding | 5923,635262 | 0,439493085 | 0,068881725 | 6,380401874 | 1,77E-10 | 1,80E-09 |
| ENSG00000128891.15 | CCDC32 | protein_coding | 3437,300188 | 0,186263479 | 0,053535361 | 3,479260724 | 0,000502799 | 0,002082721 |
| ENSG00000128908.17 | INO80 | protein_coding | 4785,360674 | 0,560063945 | 0,057320021 | 9,770825851 | 1,50E-22 | 3,48E-21 |
| ENSG00000128923.11 | MINDY2 | protein_coding | 3891,916758 | 0,448228487 | 0,055317944 | 8,102768363 | 5,37E-16 | 8,63E-15 |
| ENSG00000128944.13 | KNSTRN | protein_coding | 4425,026238 | -0,655261587 | 0,057030111 | -11,4897478 | 1,49E-30 | 4,84E-29 |
| ENSG00000128951.14 | DUT | protein_coding | 9684,65808 | -0,326663978 | 0,045906725 | -7,115819777 | 1,11E-12 | 1,40E-11 |
| ENSG00000128965.13 | CHAC1 | protein_coding | 302,7489161 | 1,882618453 | 0,163486147 | 11,51546165 | 1,10E-30 | 3,61E-29 |
| ENSG00000128973.13 | CLN6 | protein_coding | 4532,651595 | -0,181607634 | 0,074192295 | -2,447796437 | 0,014373284 | 0,040932216 |
| ENSG00000128989.10 | ARPP19 | protein_coding | 42178,0986 | -0,28499964 | 0,040267901 | -7,077588749 | 1,47E-12 | 1,83E-11 |
| ENSG00000129003.18 | VPS13C | protein_coding | 19756,89254 | 0,459670618 | 0,051463021 | 8,932056647 | 4,18E-19 | 8,10E-18 |
| ENSG00000129083.12 | COPB1 | protein_coding | 26723,86894 | 0,154651505 | 0,046866108 | 3,29985808 | 0,000967337 | 0,003764488 |
| ENSG00000129167.9 | TPH1 | protein_coding | 463,9937374 | -0,889106738 | 0,13032497 | -6,822228598 | 8,96E-12 | 1,03E-10 |
| ENSG00000129173.13 | E2F8 | protein_coding | 1916,03516 | -0,478941991 | 0,082565657 | -5,800740994 | 6,60E-09 | 5,63E-08 |

| | | | | | | | | |
|---|---|---|---|---|---|---|---|---|
| ENSG00000129195.16 | PIMREG | protein_coding | 1097,062489 | -0,86148693 | 0,086821104 | -9,922552082 | 3,32E-23 | 7,91E-22 |
| ENSG00000129197.14 | RPAIN | protein_coding | 7391,031831 | -0,133779778 | 0,054082066 | -2,473644021 | 0,013374289 | 0,038437418 |
| ENSG00000129226.14 | CD68 | protein_coding | 11872,31049 | 2,982007765 | 0,061331025 | 48,621522 | 0 | 0 |
| ENSG00000129235.10 | TXNDC17 | protein_coding | 6216,13517 | -0,347982364 | 0,051897938 | -6,705128913 | 2,01E-11 | 2,24E-10 |
| ENSG00000129315.11 | CCNT1 | protein_coding | 10035,67954 | 0,244291732 | 0,044974501 | 5,431783018 | 5,58E-08 | 4,25E-07 |
| ENSG00000129317.14 | PUS7L | protein_coding | 5234,12408 | 0,149880848 | 0,061230562 | 2,447811088 | 0,014372699 | 0,040932216 |
| ENSG00000129347.20 | KRI1 | protein_coding | 2346,770995 | -0,431389989 | 0,081059758 | -5,321876109 | 1,03E-07 | 7,56E-07 |
| ENSG00000129351.17 | ILF3 | protein_coding | 34449,0734 | -0,166385079 | 0,038742948 | -4,294589998 | 1,75E-05 | 9,42E-05 |
| ENSG00000129353.15 | SLC44A2 | protein_coding | 1664,709034 | -0,453051342 | 0,074874731 | -6,050790924 | 1,44E-09 | 1,32E-08 |
| ENSG00000129354.11 | AP1M2 | protein_coding | 8002,993865 | -0,360627997 | 0,054415511 | -6,627301442 | 3,42E-11 | 3,72E-10 |
| ENSG00000129422.14 | MTUS1 | protein_coding | 5968,336104 | 0,229020098 | 0,052572442 | 4,356276623 | 1,32E-05 | 7,27E-05 |
| ENSG00000129460.16 | NGDN | protein_coding | 4370,725074 | 0,259906841 | 0,056078308 | 4,634712576 | 3,57E-06 | 2,14E-05 |
| ENSG00000129465.16 | RIPK3 | protein_coding | 64,30020475 | 1,271803618 | 0,314842531 | 4,03949115 | 5,36E-05 | 0,000267231 |
| ENSG00000129472.15 | RAB2B | protein_coding | 3537,690093 | 0,282984584 | 0,067786397 | 4,174651493 | 2,98E-05 | 0,000154891 |
| ENSG00000129473.9 | BCL2L2 | protein_coding | 12786,63565 | 0,287876916 | 0,048067085 | 5,989065337 | 2,11E-09 | 1,90E-08 |
| ENSG00000129474.16 | AJUBA | protein_coding | 13,05095724 | 1,743675123 | 0,659311702 | 2,644690088 | 0,008176577 | 0,025146409 |
| ENSG00000129480.13 | DTD2 | protein_coding | 1413,697355 | -0,249738297 | 0,083691421 | -2,984037013 | 0,002844724 | 0,00988604 |
| ENSG00000129484.13 | PARP2 | protein_coding | 6590,155365 | -0,255942114 | 0,050867641 | -5,03153102 | 4,87E-07 | 3,30E-06 |
| ENSG00000129515.19 | SNX6 | protein_coding | 12847,92547 | 0,495026564 | 0,048745729 | 10,15528083 | 3,14E-24 | 7,89E-23 |
| ENSG00000129534.14 | MIS18BP1 | protein_coding | 5607,56195 | -0,193356808 | 0,051922725 | -3,723934116 | 0,000196142 | 0,000881582 |
| ENSG00000129559.13 | NEDD8 | protein_coding | 9089,030973 | -0,247164568 | 0,047961132 | -5,153434817 | 2,56E-07 | 1,80E-06 |
| ENSG00000129562.11 | DAD1 | protein_coding | 8689,547379 | -0,16476226 | 0,043436151 | -3,793205849 | 0,000148715 | 0,000685806 |
| ENSG00000129566.13 | TEP1 | protein_coding | 4681,81688 | 0,689306531 | 0,067202175 | 10,25720563 | 1,10E-24 | 2,82E-23 |
| ENSG00000129636.12 | ITFG1 | protein_coding | 18287,0219 | 0,155952461 | 0,041222001 | 3,783233632 | 0,000154804 | 0,000710684 |
| ENSG00000129667.12 | RHBDF2 | protein_coding | 348,3307185 | 0,386088322 | 0,157301276 | 2,454451312 | 0,014109977 | 0,040299641 |
| ENSG00000129757.13 | CDKN1C | protein_coding | 6459,619294 | -0,762444216 | 0,059030798 | -12,91604109 | 3,65E-38 | 1,55E-36 |
| ENSG00000129810.15 | SGO1 | protein_coding | 2691,222796 | -0,26406348 | 0,066003333 | -4,000759762 | 6,31E-05 | 0,000311395 |
| ENSG00000129824.16 | RPS4Y1 | protein_coding | 17874,13003 | -0,505067348 | 0,040227757 | -12,55519527 | 3,72E-36 | 1,49E-34 |
| ENSG00000129933.21 | MAU2 | protein_coding | 5807,798328 | 0,208102174 | 0,05246998 | 3,966118773 | 7,31E-05 | 0,000356535 |
| ENSG00000129993.14 | CBFA2T3 | protein_coding | 1590,99907 | -0,731602554 | 0,102151674 | -7,161924255 | 7,96E-13 | 1,01E-11 |
| ENSG00000130005.12 | GAMT | protein_coding | 1070,675712 | -0,5767355 | 0,110720959 | -5,208909893 | 1,90E-07 | 1,36E-06 |
| ENSG00000130024.15 | PHF10 | protein_coding | 12316,5789 | -0,291607102 | 0,048333919 | -6,03317726 | 1,61E-09 | 1,47E-08 |
| ENSG00000130066.16 | SAT1 | protein_coding | 16142,1625 | 0,850744011 | 0,054837778 | 15,51383081 | 2,80E-54 | 1,73E-52 |
| ENSG00000130119.16 | GNL3L | protein_coding | 3633,231587 | -0,262661377 | 0,068489138 | -3,835080764 | 0,000125523 | 0,000587279 |
| ENSG00000130147.16 | SH3BP4 | protein_coding | 251,8885795 | -0,477925598 | 0,175853626 | -2,717746624 | 0,006572815 | 0,020782061 |
| ENSG00000130150.12 | MOSPD2 | protein_coding | 10005,4496 | 0,122151655 | 0,047663742 | 2,562779346 | 0,010383802 | 0,030887934 |

| | | | | | | | | |
|---|---|---|---|---|---|---|---|---|
| ENSG00000130175.9 | PRKCSH | protein_coding | 24246,22661 | -0,129767663 | 0,045411378 | -2,857602384 | 0,004268549 | 0,014196331 |
| ENSG00000130202.10 | NECTIN2 | protein_coding | 1470,816141 | 0,873385323 | 0,089959276 | 9,708674441 | 2,77E-22 | 6,34E-21 |
| ENSG00000130204.13 | TOMM40 | protein_coding | 2931,677967 | -0,25763103 | 0,066127102 | -3,895997558 | 9,78E-05 | 0,000466072 |
| ENSG00000130208.9 | APOC1 | protein_coding | 350,0872353 | 0,330151813 | 0,134980753 | 2,445917711 | 0,014448397 | 0,041094134 |
| ENSG00000130224.15 | LRCH2 | protein_coding | 4157,602853 | 1,213841303 | 0,073725534 | 16,46432709 | 6,62E-61 | 4,67E-59 |
| ENSG00000130226.17 | DPP6 | protein_coding | 12471,29331 | -0,198142341 | 0,042886089 | -4,620200719 | 3,83E-06 | 2,28E-05 |
| ENSG00000130234.10 | ACE2 | protein_coding | 381,6846406 | 6,036114211 | 0,509164523 | 11,85493872 | 2,03E-32 | 7,24E-31 |
| ENSG00000130294.16 | KIF1A | protein_coding | 43349,81257 | -0,134493971 | 0,055443126 | -2,425800646 | 0,015274658 | 0,043074077 |
| ENSG00000130303.13 | BST2 | protein_coding | 4947,468225 | 13,86348005 | 1,403991668 | 9,874332139 | 5,38E-23 | 1,27E-21 |
| ENSG00000130305.16 | NSUN5 | protein_coding | 2425,075289 | 0,272684063 | 0,060212237 | 4,528715023 | 5,93E-06 | 3,44E-05 |
| ENSG00000130332.15 | LSM7 | protein_coding | 2873,050236 | -0,179883944 | 0,059811407 | -3,007519024 | 0,002633896 | 0,009227451 |
| ENSG00000130338.13 | TULP4 | protein_coding | 5085,745738 | 0,321502074 | 0,065205749 | 4,930578643 | 8,20E-07 | 5,39E-06 |
| ENSG00000130340.15 | SNX9 | protein_coding | 11168,21666 | 0,351285095 | 0,04306129 | 8,157793069 | 3,41E-16 | 5,54E-15 |
| ENSG00000130348.11 | QRSL1 | protein_coding | 4475,326072 | 0,228741527 | 0,050777091 | 4,504817465 | 6,64E-06 | 3,81E-05 |
| ENSG00000130414.12 | NDUFA10 | protein_coding | 8797,898786 | -0,196941745 | 0,045992898 | -4,282003407 | 1,85E-05 | 9,94E-05 |
| ENSG00000130449.6 | ZSWIM6 | protein_coding | 3364,860966 | 0,187632256 | 0,061874632 | 3,03245851 | 0,002425704 | 0,008568767 |
| ENSG00000130477.15 | UNC13A | protein_coding | 16103,70312 | -0,232329478 | 0,06460593 | -3,596101444 | 0,000323022 | 0,001394763 |
| ENSG00000130487.8 | KLHDC7B | protein_coding | 67,6386655 | 2,016845289 | 0,359797059 | 5,605507982 | 2,08E-08 | 1,67E-07 |
| ENSG00000130508.11 | PXDN | protein_coding | 2540,62229 | -0,449804941 | 0,082723259 | -5,437466377 | 5,40E-08 | 4,13E-07 |
| ENSG00000130511.16 | SSBP4 | protein_coding | 2064,128184 | -0,400261234 | 0,088935346 | -4,50058669 | 6,78E-06 | 3,89E-05 |
| ENSG00000130517.14 | PGPEP1 | protein_coding | 9367,672185 | -0,262366381 | 0,044756747 | -5,862052078 | 4,57E-09 | 3,96E-08 |
| ENSG00000130520.10 | LSM4 | protein_coding | 6711,472059 | -0,477829081 | 0,049962992 | -9,563660172 | 1,14E-21 | 2,52E-20 |
| ENSG00000130522.5 | JUND | protein_coding | 3301,058655 | 0,408609788 | 0,069415793 | 5,886409549 | 3,95E-09 | 3,45E-08 |
| ENSG00000130529.16 | TRPM4 | protein_coding | 343,9962022 | 0,379986783 | 0,14816839 | 2,564560392 | 0,01033066 | 0,030766449 |
| ENSG00000130544.12 | ZNF557 | protein_coding | 1354,799912 | 0,576341743 | 0,082190822 | 7,012239695 | 2,35E-12 | 2,87E-11 |
| ENSG00000130558.19 | OLFM1 | protein_coding | 3797,984423 | -0,619280778 | 0,056549953 | -10,95103959 | 6,57E-28 | 1,90E-26 |
| ENSG00000130584.11 | ZBTB46 | protein_coding | 1715,563125 | -0,498925058 | 0,095162353 | -5,242882717 | 1,58E-07 | 1,14E-06 |
| ENSG00000130589.16 | HELZ2 | protein_coding | 39134,4164 | 5,200794836 | 0,06152066 | 84,53737018 | 0 | 0 |
| ENSG00000130595.19 | TNNT3 | protein_coding | 71,03074995 | -2,094254202 | 0,383048062 | -5,467340557 | 4,57E-08 | 3,52E-07 |
| ENSG00000130600.19 | H19 | lncRNA | 20121,90961 | -1,011846003 | 0,065082576 | -15,54711047 | 1,66E-54 | 1,04E-52 |
| ENSG00000130635.15 | COL5A1 | protein_coding | 116,9580881 | -0,848856761 | 0,228811404 | -3,709853388 | 0,000207379 | 0,000926906 |
| ENSG00000130638.17 | ATXN10 | protein_coding | 15148,53561 | -0,170250973 | 0,052078627 | -3,269114095 | 0,001078848 | 0,004149656 |
| ENSG00000130684.14 | ZNF337 | protein_coding | 2592,146264 | 0,146047232 | 0,0571525 | 2,555395342 | 0,010606725 | 0,031472025 |
| ENSG00000130695.15 | CEP85 | protein_coding | 7409,339388 | -0,136040266 | 0,047670226 | -2,853778508 | 0,004320264 | 0,014336551 |
| ENSG00000130702.15 | LAMA5 | protein_coding | 6072,109931 | -0,60781025 | 0,07651393 | -7,943785562 | 1,96E-15 | 3,04E-14 |
| ENSG00000130703.16 | OSBPL2 | protein_coding | 10076,23423 | 0,118294813 | 0,046273965 | 2,556401047 | 0,010576115 | 0,031394354 |

| | | | | | | | | |
|---|---|---|---|---|---|---|---|---|
| ENSG00000130707.17 | ASS1 | protein_coding | 978,8371042 | -1,475516608 | 0,091308413 | -16,15970059 | 9,70E-59 | 6,68E-57 |
| ENSG00000130713.16 | EXOSC2 | protein_coding | 4024,191103 | -0,166578726 | 0,054029644 | -3,083098727 | 0,002048572 | 0,007378989 |
| ENSG00000130723.20 | PRRC2B | protein_coding | 28467,66354 | 0,115620952 | 0,043949725 | 2,630754862 | 0,008519546 | 0,026069395 |
| ENSG00000130724.8 | CHMP2A | protein_coding | 5803,576462 | -0,379834569 | 0,049259541 | -7,710883232 | 1,25E-14 | 1,82E-13 |
| ENSG00000130726.12 | TRIM28 | protein_coding | 27985,9151 | -0,145349569 | 0,042631547 | -3,40943689 | 0,000650971 | 0,002633013 |
| ENSG00000130731.16 | METTL26 | protein_coding | 1007,056468 | -0,502262288 | 0,126337886 | -3,975547655 | 7,02E-05 | 0,000343518 |
| ENSG00000130748.7 | TMEM160 | protein_coding | 391,6021598 | -0,531837871 | 0,205436229 | -2,588822201 | 0,00963048 | 0,029007425 |
| ENSG00000130749.10 | AC008755.1 | protein_coding | 4221,533516 | 0,167879902 | 0,050999673 | 3,291783897 | 0,000995541 | 0,003858899 |
| ENSG00000130758.8 | MAP3K10 | protein_coding | 1230,465729 | 0,379944677 | 0,082101696 | 4,627732353 | 3,70E-06 | 2,21E-05 |
| ENSG00000130762.15 | ARHGEF16 | protein_coding | 1430,11563 | 0,240333555 | 0,075736066 | 3,173303908 | 0,001507147 | 0,005622202 |
| ENSG00000130764.10 | LRRC47 | protein_coding | 5247,293883 | 0,143911719 | 0,054262463 | 2,652141298 | 0,007998306 | 0,024672314 |
| ENSG00000130768.15 | SMPDL3B | protein_coding | 4222,687276 | 0,338854157 | 0,051763013 | 6,546260257 | 5,90E-11 | 6,28E-10 |
| ENSG00000130803.15 | ZNF317 | protein_coding | 4689,77929 | 0,178147454 | 0,051244216 | 3,476440245 | 0,000508117 | 0,002100885 |
| ENSG00000130810.20 | PPAN | protein_coding | 1623,071025 | 0,233321498 | 0,068571682 | 3,402592605 | 0,000667497 | 0,002693081 |
| ENSG00000130812.10 | ANGPTL6 | protein_coding | 39,66668608 | 2,439661885 | 0,444589279 | 5,487451006 | 4,08E-08 | 3,15E-07 |
| ENSG00000130813.18 | C19orf66 | protein_coding | 8515,649376 | 4,423092563 | 0,062211301 | 71,09789559 | 0 | 0 |
| ENSG00000130818.12 | ZNF426 | protein_coding | 2746,114155 | 0,474711126 | 0,062156796 | 7,637316506 | 2,22E-14 | 3,18E-13 |
| ENSG00000130826.18 | DKC1 | protein_coding | 11161,62071 | -0,2060324 | 0,048330912 | -4,26295285 | 2,02E-05 | 0,000107658 |
| ENSG00000130829.18 | DUSP9 | protein_coding | 1244,447835 | -0,8746599 | 0,104861871 | -8,34106712 | 7,36E-17 | 1,25E-15 |
| ENSG00000130830.15 | MPP1 | protein_coding | 6027,207691 | -0,202232475 | 0,049847458 | -4,057026858 | 4,97E-05 | 0,000248938 |
| ENSG00000130876.11 | SLC7A10 | protein_coding | 27,06333645 | -2,281411507 | 0,470708948 | -4,846756188 | 1,25E-06 | 8,05E-06 |
| ENSG00000130881.14 | LRP3 | protein_coding | 2801,024795 | -0,418023137 | 0,06593732 | -6,339704718 | 2,30E-10 | 2,31E-09 |
| ENSG00000130939.19 | UBE4B | protein_coding | 8701,668608 | -0,45216207 | 0,04969125 | -9,099430339 | 9,08E-20 | 1,82E-18 |
| ENSG00000130940.15 | CASZ1 | protein_coding | 2255,973815 | 0,524306885 | 0,060498332 | 8,666468422 | 4,46E-18 | 8,16E-17 |
| ENSG00000130962.17 | PRRG1 | protein_coding | 2915,941103 | 0,864668278 | 0,057633968 | 15,00275471 | 7,04E-51 | 4,13E-49 |
| ENSG00000130988.13 | RGN | protein_coding | 331,5197105 | -0,477401508 | 0,140620767 | -3,394957355 | 0,000686393 | 0,002760411 |
| ENSG00000131002.12 | TXLNGY | transcribed_unproc | 3421,411089 | 0,219814348 | 0,059756068 | 3,678527633 | 0,000234584 | 0,00103983 |
| ENSG00000131015.5 | ULBP2 | protein_coding | 1729,940911 | 0,23710622 | 0,074190738 | 3,195900534 | 0,001393952 | 0,005230281 |
| ENSG00000131018.23 | SYNE1 | protein_coding | 7666,983134 | 0,145982561 | 0,056437221 | 2,586636214 | 0,009691785 | 0,029143045 |
| ENSG00000131019.11 | ULBP3 | protein_coding | 824,5029953 | 0,289395742 | 0,087641644 | 3,302034608 | 0,000959862 | 0,003737983 |
| ENSG00000131023.13 | LATS1 | protein_coding | 8672,880166 | 0,415322421 | 0,046766541 | 8,880759894 | 6,64E-19 | 1,27E-17 |
| ENSG00000131043.12 | AAR2 | protein_coding | 6461,415445 | 0,132867112 | 0,048534269 | 2,737593757 | 0,006189048 | 0,019701495 |
| ENSG00000131069.20 | ACSS2 | protein_coding | 4154,939872 | 0,213202393 | 0,081143609 | 2,627469925 | 0,008602245 | 0,026300995 |
| ENSG00000131115.16 | ZNF227 | protein_coding | 5418,541558 | 0,304809743 | 0,053381189 | 5,7100591 | 1,13E-08 | 9,33E-08 |
| ENSG00000131116.12 | ZNF428 | protein_coding | 2521,172279 | -0,206998314 | 0,062720632 | -3,300322516 | 0,000965738 | 0,003758913 |
| ENSG00000131143.8 | COX4I1 | protein_coding | 18792,7648 | -0,280929638 | 0,047607633 | -5,900936959 | 3,61E-09 | 3,16E-08 |

| | | | | | | | | |
|---|---|---|---|---|---|---|---|---|
| ENSG00000131149.19 | GSE1 | protein_coding | 19854,88224 | -0,289162576 | 0,050522752 | -5,723412954 | 1,04E-08 | 8,66E-08 |
| ENSG00000131153.9 | GINS2 | protein_coding | 9266,717758 | -0,432266908 | 0,04605249 | -9,386396088 | 6,21E-21 | 1,33E-19 |
| ENSG00000131171.13 | SH3BGRL | protein_coding | 781,5845619 | 1,424280464 | 0,102552563 | 13,88829711 | 7,46E-44 | 3,73E-42 |
| ENSG00000131236.17 | CAP1 | protein_coding | 23511,37759 | -0,174227505 | 0,040754578 | -4,275041309 | 1,91E-05 | 0,000102344 |
| ENSG00000131242.18 | RAB11FIP4 | protein_coding | 10584,66865 | -0,584041393 | 0,063154557 | -9,247810738 | 2,29E-20 | 4,76E-19 |
| ENSG00000131374.14 | TBC1D5 | protein_coding | 13458,70581 | -0,605628603 | 0,05907396 | -10,25204006 | 1,16E-24 | 2,97E-23 |
| ENSG00000131378.14 | RFTN1 | protein_coding | 9,083436424 | 2,923446811 | 0,932807972 | 3,134028546 | 0,001724241 | 0,006336479 |
| ENSG00000131381.12 | RBSN | protein_coding | 4262,038697 | 0,164129196 | 0,060436088 | 2,715748203 | 0,006612619 | 0,020878586 |
| ENSG00000131389.17 | SLC6A6 | protein_coding | 26270,58494 | 0,230337513 | 0,042676822 | 5,397250818 | 6,77E-08 | 5,10E-07 |
| ENSG00000131398.14 | KCNC3 | protein_coding | 2106,645705 | 0,268632735 | 0,093689699 | 2,867260101 | 0,004140426 | 0,013819224 |
| ENSG00000131437.15 | KIF3A | protein_coding | 11122,42044 | -0,183704172 | 0,045132965 | -4,070288122 | 4,70E-05 | 0,000236287 |
| ENSG00000131462.8 | TUBG1 | protein_coding | 6600,113158 | -0,314669246 | 0,062836198 | -5,007770274 | 5,51E-07 | 3,72E-06 |
| ENSG00000131469.14 | RPL27 | protein_coding | 25021,49721 | -0,490052307 | 0,045785306 | -10,70326588 | 9,83E-27 | 2,73E-25 |
| ENSG00000131473.17 | ACLY | protein_coding | 87019,76692 | -0,1124658 | 0,042916131 | -2,620595058 | 0,008777646 | 0,026764566 |
| ENSG00000131475.7 | VPS25 | protein_coding | 9095,011056 | -0,497859396 | 0,055796092 | -8,922836304 | 4,54E-19 | 8,79E-18 |
| ENSG00000131495.8 | NDUFA2 | protein_coding | 4735,492258 | -0,300653466 | 0,054385998 | -5,528140977 | 3,24E-08 | 2,54E-07 |
| ENSG00000131503.21 | ANKHD1 | protein_coding | 21162,39189 | 0,255973969 | 0,04166138 | 6,144154865 | 8,04E-10 | 7,61E-09 |
| ENSG00000131508.16 | UBE2D2 | protein_coding | 14088,32248 | -0,10217183 | 0,041782138 | -2,445347069 | 0,014471281 | 0,041143622 |
| ENSG00000131584.19 | ACAP3 | protein_coding | 18309,22839 | -0,257661425 | 0,064054777 | -4,02251693 | 5,76E-05 | 0,000285349 |
| ENSG00000131650.14 | KREMEN2 | protein_coding | 73,20732866 | 0,94366609 | 0,322359829 | 2,927368749 | 0,003418433 | 0,011661945 |
| ENSG00000131669.10 | NINJ1 | protein_coding | 784,073164 | 0,264322153 | 0,103896829 | 2,544082955 | 0,010956506 | 0,032360582 |
| ENSG00000131725.14 | WDR44 | protein_coding | 2273,441963 | 0,193876237 | 0,062007139 | 3,12667604 | 0,001767946 | 0,006479111 |
| ENSG00000131747.15 | TOP2A | protein_coding | 80866,88077 | -0,879254458 | 0,04850863 | -18,12573275 | 2,00E-73 | 1,71E-71 |
| ENSG00000131759.18 | RARA | protein_coding | 436,3761445 | 0,303397297 | 0,11813883 | 2,568142056 | 0,010224525 | 0,030498784 |
| ENSG00000131773.14 | KHDRBS3 | protein_coding | 3908,684566 | 0,14216871 | 0,059903566 | 2,373292929 | 0,017630276 | 0,048727882 |
| ENSG00000131778.19 | CHD1L | protein_coding | 4235,253298 | 0,161867042 | 0,05649847 | 2,864980983 | 0,004170343 | 0,013910825 |
| ENSG00000131791.8 | PRKAB2 | protein_coding | 4910,989377 | -0,243020687 | 0,06087867 | -3,99188559 | 6,55E-05 | 0,000322224 |
| ENSG00000131797.12 | CLUHP3 | transcribed_unproc | 2428,569797 | 0,261995641 | 0,072535795 | 3,611949688 | 0,000303904 | 0,001318028 |
| ENSG00000131844.16 | MCCC2 | protein_coding | 6984,972624 | 0,152756694 | 0,057911459 | 2,637762827 | 0,008345493 | 0,025599441 |
| ENSG00000131845.14 | AC005261.1 | protein_coding | 2999,643519 | 0,199172202 | 0,057716827 | 3,450851569 | 0,000558821 | 0,00229409 |
| ENSG00000131849.12 | ZNF132 | protein_coding | 1322,752536 | 0,18705263 | 0,072056632 | 2,595911364 | 0,00943404 | 0,028502397 |
| ENSG00000131871.15 | SELENOS | protein_coding | 4638,988175 | 0,261827592 | 0,053695304 | 4,876172963 | 1,08E-06 | 6,99E-06 |
| ENSG00000131876.17 | SNRPA1 | protein_coding | 4831,843243 | -0,181410236 | 0,056275568 | -3,223605619 | 0,001265876 | 0,004789646 |
| ENSG00000131899.11 | LLGL1 | protein_coding | 11668,39729 | -0,244601299 | 0,060576718 | -4,037876374 | 5,39E-05 | 0,000268957 |
| ENSG00000131910.5 | NR0B2 | protein_coding | 1196,0656 | 0,41027328 | 0,081108205 | 5,058344948 | 4,23E-07 | 2,90E-06 |
| ENSG00000131941.8 | RHPN2 | protein_coding | 1394,242825 | 0,21333796 | 0,087180283 | 2,447089557 | 0,014401505 | 0,040986658 |

| | | | | | | | | |
|---|---|---|---|---|---|---|---|---|
| ENSG00000131979.19 | GCH1 | protein_coding | 16213,35724 | 0,298256267 | 0,051995873 | 5,73615269 | 9,69E-09 | 8,06E-08 |
| ENSG00000132109.10 | TRIM21 | protein_coding | 4928,337002 | 3,704755983 | 0,061127694 | 60,60683364 | 0 | 0 |
| ENSG00000132122.12 | SPATA6 | protein_coding | 915,9503274 | -0,301609784 | 0,084621096 | -3,564238688 | 0,000364914 | 0,001558604 |
| ENSG00000132128.16 | LRRC41 | protein_coding | 6841,655447 | 0,290440386 | 0,0481883 | 6,027197216 | 1,67E-09 | 1,52E-08 |
| ENSG00000132155.11 | RAF1 | protein_coding | 8382,597941 | 0,120564227 | 0,049196082 | 2,450687579 | 0,014258365 | 0,040671846 |
| ENSG00000132199.20 | ENOSF1 | protein_coding | 3474,351122 | -0,415698087 | 0,065509794 | -6,34558687 | 2,22E-10 | 2,23E-09 |
| ENSG00000132256.18 | TRIM5 | protein_coding | 1506,54201 | 1,847101545 | 0,099410765 | 18,58049829 | 4,62E-77 | 4,21E-75 |
| ENSG00000132274.16 | TRIM22 | protein_coding | 951,4561181 | 4,522472391 | 0,337751513 | 13,38993971 | 6,92E-41 | 3,17E-39 |
| ENSG00000132294.15 | EFR3A | protein_coding | 18425,30884 | 0,474780047 | 0,043719296 | 10,85973691 | 1,79E-27 | 5,12E-26 |
| ENSG00000132305.20 | IMMT | protein_coding | 12579,44166 | -0,161498555 | 0,052551154 | -3,073168561 | 0,002117988 | 0,007602264 |
| ENSG00000132326.12 | PER2 | protein_coding | 2478,114253 | 0,244840194 | 0,064830743 | 3,776606342 | 0,00015898 | 0,000727628 |
| ENSG00000132329.11 | RAMP1 | protein_coding | 2696,166981 | 0,270136398 | 0,068490189 | 3,944161986 | 8,01E-05 | 0,000388005 |
| ENSG00000132330.17 | SCLY | protein_coding | 2048,044551 | 0,774409775 | 0,094140903 | 8,226071277 | 1,93E-16 | 3,20E-15 |
| ENSG00000132334.16 | PTPRE | protein_coding | 1326,059227 | -0,462588339 | 0,073637404 | -6,281975121 | 3,34E-10 | 3,31E-09 |
| ENSG00000132341.12 | RAN | protein_coding | 57150,44982 | -0,271188819 | 0,04425778 | -6,127483571 | 8,93E-10 | 8,38E-09 |
| ENSG00000132357.14 | CARD6 | protein_coding | 96,98389376 | 0,866488307 | 0,235983659 | 3,671814864 | 0,000240834 | 0,001065017 |
| ENSG00000132383.11 | RPA1 | protein_coding | 11748,80296 | -0,267885953 | 0,054461764 | -4,918789473 | 8,71E-07 | 5,70E-06 |
| ENSG00000132405.19 | TBC1D14 | protein_coding | 8643,509495 | -0,703420374 | 0,045631661 | -15,41518243 | 1,29E-53 | 7,98E-52 |
| ENSG00000132423.12 | COQ3 | protein_coding | 844,1042617 | -0,22962946 | 0,094167349 | -2,438525262 | 0,014747328 | 0,041817536 |
| ENSG00000132436.11 | FIGNL1 | protein_coding | 5055,482527 | -0,138742533 | 0,05517394 | -2,514638842 | 0,011915438 | 0,034755032 |
| ENSG00000132437.18 | DDC | protein_coding | 17778,34976 | -0,471000281 | 0,050937809 | -9,24657525 | 2,32E-20 | 4,81E-19 |
| ENSG00000132463.14 | GRSF1 | protein_coding | 29045,91411 | 0,167806277 | 0,041015054 | 4,091333799 | 4,29E-05 | 0,000217631 |
| ENSG00000132466.18 | ANKRD17 | protein_coding | 27944,28097 | 0,243924239 | 0,040573638 | 6,011889772 | 1,83E-09 | 1,66E-08 |
| ENSG00000132478.10 | UNK | protein_coding | 3520,00406 | -0,204251562 | 0,053835429 | -3,793998962 | 0,00014824 | 0,000683758 |
| ENSG00000132507.18 | EIF5A | protein_coding | 21523,98752 | -0,113154903 | 0,042525906 | -2,660846344 | 0,007794452 | 0,024109567 |
| ENSG00000132522.16 | GPS2 | protein_coding | 4856,696669 | -0,144044771 | 0,048458652 | -2,972529476 | 0,002953568 | 0,010213838 |
| ENSG00000132530.17 | XAF1 | protein_coding | 7693,389562 | 9,795567955 | 0,297447782 | 32,93205918 | 7,64E-238 | 1,74E-235 |
| ENSG00000132541.11 | RIDA | protein_coding | 4254,163354 | -0,173013477 | 0,05262692 | -3,287547087 | 0,001010643 | 0,003912721 |
| ENSG00000132554.20 | RGS22 | protein_coding | 850,0322301 | 2,513441561 | 0,273519114 | 9,189272083 | 3,96E-20 | 8,07E-19 |
| ENSG00000132561.14 | MATN2 | protein_coding | 9484,408046 | -0,158985017 | 0,044779432 | -3,550402722 | 0,000384642 | 0,001633877 |
| ENSG00000132563.16 | REEP2 | protein_coding | 965,2916585 | -0,301033627 | 0,095616375 | -3,148348052 | 0,001641961 | 0,0060648 |
| ENSG00000132581.9 | SDF2 | protein_coding | 7084,035014 | 0,152378487 | 0,050677326 | 3,006837579 | 0,002639808 | 0,009244045 |
| ENSG00000132600.16 | PRMT7 | protein_coding | 2607,989941 | 0,386049915 | 0,064664124 | 5,9700788 | 2,37E-09 | 2,12E-08 |
| ENSG00000132646.11 | PCNA | protein_coding | 47722,75155 | -0,248431318 | 0,039587752 | -6,275459151 | 3,49E-10 | 3,44E-09 |
| ENSG00000132664.12 | POLR3F | protein_coding | 2750,351721 | -0,1701105 | 0,060092631 | -2,830804647 | 0,004643107 | 0,015270558 |
| ENSG00000132669.13 | RIN2 | protein_coding | 4957,979305 | 0,187625854 | 0,054295076 | 3,455669779 | 0,000548927 | 0,002255944 |

| | | | | | | | | |
|---|---|---|---|---|---|---|---|---|
| ENSG00000132677.13 | RHBG | protein_coding | 131,0849964 | 0,574386056 | 0,205815371 | 2,790783077 | 0,00525807 | 0,017053875 |
| ENSG00000132694.18 | ARHGEF11 | protein_coding | 6914,732166 | 0,358945182 | 0,061607741 | 5,826299999 | 5,67E-09 | 4,87E-08 |
| ENSG00000132716.18 | DCAF8 | protein_coding | 12068,12075 | 0,13436598 | 0,051910046 | 2,588438847 | 0,009641206 | 0,029024196 |
| ENSG00000132740.8 | IGHMBP2 | protein_coding | 1910,727377 | 0,232944991 | 0,088090534 | 2,644381645 | 0,008184032 | 0,025165899 |
| ENSG00000132744.8 | ACY3 | protein_coding | 300,8082558 | 1,757134191 | 0,149042385 | 11,78949325 | 4,42E-32 | 1,56E-30 |
| ENSG00000132792.19 | CTNNBL1 | protein_coding | 9183,822565 | 0,27399294 | 0,046585669 | 5,881485562 | 4,07E-09 | 3,55E-08 |
| ENSG00000132793.11 | LPIN3 | protein_coding | 5517,918745 | 0,226755809 | 0,056971429 | 3,980167108 | 6,89E-05 | 0,00033735 |
| ENSG00000132801.7 | ZSWIM3 | protein_coding | 1529,026924 | 0,161699137 | 0,067988714 | 2,378323217 | 0,017391574 | 0,048186477 |
| ENSG00000132819.17 | RBM38 | protein_coding | 1112,327566 | -0,210968797 | 0,082956105 | -2,543137686 | 0,010986193 | 0,032435495 |
| ENSG00000132824.14 | SERINC3 | protein_coding | 24703,39955 | -0,156815637 | 0,047192577 | -3,322887793 | 0,000890907 | 0,003491197 |
| ENSG00000132825.7 | PPP1R3D | protein_coding | 2708,500741 | 0,584139628 | 0,057190876 | 10,21386053 | 1,72E-24 | 4,36E-23 |
| ENSG00000132837.14 | DMGDH | protein_coding | 394,7158206 | -0,527806272 | 0,135595409 | -3,892508417 | 9,92E-05 | 0,000472232 |
| ENSG00000132840.10 | BHMT2 | protein_coding | 375,9143551 | -0,695814926 | 0,139626906 | -4,983387113 | 6,25E-07 | 4,18E-06 |
| ENSG00000132842.13 | AP3B1 | protein_coding | 17816,00714 | 0,339364954 | 0,0477936 | 7,100635988 | 1,24E-12 | 1,56E-11 |
| ENSG00000132846.6 | ZBED3 | protein_coding | 4040,659994 | 0,175941924 | 0,056364008 | 3,12152967 | 0,001799141 | 0,006583786 |
| ENSG00000132855.4 | ANGPTL3 | protein_coding | 221,2195691 | -0,781309635 | 0,197279854 | -3,960412688 | 7,48E-05 | 0,000364294 |
| ENSG00000132879.14 | FBXO44 | protein_coding | 1417,119385 | 0,272696666 | 0,0803878 | 3,392264316 | 0,000693175 | 0,002786194 |
| ENSG00000132932.17 | ATP8A2 | protein_coding | 305,8651459 | -0,660561866 | 0,214438053 | -3,080432121 | 0,002067005 | 0,007440622 |
| ENSG00000132950.19 | ZMYM5 | protein_coding | 2536,070885 | 0,306531215 | 0,063298285 | 4,842646452 | 1,28E-06 | 8,20E-06 |
| ENSG00000132952.12 | USPL1 | protein_coding | 4702,756486 | 0,253480164 | 0,052902427 | 4,791465741 | 1,66E-06 | 1,04E-05 |
| ENSG00000132953.16 | XPO4 | protein_coding | 6349,973041 | -0,180073888 | 0,05401104 | -3,334020019 | 0,000856005 | 0,003369088 |
| ENSG00000132965.9 | ALOX5AP | protein_coding | 49,60128889 | -0,953668701 | 0,393046722 | -2,426349458 | 0,015251576 | 0,043041341 |
| ENSG00000132967.9 | HMGB1P5 | transcribed_process | 6705,993629 | -0,345698649 | 0,048706026 | -7,097656693 | 1,27E-12 | 1,59E-11 |
| ENSG00000132970.13 | WASF3 | protein_coding | 6595,095314 | 0,178538223 | 0,048463963 | 3,683937729 | 0,000229658 | 0,001020004 |
| ENSG00000133028.12 | SCO1 | protein_coding | 5171,64246 | 0,228519175 | 0,056731 | 4,028118205 | 5,62E-05 | 0,00027956 |
| ENSG00000133065.11 | SLC41A1 | protein_coding | 3182,105234 | 0,315887592 | 0,067700754 | 4,665939022 | 3,07E-06 | 1,86E-05 |
| ENSG00000133069.17 | TMCC2 | protein_coding | 1637,09886 | -0,295227595 | 0,068078964 | -4,33654656 | 1,45E-05 | 7,89E-05 |
| ENSG00000133083.14 | DCLK1 | protein_coding | 142,2318928 | -1,192481921 | 0,250337692 | -4,763493304 | 1,90E-06 | 1,19E-05 |
| ENSG00000133101.10 | CCNA1 | protein_coding | 317,0614695 | -0,707156126 | 0,148235134 | -4,770502815 | 1,84E-06 | 1,15E-05 |
| ENSG00000133104.14 | SPART | protein_coding | 2929,69617 | 0,15332625 | 0,058174295 | 2,635635701 | 0,008397984 | 0,025742914 |
| ENSG00000133106.14 | EPSTI1 | protein_coding | 19406,05059 | 4,829209892 | 0,059034661 | 81,80295791 | 0 | 0 |
| ENSG00000133107.14 | TRPC4 | protein_coding | 424,0670046 | 0,471925448 | 0,121445812 | 3,885893129 | 0,000101954 | 0,000484563 |
| ENSG00000133112.16 | TPT1 | protein_coding | 116964,2756 | -0,764179647 | 0,039552369 | -19,3207049 | 3,60E-83 | 3,45E-81 |
| ENSG00000133114.17 | GPALPP1 | protein_coding | 5786,602071 | -0,372490722 | 0,052894167 | -7,042189083 | 1,89E-12 | 2,34E-11 |
| ENSG00000133116.8 | KL | protein_coding | 2434,141973 | -0,403803734 | 0,097074626 | -4,159724865 | 3,19E-05 | 0,000164762 |
| ENSG00000133119.13 | RFC3 | protein_coding | 9599,791512 | -0,210711823 | 0,046576659 | -4,523978945 | 6,07E-06 | 3,51E-05 |

| | | | | | | | | |
|---|---|---|---|---|---|---|---|---|
| ENSG00000133135.14 | RNF128 | protein_coding | 13840,97971 | 0,186959372 | 0,040721239 | 4,591200495 | 4,41E-06 | 2,60E-05 |
| ENSG00000133142.17 | TCEAL4 | protein_coding | 8631,417448 | 0,112085688 | 0,045829495 | 2,445710742 | 0,014456693 | 0,041112534 |
| ENSG00000133216.16 | EPHB2 | protein_coding | 2140,268253 | -0,234136981 | 0,074216913 | -3,154765839 | 0,00160627 | 0,00594761 |
| ENSG00000133243.9 | BTBD2 | protein_coding | 2497,067607 | 0,180175568 | 0,063778825 | 2,825006062 | 0,004727972 | 0,015515662 |
| ENSG00000133313.15 | CNDP2 | protein_coding | 12573,90228 | 0,345662042 | 0,046632772 | 7,412427533 | 1,24E-13 | 1,69E-12 |
| ENSG00000133315.11 | MACROD1 | protein_coding | 236,2438483 | -0,449771869 | 0,154202242 | -2,916766072 | 0,003536809 | 0,012009183 |
| ENSG00000133318.13 | RTN3 | protein_coding | 47387,65569 | -0,283728768 | 0,043207145 | -6,566709425 | 5,14E-11 | 5,51E-10 |
| ENSG00000133321.11 | PLAAT4 | protein_coding | 1416,001229 | 7,338373188 | 0,386854878 | 18,96931797 | 3,06E-80 | 2,82E-78 |
| ENSG00000133393.13 | FOPNL | protein_coding | 6082,480832 | -0,502259838 | 0,074321172 | -6,757964462 | 1,40E-11 | 1,58E-10 |
| ENSG00000133398.4 | MED10 | protein_coding | 3787,081508 | 0,137652454 | 0,052675558 | 2,613213033 | 0,008969537 | 0,027272095 |
| ENSG00000133422.13 | MORC2 | protein_coding | 5903,392326 | 0,203628277 | 0,066507284 | 3,061744009 | 0,002200516 | 0,007857139 |
| ENSG00000133424.20 | LARGE1 | protein_coding | 5316,586402 | 0,249620842 | 0,053217542 | 4,690574411 | 2,72E-06 | 1,66E-05 |
| ENSG00000133433.10 | GSTT2B | protein_coding | 1649,899959 | 0,190683957 | 0,073584777 | 2,591350626 | 0,009560003 | 0,028826004 |
| ENSG00000133466.14 | C1QTNF6 | protein_coding | 3260,216361 | -0,384343487 | 0,062967245 | -6,103863769 | 1,04E-09 | 9,65E-09 |
| ENSG00000133477.17 | FAM83F | protein_coding | 1841,950012 | 0,493873949 | 0,075183261 | 6,568934911 | 5,07E-11 | 5,43E-10 |
| ENSG00000133597.11 | ADCK2 | protein_coding | 3860,128166 | -0,179453598 | 0,057544685 | -3,118508631 | 0,001817688 | 0,006640861 |
| ENSG00000133606.11 | MKRN1 | protein_coding | 13919,11593 | 0,19698342 | 0,051933356 | 3,793003823 | 0,000148836 | 0,000686149 |
| ENSG00000133627.18 | ACTR3B | protein_coding | 2957,680513 | -0,29123794 | 0,066381951 | -4,387306124 | 1,15E-05 | 6,37E-05 |
| ENSG00000133639.5 | BTG1 | protein_coding | 36866,12849 | 0,381499165 | 0,041475758 | 9,198124024 | 3,64E-20 | 7,45E-19 |
| ENSG00000133640.20 | LRRIQ1 | protein_coding | 263,5187492 | -0,45996927 | 0,177542792 | -2,590751588 | 0,009576659 | 0,028856897 |
| ENSG00000133657.15 | ATP13A3 | protein_coding | 46460,07226 | -0,430630653 | 0,045558262 | -9,452306497 | 3,31E-21 | 7,19E-20 |
| ENSG00000133665.13 | DYDC2 | protein_coding | 105,3808772 | -0,649167451 | 0,262418198 | -2,473789754 | 0,013368835 | 0,03842665 |
| ENSG00000133687.16 | TMTC1 | protein_coding | 28741,46746 | -0,157325913 | 0,053930798 | -2,917181242 | 0,003532105 | 0,011998814 |
| ENSG00000133703.11 | KRAS | protein_coding | 8122,655513 | -0,147434677 | 0,053938869 | -2,733366101 | 0,006269061 | 0,019930211 |
| ENSG00000133704.10 | IPO8 | protein_coding | 14328,88377 | -0,324118736 | 0,048375747 | -6,700025449 | 2,08E-11 | 2,32E-10 |
| ENSG00000133710.15 | SPINK5 | protein_coding | 184,9262603 | 0,512400659 | 0,180733548 | 2,835116469 | 0,004580898 | 0,015092423 |
| ENSG00000133731.10 | IMPA1 | protein_coding | 6026,39059 | 0,381757893 | 0,053679332 | 7,111822722 | 1,15E-12 | 1,44E-11 |
| ENSG00000133739.16 | LRRCC1 | protein_coding | 3131,748792 | 0,170825563 | 0,057475396 | 2,972151134 | 0,002957211 | 0,010221722 |
| ENSG00000133789.15 | SWAP70 | protein_coding | 4623,420044 | 0,136472827 | 0,051727798 | 2,638287946 | 0,00833258 | 0,025566798 |
| ENSG00000133812.15 | SBF2 | protein_coding | 8094,88352 | 0,401452158 | 0,047503342 | 8,451029818 | 2,89E-17 | 5,00E-16 |
| ENSG00000133818.14 | RRAS2 | protein_coding | 1342,245811 | 0,266774436 | 0,084948624 | 3,140420926 | 0,001687052 | 0,006210969 |
| ENSG00000133835.16 | HSD17B4 | protein_coding | 14364,1469 | -0,291469218 | 0,055743734 | -5,228735036 | 1,71E-07 | 1,23E-06 |
| ENSG00000133874.2 | RNF122 | protein_coding | 258,8493796 | 0,591287546 | 0,154948382 | 3,816029158 | 0,000135616 | 0,000629275 |
| ENSG00000133884.10 | DPF2 | protein_coding | 6333,628812 | 0,289235748 | 0,055274463 | 5,232719313 | 1,67E-07 | 1,20E-06 |
| ENSG00000133935.6 | ERG28 | protein_coding | 6485,020815 | 0,163575196 | 0,046165449 | 3,543238504 | 0,000395245 | 0,001672602 |
| ENSG00000133943.20 | DGLUCY | protein_coding | 2858,130958 | 1,434644202 | 0,061667052 | 23,26435525 | 1,02E-119 | 1,37E-117 |

| | | | | | | | | |
|---|---|---|---|---|---|---|---|---|
| ENSG00000133958.13 | UNC79 | protein_coding | 7139,609134 | 0,288080361 | 0,046863683 | 6,147198432 | 7,89E-10 | 7,47E-09 |
| ENSG00000133961.20 | NUMB | protein_coding | 7470,787273 | 0,340264912 | 0,04824216 | 7,053268585 | 1,75E-12 | 2,17E-11 |
| ENSG00000133983.16 | COX16 | protein_coding | 7258,906803 | -0,289904579 | 0,054008163 | -5,367791914 | 7,97E-08 | 5,94E-07 |
| ENSG00000134030.14 | CTIF | protein_coding | 3123,266987 | 0,171319585 | 0,064875664 | 2,640737291 | 0,008272584 | 0,025410425 |
| ENSG00000134042.13 | MRO | protein_coding | 237,6789502 | 0,980515073 | 0,159528515 | 6,14633111 | 7,93E-10 | 7,51E-09 |
| ENSG00000134057.15 | CCNB1 | protein_coding | 8607,977154 | -0,963277841 | 0,051963626 | -18,53754079 | 1,03E-76 | 9,26E-75 |
| ENSG00000134070.5 | IRAK2 | protein_coding | 529,43313 | 1,313436924 | 0,116712761 | 11,25358453 | 2,22E-29 | 6,81E-28 |
| ENSG00000134072.11 | CAMK1 | protein_coding | 427,350525 | -0,47560233 | 0,120150532 | -3,958387223 | 7,55E-05 | 0,000366841 |
| ENSG00000134109.11 | EDEM1 | protein_coding | 4958,032118 | 0,420288114 | 0,057649644 | 7,290385273 | 3,09E-13 | 4,07E-12 |
| ENSG00000134121.10 | CHL1 | protein_coding | 40,41580694 | 1,482074602 | 0,430987725 | 3,438786105 | 0,000584329 | 0,002389684 |
| ENSG00000134138.20 | MEIS2 | protein_coding | 71103,57917 | -0,359297491 | 0,043505091 | -8,258745837 | 1,47E-16 | 2,45E-15 |
| ENSG00000134153.10 | EMC7 | protein_coding | 10182,34641 | 0,405320353 | 0,04787031 | 8,467050854 | 2,52E-17 | 4,38E-16 |
| ENSG00000134184.12 | GSTM1 | protein_coding | 1655,222075 | -0,310737144 | 0,071665368 | -4,335945705 | 1,45E-05 | 7,91E-05 |
| ENSG00000134198.10 | TSPAN2 | protein_coding | 10319,54441 | 0,304387927 | 0,047689615 | 6,382687873 | 1,74E-10 | 1,77E-09 |
| ENSG00000134202.10 | GSTM3 | protein_coding | 14022,63258 | -0,165254514 | 0,043922401 | -3,762419911 | 0,000168277 | 0,000765821 |
| ENSG00000134222.16 | PSRC1 | protein_coding | 1795,86957 | -0,532499519 | 0,069359559 | -7,67737748 | 1,62E-14 | 2,35E-13 |
| ENSG00000134240.11 | HMGCS2 | protein_coding | 298,5230399 | -0,43044535 | 0,164010965 | -2,624491306 | 0,008677851 | 0,02651775 |
| ENSG00000134243.12 | SORT1 | protein_coding | 25180,61788 | -0,486913527 | 0,045739619 | -10,64533409 | 1,83E-26 | 5,06E-25 |
| ENSG00000134248.13 | LAMTOR5 | protein_coding | 4966,397481 | -0,137717055 | 0,053230089 | -2,587203195 | 0,009675851 | 0,029101248 |
| ENSG00000134258.17 | VTCN1 | protein_coding | 16,81193298 | -2,462740638 | 0,722363568 | -3,409281345 | 0,000651343 | 0,0026339 |
| ENSG00000134265.13 | NAPG | protein_coding | 4788,898309 | 0,384512779 | 0,050110105 | 7,673358017 | 1,68E-14 | 2,42E-13 |
| ENSG00000134283.17 | PPHLN1 | protein_coding | 5839,923834 | -0,155355306 | 0,06077619 | -2,556186966 | 0,010582624 | 0,031404652 |
| ENSG00000134285.11 | FKBP11 | protein_coding | 1716,151209 | 1,824730072 | 0,090733007 | 20,11098416 | 5,91E-90 | 6,19E-88 |
| ENSG00000134287.10 | ARF3 | protein_coding | 28595,24465 | 0,178482231 | 0,045482357 | 3,924208081 | 8,70E-05 | 0,000419048 |
| ENSG00000134291.12 | TMEM106C | protein_coding | 21296,68764 | -0,707206947 | 0,050208149 | -14,08550138 | 4,66E-45 | 2,40E-43 |
| ENSG00000134297.6 | PLEKHA8P1 | transcribed_process | 486,9877102 | 0,456956509 | 0,123618371 | 3,696509709 | 0,000218584 | 0,0009737 |
| ENSG00000134308.14 | YWHAQ | protein_coding | 91884,64956 | -0,455630057 | 0,040375061 | -11,28493807 | 1,56E-29 | 4,79E-28 |
| ENSG00000134317.18 | GRHL1 | protein_coding | 4486,782073 | 0,212676401 | 0,055636382 | 3,822613801 | 0,000132045 | 0,00061447 |
| ENSG00000134318.14 | ROCK2 | protein_coding | 19557,59923 | 0,187210164 | 0,040455109 | 4,62760252 | 3,70E-06 | 2,21E-05 |
| ENSG00000134321.12 | RSAD2 | protein_coding | 140202,6704 | 8,769268345 | 0,063703822 | 137,656864 | 0 | 0 |
| ENSG00000134324.11 | LPIN1 | protein_coding | 17403,79571 | 0,241750406 | 0,057708648 | 4,189153867 | 2,80E-05 | 0,000145789 |
| ENSG00000134326.11 | CMPK2 | protein_coding | 45648,46278 | 4,877817571 | 0,052533096 | 92,85227753 | 0 | 0 |
| ENSG00000134330.18 | IAH1 | protein_coding | 6133,476552 | -0,144809887 | 0,049098643 | -2,949366417 | 0,003184262 | 0,010927665 |
| ENSG00000134333.14 | LDHA | protein_coding | 79782,26406 | -0,367768076 | 0,067443032 | -5,453018094 | 4,95E-08 | 3,80E-07 |
| ENSG00000134339.8 | SAA2 | protein_coding | 42,00534553 | 3,740849283 | 0,483388926 | 7,738798063 | 1,00E-14 | 1,48E-13 |
| ENSG00000134352.20 | IL6ST | protein_coding | 4865,466653 | 0,353872481 | 0,064175077 | 5,514173043 | 3,50E-08 | 2,74E-07 |

| | | | | | | | | |
|---|---|---|---|---|---|---|---|---|
| ENSG00000134419.15 | RPS15A | protein_coding | 50881,20833 | -0,565953478 | 0,043410885 | -13,03713292 | 7,52E-39 | 3,26E-37 |
| ENSG00000134444.14 | RELCH | protein_coding | 6958,110485 | 0,374181866 | 0,049730845 | 7,524140521 | 5,31E-14 | 7,45E-13 |
| ENSG00000134480.15 | CCNH | protein_coding | 3551,311243 | -0,191448851 | 0,059285662 | -3,229260571 | 0,001241108 | 0,004706215 |
| ENSG00000134504.13 | KCTD1 | protein_coding | 1818,926467 | 0,310813762 | 0,076884843 | 4,042588241 | 5,29E-05 | 0,000264017 |
| ENSG00000134569.10 | LRP4 | protein_coding | 9273,855595 | 1,2647201 | 0,055199348 | 22,9118668 | 3,54E-116 | 4,71E-114 |
| ENSG00000134574.11 | DDB2 | protein_coding | 2493,268913 | -0,161512888 | 0,062810038 | -2,571450237 | 0,010127357 | 0,030241002 |
| ENSG00000134575.11 | ACP2 | protein_coding | 9351,241901 | 0,827456646 | 0,054272138 | 15,24643551 | 1,74E-52 | 1,06E-50 |
| ENSG00000134590.13 | RTL8C | protein_coding | 3658,544013 | -0,140466856 | 0,056267531 | -2,496410503 | 0,012545732 | 0,036371959 |
| ENSG00000134627.12 | PIWIL4 | protein_coding | 295,9211481 | 1,064412109 | 0,153734341 | 6,923710746 | 4,40E-12 | 5,24E-11 |
| ENSG00000134684.10 | YARS | protein_coding | 7165,351612 | 0,516460604 | 0,054688435 | 9,443689652 | 3,60E-21 | 7,78E-20 |
| ENSG00000134686.18 | PHC2 | protein_coding | 17921,55788 | 0,318408852 | 0,045540505 | 6,991772492 | 2,71E-12 | 3,30E-11 |
| ENSG00000134690.11 | CDCA8 | protein_coding | 8658,222222 | -0,606053896 | 0,048091994 | -12,60197053 | 2,06E-36 | 8,37E-35 |
| ENSG00000134697.13 | GNL2 | protein_coding | 7488,887796 | 0,150016077 | 0,046495414 | 3,226470385 | 0,001253272 | 0,004749941 |
| ENSG00000134698.11 | AGO4 | protein_coding | 2815,851951 | 0,345564232 | 0,07746338 | 4,461001227 | 8,16E-06 | 4,62E-05 |
| ENSG00000134709.11 | HOOK1 | protein_coding | 18010,60348 | -0,25388762 | 0,048866039 | -5,195584196 | 2,04E-07 | 1,45E-06 |
| ENSG00000134716.11 | CYP2J2 | protein_coding | 1863,026223 | 3,065242789 | 0,077853751 | 39,37180618 | 0 | 0 |
| ENSG00000134755.16 | DSC2 | protein_coding | 4072,869356 | 0,156724318 | 0,055903809 | 2,803464025 | 0,005055687 | 0,016475816 |
| ENSG00000134769.21 | DTNA | protein_coding | 2548,549404 | 0,702821139 | 0,099407475 | 7,07010355 | 1,55E-12 | 1,93E-11 |
| ENSG00000134775.15 | FHOD3 | protein_coding | 10027,26489 | 0,156475872 | 0,054570156 | 2,867425768 | 0,004138259 | 0,013816088 |
| ENSG00000134779.15 | TPGS2 | protein_coding | 11958,97571 | -0,349477315 | 0,046185183 | -7,566870895 | 3,82E-14 | 5,41E-13 |
| ENSG00000134780.10 | DAGLA | protein_coding | 3039,287671 | 0,258349878 | 0,058548682 | 4,412565219 | 1,02E-05 | 5,72E-05 |
| ENSG00000134809.9 | TIMM10 | protein_coding | 2018,670306 | -0,281562836 | 0,072502867 | -3,883471773 | 0,000102975 | 0,000489106 |
| ENSG00000134815.19 | DHX34 | protein_coding | 2233,235368 | 0,265226484 | 0,065119733 | 4,072904965 | 4,64E-05 | 0,000234013 |
| ENSG00000134824.14 | FADS2 | protein_coding | 83053,86249 | -0,171338222 | 0,043545315 | -3,934710817 | 8,33E-05 | 0,000402691 |
| ENSG00000134825.15 | TMEM258 | protein_coding | 8214,757918 | -0,254842001 | 0,047193677 | -5,399918315 | 6,67E-08 | 5,03E-07 |
| ENSG00000134852.15 | CLOCK | protein_coding | 6506,671937 | 0,222497497 | 0,078971159 | 2,8174526 | 0,004840626 | 0,015848385 |
| ENSG00000134871.18 | COL4A2 | protein_coding | 4825,652925 | -0,187863259 | 0,070986736 | -2,646455817 | 0,008134013 | 0,025036032 |
| ENSG00000134873.10 | CLDN10 | protein_coding | 34,35392713 | 1,405124113 | 0,470574038 | 2,985978828 | 0,002826722 | 0,009826516 |
| ENSG00000134874.17 | DZIP1 | protein_coding | 5231,753248 | 0,451164211 | 0,051080051 | 8,832493401 | 1,02E-18 | 1,94E-17 |
| ENSG00000134884.15 | ARGLU1 | protein_coding | 12599,75356 | -0,148380468 | 0,054260177 | -2,734610842 | 0,006245406 | 0,019866853 |
| ENSG00000134897.14 | BIVM | protein_coding | 8150,664057 | -0,107844583 | 0,045065299 | -2,393073754 | 0,016707882 | 0,04656156 |
| ENSG00000134905.17 | CARS2 | protein_coding | 2335,290522 | -0,29743876 | 0,064598211 | -4,604442667 | 4,14E-06 | 2,45E-05 |
| ENSG00000134917.10 | ADAMTS8 | protein_coding | 96,29610458 | -0,857974229 | 0,260437169 | -3,294361675 | 0,000986455 | 0,003828168 |
| ENSG00000134954.14 | ETS1 | protein_coding | 6044,740729 | -0,175539459 | 0,057558892 | -3,049736568 | 0,002290422 | 0,00814714 |
| ENSG00000134962.7 | KLB | protein_coding | 836,1566682 | 0,886616951 | 0,111132465 | 7,978019306 | 1,49E-15 | 2,31E-14 |
| ENSG00000134970.14 | TMED7 | protein_coding | 14568,56744 | 0,194173407 | 0,047656313 | 4,074453007 | 4,61E-05 | 0,000232566 |

| | | | | | | | | |
|---|---|---|---|---|---|---|---|---|
| ENSG00000134996.11 | OSTF1 | protein_coding | 4001,583995 | 0,125423335 | 0,053039391 | 2,364720509 | 0,018043682 | 0,049735917 |
| ENSG00000135047.15 | CTSL | protein_coding | 15052,45785 | 0,415193174 | 0,040569232 | 10,23418859 | 1,39E-24 | 3,55E-23 |
| ENSG00000135048.14 | CEMIP2 | protein_coding | 12927,67494 | 0,26094928 | 0,058167778 | 4,486148342 | 7,25E-06 | 4,14E-05 |
| ENSG00000135052.16 | GOLM1 | protein_coding | 39477,5207 | 0,279670483 | 0,043419546 | 6,441119424 | 1,19E-10 | 1,23E-09 |
| ENSG00000135069.14 | PSAT1 | protein_coding | 3887,120716 | 0,530424147 | 0,076697249 | 6,915817154 | 4,65E-12 | 5,53E-11 |
| ENSG00000135074.15 | ADAM19 | protein_coding | 877,9364575 | -0,329108877 | 0,112465078 | -2,926320628 | 0,003429972 | 0,011697765 |
| ENSG00000135090.14 | TAOK3 | protein_coding | 3117,882083 | 0,253809459 | 0,067499654 | 3,760159415 | 0,000169805 | 0,000772306 |
| ENSG00000135093.13 | USP30 | protein_coding | 4187,776421 | 0,276431187 | 0,059024767 | 4,683308401 | 2,82E-06 | 1,72E-05 |
| ENSG00000135108.14 | FBXO21 | protein_coding | 14548,70884 | -0,281554087 | 0,052086374 | -5,405522914 | 6,46E-08 | 4,89E-07 |
| ENSG00000135114.12 | OASL | protein_coding | 5944,199807 | 9,882103669 | 0,446143119 | 22,1500753 | 1,04E-108 | 1,30E-106 |
| ENSG00000135116.9 | HRK | protein_coding | 83,60202819 | 0,71049508 | 0,265130277 | 2,679796095 | 0,007366702 | 0,022944111 |
| ENSG00000135119.14 | RNFT2 | protein_coding | 1468,98914 | -0,214193472 | 0,084597712 | -2,531906217 | 0,011344433 | 0,033318351 |
| ENSG00000135124.14 | P2RX4 | protein_coding | 2956,603193 | 0,419947555 | 0,168063859 | 2,498738021 | 0,012463641 | 0,036175899 |
| ENSG00000135127.11 | BICDL1 | protein_coding | 6391,42138 | 0,270034221 | 0,051655059 | 5,227643296 | 1,72E-07 | 1,23E-06 |
| ENSG00000135144.7 | DTX1 | protein_coding | 2270,777981 | -0,187258295 | 0,06694134 | -2,797349085 | 0,005152383 | 0,016769085 |
| ENSG00000135148.12 | TRAFD1 | protein_coding | 11929,97899 | 0,909088553 | 0,068820477 | 13,20956486 | 7,73E-40 | 3,49E-38 |
| ENSG00000135241.17 | PNPLA8 | protein_coding | 10582,28075 | 0,176160054 | 0,043789451 | 4,022887926 | 5,75E-05 | 0,000285141 |
| ENSG00000135249.8 | RINT1 | protein_coding | 3969,303002 | 0,200679848 | 0,062759204 | 3,197616221 | 0,001385686 | 0,005204471 |
| ENSG00000135272.10 | MDFIC | protein_coding | 2551,742061 | -0,222015387 | 0,066303981 | -3,348447305 | 0,000812657 | 0,003217043 |
| ENSG00000135299.17 | ANKRD6 | protein_coding | 728,734576 | 0,630162509 | 0,12978292 | 4,855511877 | 1,20E-06 | 7,71E-06 |
| ENSG00000135315.12 | CEP162 | protein_coding | 2738,509956 | 0,367358178 | 0,062051323 | 5,920231241 | 3,21E-09 | 2,84E-08 |
| ENSG00000135316.17 | SYNCRIP | protein_coding | 30747,9555 | -0,099497726 | 0,040639481 | -2,448302069 | 0,014353127 | 0,040900687 |
| ENSG00000135318.12 | NT5E | protein_coding | 3250,388894 | 0,805384234 | 0,06632864 | 12,14233007 | 6,30E-34 | 2,31E-32 |
| ENSG00000135324.6 | MRAP2 | protein_coding | 724,3351092 | -0,74544303 | 0,119360743 | -6,245294825 | 4,23E-10 | 4,14E-09 |
| ENSG00000135346.8 | CGA | protein_coding | 10,44947146 | 2,405572432 | 0,813509525 | 2,957030442 | 0,003106174 | 0,010682488 |
| ENSG00000135365.15 | PHF21A | protein_coding | 4663,656972 | 0,520053439 | 0,064343753 | 8,082423214 | 6,35E-16 | 1,01E-14 |
| ENSG00000135372.9 | NAT10 | protein_coding | 3597,707195 | 0,190917784 | 0,061848563 | 3,086858831 | 0,002022836 | 0,00729563 |
| ENSG00000135373.12 | EHF | protein_coding | 446,0296678 | 1,458293934 | 0,11766796 | 12,39329664 | 2,84E-35 | 1,10E-33 |
| ENSG00000135378.4 | PRRG4 | protein_coding | 5602,13531 | 0,808281612 | 0,058348954 | 13,85254667 | 1,23E-43 | 6,13E-42 |
| ENSG00000135390.19 | ATP5MC2 | protein_coding | 16035,59539 | -0,533997905 | 0,043365077 | -12,31400797 | 7,61E-35 | 2,89E-33 |
| ENSG00000135406.14 | PRPH | protein_coding | 317,0937563 | 1,622972287 | 0,141498153 | 11,46991851 | 1,87E-30 | 6,06E-29 |
| ENSG00000135414.9 | GDF11 | protein_coding | 6856,835543 | -0,257964272 | 0,081214508 | -3,176332405 | 0,001491499 | 0,005573068 |
| ENSG00000135437.9 | RDH5 | protein_coding | 473,9867943 | -0,420987568 | 0,122423431 | -3,438782643 | 0,000584336 | 0,002389684 |
| ENSG00000135441.7 | BLOC1S1 | protein_coding | 4152,645096 | -0,344097506 | 0,055167921 | -6,237275239 | 4,45E-10 | 4,35E-09 |
| ENSG00000135446.17 | CDK4 | protein_coding | 15666,27356 | -0,254683569 | 0,051642089 | -4,931705429 | 8,15E-07 | 5,36E-06 |
| ENSG00000135447.17 | PPP1R1A | protein_coding | 11268,21529 | 0,488890234 | 0,051195481 | 9,549480295 | 1,30E-21 | 2,88E-20 |

| | | | | | | | | |
|---|---|---|---|---|---|---|---|---|
| ENSG00000135451.13 | TROAP | protein_coding | 2580,360605 | -0,607186374 | 0,087337996 | -6,952144569 | 3,60E-12 | 4,32E-11 |
| ENSG00000135452.10 | TSPAN31 | protein_coding | 2366,324002 | 0,441400359 | 0,071407017 | 6,181470399 | 6,35E-10 | 6,08E-09 |
| ENSG00000135469.13 | COQ10A | protein_coding | 1807,679147 | 0,18794262 | 0,073759944 | 2,548030953 | 0,010833286 | 0,032038685 |
| ENSG00000135476.11 | ESPL1 | protein_coding | 5028,373879 | -0,554642104 | 0,057101618 | -9,713246805 | 2,65E-22 | 6,08E-21 |
| ENSG00000135480.16 | KRT7 | protein_coding | 103,894589 | -0,549013126 | 0,22672062 | -2,421540335 | 0,015454886 | 0,043522335 |
| ENSG00000135486.17 | HNRNPA1 | protein_coding | 183072,916 | -0,421228011 | 0,052537418 | -8,017676305 | 1,08E-15 | 1,70E-14 |
| ENSG00000135503.13 | ACVR1B | protein_coding | 32035,40307 | 0,118610103 | 0,041165986 | 2,881264743 | 0,003960828 | 0,013286777 |
| ENSG00000135506.16 | OS9 | protein_coding | 36077,47584 | 0,46113964 | 0,057492381 | 8,02088257 | 1,05E-15 | 1,66E-14 |
| ENSG00000135525.18 | MAP7 | protein_coding | 7765,826071 | 0,17613772 | 0,046989816 | 3,748423289 | 0,00017795 | 0,000805753 |
| ENSG00000135535.17 | CD164 | protein_coding | 63492,59178 | 0,389559838 | 0,038511535 | 10,11540659 | 4,72E-24 | 1,18E-22 |
| ENSG00000135540.11 | NHSL1 | protein_coding | 4147,799534 | 0,349132904 | 0,063807528 | 5,471656973 | 4,46E-08 | 3,43E-07 |
| ENSG00000135549.15 | PKIB | protein_coding | 7344,121389 | 0,213396355 | 0,049411085 | 4,318795209 | 1,57E-05 | 8,51E-05 |
| ENSG00000135596.18 | MICAL1 | protein_coding | 2316,174709 | 0,465622971 | 0,071308344 | 6,52971226 | 6,59E-11 | 6,97E-10 |
| ENSG00000135604.9 | STX11 | protein_coding | 74,7791623 | 1,59464238 | 0,275779522 | 5,782308887 | 7,37E-09 | 6,22E-08 |
| ENSG00000135605.13 | TEC | protein_coding | 670,7566979 | 0,679802176 | 0,108519049 | 6,264358026 | 3,74E-10 | 3,67E-09 |
| ENSG00000135617.4 | PRADC1 | protein_coding | 1431,515789 | -1,027357068 | 0,089598707 | -11,46620423 | 1,95E-30 | 6,31E-29 |
| ENSG00000135625.7 | EGR4 | protein_coding | 57,50601832 | -0,841417684 | 0,314862081 | -2,672337304 | 0,007532489 | 0,023386041 |
| ENSG00000135631.16 | RAB11FIP5 | protein_coding | 2931,974146 | 0,170560518 | 0,066110765 | 2,579920523 | 0,009882306 | 0,029619283 |
| ENSG00000135655.16 | USP15 | protein_coding | 9819,759039 | 0,573343041 | 0,057112972 | 10,03875343 | 1,03E-23 | 2,52E-22 |
| ENSG00000135677.11 | GNS | protein_coding | 26923,77748 | -0,257779126 | 0,048439372 | -5,321685968 | 1,03E-07 | 7,57E-07 |
| ENSG00000135686.13 | KLHL36 | protein_coding | 4970,586501 | -0,350107462 | 0,053846464 | -6,501958302 | 7,93E-11 | 8,33E-10 |
| ENSG00000135697.10 | BCO1 | protein_coding | 1116,379286 | 2,697878247 | 0,097947786 | 27,54404534 | 5,22E-167 | 9,24E-165 |
| ENSG00000135720.13 | DYNC1LI2 | protein_coding | 28874,51304 | -0,147728267 | 0,042263182 | -3,495436472 | 0,000473287 | 0,001973184 |
| ENSG00000135736.6 | CCDC102A | protein_coding | 63,13825476 | 0,843468547 | 0,303117076 | 2,782649395 | 0,005391704 | 0,017444568 |
| ENSG00000135744.8 | AGT | protein_coding | 1365,358201 | 0,407127496 | 0,092014062 | 4,424622573 | 9,66E-06 | 5,43E-05 |
| ENSG00000135749.19 | PCNX2 | protein_coding | 2164,209413 | 0,488394085 | 0,07754619 | 6,298105506 | 3,01E-10 | 3,00E-09 |
| ENSG00000135750.14 | KCNK1 | protein_coding | 415,1958682 | 0,554255236 | 0,12301543 | 4,505574912 | 6,62E-06 | 3,80E-05 |
| ENSG00000135763.10 | URB2 | protein_coding | 2601,796602 | -0,15790105 | 0,066475017 | -2,375344256 | 0,01753259 | 0,048506282 |
| ENSG00000135776.5 | ABCB10 | protein_coding | 5745,097484 | -0,16956233 | 0,051937314 | -3,26474967 | 0,001095609 | 0,004210167 |
| ENSG00000135801.9 | TAF5L | protein_coding | 4716,427718 | -0,167081342 | 0,049083942 | -3,403991883 | 0,000664087 | 0,002680992 |
| ENSG00000135828.11 | RNASEL | protein_coding | 3385,304206 | 1,13162525 | 0,058166828 | 19,45482129 | 2,65E-84 | 2,60E-82 |
| ENSG00000135829.17 | DHX9 | protein_coding | 50576,72225 | -0,113141626 | 0,038487606 | -2,939689841 | 0,003285409 | 0,011247343 |
| ENSG00000135835.12 | KIAA1614 | protein_coding | 1139,636553 | -0,931409901 | 0,10478375 | -8,888877342 | 6,17E-19 | 1,19E-17 |
| ENSG00000135837.16 | CEP350 | protein_coding | 13480,90424 | 0,216898974 | 0,065809881 | 3,295842072 | 0,000981272 | 0,003811466 |
| ENSG00000135838.13 | NPL | protein_coding | 1032,12922 | 0,342659818 | 0,088105706 | 3,889189875 | 0,000100579 | 0,000478432 |
| ENSG00000135842.17 | FAM129A | protein_coding | 562,5363428 | -0,435564152 | 0,122945148 | -3,542751855 | 0,000395975 | 0,001675377 |

| | | | | | | | | |
|---|---|---|---|---|---|---|---|---|
| ENSG00000135862.6 | LAMC1 | protein_coding | 18584,70662 | 0,126176466 | 0,051958962 | 2,428386954 | 0,015166152 | 0,042821741 |
| ENSG00000135870.11 | RC3H1 | protein_coding | 4092,816046 | 0,238838988 | 0,058670344 | 4,070863952 | 4,68E-05 | 0,000235756 |
| ENSG00000135899.17 | SP110 | protein_coding | 7001,856112 | 12,40305233 | 0,628136732 | 19,74578415 | 8,72E-87 | 8,96E-85 |
| ENSG00000135900.4 | MRPL44 | protein_coding | 4469,136599 | 0,12554383 | 0,049869561 | 2,517444046 | 0,011820974 | 0,034533201 |
| ENSG00000135902.10 | CHRND | protein_coding | 46,15513191 | -2,064174516 | 0,356941645 | -5,782946727 | 7,34E-09 | 6,21E-08 |
| ENSG00000135905.19 | DOCK10 | protein_coding | 4436,783185 | 0,31341886 | 0,071282098 | 4,396880383 | 1,10E-05 | 6,11E-05 |
| ENSG00000135913.11 | USP37 | protein_coding | 12540,66614 | 0,297972009 | 0,041419001 | 7,194089743 | 6,29E-13 | 8,08E-12 |
| ENSG00000135916.16 | ITM2C | protein_coding | 14684,06787 | -0,192986114 | 0,044679193 | -4,319373318 | 1,56E-05 | 8,49E-05 |
| ENSG00000135924.15 | DNAJB2 | protein_coding | 6354,172024 | 0,136102948 | 0,048542976 | 2,803761932 | 0,005051019 | 0,016462985 |
| ENSG00000135932.11 | CAB39 | protein_coding | 8467,676604 | 0,119612314 | 0,05020373 | 2,382538399 | 0,017193739 | 0,047732346 |
| ENSG00000135945.10 | REV1 | protein_coding | 6332,333967 | 0,234563775 | 0,047887104 | 4,898266072 | 9,67E-07 | 6,29E-06 |
| ENSG00000135953.11 | MFSD9 | protein_coding | 3930,553347 | 0,598881547 | 0,072670965 | 8,241001693 | 1,71E-16 | 2,83E-15 |
| ENSG00000135956.8 | TMEM127 | protein_coding | 5705,475153 | 0,222688433 | 0,048135834 | 4,626250699 | 3,72E-06 | 2,22E-05 |
| ENSG00000135966.13 | TGFBRAP1 | protein_coding | 4528,862065 | 0,130991532 | 0,051692756 | 2,534040381 | 0,011275574 | 0,033153955 |
| ENSG00000135968.20 | GCC2 | protein_coding | 11656,55856 | 0,249883313 | 0,049930028 | 5,004670029 | 5,60E-07 | 3,77E-06 |
| ENSG00000135976.19 | ANKRD36 | protein_coding | 1138,806954 | 0,312373978 | 0,090232066 | 3,461895455 | 0,000536385 | 0,002208433 |
| ENSG00000135999.12 | EPC2 | protein_coding | 7003,419624 | 0,223352537 | 0,049814445 | 4,483690188 | 7,34E-06 | 4,18E-05 |
| ENSG00000136021.18 | SCYL2 | protein_coding | 10303,76732 | 0,120435722 | 0,046914785 | 2,567116574 | 0,010254813 | 0,030581026 |
| ENSG00000136026.14 | CKAP4 | protein_coding | 2951,540632 | 0,357637756 | 0,05892177 | 6,069704907 | 1,28E-09 | 1,18E-08 |
| ENSG00000136040.9 | PLXNC1 | protein_coding | 30771,12166 | 0,20658908 | 0,042604319 | 4,849017311 | 1,24E-06 | 7,96E-06 |
| ENSG00000136044.12 | APPL2 | protein_coding | 8191,71494 | -0,301232204 | 0,049491606 | -6,086531134 | 1,15E-09 | 1,07E-08 |
| ENSG00000136048.14 | DRAM1 | protein_coding | 657,6265141 | 1,394988179 | 0,122061433 | 11,42857445 | 3,01E-30 | 9,61E-29 |
| ENSG00000136051.14 | WASHC4 | protein_coding | 23054,14232 | 0,46815443 | 0,039949953 | 11,7185226 | 1,02E-31 | 3,55E-30 |
| ENSG00000136052.9 | SLC41A2 | protein_coding | 3140,396124 | -0,156375434 | 0,063646831 | -2,456924116 | 0,014013228 | 0,040058897 |
| ENSG00000136059.14 | VILL | protein_coding | 955,6347573 | 0,545365144 | 0,106166866 | 5,136867697 | 2,79E-07 | 1,95E-06 |
| ENSG00000136098.17 | NEK3 | protein_coding | 849,9845552 | 0,489721132 | 0,094378127 | 5,188926152 | 2,12E-07 | 1,50E-06 |
| ENSG00000136100.14 | VPS36 | protein_coding | 7991,81172 | -0,164508964 | 0,050257766 | -3,273304377 | 0,001062979 | 0,004097021 |
| ENSG00000136104.20 | RNASEH2B | protein_coding | 4002,997019 | -0,166618597 | 0,057055397 | -2,920295125 | 0,003497 | 0,011899317 |
| ENSG00000136108.15 | CKAP2 | protein_coding | 17110,86878 | -0,334043905 | 0,046558797 | -7,174667833 | 7,25E-13 | 9,29E-12 |
| ENSG00000136111.13 | TBC1D4 | protein_coding | 15709,23162 | -0,156343516 | 0,052231049 | -2,993306074 | 0,002759728 | 0,00961739 |
| ENSG00000136114.17 | THSD1 | protein_coding | 133,667008 | 1,387202715 | 0,232814862 | 5,958394163 | 2,55E-09 | 2,27E-08 |
| ENSG00000136122.16 | BORA | protein_coding | 1712,709287 | -0,385908704 | 0,085058532 | -4,536978209 | 5,71E-06 | 3,31E-05 |
| ENSG00000136141.15 | LRCH1 | protein_coding | 2295,065205 | 0,566024241 | 0,069333414 | 8,163801656 | 3,25E-16 | 5,28E-15 |
| ENSG00000136143.16 | SUCLA2 | protein_coding | 9362,532942 | -0,14246234 | 0,047921633 | -2,972818959 | 0,002950784 | 0,01020761 |
| ENSG00000136146.14 | MED4 | protein_coding | 10269,0915 | -0,121361184 | 0,046472292 | -2,611474062 | 0,009015283 | 0,027392683 |
| ENSG00000136147.18 | PHF11 | protein_coding | 1622,720117 | 3,759560465 | 0,11212349 | 33,53053367 | 1,73E-246 | 4,10E-244 |

| | | | | | | | | |
|---|---|---|---|---|---|---|---|---|
| ENSG00000136152.15 | COG3 | protein_coding | 6007,49047 | 0,225375129 | 0,061196698 | 3,682798875 | 0,000230687 | 0,001023765 |
| ENSG00000136153.20 | LMO7 | protein_coding | 8485,946188 | 0,265158039 | 0,055487695 | 4,778681777 | 1,76E-06 | 1,11E-05 |
| ENSG00000136156.15 | ITM2B | protein_coding | 36358,99089 | 0,322040559 | 0,043853179 | 7,343608001 | 2,08E-13 | 2,77E-12 |
| ENSG00000136159.4 | NUDT15 | protein_coding | 5549,173675 | -0,419554611 | 0,052389834 | -8,008321074 | 1,16E-15 | 1,82E-14 |
| ENSG00000136169.16 | SETDB2 | protein_coding | 2123,109134 | 0,428014333 | 0,060986341 | 7,018199931 | 2,25E-12 | 2,75E-11 |
| ENSG00000136193.17 | SCRN1 | protein_coding | 46461,2936 | -0,186031624 | 0,051777133 | -3,592930192 | 0,00032698 | 0,001410344 |
| ENSG00000136213.10 | CHST12 | protein_coding | 900,1210741 | -0,217102229 | 0,091400237 | -2,375291747 | 0,017535084 | 0,048506478 |
| ENSG00000136231.14 | IGF2BP3 | protein_coding | 7465,994751 | -0,164518638 | 0,054281246 | -3,030855978 | 0,002438615 | 0,00860762 |
| ENSG00000136237.19 | RAPGEF5 | protein_coding | 3583,114919 | 0,409315412 | 0,057091233 | 7,169496744 | 7,53E-13 | 9,62E-12 |
| ENSG00000136240.10 | KDELR2 | protein_coding | 12324,58657 | -0,175516629 | 0,051562352 | -3,403968658 | 0,000664144 | 0,002680992 |
| ENSG00000136261.15 | BZW2 | protein_coding | 2275,824733 | -0,230464783 | 0,062194421 | -3,705553935 | 0,000210929 | 0,000941838 |
| ENSG00000136295.15 | TTYH3 | protein_coding | 4693,971367 | -0,143691821 | 0,056063548 | -2,563016908 | 0,0103767 | 0,030882653 |
| ENSG00000136319.12 | TTC5 | protein_coding | 3836,079146 | -0,212572559 | 0,055728263 | -3,814447918 | 0,000136488 | 0,000633052 |
| ENSG00000136367.14 | ZFHX2 | protein_coding | 993,2129252 | 0,355668563 | 0,092613215 | 3,840365139 | 0,000122851 | 0,000575737 |
| ENSG00000136371.10 | MTHFS | protein_coding | 3513,1915 | 0,206954534 | 0,055817752 | 3,707683046 | 0,000209164 | 0,000934328 |
| ENSG00000136379.12 | ABHD17C | protein_coding | 1718,7073 | -0,292931934 | 0,071237335 | -4,112056306 | 3,92E-05 | 0,000199975 |
| ENSG00000136383.6 | ALPK3 | protein_coding | 1843,086526 | -1,153319665 | 0,108980795 | -10,58277893 | 3,58E-26 | 9,76E-25 |
| ENSG00000136425.13 | CIB2 | protein_coding | 1410,965016 | -0,369594788 | 0,084196697 | -4,389658976 | 1,14E-05 | 6,31E-05 |
| ENSG00000136436.14 | CALCOCO2 | protein_coding | 18338,40135 | 0,275820801 | 0,050826307 | 5,426733121 | 5,74E-08 | 4,37E-07 |
| ENSG00000136450.13 | SRSF1 | protein_coding | 41924,11034 | -0,107543819 | 0,037719221 | -2,851167566 | 0,004355901 | 0,01443353 |
| ENSG00000136451.9 | VEZF1 | protein_coding | 8858,702835 | 0,14266263 | 0,04667505 | 3,056507284 | 0,00223932 | 0,007985561 |
| ENSG00000136463.8 | TACO1 | protein_coding | 1384,584517 | 0,240475379 | 0,072198614 | 3,330747882 | 0,00086613 | 0,003403583 |
| ENSG00000136485.15 | DCAF7 | protein_coding | 29303,59718 | -0,461757171 | 0,046556157 | -9,918283736 | 3,47E-23 | 8,24E-22 |
| ENSG00000136518.17 | ACTL6A | protein_coding | 10255,5434 | -0,325168358 | 0,055795258 | -5,827885154 | 5,61E-09 | 4,82E-08 |
| ENSG00000136521.13 | NDUFB5 | protein_coding | 10119,15004 | -0,136955447 | 0,045532285 | -3,007875523 | 0,002630809 | 0,009222376 |
| ENSG00000136522.13 | MRPL47 | protein_coding | 5866,816926 | -0,144270345 | 0,05081688 | -2,839024068 | 0,004525174 | 0,014930687 |
| ENSG00000136527.18 | TRA2B | protein_coding | 23667,35455 | -0,156001852 | 0,03919847 | -3,979794453 | 6,90E-05 | 0,000337806 |
| ENSG00000136531.16 | SCN2A | protein_coding | 325,8209598 | 0,907515122 | 0,170041741 | 5,337013816 | 9,45E-08 | 6,97E-07 |
| ENSG00000136541.14 | ERMN | protein_coding | 1275,666045 | 0,458959053 | 0,091888526 | 4,994737374 | 5,89E-07 | 3,96E-06 |
| ENSG00000136546.16 | SCN7A | protein_coding | 191,6235099 | -1,652858083 | 0,194820111 | -8,484021857 | 2,18E-17 | 3,80E-16 |
| ENSG00000136560.13 | TANK | protein_coding | 6104,682886 | 0,590512946 | 0,064623692 | 9,137716005 | 6,38E-20 | 1,28E-18 |
| ENSG00000136603.14 | SKIL | protein_coding | 7613,684186 | -0,442726961 | 0,054961738 | -8,055184878 | 7,94E-16 | 1,26E-14 |
| ENSG00000136643.12 | RPS6KC1 | protein_coding | 6222,865589 | 0,738158766 | 0,054037912 | 13,66001654 | 1,76E-42 | 8,52E-41 |
| ENSG00000136682.14 | CBWD2 | protein_coding | 6171,282629 | -0,229904363 | 0,058563335 | -3,925738916 | 8,65E-05 | 0,000416927 |
| ENSG00000136698.8 | CFC1 | protein_coding | 6938,208903 | 0,277752516 | 0,063897833 | 4,346822129 | 1,38E-05 | 7,57E-05 |
| ENSG00000136699.19 | SMPD4 | protein_coding | 14136,39142 | -0,128493935 | 0,04630418 | -2,774996434 | 0,005520231 | 0,017809214 |

| | | | | | | | | |
|---|---|---|---|---|---|---|---|---|
| ENSG00000136715.18 | SAP130 | protein_coding | 3786,386256 | 0,41169184 | 0,057584745 | 7,149321291 | 8,72E-13 | 1,11E-11 |
| ENSG00000136717.15 | BIN1 | protein_coding | 3081,099092 | 0,296005842 | 0,059264276 | 4,994675701 | 5,89E-07 | 3,96E-06 |
| ENSG00000136750.13 | GAD2 | protein_coding | 8682,595936 | -0,251293159 | 0,052145711 | -4,819057079 | 1,44E-06 | 9,15E-06 |
| ENSG00000136802.11 | LRRC8A | protein_coding | 4054,604711 | -0,144060101 | 0,05905096 | -2,439589498 | 0,01470396 | 0,041710326 |
| ENSG00000136810.13 | TXN | protein_coding | 5824,928953 | -0,145664816 | 0,060564047 | -2,405136754 | 0,016166415 | 0,045265559 |
| ENSG00000136811.16 | ODF2 | protein_coding | 5374,640504 | -0,287385658 | 0,048648557 | -5,907382977 | 3,48E-09 | 3,05E-08 |
| ENSG00000136813.14 | ECPAS | protein_coding | 15428,0247 | 0,470963715 | 0,045245933 | 10,40897344 | 2,26E-25 | 5,94E-24 |
| ENSG00000136819.15 | C9orf78 | protein_coding | 6183,139887 | -0,184683115 | 0,046639818 | -3,959773506 | 7,50E-05 | 0,000365113 |
| ENSG00000136824.19 | SMC2 | protein_coding | 39671,7449 | -0,279113376 | 0,047516509 | -5,874029514 | 4,25E-09 | 3,70E-08 |
| ENSG00000136826.15 | KLF4 | protein_coding | 477,03375 | 0,781353914 | 0,115735849 | 6,751183146 | 1,47E-11 | 1,65E-10 |
| ENSG00000136827.12 | TOR1A | protein_coding | 5603,919818 | 0,155779802 | 0,047307396 | 3,292926979 | 0,000991502 | 0,003843907 |
| ENSG00000136828.19 | RALGPS1 | protein_coding | 7271,44304 | -0,30195743 | 0,050775305 | -5,946934804 | 2,73E-09 | 2,42E-08 |
| ENSG00000136830.12 | FAM129B | protein_coding | 4317,564862 | -0,155767527 | 0,05745951 | -2,710909436 | 0,006709895 | 0,021150123 |
| ENSG00000136842.14 | TMOD1 | protein_coding | 5016,867849 | 0,613563557 | 0,070058032 | 8,757933033 | 1,99E-18 | 3,72E-17 |
| ENSG00000136848.17 | DAB2IP | protein_coding | 4107,622327 | 1,556431909 | 0,091145755 | 17,07629619 | 2,23E-65 | 1,68E-63 |
| ENSG00000136868.11 | SLC31A1 | protein_coding | 11334,80843 | -0,14686062 | 0,060018465 | -2,446923946 | 0,014408124 | 0,040995128 |
| ENSG00000136870.10 | ZNF189 | protein_coding | 6060,387246 | 0,369851784 | 0,05166223 | 7,159036357 | 8,12E-13 | 1,03E-11 |
| ENSG00000136874.11 | STX17 | protein_coding | 12540,08556 | 0,695177727 | 0,047647131 | 14,59012787 | 3,25E-48 | 1,77E-46 |
| ENSG00000136878.13 | USP20 | protein_coding | 3705,419344 | 0,212082839 | 0,069321355 | 3,059415648 | 0,002217692 | 0,007915959 |
| ENSG00000136891.14 | TEX10 | protein_coding | 4170,585877 | 0,198089896 | 0,057136696 | 3,46694696 | 0,000526406 | 0,002169328 |
| ENSG00000136895.19 | GARNL3 | protein_coding | 433,0974201 | 0,295563234 | 0,123790561 | 2,387607184 | 0,016958457 | 0,047166429 |
| ENSG00000136925.15 | TSTD2 | protein_coding | 2700,904022 | 0,263609347 | 0,058477623 | 4,507867 | 6,55E-06 | 3,77E-05 |
| ENSG00000136928.7 | GABBR2 | protein_coding | 14,93261186 | -1,485880434 | 0,628692258 | -2,36344637 | 0,018105847 | 0,049889155 |
| ENSG00000136930.13 | PSMB7 | protein_coding | 9180,203456 | -0,123604816 | 0,047972623 | -2,576569906 | 0,009978603 | 0,029876079 |
| ENSG00000136935.13 | GOLGA1 | protein_coding | 5027,633148 | 0,202403304 | 0,054876966 | 3,688310778 | 0,000225748 | 0,001003626 |
| ENSG00000136936.10 | XPA | protein_coding | 1777,258742 | 0,192429354 | 0,070486315 | 2,730024308 | 0,006332966 | 0,020099933 |
| ENSG00000136937.13 | NCBP1 | protein_coding | 13205,50385 | -0,279177814 | 0,046659176 | -5,983342177 | 2,19E-09 | 1,96E-08 |
| ENSG00000136938.9 | ANP32B | protein_coding | 16638,57296 | -0,369824549 | 0,046110727 | -8,020358196 | 1,05E-15 | 1,66E-14 |
| ENSG00000136940.13 | PDCL | protein_coding | 4071,270639 | -0,178425532 | 0,05428756 | -3,286674383 | 0,00101378 | 0,003922842 |
| ENSG00000136942.15 | RPL35 | protein_coding | 16434,12866 | -0,248918992 | 0,050039711 | -4,974429047 | 6,54E-07 | 4,37E-06 |
| ENSG00000136960.12 | ENPP2 | protein_coding | 35828,28953 | 0,256036185 | 0,041361525 | 6,190201802 | 6,01E-10 | 5,77E-09 |
| ENSG00000136982.6 | DSCC1 | protein_coding | 3321,017081 | -0,457251452 | 0,058130682 | -7,865922725 | 3,66E-15 | 5,58E-14 |
| ENSG00000136986.10 | DERL1 | protein_coding | 6066,342273 | 0,370429045 | 0,05185155 | 7,1440303 | 9,06E-13 | 1,15E-11 |
| ENSG00000137038.8 | DMAC1 | protein_coding | 2770,051297 | -0,394468941 | 0,070079129 | -5,628907597 | 1,81E-08 | 1,47E-07 |
| ENSG00000137054.16 | POLR1E | protein_coding | 1704,663816 | -0,429318802 | 0,073466784 | -5,843713028 | 5,10E-09 | 4,41E-08 |
| ENSG00000137055.15 | PLAA | protein_coding | 6253,499804 | 0,159825652 | 0,059404557 | 2,690461125 | 0,007135335 | 0,022331653 |

| | | | | | | | | |
|---|---|---|---|---|---|---|---|---|
| ENSG00000137100.16 | DCTN3 | protein_coding | 6427,798557 | -0,466814085 | 0,053439365 | -8,735397327 | 2,43E-18 | 4,52E-17 |
| ENSG00000137103.20 | TMEM8B | protein_coding | 1635,121756 | -0,263942295 | 0,07689589 | -3,43246298 | 0,000598126 | 0,002437343 |
| ENSG00000137124.8 | ALDH1B1 | protein_coding | 2609,896181 | -0,242289011 | 0,064226384 | -3,772421777 | 0,000161671 | 0,000738257 |
| ENSG00000137135.18 | ARHGEF39 | protein_coding | 1940,179447 | -0,544455735 | 0,063006795 | -8,641222531 | 5,56E-18 | 1,01E-16 |
| ENSG00000137142.5 | IGFBPL1 | protein_coding | 11694,1429 | -0,487343379 | 0,052997854 | -9,195530505 | 3,73E-20 | 7,62E-19 |
| ENSG00000137145.20 | DENND4C | protein_coding | 17623,49262 | 0,231362942 | 0,040940287 | 5,651229152 | 1,59E-08 | 1,30E-07 |
| ENSG00000137154.12 | RPS6 | protein_coding | 55810,87292 | -0,83212985 | 0,042951341 | -19,37378053 | 1,28E-83 | 1,24E-81 |
| ENSG00000137161.16 | CNPY3 | protein_coding | 3739,581774 | 0,131469197 | 0,052868842 | 2,486704682 | 0,012893238 | 0,037218104 |
| ENSG00000137168.8 | PPIL1 | protein_coding | 5436,106121 | -0,154565511 | 0,053211289 | -2,904750382 | 0,003675461 | 0,012433266 |
| ENSG00000137177.20 | KIF13A | protein_coding | 9629,569939 | 0,161258198 | 0,04561628 | 3,535101913 | 0,000407618 | 0,001722048 |
| ENSG00000137198.9 | GMPR | protein_coding | 538,9352543 | 1,505135953 | 0,107516892 | 13,99906493 | 1,58E-44 | 8,06E-43 |
| ENSG00000137200.13 | CMTR1 | protein_coding | 16076,35024 | 1,747483718 | 0,049274539 | 35,46423246 | 1,75E-275 | 4,48E-273 |
| ENSG00000137210.13 | TMEM14B | protein_coding | 7506,057262 | -0,187276181 | 0,048134968 | -3,890647273 | 1,00E-04 | 0,000475668 |
| ENSG00000137216.19 | TMEM63B | protein_coding | 5689,27628 | 0,134600567 | 0,049311414 | 2,729602645 | 0,006341071 | 0,020117147 |
| ENSG00000137221.14 | TJAP1 | protein_coding | 1885,17677 | -0,248595503 | 0,072262182 | -3,440188159 | 0,00058131 | 0,002378605 |
| ENSG00000137266.14 | SLC22A23 | protein_coding | 2037,561803 | 0,338554101 | 0,069360372 | 4,881088328 | 1,06E-06 | 6,84E-06 |
| ENSG00000137267.6 | TUBB2A | protein_coding | 4282,101832 | 0,319491745 | 0,064761636 | 4,933348899 | 8,08E-07 | 5,32E-06 |
| ENSG00000137269.15 | LRRC1 | protein_coding | 4790,407076 | 0,176730982 | 0,051397003 | 3,438546426 | 0,000584846 | 0,002390933 |
| ENSG00000137274.13 | BPHL | protein_coding | 2603,194572 | -0,242498513 | 0,068848801 | -3,522189356 | 0,000427998 | 0,00180105 |
| ENSG00000137275.14 | RIPK1 | protein_coding | 7544,722724 | 0,907366095 | 0,048900873 | 18,55521268 | 7,40E-77 | 6,72E-75 |
| ENSG00000137310.12 | TCF19 | protein_coding | 10779,57944 | -0,277322815 | 0,043843049 | -6,325354201 | 2,53E-10 | 2,53E-09 |
| ENSG00000137331.12 | IER3 | protein_coding | 1208,337999 | 1,067278877 | 0,091062512 | 11,72028811 | 1,00E-31 | 3,48E-30 |
| ENSG00000137343.18 | ATAT1 | protein_coding | 1917,043969 | -0,213857257 | 0,064891349 | -3,295620477 | 0,000982046 | 0,003813815 |
| ENSG00000137364.5 | TPMT | protein_coding | 5083,060895 | -0,145540978 | 0,050084831 | -2,905889355 | 0,003662109 | 0,012395551 |
| ENSG00000137404.14 | NRM | protein_coding | 1766,601123 | -0,353418637 | 0,067133333 | -5,26442856 | 1,41E-07 | 1,02E-06 |
| ENSG00000137409.19 | MTCH1 | protein_coding | 12215,99659 | -0,19133219 | 0,04828602 | -3,962475892 | 7,42E-05 | 0,00036155 |
| ENSG00000137449.15 | CPEB2 | protein_coding | 5207,795274 | 1,266234898 | 0,053757819 | 23,55443203 | 1,13E-122 | 1,56E-120 |
| ENSG00000137462.8 | TLR2 | protein_coding | 11,43237264 | 1,885566196 | 0,754990036 | 2,49747163 | 0,012508247 | 0,036272628 |
| ENSG00000137474.22 | MYO7A | protein_coding | 2225,670358 | -0,472315702 | 0,071228171 | -6,631023899 | 3,33E-11 | 3,64E-10 |
| ENSG00000137486.17 | ARRB1 | protein_coding | 2773,505862 | -0,20410707 | 0,084573597 | -2,413366314 | 0,015805925 | 0,044377643 |
| ENSG00000137492.8 | THAP12 | protein_coding | 9079,600894 | -0,107820694 | 0,04429982 | -2,433885619 | 0,014937715 | 0,042261566 |
| ENSG00000137496.17 | IL18BP | protein_coding | 1239,203318 | 0,39608152 | 0,094442626 | 4,193885106 | 2,74E-05 | 0,000143044 |
| ENSG00000137497.17 | NUMA1 | protein_coding | 21630,38788 | -0,114137886 | 0,04282958 | -2,664931238 | 0,007700407 | 0,023861296 |
| ENSG00000137501.17 | SYTL2 | protein_coding | 334,3993401 | 0,494783741 | 0,163467024 | 3,02681073 | 0,002471487 | 0,008712716 |
| ENSG00000137509.11 | PRCP | protein_coding | 10820,86983 | 0,257754123 | 0,051315431 | 5,02293593 | 5,09E-07 | 3,45E-06 |
| ENSG00000137513.10 | NARS2 | protein_coding | 4308,958924 | -0,22099018 | 0,061357406 | -3,601687136 | 0,000316159 | 0,001368015 |

| | | | | | | | | |
|---|---|---|---|---|---|---|---|---|
| ENSG00000137571.11 | SLCO5A1 | protein_coding | 1249,589104 | 0,868262591 | 0,098512582 | 8,813722843 | 1,21E-18 | 2,29E-17 |
| ENSG00000137574.10 | TGS1 | protein_coding | 2769,660616 | 0,185282932 | 0,060404337 | 3,067377935 | 0,002159456 | 0,007727695 |
| ENSG00000137601.16 | NEK1 | protein_coding | 4533,596 | 0,320450122 | 0,055175844 | 5,807797336 | 6,33E-09 | 5,40E-08 |
| ENSG00000137628.17 | DDX60 | protein_coding | 20713,46226 | 6,782531463 | 0,13535367 | 50,10969751 | 0 | 0 |
| ENSG00000137642.13 | SORL1 | protein_coding | 10265,1459 | -0,628151956 | 0,069056481 | -9,096205714 | 9,35E-20 | 1,87E-18 |
| ENSG00000137699.17 | TRIM29 | protein_coding | 1170,294205 | -1,030255933 | 0,100541595 | -10,24706179 | 1,22E-24 | 3,12E-23 |
| ENSG00000137700.18 | SLC37A4 | protein_coding | 28745,30722 | 0,280577983 | 0,046568321 | 6,025082674 | 1,69E-09 | 1,54E-08 |
| ENSG00000137710.16 | RDX | protein_coding | 18974,61575 | -0,283655981 | 0,041360881 | -6,858073986 | 6,98E-12 | 8,15E-11 |
| ENSG00000137713.15 | PPP2R1B | protein_coding | 10361,26881 | 0,31752108 | 0,045200233 | 7,024766509 | 2,14E-12 | 2,64E-11 |
| ENSG00000137720.8 | C11orf1 | protein_coding | 1280,754283 | -0,203985555 | 0,082444922 | -2,474203996 | 0,013353342 | 0,038390965 |
| ENSG00000137727.12 | ARHGAP20 | protein_coding | 3224,210931 | 0,279031036 | 0,055499062 | 5,027671208 | 4,96E-07 | 3,37E-06 |
| ENSG00000137767.14 | SQOR | protein_coding | 392,1030348 | 0,390942477 | 0,130441157 | 2,997079191 | 0,002725799 | 0,009512404 |
| ENSG00000137770.14 | CTDSPL2 | protein_coding | 10055,39051 | 0,169562248 | 0,051358284 | 3,301555937 | 0,000961502 | 0,003743719 |
| ENSG00000137776.17 | SLTM | protein_coding | 14174,3517 | 0,131436914 | 0,050428711 | 2,60639052 | 0,009150207 | 0,027746465 |
| ENSG00000137802.14 | MAPKBP1 | protein_coding | 3411,646855 | 0,273081006 | 0,062678599 | 4,356846038 | 1,32E-05 | 7,26E-05 |
| ENSG00000137804.14 | NUSAP1 | protein_coding | 18578,1611 | -0,497864712 | 0,048377369 | -10,29127302 | 7,71E-25 | 1,99E-23 |
| ENSG00000137807.15 | AC027237.1 | protein_coding | 4532,184799 | -0,917343456 | 0,059257475 | -15,48063694 | 4,69E-54 | 2,90E-52 |
| ENSG00000137812.19 | KNL1 | protein_coding | 8276,396865 | -0,172571926 | 0,0585932 | -2,945255173 | 0,003226884 | 0,01105876 |
| ENSG00000137814.11 | HAUS2 | protein_coding | 3686,938944 | -0,479445197 | 0,067094822 | -7,145785362 | 8,95E-13 | 1,13E-11 |
| ENSG00000137817.17 | PARP6 | protein_coding | 12886,77057 | 0,451509207 | 0,048457887 | 9,317558659 | 1,19E-20 | 2,52E-19 |
| ENSG00000137818.12 | RPLP1 | protein_coding | 43345,46594 | -0,649728391 | 0,052316171 | -12,41926508 | 2,05E-35 | 8,03E-34 |
| ENSG00000137819.13 | PAQR5 | protein_coding | 12,66830568 | -2,07575839 | 0,845470865 | -2,455150704 | 0,014082554 | 0,040231527 |
| ENSG00000137821.11 | LRRC49 | protein_coding | 2180,017944 | -0,551774479 | 0,089015064 | -6,198664086 | 5,69E-10 | 5,49E-09 |
| ENSG00000137824.16 | RMDN3 | protein_coding | 2618,290258 | 0,213182069 | 0,066430114 | 3,209117905 | 0,001331429 | 0,005019955 |
| ENSG00000137831.15 | UACA | protein_coding | 738,4785732 | -0,288115737 | 0,118628717 | -2,428718303 | 0,0151523 | 0,042787997 |
| ENSG00000137842.7 | TMEM62 | protein_coding | 11218,08449 | 2,019624084 | 0,047254009 | 42,73974028 | 0 | 0 |
| ENSG00000137845.15 | ADAM10 | protein_coding | 20240,89246 | 0,226734826 | 0,045423323 | 4,99159492 | 5,99E-07 | 4,02E-06 |
| ENSG00000137868.19 | STRA6 | protein_coding | 12636,86964 | -0,517829571 | 0,04183345 | -12,37836165 | 3,42E-35 | 1,32E-33 |
| ENSG00000137872.16 | SEMA6D | protein_coding | 4003,445407 | 1,020907486 | 0,078497807 | 13,00555423 | 1,14E-38 | 4,91E-37 |
| ENSG00000137876.10 | RSL24D1 | protein_coding | 7447,908134 | -0,111760563 | 0,046273632 | -2,415210541 | 0,015726117 | 0,044164586 |
| ENSG00000137877.10 | SPTBN5 | protein_coding | 67,45841576 | 2,412550613 | 0,337509773 | 7,148091124 | 8,80E-13 | 1,12E-11 |
| ENSG00000137878.17 | GCOM1 | protein_coding | 325,932611 | 0,413313348 | 0,132825231 | 3,111708114 | 0,001860083 | 0,006777049 |
| ENSG00000137880.6 | GCHFR | protein_coding | 5810,155638 | -0,487227875 | 0,052757564 | -9,235223094 | 2,58E-20 | 5,33E-19 |
| ENSG00000137941.17 | TTLL7 | protein_coding | 4683,429178 | -0,327972925 | 0,063358448 | -5,176467172 | 2,26E-07 | 1,60E-06 |
| ENSG00000137944.18 | KYAT3 | protein_coding | 5511,4315 | 0,27645519 | 0,050469346 | 5,477685167 | 4,31E-08 | 3,32E-07 |
| ENSG00000137947.12 | GTF2B | protein_coding | 5352,868896 | 0,618885996 | 0,055908202 | 11,06968169 | 1,76E-28 | 5,24E-27 |

| | | | | | | | | |
|---|---|---|---|---|---|---|---|---|
| ENSG00000137955.16 | RABGGTB | protein_coding | 9124,887294 | -0,303839902 | 0,044365741 | -6,848525374 | 7,46E-12 | 8,67E-11 |
| ENSG00000137960.6 | GIPC2 | protein_coding | 8197,725048 | -0,332394691 | 0,046726549 | -7,113615279 | 1,13E-12 | 1,42E-11 |
| ENSG00000137968.16 | SLC44A5 | protein_coding | 273,8220148 | -0,398450027 | 0,148736772 | -2,678893873 | 0,00738658 | 0,023002841 |
| ENSG00000137996.12 | RTCA | protein_coding | 6020,981466 | 0,153743701 | 0,049370783 | 3,114062448 | 0,001845304 | 0,006727679 |
| ENSG00000138018.18 | SELENOI | protein_coding | 27677,40872 | -0,52157641 | 0,050425426 | -10,3435201 | 4,48E-25 | 1,17E-23 |
| ENSG00000138030.13 | KHK | protein_coding | 8829,11853 | -0,140974069 | 0,049291931 | -2,85998268 | 0,004236641 | 0,014102716 |
| ENSG00000138031.14 | ADCY3 | protein_coding | 925,9816529 | -0,27294366 | 0,113572559 | -2,403253581 | 0,016249915 | 0,045459773 |
| ENSG00000138035.15 | PNPT1 | protein_coding | 27761,35311 | 2,767715318 | 0,045882759 | 60,32146694 | 0 | 0 |
| ENSG00000138050.15 | THUMPD2 | protein_coding | 1670,371607 | 0,23832853 | 0,073537535 | 3,240909971 | 0,001191488 | 0,004534867 |
| ENSG00000138071.13 | ACTR2 | protein_coding | 47859,22286 | -0,412765003 | 0,044411095 | -9,294186509 | 1,48E-20 | 3,11E-19 |
| ENSG00000138074.15 | SLC5A6 | protein_coding | 4457,637559 | -0,205946253 | 0,056052204 | -3,674186559 | 0,000238608 | 0,001056211 |
| ENSG00000138078.15 | PREPL | protein_coding | 20091,97773 | -0,361850052 | 0,040782886 | -8,872595502 | 7,15E-19 | 1,37E-17 |
| ENSG00000138079.13 | SLC3A1 | protein_coding | 226,9217512 | -0,518996623 | 0,176369091 | -2,942673344 | 0,003253916 | 0,011148004 |
| ENSG00000138083.5 | SIX3 | protein_coding | 788,5109386 | 0,294356146 | 0,091493473 | 3,217236562 | 0,001294318 | 0,004889864 |
| ENSG00000138085.16 | ATRAID | protein_coding | 10139,82228 | 0,202963518 | 0,042997737 | 4,720330198 | 2,35E-06 | 1,45E-05 |
| ENSG00000138092.11 | CENPO | protein_coding | 6200,709769 | -0,135334743 | 0,057231816 | -2,364676743 | 0,018045814 | 0,049735917 |
| ENSG00000138095.19 | LRPPRC | protein_coding | 24834,50805 | -0,23599956 | 0,044607234 | -5,290611832 | 1,22E-07 | 8,92E-07 |
| ENSG00000138107.13 | ACTR1A | protein_coding | 17982,9135 | -0,172602753 | 0,042862017 | -4,026939589 | 5,65E-05 | 0,000280903 |
| ENSG00000138131.4 | LOXL4 | protein_coding | 363,5000374 | 0,430707765 | 0,127082155 | 3,389207288 | 0,00070095 | 0,002815432 |
| ENSG00000138138.13 | ATAD1 | protein_coding | 11444,66923 | 0,222179382 | 0,049599152 | 4,479499575 | 7,48E-06 | 4,26E-05 |
| ENSG00000138160.6 | KIF11 | protein_coding | 12152,02824 | -0,522849575 | 0,045888264 | -11,39397156 | 4,48E-30 | 1,42E-28 |
| ENSG00000138166.6 | DUSP5 | protein_coding | 1369,025593 | 0,810275274 | 0,09705614 | 8,34852152 | 6,91E-17 | 1,17E-15 |
| ENSG00000138175.9 | ARL3 | protein_coding | 7902,776715 | -0,466246988 | 0,057911956 | -8,05096251 | 8,21E-16 | 1,30E-14 |
| ENSG00000138180.15 | CEP55 | protein_coding | 5595,723768 | -0,688520676 | 0,055924291 | -12,31165669 | 7,84E-35 | 2,97E-33 |
| ENSG00000138182.14 | KIF20B | protein_coding | 7354,206507 | -0,390375276 | 0,051215476 | -7,622213197 | 2,49E-14 | 3,55E-13 |
| ENSG00000138185.20 | ENTPD1 | protein_coding | 968,7624489 | 0,864410998 | 0,106789193 | 8,094555011 | 5,75E-16 | 9,21E-15 |
| ENSG00000138207.14 | RBP4 | protein_coding | 3060,767989 | 0,297739405 | 0,059849791 | 4,974777722 | 6,53E-07 | 4,36E-06 |
| ENSG00000138231.12 | DBR1 | protein_coding | 3823,921883 | 0,218708419 | 0,05694024 | 3,841016781 | 0,000122526 | 0,000574569 |
| ENSG00000138246.17 | DNAJC13 | protein_coding | 12164,02564 | 0,378345285 | 0,043328543 | 8,732010382 | 2,50E-18 | 4,65E-17 |
| ENSG00000138279.16 | ANXA7 | protein_coding | 24928,82735 | 0,322850684 | 0,045648042 | 7,072607499 | 1,52E-12 | 1,90E-11 |
| ENSG00000138286.14 | FAM149B1 | protein_coding | 2235,561479 | 0,215391971 | 0,0672512 | 3,202797443 | 0,001360997 | 0,005119431 |
| ENSG00000138326.20 | RPS24 | protein_coding | 80712,64675 | -0,571534362 | 0,042090296 | -13,57876795 | 5,35E-42 | 2,56E-40 |
| ENSG00000138336.9 | TET1 | protein_coding | 2419,344516 | 0,149966582 | 0,062737403 | 2,390385538 | 0,016830694 | 0,046874796 |
| ENSG00000138346.15 | DNA2 | protein_coding | 5486,012066 | -0,366350943 | 0,050444959 | -7,262389502 | 3,80E-13 | 4,97E-12 |
| ENSG00000138356.14 | AOX1 | protein_coding | 557,4114788 | 0,31011574 | 0,111280054 | 2,786804363 | 0,00532306 | 0,017242304 |
| ENSG00000138375.12 | SMARCAL1 | protein_coding | 3521,264681 | 0,193286425 | 0,052955077 | 3,650007464 | 0,000262233 | 0,001151277 |

| | | | | | | | | |
|---|---|---|---|---|---|---|---|---|
| ENSG00000138376.11 | BARD1 | protein_coding | 2370,709509 | 0,650996832 | 0,083940471 | 7,755458443 | 8,80E-15 | 1,30E-13 |
| ENSG00000138378.19 | STAT4 | protein_coding | 221,7851746 | 2,129269164 | 0,186437524 | 11,42081869 | 3,29E-30 | 1,05E-28 |
| ENSG00000138386.17 | NAB1 | protein_coding | 5357,237496 | -0,263504739 | 0,052367692 | -5,031818851 | 4,86E-07 | 3,30E-06 |
| ENSG00000138400.13 | MDH1B | protein_coding | 557,0723398 | -0,349811801 | 0,10505397 | -3,329829438 | 0,000868992 | 0,003413041 |
| ENSG00000138411.13 | HECW2 | protein_coding | 3959,13438 | 0,355562238 | 0,070432432 | 5,048274342 | 4,46E-07 | 3,05E-06 |
| ENSG00000138430.16 | OLA1 | protein_coding | 14131,27103 | -0,229119316 | 0,046380336 | -4,940009856 | 7,81E-07 | 5,15E-06 |
| ENSG00000138433.16 | CIR1 | protein_coding | 5987,925427 | 0,368058544 | 0,051589667 | 7,134346222 | 9,72E-13 | 1,23E-11 |
| ENSG00000138434.17 | ITPRID2 | protein_coding | 31257,19224 | 0,252066385 | 0,045018936 | 5,599119097 | 2,15E-08 | 1,73E-07 |
| ENSG00000138435.15 | CHRNA1 | protein_coding | 17,88420276 | 3,011825581 | 0,667247171 | 4,513807941 | 6,37E-06 | 3,67E-05 |
| ENSG00000138439.12 | FAM117B | protein_coding | 3272,897421 | -0,226766636 | 0,063850561 | -3,551521426 | 0,000383011 | 0,001627868 |
| ENSG00000138442.9 | WDR12 | protein_coding | 3838,934055 | -0,294753401 | 0,071196831 | -4,139979231 | 3,47E-05 | 0,000178376 |
| ENSG00000138443.16 | ABI2 | protein_coding | 11731,37702 | -0,116555872 | 0,043191709 | -2,6985705 | 0,006963799 | 0,021873869 |
| ENSG00000138448.12 | ITGAV | protein_coding | 7034,641962 | -0,461299505 | 0,048092296 | -9,591962702 | 8,64E-22 | 1,93E-20 |
| ENSG00000138463.9 | SLC49A4 | protein_coding | 1873,267354 | 0,207050742 | 0,071888225 | 2,880176039 | 0,003974532 | 0,013326789 |
| ENSG00000138495.6 | COX17 | protein_coding | 7527,395357 | -0,39395398 | 0,060962287 | -6,462257278 | 1,03E-10 | 1,07E-09 |
| ENSG00000138496.16 | PARP9 | protein_coding | 38406,66301 | 5,492892411 | 0,157157743 | 34,951459 | 1,23E-267 | 3,11E-265 |
| ENSG00000138592.14 | USP8 | protein_coding | 11895,30419 | 0,175598551 | 0,04563498 | 3,847893696 | 0,000119138 | 0,000559965 |
| ENSG00000138593.9 | SECISBP2L | protein_coding | 9874,313954 | 0,218837161 | 0,046830696 | 4,672942747 | 2,97E-06 | 1,80E-05 |
| ENSG00000138594.14 | TMOD3 | protein_coding | 10410,94041 | -0,284895458 | 0,050241761 | -5,670491037 | 1,42E-08 | 1,17E-07 |
| ENSG00000138613.14 | APH1B | protein_coding | 5871,335717 | 0,15347252 | 0,056991819 | 2,692886845 | 0,007083631 | 0,022197611 |
| ENSG00000138629.16 | UBL7 | protein_coding | 3263,572631 | -0,174702005 | 0,058509697 | -2,985864131 | 0,002827783 | 0,009828684 |
| ENSG00000138639.18 | ARHGAP24 | protein_coding | 185,976845 | 0,829793931 | 0,185456099 | 4,474341567 | 7,66E-06 | 4,36E-05 |
| ENSG00000138642.14 | HERC6 | protein_coding | 6894,740353 | 3,307363607 | 0,059378493 | 55,69968937 | 0 | 0 |
| ENSG00000138646.9 | HERC5 | protein_coding | 33270,37297 | 3,389692371 | 0,043023168 | 78,78760556 | 0 | 0 |
| ENSG00000138663.9 | COPS4 | protein_coding | 6317,393706 | -0,252755564 | 0,056912133 | -4,441154292 | 8,95E-06 | 5,05E-05 |
| ENSG00000138669.9 | PRKG2 | protein_coding | 337,8510535 | -0,42463296 | 0,145964696 | -2,90914839 | 0,003624148 | 0,012279986 |
| ENSG00000138670.17 | RASGEF1B | protein_coding | 1699,114908 | 0,257456539 | 0,07157911 | 3,596811147 | 0,000322142 | 0,001391231 |
| ENSG00000138675.16 | FGF5 | protein_coding | 794,095242 | 0,232155206 | 0,089984784 | 2,579938472 | 0,009881792 | 0,029619283 |
| ENSG00000138678.11 | GPAT3 | protein_coding | 605,6061044 | 0,474341424 | 0,107694464 | 4,40451076 | 1,06E-05 | 5,92E-05 |
| ENSG00000138686.9 | BBS7 | protein_coding | 2733,360297 | -0,281357674 | 0,073206688 | -3,84333291 | 0,000121375 | 0,000569884 |
| ENSG00000138722.10 | MMRN1 | protein_coding | 6031,446528 | -0,891302884 | 0,052243946 | -17,06040526 | 2,93E-65 | 2,20E-63 |
| ENSG00000138735.16 | PDE5A | protein_coding | 13230,10518 | -0,164791378 | 0,052818598 | -3,11994986 | 0,001808818 | 0,006612748 |
| ENSG00000138756.17 | BMP2K | protein_coding | 9607,259873 | 0,488613711 | 0,050422762 | 9,690339991 | 3,31E-22 | 7,56E-21 |
| ENSG00000138758.11 | sept-11 | protein_coding | 15333,49941 | -0,376146892 | 0,046162175 | -8,14837884 | 3,69E-16 | 5,98E-15 |
| ENSG00000138759.19 | FRAS1 | protein_coding | 1493,258208 | 0,410942175 | 0,09596523 | 4,282198616 | 1,85E-05 | 9,93E-05 |
| ENSG00000138760.10 | SCARB2 | protein_coding | 109801,1125 | 0,516750208 | 0,038774128 | 13,32719103 | 1,61E-40 | 7,34E-39 |

| | | | | | | | | |
|---|---|---|---|---|---|---|---|---|
| ENSG00000138764.15 | CCNG2 | protein_coding | 6096,946198 | -0,134229978 | 0,055847609 | -2,403504462 | 0,016238769 | 0,045445536 |
| ENSG00000138768.14 | USO1 | protein_coding | 17080,27735 | 0,165247239 | 0,04267758 | 3,871991761 | 0,00010795 | 0,000510363 |
| ENSG00000138769.11 | CDKL2 | protein_coding | 788,7948792 | -0,337002744 | 0,095067731 | -3,54486996 | 0,000392807 | 0,001664142 |
| ENSG00000138771.16 | SHROOM3 | protein_coding | 5289,557681 | 0,352556875 | 0,075419459 | 4,674614181 | 2,95E-06 | 1,79E-05 |
| ENSG00000138777.20 | PPA2 | protein_coding | 4169,820197 | -0,135005987 | 0,051349771 | -2,62914485 | 0,008559989 | 0,026186028 |
| ENSG00000138778.11 | CENPE | protein_coding | 7223,168093 | -1,151638449 | 0,056482995 | -20,38911785 | 2,09E-92 | 2,25E-90 |
| ENSG00000138780.14 | GSTCD | protein_coding | 3748,429657 | -0,252207304 | 0,059643516 | -4,228578762 | 2,35E-05 | 0,000124436 |
| ENSG00000138785.15 | INTS12 | protein_coding | 2626,513985 | 0,362337309 | 0,069575313 | 5,207843053 | 1,91E-07 | 1,36E-06 |
| ENSG00000138796.16 | HADH | protein_coding | 86096,15542 | -0,176732017 | 0,040109749 | -4,406211005 | 1,05E-05 | 5,87E-05 |
| ENSG00000138801.9 | PAPSS1 | protein_coding | 13501,98636 | 0,138657808 | 0,043825482 | 3,163862709 | 0,001556902 | 0,005788621 |
| ENSG00000138821.13 | SLC39A8 | protein_coding | 2325,243632 | -0,177665457 | 0,066122002 | -2,68693403 | 0,007211119 | 0,022524991 |
| ENSG00000138834.12 | MAPK8IP3 | protein_coding | 5053,455726 | -0,165607717 | 0,06538903 | -2,532652897 | 0,011320299 | 0,033264831 |
| ENSG00000138835.22 | RGS3 | protein_coding | 2277,134965 | -0,497576644 | 0,071597394 | -6,949647417 | 3,66E-12 | 4,39E-11 |
| ENSG00000138867.16 | GUCD1 | protein_coding | 7849,595379 | 0,18449203 | 0,047550262 | 3,879937163 | 0,000104483 | 0,000495328 |
| ENSG00000138944.8 | SHISAL1 | protein_coding | 3231,49829 | -0,450178644 | 0,078496383 | -5,735024023 | 9,75E-09 | 8,11E-08 |
| ENSG00000139083.10 | ETV6 | protein_coding | 2467,219448 | 1,245904223 | 0,060530484 | 20,58308696 | 3,89E-94 | 4,27E-92 |
| ENSG00000139112.11 | GABARAPL1 | protein_coding | 136,6870387 | 0,609775733 | 0,201942313 | 3,019554073 | 0,002531471 | 0,008910224 |
| ENSG00000139116.18 | KIF21A | protein_coding | 24998,84984 | -0,125186474 | 0,040390021 | -3,099440703 | 0,001938864 | 0,007029943 |
| ENSG00000139144.9 | PIK3C2G | protein_coding | 71,24652702 | 0,716786331 | 0,301702155 | 2,375807792 | 0,017510582 | 0,04847884 |
| ENSG00000139146.14 | SINHCAF | protein_coding | 13046,92306 | -0,199009289 | 0,043230851 | -4,603408964 | 4,16E-06 | 2,46E-05 |
| ENSG00000139168.8 | ZCRB1 | protein_coding | 4779,479021 | -0,409274809 | 0,054979608 | -7,444120115 | 9,76E-14 | 1,34E-12 |
| ENSG00000139178.11 | C1RL | protein_coding | 2052,691635 | 0,242391971 | 0,063592389 | 3,811650642 | 0,000138042 | 0,000639865 |
| ENSG00000139180.11 | NDUFA9 | protein_coding | 10533,6785 | 0,503115636 | 0,04792323 | 10,49836653 | 8,79E-26 | 2,36E-24 |
| ENSG00000139182.14 | CLSTN3 | protein_coding | 19698,30524 | 0,469941048 | 0,055476182 | 8,471041576 | 2,43E-17 | 4,24E-16 |
| ENSG00000139192.12 | TAPBPL | protein_coding | 1285,511726 | 2,395393254 | 0,101752636 | 23,5413385 | 1,54E-122 | 2,10E-120 |
| ENSG00000139197.10 | PEX5 | protein_coding | 6980,949512 | -0,128810191 | 0,047579432 | -2,707266296 | 0,006783982 | 0,021368688 |
| ENSG00000139211.6 | AMIGO2 | protein_coding | 8900,53854 | 0,177325213 | 0,045871308 | 3,865710849 | 0,000110766 | 0,00052257 |
| ENSG00000139218.18 | SCAF11 | protein_coding | 12313,79329 | 0,167421672 | 0,046998312 | 3,562291153 | 0,000367632 | 0,001569025 |
| ENSG00000139220.16 | PPFIA2 | protein_coding | 1435,052387 | 0,976188121 | 0,114307758 | 8,539998829 | 1,34E-17 | 2,38E-16 |
| ENSG00000139263.11 | LRIG3 | protein_coding | 2782,979252 | 0,178440061 | 0,066166655 | 2,696827587 | 0,007000351 | 0,02197335 |
| ENSG00000139266.6 | mars-09 | protein_coding | 3385,589835 | 0,129679053 | 0,054045023 | 2,399463372 | 0,016419122 | 0,045859043 |
| ENSG00000139269.3 | INHBE | protein_coding | 82,11700117 | 2,695468218 | 0,39113817 | 6,891345366 | 5,53E-12 | 6,50E-11 |
| ENSG00000139287.13 | TPH2 | protein_coding | 592,2298497 | -0,559130032 | 0,118747188 | -4,708574921 | 2,49E-06 | 1,53E-05 |
| ENSG00000139291.13 | TMEM19 | protein_coding | 5830,115388 | -0,675039889 | 0,065626228 | -10,28612956 | 8,14E-25 | 2,10E-23 |
| ENSG00000139324.12 | TMTC3 | protein_coding | 6811,727346 | -0,127196908 | 0,047775092 | -2,662410533 | 0,007758319 | 0,024020909 |
| ENSG00000139343.10 | SNRPF | protein_coding | 6492,847823 | -0,237224815 | 0,06651879 | -3,566282768 | 0,000362081 | 0,001546797 |

| | | | | | | | | |
|---|---|---|---|---|---|---|---|---|
| ENSG00000139372.15 | TDG | protein_coding | 13083,79717 | -0,239809845 | 0,044683804 | -5,366818032 | 8,01E-08 | 5,96E-07 |
| ENSG00000139410.15 | SDSL | protein_coding | 491,9057357 | 0,460705246 | 0,112097145 | 4,109874915 | 3,96E-05 | 0,000201646 |
| ENSG00000139433.10 | GLTP | protein_coding | 1597,658198 | -0,432045124 | 0,076624996 | -5,638435819 | 1,72E-08 | 1,39E-07 |
| ENSG00000139436.21 | GIT2 | protein_coding | 3833,603894 | 0,187900395 | 0,058101499 | 3,234002539 | 0,001220684 | 0,004635015 |
| ENSG00000139505.11 | MTMR6 | protein_coding | 9325,39896 | -0,169620924 | 0,050161124 | -3,381521591 | 0,000720856 | 0,002887142 |
| ENSG00000139514.13 | SLC7A1 | protein_coding | 23764,96249 | 0,13009695 | 0,053502692 | 2,431596337 | 0,01503245 | 0,042492204 |
| ENSG00000139515.6 | PDX1 | protein_coding | 25226,88701 | 0,311718709 | 0,047445793 | 6,569996764 | 5,03E-11 | 5,40E-10 |
| ENSG00000139517.9 | LNX2 | protein_coding | 7254,044375 | 0,283719091 | 0,04731103 | 5,996891049 | 2,01E-09 | 1,82E-08 |
| ENSG00000139531.13 | SUOX | protein_coding | 2518,488587 | 0,431608184 | 0,066240608 | 6,515764197 | 7,23E-11 | 7,63E-10 |
| ENSG00000139546.10 | TARBP2 | protein_coding | 1807,36045 | -0,200026638 | 0,080873426 | -2,473329616 | 0,013386062 | 0,038461433 |
| ENSG00000139579.13 | NABP2 | protein_coding | 4607,378533 | -0,2693847 | 0,054112035 | -4,978277021 | 6,42E-07 | 4,29E-06 |
| ENSG00000139597.18 | N4BP2L1 | protein_coding | 444,0226747 | 0,883344067 | 0,127504369 | 6,927951336 | 4,27E-12 | 5,09E-11 |
| ENSG00000139618.15 | BRCA2 | protein_coding | 6323,168542 | 0,856471394 | 0,159003464 | 5,386495199 | 7,18E-08 | 5,38E-07 |
| ENSG00000139620.12 | KANSL2 | protein_coding | 3999,677195 | 0,180299024 | 0,058768584 | 3,067949096 | 0,002155333 | 0,007714167 |
| ENSG00000139629.16 | GALNT6 | protein_coding | 225,3404331 | 0,627477081 | 0,161979742 | 3,873799721 | 0,000107151 | 0,000507015 |
| ENSG00000139641.12 | ESYT1 | protein_coding | 30216,46486 | 0,183370188 | 0,047934387 | 3,825441362 | 0,000130538 | 0,000608718 |
| ENSG00000139644.13 | TMBIM6 | protein_coding | 86388,9706 | -0,166159766 | 0,046590254 | -3,566406064 | 0,00036191 | 0,001546545 |
| ENSG00000139651.11 | ZNF740 | protein_coding | 7618,081079 | 0,186860541 | 0,054389179 | 3,435619802 | 0,0005912 | 0,002412495 |
| ENSG00000139675.12 | HNRNPA1L2 | protein_coding | 768,712558 | 0,31649087 | 0,102376398 | 3,091443684 | 0,001991858 | 0,007201208 |
| ENSG00000139679.15 | LPAR6 | protein_coding | 609,7847565 | 0,684892934 | 0,129433786 | 5,291454085 | 1,21E-07 | 8,89E-07 |
| ENSG00000139684.14 | ESD | protein_coding | 16838,70175 | -0,196101995 | 0,041998026 | -4,669314622 | 3,02E-06 | 1,83E-05 |
| ENSG00000139687.15 | RB1 | protein_coding | 21852,26774 | -0,122168061 | 0,045342768 | -2,694322987 | 0,007053178 | 0,022117579 |
| ENSG00000139697.14 | SBNO1 | protein_coding | 11240,45204 | -0,228808135 | 0,047852169 | -4,781562496 | 1,74E-06 | 1,09E-05 |
| ENSG00000139718.10 | SETD1B | protein_coding | 2378,740179 | 0,229486416 | 0,06296871 | 3,644451589 | 0,000267963 | 0,001174828 |
| ENSG00000139719.10 | VPS33A | protein_coding | 6957,680256 | -0,194090993 | 0,053587403 | -3,621951811 | 0,000292389 | 0,001272501 |
| ENSG00000139726.11 | DENR | protein_coding | 18783,81456 | -0,287456805 | 0,04990659 | -5,759896715 | 8,42E-09 | 7,06E-08 |
| ENSG00000139734.18 | DIAPH3 | protein_coding | 4087,723312 | -0,379555434 | 0,061206171 | -6,201260832 | 5,60E-10 | 5,41E-09 |
| ENSG00000139737.22 | SLAIN1 | protein_coding | 2659,49289 | -0,189901816 | 0,06027895 | -3,150383608 | 0,001630562 | 0,006025665 |
| ENSG00000139835.13 | GRTP1 | protein_coding | 231,0664632 | 0,391543544 | 0,155317394 | 2,52092527 | 0,011704671 | 0,034220085 |
| ENSG00000139874.6 | SSTR1 | protein_coding | 7965,178742 | 0,173625373 | 0,051516645 | 3,3702772 | 0,000750926 | 0,00299446 |
| ENSG00000139921.13 | TMX1 | protein_coding | 12461,59299 | 0,230513277 | 0,043101919 | 5,348097793 | 8,89E-08 | 6,59E-07 |
| ENSG00000139946.10 | PELI2 | protein_coding | 2010,93259 | -0,267388098 | 0,06569841 | -4,069932586 | 4,70E-05 | 0,000236445 |
| ENSG00000139971.15 | ARMH4 | protein_coding | 1072,278592 | -0,695579755 | 0,128018618 | -5,433426521 | 5,53E-08 | 4,22E-07 |
| ENSG00000139973.16 | SYT16 | protein_coding | 346,8193419 | -1,322658785 | 0,418253496 | -3,16233767 | 0,00156508 | 0,005814223 |
| ENSG00000139990.17 | DCAF5 | protein_coding | 11774,87115 | 0,15603863 | 0,048708957 | 3,203489436 | 0,00135773 | 0,005109706 |
| ENSG00000140006.11 | WDR89 | protein_coding | 2365,385113 | 0,154902285 | 0,059615498 | 2,598355993 | 0,009367133 | 0,028324125 |

| | | | | | | | | |
|---|---|---|---|---|---|---|---|---|
| ENSG00000140025.16 | EFCAB11 | protein_coding | 1234,366878 | -0,640910794 | 0,083357961 | -7,688657259 | 1,49E-14 | 2,15E-13 |
| ENSG00000140043.11 | PTGR2 | protein_coding | 2673,697374 | -0,335371251 | 0,070115545 | -4,783122652 | 1,73E-06 | 1,08E-05 |
| ENSG00000140044.13 | JDP2 | protein_coding | 41,75890964 | 1,193220258 | 0,476962529 | 2,501706495 | 0,012359634 | 0,035920336 |
| ENSG00000140093.10 | SERPINA10 | protein_coding | 5256,979853 | 0,519107026 | 0,056055601 | 9,26057377 | 2,03E-20 | 4,23E-19 |
| ENSG00000140105.18 | WARS | protein_coding | 9718,646839 | 0,67171676 | 0,058891111 | 11,40608069 | 3,90E-30 | 1,24E-28 |
| ENSG00000140199.11 | SLC12A6 | protein_coding | 6804,013491 | -0,23000097 | 0,072481944 | -3,173217438 | 0,001507596 | 0,005622945 |
| ENSG00000140264.19 | SERF2 | protein_coding | 26482,61039 | -0,148908899 | 0,040146116 | -3,709173291 | 0,000207937 | 0,000929215 |
| ENSG00000140265.12 | ZSCAN29 | protein_coding | 7140,507545 | 0,518394677 | 0,050611262 | 10,24267429 | 1,28E-24 | 3,26E-23 |
| ENSG00000140280.14 | LYSMD2 | protein_coding | 2875,79106 | 0,679745435 | 0,06985225 | 9,731188804 | 2,22E-22 | 5,10E-21 |
| ENSG00000140285.10 | FGF7 | protein_coding | 1043,739424 | -0,210366711 | 0,081281104 | -2,588137967 | 0,009649632 | 0,029031727 |
| ENSG00000140297.13 | GCNT3 | protein_coding | 1713,941688 | 1,09352801 | 0,074171177 | 14,74330132 | 3,40E-49 | 1,90E-47 |
| ENSG00000140307.11 | GTF2A2 | protein_coding | 6436,727204 | -0,22950267 | 0,062579769 | -3,667362037 | 0,000245066 | 0,001081605 |
| ENSG00000140319.10 | SRP14 | protein_coding | 26712,15114 | -0,140266982 | 0,04815656 | -2,912728426 | 0,00358286 | 0,012151131 |
| ENSG00000140320.12 | BAHD1 | protein_coding | 9728,192628 | -0,139593587 | 0,046723727 | -2,987638103 | 0,002811422 | 0,00977786 |
| ENSG00000140323.6 | DISP2 | protein_coding | 2277,54655 | -0,256830814 | 0,073361534 | -3,500892063 | 0,000463704 | 0,001938975 |
| ENSG00000140332.16 | TLE3 | protein_coding | 1078,993583 | -0,452372842 | 0,123719048 | -3,656452643 | 0,00025573 | 0,00112536 |
| ENSG00000140365.15 | COMMD4 | protein_coding | 6045,442815 | -0,304261589 | 0,054975946 | -5,534449339 | 3,12E-08 | 2,46E-07 |
| ENSG00000140368.12 | PSTPIP1 | protein_coding | 72,52296015 | -1,968889657 | 0,284012503 | -6,932404868 | 4,14E-12 | 4,94E-11 |
| ENSG00000140374.16 | ETFA | protein_coding | 9854,016844 | -0,162426996 | 0,049370923 | -3,289932326 | 0,001002115 | 0,003882376 |
| ENSG00000140391.14 | TSPAN3 | protein_coding | 31463,2391 | 0,112305774 | 0,038375648 | 2,926485408 | 0,003428156 | 0,011693341 |
| ENSG00000140395.9 | WDR61 | protein_coding | 4869,915198 | -0,178952964 | 0,052232862 | -3,42606083 | 0,000612403 | 0,00248999 |
| ENSG00000140396.13 | NCOA2 | protein_coding | 8792,933752 | 0,275466172 | 0,054155537 | 5,086574508 | 3,65E-07 | 2,52E-06 |
| ENSG00000140406.3 | TLNRD1 | protein_coding | 1296,97347 | 0,216047085 | 0,081203507 | 2,660563484 | 0,007801002 | 0,024126512 |
| ENSG00000140416.21 | TPM1 | protein_coding | 13825,77967 | 0,284441802 | 0,045321495 | 6,276090476 | 3,47E-10 | 3,43E-09 |
| ENSG00000140443.15 | IGF1R | protein_coding | 7020,77857 | -0,277429639 | 0,051443941 | -5,392853537 | 6,93E-08 | 5,21E-07 |
| ENSG00000140451.13 | PIF1 | protein_coding | 582,2756021 | -1,151676243 | 0,146991388 | -7,834991296 | 4,69E-15 | 7,09E-14 |
| ENSG00000140455.17 | USP3 | protein_coding | 4204,959536 | 0,148370844 | 0,057818982 | 2,566126886 | 0,01028412 | 0,030652179 |
| ENSG00000140463.14 | BBS4 | protein_coding | 3701,53806 | -0,276970558 | 0,065832794 | -4,207182193 | 2,59E-05 | 0,000135639 |
| ENSG00000140464.19 | PML | protein_coding | 4615,260116 | 2,925074105 | 0,070229032 | 41,65049707 | 0 | 0 |
| ENSG00000140465.14 | CYP1A1 | protein_coding | 12499,42496 | 0,367333642 | 0,050252244 | 7,309795741 | 2,68E-13 | 3,54E-12 |
| ENSG00000140474.14 | ULK3 | protein_coding | 4699,475098 | -0,294461175 | 0,056020684 | -5,2562938 | 1,47E-07 | 1,06E-06 |
| ENSG00000140481.15 | CCDC33 | protein_coding | 1058,170266 | -0,317605697 | 0,084758368 | -3,747189852 | 0,000178827 | 0,000809088 |
| ENSG00000140497.16 | SCAMP2 | protein_coding | 6843,451859 | 0,21974365 | 0,06574304 | 3,342462572 | 0,000830386 | 0,003277424 |
| ENSG00000140511.11 | HAPLN3 | protein_coding | 33,51319327 | 2,640998573 | 0,43969995 | 6,006365414 | 1,90E-09 | 1,72E-08 |
| ENSG00000140525.17 | FANCI | protein_coding | 17573,15834 | -0,306765555 | 0,04865895 | -6,304401449 | 2,89E-10 | 2,88E-09 |
| ENSG00000140526.18 | ABHD2 | protein_coding | 25331,84564 | -0,437677977 | 0,057239327 | -7,646455674 | 2,07E-14 | 2,96E-13 |

| | | | | | | | | |
|---|---|---|---|---|---|---|---|---|
| ENSG00000140575.13 | IQGAP1 | protein_coding | 33492,00199 | -0,296256747 | 0,043593403 | -6,79590781 | 1,08E-11 | 1,23E-10 |
| ENSG00000140577.16 | CRTC3 | protein_coding | 6271,277506 | 0,135127133 | 0,053410241 | 2,529985457 | 0,011406725 | 0,03344893 |
| ENSG00000140600.17 | SH3GL3 | protein_coding | 555,947896 | -0,780832787 | 0,110464295 | -7,068644101 | 1,56E-12 | 1,95E-11 |
| ENSG00000140612.14 | SEC11A | protein_coding | 17316,18903 | -0,34938419 | 0,042997221 | -8,125738797 | 4,45E-16 | 7,19E-15 |
| ENSG00000140650.12 | PMM2 | protein_coding | 7219,879827 | 0,172073225 | 0,050090651 | 3,435236365 | 0,000592037 | 0,002415473 |
| ENSG00000140718.21 | FTO | protein_coding | 6951,507365 | -0,294740431 | 0,048757057 | -6,045082524 | 1,49E-09 | 1,37E-08 |
| ENSG00000140750.17 | ARHGAP17 | protein_coding | 4401,166378 | 0,195248555 | 0,054809437 | 3,562316354 | 0,000367597 | 0,001569025 |
| ENSG00000140807.7 | NKD1 | protein_coding | 1092,067651 | -0,46279592 | 0,095419949 | -4,8500961 | 1,23E-06 | 7,92E-06 |
| ENSG00000140830.9 | TXNL4B | protein_coding | 2876,258763 | 0,369851544 | 0,064010629 | 5,77797082 | 7,56E-09 | 6,38E-08 |
| ENSG00000140832.9 | MARVELD3 | protein_coding | 3462,520371 | 0,145725226 | 0,061181293 | 2,381859223 | 0,017225481 | 0,047808675 |
| ENSG00000140836.17 | ZFHX3 | protein_coding | 3821,578523 | 0,433081776 | 0,057547657 | 7,525619557 | 5,25E-14 | 7,37E-13 |
| ENSG00000140848.17 | CPNE2 | protein_coding | 623,9423611 | -0,33390945 | 0,109798017 | -3,04112459 | 0,002356963 | 0,008353029 |
| ENSG00000140853.15 | NLRC5 | protein_coding | 6173,080011 | 7,459927252 | 0,495387676 | 15,05876632 | 3,02E-51 | 1,80E-49 |
| ENSG00000140859.16 | KIFC3 | protein_coding | 2783,647215 | 0,322874408 | 0,064049991 | 5,040975068 | 4,63E-07 | 3,16E-06 |
| ENSG00000140876.11 | AC092724.1 | protein_coding | 922,3788321 | 0,312244811 | 0,087524083 | 3,567530215 | 0,000360362 | 0,001540331 |
| ENSG00000140905.10 | GCSH | protein_coding | 2776,806858 | -0,545280685 | 0,06359044 | -8,574884631 | 9,92E-18 | 1,77E-16 |
| ENSG00000140941.13 | MAP1LC3B | protein_coding | 8128,699084 | 0,196027283 | 0,051664961 | 3,794201697 | 0,000148119 | 0,000683435 |
| ENSG00000140961.13 | OSGIN1 | protein_coding | 360,1937473 | 0,435554872 | 0,124277694 | 3,504690642 | 0,000457138 | 0,00191508 |
| ENSG00000140968.11 | IRF8 | protein_coding | 2686,362865 | 0,589634083 | 0,060715146 | 9,711482562 | 2,69E-22 | 6,17E-21 |
| ENSG00000140983.14 | RHOT2 | protein_coding | 7653,178227 | -0,160366341 | 0,048893736 | -3,279895413 | 0,001038456 | 0,004009365 |
| ENSG00000140986.8 | RPL3L | protein_coding | 16,8879583 | 1,541817627 | 0,617203502 | 2,498070121 | 0,012487149 | 0,036225446 |
| ENSG00000140987.20 | ZSCAN32 | protein_coding | 4115,647522 | 0,244717365 | 0,05548752 | 4,410313644 | 1,03E-05 | 5,78E-05 |
| ENSG00000140988.16 | RPS2 | protein_coding | 41769,33722 | -0,637214083 | 0,155597982 | -4,095259294 | 4,22E-05 | 0,000214217 |
| ENSG00000140990.15 | NDUFB10 | protein_coding | 4927,549242 | -0,223576657 | 0,051905787 | -4,30735514 | 1,65E-05 | 8,94E-05 |
| ENSG00000140993.11 | TIGD7 | protein_coding | 2428,352297 | 0,342508792 | 0,066276436 | 5,167881898 | 2,37E-07 | 1,67E-06 |
| ENSG00000141013.17 | GAS8 | protein_coding | 3223,32496 | 0,157486191 | 0,061357434 | 2,566701059 | 0,010267108 | 0,030609582 |
| ENSG00000141027.21 | NCOR1 | protein_coding | 22470,67101 | 0,202015373 | 0,044556198 | 4,533945532 | 5,79E-06 | 3,36E-05 |
| ENSG00000141030.13 | COPS3 | protein_coding | 10727,78435 | -0,292118127 | 0,046332189 | -6,304863504 | 2,88E-10 | 2,88E-09 |
| ENSG00000141086.18 | CTRL | protein_coding | 47,5533678 | 1,92211657 | 0,502603349 | 3,824321057 | 0,000131133 | 0,000611239 |
| ENSG00000141179.13 | PCTP | protein_coding | 3861,23216 | 0,395833256 | 0,055841572 | 7,08850494 | 1,36E-12 | 1,70E-11 |
| ENSG00000141258.13 | SGSM2 | protein_coding | 6012,430856 | -0,215694983 | 0,060121987 | -3,587622335 | 0,000333707 | 0,001435899 |
| ENSG00000141298.19 | SSH2 | protein_coding | 5673,134215 | -0,131161412 | 0,054226283 | -2,418779339 | 0,015572684 | 0,043760991 |
| ENSG00000141376.22 | BCAS3 | protein_coding | 769,3039054 | -0,277682741 | 0,100575831 | -2,76092911 | 0,005763719 | 0,01850457 |
| ENSG00000141385.9 | AFG3L2 | protein_coding | 9915,528035 | 0,132977977 | 0,044255035 | 3,004810156 | 0,002657467 | 0,009299894 |
| ENSG00000141401.12 | IMPA2 | protein_coding | 1577,848991 | -0,458952195 | 0,074865489 | -6,130357236 | 8,77E-10 | 8,25E-09 |
| ENSG00000141425.18 | RPRD1A | protein_coding | 14536,36362 | -0,119446819 | 0,042749748 | -2,79409412 | 0,005204533 | 0,016914356 |

| | | | | | | | | |
|---|---|---|---|---|---|---|---|---|
| ENSG00000141429.13 | GALNT1 | protein_coding | 20873,73188 | -0,117649409 | 0,047315076 | -2,486509987 | 0,012900295 | 0,037232047 |
| ENSG00000141446.11 | ESCO1 | protein_coding | 4886,29772 | 0,600091064 | 0,060388106 | 9,937239353 | 2,87E-23 | 6,84E-22 |
| ENSG00000141452.9 | RMC1 | protein_coding | 4111,391104 | 0,529635093 | 0,060899948 | 8,696806961 | 3,41E-18 | 6,30E-17 |
| ENSG00000141458.13 | NPC1 | protein_coding | 7248,768809 | 0,516627402 | 0,05495171 | 9,401480053 | 5,38E-21 | 1,15E-19 |
| ENSG00000141480.18 | ARRB2 | protein_coding | 2858,027045 | -0,230021008 | 0,064081303 | -3,589518257 | 0,00033129 | 0,001426631 |
| ENSG00000141497.14 | ZMYND15 | protein_coding | 27,05483051 | 1,906507443 | 0,478961877 | 3,980499353 | 6,88E-05 | 0,000336995 |
| ENSG00000141499.16 | WRAP53 | protein_coding | 1795,361225 | -0,242206512 | 0,07005268 | -3,457491028 | 0,00054523 | 0,002241979 |
| ENSG00000141510.17 | TP53 | protein_coding | 3339,635502 | 0,316314462 | 0,068504017 | 4,617458596 | 3,88E-06 | 2,31E-05 |
| ENSG00000141519.15 | CCDC40 | protein_coding | 585,5487919 | -0,576061192 | 0,104522532 | -5,511358952 | 3,56E-08 | 2,78E-07 |
| ENSG00000141522.12 | ARHGDIA | protein_coding | 12629,01043 | -0,139113527 | 0,055529077 | -2,505237529 | 0,012236918 | 0,035595864 |
| ENSG00000141542.11 | RAB40B | protein_coding | 937,9753816 | -0,767349768 | 0,109307389 | -7,020108817 | 2,22E-12 | 2,72E-11 |
| ENSG00000141543.12 | EIF4A3 | protein_coding | 7436,127432 | 0,189308146 | 0,061232097 | 3,091648913 | 0,001990481 | 0,007197388 |
| ENSG00000141551.14 | CSNK1D | protein_coding | 21317,96457 | 0,12671837 | 0,040730291 | 3,111157986 | 0,001863552 | 0,006788589 |
| ENSG00000141552.17 | ANAPC11 | protein_coding | 4744,823358 | -0,578270484 | 0,054258573 | -10,65767947 | 1,61E-26 | 4,45E-25 |
| ENSG00000141556.21 | TBCD | protein_coding | 5509,11903 | -0,438003691 | 0,065760203 | -6,660619461 | 2,73E-11 | 3,00E-10 |
| ENSG00000141562.18 | NARF | protein_coding | 11316,6023 | -0,276176831 | 0,050097849 | -5,512748318 | 3,53E-08 | 2,76E-07 |
| ENSG00000141568.21 | FOXK2 | protein_coding | 7075,221207 | 0,171389587 | 0,047053864 | 3,642412604 | 0,000270095 | 0,001182794 |
| ENSG00000141570.11 | CBX8 | protein_coding | 2643,210716 | -0,369564155 | 0,06586321 | -5,611086309 | 2,01E-08 | 1,62E-07 |
| ENSG00000141574.7 | SECTM1 | protein_coding | 1189,451873 | 2,4474336 | 0,116584902 | 20,99271479 | 7,65E-98 | 8,74E-96 |
| ENSG00000141577.14 | CEP131 | protein_coding | 1866,301957 | -0,409423805 | 0,08170628 | -5,010922127 | 5,42E-07 | 3,66E-06 |
| ENSG00000141582.15 | CBX4 | protein_coding | 1711,843961 | 0,373342413 | 0,066187076 | 5,640714693 | 1,69E-08 | 1,38E-07 |
| ENSG00000141646.13 | SMAD4 | protein_coding | 12778,35805 | 0,283741634 | 0,045707561 | 6,207761382 | 5,37E-10 | 5,20E-09 |
| ENSG00000141655.17 | TNFRSF11A | protein_coding | 11013,32194 | 0,595570784 | 0,044706163 | 13,3218944 | 1,73E-40 | 7,87E-39 |
| ENSG00000141664.9 | ZCCHC2 | protein_coding | 5534,658815 | 1,646555531 | 0,074905373 | 21,98180795 | 4,30E-107 | 5,20E-105 |
| ENSG00000141668.10 | CBLN2 | protein_coding | 1751,827089 | -0,829099636 | 0,071088539 | -11,66291562 | 1,97E-31 | 6,68E-30 |
| ENSG00000141682.11 | PMAIP1 | protein_coding | 15906,40707 | 0,348235921 | 0,059531249 | 5,849632374 | 4,93E-09 | 4,26E-08 |
| ENSG00000141696.13 | P3H4 | protein_coding | 3426,921135 | 0,365454858 | 0,054226048 | 6,739470633 | 1,59E-11 | 1,78E-10 |
| ENSG00000141736.13 | ERBB2 | protein_coding | 1150,020024 | -0,192291668 | 0,076780982 | -2,504417926 | 0,012265306 | 0,035664613 |
| ENSG00000141753.7 | IGFBP4 | protein_coding | 2245,364736 | 0,695095996 | 0,064429756 | 10,78843133 | 3,90E-27 | 1,10E-25 |
| ENSG00000141756.18 | FKBP10 | protein_coding | 2342,553929 | 0,179721717 | 0,066058865 | 2,720629795 | 0,006515769 | 0,02061617 |
| ENSG00000141759.15 | TXNL4A | protein_coding | 5241,496829 | -0,385495844 | 0,05853709 | -6,585497217 | 4,53E-11 | 4,88E-10 |
| ENSG00000141854.9 | MISP3 | protein_coding | 924,4702658 | -0,424970632 | 0,082117165 | -5,175174205 | 2,28E-07 | 1,61E-06 |
| ENSG00000141858.11 | SAMD1 | protein_coding | 2521,638906 | -0,424599995 | 0,073593986 | -5,769493077 | 7,95E-09 | 6,69E-08 |
| ENSG00000141959.17 | PFKL | protein_coding | 11794,14219 | -0,216843713 | 0,065556191 | -3,307753402 | 0,000940476 | 0,003668198 |
| ENSG00000141971.13 | MVB12A | protein_coding | 1397,43333 | 0,536110552 | 0,108841048 | 4,92562836 | 8,41E-07 | 5,51E-06 |
| ENSG00000142039.4 | CCDC97 | protein_coding | 3018,859623 | 0,201164676 | 0,063950852 | 3,145613673 | 0,001657388 | 0,00611877 |

| | | | | | | | | |
|---|---|---|---|---|---|---|---|---|
| ENSG00000142089.16 | IFITM3 | protein_coding | 310,8258487 | 8,650343909 | 0,816275448 | 10,59733443 | 3,07E-26 | 8,41E-25 |
| ENSG00000142102.15 | PGGHG | protein_coding | 382,2757369 | 1,499900332 | 0,129850405 | 11,55098694 | 7,30E-31 | 2,42E-29 |
| ENSG00000142149.9 | HUNK | protein_coding | 1818,324867 | 0,162486128 | 0,068520318 | 2,371356871 | 0,01772291 | 0,048941844 |
| ENSG00000142173.15 | COL6A2 | protein_coding | 1100,423437 | 0,350338005 | 0,095271948 | 3,67724195 | 0,000235769 | 0,001044672 |
| ENSG00000142178.8 | SIK1 | protein_coding | 555,8678047 | 0,69287534 | 0,166703895 | 4,156323648 | 3,23E-05 | 0,000167002 |
| ENSG00000142185.16 | TRPM2 | protein_coding | 3629,438958 | -0,308167684 | 0,061272104 | -5,029494057 | 4,92E-07 | 3,34E-06 |
| ENSG00000142188.17 | TMEM50B | protein_coding | 5059,892782 | 0,424615827 | 0,055477994 | 7,653770405 | 1,95E-14 | 2,80E-13 |
| ENSG00000142192.21 | APP | protein_coding | 88823,13413 | 0,107077139 | 0,037052736 | 2,889857775 | 0,003854162 | 0,01296955 |
| ENSG00000142208.16 | AKT1 | protein_coding | 21315,03704 | -0,30950239 | 0,04800572 | -6,447198176 | 1,14E-10 | 1,18E-09 |
| ENSG00000142230.12 | SAE1 | protein_coding | 17259,03022 | -0,160945542 | 0,047758991 | -3,369952692 | 0,000751811 | 0,002996726 |
| ENSG00000142235.9 | LMTK3 | protein_coding | 128,042074 | 0,710195047 | 0,250898383 | 2,830608305 | 0,004645958 | 0,015277701 |
| ENSG00000142396.10 | ERVK3-1 | protein_coding | 8231,578623 | 0,628329942 | 0,053784746 | 11,68230743 | 1,57E-31 | 5,35E-30 |
| ENSG00000142484.7 | TM4SF5 | protein_coding | 2552,742138 | 0,439599525 | 0,067623521 | 6,50068969 | 8,00E-11 | 8,39E-10 |
| ENSG00000142507.10 | PSMB6 | protein_coding | 7586,388387 | -0,148364343 | 0,046881492 | -3,164667653 | 0,001552602 | 0,00577454 |
| ENSG00000142534.7 | RPS11 | protein_coding | 72520,46037 | -0,510243214 | 0,04026005 | -12,67368561 | 8,27E-37 | 3,37E-35 |
| ENSG00000142541.17 | RPL13A | protein_coding | 72771,0962 | -0,572876735 | 0,044338987 | -12,92038384 | 3,45E-38 | 1,46E-36 |
| ENSG00000142546.13 | NOSIP | protein_coding | 4816,571329 | 0,323774951 | 0,051224181 | 6,320744284 | 2,60E-10 | 2,60E-09 |
| ENSG00000142556.19 | ZNF614 | protein_coding | 1687,848689 | 0,184324453 | 0,072275242 | 2,550312484 | 0,01076264 | 0,031867439 |
| ENSG00000142621.19 | FHAD1 | protein_coding | 434,1486999 | -0,572045926 | 0,126795226 | -4,511573075 | 6,43E-06 | 3,71E-05 |
| ENSG00000142634.13 | EFHD2 | protein_coding | 1963,62961 | -0,341060635 | 0,070005441 | -4,871916069 | 1,11E-06 | 7,14E-06 |
| ENSG00000142655.13 | PEX14 | protein_coding | 1406,710295 | 0,202432304 | 0,082236862 | 2,461576214 | 0,0138328 | 0,039623633 |
| ENSG00000142657.21 | PGD | protein_coding | 19563,98607 | -0,14262175 | 0,046400889 | -3,0736857 | 0,002114321 | 0,007592029 |
| ENSG00000142661.19 | MYOM3 | protein_coding | 5267,781225 | -0,251750178 | 0,054611141 | -4,609868502 | 4,03E-06 | 2,39E-05 |
| ENSG00000142676.14 | RPL11 | protein_coding | 45828,86724 | -0,444612645 | 0,042191084 | -10,53807113 | 5,77E-26 | 1,56E-24 |
| ENSG00000142677.4 | IL22RA1 | protein_coding | 1782,848105 | 2,342442412 | 0,091131719 | 25,7039199 | 1,06E-145 | 1,70E-143 |
| ENSG00000142686.8 | C1orf216 | protein_coding | 870,1813071 | 0,346347497 | 0,094118364 | 3,679914138 | 0,000233312 | 0,0010346 |
| ENSG00000142687.18 | KIAA0319L | protein_coding | 7324,978439 | 0,234084374 | 0,060514188 | 3,86825603 | 0,000109617 | 0,000517809 |
| ENSG00000142694.6 | EVA1B | protein_coding | 1070,127385 | -0,427965583 | 0,107741344 | -3,972157441 | 7,12E-05 | 0,000348067 |
| ENSG00000142731.11 | PLK4 | protein_coding | 4916,16286 | -0,409642957 | 0,059003561 | -6,942681984 | 3,85E-12 | 4,61E-11 |
| ENSG00000142751.15 | GPN2 | protein_coding | 4353,887553 | 0,23841675 | 0,076698543 | 3,108491262 | 0,001880452 | 0,006840255 |
| ENSG00000142765.18 | SYTL1 | protein_coding | 3633,481545 | -0,337344353 | 0,058511236 | -5,765462819 | 8,14E-09 | 6,84E-08 |
| ENSG00000142856.16 | ITGB3BP | protein_coding | 4915,185053 | -0,373670387 | 0,058996694 | -6,333751243 | 2,39E-10 | 2,40E-09 |
| ENSG00000142864.14 | SERBP1 | protein_coding | 41138,08449 | -0,215368079 | 0,04555884 | -4,72725117 | 2,28E-06 | 1,40E-05 |
| ENSG00000142867.14 | BCL10 | protein_coding | 1687,129034 | 0,190700736 | 0,074020413 | 2,576326301 | 0,009985636 | 0,029885048 |
| ENSG00000142892.15 | PIGK | protein_coding | 8094,539758 | -0,144975506 | 0,05561755 | -2,606650358 | 0,009143267 | 0,027729156 |
| ENSG00000142920.17 | AZIN2 | protein_coding | 510,4293641 | 0,359554157 | 0,15047609 | 2,389443778 | 0,016873906 | 0,046966088 |

| | | | | | | | | |
|---|---|---|---|---|---|---|---|---|
| ENSG00000142937.12 | RPS8 | protein_coding | 38386,31467 | -0,676815184 | 0,04643738 | -14,57479254 | 4,06E-48 | 2,21E-46 |
| ENSG00000142945.13 | KIF2C | protein_coding | 5521,614682 | -0,554163674 | 0,056317278 | -9,840029367 | 7,57E-23 | 1,78E-21 |
| ENSG00000142949.17 | PTPRF | protein_coding | 12836,07491 | -0,540362897 | 0,071697776 | -7,536675909 | 4,82E-14 | 6,79E-13 |
| ENSG00000142961.14 | MOB3C | protein_coding | 2258,779628 | 1,630400015 | 0,073040748 | 22,32178708 | 2,27E-110 | 2,90E-108 |
| ENSG00000143013.13 | LMO4 | protein_coding | 14750,26259 | 0,440311226 | 0,050598878 | 8,70199584 | 3,26E-18 | 6,03E-17 |
| ENSG00000143033.18 | MTF2 | protein_coding | 9079,429105 | 0,213166034 | 0,048621014 | 4,384236699 | 1,16E-05 | 6,46E-05 |
| ENSG00000143036.17 | SLC44A3 | protein_coding | 3954,586741 | -0,19512 | 0,059610822 | -3,273231147 | 0,001063255 | 0,004097381 |
| ENSG00000143079.15 | CTTNBP2NL | protein_coding | 5278,41389 | 0,303154779 | 0,05508853 | 5,50304716 | 3,73E-08 | 2,91E-07 |
| ENSG00000143106.13 | PSMA5 | protein_coding | 11422,99741 | 0,478624367 | 0,042921358 | 11,15119354 | 7,07E-29 | 2,13E-27 |
| ENSG00000143127.13 | ITGA10 | protein_coding | 20,77902825 | -1,644018795 | 0,509530396 | -3,226537234 | 0,001252979 | 0,004749631 |
| ENSG00000143149.12 | ALDH9A1 | protein_coding | 15264,43054 | -0,228791152 | 0,041487519 | -5,514698391 | 3,49E-08 | 2,73E-07 |
| ENSG00000143153.12 | ATP1B1 | protein_coding | 15793,35125 | 0,241063801 | 0,052745145 | 4,570350512 | 4,87E-06 | 2,86E-05 |
| ENSG00000143155.13 | TIPRL | protein_coding | 9341,60232 | -0,123400822 | 0,045070203 | -2,7379691 | 0,006181988 | 0,019687377 |
| ENSG00000143157.11 | POGK | protein_coding | 12392,06076 | 0,156348652 | 0,048188484 | 3,244523146 | 0,001176475 | 0,004483031 |
| ENSG00000143162.9 | CREG1 | protein_coding | 6848,687558 | 0,249121663 | 0,048807314 | 5,104187094 | 3,32E-07 | 2,31E-06 |
| ENSG00000143164.15 | DCAF6 | protein_coding | 4689,101231 | 0,425224053 | 0,055563342 | 7,652960323 | 1,96E-14 | 2,82E-13 |
| ENSG00000143171.13 | RXRG | protein_coding | 1350,430688 | -0,711464345 | 0,08325344 | -8,545765087 | 1,28E-17 | 2,27E-16 |
| ENSG00000143178.13 | TBX19 | protein_coding | 138,0894825 | 0,643055058 | 0,22018119 | 2,920572183 | 0,003493892 | 0,011892337 |
| ENSG00000143190.23 | POU2F1 | protein_coding | 6012,885858 | -0,497889812 | 0,065449644 | -7,607219616 | 2,80E-14 | 3,98E-13 |
| ENSG00000143195.13 | ILDR2 | protein_coding | 2367,878743 | -0,805808962 | 0,091861691 | -8,771980478 | 1,76E-18 | 3,30E-17 |
| ENSG00000143198.13 | MGST3 | protein_coding | 6548,084104 | -0,210342487 | 0,048456813 | -4,340823803 | 1,42E-05 | 7,75E-05 |
| ENSG00000143222.12 | UFC1 | protein_coding | 7087,806494 | -0,260103436 | 0,047958857 | -5,423470274 | 5,85E-08 | 4,44E-07 |
| ENSG00000143228.13 | NUF2 | protein_coding | 5396,867295 | -0,745787205 | 0,054835031 | -13,60056133 | 3,97E-42 | 1,91E-40 |
| ENSG00000143252.14 | SDHC | protein_coding | 18725,59902 | -0,324938019 | 0,044203182 | -7,351009752 | 1,97E-13 | 2,63E-12 |
| ENSG00000143303.12 | RRNAD1 | protein_coding | 1913,19323 | 0,260524922 | 0,069162911 | 3,766829929 | 0,000165334 | 0,000753491 |
| ENSG00000143314.12 | MRPL24 | protein_coding | 3627,979697 | -0,361632392 | 0,061735647 | -5,857756579 | 4,69E-09 | 4,06E-08 |
| ENSG00000143315.7 | PIGM | protein_coding | 6624,445584 | -0,251951018 | 0,048194513 | -5,227794658 | 1,72E-07 | 1,23E-06 |
| ENSG00000143321.19 | HDGF | protein_coding | 20763,41015 | -0,183387453 | 0,040912699 | -4,482409119 | 7,38E-06 | 4,21E-05 |
| ENSG00000143322.21 | ABL2 | protein_coding | 4766,654851 | 0,324861679 | 0,054038645 | 6,011654736 | 1,84E-09 | 1,66E-08 |
| ENSG00000143324.13 | XPR1 | protein_coding | 16312,16833 | 0,238253317 | 0,047579954 | 5,007430599 | 5,52E-07 | 3,73E-06 |
| ENSG00000143333.7 | RGS16 | protein_coding | 226,4799567 | 0,67742909 | 0,164211232 | 4,125351722 | 3,70E-05 | 0,000189366 |
| ENSG00000143337.18 | TOR1AIP1 | protein_coding | 7611,105582 | 0,160179014 | 0,045010854 | 3,558675299 | 0,00037273 | 0,001589274 |
| ENSG00000143344.15 | RGL1 | protein_coding | 7727,399119 | 0,376690586 | 0,045884198 | 8,209592903 | 2,22E-16 | 3,65E-15 |
| ENSG00000143367.16 | TUFT1 | protein_coding | 4133,431384 | 0,57868918 | 0,056447486 | 10,25181501 | 1,16E-24 | 2,97E-23 |
| ENSG00000143373.18 | ZNF687 | protein_coding | 5854,533509 | 0,69698472 | 0,062981364 | 11,06652306 | 1,82E-28 | 5,42E-27 |
| ENSG00000143375.15 | CGN | protein_coding | 8593,275718 | 0,287030444 | 0,055360178 | 5,184781799 | 2,16E-07 | 1,54E-06 |

| | | | | | | | | |
|---|---|---|---|---|---|---|---|---|
| ENSG00000143384.13 | MCL1 | protein_coding | 29541,0477 | 0,410329861 | 0,052856826 | 7,763043892 | 8,29E-15 | 1,23E-13 |
| ENSG00000143390.17 | RFX5 | protein_coding | 7532,299819 | 1,249370662 | 0,071409336 | 17,49590092 | 1,54E-68 | 1,23E-66 |
| ENSG00000143393.16 | PI4KB | protein_coding | 7166,376168 | 0,263831042 | 0,051956409 | 5,077930641 | 3,82E-07 | 2,62E-06 |
| ENSG00000143398.20 | PIP5K1A | protein_coding | 10477,34428 | 0,126727711 | 0,044961042 | 2,818611542 | 0,004823185 | 0,015795878 |
| ENSG00000143401.15 | ANP32E | protein_coding | 11043,66014 | -0,609319754 | 0,044943425 | -13,55748379 | 7,15E-42 | 3,39E-40 |
| ENSG00000143412.10 | ANXA9 | protein_coding | 174,5458999 | -0,524879786 | 0,208481798 | -2,517628834 | 0,011814775 | 0,034519571 |
| ENSG00000143420.18 | ENSA | protein_coding | 34091,78031 | -0,212671107 | 0,043357243 | -4,905088383 | 9,34E-07 | 6,09E-06 |
| ENSG00000143442.22 | POGZ | protein_coding | 11607,36511 | 0,197608485 | 0,057935983 | 3,410807484 | 0,000647708 | 0,002622022 |
| ENSG00000143457.11 | GOLPH3L | protein_coding | 3045,973323 | 0,36245126 | 0,075898344 | 4,775483127 | 1,79E-06 | 1,12E-05 |
| ENSG00000143458.12 | GABPB2 | protein_coding | 1127,910821 | -0,443436782 | 0,099715123 | -4,4470364 | 8,71E-06 | 4,92E-05 |
| ENSG00000143473.13 | KCNH1 | protein_coding | 871,7243386 | -0,233008363 | 0,095875226 | -2,430329215 | 0,015085114 | 0,042625009 |
| ENSG00000143476.18 | DTL | protein_coding | 9668,4733 | -0,154012814 | 0,04619351 | -3,334079087 | 0,000855823 | 0,003368962 |
| ENSG00000143479.17 | DYRK3 | protein_coding | 3416,787055 | 0,390127866 | 0,057644888 | 6,767779018 | 1,31E-11 | 1,48E-10 |
| ENSG00000143486.16 | EIF2D | protein_coding | 3506,05874 | -0,314328895 | 0,054236922 | -5,795478079 | 6,81E-09 | 5,79E-08 |
| ENSG00000143493.13 | INTS7 | protein_coding | 10870,90152 | 0,232589449 | 0,053776858 | 4,325084357 | 1,52E-05 | 8,29E-05 |
| ENSG00000143494.15 | VASH2 | protein_coding | 1083,950456 | -0,601398789 | 0,088433602 | -6,800568704 | 1,04E-11 | 1,19E-10 |
| ENSG00000143507.18 | DUSP10 | protein_coding | 1342,278328 | 0,694278323 | 0,080930092 | 8,578741345 | 9,59E-18 | 1,72E-16 |
| ENSG00000143514.17 | TP53BP2 | protein_coding | 6488,716922 | 0,153875673 | 0,050231677 | 3,063319448 | 0,002188963 | 0,007824576 |
| ENSG00000143515.17 | ATP8B2 | protein_coding | 6879,634333 | 0,262610672 | 0,052445991 | 5,007259236 | 5,52E-07 | 3,73E-06 |
| ENSG00000143537.13 | ADAM15 | protein_coding | 823,1084324 | 0,29921995 | 0,112358248 | 2,663088425 | 0,007742706 | 0,023979166 |
| ENSG00000143545.8 | RAB13 | protein_coding | 4017,289202 | -0,237832188 | 0,052393429 | -4,53935145 | 5,64E-06 | 3,28E-05 |
| ENSG00000143549.21 | TPM3 | protein_coding | 27585,44447 | -0,195162976 | 0,04345595 | -4,491053081 | 7,09E-06 | 4,06E-05 |
| ENSG00000143614.10 | GATAD2B | protein_coding | 5115,267218 | 0,171165185 | 0,060662313 | 2,821606619 | 0,004778375 | 0,015671928 |
| ENSG00000143621.17 | ILF2 | protein_coding | 28172,37847 | -0,192811161 | 0,046209865 | -4,17251083 | 3,01E-05 | 0,000156174 |
| ENSG00000143622.11 | RIT1 | protein_coding | 5703,372126 | 0,294079077 | 0,049775529 | 5,908105511 | 3,46E-09 | 3,04E-08 |
| ENSG00000143627.19 | PKLR | protein_coding | 1155,156096 | -0,372265192 | 0,079027426 | -4,710582268 | 2,47E-06 | 1,52E-05 |
| ENSG00000143630.10 | HCN3 | protein_coding | 4038,024597 | 0,481435661 | 0,068651888 | 7,012708197 | 2,34E-12 | 2,86E-11 |
| ENSG00000143633.13 | C1orf131 | protein_coding | 2563,861166 | 0,236338176 | 0,058619068 | 4,031762771 | 5,54E-05 | 0,000275503 |
| ENSG00000143643.13 | TTC13 | protein_coding | 4435,764783 | -0,46462091 | 0,062998052 | -7,375163095 | 1,64E-13 | 2,21E-12 |
| ENSG00000143653.10 | SCCPDH | protein_coding | 9209,393934 | -0,226902764 | 0,053911955 | -4,208765293 | 2,57E-05 | 0,000134881 |
| ENSG00000143669.13 | LYST | protein_coding | 4672,15636 | 0,40816707 | 0,052519964 | 7,771655586 | 7,75E-15 | 1,16E-13 |
| ENSG00000143727.16 | ACP1 | protein_coding | 27353,83376 | -0,14859744 | 0,047624543 | -3,120186169 | 0,001807368 | 0,006608519 |
| ENSG00000143742.13 | SRP9 | protein_coding | 74606,02807 | -0,441701358 | 0,049832464 | -8,863726945 | 7,74E-19 | 1,48E-17 |
| ENSG00000143751.10 | SDE2 | protein_coding | 3572,52386 | 0,267070996 | 0,057617596 | 4,635233242 | 3,57E-06 | 2,13E-05 |
| ENSG00000143753.13 | DEGS1 | protein_coding | 13261,96773 | 0,50657492 | 0,053674954 | 9,437826823 | 3,81E-21 | 8,20E-20 |
| ENSG00000143756.12 | FBXO28 | protein_coding | 5909,976982 | 0,23971594 | 0,049080567 | 4,884131485 | 1,04E-06 | 6,74E-06 |

| | | | | | | | | |
|---|---|---|---|---|---|---|---|---|
| ENSG00000143774.16 | GUK1 | protein_coding | 6935,576153 | -0,444892073 | 0,061541436 | -7,229146745 | 4,86E-13 | 6,30E-12 |
| ENSG00000143776.18 | CDC42BPA | protein_coding | 17615,78177 | -0,386066335 | 0,054639918 | -7,065646352 | 1,60E-12 | 1,99E-11 |
| ENSG00000143797.12 | MBOAT2 | protein_coding | 25935,34345 | -0,142985488 | 0,04147012 | -3,447915961 | 0,00056493 | 0,002316634 |
| ENSG00000143799.13 | PARP1 | protein_coding | 59200,45435 | -0,248534245 | 0,037915043 | -6,555030018 | 5,56E-11 | 5,94E-10 |
| ENSG00000143801.17 | PSEN2 | protein_coding | 2470,549763 | 0,410054456 | 0,064614796 | 6,346138632 | 2,21E-10 | 2,22E-09 |
| ENSG00000143815.15 | LBR | protein_coding | 10733,31041 | -0,293968111 | 0,046536501 | -6,316936295 | 2,67E-10 | 2,66E-09 |
| ENSG00000143816.8 | WNT9A | protein_coding | 902,550553 | 0,508001479 | 0,087176856 | 5,827251664 | 5,63E-09 | 4,84E-08 |
| ENSG00000143845.15 | ETNK2 | protein_coding | 2179,588191 | -0,337736743 | 0,079967542 | -4,223422849 | 2,41E-05 | 0,000127019 |
| ENSG00000143847.15 | PPFIA4 | protein_coding | 711,5226429 | -0,5619249 | 0,128876862 | -4,360169019 | 1,30E-05 | 7,16E-05 |
| ENSG00000143858.12 | SYT2 | protein_coding | 231,888884 | -0,793906204 | 0,168604291 | -4,708695125 | 2,49E-06 | 1,53E-05 |
| ENSG00000143862.8 | ARL8A | protein_coding | 1560,43864 | 0,281020736 | 0,069386636 | 4,050070044 | 5,12E-05 | 0,000256114 |
| ENSG00000143870.12 | PDIA6 | protein_coding | 63162,95773 | 0,467958879 | 0,040542971 | 11,5422937 | 8,07E-31 | 2,67E-29 |
| ENSG00000143891.17 | GALM | protein_coding | 1249,105462 | 0,57971202 | 0,084438607 | 6,865485316 | 6,63E-12 | 7,75E-11 |
| ENSG00000143924.19 | EML4 | protein_coding | 14707,50602 | 0,218779691 | 0,043075259 | 5,079010437 | 3,79E-07 | 2,61E-06 |
| ENSG00000143933.18 | CALM2 | protein_coding | 96301,19008 | -0,196134186 | 0,039670084 | -4,944133421 | 7,65E-07 | 5,05E-06 |
| ENSG00000143947.13 | RPS27A | protein_coding | 45892,59584 | -0,668484048 | 0,051127108 | -13,07494354 | 4,58E-39 | 2,00E-37 |
| ENSG00000143951.16 | WDPCP | protein_coding | 878,2105424 | -0,392825249 | 0,093663107 | -4,194023269 | 2,74E-05 | 0,00014299 |
| ENSG00000143952.20 | VPS54 | protein_coding | 12924,97367 | 0,258203124 | 0,045945377 | 5,619784652 | 1,91E-08 | 1,55E-07 |
| ENSG00000143977.14 | SNRPG | protein_coding | 11400,88962 | -0,197931609 | 0,047043282 | -4,20743628 | 2,58E-05 | 0,00013555 |
| ENSG00000143995.20 | MEIS1 | protein_coding | 1914,588859 | 0,326368823 | 0,085769624 | 3,805179589 | 0,000141701 | 0,000655882 |
| ENSG00000144034.14 | TPRKB | protein_coding | 3081,581895 | -0,379311381 | 0,062325141 | -6,086009226 | 1,16E-09 | 1,07E-08 |
| ENSG00000144036.15 | EXOC6B | protein_coding | 4670,347171 | 0,13997709 | 0,05415754 | 2,584627925 | 0,009748413 | 0,02927255 |
| ENSG00000144040.12 | SFXN5 | protein_coding | 3257,66278 | 0,138495797 | 0,054286853 | 2,551184852 | 0,010735737 | 0,031796145 |
| ENSG00000144043.12 | TEX261 | protein_coding | 12868,01408 | -0,30009826 | 0,049919064 | -6,011696475 | 1,84E-09 | 1,66E-08 |
| ENSG00000144048.10 | DUSP11 | protein_coding | 2662,588268 | 0,150462326 | 0,061338684 | 2,45297608 | 0,014167976 | 0,040449895 |
| ENSG00000144061.12 | NPHP1 | protein_coding | 415,7888076 | -0,509878653 | 0,126393634 | -4,034053266 | 5,48E-05 | 0,000273071 |
| ENSG00000144063.3 | MALL | protein_coding | 5255,531082 | 0,319158853 | 0,059184446 | 5,392613716 | 6,94E-08 | 5,22E-07 |
| ENSG00000144161.13 | ZC3H8 | protein_coding | 2518,011875 | 0,289831671 | 0,073866798 | 3,923706986 | 8,72E-05 | 0,000419832 |
| ENSG00000144182.17 | LIPT1 | protein_coding | 368,5425811 | 0,374373842 | 0,145156379 | 2,579107066 | 0,009905608 | 0,02967727 |
| ENSG00000144191.11 | CNGA3 | protein_coding | 8638,337953 | 0,154469282 | 0,048345513 | 3,195111015 | 0,001397771 | 0,005242863 |
| ENSG00000144224.17 | UBXN4 | protein_coding | 47805,6135 | -0,109191188 | 0,03843874 | -2,840654708 | 0,004502103 | 0,014861099 |
| ENSG00000144227.5 | NXPH2 | protein_coding | 20,69324597 | -1,561529364 | 0,546819055 | -2,855660111 | 0,004294746 | 0,014275021 |
| ENSG00000144231.11 | POLR2D | protein_coding | 9377,361105 | -0,11734504 | 0,049328933 | -2,378827853 | 0,017367785 | 0,048126489 |
| ENSG00000144278.15 | GALNT13 | protein_coding | 11090,66878 | -0,263613389 | 0,048678054 | -5,415446317 | 6,11E-08 | 4,64E-07 |
| ENSG00000144283.21 | PKP4 | protein_coding | 4194,311079 | -0,409379955 | 0,06376994 | -6,419638413 | 1,37E-10 | 1,40E-09 |
| ENSG00000144290.17 | SLC4A10 | protein_coding | 2648,694111 | 0,218306513 | 0,062484368 | 3,49377806 | 0,000476237 | 0,001982913 |

| | | | | | | | | |
|---|---|---|---|---|---|---|---|---|
| ENSG00000144306.14 | SCRN3 | protein_coding | 3631,04983 | -0,295497166 | 0,066429305 | -4,448295308 | 8,66E-06 | 4,89E-05 |
| ENSG00000144320.14 | LNPK | protein_coding | 12897,66456 | -0,472271713 | 0,070417208 | -6,706765646 | 1,99E-11 | 2,22E-10 |
| ENSG00000144339.12 | TMEFF2 | protein_coding | 985,7768608 | -0,350155906 | 0,085440704 | -4,098232939 | 4,16E-05 | 0,00021158 |
| ENSG00000144354.14 | CDCA7 | protein_coding | 2774,845462 | -0,393967768 | 0,06533923 | -6,029574687 | 1,64E-09 | 1,50E-08 |
| ENSG00000144369.13 | FAM171B | protein_coding | 2748,743914 | -0,157718803 | 0,059331865 | -2,65824787 | 0,00785481 | 0,02427468 |
| ENSG00000144381.17 | HSPD1 | protein_coding | 75412,03022 | -0,181076441 | 0,0417941 | -4,332583831 | 1,47E-05 | 8,03E-05 |
| ENSG00000144395.18 | CCDC150 | protein_coding | 2653,848542 | 0,251417127 | 0,093650488 | 2,684632311 | 0,007260964 | 0,022664961 |
| ENSG00000144401.14 | METTL21A | protein_coding | 2294,632404 | -0,367983403 | 0,093298751 | -3,944140723 | 8,01E-05 | 0,000388005 |
| ENSG00000144407.9 | PTH2R | protein_coding | 4559,220673 | -0,347754906 | 0,051456245 | -6,758264385 | 1,40E-11 | 1,58E-10 |
| ENSG00000144445.17 | KANSL1L | protein_coding | 3200,759099 | 0,253868306 | 0,062402189 | 4,068259622 | 4,74E-05 | 0,000238087 |
| ENSG00000144451.19 | SPAG16 | protein_coding | 4792,874795 | -0,182896334 | 0,052364591 | -3,492748234 | 0,000478077 | 0,001989469 |
| ENSG00000144455.14 | SUMF1 | protein_coding | 2265,401915 | -0,681311942 | 0,067472154 | -10,09767581 | 5,66E-24 | 1,41E-22 |
| ENSG00000144468.17 | RHBDD1 | protein_coding | 3321,404181 | 0,597267859 | 0,061691109 | 9,681587299 | 3,61E-22 | 8,18E-21 |
| ENSG00000144481.17 | TRPM8 | protein_coding | 162,2031144 | -1,875882256 | 0,275810554 | -6,801343273 | 1,04E-11 | 1,19E-10 |
| ENSG00000144504.15 | ANKMY1 | protein_coding | 495,0466233 | -0,458857983 | 0,154179067 | -2,976136733 | 0,002919047 | 0,010111546 |
| ENSG00000144535.19 | DIS3L2 | protein_coding | 2967,896769 | -0,141662171 | 0,057188319 | -2,477117267 | 0,013244838 | 0,038133545 |
| ENSG00000144550.13 | CPNE9 | protein_coding | 33,64223493 | 1,531205228 | 0,455714932 | 3,360006709 | 0,000779406 | 0,003096302 |
| ENSG00000144554.10 | FANCD2 | protein_coding | 4175,506814 | -0,232655486 | 0,057208113 | -4,066826809 | 4,77E-05 | 0,000239448 |
| ENSG00000144566.11 | RAB5A | protein_coding | 9974,971459 | 0,128539965 | 0,046256818 | 2,778832833 | 0,005455459 | 0,017620452 |
| ENSG00000144583.5 | mars-04 | protein_coding | 1410,551463 | 0,220155559 | 0,071711716 | 3,070008219 | 0,002140529 | 0,007670938 |
| ENSG00000144591.18 | GMPPA | protein_coding | 4821,069942 | 0,211393464 | 0,051839667 | 4,077832219 | 4,55E-05 | 0,000229418 |
| ENSG00000144596.13 | GRIP2 | protein_coding | 9,232328008 | 3,808323058 | 0,926320997 | 4,111234734 | 3,94E-05 | 0,000200643 |
| ENSG00000144597.14 | EAF1 | protein_coding | 9558,151137 | 0,155314689 | 0,046394781 | 3,347675877 | 0,000814923 | 0,003223742 |
| ENSG00000144645.13 | OSBPL10 | protein_coding | 2919,70955 | 0,226844365 | 0,077653999 | 2,921219351 | 0,003486643 | 0,01187125 |
| ENSG00000144655.15 | CSRNP1 | protein_coding | 1802,28049 | 0,422595282 | 0,07071018 | 5,976441878 | 2,28E-09 | 2,04E-08 |
| ENSG00000144659.13 | SLC25A38 | protein_coding | 3566,483953 | 0,375367029 | 0,058602124 | 6,405348526 | 1,50E-10 | 1,54E-09 |
| ENSG00000144674.16 | GOLGA4 | protein_coding | 20200,18511 | 0,238037675 | 0,043201048 | 5,509997644 | 3,59E-08 | 2,80E-07 |
| ENSG00000144711.16 | IQSEC1 | protein_coding | 2529,446479 | -0,480714115 | 0,078566816 | -6,118538861 | 9,44E-10 | 8,84E-09 |
| ENSG00000144712.12 | CAND2 | protein_coding | 740,4420011 | -0,295744234 | 0,119281772 | -2,479374919 | 0,013161288 | 0,037925364 |
| ENSG00000144713.12 | RPL32 | protein_coding | 31548,97245 | -0,523643115 | 0,054559471 | -9,597657452 | 8,18E-22 | 1,83E-20 |
| ENSG00000144746.7 | ARL6IP5 | protein_coding | 14256,63774 | -0,182064022 | 0,04722045 | -3,855618109 | 0,000115438 | 0,000543141 |
| ENSG00000144747.17 | TMF1 | protein_coding | 10441,2981 | 0,179105044 | 0,05199177 | 3,444872981 | 0,000571328 | 0,002341591 |
| ENSG00000144749.13 | LRIG1 | protein_coding | 3465,564747 | -0,141130298 | 0,056933682 | -2,478854214 | 0,013180517 | 0,037972642 |
| ENSG00000144791.10 | LIMD1 | protein_coding | 3814,320231 | 0,430129981 | 0,059721941 | 7,202210377 | 5,92E-13 | 7,62E-12 |
| ENSG00000144802.11 | NFKBIZ | protein_coding | 5220,75803 | 1,158343594 | 0,059314544 | 19,52882925 | 6,25E-85 | 6,19E-83 |
| ENSG00000144815.16 | NXPE3 | protein_coding | 2923,328233 | 0,292441054 | 0,075417102 | 3,877649051 | 0,000105471 | 0,000499588 |

| | | | | | | | | |
|---|---|---|---|---|---|---|---|---|
| ENSG00000144827.9 | ABHD10 | protein_coding | 7624,747264 | -0,202190396 | 0,052450944 | -3,854847642 | 0,000115802 | 0,00054474 |
| ENSG00000144837.9 | PLA1A | protein_coding | 523,5667605 | 2,175746405 | 0,125310499 | 17,36284207 | 1,58E-67 | 1,22E-65 |
| ENSG00000144840.9 | RABL3 | protein_coding | 3183,663156 | -0,443051728 | 0,056994387 | -7,773602873 | 7,63E-15 | 1,14E-13 |
| ENSG00000144843.11 | ADPRH | protein_coding | 500,0724542 | 0,64756987 | 0,127096118 | 5,095119211 | 3,49E-07 | 2,41E-06 |
| ENSG00000144847.12 | IGSF11 | protein_coding | 5312,375997 | 0,190493762 | 0,048369967 | 3,93826524 | 8,21E-05 | 0,000397029 |
| ENSG00000144857.14 | BOC | protein_coding | 177,4300516 | -1,033509689 | 0,207546029 | -4,979664963 | 6,37E-07 | 4,26E-06 |
| ENSG00000144891.17 | AGTR1 | protein_coding | 41,09702289 | 1,451739776 | 0,404890298 | 3,585513862 | 0,000336415 | 0,001446765 |
| ENSG00000144893.12 | MED12L | protein_coding | 3810,782299 | 0,397108297 | 0,056543772 | 7,023024572 | 2,17E-12 | 2,66E-11 |
| ENSG00000144959.10 | NCEH1 | protein_coding | 1644,891501 | 0,499475445 | 0,073849418 | 6,763430991 | 1,35E-11 | 1,53E-10 |
| ENSG00000145016.16 | RUBCN | protein_coding | 6420,526119 | 0,203621078 | 0,060503233 | 3,365457797 | 0,000764168 | 0,003041671 |
| ENSG00000145050.17 | MANF | protein_coding | 13593,97651 | 0,35214185 | 0,053883235 | 6,535276747 | 6,35E-11 | 6,74E-10 |
| ENSG00000145087.12 | STXBP5L | protein_coding | 12420,16558 | -0,347477933 | 0,049696304 | -6,992027659 | 2,71E-12 | 3,30E-11 |
| ENSG00000145088.9 | EAF2 | protein_coding | 1178,863887 | 0,482472428 | 0,076800842 | 6,282124181 | 3,34E-10 | 3,31E-09 |
| ENSG00000145103.15 | ILDR1 | protein_coding | 686,0213772 | 0,620647525 | 0,106565556 | 5,824091279 | 5,74E-09 | 4,93E-08 |
| ENSG00000145214.14 | DGKQ | protein_coding | 2218,621905 | 0,250457472 | 0,062595238 | 4,001222431 | 6,30E-05 | 0,000310923 |
| ENSG00000145216.16 | FIP1L1 | protein_coding | 7309,744583 | 0,174823005 | 0,049246548 | 3,549954509 | 0,000385298 | 0,001636352 |
| ENSG00000145246.13 | ATP10D | protein_coding | 11178,583 | 0,621512669 | 0,059590132 | 10,42979175 | 1,81E-25 | 4,80E-24 |
| ENSG00000145247.11 | OCIAD2 | protein_coding | 8159,68542 | -0,167005455 | 0,054334293 | -3,073665723 | 0,002114462 | 0,007592029 |
| ENSG00000145248.7 | SLC10A4 | protein_coding | 43,00857055 | -1,069741806 | 0,424957322 | -2,517292326 | 0,011826066 | 0,034543593 |
| ENSG00000145284.12 | SCD5 | protein_coding | 1376,210949 | -0,791999473 | 0,073930339 | -10,71278019 | 8,87E-27 | 2,48E-25 |
| ENSG00000145287.10 | PLAC8 | protein_coding | 755,666871 | 0,334156625 | 0,108578658 | 3,077553448 | 0,002087074 | 0,007502068 |
| ENSG00000145293.16 | ENOPH1 | protein_coding | 10928,73782 | -0,212150555 | 0,045161549 | -4,697592508 | 2,63E-06 | 1,61E-05 |
| ENSG00000145321.13 | GC | protein_coding | 25454,68544 | 0,712965316 | 0,048962013 | 14,56160138 | 4,93E-48 | 2,67E-46 |
| ENSG00000145331.14 | TRMT10A | protein_coding | 1028,893516 | -0,457348906 | 0,087020158 | -5,255666239 | 1,47E-07 | 1,07E-06 |
| ENSG00000145332.14 | KLHL8 | protein_coding | 6184,547738 | -0,178915906 | 0,050472446 | -3,544823401 | 0,000392876 | 0,001664142 |
| ENSG00000145335.15 | SNCA | protein_coding | 3121,673919 | 0,260417963 | 0,057783403 | 4,506795177 | 6,58E-06 | 3,78E-05 |
| ENSG00000145362.18 | ANK2 | protein_coding | 9896,183516 | -0,211331073 | 0,068654185 | -3,078196526 | 0,002082575 | 0,007488289 |
| ENSG00000145375.8 | SPATA5 | protein_coding | 1460,246365 | -0,571846216 | 0,082124049 | -6,963200484 | 3,33E-12 | 4,01E-11 |
| ENSG00000145386.10 | CCNA2 | protein_coding | 8988,037538 | -0,721294595 | 0,045114499 | -15,98808842 | 1,55E-57 | 1,05E-55 |
| ENSG00000145388.15 | METTL14 | protein_coding | 10379,63681 | 0,232287018 | 0,050432672 | 4,605883618 | 4,11E-06 | 2,44E-05 |
| ENSG00000145390.11 | USP53 | protein_coding | 3431,638859 | 0,562512952 | 0,076488271 | 7,354238089 | 1,92E-13 | 2,57E-12 |
| ENSG00000145425.10 | RPS3A | protein_coding | 86151,10296 | -0,992828455 | 0,042061549 | -23,60418189 | 3,49E-123 | 4,85E-121 |
| ENSG00000145439.12 | CBR4 | protein_coding | 8149,585632 | -0,330537903 | 0,049852576 | -6,63030738 | 3,35E-11 | 3,65E-10 |
| ENSG00000145476.16 | CYP4V2 | protein_coding | 3841,505524 | 0,313031359 | 0,057092349 | 5,482895043 | 4,18E-08 | 3,23E-07 |
| ENSG00000145545.12 | SRD5A1 | protein_coding | 4629,046419 | -0,455364865 | 0,054744897 | -8,317941776 | 8,95E-17 | 1,51E-15 |
| ENSG00000145555.15 | MYO10 | protein_coding | 16304,56815 | 0,190554401 | 0,046803797 | 4,071344943 | 4,67E-05 | 0,000235446 |

| | | | | | | | | |
|---|---|---|---|---|---|---|---|---|
| ENSG00000145569.6 | OTULINL | protein_coding | 13111,82985 | -0,205854171 | 0,055373338 | -3,717568357 | 0,00020115 | 0,000901394 |
| ENSG00000145592.14 | RPL37 | protein_coding | 32596,1004 | -0,442176589 | 0,039474919 | -11,20145638 | 4,01E-29 | 1,22E-27 |
| ENSG00000145604.15 | SKP2 | protein_coding | 7305,353626 | -0,397632292 | 0,059867771 | -6,64184232 | 3,10E-11 | 3,39E-10 |
| ENSG00000145632.15 | PLK2 | protein_coding | 6881,877918 | 0,468774175 | 0,063473794 | 7,385318349 | 1,52E-13 | 2,06E-12 |
| ENSG00000145642.12 | SHISAL2B | protein_coding | 683,5382382 | -0,371243459 | 0,098732356 | -3,760099252 | 0,000169846 | 0,000772336 |
| ENSG00000145675.15 | PIK3R1 | protein_coding | 10758,84153 | 0,178480224 | 0,055102283 | 3,239071308 | 0,001199196 | 0,004560348 |
| ENSG00000145687.16 | SSBP2 | protein_coding | 5679,106373 | 0,145172866 | 0,054708185 | 2,653585864 | 0,007964151 | 0,024583799 |
| ENSG00000145703.16 | IQGAP2 | protein_coding | 1285,414185 | 0,250990095 | 0,077786425 | 3,226656752 | 0,001252456 | 0,004748448 |
| ENSG00000145708.11 | CRHBP | protein_coding | 236,3895375 | 0,408914537 | 0,168134177 | 2,43207267 | 0,015012695 | 0,042449228 |
| ENSG00000145725.19 | PPIP5K2 | protein_coding | 10490,00575 | -0,475580624 | 0,049459852 | -9,615488304 | 6,88E-22 | 1,55E-20 |
| ENSG00000145730.20 | PAM | protein_coding | 23546,87619 | -0,408709129 | 0,056310853 | -7,258088003 | 3,93E-13 | 5,13E-12 |
| ENSG00000145734.19 | BDP1 | protein_coding | 19211,19466 | 0,336539675 | 0,045613049 | 7,37814465 | 1,61E-13 | 2,17E-12 |
| ENSG00000145741.15 | BTF3 | protein_coding | 29627,42379 | -0,306953043 | 0,039584682 | -7,754338976 | 8,88E-15 | 1,31E-13 |
| ENSG00000145743.15 | FBXL17 | protein_coding | 3591,307732 | -0,249356728 | 0,063594307 | -3,921054262 | 8,82E-05 | 0,000424027 |
| ENSG00000145779.8 | TNFAIP8 | protein_coding | 1248,193907 | -0,345305766 | 0,091995141 | -3,753521767 | 0,000174367 | 0,00079162 |
| ENSG00000145780.8 | FEM1C | protein_coding | 4110,159592 | 0,421963159 | 0,056218367 | 7,505788196 | 6,11E-14 | 8,52E-13 |
| ENSG00000145781.9 | COMMD10 | protein_coding | 2256,718565 | -0,267654324 | 0,070024914 | -3,822272792 | 0,000132227 | 0,000615066 |
| ENSG00000145819.18 | ARHGAP26 | protein_coding | 2893,090593 | 0,294027581 | 0,073090846 | 4,022768892 | 5,75E-05 | 0,00028517 |
| ENSG00000145833.16 | DDX46 | protein_coding | 17420,54604 | -0,295614846 | 0,047186247 | -6,264851862 | 3,73E-10 | 3,67E-09 |
| ENSG00000145882.11 | PCYOX1L | protein_coding | 5922,887403 | -0,255529765 | 0,050452805 | -5,064728602 | 4,09E-07 | 2,81E-06 |
| ENSG00000145901.15 | TNIP1 | protein_coding | 4465,28963 | 0,510210292 | 0,070893278 | 7,196878285 | 6,16E-13 | 7,92E-12 |
| ENSG00000145912.8 | NHP2 | protein_coding | 3959,087033 | -0,327216319 | 0,059147972 | -5,532164613 | 3,16E-08 | 2,49E-07 |
| ENSG00000145949.10 | MYLK4 | protein_coding | 241,7505218 | 0,409894448 | 0,14908337 | 2,749431054 | 0,005969882 | 0,01907938 |
| ENSG00000145979.18 | TBC1D7 | protein_coding | 2303,263399 | -0,220223058 | 0,060801984 | -3,621971568 | 0,000292366 | 0,001272501 |
| ENSG00000145990.11 | GFOD1 | protein_coding | 1890,202605 | 0,409362269 | 0,073733039 | 5,551951689 | 2,82E-08 | 2,24E-07 |
| ENSG00000146006.8 | LRRTM2 | protein_coding | 392,4531645 | 2,861246085 | 0,138941841 | 20,59312058 | 3,16E-94 | 3,51E-92 |
| ENSG00000146066.3 | HIGD2A | protein_coding | 3128,302954 | -0,166376943 | 0,056093585 | -2,966060045 | 0,003016416 | 0,010408785 |
| ENSG00000146072.6 | TNFRSF21 | protein_coding | 7694,923232 | 0,301071276 | 0,046929187 | 6,415437674 | 1,40E-10 | 1,44E-09 |
| ENSG00000146083.12 | RNF44 | protein_coding | 4621,52263 | -0,317799821 | 0,052990107 | -5,997342531 | 2,01E-09 | 1,81E-08 |
| ENSG00000146094.14 | DOK3 | protein_coding | 937,6315032 | 0,281320123 | 0,08909571 | 3,157504707 | 0,001591257 | 0,005898814 |
| ENSG00000146109.5 | ABT1 | protein_coding | 3405,868778 | -0,166263974 | 0,064238971 | -2,588210405 | 0,009647603 | 0,029031727 |
| ENSG00000146151.13 | HMGCLL1 | protein_coding | 14179,29514 | -0,466801513 | 0,053537244 | -8,719192055 | 2,80E-18 | 5,19E-17 |
| ENSG00000146223.14 | RPL7L1 | protein_coding | 28459,28935 | -0,143123848 | 0,042905352 | -3,335804043 | 0,000850531 | 0,003351059 |
| ENSG00000146232.16 | NFKBIE | protein_coding | 1058,312597 | 0,363819709 | 0,095860034 | 3,795322138 | 0,000147452 | 0,000680819 |
| ENSG00000146242.8 | TPBG | protein_coding | 69,27675794 | 0,964162369 | 0,335806097 | 2,871187802 | 0,004089325 | 0,013668936 |
| ENSG00000146243.14 | IRAK1BP1 | protein_coding | 1562,782612 | -0,562983484 | 0,080198637 | -7,019863486 | 2,22E-12 | 2,72E-11 |

| | | | | | | | | |
|---|---|---|---|---|---|---|---|---|
| ENSG00000146247.14 | PHIP | protein_coding | 15993,83472 | -0,211025092 | 0,049504245 | -4,262767637 | 2,02E-05 | 0,000107722 |
| ENSG00000146263.11 | MMS22L | protein_coding | 5985,307674 | -0,187516231 | 0,055676566 | -3,367956098 | 0,000757277 | 0,003015842 |
| ENSG00000146267.12 | FAXC | protein_coding | 2229,338619 | 0,158954237 | 0,061972398 | 2,564919899 | 0,010319963 | 0,030738658 |
| ENSG00000146278.10 | PNRC1 | protein_coding | 2931,845865 | 0,309821812 | 0,064282819 | 4,819667465 | 1,44E-06 | 9,13E-06 |
| ENSG00000146281.6 | PM20D2 | protein_coding | 5986,441365 | -0,565600595 | 0,0568888 | -9,942213569 | 2,73E-23 | 6,52E-22 |
| ENSG00000146285.14 | SCML4 | protein_coding | 303,7929025 | 2,467228894 | 0,158512641 | 15,56487149 | 1,26E-54 | 7,95E-53 |
| ENSG00000146350.14 | TBC1D32 | protein_coding | 2082,818578 | 0,201555734 | 0,082718394 | 2,436649498 | 0,01482404 | 0,041985925 |
| ENSG00000146376.11 | ARHGAP18 | protein_coding | 2832,281098 | 0,308925733 | 0,060679761 | 5,091083571 | 3,56E-07 | 2,46E-06 |
| ENSG00000146386.8 | ABRACL | protein_coding | 2613,872157 | 0,265455402 | 0,066887285 | 3,968697536 | 7,23E-05 | 0,000352852 |
| ENSG00000146409.12 | SLC18B1 | protein_coding | 11083,68766 | 0,580469707 | 0,047666895 | 12,17762782 | 4,09E-34 | 1,51E-32 |
| ENSG00000146411.5 | SLC2A12 | protein_coding | 175,0910531 | 1,931294963 | 0,252327253 | 7,653929335 | 1,95E-14 | 2,80E-13 |
| ENSG00000146416.18 | AIG1 | protein_coding | 7400,092187 | -0,238533473 | 0,052780747 | -4,519327369 | 6,20E-06 | 3,58E-05 |
| ENSG00000146425.11 | DYNLT1 | protein_coding | 17744,44587 | 0,627183366 | 0,061812579 | 10,1465329 | 3,43E-24 | 8,60E-23 |
| ENSG00000146433.8 | TMEM181 | protein_coding | 12648,77852 | -0,232211134 | 0,050043844 | -4,640153843 | 3,48E-06 | 2,09E-05 |
| ENSG00000146463.11 | ZMYM4 | protein_coding | 16177,74851 | 0,168097645 | 0,045105686 | 3,726750658 | 0,000193964 | 0,00087249 |
| ENSG00000146530.14 | VWDE | protein_coding | 550,7490793 | 0,724877497 | 0,117305767 | 6,179384981 | 6,44E-10 | 6,16E-09 |
| ENSG00000146587.18 | RBAK | protein_coding | 3332,674441 | 0,228793066 | 0,054032741 | 4,234341294 | 2,29E-05 | 0,00012143 |
| ENSG00000146592.17 | CREB5 | protein_coding | 27,07312368 | 2,457213642 | 0,746642506 | 3,291017621 | 0,000998257 | 0,00386876 |
| ENSG00000146676.9 | PURB | protein_coding | 11392,38545 | 0,327610447 | 0,054114744 | 6,053996052 | 1,41E-09 | 1,30E-08 |
| ENSG00000146701.12 | MDH2 | protein_coding | 8797,409243 | -0,202197897 | 0,057706574 | -3,503897083 | 0,000458503 | 0,001920081 |
| ENSG00000146729.10 | NIPSNAP2 | protein_coding | 5789,998583 | -0,227866032 | 0,048520519 | -4,696281825 | 2,65E-06 | 1,62E-05 |
| ENSG00000146731.11 | CCT6A | protein_coding | 38872,3957 | -0,14139188 | 0,038855022 | -3,638960226 | 0,000273741 | 0,001197132 |
| ENSG00000146733.14 | PSPH | protein_coding | 3268,93752 | -0,223352127 | 0,071423213 | -3,127164381 | 0,001765012 | 0,006470465 |
| ENSG00000146757.13 | ZNF92 | protein_coding | 4898,335113 | 0,235827078 | 0,048948533 | 4,817857956 | 1,45E-06 | 9,20E-06 |
| ENSG00000146802.12 | TMEM168 | protein_coding | 3154,572017 | -0,353492335 | 0,054976941 | -6,429829053 | 1,28E-10 | 1,32E-09 |
| ENSG00000146830.10 | GIGYF1 | protein_coding | 7127,079163 | -0,139302105 | 0,05320904 | -2,618015774 | 0,008844272 | 0,026953115 |
| ENSG00000146833.15 | TRIM4 | protein_coding | 6093,269301 | 0,212256951 | 0,050089866 | 4,237522868 | 2,26E-05 | 0,000119807 |
| ENSG00000146842.17 | TMEM209 | protein_coding | 4983,103973 | -0,291232675 | 0,055723899 | -5,22635134 | 1,73E-07 | 1,24E-06 |
| ENSG00000146858.8 | ZC3HAV1L | protein_coding | 1415,846981 | -0,26802155 | 0,073804068 | -3,631528149 | 0,000281748 | 0,001229282 |
| ENSG00000146859.6 | TMEM140 | protein_coding | 5175,490306 | 6,069416244 | 0,204044099 | 29,74561023 | 1,98E-194 | 3,97E-192 |
| ENSG00000146872.17 | TLK2 | protein_coding | 9256,562112 | 0,625398278 | 0,047668616 | 13,11970714 | 2,54E-39 | 1,12E-37 |
| ENSG00000146918.19 | NCAPG2 | protein_coding | 14436,42069 | -0,463622945 | 0,042183948 | -10,99050639 | 4,25E-28 | 1,24E-26 |
| ENSG00000146950.13 | SHROOM2 | protein_coding | 3877,31076 | 0,309229605 | 0,05593576 | 5,528298969 | 3,23E-08 | 2,54E-07 |
| ENSG00000146955.10 | RAB19 | protein_coding | 491,9694255 | 1,551280768 | 0,114286249 | 13,57364322 | 5,74E-42 | 2,74E-40 |
| ENSG00000147003.7 | CLTRN | protein_coding | 1372,671289 | 1,702727006 | 0,115835409 | 14,69953809 | 6,49E-49 | 3,62E-47 |
| ENSG00000147044.21 | CASK | protein_coding | 30243,41006 | -0,130407501 | 0,041348073 | -3,153895473 | 0,001611068 | 0,005962434 |

| | | | | | | | | |
|---|---|---|---|---|---|---|---|---|
| ENSG00000147050.14 | KDM6A | protein_coding | 10233,43851 | 0,34991651 | 0,053348011 | 6,559129423 | 5,41E-11 | 5,79E-10 |
| ENSG00000147099.21 | HDAC8 | protein_coding | 3045,143422 | -0,233363782 | 0,071841185 | -3,248328679 | 0,001160851 | 0,004430998 |
| ENSG00000147118.11 | ZNF182 | protein_coding | 4523,144986 | 0,19638816 | 0,050174501 | 3,914102925 | 9,07E-05 | 0,00043522 |
| ENSG00000147121.16 | KRBOX4 | protein_coding | 3528,87786 | 0,136217349 | 0,053496089 | 2,546304793 | 0,010887008 | 0,03216798 |
| ENSG00000147124.12 | ZNF41 | protein_coding | 4302,376759 | 0,220159511 | 0,05449771 | 4,03979376 | 5,35E-05 | 0,000266946 |
| ENSG00000147140.16 | NONO | protein_coding | 67422,19214 | -0,165750973 | 0,041385195 | -4,005078896 | 6,20E-05 | 0,000306363 |
| ENSG00000147144.13 | CCDC120 | protein_coding | 2039,803829 | 0,189888407 | 0,061470258 | 3,08911027 | 0,002007569 | 0,007250386 |
| ENSG00000147155.11 | EBP | protein_coding | 13082,60312 | -0,194308771 | 0,05228175 | -3,716569773 | 0,000201946 | 0,000904417 |
| ENSG00000147180.16 | ZNF711 | protein_coding | 5976,127951 | 0,142546024 | 0,05102775 | 2,793500092 | 0,005214101 | 0,016935672 |
| ENSG00000147257.14 | GPC3 | protein_coding | 2365,938565 | 0,183088758 | 0,059010888 | 3,102626701 | 0,001918114 | 0,00696256 |
| ENSG00000147262.4 | GPR119 | protein_coding | 1912,562972 | -0,270794239 | 0,080238454 | -3,3748686 | 0,000738509 | 0,002949975 |
| ENSG00000147274.14 | RBMX | protein_coding | 35992,1561 | -0,361204227 | 0,041825404 | -8,636000988 | 5,82E-18 | 1,06E-16 |
| ENSG00000147364.16 | FBXO25 | protein_coding | 5911,908248 | -0,122840175 | 0,050112716 | -2,451277529 | 0,014235015 | 0,040620684 |
| ENSG00000147394.18 | ZNF185 | protein_coding | 394,3819535 | -0,613444158 | 0,153601073 | -3,993749176 | 6,50E-05 | 0,00031996 |
| ENSG00000147400.8 | CETN2 | protein_coding | 4716,613506 | -0,285985463 | 0,057615889 | -4,963656159 | 6,92E-07 | 4,60E-06 |
| ENSG00000147403.16 | RPL10 | protein_coding | 54055,09645 | -0,348759933 | 0,044939192 | -7,760707738 | 8,45E-15 | 1,25E-13 |
| ENSG00000147419.18 | CCDC25 | protein_coding | 11324,16227 | -0,206396641 | 0,051443765 | -4,012082711 | 6,02E-05 | 0,000297805 |
| ENSG00000147439.12 | BIN3 | protein_coding | 1802,607744 | 0,196867522 | 0,068717033 | 2,864901368 | 0,004171391 | 0,013912261 |
| ENSG00000147454.14 | SLC25A37 | protein_coding | 3207,543161 | -0,40236192 | 0,069490822 | -5,790144793 | 7,03E-09 | 5,96E-08 |
| ENSG00000147475.16 | ERLIN2 | protein_coding | 5383,735538 | -0,160900617 | 0,065003278 | -2,475269268 | 0,013313576 | 0,038306951 |
| ENSG00000147488.11 | ST18 | protein_coding | 13901,13584 | 0,144065102 | 0,046607264 | 3,091043983 | 0,001994541 | 0,007209751 |
| ENSG00000147536.12 | GINS4 | protein_coding | 3389,834791 | -0,355563687 | 0,072158674 | -4,927525205 | 8,33E-07 | 5,46E-06 |
| ENSG00000147548.17 | NSD3 | protein_coding | 18822,40979 | 0,131674987 | 0,04573152 | 2,879304812 | 0,003985529 | 0,013357695 |
| ENSG00000147571.5 | CRH | protein_coding | 36,6913967 | 1,174249841 | 0,394069353 | 2,97980503 | 0,002884319 | 0,010006649 |
| ENSG00000147586.10 | MRPS28 | protein_coding | 2450,866467 | -0,241390788 | 0,07013486 | -3,441808953 | 0,000577838 | 0,002364829 |
| ENSG00000147592.9 | LACTB2 | protein_coding | 1407,451085 | 0,194794603 | 0,074301763 | 2,621668641 | 0,008750046 | 0,026698493 |
| ENSG00000147604.14 | RPL7 | protein_coding | 61081,66015 | -0,910149398 | 0,041781211 | -21,78370056 | 3,31E-105 | 3,97E-103 |
| ENSG00000147654.15 | EBAG9 | protein_coding | 2771,487787 | 0,146476371 | 0,058840054 | 2,489399001 | 0,012795927 | 0,036982989 |
| ENSG00000147677.11 | EIF3H | protein_coding | 14822,69313 | -0,311577521 | 0,046249312 | -6,736911539 | 1,62E-11 | 1,81E-10 |
| ENSG00000147684.9 | NDUFB9 | protein_coding | 8442,702347 | -0,180673606 | 0,051955876 | -3,477443131 | 0,00050622 | 0,00209458 |
| ENSG00000147724.12 | FAM135B | protein_coding | 4190,981931 | -0,271685794 | 0,056352891 | -4,821150935 | 1,43E-06 | 9,07E-06 |
| ENSG00000147804.9 | SLC39A4 | protein_coding | 616,6778543 | -0,564338552 | 0,121528011 | -4,643691183 | 3,42E-06 | 2,05E-05 |
| ENSG00000147852.16 | VLDLR | protein_coding | 15910,69211 | 0,397306201 | 0,041576839 | 9,555950049 | 1,22E-21 | 2,71E-20 |
| ENSG00000147853.17 | AK3 | protein_coding | 9303,090444 | -0,186073962 | 0,044714533 | -4,161375516 | 3,16E-05 | 0,00016365 |
| ENSG00000147872.10 | PLIN2 | protein_coding | 14851,55578 | 0,414825627 | 0,043508709 | 9,534312445 | 1,51E-21 | 3,32E-20 |
| ENSG00000147883.12 | CDKN2B | protein_coding | 5035,324864 | 0,311994416 | 0,048796359 | 6,393805227 | 1,62E-10 | 1,65E-09 |

| | | | | | | | | |
|---|---|---|---|---|---|---|---|---|
| ENSG00000147889.17 | CDKN2A | protein_coding | 5107,712652 | 0,200632106 | 0,06646818 | 3,018468473 | 0,002540558 | 0,008934954 |
| ENSG00000147912.13 | FBXO10 | protein_coding | 1596,325179 | 0,381149708 | 0,093139215 | 4,092258107 | 4,27E-05 | 0,000216814 |
| ENSG00000147955.17 | SIGMAR1 | protein_coding | 7427,60345 | -0,144548627 | 0,056778628 | -2,545828099 | 0,010901886 | 0,032207711 |
| ENSG00000147996.16 | CBWD5 | protein_coding | 4377,078748 | -0,176506713 | 0,057857252 | -3,050727542 | 0,002282876 | 0,008122868 |
| ENSG00000148019.13 | CEP78 | protein_coding | 8208,370417 | -0,616046724 | 0,055385391 | -11,12291001 | 9,71E-29 | 2,91E-27 |
| ENSG00000148082.10 | SHC3 | protein_coding | 1197,633442 | -0,479223992 | 0,110727488 | -4,327958668 | 1,50E-05 | 8,19E-05 |
| ENSG00000148143.13 | ZNF462 | protein_coding | 5060,634083 | 0,384907817 | 0,153063989 | 2,51468565 | 0,011913856 | 0,034754922 |
| ENSG00000148158.17 | SNX30 | protein_coding | 5510,943103 | -0,439318584 | 0,058894893 | -7,459366373 | 8,69E-14 | 1,20E-12 |
| ENSG00000148175.12 | STOM | protein_coding | 406,9489331 | 0,385044777 | 0,125215972 | 3,075045234 | 0,002104706 | 0,007559412 |
| ENSG00000148225.16 | WDR31 | protein_coding | 1815,81128 | -0,583796614 | 0,070901821 | -8,23387332 | 1,81E-16 | 3,00E-15 |
| ENSG00000148290.10 | SURF1 | protein_coding | 3073,502334 | -0,213158917 | 0,06749259 | -3,158256581 | 0,001587158 | 0,005885558 |
| ENSG00000148297.15 | MED22 | protein_coding | 4181,765274 | 0,180287263 | 0,054737665 | 3,293660106 | 0,00098892 | 0,003835878 |
| ENSG00000148300.12 | REXO4 | protein_coding | 3666,166766 | 0,173815389 | 0,055345317 | 3,140561787 | 0,001686241 | 0,006208998 |
| ENSG00000148303.17 | RPL7A | protein_coding | 72333,05516 | -0,620026055 | 0,044500594 | -13,9329837 | 3,99E-44 | 2,01E-42 |
| ENSG00000148343.18 | MIGA2 | protein_coding | 2080,876637 | 0,231738257 | 0,074285755 | 3,119551734 | 0,001811265 | 0,006620616 |
| ENSG00000148356.13 | LRSAM1 | protein_coding | 1570,943144 | 0,591706589 | 0,078104922 | 7,575791317 | 3,57E-14 | 5,06E-13 |
| ENSG00000148362.11 | PAXX | protein_coding | 2885,742551 | -0,442283097 | 0,070134934 | -6,306174031 | 2,86E-10 | 2,85E-09 |
| ENSG00000148400.12 | NOTCH1 | protein_coding | 462,9590899 | 0,724368881 | 0,12546191 | 5,773615907 | 7,76E-09 | 6,54E-08 |
| ENSG00000148429.14 | USP6NL | protein_coding | 2770,024031 | 0,366805546 | 0,067405041 | 5,441811752 | 5,27E-08 | 4,04E-07 |
| ENSG00000148444.16 | COMMD3 | protein_coding | 2911,524528 | -0,402386342 | 0,055252742 | -7,282649304 | 3,27E-13 | 4,31E-12 |
| ENSG00000148459.16 | PDSS1 | protein_coding | 614,1684505 | -0,455708576 | 0,106064372 | -4,29652829 | 1,73E-05 | 9,35E-05 |
| ENSG00000148468.17 | FAM171A1 | protein_coding | 87,23404487 | -0,685956305 | 0,255022604 | -2,689786299 | 0,007149779 | 0,022367529 |
| ENSG00000148483.9 | TMEM236 | protein_coding | 286,0565003 | 1,26416153 | 0,161080986 | 7,847987271 | 4,23E-15 | 6,41E-14 |
| ENSG00000148498.16 | PARD3 | protein_coding | 5774,486505 | 0,314082513 | 0,048001036 | 6,543244511 | 6,02E-11 | 6,41E-10 |
| ENSG00000148516.21 | ZEB1 | protein_coding | 431,981492 | 0,439852314 | 0,129349177 | 3,400503379 | 0,000672619 | 0,0027118 |
| ENSG00000148541.12 | FAM13C | protein_coding | 1342,38351 | 0,299432734 | 0,079314455 | 3,77526058 | 0,00015984 | 0,000731122 |
| ENSG00000148572.16 | NRBF2 | protein_coding | 3573,755521 | 0,172021529 | 0,05660129 | 3,039180385 | 0,002372228 | 0,008399634 |
| ENSG00000148584.15 | A1CF | protein_coding | 34774,24572 | 0,156362928 | 0,046609415 | 3,354749875 | 0,000794368 | 0,003150735 |
| ENSG00000148634.15 | HERC4 | protein_coding | 6339,845355 | 0,143046549 | 0,047073284 | 3,038805383 | 0,002375182 | 0,008408773 |
| ENSG00000148688.13 | RPP30 | protein_coding | 4221,718982 | -0,139285664 | 0,052313068 | -2,662540548 | 0,007755323 | 0,024014934 |
| ENSG00000148690.12 | FRA10AC1 | protein_coding | 1645,924248 | -0,437923789 | 0,069805688 | -6,273468563 | 3,53E-10 | 3,48E-09 |
| ENSG00000148702.15 | HABP2 | protein_coding | 422,9937797 | 1,736862106 | 0,146737047 | 11,83656163 | 2,53E-32 | 8,98E-31 |
| ENSG00000148719.15 | DNAJB12 | protein_coding | 3952,443984 | 0,219949309 | 0,052010529 | 4,228938105 | 2,35E-05 | 0,000124266 |
| ENSG00000148730.7 | EIF4EBP2 | protein_coding | 28749,84323 | -0,774799845 | 0,052968785 | -14,62748001 | 1,88E-48 | 1,03E-46 |
| ENSG00000148773.14 | MKI67 | protein_coding | 15569,06461 | -0,284297977 | 0,05108638 | -5,565044525 | 2,62E-08 | 2,09E-07 |
| ENSG00000148814.18 | LRRC27 | protein_coding | 355,2551578 | 0,908068313 | 0,136235708 | 6,665420744 | 2,64E-11 | 2,90E-10 |

| | | | | | | | | |
|---|---|---|---|---|---|---|---|---|
| ENSG00000148834.13 | GSTO1 | protein_coding | 6226,143757 | 0,321103052 | 0,049995972 | 6,422578482 | 1,34E-10 | 1,38E-09 |
| ENSG00000148835.11 | TAF5 | protein_coding | 2522,846592 | -0,331427709 | 0,063737345 | -5,19989823 | 1,99E-07 | 1,42E-06 |
| ENSG00000148908.15 | RGS10 | protein_coding | 78,65694724 | -0,873413116 | 0,274229537 | -3,18497098 | 0,001447685 | 0,005414747 |
| ENSG00000148926.10 | ADM | protein_coding | 603,1411258 | 1,054380669 | 0,14488872 | 7,27717568 | 3,41E-13 | 4,48E-12 |
| ENSG00000148935.11 | GAS2 | protein_coding | 2475,829037 | -0,385922227 | 0,060793889 | -6,348043137 | 2,18E-10 | 2,20E-09 |
| ENSG00000148943.12 | LIN7C | protein_coding | 6984,261202 | -0,157355233 | 0,05055355 | -3,112644559 | 0,001854192 | 0,006757773 |
| ENSG00000148950.11 | IMMP1L | protein_coding | 3923,307045 | -0,980479203 | 0,076179331 | -12,87067228 | 6,58E-38 | 2,76E-36 |
| ENSG00000149089.13 | APIP | protein_coding | 3178,80572 | -0,198855041 | 0,060940743 | -3,263088574 | 0,001102051 | 0,004230228 |
| ENSG00000149091.15 | DGKZ | protein_coding | 4220,632237 | 0,19261398 | 0,057026772 | 3,377606212 | 0,000731197 | 0,002924399 |
| ENSG00000149115.14 | TNKS1BP1 | protein_coding | 1329,614502 | 0,243742292 | 0,095326707 | 2,556915056 | 0,0105605 | 0,031355533 |
| ENSG00000149136.9 | SSRP1 | protein_coding | 18999,40855 | -0,274037481 | 0,041393031 | -6,620377331 | 3,58E-11 | 3,89E-10 |
| ENSG00000149179.13 | C11orf49 | protein_coding | 6108,381539 | 0,207902182 | 0,049489866 | 4,200904126 | 2,66E-05 | 0,000139002 |
| ENSG00000149182.15 | ARFGAP2 | protein_coding | 8119,065703 | 0,168880045 | 0,065226031 | 2,589151037 | 0,009621288 | 0,028987496 |
| ENSG00000149196.16 | HIKESHI | protein_coding | 2368,217811 | -0,180134549 | 0,065035935 | -2,769769475 | 0,005609598 | 0,018067631 |
| ENSG00000149212.11 | SESN3 | protein_coding | 20737,05969 | -0,1687452 | 0,044751738 | -3,770696035 | 0,000162793 | 0,000742965 |
| ENSG00000149218.5 | ENDOD1 | protein_coding | 1837,031401 | 0,822460944 | 0,070669862 | 11,63807204 | 2,64E-31 | 8,91E-30 |
| ENSG00000149256.15 | TENM4 | protein_coding | 4434,276107 | 0,231510466 | 0,067637477 | 3,422813465 | 0,000619766 | 0,002516744 |
| ENSG00000149257.14 | SERPINH1 | protein_coding | 1203,386533 | 0,655743329 | 0,095490136 | 6,867131613 | 6,55E-12 | 7,67E-11 |
| ENSG00000149269.9 | PAK1 | protein_coding | 5429,692222 | 0,142051991 | 0,047979702 | 2,960668475 | 0,003069722 | 0,010576514 |
| ENSG00000149273.15 | RPS3 | protein_coding | 87891,08777 | -0,611297664 | 0,04693289 | -13,02493124 | 8,83E-39 | 3,81E-37 |
| ENSG00000149289.11 | ZC3H12C | protein_coding | 5400,822556 | 0,536432468 | 0,057689891 | 9,298552353 | 1,42E-20 | 3,00E-19 |
| ENSG00000149294.16 | NCAM1 | protein_coding | 16880,25289 | -0,318757965 | 0,051707593 | -6,164625881 | 7,06E-10 | 6,71E-09 |
| ENSG00000149300.10 | C11orf52 | protein_coding | 994,8952097 | 0,924965052 | 0,093293323 | 9,914590088 | 3,60E-23 | 8,54E-22 |
| ENSG00000149328.14 | GLB1L2 | protein_coding | 4351,148465 | -0,431054781 | 0,063013553 | -6,840667745 | 7,88E-12 | 9,13E-11 |
| ENSG00000149357.10 | LAMTOR1 | protein_coding | 6266,776491 | -0,132321769 | 0,048319394 | -2,73848154 | 0,006172363 | 0,019662267 |
| ENSG00000149380.12 | P4HA3 | protein_coding | 164,6883011 | -0,793037347 | 0,193280701 | -4,103034316 | 4,08E-05 | 0,000207375 |
| ENSG00000149418.11 | ST14 | protein_coding | 2716,541653 | 0,912406764 | 0,060755954 | 15,01756943 | 5,63E-51 | 3,31E-49 |
| ENSG00000149428.19 | HYOU1 | protein_coding | 37701,01015 | 0,609284973 | 0,049793562 | 12,23621993 | 1,99E-34 | 7,46E-33 |
| ENSG00000149483.12 | TMEM138 | protein_coding | 2479,17547 | -0,325469802 | 0,061189526 | -5,319044378 | 1,04E-07 | 7,67E-07 |
| ENSG00000149485.18 | FADS1 | protein_coding | 60660,36195 | -0,111463688 | 0,04559124 | -2,444848784 | 0,014491289 | 0,041184901 |
| ENSG00000149503.13 | INCENP | protein_coding | 4666,386323 | -0,885101237 | 0,194916631 | -4,540922106 | 5,60E-06 | 3,26E-05 |
| ENSG00000149527.18 | PLCH2 | protein_coding | 602,8247207 | -1,503689685 | 0,123647512 | -12,16109942 | 5,01E-34 | 1,85E-32 |
| ENSG00000149531.15 | FRG1BP | unprocessed_pseud | 1405,056327 | 0,198203772 | 0,072307969 | 2,741105505 | 0,006123284 | 0,019516321 |
| ENSG00000149547.15 | EI24 | protein_coding | 15261,18119 | -0,370064167 | 0,050035786 | -7,395989858 | 1,40E-13 | 1,91E-12 |
| ENSG00000149548.15 | CCDC15 | protein_coding | 1067,107688 | -0,332375098 | 0,083825526 | -3,965082179 | 7,34E-05 | 0,00035801 |
| ENSG00000149554.13 | CHEK1 | protein_coding | 6550,45302 | -0,153238018 | 0,052501751 | -2,91872203 | 0,003514695 | 0,011946884 |

| | | | | | | | | |
|---|---|---|---|---|---|---|---|---|
| ENSG00000149557.13 | FEZ1 | protein_coding | 86,26753871 | -0,825738784 | 0,289632712 | -2,850985922 | 0,00435839 | 0,014439652 |
| ENSG00000149564.12 | ESAM | protein_coding | 263,6991062 | -0,35898964 | 0,143556171 | -2,500691113 | 0,012395124 | 0,036006488 |
| ENSG00000149571.12 | KIRREL3 | protein_coding | 674,0683636 | 0,670263266 | 0,110496028 | 6,065948978 | 1,31E-09 | 1,21E-08 |
| ENSG00000149573.9 | MPZL2 | protein_coding | 46,44686803 | 0,908507987 | 0,355393067 | 2,556346957 | 0,010577759 | 0,031394354 |
| ENSG00000149636.15 | DSN1 | protein_coding | 12642,5481 | -0,232489114 | 0,044924649 | -5,175090238 | 2,28E-07 | 1,61E-06 |
| ENSG00000149639.15 | SOGA1 | protein_coding | 9746,44683 | -0,40491133 | 0,054732309 | -7,398031136 | 1,38E-13 | 1,88E-12 |
| ENSG00000149646.12 | CNBD2 | protein_coding | 1239,167358 | -0,2246513 | 0,089793856 | -2,501856044 | 0,012354415 | 0,035909804 |
| ENSG00000149679.11 | CABLES2 | protein_coding | 2408,214211 | -0,481713699 | 0,072971409 | -6,6014033 | 4,07E-11 | 4,41E-10 |
| ENSG00000149792.8 | MRPL49 | protein_coding | 11197,18903 | 0,29938402 | 0,054214244 | 5,522239152 | 3,35E-08 | 2,62E-07 |
| ENSG00000149806.11 | FAU | protein_coding | 12358,3712 | -0,210630235 | 0,060530019 | -3,479764879 | 0,000501854 | 0,002079189 |
| ENSG00000149823.9 | VPS51 | protein_coding | 2837,011143 | -0,321979717 | 0,082137524 | -3,92000758 | 8,85E-05 | 0,000425509 |
| ENSG00000149925.20 | ALDOA | protein_coding | 23976,52411 | 0,208006244 | 0,054133767 | 3,842449066 | 0,000121813 | 0,000571464 |
| ENSG00000149927.18 | DOC2A | protein_coding | 482,7618798 | 0,592346456 | 0,128591093 | 4,606434572 | 4,10E-06 | 2,43E-05 |
| ENSG00000149929.15 | HIRIP3 | protein_coding | 4915,64598 | -0,190341974 | 0,062080721 | -3,066039983 | 0,002169143 | 0,007759892 |
| ENSG00000149932.17 | TMEM219 | protein_coding | 3009,30224 | 0,434620647 | 0,064024599 | 6,788338484 | 1,13E-11 | 1,29E-10 |
| ENSG00000149970.16 | CNKSR2 | protein_coding | 572,4229926 | -0,31441359 | 0,117522972 | -2,675337287 | 0,00746541 | 0,02320659 |
| ENSG00000150093.19 | ITGB1 | protein_coding | 131871,1303 | -0,112379266 | 0,039372275 | -2,85427416 | 0,004313529 | 0,01432476 |
| ENSG00000150347.16 | ARID5B | protein_coding | 4816,909135 | 0,473294631 | 0,056070414 | 8,441076024 | 3,14E-17 | 5,44E-16 |
| ENSG00000150361.12 | KLHL1 | protein_coding | 290,4948645 | -0,842598201 | 0,160953336 | -5,235046501 | 1,65E-07 | 1,19E-06 |
| ENSG00000150403.18 | TMCO3 | protein_coding | 11373,66225 | -0,159643426 | 0,044461919 | -3,59056537 | 0,000329961 | 0,001421727 |
| ENSG00000150433.9 | TMEM218 | protein_coding | 1323,279485 | -0,481689786 | 0,074760194 | -6,443131845 | 1,17E-10 | 1,21E-09 |
| ENSG00000150456.10 | EEF1AKMT1 | protein_coding | 690,3990124 | -0,362748093 | 0,095989074 | -3,779056055 | 0,000157424 | 0,000720947 |
| ENSG00000150457.9 | LATS2 | protein_coding | 71,07405001 | 1,089305917 | 0,317108074 | 3,435125136 | 0,00059228 | 0,002415674 |
| ENSG00000150527.17 | MIA2 | protein_coding | 21076,05353 | 0,423857804 | 0,045342577 | 9,347898357 | 8,94E-21 | 1,90E-19 |
| ENSG00000150593.18 | PDCD4 | protein_coding | 11585,12055 | -0,450016269 | 0,043337209 | -10,38406209 | 2,93E-25 | 7,69E-24 |
| ENSG00000150594.6 | ADRA2A | protein_coding | 973,2379485 | -0,241573187 | 0,089420306 | -2,701547316 | 0,006901766 | 0,021701538 |
| ENSG00000150625.16 | GPM6A | protein_coding | 290,6724507 | 1,349081465 | 0,154439933 | 8,735315005 | 2,43E-18 | 4,52E-17 |
| ENSG00000150627.15 | WDR17 | protein_coding | 3748,396859 | -0,459176978 | 0,068984635 | -6,656221072 | 2,81E-11 | 3,08E-10 |
| ENSG00000150672.17 | DLG2 | protein_coding | 6524,609339 | 0,326998433 | 0,067565531 | 4,839722707 | 1,30E-06 | 8,31E-06 |
| ENSG00000150687.12 | PRSS23 | protein_coding | 3471,959966 | 0,509393452 | 0,07177262 | 7,097322835 | 1,27E-12 | 1,60E-11 |
| ENSG00000150753.12 | CCT5 | protein_coding | 34121,19295 | -0,171677959 | 0,044056565 | -3,896762243 | 9,75E-05 | 0,000464706 |
| ENSG00000150756.14 | ATPSCKMT | protein_coding | 2705,852034 | -0,806814989 | 0,067409131 | -11,96892732 | 5,17E-33 | 1,87E-31 |
| ENSG00000150779.11 | TIMM8B | protein_coding | 5650,399447 | -0,220109821 | 0,063367016 | -3,473570843 | 0,000513582 | 0,002120362 |
| ENSG00000150873.11 | C2orf50 | protein_coding | 25,10130721 | 2,133829721 | 0,563354936 | 3,787718159 | 0,000152037 | 0,000699963 |
| ENSG00000150893.11 | FREM2 | protein_coding | 2429,676195 | -0,434136477 | 0,069990875 | -6,202758267 | 5,55E-10 | 5,36E-09 |
| ENSG00000150938.10 | CRIM1 | protein_coding | 2657,449423 | 0,345488102 | 0,063475705 | 5,442839956 | 5,24E-08 | 4,01E-07 |

| | | | | | | | | |
|---|---|---|---|---|---|---|---|---|
| ENSG00000150961.15 | SEC24D | protein_coding | 28633,72597 | 0,19990078 | 0,043561939 | 4,588886197 | 4,46E-06 | 2,63E-05 |
| ENSG00000150990.8 | DHX37 | protein_coding | 1748,713456 | 0,458323939 | 0,085593231 | 5,35467504 | 8,57E-08 | 6,36E-07 |
| ENSG00000150991.15 | UBC | protein_coding | 187681,7178 | 0,453956152 | 0,044325381 | 10,24144944 | 1,29E-24 | 3,30E-23 |
| ENSG00000150995.19 | ITPR1 | protein_coding | 4205,705146 | 0,340114909 | 0,059282728 | 5,737166992 | 9,63E-09 | 8,02E-08 |
| ENSG00000151006.7 | PRSS53 | protein_coding | 226,3996154 | -0,585935766 | 0,202791626 | -2,889348915 | 0,003860405 | 0,012988617 |
| ENSG00000151012.13 | SLC7A11 | protein_coding | 749,1168882 | 1,585422655 | 0,10663546 | 14,86768711 | 5,34E-50 | 3,08E-48 |
| ENSG00000151014.6 | NOCT | protein_coding | 1750,532589 | 0,251363543 | 0,069574991 | 3,612843324 | 0,000302858 | 0,001313745 |
| ENSG00000151023.17 | ENKUR | protein_coding | 54,54339261 | 0,95038164 | 0,330855219 | 2,872500073 | 0,00407238 | 0,013618362 |
| ENSG00000151025.11 | GPR158 | protein_coding | 1062,167295 | 0,851744914 | 0,090574114 | 9,40384484 | 5,26E-21 | 1,13E-19 |
| ENSG00000151065.14 | DCP1B | protein_coding | 1414,863954 | 0,214454219 | 0,082015536 | 2,614799935 | 0,008927973 | 0,027171411 |
| ENSG00000151092.17 | NGLY1 | protein_coding | 3240,264291 | 0,202835624 | 0,060056086 | 3,377436638 | 0,000731648 | 0,002925682 |
| ENSG00000151150.22 | ANK3 | protein_coding | 2139,589278 | -0,190064531 | 0,071753114 | -2,648868048 | 0,008076186 | 0,024888673 |
| ENSG00000151176.8 | PLBD2 | protein_coding | 6874,922879 | 0,172054153 | 0,049460795 | 3,478596569 | 0,000504047 | 0,002087505 |
| ENSG00000151229.13 | SLC2A13 | protein_coding | 8055,670534 | -0,163287578 | 0,05259972 | -3,104343107 | 0,00190702 | 0,006925641 |
| ENSG00000151233.11 | GXYLT1 | protein_coding | 8724,863828 | -0,332192321 | 0,052967074 | -6,271675897 | 3,57E-10 | 3,52E-09 |
| ENSG00000151287.17 | TEX30 | protein_coding | 3359,07964 | -0,260632057 | 0,057664323 | -4,51981474 | 6,19E-06 | 3,57E-05 |
| ENSG00000151304.5 | SRFBP1 | protein_coding | 2716,796309 | 0,259801865 | 0,068689061 | 3,78228879 | 0,000155393 | 0,000712661 |
| ENSG00000151320.11 | AKAP6 | protein_coding | 734,7364415 | -0,364853434 | 0,142138684 | -2,566883433 | 0,01026171 | 0,030597541 |
| ENSG00000151322.18 | NPAS3 | protein_coding | 2506,733127 | -0,306234434 | 0,072133844 | -4,245364148 | 2,18E-05 | 0,000115912 |
| ENSG00000151338.18 | MIPOL1 | protein_coding | 822,6545049 | 0,584475193 | 0,092299203 | 6,332396958 | 2,41E-10 | 2,42E-09 |
| ENSG00000151364.17 | KCTD14 | protein_coding | 148,6854545 | 0,567369979 | 0,208998774 | 2,714704827 | 0,006633487 | 0,020941536 |
| ENSG00000151366.13 | NDUFC2 | protein_coding | 11628,77972 | -0,148584232 | 0,053180756 | -2,793947329 | 0,005206896 | 0,016919593 |
| ENSG00000151376.16 | ME3 | protein_coding | 639,5874474 | 0,506336393 | 0,098969922 | 5,116063379 | 3,12E-07 | 2,17E-06 |
| ENSG00000151413.17 | NUBPL | protein_coding | 1684,264682 | -0,423821225 | 0,090424554 | -4,687014801 | 2,77E-06 | 1,69E-05 |
| ENSG00000151414.15 | NEK7 | protein_coding | 7562,161529 | 0,299650336 | 0,044556327 | 6,725202741 | 1,75E-11 | 1,96E-10 |
| ENSG00000151422.13 | FER | protein_coding | 5198,966432 | -0,277533834 | 0,072577076 | -3,82398754 | 0,000131311 | 0,00061194 |
| ENSG00000151461.20 | UPF2 | protein_coding | 4238,986778 | 0,284399641 | 0,0571205 | 4,978941753 | 6,39E-07 | 4,27E-06 |
| ENSG00000151466.12 | SCLT1 | protein_coding | 3938,434243 | 0,765383204 | 0,052638185 | 14,54045595 | 6,71E-48 | 3,61E-46 |
| ENSG00000151468.11 | CCDC3 | protein_coding | 61,503778 | -1,779493379 | 0,362408022 | -4,910193124 | 9,10E-07 | 5,94E-06 |
| ENSG00000151470.13 | C4orf33 | protein_coding | 5313,364692 | 1,759490965 | 0,060773186 | 28,95176434 | 2,67E-184 | 5,00E-182 |
| ENSG00000151491.14 | EPS8 | protein_coding | 6216,581559 | 0,210678876 | 0,054987623 | 3,831387236 | 0,000127423 | 0,000595424 |
| ENSG00000151502.11 | VPS26B | protein_coding | 9200,75739 | -0,212406341 | 0,046580598 | -4,559974564 | 5,12E-06 | 2,99E-05 |
| ENSG00000151503.12 | NCAPD3 | protein_coding | 14310,39529 | -0,385390578 | 0,051902567 | -7,425270062 | 1,13E-13 | 1,54E-12 |
| ENSG00000151553.15 | FAM160B1 | protein_coding | 1761,374563 | 0,199577628 | 0,064342509 | 3,101800536 | 0,001923475 | 0,006978643 |
| ENSG00000151572.18 | ANO4 | protein_coding | 509,3183393 | 0,29594063 | 0,124202229 | 2,38273204 | 0,017184698 | 0,047714722 |
| ENSG00000151611.16 | MMAA | protein_coding | 2862,330896 | 0,493590997 | 0,06006256 | 8,217948053 | 2,07E-16 | 3,41E-15 |

| | | | | | | | | |
|---|---|---|---|---|---|---|---|---|
| ENSG00000151612.17 | ZNF827 | protein_coding | 4728,487135 | 0,33841865 | 0,059093255 | 5,726857452 | 1,02E-08 | 8,49E-08 |
| ENSG00000151615.3 | POU4F2 | protein_coding | 202,2768119 | 0,714604124 | 0,162403033 | 4,400189518 | 1,08E-05 | 6,03E-05 |
| ENSG00000151632.17 | AKR1C2 | protein_coding | 3002,310289 | -0,202532053 | 0,064044615 | -3,162358811 | 0,001564966 | 0,005814223 |
| ENSG00000151655.19 | ITIH2 | protein_coding | 112,1981488 | -0,906670361 | 0,230035976 | -3,941428546 | 8,10E-05 | 0,000391997 |
| ENSG00000151657.12 | KIN | protein_coding | 1378,278849 | 0,4229639 | 0,082341708 | 5,136690887 | 2,80E-07 | 1,96E-06 |
| ENSG00000151665.12 | PIGF | protein_coding | 2755,630599 | -0,228532956 | 0,057812154 | -3,953026142 | 7,72E-05 | 0,000374676 |
| ENSG00000151689.13 | INPP1 | protein_coding | 4993,046243 | 1,225918053 | 0,066828392 | 18,34426994 | 3,67E-75 | 3,24E-73 |
| ENSG00000151690.15 | MFSD6 | protein_coding | 7358,822143 | -0,252798799 | 0,064197725 | -3,937815546 | 8,22E-05 | 0,000397688 |
| ENSG00000151694.14 | ADAM17 | protein_coding | 8558,256157 | 0,352459778 | 0,04792559 | 7,35431281 | 1,92E-13 | 2,57E-12 |
| ENSG00000151718.16 | WWC2 | protein_coding | 633,1039371 | 0,452575094 | 0,108430195 | 4,173884331 | 2,99E-05 | 0,000155378 |
| ENSG00000151726.14 | ACSL1 | protein_coding | 9340,86621 | 0,204688014 | 0,056840801 | 3,601075441 | 0,000316904 | 0,001370975 |
| ENSG00000151778.11 | SERP2 | protein_coding | 645,4679857 | -0,314225863 | 0,103269218 | -3,042783418 | 0,00234401 | 0,008314106 |
| ENSG00000151789.12 | ZNF385D | protein_coding | 496,4347163 | 0,916018365 | 0,321086634 | 2,852869816 | 0,004332637 | 0,014373371 |
| ENSG00000151835.16 | SACS | protein_coding | 5882,33973 | -0,689915415 | 0,070420323 | -9,79710664 | 1,16E-22 | 2,70E-21 |
| ENSG00000151849.15 | CENPJ | protein_coding | 3300,550335 | -0,279051278 | 0,078752342 | -3,543402926 | 0,000394999 | 0,001672188 |
| ENSG00000151883.18 | PARP8 | protein_coding | 3583,757952 | 1,261944295 | 0,065015131 | 19,41000936 | 6,35E-84 | 6,19E-82 |
| ENSG00000151893.15 | CACUL1 | protein_coding | 8915,499471 | 0,146068345 | 0,052926954 | 2,759810135 | 0,005783496 | 0,01855219 |
| ENSG00000151952.16 | TMEM132D | protein_coding | 3471,017701 | 0,231012141 | 0,065026084 | 3,552607311 | 0,000381433 | 0,001623003 |
| ENSG00000151967.18 | SCHIP1 | protein_coding | 486,5396017 | 0,340167453 | 0,111882993 | 3,0403857 | 0,002362754 | 0,008370038 |
| ENSG00000152061.23 | RABGAP1L | protein_coding | 11673,78687 | 1,604435827 | 0,061247112 | 26,19610584 | 2,94E-151 | 4,91E-149 |
| ENSG00000152076.18 | CCDC74B | protein_coding | 796,055421 | -0,579548376 | 0,137082374 | -4,227738118 | 2,36E-05 | 0,000124843 |
| ENSG00000152082.14 | MZT2B | protein_coding | 3200,244809 | -0,478092382 | 0,065952637 | -7,249026022 | 4,20E-13 | 5,46E-12 |
| ENSG00000152092.16 | ASTN1 | protein_coding | 521,4547062 | 0,471578133 | 0,120524915 | 3,912702462 | 9,13E-05 | 0,000437659 |
| ENSG00000152102.18 | FAM168B | protein_coding | 16897,72862 | -1,122867229 | 0,06102738 | -18,39940074 | 1,33E-75 | 1,18E-73 |
| ENSG00000152117.17 | AC073869.1 | transcribed_process | 3114,705852 | 0,197419387 | 0,056409378 | 3,49976179 | 0,000465674 | 0,001945769 |
| ENSG00000152127.9 | MGAT5 | protein_coding | 7409,254352 | -0,456520077 | 0,078030193 | -5,850556823 | 4,90E-09 | 4,24E-08 |
| ENSG00000152133.14 | GPATCH11 | protein_coding | 5275,529826 | -0,522247965 | 0,04954346 | -10,54120905 | 5,58E-26 | 1,51E-24 |
| ENSG00000152193.8 | RNF219 | protein_coding | 5459,790879 | 0,216736445 | 0,049862984 | 4,346640046 | 1,38E-05 | 7,57E-05 |
| ENSG00000152207.7 | CYSLTR2 | protein_coding | 383,3512802 | 2,101732266 | 0,179498551 | 11,70890936 | 1,15E-31 | 3,97E-30 |
| ENSG00000152208.13 | GRID2 | protein_coding | 1464,842164 | -0,503594247 | 0,081530398 | -6,176766709 | 6,54E-10 | 6,25E-09 |
| ENSG00000152213.3 | ARL11 | protein_coding | 120,1155653 | 1,32223094 | 0,222257093 | 5,94910571 | 2,70E-09 | 2,39E-08 |
| ENSG00000152223.15 | EPG5 | protein_coding | 7772,727685 | 0,135328881 | 0,051745758 | 2,615265215 | 0,008915819 | 0,027138091 |
| ENSG00000152234.16 | ATP5F1A | protein_coding | 57063,78272 | -0,151734657 | 0,041154315 | -3,686968314 | 0,000226942 | 0,001008734 |
| ENSG00000152240.13 | HAUS1 | protein_coding | 4447,117731 | -0,296789181 | 0,054895887 | -5,406401031 | 6,43E-08 | 4,87E-07 |
| ENSG00000152253.9 | SPC25 | protein_coding | 4091,687831 | -0,278913159 | 0,052958759 | -5,266610497 | 1,39E-07 | 1,01E-06 |
| ENSG00000152291.14 | TGOLN2 | protein_coding | 41019,16429 | -0,168915082 | 0,04422604 | -3,819358065 | 0,000133799 | 0,000621865 |

| | | | | | | | | |
|---|---|---|---|---|---|---|---|---|
| ENSG00000152315.5 | KCNK13 | protein_coding | 134,4398329 | -0,591454409 | 0,200231811 | -2,953848378 | 0,003138382 | 0,010780075 |
| ENSG00000152332.16 | UHMK1 | protein_coding | 16726,99155 | 0,123067143 | 0,050815212 | 2,421856319 | 0,015441454 | 0,043495395 |
| ENSG00000152348.16 | ATG10 | protein_coding | 1907,348956 | -0,589359433 | 0,084397969 | -6,983099694 | 2,89E-12 | 3,50E-11 |
| ENSG00000152359.14 | POC5 | protein_coding | 1888,073398 | 0,230345682 | 0,079764645 | 2,887816807 | 0,003879257 | 0,013038406 |
| ENSG00000152402.10 | GUCY1A2 | protein_coding | 364,2216824 | -0,566645058 | 0,143127281 | -3,959029009 | 7,53E-05 | 0,000366094 |
| ENSG00000152439.12 | ZNF773 | protein_coding | 1348,31002 | 0,294295886 | 0,075385212 | 3,903894115 | 9,47E-05 | 0,000452269 |
| ENSG00000152443.13 | ZNF776 | protein_coding | 6112,156795 | 0,31370485 | 0,050749469 | 6,181440992 | 6,35E-10 | 6,08E-09 |
| ENSG00000152454.4 | ZNF256 | protein_coding | 1057,649914 | 0,303474028 | 0,080888046 | 3,751778474 | 0,000175585 | 0,00079634 |
| ENSG00000152457.18 | DCLRE1C | protein_coding | 2066,300281 | 1,42144498 | 0,074522812 | 19,07395782 | 4,16E-81 | 3,88E-79 |
| ENSG00000152484.14 | USP12 | protein_coding | 6230,064827 | -0,268093639 | 0,048692812 | -5,505815542 | 3,67E-08 | 2,86E-07 |
| ENSG00000152520.14 | PAN3 | protein_coding | 3999,501679 | 0,162090454 | 0,05350055 | 3,029696998 | 0,002447992 | 0,008638008 |
| ENSG00000152527.14 | PLEKHH2 | protein_coding | 689,9450926 | 0,473357722 | 0,115879452 | 4,08491509 | 4,41E-05 | 0,000223232 |
| ENSG00000152556.17 | PFKM | protein_coding | 10399,55806 | -0,132888224 | 0,054537998 | -2,436617191 | 0,014825365 | 0,041985925 |
| ENSG00000152558.15 | TMEM123 | protein_coding | 5663,59561 | 0,186402821 | 0,054555212 | 3,416773837 | 0,000633679 | 0,002569996 |
| ENSG00000152601.17 | MBNL1 | protein_coding | 14729,59087 | 0,283268455 | 0,047541019 | 5,958401036 | 2,55E-09 | 2,27E-08 |
| ENSG00000152642.10 | GPD1L | protein_coding | 6098,239155 | -0,382289534 | 0,069745701 | -5,481191336 | 4,22E-08 | 3,26E-07 |
| ENSG00000152684.11 | PELO | protein_coding | 3642,980076 | 0,443393989 | 0,057095022 | 7,765895732 | 8,11E-15 | 1,20E-13 |
| ENSG00000152689.18 | RASGRP3 | protein_coding | 746,3129049 | 4,098520601 | 0,280788963 | 14,59644483 | 2,96E-48 | 1,62E-46 |
| ENSG00000152749.8 | GPR180 | protein_coding | 11382,58833 | 0,772661114 | 0,044890327 | 17,21219618 | 2,15E-66 | 1,65E-64 |
| ENSG00000152767.16 | FARP1 | protein_coding | 14130,8475 | -0,600827716 | 0,070186438 | -8,560453168 | 1,12E-17 | 2,01E-16 |
| ENSG00000152778.9 | IFIT5 | protein_coding | 12460,77823 | 3,961554316 | 0,049018503 | 80,81753009 | 0 | 0 |
| ENSG00000152779.14 | SLC16A12 | protein_coding | 7494,286787 | 0,287452852 | 0,05100746 | 5,635506114 | 1,75E-08 | 1,42E-07 |
| ENSG00000152782.16 | PANK1 | protein_coding | 4843,290093 | 0,191606235 | 0,050776601 | 3,773514393 | 0,000160964 | 0,000735364 |
| ENSG00000152795.17 | HNRNPDL | protein_coding | 43982,3621 | 0,137607906 | 0,040408797 | 3,405394804 | 0,000660685 | 0,002668464 |
| ENSG00000152818.18 | UTRN | protein_coding | 10160,93793 | 0,659434554 | 0,073562824 | 8,964236527 | 3,12E-19 | 6,09E-18 |
| ENSG00000152822.14 | GRM1 | protein_coding | 989,6673048 | 0,601641891 | 0,099735126 | 6,032397185 | 1,62E-09 | 1,48E-08 |
| ENSG00000152894.14 | PTPRK | protein_coding | 6824,597841 | 0,272148478 | 0,064412783 | 4,225069416 | 2,39E-05 | 0,000126242 |
| ENSG00000152904.11 | GGPS1 | protein_coding | 5702,877485 | 0,285824984 | 0,050372399 | 5,67423811 | 1,39E-08 | 1,14E-07 |
| ENSG00000152910.19 | CNTNAP4 | protein_coding | 140,8209669 | -1,078229494 | 0,22416322 | -4,810019647 | 1,51E-06 | 9,54E-06 |
| ENSG00000152932.8 | RAB3C | protein_coding | 7829,14791 | -0,296181347 | 0,045288854 | -6,539828766 | 6,16E-11 | 6,55E-10 |
| ENSG00000152939.17 | MARVELD2 | protein_coding | 6345,378599 | 0,199443312 | 0,048549977 | 4,108000154 | 3,99E-05 | 0,000203197 |
| ENSG00000152952.12 | PLOD2 | protein_coding | 33154,82469 | 0,174645751 | 0,04567618 | 3,823562949 | 0,000131537 | 0,00061237 |
| ENSG00000152954.12 | NRSN1 | protein_coding | 1078,697921 | -0,582623886 | 0,085063901 | -6,849249583 | 7,42E-12 | 8,63E-11 |
| ENSG00000152990.14 | ADGRA3 | protein_coding | 3189,222382 | -0,422962677 | 0,056926566 | -7,429970052 | 1,09E-13 | 1,49E-12 |
| ENSG00000153006.16 | SREK1IP1 | protein_coding | 10940,43457 | -0,149147739 | 0,048466254 | -3,077352318 | 0,002088483 | 0,007504735 |
| ENSG00000153012.12 | LGI2 | protein_coding | 794,3144874 | -0,791836385 | 0,114527818 | -6,913921844 | 4,71E-12 | 5,60E-11 |

| | | | | | | | | |
|---|---|---|---|---|---|---|---|---|
| ENSG00000153029.14 | MR1 | protein_coding | 1931,893733 | 1,591319917 | 0,08978113 | 17,72443625 | 2,72E-70 | 2,20E-68 |
| ENSG00000153048.11 | CARHSP1 | protein_coding | 36745,27018 | -0,097444519 | 0,038806195 | -2,511055762 | 0,012037068 | 0,035082528 |
| ENSG00000153066.12 | TXNDC11 | protein_coding | 9353,159127 | 0,872881262 | 0,04438122 | 19,66780702 | 4,07E-86 | 4,15E-84 |
| ENSG00000153093.18 | ACOXL | protein_coding | 11,38343088 | 2,637100305 | 0,803110876 | 3,283606767 | 0,001024878 | 0,003961701 |
| ENSG00000153094.23 | BCL2L11 | protein_coding | 6152,892174 | 0,236765549 | 0,062974601 | 3,759699093 | 0,000170118 | 0,000773416 |
| ENSG00000153113.23 | CAST | protein_coding | 14970,84108 | 0,456639474 | 0,046389836 | 9,843524167 | 7,31E-23 | 1,72E-21 |
| ENSG00000153130.17 | SCOC | protein_coding | 34907,06117 | -0,283056538 | 0,045272782 | -6,252245297 | 4,05E-10 | 3,96E-09 |
| ENSG00000153132.13 | CLGN | protein_coding | 14976,23427 | -0,339363234 | 0,045924336 | -7,389616626 | 1,47E-13 | 2,00E-12 |
| ENSG00000153140.8 | CETN3 | protein_coding | 4047,221699 | -0,134449086 | 0,053057916 | -2,534006158 | 0,011276675 | 0,033153955 |
| ENSG00000153147.6 | SMARCA5 | protein_coding | 37305,61922 | 0,211087524 | 0,040299676 | 5,237945978 | 1,62E-07 | 1,17E-06 |
| ENSG00000153214.11 | TMEM87B | protein_coding | 4742,481749 | 0,210182294 | 0,058500003 | 3,592859571 | 0,000327069 | 0,001410344 |
| ENSG00000153234.14 | NR4A2 | protein_coding | 343,7673145 | 0,691811452 | 0,128861686 | 5,368635746 | 7,93E-08 | 5,91E-07 |
| ENSG00000153246.13 | PLA2R1 | protein_coding | 22,11795859 | 1,655023087 | 0,615549894 | 2,688690394 | 0,007173292 | 0,022428618 |
| ENSG00000153250.20 | RBMS1 | protein_coding | 6698,85633 | -0,135299693 | 0,046092538 | -2,935392586 | 0,003331259 | 0,011388717 |
| ENSG00000153253.18 | SCN3A | protein_coding | 1379,32174 | 0,314416731 | 0,081020854 | 3,880688943 | 0,000104161 | 0,000494216 |
| ENSG00000153310.19 | FAM49B | protein_coding | 7702,891381 | -0,1508696 | 0,044982533 | -3,353959644 | 0,00079664 | 0,003158634 |
| ENSG00000153317.15 | ASAP1 | protein_coding | 11625,70346 | 0,108170676 | 0,042560709 | 2,541561885 | 0,011035841 | 0,032552187 |
| ENSG00000153363.13 | LINC00467 | lncRNA | 1437,209582 | -0,213039244 | 0,077695665 | -2,741970787 | 0,006107177 | 0,019471169 |
| ENSG00000153395.10 | LPCAT1 | protein_coding | 13644,45975 | -0,170888172 | 0,049005054 | -3,487154009 | 0,00048819 | 0,002028179 |
| ENSG00000153404.14 | PLEKHG4B | protein_coding | 35,30554458 | 1,091388065 | 0,391905601 | 2,784823852 | 0,005355681 | 0,017340481 |
| ENSG00000153487.12 | ING1 | protein_coding | 2049,15869 | -0,16318554 | 0,065632836 | -2,486339928 | 0,012906462 | 0,03724507 |
| ENSG00000153531.13 | ADPRHL1 | protein_coding | 551,0588442 | -0,370641975 | 0,114941688 | -3,224608755 | 0,001261449 | 0,004776108 |
| ENSG00000153558.15 | FBXL2 | protein_coding | 1546,793336 | 0,295088125 | 0,079657965 | 3,704439645 | 0,000211859 | 0,000945613 |
| ENSG00000153560.12 | UBP1 | protein_coding | 11638,15319 | 0,217290229 | 0,048342167 | 4,494838438 | 6,96E-06 | 3,99E-05 |
| ENSG00000153561.13 | RMND5A | protein_coding | 11805,0961 | -0,399125341 | 0,048325008 | -8,25918837 | 1,47E-16 | 2,44E-15 |
| ENSG00000153574.9 | RPIA | protein_coding | 1942,480724 | -0,244150973 | 0,067769882 | -3,602647177 | 0,000314993 | 0,001363857 |
| ENSG00000153714.6 | LURAP1L | protein_coding | 368,2682623 | 0,994685643 | 0,148407012 | 6,702416748 | 2,05E-11 | 2,28E-10 |
| ENSG00000153721.19 | CNKSR3 | protein_coding | 1632,112867 | 0,478145102 | 0,068544009 | 6,975738738 | 3,04E-12 | 3,68E-11 |
| ENSG00000153774.9 | CFDP1 | protein_coding | 8351,156205 | -0,285053921 | 0,049465678 | -5,762660736 | 8,28E-09 | 6,95E-08 |
| ENSG00000153789.13 | FAM92B | protein_coding | 1031,313004 | -1,628848225 | 0,110427202 | -14,75042551 | 3,06E-49 | 1,72E-47 |
| ENSG00000153790.12 | C7orf31 | protein_coding | 766,14643 | 0,561420503 | 0,100881885 | 5,565126998 | 2,62E-08 | 2,09E-07 |
| ENSG00000153822.13 | KCNJ16 | protein_coding | 4098,954638 | 0,234151195 | 0,05704202 | 4,104889594 | 4,05E-05 | 0,000205764 |
| ENSG00000153827.13 | TRIP12 | protein_coding | 28854,42326 | 0,633018573 | 0,050484412 | 12,53889178 | 4,57E-36 | 1,83E-34 |
| ENSG00000153898.13 | MCOLN2 | protein_coding | 42,50352259 | 1,625754158 | 0,440983558 | 3,686654817 | 0,000227221 | 0,001009777 |
| ENSG00000153904.20 | DDAH1 | protein_coding | 9000,9062 | -0,428203439 | 0,051981282 | -8,237646778 | 1,76E-16 | 2,91E-15 |
| ENSG00000153922.10 | CHD1 | protein_coding | 9411,476206 | 0,445997608 | 0,056361161 | 7,913208381 | 2,51E-15 | 3,86E-14 |

| | | | | | | | | |
|---|---|---|---|---|---|---|---|---|
| ENSG00000153933.10 | DGKE | protein_coding | 2906,418834 | 0,711777221 | 0,063564427 | 11,19772893 | 4,18E-29 | 1,27E-27 |
| ENSG00000153936.17 | HS2ST1 | protein_coding | 14695,81699 | -0,286700648 | 0,042137864 | -6,803872282 | 1,02E-11 | 1,17E-10 |
| ENSG00000153944.11 | MSI2 | protein_coding | 8639,867407 | -0,141268114 | 0,055175353 | -2,560348186 | 0,010456734 | 0,03107203 |
| ENSG00000153956.16 | CACNA2D1 | protein_coding | 73059,46472 | -0,606974627 | 0,055919214 | -10,85449132 | 1,90E-27 | 5,41E-26 |
| ENSG00000153975.10 | ZUP1 | protein_coding | 3528,128957 | 0,231892226 | 0,05304919 | 4,371267998 | 1,24E-05 | 6,82E-05 |
| ENSG00000153989.8 | NUS1 | protein_coding | 11924,84609 | -0,23657749 | 0,048540368 | -4,873829797 | 1,09E-06 | 7,08E-06 |
| ENSG00000154001.14 | PPP2R5E | protein_coding | 7736,229031 | -0,146328932 | 0,050490589 | -2,898142694 | 0,003753798 | 0,012664003 |
| ENSG00000154016.13 | GRAP | protein_coding | 294,009281 | -0,449103323 | 0,138910833 | -3,233033113 | 0,001224834 | 0,004649204 |
| ENSG00000154059.11 | IMPACT | protein_coding | 2942,400999 | -0,367747281 | 0,060855546 | -6,042954308 | 1,51E-09 | 1,39E-08 |
| ENSG00000154065.17 | ANKRD29 | protein_coding | 265,4560316 | 0,903912587 | 0,167575688 | 5,394055642 | 6,89E-08 | 5,18E-07 |
| ENSG00000154079.6 | SDHAF4 | protein_coding | 1213,56967 | -0,446932986 | 0,090369543 | -4,945615188 | 7,59E-07 | 5,01E-06 |
| ENSG00000154080.14 | CHST9 | protein_coding | 26275,14359 | -0,211835879 | 0,044238683 | -4,788476168 | 1,68E-06 | 1,06E-05 |
| ENSG00000154114.12 | TBCEL | protein_coding | 4778,706207 | 0,25242165 | 0,051874632 | 4,86599409 | 1,14E-06 | 7,34E-06 |
| ENSG00000154122.14 | ANKH | protein_coding | 8319,584614 | -0,175084932 | 0,060341747 | -2,901555555 | 0,003713149 | 0,012540328 |
| ENSG00000154127.10 | UBASH3B | protein_coding | 386,9804032 | 0,364855446 | 0,130558255 | 2,79457967 | 0,005196723 | 0,016898736 |
| ENSG00000154144.13 | TBRG1 | protein_coding | 5469,820639 | 0,20257601 | 0,055387567 | 3,657427508 | 0,000254759 | 0,001121309 |
| ENSG00000154162.14 | CDH12 | protein_coding | 7290,570127 | 0,194618622 | 0,047598403 | 4,088763714 | 4,34E-05 | 0,000219857 |
| ENSG00000154222.14 | CC2D1B | protein_coding | 5895,026977 | 0,25994761 | 0,059926343 | 4,337785319 | 1,44E-05 | 7,86E-05 |
| ENSG00000154229.12 | PRKCA | protein_coding | 6259,000451 | -0,712395402 | 0,067253559 | -10,59267952 | 3,22E-26 | 8,81E-25 |
| ENSG00000154262.13 | ABCA6 | protein_coding | 48,11705894 | 1,178540935 | 0,406194607 | 2,901419452 | 0,003714763 | 0,012543593 |
| ENSG00000154265.16 | ABCA5 | protein_coding | 11743,52698 | 0,2028852 | 0,046034617 | 4,407231204 | 1,05E-05 | 5,85E-05 |
| ENSG00000154274.15 | C4orf19 | protein_coding | 3954,555265 | 0,379134084 | 0,052243787 | 7,257017609 | 3,96E-13 | 5,16E-12 |
| ENSG00000154305.18 | MIA3 | protein_coding | 12740,29547 | 0,480615622 | 0,041991277 | 11,44560626 | 2,47E-30 | 7,94E-29 |
| ENSG00000154316.16 | TDH | transcribed_unitary | 30,52240228 | 1,136365527 | 0,437830105 | 2,595448587 | 0,009446753 | 0,028530387 |
| ENSG00000154319.16 | FAM167A | protein_coding | 11359,12132 | 0,13671209 | 0,045392997 | 3,011744081 | 0,002597515 | 0,009118444 |
| ENSG00000154328.16 | NEIL2 | protein_coding | 953,6714459 | -0,221106191 | 0,093334049 | -2,368976739 | 0,017837375 | 0,049215675 |
| ENSG00000154330.13 | PGM5 | protein_coding | 2796,324076 | -0,505808913 | 0,069838919 | -7,242507759 | 4,40E-13 | 5,72E-12 |
| ENSG00000154370.16 | TRIM11 | protein_coding | 1622,758088 | 0,229366071 | 0,069914579 | 3,280661552 | 0,001035639 | 0,003999178 |
| ENSG00000154429.11 | CCSAP | protein_coding | 5784,776737 | 0,345144476 | 0,054813679 | 6,296685142 | 3,04E-10 | 3,02E-09 |
| ENSG00000154451.14 | GBP5 | protein_coding | 24,12750304 | 3,148418442 | 0,689854498 | 4,56388768 | 5,02E-06 | 2,94E-05 |
| ENSG00000154473.18 | BUB3 | protein_coding | 8105,502939 | -0,370090184 | 0,046289465 | -7,995127692 | 1,29E-15 | 2,02E-14 |
| ENSG00000154511.12 | DIPK1A | protein_coding | 1001,253502 | 0,236730069 | 0,089781206 | 2,63674415 | 0,008370594 | 0,025669441 |
| ENSG00000154518.9 | ATP5MC3 | protein_coding | 21478,81907 | -0,246343258 | 0,045952019 | -5,360879904 | 8,28E-08 | 6,15E-07 |
| ENSG00000154553.15 | PDLIM3 | protein_coding | 41,80580006 | 1,187796416 | 0,443549689 | 2,677933148 | 0,0074078 | 0,023059352 |
| ENSG00000154556.18 | SORBS2 | protein_coding | 12291,33476 | -0,146553785 | 0,055482889 | -2,641423116 | 0,008255854 | 0,025369423 |
| ENSG00000154639.19 | CXADR | protein_coding | 4817,220968 | -0,155250347 | 0,059570127 | -2,606177874 | 0,00915589 | 0,027759958 |

| | | | | | | | | |
|---|---|---|---|---|---|---|---|---|
| ENSG00000154642.11 | C21orf91 | protein_coding | 3182,803983 | 0,411117852 | 0,065564098 | 6,270472129 | 3,60E-10 | 3,55E-09 |
| ENSG00000154645.14 | CHODL | protein_coding | 1933,730177 | 0,34574003 | 0,064154743 | 5,389157723 | 7,08E-08 | 5,31E-07 |
| ENSG00000154655.15 | L3MBTL4 | protein_coding | 79,57983753 | 1,438515956 | 0,281315597 | 5,113530743 | 3,16E-07 | 2,20E-06 |
| ENSG00000154723.12 | ATP5PF | protein_coding | 13007,68419 | -0,272977482 | 0,047916539 | -5,696936532 | 1,22E-08 | 1,00E-07 |
| ENSG00000154727.10 | GABPA | protein_coding | 4723,628595 | -0,484091807 | 0,056283066 | -8,601020521 | 7,90E-18 | 1,42E-16 |
| ENSG00000154736.6 | ADAMTS5 | protein_coding | 924,1222765 | -0,413410639 | 0,093935059 | -4,401026038 | 1,08E-05 | 6,00E-05 |
| ENSG00000154767.14 | XPC | protein_coding | 6688,289816 | 0,229329158 | 0,054135262 | 4,236225181 | 2,27E-05 | 0,000120445 |
| ENSG00000154781.16 | CCDC174 | protein_coding | 2539,722572 | 0,236866145 | 0,071765761 | 3,300545299 | 0,000964971 | 0,003756579 |
| ENSG00000154803.13 | FLCN | protein_coding | 6593,802091 | 0,245216788 | 0,063872645 | 3,839151915 | 0,00012346 | 0,000578468 |
| ENSG00000154813.10 | DPH3 | protein_coding | 5824,964396 | -0,230464068 | 0,052701547 | -4,373003828 | 1,23E-05 | 6,77E-05 |
| ENSG00000154822.18 | PLCL2 | protein_coding | 5888,060186 | 0,304826588 | 0,051933688 | 5,869534786 | 4,37E-09 | 3,80E-08 |
| ENSG00000154832.14 | CXXC1 | protein_coding | 4914,129396 | -0,282789303 | 0,05231173 | -5,405848796 | 6,45E-08 | 4,88E-07 |
| ENSG00000154856.13 | APCDD1 | protein_coding | 154,3299061 | 0,490245403 | 0,185126714 | 2,648161327 | 0,008093089 | 0,024927114 |
| ENSG00000154874.15 | CCDC144B | transcribed_unproc | 1759,438559 | 0,264600565 | 0,069161949 | 3,825811262 | 0,000130342 | 0,00060793 |
| ENSG00000154889.16 | MPPE1 | protein_coding | 2729,128046 | 0,432747693 | 0,066887067 | 6,469826142 | 9,81E-11 | 1,02E-09 |
| ENSG00000154914.17 | USP43 | protein_coding | 805,3946941 | 0,320438206 | 0,094553623 | 3,388957464 | 0,000701589 | 0,002817496 |
| ENSG00000154917.11 | RAB6B | protein_coding | 2842,126926 | -0,258624511 | 0,083723397 | -3,0890351 | 0,002008077 | 0,007250531 |
| ENSG00000154920.14 | EME1 | protein_coding | 1344,071017 | -0,364230988 | 0,08845924 | -4,11750077 | 3,83E-05 | 0,000195532 |
| ENSG00000154930.15 | ACSS1 | protein_coding | 44,69603341 | 2,538449737 | 0,427697731 | 5,93514895 | 2,94E-09 | 2,60E-08 |
| ENSG00000154957.14 | ZNF18 | protein_coding | 680,8013738 | 0,304881171 | 0,095383036 | 3,196387767 | 0,0013916 | 0,005222326 |
| ENSG00000154975.14 | CA10 | protein_coding | 382,838991 | -0,324314553 | 0,125335773 | -2,587565747 | 0,009665675 | 0,029074525 |
| ENSG00000155008.14 | APOOL | protein_coding | 5246,286709 | -0,525396331 | 0,054266303 | -9,681815533 | 3,60E-22 | 8,17E-21 |
| ENSG00000155016.18 | CYP2U1 | protein_coding | 9980,979611 | 0,256701503 | 0,04728641 | 5,428652833 | 5,68E-08 | 4,32E-07 |
| ENSG00000155034.19 | FBXL18 | protein_coding | 948,9910777 | -0,521845948 | 0,08967177 | -5,819512047 | 5,90E-09 | 5,05E-08 |
| ENSG00000155085.15 | AK9 | protein_coding | 2952,132511 | -0,317780018 | 0,064823998 | -4,902197123 | 9,48E-07 | 6,18E-06 |
| ENSG00000155090.15 | KLF10 | protein_coding | 1718,452927 | -0,255477398 | 0,06897036 | -3,70416218 | 0,000212091 | 0,00094646 |
| ENSG00000155093.19 | PTPRN2 | protein_coding | 22998,52495 | -0,316675222 | 0,042884569 | -7,384362845 | 1,53E-13 | 2,07E-12 |
| ENSG00000155096.14 | AZIN1 | protein_coding | 30034,16403 | -0,171733875 | 0,038689602 | -4,438760423 | 9,05E-06 | 5,11E-05 |
| ENSG00000155097.12 | ATP6V1C1 | protein_coding | 14752,38657 | -0,175755472 | 0,044432819 | -3,955532767 | 7,64E-05 | 0,000370889 |
| ENSG00000155099.8 | PIP4P2 | protein_coding | 3446,377345 | 0,150960797 | 0,05567837 | 2,711300593 | 0,006701984 | 0,021137026 |
| ENSG00000155100.10 | OTUD6B | protein_coding | 5559,381426 | -0,142143964 | 0,052984196 | -2,682761574 | 0,007301703 | 0,022763715 |
| ENSG00000155111.15 | CDK19 | protein_coding | 5583,477708 | -0,41093673 | 0,063596763 | -6,461598173 | 1,04E-10 | 1,08E-09 |
| ENSG00000155115.7 | GTF3C6 | protein_coding | 5587,863978 | -0,213667416 | 0,058237391 | -3,668904334 | 0,000243592 | 0,001075312 |
| ENSG00000155158.20 | TTC39B | protein_coding | 2524,159611 | 1,074679928 | 0,060947999 | 17,63273532 | 1,38E-69 | 1,11E-67 |
| ENSG00000155252.13 | PI4K2A | protein_coding | 4510,410416 | 0,227121908 | 0,051009167 | 4,452570439 | 8,48E-06 | 4,80E-05 |
| ENSG00000155256.17 | ZFYVE27 | protein_coding | 5052,436738 | 0,27586135 | 0,056678145 | 4,867155618 | 1,13E-06 | 7,30E-06 |

| | | | | | | | | |
|---|---|---|---|---|---|---|---|---|
| ENSG00000155265.11 | GOLGA7B | protein_coding | 765,9853385 | 0,390875327 | 0,109185319 | 3,579925689 | 0,000343692 | 0,001475522 |
| ENSG00000155287.11 | SLC25A28 | protein_coding | 2478,844606 | 1,568247957 | 0,073290439 | 21,39771538 | 1,40E-101 | 1,64E-99 |
| ENSG00000155307.18 | SAMSN1 | protein_coding | 51,75017794 | 0,884008939 | 0,351095924 | 2,517855888 | 0,011807162 | 0,034501806 |
| ENSG00000155313.15 | USP25 | protein_coding | 5588,237069 | 1,277365095 | 0,052870947 | 24,16005718 | 5,85E-129 | 8,61E-127 |
| ENSG00000155363.18 | MOV10 | protein_coding | 15548,20928 | 1,709926792 | 0,054816763 | 31,19350189 | 1,31E-213 | 2,75E-211 |
| ENSG00000155368.16 | DBI | protein_coding | 36414,18374 | -0,206533425 | 0,045307286 | -4,558503515 | 5,15E-06 | 3,01E-05 |
| ENSG00000155465.18 | SLC7A7 | protein_coding | 2058,716688 | -0,456688019 | 0,078979607 | -5,782353614 | 7,37E-09 | 6,22E-08 |
| ENSG00000155506.17 | LARP1 | protein_coding | 35379,21764 | 0,15412567 | 0,04500747 | 3,424446392 | 0,000616053 | 0,002503927 |
| ENSG00000155511.18 | GRIA1 | protein_coding | 68,50266935 | 6,309711857 | 1,25973338 | 5,008767694 | 5,48E-07 | 3,70E-06 |
| ENSG00000155542.11 | SETD9 | protein_coding | 745,9996506 | -0,428926915 | 0,11519837 | -3,723376599 | 0,000196576 | 0,000883356 |
| ENSG00000155545.19 | MIER3 | protein_coding | 6066,221685 | 0,292477544 | 0,085827407 | 3,407740644 | 0,000655031 | 0,002647055 |
| ENSG00000155592.16 | ZKSCAN2 | protein_coding | 4078,089175 | 0,308153756 | 0,053201388 | 5,792212755 | 6,95E-09 | 5,90E-08 |
| ENSG00000155621.15 | C9orf85 | protein_coding | 1562,246117 | 0,248207655 | 0,076494687 | 3,244769865 | 0,001175456 | 0,004479908 |
| ENSG00000155629.15 | PIK3AP1 | protein_coding | 3914,828018 | 4,51636627 | 0,219803303 | 20,54730845 | 8,14E-94 | 8,89E-92 |
| ENSG00000155660.11 | PDIA4 | protein_coding | 26698,11145 | 0,677994393 | 0,050028851 | 13,55206798 | 7,70E-42 | 3,63E-40 |
| ENSG00000155666.11 | KDM8 | protein_coding | 612,3450157 | -0,317204597 | 0,103303242 | -3,0706161 | 0,002136176 | 0,007656559 |
| ENSG00000155744.9 | FAM126B | protein_coding | 7919,114881 | 0,141980995 | 0,049047959 | 2,894738091 | 0,00379475 | 0,012783004 |
| ENSG00000155760.2 | FZD7 | protein_coding | 570,4292451 | 0,496401082 | 0,105104976 | 4,722907539 | 2,32E-06 | 1,43E-05 |
| ENSG00000155792.10 | DEPTOR | protein_coding | 1406,791742 | 1,135013915 | 0,091550611 | 12,39766618 | 2,69E-35 | 1,04E-33 |
| ENSG00000155816.20 | FMN2 | protein_coding | 2627,09217 | -0,203008115 | 0,071473228 | -2,840337847 | 0,004506578 | 0,014873689 |
| ENSG00000155849.15 | ELMO1 | protein_coding | 357,4525242 | 0,84107532 | 0,147053256 | 5,719528704 | 1,07E-08 | 8,86E-08 |
| ENSG00000155850.7 | SLC26A2 | protein_coding | 3212,171248 | -0,547549472 | 0,055146769 | -9,928949284 | 3,12E-23 | 7,43E-22 |
| ENSG00000155858.6 | LSM11 | protein_coding | 2462,208651 | -0,250936692 | 0,066816982 | -3,755582558 | 0,000172939 | 0,000785923 |
| ENSG00000155868.8 | MED7 | protein_coding | 2789,811354 | -0,238957948 | 0,068378882 | -3,494616181 | 0,000474744 | 0,001977426 |
| ENSG00000155886.11 | SLC24A2 | protein_coding | 56,69648566 | 1,045108499 | 0,308052861 | 3,392627154 | 0,000692258 | 0,002783036 |
| ENSG00000155893.13 | PXYLP1 | protein_coding | 796,0841794 | 0,398827975 | 0,102069422 | 3,907418758 | 9,33E-05 | 0,000446482 |
| ENSG00000155897.10 | ADCY8 | protein_coding | 9602,697106 | 0,644460341 | 0,055191296 | 11,67684748 | 1,67E-31 | 5,69E-30 |
| ENSG00000155903.11 | RASA2 | protein_coding | 2413,661089 | 0,432092344 | 0,061153082 | 7,065749256 | 1,60E-12 | 1,99E-11 |
| ENSG00000155959.11 | VBP1 | protein_coding | 11367,36335 | -0,264414132 | 0,046640436 | -5,669203729 | 1,43E-08 | 1,17E-07 |
| ENSG00000155961.5 | RAB39B | protein_coding | 3934,234866 | 0,153728626 | 0,054714245 | 2,80966367 | 0,00495933 | 0,016204028 |
| ENSG00000155970.12 | MICU3 | protein_coding | 1305,048614 | 0,253948031 | 0,077006686 | 3,297740022 | 0,000974663 | 0,003789067 |
| ENSG00000156011.17 | PSD3 | protein_coding | 14231,19351 | -0,67503192 | 0,063035957 | -10,70868051 | 9,27E-27 | 2,58E-25 |
| ENSG00000156017.13 | CARNMT1 | protein_coding | 2738,692821 | -0,18544485 | 0,06707938 | -2,764558211 | 0,005699994 | 0,018326001 |
| ENSG00000156030.13 | ELMSAN1 | protein_coding | 1957,563813 | 0,376685708 | 0,078351414 | 4,807644047 | 1,53E-06 | 9,65E-06 |
| ENSG00000156042.18 | CFAP70 | protein_coding | 638,3624991 | 0,387349894 | 0,122132658 | 3,17155052 | 0,001516275 | 0,005650634 |
| ENSG00000156050.9 | FAM161B | protein_coding | 1151,258617 | 0,585410252 | 0,076473246 | 7,655098738 | 1,93E-14 | 2,78E-13 |

| | | | | | | | | |
|---|---|---|---|---|---|---|---|---|
| ENSG00000156052.10 | GNAQ | protein_coding | 11080,97701 | -0,41221295 | 0,050111433 | -8,22592626 | 1,94E-16 | 3,20E-15 |
| ENSG00000156096.14 | UGT2B4 | protein_coding | 194,8947326 | 0,45063605 | 0,185131966 | 2,434134199 | 0,01492746 | 0,042248483 |
| ENSG00000156103.16 | MMP16 | protein_coding | 1164,575882 | -0,494492869 | 0,085968262 | -5,752039872 | 8,82E-09 | 7,38E-08 |
| ENSG00000156110.13 | ADK | protein_coding | 3363,814364 | -0,50951912 | 0,058802732 | -8,664888532 | 4,52E-18 | 8,26E-17 |
| ENSG00000156113.23 | KCNMA1 | protein_coding | 7684,224042 | -0,64063266 | 0,047488403 | -13,49029706 | 1,78E-41 | 8,31E-40 |
| ENSG00000156136.10 | DCK | protein_coding | 14238,83779 | -0,215326877 | 0,04498489 | -4,786648983 | 1,70E-06 | 1,07E-05 |
| ENSG00000156162.16 | DPY19L4 | protein_coding | 7168,027854 | -0,274052451 | 0,058534852 | -4,681867972 | 2,84E-06 | 1,73E-05 |
| ENSG00000156170.13 | NDUFAF6 | protein_coding | 1133,777937 | -0,233598958 | 0,088213913 | -2,648096557 | 0,00809464 | 0,024928479 |
| ENSG00000156171.14 | DRAM2 | protein_coding | 3749,854033 | 0,131531761 | 0,053853602 | 2,442394854 | 0,014590179 | 0,04142411 |
| ENSG00000156172.6 | C8orf37 | protein_coding | 1291,513211 | -0,190175957 | 0,075624415 | -2,51474287 | 0,011911923 | 0,034753787 |
| ENSG00000156232.7 | WHAMM | protein_coding | 2827,092737 | 1,140660593 | 0,07046235 | 16,18822817 | 6,11E-59 | 4,22E-57 |
| ENSG00000156261.13 | CCT8 | protein_coding | 27394,08982 | -0,147761536 | 0,039747809 | -3,717476212 | 0,000201223 | 0,000901538 |
| ENSG00000156298.12 | TSPAN7 | protein_coding | 27827,57911 | -0,221809147 | 0,048217268 | -4,600201447 | 4,22E-06 | 2,50E-05 |
| ENSG00000156304.14 | SCAF4 | protein_coding | 6091,223594 | 0,216326541 | 0,049170923 | 4,399481036 | 1,09E-05 | 6,04E-05 |
| ENSG00000156381.9 | ANKRD9 | protein_coding | 1389,631298 | -0,358282046 | 0,078348822 | -4,57290913 | 4,81E-06 | 2,83E-05 |
| ENSG00000156398.13 | SFXN2 | protein_coding | 3997,992179 | -0,189296528 | 0,058833011 | -3,217522344 | 0,00129303 | 0,004886636 |
| ENSG00000156411.9 | ATP5MPL | protein_coding | 9284,240284 | -0,588772094 | 0,050297319 | -11,70583447 | 1,19E-31 | 4,11E-30 |
| ENSG00000156427.8 | FGF18 | protein_coding | 640,5495158 | 0,363029241 | 0,111769263 | 3,248023917 | 0,001162095 | 0,004434994 |
| ENSG00000156453.13 | PCDH1 | protein_coding | 2827,546447 | 0,332013949 | 0,065155435 | 5,095721443 | 3,47E-07 | 2,41E-06 |
| ENSG00000156467.9 | UQCRB | protein_coding | 31575,21796 | -0,509089133 | 0,044943443 | -11,32732837 | 9,61E-30 | 2,98E-28 |
| ENSG00000156475.18 | PPP2R2B | protein_coding | 2044,350512 | 0,426124497 | 0,073500287 | 5,797589551 | 6,73E-09 | 5,73E-08 |
| ENSG00000156482.11 | RPL30 | protein_coding | 40012,65365 | -0,47764019 | 0,046720814 | -10,2232849 | 1,56E-24 | 3,96E-23 |
| ENSG00000156500.15 | FAM122C | protein_coding | 901,7383626 | 1,147449693 | 0,113971302 | 10,06788262 | 7,66E-24 | 1,89E-22 |
| ENSG00000156504.16 | FAM122B | protein_coding | 5225,277426 | -0,209355177 | 0,056401902 | -3,711846023 | 0,000205753 | 0,000919821 |
| ENSG00000156508.18 | EEF1A1 | protein_coding | 516427,2192 | -1,083435627 | 0,038726301 | -27,97673908 | 3,12E-172 | 5,61E-170 |
| ENSG00000156509.14 | FBXO43 | protein_coding | 669,2066453 | -0,8152917 | 0,108112748 | -7,541124562 | 4,66E-14 | 6,56E-13 |
| ENSG00000156531.16 | PHF6 | protein_coding | 10467,99749 | -0,283126303 | 0,044323939 | -6,387661155 | 1,68E-10 | 1,72E-09 |
| ENSG00000156599.11 | ZDHHC5 | protein_coding | 8335,182316 | 0,134632046 | 0,048381926 | 2,782693008 | 0,005390979 | 0,017444568 |
| ENSG00000156642.16 | NPTN | protein_coding | 19221,26956 | 0,148229796 | 0,043216547 | 3,429931539 | 0,000603734 | 0,002457847 |
| ENSG00000156650.14 | KAT6B | protein_coding | 7451,175305 | -0,280427431 | 0,048464167 | -5,786283922 | 7,20E-09 | 6,10E-08 |
| ENSG00000156671.14 | SAMD8 | protein_coding | 4233,802467 | 0,198290156 | 0,050467002 | 3,929105117 | 8,53E-05 | 0,000411752 |
| ENSG00000156675.15 | RAB11FIP1 | protein_coding | 19753,67148 | 0,229294585 | 0,04501221 | 5,094053058 | 3,50E-07 | 2,43E-06 |
| ENSG00000156687.11 | UNC5D | protein_coding | 4488,923375 | -0,171749851 | 0,059998605 | -2,862564093 | 0,004202283 | 0,013996623 |
| ENSG00000156711.17 | MAPK13 | protein_coding | 14212,79051 | -0,201534107 | 0,044049349 | -4,575189266 | 4,76E-06 | 2,80E-05 |
| ENSG00000156802.13 | ATAD2 | protein_coding | 20967,99117 | -0,159100886 | 0,040066501 | -3,970920418 | 7,16E-05 | 0,000349727 |
| ENSG00000156804.7 | FBXO32 | protein_coding | 924,9372399 | -0,395106197 | 0,128712751 | -3,069674079 | 0,002142925 | 0,007676121 |

| | | | | | | | | |
|---|---|---|---|---|---|---|---|---|
| ENSG00000156869.13 | FRRS1 | protein_coding | 805,1530897 | 0,566439007 | 0,103471594 | 5,474343083 | 4,39E-08 | 3,39E-07 |
| ENSG00000156875.14 | MFSD14A | protein_coding | 11097,26533 | 0,200578282 | 0,045290344 | 4,428720636 | 9,48E-06 | 5,33E-05 |
| ENSG00000156931.15 | VPS8 | protein_coding | 6838,393534 | 0,261964618 | 0,049576643 | 5,284033052 | 1,26E-07 | 9,23E-07 |
| ENSG00000156958.15 | GALK2 | protein_coding | 4519,717133 | 0,2578295 | 0,056705214 | 4,546839336 | 5,45E-06 | 3,17E-05 |
| ENSG00000156959.9 | LHFPL4 | protein_coding | 2586,208118 | 0,198383418 | 0,071900371 | 2,759143192 | 0,005795314 | 0,018582153 |
| ENSG00000156966.7 | B3GNT7 | protein_coding | 25,725222 | 1,268656638 | 0,470055584 | 2,698950255 | 0,006955857 | 0,021851974 |
| ENSG00000156970.12 | BUB1B | protein_coding | 8083,004125 | -0,954722955 | 0,056092092 | -17,02063366 | 5,77E-65 | 4,33E-63 |
| ENSG00000156976.17 | EIF4A2 | protein_coding | 67768,08762 | 0,120757264 | 0,039431875 | 3,062427628 | 0,002195496 | 0,007841703 |
| ENSG00000157005.4 | SST | protein_coding | 7990,745505 | 0,35105858 | 0,052565503 | 6,678497469 | 2,41E-11 | 2,67E-10 |
| ENSG00000157014.11 | TATDN2 | protein_coding | 11280,38064 | 0,256571682 | 0,048593137 | 5,279998336 | 1,29E-07 | 9,42E-07 |
| ENSG00000157020.18 | SEC13 | protein_coding | 10883,94043 | -0,15525233 | 0,051161258 | -3,03456826 | 0,002408802 | 0,008518418 |
| ENSG00000157036.13 | EXOG | protein_coding | 2060,358269 | 0,391231389 | 0,098082353 | 3,988805097 | 6,64E-05 | 0,000326081 |
| ENSG00000157064.11 | NMNAT2 | protein_coding | 7273,516308 | 0,395403641 | 0,047401685 | 8,341552381 | 7,33E-17 | 1,24E-15 |
| ENSG00000157077.14 | ZFYVE9 | protein_coding | 6261,314277 | 0,401586682 | 0,046684401 | 8,602159906 | 7,82E-18 | 1,41E-16 |
| ENSG00000157106.16 | SMG1 | protein_coding | 23171,64435 | 0,346546506 | 0,04114721 | 8,422114236 | 3,70E-17 | 6,37E-16 |
| ENSG00000157212.18 | PAXIP1 | protein_coding | 4906,114952 | -0,150114899 | 0,052829117 | -2,841518243 | 0,004489928 | 0,014827435 |
| ENSG00000157240.3 | FZD1 | protein_coding | 7132,725652 | 0,150814122 | 0,054203177 | 2,782385305 | 0,005396093 | 0,017456262 |
| ENSG00000157259.8 | GATAD1 | protein_coding | 6955,830282 | 0,189248991 | 0,05052828 | 3,745407359 | 0,000180101 | 0,000813873 |
| ENSG00000157303.11 | SUSD3 | protein_coding | 12,99755394 | 2,324706169 | 0,669108444 | 3,474333926 | 0,000512123 | 0,002115595 |
| ENSG00000157315.5 | TMED6 | protein_coding | 4807,319617 | 0,169608552 | 0,055949781 | 3,031442663 | 0,002433881 | 0,008594954 |
| ENSG00000157326.19 | DHRS4 | protein_coding | 2172,049342 | -0,245841817 | 0,068798824 | -3,573343308 | 0,000352452 | 0,001508814 |
| ENSG00000157350.13 | ST3GAL2 | protein_coding | 2408,285777 | 0,470687739 | 0,184393211 | 2,552630519 | 0,010691284 | 0,031681657 |
| ENSG00000157399.14 | ARSE | protein_coding | 3870,875595 | 0,254841298 | 0,053876929 | 4,730063517 | 2,24E-06 | 1,39E-05 |
| ENSG00000157445.15 | CACNA2D3 | protein_coding | 1264,562106 | -0,290346767 | 0,084855203 | -3,421673131 | 0,000622371 | 0,002525955 |
| ENSG00000157450.15 | RNF111 | protein_coding | 5677,941191 | 0,330123904 | 0,072336902 | 4,563699776 | 5,03E-06 | 2,94E-05 |
| ENSG00000157456.8 | CCNB2 | protein_coding | 5637,189156 | -0,965435373 | 0,053022459 | -18,20804615 | 4,46E-74 | 3,84E-72 |
| ENSG00000157470.12 | FAM81A | protein_coding | 899,2177411 | -0,567778458 | 0,090817533 | -6,251859519 | 4,06E-10 | 3,97E-09 |
| ENSG00000157500.12 | APPL1 | protein_coding | 13799,25121 | -0,292703073 | 0,051185076 | -5,718523769 | 1,07E-08 | 8,91E-08 |
| ENSG00000157538.14 | VPS26C | protein_coding | 8245,510499 | 0,19144477 | 0,051122059 | 3,744856376 | 0,000180497 | 0,000815497 |
| ENSG00000157540.21 | DYRK1A | protein_coding | 10302,02415 | 0,232017872 | 0,049129395 | 4,722587563 | 2,33E-06 | 1,43E-05 |
| ENSG00000157557.13 | ETS2 | protein_coding | 4255,590725 | 0,932113128 | 0,06104954 | 15,26814347 | 1,25E-52 | 7,65E-51 |
| ENSG00000157578.13 | LCA5L | protein_coding | 532,9353208 | 0,434603371 | 0,135217732 | 3,214100437 | 0,001308539 | 0,00494193 |
| ENSG00000157600.12 | TMEM164 | protein_coding | 7549,211043 | -0,368067322 | 0,064843858 | -5,676209486 | 1,38E-08 | 1,13E-07 |
| ENSG00000157601.14 | MX1 | protein_coding | 77131,05376 | 10,20495779 | 0,091731278 | 111,2483982 | 0 | 0 |
| ENSG00000157637.13 | SLC38A10 | protein_coding | 5989,627986 | 0,193873549 | 0,049388447 | 3,925483791 | 8,66E-05 | 0,00041728 |
| ENSG00000157654.17 | PALM2-AKAP2 | protein_coding | 153,3827452 | 1,642508319 | 0,568716939 | 2,888094595 | 0,003875833 | 0,013030788 |

| | | | | | | | | |
|---|---|---|---|---|---|---|---|---|
| ENSG00000157657.14 | ZNF618 | protein_coding | 3535,920792 | 0,478559673 | 0,057764921 | 8,2846071 | 1,18E-16 | 1,98E-15 |
| ENSG00000157693.15 | TMEM268 | protein_coding | 1008,388323 | 0,52922405 | 0,11373232 | 4,653242361 | 3,27E-06 | 1,97E-05 |
| ENSG00000157734.14 | SNX22 | protein_coding | 828,2109295 | -0,565252192 | 0,100736975 | -5,611168981 | 2,01E-08 | 1,62E-07 |
| ENSG00000157764.13 | BRAF | protein_coding | 11261,60434 | 0,243264884 | 0,048365508 | 5,029718388 | 4,91E-07 | 3,33E-06 |
| ENSG00000157800.18 | SLC37A3 | protein_coding | 6588,558314 | 0,489244226 | 0,0580195 | 8,432410303 | 3,39E-17 | 5,84E-16 |
| ENSG00000157833.13 | GAREM2 | protein_coding | 1068,701343 | -0,610664409 | 0,09119477 | -6,696265707 | 2,14E-11 | 2,37E-10 |
| ENSG00000157851.17 | DPYSL5 | protein_coding | 10439,80802 | -0,596603981 | 0,071311282 | -8,366193405 | 5,95E-17 | 1,01E-15 |
| ENSG00000157856.12 | DRC1 | protein_coding | 50,83521167 | 1,467128347 | 0,37660286 | 3,89569093 | 9,79E-05 | 0,000466336 |
| ENSG00000157870.16 | PRXL2B | protein_coding | 3425,643499 | -0,646996214 | 0,05987163 | -10,80639057 | 3,21E-27 | 9,08E-26 |
| ENSG00000157873.17 | TNFRSF14 | protein_coding | 1723,110958 | 0,885836906 | 0,093450751 | 9,479184493 | 2,56E-21 | 5,57E-20 |
| ENSG00000157911.10 | PEX10 | protein_coding | 1488,784486 | -0,185156212 | 0,075709941 | -2,445599757 | 0,014461144 | 0,041119995 |
| ENSG00000157954.15 | WIPI2 | protein_coding | 3039,477178 | -0,153956866 | 0,062469633 | -2,464507286 | 0,013720177 | 0,039341079 |
| ENSG00000157985.19 | AGAP1 | protein_coding | 3048,174245 | -0,89558823 | 0,076904883 | -11,64540134 | 2,42E-31 | 8,20E-30 |
| ENSG00000158006.14 | PAFAH2 | protein_coding | 2692,170348 | -0,393446173 | 0,077105192 | -5,10271961 | 3,35E-07 | 2,32E-06 |
| ENSG00000158019.20 | BABAM2 | protein_coding | 2810,548097 | -0,185357943 | 0,060666865 | -3,055340731 | 0,00224805 | 0,008010345 |
| ENSG00000158023.10 | WDR66 | protein_coding | 554,4665075 | -0,525689859 | 0,146064753 | -3,599019268 | 0,00031942 | 0,001380128 |
| ENSG00000158042.8 | MRPL17 | protein_coding | 5978,224279 | 0,511851669 | 0,051004933 | 10,03533654 | 1,07E-23 | 2,60E-22 |
| ENSG00000158055.15 | GRHL3 | protein_coding | 932,1291028 | -0,648713899 | 0,116495769 | -5,568561871 | 2,57E-08 | 2,05E-07 |
| ENSG00000158062.20 | UBXN11 | protein_coding | 1329,338442 | -0,248310348 | 0,07535507 | -3,295204265 | 0,000983502 | 0,003818809 |
| ENSG00000158079.16 | PTPDC1 | protein_coding | 1600,298374 | -0,294684911 | 0,070319939 | -4,190630946 | 2,78E-05 | 0,000144943 |
| ENSG00000158089.15 | GALNT14 | protein_coding | 3414,833665 | 0,336066301 | 0,058146079 | 5,779689842 | 7,48E-09 | 6,32E-08 |
| ENSG00000158109.15 | TPRG1L | protein_coding | 3899,976668 | -0,296301405 | 0,054787757 | -5,408168218 | 6,37E-08 | 4,82E-07 |
| ENSG00000158122.12 | PRXL2C | protein_coding | 942,064344 | 0,258909602 | 0,085721245 | 3,020366784 | 0,002524687 | 0,008889127 |
| ENSG00000158158.12 | CNNM4 | protein_coding | 3291,927008 | 0,382832049 | 0,061193858 | 6,256053532 | 3,95E-10 | 3,87E-09 |
| ENSG00000158164.7 | TMSB15A | protein_coding | 6073,896281 | -0,907847737 | 0,054896557 | -16,53742563 | 1,97E-61 | 1,40E-59 |
| ENSG00000158169.13 | FANCC | protein_coding | 4187,076609 | -0,226236449 | 0,06785216 | -3,334255646 | 0,00085528 | 0,003368002 |
| ENSG00000158186.13 | MRAS | protein_coding | 149,7802251 | -0,644322812 | 0,227846942 | -2,827875617 | 0,004685801 | 0,015399721 |
| ENSG00000158201.10 | ABHD3 | protein_coding | 8881,406496 | 0,144943672 | 0,050026554 | 2,897334692 | 0,00376348 | 0,012694766 |
| ENSG00000158258.16 | CLSTN2 | protein_coding | 4625,432893 | -0,718117429 | 0,05393233 | -13,31515685 | 1,89E-40 | 8,59E-39 |
| ENSG00000158290.16 | CUL4B | protein_coding | 15009,75404 | 0,333610404 | 0,042246654 | 7,896729543 | 2,86E-15 | 4,38E-14 |
| ENSG00000158292.7 | GPR153 | protein_coding | 5830,604963 | -0,304591378 | 0,052861212 | -5,762095945 | 8,31E-09 | 6,97E-08 |
| ENSG00000158352.15 | SHROOM4 | protein_coding | 114,509505 | 1,293102106 | 0,266175095 | 4,858088262 | 1,19E-06 | 7,62E-06 |
| ENSG00000158373.8 | HIST1H2BD | protein_coding | 885,9423037 | 0,600537466 | 0,0990667 | 6,061950867 | 1,34E-09 | 1,24E-08 |
| ENSG00000158402.20 | CDC25C | protein_coding | 2024,384877 | -0,497210317 | 0,071066752 | -6,996384424 | 2,63E-12 | 3,20E-11 |
| ENSG00000158423.17 | RIBC1 | protein_coding | 317,6320995 | 0,588553068 | 0,138541152 | 4,248218385 | 2,15E-05 | 0,000114526 |
| ENSG00000158427.15 | TMSB15B | protein_coding | 4893,506602 | -0,656648628 | 0,050241471 | -13,06985275 | 4,90E-39 | 2,14E-37 |

| | | | | | | | | |
|---|---|---|---|---|---|---|---|---|
| ENSG00000158435.8 | CNOT11 | protein_coding | 8338,116151 | -0,147605043 | 0,053551966 | -2,756295518 | 0,005846015 | 0,018728716 |
| ENSG00000158458.20 | NRG2 | protein_coding | 313,9341075 | 2,316687612 | 0,170799796 | 13,56376097 | 6,57E-42 | 3,13E-40 |
| ENSG00000158467.16 | AHCYL2 | protein_coding | 5846,716349 | -0,318484564 | 0,054692791 | -5,823154322 | 5,77E-09 | 4,95E-08 |
| ENSG00000158470.5 | B4GALT5 | protein_coding | 25897,95417 | 0,15601985 | 0,03869972 | 4,031549875 | 5,54E-05 | 0,000275692 |
| ENSG00000158480.11 | SPATA2 | protein_coding | 1454,309798 | 0,197282951 | 0,074560359 | 2,645949596 | 0,008146195 | 0,025066673 |
| ENSG00000158545.15 | ZC3H18 | protein_coding | 6255,866237 | -0,163075267 | 0,049373878 | -3,302865308 | 0,000957023 | 0,003727572 |
| ENSG00000158555.15 | GDPD5 | protein_coding | 299,2249127 | 0,475451212 | 0,144160271 | 3,298073802 | 0,000973505 | 0,003785873 |
| ENSG00000158604.15 | TMED4 | protein_coding | 13122,43497 | -0,111050048 | 0,041692255 | -2,663565419 | 0,007731738 | 0,023951787 |
| ENSG00000158615.9 | PPP1R15B | protein_coding | 14318,28896 | 0,273063652 | 0,043169747 | 6,325347482 | 2,53E-10 | 2,53E-09 |
| ENSG00000158623.14 | COPG2 | protein_coding | 6403,885347 | -0,178767139 | 0,049775952 | -3,59143589 | 0,000328861 | 0,0014178 |
| ENSG00000158636.16 | EMSY | protein_coding | 3722,612597 | 0,172493349 | 0,064548938 | 2,672287936 | 0,007533597 | 0,023386256 |
| ENSG00000158669.11 | GPAT4 | protein_coding | 45217,83781 | 0,117316211 | 0,039817214 | 2,946369177 | 0,003215284 | 0,011024041 |
| ENSG00000158691.14 | ZSCAN12 | protein_coding | 3707,816215 | 0,498716217 | 0,054912061 | 9,082088903 | 1,07E-19 | 2,12E-18 |
| ENSG00000158711.13 | ELK4 | protein_coding | 6277,195068 | 0,93353941 | 0,055465289 | 16,83105655 | 1,45E-63 | 1,06E-61 |
| ENSG00000158716.8 | DUSP23 | protein_coding | 798,5412943 | -0,530147107 | 0,113862882 | -4,656013407 | 3,22E-06 | 1,94E-05 |
| ENSG00000158773.14 | USF1 | protein_coding | 4572,3736 | 0,474299129 | 0,061157925 | 7,755317553 | 8,81E-15 | 1,30E-13 |
| ENSG00000158796.17 | DEDD | protein_coding | 3197,555968 | 0,3315257 | 0,05847875 | 5,669165288 | 1,43E-08 | 1,17E-07 |
| ENSG00000158805.12 | ZNF276 | protein_coding | 1670,568754 | 0,207895446 | 0,074415203 | 2,793722763 | 0,005210513 | 0,016928902 |
| ENSG00000158813.18 | EDA | protein_coding | 2105,123488 | -0,617620651 | 0,074641576 | -8,274485646 | 1,29E-16 | 2,16E-15 |
| ENSG00000158856.18 | DMTN | protein_coding | 13434,55223 | -0,385630147 | 0,068556536 | -5,624994607 | 1,86E-08 | 1,50E-07 |
| ENSG00000158864.12 | NDUFS2 | protein_coding | 9574,263004 | -0,196946634 | 0,045823769 | -4,297914383 | 1,72E-05 | 9,30E-05 |
| ENSG00000158882.15 | TOMM40L | protein_coding | 1994,294889 | 0,21881886 | 0,072715127 | 3,009261887 | 0,002618833 | 0,009187548 |
| ENSG00000158887.16 | MPZ | protein_coding | 167,4650699 | 0,579521091 | 0,186424356 | 3,108612534 | 0,00187968 | 0,006839594 |
| ENSG00000159055.4 | MIS18A | protein_coding | 2454,839839 | -0,573291304 | 0,062987371 | -9,101686434 | 8,89E-20 | 1,78E-18 |
| ENSG00000159063.13 | ALG8 | protein_coding | 3972,664887 | -0,170114556 | 0,053054516 | -3,206410457 | 0,001344021 | 0,005062345 |
| ENSG00000159082.17 | SYNJ1 | protein_coding | 4757,077335 | 0,461671619 | 0,051177756 | 9,020942946 | 1,86E-19 | 3,67E-18 |
| ENSG00000159110.19 | IFNAR2 | protein_coding | 3303,940519 | 0,258756841 | 0,065723099 | 3,937076056 | 8,25E-05 | 0,00039883 |
| ENSG00000159128.14 | IFNGR2 | protein_coding | 6264,466905 | 0,309748757 | 0,051234521 | 6,045704224 | 1,49E-09 | 1,36E-08 |
| ENSG00000159131.17 | GART | protein_coding | 12112,9644 | 0,136099764 | 0,045651263 | 2,981292428 | 0,002870346 | 0,009964314 |
| ENSG00000159140.20 | SON | protein_coding | 53914,38038 | 0,130543315 | 0,039284592 | 3,323015685 | 0,000890499 | 0,003490204 |
| ENSG00000159147.18 | DONSON | protein_coding | 6806,854165 | -0,327353424 | 0,053838273 | -6,080310666 | 1,20E-09 | 1,11E-08 |
| ENSG00000159164.10 | SV2A | protein_coding | 6262,97452 | 0,333052402 | 0,056392471 | 5,905972862 | 3,51E-09 | 3,08E-08 |
| ENSG00000159167.12 | STC1 | protein_coding | 746,1591198 | -0,385107485 | 0,145801047 | -2,641321814 | 0,008258323 | 0,025373547 |
| ENSG00000159199.14 | ATP5MC1 | protein_coding | 5414,829607 | -0,356275224 | 0,052286903 | -6,813852152 | 9,50E-12 | 1,09E-10 |
| ENSG00000159200.18 | RCAN1 | protein_coding | 2589,321062 | -0,321056304 | 0,063087723 | -5,089045704 | 3,60E-07 | 2,48E-06 |
| ENSG00000159202.18 | UBE2Z | protein_coding | 12633,23636 | 0,138460775 | 0,046743445 | 2,96214315 | 0,003055057 | 0,010530824 |

| | | | | | | | | |
|---|---|---|---|---|---|---|---|---|
| ENSG00000159208.16 | CIART | protein_coding | 333,8815283 | 0,376279898 | 0,137078242 | 2,745000898 | 0,006051073 | 0,019308705 |
| ENSG00000159214.13 | CCDC24 | protein_coding | 820,3824301 | 0,307628073 | 0,091783747 | 3,351661744 | 0,000803281 | 0,003183844 |
| ENSG00000159228.13 | CBR1 | protein_coding | 4816,2928 | 0,207356176 | 0,077584636 | 2,672644836 | 0,007525588 | 0,023367838 |
| ENSG00000159231.6 | CBR3 | protein_coding | 13,99249565 | 2,364304948 | 0,665098187 | 3,554820916 | 0,000378237 | 0,001610009 |
| ENSG00000159247.13 | TUBBP5 | transcribed_unproc | 389,1867347 | 0,654333707 | 0,132735661 | 4,929599932 | 8,24E-07 | 5,41E-06 |
| ENSG00000159248.5 | GJD2 | protein_coding | 2020,073106 | 0,324289758 | 0,073440897 | 4,415656289 | 1,01E-05 | 5,65E-05 |
| ENSG00000159256.13 | MORC3 | protein_coding | 3730,399471 | 0,288717185 | 0,05887805 | 4,903647216 | 9,41E-07 | 6,13E-06 |
| ENSG00000159259.8 | CHAF1B | protein_coding | 5782,003841 | -0,183264618 | 0,05198058 | -3,525636258 | 0,000422467 | 0,001779103 |
| ENSG00000159267.14 | HLCS | protein_coding | 4723,654818 | -0,390841816 | 0,056002882 | -6,978958972 | 2,97E-12 | 3,60E-11 |
| ENSG00000159314.11 | ARHGAP27 | protein_coding | 419,1670643 | 0,890799744 | 0,136879039 | 6,507933932 | 7,62E-11 | 8,02E-10 |
| ENSG00000159322.17 | ADPGK | protein_coding | 4331,491518 | 0,155874232 | 0,049998223 | 3,117595421 | 0,001823329 | 0,006659308 |
| ENSG00000159335.16 | PTMS | protein_coding | 6547,275267 | -0,232922074 | 0,053166145 | -4,381022404 | 1,18E-05 | 6,55E-05 |
| ENSG00000159346.13 | ADIPOR1 | protein_coding | 13057,05275 | -0,149786733 | 0,045239737 | -3,310955002 | 0,000929782 | 0,003630891 |
| ENSG00000159403.18 | C1R | protein_coding | 214,8684458 | 2,746312879 | 0,183700831 | 14,94992081 | 1,56E-50 | 9,05E-49 |
| ENSG00000159459.12 | UBR1 | protein_coding | 12975,14365 | 0,150651397 | 0,05227303 | 2,882010015 | 0,003951472 | 0,013261321 |
| ENSG00000159461.15 | AMFR | protein_coding | 8621,812078 | 0,154401697 | 0,047433924 | 3,255090132 | 0,001133564 | 0,004340828 |
| ENSG00000159479.17 | MED8 | protein_coding | 4804,702201 | 0,243582097 | 0,053871355 | 4,521551325 | 6,14E-06 | 3,55E-05 |
| ENSG00000159579.14 | RSPRY1 | protein_coding | 8931,85198 | 0,222189469 | 0,05660685 | 3,925133924 | 8,67E-05 | 0,000417619 |
| ENSG00000159592.11 | GPBP1L1 | protein_coding | 12255,89063 | 0,210592734 | 0,042737696 | 4,927564054 | 8,33E-07 | 5,46E-06 |
| ENSG00000159593.15 | NAE1 | protein_coding | 9403,413501 | -0,140993901 | 0,050605116 | -2,786159041 | 0,00533367 | 0,017271697 |
| ENSG00000159658.11 | EFCAB14 | protein_coding | 13274,80978 | 0,187896405 | 0,042445243 | 4,426795407 | 9,56E-06 | 5,38E-05 |
| ENSG00000159685.10 | CHCHD6 | protein_coding | 533,9160048 | -0,369575108 | 0,119519766 | -3,092167266 | 0,001987009 | 0,007185986 |
| ENSG00000159784.17 | FAM131B | protein_coding | 852,6961642 | -0,404998558 | 0,096059664 | -4,216114681 | 2,49E-05 | 0,000130899 |
| ENSG00000159788.19 | RGS12 | protein_coding | 2949,881762 | -0,235337963 | 0,069553936 | -3,383531951 | 0,000715599 | 0,00286864 |
| ENSG00000159792.10 | PSKH1 | protein_coding | 3281,084058 | -0,251132733 | 0,064936179 | -3,867377744 | 0,000110012 | 0,000519568 |
| ENSG00000159871.15 | LYPD5 | protein_coding | 440,8280417 | 0,417470363 | 0,140970017 | 2,961412447 | 0,003062315 | 0,010552612 |
| ENSG00000159873.10 | CCDC117 | protein_coding | 8822,551843 | 0,265199233 | 0,05276647 | 5,025904413 | 5,01E-07 | 3,40E-06 |
| ENSG00000159882.13 | ZNF230 | protein_coding | 999,720816 | 0,35207479 | 0,095594159 | 3,683015724 | 0,000230491 | 0,001023096 |
| ENSG00000159885.14 | ZNF222 | protein_coding | 899,1875884 | 0,338696113 | 0,090252456 | 3,752763388 | 0,000174896 | 0,000793575 |
| ENSG00000159905.14 | ZNF221 | protein_coding | 834,1066954 | 0,362036139 | 0,097892233 | 3,6983132 | 0,000217037 | 0,000967 |
| ENSG00000159915.12 | ZNF233 | protein_coding | 940,8715608 | 0,252608887 | 0,093765882 | 2,694038394 | 0,007059203 | 0,022130307 |
| ENSG00000159921.16 | GNE | protein_coding | 7390,75263 | -0,149812772 | 0,046487494 | -3,22264679 | 0,001270121 | 0,004803284 |
| ENSG00000160014.17 | CALM3 | protein_coding | 37708,10763 | 0,203751043 | 0,040967944 | 4,973426079 | 6,58E-07 | 4,39E-06 |
| ENSG00000160049.12 | DFFA | protein_coding | 5567,482312 | -0,506969271 | 0,05325217 | -9,520161771 | 1,73E-21 | 3,79E-20 |
| ENSG00000160058.18 | BSDC1 | protein_coding | 7518,85598 | 0,131895111 | 0,054335267 | 2,427430979 | 0,01520618 | 0,042918609 |
| ENSG00000160075.12 | SSU72 | protein_coding | 6681,322028 | -0,186778983 | 0,047077422 | -3,967485369 | 7,26E-05 | 0,000354574 |

| | | | | | | | | |
|---|---|---|---|---|---|---|---|---|
| ENSG00000160087.20 | UBE2J2 | protein_coding | 2937,345353 | -0,181201717 | 0,067324563 | -2,69146518 | 0,007113892 | 0,022270738 |
| ENSG00000160097.18 | FNDC5 | protein_coding | 1986,305651 | 0,324139917 | 0,079636885 | 4,070223452 | 4,70E-05 | 0,0002363 |
| ENSG00000160111.13 | CPAMD8 | protein_coding | 46,93494385 | -0,971127038 | 0,386003868 | -2,515847943 | 0,01187464 | 0,034667481 |
| ENSG00000160124.9 | CCDC58 | protein_coding | 1766,656968 | -0,182758195 | 0,065534061 | -2,788751243 | 0,005291169 | 0,017141468 |
| ENSG00000160131.13 | VMA21 | protein_coding | 5727,986247 | 0,147489614 | 0,048369492 | 3,049228087 | 0,002294302 | 0,008158365 |
| ENSG00000160182.3 | TFF1 | protein_coding | 750,6988111 | -0,403780855 | 0,102863439 | -3,925406923 | 8,66E-05 | 0,000417324 |
| ENSG00000160190.14 | SLC37A1 | protein_coding | 4960,757353 | 0,329365754 | 0,056798103 | 5,798886477 | 6,68E-09 | 5,69E-08 |
| ENSG00000160191.18 | PDE9A | protein_coding | 1002,098719 | 0,502970314 | 0,083637844 | 6,013669045 | 1,81E-09 | 1,65E-08 |
| ENSG00000160200.17 | CBS | protein_coding | 340,1732964 | 0,84016478 | 0,143112242 | 5,870670252 | 4,34E-09 | 3,78E-08 |
| ENSG00000160208.13 | RRP1B | protein_coding | 5989,285641 | -0,221786784 | 0,046507602 | -4,768828651 | 1,85E-06 | 1,16E-05 |
| ENSG00000160209.19 | PDXK | protein_coding | 4203,89345 | 0,662045689 | 0,06741804 | 9,820007982 | 9,23E-23 | 2,16E-21 |
| ENSG00000160211.18 | G6PD | protein_coding | 3014,868999 | -0,153361253 | 0,063447452 | -2,417138083 | 0,015643082 | 0,043947844 |
| ENSG00000160216.19 | AGPAT3 | protein_coding | 5478,492896 | 0,435124304 | 0,058573912 | 7,42863649 | 1,10E-13 | 1,50E-12 |
| ENSG00000160218.13 | TRAPPC10 | protein_coding | 6318,011711 | 0,324780845 | 0,067905124 | 4,78286213 | 1,73E-06 | 1,09E-05 |
| ENSG00000160282.14 | FTCD | protein_coding | 3266,484161 | -0,481944271 | 0,062740163 | -7,681591021 | 1,57E-14 | 2,27E-13 |
| ENSG00000160284.15 | SPATC1L | protein_coding | 1467,864234 | -0,357911979 | 0,102640581 | -3,487041649 | 0,000488395 | 0,002028657 |
| ENSG00000160294.11 | MCM3AP | protein_coding | 10797,59353 | 0,152655789 | 0,054630417 | 2,794336886 | 0,005200627 | 0,016904103 |
| ENSG00000160298.17 | C21orf58 | protein_coding | 2743,788123 | -0,265315752 | 0,060189278 | -4,408023511 | 1,04E-05 | 5,83E-05 |
| ENSG00000160299.17 | PCNT | protein_coding | 12744,03769 | -0,15509922 | 0,051636444 | -3,003677436 | 0,00266738 | 0,009328783 |
| ENSG00000160325.14 | CACFD1 | protein_coding | 12232,48342 | -0,280906579 | 0,048366934 | -5,807822739 | 6,33E-09 | 5,40E-08 |
| ENSG00000160336.15 | ZNF761 | protein_coding | 3797,929704 | 0,658192532 | 0,056642292 | 11,62016059 | 3,26E-31 | 1,09E-29 |
| ENSG00000160360.13 | GPSM1 | protein_coding | 3140,528777 | -0,561177166 | 0,066342622 | -8,458772788 | 2,70E-17 | 4,69E-16 |
| ENSG00000160401.15 | CFAP157 | protein_coding | 264,637092 | -2,048191756 | 0,190236078 | -10,76657898 | 4,95E-27 | 1,39E-25 |
| ENSG00000160410.15 | SHKBP1 | protein_coding | 3472,260953 | 0,212390841 | 0,063188537 | 3,361224206 | 0,000775978 | 0,003083773 |
| ENSG00000160446.19 | ZDHHC12 | protein_coding | 2452,155625 | -0,4116998 | 0,079226942 | -5,196462092 | 2,03E-07 | 1,45E-06 |
| ENSG00000160469.16 | BRSK1 | protein_coding | 347,4517216 | 0,37058139 | 0,128746926 | 2,87837077 | 0,00399735 | 0,013393325 |
| ENSG00000160551.11 | TAOK1 | protein_coding | 22226,87029 | -0,217751807 | 0,052397648 | -4,155755351 | 3,24E-05 | 0,000167341 |
| ENSG00000160563.14 | MED27 | protein_coding | 7049,648956 | -0,150067388 | 0,049102258 | -3,056221748 | 0,002241454 | 0,007990639 |
| ENSG00000160570.14 | DEDD2 | protein_coding | 2071,504858 | 0,288299053 | 0,071096713 | 4,055026452 | 5,01E-05 | 0,000251022 |
| ENSG00000160584.16 | SIK3 | protein_coding | 8514,84616 | 0,289842458 | 0,057512745 | 5,039621362 | 4,66E-07 | 3,18E-06 |
| ENSG00000160602.14 | AC010761.1 | protein_coding | 891,842178 | 0,534975469 | 0,092015053 | 5,813999476 | 6,10E-09 | 5,21E-08 |
| ENSG00000160613.12 | PCSK7 | protein_coding | 5430,826606 | 0,150848003 | 0,054530559 | 2,766302159 | 0,005669598 | 0,018241419 |
| ENSG00000160685.13 | ZBTB7B | protein_coding | 1926,094049 | 0,225475054 | 0,074488449 | 3,026980108 | 0,002470102 | 0,008709199 |
| ENSG00000160691.18 | SHC1 | protein_coding | 1802,419069 | 0,211906491 | 0,067480324 | 3,140270783 | 0,001687917 | 0,006212612 |
| ENSG00000160695.14 | VPS11 | protein_coding | 4590,471876 | 0,163591229 | 0,05428657 | 3,013475128 | 0,002582742 | 0,009069414 |
| ENSG00000160710.17 | ADAR | protein_coding | 73799,61049 | 2,254210393 | 0,049006121 | 45,99854736 | 0 | 0 |

| | | | | | | | | |
|---|---|---|---|---|---|---|---|---|
| ENSG00000160714.10 | UBE2Q1 | protein_coding | 12063,98524 | 0,131362729 | 0,048103608 | 2,730829041 | 0,006317523 | 0,020065067 |
| ENSG00000160752.14 | FDPS | protein_coding | 20778,80732 | -0,316584177 | 0,046017025 | -6,879718535 | 6,00E-12 | 7,04E-11 |
| ENSG00000160753.16 | RUSC1 | protein_coding | 5229,1367 | 0,594415487 | 0,04890496 | 12,15450301 | 5,43E-34 | 2,00E-32 |
| ENSG00000160767.21 | FAM189B | protein_coding | 2515,811544 | 0,271725004 | 0,065817927 | 4,12843458 | 3,65E-05 | 0,000186888 |
| ENSG00000160781.17 | PAQR6 | protein_coding | 180,345539 | -0,463054191 | 0,176945169 | -2,616936054 | 0,008872296 | 0,027023885 |
| ENSG00000160796.17 | NBEAL2 | protein_coding | 5543,179913 | -0,233547956 | 0,051880705 | -4,501634227 | 6,74E-06 | 3,87E-05 |
| ENSG00000160862.13 | AZGP1 | protein_coding | 41,95731275 | 0,947584663 | 0,383569028 | 2,470441025 | 0,013494657 | 0,038740091 |
| ENSG00000160908.15 | ZNF394 | protein_coding | 1801,930116 | 0,193860998 | 0,075918011 | 2,553557404 | 0,01066287 | 0,031609448 |
| ENSG00000160932.11 | LY6E | protein_coding | 9574,313648 | 8,37842703 | 0,21180942 | 39,55644193 | 0 | 0 |
| ENSG00000160948.14 | VPS28 | protein_coding | 4581,276231 | -0,139203942 | 0,051877495 | -2,683320414 | 0,007289511 | 0,022741457 |
| ENSG00000160949.17 | TONSL | protein_coding | 1425,029522 | -0,20090171 | 0,078375702 | -2,563316228 | 0,010367757 | 0,030860598 |
| ENSG00000160953.16 | PWWP3A | protein_coding | 4478,477723 | -0,283631797 | 0,055954918 | -5,068934229 | 4,00E-07 | 2,75E-06 |
| ENSG00000160957.13 | RECQL4 | protein_coding | 3449,491049 | -0,398324789 | 0,066819052 | -5,961245766 | 2,50E-09 | 2,23E-08 |
| ENSG00000160972.9 | PPP1R16A | protein_coding | 3035,960328 | -0,502326648 | 0,067743455 | -7,415131823 | 1,22E-13 | 1,66E-12 |
| ENSG00000160991.16 | ORAI2 | protein_coding | 6127,007856 | -0,457339083 | 0,067852607 | -6,740184429 | 1,58E-11 | 1,77E-10 |
| ENSG00000161010.15 | MRNIP | protein_coding | 4029,050318 | 0,461565627 | 0,069006581 | 6,688718984 | 2,25E-11 | 2,49E-10 |
| ENSG00000161011.20 | SQSTM1 | protein_coding | 8980,099406 | 0,380268677 | 0,046882896 | 8,111032214 | 5,02E-16 | 8,09E-15 |
| ENSG00000161013.17 | MGAT4B | protein_coding | 12509,83578 | -0,397545915 | 0,047226656 | -8,417828955 | 3,84E-17 | 6,60E-16 |
| ENSG00000161016.17 | RPL8 | protein_coding | 33980,50976 | -0,391354253 | 0,048982183 | -7,989726632 | 1,35E-15 | 2,11E-14 |
| ENSG00000161082.13 | CELF5 | protein_coding | 491,5428049 | -0,697746679 | 0,149592347 | -4,664320695 | 3,10E-06 | 1,87E-05 |
| ENSG00000161103.11 | AC008132.1 | transcribed_process | 146,7296964 | -1,018097346 | 0,219042116 | -4,64795247 | 3,35E-06 | 2,02E-05 |
| ENSG00000161133.16 | USP41 | protein_coding | 647,9325407 | 2,265452901 | 0,163968925 | 13,81635518 | 2,03E-43 | 1,01E-41 |
| ENSG00000161249.21 | DMKN | protein_coding | 424,4308208 | 0,28953254 | 0,118588424 | 2,441490751 | 0,014626763 | 0,041512269 |
| ENSG00000161270.19 | NPHS1 | protein_coding | 2314,058877 | -0,492891673 | 0,070427287 | -6,998589536 | 2,59E-12 | 3,15E-11 |
| ENSG00000161526.15 | SAP30BP | protein_coding | 5464,717576 | 0,25314834 | 0,055889033 | 4,529481516 | 5,91E-06 | 3,42E-05 |
| ENSG00000161533.12 | ACOX1 | protein_coding | 5277,278511 | -0,243568852 | 0,052981324 | -4,597258728 | 4,28E-06 | 2,53E-05 |
| ENSG00000161542.16 | PRPSAP1 | protein_coding | 3544,046597 | -0,127032143 | 0,053009564 | -2,396400436 | 0,016556991 | 0,046203981 |
| ENSG00000161547.16 | SRSF2 | protein_coding | 18169,76139 | -0,146453206 | 0,043431832 | -3,372024609 | 0,000746178 | 0,002978917 |
| ENSG00000161551.14 | ZNF577 | protein_coding | 535,0150658 | -0,365931225 | 0,119769974 | -3,055283507 | 0,002248479 | 0,008010606 |
| ENSG00000161558.11 | TMEM143 | protein_coding | 700,0616463 | -0,280817469 | 0,094290172 | -2,978226289 | 0,002899218 | 0,010049047 |
| ENSG00000161647.18 | MPP3 | protein_coding | 525,2632459 | -0,336690285 | 0,116803854 | -2,882527193 | 0,003944991 | 0,013244405 |
| ENSG00000161654.9 | LSM12 | protein_coding | 5780,532736 | 0,149378665 | 0,05537644 | 2,697512945 | 0,006985957 | 0,021931228 |
| ENSG00000161682.14 | FAM171A2 | protein_coding | 1661,097187 | -0,241549558 | 0,072729999 | -3,321181946 | 0,000896371 | 0,003510075 |
| ENSG00000161692.18 | DBF4B | protein_coding | 1301,864736 | 0,62008303 | 0,073358158 | 8,452816253 | 2,84E-17 | 4,93E-16 |
| ENSG00000161791.14 | FMNL3 | protein_coding | 857,1549219 | -0,260393443 | 0,093823637 | -2,775350138 | 0,005514231 | 0,017793666 |
| ENSG00000161800.13 | RACGAP1 | protein_coding | 14434,00695 | -0,466311675 | 0,042156282 | -11,06149918 | 1,93E-28 | 5,73E-27 |

| | | | | | | | | |
|---|---|---|---|---|---|---|---|---|
| ENSG00000161813.22 | LARP4 | protein_coding | 11856,02139 | 0,157129595 | 0,049774317 | 3,156840827 | 0,001594884 | 0,005910534 |
| ENSG00000161888.11 | SPC24 | protein_coding | 1521,760802 | -0,408544919 | 0,080702223 | -5,062375051 | 4,14E-07 | 2,84E-06 |
| ENSG00000161904.12 | LEMD2 | protein_coding | 6485,325819 | 0,196480751 | 0,052206324 | 3,763543131 | 0,000167523 | 0,000763005 |
| ENSG00000161921.15 | CXCL16 | protein_coding | 1580,545238 | 1,766675682 | 0,083961445 | 21,04151127 | 2,74E-98 | 3,14E-96 |
| ENSG00000161955.16 | TNFSF13 | protein_coding | 204,0438502 | 1,005789402 | 0,175063396 | 5,745286704 | 9,18E-09 | 7,65E-08 |
| ENSG00000161956.13 | SENP3 | protein_coding | 4538,27272 | -0,30228926 | 0,067941064 | -4,449286547 | 8,62E-06 | 4,87E-05 |
| ENSG00000161960.15 | EIF4A1 | protein_coding | 61970,60256 | -0,131354706 | 0,04374768 | -3,002552514 | 0,002677258 | 0,009360422 |
| ENSG00000161970.15 | RPL26 | protein_coding | 49294,2279 | -0,531811916 | 0,039599461 | -13,42977655 | 4,05E-41 | 1,87E-39 |
| ENSG00000161980.5 | POLR3K | protein_coding | 2939,448517 | -0,149259158 | 0,062051039 | -2,405425618 | 0,01615364 | 0,045235417 |
| ENSG00000161981.10 | SNRNP25 | protein_coding | 4282,873791 | -0,470934932 | 0,055443036 | -8,494032113 | 2,00E-17 | 3,50E-16 |
| ENSG00000161996.19 | WDR90 | protein_coding | 4394,661795 | -0,450979156 | 0,059954504 | -7,52202297 | 5,39E-14 | 7,56E-13 |
| ENSG00000162032.16 | SPSB3 | protein_coding | 1750,393855 | -0,345559507 | 0,065041687 | -5,312892769 | 1,08E-07 | 7,93E-07 |
| ENSG00000162062.15 | TEDC2 | protein_coding | 1843,205578 | -0,182981759 | 0,07048935 | -2,5958781 | 0,009434953 | 0,028502397 |
| ENSG00000162063.13 | CCNF | protein_coding | 6255,482958 | -0,594145639 | 0,05614178 | -10,5829498 | 3,58E-26 | 9,75E-25 |
| ENSG00000162065.14 | TBC1D24 | protein_coding | 9616,571794 | -0,177530665 | 0,047339049 | -3,750194986 | 0,000176697 | 0,00080074 |
| ENSG00000162068.1 | NTN3 | protein_coding | 670,5971148 | -0,631578359 | 0,115756079 | -5,456113966 | 4,87E-08 | 3,74E-07 |
| ENSG00000162069.15 | BICDL2 | protein_coding | 320,0486172 | 0,733483776 | 0,168462612 | 4,353985528 | 1,34E-05 | 7,34E-05 |
| ENSG00000162076.13 | FLYWCH2 | protein_coding | 1936,780862 | -0,277547603 | 0,077148595 | -3,597571708 | 0,000321202 | 0,001387437 |
| ENSG00000162086.15 | ZNF75A | protein_coding | 5874,447532 | 0,538121988 | 0,048801945 | 11,02665033 | 2,84E-28 | 8,37E-27 |
| ENSG00000162104.10 | ADCY9 | protein_coding | 4703,149889 | 0,212023973 | 0,059913967 | 3,538807123 | 0,000401939 | 0,001699653 |
| ENSG00000162144.9 | CYB561A3 | protein_coding | 4121,349084 | -0,31649241 | 0,065642503 | -4,821455544 | 1,43E-06 | 9,06E-06 |
| ENSG00000162174.12 | ASRGL1 | protein_coding | 6071,849212 | -0,216936836 | 0,050788317 | -4,271392507 | 1,94E-05 | 0,000103885 |
| ENSG00000162194.12 | LBHD1 | protein_coding | 762,6167463 | 0,424737314 | 0,090168363 | 4,710491592 | 2,47E-06 | 1,52E-05 |
| ENSG00000162236.11 | STX5 | protein_coding | 2282,695633 | 0,333136248 | 0,0631108 | 5,278593311 | 1,30E-07 | 9,49E-07 |
| ENSG00000162244.11 | RPL29 | protein_coding | 30671,801 | -0,390244843 | 0,044654778 | -8,739150931 | 2,35E-18 | 4,38E-17 |
| ENSG00000162298.18 | SYVN1 | protein_coding | 5234,658638 | 0,218498981 | 0,050514127 | 4,325502479 | 1,52E-05 | 8,28E-05 |
| ENSG00000162368.13 | CMPK1 | protein_coding | 19850,26901 | -0,11098635 | 0,045330547 | -2,448378779 | 0,014350071 | 0,040897157 |
| ENSG00000162377.6 | COA7 | protein_coding | 4617,740005 | 0,208984439 | 0,062606717 | 3,338051405 | 0,000843681 | 0,003326984 |
| ENSG00000162378.13 | ZYG11B | protein_coding | 7235,768761 | 0,125633563 | 0,052275633 | 2,403291086 | 0,016248248 | 0,045459773 |
| ENSG00000162384.14 | CZIB | protein_coding | 2678,667377 | -0,247166161 | 0,06376043 | -3,876482054 | 0,000105978 | 0,000501778 |
| ENSG00000162402.14 | USP24 | protein_coding | 17923,09629 | 0,247418832 | 0,04416918 | 5,601617051 | 2,12E-08 | 1,71E-07 |
| ENSG00000162407.9 | PLPP3 | protein_coding | 1820,924201 | 1,321754155 | 0,07604885 | 17,38033059 | 1,16E-67 | 9,06E-66 |
| ENSG00000162408.11 | NOL9 | protein_coding | 2580,192461 | -0,222064483 | 0,062905244 | -3,530142641 | 0,000415336 | 0,001752351 |
| ENSG00000162415.7 | ZSWIM5 | protein_coding | 4520,632534 | 0,194260971 | 0,0501526 | 3,873397817 | 0,000107328 | 0,000507746 |
| ENSG00000162419.12 | GMEB1 | protein_coding | 1503,479808 | -0,328165945 | 0,082037885 | -4,000175606 | 6,33E-05 | 0,000312096 |
| ENSG00000162430.17 | SELENON | protein_coding | 7003,36443 | -0,147662677 | 0,058237347 | -2,535532366 | 0,011227655 | 0,033035724 |

| | | | | | | | | |
|---|---|---|---|---|---|---|---|---|
| ENSG00000162433.15 | AK4 | protein_coding | 1687,602566 | -0,570188399 | 0,234259772 | -2,434000491 | 0,014932975 | 0,042258779 |
| ENSG00000162437.14 | RAVER2 | protein_coding | 5524,320349 | -0,177461018 | 0,050421003 | -3,519585211 | 0,000432222 | 0,001817804 |
| ENSG00000162438.11 | CTRC | protein_coding | 124,0231433 | -1,336072562 | 0,216733371 | -6,164590883 | 7,07E-10 | 6,71E-09 |
| ENSG00000162441.12 | LZIC | protein_coding | 7625,367447 | -0,333042743 | 0,05424146 | -6,140003292 | 8,25E-10 | 7,80E-09 |
| ENSG00000162496.9 | DHRS3 | protein_coding | 320,7719272 | 0,682756598 | 0,143099362 | 4,771206434 | 1,83E-06 | 1,14E-05 |
| ENSG00000162510.6 | MATN1 | protein_coding | 159,0283287 | -1,494371675 | 0,191753558 | -7,793188776 | 6,53E-15 | 9,79E-14 |
| ENSG00000162511.8 | LAPTM5 | protein_coding | 90,37109843 | -1,421785733 | 0,263341892 | -5,399010853 | 6,70E-08 | 5,05E-07 |
| ENSG00000162512.16 | SDC3 | protein_coding | 3434,431533 | -0,175868156 | 0,06088063 | -2,88873747 | 0,003867919 | 0,013008065 |
| ENSG00000162521.19 | RBBP4 | protein_coding | 53263,94625 | -0,115532722 | 0,042426443 | -2,723130043 | 0,00646666 | 0,020478059 |
| ENSG00000162542.13 | TMCO4 | protein_coding | 347,9731325 | 0,777629312 | 0,152656543 | 5,093979588 | 3,51E-07 | 2,43E-06 |
| ENSG00000162543.6 | UBXN10 | protein_coding | 914,8365405 | 0,264248716 | 0,087373065 | 3,024372748 | 0,002491493 | 0,008777744 |
| ENSG00000162545.6 | CAMK2N1 | protein_coding | 11886,59803 | -0,124716098 | 0,049905521 | -2,499044083 | 0,012452882 | 0,036149332 |
| ENSG00000162585.17 | FAAP20 | protein_coding | 1810,531501 | -0,210469449 | 0,079152641 | -2,659032558 | 0,007836539 | 0,024226433 |
| ENSG00000162601.11 | MYSM1 | protein_coding | 5868,507306 | 0,22642948 | 0,081555976 | 2,776368945 | 0,005496979 | 0,017744372 |
| ENSG00000162627.17 | SNX7 | protein_coding | 7032,517868 | 0,117365616 | 0,046768968 | 2,509476294 | 0,012091033 | 0,035217011 |
| ENSG00000162630.5 | B3GALT2 | protein_coding | 2449,887728 | -0,207355017 | 0,061027071 | -3,397754679 | 0,000679413 | 0,002735503 |
| ENSG00000162636.16 | FAM102B | protein_coding | 3880,555151 | -0,572580837 | 0,055622326 | -10,29408286 | 7,49E-25 | 1,94E-23 |
| ENSG00000162639.16 | HENMT1 | protein_coding | 3210,390379 | -0,400512159 | 0,058557312 | -6,839660966 | 7,94E-12 | 9,19E-11 |
| ENSG00000162642.14 | C1orf52 | protein_coding | 2455,44266 | 0,462197221 | 0,067291376 | 6,868595181 | 6,48E-12 | 7,60E-11 |
| ENSG00000162643.13 | WDR63 | protein_coding | 118,857469 | 0,767931034 | 0,282378767 | 2,719507002 | 0,006537931 | 0,020683386 |
| ENSG00000162645.13 | GBP2 | protein_coding | 153,8548445 | 4,247688039 | 0,294051583 | 14,44538403 | 2,68E-47 | 1,43E-45 |
| ENSG00000162654.9 | GBP4 | protein_coding | 168,7383585 | 4,181557128 | 0,283158143 | 14,76756797 | 2,37E-49 | 1,34E-47 |
| ENSG00000162687.17 | KCNT2 | protein_coding | 4745,140997 | 1,456370744 | 0,060989326 | 23,87910875 | 5,05E-126 | 7,15E-124 |
| ENSG00000162688.17 | AGL | protein_coding | 8423,659769 | -0,40066405 | 0,049749416 | -8,053643322 | 8,04E-16 | 1,27E-14 |
| ENSG00000162695.12 | SLC30A7 | protein_coding | 4067,953295 | 0,343547691 | 0,051017183 | 6,733960426 | 1,65E-11 | 1,85E-10 |
| ENSG00000162702.8 | ZNF281 | protein_coding | 4842,076097 | 0,185820224 | 0,053242215 | 3,490091911 | 0,000482854 | 0,002008607 |
| ENSG00000162706.13 | CADM3 | protein_coding | 612,9112289 | 0,55508364 | 0,110328522 | 5,031188963 | 4,87E-07 | 3,31E-06 |
| ENSG00000162714.12 | ZNF496 | protein_coding | 2621,58017 | 0,159513156 | 0,065079362 | 2,451055914 | 0,014243783 | 0,040635399 |
| ENSG00000162729.14 | IGSF8 | protein_coding | 2018,869771 | 0,615993503 | 0,076815961 | 8,019082145 | 1,07E-15 | 1,68E-14 |
| ENSG00000162734.12 | PEA15 | protein_coding | 7406,643087 | 0,275548763 | 0,046307831 | 5,950370771 | 2,68E-09 | 2,38E-08 |
| ENSG00000162735.18 | PEX19 | protein_coding | 7177,479292 | -0,134511808 | 0,056501659 | -2,380670068 | 0,017281182 | 0,047933719 |
| ENSG00000162736.17 | NCSTN | protein_coding | 14896,22565 | 0,370568581 | 0,05969843 | 6,207342127 | 5,39E-10 | 5,21E-09 |
| ENSG00000162755.14 | KLHDC9 | protein_coding | 258,5031428 | -0,535555939 | 0,15690567 | -3,413235092 | 0,000641965 | 0,002600324 |
| ENSG00000162769.13 | FLVCR1 | protein_coding | 16742,94561 | -0,148917667 | 0,054548018 | -2,730028912 | 0,006332877 | 0,020099933 |
| ENSG00000162772.17 | ATF3 | protein_coding | 459,770994 | 2,519686099 | 0,131481999 | 19,16373429 | 7,43E-82 | 7,00E-80 |
| ENSG00000162777.17 | DENND2D | protein_coding | 2542,243175 | 0,677801855 | 0,072022472 | 9,41097743 | 4,92E-21 | 1,06E-19 |

| | | | | | | | | |
|---|---|---|---|---|---|---|---|---|
| ENSG00000162804.14 | SNED1 | protein_coding | 3065,405974 | 0,206420294 | 0,066689082 | 3,095263678 | 0,00196638 | 0,007118245 |
| ENSG00000162817.7 | C1orf115 | protein_coding | 2517,009986 | 0,348025229 | 0,05932836 | 5,866085427 | 4,46E-09 | 3,87E-08 |
| ENSG00000162819.12 | BROX | protein_coding | 5720,352686 | -0,212947228 | 0,056221112 | -3,787673714 | 0,000152064 | 0,000699963 |
| ENSG00000162849.16 | KIF26B | protein_coding | 901,2728583 | -0,501431993 | 0,119271849 | -4,204110185 | 2,62E-05 | 0,000137174 |
| ENSG00000162852.14 | CNST | protein_coding | 4587,884686 | 0,14698459 | 0,055225192 | 2,661549657 | 0,007778187 | 0,024075798 |
| ENSG00000162869.16 | PPP1R21 | protein_coding | 3517,720505 | 0,237556128 | 0,082888716 | 2,865964615 | 0,004157407 | 0,013869731 |
| ENSG00000162878.13 | PKDCC | protein_coding | 7522,71056 | 0,154775037 | 0,046855085 | 3,303270836 | 0,00095564 | 0,003723474 |
| ENSG00000162882.15 | HAAO | protein_coding | 2769,360585 | -0,222099759 | 0,072609646 | -3,05881891 | 0,002222114 | 0,007927971 |
| ENSG00000162889.10 | MAPKAPK2 | protein_coding | 9034,10588 | 0,264306739 | 0,046816396 | 5,645602045 | 1,65E-08 | 1,34E-07 |
| ENSG00000162909.18 | CAPN2 | protein_coding | 7410,591418 | -0,204064701 | 0,049484263 | -4,123830289 | 3,73E-05 | 0,000190449 |
| ENSG00000162923.16 | WDR26 | protein_coding | 12400,07928 | 0,124717713 | 0,047970499 | 2,599883567 | 0,00932554 | 0,028213517 |
| ENSG00000162924.14 | REL | protein_coding | 5206,058593 | 0,461881287 | 0,059828891 | 7,720037542 | 1,16E-14 | 1,70E-13 |
| ENSG00000162929.13 | KIAA1841 | protein_coding | 3906,534169 | -0,184318093 | 0,060096428 | -3,06703906 | 0,002161906 | 0,007735232 |
| ENSG00000162946.22 | DISC1 | protein_coding | 239,3343635 | 0,632814584 | 0,162431036 | 3,895896982 | 9,78E-05 | 0,000466072 |
| ENSG00000162961.14 | DPY30 | protein_coding | 8060,515002 | -0,206520915 | 0,052691046 | -3,919468895 | 8,87E-05 | 0,000426225 |
| ENSG00000162971.11 | TYW5 | protein_coding | 2275,170712 | 0,294776635 | 0,064085863 | 4,599713918 | 4,23E-06 | 2,50E-05 |
| ENSG00000162972.10 | MAIP1 | protein_coding | 2235,827414 | 0,183334373 | 0,067262969 | 2,725636028 | 0,006417773 | 0,020340417 |
| ENSG00000162981.14 | LRATD1 | protein_coding | 16401,09175 | 0,36705856 | 0,043145709 | 8,507417414 | 1,78E-17 | 3,13E-16 |
| ENSG00000162992.3 | NEUROD1 | protein_coding | 22021,22234 | 0,158153599 | 0,039530297 | 4,000819847 | 6,31E-05 | 0,000311384 |
| ENSG00000162997.15 | PRORSD1P | transcribed_unitary | 191,8079888 | 0,490179312 | 0,184204472 | 2,661060872 | 0,007789487 | 0,024104148 |
| ENSG00000163001.11 | CFAP36 | protein_coding | 6035,654439 | -0,228972458 | 0,04887519 | -4,684840247 | 2,80E-06 | 1,71E-05 |
| ENSG00000163002.13 | NUP35 | protein_coding | 2686,380718 | -0,175378687 | 0,058346869 | -3,005794293 | 0,002648881 | 0,009272732 |
| ENSG00000163006.12 | CCDC138 | protein_coding | 2133,379221 | -0,339041307 | 0,070101985 | -4,836400947 | 1,32E-06 | 8,44E-06 |
| ENSG00000163016.10 | ALMS1P1 | transcribed_unproc | 425,0873935 | 3,006851025 | 0,152257705 | 19,74843265 | 8,27E-87 | 8,54E-85 |
| ENSG00000163026.12 | WDCP | protein_coding | 3129,850261 | 0,202219748 | 0,059079648 | 3,42283267 | 0,000619722 | 0,002516744 |
| ENSG00000163040.14 | CCDC74A | protein_coding | 526,45475 | -0,629260085 | 0,121576183 | -5,175850003 | 2,27E-07 | 1,61E-06 |
| ENSG00000163041.11 | H3F3A | protein_coding | 60669,71079 | -0,372993198 | 0,040754723 | -9,152146582 | 5,58E-20 | 1,13E-18 |
| ENSG00000163050.17 | COQ8A | protein_coding | 3194,176454 | -0,283995021 | 0,056230289 | -5,050570155 | 4,40E-07 | 3,01E-06 |
| ENSG00000163071.11 | SPATA18 | protein_coding | 56,35460351 | 1,421153434 | 0,390147003 | 3,642610154 | 0,000269887 | 0,001182346 |
| ENSG00000163082.10 | SGPP2 | protein_coding | 4271,868683 | 0,399756053 | 0,052898837 | 7,556991269 | 4,12E-14 | 5,82E-13 |
| ENSG00000163083.6 | INHBB | protein_coding | 137,11466 | -1,059142869 | 0,21885177 | -4,839544454 | 1,30E-06 | 8,32E-06 |
| ENSG00000163104.17 | SMARCAD1 | protein_coding | 15178,65381 | -0,192313755 | 0,044791092 | -4,293571482 | 1,76E-05 | 9,46E-05 |
| ENSG00000163110.15 | PDLIM5 | protein_coding | 10963,66962 | 0,23241596 | 0,043699298 | 5,318528492 | 1,05E-07 | 7,69E-07 |
| ENSG00000163121.10 | NEURL3 | protein_coding | 242,2927391 | 3,825545785 | 0,201205489 | 19,01312833 | 1,33E-80 | 1,24E-78 |
| ENSG00000163131.11 | CTSS | protein_coding | 560,7737707 | 4,680528436 | 0,400587583 | 11,68415755 | 1,54E-31 | 5,25E-30 |
| ENSG00000163132.8 | MSX1 | protein_coding | 287,9987249 | 0,868840322 | 0,160355535 | 5,418212251 | 6,02E-08 | 4,57E-07 |

| | | | | | | | | |
|---|---|---|---|---|---|---|---|---|
| ENSG00000163138.19 | PACRGL | protein_coding | 1981,577156 | 0,176805667 | 0,072250929 | 2,447105789 | 0,014400856 | 0,040986658 |
| ENSG00000163156.11 | SCNM1 | protein_coding | 1847,218353 | 0,242866322 | 0,063837917 | 3,804421141 | 0,000142136 | 0,000657625 |
| ENSG00000163162.8 | RNF149 | protein_coding | 3684,158718 | 0,341560647 | 0,056996284 | 5,992682714 | 2,06E-09 | 1,86E-08 |
| ENSG00000163171.7 | CDC42EP3 | protein_coding | 6431,625241 | 0,171692205 | 0,055363791 | 3,101164167 | 0,001927614 | 0,006991405 |
| ENSG00000163191.6 | S100A11 | protein_coding | 5360,24953 | 0,292082587 | 0,049670526 | 5,880400511 | 4,09E-09 | 3,57E-08 |
| ENSG00000163214.21 | DHX57 | protein_coding | 3993,274932 | -0,143625311 | 0,053877018 | -2,665799165 | 0,007680556 | 0,023813086 |
| ENSG00000163219.11 | ARHGAP25 | protein_coding | 480,6014402 | 0,425657899 | 0,11392603 | 3,736265533 | 0,000186773 | 0,000842502 |
| ENSG00000163249.12 | CCNYL1 | protein_coding | 4010,822778 | 0,56459028 | 0,066891186 | 8,440428654 | 3,16E-17 | 5,46E-16 |
| ENSG00000163291.14 | PAQR3 | protein_coding | 1794,147337 | -0,227532411 | 0,065103477 | -3,494934871 | 0,000474177 | 0,00197612 |
| ENSG00000163312.11 | HELQ | protein_coding | 1857,659011 | 0,49053908 | 0,084507821 | 5,804658925 | 6,45E-09 | 5,50E-08 |
| ENSG00000163322.14 | ABRAXAS1 | protein_coding | 2270,343587 | 0,446185476 | 0,06346028 | 7,030940847 | 2,05E-12 | 2,52E-11 |
| ENSG00000163328.13 | GPR155 | protein_coding | 873,6664309 | -0,634705402 | 0,089999953 | -7,05228592 | 1,76E-12 | 2,18E-11 |
| ENSG00000163331.11 | DAPL1 | protein_coding | 298,6994504 | -0,507493573 | 0,144119546 | -3,521337579 | 0,000429376 | 0,00180617 |
| ENSG00000163344.6 | PMVK | protein_coding | 4062,153913 | -0,257946508 | 0,05489331 | -4,699051838 | 2,61E-06 | 1,60E-05 |
| ENSG00000163374.19 | YY1AP1 | protein_coding | 7621,219889 | 0,290027993 | 0,046284598 | 6,266188054 | 3,70E-10 | 3,64E-09 |
| ENSG00000163376.11 | KBTBD8 | protein_coding | 4157,647765 | 0,404891965 | 0,070381625 | 5,752807864 | 8,78E-09 | 7,34E-08 |
| ENSG00000163377.16 | TAFA4 | protein_coding | 2161,106894 | -0,244457065 | 0,064588142 | -3,784859835 | 0,000153795 | 0,000706774 |
| ENSG00000163412.13 | EIF4E3 | protein_coding | 3519,225095 | -0,347494829 | 0,058886136 | -5,901131406 | 3,61E-09 | 3,16E-08 |
| ENSG00000163428.4 | LRRC58 | protein_coding | 14857,98248 | -0,244664171 | 0,052207273 | -4,686400099 | 2,78E-06 | 1,69E-05 |
| ENSG00000163435.16 | ELF3 | protein_coding | 4402,663268 | 1,058734401 | 0,08171749 | 12,95603182 | 2,17E-38 | 9,28E-37 |
| ENSG00000163462.18 | TRIM46 | protein_coding | 874,6779275 | 0,496888295 | 0,08983259 | 5,531269868 | 3,18E-08 | 2,50E-07 |
| ENSG00000163463.11 | KRTCAP2 | protein_coding | 5478,896744 | 0,33186822 | 0,049894183 | 6,651441078 | 2,90E-11 | 3,18E-10 |
| ENSG00000163479.14 | SSR2 | protein_coding | 25494,46697 | 0,156887583 | 0,051592474 | 3,040900528 | 0,002358717 | 0,008357055 |
| ENSG00000163481.8 | RNF25 | protein_coding | 2024,185256 | 0,183474441 | 0,066294292 | 2,767575263 | 0,0056475 | 0,018176427 |
| ENSG00000163492.15 | CCDC141 | protein_coding | 505,8411169 | 0,700222228 | 0,115468667 | 6,064175233 | 1,33E-09 | 1,22E-08 |
| ENSG00000163507.14 | CIP2A | protein_coding | 5312,005553 | -0,301921088 | 0,052460697 | -5,75518639 | 8,65E-09 | 7,25E-08 |
| ENSG00000163512.14 | AZI2 | protein_coding | 8415,163572 | 1,300989389 | 0,054033427 | 24,07749174 | 4,30E-128 | 6,21E-126 |
| ENSG00000163517.15 | HDAC11 | protein_coding | 1574,774289 | 0,257315534 | 0,075639407 | 3,401871374 | 0,000669261 | 0,00269923 |
| ENSG00000163521.16 | GLB1L | protein_coding | 849,4424441 | -0,354491505 | 0,096181119 | -3,685666257 | 0,000228105 | 0,001013305 |
| ENSG00000163527.9 | STT3B | protein_coding | 36349,04442 | 0,106984997 | 0,041917378 | 2,552282668 | 0,010701965 | 0,03170864 |
| ENSG00000163531.15 | NFASC | protein_coding | 10437,39238 | 0,47287464 | 0,05217708 | 9,062880493 | 1,27E-19 | 2,52E-18 |
| ENSG00000163535.18 | SGO2 | protein_coding | 5254,082552 | -0,60631885 | 0,055543048 | -10,91619702 | 9,65E-28 | 2,78E-26 |
| ENSG00000163536.12 | SERPINI1 | protein_coding | 48696,86959 | 0,544845961 | 0,038669967 | 14,08964134 | 4,40E-45 | 2,27E-43 |
| ENSG00000163558.13 | PRKCI | protein_coding | 6769,571125 | -0,25229161 | 0,054366967 | -4,64053125 | 3,48E-06 | 2,08E-05 |
| ENSG00000163577.8 | EIF5A2 | protein_coding | 3181,318205 | 0,378453187 | 0,064430789 | 5,873794047 | 4,26E-09 | 3,71E-08 |
| ENSG00000163584.18 | RPL22L1 | protein_coding | 2047,627113 | 0,211164188 | 0,078871294 | 2,677326287 | 0,007421232 | 0,023091584 |

| | | | | | | | | |
|---|---|---|---|---|---|---|---|---|
| ENSG00000163590.14 | PPM1L | protein_coding | 10216,08678 | -0,14514829 | 0,045225587 | -3,209428567 | 0,001329991 | 0,005017055 |
| ENSG00000163597.15 | SNHG16 | lncRNA | 3942,123374 | -0,21153402 | 0,060665475 | -3,486893022 | 0,000488667 | 0,00202941 |
| ENSG00000163605.14 | PPP4R2 | protein_coding | 19299,57575 | 0,216701395 | 0,050940728 | 4,253990923 | 2,10E-05 | 0,00011177 |
| ENSG00000163618.18 | CADPS | protein_coding | 41361,79841 | -0,389617236 | 0,0527931 | -7,380078799 | 1,58E-13 | 2,14E-12 |
| ENSG00000163624.6 | CDS1 | protein_coding | 7224,253163 | 0,264135281 | 0,047733794 | 5,533506942 | 3,14E-08 | 2,47E-07 |
| ENSG00000163625.15 | WDFY3 | protein_coding | 14997,5957 | 0,391669448 | 0,05676272 | 6,900117674 | 5,20E-12 | 6,14E-11 |
| ENSG00000163626.16 | COX18 | protein_coding | 2155,237836 | -0,386121918 | 0,065119615 | -5,929425699 | 3,04E-09 | 2,69E-08 |
| ENSG00000163629.13 | PTPN13 | protein_coding | 3178,667581 | 0,363491474 | 0,058535502 | 6,209760979 | 5,31E-10 | 5,14E-09 |
| ENSG00000163630.11 | SYNPR | protein_coding | 135,5889887 | -0,763138643 | 0,258394643 | -2,953384147 | 0,003143106 | 0,010794654 |
| ENSG00000163634.12 | THOC7 | protein_coding | 5709,769854 | -0,174643825 | 0,05098248 | -3,425565485 | 0,000613521 | 0,002494084 |
| ENSG00000163635.18 | ATXN7 | protein_coding | 2496,173362 | 0,786714008 | 0,095913538 | 8,202324998 | 2,36E-16 | 3,87E-15 |
| ENSG00000163636.10 | PSMD6 | protein_coding | 9521,231564 | -0,166505647 | 0,047643654 | -3,49481273 | 0,000474394 | 0,001976336 |
| ENSG00000163638.13 | ADAMTS9 | protein_coding | 163,606334 | 0,708126164 | 0,196558208 | 3,602628315 | 0,000315016 | 0,001363857 |
| ENSG00000163644.15 | PPM1K | protein_coding | 13904,2454 | 2,826999072 | 0,049694831 | 56,88718539 | 0 | 0 |
| ENSG00000163659.13 | TIPARP | protein_coding | 7196,571716 | 0,556138332 | 0,056904673 | 9,773157498 | 1,47E-22 | 3,41E-21 |
| ENSG00000163660.11 | CCNL1 | protein_coding | 7522,090272 | 0,237639206 | 0,048065053 | 4,944116213 | 7,65E-07 | 5,05E-06 |
| ENSG00000163673.7 | DCLK3 | protein_coding | 166,1574225 | -1,16107389 | 0,210543357 | -5,514654592 | 3,49E-08 | 2,73E-07 |
| ENSG00000163682.16 | RPL9 | protein_coding | 38287,92679 | -0,718090895 | 0,050864199 | -14,11780609 | 2,95E-45 | 1,53E-43 |
| ENSG00000163683.12 | SMIM14 | protein_coding | 2777,418992 | 0,348315588 | 0,093571565 | 3,722451224 | 0,000197298 | 0,000886247 |
| ENSG00000163686.14 | ABHD6 | protein_coding | 1318,414046 | 0,220964596 | 0,08062017 | 2,74081036 | 0,006128787 | 0,019529001 |
| ENSG00000163689.20 | C3orf67 | protein_coding | 1688,330496 | -0,473576989 | 0,088655787 | -5,341749294 | 9,21E-08 | 6,81E-07 |
| ENSG00000163694.15 | RBM47 | protein_coding | 11515,29688 | 0,120271118 | 0,046956207 | 2,561346539 | 0,010426729 | 0,030995147 |
| ENSG00000163702.20 | IL17RC | protein_coding | 753,1102372 | 1,797701397 | 0,11308806 | 15,89647399 | 6,70E-57 | 4,46E-55 |
| ENSG00000163703.17 | CRELD1 | protein_coding | 5011,080593 | 0,287516968 | 0,060159118 | 4,77927498 | 1,76E-06 | 1,10E-05 |
| ENSG00000163728.11 | TTC14 | protein_coding | 5171,774061 | -0,136478396 | 0,053587911 | -2,546813169 | 0,010871162 | 0,032133812 |
| ENSG00000163734.4 | CXCL3 | protein_coding | 138,5606994 | 2,779226869 | 0,274103914 | 10,13931844 | 3,70E-24 | 9,25E-23 |
| ENSG00000163735.7 | CXCL5 | protein_coding | 49,16494421 | 2,555081996 | 0,376796861 | 6,781059666 | 1,19E-11 | 1,36E-10 |
| ENSG00000163738.18 | MTHFD2L | protein_coding | 1671,599621 | -0,1657703 | 0,0680107 | -2,43741499 | 0,014792692 | 0,041919766 |
| ENSG00000163739.4 | CXCL1 | protein_coding | 1620,362495 | 3,850622102 | 0,088927156 | 43,30085748 | 0 | 0 |
| ENSG00000163743.13 | RCHY1 | protein_coding | 4144,429197 | 0,29248325 | 0,065191329 | 4,48653608 | 7,24E-06 | 4,14E-05 |
| ENSG00000163754.17 | GYG1 | protein_coding | 3057,336125 | 0,387421523 | 0,060878744 | 6,363822547 | 1,97E-10 | 1,99E-09 |
| ENSG00000163755.8 | HPS3 | protein_coding | 5287,256148 | 0,227153411 | 0,049554491 | 4,583911727 | 4,56E-06 | 2,69E-05 |
| ENSG00000163785.12 | RYK | protein_coding | 3902,729293 | 0,150535234 | 0,054448705 | 2,764716505 | 0,005697229 | 0,018320173 |
| ENSG00000163788.14 | SNRK | protein_coding | 6281,180088 | 0,49605065 | 0,053727736 | 9,232673623 | 2,64E-20 | 5,45E-19 |
| ENSG00000163795.14 | ZNF513 | protein_coding | 2552,627962 | 0,235953962 | 0,057380284 | 4,112108649 | 3,92E-05 | 0,000199975 |
| ENSG00000163807.6 | KIAA1143 | protein_coding | 8552,558485 | -0,337272075 | 0,048305591 | -6,982050539 | 2,91E-12 | 3,52E-11 |

| | | | | | | | | |
|---|---|---|---|---|---|---|---|---|
| ENSG00000163808.17 | KIF15 | protein_coding | 5955,934826 | -0,487201316 | 0,050061337 | -9,732087483 | 2,20E-22 | 5,06E-21 |
| ENSG00000163812.15 | ZDHHC3 | protein_coding | 8017,13844 | 0,203985543 | 0,048115988 | 4,239454488 | 2,24E-05 | 0,000118809 |
| ENSG00000163814.8 | CDCP1 | protein_coding | 7425,983362 | 0,49780999 | 0,044931244 | 11,07937241 | 1,58E-28 | 4,71E-27 |
| ENSG00000163820.15 | FYCO1 | protein_coding | 3066,050352 | 0,688535811 | 0,089904583 | 7,658517396 | 1,88E-14 | 2,71E-13 |
| ENSG00000163832.16 | ELP6 | protein_coding | 3901,923582 | -0,262348747 | 0,06134841 | -4,276374004 | 1,90E-05 | 0,000101782 |
| ENSG00000163840.10 | DTX3L | protein_coding | 35262,86893 | 4,6293092 | 0,157037627 | 29,47898082 | 5,35E-191 | 1,06E-188 |
| ENSG00000163870.15 | TPRA1 | protein_coding | 1820,606125 | 0,232927109 | 0,070627433 | 3,297969352 | 0,000973868 | 0,003786628 |
| ENSG00000163872.16 | YEATS2 | protein_coding | 17014,10512 | 0,717914927 | 0,048686666 | 14,74561694 | 3,28E-49 | 1,84E-47 |
| ENSG00000163873.10 | GRIK3 | protein_coding | 6657,793984 | -0,180889541 | 0,071265638 | -2,538243486 | 0,011141044 | 0,032819495 |
| ENSG00000163874.11 | ZC3H12A | protein_coding | 320,391452 | 1,33832762 | 0,174014979 | 7,690875985 | 1,46E-14 | 2,12E-13 |
| ENSG00000163875.15 | MEAF6 | protein_coding | 16162,24504 | -0,162304675 | 0,041580322 | -3,903401083 | 9,49E-05 | 0,000453095 |
| ENSG00000163879.11 | DNALI1 | protein_coding | 7515,284921 | 0,253627276 | 0,048563405 | 5,222600769 | 1,76E-07 | 1,27E-06 |
| ENSG00000163882.9 | POLR2H | protein_coding | 5536,706233 | -0,150285718 | 0,049981432 | -3,006830968 | 0,002639865 | 0,009244045 |
| ENSG00000163902.12 | RPN1 | protein_coding | 38002,10861 | 0,242661635 | 0,047252355 | 5,13544006 | 2,81E-07 | 1,97E-06 |
| ENSG00000163909.8 | HEYL | protein_coding | 612,5805457 | 0,482177104 | 0,112172124 | 4,298546626 | 1,72E-05 | 9,28E-05 |
| ENSG00000163913.11 | IFT122 | protein_coding | 3298,775723 | -0,3488272 | 0,056284736 | -6,197545305 | 5,74E-10 | 5,53E-09 |
| ENSG00000163918.10 | RFC4 | protein_coding | 8875,559619 | -0,265066891 | 0,045630312 | -5,809008975 | 6,28E-09 | 5,37E-08 |
| ENSG00000163932.15 | PRKCD | protein_coding | 4181,492665 | 0,150310996 | 0,056239376 | 2,67270027 | 0,007524345 | 0,023367201 |
| ENSG00000163939.18 | PBRM1 | protein_coding | 21297,83579 | -0,394467609 | 0,042415443 | -9,300094088 | 1,40E-20 | 2,96E-19 |
| ENSG00000163945.16 | UVSSA | protein_coding | 2027,971548 | -0,370905676 | 0,098127777 | -3,779823475 | 0,00015694 | 0,000718875 |
| ENSG00000163946.13 | TASOR | protein_coding | 15928,52306 | 0,12059199 | 0,044389297 | 2,716690664 | 0,00659382 | 0,020827996 |
| ENSG00000163947.11 | ARHGEF3 | protein_coding | 2118,257413 | 0,353999811 | 0,060599626 | 5,841617106 | 5,17E-09 | 4,46E-08 |
| ENSG00000163956.13 | LRPAP1 | protein_coding | 16744,8576 | 0,511403342 | 0,065294982 | 7,832199742 | 4,79E-15 | 7,24E-14 |
| ENSG00000163960.12 | UBXN7 | protein_coding | 12155,76701 | -0,343429382 | 0,046563743 | -7,37546766 | 1,64E-13 | 2,21E-12 |
| ENSG00000163964.15 | PIGX | protein_coding | 7759,580848 | -0,263405482 | 0,045670459 | -5,767524356 | 8,04E-09 | 6,76E-08 |
| ENSG00000164010.15 | ERMAP | protein_coding | 1874,664083 | -0,285064761 | 0,071133538 | -4,007459331 | 6,14E-05 | 0,00030356 |
| ENSG00000164023.14 | SGMS2 | protein_coding | 7274,335084 | 0,627072984 | 0,058548394 | 10,71033613 | 9,10E-27 | 2,54E-25 |
| ENSG00000164031.17 | DNAJB14 | protein_coding | 7028,763334 | 0,216853554 | 0,04988801 | 4,346807069 | 1,38E-05 | 7,57E-05 |
| ENSG00000164032.12 | H2AFZ | protein_coding | 40152,66985 | -0,276480105 | 0,040086425 | -6,897100653 | 5,31E-12 | 6,26E-11 |
| ENSG00000164039.15 | BDH2 | protein_coding | 1405,02476 | -0,60393695 | 0,07390972 | -8,17127914 | 3,05E-16 | 4,97E-15 |
| ENSG00000164040.17 | PGRMC2 | protein_coding | 22057,69742 | 0,257970049 | 0,041633123 | 6,196269431 | 5,78E-10 | 5,57E-09 |
| ENSG00000164045.11 | CDC25A | protein_coding | 5685,997178 | -0,233818491 | 0,04731473 | -4,941769475 | 7,74E-07 | 5,10E-06 |
| ENSG00000164048.14 | ZNF589 | protein_coding | 2754,530787 | 0,340338424 | 0,058731498 | 5,794819434 | 6,84E-09 | 5,81E-08 |
| ENSG00000164050.13 | PLXNB1 | protein_coding | 14407,0137 | 0,160343322 | 0,066268142 | 2,419613986 | 0,01553699 | 0,043709801 |
| ENSG00000164054.15 | SHISA5 | protein_coding | 26572,23255 | 1,432212702 | 0,049735383 | 28,79665567 | 2,36E-182 | 4,36E-180 |
| ENSG00000164056.11 | SPRY1 | protein_coding | 285,8825845 | 0,435777622 | 0,150096066 | 2,903324741 | 0,003692236 | 0,012480632 |

| | | | | | | | | |
|---|---|---|---|---|---|---|---|---|
| ENSG00000164061.5 | BSN | protein_coding | 6174,294502 | 0,299599964 | 0,056099697 | 5,340491648 | 9,27E-08 | 6,85E-07 |
| ENSG00000164062.13 | APEH | protein_coding | 9094,235639 | -0,138766414 | 0,057875989 | -2,397650828 | 0,016500586 | 0,046069424 |
| ENSG00000164073.10 | MFSD8 | protein_coding | 5291,115916 | 0,378174644 | 0,058480432 | 6,466686892 | 1,00E-10 | 1,04E-09 |
| ENSG00000164074.15 | ABHD18 | protein_coding | 3240,595379 | 0,266451914 | 0,055053789 | 4,839847013 | 1,30E-06 | 8,31E-06 |
| ENSG00000164076.17 | CAMKV | protein_coding | 5086,666732 | -0,767077352 | 0,049091142 | -15,62557556 | 4,87E-55 | 3,11E-53 |
| ENSG00000164078.13 | MST1R | protein_coding | 314,4516152 | 0,755910154 | 0,173904844 | 4,346688328 | 1,38E-05 | 7,57E-05 |
| ENSG00000164081.12 | TEX264 | protein_coding | 3484,128013 | 0,200072302 | 0,057465252 | 3,48162229 | 0,000498386 | 0,002066343 |
| ENSG00000164086.10 | DUSP7 | protein_coding | 1081,000621 | -0,446095175 | 0,081667698 | -5,462320891 | 4,70E-08 | 3,61E-07 |
| ENSG00000164087.7 | POC1A | protein_coding | 1409,13541 | -0,483322285 | 0,077241515 | -6,257286462 | 3,92E-10 | 3,84E-09 |
| ENSG00000164096.13 | C4orf3 | protein_coding | 10688,26393 | 0,196473943 | 0,049837604 | 3,942283047 | 8,07E-05 | 0,000390687 |
| ENSG00000164099.3 | PRSS12 | protein_coding | 3354,704888 | -0,173539844 | 0,063225573 | -2,744772994 | 0,006055277 | 0,01931938 |
| ENSG00000164104.11 | HMGB2 | protein_coding | 20379,35086 | -0,441596868 | 0,045230845 | -9,7631797 | 1,62E-22 | 3,75E-21 |
| ENSG00000164109.14 | MAD2L1 | protein_coding | 8579,179067 | -0,614519836 | 0,047058525 | -13,05862948 | 5,67E-39 | 2,47E-37 |
| ENSG00000164111.15 | ANXA5 | protein_coding | 23450,58338 | -0,140419264 | 0,04748222 | -2,957302002 | 0,00310344 | 0,010676346 |
| ENSG00000164114.19 | MAP9 | protein_coding | 8461,065983 | -0,165016523 | 0,044582069 | -3,701410199 | 0,000214405 | 0,000955839 |
| ENSG00000164116.16 | GUCY1A1 | protein_coding | 286,2843893 | 0,420038676 | 0,163398343 | 2,570642201 | 0,010151014 | 0,030307624 |
| ENSG00000164120.14 | HPGD | protein_coding | 19583,95761 | -0,240703931 | 0,055298967 | -4,352774441 | 1,34E-05 | 7,38E-05 |
| ENSG00000164124.10 | TMEM144 | protein_coding | 1336,613279 | 0,334599715 | 0,090005031 | 3,717566819 | 0,000201151 | 0,000901394 |
| ENSG00000164136.17 | IL15 | protein_coding | 330,5868005 | 2,135160717 | 0,185980445 | 11,4805657 | 1,65E-30 | 5,37E-29 |
| ENSG00000164163.11 | ABCE1 | protein_coding | 13475,54095 | -0,119045672 | 0,047325102 | -2,515486832 | 0,011886812 | 0,034694015 |
| ENSG00000164164.16 | OTUD4 | protein_coding | 7889,940405 | 0,710555683 | 0,060246049 | 11,79422869 | 4,18E-32 | 1,48E-30 |
| ENSG00000164172.19 | MOCS2 | protein_coding | 7129,740997 | -0,298169365 | 0,053513765 | -5,571825536 | 2,52E-08 | 2,01E-07 |
| ENSG00000164175.15 | SLC45A2 | protein_coding | 281,1907894 | 0,651322091 | 0,155585822 | 4,186256059 | 2,84E-05 | 0,000147593 |
| ENSG00000164176.13 | EDIL3 | protein_coding | 12704,45783 | -0,388650089 | 0,045790572 | -8,487557003 | 2,11E-17 | 3,69E-16 |
| ENSG00000164181.14 | ELOVL7 | protein_coding | 2236,39886 | 0,563092951 | 0,066782402 | 8,431756437 | 3,41E-17 | 5,87E-16 |
| ENSG00000164182.11 | NDUFAF2 | protein_coding | 1038,88274 | -0,231561727 | 0,082055894 | -2,821999947 | 0,004772519 | 0,015655001 |
| ENSG00000164187.7 | LMBRD2 | protein_coding | 7663,870418 | -0,343206203 | 0,049773243 | -6,895395649 | 5,37E-12 | 6,33E-11 |
| ENSG00000164190.19 | NIPBL | protein_coding | 16429,27341 | 0,234991385 | 0,051166225 | 4,592705109 | 4,38E-06 | 2,58E-05 |
| ENSG00000164211.13 | STARD4 | protein_coding | 16280,73925 | 0,715667194 | 0,048377357 | 14,79343298 | 1,62E-49 | 9,18E-48 |
| ENSG00000164237.9 | CMBL | protein_coding | 1044,207968 | -0,392481155 | 0,088139431 | -4,452957654 | 8,47E-06 | 4,79E-05 |
| ENSG00000164241.13 | C5orf63 | protein_coding | 1061,882119 | -0,205695561 | 0,081225832 | -2,532390924 | 0,011328761 | 0,033281008 |
| ENSG00000164244.20 | PRRC1 | protein_coding | 10643,54504 | -0,124924324 | 0,050090313 | -2,493981672 | 0,012631907 | 0,036588803 |
| ENSG00000164251.5 | F2RL1 | protein_coding | 3280,050192 | 0,470015482 | 0,058617027 | 8,018412166 | 1,07E-15 | 1,69E-14 |
| ENSG00000164252.13 | AGGF1 | protein_coding | 11534,10839 | -0,298831439 | 0,043618669 | -6,85099849 | 7,33E-12 | 8,53E-11 |
| ENSG00000164266.10 | SPINK1 | protein_coding | 3393,093312 | 1,368042393 | 0,067573333 | 20,24529978 | 3,91E-91 | 4,13E-89 |
| ENSG00000164291.16 | ARSK | protein_coding | 2248,544814 | 0,254602189 | 0,067139293 | 3,792148799 | 0,000149349 | 0,000688028 |

| | | | | | | | | |
|---|---|---|---|---|---|---|---|---|
| ENSG00000164292.13 | RHOBTB3 | protein_coding | 34134,5262 | -0,464884034 | 0,048221073 | -9,640682036 | 5,38E-22 | 1,22E-20 |
| ENSG00000164300.17 | SERINC5 | protein_coding | 4394,338086 | -0,542140639 | 0,061242648 | -8,852338376 | 8,57E-19 | 1,63E-17 |
| ENSG00000164305.19 | CASP3 | protein_coding | 16343,42367 | 0,209143217 | 0,042685228 | 4,899662587 | 9,60E-07 | 6,25E-06 |
| ENSG00000164307.13 | ERAP1 | protein_coding | 9687,083388 | 1,717648002 | 0,046126867 | 37,23747341 | 1,69E-303 | 4,48E-301 |
| ENSG00000164308.16 | ERAP2 | protein_coding | 2999,968155 | 3,578337479 | 0,079680095 | 44,90880031 | 0 | 0 |
| ENSG00000164323.14 | CFAP97 | protein_coding | 4417,76355 | -0,224906857 | 0,058281933 | -3,85894645 | 0,000113877 | 0,000536246 |
| ENSG00000164326.5 | CARTPT | protein_coding | 31,15858877 | 1,214687276 | 0,42749715 | 2,841392686 | 0,004491697 | 0,014831099 |
| ENSG00000164327.13 | RICTOR | protein_coding | 6510,206301 | 0,500900875 | 0,046181502 | 10,8463531 | 2,08E-27 | 5,90E-26 |
| ENSG00000164329.13 | TENT2 | protein_coding | 4271,743243 | 0,160518189 | 0,062065833 | 2,586256899 | 0,009702459 | 0,029169579 |
| ENSG00000164338.10 | UTP15 | protein_coding | 2106,999612 | -0,203884463 | 0,081621816 | -2,497916296 | 0,012492569 | 0,036236499 |
| ENSG00000164342.12 | TLR3 | protein_coding | 854,2723286 | 6,761812034 | 0,179134185 | 37,74718965 | 0 | 0 |
| ENSG00000164344.16 | KLKB1 | protein_coding | 76,88335736 | 1,549050761 | 0,307601759 | 5,035896958 | 4,76E-07 | 3,24E-06 |
| ENSG00000164346.10 | NSA2 | protein_coding | 13971,87793 | -0,421949068 | 0,043058993 | -9,799325141 | 1,13E-22 | 2,65E-21 |
| ENSG00000164347.18 | GFM2 | protein_coding | 5670,632303 | -0,186649888 | 0,054637914 | -3,416123971 | 0,000635193 | 0,002575674 |
| ENSG00000164366.3 | CCDC127 | protein_coding | 3378,062716 | -0,166782332 | 0,059954934 | -2,781794949 | 0,005405918 | 0,01748302 |
| ENSG00000164405.11 | UQCRQ | protein_coding | 12410,19347 | -0,349850991 | 0,044915437 | -7,78910366 | 6,75E-15 | 1,01E-13 |
| ENSG00000164440.15 | TXLNB | protein_coding | 87,73239703 | 0,811867535 | 0,254917023 | 3,1848306 | 0,001448388 | 0,005416474 |
| ENSG00000164463.12 | CREBRF | protein_coding | 1822,197052 | 0,903564209 | 0,070637942 | 12,7914855 | 1,83E-37 | 7,54E-36 |
| ENSG00000164465.18 | DCBLD1 | protein_coding | 3660,935433 | 0,562560426 | 0,053745995 | 10,46702041 | 1,22E-25 | 3,26E-24 |
| ENSG00000164466.13 | SFXN1 | protein_coding | 15406,42429 | -0,267593678 | 0,057640285 | -4,642476652 | 3,44E-06 | 2,06E-05 |
| ENSG00000164494.12 | PDSS2 | protein_coding | 3143,255671 | -0,303118803 | 0,07897884 | -3,837974845 | 0,000124053 | 0,000580885 |
| ENSG00000164542.12 | KIAA0895 | protein_coding | 5084,55148 | -0,507281596 | 0,054766397 | -9,262643185 | 1,99E-20 | 4,16E-19 |
| ENSG00000164543.7 | STK17A | protein_coding | 866,9853268 | 1,023864714 | 0,085507283 | 11,97400589 | 4,86E-33 | 1,76E-31 |
| ENSG00000164587.13 | RPS14 | protein_coding | 34879,10515 | -0,37518854 | 0,04255085 | -8,817415892 | 1,17E-18 | 2,22E-17 |
| ENSG00000164603.12 | BMT2 | protein_coding | 3177,513391 | 0,201072345 | 0,059285293 | 3,391605795 | 0,000694843 | 0,002792399 |
| ENSG00000164604.12 | GPR85 | protein_coding | 330,3810105 | -0,545087358 | 0,161795689 | -3,368985665 | 0,000754454 | 0,003005131 |
| ENSG00000164609.10 | SLU7 | protein_coding | 8777,686591 | 0,111160302 | 0,046385489 | 2,396445608 | 0,01655495 | 0,046203981 |
| ENSG00000164611.13 | PTTG1 | protein_coding | 6602,593931 | -0,718569198 | 0,04884661 | -14,71072798 | 5,50E-49 | 3,07E-47 |
| ENSG00000164616.16 | FBXL21P | transcribed_unitary | 113,8773634 | -0,849498768 | 0,2267712 | -3,746061078 | 0,000179633 | 0,000812408 |
| ENSG00000164620.9 | RELL2 | protein_coding | 1132,889248 | 0,55630127 | 0,082480591 | 6,744632477 | 1,53E-11 | 1,72E-10 |
| ENSG00000164627.18 | KIF6 | protein_coding | 2003,710732 | 1,167275293 | 0,070462875 | 16,56581976 | 1,23E-61 | 8,80E-60 |
| ENSG00000164631.19 | ZNF12 | protein_coding | 3444,231834 | 0,16969449 | 0,063564794 | 2,669630154 | 0,007593484 | 0,023555916 |
| ENSG00000164638.10 | SLC29A4 | protein_coding | 8663,618937 | -0,183426346 | 0,066842196 | -2,744169958 | 0,006066412 | 0,019349422 |
| ENSG00000164649.20 | CDCA7L | protein_coding | 3817,928959 | -0,140725236 | 0,057075563 | -2,465595237 | 0,013678581 | 0,039226803 |
| ENSG00000164651.16 | SP8 | protein_coding | 601,8396066 | -0,414697281 | 0,114783237 | -3,61287321 | 0,000302823 | 0,001313745 |
| ENSG00000164663.14 | USP49 | protein_coding | 3971,933134 | -0,408261002 | 0,064270911 | -6,352189408 | 2,12E-10 | 2,14E-09 |

| | | | | | | | | |
|---|---|---|---|---|---|---|---|---|
| ENSG00000164674.15 | SYTL3 | protein_coding | 92,73542785 | 1,760937873 | 0,267684002 | 6,578420309 | 4,75E-11 | 5,11E-10 |
| ENSG00000164695.5 | CHMP4C | protein_coding | 3254,457183 | 0,2372745 | 0,057583436 | 4,120533909 | 3,78E-05 | 0,000193063 |
| ENSG00000164713.9 | BRI3 | protein_coding | 2035,311698 | 0,20188341 | 0,068746754 | 2,936624628 | 0,003318055 | 0,011347021 |
| ENSG00000164733.21 | CTSB | protein_coding | 9680,577467 | 0,493954434 | 0,050719688 | 9,738909183 | 2,06E-22 | 4,74E-21 |
| ENSG00000164742.16 | ADCY1 | protein_coding | 16049,45026 | -0,338647712 | 0,051934313 | -6,520692987 | 7,00E-11 | 7,39E-10 |
| ENSG00000164749.11 | HNF4G | protein_coding | 2909,381145 | 0,423770134 | 0,067943159 | 6,237127368 | 4,46E-10 | 4,35E-09 |
| ENSG00000164754.14 | RAD21 | protein_coding | 75250,48001 | -0,314368958 | 0,042084156 | -7,470007422 | 8,02E-14 | 1,11E-12 |
| ENSG00000164756.12 | SLC30A8 | protein_coding | 52663,80975 | -0,156137749 | 0,054792777 | -2,84960458 | 0,004377361 | 0,014496105 |
| ENSG00000164818.16 | DNAAF5 | protein_coding | 699,5676313 | -0,347344847 | 0,099567013 | -3,488553463 | 0,000485642 | 0,002019082 |
| ENSG00000164850.15 | GPER1 | protein_coding | 2850,190336 | -0,196247099 | 0,069795193 | -2,81175665 | 0,004927177 | 0,016117689 |
| ENSG00000164879.7 | CA3 | protein_coding | 39,78146978 | 1,397332143 | 0,389222268 | 3,590062176 | 0,000330599 | 0,001423962 |
| ENSG00000164885.13 | CDK5 | protein_coding | 2329,414482 | -0,151180325 | 0,060718777 | -2,489844697 | 0,012779893 | 0,036946134 |
| ENSG00000164889.14 | SLC4A2 | protein_coding | 11363,80013 | 0,189401234 | 0,045675657 | 4,146655916 | 3,37E-05 | 0,000173691 |
| ENSG00000164896.20 | FASTK | protein_coding | 5581,514522 | -0,142730916 | 0,056246136 | -2,537612817 | 0,011161139 | 0,032874388 |
| ENSG00000164898.13 | FMC1 | protein_coding | 1920,878371 | -0,227121855 | 0,063499947 | -3,576725093 | 0,000347926 | 0,001491423 |
| ENSG00000164902.14 | PHAX | protein_coding | 6085,380648 | -0,14754098 | 0,046861528 | -3,148445777 | 0,001641412 | 0,006063768 |
| ENSG00000164904.18 | ALDH7A1 | protein_coding | 20537,33408 | -0,372509338 | 0,049087435 | -7,588690174 | 3,23E-14 | 4,58E-13 |
| ENSG00000164916.11 | FOXK1 | protein_coding | 6566,920445 | -0,264465168 | 0,050943856 | -5,191306481 | 2,09E-07 | 1,49E-06 |
| ENSG00000164919.11 | COX6C | protein_coding | 14868,5 | -0,209642363 | 0,041968144 | -4,995273578 | 5,88E-07 | 3,95E-06 |
| ENSG00000164929.17 | BAALC | protein_coding | 1115,061725 | -0,208029116 | 0,084420272 | -2,46420808 | 0,013731637 | 0,039368923 |
| ENSG00000164930.12 | FZD6 | protein_coding | 2795,085585 | 0,28231631 | 0,058614668 | 4,816478857 | 1,46E-06 | 9,26E-06 |
| ENSG00000164933.12 | SLC25A32 | protein_coding | 1904,717418 | 0,219348915 | 0,069774095 | 3,143701329 | 0,001668256 | 0,006155865 |
| ENSG00000164934.14 | DCAF13 | protein_coding | 6695,372924 | -0,197012055 | 0,052361523 | -3,762534869 | 0,0001682 | 0,000765624 |
| ENSG00000164938.14 | TP53INP1 | protein_coding | 11405,92405 | 0,109566307 | 0,04543674 | 2,411403352 | 0,015891263 | 0,04457832 |
| ENSG00000164946.19 | FREM1 | protein_coding | 338,3754418 | 2,077259341 | 0,214291754 | 9,693603713 | 3,21E-22 | 7,33E-21 |
| ENSG00000164951.16 | PDP1 | protein_coding | 2492,239229 | -0,180435563 | 0,060276412 | -2,99346888 | 0,002758256 | 0,009615238 |
| ENSG00000164953.15 | TMEM67 | protein_coding | 877,0439164 | -0,287292884 | 0,096946039 | -2,963430873 | 0,003042304 | 0,010490076 |
| ENSG00000164961.16 | WASHC5 | protein_coding | 7087,744391 | -0,183581958 | 0,050445811 | -3,639191323 | 0,000273496 | 0,001196523 |
| ENSG00000164967.10 | RPP25L | protein_coding | 1170,769726 | 0,220718448 | 0,077543482 | 2,846382969 | 0,004421898 | 0,01462208 |
| ENSG00000164970.15 | FAM219A | protein_coding | 4181,970049 | 0,268582977 | 0,055030841 | 4,880590089 | 1,06E-06 | 6,85E-06 |
| ENSG00000164975.15 | SNAPC3 | protein_coding | 8407,519017 | 0,130455915 | 0,053259421 | 2,449443005 | 0,014307735 | 0,040797164 |
| ENSG00000164978.18 | NUDT2 | protein_coding | 1740,398243 | -0,249819061 | 0,071481211 | -3,494891298 | 0,000474255 | 0,00197612 |
| ENSG00000164983.7 | TMEM65 | protein_coding | 9038,586234 | -0,150195188 | 0,047561057 | -3,157944726 | 0,001588857 | 0,005890887 |
| ENSG00000164985.15 | PSIP1 | protein_coding | 22735,76276 | -0,172510025 | 0,039109666 | -4,4109307 | 1,03E-05 | 5,76E-05 |
| ENSG00000164989.17 | CCDC171 | protein_coding | 1057,292577 | -0,314509372 | 0,083064668 | -3,786319502 | 0,000152895 | 0,000703211 |
| ENSG00000165006.14 | UBAP1 | protein_coding | 5962,652678 | 0,250187687 | 0,049630905 | 5,040965623 | 4,63E-07 | 3,16E-06 |

| | | | | | | | | |
|---|---|---|---|---|---|---|---|---|
| ENSG00000165028.12 | NIPSNAP3B | protein_coding | 557,8499181 | -0,393772047 | 0,11823409 | -3,330444267 | 0,000867075 | 0,003406107 |
| ENSG00000165029.16 | ABCA1 | protein_coding | 409,4500237 | 0,456348315 | 0,136281613 | 3,348568498 | 0,000812302 | 0,003216201 |
| ENSG00000165030.4 | NFIL3 | protein_coding | 2584,146307 | 0,195901308 | 0,063023664 | 3,108376995 | 0,001881179 | 0,00684173 |
| ENSG00000165055.15 | METTL2B | protein_coding | 5975,16877 | -0,218769411 | 0,053450908 | -4,092903558 | 4,26E-05 | 0,000216259 |
| ENSG00000165066.12 | NKX6-3 | protein_coding | 57572,2559 | 0,273579402 | 0,055819698 | 4,90112648 | 9,53E-07 | 6,21E-06 |
| ENSG00000165092.13 | ALDH1A1 | protein_coding | 147120,1208 | -0,121799668 | 0,039941147 | -3,049478502 | 0,00229239 | 0,008152854 |
| ENSG00000165097.15 | KDM1B | protein_coding | 3695,216557 | -0,286492582 | 0,062248499 | -4,602401474 | 4,18E-06 | 2,47E-05 |
| ENSG00000165115.15 | KIF27 | protein_coding | 1370,827209 | 0,558139131 | 0,091680674 | 6,087860254 | 1,14E-09 | 1,06E-08 |
| ENSG00000165119.21 | HNRNPK | protein_coding | 101756,6748 | -0,132840917 | 0,039020465 | -3,404390919 | 0,000663118 | 0,002677811 |
| ENSG00000165121.11 | AL353743.1 | transcribed_unproc | 830,2948781 | 0,756390316 | 0,090124226 | 8,392752404 | 4,75E-17 | 8,13E-16 |
| ENSG00000165156.15 | ZHX1 | protein_coding | 12899,15391 | 0,133195753 | 0,048954961 | 2,720781501 | 0,006512779 | 0,020609609 |
| ENSG00000165169.11 | DYNLT3 | protein_coding | 25324,51863 | -0,273648338 | 0,046468327 | -5,888921719 | 3,89E-09 | 3,40E-08 |
| ENSG00000165175.15 | MID1IP1 | protein_coding | 7419,213431 | 0,415469678 | 0,054411393 | 7,635711179 | 2,25E-14 | 3,21E-13 |
| ENSG00000165197.5 | VEGFD | protein_coding | 147,7264313 | -1,887882168 | 0,201537177 | -9,367413967 | 7,43E-21 | 1,59E-19 |
| ENSG00000165209.18 | STRBP | protein_coding | 9378,147978 | -0,213821082 | 0,04770537 | -4,482117707 | 7,39E-06 | 4,21E-05 |
| ENSG00000165240.20 | ATP7A | protein_coding | 2762,482498 | -0,166259314 | 0,059817852 | -2,779426339 | 0,0054455 | 0,017595861 |
| ENSG00000165244.7 | ZNF367 | protein_coding | 4833,151614 | -0,236873226 | 0,048919539 | -4,842098486 | 1,28E-06 | 8,22E-06 |
| ENSG00000165246.14 | NLGN4Y | protein_coding | 2865,198927 | 0,45439748 | 0,058421443 | 7,777922959 | 7,37E-15 | 1,10E-13 |
| ENSG00000165259.14 | HDX | protein_coding | 2097,817944 | 1,385547557 | 0,092621819 | 14,95919183 | 1,36E-50 | 7,89E-49 |
| ENSG00000165264.11 | NDUFB6 | protein_coding | 6139,306226 | -0,274966211 | 0,057888222 | -4,749950837 | 2,03E-06 | 1,26E-05 |
| ENSG00000165269.13 | AQP7 | protein_coding | 828,1885055 | 0,277881222 | 0,094785868 | 2,931673546 | 0,003371409 | 0,011517234 |
| ENSG00000165271.17 | NOL6 | protein_coding | 3783,834391 | 0,202765887 | 0,057207013 | 3,544423569 | 0,000393473 | 0,001666354 |
| ENSG00000165275.10 | TRMT10B | protein_coding | 1321,282963 | 0,384373202 | 0,072558617 | 5,297416299 | 1,17E-07 | 8,61E-07 |
| ENSG00000165288.10 | BRWD3 | protein_coding | 6368,974439 | 0,270037217 | 0,047349411 | 5,703074489 | 1,18E-08 | 9,70E-08 |
| ENSG00000165304.8 | MELK | protein_coding | 8162,429849 | -0,237004315 | 0,056396558 | -4,202460662 | 2,64E-05 | 0,000138114 |
| ENSG00000165312.6 | OTUD1 | protein_coding | 408,906104 | 0,333379138 | 0,122866376 | 2,713347213 | 0,006660728 | 0,021015744 |
| ENSG00000165322.17 | ARHGAP12 | protein_coding | 3341,347678 | 0,323348205 | 0,071753446 | 4,506378768 | 6,59E-06 | 3,79E-05 |
| ENSG00000165325.13 | DEUP1 | protein_coding | 2499,921663 | -0,342494679 | 0,071529972 | -4,788128228 | 1,68E-06 | 1,06E-05 |
| ENSG00000165359.15 | INTS6L | protein_coding | 576,1605317 | 0,492779615 | 0,105576084 | 4,667530709 | 3,05E-06 | 1,85E-05 |
| ENSG00000165406.16 | mars-08 | protein_coding | 12858,65114 | 0,388721533 | 0,045791029 | 8,489032481 | 2,08E-17 | 3,65E-16 |
| ENSG00000165410.14 | CFL2 | protein_coding | 3623,761753 | 0,206057729 | 0,059747016 | 3,448837129 | 0,000563006 | 0,00231043 |
| ENSG00000165417.12 | GTF2A1 | protein_coding | 12052,60372 | -0,20338011 | 0,045635903 | -4,456581291 | 8,33E-06 | 4,72E-05 |
| ENSG00000165434.8 | PGM2L1 | protein_coding | 3576,359417 | -0,144184002 | 0,05363749 | -2,688119842 | 0,00718556 | 0,022460738 |
| ENSG00000165443.12 | PHYHIPL | protein_coding | 7062,12608 | 0,157166486 | 0,050324473 | 3,123062715 | 0,001789796 | 0,006553851 |
| ENSG00000165449.11 | SLC16A9 | protein_coding | 7736,007365 | 0,136263002 | 0,049883239 | 2,731639018 | 0,006302015 | 0,020024287 |
| ENSG00000165475.15 | CRYL1 | protein_coding | 287,6699947 | -0,612479154 | 0,145404362 | -4,212247471 | 2,53E-05 | 0,000132942 |

| | | | | | | | | |
|---|---|---|---|---|---|---|---|---|
| ENSG00000165476.14 | REEP3 | protein_coding | 15289,44149 | 0,145236642 | 0,04470409 | 3,248844632 | 0,001158747 | 0,004424469 |
| ENSG00000165480.16 | SKA3 | protein_coding | 2042,980564 | -0,398705226 | 0,079702788 | -5,002400023 | 5,66E-07 | 3,81E-06 |
| ENSG00000165507.8 | DEPP1 | protein_coding | 4372,106438 | 0,361870057 | 0,05542441 | 6,529073699 | 6,62E-11 | 7,00E-10 |
| ENSG00000165511.6 | C10orf25 | lncRNA | 282,0940994 | 0,499946391 | 0,139619551 | 3,580776373 | 0,000342575 | 0,001471849 |
| ENSG00000165516.11 | KLHDC2 | protein_coding | 7669,842694 | -0,182062606 | 0,043971279 | -4,140489179 | 3,47E-05 | 0,00017802 |
| ENSG00000165521.15 | AL121768.1 | protein_coding | 7207,740688 | -0,185649899 | 0,055199375 | -3,36326087 | 0,000770275 | 0,003062732 |
| ENSG00000165548.11 | TMEM63C | protein_coding | 5275,901242 | 0,201171796 | 0,079978572 | 2,515321187 | 0,011892399 | 0,034705822 |
| ENSG00000165566.12 | AMER2 | protein_coding | 119,7280872 | -1,371503859 | 0,263060141 | -5,213651347 | 1,85E-07 | 1,33E-06 |
| ENSG00000165572.7 | KBTBD6 | protein_coding | 4613,586017 | -0,135100932 | 0,050780792 | -2,660473089 | 0,007803096 | 0,024129674 |
| ENSG00000165591.6 | FAAH2 | protein_coding | 1651,670844 | 0,341315885 | 0,07451557 | 4,580463968 | 4,64E-06 | 2,73E-05 |
| ENSG00000165629.20 | ATP5F1C | protein_coding | 11136,5927 | -0,115817197 | 0,047517571 | -2,43735516 | 0,01479514 | 0,041921426 |
| ENSG00000165637.13 | VDAC2 | protein_coding | 12623,99512 | -0,109891657 | 0,042006675 | -2,616052251 | 0,008895294 | 0,027086605 |
| ENSG00000165644.10 | COMTD1 | protein_coding | 655,0615493 | -0,408984641 | 0,136900222 | -2,98746513 | 0,002813014 | 0,009780372 |
| ENSG00000165646.14 | SLC18A2 | protein_coding | 1824,897036 | -0,534593151 | 0,074641048 | -7,162187107 | 7,94E-13 | 1,01E-11 |
| ENSG00000165650.12 | PDZD8 | protein_coding | 6167,198959 | 0,176136797 | 0,067603417 | 2,605442241 | 0,009175574 | 0,027815892 |
| ENSG00000165655.17 | ZNF503 | protein_coding | 1724,383168 | 0,869213034 | 0,082117849 | 10,58494649 | 3,50E-26 | 9,56E-25 |
| ENSG00000165671.20 | NSD1 | protein_coding | 17560,53058 | 0,218658334 | 0,051011078 | 4,286487233 | 1,82E-05 | 9,75E-05 |
| ENSG00000165672.7 | PRDX3 | protein_coding | 21516,09668 | -0,298538355 | 0,047611682 | -6,270275275 | 3,60E-10 | 3,55E-09 |
| ENSG00000165678.21 | GHITM | protein_coding | 27165,24087 | -0,171325175 | 0,040666021 | -4,212981063 | 2,52E-05 | 0,000132603 |
| ENSG00000165688.12 | PMPCA | protein_coding | 4866,250504 | -0,171351318 | 0,069173612 | -2,477119706 | 0,013244747 | 0,038133545 |
| ENSG00000165699.14 | TSC1 | protein_coding | 5265,308551 | -0,195326711 | 0,065842125 | -2,966591855 | 0,003011204 | 0,010393987 |
| ENSG00000165704.15 | HPRT1 | protein_coding | 6726,319319 | -0,274360626 | 0,048469077 | -5,660529149 | 1,51E-08 | 1,23E-07 |
| ENSG00000165714.11 | BORCS5 | protein_coding | 676,6292057 | -0,311802188 | 0,112495571 | -2,771684116 | 0,005576712 | 0,017978548 |
| ENSG00000165716.11 | DIPK1B | protein_coding | 2007,271589 | -0,259845956 | 0,069397944 | -3,744288956 | 0,000180905 | 0,000817178 |
| ENSG00000165731.19 | RET | protein_coding | 2067,448262 | 0,54255529 | 0,077843701 | 6,969803407 | 3,17E-12 | 3,83E-11 |
| ENSG00000165752.17 | STK32C | protein_coding | 595,502183 | -0,417218687 | 0,109241282 | -3,819240138 | 0,000133863 | 0,000622034 |
| ENSG00000165775.18 | FUNDC2 | protein_coding | 5853,538186 | -0,256905538 | 0,053268029 | -4,822884222 | 1,41E-06 | 8,99E-06 |
| ENSG00000165792.17 | METTL17 | protein_coding | 4805,597276 | -0,119619836 | 0,04952974 | -2,415111319 | 0,015730401 | 0,044171108 |
| ENSG00000165801.10 | ARHGEF40 | protein_coding | 2254,624749 | 0,400171435 | 0,100208914 | 3,993371638 | 6,51E-05 | 0,00032035 |
| ENSG00000165802.22 | NSMF | protein_coding | 4157,372207 | -0,790299993 | 0,055728931 | -14,18114388 | 1,20E-45 | 6,26E-44 |
| ENSG00000165806.19 | CASP7 | protein_coding | 2096,072933 | 2,83801629 | 0,096940478 | 29,27586427 | 2,10E-188 | 4,01E-186 |
| ENSG00000165813.20 | CCDC186 | protein_coding | 4202,50228 | 0,152340375 | 0,057714598 | 2,639546657 | 0,0083017 | 0,025489422 |
| ENSG00000165861.14 | ZFYVE1 | protein_coding | 2996,500814 | 0,267024774 | 0,063736127 | 4,189535637 | 2,80E-05 | 0,000145577 |
| ENSG00000165898.13 | ISCA2 | protein_coding | 2978,242479 | -0,249435057 | 0,062412324 | -3,996567387 | 6,43E-05 | 0,000316473 |
| ENSG00000165905.18 | LARGE2 | protein_coding | 2956,039957 | -0,20987048 | 0,068934177 | -3,044505481 | 0,002330632 | 0,008271869 |
| ENSG00000165912.16 | PACSIN3 | protein_coding | 3312,062736 | -0,530372765 | 0,076232249 | -6,957328036 | 3,47E-12 | 4,17E-11 |

| | | | | | | | | |
|---|---|---|---|---|---|---|---|---|
| ENSG00000165916.8 | PSMC3 | protein_coding | 10657,80906 | -0,206549958 | 0,043968181 | -4,697714384 | 2,63E-06 | 1,61E-05 |
| ENSG00000165923.16 | AGBL2 | protein_coding | 372,0864014 | 1,464339762 | 0,183033653 | 8,000385377 | 1,24E-15 | 1,94E-14 |
| ENSG00000165929.13 | TC2N | protein_coding | 4379,905733 | -0,194357892 | 0,052491947 | -3,70262304 | 0,000213382 | 0,000952034 |
| ENSG00000165935.9 | SMCO2 | protein_coding | 29,69690891 | 1,471322898 | 0,481155648 | 3,057893853 | 0,002228985 | 0,007949965 |
| ENSG00000165948.11 | IFI27L1 | protein_coding | 1664,388328 | 0,261693457 | 0,067612246 | 3,870503803 | 0,000108611 | 0,00051338 |
| ENSG00000165949.12 | IFI27 | protein_coding | 10108,19184 | 11,37277908 | 0,620060068 | 18,34141508 | 3,87E-75 | 3,40E-73 |
| ENSG00000165997.5 | ARL5B | protein_coding | 7104,84396 | 0,293444882 | 0,051193028 | 5,732125922 | 9,92E-09 | 8,24E-08 |
| ENSG00000166004.15 | CEP295 | protein_coding | 9414,220012 | 0,147098187 | 0,048017319 | 3,063440262 | 0,002188079 | 0,00782266 |
| ENSG00000166006.13 | KCNC2 | protein_coding | 1154,572717 | 1,880697883 | 0,097348518 | 19,31922464 | 3,70E-83 | 3,53E-81 |
| ENSG00000166012.16 | TAF1D | protein_coding | 6484,520545 | 0,143846395 | 0,047875545 | 3,004590232 | 0,002659389 | 0,009305173 |
| ENSG00000166016.6 | ABTB2 | protein_coding | 2676,22268 | 1,14805467 | 0,077501025 | 14,81341268 | 1,20E-49 | 6,84E-48 |
| ENSG00000166024.13 | R3HCC1L | protein_coding | 3946,771756 | 0,543818953 | 0,059356769 | 9,161869163 | 5,10E-20 | 1,03E-18 |
| ENSG00000166025.18 | AMOTL1 | protein_coding | 4730,213492 | -0,758615648 | 0,065534138 | -11,57588507 | 5,46E-31 | 1,82E-29 |
| ENSG00000166073.10 | GPR176 | protein_coding | 1253,124565 | -0,35178152 | 0,078865301 | -4,460536103 | 8,18E-06 | 4,63E-05 |
| ENSG00000166104.15 | AC126323.1 | transcribed_unproc | 42,07882225 | 1,121385224 | 0,369940205 | 3,031260756 | 0,002435348 | 0,008598785 |
| ENSG00000166106.3 | ADAMTS15 | protein_coding | 197,3025769 | 0,514820528 | 0,184967082 | 2,783308919 | 0,005380755 | 0,017414149 |
| ENSG00000166123.14 | GPT2 | protein_coding | 4696,402133 | 0,236048803 | 0,053319135 | 4,427093589 | 9,55E-06 | 5,37E-05 |
| ENSG00000166128.13 | RAB8B | protein_coding | 4958,151842 | 0,598889213 | 0,065420548 | 9,154451161 | 5,46E-20 | 1,10E-18 |
| ENSG00000166130.15 | IKBIP | protein_coding | 2787,651178 | -0,272867457 | 0,068273801 | -3,996664231 | 6,42E-05 | 0,000316413 |
| ENSG00000166143.9 | PPP1R14D | protein_coding | 495,3474935 | 0,347242075 | 0,115551171 | 3,005093512 | 0,002654992 | 0,009292679 |
| ENSG00000166145.14 | SPINT1 | protein_coding | 13390,32251 | 0,217738551 | 0,044602839 | 4,881719524 | 1,05E-06 | 6,82E-06 |
| ENSG00000166164.15 | BRD7 | protein_coding | 16195,46938 | 0,154168572 | 0,046972593 | 3,282096226 | 0,001030384 | 0,003980251 |
| ENSG00000166169.17 | POLL | protein_coding | 1906,817726 | 0,222167127 | 0,080633582 | 2,755267993 | 0,005864408 | 0,018774151 |
| ENSG00000166170.9 | BAG5 | protein_coding | 7253,902604 | -0,156423313 | 0,051956716 | -3,010646654 | 0,00260692 | 0,009148607 |
| ENSG00000166171.13 | DPCD | protein_coding | 1107,294493 | -0,294240837 | 0,085335447 | -3,448049424 | 0,000564651 | 0,002315912 |
| ENSG00000166197.16 | NOLC1 | protein_coding | 14927,79409 | -0,150927826 | 0,043123771 | -3,499875408 | 0,000465476 | 0,001945301 |
| ENSG00000166225.8 | FRS2 | protein_coding | 6008,128122 | 0,238671909 | 0,048451482 | 4,92599814 | 8,39E-07 | 5,50E-06 |
| ENSG00000166226.13 | CCT2 | protein_coding | 29720,03529 | -0,19160965 | 0,052205771 | -3,670277179 | 0,000242288 | 0,001070153 |
| ENSG00000166228.9 | PCBD1 | protein_coding | 12553,95002 | -0,407049227 | 0,049634983 | -8,200853584 | 2,39E-16 | 3,91E-15 |
| ENSG00000166233.15 | ARIH1 | protein_coding | 11558,83228 | 0,203446796 | 0,049637841 | 4,098623027 | 4,16E-05 | 0,000211271 |
| ENSG00000166250.12 | CLMP | protein_coding | 527,6667609 | 0,877709791 | 0,127862462 | 6,864483741 | 6,67E-12 | 7,80E-11 |
| ENSG00000166261.11 | ZNF202 | protein_coding | 1705,030119 | 0,182877518 | 0,066500169 | 2,75003087 | 0,005958965 | 0,019051492 |
| ENSG00000166265.12 | CYYR1 | protein_coding | 803,7393455 | -0,264484816 | 0,111362225 | -2,374995803 | 0,01754915 | 0,048539421 |
| ENSG00000166266.14 | CUL5 | protein_coding | 6835,557427 | 0,23660979 | 0,051485561 | 4,595653318 | 4,31E-06 | 2,55E-05 |
| ENSG00000166272.18 | WBP1L | protein_coding | 2849,692703 | 0,237690294 | 0,056779555 | 4,186195112 | 2,84E-05 | 0,000147599 |
| ENSG00000166275.15 | BORCS7 | protein_coding | 7845,385867 | -0,335447693 | 0,051926104 | -6,460097518 | 1,05E-10 | 1,09E-09 |

| | | | | | | | | |
|---|---|---|---|---|---|---|---|---|
| ENSG00000166278.15 | C2 | protein_coding | 127,3341685 | 3,31316755 | 0,256029955 | 12,94054655 | 2,66E-38 | 1,13E-36 |
| ENSG00000166295.9 | ANAPC16 | protein_coding | 11002,23707 | -0,355967746 | 0,05210777 | -6,831375585 | 8,41E-12 | 9,72E-11 |
| ENSG00000166313.19 | APBB1 | protein_coding | 5392,857037 | -0,134139419 | 0,054724917 | -2,451158003 | 0,014239743 | 0,040629025 |
| ENSG00000166317.12 | SYNPO2L | protein_coding | 48,21872891 | -0,918101717 | 0,342884868 | -2,677580149 | 0,007415611 | 0,023080473 |
| ENSG00000166326.7 | TRIM44 | protein_coding | 21649,94027 | -0,173196972 | 0,051993914 | -3,331100837 | 0,000865033 | 0,003400457 |
| ENSG00000166343.10 | MSS51 | protein_coding | 685,7481784 | 1,317066852 | 0,110219614 | 11,94947804 | 6,53E-33 | 2,36E-31 |
| ENSG00000166347.19 | CYB5A | protein_coding | 7591,490066 | -0,194643033 | 0,048575485 | -4,007021902 | 6,15E-05 | 0,000303988 |
| ENSG00000166352.16 | C11orf74 | protein_coding | 1873,340257 | -0,305777642 | 0,079297381 | -3,856087528 | 0,000115216 | 0,000542213 |
| ENSG00000166387.13 | PPFIBP2 | protein_coding | 1263,448937 | 0,238680815 | 0,08210975 | 2,906851089 | 0,003650869 | 0,012359365 |
| ENSG00000166401.14 | SERPINB8 | protein_coding | 1218,971842 | 0,468032443 | 0,080743253 | 5,796551744 | 6,77E-09 | 5,76E-08 |
| ENSG00000166407.14 | LMO1 | protein_coding | 513,02677 | -0,319460044 | 0,13504607 | -2,365563437 | 0,018002659 | 0,049633763 |
| ENSG00000166432.15 | ZMAT1 | protein_coding | 467,8998151 | 0,309913726 | 0,118847718 | 2,607653995 | 0,009116505 | 0,027659175 |
| ENSG00000166435.15 | XRRA1 | protein_coding | 835,2691345 | 0,355470207 | 0,106285627 | 3,344480511 | 0,000824368 | 0,00325705 |
| ENSG00000166436.16 | TRIM66 | protein_coding | 1372,847964 | 0,194725619 | 0,079904506 | 2,436979207 | 0,014810531 | 0,041959754 |
| ENSG00000166439.6 | RNF169 | protein_coding | 3889,40791 | 0,414888438 | 0,053339749 | 7,778222495 | 7,36E-15 | 1,10E-13 |
| ENSG00000166441.13 | RPL27A | protein_coding | 43266,37234 | -0,506194619 | 0,043911422 | -11,52762973 | 9,57E-31 | 3,16E-29 |
| ENSG00000166444.19 | ST5 | protein_coding | 2943,941534 | 0,73215991 | 0,089375768 | 8,191928571 | 2,57E-16 | 4,20E-15 |
| ENSG00000166446.15 | CDYL2 | protein_coding | 4525,734568 | -0,589010932 | 0,058300266 | -10,10305741 | 5,35E-24 | 1,33E-22 |
| ENSG00000166451.13 | CENPN | protein_coding | 4974,794178 | -0,188625683 | 0,053938294 | -3,497064303 | 0,000470408 | 0,001964093 |
| ENSG00000166478.10 | ZNF143 | protein_coding | 2404,211701 | -0,507346569 | 0,082715665 | -6,133621359 | 8,59E-10 | 8,09E-09 |
| ENSG00000166479.10 | TMX3 | protein_coding | 6918,105387 | 0,181799485 | 0,049495998 | 3,673013819 | 0,000239707 | 0,001060864 |
| ENSG00000166482.11 | MFAP4 | protein_coding | 10006,53682 | -0,248390289 | 0,055476919 | -4,477362706 | 7,56E-06 | 4,30E-05 |
| ENSG00000166484.20 | MAPK7 | protein_coding | 2551,858423 | 0,374776126 | 0,073114825 | 5,125856868 | 2,96E-07 | 2,07E-06 |
| ENSG00000166508.17 | MCM7 | protein_coding | 30386,76736 | -0,101475565 | 0,042626197 | -2,380591557 | 0,017284865 | 0,047938027 |
| ENSG00000166510.14 | CCDC68 | protein_coding | 3819,534597 | 0,304656016 | 0,05843777 | 5,213340855 | 1,85E-07 | 1,33E-06 |
| ENSG00000166526.17 | ZNF3 | protein_coding | 10112,7961 | 0,228253488 | 0,047209387 | 4,834917445 | 1,33E-06 | 8,50E-06 |
| ENSG00000166532.16 | RIMKLB | protein_coding | 1509,338629 | 0,776365321 | 0,101588976 | 7,642220138 | 2,14E-14 | 3,06E-13 |
| ENSG00000166548.15 | TK2 | protein_coding | 2264,252356 | 0,295780528 | 0,073280468 | 4,036280569 | 5,43E-05 | 0,000270673 |
| ENSG00000166557.13 | TMED3 | protein_coding | 12957,57663 | 0,241014127 | 0,055430938 | 4,34800734 | 1,37E-05 | 7,53E-05 |
| ENSG00000166579.15 | NDEL1 | protein_coding | 3185,748364 | 0,239985791 | 0,058032793 | 4,135347959 | 3,54E-05 | 0,000181722 |
| ENSG00000166582.10 | CENPV | protein_coding | 4305,874533 | -0,13770173 | 0,056486254 | -2,437791868 | 0,014777279 | 0,041881363 |
| ENSG00000166589.13 | CDH16 | protein_coding | 4889,315029 | 0,181011089 | 0,063483906 | 2,851290984 | 0,00435421 | 0,014432178 |
| ENSG00000166598.15 | HSP90B1 | protein_coding | 254536,8424 | 0,499263266 | 0,042167858 | 11,83990098 | 2,43E-32 | 8,65E-31 |
| ENSG00000166619.15 | BLCAP | protein_coding | 6347,582184 | 0,343474693 | 0,050170439 | 6,846156836 | 7,59E-12 | 8,81E-11 |
| ENSG00000166669.13 | ATF7IP2 | protein_coding | 1538,721161 | -0,742591081 | 0,074426466 | -9,977513699 | 1,91E-23 | 4,61E-22 |
| ENSG00000166681.13 | BEX3 | protein_coding | 10811,01451 | 0,322607577 | 0,042734514 | 7,549110625 | 4,38E-14 | 6,18E-13 |

| | | | | | | | | |
|---|---|---|---|---|---|---|---|---|
| ENSG00000166689.16 | PLEKHA7 | protein_coding | 5338,770856 | 1,010113342 | 0,059450178 | 16,9909221 | 9,59E-65 | 7,17E-63 |
| ENSG00000166707.11 | ZCCHC18 | protein_coding | 831,4158538 | -0,281114488 | 0,105115635 | -2,67433562 | 0,007487747 | 0,023263176 |
| ENSG00000166710.20 | B2M | protein_coding | 459254,0089 | 3,455168087 | 0,247666793 | 13,95087344 | 3,11E-44 | 1,57E-42 |
| ENSG00000166716.10 | ZNF592 | protein_coding | 7975,574889 | 0,320997732 | 0,055316938 | 5,802883197 | 6,52E-09 | 5,56E-08 |
| ENSG00000166734.20 | CASC4 | protein_coding | 16670,43696 | -0,141909478 | 0,044904816 | -3,160228457 | 0,001576455 | 0,005849723 |
| ENSG00000166743.9 | ACSM1 | protein_coding | 103,3631557 | 1,693232043 | 0,29666125 | 5,707627948 | 1,15E-08 | 9,46E-08 |
| ENSG00000166750.10 | SLFN5 | protein_coding | 1109,110566 | 1,748030074 | 0,086108223 | 20,30038491 | 1,28E-91 | 1,36E-89 |
| ENSG00000166770.11 | ZNF667-AS1 | lncRNA | 4911,555919 | -0,30120259 | 0,052060467 | -5,785629811 | 7,22E-09 | 6,12E-08 |
| ENSG00000166788.10 | SAAL1 | protein_coding | 1845,928069 | -0,255509452 | 0,075571001 | -3,381051549 | 0,00072209 | 0,002891057 |
| ENSG00000166794.4 | PPIB | protein_coding | 54117,46948 | 0,763685149 | 0,040858463 | 18,69099075 | 5,86E-78 | 5,36E-76 |
| ENSG00000166801.15 | FAM111A | protein_coding | 7706,309403 | 0,608412519 | 0,057382028 | 10,60284093 | 2,89E-26 | 7,93E-25 |
| ENSG00000166813.15 | KIF7 | protein_coding | 27,19516584 | 1,124639583 | 0,457225602 | 2,459703874 | 0,01390517 | 0,039795481 |
| ENSG00000166821.9 | PEX11A | protein_coding | 1434,665117 | -0,259314819 | 0,089432186 | -2,899569265 | 0,003736758 | 0,012610297 |
| ENSG00000166822.13 | TMEM170A | protein_coding | 4212,434429 | 0,240145349 | 0,061181013 | 3,925161383 | 8,67E-05 | 0,000417619 |
| ENSG00000166823.5 | MESP1 | protein_coding | 219,3256243 | -0,733697613 | 0,232128478 | -3,16073934 | 0,001573693 | 0,005842363 |
| ENSG00000166833.21 | NAV2 | protein_coding | 5423,8309 | -0,217167809 | 0,06163545 | -3,523423746 | 0,00042601 | 0,001793352 |
| ENSG00000166839.17 | ANKDD1A | protein_coding | 442,5849414 | -0,432736113 | 0,157555263 | -2,746567165 | 0,006022255 | 0,019224925 |
| ENSG00000166848.7 | TERF2IP | protein_coding | 15394,58343 | 0,142415293 | 0,042514869 | 3,349776149 | 0,000808769 | 0,00320334 |
| ENSG00000166851.15 | PLK1 | protein_coding | 3483,811733 | -1,248391481 | 0,069606656 | -17,93494414 | 6,29E-72 | 5,21E-70 |
| ENSG00000166860.3 | ZBTB39 | protein_coding | 3260,97739 | 0,189525045 | 0,059683428 | 3,175505341 | 0,001495758 | 0,005587124 |
| ENSG00000166886.13 | NAB2 | protein_coding | 523,952162 | -0,430222073 | 0,108282266 | -3,973153582 | 7,09E-05 | 0,000346764 |
| ENSG00000166887.15 | VPS39 | protein_coding | 10861,38965 | 0,162705682 | 0,04991833 | 3,259437634 | 0,001116333 | 0,004281401 |
| ENSG00000166889.14 | PATL1 | protein_coding | 14265,10519 | 0,521946905 | 0,05201943 | 10,03369134 | 1,08E-23 | 2,64E-22 |
| ENSG00000166896.8 | ATP23 | protein_coding | 553,385671 | -0,2702783 | 0,113704286 | -2,377028253 | 0,017452751 | 0,048344078 |
| ENSG00000166897.15 | ELFN2 | protein_coding | 380,098364 | -0,619083656 | 0,139434277 | -4,439967474 | 9,00E-06 | 5,08E-05 |
| ENSG00000166900.17 | STX3 | protein_coding | 25236,11249 | -0,230495467 | 0,044375923 | -5,19415598 | 2,06E-07 | 1,46E-06 |
| ENSG00000166902.5 | MRPL16 | protein_coding | 1555,392686 | -0,178076411 | 0,073555805 | -2,420970182 | 0,015479147 | 0,043579752 |
| ENSG00000166908.18 | PIP4K2C | protein_coding | 8933,507266 | 0,252705013 | 0,046935739 | 5,384063759 | 7,28E-08 | 5,45E-07 |
| ENSG00000166912.17 | MTMR10 | protein_coding | 6723,521935 | 0,697210825 | 0,059286739 | 11,75997924 | 6,27E-32 | 2,19E-30 |
| ENSG00000166913.13 | YWHAB | protein_coding | 51549,33472 | -0,130134388 | 0,039015192 | -3,335479879 | 0,000851523 | 0,003354381 |
| ENSG00000166922.8 | SCG5 | protein_coding | 30017,95019 | 0,285631456 | 0,047176409 | 6,054539998 | 1,41E-09 | 1,30E-08 |
| ENSG00000166923.12 | GREM1 | protein_coding | 3701,777348 | 0,51056619 | 0,052239682 | 9,773531696 | 1,46E-22 | 3,40E-21 |
| ENSG00000166925.9 | TSC22D4 | protein_coding | 2467,990423 | 0,327979236 | 0,069910567 | 4,69141148 | 2,71E-06 | 1,66E-05 |
| ENSG00000166946.14 | CCNDBP1 | protein_coding | 2962,899469 | -0,392951065 | 0,059624353 | -6,590445751 | 4,39E-11 | 4,73E-10 |
| ENSG00000166963.13 | MAP1A | protein_coding | 5132,26353 | -0,455295615 | 0,056951948 | -7,99438179 | 1,30E-15 | 2,04E-14 |
| ENSG00000166965.12 | RCCD1 | protein_coding | 2374,756628 | -0,492472018 | 0,067055074 | -7,344291571 | 2,07E-13 | 2,76E-12 |

| | | | | | | | | |
|---|---|---|---|---|---|---|---|---|
| ENSG00000166971.17 | AKTIP | protein_coding | 2178,794593 | -0,165367554 | 0,064437876 | -2,566309837 | 0,010278697 | 0,030640072 |
| ENSG00000166974.12 | MAPRE2 | protein_coding | 12580,7295 | -0,137883543 | 0,057638068 | -2,392230492 | 0,016746322 | 0,046651357 |
| ENSG00000166986.15 | MARS | protein_coding | 12431,95598 | 0,352362512 | 0,057840702 | 6,091947401 | 1,12E-09 | 1,04E-08 |
| ENSG00000166987.15 | MBD6 | protein_coding | 4757,716408 | -0,135977217 | 0,05182307 | -2,623874187 | 0,008693589 | 0,026548939 |
| ENSG00000166997.8 | CNPY4 | protein_coding | 2793,555091 | 0,174526608 | 0,063011723 | 2,769748237 | 0,005609964 | 0,018067631 |
| ENSG00000167004.13 | PDIA3 | protein_coding | 124856,0529 | 0,502001114 | 0,045506346 | 11,03145303 | 2,69E-28 | 7,95E-27 |
| ENSG00000167005.14 | NUDT21 | protein_coding | 23749,66566 | -0,36005788 | 0,044938507 | -8,012235097 | 1,13E-15 | 1,77E-14 |
| ENSG00000167034.10 | NKX3-1 | protein_coding | 373,3593177 | 2,374914684 | 0,135950056 | 17,46902323 | 2,47E-68 | 1,96E-66 |
| ENSG00000167037.18 | SGSM1 | protein_coding | 2264,762071 | -0,491749941 | 0,102869768 | -4,780315472 | 1,75E-06 | 1,10E-05 |
| ENSG00000167065.13 | DUSP18 | protein_coding | 1758,926776 | -0,32122297 | 0,067761057 | -4,740524752 | 2,13E-06 | 1,32E-05 |
| ENSG00000167074.15 | TEF | protein_coding | 1701,254803 | 0,211452654 | 0,066654986 | 3,172345641 | 0,001512129 | 0,005636118 |
| ENSG00000167081.17 | PBX3 | protein_coding | 13700,914 | -0,136574144 | 0,045923123 | -2,973973368 | 0,002939706 | 0,010175279 |
| ENSG00000167085.11 | PHB | protein_coding | 10848,24289 | -0,196108311 | 0,043145966 | -4,545229317 | 5,49E-06 | 3,20E-05 |
| ENSG00000167088.11 | SNRPD1 | protein_coding | 15038,94732 | -0,202764941 | 0,044709357 | -4,535179088 | 5,76E-06 | 3,34E-05 |
| ENSG00000167094.16 | TTC16 | protein_coding | 30,16789806 | 2,324511567 | 0,481066635 | 4,831994981 | 1,35E-06 | 8,61E-06 |
| ENSG00000167105.8 | TMEM92 | protein_coding | 31,70354445 | 2,068552091 | 0,519947287 | 3,978388087 | 6,94E-05 | 0,000339736 |
| ENSG00000167106.12 | FAM102A | protein_coding | 4041,222785 | -0,44982131 | 0,085704873 | -5,248491677 | 1,53E-07 | 1,11E-06 |
| ENSG00000167107.12 | ACSF2 | protein_coding | 2295,570379 | -0,203390016 | 0,071046931 | -2,862755856 | 0,00419974 | 0,013990226 |
| ENSG00000167114.13 | SLC27A4 | protein_coding | 3097,752916 | -0,17366325 | 0,066237593 | -2,621823087 | 0,008746082 | 0,026690016 |
| ENSG00000167117.9 | ANKRD40CL | protein_coding | 27,78147032 | -2,308279098 | 0,560700988 | -4,116773729 | 3,84E-05 | 0,000196061 |
| ENSG00000167123.19 | CERCAM | protein_coding | 1932,391893 | -0,168804409 | 0,068561115 | -2,46210127 | 0,013812566 | 0,039575744 |
| ENSG00000167131.17 | CCDC103 | protein_coding | 428,7643226 | 0,574904905 | 0,12863006 | 4,469444438 | 7,84E-06 | 4,46E-05 |
| ENSG00000167186.11 | COQ7 | protein_coding | 4391,611035 | -0,331695459 | 0,057378191 | -5,780862962 | 7,43E-09 | 6,27E-08 |
| ENSG00000167191.12 | GPRC5B | protein_coding | 364,8566337 | -0,640406499 | 0,15680637 | -4,084059197 | 4,43E-05 | 0,000223955 |
| ENSG00000167193.8 | CRK | protein_coding | 9101,464492 | 0,213512638 | 0,044011503 | 4,851291647 | 1,23E-06 | 7,87E-06 |
| ENSG00000167216.16 | KATNAL2 | protein_coding | 186,9306753 | -0,915827143 | 0,218498395 | -4,191459365 | 2,77E-05 | 0,000144448 |
| ENSG00000167232.14 | ZNF91 | protein_coding | 4171,335605 | 0,266323211 | 0,065212672 | 4,083918108 | 4,43E-05 | 0,000223975 |
| ENSG00000167244.21 | IGF2 | protein_coding | 128634,8351 | -0,711968898 | 0,04581573 | -15,5398354 | 1,86E-54 | 1,16E-52 |
| ENSG00000167258.14 | CDK12 | protein_coding | 13109,81651 | 0,31191923 | 0,049965335 | 6,242712731 | 4,30E-10 | 4,20E-09 |
| ENSG00000167272.10 | POP5 | protein_coding | 2022,713914 | -0,361243494 | 0,074509454 | -4,848290688 | 1,25E-06 | 7,99E-06 |
| ENSG00000167283.8 | ATP5MG | protein_coding | 24704,6703 | -0,411633841 | 0,059187373 | -6,954757726 | 3,53E-12 | 4,24E-11 |
| ENSG00000167291.16 | TBC1D16 | protein_coding | 8285,906945 | -0,50056709 | 0,051295265 | -9,758543848 | 1,70E-22 | 3,92E-21 |
| ENSG00000167306.20 | MYO5B | protein_coding | 6434,262699 | 0,248457364 | 0,054021541 | 4,599227621 | 4,24E-06 | 2,51E-05 |
| ENSG00000167315.18 | ACAA2 | protein_coding | 6828,903076 | -0,601581035 | 0,059042985 | -10,18886549 | 2,22E-24 | 5,62E-23 |
| ENSG00000167323.11 | STIM1 | protein_coding | 5264,161373 | 0,150135143 | 0,062626609 | 2,397305965 | 0,016516126 | 0,046107091 |
| ENSG00000167325.15 | RRM1 | protein_coding | 28862,06805 | -0,472144541 | 0,045233409 | -10,43796063 | 1,66E-25 | 4,41E-24 |

| | | | | | | | | |
|---|---|---|---|---|---|---|---|---|
| ENSG00000167333.13 | TRIM68 | protein_coding | 990,674685 | 0,416290624 | 0,093235936 | 4,464916024 | 8,01E-06 | 4,55E-05 |
| ENSG00000167363.14 | FN3K | protein_coding | 1299,148847 | -0,394643034 | 0,092787003 | -4,253214601 | 2,11E-05 | 0,000112132 |
| ENSG00000167371.21 | PRRT2 | protein_coding | 111,5733843 | -0,941014963 | 0,223061604 | -4,218632638 | 2,46E-05 | 0,000129596 |
| ENSG00000167378.8 | IRGQ | protein_coding | 4645,298076 | -0,504382372 | 0,175743696 | -2,869988411 | 0,004104868 | 0,013716818 |
| ENSG00000167393.17 | PPP2R3B | protein_coding | 1525,330867 | -0,452672794 | 0,088033821 | -5,142032783 | 2,72E-07 | 1,91E-06 |
| ENSG00000167395.10 | ZNF646 | protein_coding | 3570,374009 | 0,178663896 | 0,059400457 | 3,007786579 | 0,002631579 | 0,009222858 |
| ENSG00000167434.10 | CA4 | protein_coding | 31,82543765 | 3,983795256 | 0,671813039 | 5,929916545 | 3,03E-09 | 2,68E-08 |
| ENSG00000167460.17 | TPM4 | protein_coding | 12855,72539 | -0,317048338 | 0,045339288 | -6,992794879 | 2,69E-12 | 3,28E-11 |
| ENSG00000167470.12 | MIDN | protein_coding | 3585,417779 | -0,206732553 | 0,05742236 | -3,600209996 | 0,00031796 | 0,00137449 |
| ENSG00000167476.10 | JSRP1 | protein_coding | 3325,880502 | 0,314390453 | 0,05391135 | 5,831618968 | 5,49E-09 | 4,73E-08 |
| ENSG00000167515.10 | TRAPPC2L | protein_coding | 5712,248486 | -0,167015562 | 0,049946476 | -3,343890807 | 0,000826122 | 0,003262887 |
| ENSG00000167523.13 | SPATA33 | protein_coding | 2160,620827 | -0,458309 | 0,066643095 | -6,877066544 | 6,11E-12 | 7,17E-11 |
| ENSG00000167526.13 | RPL13 | protein_coding | 21013,68745 | -0,530509694 | 0,124188461 | -4,271811472 | 1,94E-05 | 0,000103714 |
| ENSG00000167536.14 | DHRS13 | protein_coding | 2652,932603 | -0,193358553 | 0,060474638 | -3,197349461 | 0,001386968 | 0,005208418 |
| ENSG00000167548.15 | KMT2D | protein_coding | 10007,0699 | 0,42139265 | 0,049316566 | 8,544646974 | 1,29E-17 | 2,29E-16 |
| ENSG00000167549.18 | CORO6 | protein_coding | 189,0260104 | -0,600167942 | 0,194309875 | -3,088715607 | 0,002010238 | 0,007256004 |
| ENSG00000167550.11 | RHEBL1 | protein_coding | 371,1758428 | 0,469985668 | 0,133410004 | 3,522866756 | 0,000426906 | 0,001796789 |
| ENSG00000167552.14 | TUBA1A | protein_coding | 132645,1638 | -0,244037967 | 0,044092453 | -5,534687971 | 3,12E-08 | 2,46E-07 |
| ENSG00000167553.15 | TUBA1C | protein_coding | 10013,3794 | -0,367642856 | 0,051358695 | -7,158337119 | 8,17E-13 | 1,04E-11 |
| ENSG00000167554.15 | ZNF610 | protein_coding | 95,15157699 | 0,703124734 | 0,273370179 | 2,572060844 | 0,010109512 | 0,030211293 |
| ENSG00000167562.12 | ZNF701 | protein_coding | 1268,206188 | 0,647346524 | 0,076100689 | 8,506447644 | 1,79E-17 | 3,16E-16 |
| ENSG00000167565.13 | SERTAD3 | protein_coding | 1002,666918 | 0,285198151 | 0,08746842 | 3,260584245 | 0,001111829 | 0,004265581 |
| ENSG00000167566.17 | NCKAP5L | protein_coding | 2118,762212 | 0,221289246 | 0,070627845 | 3,133172851 | 0,001729276 | 0,006352908 |
| ENSG00000167588.13 | GPD1 | protein_coding | 8999,454367 | 0,249281795 | 0,06700904 | 3,720121891 | 0,000199127 | 0,000893569 |
| ENSG00000167595.15 | PROSER3 | protein_coding | 1305,135154 | -0,327580247 | 0,091765678 | -3,569746932 | 0,000357326 | 0,001528227 |
| ENSG00000167608.11 | TMC4 | protein_coding | 2054,123189 | 0,219854065 | 0,069751771 | 3,151949562 | 0,001621843 | 0,005998367 |
| ENSG00000167619.12 | TMEM145 | protein_coding | 642,3114178 | -0,367848109 | 0,113775058 | -3,233117312 | 0,001224473 | 0,004648618 |
| ENSG00000167625.10 | ZNF526 | protein_coding | 1441,690421 | 0,24573378 | 0,076183621 | 3,225546079 | 0,001257326 | 0,0047629 |
| ENSG00000167632.16 | TRAPPC9 | protein_coding | 4556,045881 | -0,281821106 | 0,060123668 | -4,68735718 | 2,77E-06 | 1,69E-05 |
| ENSG00000167642.13 | SPINT2 | protein_coding | 46482,95838 | 0,321211599 | 0,039147492 | 8,205164145 | 2,30E-16 | 3,78E-15 |
| ENSG00000167658.16 | EEF2 | protein_coding | 99041,36196 | -0,635519649 | 0,200175945 | -3,174805292 | 0,001499371 | 0,005598762 |
| ENSG00000167671.12 | UBXN6 | protein_coding | 3384,152427 | -0,244940838 | 0,061882335 | -3,958170564 | 7,55E-05 | 0,000367095 |
| ENSG00000167700.9 | MFSD3 | protein_coding | 764,4202527 | -0,279418678 | 0,091881004 | -3,041093001 | 0,00235721 | 0,008353029 |
| ENSG00000167701.14 | GPT | protein_coding | 73,36860816 | -1,335029861 | 0,299223173 | -4,461652636 | 8,13E-06 | 4,61E-05 |
| ENSG00000167702.12 | KIFC2 | protein_coding | 3951,628475 | -0,192463296 | 0,076647976 | -2,511002975 | 0,012038868 | 0,035083232 |
| ENSG00000167716.18 | WDR81 | protein_coding | 1932,024099 | 0,427373813 | 0,080570675 | 5,304334507 | 1,13E-07 | 8,30E-07 |

| | | | | | | | | |
|---|---|---|---|---|---|---|---|---|
| ENSG00000167740.9 | CYB5D2 | protein_coding | 2909,082607 | -0,275704917 | 0,055428961 | -4,974022847 | 6,56E-07 | 4,38E-06 |
| ENSG00000167766.18 | ZNF83 | protein_coding | 862,1223759 | 0,776157335 | 0,102093483 | 7,602418014 | 2,91E-14 | 4,13E-13 |
| ENSG00000167767.14 | KRT80 | protein_coding | 1443,037336 | -0,311147612 | 0,077696783 | -4,004639562 | 6,21E-05 | 0,000306799 |
| ENSG00000167771.6 | RCOR2 | protein_coding | 3893,354255 | -0,42827211 | 0,053612279 | -7,988321358 | 1,37E-15 | 2,14E-14 |
| ENSG00000167775.11 | CD320 | protein_coding | 3666,87302 | 0,239700177 | 0,06339387 | 3,781125479 | 0,000156121 | 0,000715417 |
| ENSG00000167778.9 | SPRYD3 | protein_coding | 3172,544695 | -0,172928212 | 0,063727979 | -2,713536714 | 0,006656919 | 0,021009618 |
| ENSG00000167780.12 | SOAT2 | protein_coding | 484,2937706 | -0,399688263 | 0,119792234 | -3,336512303 | 0,000848367 | 0,003344288 |
| ENSG00000167792.12 | NDUFV1 | protein_coding | 5462,462695 | -0,264798171 | 0,074026876 | -3,577054491 | 0,000347488 | 0,001490113 |
| ENSG00000167815.12 | PRDX2 | protein_coding | 16008,88695 | -0,309673344 | 0,045423241 | -6,817508849 | 9,26E-12 | 1,07E-10 |
| ENSG00000167840.13 | ZNF232 | protein_coding | 1637,406012 | 0,222961169 | 0,079821379 | 2,793251283 | 0,005218114 | 0,016940252 |
| ENSG00000167863.12 | ATP5PD | protein_coding | 11283,80662 | -0,252179186 | 0,049236688 | -5,121773992 | 3,03E-07 | 2,11E-06 |
| ENSG00000167880.7 | EVPL | protein_coding | 15,61390424 | -3,279176982 | 0,747817067 | -4,384998856 | 1,16E-05 | 6,44E-05 |
| ENSG00000167920.9 | TMEM99 | protein_coding | 1560,441464 | -0,380434259 | 0,079993523 | -4,755813283 | 1,98E-06 | 1,23E-05 |
| ENSG00000167925.16 | GHDC | protein_coding | 2318,609198 | 0,190120716 | 0,062417032 | 3,045974952 | 0,002319272 | 0,008240645 |
| ENSG00000167930.17 | FAM234A | protein_coding | 12635,02351 | 0,175461543 | 0,043962646 | 3,99115064 | 6,58E-05 | 0,000323043 |
| ENSG00000167964.12 | RAB26 | protein_coding | 1765,454348 | -0,855717463 | 0,080992628 | -10,56537464 | 4,31E-26 | 1,17E-24 |
| ENSG00000167965.18 | MLST8 | protein_coding | 3843,316384 | -0,265068726 | 0,053096035 | -4,992250882 | 5,97E-07 | 4,00E-06 |
| ENSG00000167971.15 | CASKIN1 | protein_coding | 615,5288267 | -0,549490682 | 0,108427362 | -5,067823046 | 4,02E-07 | 2,76E-06 |
| ENSG00000167972.14 | ABCA3 | protein_coding | 9669,979528 | -0,202426485 | 0,050674666 | -3,994628913 | 6,48E-05 | 0,000318864 |
| ENSG00000167977.9 | KCTD5 | protein_coding | 7102,367365 | 0,189922483 | 0,053148703 | 3,573417061 | 0,000352353 | 0,001508676 |
| ENSG00000167981.7 | ZNF597 | protein_coding | 720,9388957 | 0,364812507 | 0,095113858 | 3,835534752 | 0,000125291 | 0,000586439 |
| ENSG00000167986.13 | DDB1 | protein_coding | 31151,06474 | -0,196709076 | 0,042359475 | -4,643803499 | 3,42E-06 | 2,05E-05 |
| ENSG00000167996.16 | FTH1 | protein_coding | 61917,11557 | -0,315423925 | 0,052541067 | -6,003378788 | 1,93E-09 | 1,75E-08 |
| ENSG00000168000.14 | BSCL2 | protein_coding | 8309,893235 | 0,114541859 | 0,044158615 | 2,593873466 | 0,009490141 | 0,028642206 |
| ENSG00000168002.12 | POLR2G | protein_coding | 6822,004153 | -0,155890549 | 0,048704504 | -3,20074195 | 0,001370742 | 0,005151785 |
| ENSG00000168010.11 | ATG16L2 | protein_coding | 2046,9423 | -0,471971469 | 0,072808228 | -6,4823919 | 9,03E-11 | 9,45E-10 |
| ENSG00000168014.16 | C2CD3 | protein_coding | 6087,145856 | 0,325143916 | 0,060915092 | 5,337657764 | 9,42E-08 | 6,95E-07 |
| ENSG00000168016.14 | TRANK1 | protein_coding | 23317,70121 | 6,189177573 | 0,209005298 | 29,61253919 | 1,03E-192 | 2,05E-190 |
| ENSG00000168026.18 | TTC21A | protein_coding | 345,7599802 | 0,509418055 | 0,191829563 | 2,655576367 | 0,0079173 | 0,02445742 |
| ENSG00000168028.14 | RPSA | protein_coding | 55631,07594 | -0,749502926 | 0,042392435 | -17,6801103 | 5,97E-70 | 4,81E-68 |
| ENSG00000168032.9 | ENTPD3 | protein_coding | 541,2428571 | -0,559367813 | 0,117283566 | -4,769362267 | 1,85E-06 | 1,15E-05 |
| ENSG00000168040.4 | FADD | protein_coding | 1932,444265 | 0,183576074 | 0,068729521 | 2,670993077 | 0,007562721 | 0,023470189 |
| ENSG00000168061.15 | SAC3D1 | protein_coding | 896,7966843 | -0,237812893 | 0,087642703 | -2,713436324 | 0,006658936 | 0,021013038 |
| ENSG00000168062.10 | BATF2 | protein_coding | 2396,087408 | 4,694264016 | 0,097023293 | 48,38285599 | 0 | 0 |
| ENSG00000168065.15 | SLC22A11 | protein_coding | 205,8498734 | 0,618031777 | 0,212547433 | 2,907735784 | 0,003640558 | 0,012331876 |
| ENSG00000168067.12 | MAP4K2 | protein_coding | 3276,63757 | -0,637202042 | 0,075713006 | -8,416018318 | 3,89E-17 | 6,70E-16 |

| | | | | | | | | |
|---|---|---|---|---|---|---|---|---|
| ENSG00000168071.22 | CCDC88B | protein_coding | 814,6888736 | -0,533703717 | 0,123799448 | -4,311034716 | 1,62E-05 | 8,80E-05 |
| ENSG00000168077.14 | SCARA3 | protein_coding | 427,5560203 | -0,779784856 | 0,123137311 | -6,332644834 | 2,41E-10 | 2,42E-09 |
| ENSG00000168078.10 | PBK | protein_coding | 7190,770614 | -0,532822732 | 0,05877627 | -9,065269585 | 1,24E-19 | 2,47E-18 |
| ENSG00000168090.10 | COPS6 | protein_coding | 11240,55943 | -0,221451159 | 0,045445644 | -4,872879756 | 1,10E-06 | 7,11E-06 |
| ENSG00000168092.14 | PAFAH1B2 | protein_coding | 12424,86746 | -0,254121805 | 0,041833282 | -6,074632328 | 1,24E-09 | 1,15E-08 |
| ENSG00000168096.14 | ANKS3 | protein_coding | 1445,429188 | 0,210184014 | 0,086002268 | 2,443935704 | 0,014528015 | 0,041273646 |
| ENSG00000168101.14 | NUDT16L1 | protein_coding | 1973,694323 | -0,268597196 | 0,070059256 | -3,833857375 | 0,000126149 | 0,000589841 |
| ENSG00000168118.11 | RAB4A | protein_coding | 3505,931264 | -0,507360779 | 0,086488572 | -5,866217551 | 4,46E-09 | 3,87E-08 |
| ENSG00000168137.16 | SETD5 | protein_coding | 15284,83912 | 0,435370411 | 0,051773932 | 8,409065966 | 4,13E-17 | 7,10E-16 |
| ENSG00000168143.9 | FAM83B | protein_coding | 1804,892223 | 1,017544182 | 0,075839895 | 13,41700402 | 4,81E-41 | 2,22E-39 |
| ENSG00000168152.13 | THAP9 | protein_coding | 1134,809887 | 0,287546625 | 0,098347023 | 2,923795915 | 0,003457914 | 0,011780563 |
| ENSG00000168209.5 | DDIT4 | protein_coding | 2326,946125 | 0,321501689 | 0,077947685 | 4,124582881 | 3,71E-05 | 0,000189957 |
| ENSG00000168214.20 | RBPJ | protein_coding | 10607,17357 | -0,180537496 | 0,053951796 | -3,346274086 | 0,000819054 | 0,003238378 |
| ENSG00000168216.13 | LMBRD1 | protein_coding | 12426,7638 | 0,108011304 | 0,042387388 | 2,548194398 | 0,010828211 | 0,032037893 |
| ENSG00000168243.11 | GNG4 | protein_coding | 13159,12334 | -0,791550564 | 0,057828053 | -13,6880031 | 1,20E-42 | 5,86E-41 |
| ENSG00000168256.18 | NKIRAS2 | protein_coding | 6346,322695 | 0,331115335 | 0,050692007 | 6,531904278 | 6,49E-11 | 6,88E-10 |
| ENSG00000168259.16 | DNAJC7 | protein_coding | 10057,49433 | 0,143476186 | 0,043073119 | 3,330991312 | 0,000865373 | 0,003401202 |
| ENSG00000168264.10 | IRF2BP2 | protein_coding | 15829,10741 | 0,235579459 | 0,041728071 | 5,645587051 | 1,65E-08 | 1,34E-07 |
| ENSG00000168268.11 | NT5DC2 | protein_coding | 17555,5356 | -1,606595021 | 0,050419057 | -31,86483686 | 8,20E-223 | 1,81E-220 |
| ENSG00000168280.17 | KIF5C | protein_coding | 22155,16964 | -0,797829394 | 0,054849708 | -14,54573647 | 6,22E-48 | 3,35E-46 |
| ENSG00000168297.15 | PXK | protein_coding | 6183,103036 | 1,211791415 | 0,069958526 | 17,32156869 | 3,23E-67 | 2,50E-65 |
| ENSG00000168301.13 | KCTD6 | protein_coding | 2428,727419 | 0,271352819 | 0,065294344 | 4,155839609 | 3,24E-05 | 0,000167318 |
| ENSG00000168303.8 | MPLKIP | protein_coding | 2369,975551 | -0,16334569 | 0,059065298 | -2,765510295 | 0,005683381 | 0,018283154 |
| ENSG00000168309.18 | FAM107A | protein_coding | 5,499127918 | 3,591247555 | 1,215293441 | 2,955045616 | 0,003126228 | 0,010741024 |
| ENSG00000168310.11 | IRF2 | protein_coding | 3727,295782 | 1,416080553 | 0,056202625 | 25,19598608 | 4,43E-140 | 6,83E-138 |
| ENSG00000168350.8 | DEGS2 | protein_coding | 431,4957628 | 0,873502669 | 0,12815446 | 6,816014593 | 9,36E-12 | 1,08E-10 |
| ENSG00000168385.18 | sept-02 | protein_coding | 62008,14595 | -0,284763609 | 0,042439096 | -6,709935724 | 1,95E-11 | 2,17E-10 |
| ENSG00000168386.18 | FILIP1L | protein_coding | 42,18591752 | 0,928799833 | 0,388657849 | 2,389762193 | 0,016859285 | 0,046931196 |
| ENSG00000168389.17 | MFSD2A | protein_coding | 379,2223167 | 0,476060495 | 0,132973573 | 3,580113586 | 0,000343445 | 0,001474743 |
| ENSG00000168393.13 | DTYMK | protein_coding | 2927,566418 | -0,546461806 | 0,063224006 | -8,643264424 | 5,46E-18 | 9,96E-17 |
| ENSG00000168394.11 | TAP1 | protein_coding | 27295,39177 | 7,929786504 | 0,347455153 | 22,8224749 | 2,74E-115 | 3,59E-113 |
| ENSG00000168398.6 | BDKRB2 | protein_coding | 327,0922019 | 1,082994575 | 0,186055137 | 5,820825981 | 5,86E-09 | 5,02E-08 |
| ENSG00000168404.13 | MLKL | protein_coding | 525,7453289 | 5,15831106 | 0,375108407 | 13,75152079 | 4,99E-43 | 2,45E-41 |
| ENSG00000168405.17 | CMAHP | transcribed_unitary | 12,2688135 | 2,863874574 | 0,748406217 | 3,826631191 | 0,000129909 | 0,000606161 |
| ENSG00000168477.19 | TNXB | protein_coding | 26,40003534 | -2,750938302 | 0,64068687 | -4,293732913 | 1,76E-05 | 9,45E-05 |
| ENSG00000168481.9 | LGI3 | protein_coding | 468,8595676 | 0,511174537 | 0,125227 | 4,081983427 | 4,47E-05 | 0,00022561 |

| | | | | | | | | |
|---|---|---|---|---|---|---|---|---|
| ENSG00000168488.18 | ATXN2L | protein_coding | 12398,44203 | 0,299042869 | 0,048354484 | 6,184387489 | 6,23E-10 | 5,98E-09 |
| ENSG00000168502.17 | MTCL1 | protein_coding | 1952,820213 | 0,302820733 | 0,07172913 | 4,221725995 | 2,42E-05 | 0,00012792 |
| ENSG00000168564.6 | CDKN2AIP | protein_coding | 5702,513903 | 0,254894169 | 0,050232701 | 5,074267602 | 3,89E-07 | 2,68E-06 |
| ENSG00000168566.13 | SNRNP48 | protein_coding | 10706,90466 | -0,13975527 | 0,046779522 | -2,987530943 | 0,002812408 | 0,009779777 |
| ENSG00000168591.16 | TMUB2 | protein_coding | 2949,118164 | 0,232643342 | 0,061174875 | 3,802923043 | 0,000142999 | 0,000661344 |
| ENSG00000168594.15 | ADAM29 | protein_coding | 26,62526219 | -1,524242389 | 0,49027231 | -3,10897099 | 0,001877402 | 0,006832406 |
| ENSG00000168610.14 | STAT3 | protein_coding | 12136,00122 | 0,629379494 | 0,044367146 | 14,18571069 | 1,12E-45 | 5,88E-44 |
| ENSG00000168653.11 | NDUFS5 | protein_coding | 12233,19318 | -0,363129802 | 0,047183944 | -7,696045925 | 1,40E-14 | 2,04E-13 |
| ENSG00000168658.18 | VWA3B | protein_coding | 103,4418951 | 1,354467604 | 0,352472649 | 3,842759457 | 0,000121659 | 0,000570979 |
| ENSG00000168661.14 | ZNF30 | protein_coding | 1158,399569 | 0,29789447 | 0,085110901 | 3,500074214 | 0,000465129 | 0,001944212 |
| ENSG00000168671.10 | UGT3A2 | protein_coding | 63,72227057 | 0,950681166 | 0,369910507 | 2,57003018 | 0,010168966 | 0,030353169 |
| ENSG00000168672.4 | LRATD2 | protein_coding | 7139,929827 | 0,82103363 | 0,055262662 | 14,85693232 | 6,27E-50 | 3,60E-48 |
| ENSG00000168701.19 | TMEM208 | protein_coding | 4897,740922 | -0,201439923 | 0,051896113 | -3,881599474 | 0,000103772 | 0,000492784 |
| ENSG00000168702.17 | LRP1B | protein_coding | 11,24892092 | 1,857817808 | 0,745398356 | 2,492382486 | 0,012688931 | 0,036730344 |
| ENSG00000168743.12 | NPNT | protein_coding | 2234,207563 | -0,989575745 | 0,091267424 | -10,84259531 | 2,16E-27 | 6,14E-26 |
| ENSG00000168763.16 | CNNM3 | protein_coding | 8146,032474 | -0,2722376 | 0,050929059 | -5,345427657 | 9,02E-08 | 6,68E-07 |
| ENSG00000168769.13 | TET2 | protein_coding | 8869,322431 | 0,333567859 | 0,043109686 | 7,737654619 | 1,01E-14 | 1,49E-13 |
| ENSG00000168772.11 | CXXC4 | protein_coding | 5988,299699 | -0,15303063 | 0,059311991 | -2,580096013 | 0,009877285 | 0,029612121 |
| ENSG00000168779.19 | SHOX2 | protein_coding | 1597,854957 | 1,484254318 | 0,081139446 | 18,29263562 | 9,47E-75 | 8,23E-73 |
| ENSG00000168781.22 | PPIP5K1 | protein_coding | 7681,814697 | -0,191825687 | 0,078678045 | -2,438109464 | 0,014764302 | 0,041860396 |
| ENSG00000168792.5 | ABHD15 | protein_coding | 11126,28074 | 0,171493262 | 0,044466531 | 3,856681815 | 0,000114937 | 0,000541123 |
| ENSG00000168807.16 | SNTB2 | protein_coding | 4438,625774 | -0,182544111 | 0,056124938 | -3,252459878 | 0,001144107 | 0,004373763 |
| ENSG00000168813.17 | ZNF507 | protein_coding | 8282,821283 | 0,153354732 | 0,049384634 | 3,105312742 | 0,001900779 | 0,006907433 |
| ENSG00000168818.10 | STX18 | protein_coding | 2924,548147 | 0,152713562 | 0,059293315 | 2,575561228 | 0,010007755 | 0,029943449 |
| ENSG00000168824.14 | NSG1 | protein_coding | 76,72487135 | -0,921631765 | 0,295244316 | -3,121590203 | 0,001798771 | 0,006583503 |
| ENSG00000168827.14 | GFM1 | protein_coding | 9679,375216 | -0,151594642 | 0,049946644 | -3,035131662 | 0,002404307 | 0,008503857 |
| ENSG00000168876.9 | ANKRD49 | protein_coding | 3535,613942 | 0,191910531 | 0,057141109 | 3,358537058 | 0,000783562 | 0,003110069 |
| ENSG00000168899.5 | VAMP5 | protein_coding | 623,7049178 | 1,525755106 | 0,118249014 | 12,90289918 | 4,33E-38 | 1,83E-36 |
| ENSG00000168906.13 | MAT2A | protein_coding | 25053,13669 | -0,141458949 | 0,045642939 | -3,099251508 | 0,001940102 | 0,007033301 |
| ENSG00000168913.7 | ENHO | protein_coding | 1579,426524 | -0,560422216 | 0,071408868 | -7,848075856 | 4,22E-15 | 6,41E-14 |
| ENSG00000168916.15 | ZNF608 | protein_coding | 9264,879868 | 0,159950667 | 0,05892153 | 2,714638701 | 0,006634811 | 0,02094278 |
| ENSG00000168917.9 | SLC35G2 | protein_coding | 4681,879987 | 0,176803503 | 0,051798746 | 3,413277696 | 0,000641865 | 0,002600324 |
| ENSG00000168918.14 | INPP5D | protein_coding | 85,128323 | 2,379899472 | 0,301341996 | 7,89766945 | 2,84E-15 | 4,35E-14 |
| ENSG00000168938.5 | PPIC | protein_coding | 1884,39814 | 1,286348568 | 0,082556111 | 15,58150642 | 9,72E-55 | 6,15E-53 |
| ENSG00000168952.15 | STXBP6 | protein_coding | 1617,418465 | 0,245831374 | 0,082002556 | 2,997850131 | 0,002718913 | 0,009489847 |
| ENSG00000168958.19 | MFF | protein_coding | 6232,639642 | -0,149774578 | 0,048747118 | -3,072480681 | 0,002122876 | 0,007617378 |

| | | | | | | | | |
|---|---|---|---|---|---|---|---|---|
| ENSG00000168993.15 | CPLX1 | protein_coding | 1179,892896 | -0,580134276 | 0,075545386 | -7,679281357 | 1,60E-14 | 2,31E-13 |
| ENSG00000168994.13 | PXDC1 | protein_coding | 572,5197414 | 1,083713821 | 0,101928068 | 10,63214329 | 2,11E-26 | 5,80E-25 |
| ENSG00000169018.6 | FEM1B | protein_coding | 17018,56903 | 0,115562522 | 0,046174598 | 2,502729344 | 0,012323975 | 0,035825953 |
| ENSG00000169019.11 | COMMD8 | protein_coding | 4767,75861 | -0,374874853 | 0,063412164 | -5,911718363 | 3,39E-09 | 2,98E-08 |
| ENSG00000169020.10 | ATP5ME | protein_coding | 10440,75526 | -0,354750226 | 0,044489365 | -7,973820818 | 1,54E-15 | 2,39E-14 |
| ENSG00000169021.6 | UQCRFS1 | protein_coding | 13600,3833 | -0,228915861 | 0,054956473 | -4,165403025 | 3,11E-05 | 0,000160965 |
| ENSG00000169047.5 | IRS1 | protein_coding | 2988,405607 | 0,75148751 | 0,068977571 | 10,89466469 | 1,22E-27 | 3,51E-26 |
| ENSG00000169062.15 | UPF3A | protein_coding | 8216,465451 | -0,183981303 | 0,043823945 | -4,198191253 | 2,69E-05 | 0,000140547 |
| ENSG00000169100.14 | SLC25A6 | protein_coding | 25535,02824 | -0,829933827 | 0,048567277 | -17,08833355 | 1,81E-65 | 1,37E-63 |
| ENSG00000169105.7 | CHST14 | protein_coding | 1128,255262 | -0,227841152 | 0,081054558 | -2,810960381 | 0,004939387 | 0,016145898 |
| ENSG00000169118.18 | CSNK1G1 | protein_coding | 5025,924479 | 0,540241961 | 0,053634299 | 10,07269554 | 7,30E-24 | 1,80E-22 |
| ENSG00000169126.15 | ARMC4 | protein_coding | 26,84213541 | 1,190578805 | 0,485908066 | 2,450214119 | 0,014277129 | 0,04071505 |
| ENSG00000169136.11 | ATF5 | protein_coding | 7982,141095 | 1,125702891 | 0,074347805 | 15,14103737 | 8,68E-52 | 5,23E-50 |
| ENSG00000169139.12 | UBE2V2 | protein_coding | 13372,12165 | -0,243272843 | 0,045260065 | -5,374999839 | 7,66E-08 | 5,72E-07 |
| ENSG00000169155.10 | ZBTB43 | protein_coding | 3281,405798 | 0,344341262 | 0,055259735 | 6,2313231 | 4,63E-10 | 4,51E-09 |
| ENSG00000169169.14 | CPT1C | protein_coding | 4006,932663 | 0,237912739 | 0,060082448 | 3,959771057 | 7,50E-05 | 0,000365113 |
| ENSG00000169174.10 | PCSK9 | protein_coding | 13491,82686 | 0,230565997 | 0,051026864 | 4,518521829 | 6,23E-06 | 3,59E-05 |
| ENSG00000169189.17 | NSMCE1 | protein_coding | 2000,096703 | -0,168618597 | 0,06277291 | -2,686168233 | 0,007227669 | 0,022570421 |
| ENSG00000169193.12 | CCDC126 | protein_coding | 3077,489918 | -0,255187321 | 0,054614868 | -4,672488134 | 2,98E-06 | 1,81E-05 |
| ENSG00000169213.7 | RAB3B | protein_coding | 49811,32647 | -1,129524736 | 0,118598302 | -9,523953724 | 1,67E-21 | 3,66E-20 |
| ENSG00000169217.9 | CD2BP2 | protein_coding | 8729,443183 | 0,270742057 | 0,045017754 | 6,014117384 | 1,81E-09 | 1,64E-08 |
| ENSG00000169220.18 | RGS14 | protein_coding | 446,3206578 | 0,389483366 | 0,120513873 | 3,231855028 | 0,001229894 | 0,004667626 |
| ENSG00000169223.15 | LMAN2 | protein_coding | 12511,38511 | 0,24076849 | 0,052299 | 4,603692068 | 4,15E-06 | 2,46E-05 |
| ENSG00000169228.14 | RAB24 | protein_coding | 2923,280666 | 0,621928601 | 0,072370881 | 8,593630425 | 8,43E-18 | 1,52E-16 |
| ENSG00000169239.13 | CA5B | protein_coding | 843,8825461 | 0,258389119 | 0,100782955 | 2,563817636 | 0,010352792 | 0,03082013 |
| ENSG00000169241.19 | SLC50A1 | protein_coding | 5426,016579 | 0,572303322 | 0,05363663 | 10,67000895 | 1,41E-26 | 3,90E-25 |
| ENSG00000169245.6 | CXCL10 | protein_coding | 4967,602979 | 9,351347889 | 0,33247943 | 28,12609453 | 4,70E-174 | 8,53E-172 |
| ENSG00000169246.16 | NPIPB3 | protein_coding | 1372,178323 | 0,404594437 | 0,094192548 | 4,295397509 | 1,74E-05 | 9,39E-05 |
| ENSG00000169248.12 | CXCL11 | protein_coding | 621,8134226 | 2,964595924 | 0,451319509 | 6,568729838 | 5,07E-11 | 5,43E-10 |
| ENSG00000169249.12 | ZRSR2 | protein_coding | 1877,206772 | 0,589643425 | 0,069577725 | 8,474600565 | 2,36E-17 | 4,12E-16 |
| ENSG00000169255.15 | B3GALNT1 | protein_coding | 4031,562101 | 0,613926776 | 0,058498625 | 10,49472156 | 9,13E-26 | 2,45E-24 |
| ENSG00000169297.7 | NR0B1 | protein_coding | 3364,499527 | 0,309033564 | 0,061030561 | 5,063587144 | 4,11E-07 | 2,82E-06 |
| ENSG00000169359.16 | SLC33A1 | protein_coding | 6756,973214 | 0,227660747 | 0,051570103 | 4,414587765 | 1,01E-05 | 5,67E-05 |
| ENSG00000169372.13 | CRADD | protein_coding | 1459,44747 | -0,23881219 | 0,073924956 | -3,230467792 | 0,001235878 | 0,004687175 |
| ENSG00000169410.9 | PTPN9 | protein_coding | 5339,977044 | 0,150319219 | 0,051478156 | 2,920058338 | 0,003499659 | 0,011904763 |
| ENSG00000169418.10 | NPR1 | protein_coding | 1325,147298 | -0,456417044 | 0,080781065 | -5,650049826 | 1,60E-08 | 1,30E-07 |

| | | | | | | | | |
|---|---|---|---|---|---|---|---|---|
| ENSG00000169429.11 | CXCL8 | protein_coding | 331,0515639 | 5,150072559 | 0,212953136 | 24,18406537 | 3,27E-129 | 4,85E-127 |
| ENSG00000169435.14 | RASSF6 | protein_coding | 20393,99583 | 0,539086333 | 0,042069939 | 12,81405067 | 1,37E-37 | 5,68E-36 |
| ENSG00000169436.17 | COL22A1 | protein_coding | 4270,029581 | 0,20699004 | 0,076385964 | 2,709791552 | 0,006732551 | 0,021218563 |
| ENSG00000169442.9 | CD52 | protein_coding | 8,121900752 | -3,19408711 | 1,127241451 | -2,833542989 | 0,004603512 | 0,015152948 |
| ENSG00000169446.5 | MMGT1 | protein_coding | 3882,367542 | -0,341588416 | 0,054202636 | -6,302062823 | 2,94E-10 | 2,92E-09 |
| ENSG00000169490.17 | TM2D2 | protein_coding | 5186,067408 | 0,293198618 | 0,053336452 | 5,497152604 | 3,86E-08 | 3,00E-07 |
| ENSG00000169499.15 | PLEKHA2 | protein_coding | 384,4895345 | 0,7427627 | 0,137576765 | 5,398896402 | 6,71E-08 | 5,05E-07 |
| ENSG00000169515.8 | CCDC8 | protein_coding | 598,7696159 | -0,370418366 | 0,104269717 | -3,552501866 | 0,000381586 | 0,001623346 |
| ENSG00000169567.11 | HINT1 | protein_coding | 23123,60929 | -0,433181993 | 0,058721623 | -7,37687361 | 1,62E-13 | 2,19E-12 |
| ENSG00000169570.10 | DTWD2 | protein_coding | 1223,491503 | -0,307149719 | 0,088148539 | -3,48445614 | 0,000493138 | 0,002046469 |
| ENSG00000169607.13 | CKAP2L | protein_coding | 7382,892493 | -0,544579766 | 0,055352648 | -9,838368882 | 7,69E-23 | 1,81E-21 |
| ENSG00000169641.13 | LUZP1 | protein_coding | 3724,094033 | 0,292216862 | 0,058376632 | 5,005716343 | 5,57E-07 | 3,75E-06 |
| ENSG00000169660.16 | HEXD | protein_coding | 2401,846499 | 0,574514742 | 0,071048473 | 8,08623626 | 6,15E-16 | 9,84E-15 |
| ENSG00000169679.15 | BUB1 | protein_coding | 8217,606518 | -0,554270992 | 0,051229798 | -10,81930846 | 2,79E-27 | 7,90E-26 |
| ENSG00000169682.18 | SPNS1 | protein_coding | 3216,749631 | -0,18018267 | 0,062513201 | -2,882313922 | 0,003947662 | 0,013250511 |
| ENSG00000169683.8 | LRRC45 | protein_coding | 2317,454336 | -0,673929447 | 0,07137479 | -9,44212161 | 3,65E-21 | 7,89E-20 |
| ENSG00000169689.15 | CENPX | protein_coding | 3659,956577 | -0,72595082 | 0,052054661 | -13,94593321 | 3,33E-44 | 1,68E-42 |
| ENSG00000169692.13 | AGPAT2 | protein_coding | 6678,859678 | -0,225375265 | 0,060126086 | -3,74837744 | 0,000177982 | 0,000805753 |
| ENSG00000169715.14 | MT1E | protein_coding | 349,5293606 | 3,512059234 | 0,181545092 | 19,34538247 | 2,23E-83 | 2,14E-81 |
| ENSG00000169717.6 | ACTRT2 | protein_coding | 122,2705969 | -1,621329515 | 0,228599193 | -7,092455123 | 1,32E-12 | 1,65E-11 |
| ENSG00000169718.17 | DUS1L | protein_coding | 5407,706318 | -0,259128111 | 0,0688827 | -3,761875078 | 0,000168644 | 0,000767337 |
| ENSG00000169727.12 | GPS1 | protein_coding | 5992,772387 | -0,237229544 | 0,047101541 | -5,036555911 | 4,74E-07 | 3,23E-06 |
| ENSG00000169750.9 | RAC3 | protein_coding | 1227,045903 | -0,49060271 | 0,08339774 | -5,882685896 | 4,04E-09 | 3,53E-08 |
| ENSG00000169756.16 | LIMS1 | protein_coding | 6409,619107 | -0,113640588 | 0,046947512 | -2,420588089 | 0,015495424 | 0,043609216 |
| ENSG00000169764.15 | UGP2 | protein_coding | 12820,87505 | 0,111553323 | 0,046986411 | 2,374161396 | 0,01758886 | 0,04863133 |
| ENSG00000169783.12 | LINGO1 | protein_coding | 2151,613002 | -0,538934605 | 0,06958742 | -7,744713165 | 9,58E-15 | 1,41E-13 |
| ENSG00000169813.16 | HNRNPF | protein_coding | 31988,52181 | -0,103993633 | 0,041097679 | -2,530401624 | 0,011393203 | 0,033413629 |
| ENSG00000169862.19 | CTNND2 | protein_coding | 4051,179324 | -0,194617274 | 0,0559903 | -3,475910556 | 0,000509122 | 0,002103879 |
| ENSG00000169871.13 | TRIM56 | protein_coding | 15381,80758 | 1,901194426 | 0,044729088 | 42,50465485 | 0 | 0 |
| ENSG00000169891.18 | REPS2 | protein_coding | 1517,352454 | -1,071477592 | 0,104433879 | -10,25986591 | 1,07E-24 | 2,75E-23 |
| ENSG00000169895.6 | SYAP1 | protein_coding | 15420,90888 | -0,336397991 | 0,045865011 | -7,334523275 | 2,23E-13 | 2,97E-12 |
| ENSG00000169903.6 | TM4SF4 | protein_coding | 4293,147008 | 0,137183336 | 0,054767299 | 2,504840258 | 0,012250671 | 0,035631264 |
| ENSG00000169905.12 | TOR1AIP2 | protein_coding | 9664,632829 | 0,265692552 | 0,048241709 | 5,5075278 | 3,64E-08 | 2,84E-07 |
| ENSG00000169914.6 | OTUD3 | protein_coding | 2610,262886 | -0,866713038 | 0,0844998 | -10,25698336 | 1,10E-24 | 2,83E-23 |
| ENSG00000169919.17 | GUSB | protein_coding | 14782,56212 | 0,141308078 | 0,042706753 | 3,308799367 | 0,00093697 | 0,003655789 |
| ENSG00000169925.16 | BRD3 | protein_coding | 9958,874655 | -0,270537495 | 0,045211533 | -5,983816028 | 2,18E-09 | 1,96E-08 |

| | | | | | | | | |
|---|---|---|---|---|---|---|---|---|
| ENSG00000169926.11 | KLF13 | protein_coding | 8009,347106 | -0,899224824 | 0,07541665 | -11,92342573 | 8,94E-33 | 3,22E-31 |
| ENSG00000169933.14 | FRMPD4 | protein_coding | 3805,686663 | 0,208842176 | 0,051731719 | 4,037023683 | 5,41E-05 | 0,000269877 |
| ENSG00000169955.7 | ZNF747 | protein_coding | 1967,709997 | 0,231187347 | 0,063444561 | 3,643926984 | 0,00026851 | 0,001176539 |
| ENSG00000169962.5 | TAS1R3 | protein_coding | 370,423434 | -0,842703167 | 0,132218446 | -6,373567327 | 1,85E-10 | 1,87E-09 |
| ENSG00000169976.6 | SF3B5 | protein_coding | 18519,65753 | 0,140688747 | 0,050567745 | 2,782183531 | 0,00539945 | 0,017464609 |
| ENSG00000169992.10 | NLGN2 | protein_coding | 4395,746119 | 0,620493024 | 0,075826595 | 8,183052707 | 2,77E-16 | 4,51E-15 |
| ENSG00000169994.18 | MYO7B | protein_coding | 3122,404346 | -0,172104577 | 0,062069044 | -2,772792467 | 0,005557755 | 0,017922566 |
| ENSG00000170006.12 | TMEM154 | protein_coding | 58,77101106 | 1,542819154 | 0,378947522 | 4,071326677 | 4,67E-05 | 0,000235446 |
| ENSG00000170017.12 | ALCAM | protein_coding | 4196,724992 | 0,21579146 | 0,053563198 | 4,028726223 | 5,61E-05 | 0,0002789 |
| ENSG00000170035.16 | UBE2E3 | protein_coding | 10174,35938 | -0,269151906 | 0,046856885 | -5,74412723 | 9,24E-09 | 7,70E-08 |
| ENSG00000170037.13 | CNTROB | protein_coding | 4105,734708 | -0,365876012 | 0,061930292 | -5,907868315 | 3,47E-09 | 3,05E-08 |
| ENSG00000170043.11 | TRAPPC1 | protein_coding | 6744,3817 | -0,25105437 | 0,046608152 | -5,386490582 | 7,18E-08 | 5,38E-07 |
| ENSG00000170044.8 | ZPLD1 | protein_coding | 2156,993648 | -0,778060883 | 0,063784322 | -12,19830917 | 3,17E-34 | 1,18E-32 |
| ENSG00000170075.9 | GPR37L1 | protein_coding | 15,43407383 | 1,683085358 | 0,623064044 | 2,701303942 | 0,006906819 | 0,02171307 |
| ENSG00000170091.11 | NSG2 | protein_coding | 455,0065402 | -0,355147994 | 0,142819633 | -2,486688885 | 0,01289381 | 0,037218104 |
| ENSG00000170099.6 | SERPINA6 | protein_coding | 4811,838355 | 0,333274409 | 0,054111115 | 6,159074854 | 7,32E-10 | 6,95E-09 |
| ENSG00000170100.13 | ZNF778 | protein_coding | 2024,43639 | -0,854360006 | 0,099333206 | -8,600950708 | 7,91E-18 | 1,42E-16 |
| ENSG00000170142.11 | UBE2E1 | protein_coding | 2763,484609 | 0,185903407 | 0,06075003 | 3,060136871 | 0,002212359 | 0,007898173 |
| ENSG00000170145.5 | SIK2 | protein_coding | 10509,97382 | -0,416132025 | 0,0447919 | -9,290340956 | 1,54E-20 | 3,23E-19 |
| ENSG00000170175.11 | CHRNB1 | protein_coding | 1200,757926 | 0,752255585 | 0,07610992 | 9,883804644 | 4,89E-23 | 1,16E-21 |
| ENSG00000170185.9 | USP38 | protein_coding | 9166,510047 | 0,24582193 | 0,054267274 | 4,529837472 | 5,90E-06 | 3,42E-05 |
| ENSG00000170190.16 | SLC16A5 | protein_coding | 23,17792278 | 1,399641864 | 0,541462177 | 2,58493007 | 0,009739875 | 0,029257038 |
| ENSG00000170222.11 | ADPRM | protein_coding | 589,4884049 | 0,457637916 | 0,11651613 | 3,927678646 | 8,58E-05 | 0,000414023 |
| ENSG00000170234.13 | PWWP2A | protein_coding | 4191,001775 | 0,169076798 | 0,055572898 | 3,042432612 | 0,002346744 | 0,008322491 |
| ENSG00000170260.8 | ZNF212 | protein_coding | 1464,273954 | 0,240231523 | 0,075317755 | 3,18957361 | 0,001424828 | 0,005332801 |
| ENSG00000170264.13 | FAM161A | protein_coding | 3335,993899 | -0,170235294 | 0,065561561 | -2,596571682 | 0,009415926 | 0,028452553 |
| ENSG00000170265.12 | ZNF282 | protein_coding | 3269,391546 | 0,265677387 | 0,055231521 | 4,810249333 | 1,51E-06 | 9,54E-06 |
| ENSG00000170266.16 | GLB1 | protein_coding | 16849,87255 | 0,230099948 | 0,049529623 | 4,645703578 | 3,39E-06 | 2,04E-05 |
| ENSG00000170270.5 | GON7 | protein_coding | 2253,425691 | -0,25821441 | 0,060825709 | -4,245152514 | 2,18E-05 | 0,000115994 |
| ENSG00000170291.15 | ELP5 | protein_coding | 4648,13445 | -0,245084342 | 0,05031526 | -4,870974356 | 1,11E-06 | 7,17E-06 |
| ENSG00000170296.10 | GABARAP | protein_coding | 25476,94548 | -0,255649469 | 0,045210691 | -5,654624289 | 1,56E-08 | 1,27E-07 |
| ENSG00000170312.15 | CDK1 | protein_coding | 20850,28231 | -0,550461577 | 0,047080797 | -11,69184921 | 1,40E-31 | 4,80E-30 |
| ENSG00000170315.13 | UBB | protein_coding | 102828,5575 | 0,112560874 | 0,037776179 | 2,979678678 | 0,002885509 | 0,010009234 |
| ENSG00000170322.14 | NFRKB | protein_coding | 4122,299621 | 0,194856426 | 0,055868843 | 3,487747627 | 0,000487108 | 0,002024055 |
| ENSG00000170325.14 | PRDM10 | protein_coding | 2353,133763 | -0,18605773 | 0,072339138 | -2,572020286 | 0,010110697 | 0,030211293 |
| ENSG00000170340.11 | B3GNT2 | protein_coding | 2836,543874 | 0,282769228 | 0,056704892 | 4,986681311 | 6,14E-07 | 4,11E-06 |

| | | | | | | | | |
|---|---|---|---|---|---|---|---|---|
| ENSG00000170348.9 | TMED10 | protein_coding | 39397,15418 | 0,400589256 | 0,038367499 | 10,44084873 | 1,61E-25 | 4,28E-24 |
| ENSG00000170356.9 | OR2A20P | transcribed_unproc | 215,0262363 | 0,807595775 | 0,169798742 | 4,756194088 | 1,97E-06 | 1,23E-05 |
| ENSG00000170365.10 | SMAD1 | protein_coding | 665,3508301 | 0,648648534 | 0,119886205 | 5,410535203 | 6,28E-08 | 4,76E-07 |
| ENSG00000170379.20 | TCAF2 | protein_coding | 798,5914221 | 1,086482157 | 0,099042239 | 10,96988691 | 5,33E-28 | 1,55E-26 |
| ENSG00000170385.10 | SLC30A1 | protein_coding | 7909,99216 | 0,874765945 | 0,05511295 | 15,87223951 | 9,87E-57 | 6,51E-55 |
| ENSG00000170390.16 | DCLK2 | protein_coding | 1102,248771 | -0,251840601 | 0,082506479 | -3,052373641 | 0,002270393 | 0,008082283 |
| ENSG00000170396.8 | ZNF804A | protein_coding | 30,89597553 | 1,976815933 | 0,446254286 | 4,429797077 | 9,43E-06 | 5,31E-05 |
| ENSG00000170421.12 | KRT8 | protein_coding | 32041,02029 | 0,371030746 | 0,047246136 | 7,853144759 | 4,06E-15 | 6,16E-14 |
| ENSG00000170448.12 | NFXL1 | protein_coding | 3551,466426 | 0,496199624 | 0,055683026 | 8,911146956 | 5,05E-19 | 9,76E-18 |
| ENSG00000170456.16 | DENND5B | protein_coding | 4815,839466 | 0,148707986 | 0,051225327 | 2,903016813 | 0,003695868 | 0,012491035 |
| ENSG00000170468.7 | RIOX1 | protein_coding | 1367,529256 | 0,21856687 | 0,076833234 | 2,844691796 | 0,004445441 | 0,014691299 |
| ENSG00000170473.17 | PYM1 | protein_coding | 1590,644926 | -0,300993423 | 0,068915271 | -4,367586748 | 1,26E-05 | 6,93E-05 |
| ENSG00000170500.12 | LONRF2 | protein_coding | 5600,87937 | -1,280478812 | 0,066820573 | -19,16294268 | 7,55E-82 | 7,08E-80 |
| ENSG00000170515.14 | PA2G4 | protein_coding | 15554,06209 | -0,20413621 | 0,043395218 | -4,704117658 | 2,55E-06 | 1,56E-05 |
| ENSG00000170522.9 | ELOVL6 | protein_coding | 5653,483855 | 0,187127897 | 0,061986468 | 3,018850763 | 0,002537355 | 0,008928141 |
| ENSG00000170540.15 | ARL6IP1 | protein_coding | 65242,22818 | -0,410661848 | 0,04513774 | -9,097971063 | 9,20E-20 | 1,84E-18 |
| ENSG00000170542.6 | SERPINB9 | protein_coding | 2136,257737 | 1,223273291 | 0,068445409 | 17,87224761 | 1,94E-71 | 1,58E-69 |
| ENSG00000170545.17 | SMAGP | protein_coding | 296,0745578 | 0,379129301 | 0,154798812 | 2,449174492 | 0,014318406 | 0,04082242 |
| ENSG00000170571.12 | EMB | protein_coding | 14951,99427 | 0,341358006 | 0,044286506 | 7,707946229 | 1,28E-14 | 1,86E-13 |
| ENSG00000170581.14 | STAT2 | protein_coding | 19724,22372 | 2,881402598 | 0,067074412 | 42,95829829 | 0 | 0 |
| ENSG00000170624.13 | SGCD | protein_coding | 8575,164308 | -0,572460923 | 0,070595041 | -8,109081189 | 5,10E-16 | 8,21E-15 |
| ENSG00000170629.14 | DPY19L2P2 | transcribed_unproc | 4297,198516 | -0,272283815 | 0,054097373 | -5,033216912 | 4,82E-07 | 3,28E-06 |
| ENSG00000170632.14 | ARMC10 | protein_coding | 3483,196032 | 0,148226389 | 0,057012168 | 2,59990796 | 0,009324877 | 0,028213517 |
| ENSG00000170633.16 | RNF34 | protein_coding | 6545,343692 | 0,171137419 | 0,058195426 | 2,940736612 | 0,003274328 | 0,011214524 |
| ENSG00000170634.13 | ACYP2 | protein_coding | 1294,047733 | -0,230048231 | 0,080239516 | -2,867019189 | 0,004143579 | 0,013827697 |
| ENSG00000170667.14 | RASA4B | protein_coding | 7871,950332 | -0,260513526 | 0,058022644 | -4,489859642 | 7,13E-06 | 4,08E-05 |
| ENSG00000170743.17 | SYT9 | protein_coding | 834,2647791 | 0,394463086 | 0,088996624 | 4,432337627 | 9,32E-06 | 5,25E-05 |
| ENSG00000170759.11 | KIF5B | protein_coding | 21430,07864 | -0,426942649 | 0,048201343 | -8,857484444 | 8,18E-19 | 1,56E-17 |
| ENSG00000170775.2 | GPR37 | protein_coding | 1064,839646 | 1,037243352 | 0,080997638 | 12,80584694 | 1,52E-37 | 6,30E-36 |
| ENSG00000170776.21 | AKAP13 | protein_coding | 10326,08254 | 0,14931778 | 0,055040822 | 2,712855197 | 0,006670625 | 0,021044022 |
| ENSG00000170779.10 | CDCA4 | protein_coding | 1843,82408 | -0,323816251 | 0,064271661 | -5,03824307 | 4,70E-07 | 3,20E-06 |
| ENSG00000170802.16 | FOXN2 | protein_coding | 4369,35139 | 0,134633984 | 0,055165559 | 2,440544162 | 0,014665153 | 0,04161073 |
| ENSG00000170832.13 | USP32 | protein_coding | 7913,346892 | 0,304157598 | 0,049871395 | 6,098838774 | 1,07E-09 | 9,95E-09 |
| ENSG00000170837.2 | GPR27 | protein_coding | 2645,485982 | -0,46190077 | 0,087469016 | -5,280735862 | 1,29E-07 | 9,38E-07 |
| ENSG00000170846.17 | AC093323.1 | protein_coding | 5962,146061 | 0,866178024 | 0,085468835 | 10,13443117 | 3,89E-24 | 9,70E-23 |
| ENSG00000170852.11 | KBTBD2 | protein_coding | 5863,205312 | 0,194323165 | 0,056440447 | 3,442977083 | 0,000575348 | 0,00235608 |

| | | | | | | | | |
|---|---|---|---|---|---|---|---|---|
| ENSG00000170855.4 | TRIAP1 | protein_coding | 3625,385187 | -0,223718389 | 0,056690714 | -3,946296873 | 7,94E-05 | 0,000384777 |
| ENSG00000170860.4 | LSM3 | protein_coding | 8384,225418 | -0,358683318 | 0,051472537 | -6,968440673 | 3,20E-12 | 3,86E-11 |
| ENSG00000170871.12 | KIAA0232 | protein_coding | 13409,47875 | -0,302507304 | 0,051722408 | -5,848670176 | 4,96E-09 | 4,28E-08 |
| ENSG00000170873.18 | MTSS1 | protein_coding | 15971,3182 | 0,357529426 | 0,04587064 | 7,794297832 | 6,48E-15 | 9,71E-14 |
| ENSG00000170876.7 | TMEM43 | protein_coding | 3606,956842 | 0,146552127 | 0,05360783 | 2,733782142 | 0,006261146 | 0,019908479 |
| ENSG00000170881.4 | RNF139 | protein_coding | 5971,871274 | 0,142332241 | 0,055392656 | 2,569514636 | 0,010184109 | 0,03039434 |
| ENSG00000170889.14 | RPS9 | protein_coding | 34189,85603 | -0,360604472 | 0,047069417 | -7,661120494 | 1,84E-14 | 2,66E-13 |
| ENSG00000170899.11 | GSTA4 | protein_coding | 7702,840113 | -0,183296112 | 0,050401836 | -3,636695157 | 0,000276158 | 0,001206297 |
| ENSG00000170903.11 | MSANTD4 | protein_coding | 7963,214827 | 0,210875877 | 0,059090419 | 3,568698303 | 0,000358759 | 0,001533772 |
| ENSG00000170917.13 | NUDT6 | protein_coding | 295,0039388 | -0,792065047 | 0,1567775 | -5,0521602 | 4,37E-07 | 2,99E-06 |
| ENSG00000170949.17 | ZNF160 | protein_coding | 3250,463907 | 0,36050273 | 0,074568412 | 4,834523382 | 1,33E-06 | 8,51E-06 |
| ENSG00000170955.10 | CAVIN3 | protein_coding | 538,1866171 | -0,368053716 | 0,112067301 | -3,284220386 | 0,001022649 | 0,003953763 |
| ENSG00000170959.14 | DCDC1 | protein_coding | 6533,554697 | -0,46326189 | 0,063692175 | -7,273450602 | 3,50E-13 | 4,60E-12 |
| ENSG00000170962.13 | PDGFD | protein_coding | 13017,10309 | -0,36487781 | 0,049535495 | -7,365986975 | 1,76E-13 | 2,36E-12 |
| ENSG00000171016.12 | PYGO1 | protein_coding | 834,4546715 | 0,643928039 | 0,094602316 | 6,806683653 | 9,99E-12 | 1,14E-10 |
| ENSG00000171033.13 | PKIA | protein_coding | 3717,736225 | -0,424522571 | 0,074078637 | -5,73070168 | 1,00E-08 | 8,30E-08 |
| ENSG00000171045.16 | TSNARE1 | protein_coding | 1048,285277 | -0,337878484 | 0,099779614 | -3,386247666 | 0,000708554 | 0,002841412 |
| ENSG00000171055.15 | FEZ2 | protein_coding | 1986,52818 | 0,361983246 | 0,067582842 | 5,356141243 | 8,50E-08 | 6,31E-07 |
| ENSG00000171097.14 | KYAT1 | protein_coding | 396,4797295 | -0,793006521 | 0,127286438 | -6,23009438 | 4,66E-10 | 4,54E-09 |
| ENSG00000171103.11 | TRMT61B | protein_coding | 2711,957658 | -0,198709236 | 0,070705013 | -2,81039813 | 0,004948025 | 0,016171787 |
| ENSG00000171105.14 | INSR | protein_coding | 7017,150959 | -0,157893021 | 0,04846414 | -3,257935081 | 0,001122261 | 0,004301936 |
| ENSG00000171121.16 | KCNMB3 | protein_coding | 494,9026613 | 0,677524159 | 0,113683576 | 5,959736531 | 2,53E-09 | 2,25E-08 |
| ENSG00000171126.8 | KCNG3 | protein_coding | 4401,920654 | 0,212559173 | 0,059237002 | 3,588283782 | 0,000332862 | 0,001432853 |
| ENSG00000171130.18 | ATP6V0E2 | protein_coding | 9519,694364 | -0,247065703 | 0,056357601 | -4,383893202 | 1,17E-05 | 6,47E-05 |
| ENSG00000171132.14 | PRKCE | protein_coding | 1495,239196 | 1,322006751 | 0,089979881 | 14,69224828 | 7,23E-49 | 4,01E-47 |
| ENSG00000171148.13 | TADA3 | protein_coding | 6345,417292 | -0,120353819 | 0,049188875 | -2,446769103 | 0,014414315 | 0,04100756 |
| ENSG00000171155.8 | C1GALT1C1 | protein_coding | 8085,186735 | 0,286250305 | 0,04954193 | 5,777940087 | 7,56E-09 | 6,38E-08 |
| ENSG00000171163.15 | ZNF692 | protein_coding | 2437,320324 | -0,457471075 | 0,067662161 | -6,761106518 | 1,37E-11 | 1,55E-10 |
| ENSG00000171206.14 | TRIM8 | protein_coding | 2067,57089 | 0,211734615 | 0,076698047 | 2,760625909 | 0,005769072 | 0,018519115 |
| ENSG00000171208.9 | NETO2 | protein_coding | 10945,80935 | -0,195376429 | 0,045561958 | -4,288148204 | 1,80E-05 | 9,69E-05 |
| ENSG00000171219.9 | CDC42BPG | protein_coding | 6430,862833 | -0,64083615 | 0,055883958 | -11,46726494 | 1,93E-30 | 6,24E-29 |
| ENSG00000171223.6 | JUNB | protein_coding | 1561,795872 | 0,214308657 | 0,089231604 | 2,401712475 | 0,016318529 | 0,045603814 |
| ENSG00000171227.7 | TMEM37 | protein_coding | 20080,97219 | 0,155252695 | 0,045898794 | 3,38250056 | 0,000718291 | 0,002878408 |
| ENSG00000171234.14 | UGT2B7 | protein_coding | 10805,65427 | -0,384303631 | 0,046203842 | -8,317568723 | 8,98E-17 | 1,51E-15 |
| ENSG00000171241.9 | SHCBP1 | protein_coding | 2782,49729 | -0,356180675 | 0,065650305 | -5,425423018 | 5,78E-08 | 4,40E-07 |
| ENSG00000171243.8 | SOSTDC1 | protein_coding | 60,08095088 | 1,022680402 | 0,30342331 | 3,370474081 | 0,00075039 | 0,002993181 |

| | | | | | | | | |
|---|---|---|---|---|---|---|---|---|
| ENSG00000171262.11 | FAM98B | protein_coding | 10543,93738 | -0,1973881 | 0,047245126 | -4,177956909 | 2,94E-05 | 0,00015287 |
| ENSG00000171295.12 | ZNF440 | protein_coding | 3214,035656 | 0,245239114 | 0,064665759 | 3,792410651 | 0,000149192 | 0,000687443 |
| ENSG00000171303.7 | KCNK3 | protein_coding | 988,3374794 | -0,221385519 | 0,089787591 | -2,465658288 | 0,013676173 | 0,039224896 |
| ENSG00000171314.9 | PGAM1 | protein_coding | 28270,28473 | 0,130308625 | 0,04846643 | 2,688636777 | 0,007174444 | 0,022429105 |
| ENSG00000171316.12 | CHD7 | protein_coding | 38409,42593 | -0,196990772 | 0,050519279 | -3,899318778 | 9,65E-05 | 0,000460119 |
| ENSG00000171345.13 | KRT19 | protein_coding | 3922,937713 | 0,39918965 | 0,076182277 | 5,239928029 | 1,61E-07 | 1,16E-06 |
| ENSG00000171365.17 | CLCN5 | protein_coding | 14718,26571 | -0,456155121 | 0,066001298 | -6,911305361 | 4,80E-12 | 5,69E-11 |
| ENSG00000171388.12 | APLN | protein_coding | 982,5883466 | -0,843853631 | 0,132706553 | -6,358793974 | 2,03E-10 | 2,05E-09 |
| ENSG00000171428.14 | NAT1 | protein_coding | 846,0004825 | 0,654128663 | 0,089754632 | 7,287965513 | 3,15E-13 | 4,14E-12 |
| ENSG00000171435.13 | KSR2 | protein_coding | 1443,943703 | -0,380007481 | 0,083000813 | -4,578358549 | 4,69E-06 | 2,76E-05 |
| ENSG00000171443.7 | ZNF524 | protein_coding | 350,1240831 | -0,321762318 | 0,133821287 | -2,404418056 | 0,016198237 | 0,045337742 |
| ENSG00000171444.18 | MCC | protein_coding | 2160,583763 | 0,353398582 | 0,076106546 | 4,643471583 | 3,43E-06 | 2,05E-05 |
| ENSG00000171446.6 | KRT27 | protein_coding | 53,58339174 | 1,293985462 | 0,335223617 | 3,860066522 | 0,000113356 | 0,000533905 |
| ENSG00000171450.5 | CDK5R2 | protein_coding | 2633,666137 | 0,168129743 | 0,062235076 | 2,701527067 | 0,006902186 | 0,021701538 |
| ENSG00000171451.14 | DSEL | protein_coding | 2335,904575 | 0,455475694 | 0,065704111 | 6,932225229 | 4,14E-12 | 4,95E-11 |
| ENSG00000171466.10 | ZNF562 | protein_coding | 7985,19995 | 0,275582309 | 0,059305097 | 4,64685705 | 3,37E-06 | 2,03E-05 |
| ENSG00000171469.11 | ZNF561 | protein_coding | 6554,221212 | 0,213748232 | 0,056055913 | 3,813125495 | 0,00013722 | 0,000636319 |
| ENSG00000171475.14 | WIPF2 | protein_coding | 5451,895184 | 0,214195601 | 0,057960606 | 3,695537613 | 0,000219422 | 0,00097724 |
| ENSG00000171502.15 | COL24A1 | protein_coding | 411,5536149 | 0,722000803 | 0,129369633 | 5,580914068 | 2,39E-08 | 1,91E-07 |
| ENSG00000171533.11 | MAP6 | protein_coding | 2856,324047 | -0,852393937 | 0,073003452 | -11,67607717 | 1,69E-31 | 5,73E-30 |
| ENSG00000171551.12 | ECEL1 | protein_coding | 3788,124523 | -0,373338097 | 0,057393035 | -6,50493736 | 7,77E-11 | 8,18E-10 |
| ENSG00000171557.16 | FGG | protein_coding | 17,71739662 | 1,705147657 | 0,629663608 | 2,70802955 | 0,0067684 | 0,02132259 |
| ENSG00000171564.11 | FGB | protein_coding | 3231,562494 | 1,338798692 | 0,094797586 | 14,12270876 | 2,75E-45 | 1,43E-43 |
| ENSG00000171587.15 | DSCAM | protein_coding | 1550,423061 | 0,413219721 | 0,094902779 | 4,354137201 | 1,34E-05 | 7,34E-05 |
| ENSG00000171603.17 | CLSTN1 | protein_coding | 11808,40995 | 0,253363684 | 0,050749749 | 4,992412534 | 5,96E-07 | 4,00E-06 |
| ENSG00000171604.12 | CXXC5 | protein_coding | 6747,15602 | -0,227429863 | 0,050294927 | -4,521924535 | 6,13E-06 | 3,54E-05 |
| ENSG00000171606.18 | ZNF274 | protein_coding | 3033,138098 | 0,244142457 | 0,06262763 | 3,898318601 | 9,69E-05 | 0,000461925 |
| ENSG00000171621.14 | SPSB1 | protein_coding | 475,7585874 | 1,827372112 | 0,125511292 | 14,55942391 | 5,09E-48 | 2,75E-46 |
| ENSG00000171634.18 | BPTF | protein_coding | 22236,99928 | 0,182396866 | 0,052415657 | 3,479816437 | 0,000501757 | 0,002079189 |
| ENSG00000171649.12 | ZIK1 | protein_coding | 2943,432762 | 0,259004612 | 0,075042757 | 3,451427162 | 0,00055763 | 0,00228962 |
| ENSG00000171657.6 | GPR82 | protein_coding | 122,4137575 | -0,502836974 | 0,21183891 | -2,37367618 | 0,017611989 | 0,048683317 |
| ENSG00000171681.12 | ATF7IP | protein_coding | 12723,23376 | 0,12144708 | 0,042539637 | 2,854915749 | 0,004304825 | 0,014302184 |
| ENSG00000171703.17 | TCEA2 | protein_coding | 1831,462395 | -0,417802899 | 0,073709898 | -5,66820616 | 1,44E-08 | 1,18E-07 |
| ENSG00000171714.11 | ANO5 | protein_coding | 4994,335392 | 0,256234704 | 0,048943006 | 5,235369171 | 1,65E-07 | 1,18E-06 |
| ENSG00000171723.15 | GPHN | protein_coding | 3383,643171 | 0,232636781 | 0,078555234 | 2,961442144 | 0,00306202 | 0,010552612 |
| ENSG00000171724.3 | VAT1L | protein_coding | 233,4160547 | 0,535112852 | 0,187251648 | 2,857720385 | 0,004266962 | 0,014193151 |

| | | | | | | | | |
|---|---|---|---|---|---|---|---|---|
| ENSG00000171729.14 | TMEM51 | protein_coding | 1773,113258 | 0,274615803 | 0,06537174 | 4,200833589 | 2,66E-05 | 0,000139013 |
| ENSG00000171766.16 | GATM | protein_coding | 1393,10365 | 0,392718317 | 0,079677052 | 4,928876084 | 8,27E-07 | 5,43E-06 |
| ENSG00000171791.13 | BCL2 | protein_coding | 240,7953504 | -0,64329868 | 0,156646942 | -4,106678821 | 4,01E-05 | 0,000204316 |
| ENSG00000171792.11 | RHNO1 | protein_coding | 4583,267843 | -0,192783675 | 0,049286225 | -3,911512264 | 9,17E-05 | 0,000439635 |
| ENSG00000171798.18 | KNDC1 | protein_coding | 1665,365082 | -0,568174843 | 0,09330045 | -6,089733181 | 1,13E-09 | 1,05E-08 |
| ENSG00000171812.13 | COL8A2 | protein_coding | 180,7760202 | -1,229411309 | 0,178878515 | -6,87288415 | 6,29E-12 | 7,38E-11 |
| ENSG00000171815.5 | PCDHB1 | protein_coding | 92,33061729 | 0,982602383 | 0,281598422 | 3,489374602 | 0,000484152 | 0,002013261 |
| ENSG00000171843.16 | MLLT3 | protein_coding | 3939,118636 | 0,416840508 | 0,054097091 | 7,705414436 | 1,30E-14 | 1,90E-13 |
| ENSG00000171848.15 | RRM2 | protein_coding | 57771,90485 | -0,49936865 | 0,042083188 | -11,86622669 | 1,77E-32 | 6,33E-31 |
| ENSG00000171858.18 | RPS21 | protein_coding | 28693,70642 | -0,740217745 | 0,049470258 | -14,9628842 | 1,28E-50 | 7,50E-49 |
| ENSG00000171863.15 | RPS7 | protein_coding | 81073,49046 | -0,794331558 | 0,044156694 | -17,98892735 | 2,38E-72 | 2,00E-70 |
| ENSG00000171865.10 | RNASEH1 | protein_coding | 6632,016535 | -0,680742013 | 0,061689198 | -11,03502783 | 2,59E-28 | 7,65E-27 |
| ENSG00000171867.16 | PRNP | protein_coding | 15194,84127 | -0,105620434 | 0,04151491 | -2,544156663 | 0,010954194 | 0,032358 |
| ENSG00000171877.21 | FRMD5 | protein_coding | 1817,026864 | 0,262186887 | 0,069722866 | 3,760414662 | 0,000169632 | 0,000771674 |
| ENSG00000171914.16 | TLN2 | protein_coding | 1850,180009 | 0,752968007 | 0,106066197 | 7,09903842 | 1,26E-12 | 1,58E-11 |
| ENSG00000171928.14 | TVP23B | protein_coding | 8814,342207 | 0,157504813 | 0,044229888 | 3,561049324 | 0,000369376 | 0,001576166 |
| ENSG00000171951.5 | SCG2 | protein_coding | 58889,09252 | 0,345195201 | 0,039507231 | 8,737519514 | 2,38E-18 | 4,44E-17 |
| ENSG00000171956.7 | FOXB1 | protein_coding | 886,4503642 | 0,242729004 | 0,100862531 | 2,406532963 | 0,016104751 | 0,045126582 |
| ENSG00000171960.11 | PPIH | protein_coding | 1983,613505 | -0,267957862 | 0,072961028 | -3,672616328 | 0,00024008 | 0,001062307 |
| ENSG00000171988.19 | JMJD1C | protein_coding | 15336,58723 | 0,230859092 | 0,046116908 | 5,005953393 | 5,56E-07 | 3,75E-06 |
| ENSG00000171992.13 | SYNPO | protein_coding | 2203,116627 | -0,636462236 | 0,263796506 | -2,412701542 | 0,01583478 | 0,044442028 |
| ENSG00000172031.7 | EPHX4 | protein_coding | 49,67446106 | -1,144932812 | 0,333561461 | -3,432449331 | 0,000598156 | 0,002437343 |
| ENSG00000172053.18 | QARS | protein_coding | 11284,47696 | -0,334735776 | 0,054272304 | -6,167708946 | 6,93E-10 | 6,59E-09 |
| ENSG00000172057.10 | ORMDL3 | protein_coding | 6618,549812 | 0,329690834 | 0,050693178 | 6,503652911 | 7,84E-11 | 8,24E-10 |
| ENSG00000172058.15 | SERF1A | protein_coding | 463,7107653 | -0,615490121 | 0,136086629 | -4,522781753 | 6,10E-06 | 3,53E-05 |
| ENSG00000172123.12 | SLFN12 | protein_coding | 79,95348276 | 0,836575319 | 0,265526214 | 3,15063175 | 0,001629178 | 0,006021537 |
| ENSG00000172167.8 | MTBP | protein_coding | 1170,173073 | -0,299566317 | 0,094807036 | -3,159747739 | 0,001579058 | 0,005858417 |
| ENSG00000172172.7 | MRPL13 | protein_coding | 7881,872885 | -0,214915817 | 0,049887599 | -4,308000777 | 1,65E-05 | 8,91E-05 |
| ENSG00000172175.15 | MALT1 | protein_coding | 3327,104355 | 0,174340176 | 0,066785361 | 2,610454959 | 0,009042188 | 0,027459607 |
| ENSG00000172183.15 | ISG20 | protein_coding | 2383,408125 | 5,173040165 | 0,244383815 | 21,16768727 | 1,90E-99 | 2,20E-97 |
| ENSG00000172201.12 | ID4 | protein_coding | 8219,959315 | 0,788507726 | 0,063626999 | 12,39265937 | 2,86E-35 | 1,11E-33 |
| ENSG00000172216.5 | CEBPB | protein_coding | 242,2475285 | 1,171706246 | 0,158374214 | 7,398339779 | 1,38E-13 | 1,88E-12 |
| ENSG00000172239.14 | PAIP1 | protein_coding | 15301,98101 | 0,182411631 | 0,043718241 | 4,172437515 | 3,01E-05 | 0,000156189 |
| ENSG00000172260.15 | NEGR1 | protein_coding | 412,3942579 | -1,271831537 | 0,18507592 | -6,871944953 | 6,33E-12 | 7,42E-11 |
| ENSG00000172262.11 | ZNF131 | protein_coding | 4153,276848 | 0,174542266 | 0,063690566 | 2,740472851 | 0,006135085 | 0,019546303 |
| ENSG00000172264.17 | MACROD2 | protein_coding | 3186,421571 | -0,596918884 | 0,076834811 | -7,768859952 | 7,92E-15 | 1,18E-13 |

| | | | | | | | | |
|---|---|---|---|---|---|---|---|---|
| ENSG00000172270.19 | BSG | protein_coding | 48923,7587 | 0,137464127 | 0,042116978 | 3,263864924 | 0,001099035 | 0,004220094 |
| ENSG00000172273.12 | HINFP | protein_coding | 1914,358579 | 0,224145659 | 0,081773043 | 2,741070288 | 0,006123941 | 0,019516321 |
| ENSG00000172296.13 | SPTLC3 | protein_coding | 68,4991439 | 0,75333844 | 0,315988939 | 2,384065853 | 0,017122538 | 0,047564014 |
| ENSG00000172318.5 | B3GALT1 | protein_coding | 1271,945863 | 0,797869991 | 0,090607415 | 8,805791367 | 1,30E-18 | 2,45E-17 |
| ENSG00000172331.12 | BPGM | protein_coding | 1832,764174 | -0,226220312 | 0,071365015 | -3,16990492 | 0,001524888 | 0,005678973 |
| ENSG00000172336.5 | POP7 | protein_coding | 2179,49612 | -0,14989248 | 0,063336882 | -2,366590747 | 0,017952772 | 0,049509796 |
| ENSG00000172345.14 | STARD5 | protein_coding | 2697,104043 | 2,868428807 | 0,099199251 | 28,91583144 | 7,55E-184 | 1,40E-181 |
| ENSG00000172348.14 | RCAN2 | protein_coding | 4770,739415 | -0,149409024 | 0,050974659 | -2,93104511 | 0,003378237 | 0,011537057 |
| ENSG00000172354.10 | GNB2 | protein_coding | 13623,70035 | -0,108769083 | 0,042022032 | -2,588382309 | 0,009642789 | 0,029025079 |
| ENSG00000172367.16 | PDZD3 | protein_coding | 3525,290407 | -0,530677733 | 0,068036724 | -7,799871921 | 6,20E-15 | 9,31E-14 |
| ENSG00000172375.13 | C2CD2L | protein_coding | 2831,287815 | 0,185055463 | 0,06228658 | 2,971032636 | 0,002968002 | 0,010255874 |
| ENSG00000172379.21 | ARNT2 | protein_coding | 14888,47977 | -0,141266047 | 0,048416055 | -2,917752113 | 0,003525645 | 0,011980487 |
| ENSG00000172380.6 | GNG12 | protein_coding | 10764,05439 | -0,317383064 | 0,043700231 | -7,262731868 | 3,79E-13 | 4,96E-12 |
| ENSG00000172382.10 | PRSS27 | protein_coding | 112,7026805 | 0,682289771 | 0,218586695 | 3,121369177 | 0,001800122 | 0,006586305 |
| ENSG00000172403.11 | SYNPO2 | protein_coding | 1512,095443 | 3,434680149 | 0,081702808 | 42,03870364 | 0 | 0 |
| ENSG00000172428.11 | COPS9 | protein_coding | 4594,358115 | -0,489688091 | 0,062757353 | -7,802879979 | 6,05E-15 | 9,10E-14 |
| ENSG00000172432.19 | GTPBP2 | protein_coding | 7864,760871 | 0,52193432 | 0,054657747 | 9,549137166 | 1,31E-21 | 2,89E-20 |
| ENSG00000172456.17 | FGGY | protein_coding | 404,7195985 | -0,599413647 | 0,183825085 | -3,26078265 | 0,001111052 | 0,004263325 |
| ENSG00000172461.11 | FUT9 | protein_coding | 1070,17825 | -0,837603341 | 0,08368915 | -10,00850583 | 1,40E-23 | 3,39E-22 |
| ENSG00000172469.16 | MANEA | protein_coding | 4162,365468 | -0,473061013 | 0,057722741 | -8,195401025 | 2,50E-16 | 4,09E-15 |
| ENSG00000172500.12 | FIBP | protein_coding | 4963,812836 | -0,162699883 | 0,049452463 | -3,290025879 | 0,001001782 | 0,003881753 |
| ENSG00000172530.20 | BANP | protein_coding | 977,5299841 | -0,201022372 | 0,083035432 | -2,420922817 | 0,015481164 | 0,04357998 |
| ENSG00000172531.15 | PPP1CA | protein_coding | 8756,376807 | -0,260290158 | 0,046921036 | -5,547408542 | 2,90E-08 | 2,30E-07 |
| ENSG00000172554.12 | SNTG2 | protein_coding | 758,6034065 | -0,311724278 | 0,108239995 | -2,879936166 | 0,003977557 | 0,013334946 |
| ENSG00000172575.12 | RASGRP1 | protein_coding | 462,8637478 | 0,821171844 | 0,119757174 | 6,85697415 | 7,03E-12 | 8,20E-11 |
| ENSG00000172586.8 | CHCHD1 | protein_coding | 3526,561829 | -0,154079609 | 0,058941675 | -2,614103002 | 0,008946206 | 0,027212183 |
| ENSG00000172590.18 | MRPL52 | protein_coding | 3562,388282 | -0,280646025 | 0,067183945 | -4,177278126 | 2,95E-05 | 0,000153291 |
| ENSG00000172602.11 | RND1 | protein_coding | 315,0985344 | 0,408907239 | 0,132124406 | 3,094865309 | 0,001969023 | 0,007126666 |
| ENSG00000172613.8 | RAD9A | protein_coding | 1910,443313 | 0,633945072 | 0,09678961 | 6,549722363 | 5,76E-11 | 6,15E-10 |
| ENSG00000172663.9 | TMEM134 | protein_coding | 729,433983 | -0,432775915 | 0,118820631 | -3,642262388 | 0,000270252 | 0,001183254 |
| ENSG00000172667.11 | ZMAT3 | protein_coding | 2410,965258 | -0,697729252 | 0,069689138 | -10,01202303 | 1,35E-23 | 3,28E-22 |
| ENSG00000172716.16 | SLFN11 | protein_coding | 23236,33575 | 0,402833002 | 0,045097386 | 8,932513341 | 4,16E-19 | 8,07E-18 |
| ENSG00000172731.14 | LRRC20 | protein_coding | 8569,407065 | -0,339239169 | 0,054789092 | -6,19172822 | 5,95E-10 | 5,72E-09 |
| ENSG00000172785.18 | CBWD1 | protein_coding | 10033,28318 | -0,296643606 | 0,053347658 | -5,560574116 | 2,69E-08 | 2,14E-07 |
| ENSG00000172809.13 | RPL38 | protein_coding | 19763,8729 | -0,292200302 | 0,045995581 | -6,352790706 | 2,11E-10 | 2,13E-09 |
| ENSG00000172817.4 | CYP7B1 | protein_coding | 2162,44824 | -0,474931713 | 0,070918194 | -6,696895216 | 2,13E-11 | 2,36E-10 |

| | | | | | | | | |
|---|---|---|---|---|---|---|---|---|
| ENSG00000172818.10 | OVOL1 | protein_coding | 463,3601694 | 1,200937815 | 0,115219263 | 10,42306458 | 1,95E-25 | 5,14E-24 |
| ENSG00000172819.17 | RARG | protein_coding | 28,22517703 | 1,294026686 | 0,456656667 | 2,83369713 | 0,004601292 | 0,01514922 |
| ENSG00000172831.12 | CES2 | protein_coding | 4102,508133 | 0,478390316 | 0,068455995 | 6,988289557 | 2,78E-12 | 3,38E-11 |
| ENSG00000172840.6 | PDP2 | protein_coding | 4771,35201 | 0,191753555 | 0,059255442 | 3,236049697 | 0,001211963 | 0,004607342 |
| ENSG00000172889.16 | EGFL7 | protein_coding | 7687,428028 | 0,170259795 | 0,052128047 | 3,266184023 | 0,001090074 | 0,004190687 |
| ENSG00000172890.12 | NADSYN1 | protein_coding | 2657,76175 | 0,204361765 | 0,059642236 | 3,426460486 | 0,000611503 | 0,002486778 |
| ENSG00000172915.18 | NBEA | protein_coding | 12526,4137 | -0,321877971 | 0,067714399 | -4,75346423 | 2,00E-06 | 1,24E-05 |
| ENSG00000172936.14 | MYD88 | protein_coding | 7940,331224 | 2,957666591 | 0,077300149 | 38,26210709 | 0 | 0 |
| ENSG00000172954.13 | LCLAT1 | protein_coding | 7154,90923 | -0,151309338 | 0,062214633 | -2,432053859 | 0,015013475 | 0,042449228 |
| ENSG00000172955.17 | ADH6 | protein_coding | 2971,570013 | 0,422650144 | 0,065862804 | 6,417129513 | 1,39E-10 | 1,43E-09 |
| ENSG00000172965.16 | MIR4435-2HG | lncRNA | 94,02318122 | 0,775957997 | 0,250895852 | 3,092749399 | 0,001983115 | 0,007173058 |
| ENSG00000172992.12 | DCAKD | protein_coding | 2024,359562 | -0,182508438 | 0,070802815 | -2,577700292 | 0,009946022 | 0,029786457 |
| ENSG00000173011.12 | TADA2B | protein_coding | 2787,755026 | 0,207514328 | 0,057973525 | 3,57946713 | 0,000344296 | 0,001477832 |
| ENSG00000173039.19 | RELA | protein_coding | 3145,457326 | 0,195724517 | 0,060132005 | 3,254914189 | 0,001134266 | 0,004342713 |
| ENSG00000173041.12 | ZNF680 | protein_coding | 2190,89097 | 0,381126904 | 0,067926497 | 5,610872354 | 2,01E-08 | 1,62E-07 |
| ENSG00000173064.13 | HECTD4 | protein_coding | 24061,01744 | 0,204188206 | 0,058773727 | 3,474140832 | 0,000512492 | 0,002116639 |
| ENSG00000173065.13 | FAM222B | protein_coding | 3573,724316 | 0,143541601 | 0,058705499 | 2,445113366 | 0,014480662 | 0,041159896 |
| ENSG00000173083.15 | HPSE | protein_coding | 6009,731313 | 0,767374431 | 0,056315682 | 13,62630096 | 2,79E-42 | 1,34E-40 |
| ENSG00000173113.6 | TRMT112 | protein_coding | 7198,032326 | -0,326430659 | 0,05772009 | -5,655408008 | 1,55E-08 | 1,27E-07 |
| ENSG00000173114.13 | LRRN3 | protein_coding | 1152,168649 | 0,715485072 | 0,09106554 | 7,856814657 | 3,94E-15 | 6,00E-14 |
| ENSG00000173120.15 | KDM2A | protein_coding | 17176,32347 | 0,419923589 | 0,040464686 | 10,37753227 | 3,14E-25 | 8,21E-24 |
| ENSG00000173145.11 | NOC3L | protein_coding | 2688,614256 | 0,177712379 | 0,060750998 | 2,925258594 | 0,003441701 | 0,011734451 |
| ENSG00000173166.18 | RAPH1 | protein_coding | 5727,705142 | 0,147107187 | 0,056720705 | 2,593535956 | 0,009499461 | 0,028666491 |
| ENSG00000173193.15 | PARP14 | protein_coding | 47247,85509 | 7,480677525 | 0,285889218 | 26,16635066 | 6,42E-151 | 1,06E-148 |
| ENSG00000173207.13 | CKS1B | protein_coding | 5387,86342 | -0,170800234 | 0,049214606 | -3,470519149 | 0,000519453 | 0,002143031 |
| ENSG00000173213.9 | TUBB8P12 | protein_coding | 49,37517207 | 1,11644057 | 0,37907098 | 2,945201898 | 0,00322744 | 0,011058981 |
| ENSG00000173221.14 | GLRX | protein_coding | 2890,186779 | 0,450433825 | 0,061182977 | 7,362077634 | 1,81E-13 | 2,43E-12 |
| ENSG00000173226.17 | IQCB1 | protein_coding | 3714,605505 | 0,248720959 | 0,060231137 | 4,12944154 | 3,64E-05 | 0,000186156 |
| ENSG00000173230.15 | GOLGB1 | protein_coding | 19969,02532 | 0,278593907 | 0,045508596 | 6,121786443 | 9,25E-10 | 8,67E-09 |
| ENSG00000173253.15 | DMRT2 | protein_coding | 496,0673964 | 0,478894216 | 0,11579577 | 4,135679706 | 3,54E-05 | 0,000181583 |
| ENSG00000173272.16 | MZT2A | protein_coding | 2035,302477 | -0,419216526 | 0,083864436 | -4,998740174 | 5,77E-07 | 3,88E-06 |
| ENSG00000173276.14 | ZBTB21 | protein_coding | 4294,525805 | 0,378524754 | 0,060215421 | 6,286176386 | 3,25E-10 | 3,22E-09 |
| ENSG00000173281.5 | PPP1R3B | protein_coding | 2711,523034 | 0,488523988 | 0,063238673 | 7,725082882 | 1,12E-14 | 1,64E-13 |
| ENSG00000173327.8 | MAP3K11 | protein_coding | 5019,272825 | 0,280439504 | 0,055615746 | 5,04244793 | 4,60E-07 | 3,13E-06 |
| ENSG00000173349.5 | SFT2D3 | protein_coding | 2938,141095 | -0,302633061 | 0,057670563 | -5,247617556 | 1,54E-07 | 1,11E-06 |
| ENSG00000173366.11 | AC097637.1 | protein_coding | 11,0360075 | 3,063496406 | 1,016106726 | 3,014935663 | 0,002570338 | 0,009030081 |

| | | | | | | | | |
|---|---|---|---|---|---|---|---|---|
| ENSG00000173404.5 | INSM1 | protein_coding | 43165,24823 | 0,148205599 | 0,061940739 | 2,392699864 | 0,016724916 | 0,046597492 |
| ENSG00000173409.14 | ARV1 | protein_coding | 1547,820277 | -0,207622623 | 0,073843206 | -2,811668589 | 0,004928526 | 0,016119759 |
| ENSG00000173436.15 | MICOS10 | protein_coding | 7248,9104 | -0,397530357 | 0,052577347 | -7,560867515 | 4,00E-14 | 5,66E-13 |
| ENSG00000173442.13 | EHBP1L1 | protein_coding | 3353,783476 | -0,264516587 | 0,063212754 | -4,184544554 | 2,86E-05 | 0,000148641 |
| ENSG00000173451.7 | THAP2 | protein_coding | 1042,703266 | -0,214301934 | 0,080913944 | -2,648516736 | 0,008084585 | 0,024907735 |
| ENSG00000173452.13 | TMEM196 | protein_coding | 6551,388624 | 0,240042696 | 0,056369326 | 4,258392152 | 2,06E-05 | 0,00010967 |
| ENSG00000173456.4 | RNF26 | protein_coding | 4004,196147 | -0,228360935 | 0,059669417 | -3,827101853 | 0,000129661 | 0,000605129 |
| ENSG00000173457.11 | PPP1R14B | protein_coding | 1642,353229 | -0,408445756 | 0,070047447 | -5,83098706 | 5,51E-09 | 4,74E-08 |
| ENSG00000173473.11 | SMARCC1 | protein_coding | 25389,26224 | -0,304336624 | 0,044428576 | -6,8500198 | 7,38E-12 | 8,59E-11 |
| ENSG00000173482.16 | PTPRM | protein_coding | 2930,457649 | 0,456006324 | 0,180350547 | 2,52844436 | 0,011456924 | 0,033575629 |
| ENSG00000173486.13 | FKBP2 | protein_coding | 4336,464073 | 0,441893722 | 0,053459572 | 8,265942073 | 1,39E-16 | 2,31E-15 |
| ENSG00000173511.9 | VEGFB | protein_coding | 1981,625231 | 0,171432721 | 0,070039453 | 2,447659338 | 0,014378753 | 0,040942613 |
| ENSG00000173517.10 | PEAK1 | protein_coding | 5287,084702 | 0,429502567 | 0,062210582 | 6,904011423 | 5,06E-12 | 5,98E-11 |
| ENSG00000173540.12 | GMPPB | protein_coding | 7051,524296 | 0,188270627 | 0,055381548 | 3,399519052 | 0,000675045 | 0,002720119 |
| ENSG00000173545.5 | ZNF622 | protein_coding | 2768,764296 | 0,173037983 | 0,064278028 | 2,692023832 | 0,007101987 | 0,022242747 |
| ENSG00000173548.8 | SNX33 | protein_coding | 389,0168843 | 0,521846739 | 0,170862685 | 3,054187856 | 0,002256707 | 0,008037377 |
| ENSG00000173559.12 | NABP1 | protein_coding | 427,15815 | 0,491616944 | 0,141026451 | 3,485991034 | 0,000490317 | 0,002035514 |
| ENSG00000173575.21 | CHD2 | protein_coding | 20353,91726 | 0,258037005 | 0,053416583 | 4,830653538 | 1,36E-06 | 8,67E-06 |
| ENSG00000173598.14 | NUDT4 | protein_coding | 44272,95956 | 0,150683684 | 0,044193979 | 3,409597577 | 0,000650588 | 0,002631935 |
| ENSG00000173599.14 | PC | protein_coding | 1705,743033 | -0,390665346 | 0,067095927 | -5,822489733 | 5,80E-09 | 4,97E-08 |
| ENSG00000173611.17 | SCAI | protein_coding | 1835,282709 | -0,332855018 | 0,076689074 | -4,340318642 | 1,42E-05 | 7,77E-05 |
| ENSG00000173638.19 | SLC19A1 | protein_coding | 813,5998103 | -0,271492463 | 0,104265156 | -2,603865701 | 0,009217887 | 0,027932878 |
| ENSG00000173660.12 | UQCRH | protein_coding | 19470,43328 | -0,450044904 | 0,043046672 | -10,45481304 | 1,39E-25 | 3,70E-24 |
| ENSG00000173674.11 | EIF1AX | protein_coding | 31446,78217 | -0,24143992 | 0,040266001 | -5,996123576 | 2,02E-09 | 1,82E-08 |
| ENSG00000173692.13 | PSMD1 | protein_coding | 20857,35938 | -0,105817197 | 0,043910584 | -2,409833544 | 0,0159598 | 0,044765002 |
| ENSG00000173702.7 | MUC13 | protein_coding | 2028,534811 | 2,319300953 | 0,079552109 | 29,15448742 | 7,33E-187 | 1,39E-184 |
| ENSG00000173706.14 | HEG1 | protein_coding | 1319,405838 | 1,929998361 | 0,08937016 | 21,59555665 | 1,98E-103 | 2,36E-101 |
| ENSG00000173715.16 | C11orf80 | protein_coding | 1848,760231 | -0,391158238 | 0,077424149 | -5,052147693 | 4,37E-07 | 2,99E-06 |
| ENSG00000173726.11 | TOMM20 | protein_coding | 20811,7313 | -0,635574766 | 0,044229331 | -14,36998366 | 7,99E-47 | 4,25E-45 |
| ENSG00000173757.9 | STAT5B | protein_coding | 4942,141718 | -0,144375608 | 0,053684374 | -2,689341381 | 0,007159316 | 0,022388031 |
| ENSG00000173786.17 | CNP | protein_coding | 14229,84182 | 1,199501495 | 0,046051324 | 26,04705797 | 1,45E-149 | 2,37E-147 |
| ENSG00000173805.15 | HAP1 | protein_coding | 8761,581487 | 0,284169301 | 0,04738687 | 5,996794079 | 2,01E-09 | 1,82E-08 |
| ENSG00000173812.11 | EIF1 | protein_coding | 34535,22211 | -0,110547511 | 0,042987747 | -2,571605135 | 0,010122827 | 0,030235499 |
| ENSG00000173818.17 | ENDOV | protein_coding | 1072,539487 | -0,214042941 | 0,080931958 | -2,644727078 | 0,008175683 | 0,025146409 |
| ENSG00000173821.19 | AC124319.1 | protein_coding | 69994,29775 | 3,948591773 | 0,126955765 | 31,10210684 | 2,26E-212 | 4,70E-210 |
| ENSG00000173826.14 | KCNH6 | protein_coding | 5144,80223 | 0,32410903 | 0,056352501 | 5,751457784 | 8,85E-09 | 7,40E-08 |

| | | | | | | | | |
|---|---|---|---|---|---|---|---|---|
| ENSG00000173875.13 | ZNF791 | protein_coding | 5608,111122 | 0,156015236 | 0,047283952 | 3,299538857 | 0,000968438 | 0,00376812 |
| ENSG00000173889.16 | PHC3 | protein_coding | 8618,473196 | 0,317303573 | 0,063802388 | 4,973224064 | 6,58E-07 | 4,39E-06 |
| ENSG00000173894.11 | CBX2 | protein_coding | 6033,850541 | -0,417314011 | 0,052480661 | -7,951767445 | 1,84E-15 | 2,85E-14 |
| ENSG00000173898.13 | SPTBN2 | protein_coding | 5962,190136 | -0,558492208 | 0,204771576 | -2,727391267 | 0,006383729 | 0,020241071 |
| ENSG00000173905.8 | GOLIM4 | protein_coding | 7369,116259 | -0,138231007 | 0,045233009 | -3,055976401 | 0,002243289 | 0,007995914 |
| ENSG00000173915.16 | ATP5MD | protein_coding | 11914,06032 | -0,243579841 | 0,049205561 | -4,950250246 | 7,41E-07 | 4,91E-06 |
| ENSG00000173918.15 | C1QTNF1 | protein_coding | 51,92033662 | 1,616179508 | 0,434847348 | 3,716659456 | 0,000201874 | 0,000904276 |
| ENSG00000173926.6 | mars-03 | protein_coding | 488,9477655 | -0,380313054 | 0,121613908 | -3,127216772 | 0,001764698 | 0,006470366 |
| ENSG00000173930.9 | SLCO4C1 | protein_coding | 6215,773287 | 0,206801666 | 0,050577501 | 4,088807497 | 4,34E-05 | 0,000219857 |
| ENSG00000173947.14 | PIFO | protein_coding | 435,3324534 | 0,437728537 | 0,14419773 | 3,03561323 | 0,002400471 | 0,008492957 |
| ENSG00000173950.16 | XXYLT1 | protein_coding | 3736,083149 | -0,455310557 | 0,057754879 | -7,883499438 | 3,18E-15 | 4,86E-14 |
| ENSG00000173960.14 | UBXN2A | protein_coding | 3688,860811 | 0,341929242 | 0,063936912 | 5,34791608 | 8,90E-08 | 6,59E-07 |
| ENSG00000173992.9 | CCS | protein_coding | 2873,98981 | -0,345344982 | 0,069874137 | -4,942386352 | 7,72E-07 | 5,09E-06 |
| ENSG00000174004.6 | NRROS | protein_coding | 24,52881706 | -1,119887026 | 0,469342602 | -2,386075802 | 0,017029241 | 0,047345748 |
| ENSG00000174013.8 | FBXO45 | protein_coding | 6027,094477 | -0,316867319 | 0,047982758 | -6,603774547 | 4,01E-11 | 4,34E-10 |
| ENSG00000174021.11 | GNG5 | protein_coding | 6031,525441 | 0,30894511 | 0,050597019 | 6,105994326 | 1,02E-09 | 9,53E-09 |
| ENSG00000174028.6 | FAM3C2 | processed_pseudog | 142,0705775 | 9,616723377 | 2,96828989 | 3,23981947 | 0,001196054 | 0,004549937 |
| ENSG00000174032.17 | SLC25A30 | protein_coding | 1119,449342 | 0,429389929 | 0,103585686 | 4,145263167 | 3,39E-05 | 0,00017467 |
| ENSG00000174125.8 | TLR1 | protein_coding | 210,0462769 | 1,525015437 | 0,179381482 | 8,50152098 | 1,87E-17 | 3,28E-16 |
| ENSG00000174136.12 | RGMB | protein_coding | 4171,833531 | 0,190192228 | 0,055688746 | 3,415272223 | 0,000637183 | 0,002581881 |
| ENSG00000174197.16 | MGA | protein_coding | 15212,62202 | 0,410639759 | 0,050051832 | 8,204290306 | 2,32E-16 | 3,81E-15 |
| ENSG00000174206.12 | C12orf66 | protein_coding | 911,1962027 | 0,267580755 | 0,105329426 | 2,54041786 | 0,01107201 | 0,032650317 |
| ENSG00000174227.16 | PIGG | protein_coding | 6042,294506 | 0,354647839 | 0,051521875 | 6,883442024 | 5,84E-12 | 6,86E-11 |
| ENSG00000174231.17 | PRPF8 | protein_coding | 53819,10439 | -0,101738681 | 0,042164319 | -2,41290938 | 0,015825754 | 0,044422234 |
| ENSG00000174233.11 | ADCY6 | protein_coding | 7280,565815 | 0,34940722 | 0,049744902 | 7,023980504 | 2,16E-12 | 2,65E-11 |
| ENSG00000174255.6 | ZNF80 | protein_coding | 12,57415929 | 2,303620769 | 0,755297985 | 3,049949578 | 0,002288798 | 0,008142651 |
| ENSG00000174282.12 | ZBTB4 | protein_coding | 7451,692624 | 0,148546889 | 0,048398208 | 3,06926422 | 0,002145867 | 0,00768273 |
| ENSG00000174306.21 | ZHX3 | protein_coding | 13714,98962 | -0,202301925 | 0,066541788 | -3,040223743 | 0,002364025 | 0,008373222 |
| ENSG00000174327.6 | SLC16A13 | protein_coding | 643,4953933 | 0,325998844 | 0,10075646 | 3,235513093 | 0,001214243 | 0,004615076 |
| ENSG00000174348.13 | PODN | protein_coding | 73,70407978 | 1,141376667 | 0,268037808 | 4,258267422 | 2,06E-05 | 0,000109705 |
| ENSG00000174370.10 | C11orf45 | protein_coding | 78,42774842 | 1,385121571 | 0,282844818 | 4,897107826 | 9,73E-07 | 6,33E-06 |
| ENSG00000174373.16 | RALGAPA1 | protein_coding | 12516,60927 | 0,13775289 | 0,047906386 | 2,875459854 | 0,004034394 | 0,01350739 |
| ENSG00000174403.16 | MIR1-1HG-AS1 | lncRNA | 1190,977147 | -0,239587946 | 0,082572456 | -2,901548019 | 0,003713239 | 0,012540328 |
| ENSG00000174405.13 | LIG4 | protein_coding | 5069,042441 | 0,169569852 | 0,051847991 | 3,270519212 | 0,001073503 | 0,00413192 |
| ENSG00000174428.17 | GTF2IRD2B | protein_coding | 2009,290329 | -0,351133251 | 0,098862003 | -3,551751329 | 0,000382676 | 0,001627127 |
| ENSG00000174442.12 | ZWILCH | protein_coding | 7779,16293 | -0,161729736 | 0,048064608 | -3,364840422 | 0,000765879 | 0,003046868 |

| | | | | | | | | |
|---|---|---|---|---|---|---|---|---|
| ENSG00000174444.15 | RPL4 | protein_coding | 69978,40495 | -0,898211451 | 0,04059565 | -22,12580519 | 1,78E-108 | 2,21E-106 |
| ENSG00000174460.4 | ZCCHC12 | protein_coding | 235,5053272 | -0,406404318 | 0,165386505 | -2,457300362 | 0,013998559 | 0,04002713 |
| ENSG00000174469.23 | CNTNAP2 | protein_coding | 5766,592281 | -0,598758062 | 0,054427566 | -11,00100751 | 3,78E-28 | 1,11E-26 |
| ENSG00000174482.10 | LINGO2 | protein_coding | 1025,592428 | -0,26978634 | 0,094336951 | -2,859816191 | 0,004238866 | 0,014108035 |
| ENSG00000174485.15 | DENND4A | protein_coding | 7530,349443 | 0,178558821 | 0,064791817 | 2,755885381 | 0,00585335 | 0,018746879 |
| ENSG00000174498.14 | IGDCC3 | protein_coding | 2672,862225 | -0,725848553 | 0,101127091 | -7,177587562 | 7,10E-13 | 9,09E-12 |
| ENSG00000174502.19 | SLC26A9 | protein_coding | 36,72362203 | 1,639605455 | 0,404133459 | 4,057089107 | 4,97E-05 | 0,000248927 |
| ENSG00000174514.13 | MFSD4A | protein_coding | 407,358835 | -0,418729866 | 0,168755621 | -2,481279521 | 0,013091167 | 0,037744214 |
| ENSG00000174529.7 | TMEM81 | protein_coding | 184,2361525 | -0,444582472 | 0,17174661 | -2,588595329 | 0,009636827 | 0,029018041 |
| ENSG00000174547.13 | MRPL11 | protein_coding | 2545,842893 | -0,346797787 | 0,077737317 | -4,461149397 | 8,15E-06 | 4,62E-05 |
| ENSG00000174567.8 | GOLT1A | protein_coding | 2733,485846 | 0,323696325 | 0,062866744 | 5,148927827 | 2,62E-07 | 1,84E-06 |
| ENSG00000174574.16 | AKIRIN1 | protein_coding | 14693,17299 | 0,263450735 | 0,042387604 | 6,215277808 | 5,12E-10 | 4,97E-09 |
| ENSG00000174606.14 | ANGEL2 | protein_coding | 4601,703023 | 0,178395207 | 0,058240336 | 3,063086823 | 0,002190665 | 0,007828175 |
| ENSG00000174695.10 | TMEM167A | protein_coding | 22953,39969 | -0,398472064 | 0,04353218 | -9,153505825 | 5,51E-20 | 1,11E-18 |
| ENSG00000174718.12 | RESF1 | protein_coding | 4342,815294 | 0,429300871 | 0,061514906 | 6,978810492 | 2,98E-12 | 3,60E-11 |
| ENSG00000174738.13 | NR1D2 | protein_coding | 7587,165416 | 0,276439355 | 0,046676049 | 5,922509783 | 3,17E-09 | 2,80E-08 |
| ENSG00000174748.21 | RPL15 | protein_coding | 54264,92118 | -0,663147762 | 0,041898834 | -15,82735592 | 2,02E-56 | 1,31E-54 |
| ENSG00000174775.17 | HRAS | protein_coding | 1973,772237 | -0,391576915 | 0,067757211 | -5,779117956 | 7,51E-09 | 6,34E-08 |
| ENSG00000174776.11 | WDR49 | protein_coding | 264,7677768 | 0,968469843 | 0,176080069 | 5,500167323 | 3,79E-08 | 2,95E-07 |
| ENSG00000174799.10 | CEP135 | protein_coding | 2704,742789 | -0,258006237 | 0,058461806 | -4,413244414 | 1,02E-05 | 5,70E-05 |
| ENSG00000174808.12 | BTC | protein_coding | 1341,279546 | 3,944624713 | 0,102758184 | 38,3874505 | 0 | 0 |
| ENSG00000174827.13 | PDZK1 | protein_coding | 18590,27369 | -0,132981473 | 0,045225758 | -2,940392349 | 0,003277969 | 0,011225285 |
| ENSG00000174839.13 | DENND6A | protein_coding | 8532,992875 | -0,442373847 | 0,055736887 | -7,936823721 | 2,07E-15 | 3,21E-14 |
| ENSG00000174842.17 | GLMN | protein_coding | 2929,41487 | -0,237733196 | 0,056035038 | -4,242581148 | 2,21E-05 | 0,000117276 |
| ENSG00000174871.11 | CNIH2 | protein_coding | 6754,540686 | -0,335802711 | 0,0557157 | -6,027075123 | 1,67E-09 | 1,52E-08 |
| ENSG00000174886.13 | NDUFA11 | protein_coding | 4277,822931 | -0,295660891 | 0,069284066 | -4,267372103 | 1,98E-05 | 0,000105624 |
| ENSG00000174899.11 | PQLC2L | protein_coding | 76,93225056 | 0,814470755 | 0,270116344 | 3,015259068 | 0,002567598 | 0,009021866 |
| ENSG00000174938.14 | SEZ6L2 | protein_coding | 15527,60125 | 0,300159409 | 0,047600652 | 6,305783601 | 2,87E-10 | 2,86E-09 |
| ENSG00000174948.6 | GPR149 | protein_coding | 162,0303049 | 0,752289669 | 0,21476391 | 3,502868186 | 0,000460277 | 0,001926437 |
| ENSG00000174953.14 | DHX36 | protein_coding | 17675,39906 | 0,321664935 | 0,048786568 | 6,593309342 | 4,30E-11 | 4,65E-10 |
| ENSG00000175054.16 | ATR | protein_coding | 5754,166207 | 0,366738336 | 0,048784389 | 7,517534672 | 5,58E-14 | 7,81E-13 |
| ENSG00000175061.18 | SNHG29 | lncRNA | 23132,44327 | -0,898007634 | 0,048591789 | -18,48064568 | 2,96E-76 | 2,64E-74 |
| ENSG00000175063.17 | UBE2C | protein_coding | 8108,996628 | -0,809035797 | 0,051412499 | -15,73616954 | 8,55E-56 | 5,53E-54 |
| ENSG00000175073.8 | VCPIP1 | protein_coding | 4005,054379 | 0,606039939 | 0,059500891 | 10,18539273 | 2,30E-24 | 5,81E-23 |
| ENSG00000175093.5 | SPSB4 | protein_coding | 431,1798063 | -0,681152633 | 0,125363411 | -5,433424544 | 5,53E-08 | 4,22E-07 |
| ENSG00000175104.15 | TRAF6 | protein_coding | 3188,013775 | 0,28316542 | 0,066252822 | 4,274012958 | 1,92E-05 | 0,000102768 |

| | | | | | | | | |
|---|---|---|---|---|---|---|---|---|
| ENSG00000175105.7 | ZNF654 | protein_coding | 4672,475034 | 0,317253812 | 0,051405179 | 6,171631271 | 6,76E-10 | 6,44E-09 |
| ENSG00000175106.16 | TVP23C | protein_coding | 403,9158029 | 0,673625216 | 0,120446168 | 5,592749258 | 2,24E-08 | 1,79E-07 |
| ENSG00000175130.7 | MARCKSL1 | protein_coding | 21808,07942 | -0,167143408 | 0,041518547 | -4,025752851 | 5,68E-05 | 0,000282261 |
| ENSG00000175147.13 | TMEM51-AS1 | lncRNA | 227,1280054 | 0,4544532 | 0,167242336 | 2,717333489 | 0,006581026 | 0,020797104 |
| ENSG00000175166.17 | PSMD2 | protein_coding | 25493,41209 | -0,162228106 | 0,041140852 | -3,943236394 | 8,04E-05 | 0,000389304 |
| ENSG00000175182.15 | FAM131A | protein_coding | 1713,948955 | 0,300225543 | 0,088139675 | 3,406247459 | 0,000658625 | 0,002660621 |
| ENSG00000175197.12 | DDIT3 | protein_coding | 2513,27401 | 1,286407096 | 0,088557831 | 14,52618121 | 8,27E-48 | 4,42E-46 |
| ENSG00000175198.17 | PCCA | protein_coding | 1641,821027 | -0,245695922 | 0,079496678 | -3,090643882 | 0,00199723 | 0,007218313 |
| ENSG00000175203.15 | DCTN2 | protein_coding | 12497,76895 | -0,216817205 | 0,051127007 | -4,24075681 | 2,23E-05 | 0,00011815 |
| ENSG00000175216.14 | CKAP5 | protein_coding | 32870,8056 | -0,221515897 | 0,040356082 | -5,489033714 | 4,04E-08 | 3,13E-07 |
| ENSG00000175224.16 | ATG13 | protein_coding | 7767,734167 | 0,15345227 | 0,047330703 | 3,242129507 | 0,001186401 | 0,004517797 |
| ENSG00000175229.7 | GAL3ST3 | protein_coding | 23,57131586 | 1,573000196 | 0,604542633 | 2,601967355 | 0,009269067 | 0,02807663 |
| ENSG00000175262.14 | C1orf127 | protein_coding | 382,7073409 | -0,412045173 | 0,142094944 | -2,899787709 | 0,003734155 | 0,012603402 |
| ENSG00000175305.17 | CCNE2 | protein_coding | 7972,596921 | -0,378956036 | 0,046940556 | -8,073105055 | 6,85E-16 | 1,09E-14 |
| ENSG00000175309.15 | PHYKPL | protein_coding | 1339,615216 | 0,219190589 | 0,084930398 | 2,580826126 | 0,009856421 | 0,029565636 |
| ENSG00000175311.7 | ANKS4B | protein_coding | 27715,03181 | 0,265094387 | 0,040494924 | 6,546360821 | 5,90E-11 | 6,28E-10 |
| ENSG00000175318.12 | GRAMD2A | protein_coding | 494,5281904 | 0,645504555 | 0,133383837 | 4,839451102 | 1,30E-06 | 8,32E-06 |
| ENSG00000175322.11 | ZNF519 | protein_coding | 1103,683745 | -0,253375718 | 0,083212118 | -3,044937717 | 0,002327285 | 0,008263899 |
| ENSG00000175329.12 | ISX | protein_coding | 18893,91384 | 0,175115686 | 0,041362798 | 4,233651828 | 2,30E-05 | 0,000121775 |
| ENSG00000175334.8 | BANF1 | protein_coding | 8747,33588 | -0,206574928 | 0,051087414 | -4,043558131 | 5,26E-05 | 0,000263043 |
| ENSG00000175336.10 | APOF | protein_coding | 73,73359395 | 1,989503119 | 0,327993669 | 6,065675366 | 1,31E-09 | 1,21E-08 |
| ENSG00000175348.11 | TMEM9B | protein_coding | 4484,020067 | 0,163152856 | 0,051221588 | 3,185236209 | 0,001446359 | 0,005410685 |
| ENSG00000175390.14 | EIF3F | protein_coding | 12666,42146 | -0,767631415 | 0,172882368 | -4,440194948 | 8,99E-06 | 5,07E-05 |
| ENSG00000175414.7 | ARL10 | protein_coding | 7760,845894 | -1,226272392 | 0,139849358 | -8,768523562 | 1,81E-18 | 3,39E-17 |
| ENSG00000175426.11 | PCSK1 | protein_coding | 44807,6305 | -0,242149362 | 0,044755878 | -5,410448214 | 6,29E-08 | 4,76E-07 |
| ENSG00000175455.15 | CCDC14 | protein_coding | 11222,86954 | -0,186307379 | 0,04963151 | -3,753812458 | 0,000174165 | 0,000791018 |
| ENSG00000175482.9 | POLD4 | protein_coding | 973,2478949 | 0,512060489 | 0,094410311 | 5,423777191 | 5,84E-08 | 4,44E-07 |
| ENSG00000175497.16 | DPP10 | protein_coding | 8853,749096 | -0,302779089 | 0,052628272 | -5,753164198 | 8,76E-09 | 7,33E-08 |
| ENSG00000175548.9 | ALG10B | protein_coding | 3698,454154 | -0,552563952 | 0,075946731 | -7,275677858 | 3,45E-13 | 4,52E-12 |
| ENSG00000175550.8 | DRAP1 | protein_coding | 2792,467136 | 0,174542718 | 0,057503438 | 3,035344056 | 0,002402614 | 0,008499206 |
| ENSG00000175567.10 | UCP2 | protein_coding | 3993,97349 | -0,304633735 | 0,070989087 | -4,291275606 | 1,78E-05 | 9,55E-05 |
| ENSG00000175581.13 | MRPL48 | protein_coding | 2885,434929 | -0,180583324 | 0,060996899 | -2,960532864 | 0,003071074 | 0,010579552 |
| ENSG00000175595.14 | ERCC4 | protein_coding | 3644,084661 | 0,146986863 | 0,055854377 | 2,631608679 | 0,008498168 | 0,026021666 |
| ENSG00000175634.15 | RPS6KB2 | protein_coding | 3996,315724 | -0,27588125 | 0,057089543 | -4,832430549 | 1,35E-06 | 8,60E-06 |
| ENSG00000175643.10 | RMI2 | protein_coding | 4991,653308 | 0,534891631 | 0,06342846 | 8,432991047 | 3,37E-17 | 5,82E-16 |
| ENSG00000175711.8 | B3GNTL1 | protein_coding | 199,9438595 | -0,45983444 | 0,174487478 | -2,635343494 | 0,008405218 | 0,02576158 |

| | | | | | | | | |
|---|---|---|---|---|---|---|---|---|
| ENSG00000175727.14 | MLXIP | protein_coding | 4559,424699 | -0,185326344 | 0,066063221 | -2,805287744 | 0,005027168 | 0,016392369 |
| ENSG00000175756.13 | AURKAIP1 | protein_coding | 2960,408252 | -0,175875173 | 0,072405473 | -2,429031466 | 0,015139218 | 0,04275642 |
| ENSG00000175768.13 | TOMM5 | protein_coding | 7043,507476 | -0,213734724 | 0,057018001 | -3,748548217 | 0,000177861 | 0,000805609 |
| ENSG00000175779.3 | C15orf53 | protein_coding | 15,80272947 | 1,437420194 | 0,600185404 | 2,394960266 | 0,016622166 | 0,046357124 |
| ENSG00000175782.10 | SLC35E3 | protein_coding | 2875,497114 | -0,2677554 | 0,097367583 | -2,749943979 | 0,005960545 | 0,019052246 |
| ENSG00000175854.12 | SWI5 | protein_coding | 1648,261657 | -0,683192038 | 0,071650245 | -9,535097072 | 1,50E-21 | 3,29E-20 |
| ENSG00000175866.15 | BAIAP2 | protein_coding | 2256,675948 | 0,184200814 | 0,064671549 | 2,848251163 | 0,004396022 | 0,014545061 |
| ENSG00000175893.11 | ZDHHC21 | protein_coding | 5430,787257 | -0,212571432 | 0,050450315 | -4,213480763 | 2,51E-05 | 0,000132341 |
| ENSG00000175894.18 | TSPEAR | protein_coding | 1241,056151 | -0,347522832 | 0,084985018 | -4,089224672 | 4,33E-05 | 0,00021952 |
| ENSG00000175895.4 | PLEKHF2 | protein_coding | 4442,066808 | 0,565656769 | 0,051733775 | 10,93399376 | 7,93E-28 | 2,29E-26 |
| ENSG00000175928.5 | LRRN1 | protein_coding | 692,6993189 | -0,332930228 | 0,106408822 | -3,128784067 | 0,001755313 | 0,006441201 |
| ENSG00000175931.13 | UBE2O | protein_coding | 4652,745571 | 0,38052961 | 0,057572338 | 6,609591071 | 3,85E-11 | 4,18E-10 |
| ENSG00000175938.7 | ORAI3 | protein_coding | 1446,121337 | -0,218604971 | 0,072983842 | -2,99525161 | 0,002742185 | 0,009563658 |
| ENSG00000175970.11 | UNC119B | protein_coding | 15162,51871 | -0,289143683 | 0,046924859 | -6,161844456 | 7,19E-10 | 6,83E-09 |
| ENSG00000175984.15 | DENND2C | protein_coding | 360,0842934 | 0,456347602 | 0,158776904 | 2,874143469 | 0,004051249 | 0,01355777 |
| ENSG00000176014.12 | TUBB6 | protein_coding | 1291,787512 | -1,041609397 | 0,237218136 | -4,390934921 | 1,13E-05 | 6,28E-05 |
| ENSG00000176018.13 | LYSMD3 | protein_coding | 5264,063632 | 0,254420139 | 0,051942325 | 4,898127699 | 9,68E-07 | 6,29E-06 |
| ENSG00000176020.8 | AMIGO3 | protein_coding | 20,51735359 | 2,585488442 | 0,769111068 | 3,361658087 | 0,00077476 | 0,003079476 |
| ENSG00000176022.7 | B3GALT6 | protein_coding | 1802,821869 | -0,294180177 | 0,069353894 | -4,241725464 | 2,22E-05 | 0,000117668 |
| ENSG00000176024.18 | ZNF613 | protein_coding | 633,875573 | 0,382392524 | 0,117633424 | 3,250713187 | 0,001151159 | 0,00439848 |
| ENSG00000176049.16 | JAKMIP2 | protein_coding | 3388,542057 | 0,175329562 | 0,063032139 | 2,781589902 | 0,005409335 | 0,017491555 |
| ENSG00000176058.13 | TPRN | protein_coding | 1728,545832 | -0,253794668 | 0,105231264 | -2,411780105 | 0,015874853 | 0,044543388 |
| ENSG00000176087.15 | SLC35A4 | protein_coding | 8493,12776 | -0,128184442 | 0,049140283 | -2,608540987 | 0,009092912 | 0,027602476 |
| ENSG00000176092.15 | CRYBG2 | protein_coding | 74,8327558 | 0,814006735 | 0,291762748 | 2,789961159 | 0,005271437 | 0,017087946 |
| ENSG00000176095.12 | IP6K1 | protein_coding | 8031,814855 | 0,208913691 | 0,058029591 | 3,60012341 | 0,000318066 | 0,001374684 |
| ENSG00000176101.12 | SSNA1 | protein_coding | 2742,925197 | -0,217083092 | 0,06950747 | -3,123162024 | 0,001789192 | 0,006552706 |
| ENSG00000176102.13 | CSTF3 | protein_coding | 7893,313032 | 0,219779478 | 0,0517369 | 4,248021767 | 2,16E-05 | 0,0001146 |
| ENSG00000176124.14 | DLEU1 | lncRNA | 3305,735147 | -0,95356242 | 0,072527083 | -13,14767372 | 1,75E-39 | 7,81E-38 |
| ENSG00000176153.12 | GPX2 | protein_coding | 725,2991926 | 0,669798684 | 0,109340349 | 6,125814391 | 9,02E-10 | 8,47E-09 |
| ENSG00000176170.13 | SPHK1 | protein_coding | 14,65994589 | 2,583420276 | 0,714326964 | 3,616579532 | 0,000298522 | 0,001296936 |
| ENSG00000176204.13 | LRRTM4 | protein_coding | 75,22349278 | -0,714451105 | 0,269696888 | -2,64908917 | 0,008070903 | 0,024875799 |
| ENSG00000176209.11 | SMIM19 | protein_coding | 1712,760519 | -0,171737344 | 0,069564422 | -2,468752536 | 0,013558495 | 0,038912171 |
| ENSG00000176244.7 | ACBD7 | protein_coding | 1704,780553 | -0,555357472 | 0,068165404 | -8,147204348 | 3,72E-16 | 6,03E-15 |
| ENSG00000176273.15 | SLC35G1 | protein_coding | 1999,11109 | -0,220645249 | 0,073961906 | -2,983228253 | 0,002852252 | 0,009907614 |
| ENSG00000176340.4 | COX8A | protein_coding | 14545,93349 | -0,393134406 | 0,065349669 | -6,015859193 | 1,79E-09 | 1,63E-08 |
| ENSG00000176381.6 | PRR18 | protein_coding | 775,5523428 | -0,57764422 | 0,098936145 | -5,838555947 | 5,27E-09 | 4,54E-08 |

| | | | | | | | | |
|---|---|---|---|---|---|---|---|---|
| ENSG00000176386.9 | CDC26 | protein_coding | 3751,16838 | 0,294696214 | 0,053698357 | 5,48799309 | 4,07E-08 | 3,15E-07 |
| ENSG00000176390.12 | CRLF3 | protein_coding | 687,2920408 | 0,264512614 | 0,10170403 | 2,600807591 | 0,00930046 | 0,028152778 |
| ENSG00000176402.5 | GJC3 | protein_coding | 182,4533405 | 1,156264106 | 0,186711276 | 6,192792052 | 5,91E-10 | 5,69E-09 |
| ENSG00000176422.14 | SPRYD4 | protein_coding | 3431,497299 | 0,293577635 | 0,053263392 | 5,511808836 | 3,55E-08 | 2,77E-07 |
| ENSG00000176438.12 | SYNE3 | protein_coding | 1092,180477 | -0,838250192 | 0,085356112 | -9,820622918 | 9,18E-23 | 2,15E-21 |
| ENSG00000176473.13 | WDR25 | protein_coding | 1441,739002 | -0,368757173 | 0,072817407 | -5,064134933 | 4,10E-07 | 2,81E-06 |
| ENSG00000176476.9 | SGF29 | protein_coding | 1326,796189 | 0,496678447 | 0,071841723 | 6,913509666 | 4,73E-12 | 5,61E-11 |
| ENSG00000176485.11 | PLAAT3 | protein_coding | 802,0787672 | 0,749731334 | 0,112811384 | 6,645883686 | 3,01E-11 | 3,30E-10 |
| ENSG00000176490.5 | DIRAS1 | protein_coding | 2353,847441 | -0,192140275 | 0,060554797 | -3,172998402 | 0,001508734 | 0,005626256 |
| ENSG00000176532.4 | PRR15 | protein_coding | 761,0226087 | 1,521757905 | 0,093899053 | 16,20631797 | 4,55E-59 | 3,16E-57 |
| ENSG00000176542.10 | USF3 | protein_coding | 3155,440007 | 0,492391422 | 0,055882771 | 8,811149073 | 1,24E-18 | 2,34E-17 |
| ENSG00000176597.12 | B3GNT5 | protein_coding | 6914,830124 | 0,761478485 | 0,058749659 | 12,9614112 | 2,02E-38 | 8,67E-37 |
| ENSG00000176619.13 | LMNB2 | protein_coding | 6932,726744 | -0,473989165 | 0,049489939 | -9,577485241 | 9,94E-22 | 2,22E-20 |
| ENSG00000176623.12 | RMDN1 | protein_coding | 3689,510048 | -0,238592925 | 0,055205765 | -4,321884197 | 1,55E-05 | 8,40E-05 |
| ENSG00000176641.11 | RNF152 | protein_coding | 2379,693876 | 0,472726024 | 0,072471012 | 6,522967103 | 6,89E-11 | 7,28E-10 |
| ENSG00000176658.17 | MYO1D | protein_coding | 17050,56375 | -0,514083283 | 0,053063359 | -9,6881029 | 3,39E-22 | 7,71E-21 |
| ENSG00000176700.20 | SCAND2P | transcribed_unproc | 894,2058683 | 0,412554644 | 0,102323073 | 4,03188285 | 5,53E-05 | 0,000275423 |
| ENSG00000176731.12 | RBIS | protein_coding | 4361,609742 | -0,151366556 | 0,059705637 | -2,535213845 | 0,01123787 | 0,033061458 |
| ENSG00000176771.17 | NCKAP5 | protein_coding | 773,1093441 | 0,333242638 | 0,112738982 | 2,955877646 | 0,003117807 | 0,010715946 |
| ENSG00000176783.15 | RUFY1 | protein_coding | 4924,440355 | -0,154774973 | 0,059969694 | -2,580886488 | 0,009854698 | 0,029565636 |
| ENSG00000176887.7 | SOX11 | protein_coding | 9613,753666 | -0,178001739 | 0,061158029 | -2,910521193 | 0,003608265 | 0,012233535 |
| ENSG00000176890.16 | TYMS | protein_coding | 7403,367959 | -0,291960014 | 0,049765309 | -5,866737668 | 4,44E-09 | 3,86E-08 |
| ENSG00000176894.10 | PXMP2 | protein_coding | 1845,302047 | -0,613705388 | 0,06892381 | -8,904112974 | 5,38E-19 | 1,04E-17 |
| ENSG00000176915.15 | ANKLE2 | protein_coding | 11657,36587 | -0,14722238 | 0,04463817 | -3,298127565 | 0,000973319 | 0,003785802 |
| ENSG00000176920.12 | FUT2 | protein_coding | 362,0839443 | 0,691899396 | 0,131350849 | 5,267566987 | 1,38E-07 | 1,01E-06 |
| ENSG00000176928.6 | GCNT4 | protein_coding | 2386,814246 | 1,736140774 | 0,071447251 | 24,29961604 | 1,98E-130 | 2,95E-128 |
| ENSG00000176953.12 | NFATC2IP | protein_coding | 6823,695243 | 0,191403757 | 0,052546552 | 3,642556015 | 0,000269944 | 0,001182365 |
| ENSG00000176973.8 | FAM89B | protein_coding | 2113,033978 | -0,50668131 | 0,070090025 | -7,229007373 | 4,87E-13 | 6,30E-12 |
| ENSG00000176974.20 | SHMT1 | protein_coding | 3939,791974 | -0,327444742 | 0,060528908 | -5,409724924 | 6,31E-08 | 4,78E-07 |
| ENSG00000176986.16 | SEC24C | protein_coding | 20590,81458 | 0,185265323 | 0,048222632 | 3,84187497 | 0,000122098 | 0,000572683 |
| ENSG00000176994.11 | SMCR8 | protein_coding | 8896,070435 | 0,355031308 | 0,043210171 | 8,216382883 | 2,10E-16 | 3,46E-15 |
| ENSG00000177000.12 | MTHFR | protein_coding | 3993,354212 | 0,19449241 | 0,074414137 | 2,613648657 | 0,00895811 | 0,027244711 |
| ENSG00000177030.16 | DEAF1 | protein_coding | 3240,69966 | 0,18172037 | 0,075542649 | 2,405533477 | 0,016148872 | 0,045227693 |
| ENSG00000177034.16 | MTX3 | protein_coding | 6111,522631 | -0,359086447 | 0,057684313 | -6,225027711 | 4,81E-10 | 4,69E-09 |
| ENSG00000177042.14 | TMEM80 | protein_coding | 2870,502925 | 0,195272599 | 0,055751973 | 3,502523588 | 0,000460873 | 0,001928214 |
| ENSG00000177045.10 | SIX5 | protein_coding | 137,1376132 | -0,935998239 | 0,21158539 | -4,423737581 | 9,70E-06 | 5,45E-05 |

| | | | | | | | | |
|---|---|---|---|---|---|---|---|---|
| ENSG00000177076.6 | ACER2 | protein_coding | 911,3279042 | -0,280106743 | 0,0847704 | -3,304298946 | 0,000952142 | 0,003711772 |
| ENSG00000177082.12 | WDR73 | protein_coding | 5442,445999 | 0,147149899 | 0,056265325 | 2,615285654 | 0,008915286 | 0,027138091 |
| ENSG00000177084.16 | POLE | protein_coding | 13989,3446 | -0,30298565 | 0,043563908 | -6,954969536 | 3,53E-12 | 4,24E-11 |
| ENSG00000177119.16 | ANO6 | protein_coding | 6954,712349 | -0,214852151 | 0,046474276 | -4,623033807 | 3,78E-06 | 2,26E-05 |
| ENSG00000177156.11 | TALDO1 | protein_coding | 10825,65869 | -0,357242195 | 0,04629246 | -7,717070891 | 1,19E-14 | 1,74E-13 |
| ENSG00000177169.10 | ULK1 | protein_coding | 6288,091218 | -0,24292265 | 0,063349082 | -3,834667248 | 0,000125734 | 0,000588145 |
| ENSG00000177189.14 | RPS6KA3 | protein_coding | 59447,19594 | -0,360998972 | 0,050767815 | -7,110784146 | 1,15E-12 | 1,45E-11 |
| ENSG00000177191.2 | B3GNT8 | protein_coding | 1365,64584 | -0,357920341 | 0,08794276 | -4,069923879 | 4,70E-05 | 0,000236445 |
| ENSG00000177200.17 | CHD9 | protein_coding | 14049,6236 | 0,366234126 | 0,045087927 | 8,122665054 | 4,56E-16 | 7,36E-15 |
| ENSG00000177225.17 | GATD1 | protein_coding | 3894,976713 | -0,379133545 | 0,057341179 | -6,611889576 | 3,79E-11 | 4,12E-10 |
| ENSG00000177272.9 | KCNA3 | protein_coding | 688,7237496 | 0,409933562 | 0,100864204 | 4,064212551 | 4,82E-05 | 0,000241823 |
| ENSG00000177335.11 | C8orf31 | lncRNA | 186,3298884 | 2,092167374 | 0,213866386 | 9,782590947 | 1,34E-22 | 3,12E-21 |
| ENSG00000177337.8 | DLGAP1-AS1 | lncRNA | 13,22938994 | -1,765080854 | 0,65950652 | -2,67636604 | 0,007442531 | 0,023141861 |
| ENSG00000177359.20 | AC024940.2 | transcribed_unproc | 31,74868353 | -1,718125881 | 0,549114235 | -3,128904282 | 0,001754595 | 0,006439617 |
| ENSG00000177374.13 | HIC1 | protein_coding | 296,1003786 | -1,173550304 | 0,157414523 | -7,455159026 | 8,98E-14 | 1,23E-12 |
| ENSG00000177380.14 | PPFIA3 | protein_coding | 2251,903894 | 0,442151605 | 0,05994371 | 7,376113408 | 1,63E-13 | 2,20E-12 |
| ENSG00000177409.12 | SAMD9L | protein_coding | 2525,559026 | 12,10817548 | 0,988802728 | 12,24528932 | 1,78E-34 | 6,69E-33 |
| ENSG00000177410.13 | ZFAS1 | lncRNA | 8441,239427 | 0,133409708 | 0,055134879 | 2,419697137 | 0,015533438 | 0,043705271 |
| ENSG00000177426.20 | TGIF1 | protein_coding | 100,7706764 | -1,063694439 | 0,25046342 | -4,246905359 | 2,17E-05 | 0,000115145 |
| ENSG00000177459.11 | ERICH5 | protein_coding | 667,0023123 | 0,521085732 | 0,104603792 | 4,981518593 | 6,31E-07 | 4,22E-06 |
| ENSG00000177463.15 | NR2C2 | protein_coding | 11135,32203 | -0,20103248 | 0,053356718 | -3,767707002 | 0,000164754 | 0,000751154 |
| ENSG00000177485.7 | ZBTB33 | protein_coding | 9820,495469 | 0,574672032 | 0,049379277 | 11,63791916 | 2,64E-31 | 8,91E-30 |
| ENSG00000177519.4 | RPRM | protein_coding | 209,9703951 | 0,622299523 | 0,160344655 | 3,881011948 | 0,000104023 | 0,000493768 |
| ENSG00000177556.12 | ATOX1 | protein_coding | 2855,370368 | 0,207290214 | 0,059950454 | 3,45769213 | 0,000544824 | 0,002240715 |
| ENSG00000177565.16 | TBL1XR1 | protein_coding | 9030,378475 | -0,337209116 | 0,061144453 | -5,5149584 | 3,49E-08 | 2,73E-07 |
| ENSG00000177595.18 | PIDD1 | protein_coding | 1189,076228 | -0,362109274 | 0,095481536 | -3,792453372 | 0,000149166 | 0,000687443 |
| ENSG00000177600.9 | RPLP2 | protein_coding | 25276,79771 | -0,402716777 | 0,051277946 | -7,853605823 | 4,04E-15 | 6,15E-14 |
| ENSG00000177606.6 | JUN | protein_coding | 2829,363355 | 0,794605637 | 0,065104013 | 12,20517143 | 2,92E-34 | 1,09E-32 |
| ENSG00000177613.8 | CSTF2T | protein_coding | 9014,321253 | 0,148622565 | 0,059677065 | 2,490446965 | 0,012758254 | 0,036902535 |
| ENSG00000177628.16 | GBA | protein_coding | 12730,93959 | 0,359468824 | 0,045409762 | 7,916113323 | 2,45E-15 | 3,78E-14 |
| ENSG00000177663.13 | IL17RA | protein_coding | 1061,206552 | 0,396717804 | 0,137081609 | 2,894026458 | 0,003803361 | 0,012808177 |
| ENSG00000177669.3 | MBOAT4 | protein_coding | 1844,025895 | 0,337794709 | 0,067535483 | 5,001736777 | 5,68E-07 | 3,82E-06 |
| ENSG00000177692.11 | DNAJC28 | protein_coding | 572,7103273 | 0,312421193 | 0,111894358 | 2,79210855 | 0,005236579 | 0,016991516 |
| ENSG00000177697.19 | CD151 | protein_coding | 8451,772232 | -0,180205455 | 0,044954158 | -4,008649364 | 6,11E-05 | 0,000302101 |
| ENSG00000177700.6 | POLR2L | protein_coding | 6643,149911 | -0,313117571 | 0,052601785 | -5,952603479 | 2,64E-09 | 2,35E-08 |
| ENSG00000177707.11 | NECTIN3 | protein_coding | 9140,39159 | 0,146466388 | 0,046136231 | 3,174650043 | 0,001500173 | 0,005600829 |

| | | | | | | | | |
|---|---|---|---|---|---|---|---|---|
| ENSG00000177721.4 | ANXA2R | protein_coding | 252,6008379 | 1,18285161 | 0,159022386 | 7,438271037 | 1,02E-13 | 1,40E-12 |
| ENSG00000177728.17 | TMEM94 | protein_coding | 4560,088482 | -0,191404481 | 0,05678749 | -3,370539568 | 0,000750211 | 0,002993 |
| ENSG00000177732.8 | SOX12 | protein_coding | 5821,452504 | 0,200308758 | 0,048992989 | 4,088518837 | 4,34E-05 | 0,00021999 |
| ENSG00000177733.6 | HNRNPA0 | protein_coding | 26824,50753 | -0,414971223 | 0,042545214 | -9,753652294 | 1,78E-22 | 4,11E-21 |
| ENSG00000177738.4 | AC025171.1 | lncRNA | 1079,523454 | 0,676822194 | 0,084822117 | 7,979312722 | 1,47E-15 | 2,29E-14 |
| ENSG00000177842.12 | ZNF620 | protein_coding | 5689,628969 | 0,599421805 | 0,05294275 | 11,32207544 | 1,02E-29 | 3,15E-28 |
| ENSG00000177853.14 | ZNF518A | protein_coding | 18729,19674 | 0,264858461 | 0,049809391 | 5,317440301 | 1,05E-07 | 7,74E-07 |
| ENSG00000177868.12 | SVBP | protein_coding | 1566,359991 | -0,24574186 | 0,069516279 | -3,535026116 | 0,000407735 | 0,001722219 |
| ENSG00000177873.13 | ZNF619 | protein_coding | 1663,327332 | 0,484252805 | 0,06796294 | 7,125248014 | 1,04E-12 | 1,31E-11 |
| ENSG00000177875.4 | CCDC184 | protein_coding | 100,3398018 | 0,72678727 | 0,260281604 | 2,792311322 | 0,005233298 | 0,016983319 |
| ENSG00000177917.10 | ARL6IP6 | protein_coding | 4696,763508 | -0,238821021 | 0,049808394 | -4,794794681 | 1,63E-06 | 1,03E-05 |
| ENSG00000177943.14 | MAMDC4 | protein_coding | 3897,502159 | -0,553021869 | 0,07724592 | -7,159237242 | 8,11E-13 | 1,03E-11 |
| ENSG00000177951.17 | BET1L | protein_coding | 4198,541342 | 0,333566024 | 0,061151347 | 5,454761626 | 4,90E-08 | 3,76E-07 |
| ENSG00000177954.14 | RPS27 | protein_coding | 50878,38096 | -0,366723094 | 0,041034496 | -8,936946459 | 4,00E-19 | 7,78E-18 |
| ENSG00000177963.14 | RIC8A | protein_coding | 9384,089185 | 0,130368934 | 0,048641532 | 2,68019791 | 0,007357864 | 0,02292293 |
| ENSG00000177981.11 | ASB8 | protein_coding | 5540,408534 | -0,274209604 | 0,08817093 | -3,109977452 | 0,001871016 | 0,006813575 |
| ENSG00000177989.13 | ODF3B | protein_coding | 361,1238244 | 4,528607829 | 0,525654322 | 8,615182336 | 6,98E-18 | 1,26E-16 |
| ENSG00000177990.11 | DPY19L2 | protein_coding | 3429,970452 | -0,245990367 | 0,058915576 | -4,175302752 | 2,98E-05 | 0,000154556 |
| ENSG00000178031.17 | ADAMTSL1 | protein_coding | 12942,39077 | 0,263851856 | 0,044466642 | 5,933703175 | 2,96E-09 | 2,62E-08 |
| ENSG00000178035.12 | IMPDH2 | protein_coding | 9703,873584 | -0,19885525 | 0,04494762 | -4,424155266 | 9,68E-06 | 5,44E-05 |
| ENSG00000178038.17 | ALS2CL | protein_coding | 28,32227686 | 1,813440101 | 0,438669309 | 4,133957093 | 3,57E-05 | 0,000182742 |
| ENSG00000178053.18 | MLF1 | protein_coding | 1808,114721 | -0,26707626 | 0,069873305 | -3,822293203 | 0,000132216 | 0,000615066 |
| ENSG00000178075.20 | GRAMD1C | protein_coding | 1969,311178 | 1,066823973 | 0,084574152 | 12,61406646 | 1,77E-36 | 7,19E-35 |
| ENSG00000178078.11 | STAP2 | protein_coding | 2723,37458 | 0,729071313 | 0,075608755 | 9,642683761 | 5,28E-22 | 1,19E-20 |
| ENSG00000178104.19 | PDE4DIP | protein_coding | 52203,88367 | 0,196135727 | 0,037259213 | 5,264086672 | 1,41E-07 | 1,02E-06 |
| ENSG00000178105.11 | DDX10 | protein_coding | 3020,94403 | 0,197759151 | 0,073513591 | 2,690103278 | 0,007142991 | 0,0223494 |
| ENSG00000178163.8 | ZNF518B | protein_coding | 4675,154336 | 0,16871261 | 0,050034338 | 3,371936453 | 0,000746417 | 0,002978917 |
| ENSG00000178171.10 | AMER3 | protein_coding | 4650,185069 | 0,258049931 | 0,053305179 | 4,840991772 | 1,29E-06 | 8,27E-06 |
| ENSG00000178184.16 | PARD6G | protein_coding | 552,6517338 | -1,147163862 | 0,126271019 | -9,084933887 | 1,04E-19 | 2,07E-18 |
| ENSG00000178202.13 | POGLUT3 | protein_coding | 7893,481603 | -0,143168479 | 0,045567595 | -3,141892366 | 0,001678597 | 0,006184899 |
| ENSG00000178209.15 | PLEC | protein_coding | 4760,555344 | 0,576583563 | 0,070593782 | 8,167625339 | 3,15E-16 | 5,12E-15 |
| ENSG00000178229.8 | ZNF543 | protein_coding | 1127,810582 | 0,281094665 | 0,078702417 | 3,571614124 | 0,000354788 | 0,001517947 |
| ENSG00000178233.17 | TMEM151B | protein_coding | 1632,463675 | -0,262418246 | 0,0769143 | -3,41182647 | 0,000645292 | 0,002613327 |
| ENSG00000178234.13 | GALNT11 | protein_coding | 6969,396328 | -0,369723419 | 0,053336424 | -6,931912443 | 4,15E-12 | 4,96E-11 |
| ENSG00000178252.18 | WDR6 | protein_coding | 42785,97651 | -0,119987551 | 0,042992961 | -2,790864995 | 0,005256739 | 0,017052017 |
| ENSG00000178295.15 | GEN1 | protein_coding | 6689,503096 | -0,279362882 | 0,049767267 | -5,613386031 | 1,98E-08 | 1,60E-07 |

| | | | | | | | | |
|---|---|---|---|---|---|---|---|---|
| ENSG00000178381.11 | ZFAND2A | protein_coding | 396,3392858 | 0,424418915 | 0,127313406 | 3,333654554 | 0,00085713 | 0,003372928 |
| ENSG00000178385.15 | PLEKHM3 | protein_coding | 3519,223089 | -0,230033719 | 0,076309071 | -3,014500311 | 0,002574029 | 0,00904164 |
| ENSG00000178386.13 | AC092072.1 | protein_coding | 1434,591725 | 0,317143653 | 0,106356188 | 2,981901264 | 0,002864644 | 0,009947588 |
| ENSG00000178473.7 | UCN3 | protein_coding | 2085,479055 | 0,455064911 | 0,062793834 | 7,246968114 | 4,26E-13 | 5,54E-12 |
| ENSG00000178498.16 | DTX3 | protein_coding | 4827,645947 | 0,205535453 | 0,053040202 | 3,875088046 | 0,000106586 | 0,000504553 |
| ENSG00000178502.6 | KLHL11 | protein_coding | 2687,541721 | 0,178850246 | 0,061871411 | 2,89067669 | 0,003844134 | 0,01293774 |
| ENSG00000178531.6 | CTXN1 | protein_coding | 519,558161 | -0,265315094 | 0,111528785 | -2,378893429 | 0,017364695 | 0,048123852 |
| ENSG00000178538.10 | CA8 | protein_coding | 5070,714485 | -0,301802009 | 0,052788824 | -5,717157251 | 1,08E-08 | 8,97E-08 |
| ENSG00000178573.7 | MAF | protein_coding | 7904,489953 | 0,551494542 | 0,059044239 | 9,340361584 | 9,60E-21 | 2,04E-19 |
| ENSG00000178585.15 | CTNNBIP1 | protein_coding | 4510,548091 | -0,3457771 | 0,06233656 | -5,546939029 | 2,91E-08 | 2,30E-07 |
| ENSG00000178607.16 | ERN1 | protein_coding | 3005,44925 | 0,483738168 | 0,073168645 | 6,611276833 | 3,81E-11 | 4,13E-10 |
| ENSG00000178631.7 | ACTG1P1 | processed_pseudog | 44,94647583 | 0,948270302 | 0,345373501 | 2,745637111 | 0,006039353 | 0,019274037 |
| ENSG00000178665.16 | ZNF713 | protein_coding | 1020,754577 | -0,20268539 | 0,085422119 | -2,372750662 | 0,017656178 | 0,048793483 |
| ENSG00000178685.14 | PARP10 | protein_coding | 12861,62042 | 5,703211589 | 0,243136079 | 23,45687078 | 1,12E-121 | 1,53E-119 |
| ENSG00000178694.10 | NSUN3 | protein_coding | 3075,8616 | 0,250829628 | 0,065347332 | 3,838406553 | 0,000123835 | 0,000579985 |
| ENSG00000178695.5 | KCTD12 | protein_coding | 12145,20277 | 0,228842187 | 0,052328763 | 4,373162605 | 1,22E-05 | 6,77E-05 |
| ENSG00000178700.7 | DHFR2 | protein_coding | 3438,911932 | -0,415267698 | 0,055574074 | -7,472327822 | 7,88E-14 | 1,09E-12 |
| ENSG00000178718.6 | RPP25 | protein_coding | 3935,625434 | 0,200691395 | 0,059649457 | 3,364513353 | 0,000766788 | 0,003049943 |
| ENSG00000178719.17 | GRINA | protein_coding | 7549,04901 | 1,075831973 | 0,048313318 | 22,26781398 | 7,58E-110 | 9,53E-108 |
| ENSG00000178761.15 | FAM219B | protein_coding | 6649,499036 | -0,341465732 | 0,061571236 | -5,545864552 | 2,93E-08 | 2,31E-07 |
| ENSG00000178796.12 | RIIAD1 | protein_coding | 759,5273294 | -0,487053984 | 0,09582379 | -5,082808611 | 3,72E-07 | 2,56E-06 |
| ENSG00000178814.17 | OPLAH | protein_coding | 111,6421319 | 0,632991223 | 0,22970378 | 2,755684835 | 0,00585694 | 0,018755707 |
| ENSG00000178826.11 | TMEM139 | protein_coding | 1004,430538 | 0,336928892 | 0,101554786 | 3,317705699 | 0,000907601 | 0,003549198 |
| ENSG00000178882.14 | RFLNA | protein_coding | 569,2533637 | -0,295631067 | 0,120929115 | -2,44466411 | 0,01449871 | 0,041200791 |
| ENSG00000178913.7 | TAF7 | protein_coding | 14620,88244 | 0,197567662 | 0,043146802 | 4,57896423 | 4,67E-06 | 2,75E-05 |
| ENSG00000178921.14 | PFAS | protein_coding | 2050,062068 | -0,431303905 | 0,090567811 | -4,762220695 | 1,91E-06 | 1,19E-05 |
| ENSG00000178927.18 | CYBC1 | protein_coding | 5563,084985 | 0,326226852 | 0,065786128 | 4,958900339 | 7,09E-07 | 4,71E-06 |
| ENSG00000178952.11 | TUFM | protein_coding | 9996,875652 | -0,189125392 | 0,053994735 | -3,502663608 | 0,000460631 | 0,001927559 |
| ENSG00000178965.14 | ERICH3 | protein_coding | 77,63819721 | 0,747988456 | 0,26378143 | 2,835637277 | 0,004573436 | 0,015074456 |
| ENSG00000178971.16 | CTC1 | protein_coding | 4704,771483 | -0,202542307 | 0,072755434 | -2,783878765 | 0,005371311 | 0,017388585 |
| ENSG00000178974.10 | FBXO34 | protein_coding | 4300,275379 | 0,224545847 | 0,050023435 | 4,488813034 | 7,16E-06 | 4,10E-05 |
| ENSG00000178977.3 | LINC00324 | lncRNA | 192,3183024 | -0,969144363 | 0,175518495 | -5,521608216 | 3,36E-08 | 2,63E-07 |
| ENSG00000178982.9 | EIF3K | protein_coding | 9056,947389 | -0,306045751 | 0,042963827 | -7,123335376 | 1,05E-12 | 1,33E-11 |
| ENSG00000178988.11 | MRFAP1L1 | protein_coding | 12706,18363 | 0,108708194 | 0,042959286 | 2,530493509 | 0,011390219 | 0,033409232 |
| ENSG00000178996.14 | SNX18 | protein_coding | 4830,105802 | -0,211475294 | 0,054657869 | -3,869073143 | 0,00010925 | 0,000516185 |
| ENSG00000178999.13 | AURKB | protein_coding | 3619,613058 | -0,49276717 | 0,061615199 | -7,997493745 | 1,27E-15 | 1,99E-14 |

| | | | | | | | | |
|---|---|---|---|---|---|---|---|---|
| ENSG00000179021.10 | C3orf38 | protein_coding | 3864,132012 | 0,204294164 | 0,053817907 | 3,796025806 | 0,000147034 | 0,00067903 |
| ENSG00000179029.14 | TMEM107 | protein_coding | 1660,86763 | -0,851018516 | 0,07418828 | -11,47106406 | 1,84E-30 | 5,99E-29 |
| ENSG00000179044.16 | EXOC3L1 | protein_coding | 87,18784413 | 1,940552603 | 0,311839409 | 6,222922915 | 4,88E-10 | 4,75E-09 |
| ENSG00000179104.9 | TMTC2 | protein_coding | 4110,491053 | 0,207134593 | 0,056656257 | 3,655987976 | 0,000256193 | 0,001127181 |
| ENSG00000179119.15 | SPTY2D1 | protein_coding | 5953,125309 | 0,171623829 | 0,052327474 | 3,27980346 | 0,001038794 | 0,004009985 |
| ENSG00000179134.15 | SAMD4B | protein_coding | 8149,009258 | -0,14244258 | 0,054754979 | -2,601454365 | 0,009282941 | 0,028110573 |
| ENSG00000179163.11 | FUCA1 | protein_coding | 9777,743051 | 0,325784905 | 0,050145269 | 6,496822313 | 8,20E-11 | 8,60E-10 |
| ENSG00000179178.11 | TMEM125 | protein_coding | 3315,80608 | 0,267856408 | 0,071042563 | 3,770365207 | 0,000163009 | 0,0007438 |
| ENSG00000179195.16 | ZNF664 | protein_coding | 47358,31292 | -0,349660262 | 0,042302823 | -8,265648385 | 1,39E-16 | 2,31E-15 |
| ENSG00000179218.13 | CALR | protein_coding | 106334,7371 | 0,675085279 | 0,040371787 | 16,72170917 | 9,11E-63 | 6,57E-61 |
| ENSG00000179222.17 | MAGED1 | protein_coding | 24223,80904 | -0,165529269 | 0,048457415 | -3,415973997 | 0,000635543 | 0,002576628 |
| ENSG00000179240.10 | GVQW3 | protein_coding | 3196,129415 | -0,21468547 | 0,068010662 | -3,15664431 | 0,001595959 | 0,005913323 |
| ENSG00000179241.13 | LDLRAD3 | protein_coding | 2719,878043 | -0,361377548 | 0,071544359 | -5,051097722 | 4,39E-07 | 3,01E-06 |
| ENSG00000179295.17 | PTPN11 | protein_coding | 34154,55367 | -0,422371736 | 0,047014868 | -8,98379074 | 2,62E-19 | 5,11E-18 |
| ENSG00000179364.13 | PACS2 | protein_coding | 3753,400642 | -0,284608488 | 0,05432746 | -5,238759312 | 1,62E-07 | 1,16E-06 |
| ENSG00000179387.10 | ELMOD2 | protein_coding | 11105,35448 | -0,60059364 | 0,047078019 | -12,75741104 | 2,83E-37 | 1,17E-35 |
| ENSG00000179397.18 | CATSPERE | protein_coding | 71,27060018 | -0,832967777 | 0,29965616 | -2,779745211 | 0,005440156 | 0,017581117 |
| ENSG00000179399.15 | GPC5 | protein_coding | 74,89611978 | 0,794472865 | 0,318292786 | 2,496044207 | 0,012558695 | 0,03640485 |
| ENSG00000179454.13 | KLHL28 | protein_coding | 4201,99816 | 0,230587965 | 0,060187148 | 3,83118279 | 0,000127529 | 0,000595796 |
| ENSG00000179456.10 | ZBTB18 | protein_coding | 3006,88958 | 0,322443525 | 0,061525306 | 5,240827669 | 1,60E-07 | 1,15E-06 |
| ENSG00000179532.12 | DNHD1 | protein_coding | 367,5455562 | 0,606832069 | 0,129513983 | 4,68545601 | 2,79E-06 | 1,70E-05 |
| ENSG00000179562.3 | GCC1 | protein_coding | 5693,834842 | 0,457367441 | 0,053335034 | 8,575366137 | 9,88E-18 | 1,77E-16 |
| ENSG00000179583.19 | CIITA | protein_coding | 91,60980889 | 1,674242996 | 0,312544111 | 5,356821438 | 8,47E-08 | 6,29E-07 |
| ENSG00000179588.9 | ZFPM1 | protein_coding | 1813,403903 | -0,336869271 | 0,081194618 | -4,148911342 | 3,34E-05 | 0,000172107 |
| ENSG00000179603.17 | GRM8 | protein_coding | 324,2117745 | 1,605519001 | 0,455644533 | 3,523621778 | 0,000425691 | 0,001792347 |
| ENSG00000179627.10 | ZBTB42 | protein_coding | 1645,055171 | 1,008452148 | 0,080873264 | 12,46953686 | 1,09E-35 | 4,32E-34 |
| ENSG00000179630.11 | LACC1 | protein_coding | 2641,252881 | 0,507570457 | 0,069971632 | 7,253946283 | 4,05E-13 | 5,28E-12 |
| ENSG00000179674.3 | ARL14 | protein_coding | 133,6749824 | 1,468485145 | 0,211948031 | 6,928515165 | 4,25E-12 | 5,07E-11 |
| ENSG00000179715.13 | PCED1B | protein_coding | 110,5226029 | 0,94349529 | 0,268737113 | 3,510848499 | 0,000446679 | 0,001875451 |
| ENSG00000179743.4 | AL450998.2 | lncRNA | 1361,895574 | -0,826176233 | 0,072207256 | -11,44173424 | 2,59E-30 | 8,29E-29 |
| ENSG00000179750.16 | APOBEC3B | protein_coding | 689,2938006 | 1,678367044 | 0,105809824 | 15,8621097 | 1,16E-56 | 7,59E-55 |
| ENSG00000179761.12 | PIPOX | protein_coding | 5192,722559 | -0,182719043 | 0,060060341 | -3,042257824 | 0,002348107 | 0,008326014 |
| ENSG00000179774.8 | ATOH7 | protein_coding | 169,6631446 | -0,557354683 | 0,187111355 | -2,978732543 | 0,002894433 | 0,010034005 |
| ENSG00000179818.14 | PCBP1-AS1 | lncRNA | 2751,797085 | -0,200274379 | 0,061781814 | -3,24163966 | 0,001188442 | 0,004524803 |
| ENSG00000179833.4 | SERTAD2 | protein_coding | 4197,489216 | 0,18723257 | 0,060623542 | 3,088446557 | 0,002012059 | 0,007261413 |
| ENSG00000179909.15 | ZNF154 | protein_coding | 815,7170329 | 0,569366017 | 0,095219922 | 5,979484175 | 2,24E-09 | 2,01E-08 |

| | | | | | | | | |
|---|---|---|---|---|---|---|---|---|
| ENSG00000179918.18 | SEPHS2 | protein_coding | 13119,91772 | 0,353358973 | 0,04152247 | 8,510066408 | 1,74E-17 | 3,07E-16 |
| ENSG00000179933.6 | C14orf119 | protein_coding | 5982,219867 | 0,162196797 | 0,050342569 | 3,221861716 | 0,001273606 | 0,004815655 |
| ENSG00000179954.16 | SSC5D | protein_coding | 445,7998611 | -0,653306812 | 0,135979274 | -4,804458737 | 1,55E-06 | 9,80E-06 |
| ENSG00000179978.11 | AC140134.1 | unprocessed_pseud | 1175,539336 | 0,403092611 | 0,102231419 | 3,942942529 | 8,05E-05 | 0,000389697 |
| ENSG00000180011.7 | ZADH2 | protein_coding | 16195,7681 | -0,260329828 | 0,049901651 | -5,216857982 | 1,82E-07 | 1,30E-06 |
| ENSG00000180035.12 | ZNF48 | protein_coding | 1746,718651 | 0,17720787 | 0,074518787 | 2,378029447 | 0,017405436 | 0,048218948 |
| ENSG00000180044.5 | C3orf80 | protein_coding | 1255,123171 | -0,360562912 | 0,082210334 | -4,385858756 | 1,16E-05 | 6,41E-05 |
| ENSG00000180096.12 | sept-01 | protein_coding | 72,5343563 | -1,036274712 | 0,286621838 | -3,615477164 | 0,000299795 | 0,001302216 |
| ENSG00000180098.9 | TRNAU1AP | protein_coding | 1947,87929 | -0,244810249 | 0,072080232 | -3,39635767 | 0,000682891 | 0,00274829 |
| ENSG00000180113.16 | TDRD6 | protein_coding | 73,46580835 | -0,993353214 | 0,345071497 | -2,878688107 | 0,003993331 | 0,013381848 |
| ENSG00000180155.20 | LYNX1 | protein_coding | 269,0917243 | 0,571329809 | 0,176786959 | 3,231741828 | 0,001230382 | 0,004668689 |
| ENSG00000180176.14 | TH | protein_coding | 79,50854418 | -0,763523618 | 0,290164272 | -2,631349522 | 0,008504652 | 0,026034437 |
| ENSG00000180228.13 | PRKRA | protein_coding | 6147,453554 | -0,213969348 | 0,047007279 | -4,551834359 | 5,32E-06 | 3,10E-05 |
| ENSG00000180251.4 | SLC9A4 | protein_coding | 106,6153109 | 0,930408385 | 0,28108844 | 3,310020092 | 0,000932893 | 0,003641776 |
| ENSG00000180257.13 | ZNF816 | protein_coding | 625,1009956 | 0,905666514 | 0,102796713 | 8,810267253 | 1,25E-18 | 2,36E-17 |
| ENSG00000180304.14 | OAZ2 | protein_coding | 17547,19054 | -0,330314337 | 0,048172954 | -6,85684205 | 7,04E-12 | 8,21E-11 |
| ENSG00000180346.3 | TIGD2 | protein_coding | 1568,509948 | 0,357921541 | 0,070068266 | 5,108183248 | 3,25E-07 | 2,26E-06 |
| ENSG00000180347.13 | ITPRID1 | protein_coding | 70,64602076 | 0,867204962 | 0,324322142 | 2,673899954 | 0,007497481 | 0,023290203 |
| ENSG00000180354.16 | MTURN | protein_coding | 1779,213031 | -1,518657757 | 0,075105701 | -20,22027268 | 6,49E-91 | 6,83E-89 |
| ENSG00000180357.9 | ZNF609 | protein_coding | 9064,579188 | 0,260558518 | 0,046435988 | 5,611133324 | 2,01E-08 | 1,62E-07 |
| ENSG00000180398.12 | MCFD2 | protein_coding | 19579,2153 | 0,306312412 | 0,042134837 | 7,269813552 | 3,60E-13 | 4,71E-12 |
| ENSG00000180423.4 | HARBI1 | protein_coding | 993,5719222 | 0,41615212 | 0,088091645 | 4,724081583 | 2,31E-06 | 1,43E-05 |
| ENSG00000180488.15 | MIGA1 | protein_coding | 6716,049035 | -0,496867081 | 0,061338084 | -8,100466308 | 5,47E-16 | 8,78E-15 |
| ENSG00000180530.11 | NRIP1 | protein_coding | 13376,30291 | 0,42859437 | 0,044634522 | 9,60230673 | 7,82E-22 | 1,75E-20 |
| ENSG00000180535.3 | BHLHA15 | protein_coding | 42,29159007 | 0,886971488 | 0,360952741 | 2,457306423 | 0,013998323 | 0,04002713 |
| ENSG00000180537.13 | RNF182 | protein_coding | 4727,796935 | -0,622574562 | 0,062063089 | -10,03131772 | 1,11E-23 | 2,70E-22 |
| ENSG00000180573.9 | HIST1H2AC | protein_coding | 542,4624823 | 0,316028041 | 0,110835155 | 2,851333945 | 0,004353622 | 0,014432178 |
| ENSG00000180611.7 | MB21D2 | protein_coding | 2744,379985 | 0,221499503 | 0,059311295 | 3,734524804 | 0,00018807 | 0,000847501 |
| ENSG00000180616.9 | SSTR2 | protein_coding | 16518,02884 | 0,284117374 | 0,045190608 | 6,287089035 | 3,23E-10 | 3,21E-09 |
| ENSG00000180626.10 | ZNF594 | protein_coding | 2372,268368 | 0,387107193 | 0,061688324 | 6,275210087 | 3,49E-10 | 3,45E-09 |
| ENSG00000180628.15 | PCGF5 | protein_coding | 10303,26769 | 1,307030914 | 0,050508706 | 25,87733929 | 1,20E-147 | 1,94E-145 |
| ENSG00000180667.10 | YOD1 | protein_coding | 3946,731706 | 0,139238082 | 0,051882709 | 2,683708771 | 0,00728105 | 0,022718208 |
| ENSG00000180720.7 | CHRM4 | protein_coding | 209,3085515 | 0,91261239 | 0,165278714 | 5,521657137 | 3,36E-08 | 2,63E-07 |
| ENSG00000180745.5 | CLRN3 | protein_coding | 4478,069775 | -0,134944643 | 0,056272452 | -2,398058696 | 0,016482224 | 0,046023865 |
| ENSG00000180747.15 | SMG1P3 | transcribed_unproc | 2901,361486 | 0,552584558 | 0,062344827 | 8,863358519 | 7,76E-19 | 1,48E-17 |
| ENSG00000180773.15 | SLC36A4 | protein_coding | 8181,3351 | -0,19243813 | 0,054389538 | -3,538146079 | 0,000402947 | 0,001703594 |

| | | | | | | | | |
|---|---|---|---|---|---|---|---|---|
| ENSG00000180787.6 | ZFP3 | protein_coding | 1988,571237 | 0,256154868 | 0,068862085 | 3,719824464 | 0,000199361 | 0,000894444 |
| ENSG00000180801.14 | ARSJ | protein_coding | 1060,339584 | 1,113153562 | 0,091350402 | 12,18553549 | 3,71E-34 | 1,38E-32 |
| ENSG00000180815.14 | MAP3K15 | protein_coding | 3721,553322 | -0,290625534 | 0,055768999 | -5,211238111 | 1,88E-07 | 1,34E-06 |
| ENSG00000180817.12 | PPA1 | protein_coding | 13752,44724 | 0,415087398 | 0,054418295 | 7,627717798 | 2,39E-14 | 3,41E-13 |
| ENSG00000180828.2 | BHLHE22 | protein_coding | 89,11381133 | 0,814507828 | 0,25379302 | 3,209338961 | 0,001330405 | 0,005017778 |
| ENSG00000180867.11 | PDIA3P1 | transcribed_process | 1399,153935 | 0,738247034 | 0,071086726 | 10,38516018 | 2,90E-25 | 7,61E-24 |
| ENSG00000180879.13 | SSR4 | protein_coding | 28678,40194 | 0,244291493 | 0,045419873 | 5,378515567 | 7,51E-08 | 5,61E-07 |
| ENSG00000180881.19 | CAPS2 | protein_coding | 1121,324901 | 0,461122397 | 0,097305377 | 4,738920022 | 2,15E-06 | 1,33E-05 |
| ENSG00000180891.13 | CUEDC1 | protein_coding | 796,5529524 | 1,157024989 | 0,106883989 | 10,82505429 | 2,62E-27 | 7,43E-26 |
| ENSG00000180900.19 | SCRIB | protein_coding | 7018,853681 | -0,198151171 | 0,047404155 | -4,180038023 | 2,91E-05 | 0,000151512 |
| ENSG00000180901.11 | KCTD2 | protein_coding | 3254,381862 | 0,228986593 | 0,057420907 | 3,987860937 | 6,67E-05 | 0,000327095 |
| ENSG00000180917.17 | CMTR2 | protein_coding | 6370,180306 | 0,340952707 | 0,056227065 | 6,063853937 | 1,33E-09 | 1,22E-08 |
| ENSG00000180921.7 | FAM83H | protein_coding | 2439,967337 | -0,148345917 | 0,059977132 | -2,473374625 | 0,013384376 | 0,038461433 |
| ENSG00000180938.6 | ZNF572 | protein_coding | 1348,5003 | 0,321198356 | 0,071696869 | 4,479949553 | 7,47E-06 | 4,25E-05 |
| ENSG00000180979.10 | LRRC57 | protein_coding | 4711,579193 | -0,139458776 | 0,052675767 | -2,647493974 | 0,00810908 | 0,024969534 |
| ENSG00000181004.10 | BBS12 | protein_coding | 1373,652965 | 0,260701221 | 0,078490056 | 3,321455413 | 0,000895493 | 0,003507334 |
| ENSG00000181007.8 | ZFP82 | protein_coding | 1342,497276 | 0,262029571 | 0,083930375 | 3,121987379 | 0,001796346 | 0,006575697 |
| ENSG00000181019.13 | NQO1 | protein_coding | 2568,501842 | 0,537340498 | 0,0733413 | 7,326574511 | 2,36E-13 | 3,14E-12 |
| ENSG00000181026.15 | AEN | protein_coding | 1845,637987 | 0,247809697 | 0,070280742 | 3,525997151 | 0,000421892 | 0,001777345 |
| ENSG00000181027.10 | FKRP | protein_coding | 2953,08593 | 0,36819876 | 0,074119732 | 4,967621283 | 6,78E-07 | 4,51E-06 |
| ENSG00000181038.13 | METTL23 | protein_coding | 1946,81111 | -0,259887251 | 0,066323335 | -3,91848888 | 8,91E-05 | 0,000427833 |
| ENSG00000181085.15 | MAPK15 | protein_coding | 37,30965759 | -1,530001815 | 0,406593771 | -3,762974062 | 0,000167905 | 0,000764435 |
| ENSG00000181126.13 | HLA-V | transcribed_unproc | 132,4045072 | 3,219918136 | 0,230255249 | 13,9841248 | 1,95E-44 | 9,90E-43 |
| ENSG00000181163.13 | NPM1 | protein_coding | 86085,27009 | -0,294418994 | 0,04614758 | -6,379944364 | 1,77E-10 | 1,80E-09 |
| ENSG00000181201.3 | HIST3H2BA | unitary_pseudogene | 47,66395793 | 0,928154151 | 0,351811926 | 2,638211168 | 0,008334466 | 0,025569103 |
| ENSG00000181218.5 | HIST3H2A | protein_coding | 762,6200977 | 1,034087187 | 0,113559938 | 9,106091525 | 8,54E-20 | 1,71E-18 |
| ENSG00000181222.16 | POLR2A | protein_coding | 20041,50313 | 0,196917165 | 0,04815108 | 4,089569023 | 4,32E-05 | 0,000219244 |
| ENSG00000181240.13 | SLC25A41 | protein_coding | 32,77240915 | 1,311479093 | 0,422889806 | 3,101231278 | 0,001927177 | 0,006990947 |
| ENSG00000181264.8 | TMEM136 | protein_coding | 1949,245497 | 0,207710458 | 0,078378857 | 2,650082775 | 0,008047205 | 0,024812951 |
| ENSG00000181274.6 | FRAT2 | protein_coding | 893,7896552 | 0,325667739 | 0,090766694 | 3,58796518 | 0,000333269 | 0,00143433 |
| ENSG00000181284.2 | TMEM102 | protein_coding | 655,5470566 | 0,544373794 | 0,098146183 | 5,546561021 | 2,91E-08 | 2,31E-07 |
| ENSG00000181315.11 | ZNF322 | protein_coding | 12842,6358 | 0,222758915 | 0,045995161 | 4,843094548 | 1,28E-06 | 8,19E-06 |
| ENSG00000181350.11 | LRRC75A | protein_coding | 1215,0989 | -0,257301719 | 0,077620421 | -3,314871484 | 0,000916853 | 0,003584755 |
| ENSG00000181378.14 | CFAP65 | protein_coding | 197,8255768 | -0,775891776 | 0,229531473 | -3,380328482 | 0,000723992 | 0,002897643 |
| ENSG00000181381.13 | DDX60L | protein_coding | 30533,32701 | 4,854323925 | 0,06494312 | 74,74731608 | 0 | 0 |
| ENSG00000181396.13 | OGFOD3 | protein_coding | 2739,5952 | 0,350981574 | 0,061432284 | 5,713308232 | 1,11E-08 | 9,16E-08 |

| | | | | | | | | |
|---|---|---|---|---|---|---|---|---|
| ENSG00000181444.13 | ZNF467 | protein_coding | 82,99205202 | 1,254053548 | 0,257398242 | 4,872036185 | 1,10E-06 | 7,14E-06 |
| ENSG00000181450.17 | ZNF678 | protein_coding | 3533,499362 | 0,368053899 | 0,071888777 | 5,119768519 | 3,06E-07 | 2,13E-06 |
| ENSG00000181472.5 | ZBTB2 | protein_coding | 1832,612823 | 0,680046252 | 0,066510896 | 10,22458415 | 1,54E-24 | 3,91E-23 |
| ENSG00000181513.14 | ACBD4 | protein_coding | 376,2895541 | 0,58405286 | 0,12776987 | 4,571131367 | 4,85E-06 | 2,85E-05 |
| ENSG00000181544.15 | FANCB | protein_coding | 1670,325363 | -0,266790918 | 0,071028697 | -3,756100409 | 0,000172581 | 0,000784458 |
| ENSG00000181555.20 | SETD2 | protein_coding | 16925,50613 | 0,489122795 | 0,044775113 | 10,92398798 | 8,85E-28 | 2,56E-26 |
| ENSG00000181585.6 | TMIE | protein_coding | 33,01230485 | 1,132358996 | 0,421257441 | 2,688045088 | 0,007187169 | 0,022462648 |
| ENSG00000181638.18 | ZFP41 | protein_coding | 1942,580647 | 0,427053488 | 0,065406326 | 6,529238268 | 6,61E-11 | 6,99E-10 |
| ENSG00000181788.4 | SIAH2 | protein_coding | 2960,729204 | 0,482168147 | 0,058024463 | 8,309739097 | 9,59E-17 | 1,62E-15 |
| ENSG00000181804.15 | SLC9A9 | protein_coding | 93,94880124 | 1,672914647 | 0,661783586 | 2,527887792 | 0,011475102 | 0,033618778 |
| ENSG00000181826.10 | RELL1 | protein_coding | 2494,369562 | 0,664977532 | 0,058676594 | 11,33292657 | 9,01E-30 | 2,80E-28 |
| ENSG00000181827.14 | RFX7 | protein_coding | 5752,954762 | 0,501547701 | 0,050853033 | 9,862689924 | 6,04E-23 | 1,43E-21 |
| ENSG00000181830.8 | SLC35C1 | protein_coding | 2065,819258 | 0,419592517 | 0,071451816 | 5,872384205 | 4,30E-09 | 3,74E-08 |
| ENSG00000181885.18 | CLDN7 | protein_coding | 12857,03413 | 0,155338675 | 0,045093829 | 3,444787887 | 0,000571508 | 0,002341655 |
| ENSG00000181904.9 | C5orf24 | protein_coding | 16878,99451 | -0,308001857 | 0,046255709 | -6,658677702 | 2,76E-11 | 3,03E-10 |
| ENSG00000181982.18 | CCDC149 | protein_coding | 1638,918227 | 0,265186295 | 0,070795593 | 3,745802284 | 0,000179818 | 0,000813083 |
| ENSG00000181984.11 | GOLGA8CP | unprocessed_pseud | 102,7561302 | -0,661666963 | 0,238410443 | -2,775327095 | 0,005514621 | 0,017793666 |
| ENSG00000182004.13 | SNRPE | protein_coding | 10458,45783 | -0,414924411 | 0,057407907 | -7,227652631 | 4,91E-13 | 6,36E-12 |
| ENSG00000182054.9 | IDH2 | protein_coding | 15824,09868 | -0,442945606 | 0,047845417 | -9,257848235 | 2,09E-20 | 4,34E-19 |
| ENSG00000182093.16 | WRB | protein_coding | 7522,53758 | -0,54704007 | 0,047223357 | -11,58409966 | 4,96E-31 | 1,65E-29 |
| ENSG00000182103.5 | FAM181B | protein_coding | 882,9776989 | 0,844041359 | 0,089955912 | 9,382833619 | 6,42E-21 | 1,37E-19 |
| ENSG00000182107.6 | TMEM30B | protein_coding | 14493,0943 | 0,210709822 | 0,041864676 | 5,033117192 | 4,83E-07 | 3,28E-06 |
| ENSG00000182108.11 | DEXI | protein_coding | 3443,947479 | 0,149782618 | 0,056982241 | 2,628584204 | 0,008574113 | 0,026222106 |
| ENSG00000182118.8 | FAM89A | protein_coding | 461,2823566 | 0,306966319 | 0,116320167 | 2,638977628 | 0,008315647 | 0,025525282 |
| ENSG00000182134.16 | TDRKH | protein_coding | 2416,571197 | 0,211664956 | 0,063099947 | 3,354439531 | 0,000795259 | 0,003153716 |
| ENSG00000182141.10 | ZNF708 | protein_coding | 2800,025593 | 0,157391358 | 0,058253899 | 2,701816708 | 0,006896177 | 0,021688702 |
| ENSG00000182149.20 | IST1 | protein_coding | 19596,6453 | 0,179850315 | 0,042486074 | 4,233159217 | 2,30E-05 | 0,000122013 |
| ENSG00000182150.16 | ERCC6L2 | protein_coding | 5899,55265 | 0,448853993 | 0,061550199 | 7,292486519 | 3,04E-13 | 4,02E-12 |
| ENSG00000182179.13 | UBA7 | protein_coding | 3484,640935 | 8,199593619 | 0,304287485 | 26,94686445 | 6,21E-160 | 1,06E-157 |
| ENSG00000182185.18 | RAD51B | protein_coding | 658,4637899 | 0,447883705 | 0,103111267 | 4,343693165 | 1,40E-05 | 7,66E-05 |
| ENSG00000182196.13 | ARL6IP4 | protein_coding | 8339,488713 | -0,39134396 | 0,054861356 | -7,133326381 | 9,80E-13 | 1,24E-11 |
| ENSG00000182197.11 | EXT1 | protein_coding | 4513,918096 | 1,030887966 | 0,064599066 | 15,95824883 | 2,50E-57 | 1,68E-55 |
| ENSG00000182208.14 | MOB2 | protein_coding | 2316,033415 | -0,159415 | 0,059200688 | -2,692789673 | 0,007085695 | 0,02220099 |
| ENSG00000182218.9 | HHIPL1 | protein_coding | 123,9789374 | -0,905018799 | 0,270597027 | -3,344526021 | 0,000824233 | 0,00325705 |
| ENSG00000182220.14 | ATP6AP2 | protein_coding | 57857,36997 | -0,105366647 | 0,040512821 | -2,600822241 | 0,009300063 | 0,028152778 |
| ENSG00000182240.16 | BACE2 | protein_coding | 1673,156902 | 1,377102509 | 0,107574852 | 12,80134238 | 1,61E-37 | 6,67E-36 |

| | | | | | | | | |
|---|---|---|---|---|---|---|---|---|
| ENSG00000182253.14 | SYNM | protein_coding | 3202,410111 | -0,398271396 | 0,062608619 | -6,361286987 | 2,00E-10 | 2,02E-09 |
| ENSG00000182263.14 | FIGN | protein_coding | 913,0754299 | -0,341278874 | 0,093900776 | -3,634462774 | 0,00027856 | 0,001216082 |
| ENSG00000182307.14 | C8orf33 | protein_coding | 9094,644977 | -0,524628506 | 0,057944082 | -9,054048063 | 1,38E-19 | 2,72E-18 |
| ENSG00000182352.8 | C17orf77 | lncRNA | 14,4438305 | 1,821111821 | 0,715103807 | 2,546639808 | 0,010876563 | 0,032143615 |
| ENSG00000182362.14 | YBEY | protein_coding | 1620,940548 | -0,34228011 | 0,071631724 | -4,77833134 | 1,77E-06 | 1,11E-05 |
| ENSG00000182372.9 | CLN8 | protein_coding | 1972,14984 | -0,498139926 | 0,077461881 | -6,430775006 | 1,27E-10 | 1,31E-09 |
| ENSG00000182378.14 | PLCXD1 | protein_coding | 18351,82809 | -0,541015921 | 0,057314392 | -9,439442678 | 3,75E-21 | 8,08E-20 |
| ENSG00000182389.19 | CACNB4 | protein_coding | 116,9817633 | 1,441701707 | 0,25081902 | 5,747976005 | 9,03E-09 | 7,54E-08 |
| ENSG00000182400.15 | TRAPPC6B | protein_coding | 5451,410456 | 0,334707727 | 0,051203449 | 6,53681995 | 6,28E-11 | 6,68E-10 |
| ENSG00000182446.14 | NPLOC4 | protein_coding | 11466,59651 | 0,252495235 | 0,051770669 | 4,877187055 | 1,08E-06 | 6,96E-06 |
| ENSG00000182481.9 | KPNA2 | protein_coding | 24092,35264 | -0,416877821 | 0,044153246 | -9,441611976 | 3,67E-21 | 7,92E-20 |
| ENSG00000182504.11 | CEP97 | protein_coding | 4678,853724 | -0,458223531 | 0,056218737 | -8,15072619 | 3,62E-16 | 5,87E-15 |
| ENSG00000182512.5 | GLRX5 | protein_coding | 3863,639726 | -0,351933804 | 0,052826997 | -6,662006576 | 2,70E-11 | 2,97E-10 |
| ENSG00000182518.13 | FAM104B | protein_coding | 1362,241646 | -0,607808084 | 0,081366713 | -7,469984481 | 8,02E-14 | 1,11E-12 |
| ENSG00000182534.13 | MXRA7 | protein_coding | 25992,22307 | -0,29624861 | 0,0495476 | -5,979070825 | 2,24E-09 | 2,01E-08 |
| ENSG00000182541.18 | LIMK2 | protein_coding | 4823,457632 | 0,175476251 | 0,064152025 | 2,735318973 | 0,006231985 | 0,019826963 |
| ENSG00000182551.14 | ADI1 | protein_coding | 7528,161469 | -0,410323646 | 0,054718082 | -7,498867484 | 6,44E-14 | 8,97E-13 |
| ENSG00000182552.15 | RWDD4 | protein_coding | 4407,674338 | -0,301553471 | 0,050234996 | -6,002856461 | 1,94E-09 | 1,75E-08 |
| ENSG00000182568.17 | SATB1 | protein_coding | 2068,517479 | 0,717604202 | 0,069410799 | 10,33850945 | 4,72E-25 | 1,23E-23 |
| ENSG00000182575.7 | NXPH3 | protein_coding | 713,7821139 | 0,264027523 | 0,100625619 | 2,62385986 | 0,008693955 | 0,026548939 |
| ENSG00000182628.13 | SKA2 | protein_coding | 8834,775568 | -0,37313261 | 0,046732612 | -7,984415913 | 1,41E-15 | 2,20E-14 |
| ENSG00000182670.13 | TTC3 | protein_coding | 49362,73046 | -0,731651594 | 0,04651241 | -15,73024485 | 9,38E-56 | 6,05E-54 |
| ENSG00000182700.4 | IGIP | protein_coding | 954,1496821 | -0,396525844 | 0,094771845 | -4,184004673 | 2,86E-05 | 0,00014896 |
| ENSG00000182712.16 | CMC4 | protein_coding | 730,3214324 | -0,541719129 | 0,10069087 | -5,380022345 | 7,45E-08 | 5,57E-07 |
| ENSG00000182718.16 | ANXA2 | protein_coding | 870,2018635 | 0,876998142 | 0,101049555 | 8,678891723 | 4,00E-18 | 7,35E-17 |
| ENSG00000182768.8 | NGRN | protein_coding | 16663,92892 | 0,299305064 | 0,043825041 | 6,829544368 | 8,52E-12 | 9,84E-11 |
| ENSG00000182774.13 | RPS17 | protein_coding | 36956,838 | -0,63056817 | 0,039735085 | -15,86930474 | 1,03E-56 | 6,80E-55 |
| ENSG00000182791.5 | CCDC87 | protein_coding | 176,9380694 | 0,45151493 | 0,176179871 | 2,562806565 | 0,010382988 | 0,030887934 |
| ENSG00000182795.13 | C1orf116 | protein_coding | 3090,913321 | 0,510553839 | 0,058967279 | 8,658256665 | 4,79E-18 | 8,75E-17 |
| ENSG00000182798.10 | MAGEB17 | protein_coding | 68,5082499 | 1,089503256 | 0,30636925 | 3,556176915 | 0,000376291 | 0,001602635 |
| ENSG00000182836.10 | PLCXD3 | protein_coding | 63562,05392 | -0,147622048 | 0,048091368 | -3,069616344 | 0,002143339 | 0,007676121 |
| ENSG00000182841.12 | RRP7BP | transcribed_unproc | 1202,817659 | 0,191049376 | 0,078690248 | 2,427865989 | 0,015187954 | 0,042872543 |
| ENSG00000182858.14 | ALG12 | protein_coding | 3546,620172 | 0,396801027 | 0,072921214 | 5,4415033 | 5,28E-08 | 4,04E-07 |
| ENSG00000182872.16 | RBM10 | protein_coding | 10229,01659 | 0,13786084 | 0,054502974 | 2,529418658 | 0,011425165 | 0,033494275 |
| ENSG00000182899.17 | RPL35A | protein_coding | 29466,3273 | -0,462006263 | 0,044043459 | -10,48978146 | 9,63E-26 | 2,58E-24 |
| ENSG00000182916.8 | TCEAL7 | protein_coding | 3721,980121 | 0,162156035 | 0,05383983 | 3,011822956 | 0,00259684 | 0,009117497 |

| | | | | | | | | |
|---|---|---|---|---|---|---|---|---|
| ENSG00000182952.5 | HMGN4 | protein_coding | 5894,398921 | 0,207321932 | 0,061722318 | 3,35894598 | 0,000782404 | 0,003106566 |
| ENSG00000182957.16 | SPATA13 | protein_coding | 11408,16912 | 0,529976554 | 0,04653852 | 11,38791174 | 4,80E-30 | 1,52E-28 |
| ENSG00000182979.18 | MTA1 | protein_coding | 2580,382153 | -0,251113046 | 0,062877097 | -3,993712494 | 6,50E-05 | 0,00031996 |
| ENSG00000182986.13 | ZNF320 | protein_coding | 161,1273917 | 1,053437589 | 0,214520052 | 4,910671897 | 9,08E-07 | 5,93E-06 |
| ENSG00000183010.16 | PYCR1 | protein_coding | 4399,6799 | -0,661641674 | 0,074669732 | -8,860908636 | 7,94E-19 | 1,51E-17 |
| ENSG00000183011.13 | NAA38 | protein_coding | 2657,944685 | -0,337667975 | 0,058271757 | -5,794710745 | 6,84E-09 | 5,81E-08 |
| ENSG00000183018.9 | AC118754.1 | protein_coding | 738,6051986 | -0,597265668 | 0,092633438 | -6,447624959 | 1,14E-10 | 1,18E-09 |
| ENSG00000183023.18 | SLC8A1 | protein_coding | 1484,537781 | 0,696118615 | 0,085825313 | 8,110877639 | 5,03E-16 | 8,09E-15 |
| ENSG00000183036.11 | PCP4 | protein_coding | 5803,447572 | -0,179415963 | 0,056593227 | -3,170272698 | 0,001522959 | 0,005673666 |
| ENSG00000183044.12 | ABAT | protein_coding | 29611,3378 | -0,776340118 | 0,052893033 | -14,6775497 | 8,98E-49 | 4,93E-47 |
| ENSG00000183048.12 | SLC25A10 | protein_coding | 3138,79429 | -0,3084782 | 0,065473863 | -4,71147087 | 2,46E-06 | 1,51E-05 |
| ENSG00000183077.15 | AFMID | protein_coding | 857,2742044 | -0,385065094 | 0,099652648 | -3,864072892 | 0,000111512 | 0,000525659 |
| ENSG00000183098.11 | GPC6 | protein_coding | 7115,926216 | -0,39333072 | 0,052362957 | -7,511621635 | 5,84E-14 | 8,16E-13 |
| ENSG00000183145.9 | RIPPLY3 | protein_coding | 1965,685154 | -0,177450168 | 0,067813046 | -2,616755588 | 0,008876988 | 0,027034518 |
| ENSG00000183161.5 | FANCF | protein_coding | 5443,048122 | 0,172052658 | 0,052949288 | 3,249385694 | 0,001156545 | 0,004416811 |
| ENSG00000183172.8 | SMDT1 | protein_coding | 4264,319781 | -0,324290807 | 0,05728485 | -5,661022233 | 1,50E-08 | 1,23E-07 |
| ENSG00000183186.8 | C2CD4C | protein_coding | 1114,562846 | -0,213259415 | 0,08528073 | -2,500675317 | 0,012395676 | 0,036006488 |
| ENSG00000183207.14 | RUVBL2 | protein_coding | 5379,534114 | -0,255727314 | 0,062687248 | -4,079415247 | 4,51E-05 | 0,000228015 |
| ENSG00000183248.12 | PRR36 | protein_coding | 1118,763916 | -0,21780588 | 0,085892179 | -2,535805724 | 0,011218895 | 0,033014265 |
| ENSG00000183258.12 | DDX41 | protein_coding | 5571,213352 | -0,226653801 | 0,082954736 | -2,732258723 | 0,006290172 | 0,019992303 |
| ENSG00000183287.14 | CCBE1 | protein_coding | 193,8938874 | 0,524166567 | 0,175111286 | 2,993334011 | 0,002759476 | 0,00961739 |
| ENSG00000183307.3 | TMEM121B | protein_coding | 3043,039897 | -0,170635658 | 0,057712085 | -2,95667116 | 0,003109796 | 0,010691675 |
| ENSG00000183323.12 | CCDC125 | protein_coding | 2763,753205 | 0,55691421 | 0,063652507 | 8,749289523 | 2,15E-18 | 4,01E-17 |
| ENSG00000183340.7 | JRKL | protein_coding | 2738,023882 | 0,214081957 | 0,059859247 | 3,576422504 | 0,000348329 | 0,001492865 |
| ENSG00000183354.12 | KIAA2026 | protein_coding | 5390,561415 | 0,311511363 | 0,054485824 | 5,717291968 | 1,08E-08 | 8,97E-08 |
| ENSG00000183421.12 | RIPK4 | protein_coding | 2949,331537 | 0,359564385 | 0,061156114 | 5,879451171 | 4,12E-09 | 3,59E-08 |
| ENSG00000183426.17 | NPIPA1 | protein_coding | 10600,18164 | 0,458374011 | 0,067831631 | 6,757526018 | 1,40E-11 | 1,58E-10 |
| ENSG00000183439.8 | TRIM61 | protein_coding | 224,0451283 | 0,723810535 | 0,198049101 | 3,654702457 | 0,00025748 | 0,001132179 |
| ENSG00000183454.17 | GRIN2A | protein_coding | 916,3944784 | -1,063508431 | 0,243611996 | -4,365583174 | 1,27E-05 | 6,99E-05 |
| ENSG00000183458.14 | AC138932.1 | transcribed_unproc | 4586,867991 | 0,759423133 | 0,068760134 | 11,04452659 | 2,33E-28 | 6,90E-27 |
| ENSG00000183475.12 | ASB7 | protein_coding | 2236,67729 | 0,207751759 | 0,064737434 | 3,209144152 | 0,001331307 | 0,005019955 |
| ENSG00000183486.13 | MX2 | protein_coding | 9207,7839 | 2,865913142 | 0,14628371 | 19,59147149 | 1,83E-85 | 1,84E-83 |
| ENSG00000183496.6 | MEX3B | protein_coding | 1309,156353 | -0,211299564 | 0,085755444 | -2,463978431 | 0,013740438 | 0,039389139 |
| ENSG00000183530.14 | PRR14L | protein_coding | 11292,49529 | 0,183519328 | 0,044799429 | 4,096465787 | 4,20E-05 | 0,000213152 |
| ENSG00000183576.13 | SETD3 | protein_coding | 6467,428142 | -0,142932001 | 0,051585551 | -2,770775868 | 0,005592291 | 0,018023607 |
| ENSG00000183604.14 | SMG1P5 | transcribed_unproc | 661,3695417 | 0,380038039 | 0,099516382 | 3,818849044 | 0,000134076 | 0,000622892 |

| | | | | | | | | |
|---|---|---|---|---|---|---|---|---|
| ENSG00000183621.15 | ZNF438 | protein_coding | 177,9955213 | 0,499490341 | 0,192777123 | 2,591024983 | 0,009569054 | 0,028841704 |
| ENSG00000183624.14 | HMCES | protein_coding | 5237,485388 | -0,137157246 | 0,050328545 | -2,725237656 | 0,006425522 | 0,02036211 |
| ENSG00000183648.10 | NDUFB1 | protein_coding | 4447,461471 | -0,184413954 | 0,062585701 | -2,946582858 | 0,003213063 | 0,011018105 |
| ENSG00000183655.13 | KLHL25 | protein_coding | 1564,926557 | 0,222733853 | 0,078025977 | 2,854611505 | 0,00430895 | 0,014311666 |
| ENSG00000183682.8 | BMP8A | protein_coding | 1152,840778 | -0,490949165 | 0,095112932 | -5,161749887 | 2,45E-07 | 1,73E-06 |
| ENSG00000183684.7 | ALYREF | protein_coding | 2910,531633 | -0,1744772 | 0,066742228 | -2,614195032 | 0,008943797 | 0,027208531 |
| ENSG00000183688.4 | RFLNB | protein_coding | 3470,050922 | -1,579402642 | 0,062167375 | -25,405651 | 2,18E-142 | 3,41E-140 |
| ENSG00000183715.13 | OPCML | protein_coding | 285,0445988 | -0,950307854 | 0,155681973 | -6,104161175 | 1,03E-09 | 9,64E-09 |
| ENSG00000183722.9 | LHFPL6 | protein_coding | 2284,925444 | 0,675110315 | 0,06775245 | 9,964367525 | 2,18E-23 | 5,25E-22 |
| ENSG00000183723.12 | CMTM4 | protein_coding | 12303,9035 | -1,77539589 | 0,082879569 | -21,42139378 | 8,44E-102 | 9,95E-100 |
| ENSG00000183726.11 | TMEM50A | protein_coding | 14355,56297 | 0,330316512 | 0,043874633 | 7,528644385 | 5,13E-14 | 7,21E-13 |
| ENSG00000183735.10 | TBK1 | protein_coding | 5836,186606 | 0,526999026 | 0,049438797 | 10,6596247 | 1,57E-26 | 4,36E-25 |
| ENSG00000183741.11 | CBX6 | protein_coding | 14623,03427 | -0,221461805 | 0,049136272 | -4,507094159 | 6,57E-06 | 3,78E-05 |
| ENSG00000183742.12 | MACC1 | protein_coding | 123,9907496 | 0,681401986 | 0,261536914 | 2,605375953 | 0,00917735 | 0,027817529 |
| ENSG00000183760.10 | ACP7 | protein_coding | 305,5952371 | 0,499826517 | 0,143780776 | 3,476309775 | 0,000508365 | 0,002101521 |
| ENSG00000183762.12 | KREMEN1 | protein_coding | 1019,281933 | -0,82893442 | 0,098785008 | -8,391297751 | 4,81E-17 | 8,23E-16 |
| ENSG00000183763.9 | TRAIP | protein_coding | 1136,166291 | -0,222704706 | 0,090839074 | -2,451639993 | 0,014220686 | 0,040584939 |
| ENSG00000183778.17 | B3GALT5 | protein_coding | 2483,474815 | 0,345991194 | 0,064232813 | 5,386517862 | 7,18E-08 | 5,38E-07 |
| ENSG00000183780.13 | SLC35F3 | protein_coding | 1280,153673 | -0,225677679 | 0,083323108 | -2,708464481 | 0,006759535 | 0,021297644 |
| ENSG00000183793.13 | NPIPA5 | protein_coding | 2293,298213 | 0,702995941 | 0,091311734 | 7,698856574 | 1,37E-14 | 2,00E-13 |
| ENSG00000183801.8 | OLFML1 | protein_coding | 300,0454016 | -0,411539629 | 0,14820629 | -2,776802716 | 0,005489649 | 0,017723251 |
| ENSG00000183808.11 | RBM12B | protein_coding | 6852,334767 | 0,117759858 | 0,047585395 | 2,474705896 | 0,013334593 | 0,038352717 |
| ENSG00000183831.6 | ANKRD45 | protein_coding | 91,69532048 | -0,869019725 | 0,248651484 | -3,494930787 | 0,000474185 | 0,00197612 |
| ENSG00000183844.16 | FAM3B | protein_coding | 2311,080962 | 1,307008923 | 0,061574683 | 21,22640107 | 5,45E-100 | 6,35E-98 |
| ENSG00000183856.11 | IQGAP3 | protein_coding | 4381,15157 | -0,333078198 | 0,061482432 | -5,41745316 | 6,05E-08 | 4,59E-07 |
| ENSG00000183864.5 | TOB2 | protein_coding | 3617,092094 | 0,124245977 | 0,052404512 | 2,370902275 | 0,017744722 | 0,048996069 |
| ENSG00000183878.15 | UTY | protein_coding | 2604,996467 | 0,279154146 | 0,059951429 | 4,656338489 | 3,22E-06 | 1,94E-05 |
| ENSG00000183889.12 | AC138969.1 | protein_coding | 11794,54229 | 0,451553158 | 0,047804567 | 9,445816307 | 3,53E-21 | 7,63E-20 |
| ENSG00000183891.6 | TTC32 | protein_coding | 2040,724854 | 0,25952411 | 0,066379448 | 3,909705765 | 9,24E-05 | 0,000442559 |
| ENSG00000183943.5 | PRKX | protein_coding | 5003,276708 | -0,472221425 | 0,070894354 | -6,660917261 | 2,72E-11 | 2,99E-10 |
| ENSG00000183955.13 | KMT5A | protein_coding | 1924,38575 | -0,254453321 | 0,07481441 | -3,401127158 | 0,000671086 | 0,002706105 |
| ENSG00000183960.9 | KCNH8 | protein_coding | 1409,183433 | 0,28308076 | 0,090217633 | 3,137754253 | 0,001702475 | 0,006262628 |
| ENSG00000183963.18 | SMTN | protein_coding | 4542,502675 | -0,401692836 | 0,072238891 | -5,560617461 | 2,69E-08 | 2,14E-07 |
| ENSG00000183978.8 | COA3 | protein_coding | 8261,806578 | -0,144499363 | 0,048464459 | -2,981553187 | 0,002867902 | 0,009957367 |
| ENSG00000184005.11 | ST6GALNAC3 | protein_coding | 322,0342711 | -1,213549184 | 0,372163924 | -3,260792102 | 0,001111015 | 0,004263325 |
| ENSG00000184007.21 | PTP4A2 | protein_coding | 24566,18444 | -0,222385506 | 0,053038835 | -4,192880649 | 2,75E-05 | 0,000143646 |

| | | | | | | | | |
|---|---|---|---|---|---|---|---|---|
| ENSG00000184009.12 | ACTG1 | protein_coding | 196385,174 | -0,190694107 | 0,045325815 | -4,207185419 | 2,59E-05 | 0,000135639 |
| ENSG00000184014.8 | DENND5A | protein_coding | 6604,599618 | 0,481741319 | 0,055472592 | 8,684312333 | 3,81E-18 | 7,02E-17 |
| ENSG00000184076.13 | UQCR10 | protein_coding | 9508,715778 | -0,260717471 | 0,048694635 | -5,354131326 | 8,60E-08 | 6,38E-07 |
| ENSG00000184083.12 | FAM120C | protein_coding | 1080,085251 | 0,434091159 | 0,102954068 | 4,216357537 | 2,48E-05 | 0,000130818 |
| ENSG00000184117.11 | NIPSNAP1 | protein_coding | 13474,59076 | -0,160363767 | 0,046505529 | -3,448273138 | 0,000564183 | 0,002314416 |
| ENSG00000184163.3 | C1QTNF12 | protein_coding | 28,20158916 | -2,048915918 | 0,4937145 | -4,150001506 | 3,32E-05 | 0,000171329 |
| ENSG00000184164.14 | CRELD2 | protein_coding | 2339,611325 | 0,500493723 | 0,064699499 | 7,735666119 | 1,03E-14 | 1,51E-13 |
| ENSG00000184208.11 | C22orf46 | protein_coding | 2414,905039 | 0,159423223 | 0,061513406 | 2,591682568 | 0,009550785 | 0,028802066 |
| ENSG00000184209.14 | SNRNP35 | protein_coding | 2048,645665 | 0,149166638 | 0,062008934 | 2,405566871 | 0,016147396 | 0,045227693 |
| ENSG00000184216.14 | IRAK1 | protein_coding | 9601,719255 | -0,228681075 | 0,058804513 | -3,888835449 | 0,000100726 | 0,000478929 |
| ENSG00000184220.12 | CMSS1 | protein_coding | 1137,257723 | -0,410672305 | 0,082679062 | -4,967065352 | 6,80E-07 | 4,52E-06 |
| ENSG00000184226.15 | PCDH9 | protein_coding | 2619,604193 | 0,178042667 | 0,060864041 | 2,925252164 | 0,003441772 | 0,011734451 |
| ENSG00000184227.8 | ACOT1 | protein_coding | 464,4904672 | 0,791395759 | 0,116527555 | 6,791490299 | 1,11E-11 | 1,27E-10 |
| ENSG00000184260.5 | HIST2H2AC | protein_coding | 52,38734585 | 0,873150523 | 0,346342741 | 2,521059116 | 0,011700219 | 0,034211514 |
| ENSG00000184277.13 | TM2D3 | protein_coding | 5711,928127 | 0,299509608 | 0,053642865 | 5,583400651 | 2,36E-08 | 1,88E-07 |
| ENSG00000184302.6 | SIX6 | protein_coding | 2381,795164 | 0,172120761 | 0,067796392 | 2,538789387 | 0,011123676 | 0,032782285 |
| ENSG00000184319.16 | RPL23AP82 | transcribed_unproc | 1894,748538 | -0,238752227 | 0,065254559 | -3,658782328 | 0,000253416 | 0,001116263 |
| ENSG00000184349.13 | EFNA5 | protein_coding | 1160,310583 | -0,220882671 | 0,087453279 | -2,525721991 | 0,011546081 | 0,033795934 |
| ENSG00000184368.16 | MAP7D2 | protein_coding | 19372,9838 | -0,168625073 | 0,042894099 | -3,93119516 | 8,45E-05 | 0,000408363 |
| ENSG00000184371.14 | CSF1 | protein_coding | 194,7465759 | 3,254754055 | 0,278269621 | 11,69640452 | 1,33E-31 | 4,56E-30 |
| ENSG00000184381.19 | PLA2G6 | protein_coding | 844,8401872 | 0,304076792 | 0,097294807 | 3,125313684 | 0,001776155 | 0,006508137 |
| ENSG00000184384.14 | MAML2 | protein_coding | 1820,796533 | 0,920019001 | 0,067306398 | 13,66911665 | 1,55E-42 | 7,53E-41 |
| ENSG00000184402.15 | SS18L1 | protein_coding | 12588,07347 | -0,443598688 | 0,043489171 | -10,20021025 | 1,98E-24 | 5,01E-23 |
| ENSG00000184445.12 | KNTC1 | protein_coding | 14482,52155 | -0,153375136 | 0,044723177 | -3,429432903 | 0,000604844 | 0,002461478 |
| ENSG00000184486.9 | POU3F2 | protein_coding | 1826,815387 | 0,426308022 | 0,067002436 | 6,362574945 | 1,98E-10 | 2,00E-09 |
| ENSG00000184489.11 | PTP4A3 | protein_coding | 207,2028943 | -0,504380951 | 0,178243005 | -2,829737695 | 0,004658618 | 0,015314858 |
| ENSG00000184524.6 | CEND1 | protein_coding | 110,6804891 | -1,002339657 | 0,232708747 | -4,307271078 | 1,65E-05 | 8,94E-05 |
| ENSG00000184545.11 | DUSP8 | protein_coding | 2914,807819 | 0,284297123 | 0,078660523 | 3,614228749 | 0,000301243 | 0,001308001 |
| ENSG00000184564.11 | SLITRK6 | protein_coding | 2447,723497 | -0,474979923 | 0,065508407 | -7,250671224 | 4,15E-13 | 5,40E-12 |
| ENSG00000184575.12 | XPOT | protein_coding | 10555,08418 | 0,180025771 | 0,044756357 | 4,022350887 | 5,76E-05 | 0,000285488 |
| ENSG00000184588.18 | PDE4B | protein_coding | 686,9851083 | 0,435749446 | 0,106683835 | 4,08449366 | 4,42E-05 | 0,000223587 |
| ENSG00000184602.6 | SNN | protein_coding | 7864,636168 | -0,298431003 | 0,047273605 | -6,312846372 | 2,74E-10 | 2,73E-09 |
| ENSG00000184611.11 | KCNH7 | protein_coding | 3459,259555 | 0,162289621 | 0,054288884 | 2,989371097 | 0,002795524 | 0,009731587 |
| ENSG00000184613.10 | NELL2 | protein_coding | 522,9017809 | -0,594916243 | 0,114649579 | -5,188996314 | 2,11E-07 | 1,50E-06 |
| ENSG00000184634.16 | MED12 | protein_coding | 5193,922022 | -0,210895718 | 0,067575423 | -3,120893787 | 0,001803031 | 0,006595874 |
| ENSG00000184635.15 | ZNF93 | protein_coding | 1384,809819 | 0,306898582 | 0,077101109 | 3,980469116 | 6,88E-05 | 0,000336995 |

| | | | | | | | | |
|---|---|---|---|---|---|---|---|---|
| ENSG00000184640.18 | sept-09 | protein_coding | 10641,28092 | -0,309297374 | 0,056333529 | -5,49046687 | 4,01E-08 | 3,11E-07 |
| ENSG00000184661.14 | CDCA2 | protein_coding | 3086,396217 | -0,553181416 | 0,058365351 | -9,477907834 | 2,59E-21 | 5,64E-20 |
| ENSG00000184675.11 | AMER1 | protein_coding | 905,6014351 | 0,941707268 | 0,088410301 | 10,65155592 | 1,71E-26 | 4,74E-25 |
| ENSG00000184677.18 | ZBTB40 | protein_coding | 8735,262168 | 0,263989197 | 0,048543067 | 5,438247164 | 5,38E-08 | 4,11E-07 |
| ENSG00000184678.10 | HIST2H2BE | protein_coding | 1900,735834 | 0,583118147 | 0,071800352 | 8,121382851 | 4,61E-16 | 7,44E-15 |
| ENSG00000184702.20 | sept-05 | protein_coding | 5628,126844 | -0,402914402 | 0,054387415 | -7,408228522 | 1,28E-13 | 1,74E-12 |
| ENSG00000184708.18 | EIF4ENIF1 | protein_coding | 4217,690168 | 0,254600098 | 0,081393903 | 3,127999624 | 0,001760004 | 0,006457365 |
| ENSG00000184730.11 | APOBR | protein_coding | 62,72141328 | 1,444518124 | 0,304155982 | 4,749267512 | 2,04E-06 | 1,27E-05 |
| ENSG00000184731.6 | FAM110C | protein_coding | 3868,928265 | 0,671557045 | 0,057121553 | 11,75663134 | 6,53E-32 | 2,28E-30 |
| ENSG00000184743.12 | ATL3 | protein_coding | 6672,13082 | -0,330164161 | 0,050103886 | -6,589591882 | 4,41E-11 | 4,76E-10 |
| ENSG00000184752.13 | NDUFA12 | protein_coding | 6347,664904 | -0,174967223 | 0,056320799 | -3,106618281 | 0,001892406 | 0,006879225 |
| ENSG00000184787.19 | UBE2G2 | protein_coding | 7874,774 | -0,2429968 | 0,057037745 | -4,260280607 | 2,04E-05 | 0,000108825 |
| ENSG00000184792.16 | OSBP2 | protein_coding | 6203,098129 | -0,748398754 | 0,073710878 | -10,15316559 | 3,21E-24 | 8,05E-23 |
| ENSG00000184831.13 | APOO | protein_coding | 5635,495338 | -0,251032305 | 0,053979945 | -4,650473489 | 3,31E-06 | 1,99E-05 |
| ENSG00000184840.11 | TMED9 | protein_coding | 13610,89712 | 0,226933979 | 0,043227076 | 5,249811039 | 1,52E-07 | 1,10E-06 |
| ENSG00000184867.14 | ARMCX2 | protein_coding | 4992,298041 | 0,464220747 | 0,064798175 | 7,164102237 | 7,83E-13 | 9,98E-12 |
| ENSG00000184887.13 | BTBD6 | protein_coding | 5280,196551 | -0,371421546 | 0,055212139 | -6,727171814 | 1,73E-11 | 1,94E-10 |
| ENSG00000184895.7 | SRY | protein_coding | 524,4087633 | 0,27810233 | 0,113062918 | 2,459713007 | 0,013904816 | 0,039795481 |
| ENSG00000184898.7 | RBM43 | protein_coding | 2331,246639 | 2,550540985 | 0,063257984 | 40,31966891 | 0 | 0 |
| ENSG00000184903.9 | IMMP2L | protein_coding | 599,2223982 | -0,541156577 | 0,114767531 | -4,715241074 | 2,41E-06 | 1,48E-05 |
| ENSG00000184905.9 | TCEAL2 | protein_coding | 3688,088999 | 0,189944949 | 0,058287333 | 3,258768913 | 0,001118968 | 0,004290043 |
| ENSG00000184916.9 | JAG2 | protein_coding | 810,773441 | -0,684221893 | 0,140638756 | -4,865101992 | 1,14E-06 | 7,37E-06 |
| ENSG00000184923.12 | NUTM2A | protein_coding | 386,5842784 | 0,74675945 | 0,130675655 | 5,714602678 | 1,10E-08 | 9,10E-08 |
| ENSG00000184924.5 | PTRHD1 | protein_coding | 1617,169413 | -0,541361783 | 0,077880057 | -6,951224774 | 3,62E-12 | 4,34E-11 |
| ENSG00000184925.12 | LCN12 | protein_coding | 430,7423435 | -0,35135617 | 0,144001188 | -2,43995327 | 0,014689162 | 0,0416736 |
| ENSG00000184949.16 | FAM227A | protein_coding | 1279,923585 | -0,456748255 | 0,096372134 | -4,739422431 | 2,14E-06 | 1,33E-05 |
| ENSG00000184979.9 | USP18 | protein_coding | 21062,66157 | 4,363581996 | 0,052182257 | 83,62194762 | 0 | 0 |
| ENSG00000184988.8 | TMEM106A | protein_coding | 123,0755948 | 1,34673153 | 0,216617945 | 6,21708201 | 5,06E-10 | 4,91E-09 |
| ENSG00000184990.13 | SIVA1 | protein_coding | 3970,62133 | -0,374599221 | 0,069739013 | -5,371444227 | 7,81E-08 | 5,83E-07 |
| ENSG00000184992.11 | BRI3BP | protein_coding | 13929,06811 | -1,034616168 | 0,057809676 | -17,89693769 | 1,25E-71 | 1,02E-69 |
| ENSG00000185000.12 | DGAT1 | protein_coding | 2476,471331 | -0,271805113 | 0,069663665 | -3,901676907 | 9,55E-05 | 0,000456142 |
| ENSG00000185002.10 | RFX6 | protein_coding | 17757,45875 | 0,340630493 | 0,044084924 | 7,726688893 | 1,10E-14 | 1,62E-13 |
| ENSG00000185008.17 | ROBO2 | protein_coding | 14607,66896 | 0,126727425 | 0,046967748 | 2,698179695 | 0,00697198 | 0,02189651 |
| ENSG00000185010.15 | F8 | protein_coding | 412,0434605 | -0,462548017 | 0,148864167 | -3,107181711 | 0,001888803 | 0,006867236 |
| ENSG00000185015.8 | CA13 | protein_coding | 559,5355057 | 0,540464495 | 0,112161005 | 4,818648816 | 1,45E-06 | 9,17E-06 |
| ENSG00000185022.12 | MAFF | protein_coding | 242,8253051 | 1,185613889 | 0,154819743 | 7,658027733 | 1,89E-14 | 2,72E-13 |

| | | | | | | | | |
|---|---|---|---|---|---|---|---|---|
| ENSG00000185046.18 | ANKS1B | protein_coding | 970,1859913 | -0,297254754 | 0,110334683 | -2,694118891 | 0,007057499 | 0,022128045 |
| ENSG00000185101.13 | ANO9 | protein_coding | 2741,266858 | 0,907384253 | 0,091182067 | 9,951345529 | 2,49E-23 | 5,96E-22 |
| ENSG00000185104.20 | FAF1 | protein_coding | 7769,338991 | -0,212603964 | 0,05735287 | -3,706945514 | 0,000209774 | 0,000936866 |
| ENSG00000185122.11 | HSF1 | protein_coding | 2537,273337 | 0,23196686 | 0,072176047 | 3,21390362 | 0,001309436 | 0,004944489 |
| ENSG00000185158.12 | LRRC37B | protein_coding | 1343,11593 | 0,431188166 | 0,084331435 | 5,113018273 | 3,17E-07 | 2,21E-06 |
| ENSG00000185164.14 | NOMO2 | protein_coding | 35842,87544 | 0,434400984 | 0,038538309 | 11,2719263 | 1,81E-29 | 5,55E-28 |
| ENSG00000185201.16 | IFITM2 | protein_coding | 8,908962914 | 3,421217375 | 0,974019613 | 3,512472778 | 0,000443957 | 0,001865417 |
| ENSG00000185215.9 | TNFAIP2 | protein_coding | 1000,055258 | 0,238350071 | 0,097725218 | 2,438982251 | 0,014728692 | 0,041769953 |
| ENSG00000185219.17 | ZNF445 | protein_coding | 12226,0177 | 0,115055777 | 0,045160783 | 2,547692272 | 0,010843808 | 0,032061378 |
| ENSG00000185222.10 | TCEAL9 | protein_coding | 8893,009258 | 0,199649572 | 0,053189948 | 3,753520751 | 0,000174368 | 0,00079162 |
| ENSG00000185236.12 | RAB11B | protein_coding | 5138,964421 | -0,119466074 | 0,04853421 | -2,461481802 | 0,013836442 | 0,03962902 |
| ENSG00000185269.12 | NOTUM | protein_coding | 176,5467133 | -1,59800805 | 0,207667848 | -7,695019087 | 1,41E-14 | 2,06E-13 |
| ENSG00000185272.14 | RBM11 | protein_coding | 600,7779344 | 1,856675837 | 0,110070705 | 16,86802892 | 7,73E-64 | 5,71E-62 |
| ENSG00000185278.15 | ZBTB37 | protein_coding | 1601,272273 | -0,250649472 | 0,084758612 | -2,957215373 | 0,003104312 | 0,010677714 |
| ENSG00000185305.11 | ARL15 | protein_coding | 5544,339806 | -0,28550009 | 0,057034723 | -5,005724158 | 5,57E-07 | 3,75E-06 |
| ENSG00000185324.22 | CDK10 | protein_coding | 6233,884679 | -0,275327931 | 0,069510106 | -3,960976975 | 7,46E-05 | 0,000363595 |
| ENSG00000185338.5 | SOCS1 | protein_coding | 97,47632869 | 2,52665729 | 0,3075513 | 8,21540111 | 2,11E-16 | 3,48E-15 |
| ENSG00000185339.8 | TCN2 | protein_coding | 5986,895938 | 0,238607009 | 0,054140681 | 4,40716678 | 1,05E-05 | 5,85E-05 |
| ENSG00000185344.14 | ATP6V0A2 | protein_coding | 3288,644772 | 0,203046956 | 0,062996907 | 3,223125798 | 0,001267999 | 0,00479687 |
| ENSG00000185352.8 | HS6ST3 | protein_coding | 23379,41389 | -0,505674978 | 0,053129522 | -9,517777696 | 1,77E-21 | 3,87E-20 |
| ENSG00000185361.9 | TNFAIP8L1 | protein_coding | 526,9569269 | -0,780891171 | 0,121286423 | -6,43840547 | 1,21E-10 | 1,25E-09 |
| ENSG00000185404.16 | SP140L | protein_coding | 177,6142807 | 1,435144199 | 0,233752575 | 6,139586683 | 8,27E-10 | 7,82E-09 |
| ENSG00000185414.20 | MRPL30 | protein_coding | 7258,606885 | -0,262603699 | 0,06058542 | -4,334437219 | 1,46E-05 | 7,96E-05 |
| ENSG00000185418.16 | TARSL2 | protein_coding | 1587,4673 | -0,364074701 | 0,071314821 | -5,105175836 | 3,30E-07 | 2,29E-06 |
| ENSG00000185420.19 | SMYD3 | protein_coding | 2575,640274 | -0,292583232 | 0,077748575 | -3,763197338 | 0,000167755 | 0,000763907 |
| ENSG00000185436.12 | IFNLR1 | protein_coding | 2415,175993 | 0,45304733 | 0,066762791 | 6,785925577 | 1,15E-11 | 1,31E-10 |
| ENSG00000185442.13 | FAM174B | protein_coding | 7164,213845 | 0,280184878 | 0,04853874 | 5,772397056 | 7,82E-09 | 6,57E-08 |
| ENSG00000185475.11 | TMEM179B | protein_coding | 2154,370235 | 0,325727385 | 0,075336507 | 4,323632709 | 1,53E-05 | 8,34E-05 |
| ENSG00000185477.5 | GPRIN3 | protein_coding | 9517,96815 | 0,176505084 | 0,0451364 | 3,910482077 | 9,21E-05 | 0,000441326 |
| ENSG00000185480.11 | PARPBP | protein_coding | 2455,031439 | -0,590896751 | 0,092314002 | -6,40094393 | 1,54E-10 | 1,58E-09 |
| ENSG00000185483.12 | ROR1 | protein_coding | 2337,637466 | -0,241235188 | 0,074302062 | -3,246682259 | 0,001167587 | 0,004452932 |
| ENSG00000185507.21 | IRF7 | protein_coding | 4154,028711 | 4,05737482 | 0,085739115 | 47,32233155 | 0 | 0 |
| ENSG00000185513.16 | L3MBTL1 | protein_coding | 1881,356882 | -0,188616266 | 0,066469099 | -2,837653414 | 0,00454465 | 0,014988356 |
| ENSG00000185515.14 | BRCC3 | protein_coding | 9436,543002 | -0,246080942 | 0,045377745 | -5,42294334 | 5,86E-08 | 4,45E-07 |
| ENSG00000185565.12 | LSAMP | protein_coding | 5095,368266 | 0,286278173 | 0,050174603 | 5,705639016 | 1,16E-08 | 9,56E-08 |
| ENSG00000185567.7 | AHNAK2 | protein_coding | 242,9070289 | -1,017704503 | 0,18396033 | -5,53219547 | 3,16E-08 | 2,49E-07 |

| | | | | | | | | |
|---|---|---|---|---|---|---|---|---|
| ENSG00000185591.10 | SP1 | protein_coding | 12223,41499 | 0,123405881 | 0,046033296 | 2,680796117 | 0,007344725 | 0,022888331 |
| ENSG00000185596.16 | WASH3P | transcribed_unproc | 2384,930801 | 0,284216152 | 0,074703592 | 3,804584804 | 0,000142042 | 0,000657325 |
| ENSG00000185608.8 | MRPL40 | protein_coding | 3745,501414 | 0,339410735 | 0,060065433 | 5,650683228 | 1,60E-08 | 1,30E-07 |
| ENSG00000185614.5 | INKA1 | protein_coding | 25,23650884 | 1,798474229 | 0,503838226 | 3,569547002 | 0,000357599 | 0,001529103 |
| ENSG00000185619.18 | PCGF3 | protein_coding | 8372,594292 | -0,171832949 | 0,057129779 | -3,007764963 | 0,002631766 | 0,009222858 |
| ENSG00000185624.15 | P4HB | protein_coding | 96858,30427 | 0,278093354 | 0,049966924 | 5,565548765 | 2,61E-08 | 2,08E-07 |
| ENSG00000185630.18 | PBX1 | protein_coding | 7079,66054 | -0,293018932 | 0,053020311 | -5,526541126 | 3,27E-08 | 2,57E-07 |
| ENSG00000185633.10 | NDUFA4L2 | protein_coding | 34,62669717 | 2,043905263 | 0,506815145 | 4,032841725 | 5,51E-05 | 0,000274362 |
| ENSG00000185650.9 | ZFP36L1 | protein_coding | 202,6386567 | 0,78278829 | 0,183333378 | 4,269753277 | 1,96E-05 | 0,000104552 |
| ENSG00000185658.13 | BRWD1 | protein_coding | 11548,42109 | 0,137377745 | 0,045917241 | 2,991855368 | 0,002772876 | 0,009657228 |
| ENSG00000185686.18 | PRAME | protein_coding | 1374,426779 | 0,628712456 | 0,093556102 | 6,720165121 | 1,82E-11 | 2,03E-10 |
| ENSG00000185697.16 | MYBL1 | protein_coding | 2416,445178 | -0,209719833 | 0,068467947 | -3,063036695 | 0,002191032 | 0,007828244 |
| ENSG00000185716.12 | MOSMO | protein_coding | 2525,886201 | 0,424554017 | 0,073524662 | 5,774307644 | 7,73E-09 | 6,51E-08 |
| ENSG00000185722.18 | ANKFY1 | protein_coding | 16192,73557 | 0,846489383 | 0,047024922 | 18,00086748 | 1,92E-72 | 1,62E-70 |
| ENSG00000185737.13 | NRG3 | protein_coding | 764,6019893 | -0,330833363 | 0,094632856 | -3,495967224 | 0,000472347 | 0,001970358 |
| ENSG00000185745.10 | IFIT1 | protein_coding | 100740,4704 | 9,495340463 | 0,128805371 | 73,71851347 | 0 | 0 |
| ENSG00000185753.13 | CXorf38 | protein_coding | 5755,02204 | 0,778539471 | 0,049960375 | 15,58313906 | 9,48E-55 | 6,01E-53 |
| ENSG00000185787.14 | MORF4L1 | protein_coding | 60375,29902 | -0,163655533 | 0,043805214 | -3,735982999 | 0,000186983 | 0,000843279 |
| ENSG00000185818.7 | NAT8L | protein_coding | 655,0199227 | -0,834875598 | 0,104932649 | -7,95629966 | 1,77E-15 | 2,75E-14 |
| ENSG00000185825.16 | BCAP31 | protein_coding | 14515,89461 | -0,253051423 | 0,046235159 | -5,473138404 | 4,42E-08 | 3,41E-07 |
| ENSG00000185829.17 | ARL17A | protein_coding | 1213,989551 | -0,54845727 | 0,104558101 | -5,245478499 | 1,56E-07 | 1,13E-06 |
| ENSG00000185875.13 | THNSL1 | protein_coding | 1456,085576 | 0,262144434 | 0,07623925 | 3,438444563 | 0,000585066 | 0,002390933 |
| ENSG00000185880.13 | TRIM69 | protein_coding | 15216,36138 | 3,210870047 | 0,066157489 | 48,53373519 | 0 | 0 |
| ENSG00000185885.16 | IFITM1 | protein_coding | 12953,03954 | 15,12583081 | 1,125007569 | 13,44509248 | 3,29E-41 | 1,53E-39 |
| ENSG00000185904.12 | LINC00839 | lncRNA | 105,4843409 | -0,668184535 | 0,245027756 | -2,726974873 | 0,006391791 | 0,020260923 |
| ENSG00000185917.13 | SETD4 | protein_coding | 1660,06473 | 0,240567953 | 0,068613791 | 3,506116644 | 0,000454696 | 0,001906268 |
| ENSG00000185947.15 | ZNF267 | protein_coding | 4331,175559 | 0,850927382 | 0,058413031 | 14,56742391 | 4,53E-48 | 2,46E-46 |
| ENSG00000185986.12 | SDHAP3 | transcribed_unproc | 869,6068493 | 0,313687043 | 0,097999458 | 3,200905911 | 0,001369963 | 0,005151435 |
| ENSG00000186001.13 | LRCH3 | protein_coding | 6727,326205 | 0,336757388 | 0,052604011 | 6,401743574 | 1,54E-10 | 1,57E-09 |
| ENSG00000186010.19 | NDUFA13 | protein_coding | 11110,23712 | -0,163068454 | 0,052286971 | -3,118720596 | 0,001816381 | 0,006638241 |
| ENSG00000186017.14 | ZNF566 | protein_coding | 2556,168399 | 0,445204956 | 0,064559231 | 6,896069688 | 5,35E-12 | 6,30E-11 |
| ENSG00000186019.11 | AC021092.1 | lncRNA | 722,1584012 | -0,379427334 | 0,103736375 | -3,657611261 | 0,000254577 | 0,001120725 |
| ENSG00000186088.16 | GSAP | protein_coding | 1147,988415 | 0,541864202 | 0,103477663 | 5,236533046 | 1,64E-07 | 1,18E-06 |
| ENSG00000186105.8 | LRRC70 | protein_coding | 12,79691536 | 2,345294449 | 0,874156902 | 2,682921617 | 0,007298209 | 0,022759129 |
| ENSG00000186130.5 | ZBTB6 | protein_coding | 3490,636265 | 0,198020992 | 0,060049234 | 3,297643927 | 0,000974997 | 0,003789709 |
| ENSG00000186132.15 | C2orf76 | protein_coding | 753,4617776 | -0,769922102 | 0,090952586 | -8,465093041 | 2,56E-17 | 4,45E-16 |

| | | | | | | | | |
|---|---|---|---|---|---|---|---|---|
| ENSG00000186141.9 | POLR3C | protein_coding | 2552,773509 | 0,345860535 | 0,06279759 | 5,507544667 | 3,64E-08 | 2,84E-07 |
| ENSG00000186162.10 | CIDECP1 | transcribed_unproc | 621,0945904 | 0,26020607 | 0,096595091 | 2,693781493 | 0,007064646 | 0,022144287 |
| ENSG00000186174.12 | BCL9L | protein_coding | 2398,232955 | 0,334282594 | 0,076915219 | 4,346117716 | 1,39E-05 | 7,59E-05 |
| ENSG00000186185.13 | KIF18B | protein_coding | 2940,895182 | -0,559158743 | 0,067577178 | -8,274372498 | 1,29E-16 | 2,16E-15 |
| ENSG00000186188.10 | FFAR4 | protein_coding | 822,2079754 | 0,297883136 | 0,094996516 | 3,135726974 | 0,001714287 | 0,006301958 |
| ENSG00000186193.9 | SAPCD2 | protein_coding | 4077,61012 | -0,768200862 | 0,083787232 | -9,168471639 | 4,80E-20 | 9,75E-19 |
| ENSG00000186197.14 | EDARADD | protein_coding | 21100,07645 | 0,256115053 | 0,041477978 | 6,174723697 | 6,63E-10 | 6,33E-09 |
| ENSG00000186204.14 | CYP4F12 | protein_coding | 660,7597575 | 0,380712497 | 0,103844929 | 3,666163577 | 0,000246216 | 0,001086258 |
| ENSG00000186205.13 | mars-01 | protein_coding | 3413,874813 | 0,144036468 | 0,060616173 | 2,376205225 | 0,017491731 | 0,048434171 |
| ENSG00000186212.3 | SOWAHB | protein_coding | 351,5684619 | 0,445556829 | 0,131552668 | 3,386908358 | 0,00070685 | 0,002836599 |
| ENSG00000186283.14 | TOR3A | protein_coding | 5391,189615 | 0,378999063 | 0,054242763 | 6,98708998 | 2,81E-12 | 3,40E-11 |
| ENSG00000186298.11 | PPP1CC | protein_coding | 25225,06894 | -0,24639535 | 0,044409955 | -5,548200876 | 2,89E-08 | 2,29E-07 |
| ENSG00000186300.12 | ZNF555 | protein_coding | 1509,660398 | 0,191399556 | 0,075632554 | 2,530650436 | 0,011385125 | 0,03340735 |
| ENSG00000186310.9 | NAP1L3 | protein_coding | 263,5648129 | -0,60576728 | 0,166661303 | -3,63472065 | 0,000278282 | 0,001215102 |
| ENSG00000186318.16 | BACE1 | protein_coding | 19597,74809 | 0,207584195 | 0,039879016 | 5,205348915 | 1,94E-07 | 1,38E-06 |
| ENSG00000186369.10 | LINC00643 | transcribed_unitary | 2955,544661 | -0,319819233 | 0,061698805 | -5,183556319 | 2,18E-07 | 1,54E-06 |
| ENSG00000186376.15 | ZNF75D | protein_coding | 1902,755175 | 0,741889702 | 0,065964009 | 11,2468862 | 2,40E-29 | 7,34E-28 |
| ENSG00000186395.8 | KRT10 | protein_coding | 1766,941979 | -0,853081087 | 0,077176443 | -11,05364608 | 2,10E-28 | 6,24E-27 |
| ENSG00000186432.9 | KPNA4 | protein_coding | 11706,74633 | 0,209146715 | 0,058686243 | 3,563811642 | 0,000365508 | 0,001560847 |
| ENSG00000186439.14 | TRDN | protein_coding | 310,3437327 | -0,682443902 | 0,138739788 | -4,918876648 | 8,70E-07 | 5,70E-06 |
| ENSG00000186468.13 | RPS23 | protein_coding | 63449,46649 | -0,691676048 | 0,039995141 | -17,29400215 | 5,22E-67 | 4,02E-65 |
| ENSG00000186470.14 | BTN3A2 | protein_coding | 5551,544914 | 4,036760734 | 0,077775882 | 51,90247439 | 0 | 0 |
| ENSG00000186479.4 | RGS7BP | protein_coding | 142,2940315 | 0,918042776 | 0,229656022 | 3,997468763 | 6,40E-05 | 0,000315409 |
| ENSG00000186480.13 | INSIG1 | protein_coding | 27355,60686 | 0,393812776 | 0,042359594 | 9,296896841 | 1,45E-20 | 3,04E-19 |
| ENSG00000186487.19 | MYT1L | protein_coding | 3645,463873 | -0,223556162 | 0,065691859 | -3,403103018 | 0,000666252 | 0,002689018 |
| ENSG00000186493.12 | C5orf38 | protein_coding | 481,4848613 | -0,469528516 | 0,168489553 | -2,786692159 | 0,005324904 | 0,017245792 |
| ENSG00000186522.15 | sept-10 | protein_coding | 4918,695904 | -0,281644879 | 0,050432792 | -5,584558544 | 2,34E-08 | 1,87E-07 |
| ENSG00000186567.12 | CEACAM19 | protein_coding | 230,2997273 | 0,571682047 | 0,172717141 | 3,309932329 | 0,000933185 | 0,003642287 |
| ENSG00000186577.14 | SMIM29 | protein_coding | 1250,70742 | 0,503772312 | 0,075120996 | 6,706145271 | 2,00E-11 | 2,23E-10 |
| ENSG00000186594.14 | MIR22HG | lncRNA | 598,3087921 | 0,52945699 | 0,110157616 | 4,806358487 | 1,54E-06 | 9,71E-06 |
| ENSG00000186603.5 | HPDL | protein_coding | 95,84419458 | -1,005451855 | 0,294801704 | -3,410603951 | 0,000648192 | 0,002623184 |
| ENSG00000186625.14 | KATNA1 | protein_coding | 1799,760878 | 0,268079947 | 0,068733347 | 3,900289448 | 9,61E-05 | 0,000458375 |
| ENSG00000186635.14 | ARAP1 | protein_coding | 9228,417997 | 0,196233026 | 0,071881409 | 2,729955191 | 0,006334293 | 0,020101313 |
| ENSG00000186642.16 | PDE2A | protein_coding | 53,8118072 | -1,55686606 | 0,353988069 | -4,398074946 | 1,09E-05 | 6,08E-05 |
| ENSG00000186687.16 | LYRM7 | protein_coding | 5905,842835 | -0,945812923 | 0,146357577 | -6,462343417 | 1,03E-10 | 1,07E-09 |
| ENSG00000186704.9 | DTX2P1 | transcribed_unproc | 12,10168556 | 2,135294504 | 0,757053561 | 2,820532935 | 0,004794395 | 0,015715311 |

| | | | | | | | | |
|---|---|---|---|---|---|---|---|---|
| ENSG00000186714.12 | CCDC73 | protein_coding | 47,1936062 | 2,029378118 | 0,474238264 | 4,279237402 | 1,88E-05 | 0,000100553 |
| ENSG00000186732.14 | MPPED1 | protein_coding | 97,6346096 | -1,102987439 | 0,239292291 | -4,609373053 | 4,04E-06 | 2,40E-05 |
| ENSG00000186767.6 | SPIN4 | protein_coding | 3926,712058 | -0,296941165 | 0,054414802 | -5,45699252 | 4,84E-08 | 3,72E-07 |
| ENSG00000186810.8 | CXCR3 | protein_coding | 15,38594113 | 1,481815025 | 0,620704132 | 2,387312971 | 0,016972036 | 0,047198364 |
| ENSG00000186812.13 | ZNF397 | protein_coding | 6054,695831 | -0,141372494 | 0,053308454 | -2,651971371 | 0,008002333 | 0,024681352 |
| ENSG00000186814.14 | ZSCAN30 | protein_coding | 5252,9483 | 0,199767889 | 0,056062828 | 3,563285951 | 0,000366241 | 0,001563681 |
| ENSG00000186815.12 | TPCN1 | protein_coding | 5246,554165 | -0,258466333 | 0,068330735 | -3,782577971 | 0,000155212 | 0,000712123 |
| ENSG00000186834.3 | HEXIM1 | protein_coding | 6319,418186 | 0,344621227 | 0,050621765 | 6,807767907 | 9,91E-12 | 1,14E-10 |
| ENSG00000186866.16 | POFUT2 | protein_coding | 3350,956166 | 0,337372927 | 0,078417219 | 4,302281211 | 1,69E-05 | 9,13E-05 |
| ENSG00000186871.7 | ERCC6L | protein_coding | 2433,002238 | -0,328636512 | 0,068844426 | -4,773611056 | 1,81E-06 | 1,13E-05 |
| ENSG00000186889.10 | TMEM17 | protein_coding | 1139,13527 | -0,300818119 | 0,102427346 | -2,936892643 | 0,003315188 | 0,011340665 |
| ENSG00000186918.14 | ZNF395 | protein_coding | 12141,85216 | -0,204738224 | 0,042779347 | -4,785912832 | 1,70E-06 | 1,07E-05 |
| ENSG00000186919.12 | ZACN | protein_coding | 28,95578067 | 1,056241035 | 0,440017627 | 2,400451643 | 0,016374854 | 0,045745337 |
| ENSG00000186951.16 | PPARA | protein_coding | 6431,571635 | -0,319133416 | 0,062806803 | -5,081191861 | 3,75E-07 | 2,58E-06 |
| ENSG00000187017.17 | ESPN | protein_coding | 2262,637623 | -0,338710018 | 0,073017631 | -4,63874288 | 3,51E-06 | 2,10E-05 |
| ENSG00000187024.14 | PTRH1 | protein_coding | 273,6932092 | -0,452870882 | 0,175124392 | -2,585995456 | 0,009709821 | 0,029180022 |
| ENSG00000187045.18 | TMPRSS6 | protein_coding | 12632,20352 | -0,357638922 | 0,055997397 | -6,386706162 | 1,69E-10 | 1,73E-09 |
| ENSG00000187051.9 | RPS19BP1 | protein_coding | 3225,656013 | -0,216821908 | 0,062059445 | -3,493777719 | 0,000476237 | 0,001982913 |
| ENSG00000187068.3 | C3orf70 | protein_coding | 1174,798965 | -0,531748452 | 0,083456466 | -6,371566871 | 1,87E-10 | 1,90E-09 |
| ENSG00000187079.17 | TEAD1 | protein_coding | 18155,14174 | -0,189794213 | 0,046472871 | -4,083978621 | 4,43E-05 | 0,000223975 |
| ENSG00000187091.13 | PLCD1 | protein_coding | 217,4074177 | 0,909553159 | 0,164617125 | 5,525264525 | 3,29E-08 | 2,58E-07 |
| ENSG00000187097.12 | ENTPD5 | protein_coding | 5932,285165 | 0,149246065 | 0,062142163 | 2,401687627 | 0,016319638 | 0,045603814 |
| ENSG00000187098.15 | MITF | protein_coding | 147,1701778 | 0,59916832 | 0,20817735 | 2,87816287 | 0,003999986 | 0,013398166 |
| ENSG00000187109.14 | NAP1L1 | protein_coding | 53439,53003 | -0,749814895 | 0,044715742 | -16,76847705 | 4,15E-63 | 3,01E-61 |
| ENSG00000187134.14 | AKR1C1 | protein_coding | 693,7533302 | -0,475874525 | 0,108033183 | -4,404892211 | 1,06E-05 | 5,91E-05 |
| ENSG00000187186.14 | AL162231.1 | protein_coding | 371,5192122 | -0,777920099 | 0,136400824 | -5,703192084 | 1,18E-08 | 9,69E-08 |
| ENSG00000187193.9 | MT1X | protein_coding | 3973,663799 | 1,724870319 | 0,081712866 | 21,10891966 | 6,59E-99 | 7,60E-97 |
| ENSG00000187210.14 | GCNT1 | protein_coding | 23042,21089 | 0,181148386 | 0,047627953 | 3,803404833 | 0,000142721 | 0,000660194 |
| ENSG00000187231.14 | SESTD1 | protein_coding | 22332,4704 | -0,422712215 | 0,050314708 | -8,401364751 | 4,41E-17 | 7,57E-16 |
| ENSG00000187239.17 | FNBP1 | protein_coding | 3582,82847 | 0,204112483 | 0,056094452 | 3,63872853 | 0,000273987 | 0,001197977 |
| ENSG00000187244.11 | BCAM | protein_coding | 10867,3086 | -0,180007162 | 0,048313635 | -3,725804533 | 0,000194693 | 0,000875594 |
| ENSG00000187257.16 | RSBN1L | protein_coding | 5937,449953 | 0,292855853 | 0,056092344 | 5,220959437 | 1,78E-07 | 1,28E-06 |
| ENSG00000187266.14 | EPOR | protein_coding | 3040,230812 | -0,488422194 | 0,0653969 | -7,468583272 | 8,11E-14 | 1,12E-12 |
| ENSG00000187325.5 | TAF9B | protein_coding | 4893,728956 | -0,417435105 | 0,05351632 | -7,800145975 | 6,18E-15 | 9,30E-14 |
| ENSG00000187391.21 | MAGI2 | protein_coding | 4444,778404 | 0,397374337 | 0,087787327 | 4,526556972 | 6,00E-06 | 3,47E-05 |
| ENSG00000187446.12 | CHP1 | protein_coding | 6888,353092 | -0,249008269 | 0,056048494 | -4,442729013 | 8,88E-06 | 5,02E-05 |

| | | | | | | | | |
|---|---|---|---|---|---|---|---|---|
| ENSG00000187498.16 | COL4A1 | protein_coding | 1350,711693 | -0,470559872 | 0,087194368 | -5,396677388 | 6,79E-08 | 5,11E-07 |
| ENSG00000187510.9 | C12orf74 | protein_coding | 448,7187428 | 3,134721956 | 0,175396675 | 17,87218575 | 1,94E-71 | 1,58E-69 |
| ENSG00000187514.16 | PTMA | protein_coding | 139195,2471 | -0,125335473 | 0,03973058 | -3,154634865 | 0,001606991 | 0,005948324 |
| ENSG00000187522.16 | HSPA14 | protein_coding | 2355,873891 | -0,259471612 | 0,063750597 | -4,070104788 | 4,70E-05 | 0,000236367 |
| ENSG00000187555.15 | USP7 | protein_coding | 26005,55043 | -0,116783282 | 0,043108494 | -2,70905502 | 0,006747515 | 0,021262749 |
| ENSG00000187595.16 | ZNF385C | protein_coding | 57,66760698 | 0,946268905 | 0,315995579 | 2,994563744 | 0,002748376 | 0,00958228 |
| ENSG00000187601.4 | MAGEH1 | protein_coding | 2314,129188 | -0,271805859 | 0,068098738 | -3,991349427 | 6,57E-05 | 0,000322883 |
| ENSG00000187605.15 | TET3 | protein_coding | 4886,812036 | -0,16078378 | 0,052796619 | -3,045342319 | 0,002324156 | 0,008254091 |
| ENSG00000187608.10 | ISG15 | protein_coding | 24102,87496 | 10,46054967 | 0,245514741 | 42,6066054 | 0 | 0 |
| ENSG00000187609.16 | EXD3 | protein_coding | 500,4832634 | -0,346472256 | 0,11707519 | -2,959399468 | 0,003082393 | 0,010615296 |
| ENSG00000187621.14 | TCL6 | lncRNA | 94,86187535 | 2,236176004 | 0,312090209 | 7,165159122 | 7,77E-13 | 9,91E-12 |
| ENSG00000187672.13 | ERC2 | protein_coding | 547,2797727 | 0,579300877 | 0,104615336 | 5,537437427 | 3,07E-08 | 2,42E-07 |
| ENSG00000187678.9 | SPRY4 | protein_coding | 1226,419333 | 0,437409818 | 0,076606153 | 5,709852258 | 1,13E-08 | 9,34E-08 |
| ENSG00000187714.7 | SLC18A3 | protein_coding | 935,3603075 | -0,71901839 | 0,110000859 | -6,536479757 | 6,30E-11 | 6,69E-10 |
| ENSG00000187741.15 | FANCA | protein_coding | 8495,048344 | -0,124940357 | 0,048180382 | -2,593179028 | 0,009509326 | 0,028688568 |
| ENSG00000187753.13 | C9orf153 | protein_coding | 28,16366393 | 1,3828698 | 0,472095212 | 2,929218017 | 0,00339816 | 0,011596298 |
| ENSG00000187764.11 | SEMA4D | protein_coding | 4275,726535 | 0,390546176 | 0,059109331 | 6,607183145 | 3,92E-11 | 4,24E-10 |
| ENSG00000187772.8 | LIN28B | protein_coding | 4420,886442 | -0,484673499 | 0,050655962 | -9,56794589 | 1,09E-21 | 2,43E-20 |
| ENSG00000187783.12 | TMEM72 | protein_coding | 183,073115 | -0,479423086 | 0,18637563 | -2,572348572 | 0,010101113 | 0,030194683 |
| ENSG00000187796.14 | CARD9 | protein_coding | 279,3188798 | -0,453779974 | 0,182457674 | -2,487042423 | 0,012881004 | 0,037195448 |
| ENSG00000187800.13 | PEAR1 | protein_coding | 274,5651867 | -2,434631903 | 0,17646821 | -13,79643335 | 2,68E-43 | 1,33E-41 |
| ENSG00000187801.15 | ZFP69B | protein_coding | 1006,596847 | -0,254055356 | 0,090526244 | -2,806427669 | 0,005009416 | 0,016344363 |
| ENSG00000187815.10 | ZFP69 | protein_coding | 397,0293989 | 0,401712551 | 0,14298816 | 2,809411289 | 0,00496322 | 0,016214384 |
| ENSG00000187837.3 | HIST1H1C | protein_coding | 1200,579249 | 0,929986899 | 0,090492527 | 10,27694698 | 8,95E-25 | 2,31E-23 |
| ENSG00000187838.17 | PLSCR3 | protein_coding | 1680,658082 | -0,413736232 | 0,08454437 | -4,893717102 | 9,89E-07 | 6,43E-06 |
| ENSG00000187867.9 | PALM3 | protein_coding | 371,9438308 | -0,672012301 | 0,135594053 | -4,95606028 | 7,19E-07 | 4,77E-06 |
| ENSG00000187908.19 | DMBT1 | protein_coding | 265,8381236 | 2,170935195 | 0,483742894 | 4,487787259 | 7,20E-06 | 4,11E-05 |
| ENSG00000187922.13 | LCN10 | protein_coding | 876,0025903 | -1,849387445 | 0,11444757 | -16,15925484 | 9,77E-59 | 6,71E-57 |
| ENSG00000187951.11 | AC091057.1 | lncRNA | 819,6316142 | -0,313540328 | 0,091563242 | -3,424303494 | 0,000616377 | 0,002504339 |
| ENSG00000187987.9 | ZSCAN23 | protein_coding | 836,3389039 | 0,556892213 | 0,092515344 | 6,019457836 | 1,75E-09 | 1,59E-08 |
| ENSG00000187994.14 | RINL | protein_coding | 1245,770545 | -0,631273711 | 0,089691451 | -7,038281879 | 1,95E-12 | 2,40E-11 |
| ENSG00000188010.14 | MORN2 | protein_coding | 1499,703156 | -0,385310107 | 0,076370565 | -5,045269821 | 4,53E-07 | 3,09E-06 |
| ENSG00000188011.5 | RTP5 | protein_coding | 130,1278815 | -0,722269901 | 0,210118299 | -3,437444069 | 0,000587232 | 0,002398477 |
| ENSG00000188033.10 | ZNF490 | protein_coding | 1197,789539 | 0,319814533 | 0,08364072 | 3,823670264 | 0,00013148 | 0,00061237 |
| ENSG00000188039.14 | NWD1 | protein_coding | 220,254076 | -0,406836387 | 0,163240629 | -2,492249562 | 0,012693681 | 0,036739369 |
| ENSG00000188060.7 | RAB42 | protein_coding | 54,32549175 | -0,894641402 | 0,343905746 | -2,601414525 | 0,00928402 | 0,028110573 |

| | | | | | | | | |
|---|---|---|---|---|---|---|---|---|
| ENSG00000188070.9 | C11orf95 | protein_coding | 10251,89615 | -0,749899028 | 0,046443002 | -16,14665271 | 1,20E-58 | 8,20E-57 |
| ENSG00000188092.14 | GPR89B | protein_coding | 3699,073337 | 0,482955714 | 0,078983016 | 6,114678024 | 9,68E-10 | 9,05E-09 |
| ENSG00000188107.14 | EYS | protein_coding | 556,2256435 | -0,422816721 | 0,139777756 | -3,024921373 | 0,002486978 | 0,008763209 |
| ENSG00000188153.13 | COL4A5 | protein_coding | 74,72395997 | 0,992400596 | 0,395361037 | 2,510112287 | 0,012069278 | 0,035162745 |
| ENSG00000188157.15 | AGRN | protein_coding | 7655,94937 | 1,52808623 | 0,082415856 | 18,541168 | 9,61E-77 | 8,69E-75 |
| ENSG00000188162.10 | OTOG | protein_coding | 24,95995783 | 1,998456479 | 0,542937697 | 3,680821006 | 0,000232484 | 0,001031334 |
| ENSG00000188175.9 | HEPACAM2 | protein_coding | 20377,51487 | -0,250607779 | 0,04310313 | -5,814143461 | 6,09E-09 | 5,21E-08 |
| ENSG00000188185.12 | LINC00265 | lncRNA | 855,5801947 | 0,628762066 | 0,097832659 | 6,426913787 | 1,30E-10 | 1,34E-09 |
| ENSG00000188199.10 | NUTM2B | protein_coding | 139,7537612 | 0,669752268 | 0,207752883 | 3,2237929 | 0,001265049 | 0,004788125 |
| ENSG00000188229.6 | TUBB4B | protein_coding | 28180,08902 | -0,411888927 | 0,040990846 | -10,04831477 | 9,35E-24 | 2,29E-22 |
| ENSG00000188243.12 | COMMD6 | protein_coding | 5523,349443 | -0,331989762 | 0,055744276 | -5,955584741 | 2,59E-09 | 2,31E-08 |
| ENSG00000188280.11 | FAM230G | lncRNA | 45,37226375 | 5,829570326 | 1,04145933 | 5,59750166 | 2,17E-08 | 1,75E-07 |
| ENSG00000188282.12 | RUFY4 | protein_coding | 705,6922799 | 9,373352093 | 0,810687058 | 11,56223231 | 6,40E-31 | 2,13E-29 |
| ENSG00000188290.10 | HES4 | protein_coding | 33,57662898 | 2,836353446 | 0,458726375 | 6,18310522 | 6,29E-10 | 6,02E-09 |
| ENSG00000188295.14 | ZNF669 | protein_coding | 1916,182429 | 0,282021397 | 0,071773842 | 3,929306104 | 8,52E-05 | 0,000411496 |
| ENSG00000188306.6 | LRRIQ4 | protein_coding | 103,0747645 | 0,762573937 | 0,237301244 | 3,213526927 | 0,001311155 | 0,004950149 |
| ENSG00000188312.14 | CENPP | protein_coding | 1734,738701 | -0,306148438 | 0,069014279 | -4,436015911 | 9,16E-06 | 5,17E-05 |
| ENSG00000188313.13 | PLSCR1 | protein_coding | 37673,95082 | 3,616006588 | 0,062510617 | 57,84627914 | 0 | 0 |
| ENSG00000188315.8 | C3orf62 | protein_coding | 1187,528837 | 0,930169999 | 0,122110736 | 7,617430102 | 2,59E-14 | 3,69E-13 |
| ENSG00000188322.6 | SBK1 | protein_coding | 4183,760097 | -0,142702056 | 0,057996785 | -2,460516679 | 0,013873712 | 0,039720604 |
| ENSG00000188338.15 | SLC38A3 | protein_coding | 5127,300177 | -0,193684109 | 0,07276686 | -2,661707676 | 0,007774536 | 0,024067809 |
| ENSG00000188343.12 | FAM92A | protein_coding | 2727,550044 | -0,211417214 | 0,061364681 | -3,445258938 | 0,000570513 | 0,002338676 |
| ENSG00000188352.12 | FOCAD | protein_coding | 14778,27533 | -0,298615053 | 0,05680539 | -5,256808478 | 1,47E-07 | 1,06E-06 |
| ENSG00000188404.9 | SELL | protein_coding | 168,2484342 | 3,708565717 | 0,314964147 | 11,77456468 | 5,28E-32 | 1,85E-30 |
| ENSG00000188419.14 | CHM | protein_coding | 12817,06289 | -0,153429823 | 0,054364402 | -2,822247953 | 0,004768829 | 0,015645179 |
| ENSG00000188487.11 | INSC | protein_coding | 24,4343049 | 2,922799185 | 0,541233358 | 5,400256917 | 6,65E-08 | 5,02E-07 |
| ENSG00000188493.15 | C19orf54 | protein_coding | 1773,174611 | 0,209234438 | 0,082302999 | 2,542245617 | 0,011014274 | 0,03249709 |
| ENSG00000188511.12 | C22orf34 | lncRNA | 13,91633658 | -2,32280165 | 0,752828802 | -3,085431436 | 0,002032571 | 0,007327216 |
| ENSG00000188522.15 | FAM83G | protein_coding | 2062,269081 | 0,278781645 | 0,068564178 | 4,06599555 | 4,78E-05 | 0,000240196 |
| ENSG00000188529.14 | SRSF10 | protein_coding | 22640,43131 | -0,11595543 | 0,040613821 | -2,855073166 | 0,004302692 | 0,014297208 |
| ENSG00000188542.9 | DUSP28 | protein_coding | 607,8515083 | -0,462314285 | 0,124339657 | -3,718156354 | 0,000200682 | 0,000899831 |
| ENSG00000188573.7 | FBLL1 | protein_coding | 561,6341205 | 0,283800181 | 0,111074174 | 2,55505102 | 0,010617223 | 0,031494871 |
| ENSG00000188580.14 | NKAIN2 | protein_coding | 1466,179684 | -0,953619645 | 0,117675745 | -8,103791038 | 5,33E-16 | 8,56E-15 |
| ENSG00000188599.17 | NPIPP1 | transcribed_unproc | 1510,901449 | 1,17681678 | 0,090567742 | 12,99377415 | 1,33E-38 | 5,70E-37 |
| ENSG00000188603.19 | CLN3 | protein_coding | 3919,032679 | 0,878277333 | 0,060422212 | 14,53566999 | 7,20E-48 | 3,86E-46 |
| ENSG00000188610.12 | FAM72B | protein_coding | 1551,832993 | -1,178027394 | 0,078948919 | -14,92138734 | 2,39E-50 | 1,38E-48 |

| | | | | | | | | |
|---|---|---|---|---|---|---|---|---|
| ENSG00000188611.15 | ASAH2 | protein_coding | 98,99150287 | 0,761478436 | 0,314186713 | 2,423649394 | 0,015365432 | 0,043302931 |
| ENSG00000188613.7 | NANOS1 | protein_coding | 1132,984979 | -0,749257169 | 0,080537776 | -9,303176855 | 1,36E-20 | 2,88E-19 |
| ENSG00000188629.12 | ZNF177 | protein_coding | 416,4406311 | 0,543080867 | 0,124930902 | 4,347049918 | 1,38E-05 | 7,56E-05 |
| ENSG00000188641.13 | DPYD | protein_coding | 376,0299387 | 0,454268812 | 0,123068189 | 3,691196046 | 0,000223202 | 0,000993092 |
| ENSG00000188659.9 | SAXO2 | protein_coding | 428,6305733 | -0,534501381 | 0,158828861 | -3,365266096 | 0,000764699 | 0,003043247 |
| ENSG00000188677.14 | PARVB | protein_coding | 2742,798577 | -0,300705794 | 0,066199247 | -4,542435281 | 5,56E-06 | 3,24E-05 |
| ENSG00000188739.15 | RBM34 | protein_coding | 4324,483212 | 0,240701149 | 0,069450371 | 3,465800777 | 0,000528655 | 0,002177799 |
| ENSG00000188761.13 | BCL2L15 | protein_coding | 57,36758804 | 1,040329322 | 0,333371543 | 3,120630252 | 0,001804645 | 0,006599634 |
| ENSG00000188786.10 | MTF1 | protein_coding | 3851,951885 | 0,731206901 | 0,05093898 | 14,35456489 | 9,97E-47 | 5,28E-45 |
| ENSG00000188803.14 | SHISA6 | protein_coding | 1114,087297 | -0,291206251 | 0,089518231 | -3,253038488 | 0,00114178 | 0,004365608 |
| ENSG00000188846.13 | RPL14 | protein_coding | 43387,55703 | -0,53796739 | 0,04561887 | -11,79265053 | 4,26E-32 | 1,51E-30 |
| ENSG00000188878.19 | FBF1 | protein_coding | 928,7891556 | 0,274608636 | 0,106588272 | 2,576349448 | 0,009984968 | 0,029885048 |
| ENSG00000188916.9 | INSYN2A | protein_coding | 41,91208667 | -1,02215657 | 0,403935218 | -2,530496291 | 0,011390129 | 0,033409232 |
| ENSG00000188917.15 | TRMT2B | protein_coding | 2112,018392 | -0,221331267 | 0,079121079 | -2,797374212 | 0,005151982 | 0,016769085 |
| ENSG00000188993.3 | LRRC66 | protein_coding | 539,9478961 | 1,081609959 | 0,117499628 | 9,205220251 | 3,41E-20 | 6,98E-19 |
| ENSG00000188994.13 | ZNF292 | protein_coding | 14147,9384 | 0,242721122 | 0,046825372 | 5,183538603 | 2,18E-07 | 1,54E-06 |
| ENSG00000189056.14 | RELN | protein_coding | 52,44637485 | 1,220869964 | 0,369131617 | 3,307410982 | 0,000941626 | 0,003672049 |
| ENSG00000189057.11 | FAM111B | protein_coding | 13845,26302 | -0,30334546 | 0,046370039 | -6,541841802 | 6,08E-11 | 6,47E-10 |
| ENSG00000189060.5 | H1F0 | protein_coding | 24667,201 | 2,212005049 | 0,05174894 | 42,74493465 | 0 | 0 |
| ENSG00000189067.12 | LITAF | protein_coding | 1103,612018 | 0,306700422 | 0,092784638 | 3,305508648 | 0,000948042 | 0,003696427 |
| ENSG00000189114.7 | BLOC1S3 | protein_coding | 648,7536988 | 0,369575224 | 0,108636581 | 3,401940851 | 0,000669091 | 0,002699027 |
| ENSG00000189152.10 | GRAPL | protein_coding | 13,36288448 | -1,807935081 | 0,713439007 | -2,534113027 | 0,011273236 | 0,033152505 |
| ENSG00000189157.14 | FAM47E | protein_coding | 1660,269227 | -0,348562086 | 0,067886768 | -5,134462781 | 2,83E-07 | 1,98E-06 |
| ENSG00000189159.16 | JPT1 | protein_coding | 8503,577042 | -0,282048265 | 0,052960756 | -5,325608705 | 1,01E-07 | 7,41E-07 |
| ENSG00000189190.10 | ZNF600 | protein_coding | 1007,836191 | 0,77798492 | 0,082737381 | 9,403064349 | 5,30E-21 | 1,14E-19 |
| ENSG00000189195.13 | BTBD8 | protein_coding | 2310,557064 | 0,574967677 | 0,071818865 | 8,005802853 | 1,19E-15 | 1,86E-14 |
| ENSG00000189212.12 | DPY19L2P1 | transcribed_unproc | 1087,039279 | -0,449065921 | 0,086298824 | -5,20361579 | 1,95E-07 | 1,39E-06 |
| ENSG00000189227.6 | C15orf61 | protein_coding | 761,1166124 | -0,460289178 | 0,090354275 | -5,094271188 | 3,50E-07 | 2,42E-06 |
| ENSG00000189233.12 | NUGGC | protein_coding | 151,3602481 | 3,196545552 | 0,278964274 | 11,45861981 | 2,13E-30 | 6,87E-29 |
| ENSG00000189241.7 | TSPYL1 | protein_coding | 16342,22667 | 0,208948158 | 0,04464333 | 4,680389189 | 2,86E-06 | 1,74E-05 |
| ENSG00000189266.13 | PNRC2 | protein_coding | 24012,38333 | 0,277329054 | 0,047118703 | 5,885753178 | 3,96E-09 | 3,46E-08 |
| ENSG00000189275.4 | LINC01164 | lncRNA | 30,5208707 | -1,246636242 | 0,41581358 | -2,998065241 | 0,002716995 | 0,009486093 |
| ENSG00000189306.11 | RRP7A | protein_coding | 4685,877861 | 0,177456584 | 0,059797486 | 2,96762616 | 0,003001091 | 0,010362257 |
| ENSG00000189319.14 | FAM53B | protein_coding | 1573,643096 | 0,373460156 | 0,067384342 | 5,542239441 | 2,99E-08 | 2,36E-07 |
| ENSG00000189332.5 | AC090099.1 | transcribed_unproc | 756,8602202 | -0,353868409 | 0,097923155 | -3,61373579 | 0,000301817 | 0,001309987 |
| ENSG00000189339.11 | SLC35E2B | protein_coding | 13155,35088 | -0,35716064 | 0,043621672 | -8,1876879 | 2,66E-16 | 4,35E-15 |

| | | | | | | | | |
|---|---|---|---|---|---|---|---|---|
| ENSG00000189350.12 | TOGARAM2 | protein_coding | 181,2658437 | -0,839785386 | 0,218361245 | -3,84585362 | 0,000120133 | 0,000564409 |
| ENSG00000189362.12 | NEMP2 | protein_coding | 1318,246038 | -0,446290145 | 0,105881831 | -4,214983244 | 2,50E-05 | 0,000131525 |
| ENSG00000189376.12 | C8orf76 | protein_coding | 2801,083563 | 0,161278866 | 0,065929651 | 2,446226595 | 0,014436024 | 0,04106413 |
| ENSG00000189377.8 | CXCL17 | protein_coding | 37,69743279 | 2,430317201 | 0,425222169 | 5,715405689 | 1,09E-08 | 9,06E-08 |
| ENSG00000189403.15 | HMGB1 | protein_coding | 55248,04351 | -0,400948298 | 0,043548867 | -9,206859392 | 3,36E-20 | 6,88E-19 |
| ENSG00000189409.13 | MMP23B | protein_coding | 128,3444461 | -1,330948772 | 0,314876747 | -4,226888094 | 2,37E-05 | 0,000125286 |
| ENSG00000189431.7 | RASSF10 | protein_coding | 1156,917822 | 0,200193702 | 0,084104702 | 2,380291449 | 0,01729895 | 0,04797118 |
| ENSG00000196072.12 | BLOC1S2 | protein_coding | 6905,082819 | 0,153306531 | 0,04981952 | 3,077238195 | 0,002089283 | 0,007506411 |
| ENSG00000196083.10 | IL1RAP | protein_coding | 4691,132246 | 0,592938038 | 0,050768002 | 11,67936514 | 1,63E-31 | 5,53E-30 |
| ENSG00000196090.12 | PTPRT | protein_coding | 18285,91673 | -0,423067821 | 0,068381412 | -6,186883394 | 6,14E-10 | 5,89E-09 |
| ENSG00000196092.14 | PAX5 | protein_coding | 3126,619589 | 0,634224415 | 0,080165274 | 7,911460713 | 2,54E-15 | 3,91E-14 |
| ENSG00000196096.3 | AC079610.1 | lncRNA | 84,47714257 | 0,884655087 | 0,281305622 | 3,144818372 | 0,0016619 | 0,006133415 |
| ENSG00000196104.11 | SPOCK3 | protein_coding | 1852,952126 | -0,363676239 | 0,068561501 | -5,30437977 | 1,13E-07 | 8,30E-07 |
| ENSG00000196110.7 | ZNF699 | protein_coding | 1182,036447 | 0,315282715 | 0,075814006 | 4,158634167 | 3,20E-05 | 0,000165474 |
| ENSG00000196116.8 | TDRD7 | protein_coding | 13748,47036 | 2,413518124 | 0,047779483 | 50,51369244 | 0 | 0 |
| ENSG00000196123.13 | KIAA0895L | protein_coding | 6852,460856 | -0,319591699 | 0,052799722 | -6,05290495 | 1,42E-09 | 1,31E-08 |
| ENSG00000196132.14 | MYT1 | protein_coding | 7579,47044 | -0,253267669 | 0,060231712 | -4,204889095 | 2,61E-05 | 0,000136798 |
| ENSG00000196141.14 | SPATS2L | protein_coding | 11553,25033 | 1,919064739 | 0,05663163 | 33,88680045 | 1,04E-251 | 2,55E-249 |
| ENSG00000196154.12 | S100A4 | protein_coding | 671,0921261 | -0,337762767 | 0,104008676 | -3,247447999 | 0,001164449 | 0,004442501 |
| ENSG00000196169.15 | KIF19 | protein_coding | 688,3281641 | 0,432066482 | 0,107410768 | 4,02256208 | 5,76E-05 | 0,000285349 |
| ENSG00000196172.9 | ZNF681 | protein_coding | 1680,636879 | 0,264792982 | 0,071564459 | 3,700062676 | 0,000215546 | 0,000960739 |
| ENSG00000196177.12 | ACADSB | protein_coding | 3874,919429 | 0,250084058 | 0,06869175 | 3,640670939 | 0,000271928 | 0,001190361 |
| ENSG00000196204.11 | RNF216P1 | transcribed_unproc | 1002,123878 | 0,266686698 | 0,102917767 | 2,591260061 | 0,00956252 | 0,028829729 |
| ENSG00000196205.8 | EEF1A1P5 | processed_pseudog | 170,6064337 | -1,347868253 | 0,206896851 | -6,51468714 | 7,28E-11 | 7,68E-10 |
| ENSG00000196208.14 | GREB1 | protein_coding | 1234,702118 | 0,388524314 | 0,087612972 | 4,434552383 | 9,23E-06 | 5,20E-05 |
| ENSG00000196214.11 | ZNF766 | protein_coding | 4082,081933 | 0,462553647 | 0,065039847 | 7,111850201 | 1,14E-12 | 1,44E-11 |
| ENSG00000196230.13 | TUBB | protein_coding | 121397,6744 | -0,391958418 | 0,04249302 | -9,224065899 | 2,86E-20 | 5,89E-19 |
| ENSG00000196247.11 | ZNF107 | protein_coding | 8357,15854 | 1,084313073 | 0,045263414 | 23,9556183 | 8,07E-127 | 1,15E-124 |
| ENSG00000196262.14 | PPIA | protein_coding | 106880,6821 | -0,297962408 | 0,038801704 | -7,679106258 | 1,60E-14 | 2,32E-13 |
| ENSG00000196268.11 | ZNF493 | protein_coding | 454,1539529 | 0,375871926 | 0,131702101 | 2,853955428 | 0,004317859 | 0,014332795 |
| ENSG00000196323.14 | ZBTB44 | protein_coding | 8480,164296 | -0,434730221 | 0,050135795 | -8,671054624 | 4,28E-18 | 7,85E-17 |
| ENSG00000196335.13 | STK31 | protein_coding | 20,20782311 | -1,451067537 | 0,526897289 | -2,753985584 | 0,005887436 | 0,018839963 |
| ENSG00000196337.11 | CGB7 | protein_coding | 94,68202349 | 0,689050742 | 0,243005903 | 2,835530885 | 0,004574959 | 0,01507727 |
| ENSG00000196352.15 | CD55 | protein_coding | 1093,53701 | 0,411793786 | 0,080082341 | 5,142129691 | 2,72E-07 | 1,91E-06 |
| ENSG00000196353.11 | CPNE4 | protein_coding | 2574,670013 | 0,666552325 | 0,072747601 | 9,162533371 | 5,07E-20 | 1,03E-18 |
| ENSG00000196358.11 | NTNG2 | protein_coding | 1233,003736 | 0,494158115 | 0,10934165 | 4,519395075 | 6,20E-06 | 3,58E-05 |

| | | | | | | | | |
|---|---|---|---|---|---|---|---|---|
| ENSG00000196361.10 | ELAVL3 | protein_coding | 5098,742537 | -0,22081049 | 0,068856539 | -3,206819451 | 0,001342112 | 0,005056 |
| ENSG00000196363.9 | WDR5 | protein_coding | 5908,090262 | -0,2388378 | 0,048159485 | -4,959309637 | 7,07E-07 | 4,70E-06 |
| ENSG00000196369.11 | SRGAP2B | protein_coding | 2698,760159 | 0,325283567 | 0,077310316 | 4,207505329 | 2,58E-05 | 0,00013554 |
| ENSG00000196376.11 | SLC35F1 | protein_coding | 176,5717354 | -0,448977888 | 0,182959557 | -2,453973415 | 0,014128743 | 0,040343 |
| ENSG00000196381.11 | ZNF781 | protein_coding | 436,8218023 | -0,569704733 | 0,146042035 | -3,900964073 | 9,58E-05 | 0,000457293 |
| ENSG00000196387.9 | ZNF140 | protein_coding | 3790,028539 | 0,233023752 | 0,069003392 | 3,376989833 | 0,000732838 | 0,002929535 |
| ENSG00000196405.13 | EVL | protein_coding | 4333,947714 | -0,247955761 | 0,067158022 | -3,692124217 | 0,000222389 | 0,000989865 |
| ENSG00000196417.13 | ZNF765 | protein_coding | 3386,310351 | 0,411493102 | 0,053894614 | 7,635143265 | 2,26E-14 | 3,23E-13 |
| ENSG00000196419.12 | XRCC6 | protein_coding | 47475,88217 | -0,176089251 | 0,040487236 | -4,349253447 | 1,37E-05 | 7,50E-05 |
| ENSG00000196428.12 | TSC22D2 | protein_coding | 1884,433615 | 0,209908024 | 0,076112947 | 2,757849124 | 0,005818305 | 0,018647902 |
| ENSG00000196440.11 | ARMCX4 | protein_coding | 3323,633995 | 0,197728822 | 0,072933869 | 2,711069966 | 0,006706647 | 0,021142847 |
| ENSG00000196453.8 | ZNF777 | protein_coding | 786,7358373 | 0,336535988 | 0,094532175 | 3,560015274 | 0,000370833 | 0,001581486 |
| ENSG00000196456.13 | ZNF775 | protein_coding | 1502,017047 | 0,210538805 | 0,068032793 | 3,094666499 | 0,001970343 | 0,007130298 |
| ENSG00000196458.11 | ZNF605 | protein_coding | 3552,049267 | -0,235737514 | 0,065419766 | -3,603460057 | 0,000314009 | 0,001360283 |
| ENSG00000196459.14 | TRAPPC2 | protein_coding | 3640,464559 | -0,170619207 | 0,056350358 | -3,027828247 | 0,00246318 | 0,008687516 |
| ENSG00000196465.10 | MYL6B | protein_coding | 6104,044042 | -0,585098969 | 0,054152744 | -10,80460436 | 3,27E-27 | 9,25E-26 |
| ENSG00000196482.17 | ESRRG | protein_coding | 1396,908473 | 0,538567802 | 0,091258972 | 5,901532633 | 3,60E-09 | 3,16E-08 |
| ENSG00000196497.17 | IPO4 | protein_coding | 2084,62512 | 0,943500889 | 0,202431008 | 4,660851616 | 3,15E-06 | 1,90E-05 |
| ENSG00000196498.13 | NCOR2 | protein_coding | 5584,944473 | -0,26185571 | 0,064896303 | -4,034986533 | 5,46E-05 | 0,000272048 |
| ENSG00000196502.11 | SULT1A1 | protein_coding | 359,3349196 | -0,473471358 | 0,137785072 | -3,436303735 | 0,000589709 | 0,002407285 |
| ENSG00000196503.4 | ARL9 | protein_coding | 136,3772265 | 0,616033381 | 0,22671419 | 2,717224625 | 0,006583191 | 0,02080026 |
| ENSG00000196511.14 | TPK1 | protein_coding | 1381,188309 | -0,479549336 | 0,084148742 | -5,698829516 | 1,21E-08 | 9,93E-08 |
| ENSG00000196526.10 | AFAP1 | protein_coding | 1646,588421 | -0,242088654 | 0,074148027 | -3,264937214 | 0,001094883 | 0,004208458 |
| ENSG00000196531.10 | NACA | protein_coding | 30695,08596 | -0,363127063 | 0,038843755 | -9,348402744 | 8,90E-21 | 1,89E-19 |
| ENSG00000196542.8 | SPTSSB | protein_coding | 1201,345895 | 0,461601698 | 0,078918286 | 5,849109547 | 4,94E-09 | 4,27E-08 |
| ENSG00000196547.15 | MAN2A2 | protein_coding | 8538,31179 | 0,164305911 | 0,063563673 | 2,584902714 | 0,009740648 | 0,029257038 |
| ENSG00000196549.10 | MME | protein_coding | 15121,7848 | -0,282793737 | 0,064049115 | -4,415263747 | 1,01E-05 | 5,66E-05 |
| ENSG00000196550.10 | FAM72A | protein_coding | 1488,409969 | -1,173399481 | 0,088523627 | -13,25521244 | 4,21E-40 | 1,90E-38 |
| ENSG00000196557.13 | CACNA1H | protein_coding | 4845,556843 | -0,181200918 | 0,063964333 | -2,832843092 | 0,004613603 | 0,015182395 |
| ENSG00000196569.12 | LAMA2 | protein_coding | 1393,593396 | -0,199615417 | 0,07733918 | -2,581038691 | 0,009850354 | 0,029558942 |
| ENSG00000196584.3 | XRCC2 | protein_coding | 3096,884888 | -0,26392817 | 0,060959692 | -4,329552203 | 1,49E-05 | 8,13E-05 |
| ENSG00000196591.12 | HDAC2 | protein_coding | 34465,48501 | -0,238661695 | 0,044170793 | -5,403156225 | 6,55E-08 | 4,94E-07 |
| ENSG00000196597.12 | ZNF782 | protein_coding | 1490,298611 | 0,188543029 | 0,073065667 | 2,580459981 | 0,009866879 | 0,029588807 |
| ENSG00000196636.7 | SDHAF3 | protein_coding | 2443,636699 | -0,377951879 | 0,070240295 | -5,380841271 | 7,41E-08 | 5,55E-07 |
| ENSG00000196642.19 | RABL6 | protein_coding | 13513,15194 | -0,267948947 | 0,046823928 | -5,722479013 | 1,05E-08 | 8,71E-08 |
| ENSG00000196646.12 | ZNF136 | protein_coding | 3584,975285 | 0,300060638 | 0,086130918 | 3,483773833 | 0,000494397 | 0,002050937 |

| | | | | | | | | |
|---|---|---|---|---|---|---|---|---|
| ENSG00000196652.11 | ZKSCAN5 | protein_coding | 4588,611173 | 0,374200367 | 0,054057475 | 6,92226873 | 4,44E-12 | 5,29E-11 |
| ENSG00000196653.12 | ZNF502 | protein_coding | 1275,588592 | 0,205175103 | 0,080115109 | 2,561003835 | 0,01043702 | 0,031021642 |
| ENSG00000196655.11 | TRAPPC4 | protein_coding | 5627,198319 | 0,237541879 | 0,063385151 | 3,74759509 | 0,000178538 | 0,000807944 |
| ENSG00000196659.9 | TTC30B | protein_coding | 2618,599479 | 0,277423761 | 0,061530059 | 4,508751774 | 6,52E-06 | 3,75E-05 |
| ENSG00000196663.16 | TECPR2 | protein_coding | 3672,308732 | 0,381813187 | 0,085668672 | 4,456858956 | 8,32E-06 | 4,71E-05 |
| ENSG00000196670.13 | ZFP62 | protein_coding | 7015,451836 | -0,120959054 | 0,046119337 | -2,622740518 | 0,008722568 | 0,026629089 |
| ENSG00000196683.10 | TOMM7 | protein_coding | 9454,688113 | -0,355925884 | 0,047109048 | -7,555361344 | 4,18E-14 | 5,89E-13 |
| ENSG00000196684.12 | HSH2D | protein_coding | 2056,655572 | 9,143509487 | 0,356941732 | 25,61625236 | 1,01E-144 | 1,61E-142 |
| ENSG00000196689.12 | TRPV1 | protein_coding | 855,2557436 | 0,879194037 | 0,118168064 | 7,440200056 | 1,01E-13 | 1,38E-12 |
| ENSG00000196696.12 | AC009022.1 | lncRNA | 3732,677273 | 0,189550829 | 0,061041884 | 3,105258484 | 0,001901128 | 0,006907585 |
| ENSG00000196704.12 | AMZ2 | protein_coding | 6401,678404 | -0,135798829 | 0,049160296 | -2,762368016 | 0,005738376 | 0,018433723 |
| ENSG00000196705.8 | ZNF431 | protein_coding | 3661,492324 | 0,317748014 | 0,073792348 | 4,305975097 | 1,66E-05 | 8,98E-05 |
| ENSG00000196712.17 | NF1 | protein_coding | 17458,73952 | 0,281497352 | 0,05129359 | 5,487963527 | 4,07E-08 | 3,15E-07 |
| ENSG00000196715.7 | VKORC1L1 | protein_coding | 8262,25856 | -0,337870728 | 0,0554252 | -6,095976661 | 1,09E-09 | 1,01E-08 |
| ENSG00000196724.12 | ZNF418 | protein_coding | 997,743174 | 0,570979948 | 0,09852782 | 5,795113969 | 6,83E-09 | 5,80E-08 |
| ENSG00000196739.15 | COL27A1 | protein_coding | 2735,524102 | 0,527562507 | 0,070484307 | 7,484822184 | 7,16E-14 | 9,95E-13 |
| ENSG00000196756.13 | SNHG17 | lncRNA | 1174,250511 | 0,206153893 | 0,083322535 | 2,474167308 | 0,013354714 | 0,038390965 |
| ENSG00000196757.8 | ZNF700 | protein_coding | 3813,277507 | 0,141631686 | 0,052787987 | 2,683028739 | 0,007295872 | 0,022754993 |
| ENSG00000196776.16 | CD47 | protein_coding | 9903,08161 | 0,970425227 | 0,050611842 | 19,17387695 | 6,12E-82 | 5,79E-80 |
| ENSG00000196781.15 | TLE1 | protein_coding | 5649,239928 | -0,133201967 | 0,051038286 | -2,60984407 | 0,00905835 | 0,027504978 |
| ENSG00000196782.12 | MAML3 | protein_coding | 2879,336422 | -0,25346361 | 0,06087782 | -4,163480384 | 3,13E-05 | 0,00016226 |
| ENSG00000196793.13 | ZNF239 | protein_coding | 461,1614889 | 0,434930081 | 0,137658736 | 3,15948043 | 0,001580507 | 0,005861861 |
| ENSG00000196814.15 | MVB12B | protein_coding | 4209,780634 | 0,263506589 | 0,057029715 | 4,620513845 | 3,83E-06 | 2,28E-05 |
| ENSG00000196843.16 | ARID5A | protein_coding | 285,6185417 | 0,589028036 | 0,143305955 | 4,11028302 | 3,95E-05 | 0,000201335 |
| ENSG00000196872.12 | KIAA1211L | protein_coding | 1246,664018 | 0,359384147 | 0,086948404 | 4,133303576 | 3,58E-05 | 0,000183221 |
| ENSG00000196876.16 | SCN8A | protein_coding | 4093,55466 | 0,708321436 | 0,058151045 | 12,18071726 | 3,94E-34 | 1,46E-32 |
| ENSG00000196878.15 | LAMB3 | protein_coding | 1430,14822 | -0,171591451 | 0,070934016 | -2,419029121 | 0,015561994 | 0,043741874 |
| ENSG00000196890.4 | HIST3H2BB | protein_coding | 154,9573128 | 1,288083981 | 0,191092519 | 6,740630083 | 1,58E-11 | 1,77E-10 |
| ENSG00000196914.9 | ARHGEF12 | protein_coding | 27192,66756 | -0,143203551 | 0,045075126 | -3,176997263 | 0,001488084 | 0,005561231 |
| ENSG00000196922.10 | ZNF252P | transcribed_unproc | 6955,202075 | 0,119279431 | 0,050350367 | 2,368988295 | 0,017836818 | 0,049215675 |
| ENSG00000196924.17 | FLNA | protein_coding | 4460,854991 | -0,648584763 | 0,167383725 | -3,874837677 | 0,000106696 | 0,000504965 |
| ENSG00000196932.11 | TMEM26 | protein_coding | 936,5059861 | -0,985543058 | 0,084225308 | -11,70126992 | 1,26E-31 | 4,33E-30 |
| ENSG00000196937.11 | FAM3C | protein_coding | 23600,26354 | 0,152501549 | 0,041346285 | 3,688397808 | 0,000225671 | 0,001003481 |
| ENSG00000196950.14 | SLC39A10 | protein_coding | 5039,453452 | -0,590883961 | 0,054259374 | -10,88998849 | 1,29E-27 | 3,69E-26 |
| ENSG00000196951.11 | SCOC-AS1 | lncRNA | 112,2053231 | -0,584384145 | 0,223004084 | -2,620508716 | 0,008779869 | 0,026767718 |
| ENSG00000196954.14 | CASP4 | protein_coding | 4417,480536 | 2,709131525 | 0,063490647 | 42,66977327 | 0 | 0 |

| | | | | | | | | |
|---|---|---|---|---|---|---|---|---|
| ENSG00000196975.16 | ANXA4 | protein_coding | 5024,377127 | 0,340859148 | 0,054208412 | 6,287938239 | 3,22E-10 | 3,19E-09 |
| ENSG00000196976.7 | LAGE3 | protein_coding | 2562,413296 | -0,342639935 | 0,061316718 | -5,588034525 | 2,30E-08 | 1,84E-07 |
| ENSG00000196981.3 | WDR5B | protein_coding | 2021,84462 | 0,160147049 | 0,064185877 | 2,495051181 | 0,012593896 | 0,036497491 |
| ENSG00000197006.14 | METTL9 | protein_coding | 10871,68819 | -0,24549065 | 0,044223338 | -5,551156058 | 2,84E-08 | 2,25E-07 |
| ENSG00000197016.12 | ZNF470 | protein_coding | 1656,590353 | 0,273214544 | 0,072151651 | 3,786670695 | 0,000152679 | 0,000702362 |
| ENSG00000197019.5 | SERTAD1 | protein_coding | 708,8320071 | 0,67074204 | 0,095946169 | 6,990816263 | 2,73E-12 | 3,32E-11 |
| ENSG00000197020.11 | ZNF100 | protein_coding | 2188,279707 | 0,29208135 | 0,06833855 | 4,274034954 | 1,92E-05 | 0,000102768 |
| ENSG00000197021.9 | CXorf40B | protein_coding | 1976,388852 | 0,198315632 | 0,063886174 | 3,104202685 | 0,001907926 | 0,006927811 |
| ENSG00000197024.8 | ZNF398 | protein_coding | 4565,752977 | 0,241401217 | 0,05199032 | 4,643195418 | 3,43E-06 | 2,06E-05 |
| ENSG00000197043.14 | ANXA6 | protein_coding | 19458,02948 | 0,79940269 | 0,04442518 | 17,99436014 | 2,16E-72 | 1,82E-70 |
| ENSG00000197054.11 | ZNF763 | protein_coding | 203,9734417 | 0,698045202 | 0,16815703 | 4,151150882 | 3,31E-05 | 0,000170549 |
| ENSG00000197056.11 | ZMYM1 | protein_coding | 2152,618706 | -0,221543306 | 0,063353678 | -3,496928877 | 0,000470647 | 0,001964726 |
| ENSG00000197081.14 | IGF2R | protein_coding | 21028,69776 | 0,27897656 | 0,059370415 | 4,698915419 | 2,62E-06 | 1,60E-05 |
| ENSG00000197102.12 | DYNC1H1 | protein_coding | 43864,95014 | 0,554014997 | 0,04970124 | 11,14690487 | 7,41E-29 | 2,23E-27 |
| ENSG00000197106.7 | SLC6A17 | protein_coding | 302,827706 | -0,523851821 | 0,144746839 | -3,619089881 | 0,000295641 | 0,001285661 |
| ENSG00000197111.15 | PCBP2 | protein_coding | 52322,63275 | -0,20290294 | 0,044127458 | -4,598110761 | 4,26E-06 | 2,52E-05 |
| ENSG00000197121.15 | PGAP1 | protein_coding | 4790,353498 | -0,261414209 | 0,072019053 | -3,629792369 | 0,000283649 | 0,001236858 |
| ENSG00000197128.11 | ZNF772 | protein_coding | 3854,897492 | 0,29497202 | 0,060687621 | 4,860497307 | 1,17E-06 | 7,53E-06 |
| ENSG00000197136.4 | PCNX3 | protein_coding | 5280,785019 | 0,323401352 | 0,05500002 | 5,880022405 | 4,10E-09 | 3,58E-08 |
| ENSG00000197147.13 | LRRC8B | protein_coding | 7211,966096 | 0,214618922 | 0,047820534 | 4,488007664 | 7,19E-06 | 4,11E-05 |
| ENSG00000197172.10 | MAGEA6 | protein_coding | 11639,07052 | -0,113139056 | 0,043183954 | -2,619932761 | 0,008794711 | 0,026809336 |
| ENSG00000197177.15 | ADGRA1 | protein_coding | 1681,212093 | -0,282687568 | 0,071371612 | -3,960784382 | 7,47E-05 | 0,000363806 |
| ENSG00000197180.2 | AC244090.1 | lncRNA | 362,1281227 | -0,482252671 | 0,139390353 | -3,459727741 | 0,000540722 | 0,002224659 |
| ENSG00000197183.14 | NOL4L | protein_coding | 11463,48611 | -0,21633481 | 0,050986077 | -4,243017395 | 2,21E-05 | 0,000117076 |
| ENSG00000197208.6 | SLC22A4 | protein_coding | 232,7180345 | 1,209429419 | 0,163928332 | 7,377793747 | 1,61E-13 | 2,17E-12 |
| ENSG00000197210.7 | AP000550.1 | lncRNA | 80,3102404 | -0,70939033 | 0,275248733 | -2,577270098 | 0,00995841 | 0,02981959 |
| ENSG00000197217.13 | ENTPD4 | protein_coding | 10979,03405 | -0,207006637 | 0,04906119 | -4,219356205 | 2,45E-05 | 0,000129241 |
| ENSG00000197249.13 | SERPINA1 | protein_coding | 40335,53531 | 0,217086464 | 0,042520467 | 5,105458197 | 3,30E-07 | 2,29E-06 |
| ENSG00000197256.10 | KANK2 | protein_coding | 3219,080752 | -0,189796733 | 0,060573053 | -3,133352606 | 0,001728217 | 0,006350055 |
| ENSG00000197261.11 | C6orf141 | protein_coding | 1384,665538 | 0,906621274 | 0,080735647 | 11,22950407 | 2,92E-29 | 8,90E-28 |
| ENSG00000197275.14 | RAD54B | protein_coding | 3750,848731 | -0,186206871 | 0,056629626 | -3,288152946 | 0,00100847 | 0,003904982 |
| ENSG00000197279.4 | ZNF165 | protein_coding | 711,816879 | 0,390217622 | 0,103094487 | 3,785048402 | 0,000153679 | 0,000706526 |
| ENSG00000197296.6 | FITM2 | protein_coding | 7562,688536 | -0,922561183 | 0,075304403 | -12,25109217 | 1,66E-34 | 6,24E-33 |
| ENSG00000197299.12 | BLM | protein_coding | 2752,1457 | -0,165375167 | 0,064468028 | -2,565227624 | 0,010310814 | 0,030715472 |
| ENSG00000197312.12 | DDI2 | protein_coding | 10411,60688 | -0,353040269 | 0,059486481 | -5,934798366 | 2,94E-09 | 2,60E-08 |
| ENSG00000197323.12 | TRIM33 | protein_coding | 11079,8891 | 0,209541459 | 0,045052644 | 4,651035764 | 3,30E-06 | 1,99E-05 |

| | | | | | | | | |
|---|---|---|---|---|---|---|---|---|
| ENSG00000197324.9 | LRP10 | protein_coding | 6632,729686 | 1,00177842 | 0,054151043 | 18,49970689 | 2,08E-76 | 1,86E-74 |
| ENSG00000197329.12 | PELI1 | protein_coding | 4388,579815 | 0,323460106 | 0,052785916 | 6,127772867 | 8,91E-10 | 8,37E-09 |
| ENSG00000197343.11 | ZNF655 | protein_coding | 10448,83002 | 0,16786965 | 0,047291189 | 3,549702476 | 0,000385667 | 0,00163761 |
| ENSG00000197355.11 | UAP1L1 | protein_coding | 2219,188019 | 0,314832747 | 0,06347096 | 4,960264477 | 7,04E-07 | 4,68E-06 |
| ENSG00000197361.7 | FBXL22 | protein_coding | 40,11194677 | -0,912529629 | 0,358759163 | -2,543571629 | 0,010972556 | 0,032403733 |
| ENSG00000197362.14 | ZNF786 | protein_coding | 1004,334034 | 0,266055738 | 0,086856194 | 3,06317517 | 0,002190018 | 0,007827107 |
| ENSG00000197372.9 | ZNF675 | protein_coding | 2924,266793 | 0,146744669 | 0,060620375 | 2,420715304 | 0,015490003 | 0,04359941 |
| ENSG00000197375.12 | SLC22A5 | protein_coding | 3516,221187 | 0,207857848 | 0,054053942 | 3,845378122 | 0,000120367 | 0,000565387 |
| ENSG00000197385.5 | ZNF860 | protein_coding | 686,5931442 | 0,264326324 | 0,1036344 | 2,550565485 | 0,010754832 | 0,031848508 |
| ENSG00000197417.7 | SHPK | protein_coding | 4552,330078 | 0,142308812 | 0,054980857 | 2,588333816 | 0,009644147 | 0,029025285 |
| ENSG00000197429.10 | IPP | protein_coding | 3031,452361 | -0,421129483 | 0,067316767 | -6,255937425 | 3,95E-10 | 3,87E-09 |
| ENSG00000197444.10 | OGDHL | protein_coding | 2692,082253 | -0,168408344 | 0,062821634 | -2,680738029 | 0,007346 | 0,022889135 |
| ENSG00000197448.14 | GSTK1 | protein_coding | 6361,0074 | 0,277118821 | 0,056957521 | 4,865359587 | 1,14E-06 | 7,36E-06 |
| ENSG00000197451.12 | HNRNPAB | protein_coding | 24621,28694 | -0,148868387 | 0,040826555 | -3,646361688 | 0,00026598 | 0,001166816 |
| ENSG00000197461.13 | PDGFA | protein_coding | 659,5276389 | 0,45456796 | 0,101609768 | 4,473663989 | 7,69E-06 | 4,37E-05 |
| ENSG00000197535.14 | MYO5A | protein_coding | 4963,708237 | -0,512549352 | 0,072982359 | -7,02292112 | 2,17E-12 | 2,67E-11 |
| ENSG00000197536.11 | C5orf56 | lncRNA | 1156,477258 | 5,81819027 | 0,151045383 | 38,51948448 | 0 | 0 |
| ENSG00000197555.9 | SIPA1L1 | protein_coding | 8983,261208 | 0,442482781 | 0,051439981 | 8,601923542 | 7,84E-18 | 1,41E-16 |
| ENSG00000197558.11 | SSPO | protein_coding | 161,7475875 | 1,113264589 | 0,291561421 | 3,818284962 | 0,000134383 | 0,00062406 |
| ENSG00000197566.10 | ZNF624 | protein_coding | 1639,26708 | -0,177892941 | 0,07457208 | -2,385516664 | 0,01705515 | 0,047411926 |
| ENSG00000197568.14 | HHLA3 | protein_coding | 235,7363178 | 0,39103461 | 0,15974543 | 2,447861019 | 0,014370708 | 0,040932216 |
| ENSG00000197579.8 | TOPORS | protein_coding | 6405,333155 | 0,319948894 | 0,048560572 | 6,588655848 | 4,44E-11 | 4,79E-10 |
| ENSG00000197580.11 | BCO2 | protein_coding | 118,7527897 | -0,591834882 | 0,232270322 | -2,548043496 | 0,010832896 | 0,032038685 |
| ENSG00000197584.12 | KCNMB2 | protein_coding | 2181,205456 | 0,74679178 | 0,065355062 | 11,42668613 | 3,08E-30 | 9,81E-29 |
| ENSG00000197586.12 | ENTPD6 | protein_coding | 8682,74906 | 0,742133766 | 0,050545041 | 14,68262259 | 8,33E-49 | 4,60E-47 |
| ENSG00000197594.13 | ENPP1 | protein_coding | 6482,03568 | 0,391694471 | 0,054145043 | 7,234170522 | 4,68E-13 | 6,07E-12 |
| ENSG00000197601.13 | FAR1 | protein_coding | 17707,25483 | 0,304635517 | 0,049654701 | 6,135079044 | 8,51E-10 | 8,02E-09 |
| ENSG00000197603.14 | CPLANE1 | protein_coding | 6211,216711 | 0,251951638 | 0,056030822 | 4,496661509 | 6,90E-06 | 3,96E-05 |
| ENSG00000197608.11 | ZNF841 | protein_coding | 2308,887936 | 0,361077071 | 0,074720597 | 4,832363317 | 1,35E-06 | 8,60E-06 |
| ENSG00000197635.10 | DPP4 | protein_coding | 24313,1239 | 0,451154784 | 0,048608794 | 9,281340788 | 1,67E-20 | 3,50E-19 |
| ENSG00000197647.11 | ZNF433 | protein_coding | 547,4942791 | 0,624038493 | 0,115002365 | 5,426310095 | 5,75E-08 | 4,38E-07 |
| ENSG00000197694.15 | SPTAN1 | protein_coding | 20351,48001 | 0,229251168 | 0,046161573 | 4,966277196 | 6,83E-07 | 4,54E-06 |
| ENSG00000197705.9 | KLHL14 | protein_coding | 752,6565612 | -0,344296852 | 0,096846423 | -3,555080735 | 0,000377863 | 0,001608723 |
| ENSG00000197712.12 | FAM114A1 | protein_coding | 797,1229289 | -0,272610556 | 0,095181766 | -2,864104831 | 0,004181896 | 0,013943163 |
| ENSG00000197714.9 | ZNF460 | protein_coding | 327,3315003 | 0,58104845 | 0,13814797 | 4,205986157 | 2,60E-05 | 0,000136168 |
| ENSG00000197721.16 | CR1L | protein_coding | 288,4379683 | 1,090079562 | 0,141193031 | 7,720491277 | 1,16E-14 | 1,70E-13 |

| | | | | | | | | |
|---|---|---|---|---|---|---|---|---|
| ENSG00000197728.11 | RPS26 | protein_coding | 18462,846 | -0,345150977 | 0,044163014 | -7,815385438 | 5,48E-15 | 8,26E-14 |
| ENSG00000197746.14 | PSAP | protein_coding | 112539,5381 | 0,148275641 | 0,037830744 | 3,919448214 | 8,88E-05 | 0,000426225 |
| ENSG00000197756.9 | RPL37A | protein_coding | 59887,24015 | -0,580779983 | 0,04060195 | -14,30423866 | 2,06E-46 | 1,09E-44 |
| ENSG00000197780.9 | TAF13 | protein_coding | 3355,032631 | -0,20785275 | 0,064004986 | -3,247446245 | 0,001164457 | 0,004442501 |
| ENSG00000197798.9 | FAM118B | protein_coding | 3233,94305 | 0,14918186 | 0,053974856 | 2,763914012 | 0,005711259 | 0,018357093 |
| ENSG00000197808.12 | ZNF461 | protein_coding | 1296,965143 | 0,219653612 | 0,075955963 | 2,891854738 | 0,00382975 | 0,012893185 |
| ENSG00000197822.11 | OCLN | protein_coding | 4087,415425 | 0,697705411 | 0,058013931 | 12,02651507 | 2,58E-33 | 9,39E-32 |
| ENSG00000197837.3 | HIST4H4 | protein_coding | 288,1255759 | 0,586666057 | 0,151743627 | 3,866166041 | 0,00011056 | 0,000521717 |
| ENSG00000197841.15 | ZNF181 | protein_coding | 2541,007043 | 0,2487993 | 0,067779276 | 3,670728179 | 0,00024186 | 0,001069135 |
| ENSG00000197857.13 | ZNF44 | protein_coding | 2129,373093 | 0,292960409 | 0,062676352 | 4,674177727 | 2,95E-06 | 1,79E-05 |
| ENSG00000197863.9 | ZNF790 | protein_coding | 1479,483914 | 0,251657083 | 0,079429278 | 3,168316375 | 0,001533246 | 0,005707264 |
| ENSG00000197872.11 | FAM49A | protein_coding | 2339,345653 | -0,901324281 | 0,075889915 | -11,87673331 | 1,56E-32 | 5,60E-31 |
| ENSG00000197885.10 | NKIRAS1 | protein_coding | 2306,598921 | 0,337363886 | 0,062388369 | 5,40748045 | 6,39E-08 | 4,84E-07 |
| ENSG00000197903.7 | HIST1H2BK | protein_coding | 1201,189648 | 0,262203613 | 0,08956638 | 2,927478068 | 0,003417232 | 0,011659613 |
| ENSG00000197937.12 | ZNF347 | protein_coding | 210,9654536 | -1,345265825 | 0,513305102 | -2,620791846 | 0,008772581 | 0,026752746 |
| ENSG00000197943.10 | PLCG2 | protein_coding | 6401,993855 | 0,533593058 | 0,046896877 | 11,37800842 | 5,38E-30 | 1,69E-28 |
| ENSG00000197948.11 | FCHSD1 | protein_coding | 1621,960854 | 0,605123104 | 0,084638621 | 7,149491562 | 8,71E-13 | 1,11E-11 |
| ENSG00000197951.9 | ZNF71 | protein_coding | 1750,162415 | -0,217635453 | 0,0646319 | -3,36730704 | 0,000759061 | 0,003022415 |
| ENSG00000197956.10 | S100A6 | protein_coding | 19045,5751 | 1,082375734 | 0,054612487 | 19,81919861 | 2,03E-87 | 2,11E-85 |
| ENSG00000197958.13 | RPL12 | protein_coding | 28488,90137 | -0,458395182 | 0,050783714 | -9,026420939 | 1,77E-19 | 3,49E-18 |
| ENSG00000197959.14 | DNM3 | protein_coding | 492,8082467 | -0,311611443 | 0,114459828 | -2,722452491 | 0,006479935 | 0,020514326 |
| ENSG00000197961.11 | ZNF121 | protein_coding | 3053,024954 | 0,199117045 | 0,059422197 | 3,350886633 | 0,000805533 | 0,003192207 |
| ENSG00000197971.15 | MBP | protein_coding | 8148,745441 | -0,221991987 | 0,047923103 | -4,632254019 | 3,62E-06 | 2,16E-05 |
| ENSG00000197976.12 | AKAP17A | protein_coding | 5914,781273 | 0,19833361 | 0,059244599 | 3,34770784 | 0,000814829 | 0,003223742 |
| ENSG00000197977.4 | ELOVL2 | protein_coding | 12776,75718 | -0,14663238 | 0,042445415 | -3,454610561 | 0,000551088 | 0,002263585 |
| ENSG00000198001.13 | IRAK4 | protein_coding | 3126,75037 | 0,386318625 | 0,069602254 | 5,550375219 | 2,85E-08 | 2,26E-07 |
| ENSG00000198015.13 | MRPL42 | protein_coding | 14046,19022 | -0,17908216 | 0,051345928 | -3,487757804 | 0,000487089 | 0,002024055 |
| ENSG00000198018.7 | ENTPD7 | protein_coding | 5230,139906 | -0,307163721 | 0,047555481 | -6,459060303 | 1,05E-10 | 1,09E-09 |
| ENSG00000198034.11 | RPS4X | protein_coding | 51405,88692 | -0,806730827 | 0,044945604 | -17,94904836 | 4,88E-72 | 4,05E-70 |
| ENSG00000198039.11 | AC092161.1 | protein_coding | 1314,314238 | 0,498067971 | 0,07552441 | 6,594794577 | 4,26E-11 | 4,60E-10 |
| ENSG00000198046.12 | ZNF667 | protein_coding | 2174,152781 | -0,281372924 | 0,071767742 | -3,92060439 | 8,83E-05 | 0,000424729 |
| ENSG00000198049.7 | AVPR1B | protein_coding | 32,01106535 | 1,370783939 | 0,441374371 | 3,105717117 | 0,001898182 | 0,006899109 |
| ENSG00000198053.11 | SIRPA | protein_coding | 5188,866934 | 0,377644461 | 0,052391152 | 7,208172447 | 5,67E-13 | 7,31E-12 |
| ENSG00000198054.12 | DSCR8 | lncRNA | 6074,538411 | -0,320669747 | 0,053218629 | -6,025516877 | 1,69E-09 | 1,53E-08 |
| ENSG00000198055.11 | GRK6 | protein_coding | 2201,638896 | -0,445454087 | 0,087467214 | -5,092812094 | 3,53E-07 | 2,44E-06 |
| ENSG00000198060.10 | mars-05 | protein_coding | 5348,883052 | -0,148430829 | 0,060040196 | -2,472190946 | 0,013428777 | 0,038574316 |

| | | | | | | | | |
|---|---|---|---|---|---|---|---|---|
| ENSG00000198081.11 | ZBTB14 | protein_coding | 2690,825149 | -0,239296896 | 0,06764155 | -3,537720479 | 0,000403597 | 0,001706022 |
| ENSG00000198087.7 | CD2AP | protein_coding | 14902,89138 | 0,481284762 | 0,043737384 | 11,00396766 | 3,66E-28 | 1,08E-26 |
| ENSG00000198088.10 | NUP62CL | protein_coding | 490,1480507 | -0,37598916 | 0,129318358 | -2,90746933 | 0,003643661 | 0,01234053 |
| ENSG00000198089.15 | SFI1 | protein_coding | 4273,635923 | 0,374086726 | 0,087413692 | 4,279498079 | 1,87E-05 | 0,000100459 |
| ENSG00000198093.11 | ZNF649 | protein_coding | 1853,272545 | -0,24438503 | 0,07701207 | -3,173334109 | 0,00150699 | 0,005622202 |
| ENSG00000198105.14 | ZNF248 | protein_coding | 5372,771514 | -0,203752493 | 0,0494024 | -4,124344067 | 3,72E-05 | 0,000190068 |
| ENSG00000198113.3 | TOR4A | protein_coding | 1316,478105 | -0,36024666 | 0,074464987 | -4,837799262 | 1,31E-06 | 8,39E-06 |
| ENSG00000198130.16 | HIBCH | protein_coding | 3220,007155 | -0,192086563 | 0,058474003 | -3,284990836 | 0,001019857 | 0,003945001 |
| ENSG00000198133.8 | TMEM229B | protein_coding | 446,5528307 | 2,99216138 | 0,36021274 | 8,306650626 | 9,84E-17 | 1,66E-15 |
| ENSG00000198160.14 | MIER1 | protein_coding | 6777,980924 | 0,54991026 | 0,05376717 | 10,22762137 | 1,49E-24 | 3,80E-23 |
| ENSG00000198162.12 | MAN1A2 | protein_coding | 11215,25916 | -0,345273308 | 0,049385572 | -6,991380134 | 2,72E-12 | 3,31E-11 |
| ENSG00000198168.9 | SVIP | protein_coding | 12383,27415 | -0,51676592 | 0,047185697 | -10,95174928 | 6,52E-28 | 1,89E-26 |
| ENSG00000198176.13 | TFDP1 | protein_coding | 19226,28475 | -0,33564204 | 0,044029458 | -7,623124462 | 2,48E-14 | 3,53E-13 |
| ENSG00000198182.13 | ZNF607 | protein_coding | 2828,589522 | 0,241811744 | 0,062538007 | 3,866636559 | 0,000110347 | 0,00052093 |
| ENSG00000198185.11 | ZNF334 | protein_coding | 5693,296773 | 0,225617385 | 0,049250706 | 4,580997985 | 4,63E-06 | 2,73E-05 |
| ENSG00000198198.16 | SZT2 | protein_coding | 5230,413283 | 0,260485196 | 0,063372339 | 4,110392633 | 3,95E-05 | 0,000201285 |
| ENSG00000198208.11 | RPS6KL1 | protein_coding | 2117,773626 | 0,415534173 | 0,089763284 | 4,629222046 | 3,67E-06 | 2,19E-05 |
| ENSG00000198242.14 | RPL23A | protein_coding | 30178,46222 | -0,494757317 | 0,044283574 | -11,17247932 | 5,56E-29 | 1,68E-27 |
| ENSG00000198258.10 | UBL5 | protein_coding | 10948,00348 | -0,241588412 | 0,051329339 | -4,706634045 | 2,52E-06 | 1,54E-05 |
| ENSG00000198300.14 | PEG3 | protein_coding | 6604,644869 | -0,488562273 | 0,048984122 | -9,973890547 | 1,98E-23 | 4,77E-22 |
| ENSG00000198301.12 | SDAD1 | protein_coding | 5811,933364 | 0,166829001 | 0,053156433 | 3,138453635 | 0,001698418 | 0,006249745 |
| ENSG00000198315.11 | ZKSCAN8 | protein_coding | 7432,112699 | 0,240761985 | 0,047048254 | 5,117341598 | 3,10E-07 | 2,16E-06 |
| ENSG00000198324.13 | PHETA1 | protein_coding | 4424,859246 | -0,379684923 | 0,075590498 | -5,022918657 | 5,09E-07 | 3,45E-06 |
| ENSG00000198331.10 | HYLS1 | protein_coding | 2724,177874 | -0,296387108 | 0,06224497 | -4,761623436 | 1,92E-06 | 1,20E-05 |
| ENSG00000198342.10 | ZNF442 | protein_coding | 189,5756043 | 0,813243366 | 0,202106718 | 4,023831442 | 5,73E-05 | 0,000284261 |
| ENSG00000198346.11 | ZNF813 | protein_coding | 1562,474096 | 0,451679844 | 0,080539276 | 5,608193528 | 2,04E-08 | 1,65E-07 |
| ENSG00000198353.8 | HOXC4 | protein_coding | 457,0197566 | 0,553880885 | 0,133253237 | 4,15660359 | 3,23E-05 | 0,000166874 |
| ENSG00000198356.11 | ASNA1 | protein_coding | 4816,578085 | -0,195659062 | 0,050427937 | -3,879973543 | 0,000104468 | 0,000495328 |
| ENSG00000198363.18 | ASPH | protein_coding | 31680,64311 | 0,401154441 | 0,042127085 | 9,522482839 | 1,69E-21 | 3,71E-20 |
| ENSG00000198373.12 | WWP2 | protein_coding | 2725,050524 | 0,156865642 | 0,061452676 | 2,552625097 | 0,010691451 | 0,031681657 |
| ENSG00000198380.12 | GFPT1 | protein_coding | 25136,9856 | -0,128644309 | 0,044126195 | -2,915372841 | 0,003552638 | 0,012057651 |
| ENSG00000198382.9 | UVRAG | protein_coding | 4125,169965 | 0,571887344 | 0,151236551 | 3,781409578 | 0,000155943 | 0,000714892 |
| ENSG00000198393.8 | ZNF26 | protein_coding | 2710,684835 | 0,264271658 | 0,060121055 | 4,395659001 | 1,10E-05 | 6,15E-05 |
| ENSG00000198408.14 | OGA | protein_coding | 15017,71166 | 0,119007826 | 0,048467563 | 2,455411827 | 0,014072327 | 0,040212519 |
| ENSG00000198417.7 | MT1F | protein_coding | 1560,595767 | 1,422402962 | 0,07825733 | 18,17597107 | 8,00E-74 | 6,87E-72 |
| ENSG00000198420.10 | TCAF1 | protein_coding | 38148,44004 | -0,216976795 | 0,047327244 | -4,584606607 | 4,55E-06 | 2,68E-05 |

| | | | | | | | | |
|---|---|---|---|---|---|---|---|---|
| ENSG00000198435.4 | NRARP | protein_coding | 246,4418362 | -1,214004816 | 0,166252745 | -7,302164059 | 2,83E-13 | 3,75E-12 |
| ENSG00000198440.9 | ZNF583 | protein_coding | 1687,09611 | 0,188938215 | 0,079712024 | 2,370259922 | 0,017775584 | 0,049057212 |
| ENSG00000198466.12 | ZNF587 | protein_coding | 5015,950765 | 0,149183512 | 0,060273509 | 2,475109117 | 0,013319547 | 0,038314338 |
| ENSG00000198467.15 | TPM2 | protein_coding | 90,37689939 | 0,811258736 | 0,268077235 | 3,026212714 | 0,00247638 | 0,008727234 |
| ENSG00000198478.8 | SH3BGRL2 | protein_coding | 4043,567773 | -0,454298367 | 0,065612824 | -6,923926456 | 4,39E-12 | 5,23E-11 |
| ENSG00000198482.12 | ZNF808 | protein_coding | 471,7462799 | 0,854802926 | 0,126098304 | 6,778861385 | 1,21E-11 | 1,38E-10 |
| ENSG00000198492.16 | YTHDF2 | protein_coding | 12783,7711 | 0,118222503 | 0,041834825 | 2,82593514 | 0,004714281 | 0,015479759 |
| ENSG00000198513.11 | ATL1 | protein_coding | 2683,730197 | -0,271892837 | 0,059851869 | -4,542762658 | 5,55E-06 | 3,23E-05 |
| ENSG00000198521.11 | ZNF43 | protein_coding | 2500,168515 | 0,236051945 | 0,059435938 | 3,971535589 | 7,14E-05 | 0,000348901 |
| ENSG00000198535.5 | C2CD4A | protein_coding | 1900,576859 | 1,183766205 | 0,096455617 | 12,27265182 | 1,27E-34 | 4,80E-33 |
| ENSG00000198538.11 | ZNF28 | protein_coding | 4906,339276 | 0,360865734 | 0,060646026 | 5,950360775 | 2,68E-09 | 2,38E-08 |
| ENSG00000198554.12 | WDHD1 | protein_coding | 8039,699685 | -0,232167859 | 0,049694997 | -4,671855818 | 2,98E-06 | 1,81E-05 |
| ENSG00000198561.13 | CTNND1 | protein_coding | 22383,19293 | 0,101698399 | 0,039525715 | 2,572968005 | 0,010083053 | 0,030144699 |
| ENSG00000198576.4 | ARC | protein_coding | 37,16982809 | 1,433903085 | 0,408673423 | 3,508677109 | 0,000450341 | 0,001889419 |
| ENSG00000198586.14 | TLK1 | protein_coding | 9254,704533 | 0,123063711 | 0,044837243 | 2,744676146 | 0,006057064 | 0,019322343 |
| ENSG00000198589.13 | LRBA | protein_coding | 22151,74748 | -0,345759623 | 0,056507984 | -6,118774728 | 9,43E-10 | 8,83E-09 |
| ENSG00000198598.6 | MMP17 | protein_coding | 1421,190453 | 0,432721781 | 0,089413276 | 4,839569687 | 1,30E-06 | 8,32E-06 |
| ENSG00000198604.11 | BAZ1A | protein_coding | 5311,586481 | 1,246371199 | 0,073686892 | 16,91442218 | 3,52E-64 | 2,62E-62 |
| ENSG00000198612.11 | COPS8 | protein_coding | 5541,357217 | -0,28162272 | 0,051612116 | -5,456523428 | 4,86E-08 | 3,73E-07 |
| ENSG00000198624.13 | CCDC69 | protein_coding | 398,3824212 | 0,884305753 | 0,142618274 | 6,200508009 | 5,63E-10 | 5,43E-09 |
| ENSG00000198625.13 | MDM4 | protein_coding | 3449,215429 | 0,339197738 | 0,053898815 | 6,293231771 | 3,11E-10 | 3,09E-09 |
| ENSG00000198626.17 | RYR2 | protein_coding | 3193,486781 | 0,470678571 | 0,068026265 | 6,919071234 | 4,55E-12 | 5,40E-11 |
| ENSG00000198646.14 | NCOA6 | protein_coding | 14837,43105 | 0,350729052 | 0,042399655 | 8,271978972 | 1,32E-16 | 2,20E-15 |
| ENSG00000198663.16 | C6orf89 | protein_coding | 15737,18969 | -0,154471285 | 0,048162829 | -3,207271868 | 0,001340003 | 0,005049745 |
| ENSG00000198668.12 | CALM1 | protein_coding | 63815,62871 | -0,404227906 | 0,052152265 | -7,750917556 | 9,12E-15 | 1,35E-13 |
| ENSG00000198682.13 | PAPSS2 | protein_coding | 8974,744832 | -0,561577606 | 0,052315758 | -10,7343873 | 7,02E-27 | 1,96E-25 |
| ENSG00000198689.11 | SLC9A6 | protein_coding | 4782,053867 | -0,166384523 | 0,05903616 | -2,818349362 | 0,004827126 | 0,015806484 |
| ENSG00000198695.2 | MT-ND6 | protein_coding | 4476,3863 | -0,792573753 | 0,281840291 | -2,812137864 | 0,004921341 | 0,016100938 |
| ENSG00000198707.15 | CEP290 | protein_coding | 4718,006766 | 0,251309779 | 0,053937802 | 4,659251424 | 3,17E-06 | 1,92E-05 |
| ENSG00000198712.1 | MT-CO2 | protein_coding | 377351,225 | -0,702875735 | 0,047517621 | -14,79189671 | 1,65E-49 | 9,37E-48 |
| ENSG00000198715.13 | GLMP | protein_coding | 4571,310166 | 0,161948731 | 0,053131266 | 3,048087204 | 0,002303031 | 0,008186816 |
| ENSG00000198718.13 | TOGARAM1 | protein_coding | 4894,493658 | 0,366184559 | 0,085275694 | 4,294125795 | 1,75E-05 | 9,44E-05 |
| ENSG00000198719.9 | DLL1 | protein_coding | 772,332817 | 0,562778871 | 0,101616739 | 5,538249658 | 3,06E-08 | 2,41E-07 |
| ENSG00000198720.13 | ANKRD13B | protein_coding | 2035,814286 | -0,461493155 | 0,080964007 | -5,699979216 | 1,20E-08 | 9,87E-08 |
| ENSG00000198722.14 | UNC13B | protein_coding | 21957,97567 | -0,139005101 | 0,046332411 | -3,000169789 | 0,002698291 | 0,009429567 |
| ENSG00000198727.2 | MT-CYB | protein_coding | 135744,6164 | -1,398118649 | 0,05185791 | -26,96056706 | 4,29E-160 | 7,37E-158 |

| | | | | | | | | |
|---|---|---|---|---|---|---|---|---|
| ENSG00000198728.10 | LDB1 | protein_coding | 5959,914115 | -0,23356526 | 0,051818451 | -4,50737633 | 6,56E-06 | 3,77E-05 |
| ENSG00000198729.5 | PPP1R14C | protein_coding | 3003,600589 | -0,396378146 | 0,061147182 | -6,482361652 | 9,03E-11 | 9,45E-10 |
| ENSG00000198734.11 | F5 | protein_coding | 2838,063851 | 0,309457362 | 0,062615025 | 4,942222101 | 7,72E-07 | 5,09E-06 |
| ENSG00000198740.8 | ZNF652 | protein_coding | 3194,854866 | 0,22798408 | 0,069019662 | 3,30317584 | 0,000955964 | 0,003724091 |
| ENSG00000198742.9 | SMURF1 | protein_coding | 3493,400168 | 0,223086781 | 0,05555676 | 4,015475019 | 5,93E-05 | 0,000293682 |
| ENSG00000198755.11 | RPL10A | protein_coding | 21665,42731 | -0,551874797 | 0,044686439 | -12,34993907 | 4,87E-35 | 1,87E-33 |
| ENSG00000198759.12 | EGFL6 | protein_coding | 46,6961006 | 1,380209121 | 0,384718663 | 3,587580364 | 0,000333761 | 0,001435899 |
| ENSG00000198763.3 | MT-ND2 | protein_coding | 181950,6317 | -0,552195636 | 0,040272387 | -13,71151984 | 8,66E-43 | 4,25E-41 |
| ENSG00000198786.2 | MT-ND5 | protein_coding | 162894,3556 | -0,918746191 | 0,171977848 | -5,342235656 | 9,18E-08 | 6,80E-07 |
| ENSG00000198791.12 | CNOT7 | protein_coding | 23529,14728 | -0,272992382 | 0,050867024 | -5,366785028 | 8,02E-08 | 5,96E-07 |
| ENSG00000198799.12 | LRIG2 | protein_coding | 3560,786349 | 0,229460717 | 0,05993022 | 3,828798169 | 0,000128771 | 0,000601223 |
| ENSG00000198804.2 | MT-CO1 | protein_coding | 572816,1858 | -0,612811258 | 0,061371089 | -9,985341078 | 1,77E-23 | 4,27E-22 |
| ENSG00000198805.11 | PNP | protein_coding | 4755,344401 | 0,422914898 | 0,066839683 | 6,327302571 | 2,49E-10 | 2,50E-09 |
| ENSG00000198814.12 | GK | protein_coding | 9884,247763 | 0,208389093 | 0,044847701 | 4,646594763 | 3,37E-06 | 2,03E-05 |
| ENSG00000198818.10 | SFT2D1 | protein_coding | 1206,28962 | 0,46327168 | 0,087646047 | 5,285711084 | 1,25E-07 | 9,15E-07 |
| ENSG00000198825.14 | INPP5F | protein_coding | 1666,009954 | -0,216145253 | 0,068722093 | -3,145207627 | 0,00165969 | 0,006126265 |
| ENSG00000198826.11 | ARHGAP11A | protein_coding | 12191,80452 | -0,422419464 | 0,046212624 | -9,140780753 | 6,20E-20 | 1,25E-18 |
| ENSG00000198830.11 | HMGN2 | protein_coding | 96196,13359 | -0,20444688 | 0,043167462 | -4,736133922 | 2,18E-06 | 1,35E-05 |
| ENSG00000198833.7 | UBE2J1 | protein_coding | 11287,29417 | 0,198641878 | 0,043364676 | 4,580730142 | 4,63E-06 | 2,73E-05 |
| ENSG00000198835.4 | GJC2 | protein_coding | 717,0974474 | -1,196984531 | 0,122793743 | -9,747927687 | 1,88E-22 | 4,34E-21 |
| ENSG00000198836.10 | OPA1 | protein_coding | 11414,79874 | -0,216894989 | 0,044872391 | -4,833595485 | 1,34E-06 | 8,55E-06 |
| ENSG00000198837.10 | DENND4B | protein_coding | 4820,248902 | 0,208448524 | 0,059016377 | 3,532045447 | 0,000412358 | 0,001740442 |
| ENSG00000198840.2 | MT-ND3 | protein_coding | 43780,87863 | -1,518931736 | 0,064488665 | -23,55346859 | 1,16E-122 | 1,59E-120 |
| ENSG00000198841.3 | KTI12 | protein_coding | 1923,769696 | 0,20316458 | 0,07538933 | 2,694871805 | 0,007041572 | 0,02208426 |
| ENSG00000198843.13 | SELENOT | protein_coding | 17379,45744 | -0,111108203 | 0,045210942 | -2,457551164 | 0,013988788 | 0,040009357 |
| ENSG00000198855.7 | FICD | protein_coding | 1050,801359 | 0,37160675 | 0,085543237 | 4,344080974 | 1,40E-05 | 7,65E-05 |
| ENSG00000198863.7 | RUNDC1 | protein_coding | 4625,363299 | 0,408081854 | 0,052171604 | 7,821915092 | 5,20E-15 | 7,85E-14 |
| ENSG00000198879.11 | SFMBT2 | protein_coding | 3170,012347 | 0,589052282 | 0,064780635 | 9,093030454 | 9,63E-20 | 1,92E-18 |
| ENSG00000198886.2 | MT-ND4 | protein_coding | 290111,0086 | -0,89788068 | 0,12739966 | -7,047747842 | 1,82E-12 | 2,25E-11 |
| ENSG00000198888.2 | MT-ND1 | protein_coding | 91597,93651 | -0,35898556 | 0,059840176 | -5,999072595 | 1,98E-09 | 1,79E-08 |
| ENSG00000198892.6 | SHISA4 | protein_coding | 134,9078302 | 0,910554808 | 0,22508865 | 4,045316406 | 5,23E-05 | 0,000261135 |
| ENSG00000198894.8 | CIPC | protein_coding | 4376,415996 | 0,142791847 | 0,057997403 | 2,46203864 | 0,013814978 | 0,039577618 |
| ENSG00000198899.2 | MT-ATP6 | protein_coding | 39457,48512 | -1,246361478 | 0,05440773 | -22,90780166 | 3,88E-116 | 5,14E-114 |
| ENSG00000198901.14 | PRC1 | protein_coding | 14041,08626 | -0,662313641 | 0,046438288 | -14,26223198 | 3,76E-46 | 1,98E-44 |
| ENSG00000198909.7 | MAP3K3 | protein_coding | 1680,419456 | 0,26997761 | 0,075201498 | 3,590056288 | 0,000330607 | 0,001423962 |
| ENSG00000198910.13 | L1CAM | protein_coding | 1006,013212 | 0,218976378 | 0,088014925 | 2,487945976 | 0,012848325 | 0,037105844 |

| | | | | | | | | |
|---|---|---|---|---|---|---|---|---|
| ENSG00000198911.12 | SREBF2 | protein_coding | 31311,44756 | -0,113370369 | 0,046557134 | -2,4350805 | 0,014888477 | 0,042143452 |
| ENSG00000198912.11 | C1orf174 | protein_coding | 1580,070168 | -0,168883842 | 0,067775257 | -2,491821482 | 0,01270899 | 0,036774219 |
| ENSG00000198915.11 | RASGEF1A | protein_coding | 137,5840579 | -0,70604922 | 0,209161875 | -3,375611456 | 0,000736519 | 0,002943589 |
| ENSG00000198918.8 | RPL39 | protein_coding | 33930,76286 | -0,414185228 | 0,039095059 | -10,59431127 | 3,17E-26 | 8,67E-25 |
| ENSG00000198919.13 | DZIP3 | protein_coding | 10817,73729 | -0,193097591 | 0,05127838 | -3,765672635 | 0,000166101 | 0,000756837 |
| ENSG00000198931.10 | APRT | protein_coding | 3648,122118 | -0,2944089 | 0,06327659 | -4,65273019 | 3,28E-06 | 1,97E-05 |
| ENSG00000198932.13 | GPRASP1 | protein_coding | 3080,282808 | 0,496967682 | 0,063173666 | 7,866690568 | 3,64E-15 | 5,55E-14 |
| ENSG00000198937.9 | CCDC167 | protein_coding | 1966,050333 | -0,90315938 | 0,067533522 | -13,37349736 | 8,64E-41 | 3,95E-39 |
| ENSG00000198938.2 | MT-CO3 | protein_coding | 169039,4124 | -1,633782319 | 0,052269806 | -31,25671302 | 1,81E-214 | 3,88E-212 |
| ENSG00000198944.5 | SOWAHA | protein_coding | 11491,77557 | -0,277618061 | 0,048441186 | -5,731033499 | 9,98E-09 | 8,29E-08 |
| ENSG00000198947.15 | DMD | protein_coding | 1744,894197 | 0,548482167 | 0,073311791 | 7,481499994 | 7,35E-14 | 1,02E-12 |
| ENSG00000198948.12 | MFAP3L | protein_coding | 215,4199498 | -0,622007285 | 0,189505839 | -3,282259207 | 0,001029789 | 0,003978634 |
| ENSG00000198952.8 | SMG5 | protein_coding | 17350,79317 | 0,319204286 | 0,047638209 | 6,700593749 | 2,08E-11 | 2,31E-10 |
| ENSG00000198960.11 | ARMCX6 | protein_coding | 3359,446071 | 0,172241593 | 0,056906943 | 3,026723706 | 0,002472198 | 0,00871386 |
| ENSG00000198964.14 | SGMS1 | protein_coding | 3009,545744 | 0,27206313 | 0,055747537 | 4,880271764 | 1,06E-06 | 6,86E-06 |
| ENSG00000198967.4 | OR10Z1 | protein_coding | 11,4278292 | -2,415028083 | 0,755361665 | -3,19718116 | 0,001387777 | 0,005210589 |
| ENSG00000203326.11 | ZNF525 | protein_coding | 3133,150624 | 0,420974006 | 0,07091635 | 5,936205235 | 2,92E-09 | 2,58E-08 |
| ENSG00000203666.12 | EFCAB2 | protein_coding | 2934,48258 | -0,259031793 | 0,061576512 | -4,20666557 | 2,59E-05 | 0,000135917 |
| ENSG00000203667.10 | COX20 | protein_coding | 4620,337962 | -0,473524596 | 0,060453571 | -7,832863893 | 4,77E-15 | 7,20E-14 |
| ENSG00000203668.2 | CHML | protein_coding | 6360,446524 | -0,280416966 | 0,048727259 | -5,754827464 | 8,67E-09 | 7,26E-08 |
| ENSG00000203697.11 | CAPN8 | protein_coding | 70,18214582 | -1,078756532 | 0,340991241 | -3,163590154 | 0,001558361 | 0,005791972 |
| ENSG00000203706.8 | SERTAD4-AS1 | lncRNA | 17,82921026 | -1,843401131 | 0,572988557 | -3,217169189 | 0,001294622 | 0,004890192 |
| ENSG00000203711.14 | C6orf99 | lncRNA | 137,2235481 | -0,791739665 | 0,23225492 | -3,408925262 | 0,000652193 | 0,002636534 |
| ENSG00000203724.11 | C1orf53 | protein_coding | 398,6232999 | -0,573814069 | 0,121766868 | -4,712399034 | 2,45E-06 | 1,50E-05 |
| ENSG00000203727.4 | SAMD5 | protein_coding | 37076,83157 | -0,330767655 | 0,048478922 | -6,822916805 | 8,92E-12 | 1,03E-10 |
| ENSG00000203760.8 | CENPW | protein_coding | 1398,510039 | -0,55627239 | 0,075608978 | -7,357226689 | 1,88E-13 | 2,52E-12 |
| ENSG00000203778.8 | FAM229B | protein_coding | 1929,10002 | 0,400410935 | 0,081584207 | 4,907946668 | 9,20E-07 | 6,01E-06 |
| ENSG00000203791.15 | EEF1AKMT2 | protein_coding | 975,0836949 | -0,225589636 | 0,094785131 | -2,380010802 | 0,017312131 | 0,048001817 |
| ENSG00000203797.10 | DDO | protein_coding | 107,9886252 | 4,025315184 | 0,321628653 | 12,51541226 | 6,15E-36 | 2,45E-34 |
| ENSG00000203799.13 | CCDC162P | transcribed_unitary | 41,54948053 | 3,186641761 | 0,444216274 | 7,173626786 | 7,30E-13 | 9,35E-12 |
| ENSG00000203811.1 | HIST2H3C | protein_coding | 9,41681782 | -2,342230123 | 0,969094139 | -2,416927344 | 0,015652141 | 0,043967808 |
| ENSG00000203814.6 | HIST2H2BF | protein_coding | 105,9623407 | 0,980489681 | 0,290805416 | 3,37163487 | 0,000747234 | 0,002981652 |
| ENSG00000203875.12 | SNHG5 | lncRNA | 2748,005807 | -0,273222008 | 0,064493159 | -4,236449414 | 2,27E-05 | 0,000120353 |
| ENSG00000203877.8 | RIPPLY2 | protein_coding | 746,6885855 | -0,252255628 | 0,100089702 | -2,52029552 | 0,011725634 | 0,034276923 |
| ENSG00000203879.12 | GDI1 | protein_coding | 17107,76353 | -0,133996821 | 0,042459094 | -3,155903939 | 0,001600015 | 0,005926403 |
| ENSG00000203880.12 | PCMTD2 | protein_coding | 16995,67011 | -0,276070683 | 0,04088899 | -6,751711928 | 1,46E-11 | 1,65E-10 |

| | | | | | | | | |
|---|---|---|---|---|---|---|---|---|
| ENSG00000203943.8 | SAMD13 | protein_coding | 135,7084377 | -0,524055727 | 0,203960856 | -2,569393642 | 0,010187666 | 0,030400924 |
| ENSG00000203985.11 | LDLRAD1 | protein_coding | 435,9961245 | -0,534575133 | 0,130965983 | -4,081786142 | 4,47E-05 | 0,000225751 |
| ENSG00000204003.8 | AL355987.1 | protein_coding | 55,58719854 | -1,072761219 | 0,360752959 | -2,973672682 | 0,002942588 | 0,010182123 |
| ENSG00000204065.3 | TCEAL5 | protein_coding | 3787,125301 | 0,29422651 | 0,052402145 | 5,614779847 | 1,97E-08 | 1,59E-07 |
| ENSG00000204084.13 | INPP5B | protein_coding | 1216,972115 | 0,457118964 | 0,093253557 | 4,90189308 | 9,49E-07 | 6,18E-06 |
| ENSG00000204103.4 | MAFB | protein_coding | 28050,42903 | -0,165880842 | 0,049219735 | -3,370209973 | 0,000751109 | 0,00299446 |
| ENSG00000204104.12 | TRAF3IP1 | protein_coding | 2538,421023 | -0,623309061 | 0,086447966 | -7,210222392 | 5,59E-13 | 7,21E-12 |
| ENSG00000204116.11 | CHIC1 | protein_coding | 2231,400834 | 0,317292508 | 0,06256534 | 5,07137834 | 3,95E-07 | 2,72E-06 |
| ENSG00000204128.6 | C2orf72 | protein_coding | 5564,621751 | -0,230376528 | 0,05888059 | -3,912605601 | 9,13E-05 | 0,000437742 |
| ENSG00000204147.10 | ASAH2B | protein_coding | 4214,149844 | 0,239634493 | 0,059799855 | 4,007275469 | 6,14E-05 | 0,000303729 |
| ENSG00000204174.7 | NPY4R | protein_coding | 152,6936146 | -0,571030129 | 0,196283986 | -2,909203862 | 0,003623505 | 0,012279656 |
| ENSG00000204177.10 | BMS1P1 | transcribed_unproc | 1585,611036 | 0,360337728 | 0,083107015 | 4,335828089 | 1,45E-05 | 7,92E-05 |
| ENSG00000204178.11 | MACO1 | protein_coding | 6364,685129 | -0,121762817 | 0,047522595 | -2,562208939 | 0,010400873 | 0,030926452 |
| ENSG00000204217.14 | BMPR2 | protein_coding | 7432,978826 | 0,524253318 | 0,060575422 | 8,654555009 | 4,95E-18 | 9,03E-17 |
| ENSG00000204219.10 | TCEA3 | protein_coding | 281,6566876 | -0,452915551 | 0,13853673 | -3,269281382 | 0,00107821 | 0,004147912 |
| ENSG00000204228.4 | HSD17B8 | protein_coding | 638,2829777 | -0,492726615 | 0,104264247 | -4,72574854 | 2,29E-06 | 1,42E-05 |
| ENSG00000204241.8 | AP000911.1 | lncRNA | 180,4960143 | 0,47289436 | 0,186975722 | 2,529175205 | 0,011433094 | 0,033513154 |
| ENSG00000204256.13 | BRD2 | protein_coding | 37031,3027 | 0,640837532 | 0,048594749 | 13,18738232 | 1,04E-39 | 4,65E-38 |
| ENSG00000204261.9 | PSMB8-AS1 | lncRNA | 2682,97255 | 6,112235495 | 0,234024554 | 26,11792392 | 2,28E-150 | 3,75E-148 |
| ENSG00000204264.11 | PSMB8 | protein_coding | 11609,06227 | 5,595071948 | 0,191094012 | 29,27915893 | 1,91E-188 | 3,68E-186 |
| ENSG00000204267.15 | TAP2 | protein_coding | 13389,11034 | 5,528256064 | 0,134869177 | 40,98976644 | 0 | 0 |
| ENSG00000204304.12 | PBX2 | protein_coding | 11058,04837 | 0,174162707 | 0,045242366 | 3,84954905 | 0,000118335 | 0,000556543 |
| ENSG00000204308.8 | RNF5 | protein_coding | 4109,327734 | -0,258096881 | 0,054952433 | -4,696732581 | 2,64E-06 | 1,62E-05 |
| ENSG00000204310.13 | AGPAT1 | protein_coding | 6292,652649 | -0,122410591 | 0,049310697 | -2,482434833 | 0,013048793 | 0,037626862 |
| ENSG00000204323.5 | SMIM5 | protein_coding | 388,834867 | -0,290786194 | 0,119615699 | -2,431003588 | 0,015057065 | 0,04255644 |
| ENSG00000204351.12 | SKIV2L | protein_coding | 4802,197414 | -0,159960674 | 0,061544537 | -2,599104343 | 0,009346736 | 0,028270045 |
| ENSG00000204386.10 | NEU1 | protein_coding | 13239,81209 | 0,503003659 | 0,044926323 | 11,1961902 | 4,26E-29 | 1,29E-27 |
| ENSG00000204387.13 | SNHG32 | protein_coding | 8551,563075 | -0,350233048 | 0,047569355 | -7,362577155 | 1,80E-13 | 2,42E-12 |
| ENSG00000204388.7 | HSPA1B | protein_coding | 10674,57644 | 0,418471946 | 0,045951788 | 9,106760877 | 8,49E-20 | 1,70E-18 |
| ENSG00000204389.9 | HSPA1A | protein_coding | 9406,926642 | 0,486808248 | 0,048370043 | 10,06425086 | 7,95E-24 | 1,96E-22 |
| ENSG00000204394.13 | VARS | protein_coding | 6111,306995 | -0,304518984 | 0,049785873 | -6,116574113 | 9,56E-10 | 8,94E-09 |
| ENSG00000204396.10 | VWA7 | protein_coding | 590,7582633 | 0,33777488 | 0,116261403 | 2,905305389 | 0,003668949 | 0,01241497 |
| ENSG00000204406.13 | MBD5 | protein_coding | 3377,083497 | 0,514889648 | 0,165639877 | 3,108488471 | 0,00188047 | 0,006840255 |
| ENSG00000204438.11 | GPANK1 | protein_coding | 2627,544911 | 0,174374307 | 0,060241849 | 2,894570965 | 0,003796771 | 0,012787897 |
| ENSG00000204439.3 | C6orf47 | protein_coding | 3926,100219 | 0,241336513 | 0,059585007 | 4,050289278 | 5,12E-05 | 0,000255931 |
| ENSG00000204442.3 | FAM155A | protein_coding | 522,3587136 | -0,41971162 | 0,121904331 | -3,44295905 | 0,000575386 | 0,00235608 |

| | | | | | | | | |
|---|---|---|---|---|---|---|---|---|
| ENSG00000204463.12 | BAG6 | protein_coding | 18216,83973 | -0,166411823 | 0,047479282 | -3,504935527 | 0,000456718 | 0,001913675 |
| ENSG00000204469.12 | PRRC2A | protein_coding | 27594,33933 | 0,198372659 | 0,039324198 | 5,04454435 | 4,55E-07 | 3,10E-06 |
| ENSG00000204472.13 | AIF1 | protein_coding | 153,2193502 | -0,551380539 | 0,189484196 | -2,909902521 | 0,003615415 | 0,012254085 |
| ENSG00000204498.11 | NFKBIL1 | protein_coding | 1672,365786 | 0,241670278 | 0,067492871 | 3,580678593 | 0,000342703 | 0,001472119 |
| ENSG00000204524.7 | ZNF805 | protein_coding | 1068,334717 | 0,210017127 | 0,086565802 | 2,426098084 | 0,015262145 | 0,043062072 |
| ENSG00000204525.16 | HLA-C | protein_coding | 62155,42076 | 6,041107761 | 0,068370911 | 88,35786561 | 0 | 0 |
| ENSG00000204568.12 | MRPS18B | protein_coding | 5772,258319 | 0,199218357 | 0,048994951 | 4,066099706 | 4,78E-05 | 0,000240142 |
| ENSG00000204576.11 | PRR3 | protein_coding | 2213,436482 | 0,283128613 | 0,072886438 | 3,884517102 | 0,000102533 | 0,000487213 |
| ENSG00000204580.13 | DDR1 | protein_coding | 39662,22268 | -0,301494682 | 0,054344095 | -5,547883025 | 2,89E-08 | 2,29E-07 |
| ENSG00000204581.3 | ACOXL-AS1 | lncRNA | 32,4364078 | -1,168130489 | 0,429764873 | -2,718068794 | 0,006566418 | 0,020764753 |
| ENSG00000204588.5 | LINC01123 | lncRNA | 515,1916209 | 0,382120162 | 0,11705063 | 3,264571584 | 0,001096298 | 0,004211017 |
| ENSG00000204590.12 | GNL1 | protein_coding | 12610,56755 | 0,174765243 | 0,043186947 | 4,046714474 | 5,19E-05 | 0,000259696 |
| ENSG00000204592.9 | HLA-E | protein_coding | 74243,03524 | 3,25786196 | 0,055160733 | 59,06125196 | 0 | 0 |
| ENSG00000204599.14 | TRIM39 | protein_coding | 2363,262118 | 0,257761093 | 0,061845325 | 4,167834741 | 3,08E-05 | 0,000159339 |
| ENSG00000204604.11 | ZNF468 | protein_coding | 5906,367054 | 0,299607261 | 0,060513686 | 4,951066145 | 7,38E-07 | 4,89E-06 |
| ENSG00000204622.11 | HLA-J | transcribed_unproc | 378,2194871 | 3,725687795 | 0,169282476 | 22,00870339 | 2,38E-107 | 2,89E-105 |
| ENSG00000204628.11 | RACK1 | protein_coding | 43466,1677 | -0,442563882 | 0,040556788 | -10,91220255 | 1,01E-27 | 2,90E-26 |
| ENSG00000204642.14 | HLA-F | protein_coding | 7482,411459 | 9,198997101 | 0,136965626 | 67,16281596 | 0 | 0 |
| ENSG00000204653.10 | ASPDH | protein_coding | 102,0179636 | -0,804897167 | 0,237895259 | -3,383409866 | 0,000715917 | 0,002869405 |
| ENSG00000204673.10 | AKT1S1 | protein_coding | 5307,694131 | -0,483415778 | 0,053788442 | -8,987354117 | 2,53E-19 | 4,95E-18 |
| ENSG00000204682.7 | MIR1915HG | protein_coding | 800,9034787 | -0,409820215 | 0,100350649 | -4,083882083 | 4,43E-05 | 0,000223975 |
| ENSG00000204706.14 | MAMDC2-AS1 | lncRNA | 50,21753077 | 0,941373424 | 0,382289727 | 2,462460687 | 0,01379873 | 0,039546169 |
| ENSG00000204740.11 | MALRD1 | protein_coding | 121,2265966 | 0,681201103 | 0,243876113 | 2,793226011 | 0,005218522 | 0,016940252 |
| ENSG00000204789.4 | ZNF204P | processed_pseudog | 3026,985756 | 0,616520811 | 0,058857681 | 10,47477229 | 1,13E-25 | 3,02E-24 |
| ENSG00000204815.10 | TTC25 | protein_coding | 937,2489348 | 0,482289601 | 0,097411732 | 4,951042245 | 7,38E-07 | 4,89E-06 |
| ENSG00000204822.7 | MRPL53 | protein_coding | 2708,865347 | -0,179525333 | 0,064338432 | -2,79032806 | 0,005265466 | 0,017075403 |
| ENSG00000204851.6 | PNMA8B | protein_coding | 171,5954253 | 0,479273996 | 0,175111228 | 2,73696897 | 0,006200814 | 0,01973616 |
| ENSG00000204852.15 | TCTN1 | protein_coding | 3345,40472 | -0,173549348 | 0,069217701 | -2,507297196 | 0,012165837 | 0,035411979 |
| ENSG00000204856.12 | FAM216A | protein_coding | 648,617171 | -0,458577211 | 0,107537993 | -4,264327399 | 2,01E-05 | 0,000107023 |
| ENSG00000204859.13 | ZBTB48 | protein_coding | 1788,093002 | -0,19068783 | 0,074370778 | -2,564015534 | 0,010346891 | 0,030810713 |
| ENSG00000204899.6 | MZT1 | protein_coding | 5979,367541 | -0,425353514 | 0,051755532 | -8,218513082 | 2,06E-16 | 3,40E-15 |
| ENSG00000204934.10 | ATP6V0E2-AS1 | lncRNA | 872,8878562 | 0,370174025 | 0,097563972 | 3,794167232 | 0,00014814 | 0,000683435 |
| ENSG00000204946.10 | ZNF783 | protein_coding | 1193,203884 | 0,4733283 | 0,090431038 | 5,234135421 | 1,66E-07 | 1,19E-06 |
| ENSG00000204954.10 | C12orf73 | protein_coding | 1669,589212 | -0,230950189 | 0,073337162 | -3,149156348 | 0,001637426 | 0,006050035 |
| ENSG00000204959.4 | ARHGEF34P | unprocessed_pseud | 4212,568954 | 0,426014566 | 0,051645226 | 8,248866281 | 1,60E-16 | 2,66E-15 |
| ENSG00000204962.6 | PCDHA8 | protein_coding | 1322,208966 | -0,552583098 | 0,075784234 | -7,291531058 | 3,06E-13 | 4,04E-12 |

| | | | | | | | | |
|---|---|---|---|---|---|---|---|---|
| ENSG00000204991.11 | SPIRE2 | protein_coding | 3563,50445 | -0,162884816 | 0,056802983 | -2,867539833 | 0,004136767 | 0,013813157 |
| ENSG00000205002.3 | AARD | protein_coding | 807,4355984 | -0,275431025 | 0,087554921 | -3,145808626 | 0,001656283 | 0,006115696 |
| ENSG00000205022.9 | PABPN1L | protein_coding | 162,3951438 | -1,123989825 | 0,199749293 | -5,62700277 | 1,83E-08 | 1,48E-07 |
| ENSG00000205060.11 | SLC35B4 | protein_coding | 3000,974253 | 0,386355221 | 0,060847703 | 6,349544895 | 2,16E-10 | 2,18E-09 |
| ENSG00000205089.7 | CCNI2 | protein_coding | 502,8689472 | -0,488785038 | 0,156290405 | -3,127415518 | 0,001763505 | 0,006467136 |
| ENSG00000205133.11 | TRIQK | protein_coding | 8011,15762 | -0,466869103 | 0,057789559 | -8,078779545 | 6,54E-16 | 1,04E-14 |
| ENSG00000205155.8 | PSENEN | protein_coding | 4024,340562 | -0,131335648 | 0,053644915 | -2,448240369 | 0,014355585 | 0,040902514 |
| ENSG00000205213.14 | LGR4 | protein_coding | 9658,8501 | 0,260660568 | 0,047784651 | 5,454901538 | 4,90E-08 | 3,76E-07 |
| ENSG00000205220.12 | PSMB10 | protein_coding | 1190,706759 | 2,450714558 | 0,098978766 | 24,76000325 | 2,42E-135 | 3,68E-133 |
| ENSG00000205269.6 | TMEM170B | protein_coding | 5749,233325 | -0,29768729 | 0,056275063 | -5,289861514 | 1,22E-07 | 8,95E-07 |
| ENSG00000205279.8 | CTXN3 | protein_coding | 13,02186568 | 2,402284338 | 0,718518759 | 3,343384298 | 0,000827632 | 0,003267784 |
| ENSG00000205293.5 | LINC01602 | lncRNA | 34,5109702 | -2,004208556 | 0,548421759 | -3,654502254 | 0,000257681 | 0,001132621 |
| ENSG00000205302.7 | SNX2 | protein_coding | 18242,71058 | 0,229490125 | 0,049289168 | 4,6559951 | 3,22E-06 | 1,94E-05 |
| ENSG00000205336.11 | ADGRG1 | protein_coding | 3232,989746 | -0,536716941 | 0,08527908 | -6,293653042 | 3,10E-10 | 3,08E-09 |
| ENSG00000205363.5 | INSYN1 | protein_coding | 666,3903968 | -1,089790395 | 0,117922163 | -9,241607901 | 2,43E-20 | 5,04E-19 |
| ENSG00000205413.8 | SAMD9 | protein_coding | 44310,09234 | 4,311072882 | 0,041773107 | 103,2021123 | 0 | 0 |
| ENSG00000205436.7 | EXOC3L4 | protein_coding | 257,1005444 | 1,269608067 | 0,17347903 | 7,31851031 | 2,51E-13 | 3,33E-12 |
| ENSG00000205502.4 | C2CD4B | protein_coding | 48,75853884 | 1,151591451 | 0,416768191 | 2,763146218 | 0,005724711 | 0,018397706 |
| ENSG00000205517.12 | RGL3 | protein_coding | 4763,345646 | -0,214491237 | 0,070424439 | -3,045693214 | 0,002321446 | 0,008245766 |
| ENSG00000205531.13 | NAP1L4 | protein_coding | 18468,95441 | -0,10912454 | 0,042156499 | -2,588557954 | 0,009637873 | 0,029018041 |
| ENSG00000205544.4 | TMEM256 | protein_coding | 1509,52171 | -0,428905725 | 0,078465899 | -5,466141753 | 4,60E-08 | 3,54E-07 |
| ENSG00000205560.12 | CPT1B | protein_coding | 3255,603259 | 0,316930446 | 0,079276406 | 3,997790266 | 6,39E-05 | 0,00031505 |
| ENSG00000205571.13 | SMN2 | protein_coding | 2623,17007 | -0,170585908 | 0,06322785 | -2,697955231 | 0,006976682 | 0,021908223 |
| ENSG00000205572.9 | SERF1B | protein_coding | 5065,992317 | -0,209254357 | 0,050402128 | -4,151696909 | 3,30E-05 | 0,000170187 |
| ENSG00000205581.11 | HMGN1 | protein_coding | 43159,0469 | -0,29409458 | 0,043097784 | -6,823891053 | 8,86E-12 | 1,02E-10 |
| ENSG00000205611.5 | LINC01597 | lncRNA | 28,30992355 | 1,241110888 | 0,464036412 | 2,67459806 | 0,007481889 | 0,023248184 |
| ENSG00000205730.6 | ITPRIPL2 | protein_coding | 754,6480536 | -0,267817301 | 0,106785303 | -2,507997766 | 0,012141743 | 0,035346418 |
| ENSG00000205746.9 | AC126755.1 | transcribed_unproc | 2988,233407 | 0,210021429 | 0,062947629 | 3,336447021 | 0,000848566 | 0,003344488 |
| ENSG00000205771.6 | CATSPER2P1 | transcribed_unproc | 179,1110133 | -0,540175957 | 0,184951952 | -2,920628578 | 0,00349326 | 0,011891983 |
| ENSG00000205795.4 | CYS1 | protein_coding | 1410,144803 | -0,85530788 | 0,089969083 | -9,506686688 | 1,97E-21 | 4,29E-20 |
| ENSG00000205885.7 | C1RL-AS1 | lncRNA | 659,5824373 | 0,801789969 | 0,27304389 | 2,936487496 | 0,003319522 | 0,011350315 |
| ENSG00000205903.7 | ZNF316 | protein_coding | 3009,757429 | -0,287187989 | 0,081377115 | -3,52910014 | 0,000416975 | 0,001758939 |
| ENSG00000205929.11 | C21orf62 | protein_coding | 308,9336047 | -0,84672577 | 0,16437502 | -5,151182768 | 2,59E-07 | 1,82E-06 |
| ENSG00000205978.6 | NYNRIN | protein_coding | 1166,790665 | -0,322247185 | 0,101923431 | -3,1616595 | 0,001568729 | 0,005825857 |
| ENSG00000206052.11 | DOK6 | protein_coding | 428,925889 | -0,600915355 | 0,149302976 | -4,02480495 | 5,70E-05 | 0,00028327 |
| ENSG00000206053.13 | JPT2 | protein_coding | 29552,21534 | -0,375979659 | 0,054842032 | -6,855684349 | 7,10E-12 | 8,27E-11 |

| | | | | | | | | |
|---|---|---|---|---|---|---|---|---|
| ENSG00000206077.11 | ZDHHC11B | protein_coding | 280,882917 | 0,702522621 | 0,150674912 | 4,6625056 | 3,12E-06 | 1,89E-05 |
| ENSG00000206120.11 | EGFEM1P | transcribed_unitary | 309,7205291 | 0,778135516 | 0,141767457 | 5,488816187 | 4,05E-08 | 3,13E-07 |
| ENSG00000206149.10 | HERC2P9 | transcribed_unproc | 6287,14995 | 0,33613582 | 0,062283007 | 5,396910577 | 6,78E-08 | 5,10E-07 |
| ENSG00000206337.11 | HCP5 | lncRNA | 4803,440908 | 8,187275979 | 0,360624961 | 22,70302074 | 4,18E-114 | 5,44E-112 |
| ENSG00000206341.7 | HLA-H | unprocessed_pseud | 2206,286694 | 5,264878875 | 0,115773044 | 45,4758613 | 0 | 0 |
| ENSG00000206417.8 | H1FX-AS1 | lncRNA | 322,7761227 | -0,884379605 | 0,136143309 | -6,495946154 | 8,25E-11 | 8,65E-10 |
| ENSG00000206418.4 | RAB12 | protein_coding | 5120,477492 | 0,217373834 | 0,06074177 | 3,578654948 | 0,000345367 | 0,001482148 |
| ENSG00000206503.13 | HLA-A | protein_coding | 106785,2296 | 3,470242112 | 0,156881658 | 22,1201265 | 2,02E-108 | 2,49E-106 |
| ENSG00000206527.10 | HACD2 | protein_coding | 14526,39294 | -0,139450377 | 0,047435482 | -2,939790463 | 0,003284343 | 0,011245402 |
| ENSG00000206530.11 | CFAP44 | protein_coding | 716,7263297 | 0,355524121 | 0,14253336 | 2,494322189 | 0,012619794 | 0,036558422 |
| ENSG00000206560.11 | ANKRD28 | protein_coding | 6192,947027 | 0,195534521 | 0,052421859 | 3,730018813 | 0,000191465 | 0,000861939 |
| ENSG00000206561.13 | COLQ | protein_coding | 181,6687793 | 0,514876709 | 0,199931782 | 2,575261942 | 0,01001642 | 0,02996539 |
| ENSG00000206562.11 | METTL6 | protein_coding | 3162,380728 | 0,153632421 | 0,06304563 | 2,43684488 | 0,014816034 | 0,04197006 |
| ENSG00000206828.1 | RF00003 | snRNA | 24,53319068 | -1,843631438 | 0,521747691 | -3,533568945 | 0,000409989 | 0,001731091 |
| ENSG00000206965.1 | RNU6-5P | snRNA | 421,3851361 | -0,92889369 | 0,164053833 | -5,662127315 | 1,50E-08 | 1,22E-07 |
| ENSG00000210049.1 | MT-TF | Mt_tRNA | 196,9257576 | 3,975152452 | 0,250555802 | 15,86533789 | 1,10E-56 | 7,23E-55 |
| ENSG00000210196.2 | MT-TP | Mt_tRNA | 8,974210299 | -2,564187773 | 0,929271001 | -2,759354128 | 0,005791574 | 0,018572807 |
| ENSG00000211451.12 | GNRHR2 | transcribed_unitary | 576,0960413 | -0,30122286 | 0,114590326 | -2,628693635 | 0,008571354 | 0,026217232 |
| ENSG00000211459.2 | MT-RNR1 | Mt_rRNA | 86308,31669 | 0,912394028 | 0,290491471 | 3,140863402 | 0,001684506 | 0,006203623 |
| ENSG00000211460.12 | TSN | protein_coding | 17419,32858 | -0,428173993 | 0,050131562 | -8,541006389 | 1,33E-17 | 2,36E-16 |
| ENSG00000212232.1 | SNORD17 | snoRNA | 92,35067661 | -1,018143815 | 0,239580777 | -4,249689086 | 2,14E-05 | 0,000113804 |
| ENSG00000212747.4 | RTL8B | protein_coding | 2337,866177 | -0,194438552 | 0,061013853 | -3,186793522 | 0,001438594 | 0,005383428 |
| ENSG00000212907.2 | MT-ND4L | protein_coding | 108728,0562 | -0,823387604 | 0,047847678 | -17,20851739 | 2,29E-66 | 1,75E-64 |
| ENSG00000212916.4 | MAP10 | protein_coding | 512,4927657 | 0,369373762 | 0,107312429 | 3,44204082 | 0,000577343 | 0,002363233 |
| ENSG00000212978.6 | AC016747.1 | lncRNA | 2275,469132 | -0,194570757 | 0,071973083 | -2,70338227 | 0,006863776 | 0,021595853 |
| ENSG00000213047.13 | DENND1B | protein_coding | 1435,927949 | 0,223297764 | 0,078673923 | 2,838269091 | 0,004535893 | 0,014961665 |
| ENSG00000213064.10 | SFT2D2 | protein_coding | 9230,823111 | -0,33528151 | 0,062705936 | -5,346886316 | 8,95E-08 | 6,63E-07 |
| ENSG00000213079.9 | SCAF8 | protein_coding | 10725,60779 | 0,149886584 | 0,047513062 | 3,154639572 | 0,001606965 | 0,005948324 |
| ENSG00000213160.10 | KLHL23 | protein_coding | 12170,05122 | -0,267348785 | 0,049064971 | -5,448872739 | 5,07E-08 | 3,88E-07 |
| ENSG00000213186.8 | TRIM59 | protein_coding | 3918,829873 | -0,20031468 | 0,064141927 | -3,122991299 | 0,00179023 | 0,006554375 |
| ENSG00000213190.3 | MLLT11 | protein_coding | 8043,228905 | -0,18120264 | 0,048510686 | -3,735313873 | 0,000187481 | 0,000845186 |
| ENSG00000213214.4 | ARHGEF35 | protein_coding | 5051,160441 | 0,315216516 | 0,050687319 | 6,218843766 | 5,01E-10 | 4,87E-09 |
| ENSG00000213281.5 | NRAS | protein_coding | 20296,09203 | -0,407488753 | 0,044431102 | -9,17125014 | 4,68E-20 | 9,52E-19 |
| ENSG00000213297.8 | ZNF625-ZNF20 | protein_coding | 575,3006827 | 0,365079165 | 0,107432224 | 3,398227764 | 0,000678239 | 0,002731524 |
| ENSG00000213339.9 | QTRT1 | protein_coding | 2350,279813 | -0,639556306 | 0,076469917 | -8,363502056 | 6,09E-17 | 1,03E-15 |
| ENSG00000213341.11 | CHUK | protein_coding | 12529,42462 | -0,258526105 | 0,04311522 | -5,996168119 | 2,02E-09 | 1,82E-08 |

| | | | | | | | | |
|---|---|---|---|---|---|---|---|---|
| ENSG00000213347.10 | MXD3 | protein_coding | 1416,721758 | -0,420582013 | 0,07789984 | -5,399009966 | 6,70E-08 | 5,05E-07 |
| ENSG00000213366.13 | GSTM2 | protein_coding | 4795,319992 | -0,278288559 | 0,051024699 | -5,453997059 | 4,93E-08 | 3,78E-07 |
| ENSG00000213380.15 | COG8 | protein_coding | 2780,543579 | -0,200638058 | 0,071822097 | -2,793542186 | 0,005213423 | 0,016935672 |
| ENSG00000213390.11 | ARHGAP19 | protein_coding | 2192,16274 | -0,842211755 | 0,097447667 | -8,642708229 | 5,49E-18 | 1,00E-16 |
| ENSG00000213412.3 | HNRNPA1P33 | processed_pseudog | 27,07327159 | -1,402336005 | 0,471429016 | -2,974649329 | 0,002933237 | 0,01015601 |
| ENSG00000213420.8 | GPC2 | protein_coding | 1103,011742 | -0,260334344 | 0,092748911 | -2,806872244 | 0,005002508 | 0,016330884 |
| ENSG00000213462.5 | ERV3-1 | protein_coding | 683,3925758 | -0,388155222 | 0,127612012 | -3,04168248 | 0,002352599 | 0,008339316 |
| ENSG00000213465.8 | ARL2 | protein_coding | 5352,027346 | -0,331786201 | 0,053749679 | -6,172803386 | 6,71E-10 | 6,40E-09 |
| ENSG00000213516.10 | RBMXL1 | protein_coding | 3511,522545 | 0,4781951 | 0,061195414 | 7,814231039 | 5,53E-15 | 8,33E-14 |
| ENSG00000213533.12 | STIMATE | protein_coding | 1267,184016 | 0,286209932 | 0,074854322 | 3,823559197 | 0,000131539 | 0,00061237 |
| ENSG00000213542.3 | AC007000.1 | processed_pseudog | 11,60320701 | -2,009541423 | 0,711100093 | -2,82596141 | 0,004713894 | 0,015479759 |
| ENSG00000213551.5 | DNAJC9 | protein_coding | 9508,930684 | -0,144815808 | 0,044732643 | -3,237363127 | 0,001206398 | 0,004586961 |
| ENSG00000213585.11 | VDAC1 | protein_coding | 19502,46154 | -0,285178843 | 0,042721837 | -6,675247629 | 2,47E-11 | 2,72E-10 |
| ENSG00000213614.9 | HEXA | protein_coding | 14027,34262 | 0,15078279 | 0,040579251 | 3,715760772 | 0,000202593 | 0,000907136 |
| ENSG00000213619.10 | NDUFS3 | protein_coding | 6148,821325 | -0,154002988 | 0,046238963 | -3,33058912 | 0,000866624 | 0,00340493 |
| ENSG00000213626.13 | LBH | protein_coding | 5424,44092 | 0,6703585 | 0,052871315 | 12,67905847 | 7,73E-37 | 3,16E-35 |
| ENSG00000213639.10 | PPP1CB | protein_coding | 35947,09001 | -0,149741452 | 0,040974482 | -3,654505092 | 0,000257679 | 0,001132621 |
| ENSG00000213676.13 | ATF6B | protein_coding | 6387,515243 | -0,116491486 | 0,047861504 | -2,433928665 | 0,014935938 | 0,042261566 |
| ENSG00000213689.14 | TREX1 | protein_coding | 1541,234388 | 3,246951148 | 0,118261635 | 27,4556593 | 5,95E-166 | 1,05E-163 |
| ENSG00000213741.10 | RPS29 | protein_coding | 20514,65431 | -0,414391827 | 0,044585089 | -9,294404025 | 1,48E-20 | 3,11E-19 |
| ENSG00000213742.7 | ZNF337-AS1 | lncRNA | 417,5290359 | -0,392629988 | 0,134798087 | -2,912726695 | 0,00358288 | 0,012151131 |
| ENSG00000213753.11 | CENPBD1P1 | transcribed_process | 3861,285496 | -0,21576057 | 0,058635088 | -3,679717649 | 0,000233492 | 0,001035194 |
| ENSG00000213799.12 | ZNF845 | protein_coding | 3231,029012 | 0,333743638 | 0,062187404 | 5,366740124 | 8,02E-08 | 5,96E-07 |
| ENSG00000213853.10 | EMP2 | protein_coding | 17106,98689 | -0,887449755 | 0,051414177 | -17,26079865 | 9,28E-67 | 7,13E-65 |
| ENSG00000213889.10 | PPM1N | protein_coding | 229,7930135 | -0,554395791 | 0,179893915 | -3,081792902 | 0,002057579 | 0,007407878 |
| ENSG00000213918.10 | DNASE1 | protein_coding | 3851,034635 | 0,393124979 | 0,052676157 | 7,463053499 | 8,45E-14 | 1,17E-12 |
| ENSG00000213923.12 | CSNK1E | protein_coding | 5725,187001 | -0,162323826 | 0,052880771 | -3,069619118 | 0,002143319 | 0,007676121 |
| ENSG00000213928.8 | IRF9 | protein_coding | 8144,003378 | 4,716027103 | 0,06165498 | 76,49061127 | 0 | 0 |
| ENSG00000213930.11 | GALT | protein_coding | 2217,83842 | 0,24901305 | 0,067021821 | 3,715402616 | 0,00020288 | 0,00090806 |
| ENSG00000213949.9 | ITGA1 | protein_coding | 13053,11618 | 0,500105058 | 0,058812093 | 8,503439181 | 1,84E-17 | 3,23E-16 |
| ENSG00000213977.8 | TAX1BP3 | protein_coding | 2795,795193 | -0,274001707 | 0,058995434 | -4,644456125 | 3,41E-06 | 2,05E-05 |
| ENSG00000213995.11 | NAXD | protein_coding | 7763,547602 | 0,120552756 | 0,05032058 | 2,395694898 | 0,016588892 | 0,046277251 |
| ENSG00000213997.3 | PGAM1P7 | processed_pseudog | 94,98195523 | 1,034655757 | 0,238177238 | 4,34405809 | 1,40E-05 | 7,65E-05 |
| ENSG00000214013.9 | GANC | protein_coding | 1077,494864 | 0,679335008 | 0,092211365 | 7,367150565 | 1,74E-13 | 2,34E-12 |
| ENSG00000214022.11 | REPIN1 | protein_coding | 9750,577629 | 0,209500346 | 0,046696114 | 4,486462155 | 7,24E-06 | 4,14E-05 |
| ENSG00000214026.11 | MRPL23 | protein_coding | 2998,767229 | -0,208867495 | 0,058878485 | -3,547433221 | 0,000389004 | 0,001650848 |

| | | | | | | | | |
|---|---|---|---|---|---|---|---|---|
| ENSG00000214039.9 | LINC02418 | lncRNA | 2033,979728 | -0,267158879 | 0,070586183 | -3,784860813 | 0,000153795 | 0,000706774 |
| ENSG00000214046.8 | SMIM7 | protein_coding | 7628,470731 | -0,117214101 | 0,048366133 | -2,423474746 | 0,015372822 | 0,043310981 |
| ENSG00000214050.8 | FBXO16 | protein_coding | 701,3812321 | 0,572763544 | 0,101695992 | 5,632115217 | 1,78E-08 | 1,44E-07 |
| ENSG00000214063.11 | TSPAN4 | protein_coding | 731,3609777 | -0,46758969 | 0,099533178 | -4,6978274 | 2,63E-06 | 1,61E-05 |
| ENSG00000214087.8 | ARL16 | protein_coding | 3108,93435 | -0,17274391 | 0,054808179 | -3,151790741 | 0,001622725 | 0,006000644 |
| ENSG00000214106.9 | PAXIP1-AS2 | lncRNA | 2075,826993 | 2,239715179 | 0,073339647 | 30,53894131 | 7,93E-205 | 1,61E-202 |
| ENSG00000214113.10 | LYRM4 | protein_coding | 1212,211189 | -0,347951788 | 0,085722839 | -4,059032497 | 4,93E-05 | 0,000246975 |
| ENSG00000214114.9 | MYCBP | protein_coding | 3817,676951 | -0,403247416 | 0,057304256 | -7,036954028 | 1,96E-12 | 2,42E-11 |
| ENSG00000214135.8 | AC132008.2 | transcribed_unproc | 2511,43727 | -0,147932026 | 0,060747488 | -2,435195757 | 0,014883735 | 0,042135328 |
| ENSG00000214194.9 | SMIM30 | protein_coding | 11699,63514 | -0,122900782 | 0,048612816 | -2,528155959 | 0,01146634 | 0,033597482 |
| ENSG00000214226.9 | C17orf67 | protein_coding | 191,0012678 | 1,116550353 | 0,180165861 | 6,197346974 | 5,74E-10 | 5,53E-09 |
| ENSG00000214253.8 | FIS1 | protein_coding | 4296,518646 | -0,199103091 | 0,057537316 | -3,460416747 | 0,00053934 | 0,002219785 |
| ENSG00000214293.9 | APTR | lncRNA | 1801,418382 | 0,175888818 | 0,073818507 | 2,382719777 | 0,01718527 | 0,047714722 |
| ENSG00000214338.10 | SOGA3 | protein_coding | 2513,989307 | 0,322820233 | 0,069848372 | 4,621728783 | 3,81E-06 | 2,27E-05 |
| ENSG00000214357.8 | NEURL1B | protein_coding | 11214,27241 | -0,188927454 | 0,056462678 | -3,346059008 | 0,000819689 | 0,003240322 |
| ENSG00000214376.5 | VSTM5 | protein_coding | 49,59519736 | 0,981557607 | 0,412512833 | 2,379459567 | 0,017338045 | 0,048061829 |
| ENSG00000214401.4 | KANSL1-AS1 | lncRNA | 62,58218807 | -0,699733487 | 0,295210165 | -2,370289269 | 0,017774173 | 0,049057212 |
| ENSG00000214425.8 | LRRC37A4P | transcribed_unproc | 3335,597781 | 0,459493866 | 0,056305444 | 8,160736045 | 3,33E-16 | 5,41E-15 |
| ENSG00000214435.8 | AS3MT | protein_coding | 1604,899709 | -0,360426995 | 0,08340277 | -4,321523061 | 1,55E-05 | 8,41E-05 |
| ENSG00000214455.4 | RCN1P2 | processed_pseudog | 514,0950695 | 0,537006899 | 0,106987558 | 5,019339694 | 5,18E-07 | 3,51E-06 |
| ENSG00000214530.9 | STARD10 | protein_coding | 14504,21032 | -0,209287713 | 0,043344241 | -4,828501023 | 1,38E-06 | 8,75E-06 |
| ENSG00000214562.14 | NUTM2D | protein_coding | 930,3468297 | 0,57683655 | 0,089410698 | 6,45153839 | 1,11E-10 | 1,15E-09 |
| ENSG00000214595.11 | EML6 | protein_coding | 3504,099177 | -0,202081812 | 0,063041873 | -3,205517267 | 0,0013482 | 0,005077233 |
| ENSG00000214706.10 | IFRD2 | protein_coding | 2174,513016 | -0,371425038 | 0,065465741 | -5,673578794 | 1,40E-08 | 1,15E-07 |
| ENSG00000214719.13 | AC005562.1 | lncRNA | 1267,48511 | 0,233216511 | 0,083111835 | 2,806056597 | 0,005015188 | 0,016355676 |
| ENSG00000214736.7 | TOMM6 | protein_coding | 6241,911429 | -0,392921414 | 0,052298837 | -7,513004755 | 5,78E-14 | 8,08E-13 |
| ENSG00000214753.3 | HNRNPUL2 | protein_coding | 30966,71299 | -0,281206126 | 0,041199323 | -6,82550355 | 8,76E-12 | 1,01E-10 |
| ENSG00000214796.8 | AC098934.1 | transcribed_unproc | 428,6981985 | -0,808127057 | 0,118974695 | -6,79242805 | 1,10E-11 | 1,26E-10 |
| ENSG00000214837.8 | LINC01347 | transcribed_unproc | 127,8338554 | 1,054290929 | 0,275519439 | 3,826557331 | 0,000129948 | 0,000606217 |
| ENSG00000214900.10 | LINC01588 | lncRNA | 75,76132821 | 0,991506795 | 0,304704661 | 3,253992867 | 0,001137951 | 0,004353926 |
| ENSG00000214941.8 | ZSWIM7 | protein_coding | 1094,593745 | -0,388537734 | 0,085954249 | -4,520285362 | 6,18E-06 | 3,57E-05 |
| ENSG00000214944.9 | ARHGEF28 | protein_coding | 5626,330033 | 0,386473832 | 0,059640697 | 6,48003549 | 9,17E-11 | 9,59E-10 |
| ENSG00000215012.9 | RTL10 | protein_coding | 6437,806848 | -0,270507055 | 0,052350521 | -5,167227534 | 2,38E-07 | 1,68E-06 |
| ENSG00000215021.8 | PHB2 | protein_coding | 8317,410284 | -0,319380518 | 0,049595902 | -6,439655374 | 1,20E-10 | 1,24E-09 |
| ENSG00000215029.9 | TCP11X2 | protein_coding | 255,7238222 | -0,596502392 | 0,165358951 | -3,607318436 | 0,000309378 | 0,00134111 |
| ENSG00000215030.5 | RPL13P12 | processed_pseudog | 194,2704737 | -0,871231449 | 0,217616372 | -4,003519768 | 6,24E-05 | 0,00030812 |

| | | | | | | | | |
|---|---|---|---|---|---|---|---|---|
| ENSG00000215039.7 | CD27-AS1 | lncRNA | 2123,872622 | 0,295653539 | 0,073573188 | 4,018495689 | 5,86E-05 | 0,000290135 |
| ENSG00000215041.10 | NEURL4 | protein_coding | 4559,434233 | -0,188227959 | 0,06523592 | -2,885342275 | 0,003909883 | 0,013131537 |
| ENSG00000215067.10 | ALOX12-AS1 | lncRNA | 370,1413254 | -0,780873433 | 0,136896385 | -5,704120184 | 1,17E-08 | 9,65E-08 |
| ENSG00000215068.8 | AC025171.2 | lncRNA | 530,0845024 | 1,779861729 | 0,119928365 | 14,84104055 | 7,95E-50 | 4,55E-48 |
| ENSG00000215105.4 | TTC3P1 | processed_pseudog | 1107,107828 | 0,372225146 | 0,08548019 | 4,354519395 | 1,33E-05 | 7,33E-05 |
| ENSG00000215114.9 | UBXN2B | protein_coding | 3913,501043 | 0,322763276 | 0,052029037 | 6,203521962 | 5,52E-10 | 5,34E-09 |
| ENSG00000215126.10 | CBWD6 | protein_coding | 1248,624038 | 0,578443505 | 0,090489624 | 6,392373865 | 1,63E-10 | 1,67E-09 |
| ENSG00000215154.6 | AC141586.1 | transcribed_unproc | 887,616728 | 0,426231623 | 0,110373209 | 3,861730824 | 0,000112587 | 0,000530614 |
| ENSG00000215193.12 | PEX26 | protein_coding | 6923,815634 | -0,218975817 | 0,062094638 | -3,526485149 | 0,000421115 | 0,001774405 |
| ENSG00000215217.7 | C5orf49 | protein_coding | 568,0990559 | -0,374123305 | 0,102826209 | -3,638404158 | 0,000274333 | 0,001199253 |
| ENSG00000215218.4 | UBE2QL1 | protein_coding | 2497,098367 | 0,824178577 | 0,078743059 | 10,46668226 | 1,23E-25 | 3,27E-24 |
| ENSG00000215256.4 | DHRS4-AS1 | lncRNA | 2954,081591 | -0,241568359 | 0,065580253 | -3,683553336 | 0,000230005 | 0,001021141 |
| ENSG00000215301.10 | DDX3X | protein_coding | 73261,08852 | -0,177058267 | 0,042683426 | -4,148173694 | 3,35E-05 | 0,000172583 |
| ENSG00000215421.9 | ZNF407 | protein_coding | 4510,020147 | -0,17673214 | 0,052152066 | -3,388785041 | 0,00070203 | 0,002818765 |
| ENSG00000215424.10 | MCM3AP-AS1 | lncRNA | 1346,719487 | -0,220754606 | 0,085687275 | -2,576282241 | 0,009986909 | 0,029885048 |
| ENSG00000215440.12 | NPEPL1 | protein_coding | 2126,883202 | -0,169160624 | 0,071177226 | -2,376611647 | 0,017472473 | 0,048387336 |
| ENSG00000215472.10 | RPL17-C18orf32 | protein_coding | 532,1235151 | -0,548963756 | 0,114358328 | -4,800382855 | 1,58E-06 | 1,00E-05 |
| ENSG00000215498.9 | FAM230B | lncRNA | 146,8517379 | 0,767947718 | 0,264687638 | 2,901335789 | 0,003715755 | 0,012545061 |
| ENSG00000215784.6 | FAM72D | protein_coding | 1699,503266 | -1,323160925 | 0,075714383 | -17,47568776 | 2,19E-68 | 1,75E-66 |
| ENSG00000215788.10 | TNFRSF25 | protein_coding | 200,749866 | -0,45695219 | 0,190166464 | -2,402906279 | 0,016265355 | 0,045491661 |
| ENSG00000215845.11 | TSTD1 | protein_coding | 1274,175617 | -0,326035736 | 0,073781153 | -4,418956906 | 9,92E-06 | 5,57E-05 |
| ENSG00000215861.6 | AC245297.1 | transcribed_unproc | 2080,431563 | -0,677298072 | 0,079038797 | -8,569184946 | 1,04E-17 | 1,86E-16 |
| ENSG00000215883.10 | CYB5RL | protein_coding | 2937,309102 | -0,366898578 | 0,058762058 | -6,243800671 | 4,27E-10 | 4,18E-09 |
| ENSG00000215908.10 | CROCCP2 | transcribed_unproc | 1865,141362 | 0,200630994 | 0,072568532 | 2,764710667 | 0,005697331 | 0,018320173 |
| ENSG00000216490.4 | IFI30 | protein_coding | 8528,842491 | 1,970374291 | 0,174212528 | 11,31017565 | 1,17E-29 | 3,60E-28 |
| ENSG00000216895.9 | AC009403.1 | lncRNA | 893,6786032 | -0,387608569 | 0,095918071 | -4,041037997 | 5,32E-05 | 0,000265592 |
| ENSG00000217128.12 | FNIP1 | protein_coding | 6475,09233 | 0,287497878 | 0,056561972 | 5,082882902 | 3,72E-07 | 2,56E-06 |
| ENSG00000217442.4 | SYCE3 | protein_coding | 94,35119419 | 0,83401326 | 0,253747379 | 3,286785716 | 0,001013379 | 0,00392264 |
| ENSG00000217555.12 | CKLF | protein_coding | 3240,49475 | -0,442432867 | 0,065802499 | -6,723648406 | 1,77E-11 | 1,98E-10 |
| ENSG00000217929.4 | CICP18 | transcribed_process | 476,2835009 | 0,688113693 | 0,117115865 | 5,875495098 | 4,22E-09 | 3,67E-08 |
| ENSG00000218052.5 | ADAMTS7P4 | transcribed_unproc | 57,63675125 | 3,731923853 | 0,392498644 | 9,508119101 | 1,94E-21 | 4,24E-20 |
| ENSG00000218336.9 | TENM3 | protein_coding | 54555,58963 | -0,256347472 | 0,05508188 | -4,653934658 | 3,26E-06 | 1,96E-05 |
| ENSG00000218739.10 | CEBPZOS | protein_coding | 6634,121768 | -0,273320376 | 0,051648028 | -5,291980853 | 1,21E-07 | 8,86E-07 |
| ENSG00000219200.11 | RNASEK | protein_coding | 9726,105989 | 0,155318456 | 0,049284791 | 3,151447971 | 0,001624631 | 0,006006705 |
| ENSG00000219438.8 | TAFA5 | protein_coding | 2713,598618 | -0,626852947 | 0,062713955 | -9,995430007 | 1,60E-23 | 3,87E-22 |
| ENSG00000219481.10 | NBPF1 | protein_coding | 11388,22453 | -0,120510865 | 0,044349141 | -2,717321299 | 0,006581268 | 0,020797104 |

| | | | | | | | | |
|---|---|---|---|---|---|---|---|---|
| ENSG00000220323.4 | HIST2H2BD | transcribed_unproc | 828,3140966 | 0,754663501 | 0,107767281 | 7,002714517 | 2,51E-12 | 3,07E-11 |
| ENSG00000220785.7 | MTMR9LP | transcribed_unproc | 392,0023579 | -0,363018129 | 0,125006677 | -2,903989908 | 0,0036844 | 0,012459761 |
| ENSG00000220848.5 | RPS18P9 | transcribed_process | 150,100563 | -0,704882497 | 0,215917712 | -3,264588582 | 0,001096232 | 0,004211017 |
| ENSG00000221817.10 | PPP3CB-AS1 | lncRNA | 812,6735369 | 0,261450645 | 0,103357117 | 2,529585317 | 0,011419741 | 0,033482733 |
| ENSG00000221818.9 | EBF2 | protein_coding | 149,2733857 | -0,613489858 | 0,210621071 | -2,912765828 | 0,003582431 | 0,012151131 |
| ENSG00000221829.9 | FANCG | protein_coding | 3874,138918 | -0,360969114 | 0,0609137 | -5,925910126 | 3,11E-09 | 2,74E-08 |
| ENSG00000221838.9 | AP4M1 | protein_coding | 2439,561368 | -0,224828616 | 0,070925835 | -3,169911457 | 0,001524854 | 0,005678973 |
| ENSG00000221869.4 | CEBPD | protein_coding | 51,83666077 | 1,56748054 | 0,353946321 | 4,428582662 | 9,49E-06 | 5,34E-05 |
| ENSG00000221883.3 | ARIH2OS | protein_coding | 262,6222271 | 0,434984803 | 0,158279879 | 2,748200245 | 0,00599234 | 0,019136917 |
| ENSG00000221914.11 | PPP2R2A | protein_coding | 16891,12393 | 0,757643339 | 0,051583252 | 14,68777775 | 7,72E-49 | 4,27E-47 |
| ENSG00000221963.6 | APOL6 | protein_coding | 26126,8188 | 5,577882601 | 0,198111566 | 28,15525975 | 2,07E-174 | 3,78E-172 |
| ENSG00000221983.7 | UBA52 | protein_coding | 20685,15771 | -0,266482812 | 0,044819711 | -5,945661085 | 2,75E-09 | 2,44E-08 |
| ENSG00000222009.8 | BTBD19 | protein_coding | 279,5152177 | -0,568100018 | 0,166774086 | -3,406404618 | 0,000658245 | 0,002659567 |
| ENSG00000222011.9 | FAM185A | protein_coding | 1008,629812 | -0,304369505 | 0,082226949 | -3,701578484 | 0,000214262 | 0,000955394 |
| ENSG00000222012.1 | AC005481.1 | lncRNA | 184,3338743 | 0,530692868 | 0,171139973 | 3,10092879 | 0,001929147 | 0,006995838 |
| ENSG00000223403.5 | MEG9 | lncRNA | 1216,785914 | -0,297753669 | 0,095615664 | -3,114067892 | 0,00184527 | 0,006727679 |
| ENSG00000223478.1 | AL441992.1 | lncRNA | 719,4098284 | -0,685077533 | 0,095378845 | -7,182698946 | 6,83E-13 | 8,77E-12 |
| ENSG00000223496.3 | EXOSC6 | protein_coding | 3620,13524 | -0,147165941 | 0,059017119 | -2,493614436 | 0,012644982 | 0,036621962 |
| ENSG00000223501.9 | VPS52 | protein_coding | 6445,911912 | -0,127433333 | 0,047939287 | -2,658223366 | 0,007855381 | 0,02427468 |
| ENSG00000223509.8 | AC135983.2 | transcribed_process | 1292,760295 | 0,29697623 | 0,083486467 | 3,557178062 | 0,00037486 | 0,001597449 |
| ENSG00000223547.10 | ZNF844 | protein_coding | 653,5448329 | 0,667551628 | 0,146364662 | 4,56087979 | 5,09E-06 | 2,98E-05 |
| ENSG00000223658.8 | C1GALT1C1L | protein_coding | 378,6915155 | 0,338003467 | 0,132937597 | 2,542572407 | 0,01100398 | 0,032475231 |
| ENSG00000223705.9 | NSUN5P1 | transcribed_unproc | 3710,480158 | -0,183372138 | 0,064527019 | -2,841788479 | 0,004486124 | 0,014817047 |
| ENSG00000223768.2 | LINC00205 | lncRNA | 5751,402051 | -0,30569504 | 0,066971133 | -4,564579167 | 5,00E-06 | 2,93E-05 |
| ENSG00000223773.7 | CD99P1 | transcribed_unproc | 563,5836053 | 0,56453382 | 0,111441149 | 5,06575737 | 4,07E-07 | 2,79E-06 |
| ENSG00000223802.7 | CERS1 | protein_coding | 960,725861 | -0,387818989 | 0,093708236 | -4,138579554 | 3,49E-05 | 0,000179345 |
| ENSG00000223837.2 | AL645941.1 | lncRNA | 105,1769096 | 0,674634433 | 0,239208203 | 2,820281344 | 0,004798156 | 0,015720771 |
| ENSG00000223839.8 | FAM95B1 | transcribed_unproc | 1063,542639 | -0,524049447 | 0,106844233 | -4,9047986 | 9,35E-07 | 6,10E-06 |
| ENSG00000223891.6 | OSER1-DT | lncRNA | 897,26823 | -0,340015289 | 0,087318715 | -3,893956617 | 9,86E-05 | 0,00046952 |
| ENSG00000223959.8 | AFG3L1P | transcribed_unitary | 3194,925567 | 0,173974734 | 0,072786624 | 2,390201999 | 0,016839108 | 0,046886627 |
| ENSG00000223960.7 | PRKRA-AS1 | lncRNA | 3669,76134 | 1,829286289 | 0,054511769 | 33,55763919 | 6,97E-247 | 1,67E-244 |
| ENSG00000223969.6 | AC002456.1 | lncRNA | 1013,975715 | -1,148414223 | 0,098866094 | -11,61585505 | 3,42E-31 | 1,15E-29 |
| ENSG00000224043.8 | CCNT2-AS1 | lncRNA | 1309,833618 | -0,844096524 | 0,07447544 | -11,33389112 | 8,92E-30 | 2,77E-28 |
| ENSG00000224046.2 | AC005076.1 | lncRNA | 231,421687 | -1,7519226 | 0,175937092 | -9,95766488 | 2,33E-23 | 5,60E-22 |
| ENSG00000224078.15 | SNHG14 | lncRNA | 17444,19689 | -0,922933326 | 0,053054862 | -17,39582935 | 8,87E-68 | 6,98E-66 |
| ENSG00000224081.9 | SLC44A3-AS1 | transcribed_process | 357,1725344 | 0,441300334 | 0,133168515 | 3,313848882 | 0,000920212 | 0,003596641 |

| | | | | | | | | |
|---|---|---|---|---|---|---|---|---|
| ENSG00000224086.6 | AC245452.1 | lncRNA | 219,6111806 | -0,481618055 | 0,168231912 | -2,862822214 | 0,004198861 | 0,013989367 |
| ENSG00000224090.1 | AC097468.1 | lncRNA | 463,2173844 | 0,290581038 | 0,117650631 | 2,469863818 | 0,01351645 | 0,038796451 |
| ENSG00000224152.1 | AC009506.1 | lncRNA | 634,4443252 | -0,52702026 | 0,103570153 | -5,088534118 | 3,61E-07 | 2,49E-06 |
| ENSG00000224165.6 | DNAJC27-AS1 | lncRNA | 115,0658391 | -0,960762861 | 0,29084966 | -3,303297173 | 0,000955551 | 0,003723474 |
| ENSG00000224281.5 | SLC25A5-AS1 | lncRNA | 382,3526911 | -0,36586557 | 0,130730578 | -2,798622763 | 0,005132106 | 0,016712756 |
| ENSG00000224292.1 | AF196972.1 | lncRNA | 61,16327904 | -1,035397761 | 0,322260455 | -3,212922168 | 0,001313919 | 0,004959752 |
| ENSG00000224308.1 | LINC01527 | lncRNA | 53,82975937 | -1,308738168 | 0,333870746 | -3,919894705 | 8,86E-05 | 0,000425618 |
| ENSG00000224389.9 | C4B | protein_coding | 2481,826489 | 0,587981072 | 0,100226661 | 5,866513633 | 4,45E-09 | 3,86E-08 |
| ENSG00000224511.2 | LINC00365 | lncRNA | 72,78761694 | 0,735447806 | 0,271165621 | 2,712172005 | 0,00668439 | 0,021084491 |
| ENSG00000224531.6 | SMIM13 | protein_coding | 9532,760589 | -0,210175186 | 0,048616169 | -4,323154035 | 1,54E-05 | 8,35E-05 |
| ENSG00000224578.5 | HNRNPA1P48 | protein_coding | 520,8214044 | -0,65732131 | 0,130096554 | -5,052565123 | 4,36E-07 | 2,99E-06 |
| ENSG00000224687.2 | RASAL2-AS1 | lncRNA | 512,7757226 | 0,319171404 | 0,123682641 | 2,580567506 | 0,009863807 | 0,029583535 |
| ENSG00000224712.12 | NPIPA3 | protein_coding | 149,7560446 | 1,166596866 | 0,236407995 | 4,934676018 | 8,03E-07 | 5,28E-06 |
| ENSG00000224717.1 | AC098936.1 | lncRNA | 70,92049725 | -0,737570622 | 0,279752527 | -2,636511026 | 0,008376348 | 0,025683587 |
| ENSG00000224790.3 | AP000704.1 | lncRNA | 428,2014434 | -0,616160982 | 0,124284795 | -4,957653774 | 7,13E-07 | 4,73E-06 |
| ENSG00000224877.4 | NDUFAF8 | protein_coding | 486,6388097 | -1,226531541 | 0,117135502 | -10,47104868 | 1,17E-25 | 3,13E-24 |
| ENSG00000224903.2 | AC005534.1 | lncRNA | 35,91044043 | -1,041431329 | 0,402763762 | -2,585712588 | 0,009717793 | 0,029200079 |
| ENSG00000224914.4 | LINC00863 | lncRNA | 2956,914332 | 0,171603213 | 0,061501761 | 2,790216247 | 0,005267285 | 0,017078841 |
| ENSG00000224945.1 | AL353150.1 | lncRNA | 100,2500055 | 0,662036557 | 0,263063681 | 2,516639907 | 0,011847985 | 0,034594149 |
| ENSG00000224957.6 | LINC01266 | lncRNA | 162,0888065 | -0,821466105 | 0,197213689 | -4,165360479 | 3,11E-05 | 0,000160965 |
| ENSG00000224992.2 | AL445645.1 | lncRNA | 13,33402131 | 2,566306467 | 0,70030256 | 3,664568164 | 0,000247756 | 0,001092623 |
| ENSG00000225075.1 | AL603832.1 | lncRNA | 11,66989129 | -4,20564925 | 1,010592725 | -4,161566916 | 3,16E-05 | 0,000163551 |
| ENSG00000225127.3 | LINC00237 | lncRNA | 340,5705299 | 0,344524201 | 0,145079241 | 2,374731207 | 0,017561734 | 0,048568259 |
| ENSG00000225138.8 | SLC9A3-AS1 | lncRNA | 1201,367218 | -0,241145185 | 0,091662288 | -2,630800413 | 0,008518405 | 0,026069395 |
| ENSG00000225190.11 | PLEKHM1 | protein_coding | 1537,420393 | 0,36165802 | 0,081364014 | 4,444938283 | 8,79E-06 | 4,97E-05 |
| ENSG00000225361.4 | PPP1R26-AS1 | lncRNA | 500,4037902 | 0,563527998 | 0,135041287 | 4,173005245 | 3,01E-05 | 0,000155872 |
| ENSG00000225377.6 | NRSN2-AS1 | lncRNA | 1090,026114 | 0,451618862 | 0,078401583 | 5,760328365 | 8,40E-09 | 7,04E-08 |
| ENSG00000225450.1 | AL021707.1 | lncRNA | 10,37093417 | 1,952969326 | 0,766563539 | 2,547693997 | 0,010843754 | 0,032061378 |
| ENSG00000225470.8 | JPX | lncRNA | 4671,861633 | -0,175534036 | 0,057893189 | -3,032032574 | 0,00242913 | 0,008579521 |
| ENSG00000225484.7 | NUTM2B-AS1 | lncRNA | 1720,348943 | -0,324860778 | 0,071257344 | -4,558979614 | 5,14E-06 | 3,01E-05 |
| ENSG00000225489.7 | AL354707.1 | lncRNA | 1770,064167 | 0,19034822 | 0,069702452 | 2,730868344 | 0,00631677 | 0,020065067 |
| ENSG00000225492.6 | GBP1P1 | transcribed_unproc | 218,8517945 | 8,69663691 | 0,919139746 | 9,46171346 | 3,03E-21 | 6,58E-20 |
| ENSG00000225526.4 | MKRN2OS | protein_coding | 90,62562841 | 0,687444944 | 0,254686908 | 2,699176601 | 0,006951128 | 0,021840164 |
| ENSG00000225611.1 | LINC02158 | lncRNA | 56,99655937 | 0,81754291 | 0,313612396 | 2,606857769 | 0,009137731 | 0,027716101 |
| ENSG00000225630.1 | MTND2P28 | unprocessed_pseud | 8980,744203 | -0,6079926 | 0,055400528 | -10,97449113 | 5,07E-28 | 1,48E-26 |
| ENSG00000225663.7 | MCRIP1 | protein_coding | 2881,262561 | -0,582497078 | 0,065192419 | -8,935043178 | 4,07E-19 | 7,90E-18 |

| | | | | | | | | |
|---|---|---|---|---|---|---|---|---|
| ENSG00000225670.4 | CADM3-AS1 | lncRNA | 33,41070126 | -1,441983388 | 0,440839548 | -3,270993711 | 0,001071703 | 0,004127111 |
| ENSG00000225697.13 | SLC26A6 | protein_coding | 3465,238644 | 0,375804531 | 0,060486812 | 6,212999446 | 5,20E-10 | 5,04E-09 |
| ENSG00000225733.6 | FGD5-AS1 | lncRNA | 30923,96114 | -0,363573852 | 0,04092084 | -8,884809026 | 6,40E-19 | 1,23E-17 |
| ENSG00000225746.12 | MEG8 | lncRNA | 1759,891924 | -0,430983342 | 0,067262757 | -6,407458776 | 1,48E-10 | 1,52E-09 |
| ENSG00000225778.5 | PROSER2-AS1 | lncRNA | 17,27631082 | -2,174443952 | 0,589261909 | -3,690114564 | 0,000224153 | 0,00099693 |
| ENSG00000225783.8 | MIAT | lncRNA | 992,3224422 | -0,880784812 | 0,127224763 | -6,923061145 | 4,42E-12 | 5,26E-11 |
| ENSG00000225849.2 | MKRN7P | transcribed_process | 85,24794505 | 0,693760941 | 0,249924896 | 2,775877683 | 0,005505292 | 0,017768656 |
| ENSG00000225855.7 | RUSC1-AS1 | lncRNA | 221,3779986 | 1,660499027 | 0,181156575 | 9,166098612 | 4,90E-20 | 9,95E-19 |
| ENSG00000225932.3 | CTAGE4 | protein_coding | 533,682157 | 0,536408306 | 0,12345969 | 4,344805228 | 1,39E-05 | 7,63E-05 |
| ENSG00000225964.6 | NRIR | lncRNA | 129,0436546 | 4,716784793 | 0,346903394 | 13,59682517 | 4,18E-42 | 2,00E-40 |
| ENSG00000225972.1 | MTND1P23 | unprocessed_pseud | 5623,915888 | -0,264418847 | 0,060522813 | -4,368912054 | 1,25E-05 | 6,89E-05 |
| ENSG00000225975.7 | LINC01534 | lncRNA | 449,429859 | -0,483546798 | 0,127895203 | -3,78080479 | 0,000156322 | 0,000716193 |
| ENSG00000226025.9 | AC005515.1 | transcribed_unproc | 57,66097386 | 2,701016549 | 0,375066078 | 7,201441849 | 5,96E-13 | 7,66E-12 |
| ENSG00000226026.6 | AC092802.1 | lncRNA | 267,2639184 | 0,603205049 | 0,169508 | 3,558563895 | 0,000372888 | 0,001589647 |
| ENSG00000226067.7 | LINC00623 | lncRNA | 1282,039403 | 0,427785254 | 0,076613835 | 5,583655407 | 2,36E-08 | 1,88E-07 |
| ENSG00000226137.5 | BAIAP2-DT | lncRNA | 3529,257776 | -0,288378285 | 0,055744903 | -5,173177605 | 2,30E-07 | 1,63E-06 |
| ENSG00000226210.3 | WASH8P | unprocessed_pseud | 847,2862888 | 0,359992244 | 0,095520879 | 3,768728359 | 0,000164081 | 0,00074839 |
| ENSG00000226268.3 | AC135977.1 | processed_pseudog | 55,53633418 | 7,902983808 | 2,921302807 | 2,705294291 | 0,00682439 | 0,021480939 |
| ENSG00000226314.8 | ZNF192P1 | transcribed_unproc | 216,4752859 | 0,555282944 | 0,174009489 | 3,191107273 | 0,001417286 | 0,005310754 |
| ENSG00000226332.2 | AL354836.1 | lncRNA | 53,18259398 | -0,88717293 | 0,341891966 | -2,594892596 | 0,009462048 | 0,028565081 |
| ENSG00000226363.3 | HAGLROS | lncRNA | 34,00814723 | -1,631887563 | 0,464743945 | -3,511369172 | 0,000445805 | 0,001872131 |
| ENSG00000226383.7 | LINC01876 | lncRNA | 357,704898 | -0,38802057 | 0,147072285 | -2,63829837 | 0,008332323 | 0,025566798 |
| ENSG00000226416.1 | MRPL23-AS1 | lncRNA | 371,7301201 | -0,365902314 | 0,139449712 | -2,623901534 | 0,008692891 | 0,026548939 |
| ENSG00000226419.8 | SLC16A1-AS1 | lncRNA | 232,60402 | 0,465013292 | 0,171160993 | 2,716818145 | 0,006591281 | 0,020822898 |
| ENSG00000226453.2 | LINC02542 | lncRNA | 379,5123724 | 1,765870976 | 0,131814599 | 13,39662674 | 6,33E-41 | 2,91E-39 |
| ENSG00000226562.4 | CYP4F26P | lncRNA | 58,51342837 | -1,201594013 | 0,320358528 | -3,750778917 | 0,000176286 | 0,000799038 |
| ENSG00000226686.8 | LINC01535 | lncRNA | 124,2548581 | -0,812277398 | 0,32477384 | -2,501055498 | 0,012382377 | 0,035981788 |
| ENSG00000226742.4 | HSBP1L1 | protein_coding | 400,3711199 | -0,525734132 | 0,122672813 | -4,28566133 | 1,82E-05 | 9,78E-05 |
| ENSG00000226752.9 | CUTALP | transcribed_unitary | 9077,205537 | 0,198389815 | 0,050863983 | 3,900398693 | 9,60E-05 | 0,000458266 |
| ENSG00000226791.7 | LINC02611 | lncRNA | 473,5840983 | 0,68001209 | 0,129976245 | 5,231818248 | 1,68E-07 | 1,21E-06 |
| ENSG00000226913.2 | BSN-DT | lncRNA | 71,59013102 | 0,802202087 | 0,323944218 | 2,476358713 | 0,013273015 | 0,038200011 |
| ENSG00000226950.7 | DANCR | lncRNA | 1731,850802 | -2,349146628 | 0,071932749 | -32,65753987 | 6,26E-234 | 1,40E-231 |
| ENSG00000226954.1 | AL023802.1 | lncRNA | 218,7449718 | -0,396629207 | 0,165559947 | -2,395683349 | 0,016589415 | 0,046277251 |
| ENSG00000227051.6 | C14orf132 | protein_coding | 39877,82727 | -0,366933907 | 0,043815386 | -8,374544685 | 5,54E-17 | 9,46E-16 |
| ENSG00000227097.5 | RPS28P7 | processed_pseudog | 2490,064573 | -0,413238346 | 0,06742791 | -6,128594876 | 8,87E-10 | 8,33E-09 |
| ENSG00000227199.1 | ST7-AS1 | lncRNA | 102,0722937 | 1,170138736 | 0,248773419 | 4,703632495 | 2,56E-06 | 1,57E-05 |

| | | | | | | | | |
|---|---|---|---|---|---|---|---|---|
| ENSG00000227345.8 | PARG | protein_coding | 5781,973197 | 0,1816104 | 0,051940271 | 3,496523926 | 0,000471362 | 0,001967345 |
| ENSG00000227403.2 | LINC01806 | lncRNA | 864,3785267 | 0,595807449 | 0,096645686 | 6,164863393 | 7,05E-10 | 6,71E-09 |
| ENSG00000227502.3 | LINC01268 | lncRNA | 302,747909 | 0,39667747 | 0,166723244 | 2,379257157 | 0,017347569 | 0,048082309 |
| ENSG00000227507.2 | LTB | protein_coding | 36,45831578 | 2,677758773 | 0,433397004 | 6,1785355 | 6,47E-10 | 6,19E-09 |
| ENSG00000227508.6 | LINC01624 | lncRNA | 62,99948279 | -1,217475673 | 0,354137332 | -3,437863122 | 0,000586324 | 0,002395203 |
| ENSG00000227627.3 | AL080276.2 | lncRNA | 341,6298279 | -0,378660245 | 0,13719826 | -2,759949317 | 0,005781033 | 0,018549575 |
| ENSG00000227630.4 | LINC01132 | lncRNA | 15,93477006 | -1,809225754 | 0,713678512 | -2,535071078 | 0,011242451 | 0,033070614 |
| ENSG00000227755.1 | AP000344.1 | transcribed_process | 64,14766535 | -1,181561954 | 0,337999921 | -3,495746241 | 0,000472738 | 0,001971625 |
| ENSG00000227766.1 | AL671277.1 | unprocessed_pseud | 25,42385351 | 3,778061134 | 0,559704785 | 6,750096186 | 1,48E-11 | 1,66E-10 |
| ENSG00000227827.3 | AC138969.2 | unprocessed_pseud | 960,9074169 | 0,397525487 | 0,117781178 | 3,375118967 | 0,000737838 | 0,002947816 |
| ENSG00000227857.2 | AL358075.2 | lncRNA | 49,63435451 | -0,941674269 | 0,326117123 | -2,887533966 | 0,003882747 | 0,013048186 |
| ENSG00000227877.6 | MRLN | protein_coding | 3843,331164 | -1,163393528 | 0,057125845 | -20,36544963 | 3,39E-92 | 3,63E-90 |
| ENSG00000227885.3 | AL590652.1 | lncRNA | 119,6076876 | 0,699537587 | 0,242135389 | 2,889034884 | 0,003864262 | 0,01299771 |
| ENSG00000227963.1 | RBM15-AS1 | lncRNA | 83,05651876 | -0,748658455 | 0,279370548 | -2,679804512 | 0,007366517 | 0,022944111 |
| ENSG00000228065.11 | LINC01515 | lncRNA | 1151,298478 | 0,220884918 | 0,086873446 | 2,542605694 | 0,011002932 | 0,032475231 |
| ENSG00000228106.6 | AL392172.1 | lncRNA | 747,7385507 | -0,401272121 | 0,113228317 | -3,543920197 | 0,000394225 | 0,001669225 |
| ENSG00000228133.3 | AC099684.1 | lncRNA | 227,3918574 | 0,877545816 | 0,155851811 | 5,630642402 | 1,80E-08 | 1,45E-07 |
| ENSG00000228141.7 | AC105339.1 | transcribed_unitary | 18,80944195 | 1,941915475 | 0,672427463 | 2,887918149 | 0,003878008 | 0,013036152 |
| ENSG00000228223.3 | HCG11 | lncRNA | 6036,513974 | -0,318499926 | 0,053041585 | -6,004721166 | 1,92E-09 | 1,74E-08 |
| ENSG00000228253.1 | MT-ATP8 | protein_coding | 30434,48722 | -1,23806945 | 0,054069001 | -22,89795316 | 4,87E-116 | 6,41E-114 |
| ENSG00000228327.3 | AL669831.1 | transcribed_unproc | 1421,021074 | 0,75738756 | 0,082083003 | 9,227093653 | 2,78E-20 | 5,73E-19 |
| ENSG00000228340.6 | MIR646HG | lncRNA | 34,94041707 | -1,393625039 | 0,506038787 | -2,753988579 | 0,005887382 | 0,018839963 |
| ENSG00000228409.6 | CCT6P1 | transcribed_unproc | 497,0732945 | 0,561534087 | 0,118747373 | 4,728812719 | 2,26E-06 | 1,39E-05 |
| ENSG00000228474.6 | OST4 | protein_coding | 8028,959165 | -0,306730294 | 0,046305704 | -6,624028356 | 3,50E-11 | 3,80E-10 |
| ENSG00000228544.1 | CCDC183-AS1 | lncRNA | 846,0702985 | 0,440752765 | 0,099481608 | 4,430494978 | 9,40E-06 | 5,30E-05 |
| ENSG00000228570.7 | NUTM2E | protein_coding | 81,20416864 | 0,825645042 | 0,329502758 | 2,505730291 | 0,012219879 | 0,035560086 |
| ENSG00000228606.3 | AL139011.1 | lncRNA | 27,58571532 | 1,124630344 | 0,428090254 | 2,627087008 | 0,008611931 | 0,026327035 |
| ENSG00000228648.1 | AL353747.2 | lncRNA | 158,1877098 | -0,911007099 | 0,190457174 | -4,783264804 | 1,72E-06 | 1,08E-05 |
| ENSG00000228716.7 | DHFR | protein_coding | 15165,0743 | -0,965717149 | 0,053687533 | -17,98773571 | 2,43E-72 | 2,03E-70 |
| ENSG00000228782.7 | MRPL45P2 | transcribed_unproc | 442,4203427 | -0,539027497 | 0,125032382 | -4,311103141 | 1,62E-05 | 8,80E-05 |
| ENSG00000228793.2 | AL033523.1 | lncRNA | 25,69491613 | 1,840757722 | 0,500968017 | 3,674401673 | 0,000238407 | 0,001055737 |
| ENSG00000228794.9 | LINC01128 | lncRNA | 5801,233513 | 0,248377662 | 0,048613944 | 5,109185603 | 3,24E-07 | 2,25E-06 |
| ENSG00000228903.7 | RASA4CP | transcribed_unproc | 1475,998082 | -0,412773436 | 0,092068975 | -4,483306528 | 7,35E-06 | 4,19E-05 |
| ENSG00000228960.6 | OR2A9P | transcribed_unproc | 372,9078115 | 0,493614887 | 0,132865976 | 3,715133856 | 0,000203096 | 0,000908845 |
| ENSG00000229086.5 | LINC01548 | lncRNA | 438,8700909 | -0,983438937 | 0,131305562 | -7,489697507 | 6,90E-14 | 9,60E-13 |
| ENSG00000229117.9 | RPL41 | protein_coding | 66977,84417 | -0,321400317 | 0,044724565 | -7,186214503 | 6,66E-13 | 8,55E-12 |

| | | | | | | | | |
|---|---|---|---|---|---|---|---|---|
| ENSG00000229156.1 | CR769776.2 | processed_pseudog | 8,345279755 | -3,559101964 | 1,167289518 | -3,049031031 | 0,002295808 | 0,008162428 |
| ENSG00000229191.1 | AL358473.1 | lncRNA | 62,12930427 | -0,850982649 | 0,303609851 | -2,802882208 | 0,005064816 | 0,016503178 |
| ENSG00000229320.3 | KRT8P12 | transcribed_process | 230,0851609 | 0,583232888 | 0,170482458 | 3,421072728 | 0,000623746 | 0,002531082 |
| ENSG00000229474.6 | PATL2 | protein_coding | 145,0139531 | 5,738851995 | 0,635611728 | 9,028864234 | 1,73E-19 | 3,42E-18 |
| ENSG00000229589.1 | ACVR2B-AS1 | lncRNA | 186,2952494 | 0,699815442 | 0,184348789 | 3,796148842 | 0,000146961 | 0,000678833 |
| ENSG00000229644.6 | NAMPTP1 | processed_pseudog | 381,1377366 | 1,659718174 | 0,134654506 | 12,32575296 | 6,58E-35 | 2,50E-33 |
| ENSG00000229659.1 | RPL26P6 | processed_pseudog | 25,30999485 | 1,201676712 | 0,478092586 | 2,51348117 | 0,011954616 | 0,034860273 |
| ENSG00000229677.1 | AC018644.1 | processed_pseudog | 6,99919617 | 3,089419384 | 1,051237802 | 2,938839697 | 0,003294434 | 0,011274809 |
| ENSG00000229833.9 | PET100 | protein_coding | 3813,789215 | -0,234604583 | 0,069047413 | -3,397731684 | 0,00067947 | 0,002735503 |
| ENSG00000229891.2 | LINC01315 | lncRNA | 183,091984 | -1,137849222 | 0,18595614 | -6,118911807 | 9,42E-10 | 8,83E-09 |
| ENSG00000229950.1 | TFAP2A-AS1 | lncRNA | 33,17665554 | 3,035759752 | 0,448476978 | 6,769042557 | 1,30E-11 | 1,47E-10 |
| ENSG00000230023.2 | AL590683.1 | lncRNA | 49,77990324 | -1,11246337 | 0,341874587 | -3,254010133 | 0,001137882 | 0,004353926 |
| ENSG00000230067.3 | HSPD1P6 | transcribed_process | 8,754049547 | -2,021974009 | 0,823020319 | -2,456772892 | 0,014019128 | 0,040070674 |
| ENSG00000230107.1 | AL022316.1 | lncRNA | 17,8755112 | 1,556892379 | 0,571563549 | 2,723918244 | 0,006451247 | 0,020435003 |
| ENSG00000230149.2 | AL021707.3 | lncRNA | 36,94525806 | 1,808385345 | 0,414151022 | 4,366487707 | 1,26E-05 | 6,96E-05 |
| ENSG00000230177.2 | AL080317.1 | lncRNA | 652,7729135 | -0,325165477 | 0,098704808 | -3,294322578 | 0,000986592 | 0,003828168 |
| ENSG00000230183.4 | CNOT6LP1 | processed_pseudog | 12,66560735 | 4,077474037 | 1,086193967 | 3,753909672 | 0,000174098 | 0,000790871 |
| ENSG00000230223.7 | ATXN8OS | lncRNA | 122,8524929 | -0,681760832 | 0,223333112 | -3,052663462 | 0,002268202 | 0,008075759 |
| ENSG00000230267.7 | HERC2P4 | transcribed_unproc | 272,9264127 | 0,394601468 | 0,142601122 | 2,767169445 | 0,005654536 | 0,01819556 |
| ENSG00000230269.7 | LINC02525 | lncRNA | 1751,538095 | -0,37137742 | 0,071150305 | -5,219618094 | 1,79E-07 | 1,28E-06 |
| ENSG00000230356.1 | NCAPD2P1 | processed_pseudog | 204,9827911 | 0,522008242 | 0,1625751 | 3,210874488 | 0,001323317 | 0,004993554 |
| ENSG00000230373.8 | GOLGA6L5P | transcribed_unproc | 752,8907204 | -0,26387005 | 0,108105471 | -2,44085751 | 0,014652435 | 0,041579886 |
| ENSG00000230424.1 | EMC1-AS1 | lncRNA | 24,4942876 | -1,313948547 | 0,48816152 | -2,69162663 | 0,00711045 | 0,022266154 |
| ENSG00000230438.7 | SERPINB9P1 | lncRNA | 17,5994521 | 4,187250578 | 0,877945341 | 4,769375022 | 1,85E-06 | 1,15E-05 |
| ENSG00000230453.9 | ANKRD18B | protein_coding | 263,9910465 | -0,560892138 | 0,175850341 | -3,189599371 | 0,001424701 | 0,005332801 |
| ENSG00000230498.2 | AL035409.1 | lncRNA | 95,61573377 | -0,915490858 | 0,272063455 | -3,364990189 | 0,000765464 | 0,003045753 |
| ENSG00000230521.1 | AL645929.1 | unprocessed_pseud | 111,6075932 | 3,31659197 | 0,293059564 | 11,31712587 | 1,08E-29 | 3,33E-28 |
| ENSG00000230615.7 | AL139220.2 | lncRNA | 163,9146655 | -1,317859202 | 0,263863698 | -4,994469546 | 5,90E-07 | 3,96E-06 |
| ENSG00000230715.3 | AC018638.2 | processed_pseudog | 112,9461386 | 0,853083027 | 0,2384555 | 3,577535543 | 0,000346849 | 0,001487941 |
| ENSG00000230724.9 | LINC01001 | lncRNA | 432,9464787 | 0,733961958 | 0,114280063 | 6,422484736 | 1,34E-10 | 1,38E-09 |
| ENSG00000230790.2 | AC012456.2 | lncRNA | 40,29164334 | 0,984468559 | 0,390769668 | 2,519306487 | 0,011758626 | 0,034364438 |
| ENSG00000230795.3 | HLA-K | unprocessed_pseud | 124,897942 | 3,066423955 | 0,243566881 | 12,58965893 | 2,41E-36 | 9,76E-35 |
| ENSG00000230847.4 | OCLNP1 | unprocessed_pseud | 146,0681939 | 0,648121157 | 0,261737585 | 2,476225019 | 0,013277986 | 0,038209434 |
| ENSG00000230896.1 | AL604028.1 | lncRNA | 115,5787804 | -0,784839765 | 0,231744271 | -3,386663069 | 0,000707482 | 0,002838124 |
| ENSG00000230989.6 | HSBP1 | protein_coding | 34349,5493 | -0,244634926 | 0,054445771 | -4,493185123 | 7,02E-06 | 4,02E-05 |
| ENSG00000231079.7 | AC105402.3 | lncRNA | 135,4416174 | 0,540762408 | 0,198442722 | 2,725030185 | 0,006429561 | 0,020372042 |

| | | | | | | | | |
|---|---|---|---|---|---|---|---|---|
| ENSG00000231107.2 | LINC01508 | lncRNA | 95,60830365 | 1,266745157 | 0,262765592 | 4,820818242 | 1,43E-06 | 9,08E-06 |
| ENSG00000231233.1 | CFAP58-DT | lncRNA | 21,15641613 | -1,739737854 | 0,599324531 | -2,902831044 | 0,003698061 | 0,01249657 |
| ENSG00000231291.3 | AC106799.1 | lncRNA | 30,0931895 | -1,415697749 | 0,483193889 | -2,929875108 | 0,003390983 | 0,011575315 |
| ENSG00000231304.2 | SGO1-AS1 | lncRNA | 255,1005748 | -0,369082291 | 0,152571976 | -2,419069998 | 0,015560246 | 0,043741874 |
| ENSG00000231322.5 | RPL13AP17 | transcribed_process | 9,087471091 | 2,66755471 | 0,945824396 | 2,820348811 | 0,004797147 | 0,015720771 |
| ENSG00000231500.7 | RPS18 | protein_coding | 68428,53093 | -0,452791541 | 0,039930288 | -11,33955116 | 8,36E-30 | 2,60E-28 |
| ENSG00000231535.7 | LINC00278 | lncRNA | 239,7117641 | 1,684448459 | 0,179156715 | 9,402095031 | 5,35E-21 | 1,15E-19 |
| ENSG00000231551.8 | AC245100.4 | transcribed_unproc | 137,1535596 | -0,551940299 | 0,208953923 | -2,641445025 | 0,00825532 | 0,025369423 |
| ENSG00000231574.6 | LINC02015 | lncRNA | 111,0854699 | 2,028459973 | 0,251338965 | 8,070614794 | 6,99E-16 | 1,11E-14 |
| ENSG00000231607.12 | DLEU2 | lncRNA | 1541,116382 | -0,241346449 | 0,070530343 | -3,421881134 | 0,000621895 | 0,002524479 |
| ENSG00000231663.1 | COA6-AS1 | lncRNA | 52,80695605 | -0,947227498 | 0,316519311 | -2,992637309 | 0,002765782 | 0,009634674 |
| ENSG00000231742.6 | LINC01273 | lncRNA | 82,43411006 | 0,78152313 | 0,268189297 | 2,914072781 | 0,003567467 | 0,012106154 |
| ENSG00000231764.9 | DLX6-AS1 | lncRNA | 1778,397937 | -0,625953217 | 0,072208718 | -8,668665373 | 4,37E-18 | 8,01E-17 |
| ENSG00000231769.3 | AL035701.1 | lncRNA | 48,70378486 | -1,090805405 | 0,343784921 | -3,172929756 | 0,00150909 | 0,005626655 |
| ENSG00000231889.7 | TRAF3IP2-AS1 | lncRNA | 540,2046951 | -0,256290614 | 0,104738613 | -2,44695443 | 0,014406905 | 0,040995128 |
| ENSG00000231925.11 | TAPBP | protein_coding | 8159,441161 | 2,243241244 | 0,058575674 | 38,2964649 | 0 | 0 |
| ENSG00000232000.3 | CLCN3P1 | transcribed_unproc | 930,865974 | 0,280706171 | 0,099660648 | 2,816619963 | 0,004853191 | 0,015887214 |
| ENSG00000232053.7 | AC078845.1 | lncRNA | 15,21571885 | -1,780414779 | 0,711240883 | -2,50325146 | 0,012305808 | 0,035777761 |
| ENSG00000232070.8 | TMEM253 | protein_coding | 31,29364463 | -1,152218905 | 0,421952341 | -2,730684943 | 0,006320286 | 0,02007101 |
| ENSG00000232104.3 | RFX3-AS1 | lncRNA | 490,2441244 | 0,454915818 | 0,11479247 | 3,962941299 | 7,40E-05 | 0,000361048 |
| ENSG00000232110.8 | AL353751.1 | lncRNA | 170,893109 | 1,889077497 | 0,204425819 | 9,240894832 | 2,44E-20 | 5,07E-19 |
| ENSG00000232112.3 | TMA7 | protein_coding | 12894,21339 | -0,823313106 | 0,05520215 | -14,91451161 | 2,65E-50 | 1,53E-48 |
| ENSG00000232119.8 | MCTS1 | protein_coding | 4828,312684 | -0,23358502 | 0,062706484 | -3,72505372 | 0,000195274 | 0,000878029 |
| ENSG00000232164.1 | LINC01873 | lncRNA | 512,2269533 | 0,266818543 | 0,109805826 | 2,429912441 | 0,015102471 | 0,042663343 |
| ENSG00000232229.6 | LINC00865 | lncRNA | 2702,675908 | 0,512709655 | 0,057645222 | 8,894226446 | 5,88E-19 | 1,13E-17 |
| ENSG00000232434.2 | AJM1 | protein_coding | 758,5287848 | -0,351651884 | 0,094675232 | -3,714296527 | 0,00020377 | 0,000911497 |
| ENSG00000232442.1 | MHENCR | lncRNA | 415,9738271 | -0,557034488 | 0,137561788 | -4,049340274 | 5,14E-05 | 0,000256857 |
| ENSG00000232485.3 | AC098820.1 | lncRNA | 297,7672214 | -0,353583743 | 0,134384656 | -2,631131805 | 0,008510102 | 0,026047579 |
| ENSG00000232573.1 | RPL3P4 | processed_pseudog | 966,8167377 | -0,758168349 | 0,101741389 | -7,451916656 | 9,20E-14 | 1,26E-12 |
| ENSG00000232591.2 | LINC02642 | lncRNA | 16,97599571 | 1,567016098 | 0,617753335 | 2,536637215 | 0,011192287 | 0,032948887 |
| ENSG00000232593.7 | KANTR | protein_coding | 863,9604639 | -0,272327946 | 0,093842519 | -2,901967559 | 0,003708269 | 0,012527305 |
| ENSG00000232645.5 | LINC01431 | lncRNA | 90,62939035 | -0,829126463 | 0,259037934 | -3,200791676 | 0,001370506 | 0,005151756 |
| ENSG00000232648.4 | AC107214.1 | lncRNA | 108,6700629 | -1,033396371 | 0,226675244 | -4,558929125 | 5,14E-06 | 3,01E-05 |
| ENSG00000232721.2 | AC239800.2 | lncRNA | 152,9993127 | 1,120700297 | 0,208990061 | 5,362457387 | 8,21E-08 | 6,10E-07 |
| ENSG00000232838.4 | PET117 | protein_coding | 3335,19844 | -0,147926718 | 0,061140409 | -2,419459088 | 0,015543609 | 0,043712031 |
| ENSG00000232906.1 | AC092685.1 | processed_pseudog | 34,46940204 | -1,011062734 | 0,400374304 | -2,525293767 | 0,011560161 | 0,033828349 |

| | | | | | | | | |
|---|---|---|---|---|---|---|---|---|
| ENSG00000233013.10 | FAM157B | transcribed_unproc | 66,26071432 | 1,400481468 | 0,324564436 | 4,314956637 | 1,60E-05 | 8,65E-05 |
| ENSG00000233024.7 | NPIPA9 | protein_coding | 1537,066589 | 0,273727573 | 0,086631043 | 3,159693831 | 0,00157935 | 0,005858535 |
| ENSG00000233041.8 | PHGR1 | protein_coding | 52,90713629 | -1,055308984 | 0,383204534 | -2,753905268 | 0,005888881 | 0,018841908 |
| ENSG00000233178.7 | AL161457.2 | lncRNA | 809,8815941 | 0,330064797 | 0,101459505 | 3,253167809 | 0,001141261 | 0,00436494 |
| ENSG00000233223.3 | AC016876.1 | lncRNA | 684,1967819 | -0,316428911 | 0,122417646 | -2,584830888 | 0,009742677 | 0,029259228 |
| ENSG00000233242.3 | AL512604.2 | lncRNA | 10,96353025 | -2,143766974 | 0,733770342 | -2,921577569 | 0,003482635 | 0,011859401 |
| ENSG00000233304.7 | LINC01346 | lncRNA | 11,3659568 | -2,092132294 | 0,84661142 | -2,471183644 | 0,013466664 | 0,038668344 |
| ENSG00000233369.7 | GTF2IP4 | transcribed_process | 39343,22147 | -0,374061495 | 0,041280354 | -9,061489444 | 1,29E-19 | 2,55E-18 |
| ENSG00000233396.8 | LINC01719 | lncRNA | 1500,515989 | 0,851124346 | 0,097595072 | 8,720976716 | 2,76E-18 | 5,11E-17 |
| ENSG00000233426.3 | EIF3FP3 | processed_pseudog | 178,8134008 | -0,470748408 | 0,174486939 | -2,697900546 | 0,006977828 | 0,021908765 |
| ENSG00000233476.3 | EEF1A1P6 | processed_pseudog | 134,7289313 | -1,033979609 | 0,208327975 | -4,963229775 | 6,93E-07 | 4,61E-06 |
| ENSG00000233492.2 | AL133325.2 | lncRNA | 274,1479877 | -1,106228133 | 0,156104789 | -7,086445825 | 1,38E-12 | 1,72E-11 |
| ENSG00000233508.3 | AL021395.1 | lncRNA | 634,6368903 | 0,772662741 | 0,109223862 | 7,074120285 | 1,50E-12 | 1,88E-11 |
| ENSG00000233593.10 | LINC02609 | lncRNA | 398,9990396 | 0,47673325 | 0,134201287 | 3,552374662 | 0,000381771 | 0,001623825 |
| ENSG00000233654.2 | AC108047.1 | lncRNA | 228,5717837 | -0,587733221 | 0,169080852 | -3,476048381 | 0,00050886 | 0,002103184 |
| ENSG00000233730.1 | LINC01765 | lncRNA | 21,78384841 | 2,191069326 | 0,557948701 | 3,927008564 | 8,60E-05 | 0,000415 |
| ENSG00000233766.7 | AC098617.1 | lncRNA | 87,29618806 | 0,890333775 | 0,263425245 | 3,379834668 | 0,000725294 | 0,002902338 |
| ENSG00000233913.7 | RPL10P9 | processed_pseudog | 449,3905057 | -0,38696041 | 0,128702176 | -3,006634549 | 0,002641571 | 0,009248581 |
| ENSG00000233926.1 | AL591368.1 | lncRNA | 30,05173482 | -2,402433176 | 0,462934368 | -5,189576194 | 2,11E-07 | 1,50E-06 |
| ENSG00000233927.5 | RPS28 | protein_coding | 15784,57619 | -0,44958126 | 0,049910408 | -9,007765627 | 2,10E-19 | 4,13E-18 |
| ENSG00000233932.6 | CTXN2 | protein_coding | 854,0372386 | -0,772688797 | 0,136086594 | -5,677920031 | 1,36E-08 | 1,12E-07 |
| ENSG00000234072.1 | AC074117.1 | lncRNA | 3714,159109 | -0,155788384 | 0,058665069 | -2,655556108 | 0,007917776 | 0,02445742 |
| ENSG00000234127.9 | TRIM26 | protein_coding | 9449,718512 | 0,990018697 | 0,05034188 | 19,66590618 | 4,23E-86 | 4,28E-84 |
| ENSG00000234129.8 | AC073529.1 | lncRNA | 189,423571 | -0,542630814 | 0,1979077 | -2,741837807 | 0,00610965 | 0,019476295 |
| ENSG00000234155.1 | LINC02535 | lncRNA | 288,1673191 | 0,594305838 | 0,147422146 | 4,031319961 | 5,55E-05 | 0,000275901 |
| ENSG00000234171.3 | RNASEH1-AS1 | lncRNA | 1564,653441 | -0,236011146 | 0,092621267 | -2,548131254 | 0,010830171 | 0,032037893 |
| ENSG00000234284.6 | ZNF879 | protein_coding | 992,321728 | 0,340092341 | 0,084499926 | 4,024764946 | 5,70E-05 | 0,00028327 |
| ENSG00000234290.2 | AC116366.1 | lncRNA | 39,98819379 | 1,923734081 | 0,392202586 | 4,904950015 | 9,35E-07 | 6,10E-06 |
| ENSG00000234327.8 | AC012146.1 | lncRNA | 92,59520919 | -1,738499669 | 0,26621668 | -6,53039348 | 6,56E-11 | 6,95E-10 |
| ENSG00000234329.1 | AL604028.2 | processed_pseudog | 60,95047846 | 1,086439576 | 0,325683992 | 3,335870362 | 0,000850328 | 0,003350846 |
| ENSG00000234420.7 | ZNF37BP | transcribed_process | 5110,715584 | -0,128773197 | 0,0539434 | -2,387190957 | 0,01697767 | 0,0472082 |
| ENSG00000234423.2 | LINC01250 | lncRNA | 64,84686216 | -0,952299343 | 0,37027586 | -2,571864506 | 0,010115247 | 0,030220878 |
| ENSG00000234585.6 | CCT6P3 | transcribed_unproc | 653,2785272 | 0,487921958 | 0,099417337 | 4,907815626 | 9,21E-07 | 6,01E-06 |
| ENSG00000234618.1 | RPSAP9 | processed_pseudog | 120,4544975 | 0,81757101 | 0,245570225 | 3,329275813 | 0,000870721 | 0,00341864 |
| ENSG00000234648.1 | AL162151.2 | processed_pseudog | 431,0709466 | -0,822001855 | 0,128244107 | -6,409665685 | 1,46E-10 | 1,50E-09 |
| ENSG00000234678.2 | ELF3-AS1 | lncRNA | 690,1407283 | 0,618144425 | 0,124930952 | 4,947888524 | 7,50E-07 | 4,96E-06 |

| | | | | | | | | |
|---|---|---|---|---|---|---|---|---|
| ENSG00000234684.7 | SDCBP2-AS1 | lncRNA | 346,5945908 | 0,39973036 | 0,145999667 | 2,737885425 | 0,006183561 | 0,0196896 |
| ENSG00000234690.7 | EPCAM-DT | lncRNA | 78,30128274 | 0,955172351 | 0,279969639 | 3,411699766 | 0,000645592 | 0,002614072 |
| ENSG00000234699.1 | AL139339.1 | lncRNA | 17,36075824 | -2,672547834 | 0,681703231 | -3,920397781 | 8,84E-05 | 0,000425002 |
| ENSG00000234719.8 | NPIPB2 | protein_coding | 238,9121925 | 0,389850044 | 0,15866909 | 2,457000565 | 0,014010247 | 0,040055461 |
| ENSG00000234741.8 | GAS5 | lncRNA | 7304,661705 | -0,579805406 | 0,046099373 | -12,57729488 | 2,82E-36 | 1,13E-34 |
| ENSG00000234745.11 | HLA-B | protein_coding | 40135,78598 | 9,177767289 | 0,190621949 | 48,14643505 | 0 | 0 |
| ENSG00000234753.5 | FOXP4-AS1 | lncRNA | 52,49858535 | -1,231207041 | 0,361815456 | -3,402859162 | 0,000666846 | 0,002690937 |
| ENSG00000234771.3 | SLC25A25-AS1 | lncRNA | 1727,577703 | -0,206681565 | 0,069726665 | -2,964168206 | 0,003035023 | 0,01046818 |
| ENSG00000234851.4 | RPL23AP42 | processed_pseudog | 24498,32559 | -0,504735742 | 0,05115777 | -9,866257683 | 5,83E-23 | 1,38E-21 |
| ENSG00000234857.2 | HNRNPUL2-BSCL | protein_coding | 2777,140974 | 0,574256993 | 0,088712563 | 6,4732319 | 9,59E-11 | 1,00E-09 |
| ENSG00000235026.6 | DPP10-AS1 | lncRNA | 866,8078081 | -0,790266886 | 0,100707945 | -7,847115632 | 4,26E-15 | 6,45E-14 |
| ENSG00000235098.8 | ANKRD65 | protein_coding | 6170,176676 | -0,336135662 | 0,054753435 | -6,139078944 | 8,30E-10 | 7,84E-09 |
| ENSG00000235106.10 | BRD3OS | protein_coding | 4829,403174 | -0,39576202 | 0,053214681 | -7,437083422 | 1,03E-13 | 1,41E-12 |
| ENSG00000235142.10 | LINC02532 | lncRNA | 2074,84959 | 0,220168066 | 0,070969754 | 3,102280257 | 0,001920361 | 0,006968466 |
| ENSG00000235169.8 | SMIM1 | protein_coding | 120,2299569 | -0,534613278 | 0,220365422 | -2,426030698 | 0,015264979 | 0,043062072 |
| ENSG00000235173.7 | HGH1 | protein_coding | 1481,054176 | 0,225436462 | 0,07384895 | 3,05266983 | 0,002268153 | 0,008075759 |
| ENSG00000235192.1 | AC009495.3 | lncRNA | 163,4595574 | 1,332586148 | 0,205739324 | 6,477060991 | 9,35E-11 | 9,76E-10 |
| ENSG00000235217.6 | TSPY26P | transcribed_process | 3275,364597 | 0,17546374 | 0,054923631 | 3,194685743 | 0,001399832 | 0,005249719 |
| ENSG00000235257.9 | ITGA9-AS1 | lncRNA | 449,8537437 | -0,532129155 | 0,1159497 | -4,589310338 | 4,45E-06 | 2,63E-05 |
| ENSG00000235284.1 | SNORD62A | snoRNA | 34,31106599 | -1,18331628 | 0,410186752 | -2,884823255 | 0,003916335 | 0,013151242 |
| ENSG00000235314.1 | LINC00957 | lncRNA | 1502,787261 | -0,233214156 | 0,074080254 | -3,14812845 | 0,001643195 | 0,006068363 |
| ENSG00000235434.1 | AL391883.1 | lncRNA | 21,28836583 | -1,911693812 | 0,530960471 | -3,6004447 | 0,000317673 | 0,001373749 |
| ENSG00000235505.7 | CASP17P | transcribed_unproc | 44,65359356 | 2,166721769 | 0,398197613 | 5,441322851 | 5,29E-08 | 4,04E-07 |
| ENSG00000235535.8 | TRDN-AS1 | lncRNA | 14,52100969 | -2,042754695 | 0,707166483 | -2,888647503 | 0,003869025 | 0,013009843 |
| ENSG00000235560.4 | AC002310.1 | lncRNA | 499,9581352 | -0,645704163 | 0,108911754 | -5,928691226 | 3,05E-09 | 2,70E-08 |
| ENSG00000235568.6 | NFAM1 | protein_coding | 240,7306562 | 1,201253753 | 0,185598831 | 6,472313147 | 9,65E-11 | 1,01E-09 |
| ENSG00000235643.2 | LINC01647 | lncRNA | 63,12501721 | 1,707212094 | 0,301950907 | 5,653939279 | 1,57E-08 | 1,28E-07 |
| ENSG00000235750.10 | KIAA0040 | protein_coding | 2609,624643 | 1,488880059 | 0,081242113 | 18,32645651 | 5,09E-75 | 4,46E-73 |
| ENSG00000235772.1 | AP001625.2 | lncRNA | 144,373363 | -0,585899193 | 0,20548991 | -2,851230961 | 0,004355032 | 0,014432777 |
| ENSG00000235902.1 | AC108472.1 | lncRNA | 85,25683978 | 0,967988242 | 0,251007382 | 3,856413445 | 0,000115063 | 0,000541604 |
| ENSG00000235919.4 | ASH1L-AS1 | lncRNA | 268,9138868 | 0,488381718 | 0,149079839 | 3,275974271 | 0,001052982 | 0,004061268 |
| ENSG00000235933.1 | LINC01779 | lncRNA | 15,89110976 | 2,715681217 | 0,695627849 | 3,903928261 | 9,46E-05 | 0,000452269 |
| ENSG00000235954.7 | TTC28-AS1 | lncRNA | 3394,291507 | -0,236613199 | 0,052979181 | -4,466154338 | 7,96E-06 | 4,52E-05 |
| ENSG00000235961.5 | PNMA6A | protein_coding | 411,1246826 | 0,496705356 | 0,142509353 | 3,485422865 | 0,00049136 | 0,002039465 |
| ENSG00000236064.2 | AL109946.1 | lncRNA | 144,0249113 | -1,361167643 | 0,264828666 | -5,13980477 | 2,75E-07 | 1,93E-06 |
| ENSG00000236104.3 | ZBTB22 | protein_coding | 1444,661656 | 0,283975625 | 0,072707322 | 3,90573629 | 9,39E-05 | 0,000449218 |

| | | | | | | | | |
|---|---|---|---|---|---|---|---|---|
| ENSG00000236144.8 | TMEM147-AS1 | lncRNA | 723,3847412 | 0,444755315 | 0,101957058 | 4,362182704 | 1,29E-05 | 7,10E-05 |
| ENSG00000236154.1 | AL450311.1 | lncRNA | 24,17042029 | -1,258140733 | 0,507765976 | -2,47779645 | 0,013219654 | 0,038075648 |
| ENSG00000236155.6 | AL355877.1 | transcribed_unproc | 376,7859776 | -0,479321059 | 0,127896638 | -3,747722122 | 0,000178448 | 0,000807698 |
| ENSG00000236289.4 | GACAT3 | lncRNA | 7,738298527 | -3,104212673 | 1,008704231 | -3,077426047 | 0,002087966 | 0,007504077 |
| ENSG00000236373.2 | LINC02653 | lncRNA | 823,7910203 | -0,605490066 | 0,097743403 | -6,194689853 | 5,84E-10 | 5,62E-09 |
| ENSG00000236397.3 | DDX11L2 | unprocessed_pseud | 169,7182987 | -1,152617908 | 0,198779157 | -5,798484744 | 6,69E-09 | 5,70E-08 |
| ENSG00000236404.10 | VLDLR-AS1 | lncRNA | 308,9192199 | 0,424372383 | 0,163214579 | 2,600088697 | 0,009319967 | 0,028204239 |
| ENSG00000236417.2 | CTSLP1 | unprocessed_pseud | 172,2365942 | 1,068575677 | 0,187773901 | 5,690757199 | 1,26E-08 | 1,04E-07 |
| ENSG00000236423.6 | LINC01134 | lncRNA | 404,0647654 | -0,588641036 | 0,134543169 | -4,375109047 | 1,21E-05 | 6,72E-05 |
| ENSG00000236438.7 | FAM157A | transcribed_unproc | 780,3521636 | 0,692731242 | 0,146047168 | 4,743202139 | 2,10E-06 | 1,30E-05 |
| ENSG00000236502.1 | SIX3-AS1 | lncRNA | 161,3634841 | -0,846217219 | 0,194919684 | -4,341363579 | 1,42E-05 | 7,74E-05 |
| ENSG00000236617.2 | AC127070.1 | lncRNA | 32,77411209 | -1,480547524 | 0,436401906 | -3,392623871 | 0,000692266 | 0,002783036 |
| ENSG00000236618.2 | PITPNA-AS1 | lncRNA | 857,410918 | -0,807726081 | 0,092576385 | -8,724968935 | 2,66E-18 | 4,94E-17 |
| ENSG00000236753.7 | MKLN1-AS | lncRNA | 344,0927708 | 0,733104978 | 0,132742834 | 5,522746162 | 3,34E-08 | 2,62E-07 |
| ENSG00000236810.6 | ELOA-AS1 | lncRNA | 349,3251426 | -0,405361036 | 0,141990781 | -2,854840532 | 0,004305844 | 0,014303461 |
| ENSG00000236824.2 | BCYRN1 | scRNA | 297,0171123 | -1,176398721 | 0,148940093 | -7,89846907 | 2,82E-15 | 4,33E-14 |
| ENSG00000236901.6 | MIR600HG | lncRNA | 749,5161034 | -1,377081588 | 0,130529106 | -10,54999635 | 5,08E-26 | 1,38E-24 |
| ENSG00000236915.2 | CLCA4-AS1 | lncRNA | 18,07933188 | -1,517864461 | 0,571867389 | -2,654224547 | 0,007949091 | 0,024544045 |
| ENSG00000236936.1 | AL031005.1 | lncRNA | 17,89221488 | -1,916590417 | 0,6525261 | -2,937185833 | 0,003312056 | 0,011331671 |
| ENSG00000237070.1 | AC005550.2 | lncRNA | 52,14600536 | -0,807401968 | 0,31906377 | -2,530534785 | 0,011388879 | 0,033409232 |
| ENSG00000237149.5 | ZNF503-AS2 | lncRNA | 454,4979326 | 0,426872325 | 0,135846094 | 3,142323136 | 0,001676129 | 0,006178305 |
| ENSG00000237188.5 | AC242426.2 | lncRNA | 68,62713863 | 1,024225646 | 0,311625303 | 3,286721705 | 0,00101361 | 0,003922842 |
| ENSG00000237190.4 | CDKN2AIPNL | protein_coding | 3829,181316 | -0,458954689 | 0,054241983 | -8,461244722 | 2,65E-17 | 4,59E-16 |
| ENSG00000237276.8 | ANO7L1 | transcribed_unproc | 244,8255121 | 2,63662109 | 0,175114536 | 15,05655186 | 3,13E-51 | 1,86E-49 |
| ENSG00000237289.9 | CKMT1B | protein_coding | 4762,166907 | -0,195182375 | 0,060115326 | -3,246798931 | 0,001167108 | 0,004451862 |
| ENSG00000237296.9 | SMG1P1 | transcribed_unproc | 1191,88155 | 1,061946367 | 0,260088249 | 4,08302324 | 4,45E-05 | 0,000224754 |
| ENSG00000237298.10 | TTN-AS1 | lncRNA | 2081,484386 | 0,540425729 | 0,091266169 | 5,921424513 | 3,19E-09 | 2,82E-08 |
| ENSG00000237310.2 | AC008267.5 | lncRNA | 91,19898774 | -0,746820793 | 0,26190482 | -2,851496942 | 0,00435139 | 0,01442708 |
| ENSG00000237429.1 | BX293535.1 | lncRNA | 65,05236742 | -1,27709386 | 0,313537653 | -4,073175421 | 4,64E-05 | 0,000233793 |
| ENSG00000237440.9 | AC008554.1 | protein_coding | 1947,523854 | 0,218796821 | 0,092106664 | 2,37547222 | 0,017526512 | 0,048500646 |
| ENSG00000237441.10 | RGL2 | protein_coding | 4867,486096 | -0,280042074 | 0,060661082 | -4,616503134 | 3,90E-06 | 2,32E-05 |
| ENSG00000237457.6 | LINC01351 | lncRNA | 21,3871216 | 2,214957067 | 0,579718226 | 3,820747681 | 0,000133048 | 0,000618754 |
| ENSG00000237505.8 | PKN2-AS1 | lncRNA | 17,54726042 | 1,568579525 | 0,562302059 | 2,789567457 | 0,00527785 | 0,017104179 |
| ENSG00000237550.5 | AC243919.1 | transcribed_process | 18455,70066 | -0,697236239 | 0,042031509 | -16,58841806 | 8,45E-62 | 6,08E-60 |
| ENSG00000237560.7 | LINC01497 | lncRNA | 123,8908558 | -0,969263169 | 0,21544635 | -4,498860945 | 6,83E-06 | 3,92E-05 |
| ENSG00000237594.2 | AP000251.1 | lncRNA | 91,21865319 | -0,581944769 | 0,245920015 | -2,366398555 | 0,017962096 | 0,04952944 |

| | | | | | | | | |
|---|---|---|---|---|---|---|---|---|
| ENSG00000237649.8 | KIFC1 | protein_coding | 6038,614335 | -0,35735541 | 0,053279051 | -6,707240551 | 1,98E-11 | 2,21E-10 |
| ENSG00000237765.7 | FAM200B | protein_coding | 5081,649627 | -0,2183657 | 0,050427393 | -4,330299254 | 1,49E-05 | 8,11E-05 |
| ENSG00000237767.2 | LINC01370 | lncRNA | 1307,785708 | 0,26942052 | 0,097326848 | 2,768203496 | 0,005636625 | 0,018148303 |
| ENSG00000237943.7 | PRKCQ-AS1 | lncRNA | 475,644892 | 0,34289375 | 0,127646938 | 2,686266941 | 0,007225534 | 0,022566885 |
| ENSG00000237973.1 | MTCO1P12 | unprocessed_pseud | 58319,36314 | -0,579715439 | 0,058292559 | -9,944930339 | 2,65E-23 | 6,35E-22 |
| ENSG00000237976.1 | AL391069.2 | lncRNA | 85,10237138 | 0,637066446 | 0,256029411 | 2,488254939 | 0,012837167 | 0,037078378 |
| ENSG00000237984.3 | PTENP1 | transcribed_process | 311,4734927 | -0,418937693 | 0,176987185 | -2,367050992 | 0,017930462 | 0,049454328 |
| ENSG00000238035.8 | AC138035.1 | lncRNA | 235,8299833 | 0,938742717 | 0,195138254 | 4,810654488 | 1,50E-06 | 9,52E-06 |
| ENSG00000238083.7 | LRRC37A2 | protein_coding | 312,5093366 | 0,501370258 | 0,134971152 | 3,714647535 | 0,000203487 | 0,000910414 |
| ENSG00000238105.7 | GOLGA2P5 | transcribed_unproc | 2425,694934 | -0,212087297 | 0,073463892 | -2,886959716 | 0,00388984 | 0,013068122 |
| ENSG00000238142.2 | BX284668.5 | lncRNA | 243,8533727 | -0,564791863 | 0,169396485 | -3,334141583 | 0,000855631 | 0,003368794 |
| ENSG00000238164.6 | TNFRSF14-AS1 | lncRNA | 43,02877836 | 1,627731689 | 0,366963618 | 4,435675932 | 9,18E-06 | 5,18E-05 |
| ENSG00000238197.6 | PAXBP1-AS1 | lncRNA | 1576,989399 | 0,373406429 | 0,070695848 | 5,281872148 | 1,28E-07 | 9,33E-07 |
| ENSG00000238279.1 | BX470102.1 | lncRNA | 41,66596112 | 1,86572988 | 0,372624878 | 5,006992256 | 5,53E-07 | 3,73E-06 |
| ENSG00000239218.2 | RPS20P22 | transcribed_process | 46,12961472 | -0,924963871 | 0,35874415 | -2,578338551 | 0,009927668 | 0,029739403 |
| ENSG00000239306.4 | RBM14 | protein_coding | 6037,850301 | -0,157087082 | 0,052391655 | -2,998322547 | 0,002714702 | 0,009479557 |
| ENSG00000239322.1 | ATP6V1B1-AS1 | lncRNA | 427,9857751 | 0,543617685 | 0,121603508 | 4,470411186 | 7,81E-06 | 4,44E-05 |
| ENSG00000239445.6 | ST3GAL6-AS1 | lncRNA | 643,0669776 | -0,654038688 | 0,100297227 | -6,521004706 | 6,98E-11 | 7,38E-10 |
| ENSG00000239474.7 | KLHL41 | protein_coding | 378,1075629 | 1,047960832 | 0,137660376 | 7,612654136 | 2,69E-14 | 3,82E-13 |
| ENSG00000239482.6 | AC112487.1 | lncRNA | 2353,355872 | -0,212853379 | 0,07431373 | -2,864253745 | 0,00417993 | 0,013938674 |
| ENSG00000239511.2 | POM121L7P | processed_pseudog | 100,1447728 | -0,840315169 | 0,237159466 | -3,543249534 | 0,000395229 | 0,001672602 |
| ENSG00000239521.8 | CASTOR3 | protein_coding | 3696,582533 | 0,654629735 | 0,072407224 | 9,040945033 | 1,55E-19 | 3,07E-18 |
| ENSG00000239523.6 | MYLK-AS1 | lncRNA | 201,9200801 | -0,53549566 | 0,16787653 | -3,189818488 | 0,001423622 | 0,005330057 |
| ENSG00000239672.7 | NME1 | protein_coding | 5380,914542 | -0,184939994 | 0,055809501 | -3,31377259 | 0,000920463 | 0,003596998 |
| ENSG00000239713.9 | APOBEC3G | protein_coding | 2645,808188 | 4,046319706 | 0,073684552 | 54,91408458 | 0 | 0 |
| ENSG00000239779.7 | WBP1 | protein_coding | 2892,487424 | -0,48346281 | 0,056228992 | -8,598105662 | 8,10E-18 | 1,46E-16 |
| ENSG00000239789.6 | MRPS17 | protein_coding | 1802,11594 | -0,197681824 | 0,076943726 | -2,569174049 | 0,010194124 | 0,030416164 |
| ENSG00000239911.2 | PRKAG2-AS1 | lncRNA | 220,4662131 | -1,238980589 | 0,165704025 | -7,477069948 | 7,60E-14 | 1,05E-12 |
| ENSG00000240005.6 | STIM2-AS1 | lncRNA | 63,00582848 | -1,582341652 | 0,330459974 | -4,788300483 | 1,68E-06 | 1,06E-05 |
| ENSG00000240024.5 | LINC00888 | transcribed_unproc | 2705,935616 | 0,208425778 | 0,061555726 | 3,385968997 | 0,000709274 | 0,002843792 |
| ENSG00000240053.8 | LY6G5B | protein_coding | 416,5948006 | -0,347214733 | 0,134399455 | -2,583453431 | 0,009781667 | 0,02936457 |
| ENSG00000240065.8 | PSMB9 | protein_coding | 5824,831126 | 9,337548017 | 0,278647691 | 33,51022927 | 3,42E-246 | 8,02E-244 |
| ENSG00000240086.7 | AC092969.1 | lncRNA | 1250,850945 | -0,706710572 | 0,072796019 | -9,708093649 | 2,78E-22 | 6,37E-21 |
| ENSG00000240184.7 | PCDHGC3 | protein_coding | 548,76128 | -0,881631577 | 0,343987396 | -2,562976395 | 0,01037791 | 0,030882653 |
| ENSG00000240204.3 | SMKR1 | protein_coding | 216,8376083 | -0,540389434 | 0,169058662 | -3,196461091 | 0,001391246 | 0,00522187 |
| ENSG00000240342.3 | RPS2P5 | processed_pseudog | 5009,285434 | -0,587909869 | 0,058824956 | -9,994225321 | 1,62E-23 | 3,91E-22 |

| | | | | | | | | |
|---|---|---|---|---|---|---|---|---|
| ENSG00000240356.6 | RPL23AP7 | transcribed_process | 584,7788589 | -0,253012567 | 0,105608875 | -2,395750992 | 0,016586354 | 0,046277251 |
| ENSG00000240429.1 | LRRFIP1P1 | processed_pseudog | 456,636557 | 0,619615292 | 0,115313789 | 5,373297503 | 7,73E-08 | 5,77E-07 |
| ENSG00000240498.9 | CDKN2B-AS1 | lncRNA | 1603,357203 | 0,187667347 | 0,074116174 | 2,532070092 | 0,011339132 | 0,033307129 |
| ENSG00000240654.6 | C1QTNF9 | protein_coding | 8,521061859 | 1,915594521 | 0,80695107 | 2,373867007 | 0,017602889 | 0,048664141 |
| ENSG00000240694.9 | PNMA2 | protein_coding | 34344,98121 | -0,162718186 | 0,052410919 | -3,104661942 | 0,001904966 | 0,006919297 |
| ENSG00000240801.1 | AC132217.1 | lncRNA | 129,594548 | -0,777614214 | 0,278114104 | -2,796025819 | 0,005173527 | 0,016830599 |
| ENSG00000240875.6 | LINC00886 | lncRNA | 475,4684372 | 0,545977256 | 0,134101283 | 4,071379823 | 4,67E-05 | 0,000235446 |
| ENSG00000240891.7 | PLCXD2 | protein_coding | 2255,059321 | -0,492360577 | 0,081295659 | -6,056419047 | 1,39E-09 | 1,28E-08 |
| ENSG00000241015.2 | TPM3P9 | transcribed_process | 3352,588848 | 0,549852663 | 0,059845181 | 9,187918809 | 4,01E-20 | 8,17E-19 |
| ENSG00000241343.9 | RPL36A | protein_coding | 16061,71638 | -0,569142412 | 0,045415144 | -12,53199621 | 4,99E-36 | 1,99E-34 |
| ENSG00000241360.2 | PDXP | protein_coding | 4574,369597 | -0,140338537 | 0,050682021 | -2,769000428 | 0,005622856 | 0,018106562 |
| ENSG00000241370.5 | RPP21 | protein_coding | 1507,123752 | -0,267043966 | 0,083607822 | -3,194006986 | 0,001403127 | 0,005260325 |
| ENSG00000241388.5 | HNF1A-AS1 | lncRNA | 1870,010035 | 0,310733274 | 0,074845026 | 4,151689023 | 3,30E-05 | 0,000170187 |
| ENSG00000241399.7 | CD302 | protein_coding | 8106,182083 | -0,424701684 | 0,050772783 | -8,364750966 | 6,02E-17 | 1,02E-15 |
| ENSG00000241404.7 | EGFL8 | protein_coding | 275,4925497 | -0,674611447 | 0,178725702 | -3,77456313 | 0,000160288 | 0,000732819 |
| ENSG00000241468.7 | ATP5MF | protein_coding | 19999,21459 | -0,121686954 | 0,045331423 | -2,684384184 | 0,007266355 | 0,022678646 |
| ENSG00000241472.7 | PTPRG-AS1 | lncRNA | 2963,874517 | 0,322488171 | 0,070127193 | 4,598617978 | 4,25E-06 | 2,52E-05 |
| ENSG00000241484.9 | ARHGAP8 | protein_coding | 39,69160319 | 1,175800652 | 0,414962534 | 2,833510391 | 0,004603981 | 0,015152948 |
| ENSG00000241489.8 | AC244197.3 | protein_coding | 1238,511466 | -0,260284312 | 0,092089321 | -2,826433185 | 0,004706956 | 0,015462475 |
| ENSG00000241553.12 | ARPC4 | protein_coding | 14645,05935 | -0,121948084 | 0,043163446 | -2,825262934 | 0,004724183 | 0,015505488 |
| ENSG00000241685.10 | ARPC1A | protein_coding | 13016,60651 | -0,48670095 | 0,049791432 | -9,774793159 | 1,44E-22 | 3,36E-21 |
| ENSG00000241697.5 | TMEFF1 | protein_coding | 1404,958461 | -0,271340618 | 0,100974078 | -2,687230446 | 0,007204723 | 0,022508134 |
| ENSG00000241754.2 | AC063938.1 | lncRNA | 25,59694293 | -1,257281514 | 0,492229687 | -2,554257794 | 0,010641444 | 0,031562558 |
| ENSG00000241837.7 | ATP5PO | protein_coding | 13294,49596 | -0,357929718 | 0,042069221 | -8,508113791 | 1,77E-17 | 3,12E-16 |
| ENSG00000241860.7 | AL627309.5 | lncRNA | 950,3400124 | 0,493790274 | 0,112009801 | 4,408455935 | 1,04E-05 | 5,82E-05 |
| ENSG00000241973.10 | PI4KA | protein_coding | 23424,11716 | -0,121307736 | 0,047472189 | -2,555343187 | 0,010608315 | 0,031472592 |
| ENSG00000242082.2 | SLC5A4-AS1 | lncRNA | 1491,858324 | 0,306640873 | 0,070239359 | 4,365655927 | 1,27E-05 | 6,99E-05 |
| ENSG00000242086.8 | MUC20-OT1 | lncRNA | 3010,84077 | 0,307929283 | 0,070106568 | 4,392302899 | 1,12E-05 | 6,24E-05 |
| ENSG00000242125.3 | SNHG3 | lncRNA | 2405,461636 | -0,809606703 | 0,083029511 | -9,750830655 | 1,83E-22 | 4,22E-21 |
| ENSG00000242265.5 | PEG10 | protein_coding | 44737,29462 | -0,156537401 | 0,044722988 | -3,500155231 | 0,000464987 | 0,001943982 |
| ENSG00000242294.6 | STAG3L5P | transcribed_unproc | 4377,79976 | 0,265466748 | 0,073274692 | 3,622898167 | 0,000291321 | 0,001268343 |
| ENSG00000242479.1 | AC109992.1 | processed_pseudog | 15,02810514 | 2,26578697 | 0,661995489 | 3,422662248 | 0,000620111 | 0,00251769 |
| ENSG00000242498.8 | ARPIN | protein_coding | 3420,608445 | -0,349416638 | 0,073187968 | -4,774236081 | 1,80E-06 | 1,13E-05 |
| ENSG00000242588.6 | AC108010.1 | lncRNA | 902,6264929 | 1,150289689 | 0,101918799 | 11,28633479 | 1,53E-29 | 4,72E-28 |
| ENSG00000242615.1 | AC022415.1 | transcribed_process | 293,0030479 | 0,602146853 | 0,161234816 | 3,734595703 | 0,000188017 | 0,000847432 |
| ENSG00000242616.4 | GNG10 | protein_coding | 8056,131207 | -0,394153154 | 0,058915047 | -6,690195012 | 2,23E-11 | 2,47E-10 |

| | | | | | | | | |
|---|---|---|---|---|---|---|---|---|
| ENSG00000242641.6 | LINC00971 | lncRNA | 17,19017963 | 3,133961202 | 0,677090939 | 4,628567626 | 3,68E-06 | 2,20E-05 |
| ENSG00000242689.3 | CNTF | protein_coding | 116,0409151 | -0,930513954 | 0,237181016 | -3,923222732 | 8,74E-05 | 0,000420587 |
| ENSG00000242732.4 | RTL5 | protein_coding | 9975,5745 | 0,123921868 | 0,047374091 | 2,615815228 | 0,008901471 | 0,027101747 |
| ENSG00000242735.1 | RPSAP26 | processed_pseudog | 7,885425685 | -2,728216498 | 1,119104104 | -2,437857647 | 0,014774591 | 0,041881363 |
| ENSG00000242802.9 | AP5Z1 | protein_coding | 1196,468909 | 0,301387175 | 0,098137821 | 3,07106038 | 0,002133 | 0,007648829 |
| ENSG00000243064.8 | ABCC13 | transcribed_unproc | 147,9491441 | 0,684608977 | 0,20371067 | 3,360692776 | 0,000777473 | 0,003089167 |
| ENSG00000243147.8 | MRPL33 | protein_coding | 8618,34308 | -0,420744552 | 0,053246436 | -7,90183506 | 2,75E-15 | 4,22E-14 |
| ENSG00000243156.8 | MICAL3 | protein_coding | 5097,332099 | 0,25220987 | 0,056704573 | 4,447787166 | 8,68E-06 | 4,90E-05 |
| ENSG00000243199.1 | AC115223.1 | processed_pseudog | 442,0598941 | -0,871916161 | 0,131280783 | -6,641613045 | 3,10E-11 | 3,39E-10 |
| ENSG00000243232.5 | PCDHAC2 | protein_coding | 2690,257289 | 1,344865775 | 0,07351015 | 18,29496715 | 9,08E-75 | 7,92E-73 |
| ENSG00000243244.6 | STON1 | protein_coding | 104,7744816 | -0,797188205 | 0,239987108 | -3,321795955 | 0,000894401 | 0,003503666 |
| ENSG00000243302.3 | AC018638.4 | processed_pseudog | 481,2423872 | 1,187433671 | 0,128622689 | 9,231914532 | 2,66E-20 | 5,48E-19 |
| ENSG00000243317.8 | STMP1 | protein_coding | 10979,5027 | -0,41970661 | 0,04863155 | -8,630335899 | 6,12E-18 | 1,11E-16 |
| ENSG00000243335.9 | KCTD7 | protein_coding | 2293,266425 | -0,531251628 | 0,069044148 | -7,694375892 | 1,42E-14 | 2,07E-13 |
| ENSG00000243364.8 | EFNA4 | protein_coding | 294,2022581 | -2,00214732 | 0,155529348 | -12,87311588 | 6,38E-38 | 2,68E-36 |
| ENSG00000243414.5 | TICAM2 | protein_coding | 206,9901071 | 0,477002427 | 0,179973586 | 2,650402418 | 0,008039595 | 0,024792881 |
| ENSG00000243422.2 | RPL23AP49 | transcribed_process | 36,17754654 | -0,942525348 | 0,398441701 | -2,365528875 | 0,018004339 | 0,049633763 |
| ENSG00000243444.8 | PALM2 | protein_coding | 590,5584551 | -0,363412601 | 0,120580408 | -3,013861114 | 0,002579459 | 0,009059297 |
| ENSG00000243646.10 | IL10RB | protein_coding | 2941,712517 | 0,212448047 | 0,065188137 | 3,258998588 | 0,001118062 | 0,004287302 |
| ENSG00000243649.8 | CFB | protein_coding | 1030,506038 | 6,039843995 | 0,570784197 | 10,58165946 | 3,62E-26 | 9,87E-25 |
| ENSG00000243660.10 | ZNF487 | protein_coding | 476,7020362 | -0,384379508 | 0,131753439 | -2,917415372 | 0,003529454 | 0,01199162 |
| ENSG00000243667.7 | WDR92 | protein_coding | 1056,245309 | -0,219256956 | 0,089988328 | -2,436504395 | 0,014829989 | 0,041993739 |
| ENSG00000243678.11 | NME2 | protein_coding | 10298,00881 | -0,221618288 | 0,048135766 | -4,604025396 | 4,14E-06 | 2,46E-05 |
| ENSG00000243679.1 | AC018638.5 | processed_pseudog | 260,7321732 | 1,175526867 | 0,151693612 | 7,749349817 | 9,24E-15 | 1,36E-13 |
| ENSG00000243696.5 | AC006254.1 | protein_coding | 138,3723701 | 0,594280913 | 0,224289785 | 2,649612033 | 0,008058425 | 0,024844143 |
| ENSG00000243701.7 | DUBR | lncRNA | 464,5323944 | 0,442752099 | 0,142022929 | 3,11746914 | 0,00182411 | 0,006661081 |
| ENSG00000243716.10 | NPIPB5 | protein_coding | 4765,50087 | 0,415055617 | 0,080784188 | 5,137832346 | 2,78E-07 | 1,95E-06 |
| ENSG00000243749.1 | TMEM35B | protein_coding | 977,7775127 | 0,282353681 | 0,091612748 | 3,082034845 | 0,002055908 | 0,007403044 |
| ENSG00000243753.5 | HLA-L | transcribed_unproc | 199,8336579 | 3,608690771 | 0,213379598 | 16,91207037 | 3,67E-64 | 2,71E-62 |
| ENSG00000243811.10 | APOBEC3D | protein_coding | 307,6197359 | 3,213154517 | 0,164235148 | 19,56435371 | 3,11E-85 | 3,10E-83 |
| ENSG00000243896.4 | OR2A7 | protein_coding | 555,207651 | 0,77378773 | 0,106262013 | 7,281884704 | 3,29E-13 | 4,33E-12 |
| ENSG00000243927.6 | MRPS6 | protein_coding | 3358,272251 | -0,156492124 | 0,056589288 | -2,765401894 | 0,00568527 | 0,018286618 |
| ENSG00000243989.9 | ACY1 | protein_coding | 1607,199021 | -0,202226899 | 0,07142185 | -2,831443012 | 0,004633849 | 0,015244564 |
| ENSG00000244005.13 | NFS1 | protein_coding | 3735,015874 | -0,32722763 | 0,070221765 | -4,65991749 | 3,16E-06 | 1,91E-05 |
| ENSG00000244055.2 | AC007566.1 | lncRNA | 388,6110023 | 0,476556382 | 0,134426696 | 3,545102231 | 0,000392461 | 0,001663322 |
| ENSG00000244165.2 | P2RY11 | protein_coding | 367,0718591 | 0,533734947 | 0,149359794 | 3,573484756 | 0,000352262 | 0,001508573 |

| | | | | | | | | |
|---|---|---|---|---|---|---|---|---|
| ENSG00000244198.7 | AC004889.1 | lncRNA | 3187,757022 | 0,49209678 | 0,067318293 | 7,310000906 | 2,67E-13 | 3,54E-12 |
| ENSG00000244257.5 | PKD1P1 | unprocessed_pseud | 830,767762 | 1,100682639 | 0,277881765 | 3,960974695 | 7,46E-05 | 0,000363595 |
| ENSG00000244457.2 | ENO1P1 | transcribed_process | 2506,624333 | 0,333911199 | 0,067177936 | 4,97054862 | 6,68E-07 | 4,45E-06 |
| ENSG00000244503.1 | AC108751.5 | processed_pseudog | 79,6300252 | 1,161356634 | 0,277403767 | 4,186520773 | 2,83E-05 | 0,000147455 |
| ENSG00000244509.4 | APOBEC3C | protein_coding | 62,26486856 | 1,421080598 | 0,30162586 | 4,711401728 | 2,46E-06 | 1,51E-05 |
| ENSG00000244687.11 | UBE2V1 | protein_coding | 25009,97496 | -0,278272711 | 0,04740711 | -5,869851866 | 4,36E-09 | 3,79E-08 |
| ENSG00000244693.1 | CTAGE8 | protein_coding | 304,1358926 | 0,689104056 | 0,139492954 | 4,940063544 | 7,81E-07 | 5,15E-06 |
| ENSG00000244694.7 | PTCHD4 | protein_coding | 228,5514781 | 0,543620643 | 0,16564388 | 3,281863733 | 0,001031234 | 0,00398285 |
| ENSG00000244731.8 | C4A | protein_coding | 2496,852534 | 0,368274031 | 0,07444828 | 4,946709733 | 7,55E-07 | 4,99E-06 |
| ENSG00000244754.8 | N4BP2L2 | protein_coding | 10972,96695 | 0,324503173 | 0,054456601 | 5,958931843 | 2,54E-09 | 2,26E-08 |
| ENSG00000244879.8 | GABPB1-AS1 | lncRNA | 2647,605971 | 0,158659072 | 0,061093381 | 2,596992826 | 0,009404389 | 0,028425324 |
| ENSG00000245017.2 | LINC02453 | lncRNA | 70,58863798 | -1,721363324 | 0,342170799 | -5,030713692 | 4,89E-07 | 3,32E-06 |
| ENSG00000245025.3 | AC107959.1 | lncRNA | 195,9914865 | 0,992727379 | 0,251304301 | 3,950300003 | 7,81E-05 | 0,000378561 |
| ENSG00000245060.8 | LINC00847 | lncRNA | 1761,221962 | 0,192694851 | 0,064276199 | 2,997919191 | 0,002718297 | 0,009489168 |
| ENSG00000245275.8 | SAP30L-AS1 | lncRNA | 142,1403224 | 0,636849087 | 0,234248386 | 2,718691465 | 0,006554071 | 0,020731533 |
| ENSG00000245281.8 | AC124242.1 | lncRNA | 171,5505491 | -0,4882968 | 0,19734672 | -2,474309182 | 0,013349411 | 0,03839043 |
| ENSG00000245293.3 | AC096564.1 | lncRNA | 2106,337479 | -0,316880173 | 0,067908492 | -4,666281981 | 3,07E-06 | 1,86E-05 |
| ENSG00000245322.7 | AC097460.1 | lncRNA | 42,34015166 | -0,901484007 | 0,360871509 | -2,498074757 | 0,012486986 | 0,036225446 |
| ENSG00000245532.9 | NEAT1 | lncRNA | 518,0155564 | 0,845069715 | 0,174439764 | 4,844478661 | 1,27E-06 | 8,13E-06 |
| ENSG00000245552.7 | AP000787.1 | lncRNA | 388,4786155 | -0,434215983 | 0,126281969 | -3,438463839 | 0,000585025 | 0,002390933 |
| ENSG00000245556.3 | SCAMP1-AS1 | lncRNA | 3428,265768 | 1,625482306 | 0,060664788 | 26,79449426 | 3,75E-158 | 6,34E-156 |
| ENSG00000245571.7 | FAM111A-DT | lncRNA | 666,9413717 | 0,500871799 | 0,11169629 | 4,484229486 | 7,32E-06 | 4,18E-05 |
| ENSG00000245614.4 | DDX11-AS1 | lncRNA | 255,6566081 | -0,651241616 | 0,207183655 | -3,143305946 | 0,001670511 | 0,006162167 |
| ENSG00000245662.4 | LINC02211 | lncRNA | 403,6397146 | -0,355240752 | 0,140006179 | -2,537321962 | 0,011170417 | 0,032893108 |
| ENSG00000245694.10 | CRNDE | lncRNA | 2104,742708 | -0,969646613 | 0,082170721 | -11,80039074 | 3,89E-32 | 1,38E-30 |
| ENSG00000245750.9 | DRAIC | lncRNA | 1275,648612 | -0,693605513 | 0,090598845 | -7,655787622 | 1,92E-14 | 2,77E-13 |
| ENSG00000245849.8 | RAD51-AS1 | lncRNA | 638,0680757 | -0,279122077 | 0,105935116 | -2,634839953 | 0,008417696 | 0,025796314 |
| ENSG00000245910.8 | SNHG6 | lncRNA | 3665,58843 | -0,406199887 | 0,065827608 | -6,170661461 | 6,80E-10 | 6,48E-09 |
| ENSG00000246082.2 | NUDT16P1 | transcribed_unitary | 1227,98363 | 0,263437028 | 0,076261128 | 3,454407705 | 0,000551503 | 0,002264875 |
| ENSG00000246095.2 | LINC01096 | lncRNA | 274,5165949 | 0,351499572 | 0,144645308 | 2,430079323 | 0,015095519 | 0,042649056 |
| ENSG00000246100.4 | LINC00900 | lncRNA | 1195,073587 | 0,396252714 | 0,083809073 | 4,728040766 | 2,27E-06 | 1,40E-05 |
| ENSG00000246283.2 | AC090510.1 | lncRNA | 44,17051146 | -2,244090769 | 0,39783927 | -5,640696976 | 1,69E-08 | 1,38E-07 |
| ENSG00000246375.3 | PPM1K-DT | lncRNA | 26,51450631 | 3,830987625 | 0,620711418 | 6,171930325 | 6,75E-10 | 6,43E-09 |
| ENSG00000246528.3 | AC079089.1 | lncRNA | 122,4052164 | 1,268842394 | 0,230853507 | 5,496309801 | 3,88E-08 | 3,01E-07 |
| ENSG00000246582.3 | AC100861.1 | lncRNA | 24,09974116 | 1,217801454 | 0,483868944 | 2,516800198 | 0,011842597 | 0,034582902 |
| ENSG00000246695.8 | RASSF8-AS1 | lncRNA | 1030,662943 | -0,308180113 | 0,110896091 | -2,778998879 | 0,005452671 | 0,017613975 |

| | | | | | | | | |
|---|---|---|---|---|---|---|---|---|
| ENSG00000246705.4 | H2AFJ | protein_coding | 7601,985801 | 0,884196263 | 0,257268028 | 3,436868038 | 0,000588482 | 0,002402711 |
| ENSG00000247092.7 | SNHG10 | lncRNA | 486,1345619 | -0,607124027 | 0,124662856 | -4,870127685 | 1,12E-06 | 7,20E-06 |
| ENSG00000247121.7 | AC009126.1 | lncRNA | 293,1638504 | 0,601388196 | 0,143192632 | 4,199854333 | 2,67E-05 | 0,000139551 |
| ENSG00000247157.7 | LINC01252 | lncRNA | 159,3643955 | 0,740737882 | 0,189647849 | 3,905859655 | 9,39E-05 | 0,000449085 |
| ENSG00000247271.7 | ZBED5-AS1 | lncRNA | 1404,282184 | -0,202377097 | 0,073639446 | -2,748215897 | 0,005992054 | 0,019136917 |
| ENSG00000247287.3 | AL359220.1 | lncRNA | 147,6653218 | 0,638464328 | 0,189344396 | 3,371973724 | 0,000746316 | 0,002978917 |
| ENSG00000247400.4 | DNAJC3-DT | lncRNA | 1669,087408 | 0,395041679 | 0,066373395 | 5,951807633 | 2,65E-09 | 2,36E-08 |
| ENSG00000247556.6 | OIP5-AS1 | lncRNA | 12661,60253 | -0,366242283 | 0,04562597 | -8,027057499 | 9,98E-16 | 1,58E-14 |
| ENSG00000247627.2 | MTND4P12 | processed_pseudog | 805,0049663 | -0,838902423 | 0,111972054 | -7,492069594 | 6,78E-14 | 9,44E-13 |
| ENSG00000247765.2 | AC068446.2 | lncRNA | 137,3115037 | 0,752760004 | 0,217795978 | 3,456262184 | 0,000547722 | 0,002251403 |
| ENSG00000247934.4 | AC022364.1 | lncRNA | 316,1448723 | 0,391433007 | 0,159951009 | 2,447205608 | 0,014396868 | 0,040983826 |
| ENSG00000248008.3 | NRAV | lncRNA | 1517,784968 | -0,180682855 | 0,070754543 | -2,553657295 | 0,010659812 | 0,031604545 |
| ENSG00000248049.7 | UBA6-AS1 | lncRNA | 2435,650835 | 0,347598065 | 0,061194494 | 5,68021796 | 1,35E-08 | 1,10E-07 |
| ENSG00000248098.12 | BCKDHA | protein_coding | 2110,472799 | -0,566929621 | 0,067615606 | -8,384597241 | 5,09E-17 | 8,70E-16 |
| ENSG00000248206.2 | AC074194.1 | lncRNA | 14,48218806 | -2,598910211 | 0,785352429 | -3,309227954 | 0,000935536 | 0,00365083 |
| ENSG00000248275.2 | TRIM52-AS1 | lncRNA | 411,8707263 | -0,477353984 | 0,128854526 | -3,704596164 | 0,000211728 | 0,000945216 |
| ENSG00000248334.6 | WHAMMP2 | transcribed_unproc | 877,1066134 | 0,749200163 | 0,089472295 | 8,373543608 | 5,59E-17 | 9,53E-16 |
| ENSG00000248360.7 | LINC00504 | lncRNA | 757,4376186 | 0,247450381 | 0,090919527 | 2,721641766 | 0,006495852 | 0,020561823 |
| ENSG00000248431.2 | AC021134.1 | lncRNA | 40,03961526 | 2,151619994 | 0,391309053 | 5,498518308 | 3,83E-08 | 2,98E-07 |
| ENSG00000248498.4 | ASNSP1 | transcribed_unproc | 11,11470174 | 3,792163583 | 0,926434638 | 4,093287778 | 4,25E-05 | 0,00021595 |
| ENSG00000248527.1 | MTATP6P1 | unprocessed_pseud | 15699,84373 | -1,275509233 | 0,058126306 | -21,94375192 | 9,94E-107 | 1,20E-104 |
| ENSG00000248685.6 | LINC02484 | lncRNA | 188,8970422 | 1,849193277 | 0,192100739 | 9,626164329 | 6,20E-22 | 1,40E-20 |
| ENSG00000248693.1 | LINC02100 | lncRNA | 11,34382707 | 2,185762723 | 0,721623298 | 3,028952544 | 0,002454032 | 0,008657964 |
| ENSG00000248698.6 | LINC01085 | transcribed_process | 12,97581721 | 1,700352299 | 0,647903942 | 2,624389494 | 0,008680446 | 0,026522078 |
| ENSG00000248710.1 | AC079594.2 | protein_coding | 349,2288078 | -0,424139599 | 0,15430391 | -2,748728783 | 0,005982687 | 0,019117589 |
| ENSG00000248791.1 | AC010627.1 | processed_pseudog | 61,27544667 | -0,991054443 | 0,329433548 | -3,008359198 | 0,002626625 | 0,009212013 |
| ENSG00000248810.2 | LINC02432 | lncRNA | 8289,725799 | 0,246206189 | 0,052997286 | 4,645637685 | 3,39E-06 | 2,04E-05 |
| ENSG00000248859.2 | LINC01574 | lncRNA | 239,0154567 | -0,476895607 | 0,178714295 | -2,668480472 | 0,007619521 | 0,023633429 |
| ENSG00000248905.8 | FMN1 | protein_coding | 1070,297311 | 0,988209353 | 0,084022366 | 11,76126545 | 6,18E-32 | 2,16E-30 |
| ENSG00000248919.7 | ATP5MF-PTCD1 | protein_coding | 1009,177428 | 0,435207979 | 0,090608224 | 4,803184076 | 1,56E-06 | 9,86E-06 |
| ENSG00000248925.1 | AC021087.1 | lncRNA | 40,68945966 | -1,591994434 | 0,448900493 | -3,546430578 | 0,000390488 | 0,001656206 |
| ENSG00000248932.6 | AC046134.2 | lncRNA | 808,994795 | -0,267044068 | 0,099288027 | -2,689589848 | 0,007153989 | 0,022377589 |
| ENSG00000248971.2 | KRT8P46 | processed_pseudog | 96,52825815 | 0,690818486 | 0,272736972 | 2,532911036 | 0,011311966 | 0,033249026 |
| ENSG00000249042.6 | AC008771.1 | lncRNA | 943,8690285 | -0,265299878 | 0,089779528 | -2,9550153 | 0,003126535 | 0,010741024 |
| ENSG00000249115.9 | HAUS5 | protein_coding | 1449,301377 | -0,361286159 | 0,080120262 | -4,509298302 | 6,50E-06 | 3,75E-05 |
| ENSG00000249158.6 | PCDHA11 | protein_coding | 2229,388836 | -0,539693253 | 0,076225448 | -7,080224061 | 1,44E-12 | 1,80E-11 |

| | | | | | | | | |
|---|---|---|---|---|---|---|---|---|
| ENSG00000249201.2 | AC114291.1 | lncRNA | 665,7820835 | -0,359917509 | 0,117169926 | -3,071756721 | 0,002128031 | 0,007632225 |
| ENSG00000249240.2 | AC069368.1 | protein_coding | 15,93670929 | -1,791508668 | 0,633884067 | -2,826240256 | 0,004709792 | 0,015469534 |
| ENSG00000249242.8 | TMEM150C | protein_coding | 1670,966187 | 0,204718295 | 0,067413365 | 3,036761265 | 0,002391348 | 0,00846334 |
| ENSG00000249319.2 | AC068533.4 | protein_coding | 23,58966652 | -1,982023373 | 0,650372491 | -3,047520304 | 0,002307379 | 0,008199685 |
| ENSG00000249348.1 | UGDH-AS1 | lncRNA | 1170,422717 | 0,27803644 | 0,076162755 | 3,650556491 | 0,000261673 | 0,001149043 |
| ENSG00000249353.2 | NPM1P27 | processed_pseudog | 13,49666294 | -1,522282785 | 0,639147508 | -2,381739372 | 0,017231088 | 0,04781834 |
| ENSG00000249395.3 | CASC9 | lncRNA | 694,5906675 | -0,336701082 | 0,092524857 | -3,639033795 | 0,000273663 | 0,001197023 |
| ENSG00000249437.7 | NAIP | protein_coding | 1916,559414 | 0,183520156 | 0,076254235 | 2,406688059 | 0,016097913 | 0,04511304 |
| ENSG00000249464.6 | LINC01091 | lncRNA | 629,8374874 | -0,253356638 | 0,104516233 | -2,424088876 | 0,015346849 | 0,043261393 |
| ENSG00000249471.8 | ZNF324B | protein_coding | 994,8539889 | 0,245596224 | 0,098206191 | 2,500822205 | 0,012390537 | 0,03600085 |
| ENSG00000249604.1 | AC096564.2 | lncRNA | 23,05040402 | 1,219925944 | 0,496352393 | 2,457781934 | 0,013979803 | 0,039988739 |
| ENSG00000249624.9 | AP000295.1 | protein_coding | 67,06345669 | 4,276511253 | 1,416799754 | 3,018430262 | 0,002540879 | 0,008934954 |
| ENSG00000249673.7 | NOP14-AS1 | lncRNA | 1365,408077 | 0,225246522 | 0,079022599 | 2,850406415 | 0,00436634 | 0,014461734 |
| ENSG00000249709.8 | ZNF564 | protein_coding | 1987,84766 | 0,469824742 | 0,091187181 | 5,152311265 | 2,57E-07 | 1,81E-06 |
| ENSG00000249715.12 | FER1L5 | protein_coding | 703,9391879 | -1,424280248 | 0,122548669 | -11,62216004 | 3,18E-31 | 1,07E-29 |
| ENSG00000249738.10 | AC008691.1 | lncRNA | 13,62482038 | 1,858615472 | 0,639639946 | 2,905721387 | 0,003664075 | 0,012400342 |
| ENSG00000249746.1 | AC099509.1 | lncRNA | 27,757888 | 1,522824242 | 0,509187345 | 2,990695385 | 0,00278343 | 0,009692485 |
| ENSG00000249790.3 | AC092490.1 | lncRNA | 134,556702 | 1,251292568 | 0,26693974 | 4,687546958 | 2,76E-06 | 1,69E-05 |
| ENSG00000249959.6 | AC024587.2 | lncRNA | 57,48334688 | -1,014361716 | 0,34032739 | -2,980546805 | 0,002877343 | 0,009985523 |
| ENSG00000250012.2 | AC079848.1 | lncRNA | 368,313884 | -0,640049217 | 0,203742806 | -3,141456769 | 0,001681096 | 0,006193093 |
| ENSG00000250056.7 | LINC01018 | lncRNA | 1566,319237 | 0,292261331 | 0,07347987 | 3,977433995 | 6,97E-05 | 0,000340984 |
| ENSG00000250159.7 | AC106791.1 | lncRNA | 1486,423706 | -0,464896174 | 0,075613515 | -6,148321141 | 7,83E-10 | 7,42E-09 |
| ENSG00000250166.3 | AC053513.1 | lncRNA | 2516,686022 | 1,192213829 | 0,068555272 | 17,39054913 | 9,73E-68 | 7,63E-66 |
| ENSG00000250182.3 | EEF1A1P13 | processed_pseudog | 23,51378382 | -1,268116466 | 0,534528101 | -2,372403741 | 0,017672767 | 0,048833332 |
| ENSG00000250251.6 | PKD1P6 | transcribed_unproc | 2750,559762 | 1,32912321 | 0,088296261 | 15,05299543 | 3,30E-51 | 1,95E-49 |
| ENSG00000250264.1 | AL669918.1 | protein_coding | 4882,0832 | 4,638423968 | 0,085592248 | 54,1921036 | 0 | 0 |
| ENSG00000250286.2 | AC021491.2 | lncRNA | 12,27042078 | 1,958570622 | 0,680706105 | 2,877263195 | 0,004011409 | 0,013434429 |
| ENSG00000250312.8 | ZNF718 | protein_coding | 2474,857092 | -0,228961491 | 0,060074961 | -3,811263222 | 0,000138258 | 0,000640605 |
| ENSG00000250317.8 | SMIM20 | protein_coding | 2587,783293 | -0,141191428 | 0,059201542 | -2,38492821 | 0,017082455 | 0,047476104 |
| ENSG00000250349.3 | AF241726.2 | protein_coding | 20,64567705 | -2,144676716 | 0,810602712 | -2,645780336 | 0,008150272 | 0,02507579 |
| ENSG00000250360.1 | AC008808.2 | lncRNA | 186,7574136 | 1,561207695 | 0,196544959 | 7,943259912 | 1,97E-15 | 3,05E-14 |
| ENSG00000250366.2 | TUNAR | protein_coding | 4621,688185 | -0,268299639 | 0,055521527 | -4,832353356 | 1,35E-06 | 8,60E-06 |
| ENSG00000250462.8 | LRRC37BP1 | transcribed_unproc | 3179,32337 | 0,346896796 | 0,061126493 | 5,675064636 | 1,39E-08 | 1,14E-07 |
| ENSG00000250508.1 | AP000808.1 | lncRNA | 184,9474474 | 0,513991783 | 0,172898535 | 2,972794321 | 0,002951021 | 0,01020761 |
| ENSG00000250519.7 | AP002784.1 | lncRNA | 1211,074289 | -0,228301695 | 0,080989832 | -2,818893289 | 0,004818954 | 0,015784317 |
| ENSG00000250575.1 | AL732372.3 | unprocessed_pseud | 80,43568048 | 1,115384044 | 0,295502725 | 3,774530487 | 0,000160309 | 0,000732819 |

| | | | | | | | | |
|---|---|---|---|---|---|---|---|---|
| ENSG00000250634.6 | LINC01182 | lncRNA | 56,21645024 | 0,913805822 | 0,31163874 | 2,932260034 | 0,003365049 | 0,011497249 |
| ENSG00000250742.4 | LINC02381 | lncRNA | 2284,82118 | 0,166661364 | 0,059067159 | 2,821557143 | 0,004779112 | 0,015672063 |
| ENSG00000250799.10 | PRODH2 | protein_coding | 4218,598741 | -0,282857511 | 0,067143295 | -4,21274395 | 2,52E-05 | 0,000132705 |
| ENSG00000250899.3 | AC125807.2 | lncRNA | 107,4355677 | -0,708387847 | 0,238504946 | -2,970118057 | 0,002976853 | 0,010280146 |
| ENSG00000250903.9 | GMDS-DT | lncRNA | 380,6581775 | 0,344505257 | 0,135632358 | 2,539993132 | 0,011085465 | 0,032681431 |
| ENSG00000250988.7 | SNHG21 | lncRNA | 183,8638496 | -0,569219499 | 0,187954519 | -3,02849595 | 0,002457744 | 0,0086697 |
| ENSG00000251131.2 | AC025171.3 | lncRNA | 57,83613081 | 1,682767094 | 0,339073985 | 4,962831614 | 6,95E-07 | 4,62E-06 |
| ENSG00000251136.9 | AF117829.1 | lncRNA | 834,6473264 | 0,252314773 | 0,088570676 | 2,84873939 | 0,004389282 | 0,014527032 |
| ENSG00000251244.2 | AC107208.1 | lncRNA | 6,407967054 | -2,940552378 | 1,227907625 | -2,394766771 | 0,01663094 | 0,04637363 |
| ENSG00000251297.1 | TUBB7P | unprocessed_pseud | 16,82766385 | 1,612813931 | 0,602588224 | 2,676477681 | 0,007440052 | 0,023137349 |
| ENSG00000251349.3 | MSANTD3-TMEF | protein_coding | 918,5115822 | -0,581022511 | 0,143063147 | -4,061301077 | 4,88E-05 | 0,000244641 |
| ENSG00000251350.2 | LINC02475 | lncRNA | 610,8987576 | 0,304126747 | 0,102261759 | 2,974002704 | 0,002939425 | 0,010175279 |
| ENSG00000251363.3 | LINC02315 | lncRNA | 144,0915684 | -0,599464717 | 0,21007513 | -2,85357299 | 0,00432306 | 0,014343713 |
| ENSG00000251369.8 | ZNF550 | protein_coding | 2391,389412 | 0,496431031 | 0,061940266 | 8,014673825 | 1,10E-15 | 1,74E-14 |
| ENSG00000251410.2 | AC007106.1 | lncRNA | 44,07328906 | 1,319431336 | 0,359542457 | 3,669751124 | 0,000242787 | 0,001071967 |
| ENSG00000251474.6 | RPL32P3 | transcribed_unproc | 923,3410281 | 0,28857075 | 0,091513962 | 3,153297533 | 0,001614372 | 0,00597368 |
| ENSG00000251562.8 | MALAT1 | lncRNA | 25754,82276 | 0,19732466 | 0,058914033 | 3,349366 | 0,000809967 | 0,003207522 |
| ENSG00000251569.1 | AC093899.2 | protein_coding | 326,1653313 | 0,697786024 | 0,148044664 | 4,713348025 | 2,44E-06 | 1,50E-05 |
| ENSG00000251573.2 | LINC02106 | lncRNA | 181,22514 | 0,654999989 | 0,171332967 | 3,822965312 | 0,000131856 | 0,000613721 |
| ENSG00000251580.3 | LINC02482 | lncRNA | 610,4783335 | -0,247170385 | 0,10148885 | -2,435443749 | 0,014873537 | 0,042111754 |
| ENSG00000251602.7 | AL928654.1 | lncRNA | 638,2700662 | -0,346009182 | 0,103392236 | -3,34656834 | 0,000818185 | 0,003235511 |
| ENSG00000251661.3 | AC136475.1 | lncRNA | 393,8276015 | 0,350033456 | 0,12209378 | 2,866922908 | 0,00414484 | 0,013829854 |
| ENSG00000251664.4 | PCDHA12 | protein_coding | 2022,859059 | -0,267378237 | 0,073859692 | -3,62008328 | 0,000294508 | 0,001280983 |
| ENSG00000252690.3 | AC105339.2 | lncRNA | 229,3370017 | -0,524272169 | 0,158315754 | -3,311560324 | 0,000927772 | 0,003623673 |
| ENSG00000253236.2 | LINC02143 | lncRNA | 112,4492057 | 1,806277139 | 0,240219254 | 7,519285446 | 5,51E-14 | 7,72E-13 |
| ENSG00000253276.3 | CCDC71L | protein_coding | 178,7167715 | 1,581651857 | 0,256058092 | 6,176925877 | 6,54E-10 | 6,24E-09 |
| ENSG00000253326.2 | AL606534.4 | processed_pseudog | 54,77927572 | 1,163587211 | 0,383287451 | 3,035808262 | 0,002398919 | 0,0084888 |
| ENSG00000253352.9 | TUG1 | lncRNA | 52011,83421 | 0,131195281 | 0,040549612 | 3,235426317 | 0,001214612 | 0,004615076 |
| ENSG00000253361.3 | AC069120.1 | lncRNA | 62,84901004 | 0,777116057 | 0,292891282 | 2,653257725 | 0,007971898 | 0,024603648 |
| ENSG00000253389.2 | AC113133.1 | lncRNA | 1009,350675 | -0,352467114 | 0,083084234 | -4,242286368 | 2,21E-05 | 0,000117402 |
| ENSG00000253417.5 | LINC02159 | lncRNA | 37,76848434 | 1,325034346 | 0,386663814 | 3,426838248 | 0,000610653 | 0,002484219 |
| ENSG00000253438.4 | PCAT1 | lncRNA | 80,98460521 | 2,058775737 | 0,304632391 | 6,758229903 | 1,40E-11 | 1,58E-10 |
| ENSG00000253452.6 | AC124290.1 | lncRNA | 201,8601261 | -0,505977551 | 0,169271767 | -2,989143194 | 0,00279761 | 0,009734519 |
| ENSG00000253457.2 | SMIM18 | protein_coding | 179,9846867 | -0,489803567 | 0,177585573 | -2,758127018 | 0,00581336 | 0,01863471 |
| ENSG00000253522.6 | MIR3142HG | lncRNA | 41,76113261 | 2,968808182 | 0,46640269 | 6,36533246 | 1,95E-10 | 1,97E-09 |
| ENSG00000253598.1 | SLC10A5 | protein_coding | 473,0163722 | -0,653779987 | 0,123675739 | -5,286242811 | 1,25E-07 | 9,12E-07 |

| | | | | | | | | |
|---|---|---|---|---|---|---|---|---|
| ENSG00000253669.4 | GASAL1 | lncRNA | 358,3375407 | -0,36351418 | 0,151585349 | -2,398082547 | 0,016481151 | 0,046023865 |
| ENSG00000253686.2 | LINC01484 | lncRNA | 1039,314369 | -0,412698644 | 0,080221739 | -5,144473953 | 2,68E-07 | 1,88E-06 |
| ENSG00000253710.4 | ALG11 | protein_coding | 10078,57415 | 0,258539256 | 0,052016119 | 4,970368096 | 6,68E-07 | 4,45E-06 |
| ENSG00000253716.6 | MINCR | lncRNA | 211,6793865 | -0,573764087 | 0,199783169 | -2,871934058 | 0,004079681 | 0,013638725 |
| ENSG00000253729.7 | PRKDC | protein_coding | 44527,85694 | -0,458582885 | 0,059096511 | -7,759897782 | 8,50E-15 | 1,26E-13 |
| ENSG00000253738.2 | OTUD6B-AS1 | lncRNA | 8558,536082 | -0,302482515 | 0,045913132 | -6,588148164 | 4,45E-11 | 4,80E-10 |
| ENSG00000253773.3 | C8orf37-AS1 | lncRNA | 20,68122418 | -2,429102585 | 0,574954885 | -4,224857719 | 2,39E-05 | 0,000126331 |
| ENSG00000253853.2 | AC246817.1 | lncRNA | 310,544393 | 0,449670615 | 0,135827383 | 3,310603543 | 0,00093095 | 0,003634823 |
| ENSG00000253873.6 | PCDHGA11 | protein_coding | 315,8052666 | -0,510680289 | 0,141520592 | -3,608522835 | 0,000307945 | 0,0013353 |
| ENSG00000253882.6 | AC099548.2 | transcribed_unproc | 1483,873567 | 0,572948635 | 0,076503015 | 7,489229484 | 6,93E-14 | 9,63E-13 |
| ENSG00000253953.2 | PCDHGB4 | protein_coding | 1330,00928 | -0,34195609 | 0,077259586 | -4,426066828 | 9,60E-06 | 5,39E-05 |
| ENSG00000253958.1 | CLDN23 | protein_coding | 634,1500083 | 1,263470456 | 0,104052292 | 12,14264895 | 6,28E-34 | 2,31E-32 |
| ENSG00000253982.2 | AC100810.1 | lncRNA | 385,7997043 | -1,225474036 | 0,143832985 | -8,520118206 | 1,59E-17 | 2,81E-16 |
| ENSG00000254004.7 | ZNF260 | protein_coding | 10746,11008 | 0,208926903 | 0,05997973 | 3,483291845 | 0,000495288 | 0,002054255 |
| ENSG00000254087.8 | LYN | protein_coding | 275,7923431 | 1,360596913 | 0,186157159 | 7,3088616 | 2,69E-13 | 3,57E-12 |
| ENSG00000254126.7 | CD8B2 | protein_coding | 64,57754801 | -1,085946701 | 0,299623419 | -3,624371909 | 0,000289665 | 0,001262599 |
| ENSG00000254154.8 | CRYZL2P-SEC16E | lncRNA | 635,7672922 | 2,827922472 | 0,329559338 | 8,580920472 | 9,41E-18 | 1,69E-16 |
| ENSG00000254389.3 | RHPN1-AS1 | lncRNA | 139,0446613 | 0,555911944 | 0,227680088 | 2,441636199 | 0,014620872 | 0,041500783 |
| ENSG00000254413.8 | CHKB-CPT1B | protein_coding | 1441,44319 | 0,364602658 | 0,103358662 | 3,527548168 | 0,000419427 | 0,001767957 |
| ENSG00000254427.2 | AC103736.1 | lncRNA | 250,6217878 | -0,839874667 | 0,172113901 | -4,879760799 | 1,06E-06 | 6,88E-06 |
| ENSG00000254445.1 | HSPB2-C11orf52 | protein_coding | 81,44642161 | 1,050494252 | 0,33528962 | 3,13309506 | 0,001729734 | 0,006353555 |
| ENSG00000254470.2 | AP5B1 | protein_coding | 2937,944521 | 0,569172927 | 0,064840354 | 8,778066263 | 1,66E-18 | 3,12E-17 |
| ENSG00000254505.10 | CHMP4A | protein_coding | 3900,99379 | 0,126689369 | 0,052747748 | 2,40179672 | 0,016314772 | 0,045601539 |
| ENSG00000254535.4 | PABPC4L | protein_coding | 190,5296328 | -0,430619188 | 0,173827828 | -2,477274171 | 0,013239016 | 0,038126537 |
| ENSG00000254539.1 | AC239804.1 | lncRNA | 233,6872515 | 1,20395629 | 0,189114776 | 6,366272978 | 1,94E-10 | 1,96E-09 |
| ENSG00000254553.1 | AL033529.1 | protein_coding | 104,6521284 | 0,981891007 | 0,302254219 | 3,248560136 | 0,001159907 | 0,004428145 |
| ENSG00000254685.6 | FPGT | protein_coding | 2288,152558 | -0,207323981 | 0,086292197 | -2,402580852 | 0,016279835 | 0,045520846 |
| ENSG00000254726.3 | MEX3A | protein_coding | 2826,536953 | -0,263855786 | 0,061564858 | -4,28581819 | 1,82E-05 | 9,78E-05 |
| ENSG00000254772.10 | EEF1G | protein_coding | 61645,62962 | -0,75723447 | 0,046752626 | -16,196619 | 5,33E-59 | 3,69E-57 |
| ENSG00000254815.6 | AP006284.1 | lncRNA | 86,40316886 | -0,60726651 | 0,248561025 | -2,443128441 | 0,014560554 | 0,041350431 |
| ENSG00000254876.5 | SUGT1P4-STRA6 | lncRNA | 302,2894339 | 0,567940583 | 0,139640901 | 4,067150656 | 4,76E-05 | 0,000239169 |
| ENSG00000254929.6 | AL591684.2 | lncRNA | 708,5338272 | 0,598525648 | 0,109361772 | 5,472896403 | 4,43E-08 | 3,41E-07 |
| ENSG00000254996.5 | ANKHD1-EIF4EBI | protein_coding | 1267,094675 | 0,445786655 | 0,07721525 | 5,773298086 | 7,77E-09 | 6,55E-08 |
| ENSG00000254999.4 | BRK1 | protein_coding | 17980,50627 | -0,189038588 | 0,046362887 | -4,07736878 | 4,55E-05 | 0,000229824 |
| ENSG00000255036.6 | STRA6LP | lncRNA | 125,2809523 | 1,077584159 | 0,271106872 | 3,974757818 | 7,05E-05 | 0,000344585 |
| ENSG00000255104.8 | AC005324.4 | protein_coding | 18,22680328 | 3,323362144 | 0,905474285 | 3,670299861 | 0,000242266 | 0,001070153 |

| | | | | | | | | |
|---|---|---|---|---|---|---|---|---|
| ENSG00000255121.2 | AP003392.4 | lncRNA | 649,8633298 | -0,281822927 | 0,10310608 | -2,73332985 | 0,006269751 | 0,019930211 |
| ENSG00000255135.4 | AP002360.1 | lncRNA | 610,6409233 | -1,26040522 | 0,104512216 | -12,0598842 | 1,72E-33 | 6,29E-32 |
| ENSG00000255198.4 | SNHG9 | lncRNA | 418,3104751 | -1,723818215 | 0,148314751 | -11,62270244 | 3,16E-31 | 1,06E-29 |
| ENSG00000255302.4 | EID1 | protein_coding | 34010,87837 | -0,138286106 | 0,040816767 | -3,387973022 | 0,000704112 | 0,002826115 |
| ENSG00000255330.9 | AL096711.2 | protein_coding | 11609,94717 | 0,353609983 | 0,050278465 | 7,033030552 | 2,02E-12 | 2,49E-11 |
| ENSG00000255406.3 | AP000911.3 | lncRNA | 339,1382712 | -0,577159388 | 0,158546188 | -3,640323329 | 0,000272296 | 0,001191506 |
| ENSG00000255455.2 | AP003486.1 | lncRNA | 963,0041713 | -0,584005107 | 0,106274667 | -5,495242895 | 3,90E-08 | 3,03E-07 |
| ENSG00000255521.2 | AL356215.1 | lncRNA | 46,79703862 | 2,347244693 | 0,403292366 | 5,820206106 | 5,88E-09 | 5,03E-08 |
| ENSG00000255561.7 | FDXACB1 | protein_coding | 355,4875121 | -0,323292675 | 0,135402762 | -2,387637218 | 0,016957071 | 0,047166429 |
| ENSG00000255583.2 | AC084357.3 | transcribed_unproc | 55,7032359 | 1,056371324 | 0,307774405 | 3,432291014 | 0,000598505 | 0,002438325 |
| ENSG00000255639.3 | AC005833.1 | protein_coding | 1655,872376 | 0,716178915 | 0,079537108 | 9,004336825 | 2,17E-19 | 4,25E-18 |
| ENSG00000255642.1 | PABPC1P4 | processed_pseudog | 251,6857419 | 0,67153936 | 0,147582628 | 4,550260198 | 5,36E-06 | 3,13E-05 |
| ENSG00000255847.5 | AP003717.1 | lncRNA | 7,568382971 | -2,524516195 | 0,99752243 | -2,530786396 | 0,011380713 | 0,03340735 |
| ENSG00000255857.6 | PXN-AS1 | lncRNA | 524,0130983 | -0,801729639 | 0,109310858 | -7,334400721 | 2,23E-13 | 2,97E-12 |
| ENSG00000255874.2 | LINC00346 | lncRNA | 1805,123398 | 0,455308863 | 0,076360285 | 5,96263963 | 2,48E-09 | 2,22E-08 |
| ENSG00000256029.6 | AL590560.1 | protein_coding | 85,04216181 | -0,753245444 | 0,291266084 | -2,586107638 | 0,009706661 | 0,029174421 |
| ENSG00000256043.3 | CTSO | protein_coding | 1231,806553 | 1,712444006 | 0,082924692 | 20,65059227 | 9,64E-95 | 1,07E-92 |
| ENSG00000256053.7 | APOPT1 | protein_coding | 2743,986759 | -0,301343996 | 0,05972843 | -5,045235528 | 4,53E-07 | 3,09E-06 |
| ENSG00000256084.1 | AC092112.1 | lncRNA | 28,67052836 | 3,583279698 | 0,554273825 | 6,46481854 | 1,01E-10 | 1,06E-09 |
| ENSG00000256092.2 | AC137767.1 | lncRNA | 122,9551049 | -1,186296882 | 0,248909846 | -4,765970091 | 1,88E-06 | 1,17E-05 |
| ENSG00000256124.6 | LINC01152 | lncRNA | 290,0047603 | 0,67874662 | 0,150579801 | 4,507554223 | 6,56E-06 | 3,77E-05 |
| ENSG00000256151.1 | ADGRD1-AS1 | lncRNA | 928,6220412 | -0,378166757 | 0,092097098 | -4,106174495 | 4,02E-05 | 0,000204716 |
| ENSG00000256162.2 | SMLR1 | protein_coding | 646,4316121 | -0,746323943 | 0,106410157 | -7,013653247 | 2,32E-12 | 2,84E-11 |
| ENSG00000256196.2 | AP003721.1 | lncRNA | 1199,510062 | -0,964112976 | 0,095874675 | -10,05597128 | 8,65E-24 | 2,12E-22 |
| ENSG00000256223.6 | ZNF10 | protein_coding | 3626,153302 | 0,164338236 | 0,058063043 | 2,830341412 | 0,004649836 | 0,01528822 |
| ENSG00000256294.8 | ZNF225 | protein_coding | 1468,809743 | 0,266280469 | 0,074511094 | 3,573702324 | 0,000351969 | 0,001507606 |
| ENSG00000256340.8 | ABCC6P1 | transcribed_unproc | 2885,120262 | 0,175576523 | 0,068480904 | 2,56387566 | 0,010351062 | 0,030819055 |
| ENSG00000256364.1 | AC069234.2 | lncRNA | 79,01632598 | -0,879747579 | 0,26844899 | -3,277149899 | 0,001048607 | 0,004045782 |
| ENSG00000256537.4 | SMIM10L1 | protein_coding | 7513,361897 | -0,450135126 | 0,052246362 | -8,615626262 | 6,96E-18 | 1,26E-16 |
| ENSG00000256574.7 | OR13A1 | protein_coding | 312,4444671 | 0,344889612 | 0,144359661 | 2,38909963 | 0,016889721 | 0,046998484 |
| ENSG00000256594.8 | AC010186.2 | transcribed_unproc | 252,8237493 | 0,603321264 | 0,176073588 | 3,42652904 | 0,000611348 | 0,002486599 |
| ENSG00000256637.7 | LINC01965 | lncRNA | 7,396055439 | -2,495713962 | 0,999522823 | -2,496905427 | 0,012528236 | 0,036325914 |
| ENSG00000256663.1 | AC112777.1 | processed_pseudog | 96,78264014 | -0,699948943 | 0,256230397 | -2,731717039 | 0,006300523 | 0,020022373 |
| ENSG00000256683.7 | ZNF350 | protein_coding | 842,4197115 | 0,435744858 | 0,091405931 | 4,767139879 | 1,87E-06 | 1,17E-05 |
| ENSG00000256771.3 | ZNF253 | protein_coding | 3349,43078 | 0,160351107 | 0,056934047 | 2,816436138 | 0,00485597 | 0,015893997 |
| ENSG00000256806.5 | C17orf100 | protein_coding | 1009,883841 | 0,269923937 | 0,08437698 | 3,19902345 | 0,00137894 | 0,005181728 |

| | | | | | | | | |
|---|---|---|---|---|---|---|---|---|
| ENSG00000256925.3 | ADGRA1-AS1 | lncRNA | 411,7102463 | -0,37796094 | 0,122510786 | -3,085123792 | 0,002034674 | 0,007332451 |
| ENSG00000257008.6 | GPR142 | protein_coding | 3751,068354 | 0,549047855 | 0,057504005 | 9,547993257 | 1,32E-21 | 2,92E-20 |
| ENSG00000257093.7 | KIAA1147 | protein_coding | 15353,67643 | -0,311965569 | 0,058984807 | -5,288913983 | 1,23E-07 | 8,99E-07 |
| ENSG00000257103.8 | LSM14A | protein_coding | 24661,67338 | -0,23618514 | 0,040831322 | -5,784410784 | 7,28E-09 | 6,16E-08 |
| ENSG00000257114.3 | LINC02450 | lncRNA | 14,01408878 | 1,973263545 | 0,71687621 | 2,752586175 | 0,005912658 | 0,018915296 |
| ENSG00000257120.1 | AL356756.1 | lncRNA | 12,03270221 | 2,225157207 | 0,748514614 | 2,972763881 | 0,002951314 | 0,01020761 |
| ENSG00000257167.2 | TMPO-AS1 | lncRNA | 921,99893 | -0,590535603 | 0,098871571 | -5,972754331 | 2,33E-09 | 2,09E-08 |
| ENSG00000257176.3 | AC009318.1 | lncRNA | 329,2563604 | 0,90654021 | 0,14795707 | 6,127048964 | 8,95E-10 | 8,40E-09 |
| ENSG00000257218.6 | GATC | protein_coding | 10014,61337 | -0,217224625 | 0,047752056 | -4,549010909 | 5,39E-06 | 3,14E-05 |
| ENSG00000257228.5 | AC090531.1 | lncRNA | 48,41490839 | -0,960279816 | 0,376404404 | -2,551191762 | 0,010735524 | 0,031796145 |
| ENSG00000257267.3 | ZNF271P | transcribed_unitary | 9337,96071 | -0,131374235 | 0,055177972 | -2,380918133 | 0,017269549 | 0,04791001 |
| ENSG00000257303.2 | AC073896.2 | lncRNA | 129,3949959 | -0,826961912 | 0,234238624 | -3,530425082 | 0,000414892 | 0,001750809 |
| ENSG00000257315.2 | ZBED6 | protein_coding | 10101,18909 | -0,488206717 | 0,048153814 | -10,13848483 | 3,73E-24 | 9,31E-23 |
| ENSG00000257355.1 | AC008770.1 | protein_coding | 10,63485674 | 2,35960766 | 0,834537124 | 2,82744481 | 0,00469211 | 0,015418206 |
| ENSG00000257556.2 | LINC02298 | lncRNA | 43,88895431 | -1,111393377 | 0,454124924 | -2,447329618 | 0,014391915 | 0,040974908 |
| ENSG00000257591.5 | ZNF625 | protein_coding | 649,8252078 | 0,453118981 | 0,095453048 | 4,7470352 | 2,06E-06 | 1,28E-05 |
| ENSG00000257621.8 | PSMA3-AS1 | lncRNA | 6877,219958 | -0,183298771 | 0,049542042 | -3,699863038 | 0,000215716 | 0,000961304 |
| ENSG00000257698.2 | GIHCG | lncRNA | 970,3249265 | -0,742265553 | 0,114887592 | -6,460798241 | 1,04E-10 | 1,08E-09 |
| ENSG00000257702.3 | LBX2-AS1 | lncRNA | 616,0011095 | 0,738790319 | 0,121291136 | 6,091049556 | 1,12E-09 | 1,04E-08 |
| ENSG00000257727.6 | CNPY2 | protein_coding | 7988,411969 | 0,363275604 | 0,064161769 | 5,661870138 | 1,50E-08 | 1,22E-07 |
| ENSG00000257743.8 | MGAM2 | protein_coding | 1322,905643 | 3,323931734 | 0,108675161 | 30,58593796 | 1,88E-205 | 3,85E-203 |
| ENSG00000257815.6 | AC025159.1 | lncRNA | 420,6144521 | -0,847186335 | 0,169556376 | -4,996487624 | 5,84E-07 | 3,93E-06 |
| ENSG00000258017.2 | AC011603.2 | lncRNA | 206,6761557 | -0,605532376 | 0,194835332 | -3,107918724 | 0,001884099 | 0,006851241 |
| ENSG00000258056.2 | AC009779.2 | lncRNA | 835,5787482 | 0,323983431 | 0,097439373 | 3,32497451 | 0,000884266 | 0,003468795 |
| ENSG00000258128.2 | MKRN9P | transcribed_process | 417,0045308 | 0,538172617 | 0,115828971 | 4,646269529 | 3,38E-06 | 2,03E-05 |
| ENSG00000258289.8 | CHURC1 | protein_coding | 11216,37372 | 0,190800745 | 0,05208458 | 3,663286653 | 0,000249 | 0,001097891 |
| ENSG00000258301.3 | VASH1-AS1 | lncRNA | 1157,272529 | 0,224688947 | 0,081412178 | 2,759893583 | 0,005782019 | 0,018550096 |
| ENSG00000258315.5 | C17orf49 | protein_coding | 1825,10807 | -0,347534667 | 0,066591444 | -5,218908665 | 1,80E-07 | 1,29E-06 |
| ENSG00000258354.1 | AC138932.2 | lncRNA | 17,3296011 | -1,889338268 | 0,767984955 | -2,460124061 | 0,0138889 | 0,039759028 |
| ENSG00000258429.2 | PDF | protein_coding | 1754,488892 | -0,560563725 | 0,069329735 | -8,085473325 | 6,19E-16 | 9,90E-15 |
| ENSG00000258441.1 | LINC00641 | lncRNA | 715,0199648 | -1,664209118 | 0,146149186 | -11,38705706 | 4,85E-30 | 1,53E-28 |
| ENSG00000258461.5 | AC012651.1 | protein_coding | 60,58561523 | 0,969252223 | 0,345393889 | 2,806222851 | 0,005012601 | 0,016349607 |
| ENSG00000258474.1 | LINC02313 | lncRNA | 274,7523951 | -0,628074867 | 0,149327302 | -4,206028354 | 2,60E-05 | 0,000136168 |
| ENSG00000258659.6 | TRIM34 | protein_coding | 370,5076862 | 6,76020439 | 0,304781791 | 22,18047334 | 5,30E-109 | 6,63E-107 |
| ENSG00000258670.1 | AL049874.3 | lncRNA | 310,0128136 | -0,576966282 | 0,137171417 | -4,206169888 | 2,60E-05 | 0,00013612 |
| ENSG00000258733.6 | LINC02328 | lncRNA | 40,13424036 | 2,437965315 | 0,40542381 | 6,01337478 | 1,82E-09 | 1,65E-08 |

| | | | | | | | | |
|---|---|---|---|---|---|---|---|---|
| ENSG00000258738.1 | AL121603.2 | lncRNA | 327,9342303 | 0,354412401 | 0,135898933 | 2,607911578 | 0,009109648 | 0,027645823 |
| ENSG00000258742.6 | AL117192.1 | lncRNA | 23,4246325 | 1,337633551 | 0,558711033 | 2,394142 | 0,016659298 | 0,046437663 |
| ENSG00000258768.2 | AL356019.2 | lncRNA | 106,9234562 | -1,966077852 | 0,261245535 | -7,525785464 | 5,24E-14 | 7,37E-13 |
| ENSG00000258793.1 | AL355102.4 | lncRNA | 121,5386253 | 0,802284562 | 0,225329658 | 3,560492516 | 0,00037016 | 0,001579213 |
| ENSG00000258818.4 | RNASE4 | protein_coding | 3240,324687 | -0,368772423 | 0,075744624 | -4,868628344 | 1,12E-06 | 7,25E-06 |
| ENSG00000258830.1 | AC137834.1 | protein_coding | 24,82457005 | 2,053359648 | 0,551695991 | 3,721904238 | 0,000197726 | 0,000887815 |
| ENSG00000258867.6 | LINC01146 | lncRNA | 1177,480407 | 0,33080005 | 0,083464266 | 3,963373377 | 7,39E-05 | 0,000360505 |
| ENSG00000258881.6 | AC007040.2 | protein_coding | 170,6692429 | 0,544868019 | 0,180943315 | 3,011263605 | 0,002601629 | 0,009131462 |
| ENSG00000258947.7 | TUBB3 | protein_coding | 8206,273924 | -0,643301962 | 0,051150942 | -12,57654194 | 2,84E-36 | 1,14E-34 |
| ENSG00000258957.2 | AL359317.2 | lncRNA | 168,2552048 | -0,773625441 | 0,252096267 | -3,068769921 | 0,002149421 | 0,007694228 |
| ENSG00000258984.5 | UBE2F-SCLY | protein_coding | 7,530875476 | -2,325323004 | 0,980347576 | -2,371937323 | 0,017695092 | 0,048889018 |
| ENSG00000259024.6 | TVP23C-CDRT4 | protein_coding | 1181,534528 | 0,806408698 | 0,095868598 | 8,411604196 | 4,04E-17 | 6,95E-16 |
| ENSG00000259112.2 | NDUFC2-KCTD14 | protein_coding | 405,5630027 | 0,467805593 | 0,150449983 | 3,109376176 | 0,001874829 | 0,006825828 |
| ENSG00000259120.3 | SMIM6 | protein_coding | 2952,748823 | -0,224817615 | 0,058969091 | -3,812465295 | 0,000137588 | 0,00063789 |
| ENSG00000259125.1 | LRP1-AS | lncRNA | 59,48524426 | -1,998017056 | 0,377494057 | -5,292843741 | 1,20E-07 | 8,83E-07 |
| ENSG00000259153.2 | AC004816.1 | lncRNA | 943,2734549 | -0,206288762 | 0,082340391 | -2,505316773 | 0,012234177 | 0,035592489 |
| ENSG00000259235.2 | AC066612.1 | lncRNA | 295,5770764 | -0,67990395 | 0,158920961 | -4,278252189 | 1,88E-05 | 0,000100951 |
| ENSG00000259251.2 | AC104590.1 | lncRNA | 68,32936028 | -0,84465834 | 0,311552353 | -2,711128101 | 0,006705472 | 0,021142847 |
| ENSG00000259291.2 | ZNF710-AS1 | lncRNA | 372,2650361 | -1,594889838 | 0,12846595 | -12,4148838 | 2,17E-35 | 8,47E-34 |
| ENSG00000259295.6 | CSPG4P12 | transcribed_unproc | 755,7042176 | 0,272813407 | 0,112290238 | 2,429538046 | 0,015118078 | 0,042702073 |
| ENSG00000259316.11 | AC087632.1 | protein_coding | 176,0809274 | 0,537595509 | 0,210024276 | 2,559682721 | 0,010476776 | 0,031127476 |
| ENSG00000259321.1 | AL136295.2 | lncRNA | 18,21235258 | 1,625428965 | 0,596924069 | 2,723007914 | 0,006469051 | 0,020482749 |
| ENSG00000259326.1 | AC116158.1 | lncRNA | 13,19259363 | -1,843205274 | 0,714010032 | -2,58148372 | 0,009837663 | 0,029524795 |
| ENSG00000259366.1 | AC108449.2 | lncRNA | 251,3210844 | -0,433767543 | 0,171114894 | -2,53494907 | 0,011246368 | 0,033077811 |
| ENSG00000259380.6 | AC087473.1 | lncRNA | 634,9963269 | 0,371207947 | 0,097724908 | 3,798498832 | 0,000145575 | 0,000672568 |
| ENSG00000259439.2 | LINC01833 | lncRNA | 356,4418761 | 0,641448448 | 0,131066232 | 4,894078666 | 9,88E-07 | 6,42E-06 |
| ENSG00000259495.3 | AC016705.2 | lncRNA | 2572,418764 | -0,278917531 | 0,077199864 | -3,612927749 | 0,000302759 | 0,001313745 |
| ENSG00000259498.1 | TPM1-AS | lncRNA | 27,42927307 | 1,217074415 | 0,463995678 | 2,623029639 | 0,008715169 | 0,026610111 |
| ENSG00000259529.2 | AL136295.5 | protein_coding | 456,5593143 | 0,387517251 | 0,143977486 | 2,691512824 | 0,007112876 | 0,022270654 |
| ENSG00000259642.2 | ST20-AS1 | lncRNA | 606,4219507 | -0,524753121 | 0,104173866 | -5,037281828 | 4,72E-07 | 3,21E-06 |
| ENSG00000259673.6 | IQCH-AS1 | lncRNA | 1428,198349 | -0,483788149 | 0,080120851 | -6,038230273 | 1,56E-09 | 1,43E-08 |
| ENSG00000259726.1 | CSPG4P11 | unprocessed_pseud | 407,443111 | 0,398010287 | 0,139063984 | 2,862065916 | 0,004208894 | 0,014016569 |
| ENSG00000259757.1 | AC091891.1 | lncRNA | 11,3935679 | 2,349798224 | 0,759337142 | 3,094538769 | 0,001971192 | 0,007132222 |
| ENSG00000259781.1 | HMGB1P6 | processed_pseudog | 3505,479982 | -0,273574409 | 0,062844366 | -4,353204989 | 1,34E-05 | 7,37E-05 |
| ENSG00000259834.1 | AL365361.1 | lncRNA | 1417,058829 | 0,5068923 | 0,087645295 | 5,783451327 | 7,32E-09 | 6,19E-08 |
| ENSG00000259881.1 | AC092384.2 | lncRNA | 307,1845995 | -0,438127509 | 0,148050211 | -2,959317018 | 0,003083217 | 0,010616512 |

| | | | | | | | | |
|---|---|---|---|---|---|---|---|---|
| ENSG00000259956.2 | RBM15B | protein_coding | 11791,81905 | -0,110727313 | 0,045986685 | -2,407812475 | 0,016048422 | 0,044985545 |
| ENSG00000259969.1 | AL049838.1 | lncRNA | 41,94521635 | -1,509373358 | 0,415172251 | -3,63553526 | 0,000277404 | 0,001211503 |
| ENSG00000259974.3 | LINC00261 | lncRNA | 3164,533219 | -0,185586043 | 0,060083785 | -3,088787485 | 0,002009751 | 0,007255412 |
| ENSG00000259976.3 | AC093010.3 | lncRNA | 594,2923584 | 0,692662396 | 0,129050108 | 5,367391048 | 7,99E-08 | 5,95E-07 |
| ENSG00000260000.2 | AL133338.1 | lncRNA | 1873,112719 | 0,857551937 | 0,069153091 | 12,4007752 | 2,59E-35 | 1,01E-33 |
| ENSG00000260032.2 | NORAD | lncRNA | 135236,7628 | -0,50585016 | 0,046856392 | -10,79575558 | 3,60E-27 | 1,02E-25 |
| ENSG00000260062.3 | GOLGA2P11 | transcribed_unproc | 32,88948879 | 1,271181778 | 0,440998811 | 2,882506133 | 0,003945255 | 0,013244405 |
| ENSG00000260070.1 | AC006960.3 | lncRNA | 46,4521669 | 1,082754893 | 0,35095542 | 3,085163617 | 0,002034402 | 0,007332451 |
| ENSG00000260075.1 | NSFP1 | unprocessed_pseud | 228,0815057 | 0,708901742 | 0,238271736 | 2,975181843 | 0,002928149 | 0,010141516 |
| ENSG00000260077.1 | AC104794.2 | lncRNA | 193,8322012 | -0,397065652 | 0,165772629 | -2,395242535 | 0,016609375 | 0,046327189 |
| ENSG00000260081.2 | AF274858.1 | transcribed_process | 86,34862623 | -0,933600645 | 0,268211447 | -3,480838179 | 0,000499847 | 0,002072019 |
| ENSG00000260103.2 | AC012435.1 | transcribed_process | 132,5257973 | 1,017598354 | 0,222524809 | 4,572965865 | 4,81E-06 | 2,83E-05 |
| ENSG00000260136.6 | AC008915.2 | lncRNA | 242,7371346 | -0,402714001 | 0,15402175 | -2,614656704 | 0,008931718 | 0,027176526 |
| ENSG00000260220.6 | CCDC187 | protein_coding | 1110,205424 | -0,264043883 | 0,097163918 | -2,717509623 | 0,006577524 | 0,02079111 |
| ENSG00000260230.3 | FRRS1L | protein_coding | 1444,717531 | -0,651863241 | 0,089250953 | -7,30371185 | 2,80E-13 | 3,70E-12 |
| ENSG00000260233.3 | SSSCA1-AS1 | lncRNA | 141,2005999 | -0,575305447 | 0,195625468 | -2,940851489 | 0,003273114 | 0,011212072 |
| ENSG00000260236.1 | AC099778.1 | lncRNA | 338,9950399 | 0,666687866 | 0,131079076 | 5,086150182 | 3,65E-07 | 2,52E-06 |
| ENSG00000260257.2 | AL035071.1 | lncRNA | 301,8810283 | -0,642051979 | 0,170067403 | -3,77527949 | 0,000159828 | 0,000731122 |
| ENSG00000260260.1 | SNHG19 | lncRNA | 756,4142312 | -1,461027149 | 0,092989912 | -15,71167362 | 1,26E-55 | 8,07E-54 |
| ENSG00000260265.1 | LINC02562 | lncRNA | 62,33118004 | -1,228400076 | 0,303031676 | -4,053701881 | 5,04E-05 | 0,000252392 |
| ENSG00000260267.1 | AC026471.1 | lncRNA | 355,221688 | 0,414613971 | 0,127939696 | 3,240698424 | 0,001192373 | 0,004536699 |
| ENSG00000260276.2 | AC022167.2 | lncRNA | 69,05378307 | 0,858469014 | 0,309399497 | 2,774629641 | 0,00552646 | 0,0178242 |
| ENSG00000260293.2 | AC106820.4 | lncRNA | 5667,165879 | -0,86121309 | 0,063528098 | -13,55641237 | 7,26E-42 | 3,43E-40 |
| ENSG00000260296.1 | AC095057.3 | lncRNA | 140,8320401 | -1,190762605 | 0,231576504 | -5,14198368 | 2,72E-07 | 1,91E-06 |
| ENSG00000260314.3 | MRC1 | protein_coding | 27,45602605 | 1,962227876 | 0,476598866 | 4,117147597 | 3,84E-05 | 0,000195788 |
| ENSG00000260342.2 | AC138811.2 | protein_coding | 121,9134232 | -1,208622908 | 0,254192927 | -4,75474641 | 1,99E-06 | 1,24E-05 |
| ENSG00000260412.1 | AL353746.1 | lncRNA | 691,9848738 | -0,665920438 | 0,124496389 | -5,348913667 | 8,85E-08 | 6,56E-07 |
| ENSG00000260469.3 | INSYN1-AS1 | lncRNA | 185,5635401 | -2,019564175 | 0,191899055 | -10,52409652 | 6,69E-26 | 1,80E-24 |
| ENSG00000260532.1 | AL031598.1 | lncRNA | 19,23152 | -2,089137403 | 0,616898018 | -3,386519883 | 0,000707851 | 0,0028391 |
| ENSG00000260552.1 | AC023043.1 | lncRNA | 16,46418869 | 2,100936186 | 0,632594831 | 3,321140297 | 0,000896505 | 0,003510075 |
| ENSG00000260565.7 | ERVK13-1 | lncRNA | 4316,430431 | 0,204407108 | 0,06134948 | 3,331847472 | 0,000862715 | 0,003392533 |
| ENSG00000260604.2 | AL590004.3 | lncRNA | 209,404701 | -0,985539129 | 0,167605199 | -5,880122655 | 4,10E-09 | 3,58E-08 |
| ENSG00000260669.2 | AL096870.3 | lncRNA | 222,160282 | 0,74750655 | 0,179650365 | 4,160896348 | 3,17E-05 | 0,000163956 |
| ENSG00000260691.6 | ANKRD20A1 | protein_coding | 509,8456606 | -0,371542965 | 0,146363432 | -2,538495855 | 0,011133012 | 0,032800127 |
| ENSG00000260708.1 | AL118516.1 | lncRNA | 348,4018456 | -0,60734213 | 0,128735959 | -4,71773494 | 2,38E-06 | 1,47E-05 |
| ENSG00000260742.1 | AC009962.1 | lncRNA | 174,4292208 | 0,704096377 | 0,179505281 | 3,922427098 | 8,77E-05 | 0,000421888 |

| | | | | | | | | |
|---|---|---|---|---|---|---|---|---|
| ENSG00000260772.1 | AC012321.1 | lncRNA | 429,4014537 | 0,416248892 | 0,135588361 | 3,069945601 | 0,002140978 | 0,007671325 |
| ENSG00000260802.1 | SERTM2 | protein_coding | 417,77 | 0,89721084 | 0,134787631 | 6,656477557 | 2,80E-11 | 3,08E-10 |
| ENSG00000260804.3 | LINC01963 | lncRNA | 1026,483199 | -2,200210666 | 0,139880031 | -15,72926927 | 9,53E-56 | 6,13E-54 |
| ENSG00000260807.7 | AC009041.2 | lncRNA | 128,4199776 | -1,16521902 | 0,232760554 | -5,006084574 | 5,55E-07 | 3,75E-06 |
| ENSG00000260838.3 | AC022893.2 | lncRNA | 1421,892126 | -0,383531846 | 0,096207918 | -3,986489389 | 6,71E-05 | 0,000328848 |
| ENSG00000260852.1 | FBXL19-AS1 | lncRNA | 685,6498343 | -0,804520906 | 0,09688307 | -8,304040186 | 1,01E-16 | 1,69E-15 |
| ENSG00000260855.1 | AL591848.4 | lncRNA | 122,6540852 | -1,234959556 | 0,239789444 | -5,150183165 | 2,60E-07 | 1,83E-06 |
| ENSG00000260903.3 | XKR7 | protein_coding | 12693,99927 | 0,222969588 | 0,044663805 | 4,992176262 | 5,97E-07 | 4,00E-06 |
| ENSG00000260912.1 | AL158206.1 | lncRNA | 844,8968156 | -0,340025728 | 0,087037707 | -3,906648518 | 9,36E-05 | 0,000447717 |
| ENSG00000260916.7 | CCPG1 | protein_coding | 8531,041399 | 0,197798192 | 0,049992287 | 3,956574188 | 7,60E-05 | 0,000369396 |
| ENSG00000260917.1 | AL158212.3 | lncRNA | 328,248945 | -1,173420004 | 0,132788477 | -8,836760737 | 9,85E-19 | 1,87E-17 |
| ENSG00000260920.2 | AL031985.3 | lncRNA | 713,6174764 | -0,436169269 | 0,100456859 | -4,341856542 | 1,41E-05 | 7,72E-05 |
| ENSG00000260941.1 | LINC00622 | lncRNA | 34,41385371 | -1,341346435 | 0,459484731 | -2,919240499 | 0,003508854 | 0,011932436 |
| ENSG00000260947.1 | AL356489.2 | lncRNA | 790,6180378 | -0,460365493 | 0,112595063 | -4,088682756 | 4,34E-05 | 0,000219885 |
| ENSG00000260948.1 | AL390195.2 | lncRNA | 53,9174787 | -1,01989974 | 0,338046667 | -3,017038297 | 0,002552576 | 0,008973282 |
| ENSG00000261008.7 | LINC01572 | lncRNA | 118,9093204 | -0,914947038 | 0,223831018 | -4,087668655 | 4,36E-05 | 0,000220699 |
| ENSG00000261051.1 | AC107021.2 | lncRNA | 132,0080929 | -0,993554937 | 0,217256079 | -4,573197408 | 4,80E-06 | 2,82E-05 |
| ENSG00000261052.5 | SULT1A3 | protein_coding | 230,8673956 | -0,615058927 | 0,229864684 | -2,675743473 | 0,007456369 | 0,023181687 |
| ENSG00000261069.3 | AC124312.3 | lncRNA | 149,2652433 | -2,20810316 | 0,204651345 | -10,78958538 | 3,86E-27 | 1,08E-25 |
| ENSG00000261115.6 | TMEM178B | protein_coding | 11306,39749 | -0,545298927 | 0,059982845 | -9,090914685 | 9,82E-20 | 1,96E-18 |
| ENSG00000261135.1 | AL137802.2 | lncRNA | 680,5862147 | 0,324318708 | 0,108000964 | 3,002924188 | 0,00267399 | 0,00935045 |
| ENSG00000261167.1 | AC107027.3 | lncRNA | 400,2398207 | -0,994057631 | 0,15174815 | -6,550706754 | 5,73E-11 | 6,11E-10 |
| ENSG00000261189.1 | AL031058.1 | lncRNA | 311,7016824 | -0,464719265 | 0,162072654 | -2,867351482 | 0,00413923 | 0,013817282 |
| ENSG00000261211.1 | AL031123.2 | lncRNA | 938,3370131 | 0,315928817 | 0,087974888 | 3,591124961 | 0,000329254 | 0,001418949 |
| ENSG00000261240.1 | AC009065.6 | lncRNA | 147,8730395 | -1,325932219 | 0,224615042 | -5,903131891 | 3,57E-09 | 3,13E-08 |
| ENSG00000261324.2 | AC010168.2 | lncRNA | 618,3812401 | 0,585577866 | 0,103047544 | 5,682598967 | 1,33E-08 | 1,09E-07 |
| ENSG00000261335.1 | AC005837.1 | lncRNA | 215,8410718 | -0,498939613 | 0,156357016 | -3,191027981 | 0,001417675 | 0,005311328 |
| ENSG00000261342.1 | AC006538.2 | lncRNA | 146,5058634 | -0,594122084 | 0,203536336 | -2,918997631 | 0,003511589 | 0,011938129 |
| ENSG00000261346.1 | AC116348.3 | lncRNA | 46,95538634 | 0,786545396 | 0,32990321 | 2,384170182 | 0,017117684 | 0,0475564 |
| ENSG00000261351.2 | AC116913.1 | lncRNA | 92,31344616 | 0,604462149 | 0,243406897 | 2,483340265 | 0,013015669 | 0,037536157 |
| ENSG00000261377.5 | PDCD6IPP2 | transcribed_unproc | 1178,690811 | -0,317879113 | 0,087025024 | -3,652732248 | 0,000259465 | 0,001140014 |
| ENSG00000261431.1 | AL023803.1 | lncRNA | 52,38181975 | -0,909356262 | 0,314411662 | -2,89224724 | 0,003824968 | 0,012879014 |
| ENSG00000261456.5 | TUBB8 | protein_coding | 48,11200151 | 1,53728675 | 0,432376738 | 3,555433527 | 0,000377356 | 0,00160687 |
| ENSG00000261465.7 | AC099518.4 | lncRNA | 9,488722625 | -2,403971754 | 0,852122913 | -2,821156101 | 0,004785091 | 0,015689384 |
| ENSG00000261488.1 | TBILA | lncRNA | 85,35606377 | 1,156608864 | 0,275256663 | 4,201928673 | 2,65E-05 | 0,000138407 |
| ENSG00000261526.2 | AC012615.1 | lncRNA | 298,6370842 | -0,78140922 | 0,146435626 | -5,336196116 | 9,49E-08 | 7,00E-07 |

| | | | | | | | | |
|---|---|---|---|---|---|---|---|---|
| ENSG00000261578.1 | AP003119.3 | lncRNA | 197,2850477 | -0,576344808 | 0,186667064 | -3,087554901 | 0,002018105 | 0,007280898 |
| ENSG00000261609.7 | GAN | protein_coding | 578,6266454 | -0,301907826 | 0,105956754 | -2,849349517 | 0,004380872 | 0,014503465 |
| ENSG00000261618.2 | LINC02605 | lncRNA | 287,5703296 | 2,017686358 | 0,156863288 | 12,86270599 | 7,30E-38 | 3,05E-36 |
| ENSG00000261644.2 | AC007728.2 | lncRNA | 21,09390844 | 2,619109123 | 0,631311767 | 4,148677818 | 3,34E-05 | 0,000172243 |
| ENSG00000261678.3 | SCRT1 | protein_coding | 47,98507259 | -1,104022089 | 0,354205197 | -3,116899746 | 0,001827637 | 0,006672877 |
| ENSG00000261716.2 | HIST2H2BC | transcribed_unproc | 1017,808403 | 0,377569562 | 0,088087904 | 4,286281603 | 1,82E-05 | 9,76E-05 |
| ENSG00000261796.1 | ISY1-RAB43 | protein_coding | 956,4438983 | 0,613796942 | 0,130024774 | 4,720615338 | 2,35E-06 | 1,45E-05 |
| ENSG00000261801.6 | LOXL1-AS1 | lncRNA | 1172,70731 | -0,461401544 | 0,098861691 | -4,667141953 | 3,05E-06 | 1,85E-05 |
| ENSG00000261824.7 | LINC00662 | lncRNA | 2865,250443 | -0,252295415 | 0,061008551 | -4,135410693 | 3,54E-05 | 0,000181713 |
| ENSG00000261840.3 | AC093249.6 | lncRNA | 144,9793913 | -0,659005997 | 0,220902181 | -2,98324804 | 0,002852068 | 0,009907614 |
| ENSG00000261884.2 | AC040162.1 | protein_coding | 204,1473551 | 1,657811362 | 0,271727465 | 6,101007714 | 1,05E-09 | 9,82E-09 |
| ENSG00000261889.1 | AC108134.2 | lncRNA | 40,89404699 | 1,785869393 | 0,381230492 | 4,684487281 | 2,81E-06 | 1,71E-05 |
| ENSG00000262209.3 | PCDHGB3 | protein_coding | 16,29320068 | -2,160385136 | 0,741589145 | -2,913183333 | 0,003577645 | 0,012138862 |
| ENSG00000262246.6 | CORO7 | protein_coding | 5661,175581 | -0,366661631 | 0,070826885 | -5,176870786 | 2,26E-07 | 1,60E-06 |
| ENSG00000262587.2 | AC133552.2 | transcribed_unproc | 199,9001958 | 0,420516467 | 0,16979039 | 2,476680017 | 0,013261073 | 0,038175406 |
| ENSG00000262655.4 | SPON1 | protein_coding | 9304,430718 | -0,329447872 | 0,049620539 | -6,639344922 | 3,15E-11 | 3,44E-10 |
| ENSG00000262714.1 | AC007342.5 | lncRNA | 368,2082921 | -0,499605821 | 0,134337393 | -3,719037629 | 0,000199983 | 0,000897055 |
| ENSG00000262879.6 | AC005670.3 | lncRNA | 5985,899645 | -0,779206518 | 0,071075337 | -10,96310695 | 5,75E-28 | 1,67E-26 |
| ENSG00000262902.1 | MTCO1P40 | processed_pseudog | 76,18977158 | -1,10854661 | 0,325244632 | -3,408347134 | 0,000653577 | 0,002641653 |
| ENSG00000262904.1 | TMPOP2 | processed_pseudog | 61,80630083 | -0,92699935 | 0,292804526 | -3,165932453 | 0,001545867 | 0,005751391 |
| ENSG00000263001.6 | GTF2I | protein_coding | 57349,78219 | -0,145418698 | 0,042482522 | -3,423024106 | 0,000619286 | 0,002515702 |
| ENSG00000263004.2 | AC007114.1 | lncRNA | 391,2176395 | -0,310316448 | 0,119697901 | -2,592496988 | 0,009528202 | 0,028737812 |
| ENSG00000263069.6 | AC124319.3 | lncRNA | 150,6120825 | -3,336666487 | 0,539347322 | -6,186489393 | 6,15E-10 | 5,91E-09 |
| ENSG00000263142.5 | LRRC37A17P | transcribed_unproc | 140,5930954 | 0,996670251 | 0,219913559 | 4,532100043 | 5,84E-06 | 3,39E-05 |
| ENSG00000263155.6 | MYZAP | protein_coding | 497,3477217 | 0,295365609 | 0,115357693 | 2,560432683 | 0,010454191 | 0,031068576 |
| ENSG00000263235.1 | AC006111.2 | lncRNA | 154,1444532 | 0,604351535 | 0,190480967 | 3,172765991 | 0,001509942 | 0,005628897 |
| ENSG00000263300.1 | AC015853.3 | lncRNA | 15,01643412 | -1,461070368 | 0,603734872 | -2,420052965 | 0,015518247 | 0,043667985 |
| ENSG00000263400.8 | TMEM220-AS1 | lncRNA | 123,1006853 | 0,697903098 | 0,215704178 | 3,23546398 | 0,001214452 | 0,004615076 |
| ENSG00000263465.4 | SRSF8 | protein_coding | 5925,939482 | -0,237853135 | 0,04876313 | -4,877724896 | 1,07E-06 | 6,94E-06 |
| ENSG00000263513.5 | FAM72C | protein_coding | 139,097959 | -1,218124437 | 0,233725926 | -5,211764302 | 1,87E-07 | 1,34E-06 |
| ENSG00000263528.8 | IKBKE | protein_coding | 530,1263997 | 1,743195575 | 0,127414368 | 13,68131089 | 1,31E-42 | 6,41E-41 |
| ENSG00000263731.1 | AC145207.5 | lncRNA | 152,8799506 | -1,188223579 | 0,212144588 | -5,601008205 | 2,13E-08 | 1,71E-07 |
| ENSG00000263753.8 | LINC00667 | lncRNA | 14524,97934 | -0,188975739 | 0,051325088 | -3,681937 | 0,000231469 | 0,001027031 |
| ENSG00000263826.1 | AC112907.3 | lncRNA | 57,51032974 | -0,859219941 | 0,307615988 | -2,793157622 | 0,005219625 | 0,01694139 |
| ENSG00000263893.2 | AC080037.1 | lncRNA | 226,032317 | 0,54314023 | 0,158275512 | 3,431612523 | 0,000600004 | 0,002443991 |
| ENSG00000263956.6 | NBPF11 | protein_coding | 2779,980914 | 0,456630805 | 0,061632769 | 7,408896419 | 1,27E-13 | 1,73E-12 |

| | | | | | | | | |
|---|---|---|---|---|---|---|---|---|
| ENSG00000263958.2 | AC091138.1 | lncRNA | 87,48826538 | -1,254948665 | 0,290852721 | -4,314722105 | 1,60E-05 | 8,66E-05 |
| ENSG00000264247.2 | LINC00909 | lncRNA | 3325,55862 | 0,166886165 | 0,056417505 | 2,958056427 | 0,003095854 | 0,010655136 |
| ENSG00000264364.3 | DYNLL2 | protein_coding | 54705,71823 | -0,129526414 | 0,043976299 | -2,945368686 | 0,0032257 | 0,011056386 |
| ENSG00000264522.6 | OTUD7B | protein_coding | 3212,209834 | 0,190496964 | 0,064091546 | 2,972263532 | 0,002956128 | 0,01022112 |
| ENSG00000264569.1 | DCXR-DT | lncRNA | 102,6464659 | -1,189142448 | 0,250378748 | -4,749374534 | 2,04E-06 | 1,27E-05 |
| ENSG00000264672.6 | SEPT4-AS1 | lncRNA | 148,9589562 | 0,622022086 | 0,196011133 | 3,173401814 | 0,001506639 | 0,005622169 |
| ENSG00000264717.5 | NPY4R2 | protein_coding | 69,0275869 | -1,350787431 | 0,293455774 | -4,603035788 | 4,16E-06 | 2,47E-05 |
| ENSG00000264727.1 | AC005725.1 | lncRNA | 22,91002101 | -1,457771742 | 0,551858842 | -2,641566341 | 0,008252364 | 0,025365625 |
| ENSG00000264743.2 | DPRXP4 | processed_pseudog | 14,94934923 | 1,615316173 | 0,680036173 | 2,375338601 | 0,017532858 | 0,048506282 |
| ENSG00000264920.2 | AC018521.5 | lncRNA | 592,0960135 | -0,465310944 | 0,109201075 | -4,26104727 | 2,03E-05 | 0,000108504 |
| ENSG00000265185.5 | SNORD3B-1 | snoRNA | 24,23177871 | -1,236709019 | 0,514024091 | -2,405935909 | 0,016131094 | 0,045194773 |
| ENSG00000265354.4 | TIMM23 | protein_coding | 5749,779171 | -0,176560821 | 0,052133873 | -3,386681442 | 0,000707435 | 0,002838124 |
| ENSG00000265491.5 | RNF115 | protein_coding | 4690,689386 | 0,130806742 | 0,051915968 | 2,519585924 | 0,011749296 | 0,034341631 |
| ENSG00000265514.2 | LINC01892 | lncRNA | 16,29023589 | 1,537273994 | 0,581501084 | 2,643630488 | 0,008202214 | 0,02521492 |
| ENSG00000265681.7 | RPL17 | protein_coding | 52667,9339 | -0,589593391 | 0,039256374 | -15,01904874 | 5,51E-51 | 3,25E-49 |
| ENSG00000265688.2 | MAFG-DT | lncRNA | 532,8481282 | -0,437971155 | 0,114652182 | -3,819998418 | 0,000133453 | 0,000620381 |
| ENSG00000266053.2 | NDUFV2-AS1 | lncRNA | 66,737446 | -0,710740195 | 0,281099408 | -2,528430071 | 0,011457391 | 0,033575629 |
| ENSG00000266074.8 | BAHCC1 | protein_coding | 2165,064763 | 0,580479443 | 0,094485053 | 6,143611314 | 8,07E-10 | 7,63E-09 |
| ENSG00000266086.3 | AC015813.2 | protein_coding | 272,5200274 | -0,381557458 | 0,144582982 | -2,639020534 | 0,008314594 | 0,025525282 |
| ENSG00000266173.7 | STRADA | protein_coding | 3727,623186 | 0,346849248 | 0,056494569 | 6,139514951 | 8,28E-10 | 7,82E-09 |
| ENSG00000266338.6 | NBPF15 | protein_coding | 3389,787046 | 0,243935164 | 0,072234773 | 3,376976963 | 0,000732872 | 0,002929535 |
| ENSG00000266402.3 | SNHG25 | lncRNA | 12,7345506 | -2,24762073 | 0,722193265 | -3,112215026 | 0,001856892 | 0,006766518 |
| ENSG00000266412.5 | NCOA4 | protein_coding | 51330,2936 | 0,094499287 | 0,037313555 | 2,532572573 | 0,011322893 | 0,03326811 |
| ENSG00000266524.3 | GDF10 | protein_coding | 98,67942356 | -0,694448151 | 0,244053344 | -2,845476892 | 0,004434498 | 0,014657285 |
| ENSG00000266916.7 | ZNF793-AS1 | lncRNA | 634,3101299 | -0,856637877 | 0,117729666 | -7,276312832 | 3,43E-13 | 4,51E-12 |
| ENSG00000266947.1 | AC022916.1 | lncRNA | 368,9760644 | 0,798366548 | 0,125128565 | 6,380370037 | 1,77E-10 | 1,80E-09 |
| ENSG00000266962.2 | AC067852.2 | lncRNA | 1106,335843 | 0,333957712 | 0,083534 | 3,997865696 | 6,39E-05 | 0,000315019 |
| ENSG00000266976.3 | AC079466.1 | lncRNA | 6885,144879 | 0,488745368 | 0,05044297 | 9,689068079 | 3,36E-22 | 7,65E-21 |
| ENSG00000267013.6 | LINC01929 | lncRNA | 10,4252707 | -2,744085744 | 0,997430125 | -2,75115587 | 0,005938538 | 0,018989991 |
| ENSG00000267072.2 | NAGPA-AS1 | lncRNA | 47,69380064 | -1,689426636 | 0,4232955 | -3,991128272 | 6,58E-05 | 0,000323043 |
| ENSG00000267106.7 | ZNF561-AS1 | lncRNA | 757,2977916 | -0,415203212 | 0,142396268 | -2,915829308 | 0,003547445 | 0,01204184 |
| ENSG00000267216.1 | AC020915.1 | lncRNA | 748,086849 | 0,277810383 | 0,101448583 | 2,738435319 | 0,00617323 | 0,019662267 |
| ENSG00000267221.2 | C17orf113 | protein_coding | 565,981312 | 0,924982602 | 0,102768142 | 9,000674593 | 2,24E-19 | 4,39E-18 |
| ENSG00000267278.6 | MAP3K14-AS1 | lncRNA | 386,6530227 | 0,559975047 | 0,140177022 | 3,994770615 | 6,48E-05 | 0,000318743 |
| ENSG00000267289.1 | AC008752.2 | lncRNA | 35,39849853 | -2,265861257 | 0,472089102 | -4,799647451 | 1,59E-06 | 1,00E-05 |
| ENSG00000267309.1 | AC092295.2 | lncRNA | 362,4722943 | 0,338090744 | 0,132963938 | 2,542725108 | 0,010999173 | 0,032469557 |

| | | | | | | | | |
|---|---|---|---|---|---|---|---|---|
| ENSG00000267321.3 | SNHG30 | lncRNA | 1563,505409 | -0,178413228 | 0,067956219 | -2,625414294 | 0,008654359 | 0,026449555 |
| ENSG00000267397.1 | AC090229.1 | lncRNA | 93,61867609 | -0,961165521 | 0,256217527 | -3,751365221 | 0,000175874 | 0,000797493 |
| ENSG00000267429.1 | AC006116.6 | lncRNA | 16,39466237 | -1,799674331 | 0,680732368 | -2,643732567 | 0,008199741 | 0,025210761 |
| ENSG00000267443.2 | AC010641.1 | lncRNA | 17,1077315 | -4,161099308 | 0,802871699 | -5,182769943 | 2,19E-07 | 1,55E-06 |
| ENSG00000267497.1 | NFE2L3P1 | processed_pseudog | 72,95799588 | 0,789150063 | 0,271468482 | 2,906967533 | 0,003649511 | 0,012356624 |
| ENSG00000267520.2 | AC010733.2 | lncRNA | 2063,521772 | 0,181048032 | 0,069346233 | 2,610783938 | 0,009033495 | 0,027440612 |
| ENSG00000267534.4 | S1PR2 | protein_coding | 708,5485077 | 0,556514011 | 0,100333022 | 5,546668455 | 2,91E-08 | 2,31E-07 |
| ENSG00000267575.7 | AC006504.5 | lncRNA | 1863,330404 | -0,571339595 | 0,073526985 | -7,770474927 | 7,82E-15 | 1,17E-13 |
| ENSG00000267586.7 | LINC00907 | lncRNA | 1879,960441 | -0,3442418 | 0,069031986 | -4,986699963 | 6,14E-07 | 4,11E-06 |
| ENSG00000267632.1 | AC067852.3 | lncRNA | 249,44278 | 0,721083246 | 0,166694071 | 4,325788198 | 1,52E-05 | 8,27E-05 |
| ENSG00000267640.6 | AC016582.3 | transcribed_unproc | 1794,048543 | 0,246455749 | 0,094016596 | 2,62140686 | 0,008756769 | 0,026715384 |
| ENSG00000267659.5 | LINC01482 | lncRNA | 25,04056445 | 3,256776467 | 0,592934713 | 5,492639231 | 3,96E-08 | 3,07E-07 |
| ENSG00000267680.5 | ZNF224 | protein_coding | 3726,506798 | 0,590946378 | 0,062018102 | 9,528611147 | 1,59E-21 | 3,50E-20 |
| ENSG00000267681.1 | AC135721.1 | transcribed_process | 39,77354209 | 1,097353516 | 0,36711378 | 2,989137365 | 0,002797663 | 0,009734519 |
| ENSG00000267740.5 | AC024592.3 | protein_coding | 393,6341603 | -0,341600755 | 0,13882512 | -2,460655219 | 0,013868357 | 0,039710322 |
| ENSG00000267751.5 | AC009005.1 | lncRNA | 126,9930125 | -0,836089853 | 0,223389569 | -3,742743476 | 0,000182022 | 0,000821892 |
| ENSG00000267904.1 | AC024075.1 | lncRNA | 75,77153835 | -1,617341215 | 0,285464591 | -5,665645643 | 1,46E-08 | 1,20E-07 |
| ENSG00000267922.1 | AC007785.1 | lncRNA | 44,55403397 | 1,065533932 | 0,35529054 | 2,999049545 | 0,002708233 | 0,009459902 |
| ENSG00000268001.1 | CARD8-AS1 | lncRNA | 751,1987914 | 0,807776124 | 0,103965294 | 7,769670917 | 7,87E-15 | 1,17E-13 |
| ENSG00000268043.7 | NBPF12 | protein_coding | 7551,850516 | 0,39837292 | 0,05305646 | 7,508471523 | 5,98E-14 | 8,36E-13 |
| ENSG00000268061.5 | NAPA-AS1 | lncRNA | 149,9346122 | -2,290516891 | 0,273528568 | -8,373958545 | 5,57E-17 | 9,50E-16 |
| ENSG00000268089.3 | GABRQ | protein_coding | 897,8344573 | 0,261354638 | 0,084354026 | 3,098306623 | 0,001946299 | 0,007052199 |
| ENSG00000268173.3 | AC007192.1 | protein_coding | 608,9742013 | 1,942912858 | 0,324161502 | 5,993657008 | 2,05E-09 | 1,85E-08 |
| ENSG00000268350.7 | FAM156A | protein_coding | 1328,171696 | -0,310445975 | 0,098965171 | -3,136921537 | 0,001707318 | 0,006277364 |
| ENSG00000268400.5 | AC008763.2 | protein_coding | 19,45881413 | -1,590297465 | 0,660770279 | -2,406732741 | 0,016095944 | 0,04511304 |
| ENSG00000268460.2 | AC006262.1 | lncRNA | 678,6304638 | 0,426385293 | 0,129656879 | 3,288566696 | 0,001006989 | 0,003899917 |
| ENSG00000268471.6 | MIR4453HG | lncRNA | 1238,568805 | 0,231663664 | 0,078100255 | 2,966234414 | 0,003014706 | 0,01040448 |
| ENSG00000268516.2 | AC020915.3 | lncRNA | 512,6562512 | -0,430675726 | 0,113117258 | -3,807338805 | 0,00014047 | 0,000650318 |
| ENSG00000268573.1 | AC011815.1 | lncRNA | 873,6715649 | 0,21906965 | 0,090303262 | 2,425932857 | 0,015269095 | 0,043063784 |
| ENSG00000268592.3 | AL355312.3 | lncRNA | 14,68098885 | -1,597467104 | 0,673853651 | -2,370643983 | 0,017757126 | 0,049018292 |
| ENSG00000268606.5 | MAGEA2 | protein_coding | 5189,056705 | 0,187229392 | 0,063207283 | 2,962149026 | 0,003054999 | 0,010530824 |
| ENSG00000268654.2 | MIMT1 | lncRNA | 103,1297982 | -0,703564769 | 0,23198891 | -3,032751729 | 0,002423349 | 0,008563133 |
| ENSG00000268670.1 | AC016586.2 | lncRNA | 25,01712283 | -1,540831279 | 0,492685818 | -3,127411474 | 0,001763529 | 0,006467136 |
| ENSG00000268798.1 | AC027307.3 | lncRNA | 53,49028817 | -1,057265409 | 0,316615316 | -3,33927437 | 0,000839976 | 0,003313531 |
| ENSG00000268858.2 | AL118506.1 | lncRNA | 1225,720211 | 0,416372943 | 0,077628047 | 5,363692127 | 8,15E-08 | 6,06E-07 |
| ENSG00000268912.1 | AC012313.5 | lncRNA | 464,0197929 | 0,36076724 | 0,121959991 | 2,95807861 | 0,003095632 | 0,010655136 |

| | | | | | | | | |
|---|---|---|---|---|---|---|---|---|
| ENSG00000268916.6 | CSAG3 | protein_coding | 3116,171557 | 0,206691008 | 0,065774648 | 3,142411448 | 0,001675624 | 0,006177988 |
| ENSG00000269001.2 | AC092070.2 | transcribed_unproc | 72,06696771 | 1,351521124 | 0,349757735 | 3,86416364 | 0,00011147 | 0,000525574 |
| ENSG00000269038.1 | AP001462.1 | lncRNA | 30,75429133 | -1,295221619 | 0,448046969 | -2,890816609 | 0,003842423 | 0,012933915 |
| ENSG00000269044.2 | AC024075.2 | lncRNA | 269,1934141 | -1,106840203 | 0,164204997 | -6,740600003 | 1,58E-11 | 1,77E-10 |
| ENSG00000269176.2 | AP001160.3 | lncRNA | 210,6190134 | 0,406502116 | 0,162007776 | 2,509151885 | 0,012102144 | 0,035244812 |
| ENSG00000269293.3 | ZSCAN16-AS1 | lncRNA | 1223,029998 | -0,608416581 | 0,083733045 | -7,266146612 | 3,70E-13 | 4,84E-12 |
| ENSG00000269335.5 | IKBKG | protein_coding | 989,3076364 | 0,252687687 | 0,096684983 | 2,613515342 | 0,008961606 | 0,02725166 |
| ENSG00000269556.8 | TMEM185A | protein_coding | 2666,669038 | 0,214769868 | 0,081890826 | 2,622636469 | 0,008725232 | 0,02663361 |
| ENSG00000269713.7 | NBPF9 | protein_coding | 7239,916217 | 0,155038232 | 0,050245985 | 3,085584528 | 0,002031525 | 0,007324618 |
| ENSG00000269743.3 | SLC25A53 | protein_coding | 4665,849725 | -0,310961069 | 0,057173688 | -5,43888418 | 5,36E-08 | 4,10E-07 |
| ENSG00000269755.1 | AC008758.6 | protein_coding | 15,20436304 | 2,101065904 | 0,866632179 | 2,424403289 | 0,015333567 | 0,043234781 |
| ENSG00000269821.1 | KCNQ1OT1 | lncRNA | 1971,827234 | 0,189177276 | 0,072466179 | 2,610559564 | 0,009039423 | 0,027454914 |
| ENSG00000269893.8 | SNHG8 | lncRNA | 2995,71168 | -0,471524241 | 0,058429018 | -8,070035366 | 7,03E-16 | 1,12E-14 |
| ENSG00000269911.1 | FAM226B | lncRNA | 726,5061709 | 0,285736009 | 0,093088753 | 3,069500874 | 0,002144168 | 0,007677867 |
| ENSG00000269973.1 | AC010969.2 | lncRNA | 358,6256292 | -0,404322652 | 0,129414459 | -3,124246344 | 0,001782611 | 0,006529667 |
| ENSG00000270006.2 | AC010531.6 | lncRNA | 28,18059105 | -1,086812252 | 0,437130889 | -2,486239886 | 0,012910091 | 0,037250767 |
| ENSG00000270025.2 | BMS1P7 | transcribed_unproc | 289,8737055 | 0,46621064 | 0,173350887 | 2,689404404 | 0,007157965 | 0,022386915 |
| ENSG00000270055.1 | AC127502.2 | lncRNA | 1521,713357 | 0,299777692 | 0,087647272 | 3,420274075 | 0,000625581 | 0,002538067 |
| ENSG00000270084.2 | GAS5-AS1 | lncRNA | 60,25845255 | -1,616084235 | 0,307693129 | -5,252259743 | 1,50E-07 | 1,09E-06 |
| ENSG00000270149.5 | AL591806.3 | protein_coding | 1352,029897 | 1,245631028 | 0,227227967 | 5,481856159 | 4,21E-08 | 3,25E-07 |
| ENSG00000270175.1 | AC023509.3 | lncRNA | 180,6900679 | -0,705739946 | 0,175349696 | -4,024757173 | 5,70E-05 | 0,00028327 |
| ENSG00000270231.4 | NBPF8 | transcribed_unproc | 4808,024473 | -0,184496516 | 0,053185793 | -3,468906024 | 0,000522582 | 0,002154754 |
| ENSG00000270276.2 | HIST2H4B | protein_coding | 1198,984817 | 0,329859255 | 0,122355867 | 2,695900603 | 0,007019861 | 0,022031519 |
| ENSG00000270362.1 | HMGN3-AS1 | lncRNA | 231,8715975 | 0,445156605 | 0,16993139 | 2,61962551 | 0,008802637 | 0,026829867 |
| ENSG00000270504.1 | AL391422.4 | lncRNA | 59,53797276 | 1,04538288 | 0,310830678 | 3,363190811 | 0,000770471 | 0,003062969 |
| ENSG00000270580.5 | AC139256.2 | lncRNA | 1491,385797 | 1,083411314 | 0,087617678 | 12,36521368 | 4,03E-35 | 1,55E-33 |
| ENSG00000270647.6 | TAF15 | protein_coding | 30501,52839 | -0,112778243 | 0,041518234 | -2,716354547 | 0,006600519 | 0,020843306 |
| ENSG00000270673.1 | YTHDF3-AS1 | lncRNA | 108,8021624 | -0,763492824 | 0,24019727 | -3,178607418 | 0,001479844 | 0,005532271 |
| ENSG00000270800.3 | RPS10-NUDT3 | protein_coding | 2123,484745 | 0,294301846 | 0,077430627 | 3,800845438 | 0,000144203 | 0,00066664 |
| ENSG00000270816.6 | LINC00221 | lncRNA | 1233,858005 | -0,278383604 | 0,079050144 | -3,521607805 | 0,000428938 | 0,001804667 |
| ENSG00000270882.2 | HIST2H4A | protein_coding | 829,2992663 | 1,513867194 | 0,097133845 | 15,58537295 | 9,15E-55 | 5,82E-53 |
| ENSG00000271079.1 | CTAGE15 | protein_coding | 94,16353098 | 0,730740589 | 0,29048297 | 2,515605608 | 0,011882807 | 0,034686825 |
| ENSG00000271092.5 | TMEM56-RWDD | protein_coding | 351,8295757 | 0,498886073 | 0,141136958 | 3,534765653 | 0,000408137 | 0,001723594 |
| ENSG00000271121.2 | NUDT4P2 | processed_pseudog | 1592,547361 | 0,364447638 | 0,09211074 | 3,956624796 | 7,60E-05 | 0,000369396 |
| ENSG00000271601.4 | LIX1L | protein_coding | 2091,404317 | -0,457219155 | 0,061978732 | -7,377033008 | 1,62E-13 | 2,18E-12 |
| ENSG00000271614.1 | ATP2B1-AS1 | lncRNA | 492,1172185 | -0,286036596 | 0,116509055 | -2,455058942 | 0,014086149 | 0,040236691 |

| | | | | | | | | |
|---|---|---|---|---|---|---|---|---|
| ENSG00000271646.1 | AC099343.3 | lncRNA | 786,8578392 | 2,12787772 | 0,104848127 | 20,29485682 | 1,43E-91 | 1,52E-89 |
| ENSG00000271725.1 | AC103858.1 | lncRNA | 13,19094128 | 2,517579064 | 0,746020758 | 3,37467696 | 0,000739024 | 0,002951506 |
| ENSG00000271824.1 | SMIM32 | protein_coding | 31,65553634 | -1,388347625 | 0,473769379 | -2,930429207 | 0,003384941 | 0,011558198 |
| ENSG00000271860.7 | AL589740.1 | lncRNA | 183,3212975 | 0,635505647 | 0,177859736 | 3,573072027 | 0,000352818 | 0,001510092 |
| ENSG00000271862.1 | AC104118.1 | lncRNA | 68,77362374 | -0,904787066 | 0,287958736 | -3,142071946 | 0,001677568 | 0,006182119 |
| ENSG00000271869.1 | AC026979.2 | lncRNA | 59,7048789 | -1,093209306 | 0,303457168 | -3,602516007 | 0,000315152 | 0,001364184 |
| ENSG00000271966.1 | AC021321.1 | lncRNA | 51,92351943 | 0,935814558 | 0,317041157 | 2,95171317 | 0,003160163 | 0,010846617 |
| ENSG00000271976.1 | AC012467.2 | lncRNA | 1175,97298 | -0,554768383 | 0,095100149 | -5,833517503 | 5,43E-09 | 4,68E-08 |
| ENSG00000272008.1 | AL139274.2 | lncRNA | 124,6029397 | -0,72674797 | 0,207104134 | -3,509094458 | 0,000449635 | 0,00188716 |
| ENSG00000272030.1 | AL162258.2 | lncRNA | 80,25846333 | 0,869428816 | 0,280783832 | 3,096434755 | 0,00195863 | 0,007091329 |
| ENSG00000272079.2 | AC004233.2 | lncRNA | 761,6501749 | -0,488132068 | 0,091396955 | -5,340791338 | 9,25E-08 | 6,84E-07 |
| ENSG00000272106.1 | AL691432.2 | lncRNA | 498,4018928 | -1,095727365 | 0,141036485 | -7,769105752 | 7,90E-15 | 1,18E-13 |
| ENSG00000272144.1 | AC025171.4 | lncRNA | 16,27668085 | 2,285615211 | 0,644329183 | 3,54727874 | 0,000389233 | 0,001651505 |
| ENSG00000272146.6 | ARF4-AS1 | lncRNA | 84,45088932 | -1,037644641 | 0,296774447 | -3,496408307 | 0,000471567 | 0,001967468 |
| ENSG00000272155.1 | AC055822.1 | lncRNA | 13,49465385 | -1,821275232 | 0,699820227 | -2,602490128 | 0,009254948 | 0,028037634 |
| ENSG00000272163.1 | AF106564.1 | lncRNA | 610,7629919 | -0,466439968 | 0,101884025 | -4,578146257 | 4,69E-06 | 2,76E-05 |
| ENSG00000272267.2 | AC021242.3 | lncRNA | 86,9641772 | 0,836554898 | 0,241402866 | 3,465389253 | 0,000529465 | 0,002180666 |
| ENSG00000272273.1 | IER3-AS1 | lncRNA | 254,827344 | -1,062276724 | 0,149877238 | -7,08764543 | 1,36E-12 | 1,71E-11 |
| ENSG00000272288.6 | AL451165.2 | lncRNA | 178,493904 | -0,955170414 | 0,181667692 | -5,257789133 | 1,46E-07 | 1,06E-06 |
| ENSG00000272316.1 | AL021368.2 | lncRNA | 702,6313237 | -0,474278002 | 0,112491804 | -4,21611162 | 2,49E-05 | 0,000130899 |
| ENSG00000272325.2 | NUDT3 | protein_coding | 12561,93293 | -1,249545558 | 0,071559763 | -17,46156647 | 2,81E-68 | 2,23E-66 |
| ENSG00000272333.5 | KMT2B | protein_coding | 5364,474163 | 0,131355726 | 0,054295478 | 2,419275604 | 0,015551452 | 0,043728625 |
| ENSG00000272335.1 | AC093297.2 | lncRNA | 610,6766022 | -0,649584959 | 0,123319172 | -5,267509892 | 1,38E-07 | 1,01E-06 |
| ENSG00000272356.1 | AL080317.3 | lncRNA | 82,95860762 | -0,741738813 | 0,268456512 | -2,762975677 | 0,005727703 | 0,018402066 |
| ENSG00000272410.5 | AC022384.1 | protein_coding | 403,9263431 | 0,664896542 | 0,165257383 | 4,023399926 | 5,74E-05 | 0,000284657 |
| ENSG00000272416.1 | AC025175.1 | lncRNA | 30,6099272 | -0,996649002 | 0,413046813 | -2,412920207 | 0,015825284 | 0,044422234 |
| ENSG00000272419.6 | LINC01145 | transcribed_unproc | 1440,709333 | 0,878390254 | 0,113930732 | 7,709862297 | 1,26E-14 | 1,84E-13 |
| ENSG00000272463.1 | AL357054.4 | lncRNA | 103,356913 | 4,131349066 | 0,305993514 | 13,50142691 | 1,53E-41 | 7,16E-40 |
| ENSG00000272501.1 | AL662844.4 | lncRNA | 647,3257639 | 0,410718358 | 0,10591896 | 3,877666061 | 0,000105463 | 0,000499588 |
| ENSG00000272514.6 | CFAP206 | protein_coding | 434,6523505 | 0,453307834 | 0,118257723 | 3,833219701 | 0,000126477 | 0,00059125 |
| ENSG00000272556.2 | GTF2IP13 | transcribed_unproc | 262,3194633 | 0,801225433 | 0,148671579 | 5,38923067 | 7,08E-08 | 5,31E-07 |
| ENSG00000272572.1 | AL138762.1 | lncRNA | 94,98186648 | -0,61056992 | 0,233653712 | -2,613140252 | 0,008971448 | 0,027274219 |
| ENSG00000272602.6 | ZNF595 | protein_coding | 1733,702188 | 0,342106166 | 0,077495014 | 4,414557101 | 1,01E-05 | 5,67E-05 |
| ENSG00000272645.3 | GTF2IP20 | transcribed_unproc | 648,8483606 | 0,615175679 | 0,128657958 | 4,781481744 | 1,74E-06 | 1,09E-05 |
| ENSG00000272655.2 | POLR2J4 | transcribed_unproc | 1696,783716 | -0,356642302 | 0,085410209 | -4,175640214 | 2,97E-05 | 0,000154363 |
| ENSG00000272668.2 | AL590560.2 | lncRNA | 109,843637 | -1,081228893 | 0,234119649 | -4,618274875 | 3,87E-06 | 2,30E-05 |

| | | | | | | | | |
|---|---|---|---|---|---|---|---|---|
| ENSG00000272669.1 | AL021707.6 | lncRNA | 76,32639455 | 1,867968464 | 0,299002319 | 6,247337712 | 4,18E-10 | 4,09E-09 |
| ENSG00000272688.1 | AP005329.3 | lncRNA | 98,62344461 | -0,688105398 | 0,239515337 | -2,872907455 | 0,004067132 | 0,013604857 |
| ENSG00000272692.2 | AC010997.4 | lncRNA | 251,7023355 | 0,376538924 | 0,155504657 | 2,421399656 | 0,015460869 | 0,043533737 |
| ENSG00000272750.1 | AL592148.3 | lncRNA | 63,71262783 | -0,849246112 | 0,307976123 | -2,757506345 | 0,005824408 | 0,018664807 |
| ENSG00000272752.6 | STAG3L5P-PVRIG | lncRNA | 3694,282934 | 0,256000143 | 0,065615193 | 3,901537591 | 9,56E-05 | 0,000456308 |
| ENSG00000272755.1 | AC245297.2 | lncRNA | 60,20561259 | 1,089126838 | 0,325109007 | 3,350035878 | 0,000808011 | 0,003200901 |
| ENSG00000272758.6 | AC083798.2 | lncRNA | 164,3443114 | -0,882810855 | 0,183442534 | -4,812465442 | 1,49E-06 | 9,44E-06 |
| ENSG00000272769.1 | AC097532.2 | lncRNA | 471,609041 | 0,340236434 | 0,114992454 | 2,95877184 | 0,003088677 | 0,010633684 |
| ENSG00000272777.1 | AC019131.2 | lncRNA | 68,69618767 | -1,338427445 | 0,326867827 | -4,094705367 | 4,23E-05 | 0,000214681 |
| ENSG00000272831.1 | AC027644.3 | lncRNA | 213,7124209 | -0,763316223 | 0,178669114 | -4,272233782 | 1,94E-05 | 0,000103567 |
| ENSG00000272842.1 | AL391834.1 | lncRNA | 8,554652661 | -2,348112612 | 0,925704428 | -2,536568412 | 0,011194487 | 0,032951053 |
| ENSG00000272870.3 | AC097534.2 | lncRNA | 482,5380145 | -0,416297137 | 0,144535011 | -2,880251187 | 0,003973585 | 0,013325598 |
| ENSG00000272886.6 | DCP1A | protein_coding | 6176,091282 | 0,613581772 | 0,050015131 | 12,26792282 | 1,35E-34 | 5,08E-33 |
| ENSG00000272909.1 | AL122035.2 | lncRNA | 75,76072281 | -0,799411979 | 0,299175649 | -2,672048953 | 0,007538965 | 0,023399691 |
| ENSG00000272918.1 | AC005070.3 | lncRNA | 21,21538315 | -1,329345522 | 0,541442993 | -2,455190186 | 0,014081007 | 0,040231527 |
| ENSG00000272923.1 | AC092139.1 | lncRNA | 67,3417183 | -0,94414424 | 0,305561728 | -3,089864184 | 0,00200248 | 0,007234965 |
| ENSG00000272941.1 | AC083862.2 | lncRNA | 339,7616532 | 5,998921786 | 0,269132804 | 22,28982012 | 4,64E-110 | 5,87E-108 |
| ENSG00000273015.2 | AC008124.1 | lncRNA | 4241,225773 | -0,671429207 | 0,065120018 | -10,31064228 | 6,31E-25 | 1,64E-23 |
| ENSG00000273032.2 | DGCR9 | lncRNA | 642,3152662 | -0,498973458 | 0,111096727 | -4,491342579 | 7,08E-06 | 4,05E-05 |
| ENSG00000273038.2 | AL365203.2 | lncRNA | 125,5459936 | -1,661853131 | 0,22254495 | -7,467494217 | 8,17E-14 | 1,13E-12 |
| ENSG00000273066.5 | AL355987.4 | lncRNA | 895,3594835 | -0,265150912 | 0,091828143 | -2,887468957 | 0,003883549 | 0,013048934 |
| ENSG00000273080.1 | AC009309.1 | lncRNA | 161,5931824 | -0,609618522 | 0,194268856 | -3,138014688 | 0,001700964 | 0,006258089 |
| ENSG00000273149.1 | AL138963.3 | lncRNA | 169,1381864 | -0,803962977 | 0,195650002 | -4,109189711 | 3,97E-05 | 0,000202199 |
| ENSG00000273154.3 | AL121845.3 | protein_coding | 513,2035384 | -0,353040103 | 0,131199782 | -2,69085891 | 0,007126833 | 0,022308146 |
| ENSG00000273156.2 | AC124016.2 | lncRNA | 105,0259675 | -1,242680997 | 0,271889598 | -4,570535265 | 4,86E-06 | 2,86E-05 |
| ENSG00000273167.1 | AL359736.1 | protein_coding | 141,0071268 | 0,846279972 | 0,241823238 | 3,499580848 | 0,00046599 | 0,001946729 |
| ENSG00000273183.1 | AC093726.2 | lncRNA | 12,54740805 | 1,835484896 | 0,707249426 | 2,595244094 | 0,009452376 | 0,028543539 |
| ENSG00000273213.3 | HIST2H3PS2 | protein_coding | 337,51557 | 1,141753899 | 0,153140509 | 7,455596889 | 8,95E-14 | 1,23E-12 |
| ENSG00000273221.1 | AL355816.2 | lncRNA | 91,78485661 | 0,681925187 | 0,241359618 | 2,825349125 | 0,004722912 | 0,015503578 |
| ENSG00000273270.1 | AC090114.2 | lncRNA | 1964,326499 | -0,401177262 | 0,077405748 | -5,182783862 | 2,19E-07 | 1,55E-06 |
| ENSG00000273274.2 | ZBTB8B | protein_coding | 1859,263986 | -0,703800673 | 0,069225176 | -10,16683105 | 2,79E-24 | 7,01E-23 |
| ENSG00000273294.1 | C1QTNF3-AMAC | protein_coding | 280,1317635 | 0,621098526 | 0,223411408 | 2,780066298 | 0,00543478 | 0,017567954 |
| ENSG00000273320.1 | AC007032.1 | lncRNA | 51,0652251 | 0,91660791 | 0,333005009 | 2,752534901 | 0,005913584 | 0,01891557 |
| ENSG00000273340.1 | MICE | transcribed_unproc | 16,21581774 | 3,932148579 | 0,780178834 | 5,040060567 | 4,65E-07 | 3,17E-06 |
| ENSG00000273382.1 | AL356488.3 | lncRNA | 240,1198633 | -0,407911654 | 0,158241068 | -2,577786282 | 0,009943548 | 0,029783009 |
| ENSG00000273398.6 | AC017083.3 | protein_coding | 128,0899073 | -0,563717869 | 0,23676694 | -2,380897727 | 0,017270506 | 0,04791001 |

| | | | | | | | | |
|---|---|---|---|---|---|---|---|---|
| ENSG00000273454.1 | AC112503.2 | lncRNA | 13,69325865 | -2,67577703 | 0,825229073 | -3,242465781 | 0,001185002 | 0,004513233 |
| ENSG00000273456.1 | AC064836.3 | lncRNA | 100,1260727 | -1,46136646 | 0,256626832 | -5,69451935 | 1,24E-08 | 1,02E-07 |
| ENSG00000273576.1 | AC009283.1 | lncRNA | 114,6584718 | 1,108702206 | 0,232339709 | 4,771901489 | 1,82E-06 | 1,14E-05 |
| ENSG00000273604.1 | EPOP | protein_coding | 226,3344382 | 0,463459513 | 0,165089159 | 2,807328562 | 0,004995426 | 0,016312498 |
| ENSG00000273611.5 | ZNHIT3 | protein_coding | 5762,157044 | -0,16108683 | 0,048528546 | -3,319424209 | 0,000902033 | 0,003529878 |
| ENSG00000273702.1 | AC091271.1 | lncRNA | 398,4902778 | -0,363719794 | 0,122175192 | -2,977034759 | 0,00291051 | 0,010083527 |
| ENSG00000273760.1 | AC245041.1 | lncRNA | 37,97136189 | -1,15386295 | 0,390912788 | -2,951714514 | 0,00316015 | 0,010846617 |
| ENSG00000273783.1 | AL136040.1 | lncRNA | 112,3140553 | -0,542877345 | 0,218002372 | -2,490235957 | 0,012765831 | 0,036910224 |
| ENSG00000273784.4 | AL137058.2 | lncRNA | 232,736149 | 0,490166899 | 0,156323908 | 3,135584987 | 0,001715117 | 0,00630398 |
| ENSG00000273841.5 | TAF9 | protein_coding | 9956,319489 | -0,15156944 | 0,047353155 | -3,200830891 | 0,001370319 | 0,005151756 |
| ENSG00000274012.1 | RN7SL2 | misc_RNA | 2941,811829 | 0,490690122 | 0,07232655 | 6,784370628 | 1,17E-11 | 1,33E-10 |
| ENSG00000274020.4 | LINC01138 | lncRNA | 1038,841052 | 1,52666965 | 0,094884948 | 16,08969273 | 3,01E-58 | 2,04E-56 |
| ENSG00000274070.1 | CASTOR2 | protein_coding | 840,6814019 | -0,249401897 | 0,09372817 | -2,660906512 | 0,007793059 | 0,024108572 |
| ENSG00000274211.6 | SOCS7 | protein_coding | 6004,385805 | -0,940625523 | 0,068909441 | -13,65016911 | 2,01E-42 | 9,73E-41 |
| ENSG00000274253.5 | AC138649.1 | lncRNA | 1885,917346 | -0,283029568 | 0,067387652 | -4,200021234 | 2,67E-05 | 0,000139481 |
| ENSG00000274265.5 | AC245297.3 | lncRNA | 881,7104474 | 0,617637592 | 0,093346145 | 6,616637387 | 3,67E-11 | 3,99E-10 |
| ENSG00000274276.4 | CBSL | protein_coding | 227,0280029 | 1,525125507 | 0,210722157 | 7,237613388 | 4,57E-13 | 5,93E-12 |
| ENSG00000274312.1 | AC015722.2 | lncRNA | 65,57304487 | 0,737087406 | 0,306677876 | 2,403458039 | 0,016240831 | 0,045445656 |
| ENSG00000274419.6 | TBC1D3D | protein_coding | 89,29494703 | -0,821940579 | 0,339093116 | -2,423937675 | 0,01535324 | 0,04327399 |
| ENSG00000274444.1 | AC078909.2 | lncRNA | 102,9143465 | -0,585603643 | 0,246403175 | -2,376607542 | 0,017472667 | 0,048387336 |
| ENSG00000274516.2 | FAM74A6 | transcribed_process | 121,2603837 | -0,716043367 | 0,210598505 | -3,400040117 | 0,00067376 | 0,002715913 |
| ENSG00000274528.1 | AC090970.2 | lncRNA | 41,52269947 | 0,95556014 | 0,378937365 | 2,5216836 | 0,011679471 | 0,034155282 |
| ENSG00000274561.1 | AC005332.3 | lncRNA | 160,3722844 | -0,99001742 | 0,197849054 | -5,003902728 | 5,62E-07 | 3,78E-06 |
| ENSG00000274828.1 | AC068473.5 | lncRNA | 911,8888172 | -0,518460495 | 0,088580666 | -5,852975784 | 4,83E-09 | 4,18E-08 |
| ENSG00000274893.1 | AL136317.2 | lncRNA | 87,98489605 | 0,594587725 | 0,249495997 | 2,383155369 | 0,017164948 | 0,04767594 |
| ENSG00000274897.3 | PANO1 | protein_coding | 181,001884 | -0,563469507 | 0,184361198 | -3,056334588 | 0,002240611 | 0,007988897 |
| ENSG00000275023.4 | MLLT6 | protein_coding | 8261,163734 | 0,450452385 | 0,071088977 | 6,336458941 | 2,35E-10 | 2,36E-09 |
| ENSG00000275066.5 | SYNRG | protein_coding | 12196,07715 | 0,285920582 | 0,042873179 | 6,668984809 | 2,58E-11 | 2,84E-10 |
| ENSG00000275074.2 | NUDT18 | protein_coding | 421,3223091 | 0,313878451 | 0,127903373 | 2,454027944 | 0,0141266 | 0,040342 |
| ENSG00000275131.3 | AC241952.1 | transcribed_unproc | 5433,269694 | 0,87032161 | 0,06437109 | 13,52038017 | 1,19E-41 | 5,56E-40 |
| ENSG00000275183.1 | LENG9 | protein_coding | 112,6943256 | 0,75105386 | 0,2221093 | 3,38146066 | 0,000721016 | 0,002887269 |
| ENSG00000275318.1 | AL136981.2 | unprocessed_pseud | 79,72809665 | 0,650548634 | 0,273373315 | 2,379707888 | 0,017326367 | 0,048035373 |
| ENSG00000275325.4 | PDCD6IPP1 | transcribed_unproc | 105,0088325 | 1,347030034 | 0,273764339 | 4,920399936 | 8,64E-07 | 5,65E-06 |
| ENSG00000275342.4 | PRAG1 | protein_coding | 382,8070286 | 0,491846077 | 0,130062105 | 3,781624745 | 0,000155808 | 0,000714419 |
| ENSG00000275395.6 | FCGBP | protein_coding | 2033,10977 | 0,241187727 | 0,079212892 | 3,044803951 | 0,00232832 | 0,008265167 |
| ENSG00000275437.1 | AL121832.3 | lncRNA | 502,8641785 | -0,367425129 | 0,109861606 | -3,344436181 | 0,0008245 | 0,00325705 |

| | | | | | | | | |
|---|---|---|---|---|---|---|---|---|
| ENSG00000275601.1 | AC011330.2 | lncRNA | 111,1712233 | -0,688423745 | 0,23735477 | -2,900399869 | 0,003726869 | 0,012580698 |
| ENSG00000275632.1 | AL035461.2 | lncRNA | 155,854819 | -0,712194538 | 0,249936851 | -2,849497918 | 0,004378829 | 0,014498833 |
| ENSG00000275700.5 | AATF | protein_coding | 6728,79625 | 0,132548088 | 0,046820225 | 2,831000644 | 0,004640263 | 0,015263433 |
| ENSG00000275764.1 | AC092747.4 | lncRNA | 676,6742947 | -0,417240888 | 0,127494339 | -3,272622849 | 0,001065546 | 0,004104803 |
| ENSG00000275854.1 | AC084824.4 | lncRNA | 41,05199113 | -1,110495598 | 0,373431049 | -2,97376343 | 0,002941718 | 0,010180677 |
| ENSG00000275897.1 | AC021491.4 | lncRNA | 13,58008786 | -1,658973476 | 0,671309116 | -2,471251225 | 0,013464119 | 0,038665969 |
| ENSG00000275993.2 | SIK1B | protein_coding | 148,3466314 | 0,805570563 | 0,259425442 | 3,105210326 | 0,001901438 | 0,006907595 |
| ENSG00000276043.5 | UHRF1 | protein_coding | 5433,975779 | -0,573956976 | 0,050386469 | -11,39109347 | 4,63E-30 | 1,47E-28 |
| ENSG00000276115.1 | AC026356.2 | lncRNA | 21,47384927 | -1,523783891 | 0,532065505 | -2,863902803 | 0,004184564 | 0,013949993 |
| ENSG00000276131.1 | AC009118.2 | lncRNA | 79,37280994 | -0,867253651 | 0,258606597 | -3,353563523 | 0,000797781 | 0,003162602 |
| ENSG00000276141.4 | WHAMMP3 | transcribed_unproc | 825,6088321 | 0,675751701 | 0,111452077 | 6,063159347 | 1,33E-09 | 1,23E-08 |
| ENSG00000276166.1 | AC092118.2 | lncRNA | 11,21402789 | -1,934967603 | 0,808102752 | -2,394457387 | 0,016644977 | 0,046403492 |
| ENSG00000276168.1 | RN7SL1 | misc_RNA | 2019,5628 | 0,203707071 | 0,084465884 | 2,411708263 | 0,015877981 | 0,044546612 |
| ENSG00000276334.1 | AL133243.2 | lncRNA | 97,61612261 | 0,669606654 | 0,243710022 | 2,747554848 | 0,006004147 | 0,019169835 |
| ENSG00000276570.1 | AC010327.6 | lncRNA | 361,5807448 | -0,33571541 | 0,130405261 | -2,574400811 | 0,010041387 | 0,030034584 |
| ENSG00000276644.5 | DACH1 | protein_coding | 3317,063266 | -0,252712046 | 0,059902318 | -4,218735644 | 2,46E-05 | 0,000129567 |
| ENSG00000276672.1 | AL161891.1 | lncRNA | 409,9688862 | -0,43636351 | 0,124613936 | -3,50172319 | 0,00046226 | 0,001933296 |
| ENSG00000276718.1 | AC005840.5 | lncRNA | 42,10968633 | 1,949819566 | 0,380047407 | 5,130464069 | 2,89E-07 | 2,02E-06 |
| ENSG00000276805.2 | AL133216.2 | transcribed_unproc | 386,8320485 | 0,533243133 | 0,147407256 | 3,617482269 | 0,000297483 | 0,001292921 |
| ENSG00000276850.5 | AC245041.2 | lncRNA | 12,77237752 | -1,956110128 | 0,704179506 | -2,777857225 | 0,005471866 | 0,017670907 |
| ENSG00000276916.1 | AL442125.2 | lncRNA | 62,29780785 | 0,840749046 | 0,308003432 | 2,729674275 | 0,006339693 | 0,020115612 |
| ENSG00000276975.3 | HYDIN2 | transcribed_unproc | 758,6014392 | 0,464515008 | 0,129662842 | 3,582483607 | 0,000340343 | 0,001462539 |
| ENSG00000277053.4 | GTF2IP1 | transcribed_unproc | 26162,14497 | -0,2636209 | 0,047411685 | -5,560251629 | 2,69E-08 | 2,14E-07 |
| ENSG00000277112.3 | ANKRD20A21P | transcribed_unproc | 53,38005017 | -0,917764522 | 0,321745046 | -2,852458903 | 0,004338242 | 0,014389846 |
| ENSG00000277150.1 | F8A3 | protein_coding | 828,166573 | 0,27144708 | 0,091377404 | 2,970614917 | 0,002972042 | 0,010266681 |
| ENSG00000277203.1 | F8A1 | protein_coding | 807,2528071 | 0,253589716 | 0,103758873 | 2,444029212 | 0,01452425 | 0,041268159 |
| ENSG00000277235.1 | AL035420.3 | lncRNA | 29,58900911 | -1,230216115 | 0,444077484 | -2,770273563 | 0,005600923 | 0,01804626 |
| ENSG00000277245.1 | AC084782.3 | lncRNA | 34,17155275 | -1,389523767 | 0,421754498 | -3,294627022 | 0,000985524 | 0,003825342 |
| ENSG00000277301.1 | AL034550.2 | lncRNA | 112,1220302 | -1,140703604 | 0,231220715 | -4,933397102 | 8,08E-07 | 5,32E-06 |
| ENSG00000277363.5 | SRCIN1 | protein_coding | 1352,266675 | 0,320683825 | 0,095319302 | 3,364311513 | 0,000767349 | 0,003051635 |
| ENSG00000277423.1 | AC069234.5 | lncRNA | 6,733803628 | -2,865339743 | 1,093270472 | -2,620888258 | 0,0087701 | 0,026748806 |
| ENSG00000277443.3 | MARCKS | protein_coding | 36766,15974 | -0,150073823 | 0,060307928 | -2,488459261 | 0,012829794 | 0,037066592 |
| ENSG00000277476.1 | AC005332.5 | lncRNA | 438,8788069 | -0,694126622 | 0,131818865 | -5,265760865 | 1,40E-07 | 1,01E-06 |
| ENSG00000277496.1 | AL357033.4 | lncRNA | 90,76145184 | 0,957863885 | 0,264300269 | 3,624150246 | 0,000289913 | 0,001263434 |
| ENSG00000277511.1 | AC116407.2 | lncRNA | 482,6571945 | 4,943412455 | 0,178957572 | 27,62337694 | 5,83E-168 | 1,04E-165 |
| ENSG00000277586.2 | NEFL | protein_coding | 740,7623295 | -0,546805082 | 0,098645706 | -5,543120945 | 2,97E-08 | 2,35E-07 |

| | | | | | | | | |
|---|---|---|---|---|---|---|---|---|
| ENSG00000277879.1 | AL391988.1 | lncRNA | 159,5600481 | 0,510837759 | 0,190331694 | 2,683934285 | 0,007276141 | 0,022706038 |
| ENSG00000277969.1 | AC006449.6 | lncRNA | 219,5758323 | -0,604937439 | 0,167835426 | -3,604348929 | 0,000312936 | 0,001355897 |
| ENSG00000278041.1 | AL133325.3 | lncRNA | 2372,839026 | -0,284309855 | 0,07099274 | -4,004773647 | 6,21E-05 | 0,000306692 |
| ENSG00000278053.5 | DDX52 | protein_coding | 7977,251402 | 0,167289413 | 0,044577809 | 3,752750932 | 0,000174905 | 0,000793575 |
| ENSG00000278129.2 | ZNF8 | protein_coding | 3091,856949 | 0,26159018 | 0,056854344 | 4,601058821 | 4,20E-06 | 2,49E-05 |
| ENSG00000278133.1 | AC135050.6 | lncRNA | 596,3291479 | -0,919071717 | 0,113410368 | -8,103947922 | 5,32E-16 | 8,55E-15 |
| ENSG00000278238.1 | AL359513.1 | lncRNA | 324,33455 | -1,186499707 | 0,137586527 | -8,623662048 | 6,48E-18 | 1,18E-16 |
| ENSG00000278266.1 | AC079949.2 | lncRNA | 5200,88648 | 0,460693905 | 0,047996457 | 9,59849807 | 8,11E-22 | 1,81E-20 |
| ENSG00000278385.1 | AC121338.2 | lncRNA | 195,3866523 | 0,611626263 | 0,187081406 | 3,269305471 | 0,001078118 | 0,004147912 |
| ENSG00000278493.1 | AC039056.2 | lncRNA | 10,06234234 | 1,991942059 | 0,812445205 | 2,451786343 | 0,014214904 | 0,040573582 |
| ENSG00000278535.5 | DHRS11 | protein_coding | 834,2851984 | -0,285659657 | 0,090602719 | -3,152881716 | 0,001616673 | 0,005981212 |
| ENSG00000278540.5 | ACACA | protein_coding | 20763,76599 | -0,357543191 | 0,043981392 | -8,129419588 | 4,31E-16 | 6,98E-15 |
| ENSG00000278619.5 | MRM1 | protein_coding | 561,59498 | 0,426546264 | 0,118287142 | 3,60602393 | 0,000310924 | 0,001347439 |
| ENSG00000278730.1 | AC005332.6 | lncRNA | 4754,890258 | -0,750553499 | 0,062429793 | -12,02236083 | 2,71E-33 | 9,86E-32 |
| ENSG00000278828.1 | HIST1H3H | protein_coding | 144,9379955 | 0,859153861 | 0,211517834 | 4,061850685 | 4,87E-05 | 0,000244175 |
| ENSG00000278845.5 | MRPL45 | protein_coding | 4961,966453 | -0,182133682 | 0,049294552 | -3,694803443 | 0,000220057 | 0,000979874 |
| ENSG00000278934.1 | AC117489.1 | TEC | 543,1613975 | -0,507067108 | 0,132211702 | -3,835266466 | 0,000125428 | 0,000586957 |
| ENSG00000278962.1 | AC092645.1 | TEC | 165,528583 | -0,657492332 | 0,232027522 | -2,83368251 | 0,004601502 | 0,01514922 |
| ENSG00000278970.2 | HEIH | lncRNA | 1969,135205 | 0,262535184 | 0,069430577 | 3,781261747 | 0,000156036 | 0,000715171 |
| ENSG00000279059.1 | AC007485.2 | TEC | 1716,034176 | 0,203062673 | 0,070809769 | 2,86772117 | 0,004134397 | 0,013807292 |
| ENSG00000279069.1 | AC015813.5 | TEC | 242,0326188 | -0,517125442 | 0,150601788 | -3,43372711 | 0,000595343 | 0,002426862 |
| ENSG00000279145.2 | AC011912.1 | lncRNA | 17,87144494 | -2,27472902 | 0,599427568 | -3,794835507 | 0,000147741 | 0,000682016 |
| ENSG00000279168.2 | AC105052.4 | lncRNA | 1226,456796 | -0,615811204 | 0,087739374 | -7,018641388 | 2,24E-12 | 2,74E-11 |
| ENSG00000279170.2 | TSTD3 | protein_coding | 441,7074308 | -0,372801222 | 0,124573677 | -2,992616355 | 0,002765972 | 0,009634674 |
| ENSG00000279186.1 | FP236315.2 | TEC | 23,95239992 | -1,468227438 | 0,493162556 | -2,977167304 | 0,002909252 | 0,010082272 |
| ENSG00000279192.1 | PWAR5 | TEC | 416,0755226 | -1,155972977 | 0,14228253 | -8,124489883 | 4,49E-16 | 7,26E-15 |
| ENSG00000279207.1 | AC015813.6 | TEC | 1558,567742 | -0,908261456 | 0,089270812 | -10,17422644 | 2,58E-24 | 6,51E-23 |
| ENSG00000279236.1 | AC064801.2 | TEC | 14,0351258 | 2,42744494 | 0,870305021 | 2,78918871 | 0,005284027 | 0,017120794 |
| ENSG00000279249.3 | AC007614.1 | lncRNA | 674,820762 | -0,40656361 | 0,111341166 | -3,651512048 | 0,000260701 | 0,001144998 |
| ENSG00000279267.1 | AL078621.3 | TEC | 149,5219796 | -0,75209428 | 0,188234598 | -3,995515634 | 6,46E-05 | 0,000317812 |
| ENSG00000279294.1 | AC092135.1 | TEC | 255,8772333 | -0,47010115 | 0,145763545 | -3,225094098 | 0,001259313 | 0,004769623 |
| ENSG00000279296.1 | PRAL | TEC | 45,40202331 | 1,784884499 | 0,364695 | 4,894184177 | 9,87E-07 | 6,42E-06 |
| ENSG00000279320.1 | AC069528.2 | TEC | 21,36428201 | 2,479575495 | 0,583017866 | 4,253000875 | 2,11E-05 | 0,000112212 |
| ENSG00000279348.1 | AC012513.3 | TEC | 703,3914378 | -2,171445721 | 0,13867122 | -15,65895013 | 2,89E-55 | 1,85E-53 |
| ENSG00000279407.1 | AC007191.1 | TEC | 574,1021674 | -0,589743453 | 0,11143053 | -5,292476426 | 1,21E-07 | 8,84E-07 |
| ENSG00000279419.1 | AC004925.1 | TEC | 53,20525785 | 0,889541877 | 0,333993589 | 2,663350157 | 0,007736686 | 0,023963818 |

| | | | | | | | | |
|---|---|---|---|---|---|---|---|---|
| ENSG00000279516.2 | FAM230C | lncRNA | 1480,093185 | -0,230781237 | 0,069987112 | -3,297481948 | 0,000975559 | 0,003791241 |
| ENSG00000279579.3 | LINC01666 | lncRNA | 108,8497993 | -0,697096421 | 0,233199636 | -2,989268902 | 0,002796459 | 0,009733338 |
| ENSG00000279712.1 | FP325330.3 | lncRNA | 7,927278946 | -2,27313125 | 0,94333795 | -2,409667977 | 0,015967044 | 0,04477974 |
| ENSG00000279717.1 | AC005336.3 | TEC | 80,02306623 | 1,35381733 | 0,296382091 | 4,567810846 | 4,93E-06 | 2,89E-05 |
| ENSG00000279741.1 | AC007342.8 | TEC | 98,05483837 | -0,663330199 | 0,233052943 | -2,846263992 | 0,004423551 | 0,014625397 |
| ENSG00000279762.3 | AC005899.8 | lncRNA | 6,864901236 | 2,791624756 | 1,105723698 | 2,52470374 | 0,011579586 | 0,033880788 |
| ENSG00000279765.3 | AC013394.1 | protein_coding | 80,92148386 | -1,123681618 | 0,290682633 | -3,865664786 | 0,000110787 | 0,00052257 |
| ENSG00000279861.1 | AC073548.1 | TEC | 189,7495606 | 1,216642526 | 0,189735519 | 6,412307677 | 1,43E-10 | 1,47E-09 |
| ENSG00000280002.1 | AC078925.4 | TEC | 752,9533916 | -0,372856433 | 0,109317256 | -3,410773802 | 0,000647788 | 0,002622022 |
| ENSG00000280007.1 | AC008079.1 | lncRNA | 195,2520511 | 1,330638764 | 0,17990343 | 7,396405768 | 1,40E-13 | 1,90E-12 |
| ENSG00000280010.1 | AP001350.2 | TEC | 49,048688 | -1,348747359 | 0,350994626 | -3,842643901 | 0,000121716 | 0,000571129 |
| ENSG00000280022.1 | AC126544.1 | TEC | 15,86966046 | -1,487292899 | 0,608823789 | -2,442895509 | 0,014569955 | 0,041371909 |
| ENSG00000280027.1 | AC007342.9 | TEC | 93,87838414 | -0,715199008 | 0,24218294 | -2,953135382 | 0,00314564 | 0,010801709 |
| ENSG00000280099.1 | AL603750.1 | TEC | 158,5901016 | 0,549554056 | 0,207675917 | 2,646209841 | 0,00813993 | 0,02505082 |
| ENSG00000280120.1 | AC073857.1 | TEC | 685,9899717 | -0,325296172 | 0,099024871 | -3,284994653 | 0,001019843 | 0,003945001 |
| ENSG00000280128.1 | AL662795.2 | TEC | 1475,798216 | -0,418834706 | 0,091131484 | -4,595938592 | 4,31E-06 | 2,55E-05 |
| ENSG00000280135.1 | AL096816.1 | TEC | 22,33706846 | 3,823139006 | 0,631770377 | 6,051469247 | 1,44E-09 | 1,32E-08 |
| ENSG00000280165.1 | PCDH20 | protein_coding | 1291,448164 | 0,273107791 | 0,09731538 | 2,806419601 | 0,005009541 | 0,016344363 |
| ENSG00000280194.1 | AD000864.1 | TEC | 50,49612103 | 0,900282657 | 0,339205863 | 2,654089315 | 0,007952278 | 0,024550516 |
| ENSG00000280206.1 | AC026401.3 | lncRNA | 261,4962429 | -0,435095037 | 0,154750325 | -2,811593693 | 0,004929673 | 0,01612117 |
| ENSG00000280237.3 | MIR4697HG | TEC | 1807,138327 | -1,317800078 | 0,085975966 | -15,32754025 | 5,01E-53 | 3,08E-51 |
| ENSG00000280279.1 | AC240565.2 | lncRNA | 88,09631618 | 1,187799261 | 0,253432234 | 4,68685156 | 2,77E-06 | 1,69E-05 |
| ENSG00000280351.2 | AC127496.7 | TEC | 56,93604888 | -1,213882678 | 0,346377826 | -3,504504575 | 0,000457458 | 0,001916062 |
| ENSG00000280395.1 | AL034546.1 | lncRNA | 20,10126528 | -1,369328022 | 0,533064571 | -2,568784525 | 0,010205589 | 0,030446337 |
| ENSG00000280433.1 | FP565260.6 | protein_coding | 1267,594138 | 0,207420785 | 0,084383644 | 2,45806858 | 0,01396865 | 0,039966993 |
| ENSG00000280441.3 | FP236383.1 | lncRNA | 146,1543794 | 0,945512254 | 0,249883371 | 3,783814228 | 0,000154443 | 0,000709316 |
| ENSG00000280571.2 | AC006059.2 | protein_coding | 13,3024515 | -3,657296931 | 0,95624551 | -3,824642201 | 0,000130962 | 0,000610569 |
| ENSG00000280634.1 | THRIL | lncRNA | 37,67968656 | -1,133403031 | 0,408866275 | -2,772062896 | 0,005570227 | 0,017960213 |
| ENSG00000280734.3 | LINC01232 | lncRNA | 631,3709586 | 1,11889533 | 0,101844261 | 10,98633659 | 4,45E-28 | 1,30E-26 |
| ENSG00000280758.1 | AL356481.3 | lncRNA | 23,64120469 | -1,382778349 | 0,489335074 | -2,825831258 | 0,00471581 | 0,015482521 |
| ENSG00000280798.1 | LINC00294 | lncRNA | 1315,656667 | -0,262250929 | 0,073377601 | -3,573991571 | 0,00035158 | 0,001506228 |
| ENSG00000280828.1 | AC090114.3 | unprocessed_pseud | 148,8457811 | 1,378491777 | 0,20347041 | 6,774900457 | 1,24E-11 | 1,42E-10 |
| ENSG00000280913.2 | CTSLP3 | transcribed_unproc | 393,07208 | 0,673914762 | 0,150787314 | 4,469306755 | 7,85E-06 | 4,46E-05 |
| ENSG00000280968.1 | AC002059.2 | miRNA | 76,27817341 | -1,012102076 | 0,262139688 | -3,860926534 | 0,000112958 | 0,000532252 |
| ENSG00000280987.4 | MATR3 | protein_coding | 4570,910332 | -0,279939038 | 0,078266493 | -3,576741838 | 0,000347903 | 0,001491423 |
| ENSG00000281100.1 | AC105749.1 | TEC | 84,04883937 | 4,534055867 | 0,387487226 | 11,70117506 | 1,26E-31 | 4,33E-30 |

| | | | | | | | | |
|---|---|---|---|---|---|---|---|---|
| ENSG00000281128.2 | PTENP1-AS | lncRNA | 229,4208651 | -0,548931718 | 0,176275411 | -3,114057232 | 0,001845337 | 0,006727679 |
| ENSG00000281332.1 | LINC00997 | lncRNA | 638,3854703 | -0,525963109 | 0,1077889 | -4,879566518 | 1,06E-06 | 6,88E-06 |
| ENSG00000281468.1 | AC006504.7 | lncRNA | 150,6488802 | -1,050837406 | 0,212009012 | -4,956569519 | 7,17E-07 | 4,76E-06 |
| ENSG00000281501.1 | SEPSECS-AS1 | lncRNA | 980,1331056 | -0,571170912 | 0,095354064 | -5,990000735 | 2,10E-09 | 1,89E-08 |
| ENSG00000281571.2 | AC241585.2 | TEC | 982,8687195 | 0,244300703 | 0,097819152 | 2,497473114 | 0,012508195 | 0,036272628 |
| ENSG00000281649.2 | EBLN3P | lncRNA | 12943,62237 | 0,160796968 | 0,049257056 | 3,264445385 | 0,001096786 | 0,004212175 |
| ENSG00000281731.3 | METTL14-DT | lncRNA | 140,9742441 | -0,724757477 | 0,258015835 | -2,808965111 | 0,004970103 | 0,016234517 |
| ENSG00000281769.1 | LINC01230 | lncRNA | 75,99481245 | 1,770541235 | 0,290521799 | 6,094349006 | 1,10E-09 | 1,02E-08 |
| ENSG00000281880.2 | PAUPAR | lncRNA | 2579,616961 | -0,273754277 | 0,064407457 | -4,250350671 | 2,13E-05 | 0,000113495 |
| ENSG00000282012.1 | AP000357.2 | lncRNA | 50,71169171 | -0,815239058 | 0,342388009 | -2,38103858 | 0,017263904 | 0,047903502 |
| ENSG00000282034.1 | AC106886.5 | protein_coding | 5965,833914 | 0,523456769 | 0,062619641 | 8,359306489 | 6,31E-17 | 1,07E-15 |
| ENSG00000282393.1 | AC016588.2 | lncRNA | 977,2265422 | -0,330322898 | 0,086070912 | -3,837799423 | 0,000124142 | 0,000581179 |
| ENSG00000282851.2 | BISPR | lncRNA | 1161,292612 | 6,908748298 | 0,438961013 | 15,73886541 | 8,19E-56 | 5,31E-54 |
| ENSG00000282872.1 | C1orf232 | protein_coding | 167,1944647 | -0,600414372 | 0,188184077 | -3,190569471 | 0,001419927 | 0,005317993 |
| ENSG00000282936.2 | AC004706.3 | protein_coding | 934,271491 | 0,226933967 | 0,085312481 | 2,660032437 | 0,007813313 | 0,024157947 |
| ENSG00000282965.1 | AP001005.3 | lncRNA | 15,43583592 | 1,746746222 | 0,645453552 | 2,706230708 | 0,006805175 | 0,021426449 |
| ENSG00000283001.2 | AL450426.1 | lncRNA | 34,86432396 | 3,00516959 | 0,607633965 | 4,945690598 | 7,59E-07 | 5,01E-06 |
| ENSG00000283041.1 | AC008038.1 | processed_pseudog | 75,02764354 | -0,954741898 | 0,283287767 | -3,370219295 | 0,000751084 | 0,00299446 |
| ENSG00000283050.2 | GTF2IP12 | transcribed_unproc | 1605,818084 | 0,312702258 | 0,079642398 | 3,926329004 | 8,63E-05 | 0,000416084 |
| ENSG00000283103.3 | AC010642.2 | lncRNA | 958,0575722 | -1,178608149 | 0,089469867 | -13,17324125 | 1,25E-39 | 5,59E-38 |
| ENSG00000283122.1 | HYMAI | lncRNA | 286,8381492 | 0,551108313 | 0,13818094 | 3,988309205 | 6,65E-05 | 0,00032662 |
| ENSG00000283228.1 | AC068547.1 | protein_coding | 383,6888893 | 0,515852118 | 0,188551987 | 2,73586148 | 0,006221721 | 0,019797105 |
| ENSG00000283297.1 | TEX52 | protein_coding | 15,28026682 | -2,858018043 | 0,687807579 | -4,155258142 | 3,25E-05 | 0,000167667 |
| ENSG00000283345.1 | AC127502.3 | unprocessed_pseud | 9,643813936 | 3,356683359 | 1,196902278 | 2,804475704 | 0,005039849 | 0,016431337 |
| ENSG00000283486.2 | FAM95C | protein_coding | 144,9010415 | -0,928044091 | 0,201309211 | -4,610042863 | 4,03E-06 | 2,39E-05 |
| ENSG00000283537.2 | AC073264.3 | transcribed_unproc | 832,917451 | 0,567290248 | 0,124584575 | 4,553454941 | 5,28E-06 | 3,08E-05 |
| ENSG00000283627.1 | AL137785.1 | lncRNA | 304,6540452 | -0,552927014 | 0,145842024 | -3,791273588 | 0,000149877 | 0,000690316 |
| ENSG00000283782.2 | AC116366.3 | protein_coding | 265,4194858 | 1,162795698 | 0,225894928 | 5,147506895 | 2,64E-07 | 1,85E-06 |
| ENSG00000283809.1 | AC007326.4 | protein_coding | 1070,254765 | -0,384127071 | 0,10106331 | -3,800855834 | 0,000144197 | 0,00066664 |
| ENSG00000283886.2 | BX664615.2 | lncRNA | 657,7142685 | -0,296276903 | 0,111933052 | -2,646911678 | 0,008123057 | 0,025005728 |
| ENSG00000283900.1 | TPTEP2-CSNK1E | protein_coding | 21,84432307 | -1,307327024 | 0,523098573 | -2,499198222 | 0,012447466 | 0,036138272 |
| ENSG00000284048.2 | AC073111.3 | lncRNA | 80,36271675 | 1,282049387 | 0,308064076 | 4,161632226 | 3,16E-05 | 0,000163541 |
| ENSG00000284052.1 | AC006460.2 | lncRNA | 68,49343454 | -0,734134765 | 0,296718118 | -2,474182468 | 0,013354147 | 0,038390965 |
| ENSG00000284194.2 | SCO2 | protein_coding | 777,4775546 | 0,472960149 | 0,100272314 | 4,716757111 | 2,40E-06 | 1,47E-05 |
| ENSG00000284292.1 | AC004922.1 | protein_coding | 266,7170909 | -0,701770549 | 0,161975508 | -4,332572 | 1,47E-05 | 8,03E-05 |
| ENSG00000284337.1 | AC013271.1 | protein_coding | 15,8789102 | -4,061268266 | 0,942981415 | -4,306838078 | 1,66E-05 | 8,95E-05 |

| | | | | | | | | |
|---|---|---|---|---|---|---|---|---|
| ENSG00000284395.1 | AL032819.3 | protein_coding | 162,6180228 | -0,532797201 | 0,222994979 | -2,389278915 | 0,016881481 | 0,046981362 |
| ENSG00000284418.2 | AL391117.1 | lncRNA | 19,24436232 | -1,854586505 | 0,627528062 | -2,955384179 | 0,003122799 | 0,010731464 |
| ENSG00000284473.1 | MIR4720 | miRNA | 8,203304923 | -2,712691756 | 0,948974304 | -2,858551328 | 0,004255803 | 0,014160216 |
| ENSG00000284691.1 | AC073111.4 | protein_coding | 1056,455896 | 1,662179557 | 0,097056351 | 17,1259227 | 9,51E-66 | 7,23E-64 |
| ENSG00000284753.1 | EEF1AKMT4 | protein_coding | 482,5700318 | -0,282937265 | 0,119313899 | -2,371368872 | 0,017722334 | 0,048941844 |
| ENSG00000284879.1 | AC133644.3 | lncRNA | 236,3849885 | 0,971873352 | 0,187302991 | 5,188776471 | 2,12E-07 | 1,50E-06 |
| ENSG00000284946.1 | AC068831.7 | protein_coding | 1418,948431 | -0,799394596 | 0,086582989 | -9,232698049 | 2,64E-20 | 5,45E-19 |
| ENSG00000284986.1 | AL449043.2 | transcribed_unproc | 64,69811289 | -1,043847887 | 0,37304852 | -2,798155817 | 0,005139531 | 0,016734516 |
| ENSG00000285053.1 | TBCE | protein_coding | 3612,598 | 0,477752763 | 0,07547713 | 6,329768545 | 2,46E-10 | 2,46E-09 |
| ENSG00000285077.2 | AC091057.6 | protein_coding | 413,8290514 | -0,585829012 | 0,170436561 | -3,437226189 | 0,000587705 | 0,002399971 |
| ENSG00000285081.1 | AC004593.2 | lncRNA | 59,42048752 | 1,3767494 | 0,314493269 | 4,377675246 | 1,20E-05 | 6,65E-05 |
| ENSG00000285219.2 | HULC | lncRNA | 748,8014957 | -0,224954477 | 0,094993083 | -2,368114281 | 0,017879012 | 0,049318467 |
| ENSG00000285258.1 | ATXN7 | protein_coding | 1784,859543 | 1,018305464 | 0,091133458 | 11,17378274 | 5,48E-29 | 1,66E-27 |
| ENSG00000285294.1 | LINC00842 | lncRNA | 121,7121881 | -1,41239777 | 0,242261771 | -5,830048071 | 5,54E-09 | 4,77E-08 |
| ENSG00000285304.1 | Z83844.3 | protein_coding | 559,4869039 | -0,34405863 | 0,10952862 | -3,141266912 | 0,001682186 | 0,006196095 |
| ENSG00000285338.1 | AC091564.7 | lncRNA | 14,25141638 | 2,092314175 | 0,746926355 | 2,801232225 | 0,005090787 | 0,016582999 |
| ENSG00000285399.2 | AC104162.2 | lncRNA | 69,21240946 | 0,785299426 | 0,296487986 | 2,648671997 | 0,008080872 | 0,024899705 |
| ENSG00000285407.1 | AL033530.1 | lncRNA | 175,7049659 | -0,510315718 | 0,18418803 | -2,770623688 | 0,005594905 | 0,01802945 |
| ENSG00000285437.1 | POLR2J3 | protein_coding | 3918,77662 | -0,157144252 | 0,055990985 | -2,806599165 | 0,00500675 | 0,016339993 |
| ENSG00000285508.1 | AL034430.1 | protein_coding | 369,444932 | -10,73954047 | 2,088934789 | -5,141156405 | 2,73E-07 | 1,91E-06 |
| ENSG00000285528.1 | AC097104.1 | protein_coding | 27,6071822 | 1,63709878 | 0,645188067 | 2,537397798 | 0,011167997 | 0,032890285 |
| ENSG00000285541.1 | AC005548.1 | lncRNA | 32,6845101 | -1,121363504 | 0,425619627 | -2,634661168 | 0,008422131 | 0,025802858 |
| ENSG00000285542.1 | AC013717.1 | protein_coding | 327,866919 | -0,359776102 | 0,133110531 | -2,702837255 | 0,00687504 | 0,021625248 |
| ENSG00000285589.1 | AC010422.8 | protein_coding | 1971,43268 | -0,316115146 | 0,075213805 | -4,202887276 | 2,64E-05 | 0,000137886 |
| ENSG00000285602.1 | AC097625.2 | protein_coding | 68,17996192 | 0,957526994 | 0,373239681 | 2,565448005 | 0,010304266 | 0,030704095 |
| ENSG00000285641.1 | AL358472.6 | protein_coding | 135,5289752 | 0,804810607 | 0,302041735 | 2,664567555 | 0,007708738 | 0,023883825 |
| ENSG00000285693.1 | AP002381.2 | lncRNA | 131,413869 | -1,256456684 | 0,222962248 | -5,635288918 | 1,75E-08 | 1,42E-07 |
| ENSG00000285735.2 | AP003062.3 | lncRNA | 132,167553 | 0,613768049 | 0,213632887 | 2,873003583 | 0,004065895 | 0,01360274 |
| ENSG00000285744.1 | AC083837.1 | lncRNA | 728,0444197 | 2,32114949 | 0,113265872 | 20,49292919 | 2,49E-93 | 2,69E-91 |
| ENSG00000285756.2 | BX890604.2 | lncRNA | 14811,56293 | 0,299627256 | 0,047360653 | 6,326501839 | 2,51E-10 | 2,51E-09 |
| ENSG00000285769.2 | AP003100.2 | lncRNA | 44,96979003 | 2,304648538 | 0,401084316 | 5,746045025 | 9,14E-09 | 7,62E-08 |
| ENSG00000285793.1 | AC125232.2 | lncRNA | 565,0452202 | 0,393139382 | 0,120686521 | 3,257525192 | 0,001123883 | 0,004305954 |
| ENSG00000285799.1 | AL645929.3 | transcribed_unproc | 145,705262 | 2,927623901 | 0,218425679 | 13,40329538 | 5,78E-41 | 2,66E-39 |
| ENSG00000285865.1 | AC010285.3 | lncRNA | 904,8918453 | -0,343673097 | 0,087665969 | -3,920256645 | 8,85E-05 | 0,000425161 |
| ENSG00000285884.1 | AL022345.4 | lncRNA | 27,20172346 | -1,18050897 | 0,469431026 | -2,514765544 | 0,011911157 | 0,034753787 |
| ENSG00000285934.1 | AC091074.1 | lncRNA | 54,66216351 | 1,094287393 | 0,318559112 | 3,435115654 | 0,000592301 | 0,002415674 |

| | | | | | | | | |
|---|---|---|---|---|---|---|---|---|
| ENSG00000285943.1 | AC112128.1 | protein_coding | 649,0854338 | 0,329434312 | 0,126457151 | 2,60510624 | 0,009184578 | 0,027835688 |
| ENSG00000285952.1 | AC020663.4 | lncRNA | 65,02549119 | -0,895215219 | 0,349351325 | -2,562507012 | 0,010391949 | 0,030904 |
| ENSG00000285967.1 | NIPBL-DT | lncRNA | 4144,204386 | -0,497921206 | 0,058176829 | -8,558754685 | 1,14E-17 | 2,03E-16 |
| ENSG00000286016.1 | AC079949.4 | lncRNA | 229,0553454 | 0,495548791 | 0,161369527 | 3,070894479 | 0,002134186 | 0,007650642 |
| ENSG00000286021.2 | AC112481.2 | lncRNA | 139,7538572 | 1,033885736 | 0,21283126 | 4,857772001 | 1,19E-06 | 7,63E-06 |
| ENSG00000286051.1 | AC244102.3 | lncRNA | 960,4287378 | -0,204937504 | 0,08243382 | -2,486085258 | 0,012915702 | 0,037262181 |
| ENSG00000286070.1 | GGT1 | protein_coding | 2714,940719 | 0,388960316 | 0,067859337 | 5,731861415 | 9,93E-09 | 8,25E-08 |
| ENSG00000286086.1 | AC099661.1 | lncRNA | 91,82996099 | -0,603349224 | 0,253037415 | -2,384426921 | 0,017105745 | 0,047529097 |
| ENSG00000286095.1 | SPEGNB | protein_coding | 114,0501095 | -0,643053794 | 0,232985729 | -2,760056579 | 0,005779135 | 0,01854613 |
| ENSG00000286104.1 | AC016629.3 | lncRNA | 293,2822047 | -0,47549211 | 0,148676693 | -3,198161734 | 0,001383067 | 0,00519637 |
| ENSG00000286112.1 | AL441992.2 | protein_coding | 473,2047396 | -0,373320384 | 0,146205235 | -2,553399572 | 0,010667704 | 0,031619613 |
| ENSG00000286147.1 | AC092445.1 | lncRNA | 2282,260681 | 0,265556837 | 0,063477523 | 4,183478284 | 2,87E-05 | 0,000149271 |
| ENSG00000286176.2 | AC116317.1 | lncRNA | 744,5654029 | 1,592610964 | 0,113973464 | 13,97352421 | 2,26E-44 | 1,15E-42 |
| ENSG00000286181.1 | AC232271.3 | lncRNA | 305,9606166 | 0,472564848 | 0,159377876 | 2,965059274 | 0,003026246 | 0,010441106 |
| ENSG00000286190.1 | AC055839.2 | protein_coding | 12649,6539 | -0,307219056 | 0,050913471 | -6,034140901 | 1,60E-09 | 1,46E-08 |
| ENSG00000286193.2 | AC025449.2 | lncRNA | 64,46421794 | -0,813953887 | 0,326940624 | -2,489607679 | 0,012788417 | 0,036966031 |
| ENSG00000286231.1 | AL445423.2 | protein_coding | 105,2072005 | 0,690669381 | 0,279452805 | 2,471506347 | 0,013454516 | 0,038643321 |
| ENSG00000286235.1 | AL035461.3 | protein_coding | 81,70492149 | 7,641686661 | 2,856965833 | 2,674756055 | 0,007478364 | 0,02324044 |
| ENSG00000286237.1 | ARMCX5-GPRASP2 | protein_coding | 264,5284043 | -0,51938924 | 0,179337538 | -2,896154623 | 0,003777662 | 0,012735791 |
| ENSG00000286257.2 | AL137100.3 | lncRNA | 53,81140031 | 1,361872839 | 0,320256687 | 4,252441539 | 2,11E-05 | 0,000112467 |
| ENSG00000286292.1 | AC099343.4 | lncRNA | 16,3801625 | 2,570237232 | 0,707269787 | 3,634026617 | 0,000279032 | 0,001217904 |
| ENSG00000286340.1 | AL356776.2 | lncRNA | 253,7074385 | -1,182916065 | 0,167756047 | -7,051406414 | 1,77E-12 | 2,19E-11 |
| ENSG00000286375.1 | AL512590.3 | lncRNA | 27,29509555 | 1,436870718 | 0,440119917 | 3,264725506 | 0,001095702 | 0,004210167 |
| ENSG00000286408.1 | AC113410.4 | lncRNA | 133,7675605 | -0,65290494 | 0,248279248 | -2,629720146 | 0,008545518 | 0,026145313 |
| ENSG00000286416.1 | AL442067.3 | lncRNA | 212,1433273 | -0,585466602 | 0,168737336 | -3,469692093 | 0,000521055 | 0,002148852 |
| ENSG00000286422.1 | AL445242.1 | lncRNA | 176,0687864 | -0,601357055 | 0,202445657 | -2,97046162 | 0,002973526 | 0,01027023 |
| ENSG00000286443.1 | AC091178.2 | lncRNA | 23,5762664 | -1,562765849 | 0,4897938 | -3,190660738 | 0,001419479 | 0,005317198 |
| ENSG00000286472.1 | AL035252.5 | lncRNA | 129,8293114 | 0,743513791 | 0,215169187 | 3,455484497 | 0,000549305 | 0,002257083 |
| ENSG00000286522.1 | HIST1H3B | protein_coding | 9,898325635 | -2,254248263 | 0,843356306 | -2,672948843 | 0,007518772 | 0,023353116 |
| ENSG00000286523.1 | AL022151.1 | lncRNA | 165,9940266 | 0,561930068 | 0,200188949 | 2,806998447 | 0,005000548 | 0,016326855 |
| ENSG00000286540.1 | AL096854.1 | lncRNA | 1056,947277 | 0,651688766 | 0,078714471 | 8,279148091 | 1,24E-16 | 2,08E-15 |
| ENSG00000286548.1 | AL596214.1 | lncRNA | 171,7105964 | 0,993313141 | 0,193210421 | 5,141095046 | 2,73E-07 | 1,91E-06 |
| ENSG00000286585.1 | AC104472.4 | lncRNA | 31,87366252 | -1,522931688 | 0,47125811 | -3,231629666 | 0,001230865 | 0,004669734 |
| ENSG00000286787.1 | AL110114.1 | lncRNA | 140,1848322 | -0,911609683 | 0,230664342 | -3,952104932 | 7,75E-05 | 0,000375959 |
| ENSG00000286791.1 | AC079949.6 | lncRNA | 396,5754723 | 0,318020335 | 0,122472729 | 2,596662445 | 0,009413438 | 0,028448856 |
| ENSG00000286802.1 | AC104459.1 | lncRNA | 374,1584284 | 1,746868098 | 0,12938127 | 13,50170773 | 1,53E-41 | 7,15E-40 |

| | | | | | | | | |
|---|---|---|---|---|---|---|---|---|
| ENSG00000286891.1 | AC080132.1 | lncRNA | 6384,252757 | 0,305828804 | 0,048805185 | 6,266317942 | 3,70E-10 | 3,64E-09 |
| ENSG00000286905.1 | AC108488.2 | protein_coding | 348,4349364 | 0,401457264 | 0,149742381 | 2,680986252 | 0,007340553 | 0,022878498 |
| ENSG00000286927.1 | AC103955.1 | lncRNA | 68,43736491 | 0,925941733 | 0,280906237 | 3,296266192 | 0,000979791 | 0,00380703 |
| ENSG00000286985.1 | AC022239.2 | lncRNA | 46,60818444 | -1,359690417 | 0,390957921 | -3,477843377 | 0,000505465 | 0,002092224 |
| ENSG00000287064.1 | AL606500.1 | lncRNA | 197,4817432 | 1,45864678 | 0,171517584 | 8,5043571 | 1,83E-17 | 3,21E-16 |
| ENSG00000287112.1 | AC068787.3 | lncRNA | 874,571695 | 0,308250491 | 0,096500767 | 3,194280223 | 0,0014018 | 0,005256224 |
| ENSG00000287126.1 | AC108488.3 | lncRNA | 2047,889446 | 0,179926041 | 0,074239457 | 2,423590488 | 0,015367924 | 0,043304532 |
| ENSG00000287189.1 | AL121956.6 | lncRNA | 377,939423 | -0,821496714 | 0,139024503 | -5,909006672 | 3,44E-09 | 3,03E-08 |
| ENSG00000287234.1 | AL592211.2 | lncRNA | 15,96010857 | -1,39097167 | 0,578394043 | -2,404885887 | 0,016177516 | 0,04529101 |
| ENSG00000287244.1 | AL020997.5 | lncRNA | 10,32093413 | 1,835132574 | 0,727423013 | 2,522785973 | 0,011642924 | 0,034061681 |
| ENSG00000287255.1 | AC007877.1 | lncRNA | 8,375073485 | -2,726827939 | 0,976224611 | -2,793238266 | 0,005218324 | 0,016940252 |
| ENSG00000287263.1 | AC008875.3 | lncRNA | 210,2518654 | 2,232943704 | 0,192180137 | 11,61901403 | 3,30E-31 | 1,11E-29 |
| ENSG00000287279.1 | AL121932.2 | lncRNA | 32,71610007 | -1,28058516 | 0,448284555 | -2,856634576 | 0,004281585 | 0,014237581 |
| ENSG00000287281.1 | AC012101.2 | lncRNA | 3566,48033 | -0,548505235 | 0,062089152 | -8,834155641 | 1,01E-18 | 1,91E-17 |
| ENSG00000287300.1 | AL080275.1 | lncRNA | 36,59902157 | 1,622975688 | 0,406487732 | 3,992680615 | 6,53E-05 | 0,000321216 |
| ENSG00000287312.1 | AC127526.4 | lncRNA | 35,81499346 | 1,490441495 | 0,406052425 | 3,670564204 | 0,000242016 | 0,001069434 |
| ENSG00000287388.1 | AC068774.1 | lncRNA | 63,54376039 | 0,961345199 | 0,322689935 | 2,979160784 | 0,002890391 | 0,010024623 |
| ENSG00000287391.1 | AC103871.1 | lncRNA | 94,16831721 | -0,701100124 | 0,269860467 | -2,598009751 | 0,009376583 | 0,028348893 |
| ENSG00000287427.1 | AL390236.1 | lncRNA | 9,182786473 | -2,34709896 | 0,953062891 | -2,462690534 | 0,013789888 | 0,039525862 |
| ENSG00000287439.1 | AC092078.3 | lncRNA | 24,48003697 | 2,223016526 | 0,640598155 | 3,470219997 | 0,000520032 | 0,002145026 |
| ENSG00000287485.1 | AL451166.1 | lncRNA | 12,69159014 | 2,5291981 | 0,760476471 | 3,32580717 | 0,000881629 | 0,003459053 |
| ENSG00000287516.1 | AL359764.2 | lncRNA | 219,5539704 | 1,501881108 | 0,164878602 | 9,109011637 | 8,31E-20 | 1,67E-18 |
| ENSG00000287545.1 | AP001994.2 | lncRNA | 48,38944234 | 1,609870994 | 0,374425695 | 4,299574036 | 1,71E-05 | 9,23E-05 |
| ENSG00000287550.1 | AL450345.2 | lncRNA | 148,7087135 | 1,096323715 | 0,196991281 | 5,565341306 | 2,62E-08 | 2,08E-07 |
| ENSG00000287620.1 | AC092053.4 | lncRNA | 155,383691 | 0,962630232 | 0,197039576 | 4,885466423 | 1,03E-06 | 6,69E-06 |
| ENSG00000287733.1 | AC083862.3 | lncRNA | 59,64632616 | 0,890461661 | 0,302970629 | 2,93910226 | 0,003291645 | 0,011266975 |
| ENSG00000287790.1 | AC025756.1 | lncRNA | 110,4615478 | 0,958622579 | 0,230956894 | 4,150655836 | 3,32E-05 | 0,000170879 |
| ENSG00000287839.1 | AL353807.5 | lncRNA | 530,4231278 | 2,901397756 | 0,12222377 | 23,73840839 | 1,45E-124 | 2,02E-122 |
| ENSG00000287879.1 | AC114341.2 | lncRNA | 48,22054654 | 1,278067861 | 0,341648264 | 3,740887914 | 0,000183371 | 0,000827653 |
| ENSG00000287887.1 | AL391560.2 | lncRNA | 16,76279115 | 2,644037009 | 0,63107715 | 4,189720717 | 2,79E-05 | 0,000145492 |
| ENSG00000287910.1 | AC104024.4 | lncRNA | 103,2472237 | -0,720022217 | 0,233753886 | -3,08025774 | 0,002068215 | 0,007442599 |
| ENSG00000287915.1 | AC037433.1 | lncRNA | 1158,804513 | 0,443233153 | 0,094897872 | 4,670633226 | 3,00E-06 | 1,82E-05 |
| ENSG00000287933.1 | AL357141.1 | lncRNA | 246,4747074 | 0,57838386 | 0,155095984 | 3,729199461 | 0,000192089 | 0,000864401 |
| ENSG00000287978.1 | AC245407.2 | lncRNA | 185,7607665 | -0,977336698 | 0,180911566 | -5,402289755 | 6,58E-08 | 4,97E-07 |
| ENSG00000288012.1 | AP003307.1 | lncRNA | 125,7136454 | -0,738329226 | 0,235343086 | -3,137246308 | 0,001705428 | 0,006272464 |

**DEG_significative_IFNγ vs NT 24h**

| Ensembl_gene_ID+IA1:I35 | gene_name | gene_type | baseMean | log2FoldChange | lfcSE | stat | pvalue | padj |
|---|---|---|---|---|---|---|---|---|
| ENSG00000130303.13 | BST2 | protein_coding | 7571,838956 | 14,71547539 | 1,658764932 | 8,87134464 | 7,23E-19 | 3,75E-17 |
| ENSG00000185885.16 | IFITM1 | protein_coding | 2094,79633 | 14,44400025 | 1,489994481 | 9,693995809 | 3,20E-22 | 1,97E-20 |
| ENSG00000138755.6 | CXCL9 | protein_coding | 2098,435669 | 14,24586117 | 2,003407159 | 7,110816741 | 1,15E-12 | 4,17E-11 |
| ENSG00000213886.4 | UBD | protein_coding | 14527,10049 | 13,80367561 | 1,909566034 | 7,228697705 | 4,88E-13 | 1,81E-11 |
| ENSG00000117228.10 | GBP1 | protein_coding | 44680,87298 | 13,71254727 | 0,330699328 | 41,46530134 | 0 | 0 |
| ENSG00000156587.16 | UBE2L6 | protein_coding | 7041,138334 | 13,24661014 | 0,661145351 | 20,0358516 | 2,68E-89 | 5,79E-87 |
| ENSG00000177409.12 | SAMD9L | protein_coding | 795,7860387 | 13,19513672 | 1,453768897 | 9,076502283 | 1,12E-19 | 6,07E-18 |
| ENSG00000225492.6 | GBP1P1 | transcribed_unproc | 692,3947326 | 13,01578854 | 1,193988902 | 10,90109675 | 1,14E-27 | 9,10E-26 |
| ENSG00000169245.6 | CXCL10 | protein_coding | 26555,21761 | 11,9124884 | 0,265720763 | 44,83085278 | 0 | 0 |
| ENSG00000154451.14 | GBP5 | protein_coding | 2868,890101 | 11,81130846 | 1,025684063 | 11,51554254 | 1,10E-30 | 9,38E-29 |
| ENSG00000133321.11 | PLAAT4 | protein_coding | 17595,24819 | 11,74903926 | 0,836685325 | 14,04236325 | 8,58E-45 | 1,04E-42 |
| ENSG00000132530.17 | XAF1 | protein_coding | 835,0609374 | 11,73114528 | 1,244714461 | 9,424768209 | 4,31E-21 | 2,50E-19 |
| ENSG00000243649.9 | CFB | protein_coding | 242,4251817 | 11,45578941 | 1,649847629 | 6,943543881 | 3,82E-12 | 1,34E-10 |
| ENSG00000067066.17 | SP100 | protein_coding | 463,605699 | 11,31223121 | 0,957697404 | 11,81190548 | 3,39E-32 | 3,01E-30 |
| ENSG00000183347.15 | GBP6 | protein_coding | 195,2503156 | 11,09599205 | 1,217790028 | 9,111580648 | 8,12E-20 | 4,41E-18 |
| ENSG00000162654.9 | GBP4 | protein_coding | 20820,26474 | 11,084712 | 0,997348101 | 11,1141857 | 1,07E-28 | 8,68E-27 |
| ENSG00000240065.8 | PSMB9 | protein_coding | 28688,01893 | 10,95178172 | 0,502827302 | 21,78040387 | 3,56E-105 | 9,09E-103 |
| ENSG00000226025.9 | LGALS17A | transcribed_unproc | 3774,974303 | 10,8300434 | 0,646985394 | 16,73923941 | 6,78E-63 | 1,15E-60 |
| ENSG00000288587.1 | AL645933.3 | lncRNA | 493,5898243 | 10,69433803 | 1,344304218 | 7,955296045 | 1,79E-15 | 7,81E-14 |
| ENSG00000237988.5 | OR2I1P | protein_coding | 349,24713 | 10,53728675 | 2,972730722 | 3,544648922 | 0,000393136 | 0,003949285 |
| ENSG00000119917.14 | IFIT3 | protein_coding | 10946,85022 | 10,05498512 | 0,249959965 | 40,22638238 | 0 | 0 |
| ENSG00000162645.13 | GBP2 | protein_coding | 2554,988615 | 9,836135026 | 0,888471286 | 11,07085302 | 1,74E-28 | 1,40E-26 |
| ENSG00000110092.4 | CCND1 | protein_coding | 186,5706728 | 9,634793418 | 1,892495611 | 5,091051923 | 3,56E-07 | 7,05E-06 |
| ENSG00000206337.12 | HCP5 | lncRNA | 7478,971135 | 9,587721773 | 0,232436623 | 41,24875693 | 0 | 0 |
| ENSG00000136514.3 | RTP4 | protein_coding | 220,6524355 | 9,567706299 | 1,059137059 | 9,033492138 | 1,66E-19 | 8,95E-18 |
| ENSG00000090339.9 | ICAM1 | protein_coding | 4832,591479 | 9,530505593 | 0,289466498 | 32,92438214 | 9,84E-238 | 5,72E-235 |
| ENSG00000105205.7 | CLC | protein_coding | 59,69816385 | 9,487655507 | 1,281711555 | 7,402332818 | 1,34E-13 | 5,19E-12 |
| ENSG00000134470.21 | IL15RA | protein_coding | 144,6553328 | 9,061168025 | 1,608128023 | 5,634606134 | 1,75E-08 | 4,14E-07 |
| ENSG00000204642.14 | HLA-F | protein_coding | 5989,360677 | 9,053157435 | 0,237904588 | 38,05373204 | 0 | 0 |
| ENSG00000224666.3 | ETV7-AS1 | lncRNA | 44,87937962 | 9,045394419 | 1,312408703 | 6,8922085 | 5,49E-12 | 1,89E-10 |
| ENSG00000166278.15 | C2 | protein_coding | 621,7832522 | 9,036178366 | 1,385833823 | 6,520390986 | 7,01E-11 | 2,14E-09 |
| ENSG00000168394.11 | TAP1 | protein_coding | 151269,4418 | 8,980267496 | 0,12733753 | 70,5233364 | 0 | 0 |
| ENSG00000128284.19 | APOL3 | protein_coding | 436,5939316 | 8,944846984 | 1,135707893 | 7,876010233 | 3,38E-15 | 1,46E-13 |
| ENSG00000256262.1 | USP30-AS1 | lncRNA | 493,1905527 | 8,935664708 | 0,588644882 | 15,18006012 | 4,79E-52 | 6,92E-50 |

| | | | | | | | | |
|---|---|---|---|---|---|---|---|---|
| ENSG00000272760.1 | AC093726.1 | lncRNA | 39,21451625 | 8,878380965 | 1,329826688 | 6,676344402 | 2,45E-11 | 7,95E-10 |
| ENSG00000261644.2 | AC007728.2 | lncRNA | 58,9634828 | 8,809362095 | 1,269443374 | 6,939547107 | 3,93E-12 | 1,38E-10 |
| ENSG00000135899.19 | SP110 | protein_coding | 1427,300675 | 8,808030215 | 0,673355942 | 13,08079377 | 4,24E-39 | 4,53E-37 |
| ENSG00000241106.8 | HLA-DOB | protein_coding | 38,47244862 | 8,774952607 | 1,334414173 | 6,57588385 | 4,84E-11 | 1,51E-09 |
| ENSG00000166741.8 | NNMT | protein_coding | 628,3403953 | 8,699746348 | 0,594104197 | 14,64346892 | 1,48E-48 | 1,99E-46 |
| ENSG00000197506.8 | SLC28A3 | protein_coding | 32,31753556 | 8,610814128 | 1,358520883 | 6,338374503 | 2,32E-10 | 6,79E-09 |
| ENSG00000203799.13 | CCDC162P | transcribed_unitary | 40,38088838 | 8,574115094 | 1,329819801 | 6,447576649 | 1,14E-10 | 3,40E-09 |
| ENSG00000165949.12 | IFI27 | protein_coding | 572,4686369 | 8,471565884 | 0,877512348 | 9,654070287 | 4,72E-22 | 2,88E-20 |
| ENSG00000187323.12 | DCC | protein_coding | 266,2870093 | 8,469834672 | 0,582206124 | 14,54782821 | 6,03E-48 | 8,04E-46 |
| ENSG00000234745.12 | HLA-B | protein_coding | 70879,78593 | 8,446617971 | 0,169376956 | 49,86875515 | 0 | 0 |
| ENSG00000073861.3 | TBX21 | protein_coding | 28,3119987 | 8,312015329 | 1,383309875 | 6,008787674 | 1,87E-09 | 5,01E-08 |
| ENSG00000197142.10 | ACSL5 | protein_coding | 75,92590956 | 8,277010517 | 2,056339798 | 4,025118089 | 5,69E-05 | 0,00072035 |
| ENSG00000140511.12 | HAPLN3 | protein_coding | 1208,819526 | 8,223941967 | 0,763307516 | 10,77408751 | 4,56E-27 | 3,54E-25 |
| ENSG00000110446.11 | SLC15A3 | protein_coding | 130,9057432 | 8,161805789 | 0,837438487 | 9,746155582 | 1,92E-22 | 1,21E-20 |
| ENSG00000010030.14 | ETV7 | protein_coding | 5554,684006 | 8,057854344 | 0,195187228 | 41,28269258 | 0 | 0 |
| ENSG00000140853.16 | NLRC5 | protein_coding | 6145,977734 | 8,015141725 | 0,23981749 | 33,42183981 | 6,60E-245 | 4,07E-242 |
| ENSG00000006659.13 | LGALS14 | protein_coding | 261,7320757 | 8,011120517 | 0,609792922 | 13,13744424 | 2,01E-39 | 2,17E-37 |
| ENSG00000119922.10 | IFIT2 | protein_coding | 2781,992138 | 7,988828744 | 0,249650808 | 32,00001155 | 1,09E-224 | 5,70E-222 |
| ENSG00000268088.1 | AC093063.1 | transcribed_unproc | 39,13523757 | 7,807609362 | 1,335467944 | 5,846347266 | 5,02E-09 | 1,27E-07 |
| ENSG00000228742.11 | LINC02577 | lncRNA | 45,05639709 | 7,762139272 | 1,270441258 | 6,109797852 | 9,98E-10 | 2,74E-08 |
| ENSG00000250673.2 | REELD1 | protein_coding | 18,0121398 | 7,755620351 | 1,48255159 | 5,231265071 | 1,68E-07 | 3,48E-06 |
| ENSG00000185291.12 | IL3RA | protein_coding | 17,10115904 | 7,657968288 | 1,49804166 | 5,111986196 | 3,19E-07 | 6,35E-06 |
| ENSG00000170989.10 | S1PR1 | protein_coding | 26,64192316 | 7,644743464 | 1,392887394 | 5,488414568 | 4,06E-08 | 9,15E-07 |
| ENSG00000182179.13 | UBA7 | protein_coding | 979,2318198 | 7,607152246 | 0,643621753 | 11,81929015 | 3,10E-32 | 2,76E-30 |
| ENSG00000111335.13 | OAS2 | protein_coding | 34,0807909 | 7,549476342 | 1,35006589 | 5,591931771 | 2,25E-08 | 5,23E-07 |
| ENSG00000000971.16 | CFH | protein_coding | 19,34031081 | 7,506306888 | 1,472667049 | 5,09708348 | 3,45E-07 | 6,83E-06 |
| ENSG00000282851.2 | BISPR | lncRNA | 677,5357973 | 7,488479301 | 0,401583629 | 18,64737197 | 1,33E-77 | 2,60E-75 |
| ENSG00000096996.16 | IL12RB1 | protein_coding | 139,8869495 | 7,455582259 | 0,805406742 | 9,256915635 | 2,10E-20 | 1,17E-18 |
| ENSG00000204287.14 | HLA-DRA | protein_coding | 15,26498281 | 7,410245022 | 1,532792399 | 4,834474013 | 1,33E-06 | 2,40E-05 |
| ENSG00000079263.19 | SP140 | protein_coding | 15,82087552 | 7,409374105 | 1,522966211 | 4,865094217 | 1,14E-06 | 2,08E-05 |
| ENSG00000117226.12 | GBP3 | protein_coding | 5023,886144 | 7,387098375 | 0,566041514 | 13,05045336 | 6,32E-39 | 6,65E-37 |
| ENSG00000271581.1 | AL671883.2 | unprocessed_pseud | 13,680163 | 7,347044527 | 1,564941194 | 4,69477355 | 2,67E-06 | 4,51E-05 |
| ENSG00000286116.1 | AL157394.3 | lncRNA | 13,25438227 | 7,332968599 | 1,575160074 | 4,655379933 | 3,23E-06 | 5,41E-05 |
| ENSG00000173193.15 | PARP14 | protein_coding | 20136,68346 | 7,11944672 | 0,129946531 | 54,78750883 | 0 | 0 |
| ENSG00000137752.24 | CASP1 | protein_coding | 12,61497333 | 7,080261829 | 1,601332058 | 4,421482598 | 9,80E-06 | 0,000149466 |
| ENSG00000137496.18 | IL18BP | protein_coding | 11914,00583 | 7,064996977 | 0,133068809 | 53,09280972 | 0 | 0 |

| | | | | | | | | |
|---|---|---|---|---|---|---|---|---|
| ENSG00000225101.6 | OR52K3P | transcribed_unproc | 21,36287332 | 7,037824755 | 1,370378159 | 5,135680768 | 2,81E-07 | 5,65E-06 |
| ENSG00000179073.6 | TAAR3P | transcribed_unproc | 20,83860107 | 7,028438654 | 1,417783252 | 4,957343547 | 7,15E-07 | 1,34E-05 |
| ENSG00000280893.1 | AC009133.6 | protein_coding | 51,25958081 | 7,025908078 | 2,64257751 | 2,658733017 | 0,007843509 | 0,047185018 |
| ENSG00000197536.11 | IRF1-AS1 | lncRNA | 1239,988114 | 7,003861485 | 0,571883854 | 12,24699987 | 1,74E-34 | 1,64E-32 |
| ENSG00000182326.15 | C1S | protein_coding | 144,4131441 | 6,940538029 | 0,713500882 | 9,727441409 | 2,30E-22 | 1,43E-20 |
| ENSG00000143226.15 | FCGR2A | protein_coding | 12,22582837 | 6,93590278 | 1,613001836 | 4,299996829 | 1,71E-05 | 0,000244744 |
| ENSG00000204267.16 | TAP2 | protein_coding | 48232,69583 | 6,922133179 | 0,128551566 | 53,84713231 | 0 | 0 |
| ENSG00000004468.13 | CD38 | protein_coding | 36,99600514 | 6,90587303 | 2,320120899 | 2,976514298 | 0,002915455 | 0,021267508 |
| ENSG00000163131.11 | CTSS | protein_coding | 5177,673519 | 6,897443902 | 0,455489155 | 15,1429377 | 8,44E-52 | 1,21E-49 |
| ENSG00000234647.2 | AL606970.3 | lncRNA | 10,17544934 | 6,89728153 | 1,677428766 | 4,111817843 | 3,93E-05 | 0,000516216 |
| ENSG00000263179.1 | HNRNPCP4 | processed_pseudog | 19,23021262 | 6,896268389 | 1,448872483 | 4,759748336 | 1,94E-06 | 3,39E-05 |
| ENSG00000204709.4 | LINC01556 | lncRNA | 10,41149028 | 6,893782115 | 1,668474966 | 4,131786364 | 3,60E-05 | 0,000477275 |
| ENSG00000204261.9 | PSMB8-AS1 | lncRNA | 2400,450126 | 6,868719756 | 0,186731722 | 36,78389346 | 3,34E-296 | 2,41E-293 |
| ENSG00000113263.13 | ITK | protein_coding | 16,2033052 | 6,852121959 | 1,438663473 | 4,762838627 | 1,91E-06 | 3,34E-05 |
| ENSG00000100342.21 | APOL1 | protein_coding | 7913,670909 | 6,840372184 | 0,183166292 | 37,34514752 | 3,04E-305 | 2,45E-302 |
| ENSG00000146859.6 | TMEM140 | protein_coding | 6126,790147 | 6,833826416 | 0,185662549 | 36,80778083 | 1,39E-296 | 1,04E-293 |
| ENSG00000273272.2 | U62317.3 | lncRNA | 9,512134929 | 6,798176536 | 1,70636519 | 3,984010326 | 6,78E-05 | 0,000842397 |
| ENSG00000073605.18 | GSDMB | protein_coding | 1075,535785 | 6,788922167 | 0,69499002 | 9,768373607 | 1,54E-22 | 9,82E-21 |
| ENSG00000221963.6 | APOL6 | protein_coding | 24354,48229 | 6,76387111 | 0,13873499 | 48,75389477 | 0 | 0 |
| ENSG00000125347.14 | IRF1 | protein_coding | 37984,71333 | 6,758489716 | 0,153381189 | 44,06335448 | 0 | 0 |
| ENSG00000254139.1 | AC104051.2 | lncRNA | 9,004754566 | 6,730442193 | 1,730891004 | 3,888426352 | 0,000100896 | 0,001201515 |
| ENSG00000141086.18 | CTRL | protein_coding | 30,10436506 | 6,704393488 | 1,13771073 | 5,892880597 | 3,80E-09 | 9,76E-08 |
| ENSG00000169248.13 | CXCL11 | protein_coding | 3098,405239 | 6,699331878 | 0,24444532 | 27,40625954 | 2,31E-165 | 9,12E-163 |
| ENSG00000188282.12 | RUFY4 | protein_coding | 61,99712364 | 6,676071108 | 0,92464796 | 7,220122031 | 5,19E-13 | 1,93E-11 |
| ENSG00000188909.5 | BSX | protein_coding | 9,661067855 | 6,655304669 | 1,707983095 | 3,896586968 | 9,76E-05 | 0,001169323 |
| ENSG00000206341.7 | HLA-H | unprocessed_pseud | 1196,674237 | 6,643263087 | 0,232961761 | 28,51653873 | 7,31E-179 | 3,25E-176 |
| ENSG00000026103.22 | FAS | protein_coding | 938,6265583 | 6,610431912 | 0,280802732 | 23,54119514 | 1,55E-122 | 4,90E-120 |
| ENSG00000187608.10 | ISG15 | protein_coding | 1953,75628 | 6,59937618 | 0,244589455 | 26,98144191 | 2,44E-160 | 9,46E-158 |
| ENSG00000111801.16 | BTN3A3 | protein_coding | 2998,394151 | 6,560874858 | 0,182879909 | 35,87531781 | 7,41E-282 | 5,17E-279 |
| ENSG00000187808.5 | SOWAHD | protein_coding | 32,37061219 | 6,552305539 | 1,158998292 | 5,653421222 | 1,57E-08 | 3,75E-07 |
| ENSG00000204614.9 | TRIM40 | protein_coding | 8,633848667 | 6,532293835 | 1,754714279 | 3,722710822 | 0,000197095 | 0,002178558 |
| ENSG00000106560.11 | GIMAP2 | protein_coding | 658,8307675 | 6,464647432 | 0,282177984 | 22,90982216 | 3,71E-116 | 1,08E-113 |
| ENSG00000257743.8 | MGAM2 | protein_coding | 1743,850262 | 6,460502837 | 0,230132593 | 28,07295895 | 2,10E-173 | 8,95E-171 |
| ENSG00000185338.6 | SOCS1 | protein_coding | 1793,241174 | 6,449163435 | 0,234125608 | 27,54574131 | 4,98E-167 | 2,00E-164 |
| ENSG00000204264.12 | PSMB8 | protein_coding | 28890,63248 | 6,407823189 | 0,154394515 | 41,50291989 | 0 | 0 |
| ENSG00000189233.12 | NUGGC | protein_coding | 267,3050094 | 6,368846682 | 1,200688788 | 5,304327603 | 1,13E-07 | 2,38E-06 |

| | | | | | | | | |
|---|---|---|---|---|---|---|---|---|
| ENSG00000234518.2 | PTGES3P1 | processed_pseudog | 46,7074319 | 6,357675989 | 1,031367955 | 6,16431406 | 7,08E-10 | 1,98E-08 |
| ENSG00000225131.2 | PSME2P2 | processed_pseudog | 6,738028805 | 6,313231524 | 1,878822663 | 3,36020618 | 0,000778843 | 0,007022 |
| ENSG00000234290.2 | AC116366.1 | lncRNA | 19,24689453 | 6,308296442 | 1,328497127 | 4,748445679 | 2,05E-06 | 3,56E-05 |
| ENSG00000167207.14 | NOD2 | protein_coding | 7,635794533 | 6,307645167 | 1,818210021 | 3,46915103 | 0,000522106 | 0,005018497 |
| ENSG00000251598.2 | AC093916.1 | lncRNA | 7,463593504 | 6,30656755 | 1,827859446 | 3,450247537 | 0,000560073 | 0,00532718 |
| ENSG00000157601.14 | MX1 | protein_coding | 4901,065601 | 6,302306205 | 0,160790448 | 39,19577488 | 0 | 0 |
| ENSG00000242574.9 | HLA-DMB | protein_coding | 45,57119314 | 6,221314502 | 1,051262019 | 5,917948514 | 3,26E-09 | 8,51E-08 |
| ENSG00000146006.8 | LRRTM2 | protein_coding | 970,240763 | 6,055549616 | 0,225361659 | 26,87036319 | 4,88E-159 | 1,82E-156 |
| ENSG00000137628.17 | DDX60 | protein_coding | 8315,222238 | 6,024716703 | 0,139959781 | 43,0460569 | 0 | 0 |
| ENSG00000112303.14 | VNN2 | protein_coding | 676,0324538 | 6,022012647 | 0,827610076 | 7,276388753 | 3,43E-13 | 1,29E-11 |
| ENSG00000270792.6 | AL050403.2 | lncRNA | 5,655048518 | 5,970561593 | 1,986958544 | 3,004874767 | 0,002656902 | 0,019640624 |
| ENSG00000248159.1 | HSPA8P11 | processed_pseudog | 6,135348187 | 5,944466865 | 1,942520084 | 3,060182962 | 0,002212018 | 0,016925658 |
| ENSG00000140287.11 | HDC | protein_coding | 5,028497336 | 5,922338216 | 2,059226386 | 2,87600152 | 0,004027478 | 0,027836001 |
| ENSG00000154930.15 | ACSS1 | protein_coding | 110,0657475 | 5,919584398 | 0,557972583 | 10,60909547 | 2,70E-26 | 2,02E-24 |
| ENSG00000229950.1 | TFAP2A-AS1 | lncRNA | 64,43488317 | 5,91446045 | 0,748937616 | 7,897133654 | 2,85E-15 | 1,23E-13 |
| ENSG00000197430.11 | OPALIN | protein_coding | 5,516300528 | 5,908745973 | 2,005242002 | 2,946649815 | 0,003212368 | 0,023040464 |
| ENSG00000137959.16 | IFI44L | protein_coding | 4,939011705 | 5,892861 | 2,071878343 | 2,844211881 | 0,004452143 | 0,030056632 |
| ENSG00000105371.10 | ICAM4 | protein_coding | 10,08426936 | 5,886222559 | 1,637342829 | 3,594984786 | 0,000324411 | 0,003329184 |
| ENSG00000184557.4 | SOCS3 | protein_coding | 33,96798414 | 5,87254454 | 0,9879762 | 5,944014179 | 2,78E-09 | 7,33E-08 |
| ENSG00000211448.13 | DIO2 | protein_coding | 110,2129665 | 5,867295426 | 0,542506599 | 10,81515955 | 2,92E-27 | 2,29E-25 |
| ENSG00000137270.11 | GCM1 | protein_coding | 4,885831893 | 5,833452292 | 2,08049615 | 2,803875552 | 0,005049239 | 0,033170228 |
| ENSG00000235659.1 | AL391987.4 | lncRNA | 4,664683715 | 5,829556613 | 2,110978284 | 2,761542674 | 0,0057529 | 0,036871992 |
| ENSG00000179520.11 | SLC17A8 | protein_coding | 1600,137102 | 5,758973135 | 0,249575454 | 23,07507831 | 8,24E-118 | 2,43E-115 |
| ENSG00000276412.2 | AL591926.2 | lncRNA | 4,895274318 | 5,736262898 | 2,082414473 | 2,754621124 | 0,005876014 | 0,037497715 |
| ENSG00000172183.15 | ISG20 | protein_coding | 3009,449924 | 5,707773967 | 0,214821689 | 26,56982163 | 1,52E-155 | 5,47E-153 |
| ENSG00000148948.8 | LRRC4C | protein_coding | 8,361354732 | 5,696376044 | 1,637993262 | 3,477655358 | 0,00050582 | 0,004893409 |
| ENSG00000168404.13 | MLKL | protein_coding | 611,6482949 | 5,682835746 | 0,283090273 | 20,0742883 | 1,24E-89 | 2,70E-87 |
| ENSG00000168062.10 | BATF2 | protein_coding | 4248,657671 | 5,677347994 | 0,165465163 | 34,31143984 | 5,30E-258 | 3,47E-255 |
| ENSG00000286647.1 | AC008459.1 | lncRNA | 4,524289258 | 5,667074725 | 2,136591669 | 2,652390163 | 0,007992413 | 0,047867513 |
| ENSG00000115956.10 | PLEK | protein_coding | 18,63861976 | 5,646516535 | 1,332241861 | 4,238356939 | 2,25E-05 | 0,000312999 |
| ENSG00000132274.16 | TRIM22 | protein_coding | 182,0770311 | 5,635310364 | 0,500470608 | 11,26002262 | 2,07E-29 | 1,70E-27 |
| ENSG00000258733.6 | LINC02328 | lncRNA | 64,62432057 | 5,61212376 | 0,72401626 | 7,75137807 | 9,09E-15 | 3,81E-13 |
| ENSG00000244509.4 | APOBEC3C | protein_coding | 186,0339212 | 5,553004718 | 0,521588476 | 10,64633321 | 1,81E-26 | 1,37E-24 |
| ENSG00000163898.10 | LIPH | protein_coding | 355,5075201 | 5,543107983 | 0,403227571 | 13,74684764 | 5,32E-43 | 6,22E-41 |
| ENSG00000026950.17 | BTN3A1 | protein_coding | 7929,564535 | 5,532752912 | 0,134376925 | 41,17338548 | 0 | 0 |
| ENSG00000269460.1 | AC005515.1 | lncRNA | 10,9305752 | 5,515862415 | 1,592360532 | 3,463953235 | 0,000532299 | 0,005090763 |

| | | | | | | | | |
|---|---|---|---|---|---|---|---|---|
| ENSG00000198959.12 | TGM2 | protein_coding | 75,6604897 | 5,504745503 | 0,67211402 | 8,190195916 | 2,61E-16 | 1,19E-14 |
| ENSG00000184371.14 | CSF1 | protein_coding | 938,5388788 | 5,504594091 | 0,222987529 | 24,68565895 | 1,52E-134 | 5,07E-132 |
| ENSG00000134321.12 | RSAD2 | protein_coding | 14495,90698 | 5,498719812 | 0,146732865 | 37,47435732 | 2,41E-307 | 2,02E-304 |
| ENSG00000205220.12 | PSMB10 | protein_coding | 6871,181127 | 5,497798538 | 0,137375532 | 40,02021655 | 0 | 0 |
| ENSG00000186470.14 | BTN3A2 | protein_coding | 7575,027326 | 5,48904849 | 0,178487268 | 30,75316557 | 1,11E-207 | 5,40E-205 |
| ENSG00000205863.11 | C1QTNF9B | protein_coding | 12,61299619 | 5,426409753 | 1,411343293 | 3,844854601 | 0,000120624 | 0,001412339 |
| ENSG00000120217.14 | CD274 | protein_coding | 4667,245258 | 5,421804796 | 0,6226875 | 8,707103964 | 3,12E-18 | 1,56E-16 |
| ENSG00000003400.15 | CASP10 | protein_coding | 74,48565467 | 5,419411015 | 1,86532061 | 2,905350955 | 0,003668415 | 0,025745314 |
| ENSG00000213512.3 | GBP7 | protein_coding | 11,36616549 | 5,411628787 | 1,634664619 | 3,31054378 | 0,000931149 | 0,008180277 |
| ENSG00000164308.17 | ERAP2 | protein_coding | 5983,90862 | 5,396419807 | 0,139783655 | 38,60551374 | 0 | 0 |
| ENSG00000287839.1 | AL353807.5 | lncRNA | 1790,177863 | 5,391793435 | 0,162161534 | 33,24952168 | 2,07E-242 | 1,24E-239 |
| ENSG00000008517.18 | IL32 | protein_coding | 155,9525975 | 5,378273329 | 1,265510309 | 4,249885038 | 2,14E-05 | 0,000299187 |
| ENSG00000120341.18 | SEC16B | protein_coding | 446,3434251 | 5,322134596 | 0,288714476 | 18,43390285 | 7,02E-76 | 1,35E-73 |
| ENSG00000261359.2 | PYCARD-AS1 | lncRNA | 6,41117137 | 5,288581253 | 1,780448913 | 2,970363943 | 0,002974471 | 0,021581825 |
| ENSG00000257114.3 | LINC02450 | lncRNA | 12,87442688 | 5,236304036 | 1,495497331 | 3,501379727 | 0,000462856 | 0,004529724 |
| ENSG00000140105.18 | WARS1 | protein_coding | 201127,3287 | 5,223689061 | 0,171555416 | 30,44898953 | 1,24E-203 | 5,88E-201 |
| ENSG00000254838.5 | GVINP1 | transcribed_unproc | 135,8560287 | 5,217628651 | 0,527785751 | 9,885883893 | 4,79E-23 | 3,18E-21 |
| ENSG00000156219.17 | ART3 | protein_coding | 7,833041386 | 5,194582641 | 1,628400445 | 3,189990924 | 0,001422773 | 0,011763722 |
| ENSG00000260918.1 | AC107398.3 | lncRNA | 23,37768568 | 5,185814938 | 1,077078747 | 4,814703618 | 1,47E-06 | 2,63E-05 |
| ENSG00000115415.20 | STAT1 | protein_coding | 177729,4252 | 5,185547182 | 0,108665769 | 47,72015352 | 0 | 0 |
| ENSG00000112782.19 | CLIC5 | protein_coding | 2439,540139 | 5,171453027 | 0,179457302 | 28,81717808 | 1,31E-182 | 5,95E-180 |
| ENSG00000257434.1 | AC073525.1 | lncRNA | 4,80165409 | 5,165809389 | 1,944243551 | 2,656976481 | 0,007884494 | 0,047377121 |
| ENSG00000196684.12 | HSH2D | protein_coding | 29,39249532 | 5,145772763 | 1,018594688 | 5,051835457 | 4,38E-07 | 8,51E-06 |
| ENSG00000234142.2 | LINC01675 | lncRNA | 10,00878255 | 5,136193611 | 1,504012984 | 3,414992866 | 0,000637837 | 0,005937356 |
| ENSG00000145824.13 | CXCL14 | protein_coding | 34,64929662 | 5,111846402 | 1,004639943 | 5,088237274 | 3,61E-07 | 7,14E-06 |
| ENSG00000130234.12 | ACE2 | protein_coding | 179,8578584 | 5,102347704 | 0,516505706 | 9,878589223 | 5,16E-23 | 3,40E-21 |
| ENSG00000277801.1 | AL138478.1 | lncRNA | 5,32735586 | 5,083436345 | 1,904607136 | 2,669020949 | 0,007607271 | 0,046041691 |
| ENSG00000198797.7 | BRINP2 | protein_coding | 258,3713612 | 5,08165801 | 0,357370787 | 14,21956743 | 6,93E-46 | 8,63E-44 |
| ENSG00000184497.13 | TMEM255B | protein_coding | 112,5553395 | 5,059985579 | 0,516409467 | 9,798398168 | 1,14E-22 | 7,32E-21 |
| ENSG00000078081.8 | LAMP3 | protein_coding | 1734,331675 | 5,054955986 | 0,162345427 | 31,13703959 | 7,60E-213 | 3,88E-210 |
| ENSG00000272941.1 | AC083862.1 | lncRNA | 434,6643154 | 5,052113896 | 0,321446062 | 15,71683246 | 1,16E-55 | 1,83E-53 |
| ENSG00000189325.7 | BNIP5 | protein_coding | 100,2652253 | 5,050035848 | 0,725267716 | 6,96299551 | 3,33E-12 | 1,18E-10 |
| ENSG00000204525.17 | HLA-C | protein_coding | 82566,90305 | 5,024808315 | 0,114489423 | 43,8888429 | 0 | 0 |
| ENSG00000135114.12 | OASL | protein_coding | 80,10480231 | 5,011022848 | 0,608084351 | 8,240670628 | 1,71E-16 | 7,97E-15 |
| ENSG00000187105.9 | HEATR4 | protein_coding | 5,696669869 | 5,001678736 | 1,790785827 | 2,793007774 | 0,005222044 | 0,034144913 |
| ENSG00000213928.9 | IRF9 | protein_coding | 4627,229466 | 4,996149089 | 0,223692158 | 22,3349318 | 1,69E-110 | 4,60E-108 |

| | | | | | | | | |
|---|---|---|---|---|---|---|---|---|
| ENSG00000248693.1 | LINC02100 | lncRNA | 17,16040511 | 4,97248063 | 1,193985293 | 4,164607939 | 3,12E-05 | 0,000420707 |
| ENSG00000286239.1 | AC093884.1 | protein_coding | 7,163533975 | 4,939259538 | 1,733068367 | 2,850008478 | 0,004371806 | 0,029657734 |
| ENSG00000235933.1 | LINC01779 | lncRNA | 10,21661447 | 4,903525056 | 1,551559049 | 3,160385717 | 0,001575604 | 0,012834736 |
| ENSG00000125148.7 | MT2A | protein_coding | 4153,062334 | 4,900822799 | 0,13185241 | 37,1690042 | 2,16E-302 | 1,68E-299 |
| ENSG00000002549.12 | LAP3 | protein_coding | 60850,80715 | 4,897383134 | 0,127797505 | 38,3214298 | 0 | 0 |
| ENSG00000285245.1 | AL162417.1 | protein_coding | 21,16112233 | 4,884845478 | 1,39213385 | 3,508890669 | 0,00044998 | 0,00443101 |
| ENSG00000182584.6 | ACTL10 | protein_coding | 16,41043866 | 4,879200005 | 1,398662821 | 3,4884748 | 0,000485785 | 0,004730186 |
| ENSG00000138496.16 | PARP9 | protein_coding | 18125,96554 | 4,850516892 | 0,123726186 | 39,20364037 | 0 | 0 |
| ENSG00000249700.9 | SRD5A3-AS1 | lncRNA | 11,59798256 | 4,843800375 | 1,39648601 | 3,46856348 | 0,000523249 | 0,005023382 |
| ENSG00000168811.7 | IL12A | protein_coding | 11,27859572 | 4,813947212 | 1,517524017 | 3,172237907 | 0,00151269 | 0,012380051 |
| ENSG00000008516.17 | MMP25 | protein_coding | 23,33649051 | 4,756926916 | 1,080001754 | 4,404554806 | 1,06E-05 | 0,000160574 |
| ENSG00000185745.10 | IFIT1 | protein_coding | 2754,196405 | 4,752513444 | 0,147181216 | 32,2902172 | 9,60E-229 | 5,29E-226 |
| ENSG00000239713.9 | APOBEC3G | protein_coding | 2796,516052 | 4,748699568 | 0,147200681 | 32,26003796 | 2,54E-228 | 1,37E-225 |
| ENSG00000173762.8 | CD7 | protein_coding | 19,72291376 | 4,74740359 | 1,108065942 | 4,284405294 | 1,83E-05 | 0,00026059 |
| ENSG00000164136.17 | IL15 | protein_coding | 645,8198008 | 4,714272019 | 0,210135633 | 22,43442467 | 1,82E-111 | 5,00E-109 |
| ENSG00000142089.16 | IFITM3 | protein_coding | 54,15569942 | 4,71343242 | 0,729765458 | 6,458831897 | 1,06E-10 | 3,17E-09 |
| ENSG00000187955.12 | COL14A1 | protein_coding | 73,65330425 | 4,710790187 | 0,721654149 | 6,527767063 | 6,68E-11 | 2,04E-09 |
| ENSG00000111262.6 | KCNA1 | protein_coding | 169,2642117 | 4,697915394 | 0,498908819 | 9,416380735 | 4,67E-21 | 2,70E-19 |
| ENSG00000243811.11 | APOBEC3D | protein_coding | 335,9326129 | 4,670908795 | 0,278804313 | 16,75335918 | 5,35E-63 | 9,26E-61 |
| ENSG00000280362.1 | AC084759.3 | TEC | 7,357704299 | 4,647263311 | 1,740286595 | 2,670401142 | 0,007576068 | 0,045892644 |
| ENSG00000137462.9 | TLR2 | protein_coding | 17,5091184 | 4,645779779 | 1,320070235 | 3,519342878 | 0,000432617 | 0,004286248 |
| ENSG00000285526.1 | AC020907.6 | protein_coding | 11,27655418 | 4,62920876 | 1,658908104 | 2,790515489 | 0,005262418 | 0,034363295 |
| ENSG00000152689.18 | RASGRP3 | protein_coding | 150,6297533 | 4,62555768 | 0,426561712 | 10,84381825 | 2,13E-27 | 1,69E-25 |
| ENSG00000133106.15 | EPSTI1 | protein_coding | 8428,08027 | 4,625307044 | 0,120135504 | 38,50075031 | 0 | 0 |
| ENSG00000169429.11 | CXCL8 | protein_coding | 269,8230641 | 4,605509047 | 0,339318846 | 13,57280653 | 5,81E-42 | 6,61E-40 |
| ENSG00000134873.10 | CLDN10 | protein_coding | 108,8481807 | 4,576968419 | 0,484175206 | 9,453124338 | 3,29E-21 | 1,93E-19 |
| ENSG00000272463.1 | AL357054.4 | lncRNA | 172,6949648 | 4,557926302 | 0,354307466 | 12,86432474 | 7,15E-38 | 7,30E-36 |
| ENSG00000162896.6 | PIGR | protein_coding | 51,40737823 | 4,536294307 | 0,706554455 | 6,420303882 | 1,36E-10 | 4,05E-09 |
| ENSG00000144837.9 | PLA1A | protein_coding | 729,0316043 | 4,532635888 | 0,200389672 | 22,61910932 | 2,81E-113 | 7,95E-111 |
| ENSG00000138435.16 | CHRNA1 | protein_coding | 72,40070186 | 4,532379016 | 0,570687562 | 7,94196215 | 1,99E-15 | 8,68E-14 |
| ENSG00000068079.8 | IFI35 | protein_coding | 9559,632691 | 4,499998874 | 0,145759779 | 30,87270656 | 2,78E-209 | 1,38E-206 |
| ENSG00000089127.14 | OAS1 | protein_coding | 8330,805788 | 4,453504054 | 0,165175918 | 26,96218744 | 4,11E-160 | 1,56E-157 |
| ENSG00000258659.7 | TRIM34 | protein_coding | 68,17093394 | 4,44304859 | 0,671082602 | 6,62071789 | 3,57E-11 | 1,13E-09 |
| ENSG00000163288.14 | GABRB1 | protein_coding | 53,60445751 | 4,413595212 | 0,648690383 | 6,803854855 | 1,02E-11 | 3,43E-10 |
| ENSG00000254087.8 | LYN | protein_coding | 2916,61089 | 4,400235599 | 0,482652535 | 9,116777148 | 7,74E-20 | 4,23E-18 |
| ENSG00000013364.19 | MVP | protein_coding | 2269,471745 | 4,38506821 | 0,194943024 | 22,49410166 | 4,74E-112 | 1,32E-109 |

| | | | | | | | | |
|---|---|---|---|---|---|---|---|---|
| ENSG00000246898.1 | LINC00920 | lncRNA | 14,10765561 | 4,375090538 | 1,246793158 | 3,509074868 | 0,000449668 | 0,00443101 |
| ENSG00000229474.7 | PATL2 | protein_coding | 46,08722815 | 4,374163837 | 0,664436258 | 6,583270836 | 4,60E-11 | 1,44E-09 |
| ENSG00000140044.13 | JDP2 | protein_coding | 59,7171929 | 4,370021303 | 0,562600411 | 7,76754019 | 8,00E-15 | 3,36E-13 |
| ENSG00000237626.2 | AL161629.1 | lncRNA | 20,32104608 | 4,340088514 | 1,062821342 | 4,083554163 | 4,44E-05 | 0,000576357 |
| ENSG00000144035.4 | NAT8 | protein_coding | 32,03353094 | 4,336042598 | 0,827113863 | 5,242376886 | 1,59E-07 | 3,28E-06 |
| ENSG00000243753.5 | HLA-L | transcribed_unproc | 171,0076629 | 4,323837733 | 0,328851142 | 13,14831297 | 1,74E-39 | 1,89E-37 |
| ENSG00000047457.14 | CP | protein_coding | 11,89893989 | 4,321904565 | 1,27414666 | 3,391999289 | 0,000693846 | 0,006354186 |
| ENSG00000177989.13 | ODF3B | protein_coding | 280,3268233 | 4,321252251 | 0,297873132 | 14,50702259 | 1,09E-47 | 1,45E-45 |
| ENSG00000163016.10 | ALMS1P1 | transcribed_unproc | 66,83471096 | 4,320967374 | 1,565026315 | 2,760955092 | 0,00576326 | 0,036908489 |
| ENSG00000229023.1 | RAB1AP1 | processed_pseudog | 8,451904525 | 4,311201889 | 1,471342633 | 2,930114164 | 0,003388375 | 0,024045976 |
| ENSG00000111962.8 | UST | protein_coding | 14,99529656 | 4,310254077 | 1,369553643 | 3,147196241 | 0,001648443 | 0,013344993 |
| ENSG00000231550.1 | PTCHD3P2 | processed_pseudog | 8,073926834 | 4,293302347 | 1,62703498 | 2,638727747 | 0,008321778 | 0,04952144 |
| ENSG00000116663.11 | FBXO6 | protein_coding | 1848,28728 | 4,287348605 | 0,151968172 | 28,21214822 | 4,15E-175 | 1,81E-172 |
| ENSG00000287485.1 | AL451166.1 | lncRNA | 9,135187836 | 4,276824311 | 1,462548556 | 2,924227228 | 0,003453126 | 0,024431474 |
| ENSG00000115414.21 | FN1 | protein_coding | 15208,60592 | 4,270142631 | 0,145929008 | 29,26178077 | 3,18E-188 | 1,48E-185 |
| ENSG00000188820.13 | CALHM6 | protein_coding | 64,33458571 | 4,252981731 | 0,694629872 | 6,122658846 | 9,20E-10 | 2,55E-08 |
| ENSG00000173369.17 | C1QB | protein_coding | 23,609417 | 4,249584676 | 0,97714356 | 4,348987038 | 1,37E-05 | 0,000200929 |
| ENSG00000274419.6 | TBC1D3D | protein_coding | 22,93719848 | 4,248819573 | 1,242661617 | 3,419128357 | 0,000628221 | 0,005863488 |
| ENSG00000196376.11 | SLC35F1 | protein_coding | 1578,370428 | 4,242582036 | 0,177207498 | 23,94132347 | 1,14E-126 | 3,66E-124 |
| ENSG00000287263.1 | AC008875.3 | lncRNA | 363,8487745 | 4,238311707 | 0,25459654 | 16,64716931 | 3,17E-62 | 5,31E-60 |
| ENSG00000181126.13 | HLA-V | transcribed_unproc | 55,30403789 | 4,232684042 | 0,603657808 | 7,011727485 | 2,35E-12 | 8,40E-11 |
| ENSG00000109684.15 | CLNK | protein_coding | 13,80886992 | 4,204920946 | 1,179748945 | 3,564250651 | 0,000364897 | 0,003692181 |
| ENSG00000149403.13 | GRIK4 | protein_coding | 105,9222559 | 4,200257522 | 0,436482042 | 9,622978994 | 6,40E-22 | 3,86E-20 |
| ENSG00000204592.9 | HLA-E | protein_coding | 200099,2028 | 4,19492686 | 0,128077814 | 32,75295476 | 2,76E-235 | 1,56E-232 |
| ENSG00000101974.14 | ATP11C | protein_coding | 81,87427864 | 4,193011783 | 0,549838279 | 7,625900091 | 2,42E-14 | 9,93E-13 |
| ENSG00000204616.11 | TRIM31 | protein_coding | 10,16554911 | 4,181410478 | 1,469682691 | 2,845111059 | 0,004439594 | 0,030000937 |
| ENSG00000189377.9 | CXCL17 | protein_coding | 19,37135447 | 4,170849525 | 1,037043722 | 4,021864687 | 5,77E-05 | 0,000728614 |
| ENSG00000081803.16 | CADPS2 | protein_coding | 362,8074892 | 4,129805999 | 0,331720812 | 12,44964395 | 1,40E-35 | 1,37E-33 |
| ENSG00000116774.12 | OLFML3 | protein_coding | 176,5831312 | 4,094770585 | 0,996826829 | 4,107805355 | 3,99E-05 | 0,000523948 |
| ENSG00000179044.16 | EXOC3L1 | protein_coding | 139,6817807 | 4,089942793 | 0,378871506 | 10,79506569 | 3,63E-27 | 2,84E-25 |
| ENSG00000123685.9 | BATF3 | protein_coding | 48,54912012 | 4,063767628 | 0,599995391 | 6,772998069 | 1,26E-11 | 4,19E-10 |
| ENSG00000170396.9 | ZNF804A | protein_coding | 31,1757076 | 4,031001596 | 0,809604815 | 4,978974335 | 6,39E-07 | 1,21E-05 |
| ENSG00000163840.10 | DTX3L | protein_coding | 12130,49956 | 4,020840665 | 0,119659311 | 33,6024054 | 1,55E-247 | 9,82E-245 |
| ENSG00000135697.10 | BCO1 | protein_coding | 1760,960396 | 4,015748637 | 0,21848284 | 18,38015582 | 1,89E-75 | 3,57E-73 |
| ENSG00000120278.16 | PLEKHG1 | protein_coding | 637,7109502 | 4,012482372 | 0,327771702 | 12,24169857 | 1,86E-34 | 1,75E-32 |
| ENSG00000128335.14 | APOL2 | protein_coding | 16232,1722 | 3,995957824 | 0,113036165 | 35,35114459 | 9,63E-274 | 6,50E-271 |

| | | | | | | | | |
|---|---|---|---|---|---|---|---|---|
| ENSG00000159403.18 | C1R | protein_coding | 359,8760121 | 3,992963868 | 0,264453017 | 15,09895371 | 1,65E-51 | 2,34E-49 |
| ENSG00000107796.13 | ACTA2 | protein_coding | 37,73341104 | 3,987755122 | 0,746108999 | 5,344735324 | 9,05E-08 | 1,95E-06 |
| ENSG00000104312.8 | RIPK2 | protein_coding | 2958,998688 | 3,946417761 | 0,195477199 | 20,18863464 | 1,23E-90 | 2,77E-88 |
| ENSG00000066382.16 | MPPED2 | protein_coding | 9,282113655 | 3,938702915 | 1,399715649 | 2,813930757 | 0,004893977 | 0,032381607 |
| ENSG00000100911.16 | PSME2 | protein_coding | 31853,96338 | 3,932262566 | 0,141085813 | 27,87142444 | 5,93E-171 | 2,43E-168 |
| ENSG00000168899.5 | VAMP5 | protein_coding | 6057,812213 | 3,927664353 | 0,158488523 | 24,78201123 | 1,40E-135 | 4,73E-133 |
| ENSG00000150625.16 | GPM6A | protein_coding | 648,5477478 | 3,906119153 | 0,199737724 | 19,55624139 | 3,65E-85 | 7,57E-83 |
| ENSG00000165409.18 | TSHR | protein_coding | 22,54267776 | 3,895728589 | 0,929999512 | 4,18895767 | 2,80E-05 | 0,0003827 |
| ENSG00000279296.1 | PRAL | TEC | 10,92030651 | 3,874011989 | 1,425514967 | 2,71762281 | 0,006575275 | 0,041055927 |
| ENSG00000023445.16 | BIRC3 | protein_coding | 1912,438808 | 3,872165917 | 0,251377878 | 15,40376565 | 1,54E-53 | 2,29E-51 |
| ENSG00000131203.13 | IDO1 | protein_coding | 281,8901622 | 3,871021732 | 1,183810601 | 3,269967113 | 0,0010756 | 0,009228559 |
| ENSG00000152583.12 | SPARCL1 | protein_coding | 6,892373262 | 3,870052827 | 1,439320759 | 2,688804982 | 0,00717083 | 0,043946523 |
| ENSG00000166573.6 | GALR1 | protein_coding | 10,26171843 | 3,869460541 | 1,310295731 | 2,953120009 | 0,003145796 | 0,022631357 |
| ENSG00000155629.15 | PIK3AP1 | protein_coding | 2121,592148 | 3,86507375 | 0,144906732 | 26,67283768 | 9,73E-157 | 3,57E-154 |
| ENSG00000025708.14 | TYMP | protein_coding | 1211,144916 | 3,836173309 | 0,245236533 | 15,64274811 | 3,72E-55 | 5,77E-53 |
| ENSG00000261618.2 | LINC02605 | lncRNA | 1149,997686 | 3,830932295 | 0,68076702 | 5,62737645 | 1,83E-08 | 4,31E-07 |
| ENSG00000285769.2 | AP003100.2 | lncRNA | 55,36023825 | 3,829325818 | 0,60395308 | 6,340435946 | 2,29E-10 | 6,71E-09 |
| ENSG00000187753.13 | C9orf153 | protein_coding | 35,94637429 | 3,823109027 | 0,812692946 | 4,704247831 | 2,55E-06 | 4,33E-05 |
| ENSG00000093134.15 | VNN3 | protein_coding | 112,9717205 | 3,80440466 | 0,519888458 | 7,317732492 | 2,52E-13 | 9,58E-12 |
| ENSG00000287665.1 | AC092428.1 | lncRNA | 50,14489978 | 3,803236395 | 0,579056695 | 6,56798622 | 5,10E-11 | 1,58E-09 |
| ENSG00000126759.14 | CFP | protein_coding | 9,911366909 | 3,788939911 | 1,307001665 | 2,898955688 | 0,003744078 | 0,02621039 |
| ENSG00000254859.1 | AC067930.3 | lncRNA | 35,27689615 | 3,785476816 | 0,824899445 | 4,589016076 | 4,45E-06 | 7,28E-05 |
| ENSG00000095585.17 | BLNK | protein_coding | 83,39665568 | 3,77736182 | 0,694163792 | 5,441600189 | 5,28E-08 | 1,17E-06 |
| ENSG00000259321.1 | AL136295.2 | lncRNA | 8,748678584 | 3,759288989 | 1,408933541 | 2,668180492 | 0,007626328 | 0,04614369 |
| ENSG00000123609.11 | NMI | protein_coding | 3522,287796 | 3,753676682 | 0,134368634 | 27,93566162 | 9,85E-172 | 4,12E-169 |
| ENSG00000160932.11 | LY6E | protein_coding | 1280,311454 | 3,748973119 | 0,203533672 | 18,41942457 | 9,18E-76 | 1,75E-73 |
| ENSG00000179583.19 | CIITA | protein_coding | 213,9562458 | 3,740643664 | 0,312525357 | 11,96908851 | 5,16E-33 | 4,70E-31 |
| ENSG00000132109.10 | TRIM21 | protein_coding | 2885,174387 | 3,732649258 | 0,189072813 | 19,74186138 | 9,42E-87 | 1,99E-84 |
| ENSG00000104518.11 | GSDMD | protein_coding | 2183,484852 | 3,728394916 | 0,160869923 | 23,17645734 | 7,87E-119 | 2,35E-116 |
| ENSG00000168386.18 | FILIP1L | protein_coding | 18,78186632 | 3,717444926 | 0,9915831 | 3,748999882 | 0,000177541 | 0,001992935 |
| ENSG00000283235.2 | AC139493.2 | lncRNA | 13,62599167 | 3,702396788 | 1,029494173 | 3,596326123 | 0,000322743 | 0,003315323 |
| ENSG00000087085.16 | ACHE | protein_coding | 30,07088299 | 3,701025872 | 0,750200216 | 4,933384172 | 8,08E-07 | 1,50E-05 |
| ENSG00000216490.4 | IFI30 | protein_coding | 20847,93346 | 3,658005643 | 0,167008046 | 21,90317015 | 2,42E-106 | 6,42E-104 |
| ENSG00000078401.7 | EDN1 | protein_coding | 15,39043881 | 3,653477835 | 1,109531462 | 3,29281139 | 0,00099191 | 0,008616446 |
| ENSG00000096968.14 | JAK2 | protein_coding | 6625,175178 | 3,642420292 | 0,147879264 | 24,63104156 | 5,88E-134 | 1,92E-131 |
| ENSG00000157404.16 | KIT | protein_coding | 19,3190071 | 3,641523874 | 1,091416989 | 3,336510163 | 0,000848373 | 0,007541696 |

| | | | | | | | | |
|---|---|---|---|---|---|---|---|---|
| ENSG00000278249.1 | SCARNA2 | scaRNA | 23,22598322 | 3,613398471 | 1,228569893 | 2,941141965 | 0,003270046 | 0,023372625 |
| ENSG00000028137.19 | TNFRSF1B | protein_coding | 46,43548605 | 3,611231222 | 0,639856353 | 5,643815533 | 1,66E-08 | 3,95E-07 |
| ENSG00000196503.5 | ARL9 | protein_coding | 184,3067832 | 3,604931592 | 0,316937732 | 11,37425819 | 5,62E-30 | 4,70E-28 |
| ENSG00000014257.16 | ACP3 | protein_coding | 9,8667827 | 3,59565714 | 1,329938827 | 2,703625962 | 0,006858745 | 0,042494176 |
| ENSG00000041880.14 | PARP3 | protein_coding | 760,7491629 | 3,583992084 | 0,180756283 | 19,82775939 | 1,72E-87 | 3,66E-85 |
| ENSG00000154655.16 | L3MBTL4 | protein_coding | 99,14665362 | 3,576369896 | 0,414872593 | 8,620405298 | 6,67E-18 | 3,29E-16 |
| ENSG00000254154.8 | CRYZL2P-SEC16E | lncRNA | 154,4757296 | 3,557427481 | 0,423966979 | 8,390812633 | 4,83E-17 | 2,30E-15 |
| ENSG00000170162.14 | VGLL2 | protein_coding | 135,5684695 | 3,549874649 | 0,393209551 | 9,027946131 | 1,75E-19 | 9,37E-18 |
| ENSG00000081985.13 | IL12RB2 | protein_coding | 149,6404602 | 3,537625386 | 0,455179912 | 7,771927752 | 7,73E-15 | 3,26E-13 |
| ENSG00000259466.1 | NPM1P47 | processed_pseudog | 20,81533252 | 3,532266628 | 0,982057543 | 3,596802095 | 0,000322153 | 0,003312515 |
| ENSG00000256870.3 | SLC5A8 | protein_coding | 122,9694536 | 3,525251075 | 0,43960947 | 8,019051707 | 1,07E-15 | 4,74E-14 |
| ENSG00000141574.8 | SECTM1 | protein_coding | 5235,749814 | 3,50674774 | 0,439534401 | 7,978323724 | 1,48E-15 | 6,57E-14 |
| ENSG00000175084.12 | DES | protein_coding | 221,0693146 | 3,483642802 | 0,283853495 | 12,27267889 | 1,27E-34 | 1,21E-32 |
| ENSG00000242268.3 | LINC02082 | lncRNA | 42,5393547 | 3,480988806 | 0,790967293 | 4,400926358 | 1,08E-05 | 0,000163048 |
| ENSG00000280285.1 | AC108215.1 | TEC | 16,03147714 | 3,47138324 | 1,126806248 | 3,080727718 | 0,002064954 | 0,015975539 |
| ENSG00000123096.12 | SSPN | protein_coding | 194,9093334 | 3,469332365 | 0,321702089 | 10,78430165 | 4,08E-27 | 3,18E-25 |
| ENSG00000248132.3 | LINC02101 | lncRNA | 22,22848947 | 3,451947101 | 0,967440861 | 3,568122085 | 0,000359549 | 0,003650416 |
| ENSG00000268173.3 | AC007192.1 | protein_coding | 2580,037433 | 3,444506638 | 0,175021971 | 19,68042422 | 3,17E-86 | 6,64E-84 |
| ENSG00000173706.14 | HEG1 | protein_coding | 1931,697497 | 3,43740401 | 0,209593544 | 16,40033342 | 1,90E-60 | 3,14E-58 |
| ENSG00000185880.13 | TRIM69 | protein_coding | 10512,31624 | 3,43619034 | 0,130783536 | 26,27387552 | 3,81E-152 | 1,35E-149 |
| ENSG00000188641.14 | DPYD | protein_coding | 658,2839585 | 3,435888675 | 0,204744981 | 16,78130842 | 3,34E-63 | 5,83E-61 |
| ENSG00000163121.10 | NEURL3 | protein_coding | 434,4494613 | 3,435865622 | 0,264550149 | 12,98757773 | 1,44E-38 | 1,50E-36 |
| ENSG00000141497.14 | ZMYND15 | protein_coding | 64,50179544 | 3,433765518 | 0,562189288 | 6,10784587 | 1,01E-09 | 2,77E-08 |
| ENSG00000152034.11 | MCHR2 | protein_coding | 17,54989131 | 3,429249686 | 0,962435159 | 3,563096851 | 0,000366505 | 0,003704872 |
| ENSG00000090554.13 | FLT3LG | protein_coding | 170,6459762 | 3,428447327 | 0,318481754 | 10,76497251 | 5,04E-27 | 3,89E-25 |
| ENSG00000169715.15 | MT1E | protein_coding | 143,3341987 | 3,42251728 | 0,379632875 | 9,015334296 | 1,96E-19 | 1,05E-17 |
| ENSG00000285744.1 | AC083837.1 | lncRNA | 2450,441398 | 3,421438078 | 0,494531172 | 6,918548859 | 4,56E-12 | 1,59E-10 |
| ENSG00000204632.11 | HLA-G | protein_coding | 35,26225451 | 3,402532387 | 0,761583964 | 4,467704872 | 7,91E-06 | 0,000122546 |
| ENSG00000204622.11 | HLA-J | transcribed_unproc | 226,4493638 | 3,386093124 | 0,32613707 | 10,38242332 | 2,98E-25 | 2,11E-23 |
| ENSG00000170890.14 | PLA2G1B | protein_coding | 14,73682227 | 3,382448031 | 1,09078697 | 3,100924492 | 0,001929175 | 0,015143335 |
| ENSG00000203797.12 | DDO | protein_coding | 118,1671095 | 3,374969316 | 0,381084231 | 8,856229263 | 8,28E-19 | 4,28E-17 |
| ENSG00000173821.19 | RNF213 | protein_coding | 23942,30604 | 3,361250736 | 0,154457247 | 21,76168999 | 5,35E-105 | 1,35E-102 |
| ENSG00000196954.14 | CASP4 | protein_coding | 2294,247718 | 3,358703587 | 0,166773843 | 20,1392708 | 3,34E-90 | 7,37E-88 |
| ENSG00000127324.9 | TSPAN8 | protein_coding | 16,20712689 | 3,34875261 | 0,976987325 | 3,427631582 | 0,000608871 | 0,005703229 |
| ENSG00000277511.1 | AC116407.2 | lncRNA | 127,9565411 | 3,345489818 | 0,3812467 | 8,775131222 | 1,71E-18 | 8,70E-17 |
| ENSG00000286802.1 | AC104459.1 | lncRNA | 541,6376679 | 3,326433268 | 0,238802346 | 13,9296507 | 4,18E-44 | 4,98E-42 |

| | | | | | | | | |
|---|---|---|---|---|---|---|---|---|
| ENSG00000260476.1 | AC104794.4 | lncRNA | 12,40315463 | 3,314710791 | 1,084169394 | 3,057373517 | 0,002232859 | 0,017071109 |
| ENSG00000178685.14 | PARP10 | protein_coding | 3833,558469 | 3,310881515 | 0,151968218 | 21,78667066 | 3,10E-105 | 8,02E-103 |
| ENSG00000147571.5 | CRH | protein_coding | 23,22476378 | 3,302057605 | 1,015179437 | 3,252683697 | 0,001143207 | 0,009725548 |
| ENSG00000256043.4 | CTSO | protein_coding | 4242,627315 | 3,299728694 | 0,170149826 | 19,39307705 | 8,83E-84 | 1,81E-81 |
| ENSG00000244731.8 | C4A | protein_coding | 3241,851226 | 3,291590394 | 0,232458838 | 14,15988489 | 1,62E-45 | 2,01E-43 |
| ENSG00000153993.13 | SEMA3D | protein_coding | 160,4464023 | 3,289755354 | 0,33730681 | 9,753006023 | 1,79E-22 | 1,14E-20 |
| ENSG00000288640.1 | AC005192.1 | protein_coding | 28,98171638 | 3,261651976 | 0,912138976 | 3,575827876 | 0,000349121 | 0,003563557 |
| ENSG00000221869.5 | CEBPD | protein_coding | 642,0083445 | 3,235471469 | 0,197523892 | 16,38015249 | 2,65E-60 | 4,34E-58 |
| ENSG00000112343.11 | TRIM38 | protein_coding | 38,0103704 | 3,234551536 | 0,650143583 | 4,975134143 | 6,52E-07 | 1,23E-05 |
| ENSG00000159860.7 | TCAF2P1 | transcribed_unproc | 44,75480425 | 3,22529949 | 0,599146389 | 5,383157685 | 7,32E-08 | 1,60E-06 |
| ENSG00000237352.4 | LINC01358 | lncRNA | 20,6994505 | 3,199415122 | 0,969814292 | 3,298997705 | 0,000970307 | 0,008463907 |
| ENSG00000153093.20 | ACOXL | protein_coding | 25,86175106 | 3,198714731 | 0,731262044 | 4,374238698 | 1,22E-05 | 0,00018196 |
| ENSG00000152229.18 | PSTPIP2 | protein_coding | 71,2061406 | 3,185571326 | 0,467089277 | 6,820048086 | 9,10E-12 | 3,08E-10 |
| ENSG00000230795.3 | HLA-K | unprocessed_pseud | 67,27557952 | 3,183638459 | 0,492050527 | 6,470145412 | 9,79E-11 | 2,96E-09 |
| ENSG00000049249.9 | TNFRSF9 | protein_coding | 24,70569084 | 3,181788471 | 0,829858351 | 3,834134423 | 0,000126007 | 0,001467164 |
| ENSG00000150551.11 | LYPD1 | protein_coding | 43,64250537 | 3,176980751 | 0,734264085 | 4,326754935 | 1,51E-05 | 0,000219995 |
| ENSG00000121858.11 | TNFSF10 | protein_coding | 6055,477779 | 3,171559374 | 0,152593378 | 20,78438409 | 5,99E-96 | 1,39E-93 |
| ENSG00000268601.1 | AC115522.1 | lncRNA | 26,94330808 | 3,159113811 | 0,810900554 | 3,895809165 | 9,79E-05 | 0,001171491 |
| ENSG00000130487.9 | KLHDC7B | protein_coding | 71,09409114 | 3,157112643 | 0,458559069 | 6,884854873 | 5,78E-12 | 1,99E-10 |
| ENSG00000172403.11 | SYNPO2 | protein_coding | 636,8165185 | 3,153212736 | 0,195283114 | 16,14687864 | 1,19E-58 | 1,91E-56 |
| ENSG00000168658.19 | VWA3B | protein_coding | 207,9753612 | 3,147191753 | 0,470580488 | 6,687892584 | 2,26E-11 | 7,38E-10 |
| ENSG00000281100.1 | AC105749.1 | TEC | 11,86049599 | 3,145928322 | 1,091434986 | 2,882378119 | 0,003946858 | 0,027387296 |
| ENSG00000230521.1 | AL645929.1 | unprocessed_pseud | 152,2085592 | 3,140090866 | 0,35601658 | 8,820069175 | 1,14E-18 | 5,88E-17 |
| ENSG00000163734.4 | CXCL3 | protein_coding | 97,78369239 | 3,134405256 | 0,639176407 | 4,903818763 | 9,40E-07 | 1,73E-05 |
| ENSG00000204952.3 | FBXO47 | protein_coding | 21,94883261 | 3,125185845 | 0,842945788 | 3,707457692 | 0,00020935 | 0,002305497 |
| ENSG00000110057.8 | UNC93B1 | protein_coding | 617,2173594 | 3,109170281 | 0,216371702 | 14,3695791 | 8,03E-47 | 1,02E-44 |
| ENSG00000166710.21 | B2M | protein_coding | 331566,2421 | 3,107124238 | 0,143973589 | 21,58120987 | 2,70E-103 | 6,64E-101 |
| ENSG00000230438.7 | SERPINB9P1 | lncRNA | 12,93683108 | 3,106915366 | 1,10001886 | 2,824420089 | 0,004736626 | 0,031569965 |
| ENSG00000181381.14 | DDX60L | protein_coding | 3907,404016 | 3,105512031 | 0,145596686 | 21,32955161 | 6,04E-101 | 1,45E-98 |
| ENSG00000150938.10 | CRIM1 | protein_coding | 6231,441092 | 3,097798869 | 0,198272891 | 15,62391539 | 5,00E-55 | 7,70E-53 |
| ENSG00000148702.15 | HABP2 | protein_coding | 557,7272097 | 3,086608034 | 0,27161535 | 11,36389396 | 6,33E-30 | 5,28E-28 |
| ENSG00000273628.1 | AL354798.1 | unprocessed_pseud | 16,08645847 | 3,069785394 | 1,121405057 | 2,737445649 | 0,006191835 | 0,039161954 |
| ENSG00000152778.9 | IFIT5 | protein_coding | 5010,457023 | 3,061073465 | 0,120158751 | 25,47524361 | 3,71E-143 | 1,29E-140 |
| ENSG00000285081.1 | AC004593.1 | lncRNA | 89,1290018 | 3,058663936 | 0,42730529 | 7,158029655 | 8,18E-13 | 2,99E-11 |
| ENSG00000128394.17 | APOBEC3F | protein_coding | 555,5441279 | 3,058483237 | 0,219290001 | 13,94720792 | 3,27E-44 | 3,91E-42 |
| ENSG00000136630.13 | HLX | protein_coding | 28,07752768 | 3,048763857 | 0,828691387 | 3,679009947 | 0,000234141 | 0,002533166 |

| | | | | | | | | |
|---|---|---|---|---|---|---|---|---|
| ENSG00000285799.1 | AL645929.2 | transcribed_unproc | 53,93591832 | 3,047938095 | 0,593988892 | 5,131304873 | 2,88E-07 | 5,78E-06 |
| ENSG00000260487.1 | AC116348.1 | lncRNA | 12,6562513 | 3,037203018 | 1,141368051 | 2,661019831 | 0,007790437 | 0,046946688 |
| ENSG00000253438.4 | PCAT1 | lncRNA | 58,04394032 | 3,034499674 | 0,601643214 | 5,04368636 | 4,57E-07 | 8,87E-06 |
| ENSG00000138759.19 | FRAS1 | protein_coding | 2433,082211 | 3,033314539 | 0,195594061 | 15,50821394 | 3,05E-54 | 4,60E-52 |
| ENSG00000136048.14 | DRAM1 | protein_coding | 1416,734834 | 3,023426011 | 0,157428805 | 19,20503686 | 3,36E-82 | 6,70E-80 |
| ENSG00000251569.1 | AC093899.2 | protein_coding | 86,94287374 | 3,013377377 | 0,549736544 | 5,48149365 | 4,22E-08 | 9,49E-07 |
| ENSG00000005108.16 | THSD7A | protein_coding | 293,8544887 | 3,003590961 | 0,308016359 | 9,751400769 | 1,82E-22 | 1,15E-20 |
| ENSG00000235505.7 | CASP4LP | transcribed_unproc | 18,45019148 | 2,998302673 | 0,918548615 | 3,26417418 | 0,001097837 | 0,009392402 |
| ENSG00000253882.6 | AC099548.2 | transcribed_unproc | 1592,490423 | 2,997629757 | 0,225449498 | 13,29623614 | 2,43E-40 | 2,67E-38 |
| ENSG00000064763.11 | FAR2 | protein_coding | 37,71232277 | 2,994679403 | 0,684936137 | 4,372202373 | 1,23E-05 | 0,000183535 |
| ENSG00000115353.11 | TACR1 | protein_coding | 48,60373556 | 2,985941507 | 0,553014955 | 5,399386539 | 6,69E-08 | 1,47E-06 |
| ENSG00000276718.1 | AC005840.4 | lncRNA | 18,46043745 | 2,983921168 | 0,941921296 | 3,167909231 | 0,001535394 | 0,012546247 |
| ENSG00000165806.21 | CASP7 | protein_coding | 1628,778901 | 2,926918392 | 0,175279092 | 16,69861679 | 1,34E-62 | 2,26E-60 |
| ENSG00000170379.20 | TCAF2 | protein_coding | 909,0944041 | 2,925267887 | 0,187295059 | 15,61850003 | 5,45E-55 | 8,32E-53 |
| ENSG00000155307.19 | SAMSN1 | protein_coding | 12,56213053 | 2,918331596 | 1,070031756 | 2,727331763 | 0,006384881 | 0,040128333 |
| ENSG00000229644.6 | NAMPTP1 | processed_pseudog | 447,1329817 | 2,913915443 | 0,207323594 | 14,05491474 | 7,19E-45 | 8,80E-43 |
| ENSG00000172345.14 | STARD5 | protein_coding | 1261,102536 | 2,905999354 | 0,189002865 | 15,37542489 | 2,39E-53 | 3,53E-51 |
| ENSG00000281769.1 | LINC01230 | lncRNA | 72,14775489 | 2,904883403 | 0,481424042 | 6,033939215 | 1,60E-09 | 4,31E-08 |
| ENSG00000140968.11 | IRF8 | protein_coding | 8292,748483 | 2,887653367 | 0,159415883 | 18,11396275 | 2,47E-73 | 4,58E-71 |
| ENSG00000132744.8 | ACY3 | protein_coding | 497,7195123 | 2,886701449 | 0,211822702 | 13,62791347 | 2,73E-42 | 3,14E-40 |
| ENSG00000081041.9 | CXCL2 | protein_coding | 151,9791454 | 2,885362368 | 0,400565122 | 7,20322917 | 5,88E-13 | 2,17E-11 |
| ENSG00000039068.19 | CDH1 | protein_coding | 66054,61508 | 2,883945054 | 0,13353378 | 21,59711994 | 1,91E-103 | 4,76E-101 |
| ENSG00000152207.7 | CYSLTR2 | protein_coding | 212,2407099 | 2,873912799 | 0,285836709 | 10,05438667 | 8,79E-24 | 5,86E-22 |
| ENSG00000092010.15 | PSME1 | protein_coding | 27735,88387 | 2,862698216 | 0,131376234 | 21,79007663 | 2,88E-105 | 7,54E-103 |
| ENSG00000134326.11 | CMPK2 | protein_coding | 12090,56735 | 2,856716012 | 0,122799028 | 23,26334381 | 1,04E-119 | 3,21E-117 |
| ENSG00000116194.13 | ANGPTL1 | protein_coding | 16,35856634 | 2,856621376 | 0,961747107 | 2,970241713 | 0,002975655 | 0,021581825 |
| ENSG00000139192.12 | TAPBPL | protein_coding | 1330,63428 | 2,856019936 | 0,147941636 | 19,30504504 | 4,87E-83 | 9,90E-81 |
| ENSG00000072858.11 | SIDT1 | protein_coding | 92,56630726 | 2,850235811 | 0,419461187 | 6,794992963 | 1,08E-11 | 3,64E-10 |
| ENSG00000059378.13 | PARP12 | protein_coding | 4473,68041 | 2,849789879 | 0,122036157 | 23,3520126 | 1,31E-120 | 4,11E-118 |
| ENSG00000115363.14 | EVA1A | protein_coding | 232,8041019 | 2,838304799 | 0,343757837 | 8,256698457 | 1,50E-16 | 7,00E-15 |
| ENSG00000248685.6 | LINC02484 | lncRNA | 93,48100138 | 2,835766796 | 0,441721388 | 6,419808662 | 1,36E-10 | 4,06E-09 |
| ENSG00000206503.13 | HLA-A | protein_coding | 191340,4722 | 2,828366012 | 0,112588848 | 25,12119142 | 2,92E-139 | 1,00E-136 |
| ENSG00000152818.18 | UTRN | protein_coding | 13451,69504 | 2,823966483 | 0,18243565 | 15,47924696 | 4,79E-54 | 7,16E-52 |
| ENSG00000238164.6 | TNFRSF14-AS1 | lncRNA | 41,07743723 | 2,806672367 | 0,696579508 | 4,029220403 | 5,60E-05 | 0,000709611 |
| ENSG00000115267.8 | IFIH1 | protein_coding | 5610,77785 | 2,802873894 | 0,125923889 | 22,25847633 | 9,34E-110 | 2,51E-107 |
| ENSG00000157873.17 | TNFRSF14 | protein_coding | 2558,980545 | 2,798673282 | 0,132287096 | 21,15605651 | 2,43E-99 | 5,77E-97 |

| | | | | | | | | |
|---|---|---|---|---|---|---|---|---|
| ENSG00000164879.7 | CA3 | protein_coding | 15,75221811 | 2,790146978 | 0,892718792 | 3,125448914 | 0,001775339 | 0,014131834 |
| ENSG00000187094.12 | CCK | protein_coding | 19,53508852 | 2,784497233 | 0,970596146 | 2,868852554 | 0,004119638 | 0,028379276 |
| ENSG00000186714.12 | CCDC73 | protein_coding | 33,09262727 | 2,777006161 | 0,668601771 | 4,153453194 | 3,27E-05 | 0,000438651 |
| ENSG00000203999.9 | LINC01270 | lncRNA | 39,86061716 | 2,773960849 | 0,719116916 | 3,857454593 | 0,000114574 | 0,001345264 |
| ENSG00000250264.1 | AL669918.1 | protein_coding | 951,486219 | 2,770416718 | 0,183887072 | 15,06585907 | 2,72E-51 | 3,84E-49 |
| ENSG00000144820.8 | ADGRG7 | protein_coding | 61,73816616 | 2,768184023 | 0,541895444 | 5,108336035 | 3,25E-07 | 6,47E-06 |
| ENSG00000108950.12 | FAM20A | protein_coding | 22,57287939 | 2,750363453 | 1,003975651 | 2,739472268 | 0,006153791 | 0,038968423 |
| ENSG00000231925.12 | TAPBP | protein_coding | 7866,980829 | 2,734554233 | 0,173240722 | 15,78470814 | 3,96E-56 | 6,29E-54 |
| ENSG00000126246.10 | IGFLR1 | protein_coding | 753,6479967 | 2,723403374 | 0,184131097 | 14,79056726 | 1,69E-49 | 2,34E-47 |
| ENSG00000251136.9 | AF117829.1 | lncRNA | 1007,09912 | 2,706381459 | 0,234711499 | 11,53067263 | 9,24E-31 | 7,92E-29 |
| ENSG00000277496.1 | AL357033.4 | lncRNA | 100,8012378 | 2,704019872 | 0,446366654 | 6,057844705 | 1,38E-09 | 3,75E-08 |
| ENSG00000174808.12 | BTC | protein_coding | 785,1165907 | 2,702273901 | 0,220493546 | 12,25556915 | 1,57E-34 | 1,49E-32 |
| ENSG00000198597.9 | ZNF536 | protein_coding | 23,23046196 | 2,691845177 | 0,858759143 | 3,134575274 | 0,001721031 | 0,013809805 |
| ENSG00000168016.15 | TRANK1 | protein_coding | 2335,005435 | 2,662144743 | 0,145830301 | 18,25508644 | 1,89E-74 | 3,52E-72 |
| ENSG00000140464.20 | PML | protein_coding | 3109,315817 | 2,660760272 | 0,188427606 | 14,12086226 | 2,83E-45 | 3,48E-43 |
| ENSG00000111331.13 | OAS3 | protein_coding | 18646,79526 | 2,647524869 | 0,140242391 | 18,87820687 | 1,72E-79 | 3,40E-77 |
| ENSG00000132842.14 | AP3B1 | protein_coding | 25462,79803 | 2,634369498 | 0,113508018 | 23,20866443 | 3,72E-119 | 1,13E-116 |
| ENSG00000162687.19 | KCNT2 | protein_coding | 3649,049415 | 2,624088966 | 0,128387623 | 20,43880018 | 7,56E-93 | 1,72E-90 |
| ENSG00000235750.10 | KIAA0040 | protein_coding | 4339,865794 | 2,611107618 | 0,12562703 | 20,78460048 | 5,97E-96 | 1,39E-93 |
| ENSG00000135604.10 | STX11 | protein_coding | 457,5559175 | 2,60686102 | 0,203146307 | 12,83243128 | 1,08E-37 | 1,10E-35 |
| ENSG00000137571.11 | SLCO5A1 | protein_coding | 944,1664442 | 2,605754784 | 0,170564926 | 15,27720176 | 1,09E-52 | 1,58E-50 |
| ENSG00000136147.18 | PHF11 | protein_coding | 454,3484585 | 2,598242925 | 0,213868968 | 12,14876073 | 5,82E-34 | 5,40E-32 |
| ENSG00000260025.1 | CRIM1-DT | lncRNA | 179,2674925 | 2,595833984 | 0,29811069 | 8,70761791 | 3,10E-18 | 1,56E-16 |
| ENSG00000146666.5 | LINC00525 | lncRNA | 45,46403326 | 2,591498969 | 0,611562083 | 4,237507592 | 2,26E-05 | 0,000313976 |
| ENSG00000112297.18 | CRYBG1 | protein_coding | 2615,507817 | 2,585918427 | 0,139125003 | 18,58701437 | 4,09E-77 | 7,93E-75 |
| ENSG00000280734.3 | LINC01232 | lncRNA | 433,8606775 | 2,58335287 | 0,208381336 | 12,39723728 | 2,70E-35 | 2,62E-33 |
| ENSG00000166349.10 | RAG1 | protein_coding | 197,1335399 | 2,58204001 | 0,394217372 | 6,549787479 | 5,76E-11 | 1,78E-09 |
| ENSG00000171476.22 | HOPX | protein_coding | 4712,836196 | 2,57851292 | 0,142653868 | 18,07531017 | 4,99E-73 | 9,16E-71 |
| ENSG00000224389.9 | C4B | protein_coding | 2370,509193 | 2,5785128 | 0,199758859 | 12,90812741 | 4,05E-38 | 4,16E-36 |
| ENSG00000185022.12 | MAFF | protein_coding | 498,4562617 | 2,5684509 | 0,214757116 | 11,95979414 | 5,77E-33 | 5,23E-31 |
| ENSG00000261884.2 | AC040162.1 | protein_coding | 135,2704543 | 2,5551034 | 0,376530295 | 6,785917194 | 1,15E-11 | 3,87E-10 |
| ENSG00000105835.12 | NAMPT | protein_coding | 18626,69265 | 2,551453911 | 0,11921153 | 21,40274438 | 1,26E-101 | 3,07E-99 |
| ENSG00000124493.14 | GRM4 | protein_coding | 2251,393384 | 2,550263332 | 0,176674809 | 14,43478757 | 3,13E-47 | 4,01E-45 |
| ENSG00000214106.9 | PAXIP1-AS2 | lncRNA | 1050,798161 | 2,549353969 | 0,166536717 | 15,30805948 | 6,75E-53 | 9,89E-51 |
| ENSG00000250241.6 | AC105383.1 | lncRNA | 28,74086451 | 2,546082891 | 0,74227378 | 3,430112933 | 0,00060333 | 0,005658923 |
| ENSG00000172159.16 | FRMD3 | protein_coding | 122,6817955 | 2,545510909 | 0,363551333 | 7,001792267 | 2,53E-12 | 9,00E-11 |

| | | | | | | | | |
|---|---|---|---|---|---|---|---|---|
| ENSG00000170581.14 | STAT2 | protein_coding | 4521,605684 | 2,542423922 | 0,147487382 | 17,23824699 | 1,37E-66 | 2,41E-64 |
| ENSG00000266850.1 | AC090912.2 | lncRNA | 16,86756756 | 2,542128568 | 0,920736356 | 2,760973378 | 0,005762937 | 0,036908489 |
| ENSG00000230392.1 | AC004835.1 | lncRNA | 42,35657786 | 2,539610073 | 0,590482855 | 4,300903999 | 1,70E-05 | 0,000243911 |
| ENSG00000100336.17 | APOL4 | protein_coding | 90,22030997 | 2,534608216 | 0,392632366 | 6,455423538 | 1,08E-10 | 3,24E-09 |
| ENSG00000205927.5 | OLIG2 | protein_coding | 24,57242624 | 2,52694663 | 0,829656542 | 3,045774367 | 0,002320819 | 0,017597375 |
| ENSG00000064012.22 | CASP8 | protein_coding | 881,8393202 | 2,524982366 | 0,1801625 | 14,01502737 | 1,26E-44 | 1,52E-42 |
| ENSG00000105971.15 | CAV2 | protein_coding | 309,7897283 | 2,521472932 | 0,234667822 | 10,74486016 | 6,27E-27 | 4,79E-25 |
| ENSG00000163739.5 | CXCL1 | protein_coding | 1391,469079 | 2,519760869 | 0,223130071 | 11,29278927 | 1,42E-29 | 1,18E-27 |
| ENSG00000166016.6 | ABTB2 | protein_coding | 3664,789174 | 2,515648831 | 0,122322032 | 20,56578676 | 5,56E-94 | 1,28E-91 |
| ENSG00000073331.18 | ALPK1 | protein_coding | 168,9386204 | 2,513260722 | 0,387069008 | 6,493055936 | 8,41E-11 | 2,55E-09 |
| ENSG00000114204.15 | SERPINI2 | protein_coding | 32,56592522 | 2,502574977 | 0,653868873 | 3,827334623 | 0,000129538 | 0,001503263 |
| ENSG00000101347.10 | SAMHD1 | protein_coding | 27927,83577 | 2,498199904 | 0,109325119 | 22,85110624 | 1,42E-115 | 4,09E-113 |
| ENSG00000159216.19 | RUNX1 | protein_coding | 36,90056911 | 2,494868786 | 0,692486191 | 3,60277045 | 0,000314844 | 0,003251727 |
| ENSG00000139597.18 | N4BP2L1 | protein_coding | 813,59352 | 2,493294952 | 0,20145886 | 12,37619905 | 3,52E-35 | 3,39E-33 |
| ENSG00000164674.16 | SYTL3 | protein_coding | 86,35846269 | 2,484389895 | 0,418244648 | 5,940039899 | 2,85E-09 | 7,49E-08 |
| ENSG00000287064.1 | AL606500.1 | lncRNA | 66,32537117 | 2,48107905 | 0,510432091 | 4,860742678 | 1,17E-06 | 2,12E-05 |
| ENSG00000167034.10 | NKX3-1 | protein_coding | 284,6098984 | 2,474677758 | 0,24306932 | 10,18095478 | 2,41E-24 | 1,65E-22 |
| ENSG00000172572.7 | PDE3A | protein_coding | 25,85870427 | 2,463580395 | 0,772675977 | 3,188374518 | 0,001430751 | 0,011818745 |
| ENSG00000278948.1 | AL031587.5 | TEC | 24,04225232 | 2,449298898 | 0,865994972 | 2,828306143 | 0,004679503 | 0,031244472 |
| ENSG00000259529.2 | AL136295.5 | protein_coding | 101,5613807 | 2,447718465 | 0,551300966 | 4,439895114 | 9,00E-06 | 0,000138341 |
| ENSG00000204257.15 | HLA-DMA | protein_coding | 42,53923795 | 2,44523999 | 0,648430676 | 3,7710122 | 0,000162587 | 0,00183788 |
| ENSG00000182389.20 | CACNB4 | protein_coding | 24,94711227 | 2,440815144 | 0,853901841 | 2,858425908 | 0,004257485 | 0,029108574 |
| ENSG00000146674.15 | IGFBP3 | protein_coding | 883,5602609 | 2,439803399 | 0,183455733 | 13,29913958 | 2,34E-40 | 2,58E-38 |
| ENSG00000103490.14 | PYCARD | protein_coding | 108,302952 | 2,435267271 | 0,366596667 | 6,642906204 | 3,08E-11 | 9,79E-10 |
| ENSG00000108771.13 | DHX58 | protein_coding | 129,563016 | 2,432830509 | 0,40110978 | 6,06524854 | 1,32E-09 | 3,59E-08 |
| ENSG00000138650.9 | PCDH10 | protein_coding | 867,8967702 | 2,424508561 | 0,187122372 | 12,95680754 | 2,15E-38 | 2,23E-36 |
| ENSG00000227766.1 | AL671277.1 | unprocessed_pseud | 32,42658995 | 2,419515938 | 0,73869059 | 3,275411885 | 0,00105508 | 0,00908228 |
| ENSG00000020577.14 | SAMD4A | protein_coding | 525,5391616 | 2,418545354 | 0,236993959 | 10,20509282 | 1,88E-24 | 1,30E-22 |
| ENSG00000205413.8 | SAMD9 | protein_coding | 5292,732857 | 2,417133287 | 0,119859267 | 20,16642815 | 1,93E-90 | 4,30E-88 |
| ENSG00000172216.6 | CEBPB | protein_coding | 1426,971265 | 2,410547093 | 0,146646143 | 16,43784859 | 1,02E-60 | 1,70E-58 |
| ENSG00000254445.1 | HSPB2-C11orf52 | protein_coding | 56,56414145 | 2,406532331 | 0,578303911 | 4,161362709 | 3,16E-05 | 0,000426182 |
| ENSG00000253417.5 | LINC02159 | lncRNA | 21,05798379 | 2,401814604 | 0,797908619 | 3,010137434 | 0,002611295 | 0,019399383 |
| ENSG00000144644.14 | GADL1 | protein_coding | 66,30976156 | 2,393193405 | 0,472731662 | 5,062477506 | 4,14E-07 | 8,07E-06 |
| ENSG00000157890.17 | MEGF11 | protein_coding | 1005,4477 | 2,389281825 | 0,162312052 | 14,72029828 | 4,78E-49 | 6,49E-47 |
| ENSG00000128683.14 | GAD1 | protein_coding | 38,59482669 | 2,383591155 | 0,732680969 | 3,253245622 | 0,001140948 | 0,009713604 |
| ENSG00000175318.12 | GRAMD2A | protein_coding | 442,2844334 | 2,36961883 | 0,213315846 | 11,10849886 | 1,14E-28 | 9,22E-27 |

| | | | | | | | | |
|---|---|---|---|---|---|---|---|---|
| ENSG00000283537.2 | AC073264.3 | transcribed_unproc | 772,1666392 | 2,356763846 | 0,180209622 | 13,07790238 | 4,40E-39 | 4,68E-37 |
| ENSG00000126790.12 | L3HYPDH | protein_coding | 85,08468793 | 2,35211911 | 0,509437965 | 4,617086421 | 3,89E-06 | 6,44E-05 |
| ENSG00000231574.6 | LINC02015 | lncRNA | 74,70905401 | 2,347495555 | 0,468427382 | 5,011439648 | 5,40E-07 | 1,03E-05 |
| ENSG00000287733.1 | AC083862.2 | lncRNA | 101,1859648 | 2,343996264 | 0,378463045 | 6,193461403 | 5,89E-10 | 1,66E-08 |
| ENSG00000162772.17 | ATF3 | protein_coding | 534,8044073 | 2,343738031 | 0,225463907 | 10,3951806 | 2,61E-25 | 1,86E-23 |
| ENSG00000214212.9 | C19orf38 | protein_coding | 262,7607624 | 2,337871403 | 0,272114913 | 8,591485767 | 8,59E-18 | 4,21E-16 |
| ENSG00000177337.8 | DLGAP1-AS1 | lncRNA | 39,30035106 | 2,331119691 | 0,708896181 | 3,288379532 | 0,001007659 | 0,008731515 |
| ENSG00000019582.15 | CD74 | protein_coding | 21912,81109 | 2,317645679 | 0,171397676 | 13,5220368 | 1,16E-41 | 1,30E-39 |
| ENSG00000130589.16 | HELZ2 | protein_coding | 7510,244647 | 2,311010924 | 0,148373904 | 15,57558887 | 1,07E-54 | 1,62E-52 |
| ENSG00000107201.10 | DDX58 | protein_coding | 2062,378659 | 2,308975482 | 0,141559375 | 16,31100362 | 8,24E-60 | 1,34E-57 |
| ENSG00000259479.6 | SORD2P | transcribed_unproc | 124,6662087 | 2,305476128 | 0,513020239 | 4,493928213 | 6,99E-06 | 0,00010973 |
| ENSG00000171815.6 | PCDHB1 | protein_coding | 57,74170418 | 2,300539524 | 0,585244836 | 3,93090102 | 8,46E-05 | 0,001032453 |
| ENSG00000224992.2 | AL445645.1 | lncRNA | 20,72597064 | 2,299469027 | 0,855193828 | 2,688827903 | 0,007170338 | 0,043946523 |
| ENSG00000227199.1 | ST7-AS1 | lncRNA | 347,5397461 | 2,29173478 | 0,237525641 | 9,648367946 | 4,99E-22 | 3,02E-20 |
| ENSG00000113494.17 | PRLR | protein_coding | 345,5229534 | 2,288234953 | 0,236199358 | 9,687727222 | 3,40E-22 | 2,09E-20 |
| ENSG00000105974.13 | CAV1 | protein_coding | 52,87070855 | 2,285843424 | 0,592152531 | 3,860227399 | 0,000113282 | 0,001336084 |
| ENSG00000106066.15 | CPVL | protein_coding | 27,8519018 | 2,285813608 | 0,758708017 | 3,012771126 | 0,002588741 | 0,019280194 |
| ENSG00000135148.12 | TRAFD1 | protein_coding | 16534,58438 | 2,276359408 | 0,118415641 | 19,22346902 | 2,35E-82 | 4,74E-80 |
| ENSG00000169126.16 | ARMC4 | protein_coding | 63,64247538 | 2,274195492 | 0,680016048 | 3,344326211 | 0,000824827 | 0,007366788 |
| ENSG00000270426.1 | AC099343.2 | lncRNA | 28,15041306 | 2,272459319 | 0,680668396 | 3,338570343 | 0,000842107 | 0,007492355 |
| ENSG00000287726.1 | AC005740.5 | lncRNA | 20,57919725 | 2,250125069 | 0,791397368 | 2,843230417 | 0,004465877 | 0,030110512 |
| ENSG00000226380.9 | AC016831.1 | lncRNA | 27,57995324 | 2,246559935 | 0,784516283 | 2,863624355 | 0,004188244 | 0,028757259 |
| ENSG00000232810.4 | TNF | protein_coding | 34,67046359 | 2,245851112 | 0,795709351 | 2,822451577 | 0,004765802 | 0,031703864 |
| ENSG00000112299.8 | VNN1 | protein_coding | 104,6136065 | 2,243645876 | 0,593168833 | 3,782474314 | 0,000155277 | 0,001759963 |
| ENSG00000197558.13 | SSPOP | transcribed_unitary | 43,57480097 | 2,2378816 | 0,659623894 | 3,392663032 | 0,000692167 | 0,006341585 |
| ENSG00000185112.6 | FAM43A | protein_coding | 249,8926089 | 2,237671388 | 0,280629457 | 7,973758038 | 1,54E-15 | 6,80E-14 |
| ENSG00000177721.4 | ANXA2R | protein_coding | 196,5879709 | 2,237430084 | 0,280985446 | 7,962797061 | 1,68E-15 | 7,41E-14 |
| ENSG00000130813.18 | SHFL | protein_coding | 1095,023996 | 2,23725315 | 0,155327264 | 14,40348006 | 4,92E-47 | 6,28E-45 |
| ENSG00000246528.3 | AC079089.1 | lncRNA | 63,52665804 | 2,233770354 | 0,512095732 | 4,362017129 | 1,29E-05 | 0,000190662 |
| ENSG00000180801.14 | ARSJ | protein_coding | 2047,377684 | 2,221766593 | 0,18410937 | 12,0676454 | 1,57E-33 | 1,44E-31 |
| ENSG00000237276.8 | ANO7L1 | transcribed_unproc | 68,34936395 | 2,212855584 | 0,450126225 | 4,91607789 | 8,83E-07 | 1,64E-05 |
| ENSG00000161944.16 | ASGR2 | protein_coding | 86,10924112 | 2,212360966 | 0,436218864 | 5,071676505 | 3,94E-07 | 7,71E-06 |
| ENSG00000123992.20 | DNPEP | protein_coding | 8994,739232 | 2,203638293 | 0,127205593 | 17,32343875 | 3,13E-67 | 5,55E-65 |
| ENSG00000172247.4 | C1QTNF4 | protein_coding | 118,7674175 | 2,19976051 | 0,349631588 | 6,291652665 | 3,14E-10 | 9,10E-09 |
| ENSG00000124201.15 | ZNFX1 | protein_coding | 6221,419779 | 2,194798695 | 0,156582251 | 14,01690598 | 1,23E-44 | 1,49E-42 |
| ENSG00000130182.8 | ZSCAN10 | protein_coding | 44,30899293 | 2,191663962 | 0,565515207 | 3,875517288 | 0,000106398 | 0,001263444 |

| | | | | | | | | |
|---|---|---|---|---|---|---|---|---|
| ENSG00000198133.8 | TMEM229B | protein_coding | 324,2203386 | 2,19092725 | 0,226954256 | 9,653607258 | 4,75E-22 | 2,88E-20 |
| ENSG00000172318.5 | B3GALT1 | protein_coding | 1516,590274 | 2,186358889 | 0,175495705 | 12,45819032 | 1,26E-35 | 1,24E-33 |
| ENSG00000176485.12 | PLAAT3 | protein_coding | 2414,262555 | 2,18431943 | 0,138991121 | 15,71553215 | 1,18E-55 | 1,85E-53 |
| ENSG00000251584.2 | AC096751.2 | lncRNA | 75,98996204 | 2,183180184 | 0,4291179 | 5,087599896 | 3,63E-07 | 7,16E-06 |
| ENSG00000105339.11 | DENND3 | protein_coding | 2486,315395 | 2,181705008 | 0,177005269 | 12,32565009 | 6,59E-35 | 6,30E-33 |
| ENSG00000204335.4 | SP5 | protein_coding | 116,9064178 | 2,151546638 | 0,365111286 | 5,89285164 | 3,80E-09 | 9,76E-08 |
| ENSG00000185404.17 | SP140L | protein_coding | 39,5412813 | 2,141602044 | 0,632421006 | 3,386355011 | 0,000708277 | 0,006460905 |
| ENSG00000196776.17 | CD47 | protein_coding | 11411,93675 | 2,139524905 | 0,131410955 | 16,28117612 | 1,34E-59 | 2,16E-57 |
| ENSG00000173068.18 | BNC2 | protein_coding | 24,83923752 | 2,135354965 | 0,770312397 | 2,77206361 | 0,005570215 | 0,035936041 |
| ENSG00000168610.16 | STAT3 | protein_coding | 21742,41617 | 2,130504851 | 0,118885812 | 17,92059807 | 8,14E-72 | 1,48E-69 |
| ENSG00000171446.7 | KRT27 | protein_coding | 63,86236581 | 2,129927307 | 0,547246343 | 3,892081391 | 9,94E-05 | 0,001186928 |
| ENSG00000260103.2 | AC012435.2 | transcribed_process | 118,0239412 | 2,12916311 | 0,355940973 | 5,981787073 | 2,21E-09 | 5,89E-08 |
| ENSG00000144802.11 | NFKBIZ | protein_coding | 2778,287822 | 2,122856908 | 0,18558225 | 11,43890061 | 2,67E-30 | 2,26E-28 |
| ENSG00000170006.12 | TMEM154 | protein_coding | 33,78630976 | 2,091997127 | 0,678425488 | 3,083606328 | 0,00204508 | 0,015844046 |
| ENSG00000159871.15 | LYPD5 | protein_coding | 699,5843458 | 2,083168934 | 0,189549989 | 10,99007678 | 4,27E-28 | 3,42E-26 |
| ENSG00000168671.10 | UGT3A2 | protein_coding | 31,89460507 | 2,06852578 | 0,671247488 | 3,081614184 | 0,002058815 | 0,015939827 |
| ENSG00000205436.7 | EXOC3L4 | protein_coding | 251,4247811 | 2,061068558 | 0,339428277 | 6,072176941 | 1,26E-09 | 3,45E-08 |
| ENSG00000118946.12 | PCDH17 | protein_coding | 3987,521905 | 2,060821947 | 0,139833101 | 14,73772613 | 3,69E-49 | 5,05E-47 |
| ENSG00000164307.13 | ERAP1 | protein_coding | 8480,69649 | 2,059861088 | 0,117454893 | 17,5374651 | 7,42E-69 | 1,34E-66 |
| ENSG00000167705.12 | RILP | protein_coding | 27,13412703 | 2,059612679 | 0,707886989 | 2,909521875 | 0,003619821 | 0,025455473 |
| ENSG00000137203.15 | TFAP2A | protein_coding | 103,9935299 | 2,050264229 | 0,4325776 | 4,739644934 | 2,14E-06 | 3,71E-05 |
| ENSG00000250360.1 | AC008808.2 | lncRNA | 216,5116146 | 2,049040254 | 0,269458733 | 7,604282232 | 2,86E-14 | 1,17E-12 |
| ENSG00000006210.7 | CX3CL1 | protein_coding | 4380,746333 | 2,047248658 | 0,163288408 | 12,537624 | 4,65E-36 | 4,61E-34 |
| ENSG00000164236.12 | ANKRD33B | protein_coding | 45,06859567 | 2,039410301 | 0,554908632 | 3,675218198 | 0,000237646 | 0,002565819 |
| ENSG00000184731.6 | FAM110C | protein_coding | 4145,143095 | 2,038634593 | 0,137204897 | 14,8583224 | 6,14E-50 | 8,58E-48 |
| ENSG00000064932.16 | SBNO2 | protein_coding | 1931,979302 | 2,026895024 | 0,156683848 | 12,93620916 | 2,81E-38 | 2,90E-36 |
| ENSG00000174776.11 | WDR49 | protein_coding | 377,8124747 | 2,019312692 | 0,338592359 | 5,963846013 | 2,46E-09 | 6,54E-08 |
| ENSG00000109705.8 | NKX3-2 | protein_coding | 281,6110532 | 2,017963328 | 0,243501933 | 8,28725794 | 1,16E-16 | 5,45E-15 |
| ENSG00000124557.12 | BTN1A1 | protein_coding | 568,2333619 | 2,017322284 | 0,224209192 | 8,997500347 | 2,31E-19 | 1,23E-17 |
| ENSG00000266094.8 | RASSF5 | protein_coding | 357,669354 | 2,012179027 | 0,220805331 | 9,112909634 | 8,02E-20 | 4,37E-18 |
| ENSG00000175336.10 | APOF | protein_coding | 97,37496992 | 2,010766752 | 0,400397948 | 5,021920716 | 5,12E-07 | 9,85E-06 |
| ENSG00000261971.8 | MMP25-AS1 | lncRNA | 115,3941428 | 2,006272303 | 0,437588939 | 4,584833214 | 4,54E-06 | 7,39E-05 |
| ENSG00000112541.17 | PDE10A | protein_coding | 125,6699757 | 2,002433364 | 0,402736511 | 4,972068115 | 6,62E-07 | 1,25E-05 |
| ENSG00000187908.20 | DMBT1 | protein_coding | 124,5957222 | 1,99189258 | 0,364872813 | 5,459142222 | 4,78E-08 | 1,07E-06 |
| ENSG00000171621.14 | SPSB1 | protein_coding | 391,5025224 | 1,990169299 | 0,220969577 | 9,006530787 | 2,13E-19 | 1,13E-17 |
| ENSG00000122679.8 | RAMP3 | protein_coding | 95,18079908 | 1,980258052 | 0,386721566 | 5,12063 | 3,05E-07 | 6,09E-06 |

| | | | | | | | | |
|---|---|---|---|---|---|---|---|---|
| ENSG00000185432.12 | METTL7A | protein_coding | 58,31779448 | 1,976313613 | 0,500102832 | 3,951814476 | 7,76E-05 | 0,000952897 |
| ENSG00000236078.2 | LINC01447 | lncRNA | 36,76785798 | 1,973565301 | 0,593433669 | 3,325671266 | 0,000882059 | 0,007801396 |
| ENSG00000128203.7 | ASPHD2 | protein_coding | 1930,128246 | 1,968684722 | 0,135817805 | 14,49504154 | 1,30E-47 | 1,70E-45 |
| ENSG00000013374.16 | NUB1 | protein_coding | 12747,07319 | 1,968610704 | 0,112587547 | 17,48515489 | 1,86E-68 | 3,33E-66 |
| ENSG00000164342.13 | TLR3 | protein_coding | 58,15328947 | 1,966333304 | 0,646566304 | 3,041193596 | 0,002356423 | 0,017841486 |
| ENSG00000099860.9 | GADD45B | protein_coding | 341,0845802 | 1,964785068 | 0,233752186 | 8,405419014 | 4,26E-17 | 2,03E-15 |
| ENSG00000231535.7 | LINC00278 | lncRNA | 181,132262 | 1,960174929 | 0,33348688 | 5,877817222 | 4,16E-09 | 1,06E-07 |
| ENSG00000183323.13 | CCDC125 | protein_coding | 1165,131502 | 1,955927734 | 0,191310237 | 10,22385298 | 1,55E-24 | 1,07E-22 |
| ENSG00000112773.16 | TENT5A | protein_coding | 31475,35305 | 1,939217629 | 0,115845699 | 16,73966012 | 6,74E-63 | 1,15E-60 |
| ENSG00000056291.17 | NPFFR2 | protein_coding | 910,7739615 | 1,924993258 | 0,181684708 | 10,59524098 | 3,14E-26 | 2,33E-24 |
| ENSG00000166343.10 | MSS51 | protein_coding | 854,9935749 | 1,923147903 | 0,204526745 | 9,402916481 | 5,31E-21 | 3,04E-19 |
| ENSG00000151883.19 | PARP8 | protein_coding | 4208,757799 | 1,92219696 | 0,132728347 | 14,48218863 | 1,57E-47 | 2,04E-45 |
| ENSG00000188290.11 | HES4 | protein_coding | 35,36800007 | 1,920710725 | 0,618597026 | 3,104946587 | 0,001903134 | 0,01496753 |
| ENSG00000187510.10 | PLEKHG7 | protein_coding | 79,03641616 | 1,919188966 | 0,427159911 | 4,492905158 | 7,03E-06 | 0,000110176 |
| ENSG00000124508.17 | BTN2A2 | protein_coding | 478,4354177 | 1,918086552 | 0,21542266 | 8,903829122 | 5,40E-19 | 2,82E-17 |
| ENSG00000158458.21 | NRG2 | protein_coding | 225,459636 | 1,911097097 | 0,256816787 | 7,441480451 | 9,96E-14 | 3,92E-12 |
| ENSG00000258499.1 | LINC02287 | lncRNA | 64,07219685 | 1,909775526 | 0,45548556 | 4,192834401 | 2,75E-05 | 0,000376462 |
| ENSG00000053328.8 | METTL24 | protein_coding | 114,4871588 | 1,908725724 | 0,359726965 | 5,306040168 | 1,12E-07 | 2,37E-06 |
| ENSG00000206120.11 | EGFEM1P | transcribed_unitary | 209,4559913 | 1,907648856 | 0,297702996 | 6,407892712 | 1,48E-10 | 4,38E-09 |
| ENSG00000139083.11 | ETV6 | protein_coding | 1925,703582 | 1,907208829 | 0,180759562 | 10,5510813 | 5,02E-26 | 3,71E-24 |
| ENSG00000177453.7 | NIM1K | protein_coding | 109,9597946 | 1,906975908 | 0,366709538 | 5,200235372 | 1,99E-07 | 4,07E-06 |
| ENSG00000104432.14 | IL7 | protein_coding | 1402,52241 | 1,904081729 | 0,212724141 | 8,950943341 | 3,52E-19 | 1,86E-17 |
| ENSG00000108821.14 | COL1A1 | protein_coding | 332,1351017 | 1,903428828 | 0,270861846 | 7,027305082 | 2,11E-12 | 7,54E-11 |
| ENSG00000137393.10 | RNF144B | protein_coding | 548,1359814 | 1,901337685 | 0,257188509 | 7,392778531 | 1,44E-13 | 5,55E-12 |
| ENSG00000287550.1 | AL450345.2 | lncRNA | 77,06575443 | 1,897878588 | 0,459337859 | 4,131770438 | 3,60E-05 | 0,000477275 |
| ENSG00000121351.8 | IAPP | protein_coding | 32833,21281 | 1,893868406 | 0,392095707 | 4,83011768 | 1,36E-06 | 2,45E-05 |
| ENSG00000114805.18 | PLCH1 | protein_coding | 1215,752682 | 1,886428967 | 0,157098473 | 12,00793951 | 3,23E-33 | 2,96E-31 |
| ENSG00000225706.2 | PTPRD-AS1 | lncRNA | 70,33415796 | 1,883202358 | 0,450901362 | 4,176528434 | 2,96E-05 | 0,000401074 |
| ENSG00000124549.14 | BTN2A3P | transcribed_unproc | 30,35505381 | 1,880143931 | 0,703465774 | 2,672687146 | 0,007524639 | 0,045700121 |
| ENSG00000185507.21 | IRF7 | protein_coding | 604,8938335 | 1,876740558 | 0,197533627 | 9,500866188 | 2,08E-21 | 1,23E-19 |
| ENSG00000136114.17 | THSD1 | protein_coding | 34,11137197 | 1,868499402 | 0,618420914 | 3,021403965 | 0,002516054 | 0,018804584 |
| ENSG00000166743.9 | ACSM1 | protein_coding | 86,81891173 | 1,866769464 | 0,506974815 | 3,682173965 | 0,000231254 | 0,002509742 |
| ENSG00000113504.21 | SLC12A7 | protein_coding | 17394,7941 | 1,857845103 | 0,129866058 | 14,30585585 | 2,01E-46 | 2,54E-44 |
| ENSG00000152785.8 | BMP3 | protein_coding | 194,5235323 | 1,855437961 | 0,325931087 | 5,692730873 | 1,25E-08 | 3,02E-07 |
| ENSG00000099250.18 | NRP1 | protein_coding | 1980,601705 | 1,853936348 | 0,222984328 | 8,314200216 | 9,24E-17 | 4,37E-15 |
| ENSG00000281741.2 | AC241377.3 | TEC | 82,13355151 | 1,853909644 | 0,546757014 | 3,39073774 | 0,000697048 | 0,006372985 |

| | | | | | | | | |
|---|---|---|---|---|---|---|---|---|
| ENSG00000154889.17 | MPPE1 | protein_coding | 3209,06506 | 1,852691788 | 0,157492774 | 11,76366219 | 6,01E-32 | 5,26E-30 |
| ENSG00000142178.9 | SIK1 | protein_coding | 537,5877383 | 1,851075979 | 0,307529156 | 6,019188561 | 1,75E-09 | 4,70E-08 |
| ENSG00000136848.17 | DAB2IP | protein_coding | 4056,142188 | 1,847525023 | 0,187106459 | 9,874191589 | 5,39E-23 | 3,54E-21 |
| ENSG00000135744.8 | AGT | protein_coding | 922,0876239 | 1,843989854 | 0,171690005 | 10,7402283 | 6,59E-27 | 5,00E-25 |
| ENSG00000185947.15 | ZNF267 | protein_coding | 2926,225007 | 1,840792375 | 0,135931441 | 13,5420647 | 8,83E-42 | 9,99E-40 |
| ENSG00000172936.16 | MYD88 | protein_coding | 3530,991021 | 1,836203323 | 0,122448739 | 14,99568992 | 7,83E-51 | 1,10E-48 |
| ENSG00000107968.10 | MAP3K8 | protein_coding | 260,4972562 | 1,830381779 | 0,274867868 | 6,659133319 | 2,75E-11 | 8,79E-10 |
| ENSG00000256646.8 | AC010132.3 | protein_coding | 621,8614977 | 1,827409903 | 0,316033856 | 5,782323225 | 7,37E-09 | 1,82E-07 |
| ENSG00000092969.12 | TGFB2 | protein_coding | 31,48470224 | 1,819972101 | 0,630327873 | 2,887341937 | 0,003885118 | 0,027066535 |
| ENSG00000261512.2 | AC092368.3 | lncRNA | 108,832118 | 1,811077271 | 0,443903205 | 4,079892304 | 4,51E-05 | 0,000584061 |
| ENSG00000105810.10 | CDK6 | protein_coding | 704,6144144 | 1,809330846 | 0,226018378 | 8,005237731 | 1,19E-15 | 5,29E-14 |
| ENSG00000167434.10 | CA4 | protein_coding | 1343,685627 | 1,808091398 | 0,269365302 | 6,712413904 | 1,91E-11 | 6,27E-10 |
| ENSG00000156515.24 | HK1 | protein_coding | 43,94541155 | 1,806735199 | 0,563897663 | 3,204012566 | 0,001355266 | 0,011303779 |
| ENSG00000167371.21 | PRRT2 | protein_coding | 106,7802852 | 1,804393789 | 0,414301838 | 4,355263775 | 1,33E-05 | 0,000196082 |
| ENSG00000115963.13 | RND3 | protein_coding | 1932,553012 | 1,799522622 | 0,184992995 | 9,727517663 | 2,30E-22 | 1,43E-20 |
| ENSG00000243064.8 | ABCC13 | transcribed_unproc | 106,4726316 | 1,794761441 | 0,441115592 | 4,068687377 | 4,73E-05 | 0,000608721 |
| ENSG00000265203.2 | RBP3 | protein_coding | 36,85607308 | 1,793969248 | 0,581648014 | 3,084286724 | 0,002040408 | 0,015820723 |
| ENSG00000198216.12 | CACNA1E | protein_coding | 32,32027129 | 1,791129802 | 0,638481725 | 2,805295331 | 0,00502705 | 0,033073942 |
| ENSG00000141452.10 | RMC1 | protein_coding | 5435,440152 | 1,791039654 | 0,125735207 | 14,24453575 | 4,85E-46 | 6,08E-44 |
| ENSG00000146648.19 | EGFR | protein_coding | 114,8542469 | 1,789183334 | 0,361808365 | 4,945113239 | 7,61E-07 | 1,42E-05 |
| ENSG00000169871.13 | TRIM56 | protein_coding | 6680,094203 | 1,787676111 | 0,159405446 | 11,21464888 | 3,46E-29 | 2,84E-27 |
| ENSG00000198535.5 | C2CD4A | protein_coding | 6045,591901 | 1,787521701 | 0,143770018 | 12,43320222 | 1,73E-35 | 1,68E-33 |
| ENSG00000135378.4 | PRRG4 | protein_coding | 7214,862931 | 1,775917795 | 0,1358025 | 13,07720985 | 4,44E-39 | 4,70E-37 |
| ENSG00000164946.20 | FREM1 | protein_coding | 100,6237747 | 1,774480066 | 0,403997824 | 4,392301053 | 1,12E-05 | 0,0001688 |
| ENSG00000128918.15 | ALDH1A2 | protein_coding | 50,56031625 | 1,77316655 | 0,547409108 | 3,239198113 | 0,001198663 | 0,01016775 |
| ENSG00000166510.14 | CCDC68 | protein_coding | 2472,682798 | 1,766452808 | 0,142994479 | 12,35329377 | 4,68E-35 | 4,49E-33 |
| ENSG00000180660.8 | MAB21L1 | protein_coding | 84,74066777 | 1,764528437 | 0,394713446 | 4,470403673 | 7,81E-06 | 0,000121235 |
| ENSG00000168938.6 | PPIC | protein_coding | 2900,511942 | 1,757666664 | 0,128353602 | 13,69394112 | 1,10E-42 | 1,28E-40 |
| ENSG00000151611.16 | MMAA | protein_coding | 2602,036534 | 1,757349152 | 0,127191789 | 13,81652985 | 2,03E-43 | 2,40E-41 |
| ENSG00000163702.20 | IL17RC | protein_coding | 2042,520924 | 1,754904006 | 0,174311945 | 10,0676061 | 7,68E-24 | 5,17E-22 |
| ENSG00000058866.15 | DGKG | protein_coding | 2097,594252 | 1,749925774 | 0,157221648 | 11,1303106 | 8,93E-29 | 7,28E-27 |
| ENSG00000034510.6 | TMSB10 | protein_coding | 67630,5711 | 1,747446298 | 0,161071856 | 10,84886174 | 2,02E-27 | 1,60E-25 |
| ENSG00000177669.4 | MBOAT4 | protein_coding | 2662,312381 | 1,745635126 | 0,127799199 | 13,65920244 | 1,78E-42 | 2,06E-40 |
| ENSG00000126456.16 | IRF3 | protein_coding | 3248,60812 | 1,744370763 | 0,151284253 | 11,5304186 | 9,27E-31 | 7,92E-29 |
| ENSG00000060491.16 | OGFR | protein_coding | 7593,196428 | 1,744028379 | 0,120298013 | 14,49756596 | 1,26E-47 | 1,65E-45 |
| ENSG00000169252.6 | ADRB2 | protein_coding | 229,6320003 | 1,742351233 | 0,283629481 | 6,143054054 | 8,09E-10 | 2,25E-08 |

| | | | | | | | | |
|---|---|---|---|---|---|---|---|---|
| ENSG00000134013.16 | LOXL2 | protein_coding | 5809,852829 | 1,740496534 | 0,180998064 | 9,616105818 | 6,84E-22 | 4,11E-20 |
| ENSG00000106785.15 | TRIM14 | protein_coding | 1882,894754 | 1,737556157 | 0,145666662 | 11,92830353 | 8,43E-33 | 7,60E-31 |
| ENSG00000168310.11 | IRF2 | protein_coding | 3335,824435 | 1,731479369 | 0,125837865 | 13,75960547 | 4,46E-43 | 5,24E-41 |
| ENSG00000119950.21 | MXI1 | protein_coding | 448,282697 | 1,731002871 | 0,236741337 | 7,311789704 | 2,64E-13 | 1,00E-11 |
| ENSG00000090104.12 | RGS1 | protein_coding | 117,3435345 | 1,730309581 | 0,355438597 | 4,86809704 | 1,13E-06 | 2,05E-05 |
| ENSG00000158406.5 | H4C8 | protein_coding | 123,5387268 | 1,728717495 | 0,386511335 | 4,472617845 | 7,73E-06 | 0,000120178 |
| ENSG00000129596.5 | CDO1 | protein_coding | 347,8030035 | 1,727280925 | 0,248439779 | 6,95251354 | 3,59E-12 | 1,26E-10 |
| ENSG00000250166.3 | AC053513.1 | lncRNA | 312,9567198 | 1,724799789 | 0,224458856 | 7,684258116 | 1,54E-14 | 6,39E-13 |
| ENSG00000245275.8 | SAP30L-AS1 | lncRNA | 45,9287219 | 1,72238803 | 0,576412434 | 2,98811741 | 0,002807017 | 0,020604804 |
| ENSG00000101384.12 | JAG1 | protein_coding | 743,731541 | 1,719750479 | 0,226600843 | 7,589338381 | 3,22E-14 | 1,31E-12 |
| ENSG00000239474.7 | KLHL41 | protein_coding | 396,9445041 | 1,709816994 | 0,208422578 | 8,203607346 | 2,33E-16 | 1,07E-14 |
| ENSG00000113083.15 | LOX | protein_coding | 71,01063903 | 1,706025486 | 0,449976809 | 3,791363134 | 0,000149823 | 0,001709285 |
| ENSG00000151572.18 | ANO4 | protein_coding | 168,0043183 | 1,704566473 | 0,361026759 | 4,721440808 | 2,34E-06 | 4,02E-05 |
| ENSG00000125845.7 | BMP2 | protein_coding | 1532,126571 | 1,701291599 | 0,146612336 | 11,60401397 | 3,93E-31 | 3,40E-29 |
| ENSG00000143333.7 | RGS16 | protein_coding | 342,4008836 | 1,698435683 | 0,23245167 | 7,306618545 | 2,74E-13 | 1,04E-11 |
| ENSG00000184232.9 | OAF | protein_coding | 32,82731922 | 1,696840253 | 0,630954881 | 2,689321063 | 0,007159752 | 0,043909153 |
| ENSG00000261796.1 | ISY1-RAB43 | protein_coding | 869,0839172 | 1,687119627 | 0,197656538 | 8,535612557 | 1,39E-17 | 6,77E-16 |
| ENSG00000126778.12 | SIX1 | protein_coding | 522,0031182 | 1,686182304 | 0,209672669 | 8,041974715 | 8,84E-16 | 3,95E-14 |
| ENSG00000085563.15 | ABCB1 | protein_coding | 23269,90303 | 1,685331444 | 0,114905797 | 14,6670706 | 1,05E-48 | 1,42E-46 |
| ENSG00000057657.17 | PRDM1 | protein_coding | 242,2873175 | 1,683053311 | 0,298478744 | 5,638771076 | 1,71E-08 | 4,05E-07 |
| ENSG00000105287.13 | PRKD2 | protein_coding | 3291,226672 | 1,682958759 | 0,142966622 | 11,77169 | 5,46E-32 | 4,82E-30 |
| ENSG00000173702.7 | MUC13 | protein_coding | 1175,86997 | 1,681501856 | 0,181250174 | 9,277242708 | 1,74E-20 | 9,79E-19 |
| ENSG00000176597.12 | B3GNT5 | protein_coding | 7459,755881 | 1,680417608 | 0,114010646 | 14,7391289 | 3,61E-49 | 4,98E-47 |
| ENSG00000159167.12 | STC1 | protein_coding | 464,707638 | 1,674644892 | 0,272909605 | 6,136262203 | 8,45E-10 | 2,35E-08 |
| ENSG00000130513.6 | GDF15 | protein_coding | 70,95140696 | 1,672499761 | 0,459510795 | 3,639739871 | 0,000272914 | 0,00287831 |
| ENSG00000185272.14 | RBM11 | protein_coding | 187,6396173 | 1,669875484 | 0,280723002 | 5,948481148 | 2,71E-09 | 7,14E-08 |
| ENSG00000227507.3 | LTB | protein_coding | 370,2543625 | 1,659155198 | 0,3519805 | 4,713770214 | 2,43E-06 | 4,15E-05 |
| ENSG00000184979.10 | USP18 | protein_coding | 5704,559988 | 1,658939962 | 0,132436274 | 12,52632617 | 5,36E-36 | 5,29E-34 |
| ENSG00000284691.1 | AC073111.3 | protein_coding | 813,4757157 | 1,65496745 | 0,201866204 | 8,198338377 | 2,44E-16 | 1,12E-14 |
| ENSG00000288093.1 | AL592295.5 | lncRNA | 54,08292007 | 1,650953806 | 0,527027119 | 3,132578473 | 0,001732781 | 0,01388812 |
| ENSG00000146425.11 | DYNLT1 | protein_coding | 22370,87765 | 1,650465298 | 0,145911769 | 11,31139253 | 1,15E-29 | 9,57E-28 |
| ENSG00000168994.14 | PXDC1 | protein_coding | 604,2865365 | 1,648277893 | 0,2474896 | 6,659988524 | 2,74E-11 | 8,75E-10 |
| ENSG00000116514.17 | RNF19B | protein_coding | 4817,480726 | 1,639588825 | 0,121471539 | 13,49771999 | 1,61E-41 | 1,80E-39 |
| ENSG00000061273.18 | HDAC7 | protein_coding | 179,6816363 | 1,637968697 | 0,290260233 | 5,643104063 | 1,67E-08 | 3,96E-07 |
| ENSG00000136826.15 | KLF4 | protein_coding | 603,684363 | 1,633080118 | 0,203375987 | 8,029857131 | 9,76E-16 | 4,35E-14 |
| ENSG00000196358.11 | NTNG2 | protein_coding | 1068,689203 | 1,62836167 | 0,177590523 | 9,169192394 | 4,77E-20 | 2,63E-18 |

| | | | | | | | | |
|---|---|---|---|---|---|---|---|---|
| ENSG00000183801.8 | OLFML1 | protein_coding | 288,7024284 | 1,625226663 | 0,249392852 | 6,516733129 | 7,19E-11 | 2,19E-09 |
| ENSG00000257594.4 | GALNT4 | protein_coding | 249,7504279 | 1,623121902 | 0,446468127 | 3,63547094 | 0,000277473 | 0,002915806 |
| ENSG00000079385.23 | CEACAM1 | protein_coding | 9344,000619 | 1,621145394 | 0,127002639 | 12,76465914 | 2,58E-37 | 2,61E-35 |
| ENSG00000179750.16 | APOBEC3B | protein_coding | 946,8764858 | 1,613997114 | 0,203972383 | 7,912821789 | 2,52E-15 | 1,09E-13 |
| ENSG00000229589.1 | ACVR2B-AS1 | lncRNA | 94,62182968 | 1,612363345 | 0,385652029 | 4,180876084 | 2,90E-05 | 0,000393991 |
| ENSG00000246339.6 | EXTL3-AS1 | lncRNA | 37,1178407 | 1,609968448 | 0,57445478 | 2,802602579 | 0,005069209 | 0,033267679 |
| ENSG00000184988.8 | TMEM106A | protein_coding | 92,47018946 | 1,609789544 | 0,452737506 | 3,555679663 | 0,000377003 | 0,003799982 |
| ENSG00000138439.12 | FAM117B | protein_coding | 4112,958103 | 1,609606989 | 0,141029654 | 11,41325206 | 3,59E-30 | 3,02E-28 |
| ENSG00000176396.11 | EID2 | protein_coding | 6760,021724 | 1,608406559 | 0,136818992 | 11,7557258 | 6,60E-32 | 5,76E-30 |
| ENSG00000250508.1 | LINC02701 | lncRNA | 265,5714435 | 1,606104375 | 0,240429996 | 6,680133097 | 2,39E-11 | 7,77E-10 |
| ENSG00000084674.15 | APOB | protein_coding | 16587,76548 | 1,60403873 | 0,197208077 | 8,133737474 | 4,16E-16 | 1,89E-14 |
| ENSG00000166923.12 | GREM1 | protein_coding | 5962,196884 | 1,598212274 | 0,172792483 | 9,249315972 | 2,26E-20 | 1,26E-18 |
| ENSG00000123240.17 | OPTN | protein_coding | 11083,71281 | 1,597632383 | 0,119759206 | 13,34037224 | 1,35E-40 | 1,49E-38 |
| ENSG00000112232.10 | KHDRBS2 | protein_coding | 185,9620948 | 1,596260107 | 0,29578173 | 5,396750191 | 6,79E-08 | 1,49E-06 |
| ENSG00000278996.1 | FP671120.4 | lncRNA | 96,35620478 | 1,595480961 | 0,372327905 | 4,285150104 | 1,83E-05 | 0,000260072 |
| ENSG00000284734.1 | AC099063.4 | lncRNA | 48,63717422 | 1,594317214 | 0,552287556 | 2,886752015 | 0,003892409 | 0,027090286 |
| ENSG00000197085.11 | NPSR1-AS1 | lncRNA | 3821,871553 | 1,593444657 | 0,137389994 | 11,59796733 | 4,22E-31 | 3,64E-29 |
| ENSG00000146955.11 | RAB19 | protein_coding | 662,1764729 | 1,587621833 | 0,221370946 | 7,171771452 | 7,40E-13 | 2,71E-11 |
| ENSG00000161921.17 | CXCL16 | protein_coding | 2270,268498 | 1,586859115 | 0,133507921 | 11,8858799 | 1,40E-32 | 1,26E-30 |
| ENSG00000197448.14 | GSTK1 | protein_coding | 6840,576031 | 1,583026614 | 0,148004336 | 10,69581241 | 1,06E-26 | 8,05E-25 |
| ENSG00000114127.11 | XRN1 | protein_coding | 8190,669332 | 1,576602848 | 0,150766553 | 10,45724546 | 1,36E-25 | 9,87E-24 |
| ENSG00000125968.9 | ID1 | protein_coding | 3506,466553 | 1,573262316 | 0,160115591 | 9,825790908 | 8,72E-23 | 5,65E-21 |
| ENSG00000139318.8 | DUSP6 | protein_coding | 2174,847934 | 1,57236111 | 0,151142475 | 10,40317161 | 2,40E-25 | 1,72E-23 |
| ENSG00000188313.13 | PLSCR1 | protein_coding | 9203,642367 | 1,56916019 | 0,119738354 | 13,10490861 | 3,09E-39 | 3,31E-37 |
| ENSG00000008441.18 | NFIX | protein_coding | 431,0749405 | 1,567077381 | 0,320775664 | 4,885275149 | 1,03E-06 | 1,89E-05 |
| ENSG00000167550.11 | RHEBL1 | protein_coding | 384,9424833 | 1,559372675 | 0,21147184 | 7,373902246 | 1,66E-13 | 6,34E-12 |
| ENSG00000183049.13 | CAMK1D | protein_coding | 1829,092474 | 1,558058424 | 0,151512126 | 10,28339091 | 8,37E-25 | 5,86E-23 |
| ENSG00000205502.4 | C2CD4B | protein_coding | 552,4382525 | 1,549098933 | 0,187731938 | 8,251653661 | 1,56E-16 | 7,28E-15 |
| ENSG00000259721.1 | AC090877.2 | lncRNA | 62,42273073 | 1,545122976 | 0,467181416 | 3,307329706 | 0,0009419 | 0,008250489 |
| ENSG00000163814.8 | CDCP1 | protein_coding | 10138,52466 | 1,542246255 | 0,152465449 | 10,11538197 | 4,72E-24 | 3,21E-22 |
| ENSG00000107566.14 | ERLIN1 | protein_coding | 14234,9582 | 1,539592826 | 0,146920506 | 10,47908742 | 1,08E-25 | 7,86E-24 |
| ENSG00000272916.5 | AC022400.5 | protein_coding | 125,9086093 | 1,538172285 | 0,329891788 | 4,662657086 | 3,12E-06 | 5,24E-05 |
| ENSG00000147251.15 | DOCK11 | protein_coding | 45,41449414 | 1,536650525 | 0,539020904 | 2,850818055 | 0,004360691 | 0,029611117 |
| ENSG00000226453.2 | LINC02542 | lncRNA | 113,6528445 | 1,53038586 | 0,362432273 | 4,222543003 | 2,42E-05 | 0,000333581 |
| ENSG00000141431.12 | ASXL3 | protein_coding | 206,0903561 | 1,528970601 | 0,271455068 | 5,632499747 | 1,78E-08 | 4,19E-07 |
| ENSG00000185650.10 | ZFP36L1 | protein_coding | 67,56148264 | 1,5227181 | 0,491928649 | 3,095404392 | 0,001965447 | 0,015358954 |

| | | | | | | | | |
|---|---|---|---|---|---|---|---|---|
| ENSG00000012504.15 | NR1H4 | protein_coding | 208,9409515 | 1,516949078 | 0,26718202 | 5,677586681 | 1,37E-08 | 3,28E-07 |
| ENSG00000143387.14 | CTSK | protein_coding | 52,5556158 | 1,51104832 | 0,529718109 | 2,85255175 | 0,004336975 | 0,029488331 |
| ENSG00000145632.15 | PLK2 | protein_coding | 8569,931107 | 1,510639401 | 0,115868471 | 13,03753629 | 7,48E-39 | 7,83E-37 |
| ENSG00000180263.14 | FGD6 | protein_coding | 1429,84102 | 1,508296891 | 0,162122255 | 9,303453664 | 1,36E-20 | 7,76E-19 |
| ENSG00000286086.1 | AC099661.1 | lncRNA | 62,63245943 | 1,508177191 | 0,450546701 | 3,347438097 | 0,000815622 | 0,007303271 |
| ENSG00000144290.17 | SLC4A10 | protein_coding | 3945,414165 | 1,506429734 | 0,171565876 | 8,780474106 | 1,63E-18 | 8,31E-17 |
| ENSG00000155792.10 | DEPTOR | protein_coding | 1395,371804 | 1,50421445 | 0,144691066 | 10,39604235 | 2,58E-25 | 1,85E-23 |
| ENSG00000255690.3 | TRIL | protein_coding | 79,85680841 | 1,498445969 | 0,452704504 | 3,309986882 | 0,000933003 | 0,008186264 |
| ENSG00000157557.13 | ETS2 | protein_coding | 9111,290853 | 1,496957398 | 0,128294495 | 11,66813427 | 1,85E-31 | 1,61E-29 |
| ENSG00000085449.15 | WDFY1 | protein_coding | 18378,27085 | 1,496841551 | 0,171558089 | 8,724983831 | 2,66E-18 | 1,35E-16 |
| ENSG00000166006.14 | KCNC2 | protein_coding | 607,4717634 | 1,496610079 | 0,181052039 | 8,266187396 | 1,38E-16 | 6,49E-15 |
| ENSG00000155363.18 | MOV10 | protein_coding | 10096,22432 | 1,495959944 | 0,113470941 | 13,18363923 | 1,09E-39 | 1,19E-37 |
| ENSG00000230772.1 | VN1R108P | unprocessed_pseud | 46,0144307 | 1,494478355 | 0,532155683 | 2,808348012 | 0,004979638 | 0,032834244 |
| ENSG00000179603.18 | GRM8 | protein_coding | 111,920106 | 1,491682078 | 0,345367822 | 4,319111345 | 1,57E-05 | 0,000226027 |
| ENSG00000144648.16 | ACKR2 | protein_coding | 96,63931337 | 1,489668734 | 0,374012908 | 3,982934012 | 6,81E-05 | 0,00084572 |
| ENSG00000271646.1 | IRF2-DT | lncRNA | 392,0345065 | 1,484413504 | 0,222547389 | 6,670100742 | 2,56E-11 | 8,28E-10 |
| ENSG00000151012.13 | SLC7A11 | protein_coding | 302,3762816 | 1,480422418 | 0,249914073 | 5,923725712 | 3,15E-09 | 8,23E-08 |
| ENSG00000142961.15 | MOB3C | protein_coding | 1250,033709 | 1,477516322 | 0,154454602 | 9,566023335 | 1,11E-21 | 6,63E-20 |
| ENSG00000145147.20 | SLIT2 | protein_coding | 259,7092215 | 1,477463011 | 0,320709475 | 4,606857996 | 4,09E-06 | 6,72E-05 |
| ENSG00000198417.7 | MT1F | protein_coding | 1090,087307 | 1,47570448 | 0,155676023 | 9,479330567 | 2,56E-21 | 1,51E-19 |
| ENSG00000197584.12 | KCNMB2 | protein_coding | 3103,884556 | 1,475457767 | 0,135828749 | 10,86263239 | 1,74E-27 | 1,38E-25 |
| ENSG00000160710.17 | ADAR | protein_coding | 24829,54074 | 1,471539867 | 0,141410572 | 10,40615176 | 2,32E-25 | 1,67E-23 |
| ENSG00000260669.2 | AL096870.2 | lncRNA | 70,37504785 | 1,468904274 | 0,488237831 | 3,008583485 | 0,002624687 | 0,019485042 |
| ENSG00000158560.14 | DYNC1I1 | protein_coding | 350,7885691 | 1,464332224 | 0,215856385 | 6,7838263 | 1,17E-11 | 3,92E-10 |
| ENSG00000280399.1 | AC022497.1 | TEC | 40,77074716 | 1,463979033 | 0,551614072 | 2,653991456 | 0,007954584 | 0,047716108 |
| ENSG00000106100.11 | NOD1 | protein_coding | 427,2551574 | 1,463892764 | 0,219494882 | 6,669370842 | 2,57E-11 | 8,30E-10 |
| ENSG00000100628.12 | ASB2 | protein_coding | 140,0802542 | 1,456900671 | 0,32841445 | 4,436164946 | 9,16E-06 | 0,000140372 |
| ENSG00000144118.14 | RALB | protein_coding | 5484,051575 | 1,45544726 | 0,135218218 | 10,76369207 | 5,11E-27 | 3,93E-25 |
| ENSG00000151689.13 | INPP1 | protein_coding | 4439,548299 | 1,453338179 | 0,123469134 | 11,77086231 | 5,52E-32 | 4,85E-30 |
| ENSG00000088387.19 | DOCK9 | protein_coding | 3688,161093 | 1,451028584 | 0,153574939 | 9,448342244 | 3,44E-21 | 2,01E-19 |
| ENSG00000231133.8 | HAR1B | lncRNA | 97,9885372 | 1,44826066 | 0,428641916 | 3,378719175 | 0,000728244 | 0,006625719 |
| ENSG00000180881.19 | CAPS2 | protein_coding | 733,201697 | 1,445143623 | 0,171737509 | 8,414839768 | 3,93E-17 | 1,88E-15 |
| ENSG00000090932.11 | DLL3 | protein_coding | 599,7099211 | 1,438743718 | 0,213048164 | 6,753138324 | 1,45E-11 | 4,78E-10 |
| ENSG00000166750.10 | SLFN5 | protein_coding | 809,371863 | 1,433699722 | 0,23420976 | 6,12143458 | 9,27E-10 | 2,56E-08 |
| ENSG00000260083.2 | MIR762HG | lncRNA | 236,8164758 | 1,431380837 | 0,258992976 | 5,526716812 | 3,26E-08 | 7,46E-07 |
| ENSG00000187634.12 | SAMD11 | protein_coding | 613,4715746 | 1,427907802 | 0,207546408 | 6,879944663 | 5,99E-12 | 2,06E-10 |

| | | | | | | | | |
|---|---|---|---|---|---|---|---|---|
| ENSG00000152061.23 | RABGAP1L | protein_coding | 8208,064856 | 1,427431286 | 0,11326737 | 12,60231681 | 2,05E-36 | 2,05E-34 |
| ENSG00000143891.17 | GALM | protein_coding | 1459,118264 | 1,419406248 | 0,14459748 | 9,816258573 | 9,58E-23 | 6,15E-21 |
| ENSG00000164211.13 | STARD4 | protein_coding | 11654,6805 | 1,414034012 | 0,112776295 | 12,53839749 | 4,60E-36 | 4,59E-34 |
| ENSG00000237976.1 | RFX5-AS1 | lncRNA | 76,10018119 | 1,414016705 | 0,41112427 | 3,439390001 | 0,000583027 | 0,005505486 |
| ENSG00000272755.1 | AC245297.2 | lncRNA | 84,5575007 | 1,412920046 | 0,403379996 | 3,502702315 | 0,000460564 | 0,004516163 |
| ENSG00000267248.3 | AC025048.2 | lncRNA | 148,2324645 | 1,411141481 | 0,315228466 | 4,476567421 | 7,59E-06 | 0,000118153 |
| ENSG00000102524.11 | TNFSF13B | protein_coding | 396,7666889 | 1,41006411 | 0,24123569 | 5,845172047 | 5,06E-09 | 1,28E-07 |
| ENSG00000013016.16 | EHD3 | protein_coding | 1284,272958 | 1,405781951 | 0,153668046 | 9,148173513 | 5,79E-20 | 3,18E-18 |
| ENSG00000183734.4 | ASCL2 | protein_coding | 8092,58024 | 1,404942007 | 0,13077203 | 10,74344417 | 6,36E-27 | 4,84E-25 |
| ENSG00000092098.17 | RNF31 | protein_coding | 2708,502999 | 1,404832241 | 0,16312758 | 8,611862215 | 7,19E-18 | 3,53E-16 |
| ENSG00000197208.6 | SLC22A4 | protein_coding | 342,430772 | 1,402601094 | 0,236681101 | 5,926122066 | 3,10E-09 | 8,12E-08 |
| ENSG00000184500.16 | PROS1 | protein_coding | 1225,363296 | 1,400680106 | 0,165774241 | 8,449323012 | 2,93E-17 | 1,40E-15 |
| ENSG00000224420.3 | ADM5 | protein_coding | 85,7838376 | 1,40038238 | 0,405746703 | 3,45137094 | 0,000557746 | 0,005309878 |
| ENSG00000272196.2 | H2AC19 | protein_coding | 2056,42979 | 1,398600929 | 0,135230291 | 10,34236429 | 4,53E-25 | 3,19E-23 |
| ENSG00000115756.13 | HPCAL1 | protein_coding | 2528,667306 | 1,39640974 | 0,136114316 | 10,25909533 | 1,08E-24 | 7,49E-23 |
| ENSG00000163412.13 | EIF4E3 | protein_coding | 5904,160794 | 1,396297872 | 0,116294788 | 12,00653871 | 3,28E-33 | 3,00E-31 |
| ENSG00000167654.18 | ATCAY | protein_coding | 1601,952655 | 1,394933073 | 0,213582387 | 6,531124078 | 6,53E-11 | 2,00E-09 |
| ENSG00000118507.17 | AKAP7 | protein_coding | 200,1260741 | 1,391919926 | 0,280370214 | 4,964578463 | 6,89E-07 | 1,30E-05 |
| ENSG00000125826.21 | RBCK1 | protein_coding | 8680,104093 | 1,390081944 | 0,114014565 | 12,19214351 | 3,42E-34 | 3,20E-32 |
| ENSG00000124875.10 | CXCL6 | protein_coding | 354,8956353 | 1,388864342 | 0,225343336 | 6,16332557 | 7,12E-10 | 1,99E-08 |
| ENSG00000260992.1 | DOCK9-DT | lncRNA | 46,74379639 | 1,387556309 | 0,526368724 | 2,636091862 | 0,008386702 | 0,049766367 |
| ENSG00000135447.17 | PPP1R1A | protein_coding | 26969,81238 | 1,386300592 | 0,108952934 | 12,72384821 | 4,36E-37 | 4,39E-35 |
| ENSG00000133943.21 | DGLUCY | protein_coding | 1218,756923 | 1,384687826 | 0,14886045 | 9,301918844 | 1,38E-20 | 7,85E-19 |
| ENSG00000260459.2 | FTLP14 | processed_pseudog | 85,83427951 | 1,381220728 | 0,413521557 | 3,34014202 | 0,000837356 | 0,007465945 |
| ENSG00000183840.7 | GPR39 | protein_coding | 72,10259852 | 1,381040861 | 0,454278971 | 3,040072182 | 0,002365215 | 0,017892872 |
| ENSG00000163874.11 | ZC3H12A | protein_coding | 302,3624217 | 1,379860236 | 0,271431868 | 5,083633858 | 3,70E-07 | 7,30E-06 |
| ENSG00000124151.19 | NCOA3 | protein_coding | 4841,920387 | 1,379350023 | 0,222386805 | 6,202481405 | 5,56E-10 | 1,57E-08 |
| ENSG00000279631.1 | AL158211.5 | TEC | 78,21664878 | 1,378632084 | 0,420900358 | 3,275435764 | 0,001054991 | 0,00908228 |
| ENSG00000063180.9 | CA11 | protein_coding | 1311,842752 | 1,374951474 | 0,181737424 | 7,565593499 | 3,86E-14 | 1,56E-12 |
| ENSG00000135605.13 | TEC | protein_coding | 429,5427632 | 1,37279544 | 0,232310343 | 5,909316911 | 3,44E-09 | 8,92E-08 |
| ENSG00000130224.15 | LRCH2 | protein_coding | 3732,781322 | 1,371995271 | 0,122765384 | 11,17575029 | 5,36E-29 | 4,38E-27 |
| ENSG00000133019.12 | CHRM3 | protein_coding | 1747,059854 | 1,368253358 | 0,138684637 | 9,865933146 | 5,85E-23 | 3,83E-21 |
| ENSG00000108679.13 | LGALS3BP | protein_coding | 38839,51174 | 1,365980102 | 0,14481009 | 9,432906947 | 3,99E-21 | 2,32E-19 |
| ENSG00000255508.7 | AP002990.1 | protein_coding | 153,8963804 | 1,357097113 | 0,30513199 | 4,447574021 | 8,68E-06 | 0,000133685 |
| ENSG00000143390.18 | RFX5 | protein_coding | 5209,698147 | 1,356861026 | 0,139343345 | 9,737537361 | 2,09E-22 | 1,31E-20 |
| ENSG00000067057.18 | PFKP | protein_coding | 3371,772344 | 1,355076542 | 0,132438071 | 10,23177499 | 1,43E-24 | 9,90E-23 |

| | | | | | | | | |
|---|---|---|---|---|---|---|---|---|
| ENSG00000133116.8 | KL | protein_coding | 1167,975972 | 1,351855904 | 0,151605144 | 8,916952729 | 4,79E-19 | 2,51E-17 |
| ENSG00000153234.15 | NR4A2 | protein_coding | 277,6058127 | 1,350331349 | 0,245988274 | 5,489413483 | 4,03E-08 | 9,11E-07 |
| ENSG00000136842.14 | TMOD1 | protein_coding | 11482,37146 | 1,347213801 | 0,138389459 | 9,734945201 | 2,14E-22 | 1,34E-20 |
| ENSG00000124762.14 | CDKN1A | protein_coding | 1307,455787 | 1,343659151 | 0,165846504 | 8,101823797 | 5,41E-16 | 2,45E-14 |
| ENSG00000038945.15 | MSR1 | protein_coding | 93,72547709 | 1,340873792 | 0,450627848 | 2,975567974 | 0,002924465 | 0,021310897 |
| ENSG00000105499.14 | PLA2G4C | protein_coding | 341,6104992 | 1,340494377 | 0,288112157 | 4,652682451 | 3,28E-06 | 5,48E-05 |
| ENSG00000104325.7 | DECR1 | protein_coding | 5545,84726 | 1,336818082 | 0,132890749 | 10,05952703 | 8,34E-24 | 5,58E-22 |
| ENSG00000103056.12 | SMPD3 | protein_coding | 1013,385669 | 1,336233651 | 0,185473524 | 7,204444166 | 5,83E-13 | 2,16E-11 |
| ENSG00000176928.7 | GCNT4 | protein_coding | 1213,981222 | 1,334881091 | 0,149879259 | 8,906376388 | 5,27E-19 | 2,76E-17 |
| ENSG00000142677.4 | IL22RA1 | protein_coding | 814,2392195 | 1,333699641 | 0,179473089 | 7,431195649 | 1,08E-13 | 4,23E-12 |
| ENSG00000172780.17 | RAB43 | protein_coding | 785,6613188 | 1,326740759 | 0,22661075 | 5,854712363 | 4,78E-09 | 1,21E-07 |
| ENSG00000121361.5 | KCNJ8 | protein_coding | 115,5316333 | 1,326179248 | 0,34024398 | 3,897730236 | 9,71E-05 | 0,001165572 |
| ENSG00000134575.13 | ACP2 | protein_coding | 13168,04046 | 1,323942853 | 0,115069406 | 11,5056026 | 1,24E-30 | 1,05E-28 |
| ENSG00000115604.11 | IL18R1 | protein_coding | 1294,47786 | 1,322532911 | 0,166216276 | 7,956699194 | 1,77E-15 | 7,74E-14 |
| ENSG00000147509.14 | RGS20 | protein_coding | 111,2709396 | 1,320866832 | 0,34900584 | 3,784655383 | 0,000153922 | 0,001746532 |
| ENSG00000145708.11 | CRHBP | protein_coding | 57,04577118 | 1,319418053 | 0,4690904 | 2,812715953 | 0,004912502 | 0,032478624 |
| ENSG00000055332.18 | EIF2AK2 | protein_coding | 5059,120439 | 1,317594294 | 0,135772225 | 9,704446489 | 2,89E-22 | 1,78E-20 |
| ENSG00000099251.14 | HSD17B7P2 | transcribed_unproc | 83,57317475 | 1,314886927 | 0,41149549 | 3,195386 | 0,001396439 | 0,011569667 |
| ENSG00000166148.4 | AVPR1A | protein_coding | 113,9758497 | 1,314336359 | 0,341452181 | 3,849254547 | 0,000118478 | 0,001387987 |
| ENSG00000263528.8 | IKBKE | protein_coding | 859,6857308 | 1,312187566 | 0,165558652 | 7,925816911 | 2,27E-15 | 9,84E-14 |
| ENSG00000006062.17 | MAP3K14 | protein_coding | 122,8788579 | 1,301405151 | 0,399789449 | 3,255226355 | 0,00113302 | 0,009669702 |
| ENSG00000116761.12 | CTH | protein_coding | 215,1508956 | 1,300788922 | 0,266191063 | 4,886673914 | 1,03E-06 | 1,88E-05 |
| ENSG00000155287.11 | SLC25A28 | protein_coding | 1013,65343 | 1,299710891 | 0,152673127 | 8,513029855 | 1,69E-17 | 8,19E-16 |
| ENSG00000114450.11 | GNB4 | protein_coding | 3992,793691 | 1,298932336 | 0,140905031 | 9,218495106 | 3,01E-20 | 1,67E-18 |
| ENSG00000099377.14 | HSD3B7 | protein_coding | 747,9788128 | 1,2966254 | 0,164932864 | 7,861534474 | 3,79E-15 | 1,63E-13 |
| ENSG00000135373.13 | EHF | protein_coding | 1602,415692 | 1,293978891 | 0,196590021 | 6,582118906 | 4,64E-11 | 1,45E-09 |
| ENSG00000152092.16 | ASTN1 | protein_coding | 720,5049443 | 1,293923464 | 0,205319596 | 6,301996918 | 2,94E-10 | 8,54E-09 |
| ENSG00000171560.16 | FGA | protein_coding | 83,70733658 | 1,290580831 | 0,439600346 | 2,935804856 | 0,003326835 | 0,023705684 |
| ENSG00000123358.20 | NR4A1 | protein_coding | 198,744621 | 1,289513013 | 0,293124913 | 4,399192823 | 1,09E-05 | 0,000164237 |
| ENSG00000283149.1 | AC068631.2 | protein_coding | 641,73508 | 1,288536807 | 0,297821293 | 4,326543593 | 1,51E-05 | 0,000220053 |
| ENSG00000152332.16 | UHMK1 | protein_coding | 15973,89422 | 1,283265806 | 0,122059859 | 10,5134138 | 7,49E-26 | 5,50E-24 |
| ENSG00000126709.15 | IFI6 | protein_coding | 9870,774876 | 1,276630888 | 0,126369919 | 10,1023321 | 5,39E-24 | 3,65E-22 |
| ENSG00000068971.14 | PPP2R5B | protein_coding | 3665,592264 | 1,274571726 | 0,13487979 | 9,449686456 | 3,40E-21 | 1,99E-19 |
| ENSG00000251201.8 | TMED7-TICAM2 | protein_coding | 1097,742661 | 1,273159081 | 0,171391391 | 7,428372421 | 1,10E-13 | 4,31E-12 |
| ENSG00000173221.14 | GLRX | protein_coding | 2227,581462 | 1,27254499 | 0,170425676 | 7,466861938 | 8,21E-14 | 3,24E-12 |
| ENSG00000165246.14 | NLGN4Y | protein_coding | 3402,807136 | 1,267766955 | 0,161727841 | 7,838891242 | 4,55E-15 | 1,94E-13 |

| | | | | | | | | |
|---|---|---|---|---|---|---|---|---|
| ENSG00000070759.17 | TESK2 | protein_coding | 553,9919499 | 1,266210725 | 0,193038694 | 6,559362274 | 5,40E-11 | 1,67E-09 |
| ENSG00000143882.12 | ATP6V1C2 | protein_coding | 196,2442705 | 1,265768637 | 0,429856833 | 2,944628394 | 0,003233428 | 0,023150413 |
| ENSG00000228133.3 | AC099684.1 | lncRNA | 168,7648489 | 1,26336903 | 0,296910052 | 4,255056439 | 2,09E-05 | 0,00029345 |
| ENSG00000137502.10 | RAB30 | protein_coding | 1628,303043 | 1,2625297 | 0,170367345 | 7,410632029 | 1,26E-13 | 4,89E-12 |
| ENSG00000172965.17 | MIR4435-2HG | lncRNA | 60,24960325 | 1,262234513 | 0,462104929 | 2,731488963 | 0,006304885 | 0,039722378 |
| ENSG00000091844.8 | RGS17 | protein_coding | 3699,365179 | 1,26103637 | 0,121009809 | 10,42094341 | 1,99E-25 | 1,44E-23 |
| ENSG00000259834.1 | AL365361.1 | lncRNA | 1210,002342 | 1,259116242 | 0,181693173 | 6,929903968 | 4,21E-12 | 1,47E-10 |
| ENSG00000187193.9 | MT1X | protein_coding | 1796,804731 | 1,25899955 | 0,136688279 | 9,210735269 | 3,24E-20 | 1,79E-18 |
| ENSG00000145362.20 | ANK2 | protein_coding | 2967,453002 | 1,257285621 | 0,201644678 | 6,235153996 | 4,51E-10 | 1,29E-08 |
| ENSG00000116833.14 | NR5A2 | protein_coding | 77,14171229 | 1,255792398 | 0,471064835 | 2,665858933 | 0,007679191 | 0,046396496 |
| ENSG00000154134.15 | ROBO3 | protein_coding | 107,3244672 | 1,255594634 | 0,443635009 | 2,830242451 | 0,004651274 | 0,031119983 |
| ENSG00000230490.3 | AL139383.1 | lncRNA | 146,9118268 | 1,252305092 | 0,365179506 | 3,429286341 | 0,000605171 | 0,005673645 |
| ENSG00000111252.11 | SH2B3 | protein_coding | 1606,230258 | 1,250294225 | 0,186436974 | 6,706256804 | 2,00E-11 | 6,53E-10 |
| ENSG00000141971.13 | MVB12A | protein_coding | 3912,975498 | 1,245662036 | 0,14065264 | 8,85630044 | 8,27E-19 | 4,28E-17 |
| ENSG00000117266.16 | CDK18 | protein_coding | 86,64233118 | 1,245526694 | 0,414797345 | 3,002735456 | 0,002675649 | 0,019765248 |
| ENSG00000163683.12 | SMIM14 | protein_coding | 3032,308814 | 1,239708571 | 0,155805754 | 7,956757285 | 1,77E-15 | 7,74E-14 |
| ENSG00000144182.17 | LIPT1 | protein_coding | 284,6075937 | 1,239528259 | 0,23447235 | 5,286458118 | 1,25E-07 | 2,62E-06 |
| ENSG00000054392.13 | HHAT | protein_coding | 935,8611806 | 1,239288732 | 0,165246045 | 7,49965744 | 6,40E-14 | 2,54E-12 |
| ENSG00000148483.9 | TMEM236 | protein_coding | 84,53798694 | 1,238999461 | 0,40120803 | 3,088172143 | 0,002013918 | 0,015685033 |
| ENSG00000178695.6 | KCTD12 | protein_coding | 21858,51823 | 1,238232846 | 0,125379455 | 9,875883114 | 5,30E-23 | 3,49E-21 |
| ENSG00000128965.13 | CHAC1 | protein_coding | 347,1347489 | 1,238106809 | 0,218567603 | 5,664640097 | 1,47E-08 | 3,53E-07 |
| ENSG00000164056.11 | SPRY1 | protein_coding | 199,8588655 | 1,236904132 | 0,313615233 | 3,944018022 | 8,01E-05 | 0,000980978 |
| ENSG00000169435.14 | RASSF6 | protein_coding | 17411,53554 | 1,235656048 | 0,119014844 | 10,38236916 | 2,98E-25 | 2,11E-23 |
| ENSG00000129667.13 | RHBDF2 | protein_coding | 501,7657479 | 1,234303252 | 0,186796242 | 6,607752045 | 3,90E-11 | 1,23E-09 |
| ENSG00000234678.2 | ELF3-AS1 | lncRNA | 678,570275 | 1,234261266 | 0,182181061 | 6,77491536 | 1,24E-11 | 4,14E-10 |
| ENSG00000232721.2 | AC239800.2 | lncRNA | 275,7122573 | 1,233124977 | 0,242781159 | 5,079162577 | 3,79E-07 | 7,44E-06 |
| ENSG00000180628.15 | PCGF5 | protein_coding | 8706,820019 | 1,23296777 | 0,116874767 | 10,54947784 | 5,11E-26 | 3,77E-24 |
| ENSG00000184898.7 | RBM43 | protein_coding | 157,8864963 | 1,226821622 | 0,30911627 | 3,968803131 | 7,22E-05 | 0,000893753 |
| ENSG00000153029.14 | MR1 | protein_coding | 1674,033287 | 1,225546999 | 0,136565888 | 8,974034564 | 2,86E-19 | 1,51E-17 |
| ENSG00000267454.6 | ZNF582-AS1 | lncRNA | 127,3171033 | 1,225373939 | 0,343636082 | 3,565905917 | 0,000362601 | 0,003676059 |
| ENSG00000185420.19 | SMYD3 | protein_coding | 2230,05966 | 1,22388051 | 0,141653115 | 8,639983044 | 5,62E-18 | 2,78E-16 |
| ENSG00000198604.11 | BAZ1A | protein_coding | 1734,694502 | 1,220279264 | 0,142119296 | 8,58630249 | 8,98E-18 | 4,39E-16 |
| ENSG00000169116.11 | PARM1 | protein_coding | 12581,55671 | 1,219494968 | 0,13137702 | 9,282406978 | 1,66E-20 | 9,38E-19 |
| ENSG00000168398.7 | BDKRB2 | protein_coding | 116,9067341 | 1,218100647 | 0,366696732 | 3,321820294 | 0,000894323 | 0,00789319 |
| ENSG00000213516.10 | RBMXL1 | protein_coding | 2973,125331 | 1,21670795 | 0,126938758 | 9,584999613 | 9,25E-22 | 5,55E-20 |
| ENSG00000260802.1 | SERTM2 | protein_coding | 264,1500554 | 1,214180535 | 0,240930734 | 5,039541924 | 4,67E-07 | 9,04E-06 |

| | | | | | | | | |
|---|---|---|---|---|---|---|---|---|
| ENSG00000075426.12 | FOSL2 | protein_coding | 951,3633638 | 1,212799413 | 0,156791317 | 7,735118477 | 1,03E-14 | 4,32E-13 |
| ENSG00000123338.13 | NCKAP1L | protein_coding | 108,9508913 | 1,211745568 | 0,360399042 | 3,362233041 | 0,000773148 | 0,006985698 |
| ENSG00000253276.4 | CCDC71L | protein_coding | 244,9982518 | 1,208966618 | 0,281812943 | 4,289961301 | 1,79E-05 | 0,000255022 |
| ENSG00000102575.13 | ACP5 | protein_coding | 1053,594924 | 1,208572164 | 0,159315543 | 7,58602798 | 3,30E-14 | 1,34E-12 |
| ENSG00000272419.6 | LINC01145 | transcribed_unproc | 1455,405632 | 1,208264927 | 0,142717859 | 8,466108865 | 2,54E-17 | 1,22E-15 |
| ENSG00000134962.7 | KLB | protein_coding | 329,7783052 | 1,206071532 | 0,220792224 | 5,462472865 | 4,70E-08 | 1,05E-06 |
| ENSG00000164761.9 | TNFRSF11B | protein_coding | 306,1547433 | 1,205315407 | 0,344031978 | 3,503498179 | 0,00045919 | 0,004508977 |
| ENSG00000183715.13 | OPCML | protein_coding | 256,4217566 | 1,205288891 | 0,261691829 | 4,605756677 | 4,11E-06 | 6,75E-05 |
| ENSG00000168461.13 | RAB31 | protein_coding | 229,6502041 | 1,204939387 | 0,261709899 | 4,604103214 | 4,14E-06 | 6,80E-05 |
| ENSG00000217442.4 | SYCE3 | protein_coding | 125,2236372 | 1,20490775 | 0,35906437 | 3,355687308 | 0,00079168 | 0,007116284 |
| ENSG00000171940.13 | ZNF217 | protein_coding | 86,89346021 | 1,203944256 | 0,386941352 | 3,111438595 | 0,001861782 | 0,014680378 |
| ENSG00000101638.14 | ST8SIA5 | protein_coding | 481,1171239 | 1,203259428 | 0,23972374 | 5,019358642 | 5,18E-07 | 9,98E-06 |
| ENSG00000139112.11 | GABARAPL1 | protein_coding | 101,0304326 | 1,201146589 | 0,366063032 | 3,281256188 | 0,001033458 | 0,008932886 |
| ENSG00000198963.11 | RORB | protein_coding | 2486,05326 | 1,200532429 | 0,164542945 | 7,296164716 | 2,96E-13 | 1,12E-11 |
| ENSG00000175106.17 | TVP23C | protein_coding | 290,5808559 | 1,200010114 | 0,267983804 | 4,477920293 | 7,54E-06 | 0,000117494 |
| ENSG00000101236.17 | RNF24 | protein_coding | 2316,372806 | 1,199299019 | 0,1572489 | 7,626756176 | 2,41E-14 | 9,88E-13 |
| ENSG00000043143.20 | JADE2 | protein_coding | 2939,557212 | 1,196502904 | 0,136456937 | 8,76835528 | 1,81E-18 | 9,19E-17 |
| ENSG00000083799.18 | CYLD | protein_coding | 11150,72151 | 1,195886251 | 0,118179229 | 10,11925923 | 4,54E-24 | 3,09E-22 |
| ENSG00000010818.10 | HIVEP2 | protein_coding | 1325,033586 | 1,19329913 | 0,245427265 | 4,86212944 | 1,16E-06 | 2,11E-05 |
| ENSG00000173531.15 | MST1 | protein_coding | 198,1025018 | 1,192027468 | 0,331425126 | 3,596671997 | 0,000322314 | 0,003312545 |
| ENSG00000249249.2 | AC010226.1 | lncRNA | 158,2989878 | 1,186885047 | 0,35184468 | 3,373320992 | 0,000742673 | 0,006739429 |
| ENSG00000159110.20 | IFNAR2 | protein_coding | 3496,435722 | 1,186781176 | 0,124180321 | 9,556918235 | 1,21E-21 | 7,22E-20 |
| ENSG00000259976.3 | AC093010.2 | lncRNA | 296,1189032 | 1,186715181 | 0,256196511 | 4,632050516 | 3,62E-06 | 6,02E-05 |
| ENSG00000143153.13 | ATP1B1 | protein_coding | 18214,78773 | 1,186688215 | 0,137672354 | 8,619655153 | 6,72E-18 | 3,31E-16 |
| ENSG00000112182.15 | BACH2 | protein_coding | 1027,410275 | 1,186458435 | 0,168902005 | 7,024537295 | 2,15E-12 | 7,67E-11 |
| ENSG00000100918.13 | REC8 | protein_coding | 11202,83124 | 1,1845635 | 0,117394023 | 10,09049244 | 6,09E-24 | 4,11E-22 |
| ENSG00000164054.15 | SHISA5 | protein_coding | 28474,11441 | 1,184418531 | 0,109392375 | 10,82724945 | 2,56E-27 | 2,01E-25 |
| ENSG00000100225.18 | FBXO7 | protein_coding | 14432,20407 | 1,18403157 | 0,115404368 | 10,25985059 | 1,07E-24 | 7,46E-23 |
| ENSG00000145675.15 | PIK3R1 | protein_coding | 10469,10325 | 1,183719668 | 0,120553183 | 9,819066054 | 9,32E-23 | 6,00E-21 |
| ENSG00000230356.1 | NCAPD2P1 | processed_pseudog | 196,8975411 | 1,181650016 | 0,267180112 | 4,422672067 | 9,75E-06 | 0,000148754 |
| ENSG00000167711.14 | SERPINF2 | protein_coding | 783,2257583 | 1,181625069 | 0,173604267 | 6,806428734 | 1,00E-11 | 3,37E-10 |
| ENSG00000273108.1 | AL121929.2 | lncRNA | 204,9256623 | 1,180649869 | 0,279603717 | 4,222582888 | 2,42E-05 | 0,000333581 |
| ENSG00000077063.11 | CTTNBP2 | protein_coding | 1003,668855 | 1,179701401 | 0,169567016 | 6,957139607 | 3,47E-12 | 1,23E-10 |
| ENSG00000227617.9 | CERS6-AS1 | lncRNA | 440,8970453 | 1,179221421 | 0,302080976 | 3,903659993 | 9,47E-05 | 0,001142452 |
| ENSG00000275993.3 | SIK1B | protein_coding | 102,0696449 | 1,178306215 | 0,427539232 | 2,756018928 | 0,005850961 | 0,037355862 |
| ENSG00000166250.12 | CLMP | protein_coding | 546,4729426 | 1,17787814 | 0,249253709 | 4,725619305 | 2,29E-06 | 3,95E-05 |

| | | | | | | | | |
|---|---|---|---|---|---|---|---|---|
| ENSG00000178075.20 | GRAMD1C | protein_coding | 838,9978393 | 1,174760879 | 0,178824744 | 6,569341887 | 5,05E-11 | 1,57E-09 |
| ENSG00000100902.11 | PSMA6 | protein_coding | 19422,68484 | 1,172875014 | 0,14279013 | 8,213978176 | 2,14E-16 | 9,89E-15 |
| ENSG00000163435.16 | ELF3 | protein_coding | 6176,043004 | 1,17084177 | 0,118846207 | 9,851738621 | 6,74E-23 | 4,39E-21 |
| ENSG00000104320.14 | NBN | protein_coding | 7607,681973 | 1,168984653 | 0,119749589 | 9,761909494 | 1,64E-22 | 1,04E-20 |
| ENSG00000065427.15 | KARS1 | protein_coding | 16734,99546 | 1,166101804 | 0,118632015 | 9,829570924 | 8,40E-23 | 5,46E-21 |
| ENSG00000100271.17 | TTLL1 | protein_coding | 403,245277 | 1,166071106 | 0,209023266 | 5,578666579 | 2,42E-08 | 5,62E-07 |
| ENSG00000162407.9 | PLPP3 | protein_coding | 1253,246547 | 1,165775779 | 0,162105378 | 7,191468874 | 6,41E-13 | 2,36E-11 |
| ENSG00000104805.16 | NUCB1 | protein_coding | 38269,51246 | 1,165380164 | 0,120547803 | 9,667369603 | 4,15E-22 | 2,54E-20 |
| ENSG00000286016.1 | AC079949.3 | lncRNA | 109,2635487 | 1,162211538 | 0,345509206 | 3,363764321 | 0,000768872 | 0,006950056 |
| ENSG00000196116.8 | TDRD7 | protein_coding | 4827,802807 | 1,15902066 | 0,1180319 | 9,819554413 | 9,28E-23 | 5,99E-21 |
| ENSG00000133740.11 | E2F5 | protein_coding | 3352,780417 | 1,158574064 | 0,15138422 | 7,653202333 | 1,96E-14 | 8,09E-13 |
| ENSG00000178764.8 | ZHX2 | protein_coding | 95,08105723 | 1,156924488 | 0,416967221 | 2,774617357 | 0,005526669 | 0,035732185 |
| ENSG00000170381.14 | SEMA3E | protein_coding | 112,9224402 | 1,155872313 | 0,358740747 | 3,222026838 | 0,001272872 | 0,010722219 |
| ENSG00000183722.9 | LHFPL6 | protein_coding | 2684,100159 | 1,153641555 | 0,148091669 | 7,790050311 | 6,70E-15 | 2,84E-13 |
| ENSG00000143507.18 | DUSP10 | protein_coding | 926,2172015 | 1,151660502 | 0,17388073 | 6,62327849 | 3,51E-11 | 1,11E-09 |
| ENSG00000188599.17 | NPIPP1 | transcribed_unproc | 117,8324313 | 1,146683379 | 0,364871753 | 3,142702522 | 0,001673959 | 0,013509763 |
| ENSG00000100600.15 | LGMN | protein_coding | 8342,07291 | 1,143533524 | 0,121085004 | 9,444055708 | 3,59E-21 | 2,09E-19 |
| ENSG00000272752.6 | STAG3L5P-PVRIG | lncRNA | 371,8363689 | 1,141835863 | 0,29351279 | 3,890242276 | 0,000100144 | 0,001194597 |
| ENSG00000164181.14 | ELOVL7 | protein_coding | 1726,949736 | 1,140009485 | 0,156693616 | 7,275404811 | 3,45E-13 | 1,30E-11 |
| ENSG00000177272.9 | KCNA3 | protein_coding | 704,2093823 | 1,137733846 | 0,218411559 | 5,20912837 | 1,90E-07 | 3,89E-06 |
| ENSG00000149932.17 | TMEM219 | protein_coding | 3940,91337 | 1,137542294 | 0,128027025 | 8,885173188 | 6,38E-19 | 3,32E-17 |
| ENSG00000138685.17 | FGF2 | protein_coding | 1336,246619 | 1,131809623 | 0,154933574 | 7,305128196 | 2,77E-13 | 1,05E-11 |
| ENSG00000119139.21 | TJP2 | protein_coding | 21098,69936 | 1,130477819 | 0,121742812 | 9,285786972 | 1,61E-20 | 9,11E-19 |
| ENSG00000197721.17 | CR1L | protein_coding | 121,9739586 | 1,130202885 | 0,34264473 | 3,298468605 | 0,000972138 | 0,008469288 |
| ENSG00000122694.16 | GLIPR2 | protein_coding | 1574,154773 | 1,129743037 | 0,165346034 | 6,832598326 | 8,34E-12 | 2,83E-10 |
| ENSG00000257900.2 | AL162632.1 | lncRNA | 91,25270393 | 1,129618378 | 0,419009296 | 2,695926772 | 0,00701931 | 0,043233082 |
| ENSG00000136153.20 | LMO7 | protein_coding | 6380,862076 | 1,128183737 | 0,12961795 | 8,703915888 | 3,21E-18 | 1,60E-16 |
| ENSG00000179456.10 | ZBTB18 | protein_coding | 1911,913466 | 1,12578002 | 0,174111413 | 6,465859988 | 1,01E-10 | 3,03E-09 |
| ENSG00000062038.14 | CDH3 | protein_coding | 744,7403195 | 1,123546883 | 0,205396173 | 5,470145162 | 4,50E-08 | 1,01E-06 |
| ENSG00000256574.8 | OR13A1 | protein_coding | 287,8735964 | 1,121608806 | 0,228595623 | 4,906519162 | 9,27E-07 | 1,71E-05 |
| ENSG00000125952.20 | MAX | protein_coding | 6321,12742 | 1,117595581 | 0,123448871 | 9,053104903 | 1,39E-19 | 7,50E-18 |
| ENSG00000003402.20 | CFLAR | protein_coding | 2732,871104 | 1,115290585 | 0,143459818 | 7,774236716 | 7,59E-15 | 3,20E-13 |
| ENSG00000164749.13 | HNF4G | protein_coding | 2812,146709 | 1,112640752 | 0,12971401 | 8,577645184 | 9,68E-18 | 4,71E-16 |
| ENSG00000164164.17 | OTUD4 | protein_coding | 4706,910732 | 1,111854071 | 0,130443713 | 8,523630961 | 1,55E-17 | 7,49E-16 |
| ENSG00000177519.4 | RPRM | protein_coding | 287,0680093 | 1,109411631 | 0,233889441 | 4,743316441 | 2,10E-06 | 3,65E-05 |
| ENSG00000145246.14 | ATP10D | protein_coding | 9897,056275 | 1,108613508 | 0,11483172 | 9,654244518 | 4,72E-22 | 2,88E-20 |

| | | | | | | | | |
|---|---|---|---|---|---|---|---|---|
| ENSG00000121060.18 | TRIM25 | protein_coding | 7999,598188 | 1,10772533 | 0,113987589 | 9,717946837 | 2,53E-22 | 1,57E-20 |
| ENSG00000118894.15 | EEF2KMT | protein_coding | 1607,497704 | 1,10731825 | 0,16198691 | 6,835850172 | 8,15E-12 | 2,77E-10 |
| ENSG00000204044.6 | SLC12A5-AS1 | lncRNA | 379,062502 | 1,106790817 | 0,27540313 | 4,018802606 | 5,85E-05 | 0,000735929 |
| ENSG00000106069.23 | CHN2 | protein_coding | 1533,320641 | 1,106536506 | 0,19757539 | 5,600578633 | 2,14E-08 | 5,01E-07 |
| ENSG00000181444.13 | ZNF467 | protein_coding | 181,4901167 | 1,105514087 | 0,36264725 | 3,048455727 | 0,002300208 | 0,017485423 |
| ENSG00000106605.11 | BLVRA | protein_coding | 664,618187 | 1,102782052 | 0,181465448 | 6,077091071 | 1,22E-09 | 3,35E-08 |
| ENSG00000137101.13 | CD72 | protein_coding | 319,9506273 | 1,102643514 | 0,236382155 | 4,664664788 | 3,09E-06 | 5,19E-05 |
| ENSG00000150687.12 | PRSS23 | protein_coding | 4358,005489 | 1,101447691 | 0,126433746 | 8,711659093 | 2,99E-18 | 1,51E-16 |
| ENSG00000095794.19 | CREM | protein_coding | 1447,703162 | 1,096134146 | 0,145675894 | 7,524471729 | 5,29E-14 | 2,12E-12 |
| ENSG00000078328.22 | RBFOX1 | protein_coding | 102,4938921 | 1,095449555 | 0,389939224 | 2,809282801 | 0,004965201 | 0,032759686 |
| ENSG00000274265.5 | AC245297.3 | lncRNA | 1236,674823 | 1,095160682 | 0,144454628 | 7,581347161 | 3,42E-14 | 1,38E-12 |
| ENSG00000151834.16 | GABRA2 | protein_coding | 1512,070113 | 1,094509413 | 0,164179215 | 6,666552856 | 2,62E-11 | 8,42E-10 |
| ENSG00000128908.17 | INO80 | protein_coding | 2894,694038 | 1,094286665 | 0,140693191 | 7,77782249 | 7,38E-15 | 3,12E-13 |
| ENSG00000170365.10 | SMAD1 | protein_coding | 517,0287728 | 1,091606596 | 0,192908286 | 5,658681754 | 1,53E-08 | 3,64E-07 |
| ENSG00000143106.13 | PSMA5 | protein_coding | 12591,85876 | 1,091498245 | 0,124404506 | 8,773783814 | 1,73E-18 | 8,78E-17 |
| ENSG00000177875.5 | CCDC184 | protein_coding | 230,646005 | 1,089900083 | 0,255570823 | 4,264571635 | 2,00E-05 | 0,000282546 |
| ENSG00000129084.18 | PSMA1 | protein_coding | 16632,64026 | 1,08552594 | 0,136427813 | 7,956778889 | 1,77E-15 | 7,74E-14 |
| ENSG00000144843.11 | ADPRH | protein_coding | 437,3556164 | 1,079844064 | 0,203851975 | 5,297196979 | 1,18E-07 | 2,47E-06 |
| ENSG00000132963.8 | POMP | protein_coding | 15352,74514 | 1,078812318 | 0,143866789 | 7,498689063 | 6,45E-14 | 2,56E-12 |
| ENSG00000075213.11 | SEMA3A | protein_coding | 7738,671178 | 1,072460105 | 0,118745338 | 9,031597528 | 1,69E-19 | 9,08E-18 |
| ENSG00000152894.15 | PTPRK | protein_coding | 5373,602808 | 1,071978351 | 0,119883973 | 8,941798649 | 3,83E-19 | 2,02E-17 |
| ENSG00000128016.7 | ZFP36 | protein_coding | 533,5214944 | 1,07038744 | 0,194663579 | 5,498652829 | 3,83E-08 | 8,68E-07 |
| ENSG00000163644.15 | PPM1K | protein_coding | 1999,075849 | 1,070303336 | 0,18262534 | 5,860650762 | 4,61E-09 | 1,17E-07 |
| ENSG00000102096.9 | PIM2 | protein_coding | 6936,981643 | 1,066436894 | 0,132133978 | 8,070875556 | 6,98E-16 | 3,14E-14 |
| ENSG00000101680.15 | LAMA1 | protein_coding | 441,7421692 | 1,066231479 | 0,289207416 | 3,686736306 | 0,000227149 | 0,00246775 |
| ENSG00000168065.16 | SLC22A11 | protein_coding | 277,7262451 | 1,063472694 | 0,241946662 | 4,395484048 | 1,11E-05 | 0,000166465 |
| ENSG00000171992.13 | SYNPO | protein_coding | 194,9072492 | 1,063258222 | 0,285324843 | 3,726483154 | 0,00019417 | 0,002150767 |
| ENSG00000140297.13 | GCNT3 | protein_coding | 1006,373969 | 1,06137872 | 0,154569641 | 6,866670002 | 6,57E-12 | 2,26E-10 |
| ENSG00000124226.11 | RNF114 | protein_coding | 12847,5265 | 1,057612327 | 0,137904758 | 7,669150379 | 1,73E-14 | 7,16E-13 |
| ENSG00000071575.11 | TRIB2 | protein_coding | 485,9429997 | 1,057061007 | 0,254831003 | 4,148086363 | 3,35E-05 | 0,000448199 |
| ENSG00000164362.21 | TERT | protein_coding | 458,175045 | 1,056374788 | 0,238476842 | 4,429674513 | 9,44E-06 | 0,00014411 |
| ENSG00000196353.11 | CPNE4 | protein_coding | 1447,637489 | 1,054476622 | 0,152084776 | 6,933479144 | 4,11E-12 | 1,44E-10 |
| ENSG00000180535.4 | BHLHA15 | protein_coding | 423,2460771 | 1,054006473 | 0,327647303 | 3,216893476 | 0,001295867 | 0,010860755 |
| ENSG00000067182.8 | TNFRSF1A | protein_coding | 1092,717734 | 1,05350035 | 0,178306424 | 5,908370128 | 3,46E-09 | 8,95E-08 |
| ENSG00000138002.16 | IFT172 | protein_coding | 599,1969211 | 1,052666409 | 0,245026318 | 4,296136092 | 1,74E-05 | 0,000248873 |
| ENSG00000108622.11 | ICAM2 | protein_coding | 172,9249956 | 1,050973583 | 0,288405912 | 3,644077805 | 0,000268352 | 0,002844534 |

| | | | | | | | | |
|---|---|---|---|---|---|---|---|---|
| ENSG00000176769.9 | TCERG1L | protein_coding | 193,1387736 | 1,049066542 | 0,268265295 | 3,910556314 | 9,21E-05 | 0,001113677 |
| ENSG00000049449.10 | RCN1 | protein_coding | 27734,38343 | 1,047009553 | 0,112974183 | 9,267688634 | 1,90E-20 | 1,06E-18 |
| ENSG00000103257.9 | SLC7A5 | protein_coding | 448,8342207 | 1,046213585 | 0,218362327 | 4,7911817 | 1,66E-06 | 2,93E-05 |
| ENSG00000145819.18 | ARHGAP26 | protein_coding | 1881,109177 | 1,044339222 | 0,174621 | 5,980604982 | 2,22E-09 | 5,93E-08 |
| ENSG00000102390.11 | PBDC1 | protein_coding | 1704,966904 | 1,037996038 | 0,17722093 | 5,857073655 | 4,71E-09 | 1,20E-07 |
| ENSG00000157617.17 | C2CD2 | protein_coding | 350,3692613 | 1,035901438 | 0,328415499 | 3,154240412 | 0,001609165 | 0,013067441 |
| ENSG00000113583.8 | C5orf15 | protein_coding | 16195,15216 | 1,033823555 | 0,120497269 | 8,579643051 | 9,52E-18 | 4,64E-16 |
| ENSG00000259522.3 | AL136295.4 | protein_coding | 242,285955 | 1,033708951 | 0,248133886 | 4,165932224 | 3,10E-05 | 0,000418542 |
| ENSG00000186479.5 | RGS7BP | protein_coding | 153,2733974 | 1,033657609 | 0,302171583 | 3,420763786 | 0,000624455 | 0,005833545 |
| ENSG00000148516.22 | ZEB1 | protein_coding | 177,6446606 | 1,033232098 | 0,28475572 | 3,628485843 | 0,000285088 | 0,002985655 |
| ENSG00000188042.8 | ARL4C | protein_coding | 4618,246237 | 1,0320884 | 0,127999115 | 8,063246352 | 7,43E-16 | 3,34E-14 |
| ENSG00000134285.11 | FKBP11 | protein_coding | 750,6948736 | 1,028917115 | 0,178687539 | 5,75819177 | 8,50E-09 | 2,09E-07 |
| ENSG00000002933.9 | TMEM176A | protein_coding | 535,4674827 | 1,02888995 | 0,187032845 | 5,501119083 | 3,77E-08 | 8,57E-07 |
| ENSG00000180817.12 | PPA1 | protein_coding | 13503,94676 | 1,02866288 | 0,142304204 | 7,228619039 | 4,88E-13 | 1,81E-11 |
| ENSG00000186594.14 | MIR22HG | lncRNA | 283,7390901 | 1,028532463 | 0,274599716 | 3,745570021 | 0,000179985 | 0,002013885 |
| ENSG00000164120.14 | HPGD | protein_coding | 20550,68901 | 1,0272763 | 0,136528189 | 7,524279861 | 5,30E-14 | 2,12E-12 |
| ENSG00000120675.6 | DNAJC15 | protein_coding | 5118,735491 | 1,027112637 | 0,126585852 | 8,113960783 | 4,90E-16 | 2,23E-14 |
| ENSG00000179104.9 | TMTC2 | protein_coding | 2934,353467 | 1,026899882 | 0,124974606 | 8,216868318 | 2,09E-16 | 9,67E-15 |
| ENSG00000085117.12 | CD82 | protein_coding | 12536,52389 | 1,026767567 | 0,112418385 | 9,133448808 | 6,64E-20 | 3,64E-18 |
| ENSG00000172057.10 | ORMDL3 | protein_coding | 7089,67689 | 1,026758503 | 0,125026349 | 8,212336936 | 2,17E-16 | 1,00E-14 |
| ENSG00000185561.10 | TLCD2 | protein_coding | 142,3124572 | 1,025663341 | 0,354816603 | 2,890685868 | 0,003844021 | 0,026815924 |
| ENSG00000112096.19 | SOD2 | protein_coding | 16998,32459 | 1,025038205 | 0,187032698 | 5,480529424 | 4,24E-08 | 9,54E-07 |
| ENSG00000101596.16 | SMCHD1 | protein_coding | 10916,08468 | 1,024414731 | 0,133132817 | 7,694682302 | 1,42E-14 | 5,92E-13 |
| ENSG00000141682.12 | PMAIP1 | protein_coding | 16189,87683 | 1,02438839 | 0,14495375 | 7,067001631 | 1,58E-12 | 5,69E-11 |
| ENSG00000275183.2 | LENG9 | protein_coding | 166,8917623 | 1,023408307 | 0,290881568 | 3,51829892 | 0,000434323 | 0,004299077 |
| ENSG00000117318.9 | ID3 | protein_coding | 2265,487972 | 1,022222598 | 0,147463467 | 6,932039632 | 4,15E-12 | 1,45E-10 |
| ENSG00000251573.2 | LINC02106 | lncRNA | 182,8674786 | 1,020418365 | 0,299450356 | 3,407637841 | 0,000655278 | 0,006056621 |
| ENSG00000149599.15 | DUSP15 | protein_coding | 1039,118288 | 1,019373257 | 0,163612072 | 6,230428146 | 4,65E-10 | 1,33E-08 |
| ENSG00000164543.7 | STK17A | protein_coding | 599,0830839 | 1,015661192 | 0,178401001 | 5,693136178 | 1,25E-08 | 3,02E-07 |
| ENSG00000233184.7 | AC093157.1 | lncRNA | 164,9121636 | 1,014876798 | 0,30562295 | 3,320682559 | 0,000897976 | 0,007918757 |
| ENSG00000100906.11 | NFKBIA | protein_coding | 4754,047541 | 1,014851788 | 0,122065623 | 8,313985268 | 9,25E-17 | 4,37E-15 |
| ENSG00000258867.6 | LINC01146 | lncRNA | 550,4290025 | 1,014718167 | 0,214833701 | 4,723272758 | 2,32E-06 | 3,98E-05 |
| ENSG00000026508.20 | CD44 | protein_coding | 13287,34409 | 1,014009271 | 0,138148902 | 7,339973434 | 2,14E-13 | 8,15E-12 |
| ENSG00000131943.18 | C19orf12 | protein_coding | 2964,629894 | 1,010715337 | 0,136784453 | 7,38910975 | 1,48E-13 | 5,70E-12 |
| ENSG00000235568.7 | NFAM1 | protein_coding | 202,9397376 | 1,008903768 | 0,314563247 | 3,207316102 | 0,001339797 | 0,011192597 |
| ENSG00000156345.18 | CDK20 | protein_coding | 835,8119595 | 1,002114913 | 0,191866646 | 5,222976127 | 1,76E-07 | 3,63E-06 |

| | | | | | | | | |
|---|---|---|---|---|---|---|---|---|
| ENSG00000143384.14 | MCL1 | protein_coding | 28231,14963 | 1,001786139 | 0,10846798 | 9,235777618 | 2,56E-20 | 1,42E-18 |
| ENSG00000169432.18 | SCN9A | protein_coding | 1515,639452 | 1,000221861 | 0,139150083 | 7,188079507 | 6,57E-13 | 2,42E-11 |
| ENSG00000111266.9 | DUSP16 | protein_coding | 1930,997998 | 0,994930992 | 0,143202499 | 6,947720882 | 3,71E-12 | 1,31E-10 |
| ENSG00000152495.11 | CAMK4 | protein_coding | 3432,218718 | 0,994723899 | 0,158215821 | 6,287132945 | 3,23E-10 | 9,35E-09 |
| ENSG00000213689.14 | TREX1 | protein_coding | 354,5652495 | 0,994533188 | 0,234172324 | 4,24701421 | 2,17E-05 | 0,000302556 |
| ENSG00000033178.13 | UBA6 | protein_coding | 7106,70601 | 0,986436015 | 0,128546192 | 7,673786391 | 1,67E-14 | 6,92E-13 |
| ENSG00000187672.14 | ERC2 | protein_coding | 259,3307384 | 0,986076327 | 0,260704998 | 3,782345318 | 0,000155358 | 0,001759963 |
| ENSG00000090975.13 | PITPNM2 | protein_coding | 454,0668797 | 0,985651651 | 0,197233684 | 4,997379909 | 5,81E-07 | 1,11E-05 |
| ENSG00000134352.20 | IL6ST | protein_coding | 6021,458034 | 0,984280611 | 0,143474956 | 6,860295598 | 6,87E-12 | 2,35E-10 |
| ENSG00000119508.18 | NR4A3 | protein_coding | 186,8425418 | 0,984187257 | 0,299413917 | 3,28704579 | 0,001012444 | 0,008769346 |
| ENSG00000103034.14 | NDRG4 | protein_coding | 2947,762055 | 0,983557311 | 0,131115706 | 7,501445425 | 6,31E-14 | 2,51E-12 |
| ENSG00000215068.8 | AC025171.2 | lncRNA | 97,44456689 | 0,982673502 | 0,367872426 | 2,67123446 | 0,007557284 | 0,045845187 |
| ENSG00000169228.14 | RAB24 | protein_coding | 2629,592838 | 0,982410341 | 0,150247119 | 6,538630149 | 6,21E-11 | 1,91E-09 |
| ENSG00000125354.24 | SEPTIN6 | protein_coding | 1975,606699 | 0,982267103 | 0,158624147 | 6,192418502 | 5,92E-10 | 1,67E-08 |
| ENSG00000163545.11 | NUAK2 | protein_coding | 370,6248851 | 0,982177871 | 0,282276786 | 3,479485099 | 0,000502378 | 0,004871371 |
| ENSG00000184905.9 | TCEAL2 | protein_coding | 3528,025126 | 0,981680907 | 0,145103796 | 6,765370287 | 1,33E-11 | 4,40E-10 |
| ENSG00000004799.8 | PDK4 | protein_coding | 235,3822694 | 0,981665933 | 0,306645463 | 3,201305913 | 0,001368062 | 0,011387825 |
| ENSG00000064042.18 | LIMCH1 | protein_coding | 1922,447675 | 0,981106469 | 0,208937991 | 4,695682518 | 2,66E-06 | 4,49E-05 |
| ENSG00000139679.15 | LPAR6 | protein_coding | 325,8430847 | 0,980621916 | 0,249930187 | 3,923583326 | 8,72E-05 | 0,001060013 |
| ENSG00000157992.12 | KRTCAP3 | protein_coding | 353,3790323 | 0,980003369 | 0,227334451 | 4,310844067 | 1,63E-05 | 0,000233519 |
| ENSG00000253669.4 | GASAL1 | lncRNA | 216,3943672 | 0,976676651 | 0,276545247 | 3,531706517 | 0,000412887 | 0,004118054 |
| ENSG00000000457.14 | SCYL3 | protein_coding | 1755,265114 | 0,975144076 | 0,149311518 | 6,530936731 | 6,54E-11 | 2,00E-09 |
| ENSG00000184226.15 | PCDH9 | protein_coding | 1859,283996 | 0,974589731 | 0,143785647 | 6,778073812 | 1,22E-11 | 4,07E-10 |
| ENSG00000124788.19 | ATXN1 | protein_coding | 202,8874548 | 0,972306061 | 0,283214974 | 3,433102591 | 0,000596716 | 0,005605757 |
| ENSG00000005884.18 | ITGA3 | protein_coding | 3570,740613 | 0,971173848 | 0,130833138 | 7,42299593 | 1,15E-13 | 4,47E-12 |
| ENSG00000166387.13 | PPFIBP2 | protein_coding | 525,7492536 | 0,967907939 | 0,183504098 | 5,274584877 | 1,33E-07 | 2,78E-06 |
| ENSG00000233593.10 | LINC02609 | lncRNA | 365,1671554 | 0,965525598 | 0,222001851 | 4,349178136 | 1,37E-05 | 0,000200895 |
| ENSG00000137842.7 | TMEM62 | protein_coding | 6966,771824 | 0,964629139 | 0,121632205 | 7,930705016 | 2,18E-15 | 9,48E-14 |
| ENSG00000148655.15 | LRMDA | protein_coding | 332,4200534 | 0,961321072 | 0,349383493 | 2,751478226 | 0,005932696 | 0,037808523 |
| ENSG00000148926.10 | ADM | protein_coding | 367,5402855 | 0,959941709 | 0,225622785 | 4,254631059 | 2,09E-05 | 0,000293811 |
| ENSG00000135315.12 | CEP162 | protein_coding | 529,7000955 | 0,958238943 | 0,220773372 | 4,34037372 | 1,42E-05 | 0,000208093 |
| ENSG00000226067.7 | LINC00623 | lncRNA | 2642,122406 | 0,955445401 | 0,143329734 | 6,666065553 | 2,63E-11 | 8,44E-10 |
| ENSG00000051108.15 | HERPUD1 | protein_coding | 9155,573105 | 0,955324764 | 0,128827022 | 7,415561992 | 1,21E-13 | 4,72E-12 |
| ENSG00000174485.16 | DENND4A | protein_coding | 5651,239713 | 0,954887606 | 0,138077112 | 6,915611106 | 4,66E-12 | 1,62E-10 |
| ENSG00000105856.14 | HBP1 | protein_coding | 4698,500931 | 0,954504063 | 0,12171735 | 7,8419721 | 4,44E-15 | 1,90E-13 |
| ENSG00000131069.20 | ACSS2 | protein_coding | 4708,377209 | 0,954231559 | 0,118209088 | 8,072404389 | 6,89E-16 | 3,11E-14 |

| | | | | | | | | |
|---|---|---|---|---|---|---|---|---|
| ENSG00000110925.7 | CSRNP2 | protein_coding | 4658,667339 | 0,95420687 | 0,127987372 | 7,455476683 | 8,95E-14 | 3,53E-12 |
| ENSG00000129003.19 | VPS13C | protein_coding | 10225,50245 | 0,952324658 | 0,120965554 | 7,872692898 | 3,47E-15 | 1,49E-13 |
| ENSG00000164078.13 | MST1R | protein_coding | 198,8979628 | 0,951457199 | 0,276702556 | 3,43855587 | 0,000584826 | 0,005519986 |
| ENSG00000168143.9 | FAM83B | protein_coding | 588,7533445 | 0,950269389 | 0,234754618 | 4,047926293 | 5,17E-05 | 0,000660026 |
| ENSG00000266947.1 | AC022916.1 | lncRNA | 456,4493066 | 0,948366176 | 0,199069002 | 4,76400729 | 1,90E-06 | 3,33E-05 |
| ENSG00000115904.14 | SOS1 | protein_coding | 7681,211027 | 0,948152904 | 0,131694071 | 7,199662811 | 6,04E-13 | 2,23E-11 |
| ENSG00000172292.15 | CERS6 | protein_coding | 53995,3275 | 0,947617987 | 0,108676498 | 8,719622012 | 2,79E-18 | 1,41E-16 |
| ENSG00000111846.19 | GCNT2 | protein_coding | 1904,502729 | 0,946164174 | 0,13597069 | 6,958589172 | 3,44E-12 | 1,22E-10 |
| ENSG00000126561.17 | STAT5A | protein_coding | 688,1109111 | 0,944847709 | 0,171690889 | 5,503190742 | 3,73E-08 | 8,49E-07 |
| ENSG00000204381.12 | LAYN | protein_coding | 204,6672616 | 0,944377724 | 0,277172509 | 3,407183949 | 0,000656369 | 0,006064025 |
| ENSG00000145451.13 | GLRA3 | protein_coding | 447,5995735 | 0,943383801 | 0,196664708 | 4,79691455 | 1,61E-06 | 2,86E-05 |
| ENSG00000105202.9 | FBL | protein_coding | 16651,97835 | 0,942883249 | 0,125429422 | 7,517241464 | 5,59E-14 | 2,23E-12 |
| ENSG00000182791.5 | CCDC87 | protein_coding | 158,7434961 | 0,941880083 | 0,313188225 | 3,007393019 | 0,002634989 | 0,019526897 |
| ENSG00000143870.13 | PDIA6 | protein_coding | 101762,645 | 0,939928018 | 0,131654265 | 7,139366244 | 9,38E-13 | 3,41E-11 |
| ENSG00000088298.13 | EDEM2 | protein_coding | 3611,23181 | 0,938605038 | 0,161986817 | 5,794329798 | 6,86E-09 | 1,71E-07 |
| ENSG00000175262.14 | C1orf127 | protein_coding | 310,2863035 | 0,938098822 | 0,30272335 | 3,098865098 | 0,001942635 | 0,015224148 |
| ENSG00000171643.14 | S100Z | protein_coding | 733,3905745 | 0,93749679 | 0,177890961 | 5,270064216 | 1,36E-07 | 2,84E-06 |
| ENSG00000148834.13 | GSTO1 | protein_coding | 7369,652888 | 0,937148534 | 0,157082545 | 5,965962259 | 2,43E-09 | 6,47E-08 |
| ENSG00000053108.17 | FSTL4 | protein_coding | 326,6851661 | 0,936845798 | 0,298283953 | 3,140785104 | 0,001684956 | 0,01358281 |
| ENSG00000092096.18 | SLC22A17 | protein_coding | 20780,03138 | 0,936837807 | 0,111780312 | 8,381062698 | 5,25E-17 | 2,49E-15 |
| ENSG00000169297.8 | NR0B1 | protein_coding | 3729,228257 | 0,935048438 | 0,1271302 | 7,355045782 | 1,91E-13 | 7,29E-12 |
| ENSG00000156920.11 | ADGRG4 | protein_coding | 191,1446728 | 0,93019271 | 0,280315651 | 3,318375937 | 0,000905425 | 0,007971017 |
| ENSG00000260000.2 | AL133338.1 | lncRNA | 882,0654327 | 0,92814066 | 0,166308874 | 5,58082464 | 2,39E-08 | 5,56E-07 |
| ENSG00000115295.20 | CLIP4 | protein_coding | 1786,343486 | 0,927457671 | 0,159728977 | 5,806445935 | 6,38E-09 | 1,60E-07 |
| ENSG00000185215.11 | TNFAIP2 | protein_coding | 389,3450953 | 0,927331398 | 0,309928503 | 2,992081682 | 0,002770821 | 0,02037483 |
| ENSG00000165805.10 | C12orf50 | protein_coding | 294,2018173 | 0,926616444 | 0,264425826 | 3,504258486 | 0,000457881 | 0,004500344 |
| ENSG00000163900.11 | TMEM41A | protein_coding | 3291,774296 | 0,925512328 | 0,149111445 | 6,206849714 | 5,41E-10 | 1,53E-08 |
| ENSG00000142920.17 | AZIN2 | protein_coding | 365,7367739 | 0,925446755 | 0,226382482 | 4,087978654 | 4,35E-05 | 0,000566532 |
| ENSG00000166206.15 | GABRB3 | protein_coding | 26868,99169 | 0,923454245 | 0,114391213 | 8,072772538 | 6,87E-16 | 3,11E-14 |
| ENSG00000026297.16 | RNASET2 | protein_coding | 6092,448719 | 0,921522401 | 0,150508398 | 6,122730782 | 9,20E-10 | 2,55E-08 |
| ENSG00000102804.15 | TSC22D1 | protein_coding | 14449,50814 | 0,919372008 | 0,132946932 | 6,915330779 | 4,67E-12 | 1,62E-10 |
| ENSG00000153707.17 | PTPRD | protein_coding | 1324,78572 | 0,914791691 | 0,251579305 | 3,636196113 | 0,000276694 | 0,002910846 |
| ENSG00000158985.14 | CDC42SE2 | protein_coding | 16883,06342 | 0,914331115 | 0,117107161 | 7,807644804 | 5,83E-15 | 2,48E-13 |
| ENSG00000184809.13 | B3GALT5-AS1 | lncRNA | 166,0592454 | 0,913862949 | 0,287055581 | 3,18357492 | 0,001454685 | 0,011970842 |
| ENSG00000172602.11 | RND1 | protein_coding | 281,8966144 | 0,913213001 | 0,25193976 | 3,624727601 | 0,000289266 | 0,003023359 |
| ENSG00000253958.2 | CLDN23 | protein_coding | 732,6295722 | 0,913093175 | 0,171201874 | 5,333429781 | 9,64E-08 | 2,07E-06 |

| | | | | | | | | |
|---|---|---|---|---|---|---|---|---|
| ENSG00000164442.10 | CITED2 | protein_coding | 2990,792289 | 0,911932517 | 0,140320492 | 6,498926143 | 8,09E-11 | 2,46E-09 |
| ENSG00000197956.10 | S100A6 | protein_coding | 11640,44119 | 0,911242073 | 0,149006154 | 6,115466038 | 9,63E-10 | 2,66E-08 |
| ENSG00000280165.1 | PCDH20 | protein_coding | 990,0017478 | 0,909183614 | 0,154126281 | 5,898952502 | 3,66E-09 | 9,44E-08 |
| ENSG00000100139.13 | MICALL1 | protein_coding | 675,8471824 | 0,909096489 | 0,191671335 | 4,742996595 | 2,11E-06 | 3,66E-05 |
| ENSG00000255639.3 | AC005833.1 | protein_coding | 2278,047083 | 0,906716165 | 0,128562086 | 7,052749313 | 1,75E-12 | 6,29E-11 |
| ENSG00000008311.15 | AASS | protein_coding | 802,2550514 | 0,90656833 | 0,176469126 | 5,137263105 | 2,79E-07 | 5,61E-06 |
| ENSG00000171564.12 | FGB | protein_coding | 1307,806431 | 0,901523002 | 0,144384778 | 6,243892297 | 4,27E-10 | 1,22E-08 |
| ENSG00000248905.10 | FMN1 | protein_coding | 1065,034914 | 0,897340878 | 0,187391481 | 4,788589499 | 1,68E-06 | 2,97E-05 |
| ENSG00000184164.15 | CRELD2 | protein_coding | 5600,604497 | 0,895804306 | 0,155612756 | 5,756625155 | 8,58E-09 | 2,10E-07 |
| ENSG00000023171.18 | GRAMD1B | protein_coding | 3903,265715 | 0,894702566 | 0,149190233 | 5,997058577 | 2,01E-09 | 5,37E-08 |
| ENSG00000136040.9 | PLXNC1 | protein_coding | 28617,83394 | 0,894257879 | 0,123401672 | 7,246724162 | 4,27E-13 | 1,60E-11 |
| ENSG00000137947.12 | GTF2B | protein_coding | 2695,861924 | 0,894197956 | 0,153757762 | 5,815628073 | 6,04E-09 | 1,52E-07 |
| ENSG00000159761.15 | C16orf86 | protein_coding | 282,2352251 | 0,892120142 | 0,232589231 | 3,835603812 | 0,000125256 | 0,001460464 |
| ENSG00000122643.22 | NT5C3A | protein_coding | 5606,270198 | 0,891830625 | 0,129061578 | 6,910117181 | 4,84E-12 | 1,68E-10 |
| ENSG00000170542.6 | SERPINB9 | protein_coding | 1092,749861 | 0,89147314 | 0,166560681 | 5,352242416 | 8,69E-08 | 1,88E-06 |
| ENSG00000100567.13 | PSMA3 | protein_coding | 9576,130653 | 0,891388088 | 0,141524915 | 6,298453453 | 3,01E-10 | 8,72E-09 |
| ENSG00000120251.21 | GRIA2 | protein_coding | 372,2282911 | 0,891119857 | 0,209066188 | 4,262381528 | 2,02E-05 | 0,000285139 |
| ENSG00000164300.17 | SERINC5 | protein_coding | 4804,544097 | 0,890772292 | 0,153945497 | 5,786283512 | 7,20E-09 | 1,78E-07 |
| ENSG00000121716.20 | PILRB | protein_coding | 438,8725225 | 0,890656361 | 0,32789733 | 2,716265977 | 0,006602285 | 0,041173322 |
| ENSG00000119537.18 | KDSR | protein_coding | 3985,616443 | 0,890536754 | 0,127615264 | 6,978293395 | 2,99E-12 | 1,06E-10 |
| ENSG00000024048.11 | UBR2 | protein_coding | 6339,462605 | 0,88961144 | 0,135496658 | 6,565560002 | 5,18E-11 | 1,61E-09 |
| ENSG00000140092.14 | FBLN5 | protein_coding | 576,8332582 | 0,888379279 | 0,213045498 | 4,169904029 | 3,05E-05 | 0,000412068 |
| ENSG00000250903.9 | GMDS-DT | lncRNA | 153,6320521 | 0,888326095 | 0,30536549 | 2,909058568 | 0,003625189 | 0,025484668 |
| ENSG00000160014.17 | CALM3 | protein_coding | 43581,75804 | 0,887070184 | 0,122075851 | 7,266549233 | 3,69E-13 | 1,38E-11 |
| ENSG00000034677.13 | RNF19A | protein_coding | 5048,272141 | 0,885623565 | 0,12005567 | 7,376774171 | 1,62E-13 | 6,24E-12 |
| ENSG00000205560.13 | CPT1B | protein_coding | 249,6333101 | 0,884865559 | 0,260429426 | 3,397717265 | 0,000679506 | 0,006241974 |
| ENSG00000149218.5 | ENDOD1 | protein_coding | 1653,874643 | 0,88440701 | 0,150397197 | 5,880475346 | 4,09E-09 | 1,05E-07 |
| ENSG00000014914.21 | MTMR11 | protein_coding | 404,1213405 | 0,883365458 | 0,217735218 | 4,05706283 | 4,97E-05 | 0,000637142 |
| ENSG00000132196.15 | HSD17B7 | protein_coding | 8187,184029 | 0,882632669 | 0,131027696 | 6,736229793 | 1,63E-11 | 5,36E-10 |
| ENSG00000135549.15 | PKIB | protein_coding | 8789,656366 | 0,881185039 | 0,136996295 | 6,432181543 | 1,26E-10 | 3,76E-09 |
| ENSG00000155660.11 | PDIA4 | protein_coding | 27379,06366 | 0,879735815 | 0,167127427 | 5,263862609 | 1,41E-07 | 2,93E-06 |
| ENSG00000198879.12 | SFMBT2 | protein_coding | 2092,785988 | 0,87968083 | 0,171994012 | 5,114601499 | 3,14E-07 | 6,27E-06 |
| ENSG00000184675.11 | AMER1 | protein_coding | 393,2270235 | 0,878247435 | 0,253407718 | 3,465748571 | 0,000528758 | 0,005063833 |
| ENSG00000067798.16 | NAV3 | protein_coding | 198,6090115 | 0,878189891 | 0,290763175 | 3,020292686 | 0,002525305 | 0,018860958 |
| ENSG00000171843.17 | MLLT3 | protein_coding | 2028,258727 | 0,873129856 | 0,154287233 | 5,659119278 | 1,52E-08 | 3,63E-07 |
| ENSG00000073910.22 | FRY | protein_coding | 2079,970602 | 0,869658148 | 0,231552734 | 3,755767136 | 0,000172811 | 0,001946103 |

| | | | | | | | | |
|---|---|---|---|---|---|---|---|---|
| ENSG00000102699.6 | PARP4 | protein_coding | 9112,69255 | 0,868828878 | 0,116315622 | 7,469580316 | 8,05E-14 | 3,18E-12 |
| ENSG00000118503.15 | TNFAIP3 | protein_coding | 1640,815201 | 0,868483157 | 0,14770322 | 5,879920287 | 4,10E-09 | 1,05E-07 |
| ENSG00000223804.5 | AC244669.1 | transcribed_unproc | 322,2881944 | 0,866953028 | 0,222065584 | 3,904040465 | 9,46E-05 | 0,00114147 |
| ENSG00000233024.7 | NPIPA9 | protein_coding | 461,8785525 | 0,865120995 | 0,236018535 | 3,665479043 | 0,000246876 | 0,002650386 |
| ENSG00000138756.18 | BMP2K | protein_coding | 6313,732325 | 0,864681179 | 0,149622005 | 5,779104356 | 7,51E-09 | 1,86E-07 |
| ENSG00000183844.17 | FAM3B | protein_coding | 2220,865919 | 0,862354787 | 0,167948395 | 5,13464143 | 2,83E-07 | 5,68E-06 |
| ENSG00000110675.13 | ELMOD1 | protein_coding | 152,9977388 | 0,858146206 | 0,304772711 | 2,815692399 | 0,004867224 | 0,032255569 |
| ENSG00000145050.19 | MANF | protein_coding | 20187,40752 | 0,857807278 | 0,12565078 | 6,826915664 | 8,68E-12 | 2,94E-10 |
| ENSG00000106565.18 | TMEM176B | protein_coding | 1995,22948 | 0,857680537 | 0,158215023 | 5,420980396 | 5,93E-08 | 1,31E-06 |
| ENSG00000131389.17 | SLC6A6 | protein_coding | 16462,7317 | 0,857243231 | 0,148509282 | 5,7723209 | 7,82E-09 | 1,93E-07 |
| ENSG00000041357.16 | PSMA4 | protein_coding | 21498,48865 | 0,856708638 | 0,128452729 | 6,669446781 | 2,57E-11 | 8,30E-10 |
| ENSG00000196497.17 | IPO4 | protein_coding | 2763,961097 | 0,855367626 | 0,124770272 | 6,855540308 | 7,10E-12 | 2,43E-10 |
| ENSG00000151718.16 | WWC2 | protein_coding | 258,4191859 | 0,851722669 | 0,270996537 | 3,14292824 | 0,001672668 | 0,013509763 |
| ENSG00000129472.15 | RAB2B | protein_coding | 1386,721357 | 0,851042394 | 0,168352513 | 5,055121405 | 4,30E-07 | 8,38E-06 |
| ENSG00000270800.3 | RPS10-NUDT3 | protein_coding | 1925,70103 | 0,850872256 | 0,22057433 | 3,857530727 | 0,000114538 | 0,001345264 |
| ENSG00000146426.18 | TIAM2 | protein_coding | 346,3460088 | 0,85072062 | 0,252230259 | 3,372793668 | 0,000744097 | 0,0067465 |
| ENSG00000105976.16 | MET | protein_coding | 8090,025039 | 0,849151527 | 0,133464643 | 6,362370667 | 1,99E-10 | 5,85E-09 |
| ENSG00000246095.2 | LINC01096 | lncRNA | 283,4950661 | 0,847530691 | 0,242130959 | 3,500298735 | 0,000464737 | 0,004544264 |
| ENSG00000120690.16 | ELF1 | protein_coding | 4996,318946 | 0,846755932 | 0,117011523 | 7,236517469 | 4,60E-13 | 1,72E-11 |
| ENSG00000134970.14 | TMED7 | protein_coding | 21739,43293 | 0,84659535 | 0,126064568 | 6,715569355 | 1,87E-11 | 6,16E-10 |
| ENSG00000134769.21 | DTNA | protein_coding | 1854,829982 | 0,845047905 | 0,194319694 | 4,348750705 | 1,37E-05 | 0,000201004 |
| ENSG00000177738.4 | AC025171.1 | lncRNA | 970,7366853 | 0,8427242 | 0,154488494 | 5,454931822 | 4,90E-08 | 1,09E-06 |
| ENSG00000140280.14 | LYSMD2 | protein_coding | 3799,771075 | 0,842389781 | 0,141746594 | 5,942927831 | 2,80E-09 | 7,37E-08 |
| ENSG00000260231.2 | KDM7A-DT | lncRNA | 3975,801875 | 0,841965031 | 0,163186894 | 5,159513784 | 2,48E-07 | 5,00E-06 |
| ENSG00000139668.9 | WDFY2 | protein_coding | 1758,363895 | 0,837724292 | 0,143699179 | 5,82970828 | 5,55E-09 | 1,40E-07 |
| ENSG00000155897.10 | ADCY8 | protein_coding | 14461,64731 | 0,83694029 | 0,124684421 | 6,712468806 | 1,91E-11 | 6,27E-10 |
| ENSG00000091409.15 | ITGA6 | protein_coding | 2140,412418 | 0,836620315 | 0,141961047 | 5,893309005 | 3,79E-09 | 9,76E-08 |
| ENSG00000107798.18 | LIPA | protein_coding | 12191,29378 | 0,834968618 | 0,125787467 | 6,637931744 | 3,18E-11 | 1,01E-09 |
| ENSG00000146278.11 | PNRC1 | protein_coding | 2446,176692 | 0,833908241 | 0,127831179 | 6,523512107 | 6,87E-11 | 2,10E-09 |
| ENSG00000146409.12 | SLC18B1 | protein_coding | 16147,50181 | 0,833558129 | 0,121666133 | 6,851192742 | 7,32E-12 | 2,50E-10 |
| ENSG00000166974.13 | MAPRE2 | protein_coding | 11066,08852 | 0,832518092 | 0,123609468 | 6,735067364 | 1,64E-11 | 5,39E-10 |
| ENSG00000179546.4 | HTR1D | protein_coding | 2319,822186 | 0,832365235 | 0,159997243 | 5,202372353 | 1,97E-07 | 4,03E-06 |
| ENSG00000131844.16 | MCCC2 | protein_coding | 5880,32256 | 0,832313993 | 0,116763753 | 7,128188057 | 1,02E-12 | 3,68E-11 |
| ENSG00000138166.6 | DUSP5 | protein_coding | 2349,26946 | 0,830929948 | 0,205127499 | 4,050797443 | 5,10E-05 | 0,000653174 |
| ENSG00000182718.18 | ANXA2 | protein_coding | 773,4330504 | 0,830897271 | 0,203842669 | 4,076169502 | 4,58E-05 | 0,00059129 |
| ENSG00000170345.10 | FOS | protein_coding | 272,9966571 | 0,830405842 | 0,241137001 | 3,443709749 | 0,000573791 | 0,005440363 |

| | | | | | | | | |
|---|---|---|---|---|---|---|---|---|
| ENSG00000175895.4 | PLEKHF2 | protein_coding | 6504,36896 | 0,830112887 | 0,138412004 | 5,997405304 | 2,00E-09 | 5,37E-08 |
| ENSG00000197822.11 | OCLN | protein_coding | 3262,914487 | 0,829948861 | 0,150356101 | 5,519888153 | 3,39E-08 | 7,74E-07 |
| ENSG00000136141.15 | LRCH1 | protein_coding | 1784,808417 | 0,828870597 | 0,144374122 | 5,741129961 | 9,40E-09 | 2,29E-07 |
| ENSG00000149428.19 | HYOU1 | protein_coding | 41896,91534 | 0,827693878 | 0,156696704 | 5,28213967 | 1,28E-07 | 2,68E-06 |
| ENSG00000181104.7 | F2R | protein_coding | 173,3299158 | 0,825635342 | 0,304858533 | 2,708257279 | 0,006763757 | 0,041992659 |
| ENSG00000100418.8 | DESI1 | protein_coding | 2733,389814 | 0,82558765 | 0,135831556 | 6,078025402 | 1,22E-09 | 3,33E-08 |
| ENSG00000135052.16 | GOLM1 | protein_coding | 47342,83871 | 0,825555789 | 0,107933492 | 7,648745268 | 2,03E-14 | 8,36E-13 |
| ENSG00000184678.10 | H2BC21 | protein_coding | 2180,261995 | 0,825113918 | 0,150644375 | 5,477230167 | 4,32E-08 | 9,68E-07 |
| ENSG00000175745.14 | NR2F1 | protein_coding | 1418,875307 | 0,824631337 | 0,157943496 | 5,221052831 | 1,78E-07 | 3,66E-06 |
| ENSG00000163947.12 | ARHGEF3 | protein_coding | 1599,343041 | 0,823227095 | 0,140913409 | 5,842077779 | 5,16E-09 | 1,30E-07 |
| ENSG00000198353.8 | HOXC4 | protein_coding | 324,6084266 | 0,820523849 | 0,23351416 | 3,513807688 | 0,000441732 | 0,00436829 |
| ENSG00000185522.9 | LMNTD2 | protein_coding | 159,2313956 | 0,81935619 | 0,300813234 | 2,723803669 | 0,006453486 | 0,040441456 |
| ENSG00000135047.16 | CTSL | protein_coding | 14732,32787 | 0,818788819 | 0,122889753 | 6,662791645 | 2,69E-11 | 8,61E-10 |
| ENSG00000134070.5 | IRAK2 | protein_coding | 424,0541283 | 0,817506571 | 0,209757644 | 3,897386318 | 9,72E-05 | 0,001166559 |
| ENSG00000183486.13 | MX2 | protein_coding | 1558,461694 | 0,816889139 | 0,152212164 | 5,366779604 | 8,02E-08 | 1,74E-06 |
| ENSG00000102393.14 | GLA | protein_coding | 3061,632522 | 0,816613842 | 0,141003127 | 5,791459074 | 6,98E-09 | 1,74E-07 |
| ENSG00000078114.19 | NEBL | protein_coding | 9049,875953 | 0,816538883 | 0,123515712 | 6,610809852 | 3,82E-11 | 1,21E-09 |
| ENSG00000175104.15 | TRAF6 | protein_coding | 2644,85252 | 0,816498936 | 0,141736102 | 5,760698411 | 8,38E-09 | 2,06E-07 |
| ENSG00000147488.11 | ST18 | protein_coding | 7004,971706 | 0,81313276 | 0,119294307 | 6,816190818 | 9,35E-12 | 3,16E-10 |
| ENSG00000091664.9 | SLC17A6 | protein_coding | 3578,603892 | 0,812563075 | 0,122490834 | 6,633664305 | 3,27E-11 | 1,04E-09 |
| ENSG00000137944.18 | KYAT3 | protein_coding | 4446,483176 | 0,811154856 | 0,12583724 | 6,446063607 | 1,15E-10 | 3,43E-09 |
| ENSG00000181192.12 | DHTKD1 | protein_coding | 2579,193272 | 0,811141938 | 0,133760288 | 6,064146161 | 1,33E-09 | 3,61E-08 |
| ENSG00000135596.18 | MICAL1 | protein_coding | 647,4655976 | 0,810452711 | 0,174347543 | 4,648489436 | 3,34E-06 | 5,59E-05 |
| ENSG00000127124.16 | HIVEP3 | protein_coding | 795,6934644 | 0,809978254 | 0,162364575 | 4,988639027 | 6,08E-07 | 1,16E-05 |
| ENSG00000128228.5 | SDF2L1 | protein_coding | 8298,973327 | 0,809954991 | 0,133938148 | 6,047231503 | 1,47E-09 | 3,99E-08 |
| ENSG00000105281.12 | SLC1A5 | protein_coding | 767,0825516 | 0,809691749 | 0,221309214 | 3,658644539 | 0,000253553 | 0,002708298 |
| ENSG00000186088.16 | GSAP | protein_coding | 621,388407 | 0,80785096 | 0,173473196 | 4,656920956 | 3,21E-06 | 5,38E-05 |
| ENSG00000203778.8 | FAM229B | protein_coding | 1256,388259 | 0,807620798 | 0,158150304 | 5,106666113 | 3,28E-07 | 6,52E-06 |
| ENSG00000112877.8 | CEP72 | protein_coding | 446,2339022 | 0,806945238 | 0,20198565 | 3,995062207 | 6,47E-05 | 0,000807887 |
| ENSG00000158773.14 | USF1 | protein_coding | 1870,582503 | 0,806617813 | 0,155744119 | 5,17912214 | 2,23E-07 | 4,54E-06 |
| ENSG00000152779.14 | SLC16A12 | protein_coding | 9729,424769 | 0,805849145 | 0,137789449 | 5,848409648 | 4,96E-09 | 1,26E-07 |
| ENSG00000151131.11 | C12orf45 | protein_coding | 464,1296304 | 0,804921775 | 0,232548952 | 3,461300378 | 0,000537573 | 0,005138851 |
| ENSG00000251350.2 | LINC02475 | lncRNA | 540,2002808 | 0,804261775 | 0,210667496 | 3,817683279 | 0,000134711 | 0,001553812 |
| ENSG00000175550.8 | DRAP1 | protein_coding | 7175,683708 | 0,803190136 | 0,134673426 | 5,963983822 | 2,46E-09 | 6,54E-08 |
| ENSG00000136237.19 | RAPGEF5 | protein_coding | 2232,790395 | 0,800753161 | 0,168434567 | 4,754090404 | 1,99E-06 | 3,47E-05 |
| ENSG00000177606.8 | JUN | protein_coding | 3391,016516 | 0,798935505 | 0,156560032 | 5,103061717 | 3,34E-07 | 6,64E-06 |

| | | | | | | | | |
|---|---|---|---|---|---|---|---|---|
| ENSG00000257702.3 | LBX2-AS1 | lncRNA | 668,0430905 | 0,798359035 | 0,179410185 | 4,449909221 | 8,59E-06 | 0,000132434 |
| ENSG00000169660.17 | HEXD | protein_coding | 1347,069313 | 0,797705969 | 0,153844551 | 5,185142811 | 2,16E-07 | 4,40E-06 |
| ENSG00000044574.8 | HSPA5 | protein_coding | 136342,229 | 0,7962309 | 0,108586987 | 7,33265491 | 2,26E-13 | 8,59E-12 |
| ENSG00000185015.8 | CA13 | protein_coding | 283,6091267 | 0,795655002 | 0,257691893 | 3,087621389 | 0,002017653 | 0,015702445 |
| ENSG00000050344.9 | NFE2L3 | protein_coding | 2984,810121 | 0,795159198 | 0,147699933 | 5,383612443 | 7,30E-08 | 1,60E-06 |
| ENSG00000139410.15 | SDSL | protein_coding | 1371,772482 | 0,795013189 | 0,148383174 | 5,357839219 | 8,42E-08 | 1,82E-06 |
| ENSG00000287721.1 | AC061975.8 | lncRNA | 260,2452053 | 0,794454789 | 0,256254933 | 3,100251693 | 0,001933563 | 0,015172089 |
| ENSG00000166794.5 | PPIB | protein_coding | 64177,87253 | 0,793759505 | 0,133920784 | 5,927082286 | 3,08E-09 | 8,08E-08 |
| ENSG00000095739.11 | BAMBI | protein_coding | 1303,136407 | 0,792296211 | 0,146839461 | 5,395662754 | 6,83E-08 | 1,50E-06 |
| ENSG00000122859.5 | NEUROG3 | protein_coding | 258,2956569 | 0,791501999 | 0,262561397 | 3,014540632 | 0,002573687 | 0,019188086 |
| ENSG00000151914.20 | DST | protein_coding | 8192,689732 | 0,791192836 | 0,157636287 | 5,01910347 | 5,19E-07 | 9,98E-06 |
| ENSG00000145391.13 | SETD7 | protein_coding | 2776,436064 | 0,791022578 | 0,20177552 | 3,920309949 | 8,84E-05 | 0,001073892 |
| ENSG00000286540.1 | AL096854.1 | lncRNA | 277,7652624 | 0,79046473 | 0,242784141 | 3,255833459 | 0,0011306 | 0,009656924 |
| ENSG00000111057.11 | KRT18 | protein_coding | 13677,252 | 0,790318507 | 0,13933548 | 5,67205502 | 1,41E-08 | 3,38E-07 |
| ENSG00000168301.13 | KCTD6 | protein_coding | 1091,496224 | 0,789564129 | 0,160825399 | 4,909449218 | 9,13E-07 | 1,69E-05 |
| ENSG00000164823.11 | OSGIN2 | protein_coding | 4841,526883 | 0,789503832 | 0,135379839 | 5,831768124 | 5,48E-09 | 1,38E-07 |
| ENSG00000139180.11 | NDUFA9 | protein_coding | 10080,09255 | 0,788872198 | 0,133506003 | 5,908889364 | 3,44E-09 | 8,93E-08 |
| ENSG00000149115.14 | TNKS1BP1 | protein_coding | 519,0294441 | 0,788686483 | 0,276909442 | 2,848174761 | 0,004397077 | 0,029752039 |
| ENSG00000151322.19 | NPAS3 | protein_coding | 2756,717062 | 0,785778308 | 0,1375917 | 5,710942669 | 1,12E-08 | 2,73E-07 |
| ENSG00000185753.13 | CXorf38 | protein_coding | 3497,833413 | 0,785332898 | 0,130024306 | 6,039893007 | 1,54E-09 | 4,16E-08 |
| ENSG00000185158.12 | LRRC37B | protein_coding | 539,222538 | 0,784820654 | 0,24427334 | 3,212878876 | 0,001314117 | 0,011004415 |
| ENSG00000101096.20 | NFATC2 | protein_coding | 304,7128334 | 0,784772813 | 0,281112132 | 2,791671809 | 0,005243651 | 0,034251431 |
| ENSG00000139263.12 | LRIG3 | protein_coding | 1261,337732 | 0,783655705 | 0,154225018 | 5,081248921 | 3,75E-07 | 7,38E-06 |
| ENSG00000197324.9 | LRP10 | protein_coding | 9860,289498 | 0,780876378 | 0,126938268 | 6,151623082 | 7,67E-10 | 2,14E-08 |
| ENSG00000183891.6 | TTC32 | protein_coding | 1220,654074 | 0,779293595 | 0,160123298 | 4,866834542 | 1,13E-06 | 2,06E-05 |
| ENSG00000074842.8 | MYDGF | protein_coding | 22706,82576 | 0,777678253 | 0,150343541 | 5,172674862 | 2,31E-07 | 4,69E-06 |
| ENSG00000166352.17 | IFTAP | protein_coding | 1676,523005 | 0,777574295 | 0,149031115 | 5,217529884 | 1,81E-07 | 3,73E-06 |
| ENSG00000116574.6 | RHOU | protein_coding | 4970,960047 | 0,776745777 | 0,128574974 | 6,041189446 | 1,53E-09 | 4,13E-08 |
| ENSG00000173281.5 | PPP1R3B | protein_coding | 1540,088251 | 0,776337272 | 0,145354515 | 5,340991789 | 9,24E-08 | 1,99E-06 |
| ENSG00000182957.16 | SPATA13 | protein_coding | 6263,472149 | 0,775911115 | 0,122047246 | 6,357465186 | 2,05E-10 | 6,03E-09 |
| ENSG00000120875.9 | DUSP4 | protein_coding | 3230,910378 | 0,774951639 | 0,139164868 | 5,568586727 | 2,57E-08 | 5,93E-07 |
| ENSG00000187243.16 | MAGED4B | protein_coding | 5688,45756 | 0,774830857 | 0,12530215 | 6,183699597 | 6,26E-10 | 1,76E-08 |
| ENSG00000167306.20 | MYO5B | protein_coding | 4518,761864 | 0,7746539 | 0,138927373 | 5,575963047 | 2,46E-08 | 5,70E-07 |
| ENSG00000186472.20 | PCLO | protein_coding | 2120,359572 | 0,773613963 | 0,23610078 | 3,276626034 | 0,001050554 | 0,009061949 |
| ENSG00000176473.13 | WDR25 | protein_coding | 1390,115717 | 0,772923269 | 0,148034972 | 5,22122077 | 1,78E-07 | 3,66E-06 |
| ENSG00000163635.20 | ATXN7 | protein_coding | 1736,28517 | 0,772195381 | 0,160679937 | 4,805798378 | 1,54E-06 | 2,74E-05 |

| | | | | | | | | |
|---|---|---|---|---|---|---|---|---|
| ENSG00000167004.13 | PDIA3 | protein_coding | 131932,6727 | 0,771213514 | 0,112988818 | 6,825573784 | 8,76E-12 | 2,97E-10 |
| ENSG00000177542.11 | SLC25A22 | protein_coding | 6429,496079 | 0,771188233 | 0,122755939 | 6,282288564 | 3,34E-10 | 9,63E-09 |
| ENSG00000008710.20 | PKD1 | protein_coding | 2189,881959 | 0,771135246 | 0,209479658 | 3,681193935 | 0,000232144 | 0,002518105 |
| ENSG00000023318.8 | ERP44 | protein_coding | 10721,05232 | 0,769156336 | 0,13222386 | 5,817076716 | 5,99E-09 | 1,51E-07 |
| ENSG00000162695.12 | SLC30A7 | protein_coding | 3623,52695 | 0,768939658 | 0,142990615 | 5,37755335 | 7,55E-08 | 1,65E-06 |
| ENSG00000130066.16 | SAT1 | protein_coding | 4677,532716 | 0,768488585 | 0,153142734 | 5,018119793 | 5,22E-07 | 1,00E-05 |
| ENSG00000196141.14 | SPATS2L | protein_coding | 4424,405423 | 0,767762241 | 0,143934116 | 5,334122735 | 9,60E-08 | 2,06E-06 |
| ENSG00000160602.14 | NEK8 | protein_coding | 377,5677736 | 0,767286905 | 0,242317951 | 3,166446815 | 0,001543136 | 0,012604582 |
| ENSG00000169241.19 | SLC50A1 | protein_coding | 4934,116614 | 0,765208639 | 0,134793311 | 5,676903644 | 1,37E-08 | 3,29E-07 |
| ENSG00000119938.9 | PPP1R3C | protein_coding | 3338,043474 | 0,764493842 | 0,122793376 | 6,225855696 | 4,79E-10 | 1,36E-08 |
| ENSG00000108375.13 | RNF43 | protein_coding | 3071,126973 | 0,764346246 | 0,138608706 | 5,514417292 | 3,50E-08 | 7,98E-07 |
| ENSG00000136960.13 | ENPP2 | protein_coding | 35007,2287 | 0,764196099 | 0,125921751 | 6,06881727 | 1,29E-09 | 3,52E-08 |
| ENSG00000150977.10 | RILPL2 | protein_coding | 256,3336571 | 0,764117097 | 0,243980439 | 3,131878516 | 0,001736917 | 0,013915945 |
| ENSG00000205856.4 | C22orf42 | protein_coding | 5255,555833 | 0,7638141 | 0,131227476 | 5,820534865 | 5,87E-09 | 1,48E-07 |
| ENSG00000069020.18 | MAST4 | protein_coding | 3295,675432 | 0,763643498 | 0,146845416 | 5,200322334 | 1,99E-07 | 4,07E-06 |
| ENSG00000102316.17 | MAGED2 | protein_coding | 23773,22807 | 0,760881237 | 0,115960325 | 6,56156526 | 5,32E-11 | 1,65E-09 |
| ENSG00000165322.18 | ARHGAP12 | protein_coding | 2525,649529 | 0,760735249 | 0,150076908 | 5,068969383 | 4,00E-07 | 7,81E-06 |
| ENSG00000152217.20 | SETBP1 | protein_coding | 677,8501687 | 0,759996082 | 0,232807385 | 3,264484431 | 0,001096635 | 0,009385957 |
| ENSG00000162374.18 | ELAVL4 | protein_coding | 6241,084892 | 0,759489016 | 0,119361101 | 6,362952479 | 1,98E-10 | 5,84E-09 |
| ENSG00000026652.15 | AGPAT4 | protein_coding | 2370,033622 | 0,759289939 | 0,155903642 | 4,870251461 | 1,11E-06 | 2,03E-05 |
| ENSG00000167578.18 | RAB4B | protein_coding | 1558,051535 | 0,759272957 | 0,150997941 | 5,028366287 | 4,95E-07 | 9,56E-06 |
| ENSG00000162729.14 | IGSF8 | protein_coding | 4094,256223 | 0,759097094 | 0,145171027 | 5,228984819 | 1,70E-07 | 3,52E-06 |
| ENSG00000005810.18 | MYCBP2 | protein_coding | 8131,172767 | 0,757251919 | 0,135001073 | 5,609228874 | 2,03E-08 | 4,77E-07 |
| ENSG00000156958.15 | GALK2 | protein_coding | 4583,569182 | 0,756387556 | 0,131337509 | 5,759113009 | 8,46E-09 | 2,08E-07 |
| ENSG00000145780.8 | FEM1C | protein_coding | 3710,888906 | 0,756052551 | 0,131495621 | 5,749640538 | 8,94E-09 | 2,18E-07 |
| ENSG00000164463.12 | CREBRF | protein_coding | 1143,8654 | 0,755722132 | 0,14933731 | 5,060504536 | 4,18E-07 | 8,15E-06 |
| ENSG00000259024.6 | TVP23C-CDRT4 | protein_coding | 900,9131178 | 0,755130298 | 0,198779654 | 3,798830931 | 0,00014538 | 0,001662465 |
| ENSG00000038274.17 | MAT2B | protein_coding | 2985,214036 | 0,754013977 | 0,143224113 | 5,264574246 | 1,41E-07 | 2,92E-06 |
| ENSG00000118804.8 | STBD1 | protein_coding | 786,5010695 | 0,752604097 | 0,169589642 | 4,437795171 | 9,09E-06 | 0,000139595 |
| ENSG00000115738.10 | ID2 | protein_coding | 9702,252923 | 0,751065438 | 0,126972146 | 5,915198427 | 3,31E-09 | 8,64E-08 |
| ENSG00000176532.4 | PRR15 | protein_coding | 856,5588575 | 0,75106122 | 0,158654291 | 4,733948364 | 2,20E-06 | 3,80E-05 |
| ENSG00000107562.16 | CXCL12 | protein_coding | 5095,543681 | 0,74957428 | 0,17340855 | 4,322591236 | 1,54E-05 | 0,000223086 |
| ENSG00000138433.16 | CIR1 | protein_coding | 3194,238835 | 0,747884512 | 0,148919658 | 5,022067089 | 5,11E-07 | 9,85E-06 |
| ENSG00000173486.14 | FKBP2 | protein_coding | 9051,986162 | 0,747788791 | 0,12730437 | 5,874022966 | 4,25E-09 | 1,09E-07 |
| ENSG00000104341.17 | LAPTM4B | protein_coding | 38273,71317 | 0,747536142 | 0,120827037 | 6,186828381 | 6,14E-10 | 1,73E-08 |
| ENSG00000006459.11 | KDM7A | protein_coding | 7563,251009 | 0,747137262 | 0,166106433 | 4,497942971 | 6,86E-06 | 0,00010776 |

| | | | | | | | | |
|---|---|---|---|---|---|---|---|---|
| ENSG00000160190.14 | SLC37A1 | protein_coding | 5162,894459 | 0,747107692 | 0,133853923 | 5,581515107 | 2,38E-08 | 5,54E-07 |
| ENSG00000164305.19 | CASP3 | protein_coding | 10364,46692 | 0,74649991 | 0,129497404 | 5,764593609 | 8,19E-09 | 2,02E-07 |
| ENSG00000175197.12 | DDIT3 | protein_coding | 521,9298404 | 0,746339252 | 0,196467356 | 3,798795216 | 0,000145401 | 0,001662465 |
| ENSG00000170421.12 | KRT8 | protein_coding | 55470,4445 | 0,746230875 | 0,119331646 | 6,253419763 | 4,02E-10 | 1,15E-08 |
| ENSG00000054219.11 | LY75 | protein_coding | 2169,938503 | 0,745892821 | 0,130750364 | 5,704709346 | 1,17E-08 | 2,83E-07 |
| ENSG00000102158.20 | MAGT1 | protein_coding | 9956,3972 | 0,745290518 | 0,139298701 | 5,350304857 | 8,78E-08 | 1,90E-06 |
| ENSG00000181191.12 | PJA1 | protein_coding | 6214,711868 | 0,744927093 | 0,117876084 | 6,319577905 | 2,62E-10 | 7,65E-09 |
| ENSG00000070540.13 | WIPI1 | protein_coding | 742,3533367 | 0,743961781 | 0,173860401 | 4,279075491 | 1,88E-05 | 0,000266005 |
| ENSG00000254319.6 | AC246817.2 | lncRNA | 258,9651193 | 0,743647482 | 0,247334948 | 3,006641352 | 0,002641512 | 0,019560286 |
| ENSG00000130768.15 | SMPDL3B | protein_coding | 7969,653977 | 0,742802335 | 0,162445061 | 4,572637224 | 4,82E-06 | 7,79E-05 |
| ENSG00000101349.17 | PAK5 | protein_coding | 1184,284141 | 0,742695785 | 0,150128043 | 4,947082289 | 7,53E-07 | 1,41E-05 |
| ENSG00000127666.10 | TICAM1 | protein_coding | 1988,318289 | 0,739227613 | 0,164235659 | 4,501017721 | 6,76E-06 | 0,000106372 |
| ENSG00000184384.14 | MAML2 | protein_coding | 658,9949177 | 0,737596058 | 0,195863323 | 3,765871249 | 0,000165969 | 0,001874092 |
| ENSG00000149300.10 | C11orf52 | protein_coding | 620,0446513 | 0,736813105 | 0,200522454 | 3,674466823 | 0,000238347 | 0,002572055 |
| ENSG00000090776.6 | EFNB1 | protein_coding | 6187,863984 | 0,73582499 | 0,129934039 | 5,663065625 | 1,49E-08 | 3,55E-07 |
| ENSG00000108176.15 | DNAJC12 | protein_coding | 4121,69523 | 0,734783042 | 0,125360965 | 5,861338414 | 4,59E-09 | 1,17E-07 |
| ENSG00000117461.15 | PIK3R3 | protein_coding | 9264,41509 | 0,734353635 | 0,118016603 | 6,222460353 | 4,89E-10 | 1,39E-08 |
| ENSG00000189060.5 | H1-0 | protein_coding | 8827,948131 | 0,73231598 | 0,116001791 | 6,312971345 | 2,74E-10 | 7,97E-09 |
| ENSG00000135655.16 | USP15 | protein_coding | 5499,186816 | 0,732113613 | 0,129755838 | 5,642240267 | 1,68E-08 | 3,98E-07 |
| ENSG00000137824.16 | RMDN3 | protein_coding | 2178,053979 | 0,731375914 | 0,130665542 | 5,59731281 | 2,18E-08 | 5,10E-07 |
| ENSG00000154217.15 | PITPNC1 | protein_coding | 1333,511442 | 0,730012875 | 0,147400229 | 4,952589822 | 7,32E-07 | 1,37E-05 |
| ENSG00000166532.16 | RIMKLB | protein_coding | 514,0482843 | 0,726728409 | 0,193482671 | 3,756038749 | 0,000172624 | 0,001945039 |
| ENSG00000166507.19 | NDST2 | protein_coding | 2233,458992 | 0,725975515 | 0,138741772 | 5,232566274 | 1,67E-07 | 3,46E-06 |
| ENSG00000091513.16 | TF | protein_coding | 444,5052285 | 0,725382943 | 0,232041723 | 3,126088421 | 0,001771483 | 0,014106501 |
| ENSG00000205155.8 | PSENEN | protein_coding | 3851,669436 | 0,725105089 | 0,123233816 | 5,883978206 | 4,01E-09 | 1,03E-07 |
| ENSG00000130202.10 | NECTIN2 | protein_coding | 2892,583899 | 0,724853215 | 0,162673996 | 4,455864075 | 8,36E-06 | 0,000129 |
| ENSG00000160209.19 | PDXK | protein_coding | 4097,170798 | 0,72377092 | 0,134404938 | 5,385002466 | 7,24E-08 | 1,59E-06 |
| ENSG00000109854.14 | HTATIP2 | protein_coding | 6618,561783 | 0,723104183 | 0,136213133 | 5,308623106 | 1,10E-07 | 2,34E-06 |
| ENSG00000179008.9 | C14orf39 | protein_coding | 223,6268089 | 0,722475943 | 0,256585513 | 2,815731626 | 0,00486663 | 0,032255569 |
| ENSG00000265763.4 | ZNF488 | protein_coding | 1929,5619 | 0,722220464 | 0,147073008 | 4,910625512 | 9,08E-07 | 1,68E-05 |
| ENSG00000175073.8 | VCPIP1 | protein_coding | 3535,854956 | 0,721061443 | 0,135516018 | 5,320857653 | 1,03E-07 | 2,20E-06 |
| ENSG00000175602.4 | CCDC85B | protein_coding | 1588,378644 | 0,720431729 | 0,154123627 | 4,674375637 | 2,95E-06 | 4,96E-05 |
| ENSG00000129116.19 | PALLD | protein_coding | 14265,53587 | 0,719040277 | 0,133762799 | 5,375487658 | 7,64E-08 | 1,66E-06 |
| ENSG00000153721.19 | CNKSR3 | protein_coding | 1018,478485 | 0,718690487 | 0,163910577 | 4,38464986 | 1,16E-05 | 0,000174718 |
| ENSG00000112308.13 | C6orf62 | protein_coding | 6772,464717 | 0,718268666 | 0,148877163 | 4,824572484 | 1,40E-06 | 2,51E-05 |
| ENSG00000103496.15 | STX4 | protein_coding | 2088,372016 | 0,717736761 | 0,134735257 | 5,32701521 | 9,98E-08 | 2,13E-06 |

| | | | | | | | | |
|---|---|---|---|---|---|---|---|---|
| ENSG00000267002.4 | AC060780.1 | lncRNA | 743,3952855 | 0,717599643 | 0,177629152 | 4,039875404 | 5,35E-05 | 0,000681019 |
| ENSG00000121297.8 | TSHZ3 | protein_coding | 655,1112124 | 0,716269802 | 0,265705882 | 2,695724299 | 0,007023577 | 0,043246644 |
| ENSG00000108387.15 | SEPTIN4 | protein_coding | 195,6234144 | 0,715912115 | 0,266698078 | 2,684354232 | 0,007267006 | 0,044405949 |
| ENSG00000113580.15 | NR3C1 | protein_coding | 21421,65346 | 0,715876718 | 0,108883792 | 6,574685764 | 4,88E-11 | 1,52E-09 |
| ENSG00000179921.15 | GPBAR1 | protein_coding | 3871,449341 | 0,715459136 | 0,120100286 | 5,957180935 | 2,57E-09 | 6,79E-08 |
| ENSG00000198648.11 | STK39 | protein_coding | 6804,820633 | 0,71406074 | 0,114991584 | 6,209678251 | 5,31E-10 | 1,51E-08 |
| ENSG00000153975.10 | ZUP1 | protein_coding | 1853,046794 | 0,714051741 | 0,134003168 | 5,328618378 | 9,90E-08 | 2,12E-06 |
| ENSG00000109756.9 | RAPGEF2 | protein_coding | 1373,337311 | 0,71343477 | 0,212543052 | 3,356660052 | 0,0007889 | 0,007094343 |
| ENSG00000197019.5 | SERTAD1 | protein_coding | 826,168566 | 0,713359009 | 0,173345311 | 4,115248377 | 3,87E-05 | 0,000508917 |
| ENSG00000239521.9 | CASTOR3 | transcribed_unproc | 2806,614421 | 0,712063262 | 0,131538843 | 5,413330758 | 6,19E-08 | 1,36E-06 |
| ENSG00000132256.19 | TRIM5 | protein_coding | 341,4924493 | 0,711639992 | 0,245462468 | 2,899180461 | 0,003741395 | 0,026204785 |
| ENSG00000105939.13 | ZC3HAV1 | protein_coding | 6063,663859 | 0,711321476 | 0,157484344 | 4,516775802 | 6,28E-06 | 9,91E-05 |
| ENSG00000112763.17 | BTN2A1 | protein_coding | 5230,902619 | 0,710886079 | 0,123877597 | 5,738616975 | 9,55E-09 | 2,33E-07 |
| ENSG00000085832.17 | EPS15 | protein_coding | 6427,47057 | 0,710537869 | 0,128298049 | 5,538181418 | 3,06E-08 | 7,02E-07 |
| ENSG00000180616.9 | SSTR2 | protein_coding | 18540,0158 | 0,710052401 | 0,110942986 | 6,400155851 | 1,55E-10 | 4,60E-09 |
| ENSG00000237036.5 | ZEB1-AS1 | lncRNA | 393,4501639 | 0,70910744 | 0,215145629 | 3,29594166 | 0,000980924 | 0,008538728 |
| ENSG00000149269.10 | PAK1 | protein_coding | 4575,306202 | 0,706207706 | 0,13137826 | 5,375377238 | 7,64E-08 | 1,66E-06 |
| ENSG00000157353.17 | FCSK | protein_coding | 1491,650337 | 0,70577731 | 0,145532714 | 4,849612768 | 1,24E-06 | 2,24E-05 |
| ENSG00000130758.8 | MAP3K10 | protein_coding | 1091,934288 | 0,705443712 | 0,202866472 | 3,477379512 | 0,000506341 | 0,004896185 |
| ENSG00000095380.11 | NANS | protein_coding | 3044,971447 | 0,705008182 | 0,16145673 | 4,36654564 | 1,26E-05 | 0,000187283 |
| ENSG00000143013.13 | LMO4 | protein_coding | 11650,68625 | 0,70482847 | 0,118399739 | 5,952956284 | 2,63E-09 | 6,96E-08 |
| ENSG00000279013.1 | AC110774.1 | TEC | 435,4619273 | 0,704828161 | 0,198641617 | 3,548240158 | 0,000387814 | 0,003897693 |
| ENSG00000144468.17 | RHBDD1 | protein_coding | 2884,292324 | 0,70444126 | 0,12618112 | 5,582778622 | 2,37E-08 | 5,51E-07 |
| ENSG00000135677.11 | GNS | protein_coding | 38725,48085 | 0,704416485 | 0,111205069 | 6,334391891 | 2,38E-10 | 6,96E-09 |
| ENSG00000110218.9 | PANX1 | protein_coding | 2803,716042 | 0,703717744 | 0,130146834 | 5,407106144 | 6,41E-08 | 1,41E-06 |
| ENSG00000168297.16 | PXK | protein_coding | 3048,118564 | 0,703443062 | 0,129520393 | 5,431137469 | 5,60E-08 | 1,24E-06 |
| ENSG00000099937.11 | SERPIND1 | protein_coding | 256,0113907 | 0,702039304 | 0,242113383 | 2,899630309 | 0,00373603 | 0,026184731 |
| ENSG00000166333.14 | ILK | protein_coding | 9423,675207 | 0,701759851 | 0,118845346 | 5,904815583 | 3,53E-09 | 9,12E-08 |
| ENSG00000162981.14 | LRATD1 | protein_coding | 19157,57611 | 0,701530879 | 0,118696067 | 5,910312777 | 3,41E-09 | 8,88E-08 |
| ENSG00000047634.15 | SCML1 | protein_coding | 4571,189291 | 0,700389282 | 0,14657685 | 4,778307643 | 1,77E-06 | 3,11E-05 |
| ENSG00000234127.9 | TRIM26 | protein_coding | 6656,852633 | 0,699884605 | 0,11562046 | 6,05329371 | 1,42E-09 | 3,84E-08 |
| ENSG00000106588.12 | PSMA2 | protein_coding | 16478,41282 | 0,698495077 | 0,127489703 | 5,478835219 | 4,28E-08 | 9,62E-07 |
| ENSG00000170142.12 | UBE2E1 | protein_coding | 2221,221175 | 0,698362017 | 0,138666598 | 5,036267031 | 4,75E-07 | 9,18E-06 |
| ENSG00000247400.4 | DNAJC3-DT | lncRNA | 1385,698253 | 0,697485999 | 0,156027712 | 4,47026999 | 7,81E-06 | 0,000121235 |
| ENSG00000197948.11 | FCHSD1 | protein_coding | 451,1058496 | 0,695922466 | 0,205799612 | 3,381553821 | 0,000720771 | 0,006566294 |
| ENSG00000118707.11 | TGIF2 | protein_coding | 1049,206258 | 0,695813067 | 0,167296681 | 4,15915644 | 3,19E-05 | 0,000430043 |

| | | | | | | | | |
|---|---|---|---|---|---|---|---|---|
| ENSG00000187678.10 | SPRY4 | protein_coding | 924,8794978 | 0,695697798 | 0,20707315 | 3,359671677 | 0,000780352 | 0,00702652 |
| ENSG00000163110.15 | PDLIM5 | protein_coding | 11010,30948 | 0,694778069 | 0,113384189 | 6,127645107 | 8,92E-10 | 2,47E-08 |
| ENSG00000103021.9 | CCDC113 | protein_coding | 235,8463753 | 0,693988208 | 0,2475427 | 2,803509083 | 0,005054981 | 0,033195114 |
| ENSG00000243414.6 | TICAM2 | protein_coding | 386,8361046 | 0,693188231 | 0,223964511 | 3,095080676 | 0,001967594 | 0,01536999 |
| ENSG00000172716.16 | SLFN11 | protein_coding | 21008,61555 | 0,692591974 | 0,121555744 | 5,697731375 | 1,21E-08 | 2,94E-07 |
| ENSG00000155313.15 | USP25 | protein_coding | 1873,674858 | 0,692430106 | 0,162722588 | 4,255279581 | 2,09E-05 | 0,000293354 |
| ENSG00000250742.4 | LINC02381 | lncRNA | 2581,288339 | 0,691218102 | 0,133192106 | 5,189632641 | 2,11E-07 | 4,30E-06 |
| ENSG00000183161.6 | FANCF | protein_coding | 5502,201648 | 0,690908714 | 0,120070881 | 5,754173763 | 8,71E-09 | 2,13E-07 |
| ENSG00000159248.5 | GJD2 | protein_coding | 3533,241681 | 0,689615631 | 0,1413232 | 4,879705737 | 1,06E-06 | 1,94E-05 |
| ENSG00000100647.8 | SUSD6 | protein_coding | 2310,145488 | 0,689606787 | 0,15988201 | 4,313223155 | 1,61E-05 | 0,000231496 |
| ENSG00000275023.5 | MLLT6 | protein_coding | 2787,412427 | 0,689245928 | 0,155434493 | 4,434317735 | 9,24E-06 | 0,000141452 |
| ENSG00000156030.13 | MIDEAS | protein_coding | 987,0158994 | 0,688382578 | 0,244096187 | 2,820128349 | 0,004800444 | 0,031883663 |
| ENSG00000120899.18 | PTK2B | protein_coding | 967,7549026 | 0,686815043 | 0,17970728 | 3,821854316 | 0,000132452 | 0,001529444 |
| ENSG00000105321.14 | CCDC9 | protein_coding | 875,5413591 | 0,684964409 | 0,204609539 | 3,347666054 | 0,000814951 | 0,007300389 |
| ENSG00000170577.8 | SIX2 | protein_coding | 930,1903846 | 0,684166802 | 0,156933891 | 4,359586042 | 1,30E-05 | 0,000192656 |
| ENSG00000150347.16 | ARID5B | protein_coding | 3791,130953 | 0,683931808 | 0,15086119 | 4,533517258 | 5,80E-06 | 9,24E-05 |
| ENSG00000104324.16 | CPQ | protein_coding | 6177,781686 | 0,680749494 | 0,129281804 | 5,265624944 | 1,40E-07 | 2,90E-06 |
| ENSG00000169122.11 | FAM110B | protein_coding | 1616,845596 | 0,679958917 | 0,157406848 | 4,319754351 | 1,56E-05 | 0,000225525 |
| ENSG00000275496.4 | CU633906.1 | lncRNA | 891,9315949 | 0,679582841 | 0,191450427 | 3,54965435 | 0,000385737 | 0,003884276 |
| ENSG00000005471.19 | ABCB4 | protein_coding | 2046,479187 | 0,679566889 | 0,132894134 | 5,113595843 | 3,16E-07 | 6,30E-06 |
| ENSG00000075420.13 | FNDC3B | protein_coding | 5448,184025 | 0,679279655 | 0,135984119 | 4,995286645 | 5,87E-07 | 1,12E-05 |
| ENSG00000148143.13 | ZNF462 | protein_coding | 2033,3448 | 0,679024116 | 0,24913632 | 2,725512346 | 0,006420178 | 0,040301774 |
| ENSG00000123570.4 | RAB9B | protein_coding | 5212,524091 | 0,678077608 | 0,124887088 | 5,429525311 | 5,65E-08 | 1,25E-06 |
| ENSG00000189067.13 | LITAF | protein_coding | 835,5372139 | 0,677970121 | 0,178182849 | 3,804912333 | 0,000141854 | 0,001627245 |
| ENSG00000123411.15 | IKZF4 | protein_coding | 642,0072089 | 0,676999299 | 0,241355676 | 2,804986023 | 0,005031876 | 0,033095296 |
| ENSG00000115306.16 | SPTBN1 | protein_coding | 10075,69757 | 0,675884636 | 0,204431687 | 3,306163765 | 0,000945828 | 0,008277971 |
| ENSG00000137872.17 | SEMA6D | protein_coding | 1026,587034 | 0,674506964 | 0,204895063 | 3,291962991 | 0,000994907 | 0,008638896 |
| ENSG00000104856.14 | RELB | protein_coding | 649,0480422 | 0,674256385 | 0,195675155 | 3,445794559 | 0,000569383 | 0,005403461 |
| ENSG00000148411.8 | NACC2 | protein_coding | 308,3206983 | 0,672805454 | 0,250700117 | 2,683706177 | 0,007281107 | 0,044440223 |
| ENSG00000178078.12 | STAP2 | protein_coding | 3434,214015 | 0,671591031 | 0,136301495 | 4,927246258 | 8,34E-07 | 1,55E-05 |
| ENSG00000112531.17 | QKI | protein_coding | 921,1889659 | 0,670784102 | 0,195090642 | 3,438320231 | 0,000585335 | 0,005522302 |
| ENSG00000077150.20 | NFKB2 | protein_coding | 1519,868241 | 0,670758894 | 0,163590491 | 4,100231561 | 4,13E-05 | 0,000538358 |
| ENSG00000100095.19 | SEZ6L | protein_coding | 10982,76849 | 0,669765014 | 0,182692364 | 3,666081058 | 0,000246296 | 0,002646922 |
| ENSG00000099308.10 | MAST3 | protein_coding | 2047,161798 | 0,668559059 | 0,203733921 | 3,281530425 | 0,001032454 | 0,00892789 |
| ENSG00000125355.16 | TMEM255A | protein_coding | 532,338469 | 0,66852331 | 0,189692389 | 3,524249512 | 0,000424684 | 0,004219631 |
| ENSG00000131979.20 | GCH1 | protein_coding | 24630,6081 | 0,667593651 | 0,121869302 | 5,47794759 | 4,30E-08 | 9,65E-07 |

| | | | | | | | | |
|---|---|---|---|---|---|---|---|---|
| ENSG00000165626.17 | BEND7 | protein_coding | 1570,100027 | 0,666759872 | 0,206669778 | 3,226208873 | 0,001254418 | 0,010597755 |
| ENSG00000188766.12 | SPRED3 | protein_coding | 337,8926846 | 0,666525381 | 0,234925764 | 2,837174471 | 0,004551473 | 0,030648149 |
| ENSG00000236753.7 | MKLN1-AS | lncRNA | 236,528877 | 0,665807859 | 0,247060376 | 2,694919641 | 0,007040561 | 0,043338472 |
| ENSG00000008086.13 | CDKL5 | protein_coding | 3154,031098 | 0,665370284 | 0,140403915 | 4,738972447 | 2,15E-06 | 3,72E-05 |
| ENSG00000128590.5 | DNAJB9 | protein_coding | 10579,61441 | 0,664891791 | 0,165166921 | 4,025574782 | 5,68E-05 | 0,000719823 |
| ENSG00000173889.16 | PHC3 | protein_coding | 4246,655007 | 0,664829866 | 0,184066057 | 3,6119091 | 0,000303951 | 0,003156357 |
| ENSG00000112304.11 | ACOT13 | protein_coding | 4065,573392 | 0,664669154 | 0,125517406 | 5,295434104 | 1,19E-07 | 2,50E-06 |
| ENSG00000196083.10 | IL1RAP | protein_coding | 2158,278565 | 0,664182275 | 0,140610248 | 4,723569471 | 2,32E-06 | 3,98E-05 |
| ENSG00000165591.6 | FAAH2 | protein_coding | 1923,884431 | 0,663829466 | 0,15564048 | 4,265146602 | 2,00E-05 | 0,0002822 |
| ENSG00000142188.17 | TMEM50B | protein_coding | 5573,441757 | 0,663810787 | 0,135922305 | 4,883751684 | 1,04E-06 | 1,90E-05 |
| ENSG00000117519.16 | CNN3 | protein_coding | 7449,739901 | 0,663061117 | 0,114564405 | 5,787671302 | 7,14E-09 | 1,77E-07 |
| ENSG00000103460.17 | TOX3 | protein_coding | 2072,385933 | 0,662511261 | 0,207335057 | 3,195365371 | 0,001396539 | 0,011569667 |
| ENSG00000165259.14 | HDX | protein_coding | 633,1059803 | 0,662404567 | 0,174734867 | 3,790912364 | 0,000150095 | 0,001709596 |
| ENSG00000164713.10 | BRI3 | protein_coding | 6216,40065 | 0,66217203 | 0,124118931 | 5,334980109 | 9,56E-08 | 2,05E-06 |
| ENSG00000111696.12 | NT5DC3 | protein_coding | 2780,931811 | 0,66205073 | 0,193118528 | 3,42820928 | 0,000607577 | 0,005693654 |
| ENSG00000178607.16 | ERN1 | protein_coding | 1227,784412 | 0,661417081 | 0,211266292 | 3,130726978 | 0,001743742 | 0,013959938 |
| ENSG00000136156.15 | ITM2B | protein_coding | 56362,65076 | 0,660932076 | 0,12299564 | 5,373622003 | 7,72E-08 | 1,67E-06 |
| ENSG00000101745.17 | ANKRD12 | protein_coding | 2383,380602 | 0,660781455 | 0,150970268 | 4,376897938 | 1,20E-05 | 0,000180089 |
| ENSG00000185551.15 | NR2F2 | protein_coding | 402,2946995 | 0,660741788 | 0,230103016 | 2,871504243 | 0,004085233 | 0,028188646 |
| ENSG00000170919.16 | TPT1-AS1 | lncRNA | 430,0086719 | 0,660015206 | 0,237700854 | 2,776663169 | 0,005492006 | 0,035551994 |
| ENSG00000168495.13 | POLR3D | protein_coding | 2579,480522 | 0,659939049 | 0,137783653 | 4,789675961 | 1,67E-06 | 2,95E-05 |
| ENSG00000120658.13 | ENOX1 | protein_coding | 552,3920859 | 0,657582986 | 0,189265948 | 3,474386129 | 0,000512024 | 0,004942008 |
| ENSG00000213930.12 | GALT | protein_coding | 1061,321217 | 0,656018746 | 0,151560582 | 4,328425888 | 1,50E-05 | 0,000218637 |
| ENSG00000105402.8 | NAPA | protein_coding | 10232,91781 | 0,655988772 | 0,116663329 | 5,622921781 | 1,88E-08 | 4,42E-07 |
| ENSG00000101294.18 | HM13 | protein_coding | 27810,33077 | 0,655733342 | 0,133692462 | 4,904789193 | 9,35E-07 | 1,73E-05 |
| ENSG00000106080.11 | FKBP14 | protein_coding | 506,9308987 | 0,654958196 | 0,21094098 | 3,104935775 | 0,001903203 | 0,01496753 |
| ENSG00000265491.5 | RNF115 | protein_coding | 2273,038971 | 0,654402607 | 0,135008131 | 4,847134778 | 1,25E-06 | 2,26E-05 |
| ENSG00000134324.12 | LPIN1 | protein_coding | 10771,56786 | 0,654188225 | 0,118370703 | 5,526605914 | 3,26E-08 | 7,46E-07 |
| ENSG00000153944.12 | MSI2 | protein_coding | 5391,720032 | 0,653858165 | 0,124233064 | 5,263157373 | 1,42E-07 | 2,94E-06 |
| ENSG00000166598.15 | HSP90B1 | protein_coding | 216888,3468 | 0,652697591 | 0,123928688 | 5,266719117 | 1,39E-07 | 2,89E-06 |
| ENSG00000155158.20 | TTC39B | protein_coding | 880,1869642 | 0,651940719 | 0,194557693 | 3,350886356 | 0,000805534 | 0,007231494 |
| ENSG00000139971.15 | ARMH4 | protein_coding | 633,9510569 | 0,650936671 | 0,178347044 | 3,649831575 | 0,000262412 | 0,002795726 |
| ENSG00000138279.16 | ANXA7 | protein_coding | 25028,35189 | 0,65025808 | 0,112122415 | 5,799536866 | 6,65E-09 | 1,66E-07 |
| ENSG00000165801.10 | ARHGEF40 | protein_coding | 2239,490209 | 0,649638909 | 0,151285032 | 4,294138675 | 1,75E-05 | 0,000250952 |
| ENSG00000075539.15 | FRYL | protein_coding | 4407,486588 | 0,649083939 | 0,156484127 | 4,147921906 | 3,36E-05 | 0,000448235 |
| ENSG00000100226.16 | GTPBP1 | protein_coding | 4532,365395 | 0,648107012 | 0,161847098 | 4,004440108 | 6,22E-05 | 0,00077929 |

| | | | | | | | | |
|---|---|---|---|---|---|---|---|---|
| ENSG00000278266.1 | AC079949.2 | lncRNA | 3067,241065 | 0,647550221 | 0,12956347 | 4,997938255 | 5,79E-07 | 1,11E-05 |
| ENSG00000137200.13 | CMTR1 | protein_coding | 6819,872768 | 0,647432377 | 0,114298385 | 5,664405289 | 1,48E-08 | 3,53E-07 |
| ENSG00000100908.14 | EMC9 | protein_coding | 1655,254212 | 0,646993689 | 0,137636231 | 4,700751287 | 2,59E-06 | 4,39E-05 |
| ENSG00000073578.17 | SDHA | protein_coding | 14000,74133 | 0,64681197 | 0,116614363 | 5,546589221 | 2,91E-08 | 6,69E-07 |
| ENSG00000198692.10 | EIF1AY | protein_coding | 2998,421232 | 0,645492124 | 0,141888698 | 4,54928499 | 5,38E-06 | 8,63E-05 |
| ENSG00000175215.11 | CTDSP2 | protein_coding | 10101,2363 | 0,644797212 | 0,125845788 | 5,123709117 | 3,00E-07 | 6,00E-06 |
| ENSG00000167565.13 | SERTAD3 | protein_coding | 866,7761386 | 0,644255124 | 0,161124763 | 3,99848608 | 6,37E-05 | 0,000797242 |
| ENSG00000163191.6 | S100A11 | protein_coding | 2516,494271 | 0,644154746 | 0,174115987 | 3,699572668 | 0,000215963 | 0,00237457 |
| ENSG00000163788.14 | SNRK | protein_coding | 5710,517721 | 0,642433923 | 0,122788006 | 5,232057614 | 1,68E-07 | 3,46E-06 |
| ENSG00000162889.11 | MAPKAPK2 | protein_coding | 10218,35011 | 0,642268482 | 0,115695051 | 5,55139117 | 2,83E-08 | 6,53E-07 |
| ENSG00000102858.13 | MGRN1 | protein_coding | 4511,415003 | 0,642245381 | 0,189066244 | 3,396933094 | 0,000681456 | 0,0062544 |
| ENSG00000144785.8 | AC073896.1 | protein_coding | 612,8997684 | 0,641109851 | 0,228181257 | 2,809651681 | 0,004959514 | 0,032732482 |
| ENSG00000188690.15 | UROS | protein_coding | 3605,692193 | 0,640615412 | 0,137839558 | 4,647544017 | 3,36E-06 | 5,61E-05 |
| ENSG00000100219.16 | XBP1 | protein_coding | 31744,39808 | 0,639538615 | 0,118975694 | 5,375371998 | 7,64E-08 | 1,66E-06 |
| ENSG00000103222.20 | ABCC1 | protein_coding | 2823,504066 | 0,639453182 | 0,141944851 | 4,504941014 | 6,64E-06 | 0,000104583 |
| ENSG00000090520.11 | DNAJB11 | protein_coding | 17209,02399 | 0,639309357 | 0,129655579 | 4,930827976 | 8,19E-07 | 1,52E-05 |
| ENSG00000102580.15 | DNAJC3 | protein_coding | 11828,70043 | 0,639054063 | 0,141889355 | 4,503890108 | 6,67E-06 | 0,000105022 |
| ENSG00000140406.4 | TLNRD1 | protein_coding | 3598,969561 | 0,637527169 | 0,129183216 | 4,935061904 | 8,01E-07 | 1,49E-05 |
| ENSG00000117523.16 | PRRC2C | protein_coding | 3949,983468 | 0,637461915 | 0,19253359 | 3,310912735 | 0,000929922 | 0,008172929 |
| ENSG00000153982.11 | GDPD1 | protein_coding | 537,9401712 | 0,636791614 | 0,225035874 | 2,829733781 | 0,004658675 | 0,031159541 |
| ENSG00000086061.16 | DNAJA1 | protein_coding | 56390,46407 | 0,633619014 | 0,117015168 | 5,414845145 | 6,13E-08 | 1,35E-06 |
| ENSG00000035928.16 | RFC1 | protein_coding | 5240,41318 | 0,632819084 | 0,152836545 | 4,140495865 | 3,47E-05 | 0,000460937 |
| ENSG00000136560.14 | TANK | protein_coding | 3953,513058 | 0,63217136 | 0,12211962 | 5,176656793 | 2,26E-07 | 4,60E-06 |
| ENSG00000226479.4 | TMEM185B | protein_coding | 3502,260447 | 0,631933939 | 0,128957465 | 4,900328469 | 9,57E-07 | 1,76E-05 |
| ENSG00000189319.14 | FAM53B | protein_coding | 1164,579836 | 0,631641006 | 0,173087911 | 3,649249704 | 0,000263007 | 0,002800089 |
| ENSG00000198625.13 | MDM4 | protein_coding | 328,6770587 | 0,631564585 | 0,235352627 | 2,683482203 | 0,007285985 | 0,044451854 |
| ENSG00000147100.11 | SLC16A2 | protein_coding | 5998,772758 | 0,631297274 | 0,126855299 | 4,97651481 | 6,47E-07 | 1,22E-05 |
| ENSG00000102081.15 | FMR1 | protein_coding | 2182,945529 | 0,630980632 | 0,165626555 | 3,809658617 | 0,000139159 | 0,001600709 |
| ENSG00000151612.18 | ZNF827 | protein_coding | 2902,039643 | 0,630784027 | 0,224141348 | 2,81422429 | 0,00488951 | 0,032380043 |
| ENSG00000223459.6 | TCAF1P1 | unprocessed_pseud | 1777,637173 | 0,629591015 | 0,1555865 | 4,046565828 | 5,20E-05 | 0,000663063 |
| ENSG00000173826.15 | KCNH6 | protein_coding | 4234,280079 | 0,629001948 | 0,137631668 | 4,570183283 | 4,87E-06 | 7,87E-05 |
| ENSG00000069974.16 | RAB27A | protein_coding | 8173,884463 | 0,628278954 | 0,119241512 | 5,268961669 | 1,37E-07 | 2,86E-06 |
| ENSG00000079257.8 | LXN | protein_coding | 928,1764403 | 0,627089159 | 0,167983489 | 3,73304045 | 0,000189182 | 0,002101077 |
| ENSG00000170175.11 | CHRNB1 | protein_coding | 810,0103798 | 0,626951489 | 0,170879022 | 3,668978684 | 0,000243521 | 0,002621141 |
| ENSG00000225335.3 | AC016027.1 | lncRNA | 297,8953576 | 0,626453002 | 0,22591516 | 2,772956898 | 0,005554948 | 0,035866945 |
| ENSG00000138069.18 | RAB1A | protein_coding | 28563,67085 | 0,62605279 | 0,117231201 | 5,340325664 | 9,28E-08 | 2,00E-06 |

| | | | | | | | | |
|---|---|---|---|---|---|---|---|---|
| ENSG00000139178.11 | C1RL | protein_coding | 924,2000795 | 0,62520637 | 0,195581399 | 3,196655583 | 0,001390308 | 0,011531737 |
| ENSG00000166780.11 | BMERB1 | protein_coding | 2245,979858 | 0,624902777 | 0,165038444 | 3,78640734 | 0,000152841 | 0,001737094 |
| ENSG00000059728.11 | MXD1 | protein_coding | 3044,029062 | 0,622460014 | 0,141546006 | 4,397580906 | 1,09E-05 | 0,000165103 |
| ENSG00000182463.16 | TSHZ2 | protein_coding | 2114,418959 | 0,622323866 | 0,190112658 | 3,273447821 | 0,00106244 | 0,009134366 |
| ENSG00000065675.15 | PRKCQ | protein_coding | 475,554541 | 0,620233465 | 0,19069145 | 3,252549952 | 0,001143745 | 0,009725548 |
| ENSG00000210082.2 | MT-RNR2 | Mt_rRNA | 180933,4298 | 0,61984888 | 0,116833077 | 5,305422873 | 1,12E-07 | 2,37E-06 |
| ENSG00000204427.12 | ABHD16A | protein_coding | 11548,22374 | 0,61960904 | 0,112752405 | 5,49530666 | 3,90E-08 | 8,83E-07 |
| ENSG00000196352.16 | CD55 | protein_coding | 734,4274048 | 0,619305343 | 0,165637143 | 3,738927944 | 0,000184807 | 0,002063428 |
| ENSG00000136874.11 | STX17 | protein_coding | 5718,517733 | 0,619028768 | 0,116194713 | 5,327512353 | 9,96E-08 | 2,13E-06 |
| ENSG00000163479.14 | SSR2 | protein_coding | 28608,78891 | 0,618669006 | 0,150768286 | 4,103442594 | 4,07E-05 | 0,000532264 |
| ENSG00000105722.10 | ERF | protein_coding | 1219,391992 | 0,616981496 | 0,14425052 | 4,277152671 | 1,89E-05 | 0,00026795 |
| ENSG00000170382.12 | LRRN2 | protein_coding | 1110,757528 | 0,616051271 | 0,21013322 | 2,931717655 | 0,003370931 | 0,023966612 |
| ENSG00000261135.1 | AL137802.2 | lncRNA | 598,9480531 | 0,615745791 | 0,186017574 | 3,310148493 | 0,000932465 | 0,008186264 |
| ENSG00000136436.15 | CALCOCO2 | protein_coding | 11001,73545 | 0,614952776 | 0,111941407 | 5,493523723 | 3,94E-08 | 8,91E-07 |
| ENSG00000138771.16 | SHROOM3 | protein_coding | 4129,701068 | 0,613937507 | 0,160518476 | 3,824715533 | 0,000130923 | 0,00151765 |
| ENSG00000178573.7 | MAF | protein_coding | 4800,163253 | 0,613876608 | 0,150423301 | 4,080994118 | 4,48E-05 | 0,00058166 |
| ENSG00000117448.14 | AKR1A1 | protein_coding | 8406,555081 | 0,613265734 | 0,130301137 | 4,706526344 | 2,52E-06 | 4,28E-05 |
| ENSG00000120129.6 | DUSP1 | protein_coding | 1250,109223 | 0,612947016 | 0,152516037 | 4,018902079 | 5,85E-05 | 0,000735929 |
| ENSG00000159202.18 | UBE2Z | protein_coding | 13792,28864 | 0,610534978 | 0,118712756 | 5,142960192 | 2,70E-07 | 5,45E-06 |
| ENSG00000154822.18 | PLCL2 | protein_coding | 5066,332728 | 0,609235576 | 0,118825435 | 5,12714785 | 2,94E-07 | 5,90E-06 |
| ENSG00000102024.18 | PLS3 | protein_coding | 10558,74904 | 0,608989585 | 0,117697295 | 5,17420203 | 2,29E-07 | 4,65E-06 |
| ENSG00000050405.13 | LIMA1 | protein_coding | 9512,770627 | 0,607127573 | 0,113595007 | 5,344667775 | 9,06E-08 | 1,95E-06 |
| ENSG00000143630.10 | HCN3 | protein_coding | 1426,312429 | 0,606662908 | 0,151038257 | 4,016617522 | 5,90E-05 | 0,000742337 |
| ENSG00000143889.16 | HNRNPLL | protein_coding | 7239,551353 | 0,604888755 | 0,114020493 | 5,305088049 | 1,13E-07 | 2,38E-06 |
| ENSG00000137331.12 | IER3 | protein_coding | 1183,896974 | 0,604746906 | 0,147956497 | 4,087329157 | 4,36E-05 | 0,000567766 |
| ENSG00000116954.8 | RRAGC | protein_coding | 4871,784388 | 0,604587236 | 0,125106564 | 4,832578067 | 1,35E-06 | 2,42E-05 |
| ENSG00000074590.13 | NUAK1 | protein_coding | 2189,404948 | 0,603905814 | 0,132811971 | 4,547073638 | 5,44E-06 | 8,71E-05 |
| ENSG00000170385.10 | SLC30A1 | protein_coding | 6122,405738 | 0,603578535 | 0,11879706 | 5,080753144 | 3,76E-07 | 7,39E-06 |
| ENSG00000166908.18 | PIP4K2C | protein_coding | 5948,403722 | 0,602260111 | 0,124997992 | 4,81815829 | 1,45E-06 | 2,59E-05 |
| ENSG00000157064.11 | NMNAT2 | protein_coding | 4863,920692 | 0,601365455 | 0,166608035 | 3,609462507 | 0,000306832 | 0,003179966 |
| ENSG00000170185.10 | USP38 | protein_coding | 9581,773446 | 0,598339284 | 0,122268155 | 4,893664116 | 9,90E-07 | 1,81E-05 |
| ENSG00000156869.14 | FRRS1 | protein_coding | 453,7509527 | 0,593485396 | 0,197451099 | 3,005733573 | 0,00264941 | 0,019606011 |
| ENSG00000162595.7 | DIRAS3 | protein_coding | 880,5999115 | 0,593418017 | 0,165976922 | 3,575304392 | 0,000349821 | 0,003568955 |
| ENSG00000270149.5 | AL591806.2 | protein_coding | 2978,792614 | 0,592507958 | 0,135719088 | 4,365693644 | 1,27E-05 | 0,000187881 |
| ENSG00000087074.8 | PPP1R15A | protein_coding | 1079,098616 | 0,592189001 | 0,153005764 | 3,870370539 | 0,00010867 | 0,001287271 |
| ENSG00000111912.20 | NCOA7 | protein_coding | 1580,358582 | 0,592033186 | 0,174218462 | 3,39822301 | 0,000678251 | 0,006233181 |

| | | | | | | | | |
|---|---|---|---|---|---|---|---|---|
| ENSG00000196911.11 | KPNA5 | protein_coding | 3507,021958 | 0,591293481 | 0,138086092 | 4,28206397 | 1,85E-05 | 0,000262991 |
| ENSG00000137713.16 | PPP2R1B | protein_coding | 6901,391776 | 0,589691191 | 0,136401021 | 4,323216852 | 1,54E-05 | 0,000222627 |
| ENSG00000145901.16 | TNIP1 | protein_coding | 7029,349336 | 0,58924054 | 0,12377437 | 4,760602202 | 1,93E-06 | 3,38E-05 |
| ENSG00000196482.18 | ESRRG | protein_coding | 1394,544001 | 0,58728456 | 0,14130923 | 4,156024053 | 3,24E-05 | 0,000435139 |
| ENSG00000183023.18 | SLC8A1 | protein_coding | 1134,448782 | 0,587038457 | 0,155410578 | 3,777339121 | 0,000158513 | 0,001793765 |
| ENSG00000197102.12 | DYNC1H1 | protein_coding | 45253,49963 | 0,586151465 | 0,156387824 | 3,748063301 | 0,000178205 | 0,001998247 |
| ENSG00000197702.14 | PARVA | protein_coding | 853,0553121 | 0,585869893 | 0,158611946 | 3,693731194 | 0,000220987 | 0,002420697 |
| ENSG00000131435.13 | PDLIM4 | protein_coding | 729,8180064 | 0,585184005 | 0,192020371 | 3,047510027 | 0,002307458 | 0,017527808 |
| ENSG00000115170.15 | ACVR1 | protein_coding | 3377,538768 | 0,584186821 | 0,135151539 | 4,322457779 | 1,54E-05 | 0,000223086 |
| ENSG00000180822.12 | PSMG4 | protein_coding | 812,7768108 | 0,584026823 | 0,190982953 | 3,058004998 | 0,002228159 | 0,01704293 |
| ENSG00000154305.18 | MIA3 | protein_coding | 5594,195462 | 0,581572869 | 0,14681403 | 3,961289443 | 7,45E-05 | 0,000919095 |
| ENSG00000179627.10 | ZBTB42 | protein_coding | 1737,709182 | 0,580468646 | 0,136350952 | 4,257166077 | 2,07E-05 | 0,000291259 |
| ENSG00000189134.4 | NKAPL | protein_coding | 628,1605471 | 0,58031292 | 0,211608388 | 2,74239091 | 0,006099371 | 0,038725762 |
| ENSG00000149418.11 | ST14 | protein_coding | 3101,759107 | 0,580090154 | 0,126647801 | 4,580341303 | 4,64E-06 | 7,55E-05 |
| ENSG00000169118.18 | CSNK1G1 | protein_coding | 3238,454938 | 0,579950531 | 0,131853495 | 4,398446406 | 1,09E-05 | 0,000164683 |
| ENSG00000109458.8 | GAB1 | protein_coding | 646,9840897 | 0,579760467 | 0,181469966 | 3,194801209 | 0,001399272 | 0,011583138 |
| ENSG00000157350.13 | ST3GAL2 | protein_coding | 2818,200938 | 0,57898199 | 0,136998775 | 4,226183698 | 2,38E-05 | 0,000328883 |
| ENSG00000166444.19 | DENND2B | protein_coding | 2575,148602 | 0,578893058 | 0,126921154 | 4,561044702 | 5,09E-06 | 8,20E-05 |
| ENSG00000156232.7 | WHAMM | protein_coding | 988,4997919 | 0,578252245 | 0,185727553 | 3,113443514 | 0,001849179 | 0,014597497 |
| ENSG00000152457.18 | DCLRE1C | protein_coding | 574,362162 | 0,577872009 | 0,182266228 | 3,170483176 | 0,001521856 | 0,012450201 |
| ENSG00000028116.18 | VRK2 | protein_coding | 1314,872238 | 0,576753678 | 0,156279839 | 3,690518763 | 0,000223797 | 0,002442777 |
| ENSG00000196739.15 | COL27A1 | protein_coding | 560,9819787 | 0,576454518 | 0,203912122 | 2,826975222 | 0,004698996 | 0,031355126 |
| ENSG00000198719.9 | DLL1 | protein_coding | 740,4108127 | 0,576231921 | 0,1999817 | 2,881423253 | 0,003958836 | 0,027443125 |
| ENSG00000151376.17 | ME3 | protein_coding | 585,4751876 | 0,576082394 | 0,177971509 | 3,236936051 | 0,001208205 | 0,01024454 |
| ENSG00000134109.11 | EDEM1 | protein_coding | 2042,840239 | 0,575176309 | 0,204619047 | 2,810961719 | 0,004939366 | 0,03262925 |
| ENSG00000286791.1 | AC079949.5 | lncRNA | 377,8135067 | 0,574676962 | 0,213819508 | 2,687673204 | 0,007195178 | 0,044057047 |
| ENSG00000068400.13 | GRIPAP1 | protein_coding | 6639,1874 | 0,574615131 | 0,151716997 | 3,787414344 | 0,000152223 | 0,001731011 |
| ENSG00000110328.6 | GALNT18 | protein_coding | 4354,388718 | 0,574167641 | 0,124458202 | 4,613337092 | 3,96E-06 | 6,53E-05 |
| ENSG00000197302.10 | ZNF720 | protein_coding | 1300,245885 | 0,574108333 | 0,157830621 | 3,637496509 | 0,000275301 | 0,002899107 |
| ENSG00000181789.14 | COPG1 | protein_coding | 13179,49332 | 0,572559735 | 0,181753769 | 3,150194563 | 0,001631618 | 0,013219007 |
| ENSG00000076108.12 | BAZ2A | protein_coding | 9862,047058 | 0,572526454 | 0,158487062 | 3,612449166 | 0,000303319 | 0,003154483 |
| ENSG00000160767.21 | FAM189B | protein_coding | 3291,836066 | 0,571362538 | 0,135863671 | 4,205410715 | 2,61E-05 | 0,00035823 |
| ENSG00000121753.12 | ADGRB2 | protein_coding | 2423,227351 | 0,57115423 | 0,147531408 | 3,871407708 | 0,000108209 | 0,001282756 |
| ENSG00000154429.11 | CCSAP | protein_coding | 4333,501544 | 0,570213408 | 0,125829877 | 4,531621748 | 5,85E-06 | 9,30E-05 |
| ENSG00000131174.7 | COX7B | protein_coding | 5468,749371 | 0,56936362 | 0,131860507 | 4,317923786 | 1,58E-05 | 0,00022709 |
| ENSG00000170561.13 | IRX2 | protein_coding | 2118,589425 | 0,569013698 | 0,177924666 | 3,198059661 | 0,001383557 | 0,011489392 |

| | | | | | | | | |
|---|---|---|---|---|---|---|---|---|
| ENSG00000181061.14 | HIGD1A | protein_coding | 28919,90391 | 0,568482879 | 0,13937768 | 4,078722482 | 4,53E-05 | 0,000586282 |
| ENSG00000257727.6 | CNPY2 | protein_coding | 11661,62484 | 0,568374547 | 0,119944236 | 4,738656631 | 2,15E-06 | 3,72E-05 |
| ENSG00000179218.14 | CALR | protein_coding | 150230,8182 | 0,56767013 | 0,113014477 | 5,022985946 | 5,09E-07 | 9,82E-06 |
| ENSG00000170017.12 | ALCAM | protein_coding | 3268,199163 | 0,567483379 | 0,153768032 | 3,690515972 | 0,0002238 | 0,002442777 |
| ENSG00000088727.13 | KIF9 | protein_coding | 1068,523545 | 0,567191698 | 0,176906837 | 3,20616041 | 0,00134519 | 0,011233166 |
| ENSG00000197077.13 | KIAA1671 | protein_coding | 3274,988767 | 0,566874149 | 0,167881574 | 3,376631137 | 0,000733794 | 0,006670419 |
| ENSG00000186212.4 | SOWAHB | protein_coding | 375,7227454 | 0,566760468 | 0,209478427 | 2,705579172 | 0,006818539 | 0,042295147 |
| ENSG00000135473.15 | PAN2 | protein_coding | 1323,220548 | 0,566500159 | 0,157022407 | 3,607766364 | 0,000308844 | 0,003196074 |
| ENSG00000169249.13 | ZRSR2 | protein_coding | 950,9814096 | 0,566177607 | 0,155525095 | 3,640426038 | 0,000272187 | 0,002872097 |
| ENSG00000108094.16 | CUL2 | protein_coding | 6573,676098 | 0,565364667 | 0,120247308 | 4,701682522 | 2,58E-06 | 4,37E-05 |
| ENSG00000138867.17 | GUCD1 | protein_coding | 6250,829766 | 0,565342983 | 0,156967253 | 3,601661956 | 0,000316189 | 0,003262407 |
| ENSG00000213626.13 | LBH | protein_coding | 3953,604664 | 0,5645727 | 0,152975968 | 3,690597329 | 0,000223728 | 0,002442777 |
| ENSG00000145012.13 | LPP | protein_coding | 2793,182091 | 0,564303427 | 0,201872337 | 2,795347962 | 0,005184388 | 0,03393845 |
| ENSG00000176153.12 | GPX2 | protein_coding | 751,9281876 | 0,563997115 | 0,171607066 | 3,286561132 | 0,001014188 | 0,008780818 |
| ENSG00000072786.13 | STK10 | protein_coding | 844,5267772 | 0,563903307 | 0,170362968 | 3,310011047 | 0,000932923 | 0,008186264 |
| ENSG00000175832.13 | ETV4 | protein_coding | 1763,158285 | 0,562672064 | 0,174144419 | 3,231065731 | 0,001233296 | 0,010450631 |
| ENSG00000165655.17 | ZNF503 | protein_coding | 1435,310403 | 0,562456368 | 0,195653431 | 2,874758526 | 0,004043366 | 0,027918162 |
| ENSG00000215252.11 | GOLGA8B | protein_coding | 465,4025024 | 0,561687541 | 0,196634185 | 2,856510128 | 0,004283263 | 0,029218025 |
| ENSG00000116117.18 | PARD3B | protein_coding | 747,9287594 | 0,56055829 | 0,178703645 | 3,136803903 | 0,001708003 | 0,013726314 |
| ENSG00000142484.7 | TM4SF5 | protein_coding | 3158,461484 | 0,558805454 | 0,157888757 | 3,539235253 | 0,000401288 | 0,004019601 |
| ENSG00000198382.9 | UVRAG | protein_coding | 3317,274043 | 0,558776636 | 0,128050206 | 4,363730874 | 1,28E-05 | 0,000189442 |
| ENSG00000172201.12 | ID4 | protein_coding | 10197,73512 | 0,558539829 | 0,116704499 | 4,785932266 | 1,70E-06 | 3,00E-05 |
| ENSG00000144834.14 | TAGLN3 | protein_coding | 2556,489459 | 0,558520322 | 0,144310227 | 3,870275405 | 0,000108712 | 0,001287271 |
| ENSG00000154310.17 | TNIK | protein_coding | 2085,146556 | 0,557753583 | 0,155414719 | 3,588807968 | 0,000332193 | 0,003402382 |
| ENSG00000184221.13 | OLIG1 | protein_coding | 551,7804395 | 0,557421067 | 0,185014094 | 3,012857309 | 0,002588006 | 0,019280194 |
| ENSG00000113916.18 | BCL6 | protein_coding | 1507,265034 | 0,557257549 | 0,143405516 | 3,885886441 | 0,000101957 | 0,001213458 |
| ENSG00000135637.14 | CCDC142 | protein_coding | 854,8434027 | 0,556225919 | 0,176294379 | 3,155097294 | 0,001604446 | 0,013034177 |
| ENSG00000104112.9 | SCG3 | protein_coding | 72173,23361 | 0,556219479 | 0,121223754 | 4,588370368 | 4,47E-06 | 7,30E-05 |
| ENSG00000117707.16 | PROX1 | protein_coding | 6357,935995 | 0,555808136 | 0,152488477 | 3,644918938 | 0,000267476 | 0,002841001 |
| ENSG00000188811.14 | NHLRC3 | protein_coding | 3330,60025 | 0,55483569 | 0,127336061 | 4,357255032 | 1,32E-05 | 0,000194444 |
| ENSG00000108039.18 | XPNPEP1 | protein_coding | 3049,3632 | 0,553033076 | 0,133146209 | 4,153577329 | 3,27E-05 | 0,000438651 |
| ENSG00000184205.14 | TSPYL2 | protein_coding | 676,6203687 | 0,552819471 | 0,179494739 | 3,07986448 | 0,002070948 | 0,01601008 |
| ENSG00000106346.11 | USP42 | protein_coding | 1267,984862 | 0,552599344 | 0,208077539 | 2,655737601 | 0,007913516 | 0,047510601 |
| ENSG00000050030.16 | NEXMIF | protein_coding | 1746,990526 | 0,552377542 | 0,186219623 | 2,966269248 | 0,003014365 | 0,021798178 |
| ENSG00000153214.11 | TMEM87B | protein_coding | 3781,593427 | 0,552129974 | 0,133792635 | 4,126759098 | 3,68E-05 | 0,000486966 |
| ENSG00000133424.22 | LARGE1 | protein_coding | 4594,091419 | 0,551568677 | 0,139837152 | 3,944364342 | 8,00E-05 | 0,000980134 |

| | | | | | | | | |
|---|---|---|---|---|---|---|---|---|
| ENSG00000112218.9 | GPR63 | protein_coding | 551,457467 | 0,551505301 | 0,187431 | 2,942444431 | 0,003256322 | 0,023290436 |
| ENSG00000076053.12 | RBM7 | protein_coding | 3175,376648 | 0,55137841 | 0,133063177 | 4,14373399 | 3,42E-05 | 0,000455341 |
| ENSG00000104205.16 | SGK3 | protein_coding | 3583,639264 | 0,551000615 | 0,137043297 | 4,020631625 | 5,80E-05 | 0,000731997 |
| ENSG00000133121.21 | STARD13 | protein_coding | 1936,374442 | 0,550057081 | 0,150823004 | 3,647037037 | 0,000265282 | 0,002820555 |
| ENSG00000005238.20 | FAM214B | protein_coding | 408,7438839 | 0,549817504 | 0,208083561 | 2,642291886 | 0,008234706 | 0,04917101 |
| ENSG00000135318.12 | NT5E | protein_coding | 4137,456518 | 0,549656309 | 0,126552437 | 4,343308758 | 1,40E-05 | 0,000205762 |
| ENSG00000130150.12 | MOSPD2 | protein_coding | 7209,886782 | 0,549608767 | 0,118329615 | 4,644727079 | 3,41E-06 | 5,67E-05 |
| ENSG00000183655.13 | KLHL25 | protein_coding | 1663,91483 | 0,549550783 | 0,136378552 | 4,029598319 | 5,59E-05 | 0,0007089 |
| ENSG00000126010.6 | GRPR | protein_coding | 579,7947954 | 0,549417065 | 0,198247566 | 2,771368527 | 0,005582121 | 0,035990669 |
| ENSG00000128881.17 | TTBK2 | protein_coding | 3074,735891 | 0,54897797 | 0,133657263 | 4,107356063 | 4,00E-05 | 0,000524639 |
| ENSG00000133739.16 | LRRCC1 | protein_coding | 1360,479405 | 0,548823501 | 0,144121916 | 3,808050257 | 0,000140067 | 0,001608502 |
| ENSG00000130787.14 | HIP1R | protein_coding | 6714,152118 | 0,54762065 | 0,13204024 | 4,14737696 | 3,36E-05 | 0,000449016 |
| ENSG00000183780.13 | SLC35F3 | protein_coding | 1701,33931 | 0,547213787 | 0,157373545 | 3,477165024 | 0,000506746 | 0,004897841 |
| ENSG00000173230.15 | GOLGB1 | protein_coding | 2593,052039 | 0,547190197 | 0,197748337 | 2,767103911 | 0,005655673 | 0,03635788 |
| ENSG00000237298.10 | TTN-AS1 | lncRNA | 724,3325833 | 0,546637837 | 0,16859183 | 3,242374411 | 0,001185382 | 0,010060111 |
| ENSG00000123191.16 | ATP7B | protein_coding | 1211,228177 | 0,545928 | 0,148351773 | 3,67995601 | 0,000233274 | 0,002525624 |
| ENSG00000163347.6 | CLDN1 | protein_coding | 1343,144705 | 0,545651167 | 0,162313815 | 3,361705027 | 0,000774628 | 0,006993033 |
| ENSG00000116044.16 | NFE2L2 | protein_coding | 6199,806678 | 0,545031564 | 0,122211211 | 4,459750952 | 8,21E-06 | 0,000126776 |
| ENSG00000270136.6 | MICOS10-NBL1 | protein_coding | 749,6311015 | 0,5449001 | 0,183467202 | 2,97001368 | 0,002977865 | 0,021586427 |
| ENSG00000105855.10 | ITGB8 | protein_coding | 1611,131493 | 0,544809779 | 0,185743119 | 2,933135736 | 0,003355572 | 0,023877937 |
| ENSG00000166199.13 | ALKBH3 | protein_coding | 2402,587819 | 0,544715971 | 0,148947812 | 3,657092805 | 0,000255092 | 0,002720505 |
| ENSG00000107897.20 | ACBD5 | protein_coding | 2216,409459 | 0,543955368 | 0,136125692 | 3,995978722 | 6,44E-05 | 0,000805248 |
| ENSG00000169359.16 | SLC33A1 | protein_coding | 5808,849756 | 0,543704531 | 0,139438123 | 3,89925308 | 9,65E-05 | 0,001159596 |
| ENSG00000178538.10 | CA8 | protein_coding | 6459,364498 | 0,543113797 | 0,12989024 | 4,18132877 | 2,90E-05 | 0,000393889 |
| ENSG00000255277.3 | ABCC6P2 | transcribed_unproc | 811,4032466 | 0,542893245 | 0,180202896 | 3,012677695 | 0,002589538 | 0,019280194 |
| ENSG00000147421.18 | HMBOX1 | protein_coding | 436,119573 | 0,542749133 | 0,196023707 | 2,768793333 | 0,005626431 | 0,036231725 |
| ENSG00000091490.11 | SEL1L3 | protein_coding | 4693,414212 | 0,542630717 | 0,170729756 | 3,178301953 | 0,001481404 | 0,012162035 |
| ENSG00000126067.12 | PSMB2 | protein_coding | 9585,859915 | 0,542466076 | 0,114019747 | 4,757650216 | 1,96E-06 | 3,42E-05 |
| ENSG00000198420.10 | TCAF1 | protein_coding | 22622,79901 | 0,542244287 | 0,112296058 | 4,828702764 | 1,37E-06 | 2,47E-05 |
| ENSG00000227036.8 | LINC00511 | lncRNA | 5194,262848 | 0,541819446 | 0,130000023 | 4,167841165 | 3,07E-05 | 0,000415321 |
| ENSG00000181284.3 | TMEM102 | protein_coding | 1243,586017 | 0,541794185 | 0,1479257 | 3,662610258 | 0,000249658 | 0,00267345 |
| ENSG00000149571.12 | KIRREL3 | protein_coding | 600,0473263 | 0,541599399 | 0,20169198 | 2,685279797 | 0,007246911 | 0,04430902 |
| ENSG00000179222.18 | MAGED1 | protein_coding | 23644,54655 | 0,541304673 | 0,139152494 | 3,89001057 | 0,00010024 | 0,001195058 |
| ENSG00000107290.14 | SETX | protein_coding | 11559,70813 | 0,541272511 | 0,116560987 | 4,643685046 | 3,42E-06 | 5,70E-05 |
| ENSG00000008735.14 | MAPK8IP2 | protein_coding | 10046,79118 | 0,540589603 | 0,126987406 | 4,257033178 | 2,07E-05 | 0,000291259 |
| ENSG00000066405.13 | CLDN18 | protein_coding | 53566,71925 | 0,539326532 | 0,11665525 | 4,623251263 | 3,78E-06 | 6,26E-05 |

| | | | | | | | | |
|---|---|---|---|---|---|---|---|---|
| ENSG00000148429.14 | USP6NL | protein_coding | 2007,291173 | 0,538407248 | 0,131825477 | 4,084242738 | 4,42E-05 | 0,000575008 |
| ENSG00000145335.17 | SNCA | protein_coding | 2746,639814 | 0,536858366 | 0,14862904 | 3,612069122 | 0,000303764 | 0,003155975 |
| ENSG00000116525.14 | TRIM62 | protein_coding | 692,4228111 | 0,536201631 | 0,193476228 | 2,77140833 | 0,005581439 | 0,035990669 |
| ENSG00000182103.5 | FAM181B | protein_coding | 1948,287892 | 0,536149873 | 0,131735313 | 4,069902442 | 4,70E-05 | 0,000605928 |
| ENSG00000132432.14 | SEC61G | protein_coding | 6518,191185 | 0,535707961 | 0,15684309 | 3,415566225 | 0,000636496 | 0,005927506 |
| ENSG00000080503.24 | SMARCA2 | protein_coding | 20818,24493 | 0,535681969 | 0,12516767 | 4,279715114 | 1,87E-05 | 0,000265422 |
| ENSG00000123983.14 | ACSL3 | protein_coding | 10378,24566 | 0,535470583 | 0,119359476 | 4,48620087 | 7,25E-06 | 0,000113359 |
| ENSG00000152492.15 | CCDC50 | protein_coding | 2701,119835 | 0,535206841 | 0,147859993 | 3,619686635 | 0,00029496 | 0,003076725 |
| ENSG00000180867.11 | PDIA3P1 | transcribed_process | 779,5574165 | 0,535090188 | 0,18030437 | 2,967705045 | 0,003000321 | 0,021723747 |
| ENSG00000114439.19 | BBX | protein_coding | 4504,274375 | 0,535062753 | 0,12703822 | 4,211825033 | 2,53E-05 | 0,000348434 |
| ENSG00000164970.15 | FAM219A | protein_coding | 2197,546406 | 0,534708163 | 0,145554261 | 3,673600201 | 0,000239157 | 0,002575488 |
| ENSG00000116260.17 | QSOX1 | protein_coding | 28099,30401 | 0,534248353 | 0,158256869 | 3,375830421 | 0,000735933 | 0,006684056 |
| ENSG00000107864.15 | CPEB3 | protein_coding | 1112,317286 | 0,534062947 | 0,176533542 | 3,025277458 | 0,002484051 | 0,01861031 |
| ENSG00000109171.15 | SLAIN2 | protein_coding | 3955,045562 | 0,532582461 | 0,17080822 | 3,118014229 | 0,00182074 | 0,014416488 |
| ENSG00000172380.6 | GNG12 | protein_coding | 10469,73179 | 0,53249257 | 0,113141998 | 4,706409459 | 2,52E-06 | 4,28E-05 |
| ENSG00000175482.9 | POLD4 | protein_coding | 869,4569145 | 0,532062069 | 0,157390157 | 3,380529496 | 0,000723463 | 0,006585087 |
| ENSG00000121039.10 | RDH10 | protein_coding | 6246,075685 | 0,531033063 | 0,154313874 | 3,441252884 | 0,000579027 | 0,005477602 |
| ENSG00000179094.16 | PER1 | protein_coding | 2132,670749 | 0,529974724 | 0,157692735 | 3,360806214 | 0,000777153 | 0,007009782 |
| ENSG00000198740.9 | ZNF652 | protein_coding | 3257,527204 | 0,5299526 | 0,128199838 | 4,133800871 | 3,57E-05 | 0,000473977 |
| ENSG00000130958.13 | SLC35D2 | protein_coding | 5425,013923 | 0,529814372 | 0,130591452 | 4,057037162 | 4,97E-05 | 0,000637142 |
| ENSG00000166689.16 | PLEKHA7 | protein_coding | 2000,869623 | 0,52957718 | 0,130840787 | 4,047493087 | 5,18E-05 | 0,000660845 |
| ENSG00000140400.17 | MAN2C1 | protein_coding | 1699,375281 | 0,528724508 | 0,141099633 | 3,747171389 | 0,00017884 | 0,002003217 |
| ENSG00000100994.12 | PYGB | protein_coding | 8319,324394 | 0,528078328 | 0,116003061 | 4,552279268 | 5,31E-06 | 8,51E-05 |
| ENSG00000185164.14 | NOMO2 | protein_coding | 45583,73438 | 0,527796805 | 0,113601209 | 4,646049181 | 3,38E-06 | 5,64E-05 |
| ENSG00000231312.7 | MAP4K3-DT | lncRNA | 726,7989434 | 0,527529546 | 0,182796748 | 2,885880365 | 0,003903205 | 0,027156396 |
| ENSG00000042493.16 | CAPG | protein_coding | 1115,471267 | 0,527282238 | 0,149127199 | 3,535788516 | 0,00040656 | 0,004066571 |
| ENSG00000123094.15 | RASSF8 | protein_coding | 1623,362278 | 0,526681442 | 0,143691888 | 3,665352641 | 0,000246998 | 0,002650386 |
| ENSG00000173540.14 | GMPPB | protein_coding | 7651,516572 | 0,526314791 | 0,124949841 | 4,212208572 | 2,53E-05 | 0,000348144 |
| ENSG00000116871.15 | MAP7D1 | protein_coding | 4945,954371 | 0,524475315 | 0,151599393 | 3,459613552 | 0,000540951 | 0,005166429 |
| ENSG00000065518.8 | NDUFB4 | protein_coding | 11591,69909 | 0,523614593 | 0,143757859 | 3,64233716 | 0,000270174 | 0,0028595 |
| ENSG00000160216.21 | AGPAT3 | protein_coding | 6413,61351 | 0,52338322 | 0,114460522 | 4,57260905 | 4,82E-06 | 7,79E-05 |
| ENSG00000177556.12 | ATOX1 | protein_coding | 1500,362158 | 0,523038427 | 0,148421807 | 3,523999867 | 0,000425084 | 0,004221605 |
| ENSG00000120008.16 | WDR11 | protein_coding | 3780,574976 | 0,522486577 | 0,12033002 | 4,34211328 | 1,41E-05 | 0,000206718 |
| ENSG00000107281.10 | NPDC1 | protein_coding | 5388,938202 | 0,522362418 | 0,120043722 | 4,351434713 | 1,35E-05 | 0,000199258 |
| ENSG00000176386.9 | CDC26 | protein_coding | 3029,382345 | 0,521588956 | 0,139879451 | 3,728846169 | 0,000192359 | 0,002132959 |
| ENSG00000143156.14 | NME7 | protein_coding | 1473,899957 | 0,521481254 | 0,14309132 | 3,644394733 | 0,000268022 | 0,002843911 |

| | | | | | | | | |
|---|---|---|---|---|---|---|---|---|
| ENSG00000136717.15 | BIN1 | protein_coding | 2895,240245 | 0,520054745 | 0,127913793 | 4,065665898 | 4,79E-05 | 0,000616283 |
| ENSG00000164733.22 | CTSB | protein_coding | 14050,28114 | 0,520004965 | 0,110205544 | 4,718500964 | 2,38E-06 | 4,06E-05 |
| ENSG00000169436.17 | COL22A1 | protein_coding | 7813,998252 | 0,519833434 | 0,122432485 | 4,245878311 | 2,18E-05 | 0,000303891 |
| ENSG00000272501.1 | AL662844.4 | lncRNA | 632,1291369 | 0,519678227 | 0,191784484 | 2,709699016 | 0,006734429 | 0,041872667 |
| ENSG00000114861.23 | FOXP1 | protein_coding | 3549,846728 | 0,519617725 | 0,146976683 | 3,535375239 | 0,000407196 | 0,004070992 |
| ENSG00000121879.6 | PIK3CA | protein_coding | 2548,042127 | 0,519208229 | 0,133091829 | 3,901127755 | 9,57E-05 | 0,001152636 |
| ENSG00000165288.11 | BRWD3 | protein_coding | 2487,453007 | 0,518801249 | 0,168144526 | 3,085448333 | 0,002032455 | 0,015770738 |
| ENSG00000101298.15 | SNPH | protein_coding | 2802,528587 | 0,518113214 | 0,176243311 | 2,939761013 | 0,003284655 | 0,023453018 |
| ENSG00000107954.10 | NEURL1 | protein_coding | 6058,071498 | 0,516647501 | 0,116281424 | 4,44307855 | 8,87E-06 | 0,000136409 |
| ENSG00000170315.14 | UBB | protein_coding | 134241,004 | 0,516283455 | 0,11784179 | 4,381157618 | 1,18E-05 | 0,00017716 |
| ENSG00000136986.10 | DERL1 | protein_coding | 7751,668381 | 0,515901466 | 0,120413369 | 4,284420175 | 1,83E-05 | 0,00026059 |
| ENSG00000163795.14 | ZNF513 | protein_coding | 1342,478339 | 0,515709967 | 0,143120699 | 3,603322032 | 0,000314176 | 0,003246432 |
| ENSG00000169223.15 | LMAN2 | protein_coding | 21255,59906 | 0,515638807 | 0,123215917 | 4,184839264 | 2,85E-05 | 0,000388691 |
| ENSG00000168672.4 | LRATD2 | protein_coding | 3753,116792 | 0,514736255 | 0,156280015 | 3,293679327 | 0,000988852 | 0,008593452 |
| ENSG00000283103.3 | AC010642.2 | lncRNA | 1340,997877 | 0,514220096 | 0,151849026 | 3,386390481 | 0,000708185 | 0,006460905 |
| ENSG00000104537.17 | ANXA13 | protein_coding | 650,8166294 | 0,513020495 | 0,19359208 | 2,650007659 | 0,008048995 | 0,048130735 |
| ENSG00000184515.11 | BEX5 | protein_coding | 1068,589445 | 0,511859657 | 0,170151172 | 3,00826406 | 0,002627447 | 0,019491711 |
| ENSG00000166925.9 | TSC22D4 | protein_coding | 3773,828081 | 0,511828805 | 0,128200297 | 3,992415122 | 6,54E-05 | 0,000815502 |
| ENSG00000137275.15 | RIPK1 | protein_coding | 4587,66325 | 0,511757937 | 0,12306777 | 4,158342479 | 3,21E-05 | 0,0004313 |
| ENSG00000156858.12 | PRR14 | protein_coding | 1453,318763 | 0,510428716 | 0,166414184 | 3,067218811 | 0,002160606 | 0,016605839 |
| ENSG00000118058.23 | KMT2A | protein_coding | 2953,417684 | 0,509857724 | 0,173758858 | 2,934283354 | 0,003343189 | 0,023806004 |
| ENSG00000079691.18 | CARMIL1 | protein_coding | 1311,690525 | 0,509738952 | 0,180065931 | 2,83084617 | 0,004642504 | 0,031071237 |
| ENSG00000163755.9 | HPS3 | protein_coding | 3805,92826 | 0,509406151 | 0,121722289 | 4,18498662 | 2,85E-05 | 0,000388691 |
| ENSG00000165929.13 | TC2N | protein_coding | 3585,016535 | 0,507902056 | 0,158854533 | 3,197277703 | 0,001387313 | 0,011511453 |
| ENSG00000225127.3 | LINC00237 | lncRNA | 573,6566084 | 0,50755186 | 0,187540079 | 2,706364747 | 0,006802429 | 0,042207718 |
| ENSG00000110330.8 | BIRC2 | protein_coding | 4252,619452 | 0,507458431 | 0,140919436 | 3,601053522 | 0,00031693 | 0,003267579 |
| ENSG00000133961.20 | NUMB | protein_coding | 4498,784228 | 0,507288094 | 0,119737561 | 4,236666342 | 2,27E-05 | 0,000314945 |
| ENSG00000109814.12 | UGDH | protein_coding | 5138,053237 | 0,506679359 | 0,126445396 | 4,00710011 | 6,15E-05 | 0,000772419 |
| ENSG00000047579.20 | DTNBP1 | protein_coding | 1902,35294 | 0,506616329 | 0,134298302 | 3,772321169 | 0,000161736 | 0,001829249 |
| ENSG00000101966.13 | XIAP | protein_coding | 5763,635233 | 0,506313093 | 0,168247953 | 3,009326904 | 0,002618272 | 0,019444317 |
| ENSG00000164938.14 | TP53INP1 | protein_coding | 8137,927928 | 0,506279567 | 0,137178739 | 3,690656239 | 0,000223676 | 0,002442777 |
| ENSG00000105699.16 | LSR | protein_coding | 7906,677871 | 0,506208262 | 0,11672589 | 4,336726512 | 1,45E-05 | 0,000211132 |
| ENSG00000099812.9 | MISP | protein_coding | 4798,06636 | 0,50561362 | 0,139474503 | 3,625132978 | 0,000288813 | 0,003020128 |
| ENSG00000181788.4 | SIAH2 | protein_coding | 2552,043476 | 0,505486227 | 0,131538359 | 3,842880731 | 0,000121599 | 0,00142216 |
| ENSG00000102871.16 | TRADD | protein_coding | 588,4089714 | 0,505137399 | 0,176884917 | 2,855740382 | 0,004293661 | 0,029260347 |
| ENSG00000143514.17 | TP53BP2 | protein_coding | 4254,967515 | 0,505055944 | 0,152014204 | 3,322425995 | 0,000892383 | 0,007879395 |

| | | | | | | | | |
|---|---|---|---|---|---|---|---|---|
| ENSG00000091262.16 | ABCC6 | protein_coding | 4273,816843 | 0,504343555 | 0,143723088 | 3,509133867 | 0,000449569 | 0,00443101 |
| ENSG00000183145.9 | RIPPLY3 | protein_coding | 831,5768235 | 0,504278652 | 0,175671991 | 2,870569452 | 0,004097332 | 0,028253505 |
| ENSG00000145349.17 | CAMK2D | protein_coding | 6467,894939 | 0,503979338 | 0,131698473 | 3,82676677 | 0,000129837 | 0,001505898 |
| ENSG00000113811.11 | SELENOK | protein_coding | 9994,326969 | 0,503625497 | 0,127752902 | 3,942184392 | 8,07E-05 | 0,000986779 |
| ENSG00000184305.15 | CCSER1 | protein_coding | 4155,175432 | 0,503472998 | 0,147634224 | 3,410272935 | 0,000648979 | 0,006006355 |
| ENSG00000099917.17 | MED15 | protein_coding | 1782,515082 | 0,503038938 | 0,178123687 | 2,824099064 | 0,004741373 | 0,031581495 |
| ENSG00000165175.16 | MID1IP1 | protein_coding | 8036,695671 | 0,501911909 | 0,124616808 | 4,027642158 | 5,63E-05 | 0,000713957 |
| ENSG00000129128.13 | SPCS3 | protein_coding | 10701,92468 | 0,501758353 | 0,132537954 | 3,785771072 | 0,000153233 | 0,001739656 |
| ENSG00000277147.6 | LINC00869 | transcribed_unproc | 1123,170479 | 0,501229809 | 0,148564561 | 3,373818136 | 0,000741333 | 0,006730187 |
| ENSG00000277791.5 | PSMB3 | protein_coding | 9508,639365 | 0,50118471 | 0,129636394 | 3,866080317 | 0,000110599 | 0,001308864 |
| ENSG00000167978.17 | SRRM2 | protein_coding | 10557,33524 | 0,501068204 | 0,11789471 | 4,250133051 | 2,14E-05 | 0,000299172 |
| ENSG00000154545.16 | MAGED4 | protein_coding | 1461,825232 | 0,500643939 | 0,163353121 | 3,064795671 | 0,002178188 | 0,016709548 |
| ENSG00000281649.2 | EBLN3P | lncRNA | 9001,751478 | 0,499506497 | 0,119792034 | 4,169780575 | 3,05E-05 | 0,000412068 |
| ENSG00000158158.12 | CNNM4 | protein_coding | 1934,544609 | 0,498937076 | 0,149104138 | 3,346232262 | 0,000819177 | 0,007328838 |
| ENSG00000144591.19 | GMPPA | protein_coding | 4690,31859 | 0,498795809 | 0,161744258 | 3,083854809 | 0,002043373 | 0,015837842 |
| ENSG00000138600.10 | SPPL2A | protein_coding | 5785,825996 | 0,49870569 | 0,120980146 | 4,122211019 | 3,75E-05 | 0,000494658 |
| ENSG00000112139.16 | MDGA1 | protein_coding | 987,3170488 | 0,498371517 | 0,167960552 | 2,967193841 | 0,003005314 | 0,021744722 |
| ENSG00000100888.15 | CHD8 | protein_coding | 8217,497393 | 0,497712874 | 0,161627224 | 3,079387624 | 0,002074266 | 0,016029812 |
| ENSG00000243279.4 | PRAF2 | protein_coding | 7628,134552 | 0,497390754 | 0,140669247 | 3,535888377 | 0,000406406 | 0,004066571 |
| ENSG00000131724.11 | IL13RA1 | protein_coding | 4138,613523 | 0,496477895 | 0,126376288 | 3,928568431 | 8,55E-05 | 0,001041303 |
| ENSG00000166801.16 | FAM111A | protein_coding | 3897,147369 | 0,496357538 | 0,148525967 | 3,341890642 | 0,000832098 | 0,007422232 |
| ENSG00000023330.15 | ALAS1 | protein_coding | 6448,648977 | 0,494918817 | 0,128253761 | 3,858902951 | 0,000113897 | 0,001341833 |
| ENSG00000278934.1 | AC117489.1 | TEC | 492,4061946 | 0,49488631 | 0,187819877 | 2,634898492 | 0,008416245 | 0,049885075 |
| ENSG00000130158.14 | DOCK6 | protein_coding | 1291,720056 | 0,49467648 | 0,16316655 | 3,031727271 | 0,002431588 | 0,018291518 |
| ENSG00000131094.4 | C1QL1 | protein_coding | 20467,00109 | 0,494551312 | 0,128751689 | 3,841124866 | 0,000122472 | 0,001430774 |
| ENSG00000074657.14 | ZNF532 | protein_coding | 546,5837406 | 0,494384637 | 0,180477673 | 2,739311899 | 0,006156793 | 0,038975649 |
| ENSG00000110492.15 | MDK | protein_coding | 10220,5976 | 0,494153712 | 0,128086928 | 3,85795583 | 0,000114339 | 0,001344772 |
| ENSG00000158411.11 | MITD1 | protein_coding | 1061,684321 | 0,494101542 | 0,174289813 | 2,834942171 | 0,004583398 | 0,030813565 |
| ENSG00000145476.16 | CYP4V2 | protein_coding | 2524,082726 | 0,493889348 | 0,133845532 | 3,689995019 | 0,000224258 | 0,002443962 |
| ENSG00000124766.7 | SOX4 | protein_coding | 14757,06592 | 0,493306101 | 0,137004765 | 3,600649237 | 0,000317424 | 0,003270306 |
| ENSG00000157483.9 | MYO1E | protein_coding | 4204,567472 | 0,492000528 | 0,157476116 | 3,124286652 | 0,001782367 | 0,014171614 |
| ENSG00000069702.11 | TGFBR3 | protein_coding | 10317,6305 | 0,491706158 | 0,125217776 | 3,926807942 | 8,61E-05 | 0,001047123 |
| ENSG00000150672.17 | DLG2 | protein_coding | 2247,180411 | 0,491172961 | 0,145773493 | 3,36942574 | 0,00075325 | 0,006823577 |
| ENSG00000131871.15 | SELENOS | protein_coding | 4630,538228 | 0,491029667 | 0,162994815 | 3,012547773 | 0,002590647 | 0,019280194 |
| ENSG00000118640.11 | VAMP8 | protein_coding | 3152,083406 | 0,490972952 | 0,160615064 | 3,05683003 | 0,002236911 | 0,017086044 |
| ENSG00000167861.16 | HID1 | protein_coding | 6915,41782 | 0,490813991 | 0,161178193 | 3,045163754 | 0,002325537 | 0,017626759 |

| | | | | | | | | |
|---|---|---|---|---|---|---|---|---|
| ENSG00000133639.6 | BTG1 | protein_coding | 33779,59015 | 0,490047371 | 0,111034768 | 4,413458773 | 1,02E-05 | 0,000154555 |
| ENSG00000065413.20 | ANKRD44 | protein_coding | 949,5238508 | 0,489418318 | 0,1544482 | 3,168818528 | 0,001530599 | 0,012516834 |
| ENSG00000198001.14 | IRAK4 | protein_coding | 1191,25525 | 0,489153778 | 0,17110226 | 2,858838794 | 0,004251948 | 0,029080213 |
| ENSG00000163093.12 | BBS5 | protein_coding | 920,7551199 | 0,488996323 | 0,158081273 | 3,093322276 | 0,001979291 | 0,01545304 |
| ENSG00000030110.13 | BAK1 | protein_coding | 4250,71606 | 0,488742177 | 0,138259186 | 3,534970734 | 0,00040782 | 0,004075283 |
| ENSG00000185475.11 | TMEM179B | protein_coding | 4437,398037 | 0,48836271 | 0,135431481 | 3,605976293 | 0,000310982 | 0,0032166 |
| ENSG00000123636.18 | BAZ2B | protein_coding | 3155,649692 | 0,486570723 | 0,182188161 | 2,670704391 | 0,007569227 | 0,045891044 |
| ENSG00000142396.10 | ERVK3-1 | protein_coding | 4551,850075 | 0,485880966 | 0,12660624 | 3,837733154 | 0,000124175 | 0,001449866 |
| ENSG00000174749.6 | FAM241A | protein_coding | 2814,490905 | 0,485822329 | 0,130401693 | 3,725582984 | 0,000194864 | 0,002156176 |
| ENSG00000110047.18 | EHD1 | protein_coding | 11905,55162 | 0,485443156 | 0,111404874 | 4,35746783 | 1,32E-05 | 0,000194392 |
| ENSG00000132330.18 | SCLY | protein_coding | 2540,331341 | 0,485085743 | 0,133329949 | 3,638235413 | 0,000274512 | 0,002892259 |
| ENSG00000178694.10 | NSUN3 | protein_coding | 1871,593935 | 0,484939637 | 0,139821768 | 3,468269961 | 0,000523821 | 0,005025751 |
| ENSG00000163872.16 | YEATS2 | protein_coding | 5372,440575 | 0,48449201 | 0,13217189 | 3,665620646 | 0,000246739 | 0,002650329 |
| ENSG00000113621.15 | TXNDC15 | protein_coding | 3805,925471 | 0,484031234 | 0,153511107 | 3,153069779 | 0,001615632 | 0,013109788 |
| ENSG00000250320.6 | EDIL3-DT | lncRNA | 546,7062522 | 0,48347505 | 0,182251323 | 2,652793095 | 0,007982879 | 0,047844709 |
| ENSG00000176887.7 | SOX11 | protein_coding | 4620,681131 | 0,483473804 | 0,16745836 | 2,887128512 | 0,003887754 | 0,027075324 |
| ENSG00000137094.15 | DNAJB5 | protein_coding | 920,1244673 | 0,482922083 | 0,155296188 | 3,109684077 | 0,001872875 | 0,014745931 |
| ENSG00000172831.14 | CES2 | protein_coding | 3982,384558 | 0,48242965 | 0,126810038 | 3,804349076 | 0,000142178 | 0,001630059 |
| ENSG00000100379.17 | KCTD17 | protein_coding | 2547,340207 | 0,48235286 | 0,130735398 | 3,689535244 | 0,000224664 | 0,002445836 |
| ENSG00000103966.11 | EHD4 | protein_coding | 2676,08459 | 0,482341889 | 0,15195556 | 3,174229937 | 0,001502346 | 0,012314651 |
| ENSG00000164096.13 | C4orf3 | protein_coding | 6597,683852 | 0,48165673 | 0,117961274 | 4,083176735 | 4,44E-05 | 0,000576936 |
| ENSG00000120688.9 | WBP4 | protein_coding | 1629,849424 | 0,481301978 | 0,143248717 | 3,359904296 | 0,000779695 | 0,007025154 |
| ENSG00000160584.16 | SIK3 | protein_coding | 3481,658605 | 0,481103092 | 0,150293327 | 3,201094163 | 0,001369068 | 0,011391667 |
| ENSG00000159377.11 | PSMB4 | protein_coding | 18022,44413 | 0,480724276 | 0,117888383 | 4,077791754 | 4,55E-05 | 0,000588269 |
| ENSG00000177873.14 | ZNF619 | protein_coding | 619,9235298 | 0,479477061 | 0,177391385 | 2,702933188 | 0,006873056 | 0,042535815 |
| ENSG00000112249.14 | ASCC3 | protein_coding | 10477,85124 | 0,478633445 | 0,117567017 | 4,071154125 | 4,68E-05 | 0,000603052 |
| ENSG00000123815.13 | COQ8B | protein_coding | 5537,558209 | 0,478401159 | 0,134183226 | 3,565282887 | 0,000363464 | 0,003681237 |
| ENSG00000011376.12 | LARS2 | protein_coding | 6592,99848 | 0,47825284 | 0,116400712 | 4,108676235 | 3,98E-05 | 0,000522303 |
| ENSG00000004961.15 | HCCS | protein_coding | 5712,944964 | 0,477768103 | 0,116243202 | 4,110073481 | 3,96E-05 | 0,000519479 |
| ENSG00000137478.15 | FCHSD2 | protein_coding | 1791,211724 | 0,476922711 | 0,137482712 | 3,46896496 | 0,000522468 | 0,005019669 |
| ENSG00000119242.9 | CCDC92 | protein_coding | 850,9347936 | 0,476840621 | 0,174789859 | 2,728079438 | 0,006370426 | 0,040085626 |
| ENSG00000166589.13 | CDH16 | protein_coding | 4913,415841 | 0,476804074 | 0,150260825 | 3,173176204 | 0,00150781 | 0,012349765 |
| ENSG00000182118.8 | FAM89A | protein_coding | 883,1353007 | 0,475466036 | 0,178588369 | 2,662357233 | 0,007759548 | 0,046792079 |
| ENSG00000186480.13 | INSIG1 | protein_coding | 23623,85891 | 0,475419092 | 0,11111856 | 4,278485005 | 1,88E-05 | 0,000266531 |
| ENSG00000132780.17 | NASP | protein_coding | 17849,06913 | 0,4751442 | 0,128731739 | 3,690963879 | 0,000223406 | 0,002442777 |
| ENSG00000145321.13 | GC | protein_coding | 19860,9522 | 0,474309178 | 0,142920708 | 3,318687571 | 0,000904416 | 0,007965478 |

| | | | | | | | | |
|---|---|---|---|---|---|---|---|---|
| ENSG00000105404.11 | RABAC1 | protein_coding | 5541,623156 | 0,472869418 | 0,135325529 | 3,494310514 | 0,000475288 | 0,004636604 |
| ENSG00000275964.1 | AL355001.2 | lncRNA | 662,5837114 | 0,472441682 | 0,173844827 | 2,717605639 | 0,006575616 | 0,041055927 |
| ENSG00000163812.15 | ZDHHC3 | protein_coding | 8808,459549 | 0,472303319 | 0,136469363 | 3,460874368 | 0,000538424 | 0,00514464 |
| ENSG00000127952.17 | STYXL1 | protein_coding | 950,8360532 | 0,472287136 | 0,156851279 | 3,011050578 | 0,002603455 | 0,01936175 |
| ENSG00000138768.15 | USO1 | protein_coding | 10605,94941 | 0,471989062 | 0,127908404 | 3,690055119 | 0,000224205 | 0,002443962 |
| ENSG00000128534.8 | LSM8 | protein_coding | 5295,015993 | 0,471424272 | 0,129322654 | 3,645334036 | 0,000267045 | 0,002837858 |
| ENSG00000285331.2 | AC090517.5 | lncRNA | 1068,178299 | 0,470208724 | 0,15345927 | 3,064062033 | 0,002183536 | 0,016732186 |
| ENSG00000197892.13 | KIF13B | protein_coding | 7824,897265 | 0,469850054 | 0,156186421 | 3,008264431 | 0,002627444 | 0,019491711 |
| ENSG00000270882.2 | H4C14 | protein_coding | 1142,257964 | 0,469438254 | 0,152121251 | 3,085947903 | 0,002029044 | 0,015762621 |
| ENSG00000065882.16 | TBC1D1 | protein_coding | 4685,176749 | 0,46941762 | 0,125718349 | 3,733883096 | 0,00018855 | 0,002095167 |
| ENSG00000219200.12 | RNASEK | protein_coding | 15406,6327 | 0,468682235 | 0,122199167 | 3,83539632 | 0,000125362 | 0,001460464 |
| ENSG00000158373.8 | H2BC5 | protein_coding | 893,2476146 | 0,467983892 | 0,173113553 | 2,703334804 | 0,006864756 | 0,042518837 |
| ENSG00000127022.15 | CANX | protein_coding | 149919,5818 | 0,467962961 | 0,125278797 | 3,735372402 | 0,000187438 | 0,002086128 |
| ENSG00000079931.15 | MOXD1 | protein_coding | 622,2017812 | 0,467429582 | 0,174827136 | 2,673667214 | 0,007502686 | 0,045593247 |
| ENSG00000111321.11 | LTBR | protein_coding | 3312,098482 | 0,466914734 | 0,131578436 | 3,548565763 | 0,000387335 | 0,003897693 |
| ENSG00000115339.14 | GALNT3 | protein_coding | 7081,024258 | 0,466477832 | 0,117351791 | 3,975038042 | 7,04E-05 | 0,000872205 |
| ENSG00000166562.9 | SEC11C | protein_coding | 38497,88776 | 0,466339349 | 0,156832744 | 2,973482048 | 0,002944416 | 0,021418122 |
| ENSG00000105669.14 | COPE | protein_coding | 12084,79314 | 0,466092071 | 0,132111999 | 3,528007112 | 0,000418701 | 0,004166114 |
| ENSG00000134287.10 | ARF3 | protein_coding | 35814,73922 | 0,465829716 | 0,109977782 | 4,23567113 | 2,28E-05 | 0,000316134 |
| ENSG00000132207.18 | SLX1A | protein_coding | 1340,624015 | 0,465570983 | 0,147761128 | 3,150835337 | 0,001628042 | 0,013205374 |
| ENSG00000013293.6 | SLC7A14 | protein_coding | 3913,13987 | 0,465431381 | 0,13604459 | 3,421167869 | 0,000623528 | 0,005827485 |
| ENSG00000139517.9 | LNX2 | protein_coding | 5698,719003 | 0,464563765 | 0,117581652 | 3,950988586 | 7,78E-05 | 0,000955631 |
| ENSG00000137601.17 | NEK1 | protein_coding | 2043,65766 | 0,464422685 | 0,143113592 | 3,245133314 | 0,001173956 | 0,009970296 |
| ENSG00000150991.15 | UBC | protein_coding | 164269,0099 | 0,462580032 | 0,106996058 | 4,323337151 | 1,54E-05 | 0,000222627 |
| ENSG00000105204.14 | DYRK1B | protein_coding | 2834,232882 | 0,461694709 | 0,159044857 | 2,902921336 | 0,003696995 | 0,025928505 |
| ENSG00000138642.15 | HERC6 | protein_coding | 1393,1203 | 0,461672825 | 0,146088599 | 3,160224875 | 0,001576474 | 0,012836828 |
| ENSG00000163703.19 | CRELD1 | protein_coding | 6887,519037 | 0,460287289 | 0,122410207 | 3,760203502 | 0,000169775 | 0,001913971 |
| ENSG00000107829.15 | FBXW4 | protein_coding | 3957,369564 | 0,460262255 | 0,122598566 | 3,754222179 | 0,000173881 | 0,001957091 |
| ENSG00000114125.14 | RNF7 | protein_coding | 5615,189988 | 0,460169224 | 0,13378924 | 3,439508468 | 0,000582772 | 0,005505486 |
| ENSG00000181827.15 | RFX7 | protein_coding | 3216,145034 | 0,459882583 | 0,1539209 | 2,987785162 | 0,00281007 | 0,020614602 |
| ENSG00000196975.16 | ANXA4 | protein_coding | 5405,208785 | 0,459598541 | 0,122207608 | 3,760801366 | 0,00016937 | 0,00191043 |
| ENSG00000088836.14 | SLC4A11 | protein_coding | 1340,033701 | 0,459341609 | 0,14272874 | 3,218283915 | 0,001289601 | 0,010825099 |
| ENSG00000168701.19 | TMEM208 | protein_coding | 7403,940428 | 0,459095919 | 0,135467646 | 3,388970974 | 0,000701554 | 0,006410755 |
| ENSG00000168758.11 | SEMA4C | protein_coding | 681,2698888 | 0,458383962 | 0,168267132 | 2,724144388 | 0,006446832 | 0,040441456 |
| ENSG00000161091.13 | MFSD12 | protein_coding | 5398,695759 | 0,457659952 | 0,11797342 | 3,879348004 | 0,000104737 | 0,001244418 |
| ENSG00000154274.15 | C4orf19 | protein_coding | 3673,334672 | 0,456663001 | 0,125305399 | 3,644400046 | 0,000268016 | 0,002843911 |

| | | | | | | | | |
|---|---|---|---|---|---|---|---|---|
| ENSG00000169641.13 | LUZP1 | protein_coding | 2282,389156 | 0,455430049 | 0,163239144 | 2,789956131 | 0,005271518 | 0,034411985 |
| ENSG00000172331.12 | BPGM | protein_coding | 1671,823429 | 0,455233175 | 0,163896805 | 2,777559782 | 0,005476876 | 0,035519156 |
| ENSG00000162777.17 | DENND2D | protein_coding | 2306,924936 | 0,455029564 | 0,132157947 | 3,443073785 | 0,000575142 | 0,005446805 |
| ENSG00000173786.17 | CNP | protein_coding | 11084,34596 | 0,454238124 | 0,114740738 | 3,958821692 | 7,53E-05 | 0,000927004 |
| ENSG00000124783.14 | SSR1 | protein_coding | 29318,69777 | 0,454018227 | 0,136269096 | 3,3317769 | 0,000862934 | 0,007651641 |
| ENSG00000120725.13 | SIL1 | protein_coding | 6614,349013 | 0,453819499 | 0,123939841 | 3,661611107 | 0,000250634 | 0,002681158 |
| ENSG00000166326.7 | TRIM44 | protein_coding | 16927,78048 | 0,4531901 | 0,136993746 | 3,308107959 | 0,000939286 | 0,008231038 |
| ENSG00000165948.11 | IFI27L1 | protein_coding | 1967,371374 | 0,452883921 | 0,146190196 | 3,097908979 | 0,001948913 | 0,015258223 |
| ENSG00000127314.18 | RAP1B | protein_coding | 24911,12187 | 0,452850098 | 0,144454147 | 3,134905491 | 0,001719095 | 0,013799559 |
| ENSG00000120910.15 | PPP3CC | protein_coding | 915,2540352 | 0,452760515 | 0,16573726 | 2,731796799 | 0,006298998 | 0,039722378 |
| ENSG00000245556.3 | SCAMP1-AS1 | lncRNA | 1190,294374 | 0,452079434 | 0,161392487 | 2,801118212 | 0,005092586 | 0,033379239 |
| ENSG00000213983.12 | AP1G2 | protein_coding | 2754,381463 | 0,451848626 | 0,135525401 | 3,334051213 | 0,000855909 | 0,007599005 |
| ENSG00000175643.10 | RMI2 | protein_coding | 4146,773152 | 0,450263802 | 0,134695646 | 3,342823736 | 0,000829306 | 0,007400474 |
| ENSG00000160752.15 | FDPS | protein_coding | 27867,30909 | 0,450209938 | 0,129502315 | 3,476462472 | 0,000508075 | 0,004908425 |
| ENSG00000204789.4 | ZNF204P | processed_pseudog | 897,7607736 | 0,449711839 | 0,156478746 | 2,873948383 | 0,004053752 | 0,027980646 |
| ENSG00000144191.12 | CNGA3 | protein_coding | 5042,588379 | 0,449627097 | 0,128468149 | 3,499911082 | 0,000465413 | 0,004548753 |
| ENSG00000119522.16 | DENND1A | protein_coding | 2940,066576 | 0,448999146 | 0,135890048 | 3,304135613 | 0,000952697 | 0,008331127 |
| ENSG00000106803.10 | SEC61B | protein_coding | 11194,38194 | 0,448671137 | 0,156515568 | 2,866623072 | 0,004148768 | 0,028533003 |
| ENSG00000181004.10 | BBS12 | protein_coding | 1019,031349 | 0,447740017 | 0,15311431 | 2,924220583 | 0,003453199 | 0,024431474 |
| ENSG00000089818.18 | NECAP1 | protein_coding | 11758,06309 | 0,44757097 | 0,123514598 | 3,623628105 | 0,000290499 | 0,003034731 |
| ENSG00000112902.12 | SEMA5A | protein_coding | 1706,532062 | 0,447552484 | 0,158837657 | 2,817672413 | 0,004837314 | 0,032094436 |
| ENSG00000150527.17 | MIA2 | protein_coding | 9769,474111 | 0,447006918 | 0,115390163 | 3,873873704 | 0,000107119 | 0,001271278 |
| ENSG00000164414.18 | SLC35A1 | protein_coding | 3053,344972 | 0,446929256 | 0,142356571 | 3,139505633 | 0,001692332 | 0,013626524 |
| ENSG00000221914.11 | PPP2R2A | protein_coding | 10233,43772 | 0,446572907 | 0,116925508 | 3,819294153 | 0,000133834 | 0,001544552 |
| ENSG00000144224.17 | UBXN4 | protein_coding | 33606,8765 | 0,446572537 | 0,111465456 | 4,006376097 | 6,17E-05 | 0,000774324 |
| ENSG00000077232.18 | DNAJC10 | protein_coding | 13866,63313 | 0,446567253 | 0,150345111 | 2,97028118 | 0,002975273 | 0,021581825 |
| ENSG00000143079.15 | CTTNBP2NL | protein_coding | 3718,422622 | 0,446288596 | 0,13170727 | 3,388488694 | 0,000702789 | 0,006419238 |
| ENSG00000067082.15 | KLF6 | protein_coding | 1511,598648 | 0,445976765 | 0,141144499 | 3,159717649 | 0,001579221 | 0,0128492 |
| ENSG00000102910.14 | LONP2 | protein_coding | 6988,817849 | 0,445864857 | 0,130574685 | 3,414634741 | 0,000638676 | 0,005941825 |
| ENSG00000243646.10 | IL10RB | protein_coding | 4004,808472 | 0,445719703 | 0,128504298 | 3,4685198 | 0,000523334 | 0,005023382 |
| ENSG00000112812.16 | PRSS16 | protein_coding | 1147,968126 | 0,445581684 | 0,150456971 | 2,96152236 | 0,003061223 | 0,022091243 |
| ENSG00000022840.16 | RNF10 | protein_coding | 10063,72528 | 0,445557634 | 0,130622227 | 3,411039952 | 0,000647156 | 0,005997438 |
| ENSG00000160685.13 | ZBTB7B | protein_coding | 1947,19946 | 0,444942108 | 0,137228376 | 3,242347699 | 0,001185493 | 0,010060111 |
| ENSG00000157800.18 | SLC37A3 | protein_coding | 4746,605689 | 0,444454453 | 0,117233605 | 3,791186443 | 0,000149929 | 0,00170957 |
| ENSG00000114978.18 | MOB1A | protein_coding | 7431,20104 | 0,444379052 | 0,113939128 | 3,900144407 | 9,61E-05 | 0,001155998 |
| ENSG00000126262.5 | FFAR2 | protein_coding | 1674,688951 | 0,44394045 | 0,152387222 | 2,913239339 | 0,003577003 | 0,025179746 |

| | | | | | | | | |
|---|---|---|---|---|---|---|---|---|
| ENSG00000079459.13 | FDFT1 | protein_coding | 25958,7942 | 0,443463153 | 0,120310024 | 3,686003347 | 0,000227803 | 0,002473581 |
| ENSG00000137449.17 | CPEB2 | protein_coding | 2163,826463 | 0,443339769 | 0,15086484 | 2,93865535 | 0,003296394 | 0,023512781 |
| ENSG00000198087.7 | CD2AP | protein_coding | 10492,71043 | 0,443283229 | 0,116383118 | 3,808827585 | 0,000139627 | 0,001605215 |
| ENSG00000174953.14 | DHX36 | protein_coding | 9037,001637 | 0,442655482 | 0,124419463 | 3,557767183 | 0,000374021 | 0,003771735 |
| ENSG00000236104.3 | ZBTB22 | protein_coding | 1122,153914 | 0,442449096 | 0,15314775 | 2,88903425 | 0,00386427 | 0,026939224 |
| ENSG00000068697.7 | LAPTM4A | protein_coding | 40156,13105 | 0,442304025 | 0,131702849 | 3,358348196 | 0,000784098 | 0,007054184 |
| ENSG00000225972.1 | MTND1P23 | unprocessed_pseud | 3351,530212 | 0,441964012 | 0,157056831 | 2,814038767 | 0,004892333 | 0,032380963 |
| ENSG00000178719.17 | GRINA | protein_coding | 8920,360806 | 0,441701663 | 0,141388602 | 3,124025953 | 0,001783947 | 0,014178791 |
| ENSG00000103512.15 | NOMO1 | protein_coding | 59274,34046 | 0,44035615 | 0,1146283 | 3,84160063 | 0,000122235 | 0,0014288 |
| ENSG00000240972.2 | MIF | protein_coding | 27790,23824 | 0,440296008 | 0,144623976 | 3,044419185 | 0,002331301 | 0,017664053 |
| ENSG00000132952.12 | USPL1 | protein_coding | 1723,105234 | 0,440083083 | 0,134344985 | 3,275768595 | 0,001053749 | 0,009078283 |
| ENSG00000134153.10 | EMC7 | protein_coding | 12910,23593 | 0,438201106 | 0,138489273 | 3,164151964 | 0,001555356 | 0,012689544 |
| ENSG00000107175.12 | CREB3 | protein_coding | 5050,402685 | 0,437372731 | 0,12235707 | 3,574560357 | 0,000350817 | 0,003577375 |
| ENSG00000213463.5 | SYNJ2BP | protein_coding | 7244,638795 | 0,437225058 | 0,125708327 | 3,478091455 | 0,000504998 | 0,004887714 |
| ENSG00000052802.13 | MSMO1 | protein_coding | 25252,08436 | 0,435920081 | 0,120566976 | 3,61558444 | 0,000299671 | 0,003118095 |
| ENSG00000257923.12 | CUX1 | protein_coding | 3378,924253 | 0,435756178 | 0,156841302 | 2,778325439 | 0,005463986 | 0,035447336 |
| ENSG00000168591.16 | TMUB2 | protein_coding | 2357,570557 | 0,43564496 | 0,140650452 | 3,097359113 | 0,001952532 | 0,015275132 |
| ENSG00000067445.21 | TRO | protein_coding | 3707,857128 | 0,435103037 | 0,127237164 | 3,41962226 | 0,000627081 | 0,005855464 |
| ENSG00000112511.18 | PHF1 | protein_coding | 2447,397757 | 0,43502539 | 0,127859093 | 3,402381331 | 0,000668014 | 0,00615802 |
| ENSG00000164576.12 | SAP30L | protein_coding | 6138,272716 | 0,435007917 | 0,136781581 | 3,180310643 | 0,001471173 | 0,012087518 |
| ENSG00000163463.13 | KRTCAP2 | protein_coding | 5633,460113 | 0,434495541 | 0,140909031 | 3,083518053 | 0,002045687 | 0,015844046 |
| ENSG00000129515.20 | SNX6 | protein_coding | 9866,268352 | 0,434331894 | 0,131412354 | 3,305107014 | 0,000949401 | 0,008305775 |
| ENSG00000007516.14 | BAIAP3 | protein_coding | 2522,061662 | 0,434176626 | 0,156267606 | 2,778417343 | 0,005462441 | 0,035447336 |
| ENSG00000170348.9 | TMED10 | protein_coding | 33802,86869 | 0,433928214 | 0,134741452 | 3,220450778 | 0,001279892 | 0,010756537 |
| ENSG00000133731.10 | IMPA1 | protein_coding | 4816,039689 | 0,433926023 | 0,127086127 | 3,414424806 | 0,000639168 | 0,005941825 |
| ENSG00000108433.17 | GOSR2 | protein_coding | 4779,140578 | 0,433809726 | 0,119330316 | 3,635368944 | 0,000277583 | 0,002915806 |
| ENSG00000163694.15 | RBM47 | protein_coding | 12557,01026 | 0,433511906 | 0,122754725 | 3,531529274 | 0,000413164 | 0,004118853 |
| ENSG00000119888.11 | EPCAM | protein_coding | 85140,10545 | 0,433245338 | 0,13080106 | 3,312246374 | 0,0009255 | 0,008137481 |
| ENSG00000128272.15 | ATF4 | protein_coding | 6406,981564 | 0,432878396 | 0,154203466 | 2,807189802 | 0,004997579 | 0,032911075 |
| ENSG00000185442.13 | FAM174B | protein_coding | 7572,738168 | 0,43264654 | 0,118725887 | 3,644079235 | 0,000268351 | 0,002844534 |
| ENSG00000179178.11 | TMEM125 | protein_coding | 3765,550017 | 0,432615733 | 0,12914973 | 3,349722316 | 0,000808926 | 0,007249515 |
| ENSG00000163162.9 | RNF149 | protein_coding | 2932,181744 | 0,432302778 | 0,127290558 | 3,396188888 | 0,000683312 | 0,006268683 |
| ENSG00000110619.18 | CARS1 | protein_coding | 2340,980204 | 0,43197291 | 0,134927687 | 3,201514232 | 0,001367073 | 0,01138412 |
| ENSG00000168818.10 | STX18 | protein_coding | 2366,870816 | 0,431823464 | 0,141396246 | 3,053995259 | 0,002258156 | 0,017215771 |
| ENSG00000122417.15 | ODF2L | protein_coding | 1020,877787 | 0,431275044 | 0,15635498 | 2,758307051 | 0,005810159 | 0,037162929 |
| ENSG00000109929.10 | SC5D | protein_coding | 20333,98924 | 0,43042085 | 0,138899508 | 3,098793202 | 0,001943106 | 0,015224148 |

| | | | | | | | | |
|---|---|---|---|---|---|---|---|---|
| ENSG00000135953.11 | MFSD9 | protein_coding | 2734,776614 | 0,430182313 | 0,12856188 | 3,346110953 | 0,000819536 | 0,007328913 |
| ENSG00000170027.7 | YWHAG | protein_coding | 26824,516 | 0,430166866 | 0,110397271 | 3,89653532 | 9,76E-05 | 0,001169323 |
| ENSG00000133313.15 | CNDP2 | protein_coding | 13711,30083 | 0,430103297 | 0,113029333 | 3,805236078 | 0,000141669 | 0,001626009 |
| ENSG00000174243.10 | DDX23 | protein_coding | 13748,66052 | 0,429655717 | 0,110632814 | 3,88361918 | 0,000102913 | 0,001223985 |
| ENSG00000100997.20 | ABHD12 | protein_coding | 17025,62902 | 0,429406413 | 0,117303377 | 3,660648361 | 0,000251578 | 0,002689878 |
| ENSG00000011638.11 | TMEM159 | protein_coding | 1139,351344 | 0,429299833 | 0,161851194 | 2,652435381 | 0,007991342 | 0,047867513 |
| ENSG00000144815.16 | NXPE3 | protein_coding | 1743,40547 | 0,428924952 | 0,154346047 | 2,778982426 | 0,005452948 | 0,03540864 |
| ENSG00000063241.8 | ISOC2 | protein_coding | 2983,812473 | 0,427905074 | 0,133934187 | 3,194890587 | 0,001398839 | 0,011583138 |
| ENSG00000174720.16 | LARP7 | protein_coding | 3114,267002 | 0,427821962 | 0,154902521 | 2,76187862 | 0,005746984 | 0,036849344 |
| ENSG00000165527.7 | ARF6 | protein_coding | 12459,3963 | 0,426559829 | 0,111434488 | 3,827897759 | 0,000129242 | 0,00150066 |
| ENSG00000119698.12 | PPP4R4 | protein_coding | 1208,087882 | 0,425934895 | 0,149760685 | 2,844103533 | 0,004453657 | 0,030057162 |
| ENSG00000183475.13 | ASB7 | protein_coding | 1881,782019 | 0,425889733 | 0,143417099 | 2,969588265 | 0,002981991 | 0,021608857 |
| ENSG00000196586.16 | MYO6 | protein_coding | 9393,815377 | 0,42457611 | 0,119183251 | 3,562380672 | 0,000367507 | 0,003713204 |
| ENSG00000153113.24 | CAST | protein_coding | 6820,923818 | 0,423063829 | 0,116774799 | 3,622903518 | 0,000291315 | 0,00304173 |
| ENSG00000104490.18 | NCALD | protein_coding | 3489,853788 | 0,422628464 | 0,131163025 | 3,222161609 | 0,001272274 | 0,010722219 |
| ENSG00000165406.16 | MARCHF8 | protein_coding | 12972,12123 | 0,4225037 | 0,117020886 | 3,610498232 | 0,000305609 | 0,003172004 |
| ENSG00000171222.10 | SCAND1 | protein_coding | 10795,0305 | 0,422377853 | 0,116713239 | 3,61893693 | 0,000295816 | 0,003084115 |
| ENSG00000204406.14 | MBD5 | protein_coding | 1643,141002 | 0,422157998 | 0,147665468 | 2,858880985 | 0,004251383 | 0,029080213 |
| ENSG00000011275.19 | RNF216 | protein_coding | 3870,994527 | 0,422126726 | 0,136756155 | 3,086710987 | 0,002023843 | 0,01573891 |
| ENSG00000073712.15 | FERMT2 | protein_coding | 3578,911089 | 0,421706986 | 0,123841368 | 3,405219063 | 0,00066111 | 0,006102441 |
| ENSG00000172086.9 | KRCC1 | protein_coding | 1064,492359 | 0,421415956 | 0,153371297 | 2,747684626 | 0,006001771 | 0,038190599 |
| ENSG00000090487.11 | SPG21 | protein_coding | 10934,32346 | 0,420966572 | 0,123721408 | 3,402536224 | 0,000667635 | 0,006157242 |
| ENSG00000157869.16 | RAB28 | protein_coding | 2526,950025 | 0,420406829 | 0,134074458 | 3,135622076 | 0,0017149 | 0,01377117 |
| ENSG00000198455.4 | ZXDB | protein_coding | 1130,427922 | 0,420378557 | 0,159043234 | 2,643171574 | 0,00821334 | 0,049057427 |
| ENSG00000198900.6 | TOP1 | protein_coding | 18829,98724 | 0,419627963 | 0,115957511 | 3,618807949 | 0,000295963 | 0,003084116 |
| ENSG00000165502.7 | RPL36AL | protein_coding | 12061,43369 | 0,417161249 | 0,141242236 | 2,953516314 | 0,00314176 | 0,022617863 |
| ENSG00000082269.17 | FAM135A | protein_coding | 5901,187176 | 0,416382507 | 0,116932353 | 3,560883674 | 0,000369609 | 0,003732639 |
| ENSG00000086544.3 | ITPKC | protein_coding | 1374,432823 | 0,416245046 | 0,144532228 | 2,87994623 | 0,00397743 | 0,027544657 |
| ENSG00000196290.16 | NIF3L1 | protein_coding | 1832,204606 | 0,41605087 | 0,141209513 | 2,946337401 | 0,003215614 | 0,023053482 |
| ENSG00000127399.15 | LRRC61 | protein_coding | 4430,975299 | 0,415223248 | 0,121666676 | 3,412793555 | 0,000643006 | 0,005972199 |
| ENSG00000082458.12 | DLG3 | protein_coding | 5110,575642 | 0,415211284 | 0,120396763 | 3,448691427 | 0,00056331 | 0,005353107 |
| ENSG00000272886.6 | DCP1A | protein_coding | 3250,015947 | 0,414839911 | 0,12888348 | 3,218720588 | 0,001287639 | 0,010812964 |
| ENSG00000067369.14 | TP53BP1 | protein_coding | 8736,267226 | 0,414342373 | 0,141330932 | 2,931717544 | 0,003370932 | 0,023966612 |
| ENSG00000272333.7 | KMT2B | protein_coding | 2146,214397 | 0,412098196 | 0,147604354 | 2,791910838 | 0,005239779 | 0,034236822 |
| ENSG00000113407.14 | TARS1 | protein_coding | 12123,08058 | 0,411669507 | 0,112059727 | 3,673661509 | 0,000239099 | 0,002575488 |
| ENSG00000167608.12 | TMC4 | protein_coding | 1734,910762 | 0,41121065 | 0,147480398 | 2,788239355 | 0,005299537 | 0,034584106 |

| | | | | | | | | |
|---|---|---|---|---|---|---|---|---|
| ENSG00000119801.13 | YPEL5 | protein_coding | 9515,205006 | 0,411118343 | 0,139813152 | 2,940484039 | 0,003276999 | 0,023406336 |
| ENSG00000074276.10 | CDHR2 | protein_coding | 8771,530312 | 0,411078279 | 0,120880302 | 3,400705269 | 0,000672123 | 0,00618515 |
| ENSG00000206418.4 | RAB12 | protein_coding | 5077,743396 | 0,411062519 | 0,11902356 | 3,453623126 | 0,00055311 | 0,005270529 |
| ENSG00000122971.9 | ACADS | protein_coding | 2131,733505 | 0,410420367 | 0,131443227 | 3,122415479 | 0,001793736 | 0,014224191 |
| ENSG00000148824.19 | MTG1 | protein_coding | 2267,849184 | 0,409999272 | 0,133579708 | 3,069323008 | 0,002145445 | 0,0165189 |
| ENSG00000101608.13 | MYL12A | protein_coding | 8223,401958 | 0,409623729 | 0,143388352 | 2,856743412 | 0,004280117 | 0,02921228 |
| ENSG00000113361.13 | CDH6 | protein_coding | 1347,818213 | 0,409569375 | 0,142654039 | 2,871067494 | 0,004090882 | 0,028218322 |
| ENSG00000130522.6 | JUND | protein_coding | 10008,20246 | 0,409351357 | 0,137096093 | 2,985871786 | 0,002827712 | 0,020734902 |
| ENSG00000183735.11 | TBK1 | protein_coding | 3053,493506 | 0,40930998 | 0,123287541 | 3,319962224 | 0,000900296 | 0,007932535 |
| ENSG00000174567.8 | GOLT1A | protein_coding | 1645,304165 | 0,4090162 | 0,15377034 | 2,659916084 | 0,007816012 | 0,047063574 |
| ENSG00000270276.2 | H4C15 | protein_coding | 1022,744714 | 0,40884262 | 0,152783876 | 2,675953962 | 0,007451688 | 0,04530964 |
| ENSG00000183421.12 | RIPK4 | protein_coding | 3526,696688 | 0,408815166 | 0,127853633 | 3,197524833 | 0,001386125 | 0,011506155 |
| ENSG00000121073.15 | SLC35B1 | protein_coding | 9048,839608 | 0,407777009 | 0,134143367 | 3,03985965 | 0,002366884 | 0,017894807 |
| ENSG00000109618.12 | SEPSECS | protein_coding | 2155,023875 | 0,405819089 | 0,130481574 | 3,110163953 | 0,001869835 | 0,014732782 |
| ENSG00000277258.5 | PCGF2 | protein_coding | 2746,530551 | 0,405794064 | 0,148407415 | 2,734324729 | 0,006250836 | 0,039451691 |
| ENSG00000149600.12 | COMMD7 | protein_coding | 3357,930463 | 0,40455508 | 0,12995548 | 3,113028246 | 0,001851783 | 0,014610589 |
| ENSG00000164327.13 | RICTOR | protein_coding | 3340,003844 | 0,40447559 | 0,143121545 | 2,826098534 | 0,004711877 | 0,031425021 |
| ENSG00000133028.12 | SCO1 | protein_coding | 3473,970111 | 0,404458647 | 0,137192411 | 2,948112398 | 0,003197208 | 0,022961766 |
| ENSG00000114354.15 | TFG | protein_coding | 6738,247311 | 0,404119109 | 0,12507492 | 3,231016325 | 0,001233509 | 0,010450631 |
| ENSG00000104517.13 | UBR5 | protein_coding | 10744,5843 | 0,40344045 | 0,115690359 | 3,487243485 | 0,000488027 | 0,004747603 |
| ENSG00000115486.12 | GGCX | protein_coding | 5064,998017 | 0,403185963 | 0,131953821 | 3,055508052 | 0,002246796 | 0,017135398 |
| ENSG00000107104.20 | KANK1 | protein_coding | 3828,927115 | 0,402950254 | 0,135153828 | 2,981419476 | 0,002869155 | 0,020987337 |
| ENSG00000035115.22 | SH3YL1 | protein_coding | 8369,146567 | 0,401092216 | 0,135425688 | 2,961714443 | 0,003059314 | 0,02208508 |
| ENSG00000148842.18 | CNNM2 | protein_coding | 3285,105229 | 0,400029994 | 0,149732457 | 2,671631799 | 0,007548342 | 0,045804216 |
| ENSG00000176986.16 | SEC24C | protein_coding | 16253,34914 | 0,399050927 | 0,124313667 | 3,210032619 | 0,001327199 | 0,011105081 |
| ENSG00000167113.11 | COQ4 | protein_coding | 3262,739658 | 0,39900381 | 0,132063305 | 3,021307161 | 0,002516859 | 0,018804584 |
| ENSG00000169567.13 | HINT1 | protein_coding | 25570,08747 | 0,397737607 | 0,147775799 | 2,691493531 | 0,007113288 | 0,043670581 |
| ENSG00000152904.11 | GGPS1 | protein_coding | 4343,804396 | 0,396790023 | 0,123462143 | 3,213859849 | 0,001309636 | 0,010971279 |
| ENSG00000113643.9 | RARS1 | protein_coding | 10072,33949 | 0,395617063 | 0,12287164 | 3,219758954 | 0,001282984 | 0,0107782 |
| ENSG00000155366.17 | RHOC | protein_coding | 3434,300631 | 0,395606763 | 0,142971608 | 2,767030232 | 0,005656951 | 0,03635788 |
| ENSG00000198846.6 | TOX | protein_coding | 5573,320088 | 0,395021833 | 0,119714754 | 3,299692152 | 0,000967909 | 0,008450035 |
| ENSG00000196655.12 | TRAPPC4 | protein_coding | 4825,826146 | 0,39482922 | 0,129975273 | 3,037725651 | 0,002383709 | 0,018002504 |
| ENSG00000254505.11 | CHMP4A | protein_coding | 3434,372231 | 0,394779113 | 0,13469958 | 2,930811761 | 0,003380776 | 0,024015713 |
| ENSG00000115520.8 | COQ10B | protein_coding | 2365,974416 | 0,394367661 | 0,148336916 | 2,658594175 | 0,007846742 | 0,047190904 |
| ENSG00000002586.20 | CD99 | protein_coding | 55620,58044 | 0,394128965 | 0,110608111 | 3,563291704 | 0,000366233 | 0,00370391 |
| ENSG00000152932.8 | RAB3C | protein_coding | 9516,202303 | 0,394066592 | 0,113379317 | 3,475647961 | 0,000509621 | 0,004921084 |

| | | | | | | | | |
|---|---|---|---|---|---|---|---|---|
| ENSG00000135334.9 | AKIRIN2 | protein_coding | 2316,076356 | 0,393308324 | 0,131306138 | 2,995353678 | 0,002741268 | 0,020178777 |
| ENSG00000172269.19 | DPAGT1 | protein_coding | 3034,747929 | 0,392970436 | 0,127956529 | 3,071124536 | 0,002132542 | 0,016437689 |
| ENSG00000267795.6 | SMIM22 | protein_coding | 4190,396156 | 0,392726262 | 0,132265112 | 2,969235467 | 0,002985417 | 0,021626198 |
| ENSG00000198517.10 | MAFK | protein_coding | 1264,039262 | 0,39181582 | 0,145195321 | 2,698543016 | 0,006964374 | 0,043008603 |
| ENSG00000162989.5 | KCNJ3 | protein_coding | 20124,9972 | 0,391182518 | 0,11344184 | 3,448309004 | 0,000564108 | 0,005358261 |
| ENSG00000160445.11 | ZER1 | protein_coding | 4445,322399 | 0,39107308 | 0,128297236 | 3,048180094 | 0,002302319 | 0,017495118 |
| ENSG00000142751.15 | GPN2 | protein_coding | 1966,665472 | 0,390833701 | 0,132969687 | 2,939269172 | 0,003289872 | 0,023482262 |
| ENSG00000179085.8 | DPM3 | protein_coding | 3659,634955 | 0,390638353 | 0,136864774 | 2,854192067 | 0,004314644 | 0,029365106 |
| ENSG00000104133.15 | SPG11 | protein_coding | 5448,183296 | 0,390583782 | 0,129105855 | 3,025298762 | 0,002483876 | 0,01861031 |
| ENSG00000005700.15 | IBTK | protein_coding | 8383,339021 | 0,389758225 | 0,136197954 | 2,861703974 | 0,004213703 | 0,028884697 |
| ENSG00000162236.13 | STX5 | protein_coding | 3424,62746 | 0,389331869 | 0,126405986 | 3,080011332 | 0,002069927 | 0,0160081 |
| ENSG00000180425.11 | C11orf71 | protein_coding | 2919,58588 | 0,388691636 | 0,140154012 | 2,773317951 | 0,005548788 | 0,035853046 |
| ENSG00000186063.13 | AIDA | protein_coding | 9657,465876 | 0,387953691 | 0,113990823 | 3,403376524 | 0,000665585 | 0,00614104 |
| ENSG00000179918.19 | SEPHS2 | protein_coding | 13907,32033 | 0,386853189 | 0,118696718 | 3,259173436 | 0,001117373 | 0,009547841 |
| ENSG00000197635.11 | DPP4 | protein_coding | 13815,41876 | 0,386644794 | 0,120160117 | 3,217746498 | 0,00129202 | 0,01083671 |
| ENSG00000104375.17 | STK3 | protein_coding | 2349,187176 | 0,386178708 | 0,139555629 | 2,767202668 | 0,005653959 | 0,03635788 |
| ENSG00000182952.5 | HMGN4 | protein_coding | 4922,803858 | 0,38533478 | 0,135132165 | 2,851540054 | 0,0043508 | 0,029553539 |
| ENSG00000133687.16 | TMTC1 | protein_coding | 32085,66395 | 0,384660297 | 0,136847357 | 2,810871211 | 0,004940756 | 0,03262925 |
| ENSG00000131507.11 | NDFIP1 | protein_coding | 25529,22221 | 0,384554706 | 0,12298682 | 3,126796088 | 0,001767225 | 0,014077947 |
| ENSG00000068366.20 | ACSL4 | protein_coding | 6800,531548 | 0,384465127 | 0,122937069 | 3,127332802 | 0,001764001 | 0,014060025 |
| ENSG00000085733.16 | CTTN | protein_coding | 6487,180668 | 0,38430417 | 0,135462637 | 2,836975407 | 0,004554312 | 0,030657402 |
| ENSG00000104412.8 | EMC2 | protein_coding | 3528,922495 | 0,383760526 | 0,135557941 | 2,830970448 | 0,004640701 | 0,031069099 |
| ENSG00000163069.13 | SGCB | protein_coding | 13918,2403 | 0,38372064 | 0,111772908 | 3,433037982 | 0,000596858 | 0,005605757 |
| ENSG00000185043.12 | CIB1 | protein_coding | 2590,335365 | 0,383020194 | 0,136308172 | 2,809957671 | 0,004954802 | 0,032711691 |
| ENSG00000106052.13 | TAX1BP1 | protein_coding | 6900,38864 | 0,382635868 | 0,114199792 | 3,350582893 | 0,000806417 | 0,00723632 |
| ENSG00000274333.4 | CU633967.1 | lncRNA | 4735,531646 | 0,382036791 | 0,123338531 | 3,09746507 | 0,001951834 | 0,015275132 |
| ENSG00000149289.11 | ZC3H12C | protein_coding | 3661,474737 | 0,382017 | 0,141209961 | 2,705311991 | 0,006824026 | 0,042316645 |
| ENSG00000165030.4 | NFIL3 | protein_coding | 1436,444249 | 0,38194173 | 0,144075011 | 2,65099219 | 0,00802557 | 0,048004372 |
| ENSG00000275895.7 | U2AF1L5 | protein_coding | 4453,240278 | 0,381812179 | 0,121359451 | 3,146126445 | 0,001654485 | 0,013388726 |
| ENSG00000171223.6 | JUNB | protein_coding | 3682,210924 | 0,38165997 | 0,120938983 | 3,155806015 | 0,001600553 | 0,013012649 |
| ENSG00000170903.11 | MSANTD4 | protein_coding | 5006,705863 | 0,381127533 | 0,12613774 | 3,021518644 | 0,002515102 | 0,018804584 |
| ENSG00000110931.19 | CAMKK2 | protein_coding | 2114,563759 | 0,381030789 | 0,133211984 | 2,860334157 | 0,004231948 | 0,028971824 |
| ENSG00000086065.14 | CHMP5 | protein_coding | 10015,93341 | 0,38089017 | 0,137144763 | 2,777285552 | 0,0054815 | 0,035519156 |
| ENSG00000094975.14 | SUCO | protein_coding | 8035,958144 | 0,380278848 | 0,123079765 | 3,089694304 | 0,002003626 | 0,015610685 |
| ENSG00000102921.8 | N4BP1 | protein_coding | 4186,33236 | 0,380146928 | 0,129106026 | 2,944455339 | 0,003235236 | 0,023153852 |
| ENSG00000183283.16 | DAZAP2 | protein_coding | 13231,68933 | 0,379756592 | 0,121701709 | 3,120388316 | 0,001806128 | 0,014315125 |

| | | | | | | | | |
|---|---|---|---|---|---|---|---|---|
| ENSG00000100897.18 | DCAF11 | protein_coding | 6917,661097 | 0,379527719 | 0,122302483 | 3,103188983 | 0,001914473 | 0,015039213 |
| ENSG00000072506.13 | HSD17B10 | protein_coding | 12708,03301 | 0,37937649 | 0,133844026 | 2,834467122 | 0,004590218 | 0,030839607 |
| ENSG00000135900.4 | MRPL44 | protein_coding | 3789,184578 | 0,379286229 | 0,130271469 | 2,911506508 | 0,003596904 | 0,025302819 |
| ENSG00000142192.21 | APP | protein_coding | 120987,7466 | 0,379132873 | 0,106848606 | 3,548318384 | 0,000387699 | 0,003897693 |
| ENSG00000198736.12 | MSRB1 | protein_coding | 14038,83399 | 0,379104746 | 0,12467191 | 3,040819273 | 0,002359354 | 0,017857222 |
| ENSG00000120742.11 | SERP1 | protein_coding | 15749,68228 | 0,37899063 | 0,138201324 | 2,742308245 | 0,006100906 | 0,038725762 |
| ENSG00000174021.11 | GNG5 | protein_coding | 4564,836911 | 0,377601354 | 0,129830189 | 2,908424897 | 0,003632544 | 0,025527799 |
| ENSG00000068796.19 | KIF2A | protein_coding | 11662,19058 | 0,377347293 | 0,11756866 | 3,209590826 | 0,00132924 | 0,011117718 |
| ENSG00000100592.16 | DAAM1 | protein_coding | 5974,744403 | 0,376908007 | 0,118279322 | 3,186592564 | 0,001439594 | 0,011879347 |
| ENSG00000185624.15 | P4HB | protein_coding | 154442,2638 | 0,376277898 | 0,11682885 | 3,220761797 | 0,001278504 | 0,010749186 |
| ENSG00000110013.13 | SIAE | protein_coding | 1983,534169 | 0,375650178 | 0,1425557 | 2,635111588 | 0,008410963 | 0,049885075 |
| ENSG00000134686.18 | PHC2 | protein_coding | 18272,79603 | 0,375399376 | 0,128417585 | 2,923270801 | 0,003463751 | 0,024489575 |
| ENSG00000135404.12 | CD63 | protein_coding | 69489,11646 | 0,375314501 | 0,135537659 | 2,769079105 | 0,005621498 | 0,036211099 |
| ENSG00000135525.18 | MAP7 | protein_coding | 6550,990968 | 0,375224835 | 0,124015972 | 3,025617028 | 0,002481264 | 0,018605034 |
| ENSG00000159579.14 | RSPRY1 | protein_coding | 5352,166611 | 0,37454924 | 0,125727364 | 2,979059045 | 0,002891351 | 0,021126613 |
| ENSG00000140262.18 | TCF12 | protein_coding | 4336,066618 | 0,374471638 | 0,136624406 | 2,740883922 | 0,006127415 | 0,038848405 |
| ENSG00000176946.12 | THAP4 | protein_coding | 2505,041404 | 0,374126088 | 0,130949015 | 2,857036291 | 0,004276169 | 0,029207702 |
| ENSG00000161813.22 | LARP4 | protein_coding | 8527,516046 | 0,374109292 | 0,117439043 | 3,185561467 | 0,001444734 | 0,01191237 |
| ENSG00000239779.7 | WBP1 | protein_coding | 4212,366141 | 0,373809537 | 0,131580091 | 2,840927784 | 0,00449825 | 0,030319014 |
| ENSG00000169047.5 | IRS1 | protein_coding | 2310,117664 | 0,373132204 | 0,134996883 | 2,764006074 | 0,005709648 | 0,036643615 |
| ENSG00000198721.13 | ECI2 | protein_coding | 3769,343814 | 0,372929769 | 0,126433875 | 2,949603256 | 0,003181822 | 0,022859111 |
| ENSG00000128563.13 | PRKRIP1 | protein_coding | 2175,215888 | 0,371773495 | 0,130463575 | 2,849634429 | 0,00437695 | 0,029663794 |
| ENSG00000163251.4 | FZD5 | protein_coding | 5948,736138 | 0,371541329 | 0,130440613 | 2,848356192 | 0,004394571 | 0,029744695 |
| ENSG00000050165.18 | DKK3 | protein_coding | 2465,149321 | 0,370181913 | 0,126692235 | 2,921898977 | 0,003479044 | 0,024564512 |
| ENSG00000075089.9 | ACTR6 | protein_coding | 2556,133285 | 0,369997289 | 0,13415533 | 2,757976797 | 0,005816033 | 0,037166863 |
| ENSG00000133794.20 | ARNTL | protein_coding | 2340,854629 | 0,369638165 | 0,135325368 | 2,731477255 | 0,006305109 | 0,039722378 |
| ENSG00000125798.15 | FOXA2 | protein_coding | 9486,57999 | 0,369614175 | 0,13721948 | 2,693598427 | 0,007068527 | 0,043485047 |
| ENSG00000066117.15 | SMARCD1 | protein_coding | 5623,173824 | 0,369281682 | 0,12917161 | 2,858845546 | 0,004251858 | 0,029080213 |
| ENSG00000258289.8 | CHURC1 | protein_coding | 6825,073797 | 0,369248042 | 0,124150712 | 2,974191908 | 0,002937613 | 0,021390931 |
| ENSG00000152684.11 | PELO | protein_coding | 3574,842639 | 0,369030827 | 0,122281668 | 3,017875309 | 0,002545536 | 0,018998503 |
| ENSG00000204619.8 | PPP1R11 | protein_coding | 3179,0885 | 0,368531604 | 0,130271806 | 2,828943695 | 0,004670191 | 0,031216621 |
| ENSG00000079950.14 | STX7 | protein_coding | 6468,371144 | 0,367096439 | 0,1346431 | 2,726440785 | 0,006402144 | 0,040224755 |
| ENSG00000078618.22 | NRDC | protein_coding | 16113,83987 | 0,366840993 | 0,117100173 | 3,132710939 | 0,001731999 | 0,013887171 |
| ENSG00000127481.15 | UBR4 | protein_coding | 15372,62602 | 0,366392708 | 0,109836198 | 3,335810179 | 0,000850512 | 0,007557501 |
| ENSG00000232229.6 | LINC00865 | lncRNA | 1757,078597 | 0,366181806 | 0,133811427 | 2,736551075 | 0,006208696 | 0,039221196 |
| ENSG00000181038.14 | METTL23 | protein_coding | 2837,653805 | 0,366141234 | 0,137652683 | 2,659891729 | 0,007816577 | 0,047063574 |

| | | | | | | | | |
|---|---|---|---|---|---|---|---|---|
| ENSG00000127837.10 | AAMP | protein_coding | 14217,38988 | 0,366129638 | 0,121621512 | 3,01040196 | 0,002609022 | 0,019389374 |
| ENSG00000048649.13 | RSF1 | protein_coding | 6407,213263 | 0,363786696 | 0,133324981 | 2,728571142 | 0,006360937 | 0,040049989 |
| ENSG00000177731.16 | FLII | protein_coding | 6413,859242 | 0,363528097 | 0,124373059 | 2,922884582 | 0,00346805 | 0,024495154 |
| ENSG00000154642.11 | C21orf91 | protein_coding | 2242,027688 | 0,362669339 | 0,129445169 | 2,801721689 | 0,00508307 | 0,033327303 |
| ENSG00000065243.20 | PKN2 | protein_coding | 5001,123719 | 0,362337169 | 0,117350073 | 3,087660368 | 0,002017389 | 0,015702445 |
| ENSG00000113328.19 | CCNG1 | protein_coding | 6914,579764 | 0,362329243 | 0,12376313 | 2,927602462 | 0,003415865 | 0,024231969 |
| ENSG00000082212.13 | ME2 | protein_coding | 9084,732449 | 0,362092762 | 0,119484854 | 3,030449045 | 0,002441904 | 0,018349339 |
| ENSG00000112972.15 | HMGCS1 | protein_coding | 44085,87157 | 0,361683753 | 0,116103891 | 3,11517341 | 0,001838368 | 0,014523107 |
| ENSG00000077254.14 | USP33 | protein_coding | 10226,36245 | 0,360568201 | 0,113242085 | 3,184047699 | 0,001452311 | 0,011960711 |
| ENSG00000171155.8 | C1GALT1C1 | protein_coding | 5993,826816 | 0,360420004 | 0,129426556 | 2,784745375 | 0,005356977 | 0,034882836 |
| ENSG00000178028.14 | DMAP1 | protein_coding | 3168,025984 | 0,360260879 | 0,125307953 | 2,875004102 | 0,004040222 | 0,027905661 |
| ENSG00000064490.14 | RFXANK | protein_coding | 2409,952203 | 0,359874493 | 0,131749613 | 2,731503237 | 0,006304612 | 0,039722378 |
| ENSG00000127947.16 | PTPN12 | protein_coding | 2055,423016 | 0,359852286 | 0,132436438 | 2,717169769 | 0,006584282 | 0,04108553 |
| ENSG00000213949.10 | ITGA1 | protein_coding | 6839,141472 | 0,359414801 | 0,114928793 | 3,127282488 | 0,001764303 | 0,014060025 |
| ENSG00000100934.15 | SEC23A | protein_coding | 3568,082338 | 0,359026536 | 0,13406236 | 2,678056219 | 0,007405079 | 0,045078606 |
| ENSG00000145730.20 | PAM | protein_coding | 26986,17505 | 0,358518356 | 0,12676333 | 2,828249742 | 0,004680328 | 0,031244472 |
| ENSG00000112234.9 | FBXL4 | protein_coding | 5042,246073 | 0,358116534 | 0,118218784 | 3,029269294 | 0,002451461 | 0,018414542 |
| ENSG00000178498.16 | DTX3 | protein_coding | 2609,685608 | 0,357460465 | 0,128379799 | 2,784398077 | 0,005362717 | 0,034909355 |
| ENSG00000167797.8 | CDK2AP2 | protein_coding | 2072,997851 | 0,357229309 | 0,131466403 | 2,717266924 | 0,00658235 | 0,04108553 |
| ENSG00000163527.10 | STT3B | protein_coding | 35727,98294 | 0,35602145 | 0,115969989 | 3,069944677 | 0,002140984 | 0,01649062 |
| ENSG00000137500.9 | CCDC90B | protein_coding | 3784,687939 | 0,35555063 | 0,121639875 | 2,922977593 | 0,003467015 | 0,024495154 |
| ENSG00000102882.12 | MAPK3 | protein_coding | 4838,908783 | 0,355276629 | 0,133171959 | 2,667803579 | 0,007634888 | 0,046182138 |
| ENSG00000185262.9 | UBALD2 | protein_coding | 2142,392388 | 0,354920605 | 0,129317911 | 2,744558749 | 0,006059231 | 0,03850941 |
| ENSG00000072682.19 | P4HA2 | protein_coding | 10824,86818 | 0,354576709 | 0,116397623 | 3,046253868 | 0,002317121 | 0,017581352 |
| ENSG00000163393.13 | SLC22A15 | protein_coding | 2437,802928 | 0,354179225 | 0,131912591 | 2,684953893 | 0,007253981 | 0,044339298 |
| ENSG00000167703.15 | SLC43A2 | protein_coding | 13727,68595 | 0,352818053 | 0,110972521 | 3,179328102 | 0,001476169 | 0,012123812 |
| ENSG00000109220.11 | CHIC2 | protein_coding | 4288,486472 | 0,352313182 | 0,130844007 | 2,692619936 | 0,007089303 | 0,043574446 |
| ENSG00000121486.12 | TRMT1L | protein_coding | 4349,89812 | 0,352006772 | 0,121154953 | 2,905426164 | 0,003667534 | 0,025745314 |
| ENSG00000163399.16 | ATP1A1 | protein_coding | 132556,7853 | 0,351365421 | 0,106650061 | 3,294563728 | 0,000985746 | 0,008577137 |
| ENSG00000141458.13 | NPC1 | protein_coding | 5832,941746 | 0,351222039 | 0,119738676 | 2,933238036 | 0,003354466 | 0,023877937 |
| ENSG00000178741.12 | COX5A | protein_coding | 15275,80609 | 0,35040284 | 0,127881971 | 2,740048781 | 0,006143007 | 0,038923681 |
| ENSG00000002834.18 | LASP1 | protein_coding | 20292,1092 | 0,350377845 | 0,117480561 | 2,982432508 | 0,002859677 | 0,020932635 |
| ENSG00000138760.10 | SCARB2 | protein_coding | 73218,2308 | 0,349109211 | 0,107251555 | 3,255050336 | 0,001133723 | 0,009671754 |
| ENSG00000014216.16 | CAPN1 | protein_coding | 11848,80109 | 0,348216745 | 0,115176839 | 3,023322647 | 0,002500155 | 0,018708735 |
| ENSG00000132507.18 | EIF5A | protein_coding | 38761,06286 | 0,348171839 | 0,128866918 | 2,701793795 | 0,006896652 | 0,042665901 |
| ENSG00000196659.10 | TTC30B | protein_coding | 2213,086612 | 0,347638935 | 0,130175579 | 2,670538793 | 0,007572962 | 0,045892392 |

| | | | | | | | | |
|---|---|---|---|---|---|---|---|---|
| ENSG00000160213.9 | CSTB | protein_coding | 5782,182358 | 0,347403205 | 0,122121906 | 2,84472471 | 0,004444982 | 0,030027654 |
| ENSG00000214548.18 | MEG3 | lncRNA | 7981,044118 | 0,347385153 | 0,123331053 | 2,816688458 | 0,004852157 | 0,032176085 |
| ENSG00000113645.15 | WWC1 | protein_coding | 2337,282913 | 0,346483823 | 0,12892976 | 2,687384373 | 0,007201403 | 0,044077633 |
| ENSG00000100258.18 | LMF2 | protein_coding | 10571,45932 | 0,346279065 | 0,124210643 | 2,787837309 | 0,005306118 | 0,034594702 |
| ENSG00000127334.11 | DYRK2 | protein_coding | 11608,34249 | 0,346145446 | 0,121713373 | 2,843939315 | 0,004455953 | 0,030062966 |
| ENSG00000143862.8 | ARL8A | protein_coding | 2485,949638 | 0,343119677 | 0,127587722 | 2,689284457 | 0,007160537 | 0,043909153 |
| ENSG00000170558.10 | CDH2 | protein_coding | 10977,5711 | 0,34299995 | 0,124208828 | 2,761478045 | 0,005754039 | 0,036871992 |
| ENSG00000244038.11 | DDOST | protein_coding | 44087,82786 | 0,342637449 | 0,129044524 | 2,655187817 | 0,007926426 | 0,047574465 |
| ENSG00000049759.19 | NEDD4L | protein_coding | 6368,260417 | 0,339222759 | 0,118353781 | 2,866175941 | 0,004154633 | 0,028559657 |
| ENSG00000003989.18 | SLC7A2 | protein_coding | 38372,6595 | 0,338662941 | 0,11622916 | 2,913751948 | 0,003571135 | 0,025163823 |
| ENSG00000124659.7 | TBCC | protein_coding | 5491,079571 | 0,337839084 | 0,12525942 | 2,697115189 | 0,006994307 | 0,043117144 |
| ENSG00000164933.12 | SLC25A32 | protein_coding | 2950,509185 | 0,337780858 | 0,12770259 | 2,645058798 | 0,008167672 | 0,048809448 |
| ENSG00000116209.12 | TMEM59 | protein_coding | 64958,21124 | 0,337001186 | 0,122744176 | 2,745557457 | 0,006040819 | 0,038415718 |
| ENSG00000100105.18 | PATZ1 | protein_coding | 4364,293551 | 0,336642504 | 0,118878757 | 2,83181378 | 0,00462848 | 0,031036906 |
| ENSG00000168792.5 | ABHD15 | protein_coding | 12386,54699 | 0,336338643 | 0,113354855 | 2,967130455 | 0,003005934 | 0,021744722 |
| ENSG00000168000.14 | BSCL2 | protein_coding | 12571,58717 | 0,334645613 | 0,1155835 | 2,89527149 | 0,003788307 | 0,026476575 |
| ENSG00000137145.20 | DENND4C | protein_coding | 8606,277682 | 0,334618988 | 0,116333564 | 2,876375273 | 0,004022711 | 0,027821428 |
| ENSG00000131067.17 | GGT7 | protein_coding | 5351,362424 | 0,333676977 | 0,126299375 | 2,641952729 | 0,008242956 | 0,04920624 |
| ENSG00000076944.16 | STXBP2 | protein_coding | 4259,558338 | 0,333500898 | 0,11998256 | 2,779578102 | 0,005442956 | 0,035354728 |
| ENSG00000107745.19 | MICU1 | protein_coding | 6110,48968 | 0,330779188 | 0,121662455 | 2,718827179 | 0,006551383 | 0,040953478 |
| ENSG00000178927.18 | CYBC1 | protein_coding | 4356,590249 | 0,330262843 | 0,119667841 | 2,759829538 | 0,005783153 | 0,037024558 |
| ENSG00000177485.7 | ZBTB33 | protein_coding | 5687,900183 | 0,32677366 | 0,116118086 | 2,814149551 | 0,004890647 | 0,032380043 |
| ENSG00000106780.9 | MEGF9 | protein_coding | 10072,79914 | 0,325272235 | 0,117066604 | 2,77852285 | 0,005460667 | 0,035447336 |
| ENSG00000055130.17 | CUL1 | protein_coding | 3947,361718 | 0,324506251 | 0,121400134 | 2,673030422 | 0,007516943 | 0,04566663 |
| ENSG00000068394.11 | GPKOW | protein_coding | 3573,670817 | 0,324134049 | 0,122947789 | 2,63635525 | 0,008380195 | 0,049766367 |
| ENSG00000100324.14 | TAB1 | protein_coding | 4226,738602 | 0,322029235 | 0,120006851 | 2,68342376 | 0,007287259 | 0,044451854 |
| ENSG00000166986.15 | MARS1 | protein_coding | 11528,36679 | 0,320437301 | 0,115202944 | 2,781502716 | 0,005410788 | 0,035167601 |
| ENSG00000156026.14 | MCU | protein_coding | 3954,417587 | 0,319630772 | 0,120546239 | 2,651520074 | 0,008013035 | 0,047943094 |
| ENSG00000095203.14 | EPB41L4B | protein_coding | 9967,380306 | 0,318834278 | 0,1140082 | 2,796590763 | 0,00516449 | 0,033818769 |
| ENSG00000141580.16 | WDR45B | protein_coding | 12216,11999 | 0,318832279 | 0,114127262 | 2,793655725 | 0,005211593 | 0,034095218 |
| ENSG00000102572.15 | STK24 | protein_coding | 13026,2157 | 0,318707898 | 0,111534214 | 2,85748997 | 0,004270061 | 0,0291755 |
| ENSG00000004478.8 | FKBP4 | protein_coding | 19039,06539 | 0,314311723 | 0,115704898 | 2,716494523 | 0,006597729 | 0,041157166 |
| ENSG00000132792.19 | CTNNBL1 | protein_coding | 7025,116549 | 0,312116621 | 0,116296114 | 2,68380955 | 0,007278856 | 0,044440223 |
| ENSG00000023909.10 | GCLM | protein_coding | 7015,522824 | 0,311331681 | 0,114715924 | 2,713936059 | 0,0066489 | 0,041439332 |
| ENSG00000164292.13 | RHOBTB3 | protein_coding | 22454,34719 | 0,310229595 | 0,110459835 | 2,808528505 | 0,004976848 | 0,032826183 |
| ENSG00000264364.3 | DYNLL2 | protein_coding | 34719,88944 | 0,30983477 | 0,113775328 | 2,723215795 | 0,006464981 | 0,040485906 |

| | | | | | | | | |
|---|---|---|---|---|---|---|---|---|
| ENSG00000105701.16 | FKBP8 | protein_coding | 18004,25099 | 0,309466371 | 0,114089333 | 2,712491723 | 0,006677945 | 0,04159559 |
| ENSG00000044115.21 | CTNNA1 | protein_coding | 16696,08874 | 0,302468461 | 0,113023131 | 2,676164249 | 0,007447014 | 0,045306576 |
| ENSG00000155189.12 | AGPAT5 | protein_coding | 21320,83705 | 0,298591961 | 0,113156843 | 2,63874417 | 0,008321375 | 0,04952144 |
| ENSG00000057704.13 | TMCC3 | protein_coding | 16127,88828 | 0,294580519 | 0,109792411 | 2,683068139 | 0,007295013 | 0,044486189 |
| ENSG00000278540.5 | ACACA | protein_coding | 16073,7263 | -0,296718119 | 0,112638042 | -2,634262046 | 0,008432038 | 0,049950403 |
| ENSG00000225733.6 | FGD5-AS1 | lncRNA | 24511,01279 | -0,298686276 | 0,109247375 | -2,734036177 | 0,006256317 | 0,039474381 |
| ENSG00000155097.12 | ATP6V1C1 | protein_coding | 13325,9921 | -0,307515119 | 0,112360336 | -2,736865429 | 0,006202766 | 0,039209909 |
| ENSG00000181904.9 | C5orf24 | protein_coding | 13850,15913 | -0,307651424 | 0,115391661 | -2,66614954 | 0,007672556 | 0,046381761 |
| ENSG00000108588.14 | CCDC47 | protein_coding | 18212,68363 | -0,308411659 | 0,109995695 | -2,803852075 | 0,005049607 | 0,033170228 |
| ENSG00000113163.18 | CERT1 | protein_coding | 7517,518943 | -0,310387458 | 0,11727387 | -2,646688973 | 0,008128408 | 0,048591722 |
| ENSG00000148730.7 | EIF4EBP2 | protein_coding | 28142,72007 | -0,311850294 | 0,112741793 | -2,766057597 | 0,005673851 | 0,03643622 |
| ENSG00000177511.6 | ST8SIA3 | protein_coding | 7213,004058 | -0,312547185 | 0,117638584 | -2,656842467 | 0,007887629 | 0,047382358 |
| ENSG00000182013.18 | PNMA8A | protein_coding | 7647,499926 | -0,312823243 | 0,115352762 | -2,711883426 | 0,006690212 | 0,041634834 |
| ENSG00000004864.14 | SLC25A13 | protein_coding | 15028,72585 | -0,313058361 | 0,11784078 | -2,656621604 | 0,007892798 | 0,047399806 |
| ENSG00000174607.11 | UGT8 | protein_coding | 13119,98939 | -0,313560899 | 0,113348451 | -2,766344808 | 0,005668856 | 0,036415312 |
| ENSG00000206527.10 | HACD2 | protein_coding | 10765,44394 | -0,314038529 | 0,118332034 | -2,653875862 | 0,007957309 | 0,047718784 |
| ENSG00000257093.7 | DENND11 | protein_coding | 8775,956534 | -0,314045651 | 0,116394946 | -2,698103833 | 0,006973569 | 0,043027309 |
| ENSG00000134897.14 | BIVM | protein_coding | 4082,359679 | -0,314477202 | 0,119281714 | -2,636424232 | 0,008378491 | 0,049766367 |
| ENSG00000128989.11 | ARPP19 | protein_coding | 27002,36324 | -0,3155899 | 0,111359707 | -2,833968496 | 0,004597386 | 0,030877858 |
| ENSG00000278195.2 | SSTR3 | protein_coding | 13341,78233 | -0,316198202 | 0,11655858 | -2,712783595 | 0,006672066 | 0,041571342 |
| ENSG00000130193.8 | THEM6 | protein_coding | 4574,95308 | -0,317625642 | 0,117674752 | -2,699182587 | 0,006951003 | 0,042951371 |
| ENSG00000163618.18 | CADPS | protein_coding | 30376,55631 | -0,318726665 | 0,110087591 | -2,895209733 | 0,003789053 | 0,026476575 |
| ENSG00000197977.4 | ELOVL2 | protein_coding | 9730,911575 | -0,318802114 | 0,112948041 | -2,822555501 | 0,004764257 | 0,031703665 |
| ENSG00000091039.17 | OSBPL8 | protein_coding | 6636,239416 | -0,320403108 | 0,118528055 | -2,703183714 | 0,006867878 | 0,042525589 |
| ENSG00000102893.16 | PHKB | protein_coding | 9343,724534 | -0,320408365 | 0,119300588 | -2,685723256 | 0,007237301 | 0,044263187 |
| ENSG00000095319.14 | NUP188 | protein_coding | 9018,532723 | -0,320834541 | 0,113104527 | -2,836619792 | 0,004559387 | 0,0306817 |
| ENSG00000067955.15 | CBFB | protein_coding | 9094,668154 | -0,322520981 | 0,112177421 | -2,875097124 | 0,004039032 | 0,027905661 |
| ENSG00000167460.17 | TPM4 | protein_coding | 9501,731683 | -0,322635741 | 0,114074133 | -2,828298868 | 0,00467961 | 0,031244472 |
| ENSG00000164187.7 | LMBRD2 | protein_coding | 6493,875502 | -0,323421449 | 0,114679795 | -2,820213005 | 0,004799178 | 0,031883663 |
| ENSG00000196363.9 | WDR5 | protein_coding | 3744,212727 | -0,324128796 | 0,121988132 | -2,657051879 | 0,007882731 | 0,047377121 |
| ENSG00000122042.10 | UBL3 | protein_coding | 15021,60711 | -0,324327872 | 0,115302809 | -2,81283582 | 0,004910672 | 0,032478624 |
| ENSG00000108828.16 | VAT1 | protein_coding | 16357,36979 | -0,324439228 | 0,118473311 | -2,73850056 | 0,006172006 | 0,039060138 |
| ENSG00000196715.7 | VKORC1L1 | protein_coding | 7093,585321 | -0,324610738 | 0,116503138 | -2,786283223 | 0,005331627 | 0,034750187 |
| ENSG00000064115.11 | TM7SF3 | protein_coding | 9777,780999 | -0,324618292 | 0,119250596 | -2,722152354 | 0,006485823 | 0,040592141 |
| ENSG00000088832.18 | FKBP1A | protein_coding | 23271,86796 | -0,324820764 | 0,113411789 | -2,864082885 | 0,004182186 | 0,028725084 |
| ENSG00000136938.9 | ANP32B | protein_coding | 11760,58957 | -0,32487521 | 0,119922957 | -2,709032673 | 0,006747969 | 0,041908129 |

| | | | | | | | | |
|---|---|---|---|---|---|---|---|---|
| ENSG00000114999.8 | TTL | protein_coding | 4147,041055 | -0,325362147 | 0,122201291 | -2,662509905 | 0,007756029 | 0,046792079 |
| ENSG00000136816.16 | TOR1B | protein_coding | 6368,605114 | -0,325640017 | 0,120205646 | -2,709024303 | 0,00674814 | 0,041908129 |
| ENSG00000158669.11 | GPAT4 | protein_coding | 37672,27208 | -0,326756308 | 0,122761156 | -2,661723938 | 0,007774161 | 0,046862096 |
| ENSG00000102753.10 | KPNA3 | protein_coding | 10283,69279 | -0,327399371 | 0,115032672 | -2,846142447 | 0,004425239 | 0,029923252 |
| ENSG00000104164.12 | BLOC1S6 | protein_coding | 8071,189915 | -0,327937394 | 0,114529406 | -2,863346677 | 0,004191916 | 0,028773039 |
| ENSG00000134758.14 | RNF138 | protein_coding | 4987,382617 | -0,330040781 | 0,119643637 | -2,758531829 | 0,005806165 | 0,037149164 |
| ENSG00000121644.19 | DESI2 | protein_coding | 6769,693827 | -0,330144272 | 0,123543815 | -2,6722849 | 0,007533666 | 0,045741673 |
| ENSG00000100714.17 | MTHFD1 | protein_coding | 11043,9976 | -0,33137478 | 0,120735169 | -2,744641705 | 0,0060577 | 0,03850941 |
| ENSG00000133789.15 | SWAP70 | protein_coding | 2760,290674 | -0,331502827 | 0,125569287 | -2,639999278 | 0,00829062 | 0,049434388 |
| ENSG00000096401.8 | CDC5L | protein_coding | 8804,863163 | -0,331883134 | 0,112839244 | -2,941203108 | 0,003269401 | 0,023372625 |
| ENSG00000082258.13 | CCNT2 | protein_coding | 3450,156921 | -0,331966603 | 0,125165053 | -2,652230761 | 0,007996187 | 0,047869654 |
| ENSG00000136908.18 | DPM2 | protein_coding | 5125,937843 | -0,333012658 | 0,11670708 | -2,853405781 | 0,004325335 | 0,029428304 |
| ENSG00000134987.11 | WDR36 | protein_coding | 2983,042174 | -0,333451719 | 0,126315245 | -2,639837494 | 0,008294579 | 0,04944391 |
| ENSG00000163781.14 | TOPBP1 | protein_coding | 8158,780602 | -0,334942057 | 0,121652591 | -2,753266948 | 0,005900376 | 0,037625458 |
| ENSG00000136875.13 | PRPF4 | protein_coding | 5454,025498 | -0,337268365 | 0,116644351 | -2,891424758 | 0,003834994 | 0,026761867 |
| ENSG00000109066.14 | TMEM104 | protein_coding | 4519,377713 | -0,338155295 | 0,126569254 | -2,671701734 | 0,007546769 | 0,045804216 |
| ENSG00000138758.12 | SEPTIN11 | protein_coding | 8289,645863 | -0,340888575 | 0,114239869 | -2,983972035 | 0,002845328 | 0,020842175 |
| ENSG00000101193.8 | GID8 | protein_coding | 11072,00646 | -0,341044487 | 0,114147298 | -2,987757852 | 0,002810321 | 0,020614602 |
| ENSG00000163013.12 | FBXO41 | protein_coding | 3856,806362 | -0,34104938 | 0,126910506 | -2,687321886 | 0,00720275 | 0,044077633 |
| ENSG00000106484.16 | MEST | protein_coding | 15942,20384 | -0,341551883 | 0,129553524 | -2,636376641 | 0,008379666 | 0,049766367 |
| ENSG00000100129.18 | EIF3L | protein_coding | 15704,27122 | -0,341786516 | 0,116292723 | -2,939018943 | 0,003292529 | 0,023493219 |
| ENSG00000099901.17 | RANBP1 | protein_coding | 8601,19924 | -0,342596503 | 0,121465771 | -2,820518891 | 0,004794605 | 0,031867008 |
| ENSG00000096060.15 | FKBP5 | protein_coding | 5433,343787 | -0,342918901 | 0,120455643 | -2,846847946 | 0,004415445 | 0,029866667 |
| ENSG00000136305.11 | CIDEB | protein_coding | 2784,410746 | -0,343014807 | 0,125192426 | -2,739900625 | 0,006145776 | 0,038929449 |
| ENSG00000102178.13 | UBL4A | protein_coding | 4291,042621 | -0,343193147 | 0,119321817 | -2,876197794 | 0,004024974 | 0,027827884 |
| ENSG00000139998.16 | RAB15 | protein_coding | 12962,50296 | -0,343231131 | 0,111717277 | -3,072319165 | 0,002124025 | 0,01637807 |
| ENSG00000112640.16 | PPP2R5D | protein_coding | 2834,826132 | -0,343371038 | 0,124903234 | -2,749096459 | 0,00597598 | 0,038061191 |
| ENSG00000128059.8 | PPAT | protein_coding | 4441,351153 | -0,344312157 | 0,119433937 | -2,88286701 | 0,003940738 | 0,027372045 |
| ENSG00000135503.13 | ACVR1B | protein_coding | 21909,24641 | -0,344717715 | 0,116863288 | -2,949751976 | 0,003180291 | 0,022855955 |
| ENSG00000111445.14 | RFC5 | protein_coding | 3744,099463 | -0,344734657 | 0,129114896 | -2,669983619 | 0,007585495 | 0,045936444 |
| ENSG00000165572.8 | KBTBD6 | protein_coding | 5462,882653 | -0,345119376 | 0,116855119 | -2,95339544 | 0,003142991 | 0,022618944 |
| ENSG00000171551.12 | ECEL1 | protein_coding | 2319,857438 | -0,345818826 | 0,129816304 | -2,663909039 | 0,007723844 | 0,046625918 |
| ENSG00000092201.10 | SUPT16H | protein_coding | 19468,86253 | -0,345970935 | 0,116037062 | -2,981555463 | 0,002867881 | 0,02098535 |
| ENSG00000149547.15 | EI24 | protein_coding | 14488,39772 | -0,346792008 | 0,122792663 | -2,824207887 | 0,004739763 | 0,031580821 |
| ENSG00000113569.16 | NUP155 | protein_coding | 4392,952736 | -0,347144194 | 0,130530775 | -2,659481592 | 0,007826101 | 0,047107365 |
| ENSG00000241399.7 | CD302 | protein_coding | 5878,332203 | -0,347782009 | 0,118117597 | -2,944370838 | 0,00323612 | 0,023153852 |

| | | | | | | | | |
|---|---|---|---|---|---|---|---|---|
| ENSG00000159147.18 | DONSON | protein_coding | 4193,479388 | -0,348786034 | 0,122668206 | -2,843328725 | 0,0044645 | 0,030110512 |
| ENSG00000225151.10 | GOLGA2P7 | transcribed_unproc | 4254,198212 | -0,348999463 | 0,130408791 | -2,676195828 | 0,007446312 | 0,045306576 |
| ENSG00000145725.20 | PPIP5K2 | protein_coding | 5341,310379 | -0,349099036 | 0,129354189 | -2,698784165 | 0,00695933 | 0,042990134 |
| ENSG00000121653.11 | MAPK8IP1 | protein_coding | 16989,44528 | -0,349457287 | 0,111402072 | -3,136901147 | 0,001707437 | 0,013726314 |
| ENSG00000136819.15 | C9orf78 | protein_coding | 5205,494395 | -0,349536661 | 0,127107418 | -2,749931253 | 0,005960777 | 0,037975917 |
| ENSG00000184887.13 | BTBD6 | protein_coding | 6839,376922 | -0,349704605 | 0,117391284 | -2,978965676 | 0,002892232 | 0,021126613 |
| ENSG00000085491.17 | SLC25A24 | protein_coding | 7394,974927 | -0,349784858 | 0,114007232 | -3,068093588 | 0,002154291 | 0,016568733 |
| ENSG00000143179.16 | UCK2 | protein_coding | 3457,723801 | -0,349946808 | 0,124072629 | -2,820499652 | 0,004794893 | 0,031867008 |
| ENSG00000145332.14 | KLHL8 | protein_coding | 5224,915628 | -0,350227776 | 0,122363234 | -2,862197766 | 0,004207143 | 0,028849177 |
| ENSG00000124299.15 | PEPD | protein_coding | 18678,97645 | -0,350494443 | 0,119994557 | -2,920919509 | 0,00349 | 0,02462526 |
| ENSG00000173473.11 | SMARCC1 | protein_coding | 15288,01196 | -0,350736474 | 0,1222645 | -2,868669758 | 0,004122019 | 0,02838634 |
| ENSG00000138078.16 | PREPL | protein_coding | 15153,10356 | -0,351254724 | 0,112424343 | -3,124365366 | 0,00178189 | 0,014171614 |
| ENSG00000126214.21 | KLC1 | protein_coding | 3662,516051 | -0,351399864 | 0,120883057 | -2,90694057 | 0,003649825 | 0,025632034 |
| ENSG00000101654.18 | RNMT | protein_coding | 4603,579216 | -0,35164544 | 0,122908833 | -2,8610266 | 0,004222716 | 0,028927541 |
| ENSG00000071794.16 | HLTF | protein_coding | 8454,543603 | -0,351836574 | 0,116847239 | -3,011081619 | 0,002603189 | 0,01936175 |
| ENSG00000086475.15 | SEPHS1 | protein_coding | 3909,087043 | -0,352061293 | 0,126759953 | -2,777385807 | 0,005479809 | 0,035519156 |
| ENSG00000128973.13 | CLN6 | protein_coding | 5280,22812 | -0,352099931 | 0,122107907 | -2,883514582 | 0,003932644 | 0,027331415 |
| ENSG00000116237.16 | ICMT | protein_coding | 28745,0456 | -0,352525073 | 0,114933633 | -3,067205514 | 0,002160702 | 0,016605839 |
| ENSG00000120438.12 | TCP1 | protein_coding | 25198,57066 | -0,353241982 | 0,11244401 | -3,141492219 | 0,001680893 | 0,013555273 |
| ENSG00000117906.14 | RCN2 | protein_coding | 6737,152581 | -0,353356989 | 0,128062677 | -2,759250367 | 0,005793413 | 0,037078906 |
| ENSG00000161048.11 | NAPEPLD | protein_coding | 5101,07186 | -0,353945165 | 0,118330324 | -2,991161965 | 0,00277918 | 0,020421953 |
| ENSG00000161800.13 | RACGAP1 | protein_coding | 7958,619911 | -0,354004204 | 0,124217403 | -2,849876062 | 0,004373627 | 0,029660472 |
| ENSG00000261236.8 | BOP1 | protein_coding | 3684,542734 | -0,35438748 | 0,121320845 | -2,921076592 | 0,003488241 | 0,024621145 |
| ENSG00000114346.14 | ECT2 | protein_coding | 9705,3976 | -0,354471794 | 0,122973001 | -2,882517221 | 0,003945116 | 0,027384283 |
| ENSG00000130826.18 | DKC1 | protein_coding | 7422,389092 | -0,354971041 | 0,121444274 | -2,922912942 | 0,003467735 | 0,024495154 |
| ENSG00000160325.15 | CACFD1 | protein_coding | 22209,63806 | -0,355285501 | 0,124463941 | -2,854525552 | 0,004310116 | 0,029343833 |
| ENSG00000163950.13 | SLBP | protein_coding | 9740,172796 | -0,355469972 | 0,129442095 | -2,746169805 | 0,006029555 | 0,038355736 |
| ENSG00000102781.14 | KATNAL1 | protein_coding | 4215,862946 | -0,356565739 | 0,133068512 | -2,679565088 | 0,007371787 | 0,044928198 |
| ENSG00000206053.13 | JPT2 | protein_coding | 21119,94009 | -0,356909151 | 0,114981815 | -3,104048674 | 0,001908919 | 0,01500121 |
| ENSG00000116922.14 | C1orf109 | protein_coding | 1986,950168 | -0,357218431 | 0,132940321 | -2,687058585 | 0,007208431 | 0,044099503 |
| ENSG00000167747.15 | C19orf48 | transcribed_unitary | 3135,137443 | -0,357448605 | 0,135381361 | -2,640308848 | 0,00828305 | 0,049403322 |
| ENSG00000142794.19 | NBPF3 | protein_coding | 2197,88316 | -0,357638541 | 0,134812611 | -2,652856709 | 0,007981375 | 0,047844709 |
| ENSG00000196123.13 | KIAA0895L | protein_coding | 2904,916796 | -0,35810937 | 0,126282663 | -2,835776198 | 0,004571447 | 0,030752697 |
| ENSG00000169504.15 | CLIC4 | protein_coding | 5847,784377 | -0,358506474 | 0,11601434 | -3,090191027 | 0,002000278 | 0,015596209 |
| ENSG00000135372.9 | NAT10 | protein_coding | 2462,612011 | -0,35899902 | 0,132939565 | -2,700467829 | 0,006924203 | 0,042823692 |
| ENSG00000171365.17 | CLCN5 | protein_coding | 7418,199661 | -0,359205544 | 0,132455915 | -2,711887525 | 0,006690129 | 0,041634834 |

| | | | | | | | | |
|---|---|---|---|---|---|---|---|---|
| ENSG00000136111.14 | TBC1D4 | protein_coding | 9404,087792 | -0,359901078 | 0,125821739 | -2,860404567 | 0,004231009 | 0,028971824 |
| ENSG00000177084.18 | POLE | protein_coding | 6639,55847 | -0,360499052 | 0,132037285 | -2,730282218 | 0,006328013 | 0,039854677 |
| ENSG00000072501.19 | SMC1A | protein_coding | 29175,60438 | -0,360786367 | 0,1277118 | -2,825004159 | 0,004728 | 0,031522511 |
| ENSG00000129484.14 | PARP2 | protein_coding | 2538,268515 | -0,361723275 | 0,134322718 | -2,692941906 | 0,007082461 | 0,043545174 |
| ENSG00000170191.5 | NANP | protein_coding | 1733,092746 | -0,362104994 | 0,134214075 | -2,697965886 | 0,006976459 | 0,04303246 |
| ENSG00000150712.11 | MTMR12 | protein_coding | 3001,53884 | -0,363029065 | 0,123870428 | -2,930716169 | 0,003381816 | 0,024015713 |
| ENSG00000180694.14 | TMEM64 | protein_coding | 3472,991332 | -0,36325042 | 0,129549051 | -2,803960497 | 0,005047909 | 0,033170228 |
| ENSG00000112029.10 | FBXO5 | protein_coding | 2876,327774 | -0,364448677 | 0,137104054 | -2,658190379 | 0,00785615 | 0,047233918 |
| ENSG00000168615.13 | ADAM9 | protein_coding | 9192,169946 | -0,36471044 | 0,114172435 | -3,194382615 | 0,001401303 | 0,011595364 |
| ENSG00000253729.8 | PRKDC | protein_coding | 35589,03528 | -0,365678398 | 0,111909006 | -3,267640472 | 0,00108448 | 0,009297132 |
| ENSG00000109511.12 | ANXA10 | protein_coding | 52903,26467 | -0,366021383 | 0,13013724 | -2,812579883 | 0,004914581 | 0,032478624 |
| ENSG00000138111.14 | MFSD13A | protein_coding | 2181,463581 | -0,366255503 | 0,131624819 | -2,782571742 | 0,005392994 | 0,035072823 |
| ENSG00000119685.20 | TTLL5 | protein_coding | 1883,195361 | -0,366430697 | 0,13451966 | -2,723993628 | 0,006449775 | 0,040441456 |
| ENSG00000119787.14 | ATL2 | protein_coding | 3662,590312 | -0,366598166 | 0,130109795 | -2,817606213 | 0,004838311 | 0,032094436 |
| ENSG00000153132.13 | CLGN | protein_coding | 8913,644554 | -0,367126133 | 0,133107807 | -2,758111195 | 0,005813642 | 0,037162929 |
| ENSG00000136144.12 | RCBTB1 | protein_coding | 5010,848304 | -0,367258667 | 0,121184716 | -3,030569185 | 0,002440933 | 0,018348626 |
| ENSG00000188229.6 | TUBB4B | protein_coding | 26957,05245 | -0,367557051 | 0,127298948 | -2,887353409 | 0,003884976 | 0,027066535 |
| ENSG00000168785.8 | TSPAN5 | protein_coding | 13932,22977 | -0,367668065 | 0,116602599 | -3,153172134 | 0,001615065 | 0,013109788 |
| ENSG00000123374.11 | CDK2 | protein_coding | 4385,403432 | -0,367719958 | 0,123541027 | -2,97650073 | 0,002915584 | 0,021267508 |
| ENSG00000171867.17 | PRNP | protein_coding | 10064,1509 | -0,367864373 | 0,12385232 | -2,970185558 | 0,002976199 | 0,021581825 |
| ENSG00000136636.13 | KCTD3 | protein_coding | 10576,62884 | -0,36790478 | 0,11138556 | -3,3029845 | 0,000956617 | 0,008358419 |
| ENSG00000101752.12 | MIB1 | protein_coding | 16127,80831 | -0,367937248 | 0,112552614 | -3,269024472 | 0,00107919 | 0,009255566 |
| ENSG00000138780.15 | GSTCD | protein_coding | 2022,067611 | -0,367952935 | 0,135094165 | -2,723677488 | 0,006455952 | 0,040441456 |
| ENSG00000225663.8 | MCRIP1 | protein_coding | 5617,630625 | -0,368268608 | 0,129212538 | -2,85009964 | 0,004370553 | 0,029657734 |
| ENSG00000122591.12 | FAM126A | protein_coding | 4385,301365 | -0,368420675 | 0,12309096 | -2,993076629 | 0,002761804 | 0,0203228 |
| ENSG00000178996.14 | SNX18 | protein_coding | 3715,526461 | -0,368636638 | 0,133158683 | -2,768401052 | 0,005633209 | 0,036253067 |
| ENSG00000175175.6 | PPM1E | protein_coding | 7838,604997 | -0,368871212 | 0,117149582 | -3,14871984 | 0,001639873 | 0,013280753 |
| ENSG00000182534.13 | MXRA7 | protein_coding | 28019,13636 | -0,36958938 | 0,123534506 | -2,991790646 | 0,002773464 | 0,020387102 |
| ENSG00000151233.11 | GXYLT1 | protein_coding | 4188,479024 | -0,371788414 | 0,125814927 | -2,955042178 | 0,003126263 | 0,022521789 |
| ENSG00000180745.5 | CLRN3 | protein_coding | 2480,981478 | -0,372054312 | 0,130589999 | -2,849026085 | 0,004385328 | 0,029701343 |
| ENSG00000093000.19 | NUP50 | protein_coding | 8535,735644 | -0,372797635 | 0,116650645 | -3,195847168 | 0,001394209 | 0,011559514 |
| ENSG00000179409.11 | GEMIN4 | protein_coding | 2829,685227 | -0,373336781 | 0,126259172 | -2,956908198 | 0,003107406 | 0,02240136 |
| ENSG00000198743.7 | SLC5A3 | protein_coding | 5986,274984 | -0,374032439 | 0,119564819 | -3,128281718 | 0,001758316 | 0,01402833 |
| ENSG00000114268.12 | PFKFB4 | protein_coding | 9039,663555 | -0,374105561 | 0,138956745 | -2,692244699 | 0,007097285 | 0,043610702 |
| ENSG00000161682.15 | FAM171A2 | protein_coding | 4160,802168 | -0,374634542 | 0,134722646 | -2,780783738 | 0,005422785 | 0,035234637 |
| ENSG00000059915.17 | PSD | protein_coding | 2496,886554 | -0,37575113 | 0,133833444 | -2,807602641 | 0,004991177 | 0,032879262 |

| | | | | | | | | |
|---|---|---|---|---|---|---|---|---|
| ENSG00000204394.13 | VARS1 | protein_coding | 7189,183472 | -0,376216256 | 0,126519198 | -2,973590275 | 0,002943378 | 0,021418012 |
| ENSG00000173599.15 | PC | protein_coding | 2069,761109 | -0,376576546 | 0,138308847 | -2,722722041 | 0,006474651 | 0,040534334 |
| ENSG00000118922.17 | KLF12 | protein_coding | 2930,794927 | -0,376955837 | 0,135824642 | -2,775312581 | 0,005514868 | 0,035677921 |
| ENSG00000204899.6 | MZT1 | protein_coding | 6026,513482 | -0,377210203 | 0,124224002 | -3,036532358 | 0,002393164 | 0,018060884 |
| ENSG00000096092.6 | TMEM14A | protein_coding | 4868,539426 | -0,377660765 | 0,135463594 | -2,787913369 | 0,005304873 | 0,034594702 |
| ENSG00000119185.13 | ITGB1BP1 | protein_coding | 6895,877881 | -0,377715956 | 0,11870352 | -3,182011418 | 0,001462561 | 0,0120262 |
| ENSG00000147955.17 | SIGMAR1 | protein_coding | 9295,307518 | -0,378078614 | 0,117101518 | -3,228639752 | 0,001243805 | 0,010525081 |
| ENSG00000168067.12 | MAP4K2 | protein_coding | 2280,606927 | -0,378486543 | 0,134804625 | -2,807667338 | 0,004990175 | 0,032879262 |
| ENSG00000173726.11 | TOMM20 | protein_coding | 14895,63792 | -0,379873209 | 0,121877446 | -3,116845839 | 0,001827971 | 0,014454849 |
| ENSG00000187325.5 | TAF9B | protein_coding | 3463,125153 | -0,380351058 | 0,125472253 | -3,031355939 | 0,002434581 | 0,018307452 |
| ENSG00000173442.13 | EHBP1L1 | protein_coding | 3503,706197 | -0,380627259 | 0,139075229 | -2,736844377 | 0,006203163 | 0,039209909 |
| ENSG00000089009.15 | RPL6 | protein_coding | 47559,40545 | -0,381581837 | 0,126270372 | -3,021942766 | 0,00251158 | 0,018785257 |
| ENSG00000153904.21 | DDAH1 | protein_coding | 7160,625983 | -0,38177899 | 0,119362031 | -3,198496097 | 0,001381464 | 0,011476567 |
| ENSG00000180488.15 | MIGA1 | protein_coding | 4997,539777 | -0,382449569 | 0,133946855 | -2,855233655 | 0,004300518 | 0,029297539 |
| ENSG00000144455.14 | SUMF1 | protein_coding | 3301,895367 | -0,38258177 | 0,124667801 | -3,068809804 | 0,002149134 | 0,016541218 |
| ENSG00000187601.5 | MAGEH1 | protein_coding | 3114,780923 | -0,382926846 | 0,141322491 | -2,709595923 | 0,006736522 | 0,041873246 |
| ENSG00000174013.8 | FBXO45 | protein_coding | 4474,450851 | -0,383009106 | 0,120965446 | -3,16626872 | 0,001544081 | 0,012607384 |
| ENSG00000135617.4 | PRADC1 | protein_coding | 2797,683231 | -0,383026927 | 0,141622296 | -2,704566564 | 0,006839357 | 0,042386598 |
| ENSG00000109466.14 | KLHL2 | protein_coding | 4986,681244 | -0,383677856 | 0,116915376 | -3,281671497 | 0,001031938 | 0,008927112 |
| ENSG00000144535.20 | DIS3L2 | protein_coding | 2289,932352 | -0,383879681 | 0,131174516 | -2,926480631 | 0,003428209 | 0,024303944 |
| ENSG00000146733.14 | PSPH | protein_coding | 3068,801955 | -0,384652412 | 0,128628759 | -2,990407557 | 0,002786054 | 0,020465281 |
| ENSG00000163377.16 | TAFA4 | protein_coding | 4096,722422 | -0,38549664 | 0,135168935 | -2,851961806 | 0,004345032 | 0,029523936 |
| ENSG00000089006.17 | SNX5 | protein_coding | 6390,044628 | -0,38569084 | 0,122713298 | -3,143023987 | 0,001672121 | 0,013509763 |
| ENSG00000184787.19 | UBE2G2 | protein_coding | 5308,057479 | -0,386541043 | 0,126577529 | -3,053788831 | 0,002259711 | 0,017221349 |
| ENSG00000082781.12 | ITGB5 | protein_coding | 4294,248314 | -0,386541226 | 0,1264572 | -3,056696066 | 0,002237911 | 0,017086044 |
| ENSG00000169255.15 | B3GALNT1 | protein_coding | 2317,53904 | -0,386815972 | 0,142617697 | -2,712257876 | 0,006682658 | 0,041612567 |
| ENSG00000113368.12 | LMNB1 | protein_coding | 11245,73735 | -0,387021441 | 0,126749033 | -3,053446904 | 0,002262288 | 0,017234712 |
| ENSG00000138386.17 | NAB1 | protein_coding | 2902,524499 | -0,387119478 | 0,125931825 | -3,074040083 | 0,002111811 | 0,016295892 |
| ENSG00000071205.12 | ARHGAP10 | protein_coding | 1536,503347 | -0,387635382 | 0,138590504 | -2,796983707 | 0,005158213 | 0,033788235 |
| ENSG00000258947.7 | TUBB3 | protein_coding | 6869,792409 | -0,387991729 | 0,119246557 | -3,253693339 | 0,001139152 | 0,009709488 |
| ENSG00000124374.9 | PAIP2B | protein_coding | 1528,957557 | -0,388316292 | 0,145755483 | -2,664162498 | 0,007718027 | 0,046617685 |
| ENSG00000160199.15 | PKNOX1 | protein_coding | 3941,177939 | -0,388920159 | 0,121987926 | -3,188185693 | 0,001431686 | 0,011818745 |
| ENSG00000116729.14 | WLS | protein_coding | 8465,040999 | -0,388960405 | 0,130404586 | -2,9827203 | 0,00285699 | 0,020920278 |
| ENSG00000177150.13 | FAM210A | protein_coding | 1189,903135 | -0,389179867 | 0,147696631 | -2,634994876 | 0,008413855 | 0,049885075 |
| ENSG00000278259.4 | MYO19 | protein_coding | 4358,137686 | -0,38918905 | 0,131685775 | -2,955437279 | 0,003122262 | 0,022500706 |
| ENSG00000149654.11 | CDH22 | protein_coding | 7043,38743 | -0,38949003 | 0,134466413 | -2,896559981 | 0,003772785 | 0,026398147 |

| | | | | | | | | |
|---|---|---|---|---|---|---|---|---|
| ENSG00000126351.13 | THRA | protein_coding | 3210,69153 | -0,389693508 | 0,131558079 | -2,962140455 | 0,003055084 | 0,022062153 |
| ENSG00000148180.21 | GSN | protein_coding | 3489,624953 | -0,390020035 | 0,122454799 | -3,18501226 | 0,001447479 | 0,011925607 |
| ENSG00000104889.7 | RNASEH2A | protein_coding | 5079,379815 | -0,390508646 | 0,140754812 | -2,774389303 | 0,005530545 | 0,035746207 |
| ENSG00000068028.17 | RASSF1 | protein_coding | 1307,728914 | -0,391375509 | 0,143486292 | -2,727616027 | 0,006379382 | 0,040105813 |
| ENSG00000104332.12 | SFRP1 | protein_coding | 10401,4833 | -0,391784618 | 0,113955443 | -3,438050931 | 0,000585917 | 0,005525307 |
| ENSG00000168118.12 | RAB4A | protein_coding | 3587,851574 | -0,391935141 | 0,133437486 | -2,937219162 | 0,0033117 | 0,023605868 |
| ENSG00000157500.12 | APPL1 | protein_coding | 10235,39486 | -0,392011086 | 0,111916248 | -3,502718253 | 0,000460536 | 0,004516163 |
| ENSG00000015171.20 | ZMYND11 | protein_coding | 5614,803878 | -0,392634755 | 0,116121178 | -3,381250192 | 0,000721568 | 0,006570695 |
| ENSG00000221890.6 | NPTXR | protein_coding | 5578,542262 | -0,393001337 | 0,139322881 | -2,820795365 | 0,004790475 | 0,031857877 |
| ENSG00000196230.13 | TUBB | protein_coding | 85304,72897 | -0,393607951 | 0,122357017 | -3,216880896 | 0,001295924 | 0,010860755 |
| ENSG00000085365.18 | SCAMP1 | protein_coding | 10125,5214 | -0,393823687 | 0,115806378 | -3,400708088 | 0,000672116 | 0,00618515 |
| ENSG00000087086.15 | FTL | protein_coding | 118403,1402 | -0,394010799 | 0,124530096 | -3,163980525 | 0,001556272 | 0,012692075 |
| ENSG00000197119.13 | SLC25A29 | protein_coding | 4950,142412 | -0,394560773 | 0,132954407 | -2,967639682 | 0,003000959 | 0,021723747 |
| ENSG00000105983.22 | LMBR1 | protein_coding | 8958,13691 | -0,394900384 | 0,117535324 | -3,359844265 | 0,000779864 | 0,007025154 |
| ENSG00000177311.11 | ZBTB38 | protein_coding | 5092,205419 | -0,395018517 | 0,124357633 | -3,17647182 | 0,001490783 | 0,012229442 |
| ENSG00000198937.9 | CCDC167 | protein_coding | 1599,481467 | -0,395244809 | 0,146533766 | -2,697295103 | 0,006990529 | 0,043106546 |
| ENSG00000157212.19 | PAXIP1 | protein_coding | 2017,905396 | -0,395458811 | 0,139644494 | -2,831896911 | 0,004627276 | 0,031036906 |
| ENSG00000100522.10 | GNPNAT1 | protein_coding | 5815,742683 | -0,395599328 | 0,122533546 | -3,228498169 | 0,001244421 | 0,010526038 |
| ENSG00000128951.14 | DUT | protein_coding | 11169,53709 | -0,395683773 | 0,141588225 | -2,794609318 | 0,005196247 | 0,034005448 |
| ENSG00000072415.10 | MPP5 | protein_coding | 5269,705519 | -0,396410008 | 0,126231962 | -3,14032994 | 0,001687577 | 0,013593465 |
| ENSG00000092439.16 | TRPM7 | protein_coding | 3998,012192 | -0,396630994 | 0,119614129 | -3,315920923 | 0,000913417 | 0,00803799 |
| ENSG00000152270.9 | PDE3B | protein_coding | 12761,10578 | -0,397285681 | 0,112955879 | -3,517175773 | 0,000436165 | 0,00431527 |
| ENSG00000092140.16 | G2E3 | protein_coding | 3191,40013 | -0,39785913 | 0,123311724 | -3,226450138 | 0,001253361 | 0,010597377 |
| ENSG00000178035.13 | IMPDH2 | protein_coding | 5560,37478 | -0,39800648 | 0,121609603 | -3,272821129 | 0,001064798 | 0,009150884 |
| ENSG00000278730.1 | AC005332.6 | lncRNA | 3596,71846 | -0,398190916 | 0,121535238 | -3,276341265 | 0,001051614 | 0,009067356 |
| ENSG00000133816.17 | MICAL2 | protein_coding | 1699,497159 | -0,398315665 | 0,149402209 | -2,666062763 | 0,007674536 | 0,046381761 |
| ENSG00000169926.11 | KLF13 | protein_coding | 4804,172548 | -0,398527037 | 0,131744824 | -3,024991994 | 0,002486397 | 0,018616854 |
| ENSG00000143845.15 | ETNK2 | protein_coding | 4933,465009 | -0,398811358 | 0,117468367 | -3,395053207 | 0,000686153 | 0,006289232 |
| ENSG00000137486.17 | ARRB1 | protein_coding | 1808,958527 | -0,398918061 | 0,136265387 | -2,927508371 | 0,003416899 | 0,024231969 |
| ENSG00000161981.11 | SNRNP25 | protein_coding | 4525,475005 | -0,399800887 | 0,135954586 | -2,940694391 | 0,003274775 | 0,023398432 |
| ENSG00000183258.12 | DDX41 | protein_coding | 5005,649937 | -0,399990057 | 0,132086998 | -3,02823187 | 0,002459893 | 0,01846463 |
| ENSG00000251664.5 | PCDHA12 | protein_coding | 1959,004951 | -0,400164065 | 0,138193568 | -2,895677922 | 0,003783405 | 0,02645477 |
| ENSG00000099810.21 | MTAP | protein_coding | 3922,017173 | -0,400449836 | 0,121572117 | -3,293928287 | 0,000987977 | 0,008589493 |
| ENSG00000106009.16 | BRAT1 | protein_coding | 1740,691214 | -0,401327944 | 0,147105238 | -2,728168951 | 0,006368698 | 0,040085626 |
| ENSG00000124422.12 | USP22 | protein_coding | 30646,64934 | -0,401680704 | 0,13089957 | -3,068617446 | 0,002150518 | 0,016545788 |
| ENSG00000095002.15 | MSH2 | protein_coding | 8039,422177 | -0,402138421 | 0,117840862 | -3,412554999 | 0,000643569 | 0,005974778 |

| | | | | | | | | |
|---|---|---|---|---|---|---|---|---|
| ENSG00000165716.11 | DIPK1B | protein_coding | 2434,687195 | -0,402867949 | 0,12926214 | -3,116673974 | 0,001829037 | 0,014454849 |
| ENSG00000163964.17 | PIGX | protein_coding | 6632,385635 | -0,40289837 | 0,141037475 | -2,856676004 | 0,004281026 | 0,02921228 |
| ENSG00000177302.15 | TOP3A | protein_coding | 6007,08471 | -0,403407946 | 0,133433591 | -3,023286292 | 0,002500456 | 0,018708735 |
| ENSG00000040531.16 | CTNS | protein_coding | 2994,521653 | -0,403515844 | 0,127145713 | -3,173648838 | 0,001505357 | 0,0123345 |
| ENSG00000224417.2 | AL606970.1 | lncRNA | 2852,284046 | -0,403594964 | 0,126898962 | -3,180443378 | 0,001470499 | 0,012086726 |
| ENSG00000132953.16 | XPO4 | protein_coding | 3545,16034 | -0,404013873 | 0,126278897 | -3,199377594 | 0,001377247 | 0,011446073 |
| ENSG00000119285.11 | HEATR1 | protein_coding | 3468,94595 | -0,404069333 | 0,131513319 | -3,072459396 | 0,002123027 | 0,016376409 |
| ENSG00000109103.12 | UNC119 | protein_coding | 2034,736616 | -0,404255701 | 0,133076674 | -3,037765295 | 0,002383395 | 0,018002504 |
| ENSG00000134278.15 | SPIRE1 | protein_coding | 1244,445175 | -0,404563448 | 0,14433412 | -2,802964735 | 0,005063521 | 0,033240766 |
| ENSG00000140525.19 | FANCI | protein_coding | 8283,990894 | -0,405621944 | 0,129470436 | -3,132931011 | 0,001730701 | 0,01388208 |
| ENSG00000160408.15 | ST6GALNAC6 | protein_coding | 1903,208877 | -0,405818522 | 0,13845756 | -2,930995763 | 0,003378774 | 0,024010398 |
| ENSG00000150401.15 | DCUN1D2 | protein_coding | 1482,871459 | -0,40637274 | 0,146420639 | -2,775378812 | 0,005513744 | 0,035677921 |
| ENSG00000276644.5 | DACH1 | protein_coding | 1664,950438 | -0,40679603 | 0,136900136 | -2,971480109 | 0,002963681 | 0,021528333 |
| ENSG00000174010.9 | KLHL15 | protein_coding | 8756,851134 | -0,407307596 | 0,115643877 | -3,522085281 | 0,000428166 | 0,004246169 |
| ENSG00000273015.2 | AC008124.1 | lncRNA | 3553,271992 | -0,407405695 | 0,124740423 | -3,266027847 | 0,001090675 | 0,009338767 |
| ENSG00000066230.12 | SLC9A3 | protein_coding | 1071,306712 | -0,407807095 | 0,151684765 | -2,688517175 | 0,007177014 | 0,043971553 |
| ENSG00000156136.10 | DCK | protein_coding | 13252,83473 | -0,408427385 | 0,116289282 | -3,512167054 | 0,000444469 | 0,004393271 |
| ENSG00000134709.12 | HOOK1 | protein_coding | 10534,87559 | -0,40843164 | 0,114407573 | -3,569970328 | 0,000357022 | 0,003630038 |
| ENSG00000151503.13 | NCAPD3 | protein_coding | 6433,238903 | -0,408443096 | 0,115667953 | -3,531169061 | 0,000413727 | 0,004122503 |
| ENSG00000011523.14 | CEP68 | protein_coding | 1951,024163 | -0,408619774 | 0,134967101 | -3,027550931 | 0,002465442 | 0,01849928 |
| ENSG00000136159.4 | NUDT15 | protein_coding | 3712,61877 | -0,409059885 | 0,132961536 | -3,076527979 | 0,002094266 | 0,016166471 |
| ENSG00000149554.13 | CHEK1 | protein_coding | 2735,882871 | -0,40947701 | 0,126951935 | -3,225449137 | 0,001257752 | 0,010621636 |
| ENSG00000100372.15 | SLC25A17 | protein_coding | 3080,530093 | -0,410373398 | 0,14314923 | -2,866752394 | 0,004147073 | 0,028533003 |
| ENSG00000187676.8 | B3GLCT | protein_coding | 1145,694029 | -0,410711968 | 0,147903007 | -2,776900725 | 0,005487994 | 0,035537009 |
| ENSG00000137154.13 | RPS6 | protein_coding | 43092,15514 | -0,410816865 | 0,137799413 | -2,98126715 | 0,002870583 | 0,020990446 |
| ENSG00000132613.15 | MTSS2 | protein_coding | 10839,74331 | -0,410856949 | 0,152374493 | -2,696363026 | 0,007010122 | 0,043189202 |
| ENSG00000089685.15 | BIRC5 | protein_coding | 10934,44659 | -0,411918976 | 0,156097493 | -2,638857084 | 0,008318604 | 0,04952144 |
| ENSG00000108840.15 | HDAC5 | protein_coding | 2372,062318 | -0,412107967 | 0,140571149 | -2,931668191 | 0,003371467 | 0,023966612 |
| ENSG00000160298.18 | C21orf58 | protein_coding | 1591,552138 | -0,412256762 | 0,147604489 | -2,792982546 | 0,005222451 | 0,034144913 |
| ENSG00000169972.12 | PUSL1 | protein_coding | 1011,515228 | -0,412499732 | 0,15217153 | -2,710754984 | 0,006713021 | 0,04175196 |
| ENSG00000186205.13 | MTARC1 | protein_coding | 1908,828573 | -0,412736332 | 0,135380342 | -3,048716873 | 0,00229821 | 0,017476578 |
| ENSG00000163807.6 | KIAA1143 | protein_coding | 3292,013197 | -0,412767551 | 0,147383749 | -2,800631382 | 0,005100274 | 0,033419167 |
| ENSG00000088325.16 | TPX2 | protein_coding | 12284,17419 | -0,413222235 | 0,134841985 | -3,064492386 | 0,002180397 | 0,016720372 |
| ENSG00000156110.15 | ADK | protein_coding | 2457,563337 | -0,413837466 | 0,137735341 | -3,004584458 | 0,002659439 | 0,019652438 |
| ENSG00000115457.10 | IGFBP2 | protein_coding | 5630,103594 | -0,413987254 | 0,155397551 | -2,664052629 | 0,007720548 | 0,046619463 |
| ENSG00000103066.13 | PLA2G15 | protein_coding | 2218,185296 | -0,414352212 | 0,128615504 | -3,221635032 | 0,001274614 | 0,010729409 |

| | | | | | | | | |
|---|---|---|---|---|---|---|---|---|
| ENSG00000160953.16 | PWWP3A | protein_coding | 2150,052329 | -0,414563829 | 0,132967194 | -3,117790305 | 0,001822124 | 0,014421991 |
| ENSG00000115461.5 | IGFBP5 | protein_coding | 21258,25969 | -0,414666924 | 0,143324564 | -2,893202066 | 0,003813359 | 0,026619762 |
| ENSG00000162882.15 | HAAO | protein_coding | 2050,644578 | -0,414698551 | 0,143857035 | -2,88271303 | 0,003942664 | 0,027376345 |
| ENSG00000106976.21 | DNM1 | protein_coding | 6400,208581 | -0,414965119 | 0,127468383 | -3,25543567 | 0,001132185 | 0,009666518 |
| ENSG00000092470.12 | WDR76 | protein_coding | 6222,028817 | -0,415655194 | 0,135088187 | -3,076917412 | 0,002091532 | 0,016151321 |
| ENSG00000100316.16 | RPL3 | protein_coding | 89711,90227 | -0,415765146 | 0,131051665 | -3,172528526 | 0,001511177 | 0,012372504 |
| ENSG00000116711.10 | PLA2G4A | protein_coding | 5398,790389 | -0,417678959 | 0,127343567 | -3,279937636 | 0,0010383 | 0,008971035 |
| ENSG00000068078.20 | FGFR3 | protein_coding | 4772,545517 | -0,417815414 | 0,120064836 | -3,479914912 | 0,000501573 | 0,004865816 |
| ENSG00000165410.15 | CFL2 | protein_coding | 2176,183727 | -0,418152183 | 0,146055354 | -2,862970585 | 0,004196895 | 0,028788336 |
| ENSG00000112578.10 | BYSL | protein_coding | 1248,677088 | -0,418336069 | 0,14523945 | -2,880319845 | 0,003972719 | 0,027523441 |
| ENSG00000099256.19 | PRTFDC1 | protein_coding | 1648,432731 | -0,418654849 | 0,143682311 | -2,913753584 | 0,003571117 | 0,025163823 |
| ENSG00000144381.18 | HSPD1 | protein_coding | 48300,63053 | -0,418696305 | 0,121855644 | -3,436002567 | 0,000590365 | 0,005554741 |
| ENSG00000170846.17 | AC093323.1 | protein_coding | 2754,343604 | -0,418698709 | 0,12674394 | -3,303500801 | 0,000954857 | 0,008346526 |
| ENSG00000146918.20 | NCAPG2 | protein_coding | 6914,869025 | -0,41901088 | 0,145475066 | -2,880293455 | 0,003973052 | 0,027523441 |
| ENSG00000160972.9 | PPP1R16A | protein_coding | 2987,381472 | -0,419801463 | 0,130673359 | -3,212601758 | 0,001315385 | 0,011010633 |
| ENSG00000121957.15 | GPSM2 | protein_coding | 3160,771247 | -0,419866549 | 0,133595951 | -3,142808948 | 0,00167335 | 0,013509763 |
| ENSG00000084444.14 | FAM234B | protein_coding | 3195,134813 | -0,42026439 | 0,125833087 | -3,339856007 | 0,000838218 | 0,007470456 |
| ENSG00000106701.13 | FSD1L | protein_coding | 5061,892195 | -0,420606107 | 0,126671454 | -3,320449027 | 0,000898728 | 0,007922048 |
| ENSG00000137404.14 | NRM | protein_coding | 1518,109074 | -0,420878088 | 0,158699405 | -2,652045787 | 0,008000569 | 0,047882194 |
| ENSG00000196557.13 | CACNA1H | protein_coding | 3323,259369 | -0,420944724 | 0,145208248 | -2,898903674 | 0,003744699 | 0,02621039 |
| ENSG00000103260.9 | METRN | protein_coding | 1909,141998 | -0,421534655 | 0,13899701 | -3,032688663 | 0,002423855 | 0,018246462 |
| ENSG00000164070.12 | HSPA4L | protein_coding | 4186,990581 | -0,421592217 | 0,134369631 | -3,137555826 | 0,001703628 | 0,013701674 |
| ENSG00000104756.16 | KCTD9 | protein_coding | 1815,763214 | -0,421830386 | 0,135650817 | -3,10967818 | 0,001872913 | 0,014745931 |
| ENSG00000120868.14 | APAF1 | protein_coding | 3586,824997 | -0,421845901 | 0,154996137 | -2,721654295 | 0,006495605 | 0,040629071 |
| ENSG00000101442.10 | ACTR5 | protein_coding | 984,7250542 | -0,422129578 | 0,155081099 | -2,721992434 | 0,006488963 | 0,040599652 |
| ENSG00000164754.15 | RAD21 | protein_coding | 64859,7326 | -0,422361422 | 0,110819395 | -3,811259041 | 0,000138261 | 0,001591253 |
| ENSG00000163918.11 | RFC4 | protein_coding | 3596,770346 | -0,423615249 | 0,15540669 | -2,725849497 | 0,006413624 | 0,040278605 |
| ENSG00000090889.12 | KIF4A | protein_coding | 3259,206719 | -0,424025884 | 0,136036315 | -3,117005071 | 0,001826984 | 0,014454849 |
| ENSG00000134802.18 | SLC43A3 | protein_coding | 946,5991 | -0,424162126 | 0,160807756 | -2,637696937 | 0,008347114 | 0,049629888 |
| ENSG00000140474.14 | ULK3 | protein_coding | 3521,122432 | -0,42456731 | 0,135631407 | -3,130302345 | 0,001746265 | 0,01396945 |
| ENSG00000170088.14 | TMEM192 | protein_coding | 2170,943668 | -0,424993039 | 0,13770092 | -3,086348581 | 0,002026311 | 0,015752256 |
| ENSG00000138095.19 | LRPPRC | protein_coding | 17170,00117 | -0,425020324 | 0,111170309 | -3,823146008 | 0,00013176 | 0,00152314 |
| ENSG00000116273.6 | PHF13 | protein_coding | 4506,326334 | -0,425374851 | 0,121226156 | -3,508936214 | 0,000449903 | 0,00443101 |
| ENSG00000164134.13 | NAA15 | protein_coding | 5633,05419 | -0,425694596 | 0,125313055 | -3,397049071 | 0,000681168 | 0,0062544 |
| ENSG00000156531.17 | PHF6 | protein_coding | 3532,343599 | -0,426578576 | 0,128311911 | -3,324543862 | 0,000885633 | 0,007829699 |
| ENSG00000145833.16 | DDX46 | protein_coding | 6216,505174 | -0,426678806 | 0,116504784 | -3,662328632 | 0,000249933 | 0,002675024 |

| | | | | | | | | |
|---|---|---|---|---|---|---|---|---|
| ENSG00000100697.15 | DICER1 | protein_coding | 11733,84298 | -0,427162867 | 0,126470064 | -3,377580863 | 0,000731264 | 0,006650314 |
| ENSG00000198554.12 | WDHD1 | protein_coding | 3842,883971 | -0,427284081 | 0,13051445 | -3,273845019 | 0,001060948 | 0,009129035 |
| ENSG00000164163.11 | ABCE1 | protein_coding | 6908,546699 | -0,427368673 | 0,120036412 | -3,560325296 | 0,000370396 | 0,003738782 |
| ENSG00000150593.18 | PDCD4 | protein_coding | 6191,198395 | -0,4273886 | 0,149636827 | -2,856172569 | 0,00428782 | 0,029239581 |
| ENSG00000008130.15 | NADK | protein_coding | 5396,276401 | -0,427709608 | 0,121561411 | -3,51846531 | 0,000434051 | 0,004298415 |
| ENSG00000197275.14 | RAD54B | protein_coding | 1544,22554 | -0,428052031 | 0,149348976 | -2,866119613 | 0,004155372 | 0,028559657 |
| ENSG00000092531.10 | SNAP23 | protein_coding | 2707,640881 | -0,428163903 | 0,132837964 | -3,223204334 | 0,001267651 | 0,010700916 |
| ENSG00000132383.12 | RPA1 | protein_coding | 11898,01901 | -0,428888644 | 0,13698187 | -3,130988385 | 0,001742191 | 0,013952853 |
| ENSG00000162073.14 | PAQR4 | protein_coding | 7628,54848 | -0,428963835 | 0,115439965 | -3,715904074 | 0,000202478 | 0,002232167 |
| ENSG00000005436.14 | GCFC2 | protein_coding | 1626,365339 | -0,429017708 | 0,159372781 | -2,691913293 | 0,007104341 | 0,04362845 |
| ENSG00000152133.16 | GPATCH11 | protein_coding | 2531,439736 | -0,429098059 | 0,128094794 | -3,349847767 | 0,00080856 | 0,007249336 |
| ENSG00000010292.13 | NCAPD2 | protein_coding | 8960,867715 | -0,42917516 | 0,141367232 | -3,035888538 | 0,00239828 | 0,01809297 |
| ENSG00000224877.4 | NDUFAF8 | protein_coding | 1978,362637 | -0,429784436 | 0,15677783 | -2,741359777 | 0,006118547 | 0,038803933 |
| ENSG00000160208.13 | RRP1B | protein_coding | 3309,77004 | -0,429960208 | 0,138761569 | -3,098553955 | 0,001944676 | 0,015230746 |
| ENSG00000152767.17 | FARP1 | protein_coding | 10170,24842 | -0,430491983 | 0,134329583 | -3,20474444 | 0,001351824 | 0,011284069 |
| ENSG00000089157.16 | RPLP0 | protein_coding | 104054,041 | -0,430586976 | 0,140526176 | -3,064105136 | 0,002183222 | 0,016732186 |
| ENSG00000186998.16 | EMID1 | protein_coding | 7049,030004 | -0,430630331 | 0,118331097 | -3,639198339 | 0,000273488 | 0,002882917 |
| ENSG00000184178.16 | SCFD2 | protein_coding | 1005,318173 | -0,430699774 | 0,162274409 | -2,65414477 | 0,007950971 | 0,047708103 |
| ENSG00000176022.7 | B3GALT6 | protein_coding | 3976,184127 | -0,431483328 | 0,119780739 | -3,602276406 | 0,000315443 | 0,003256308 |
| ENSG00000146143.19 | PRIM2 | protein_coding | 1840,435111 | -0,431694064 | 0,137028226 | -3,150402488 | 0,001630457 | 0,013214716 |
| ENSG00000129810.15 | SGO1 | protein_coding | 1302,522847 | -0,431736738 | 0,160871182 | -2,683741937 | 0,007280328 | 0,044440223 |
| ENSG00000137310.12 | TCF19 | protein_coding | 9736,956836 | -0,432077772 | 0,136571483 | -3,163748117 | 0,001557515 | 0,012697267 |
| ENSG00000091127.14 | PUS7 | protein_coding | 1077,81973 | -0,432169949 | 0,151523554 | -2,852163485 | 0,004342276 | 0,029514789 |
| ENSG00000103249.18 | CLCN7 | protein_coding | 5236,24024 | -0,43228697 | 0,125300243 | -3,450009032 | 0,000560568 | 0,005329467 |
| ENSG00000075711.21 | DLG1 | protein_coding | 7947,665666 | -0,43254775 | 0,115306634 | -3,751282433 | 0,000175932 | 0,001976996 |
| ENSG00000162783.11 | IER5 | protein_coding | 1362,806181 | -0,432747154 | 0,145581057 | -2,972551262 | 0,002953359 | 0,021460801 |
| ENSG00000182307.14 | C8orf33 | protein_coding | 10163,62734 | -0,433192251 | 0,121372224 | -3,569121793 | 0,00035818 | 0,003640047 |
| ENSG00000171451.14 | DSEL | protein_coding | 874,4888665 | -0,433482042 | 0,164237562 | -2,639359949 | 0,008306274 | 0,049494671 |
| ENSG00000213551.7 | DNAJC9 | protein_coding | 5581,967595 | -0,433772404 | 0,140247585 | -3,092904621 | 0,001982078 | 0,015465825 |
| ENSG00000114541.15 | FRMD4B | protein_coding | 3269,601856 | -0,433816975 | 0,159038137 | -2,727754387 | 0,006376707 | 0,040102684 |
| ENSG00000131238.18 | PPT1 | protein_coding | 18812,27684 | -0,434126683 | 0,129143529 | -3,361582937 | 0,000774971 | 0,006993109 |
| ENSG00000247746.4 | USP51 | protein_coding | 868,3478893 | -0,43439171 | 0,163299045 | -2,660099504 | 0,007811757 | 0,047061621 |
| ENSG00000213186.8 | TRIM59 | protein_coding | 2815,618044 | -0,434658837 | 0,135855194 | -3,199427452 | 0,001377008 | 0,011446073 |
| ENSG00000082805.20 | ERC1 | protein_coding | 6241,92315 | -0,435311113 | 0,161422001 | -2,696727276 | 0,007002459 | 0,043154692 |
| ENSG00000128050.9 | PAICS | protein_coding | 8520,904504 | -0,435464337 | 0,12295497 | -3,541657051 | 0,000397622 | 0,003988605 |
| ENSG00000088812.18 | ATRN | protein_coding | 11652,2069 | -0,435544053 | 0,126500494 | -3,443022552 | 0,000575251 | 0,005446805 |

| | | | | | | | | |
|---|---|---|---|---|---|---|---|---|
| ENSG00000083312.17 | TNPO1 | protein_coding | 24788,78963 | -0,435601964 | 0,10883361 | -4,002458118 | 6,27E-05 | 0,000785379 |
| ENSG00000186767.7 | SPIN4 | protein_coding | 2322,39916 | -0,43573129 | 0,136128765 | -3,200875964 | 0,001370105 | 0,011395768 |
| ENSG00000133706.19 | LARS1 | protein_coding | 8086,997566 | -0,436331655 | 0,114965033 | -3,795342304 | 0,00014744 | 0,001683936 |
| ENSG00000167114.13 | SLC27A4 | protein_coding | 5720,470326 | -0,436922745 | 0,120350123 | -3,630430406 | 0,000282949 | 0,002967704 |
| ENSG00000131153.9 | GINS2 | protein_coding | 7411,276268 | -0,437171117 | 0,146995032 | -2,974053694 | 0,002938936 | 0,021393127 |
| ENSG00000162636.16 | FAM102B | protein_coding | 1860,154051 | -0,43720567 | 0,141059266 | -3,099446655 | 0,001938825 | 0,015201985 |
| ENSG00000156162.16 | DPY19L4 | protein_coding | 5020,213029 | -0,437213225 | 0,131204309 | -3,332308417 | 0,000861287 | 0,007640277 |
| ENSG00000164484.12 | TMEM200A | protein_coding | 22018,03743 | -0,438067211 | 0,126805578 | -3,454636764 | 0,000551035 | 0,005253147 |
| ENSG00000112984.12 | KIF20A | protein_coding | 2910,108 | -0,438600095 | 0,166378884 | -2,636152409 | 0,008385206 | 0,049766367 |
| ENSG00000265688.2 | MAFG-DT | lncRNA | 939,893237 | -0,438688892 | 0,161340877 | -2,719018882 | 0,006547587 | 0,040941976 |
| ENSG00000166197.17 | NOLC1 | protein_coding | 6716,418128 | -0,439887491 | 0,128020125 | -3,436080775 | 0,000590195 | 0,005554741 |
| ENSG00000100167.20 | SEPTIN3 | protein_coding | 7492,279335 | -0,440761825 | 0,130547361 | -3,376259934 | 0,000734785 | 0,006676526 |
| ENSG00000172469.16 | MANEA | protein_coding | 3069,590437 | -0,440910445 | 0,125728475 | -3,506846358 | 0,000453451 | 0,004460994 |
| ENSG00000083444.17 | PLOD1 | protein_coding | 6333,149245 | -0,441398206 | 0,125778623 | -3,509326099 | 0,000449244 | 0,00443101 |
| ENSG00000167325.15 | RRM1 | protein_coding | 22251,03383 | -0,441783088 | 0,125233094 | -3,527686448 | 0,000419208 | 0,004169182 |
| ENSG00000184517.12 | ZFP1 | protein_coding | 2057,284151 | -0,442535696 | 0,147190482 | -3,006551025 | 0,002642297 | 0,019560286 |
| ENSG00000165304.8 | MELK | protein_coding | 3955,088856 | -0,44347988 | 0,129717297 | -3,41881839 | 0,000628937 | 0,005864942 |
| ENSG00000123739.11 | PLA2G12A | protein_coding | 6942,420289 | -0,443598887 | 0,128281359 | -3,458015173 | 0,000544171 | 0,005194808 |
| ENSG00000147536.12 | GINS4 | protein_coding | 1297,128383 | -0,443789395 | 0,143839135 | -3,085317467 | 0,00203335 | 0,015771834 |
| ENSG00000114166.8 | KAT2B | protein_coding | 4043,523708 | -0,444386341 | 0,120954725 | -3,673989109 | 0,000238793 | 0,002575488 |
| ENSG00000155016.18 | CYP2U1 | protein_coding | 9983,741965 | -0,444434727 | 0,111657501 | -3,980339193 | 6,88E-05 | 0,000853973 |
| ENSG00000108395.14 | TRIM37 | protein_coding | 6167,39336 | -0,444533857 | 0,125764726 | -3,534646572 | 0,000408321 | 0,004078339 |
| ENSG00000176124.15 | DLEU1 | lncRNA | 2372,720635 | -0,444878505 | 0,140568895 | -3,164843147 | 0,001551666 | 0,012664375 |
| ENSG00000146281.6 | PM20D2 | protein_coding | 3879,92141 | -0,445967601 | 0,128714955 | -3,464769105 | 0,000530687 | 0,005077665 |
| ENSG00000101811.14 | CSTF2 | protein_coding | 1395,768248 | -0,446911892 | 0,144630404 | -3,090027257 | 0,002001381 | 0,015599001 |
| ENSG00000160271.16 | RALGDS | protein_coding | 2579,526105 | -0,447345143 | 0,142961371 | -3,129133002 | 0,001753229 | 0,01400376 |
| ENSG00000102057.10 | KCND1 | protein_coding | 3037,791788 | -0,447883465 | 0,149433016 | -2,997218944 | 0,00272455 | 0,02007689 |
| ENSG00000174442.12 | ZWILCH | protein_coding | 3557,965637 | -0,447931489 | 0,129629742 | -3,455468484 | 0,000549337 | 0,005239352 |
| ENSG00000158164.7 | TMSB15A | protein_coding | 3182,578645 | -0,448259296 | 0,149922404 | -2,989942032 | 0,002790304 | 0,020489307 |
| ENSG00000149483.12 | TMEM138 | protein_coding | 1442,038784 | -0,449230456 | 0,145228774 | -3,09326067 | 0,001979702 | 0,01545304 |
| ENSG00000185480.12 | PARPBP | protein_coding | 1352,865041 | -0,449502197 | 0,148183153 | -3,033423088 | 0,002417963 | 0,018211823 |
| ENSG00000100150.20 | DEPDC5 | protein_coding | 967,1525681 | -0,449627032 | 0,166240397 | -2,704679729 | 0,006837027 | 0,04238471 |
| ENSG00000152620.13 | NADK2 | protein_coding | 1446,740128 | -0,44968983 | 0,139752306 | -3,217763211 | 0,001291944 | 0,01083671 |
| ENSG00000119969.15 | HELLS | protein_coding | 3727,09753 | -0,450064963 | 0,122287496 | -3,680384147 | 0,000232883 | 0,0025235 |
| ENSG00000161888.11 | SPC24 | protein_coding | 952,8242503 | -0,45016445 | 0,159468421 | -2,822906556 | 0,004759044 | 0,031679042 |
| ENSG00000187109.15 | NAP1L1 | protein_coding | 27180,14687 | -0,450602341 | 0,122893281 | -3,666614945 | 0,000245782 | 0,002642761 |

| | | | | | | | | |
|---|---|---|---|---|---|---|---|---|
| ENSG00000101464.11 | PIGU | protein_coding | 4747,962515 | -0,450920181 | 0,124225878 | -3,629840955 | 0,000283596 | 0,002971512 |
| ENSG00000114544.17 | SLC41A3 | protein_coding | 2026,597289 | -0,451407316 | 0,150496276 | -2,99945839 | 0,002704601 | 0,019965026 |
| ENSG00000024526.17 | DEPDC1 | protein_coding | 4851,286984 | -0,452132504 | 0,135781191 | -3,329861093 | 0,000868893 | 0,00769145 |
| ENSG00000164291.17 | ARSK | protein_coding | 1695,7912 | -0,452671115 | 0,148700779 | -3,044174462 | 0,002333198 | 0,017672033 |
| ENSG00000101003.10 | GINS1 | protein_coding | 2759,496455 | -0,453467961 | 0,127070506 | -3,568632677 | 0,000358849 | 0,003645078 |
| ENSG00000117593.12 | DARS2 | protein_coding | 4011,574199 | -0,45373837 | 0,122952564 | -3,690353062 | 0,000223943 | 0,002443068 |
| ENSG00000130695.16 | CEP85 | protein_coding | 3298,205142 | -0,454195947 | 0,131440396 | -3,455527832 | 0,000549216 | 0,005239352 |
| ENSG00000112118.20 | MCM3 | protein_coding | 15683,51218 | -0,45456698 | 0,116605417 | -3,89833501 | 9,69E-05 | 0,001163332 |
| ENSG00000141198.16 | TOM1L1 | protein_coding | 12525,77745 | -0,454931181 | 0,111119865 | -4,094058068 | 4,24E-05 | 0,000552559 |
| ENSG00000079689.14 | SCGN | protein_coding | 35280,62586 | -0,455013511 | 0,12977564 | -3,506155015 | 0,00045463 | 0,004470496 |
| ENSG00000137142.5 | IGFBPL1 | protein_coding | 12798,1712 | -0,455350972 | 0,125057658 | -3,641128263 | 0,000271446 | 0,002865718 |
| ENSG00000122952.17 | ZWINT | protein_coding | 7686,450797 | -0,455545179 | 0,160385491 | -2,840314145 | 0,004506913 | 0,030367627 |
| ENSG00000187017.17 | ESPN | protein_coding | 2368,267085 | -0,455762753 | 0,140123298 | -3,252583673 | 0,001143609 | 0,009725548 |
| ENSG00000171316.12 | CHD7 | protein_coding | 7031,493909 | -0,456423879 | 0,172083317 | -2,652342414 | 0,007993543 | 0,047867513 |
| ENSG00000125967.17 | NECAB3 | protein_coding | 2514,888703 | -0,456474985 | 0,129322886 | -3,529730882 | 0,000415983 | 0,004143004 |
| ENSG00000089060.11 | SLC8B1 | protein_coding | 952,8201193 | -0,456599553 | 0,172628511 | -2,644983438 | 0,008169492 | 0,048809448 |
| ENSG00000147202.18 | DIAPH2 | protein_coding | 2292,436606 | -0,456772638 | 0,14039479 | -3,253487091 | 0,001139979 | 0,009709488 |
| ENSG00000180198.16 | RCC1 | protein_coding | 3182,611522 | -0,456820311 | 0,147101237 | -3,10548246 | 0,001899689 | 0,014951123 |
| ENSG00000180769.10 | WDFY3-AS2 | lncRNA | 755,2807041 | -0,456922365 | 0,169783598 | -2,691204399 | 0,007119456 | 0,043695635 |
| ENSG00000005100.13 | DHX33 | protein_coding | 1780,238665 | -0,457486456 | 0,13734624 | -3,330899022 | 0,00086566 | 0,007672562 |
| ENSG00000275835.5 | TUBGCP5 | protein_coding | 1816,991447 | -0,457595663 | 0,133528055 | -3,426962694 | 0,000610373 | 0,005714739 |
| ENSG00000167670.16 | CHAF1A | protein_coding | 2803,995067 | -0,457597701 | 0,162590844 | -2,814412476 | 0,004886648 | 0,032374045 |
| ENSG00000146414.16 | SHPRH | protein_coding | 1689,918207 | -0,457614588 | 0,147003208 | -3,112956484 | 0,001852233 | 0,014610589 |
| ENSG00000109805.10 | NCAPG | protein_coding | 8308,392547 | -0,458057167 | 0,14985323 | -3,056705342 | 0,002237841 | 0,017086044 |
| ENSG00000106692.15 | FKTN | protein_coding | 2554,604 | -0,458167383 | 0,134806912 | -3,398693549 | 0,000677085 | 0,006227936 |
| ENSG00000101624.11 | CEP76 | protein_coding | 1420,487427 | -0,458365933 | 0,146954108 | -3,119109346 | 0,001813986 | 0,014368446 |
| ENSG00000147224.13 | PRPS1 | protein_coding | 10642,9178 | -0,45849071 | 0,134161084 | -3,417464255 | 0,000632074 | 0,005888949 |
| ENSG00000169679.15 | BUB1 | protein_coding | 4422,769664 | -0,458742611 | 0,146852242 | -3,12383798 | 0,001785087 | 0,014182467 |
| ENSG00000121988.18 | ZRANB3 | protein_coding | 1145,292737 | -0,458995273 | 0,150328833 | -3,053275035 | 0,002263584 | 0,017238314 |
| ENSG00000235194.9 | PPP1R3E | protein_coding | 2620,658638 | -0,459436203 | 0,157105502 | -2,924380089 | 0,00345143 | 0,024431474 |
| ENSG00000122406.14 | RPL5 | protein_coding | 33961,34933 | -0,459505923 | 0,125075669 | -3,67382342 | 0,000238948 | 0,002575488 |
| ENSG00000188175.10 | HEPACAM2 | protein_coding | 13338,22978 | -0,459798807 | 0,12144951 | -3,78592558 | 0,000153138 | 0,001739519 |
| ENSG00000116191.17 | RALGPS2 | protein_coding | 5811,079101 | -0,460179839 | 0,115617163 | -3,980203511 | 6,89E-05 | 0,000853973 |
| ENSG00000116005.12 | PCYOX1 | protein_coding | 24663,10213 | -0,460661563 | 0,114019305 | -4,040206745 | 5,34E-05 | 0,000680472 |
| ENSG00000130309.11 | COLGALT1 | protein_coding | 11275,46817 | -0,460751957 | 0,11463218 | -4,019394519 | 5,83E-05 | 0,000735408 |
| ENSG00000186815.13 | TPCN1 | protein_coding | 2820,436991 | -0,460832008 | 0,138006075 | -3,33921538 | 0,000840154 | 0,007481337 |

| | | | | | | | | |
|---|---|---|---|---|---|---|---|---|
| ENSG00000132646.11 | PCNA | protein_coding | 31076,74481 | -0,461183473 | 0,138331817 | -3,333892989 | 0,000856396 | 0,007600106 |
| ENSG00000159423.17 | ALDH4A1 | protein_coding | 1924,665983 | -0,461233796 | 0,136218301 | -3,385989936 | 0,00070922 | 0,006466689 |
| ENSG00000196177.13 | ACADSB | protein_coding | 2176,291433 | -0,461243621 | 0,130432868 | -3,536252993 | 0,000405846 | 0,004063311 |
| ENSG00000133302.13 | SLF1 | protein_coding | 1887,936334 | -0,461317523 | 0,140697332 | -3,278793682 | 0,001042518 | 0,009000049 |
| ENSG00000178971.16 | CTC1 | protein_coding | 1706,820443 | -0,461338 | 0,144699481 | -3,188249175 | 0,001431371 | 0,011818745 |
| ENSG00000053371.12 | AKR7A2 | protein_coding | 5146,287308 | -0,461490772 | 0,120897531 | -3,817205942 | 0,000134972 | 0,001555963 |
| ENSG00000267575.7 | AC006504.5 | lncRNA | 901,4386879 | -0,461609452 | 0,155796719 | -2,962895843 | 0,003047597 | 0,022015679 |
| ENSG00000115687.14 | PASK | protein_coding | 1177,707456 | -0,461815632 | 0,163234962 | -2,829146572 | 0,004667232 | 0,031206802 |
| ENSG00000258429.2 | PDF | protein_coding | 770,7618658 | -0,462090578 | 0,168152751 | -2,748040542 | 0,00599526 | 0,038172372 |
| ENSG00000203880.12 | PCMTD2 | protein_coding | 7129,525385 | -0,462223037 | 0,118865584 | -3,888619563 | 0,000100816 | 0,001201242 |
| ENSG00000029725.17 | RABEP1 | protein_coding | 10063,34699 | -0,462225986 | 0,121743142 | -3,796731209 | 0,000146617 | 0,001675447 |
| ENSG00000005059.16 | MCUB | protein_coding | 2619,727745 | -0,462474692 | 0,162263958 | -2,850138117 | 0,004370025 | 0,029657734 |
| ENSG00000143228.13 | NUF2 | protein_coding | 2389,034553 | -0,462478272 | 0,171104012 | -2,702907238 | 0,006873593 | 0,042535815 |
| ENSG00000126106.14 | TMEM53 | protein_coding | 1293,032339 | -0,462539324 | 0,164909883 | -2,804800519 | 0,005034773 | 0,033103948 |
| ENSG00000112039.5 | FANCE | protein_coding | 1562,717769 | -0,462611174 | 0,140443725 | -3,293925549 | 0,000987987 | 0,008589493 |
| ENSG00000237037.9 | NDUFA6-DT | lncRNA | 681,5659197 | -0,462863377 | 0,174050661 | -2,659360063 | 0,007828925 | 0,047110818 |
| ENSG00000163597.15 | SNHG16 | lncRNA | 1525,590846 | -0,463215594 | 0,148102642 | -3,127665985 | 0,001762003 | 0,014052393 |
| ENSG00000171865.10 | RNASEH1 | protein_coding | 5556,242942 | -0,463557299 | 0,12085845 | -3,83553901 | 0,000125289 | 0,001460464 |
| ENSG00000123213.23 | NLN | protein_coding | 1540,318624 | -0,463646304 | 0,141254574 | -3,282345422 | 0,001029474 | 0,008909484 |
| ENSG00000107130.10 | NCS1 | protein_coding | 2371,659686 | -0,46401738 | 0,164573313 | -2,819517773 | 0,004809587 | 0,031934253 |
| ENSG00000182093.16 | GET1 | protein_coding | 7674,118273 | -0,464615826 | 0,133031976 | -3,492512399 | 0,000478499 | 0,004665759 |
| ENSG00000140465.14 | CYP1A1 | protein_coding | 2128,813083 | -0,464719895 | 0,13239946 | -3,50998331 | 0,000448135 | 0,004423245 |
| ENSG00000115163.15 | CENPA | protein_coding | 1327,422664 | -0,465134318 | 0,160668265 | -2,894998079 | 0,003791609 | 0,026485594 |
| ENSG00000100578.18 | KIAA0586 | protein_coding | 1784,231098 | -0,465478315 | 0,141474704 | -3,29018758 | 0,001001206 | 0,008682788 |
| ENSG00000182010.11 | RTKN2 | protein_coding | 2063,712197 | -0,466816866 | 0,15966719 | -2,92368687 | 0,003459125 | 0,024465131 |
| ENSG00000136108.15 | CKAP2 | protein_coding | 9436,746412 | -0,46755599 | 0,115072682 | -4,063136296 | 4,84E-05 | 0,00062262 |
| ENSG00000168078.10 | PBK | protein_coding | 5323,623748 | -0,467896754 | 0,157049985 | -2,979285571 | 0,002889214 | 0,021119305 |
| ENSG00000176894.10 | PXMP2 | protein_coding | 3602,182408 | -0,467943135 | 0,171120946 | -2,734575436 | 0,006246078 | 0,039433548 |
| ENSG00000138686.10 | BBS7 | protein_coding | 1335,991932 | -0,468120396 | 0,153421375 | -3,051207139 | 0,002279233 | 0,01734487 |
| ENSG00000116120.11 | FARSB | protein_coding | 4851,661476 | -0,468759066 | 0,126579061 | -3,703290742 | 0,000212821 | 0,002341252 |
| ENSG00000065150.21 | IPO5 | protein_coding | 15663,92726 | -0,469557466 | 0,116501474 | -4,030485208 | 5,57E-05 | 0,000707088 |
| ENSG00000138074.15 | SLC5A6 | protein_coding | 3828,591178 | -0,470249131 | 0,123469467 | -3,80862688 | 0,000139741 | 0,001605637 |
| ENSG00000144554.12 | FANCD2 | protein_coding | 2295,809065 | -0,470555518 | 0,151813759 | -3,099557777 | 0,001938098 | 0,015201977 |
| ENSG00000152127.9 | MGAT5 | protein_coding | 7005,120917 | -0,470658006 | 0,150698671 | -3,123172896 | 0,001789126 | 0,014206785 |
| ENSG00000166881.10 | NEMP1 | protein_coding | 2964,829926 | -0,470732828 | 0,143800144 | -3,27352127 | 0,001062164 | 0,009134366 |
| ENSG00000145912.9 | NHP2 | protein_coding | 5601,014213 | -0,470987733 | 0,157807233 | -2,984576343 | 0,002839713 | 0,020815615 |

| | | | | | | | | |
|---|---|---|---|---|---|---|---|---|
| ENSG00000107164.16 | FUBP3 | protein_coding | 6452,510378 | -0,471995432 | 0,124232721 | -3,799284333 | 0,000145115 | 0,001661002 |
| ENSG00000162885.14 | B3GALNT2 | protein_coding | 1929,694238 | -0,472039963 | 0,140111602 | -3,369028377 | 0,000754337 | 0,006827513 |
| ENSG00000142765.18 | SYTL1 | protein_coding | 2158,848607 | -0,472068995 | 0,13068702 | -3,612210252 | 0,000303598 | 0,003155824 |
| ENSG00000104147.9 | OIP5 | protein_coding | 1464,998839 | -0,472133063 | 0,165206616 | -2,85783387 | 0,004265436 | 0,029153416 |
| ENSG00000219481.10 | NBPF1 | protein_coding | 4127,434481 | -0,472233035 | 0,128015361 | -3,688877897 | 0,000225245 | 0,002450889 |
| ENSG00000174444.15 | RPL4 | protein_coding | 51967,74477 | -0,472551178 | 0,134920177 | -3,502450042 | 0,000461 | 0,004516163 |
| ENSG00000135045.7 | C9orf40 | protein_coding | 922,1017774 | -0,472910465 | 0,17186631 | -2,751618185 | 0,005930162 | 0,037803878 |
| ENSG00000120254.16 | MTHFD1L | protein_coding | 984,8047836 | -0,473004936 | 0,160512799 | -2,94683626 | 0,003210432 | 0,023040464 |
| ENSG00000099822.3 | HCN2 | protein_coding | 1555,217633 | -0,473166306 | 0,179117068 | -2,641659506 | 0,008250095 | 0,049234817 |
| ENSG00000099960.13 | SLC7A4 | protein_coding | 1389,923256 | -0,473335206 | 0,141956476 | -3,33436852 | 0,000854933 | 0,007593561 |
| ENSG00000005469.12 | CROT | protein_coding | 1748,232184 | -0,473638322 | 0,134467015 | -3,522338336 | 0,000427758 | 0,004244128 |
| ENSG00000164045.12 | CDC25A | protein_coding | 2991,95217 | -0,473725847 | 0,133120071 | -3,558635772 | 0,000372786 | 0,003761097 |
| ENSG00000058804.12 | NDC1 | protein_coding | 4103,324512 | -0,474054496 | 0,126547776 | -3,746051572 | 0,00017964 | 0,0020111 |
| ENSG00000101444.13 | AHCY | protein_coding | 35767,9455 | -0,474364681 | 0,131827564 | -3,598372498 | 0,000320215 | 0,003295819 |
| ENSG00000035499.13 | DEPDC1B | protein_coding | 1708,466017 | -0,475560558 | 0,152582733 | -3,116739022 | 0,001828634 | 0,014454849 |
| ENSG00000082516.9 | GEMIN5 | protein_coding | 2860,464998 | -0,476209043 | 0,129122876 | -3,688030036 | 0,000225997 | 0,00245697 |
| ENSG00000118193.12 | KIF14 | protein_coding | 1564,692349 | -0,477128947 | 0,149800974 | -3,185085749 | 0,001447111 | 0,011925607 |
| ENSG00000117394.24 | SLC2A1 | protein_coding | 3603,552249 | -0,477635509 | 0,120789651 | -3,954275095 | 7,68E-05 | 0,000944253 |
| ENSG00000058085.15 | LAMC2 | protein_coding | 668,8989467 | -0,478080727 | 0,178507961 | -2,67820395 | 0,007401813 | 0,045071833 |
| ENSG00000105176.18 | URI1 | protein_coding | 3995,205631 | -0,478103969 | 0,128258291 | -3,727665206 | 0,000193262 | 0,002141841 |
| ENSG00000163507.14 | CIP2A | protein_coding | 2116,524581 | -0,479007776 | 0,134779663 | -3,554006338 | 0,00037941 | 0,003822401 |
| ENSG00000198908.12 | BHLHB9 | protein_coding | 1081,80454 | -0,479480424 | 0,158299727 | -3,028940308 | 0,002454132 | 0,018427994 |
| ENSG00000104419.17 | NDRG1 | protein_coding | 4265,027981 | -0,479594417 | 0,152837466 | -3,137937502 | 0,001701412 | 0,013689104 |
| ENSG00000065328.17 | MCM10 | protein_coding | 677,0688307 | -0,479681887 | 0,178065685 | -2,693847987 | 0,007063237 | 0,043465276 |
| ENSG00000154727.11 | GABPA | protein_coding | 3321,828871 | -0,480038233 | 0,124583539 | -3,853143324 | 0,000116611 | 0,001366883 |
| ENSG00000171219.9 | CDC42BPG | protein_coding | 4458,874727 | -0,480252147 | 0,162983498 | -2,946630515 | 0,003212568 | 0,023040464 |
| ENSG00000166845.15 | C18orf54 | protein_coding | 1512,382626 | -0,480383238 | 0,154911913 | -3,101009015 | 0,001928624 | 0,015143335 |
| ENSG00000070718.12 | AP3M2 | protein_coding | 3290,628862 | -0,480648979 | 0,14536068 | -3,306595553 | 0,000944371 | 0,00826868 |
| ENSG00000184368.16 | MAP7D2 | protein_coding | 10580,16203 | -0,48132528 | 0,118833427 | -4,050419905 | 5,11E-05 | 0,000653829 |
| ENSG00000178718.7 | RPP25 | protein_coding | 5654,776793 | -0,481630276 | 0,12024052 | -4,005557158 | 6,19E-05 | 0,000776081 |
| ENSG00000150756.14 | ATPSCKMT | protein_coding | 2330,640365 | -0,482404241 | 0,135030251 | -3,572564211 | 0,000353503 | 0,003599504 |
| ENSG00000152253.9 | SPC25 | protein_coding | 2606,791264 | -0,482570617 | 0,154233553 | -3,128830324 | 0,001755036 | 0,014007504 |
| ENSG00000117650.13 | NEK2 | protein_coding | 3117,283579 | -0,482885556 | 0,167746074 | -2,878669791 | 0,003993562 | 0,027638092 |
| ENSG00000118276.11 | B4GALT6 | protein_coding | 3423,457177 | -0,482999903 | 0,14240378 | -3,391763229 | 0,000694444 | 0,006356882 |
| ENSG00000105185.12 | PDCD5 | protein_coding | 7246,781553 | -0,483036839 | 0,154665303 | -3,12311055 | 0,001789505 | 0,014206785 |
| ENSG00000156687.11 | UNC5D | protein_coding | 1607,864553 | -0,483037307 | 0,149921152 | -3,221942344 | 0,001273248 | 0,010722219 |

| | | | | | | | | |
|---|---|---|---|---|---|---|---|---|
| ENSG00000173542.9 | MOB1B | protein_coding | 22007,48483 | -0,483385451 | 0,135563184 | -3,565757589 | 0,000362807 | 0,003676358 |
| ENSG00000148935.11 | GAS2 | protein_coding | 1231,598183 | -0,483456508 | 0,179582104 | -2,692119632 | 0,007099947 | 0,043614259 |
| ENSG00000130021.15 | PUDP | protein_coding | 3375,683837 | -0,483527246 | 0,132503363 | -3,649169603 | 0,000263089 | 0,002800089 |
| ENSG00000115758.13 | ODC1 | protein_coding | 21908,21414 | -0,483916237 | 0,123608844 | -3,91489979 | 9,04E-05 | 0,001095717 |
| ENSG00000171488.15 | LRRC8C | protein_coding | 1251,184305 | -0,483941798 | 0,154593291 | -3,130419153 | 0,00174557 | 0,013969234 |
| ENSG00000076382.17 | SPAG5 | protein_coding | 3874,77889 | -0,48447301 | 0,141400649 | -3,42624318 | 0,000611992 | 0,005727339 |
| ENSG00000168496.4 | FEN1 | protein_coding | 11533,78833 | -0,48449293 | 0,124115862 | -3,903553684 | 9,48E-05 | 0,001142452 |
| ENSG00000053372.5 | MRTO4 | protein_coding | 1860,663427 | -0,484811617 | 0,139736101 | -3,469480052 | 0,000521467 | 0,005016961 |
| ENSG00000129993.15 | CBFA2T3 | protein_coding | 1184,570232 | -0,485645655 | 0,182408806 | -2,662402467 | 0,007758505 | 0,046792079 |
| ENSG00000162063.13 | CCNF | protein_coding | 4378,477101 | -0,485710455 | 0,151679675 | -3,202211861 | 0,001363767 | 0,011361104 |
| ENSG00000189227.6 | C15orf61 | protein_coding | 823,9886532 | -0,486009069 | 0,168551856 | -2,883439439 | 0,003933583 | 0,027331415 |
| ENSG00000171208.10 | NETO2 | protein_coding | 6872,126499 | -0,486449838 | 0,135087651 | -3,600994137 | 0,000317003 | 0,003267579 |
| ENSG00000094916.16 | CBX5 | protein_coding | 31997,41431 | -0,486464267 | 0,138054034 | -3,523723668 | 0,000425528 | 0,004224004 |
| ENSG00000196569.13 | LAMA2 | protein_coding | 907,1695854 | -0,48669075 | 0,174744294 | -2,785159617 | 0,005350138 | 0,034859987 |
| ENSG00000154328.16 | NEIL2 | protein_coding | 634,6142215 | -0,487013035 | 0,179147603 | -2,718501537 | 0,006557835 | 0,040981576 |
| ENSG00000205269.6 | TMEM170B | protein_coding | 3978,934113 | -0,487685871 | 0,12359045 | -3,945983469 | 7,95E-05 | 0,000974104 |
| ENSG00000103494.15 | RPGRIP1L | protein_coding | 1185,85832 | -0,488311614 | 0,147592121 | -3,308520887 | 0,000937902 | 0,008222352 |
| ENSG00000101546.13 | RBFA | protein_coding | 838,1154533 | -0,488397409 | 0,181667538 | -2,688413215 | 0,007179249 | 0,043972378 |
| ENSG00000094804.12 | CDC6 | protein_coding | 4263,20698 | -0,488959494 | 0,13202401 | -3,703564939 | 0,000212591 | 0,002339951 |
| ENSG00000101868.13 | POLA1 | protein_coding | 11307,15862 | -0,489268095 | 0,134364663 | -3,641345014 | 0,000271217 | 0,002864751 |
| ENSG00000120800.5 | UTP20 | protein_coding | 1490,561885 | -0,489597446 | 0,160183317 | -3,056482134 | 0,002239508 | 0,017086044 |
| ENSG00000170837.3 | GPR27 | protein_coding | 2185,076359 | -0,489599679 | 0,140476312 | -3,485282837 | 0,000491617 | 0,004778089 |
| ENSG00000149636.16 | DSN1 | protein_coding | 5395,555539 | -0,490507636 | 0,12379967 | -3,962107767 | 7,43E-05 | 0,000917003 |
| ENSG00000065802.12 | ASB1 | protein_coding | 4334,104467 | -0,490812483 | 0,180061733 | -2,725801168 | 0,006414563 | 0,040278605 |
| ENSG00000237649.8 | KIFC1 | protein_coding | 3592,412992 | -0,490842258 | 0,141600975 | -3,466376259 | 0,000527524 | 0,005054336 |
| ENSG00000166900.17 | STX3 | protein_coding | 15527,03709 | -0,491081463 | 0,125674779 | -3,907557788 | 9,32E-05 | 0,001125633 |
| ENSG00000214595.12 | EML6 | protein_coding | 993,5084042 | -0,491527889 | 0,163571586 | -3,004971108 | 0,002656061 | 0,019640624 |
| ENSG00000154122.14 | ANKH | protein_coding | 7089,683813 | -0,491731532 | 0,133107302 | -3,694249104 | 0,000220538 | 0,002419071 |
| ENSG00000139880.19 | CDH24 | protein_coding | 1207,09914 | -0,491760647 | 0,152649807 | -3,221495382 | 0,001275235 | 0,010730325 |
| ENSG00000184162.15 | NR2C2AP | protein_coding | 1143,971017 | -0,492284838 | 0,146899951 | -3,351157262 | 0,000804746 | 0,007227522 |
| ENSG00000073464.13 | CLCN4 | protein_coding | 10770,52348 | -0,492290394 | 0,117585065 | -4,186674504 | 2,83E-05 | 0,000386317 |
| ENSG00000135698.10 | MPHOSPH6 | protein_coding | 2559,009943 | -0,492331952 | 0,152382455 | -3,230896564 | 0,001234026 | 0,010450781 |
| ENSG00000278535.5 | DHRS11 | protein_coding | 794,5245707 | -0,492434874 | 0,183766323 | -2,679679631 | 0,007369265 | 0,044925908 |
| ENSG00000123485.12 | HJURP | protein_coding | 1953,496208 | -0,492616041 | 0,139952844 | -3,519871602 | 0,000431756 | 0,004279738 |
| ENSG00000079482.13 | OPHN1 | protein_coding | 981,0328217 | -0,492671639 | 0,177674383 | -2,772890666 | 0,005556078 | 0,035866945 |
| ENSG00000079112.10 | CDH17 | protein_coding | 41884,85899 | -0,49280606 | 0,115128382 | -4,280491489 | 1,86E-05 | 0,000264677 |

| | | | | | | | | |
|---|---|---|---|---|---|---|---|---|
| ENSG00000184661.14 | CDCA2 | protein_coding | 1563,172137 | -0,493268844 | 0,16305295 | -3,025206502 | 0,002484634 | 0,01861031 |
| ENSG00000138821.13 | SLC39A8 | protein_coding | 3170,909431 | -0,493313692 | 0,136262116 | -3,620328998 | 0,000294229 | 0,003070627 |
| ENSG00000117598.13 | PLPPR5 | protein_coding | 951,4584645 | -0,493535838 | 0,164637845 | -2,997705899 | 0,0027202 | 0,020057103 |
| ENSG00000134690.11 | CDCA8 | protein_coding | 5213,494657 | -0,493942122 | 0,147996567 | -3,337524184 | 0,000845284 | 0,007517423 |
| ENSG00000145241.11 | CENPC | protein_coding | 632,6310668 | -0,493994159 | 0,178257481 | -2,771239425 | 0,005584335 | 0,035993858 |
| ENSG00000213390.11 | ARHGAP19 | protein_coding | 1049,622906 | -0,49409333 | 0,150153624 | -3,290585456 | 0,000999791 | 0,008677708 |
| ENSG00000164182.12 | NDUFAF2 | protein_coding | 793,450018 | -0,494141214 | 0,176978526 | -2,792097013 | 0,005236765 | 0,034227812 |
| ENSG00000172731.14 | LRRC20 | protein_coding | 8420,478809 | -0,494385566 | 0,139323651 | -3,548468344 | 0,000387479 | 0,003897693 |
| ENSG00000213339.9 | QTRT1 | protein_coding | 2117,340234 | -0,494414772 | 0,182028137 | -2,716144765 | 0,006604703 | 0,041176135 |
| ENSG00000185065.6 | AC000068.1 | lncRNA | 650,1641783 | -0,49553938 | 0,176187855 | -2,812562639 | 0,004914845 | 0,032478624 |
| ENSG00000245848.3 | CEBPA | protein_coding | 4378,335734 | -0,495917127 | 0,124414588 | -3,986004655 | 6,72E-05 | 0,000836344 |
| ENSG00000203667.10 | COX20 | protein_coding | 5081,105318 | -0,496023569 | 0,144389659 | -3,435312284 | 0,000591871 | 0,005566409 |
| ENSG00000138796.17 | HADH | protein_coding | 81832,10615 | -0,496722336 | 0,121121869 | -4,101012798 | 4,11E-05 | 0,000537213 |
| ENSG00000159685.11 | CHCHD6 | protein_coding | 671,1936933 | -0,497138817 | 0,18199288 | -2,731638829 | 0,006302018 | 0,039722378 |
| ENSG00000144136.11 | SLC20A1 | protein_coding | 3577,698961 | -0,497206229 | 0,129633344 | -3,835481014 | 0,000125319 | 0,001460464 |
| ENSG00000137343.18 | ATAT1 | protein_coding | 675,7284707 | -0,497948526 | 0,178607799 | -2,787943911 | 0,005304373 | 0,034594702 |
| ENSG00000162302.13 | RPS6KA4 | protein_coding | 1781,984518 | -0,498746564 | 0,148351623 | -3,361921846 | 0,00077402 | 0,006990558 |
| ENSG00000239672.8 | NME1 | protein_coding | 4876,720323 | -0,498921869 | 0,155539259 | -3,207690922 | 0,001338052 | 0,011182484 |
| ENSG00000118965.16 | WDR35 | protein_coding | 1977,651589 | -0,499879217 | 0,137265042 | -3,641708112 | 0,000270835 | 0,002863602 |
| ENSG00000170779.11 | CDCA4 | protein_coding | 1150,121236 | -0,500342129 | 0,170297323 | -2,938050463 | 0,003302833 | 0,023550681 |
| ENSG00000131747.15 | TOP2A | protein_coding | 31946,4282 | -0,500663772 | 0,137924509 | -3,629984092 | 0,000283439 | 0,002971351 |
| ENSG00000249158.7 | PCDHA11 | protein_coding | 2024,989476 | -0,501194109 | 0,156413765 | -3,204283898 | 0,001353989 | 0,011297632 |
| ENSG00000123552.18 | USP45 | protein_coding | 1175,496459 | -0,501823653 | 0,153606298 | -3,266947119 | 0,00108714 | 0,009315434 |
| ENSG00000235954.7 | TTC28-AS1 | lncRNA | 1811,279574 | -0,502065591 | 0,144335504 | -3,478462175 | 0,0005043 | 0,004883216 |
| ENSG00000146670.10 | CDCA5 | protein_coding | 4673,073816 | -0,502086817 | 0,162621444 | -3,087457629 | 0,002018765 | 0,01570526 |
| ENSG00000102098.19 | SCML2 | protein_coding | 7244,825669 | -0,502157296 | 0,115869132 | -4,333831513 | 1,47E-05 | 0,000213769 |
| ENSG00000162377.6 | COA7 | protein_coding | 2123,095125 | -0,503958211 | 0,129535898 | -3,890490734 | 0,000100042 | 0,001194055 |
| ENSG00000163689.20 | CFAP20DC | protein_coding | 802,6750477 | -0,504243744 | 0,182848097 | -2,757719397 | 0,005820614 | 0,037184788 |
| ENSG00000178105.11 | DDX10 | protein_coding | 1243,180081 | -0,506974717 | 0,160461386 | -3,1594811 | 0,001580504 | 0,012854639 |
| ENSG00000122912.15 | SLC25A16 | protein_coding | 687,9886051 | -0,507282579 | 0,170348056 | -2,977918211 | 0,002902134 | 0,021191552 |
| ENSG00000132329.11 | RAMP1 | protein_coding | 4169,269478 | -0,50775109 | 0,125511681 | -4,045448881 | 5,22E-05 | 0,000665827 |
| ENSG00000121410.12 | A1BG | protein_coding | 748,2720205 | -0,507928065 | 0,183902973 | -2,761935047 | 0,005745991 | 0,036849344 |
| ENSG00000127863.16 | TNFRSF19 | protein_coding | 3365,077142 | -0,50892387 | 0,127045306 | -4,005845513 | 6,18E-05 | 0,000775599 |
| ENSG00000160818.17 | GPATCH4 | protein_coding | 1332,690803 | -0,509091994 | 0,143829842 | -3,539543573 | 0,00040082 | 0,00401683 |
| ENSG00000167900.12 | TK1 | protein_coding | 2937,157666 | -0,50923269 | 0,181428109 | -2,806801503 | 0,005003606 | 0,032940408 |
| ENSG00000180089.6 | TMEM86B | protein_coding | 1979,578832 | -0,509333581 | 0,146646635 | -3,473203335 | 0,000514286 | 0,004956984 |

| | | | | | | | | |
|---|---|---|---|---|---|---|---|---|
| ENSG00000166582.10 | CENPV | protein_coding | 3457,201135 | -0,509532831 | 0,135401768 | -3,763117999 | 0,000167808 | 0,001893833 |
| ENSG00000012817.16 | KDM5D | protein_coding | 2961,489147 | -0,509541273 | 0,126673185 | -4,022487248 | 5,76E-05 | 0,000727567 |
| ENSG00000135423.13 | GLS2 | protein_coding | 555,0455778 | -0,509638962 | 0,183828812 | -2,772356281 | 0,005565209 | 0,035914811 |
| ENSG00000140285.12 | FGF7 | protein_coding | 525,0562103 | -0,510982647 | 0,18554607 | -2,753939484 | 0,005888265 | 0,03755967 |
| ENSG00000187714.7 | SLC18A3 | protein_coding | 2599,159997 | -0,51103155 | 0,148109243 | -3,450369068 | 0,000559821 | 0,00532718 |
| ENSG00000198157.11 | HMGN5 | protein_coding | 1025,258872 | -0,511149382 | 0,160541299 | -3,183912085 | 0,001452991 | 0,01196161 |
| ENSG00000261801.6 | LOXL1-AS1 | lncRNA | 945,7623628 | -0,511285852 | 0,168334949 | -3,037312539 | 0,002386978 | 0,018020696 |
| ENSG00000013810.20 | TACC3 | protein_coding | 4718,208337 | -0,511319612 | 0,133566156 | -3,828212372 | 0,000129077 | 0,001499575 |
| ENSG00000184402.15 | SS18L1 | protein_coding | 4104,541673 | -0,512162874 | 0,129970041 | -3,940622551 | 8,13E-05 | 0,000992647 |
| ENSG00000153574.9 | RPIA | protein_coding | 1097,469588 | -0,512825936 | 0,150396354 | -3,409829582 | 0,000650035 | 0,006013469 |
| ENSG00000101412.13 | E2F1 | protein_coding | 11516,94588 | -0,512932924 | 0,13684554 | -3,748261907 | 0,000178064 | 0,001997736 |
| ENSG00000147454.14 | SLC25A37 | protein_coding | 920,6390512 | -0,513848221 | 0,161103876 | -3,189545984 | 0,001424965 | 0,011777195 |
| ENSG00000113269.14 | RNF130 | protein_coding | 7742,93975 | -0,514425857 | 0,118845484 | -4,328526746 | 1,50E-05 | 0,000218637 |
| ENSG00000258986.7 | TMEM179 | protein_coding | 15405,97319 | -0,514582548 | 0,112796039 | -4,562062217 | 5,07E-06 | 8,17E-05 |
| ENSG00000164659.15 | ELAPOR2 | protein_coding | 2388,700757 | -0,515444716 | 0,127673046 | -4,037224227 | 5,41E-05 | 0,00068834 |
| ENSG00000133805.15 | AMPD3 | protein_coding | 765,3521615 | -0,515627413 | 0,170317476 | -3,027448651 | 0,002466276 | 0,01849928 |
| ENSG00000120885.22 | CLU | protein_coding | 86999,35712 | -0,516123909 | 0,148691583 | -3,471103736 | 0,000518324 | 0,004989015 |
| ENSG00000066926.12 | FECH | protein_coding | 2306,096542 | -0,516524722 | 0,149075432 | -3,464854764 | 0,000530518 | 0,005077665 |
| ENSG00000198176.13 | TFDP1 | protein_coding | 14970,1487 | -0,516654321 | 0,112869101 | -4,577464675 | 4,71E-06 | 7,64E-05 |
| ENSG00000004660.15 | CAMKK1 | protein_coding | 1589,931495 | -0,517981221 | 0,140798551 | -3,678881745 | 0,000234259 | 0,002533166 |
| ENSG00000137807.16 | KIF23 | protein_coding | 1989,152772 | -0,518591828 | 0,162428262 | -3,192743816 | 0,001409279 | 0,01165676 |
| ENSG00000135763.10 | URB2 | protein_coding | 1897,553681 | -0,518622139 | 0,133935247 | -3,872185635 | 0,000107864 | 0,001279392 |
| ENSG00000105676.14 | ARMC6 | protein_coding | 3071,671366 | -0,518981487 | 0,12506255 | -4,149775357 | 3,33E-05 | 0,000445473 |
| ENSG00000145569.6 | OTULINL | protein_coding | 11570,23803 | -0,519573503 | 0,127480089 | -4,075722791 | 4,59E-05 | 0,000592061 |
| ENSG00000134504.14 | KCTD1 | protein_coding | 1355,389058 | -0,519629695 | 0,155307629 | -3,345809216 | 0,000820428 | 0,00733376 |
| ENSG00000186020.13 | ZNF529 | protein_coding | 678,5543627 | -0,521161267 | 0,178642874 | -2,917335891 | 0,003530354 | 0,024893215 |
| ENSG00000059769.20 | DNAJC25 | protein_coding | 1035,976598 | -0,521835123 | 0,174871686 | -2,984103008 | 0,00284411 | 0,020840548 |
| ENSG00000261824.7 | LINC00662 | lncRNA | 1485,358779 | -0,521873488 | 0,158630957 | -3,289859046 | 0,001002376 | 0,008689332 |
| ENSG00000174177.13 | CTU2 | protein_coding | 896,4680354 | -0,521883554 | 0,159308287 | -3,275934761 | 0,001053129 | 0,009076679 |
| ENSG00000009830.12 | POMT2 | protein_coding | 1704,272339 | -0,521884066 | 0,147966744 | -3,527036236 | 0,000420239 | 0,004177449 |
| ENSG00000204103.4 | MAFB | protein_coding | 36940,42306 | -0,521994243 | 0,116342875 | -4,486688544 | 7,23E-06 | 0,000113185 |
| ENSG00000228623.6 | ZNF883 | transcribed_unproc | 790,4515867 | -0,522061312 | 0,175403564 | -2,97634381 | 0,002917076 | 0,021270983 |
| ENSG00000130054.5 | FAM155B | protein_coding | 1039,576025 | -0,522082702 | 0,153766757 | -3,395289809 | 0,00068556 | 0,006286551 |
| ENSG00000138448.12 | ITGAV | protein_coding | 6197,21359 | -0,522140375 | 0,1248188 | -4,183186956 | 2,87E-05 | 0,000391273 |
| ENSG00000197056.11 | ZMYM1 | protein_coding | 664,5649306 | -0,522724315 | 0,179411644 | -2,913547319 | 0,003573477 | 0,025171849 |
| ENSG00000204271.13 | SPIN3 | protein_coding | 1584,327815 | -0,523007092 | 0,156260638 | -3,347017515 | 0,00081686 | 0,007311233 |

| | | | | | | | | |
|---|---|---|---|---|---|---|---|---|
| ENSG00000181666.18 | ZNF875 | protein_coding | 1175,11951 | -0,523241983 | 0,166078019 | -3,150579387 | 0,00162947 | 0,013211831 |
| ENSG00000118762.8 | PKD2 | protein_coding | 6439,819058 | -0,52341977 | 0,123776214 | -4,228758934 | 2,35E-05 | 0,00032567 |
| ENSG00000271303.2 | SRXN1 | protein_coding | 3726,780262 | -0,524197228 | 0,123281293 | -4,252041919 | 2,12E-05 | 0,000297016 |
| ENSG00000144401.14 | METTL21A | protein_coding | 1856,791153 | -0,524243652 | 0,14341893 | -3,655330933 | 0,00025685 | 0,002737863 |
| ENSG00000166803.14 | PCLAF | protein_coding | 2654,754222 | -0,524361662 | 0,180329783 | -2,907792895 | 0,003639893 | 0,02557086 |
| ENSG00000274211.6 | SOCS7 | protein_coding | 2756,176754 | -0,524971119 | 0,154809757 | -3,391072551 | 0,000696197 | 0,006370139 |
| ENSG00000156011.17 | PSD3 | protein_coding | 10094,3931 | -0,52619635 | 0,134277649 | -3,918718811 | 8,90E-05 | 0,001079752 |
| ENSG00000130005.13 | GAMT | protein_coding | 3394,020993 | -0,527033201 | 0,146040205 | -3,608822656 | 0,00030759 | 0,003184665 |
| ENSG00000146243.14 | IRAK1BP1 | protein_coding | 927,2080805 | -0,527365313 | 0,15773045 | -3,343459142 | 0,000827409 | 0,007386696 |
| ENSG00000164237.9 | CMBL | protein_coding | 735,2230209 | -0,527778431 | 0,168675705 | -3,128953464 | 0,001754301 | 0,014006975 |
| ENSG00000175894.18 | TSPEAR | protein_coding | 918,5982618 | -0,527912177 | 0,180522315 | -2,924359674 | 0,003451657 | 0,024431474 |
| ENSG00000105926.16 | MPP6 | protein_coding | 6232,618516 | -0,52851561 | 0,117291495 | -4,506001158 | 6,61E-06 | 0,00010414 |
| ENSG00000076351.13 | SLC46A1 | protein_coding | 5720,868274 | -0,528621717 | 0,126431792 | -4,181082222 | 2,90E-05 | 0,000393889 |
| ENSG00000138160.7 | KIF11 | protein_coding | 6047,121816 | -0,528700733 | 0,1448901 | -3,648977617 | 0,000263286 | 0,002800759 |
| ENSG00000021488.13 | SLC7A9 | protein_coding | 2679,156637 | -0,528930416 | 0,133954598 | -3,948579768 | 7,86E-05 | 0,000964166 |
| ENSG00000170264.13 | FAM161A | protein_coding | 1046,073166 | -0,529942628 | 0,161063441 | -3,2902726 | 0,001000904 | 0,008682788 |
| ENSG00000101190.13 | TCFL5 | protein_coding | 1301,612629 | -0,53093146 | 0,170035526 | -3,122473708 | 0,001793381 | 0,014224191 |
| ENSG00000126803.9 | HSPA2 | protein_coding | 13060,60928 | -0,531082653 | 0,113458534 | -4,680852426 | 2,86E-06 | 4,82E-05 |
| ENSG00000182759.4 | MAFA | protein_coding | 10192,84479 | -0,531106281 | 0,136315312 | -3,896160114 | 9,77E-05 | 0,001170465 |
| ENSG00000129347.21 | KRI1 | protein_coding | 1036,506364 | -0,532098002 | 0,152862123 | -3,480901548 | 0,000499729 | 0,004850176 |
| ENSG00000221887.6 | HMSD | protein_coding | 1122,817187 | -0,532764686 | 0,149092876 | -3,573374538 | 0,00035241 | 0,003590125 |
| ENSG00000142185.16 | TRPM2 | protein_coding | 4194,454362 | -0,533176451 | 0,13461893 | -3,960635047 | 7,48E-05 | 0,000921075 |
| ENSG00000049089.15 | COL9A2 | protein_coding | 1303,201945 | -0,533287245 | 0,160160944 | -3,32969594 | 0,000869409 | 0,00769276 |
| ENSG00000166166.13 | TRMT61A | protein_coding | 961,8122514 | -0,533304609 | 0,157416151 | -3,387864617 | 0,00070439 | 0,006431055 |
| ENSG00000117597.18 | UTP25 | protein_coding | 838,3328639 | -0,533436187 | 0,162658166 | -3,279492202 | 0,001039941 | 0,008981502 |
| ENSG00000133027.18 | PEMT | protein_coding | 765,6355188 | -0,53394524 | 0,173282152 | -3,081363154 | 0,002060552 | 0,015947375 |
| ENSG00000076003.5 | MCM6 | protein_coding | 25550,99552 | -0,53395352 | 0,129450172 | -4,124780318 | 3,71E-05 | 0,000490142 |
| ENSG00000183579.16 | ZNRF3 | protein_coding | 1101,76694 | -0,534446137 | 0,19315286 | -2,766959477 | 0,005658179 | 0,03635788 |
| ENSG00000118412.13 | CASP8AP2 | protein_coding | 2484,792043 | -0,534660846 | 0,136696307 | -3,911304259 | 9,18E-05 | 0,001110874 |
| ENSG00000103202.13 | NME4 | protein_coding | 4256,353789 | -0,534755971 | 0,153377069 | -3,486544459 | 0,000489304 | 0,004757817 |
| ENSG00000152076.19 | CCDC74B | protein_coding | 586,094292 | -0,535238787 | 0,198257829 | -2,699710718 | 0,006939979 | 0,042895915 |
| ENSG00000123219.13 | CENPK | protein_coding | 2336,234043 | -0,535442828 | 0,147971977 | -3,618542101 | 0,000296267 | 0,003085749 |
| ENSG00000148158.17 | SNX30 | protein_coding | 2408,362651 | -0,53549871 | 0,154271326 | -3,471148689 | 0,000518237 | 0,004989015 |
| ENSG00000255717.7 | SNHG1 | lncRNA | 1846,843518 | -0,535647312 | 0,14994416 | -3,572311933 | 0,000353844 | 0,003601223 |
| ENSG00000138031.14 | ADCY3 | protein_coding | 619,6206227 | -0,535826378 | 0,183131825 | -2,925905299 | 0,003434555 | 0,024340692 |
| ENSG00000102384.13 | CENPI | protein_coding | 874,6238371 | -0,53672952 | 0,195880002 | -2,740093495 | 0,006142171 | 0,038923681 |

| | | | | | | | | |
|---|---|---|---|---|---|---|---|---|
| ENSG00000170312.16 | CDK1 | protein_coding | 10707,91474 | -0,536813108 | 0,139251282 | -3,854995797 | 0,000115732 | 0,001358095 |
| ENSG00000107518.18 | ATRNL1 | protein_coding | 3281,395706 | -0,537110095 | 0,124182386 | -4,325171315 | 1,52E-05 | 0,000221121 |
| ENSG00000089057.15 | SLC23A2 | protein_coding | 5344,57007 | -0,538353516 | 0,123737291 | -4,350778257 | 1,36E-05 | 0,000199574 |
| ENSG00000223891.6 | OSER1-DT | lncRNA | 561,3627618 | -0,539376159 | 0,199206651 | -2,707621235 | 0,006776731 | 0,042060739 |
| ENSG00000167523.14 | SPATA33 | protein_coding | 1457,836053 | -0,540569342 | 0,155614057 | -3,473782199 | 0,000513177 | 0,004950861 |
| ENSG00000068383.19 | INPP5A | protein_coding | 526,6384799 | -0,542446832 | 0,197820113 | -2,742121738 | 0,006104371 | 0,038725762 |
| ENSG00000156802.13 | ATAD2 | protein_coding | 10336,15458 | -0,542636733 | 0,14546916 | -3,730252737 | 0,000191288 | 0,002123335 |
| ENSG00000073536.18 | NLE1 | protein_coding | 801,6613959 | -0,543038916 | 0,182439361 | -2,976544721 | 0,002915166 | 0,021267508 |
| ENSG00000156475.18 | PPP2R2B | protein_coding | 1062,554608 | -0,543086125 | 0,166924602 | -3,253481625 | 0,001140001 | 0,009709488 |
| ENSG00000167536.14 | DHRS13 | protein_coding | 4851,301718 | -0,543170185 | 0,123904291 | -4,383788335 | 1,17E-05 | 0,000175285 |
| ENSG00000145882.11 | PCYOX1L | protein_coding | 3833,617751 | -0,543418876 | 0,130022866 | -4,179410088 | 2,92E-05 | 0,000396282 |
| ENSG00000164338.10 | UTP15 | protein_coding | 1117,684407 | -0,543779275 | 0,160725738 | -3,383274396 | 0,00071627 | 0,006528132 |
| ENSG00000135363.12 | LMO2 | protein_coding | 1763,709585 | -0,544247838 | 0,177470677 | -3,066691624 | 0,00216442 | 0,016628308 |
| ENSG00000279516.2 | FAM230C | lncRNA | 646,1205046 | -0,5444375 | 0,181593018 | -2,998119122 | 0,002716515 | 0,020045906 |
| ENSG00000154917.11 | RAB6B | protein_coding | 1469,631691 | -0,545342637 | 0,140073027 | -3,893273743 | 9,89E-05 | 0,001182456 |
| ENSG00000130508.11 | PXDN | protein_coding | 1210,150568 | -0,546030376 | 0,174992008 | -3,120316069 | 0,001806571 | 0,014315125 |
| ENSG00000077514.9 | POLD3 | protein_coding | 2089,054677 | -0,546073705 | 0,152388802 | -3,5834241 | 0,000339119 | 0,003466535 |
| ENSG00000084774.14 | CAD | protein_coding | 3817,753003 | -0,546154613 | 0,156075948 | -3,499287481 | 0,000466503 | 0,004555152 |
| ENSG00000196549.12 | MME | protein_coding | 8263,573267 | -0,546224209 | 0,143261882 | -3,812767238 | 0,00013742 | 0,001582441 |
| ENSG00000076248.11 | UNG | protein_coding | 5426,081839 | -0,546258126 | 0,139747804 | -3,908885231 | 9,27E-05 | 0,001120113 |
| ENSG00000197763.19 | TXNRD3 | protein_coding | 895,155302 | -0,546919643 | 0,157722548 | -3,46760592 | 0,000525117 | 0,005033571 |
| ENSG00000113391.19 | FAM172A | protein_coding | 2940,818196 | -0,547035194 | 0,123745413 | -4,420650284 | 9,84E-06 | 0,000149934 |
| ENSG00000181544.15 | FANCB | protein_coding | 511,2874594 | -0,547071126 | 0,193226051 | -2,831249329 | 0,004636656 | 0,031069023 |
| ENSG00000166263.13 | STXBP4 | protein_coding | 6493,709564 | -0,547227071 | 0,11827023 | -4,626921498 | 3,71E-06 | 6,16E-05 |
| ENSG00000177990.12 | DPY19L2 | protein_coding | 1128,063835 | -0,548628404 | 0,160835495 | -3,411115223 | 0,000646977 | 0,005997438 |
| ENSG00000137814.12 | HAUS2 | protein_coding | 1960,923284 | -0,548673915 | 0,144627175 | -3,793712454 | 0,000148412 | 0,001694109 |
| ENSG00000162066.16 | AMDHD2 | protein_coding | 2329,750601 | -0,549422188 | 0,160929187 | -3,414061799 | 0,000640021 | 0,005947106 |
| ENSG00000165731.20 | RET | protein_coding | 1832,778818 | -0,549544402 | 0,178997556 | -3,070122384 | 0,002139711 | 0,016486877 |
| ENSG00000258754.8 | LINC01579 | lncRNA | 410,247442 | -0,54975917 | 0,200890805 | -2,736606936 | 0,006207642 | 0,039221196 |
| ENSG00000184602.6 | SNN | protein_coding | 5271,289741 | -0,550194134 | 0,131589035 | -4,181154871 | 2,90E-05 | 0,000393889 |
| ENSG00000051180.17 | RAD51 | protein_coding | 988,0087847 | -0,550319823 | 0,199027048 | -2,765050417 | 0,0056914 | 0,036537705 |
| ENSG00000071564.17 | TCF3 | protein_coding | 2462,961494 | -0,550606729 | 0,17594257 | -3,12946849 | 0,001751229 | 0,013998461 |
| ENSG00000163704.12 | PRRT3 | protein_coding | 2618,437695 | -0,550732608 | 0,1448981 | -3,800826982 | 0,000144214 | 0,001651597 |
| ENSG00000183775.11 | KCTD16 | protein_coding | 783,2995959 | -0,550757236 | 0,170893726 | -3,222805475 | 0,001269417 | 0,010711508 |
| ENSG00000131470.15 | PSMC3IP | protein_coding | 1604,49978 | -0,551203104 | 0,154357485 | -3,570951572 | 0,000355687 | 0,003618222 |
| ENSG00000118495.20 | PLAGL1 | protein_coding | 42038,0122 | -0,552490903 | 0,121224585 | -4,557581306 | 5,17E-06 | 8,33E-05 |

| | | | | | | | | |
|---|---|---|---|---|---|---|---|---|
| ENSG00000068654.16 | POLR1A | protein_coding | 3660,342423 | -0,552931314 | 0,135484807 | -4,081131512 | 4,48E-05 | 0,00058166 |
| ENSG00000107262.26 | BAG1 | protein_coding | 4790,370626 | -0,553210291 | 0,146059856 | -3,787558786 | 0,000152135 | 0,001730946 |
| ENSG00000136492.9 | BRIP1 | protein_coding | 1342,093699 | -0,553667742 | 0,186733296 | -2,965018849 | 0,003026644 | 0,021871864 |
| ENSG00000266338.6 | NBPF15 | protein_coding | 1205,10802 | -0,554397663 | 0,150766126 | -3,677203089 | 0,000235805 | 0,002548572 |
| ENSG00000104738.18 | MCM4 | protein_coding | 15074,60895 | -0,55441571 | 0,144794759 | -3,828976374 | 0,000128677 | 0,001495758 |
| ENSG00000177191.2 | B3GNT8 | protein_coding | 1585,021727 | -0,554605324 | 0,166534737 | -3,330268111 | 0,000867624 | 0,007686715 |
| ENSG00000240445.6 | FOXO3B | protein_coding | 583,3194697 | -0,555042249 | 0,184238581 | -3,01262769 | 0,002589965 | 0,019280194 |
| ENSG00000122966.17 | CIT | protein_coding | 1333,525251 | -0,555454062 | 0,193765044 | -2,8666371 | 0,004148584 | 0,028533003 |
| ENSG00000066032.18 | CTNNA2 | protein_coding | 1185,386264 | -0,555649224 | 0,149458116 | -3,717758796 | 0,000200998 | 0,002217015 |
| ENSG00000070950.10 | RAD18 | protein_coding | 1461,795207 | -0,555876541 | 0,154171527 | -3,605571993 | 0,000311466 | 0,003220021 |
| ENSG00000187231.14 | SESTD1 | protein_coding | 12802,55553 | -0,556373831 | 0,132551263 | -4,197423829 | 2,70E-05 | 0,000370124 |
| ENSG00000100483.14 | VCPKMT | protein_coding | 831,0376649 | -0,556690073 | 0,168830571 | -3,297329797 | 0,000976088 | 0,008500169 |
| ENSG00000174358.16 | SLC6A19 | protein_coding | 3200,47408 | -0,557143301 | 0,12264717 | -4,5426511 | 5,56E-06 | 8,89E-05 |
| ENSG00000165905.18 | LARGE2 | protein_coding | 2775,769941 | -0,557168917 | 0,131104028 | -4,249823013 | 2,14E-05 | 0,000299187 |
| ENSG00000185818.8 | NAT8L | protein_coding | 506,4793607 | -0,557304051 | 0,200290455 | -2,782479329 | 0,00539453 | 0,035072823 |
| ENSG00000148019.14 | CEP78 | protein_coding | 3347,398729 | -0,557599129 | 0,131388961 | -4,243881111 | 2,20E-05 | 0,000305794 |
| ENSG00000141480.18 | ARRB2 | protein_coding | 2181,802013 | -0,557747251 | 0,141640637 | -3,937762942 | 8,22E-05 | 0,001003962 |
| ENSG00000160180.15 | TFF3 | protein_coding | 1051,415539 | -0,558322297 | 0,211380962 | -2,641308342 | 0,008258652 | 0,049271837 |
| ENSG00000179743.4 | FLJ37453 | lncRNA | 1227,400991 | -0,558626776 | 0,16930966 | -3,299438302 | 0,000968785 | 0,008454155 |
| ENSG00000184307.16 | ZDHHC23 | protein_coding | 2685,37578 | -0,558826617 | 0,126036198 | -4,433858102 | 9,26E-06 | 0,00014165 |
| ENSG00000170629.14 | DPY19L2P2 | transcribed_unproc | 2155,206037 | -0,559047821 | 0,132522284 | -4,218519366 | 2,46E-05 | 0,000339141 |
| ENSG00000188312.14 | CENPP | protein_coding | 1068,658955 | -0,559205638 | 0,190504089 | -2,935399666 | 0,003331183 | 0,023728588 |
| ENSG00000015532.10 | XYLT2 | protein_coding | 752,9737215 | -0,561151132 | 0,18638554 | -3,010701002 | 0,002606454 | 0,019377169 |
| ENSG00000168393.13 | DTYMK | protein_coding | 4297,721972 | -0,561476771 | 0,146092386 | -3,843299347 | 0,000121391 | 0,001420529 |
| ENSG00000166228.9 | PCBD1 | protein_coding | 11125,41021 | -0,561606164 | 0,124208168 | -4,521491425 | 6,14E-06 | 9,72E-05 |
| ENSG00000153395.10 | LPCAT1 | protein_coding | 12683,45193 | -0,561710295 | 0,113595735 | -4,944818533 | 7,62E-07 | 1,42E-05 |
| ENSG00000115652.15 | UXS1 | protein_coding | 3477,349611 | -0,561931298 | 0,129089317 | -4,353042628 | 1,34E-05 | 0,00019794 |
| ENSG00000091129.21 | NRCAM | protein_coding | 14042,71725 | -0,562137872 | 0,132551769 | -4,240893015 | 2,23E-05 | 0,000309688 |
| ENSG00000143816.8 | WNT9A | protein_coding | 560,6845313 | -0,562602499 | 0,17941925 | -3,135686384 | 0,001714524 | 0,01377117 |
| ENSG00000101447.15 | FAM83D | protein_coding | 1616,904056 | -0,56303972 | 0,161662761 | -3,482804053 | 0,000496191 | 0,004818074 |
| ENSG00000179029.15 | TMEM107 | protein_coding | 1689,391517 | -0,564255487 | 0,166412162 | -3,390710637 | 0,000697117 | 0,006372985 |
| ENSG00000227372.12 | TP73-AS1 | transcribed_unitary | 2988,376303 | -0,56431293 | 0,128778099 | -4,382056685 | 1,18E-05 | 0,000176557 |
| ENSG00000187498.16 | COL4A1 | protein_coding | 507,4851047 | -0,564610477 | 0,187727165 | -3,00761202 | 0,002633091 | 0,019519742 |
| ENSG00000248383.5 | PCDHAC1 | protein_coding | 392,001736 | -0,564877066 | 0,210849703 | -2,679050804 | 0,007383119 | 0,044984167 |
| ENSG00000161267.12 | BDH1 | protein_coding | 1663,761561 | -0,565467233 | 0,137848923 | -4,102079427 | 4,09E-05 | 0,000535076 |
| ENSG00000134901.13 | POGLUT2 | protein_coding | 1845,095307 | -0,566663212 | 0,136807256 | -4,142055246 | 3,44E-05 | 0,000458104 |

| | | | | | | | | |
|---|---|---|---|---|---|---|---|---|
| ENSG00000153558.16 | FBXL2 | protein_coding | 679,4595857 | -0,567551528 | 0,195956597 | -2,896312428 | 0,003775763 | 0,026410156 |
| ENSG00000234171.3 | RNASEH1-AS1 | lncRNA | 1643,079097 | -0,568594265 | 0,185979768 | -3,057290974 | 0,002233474 | 0,017071109 |
| ENSG00000112759.19 | SLC29A1 | protein_coding | 39640,50666 | -0,568629006 | 0,139493912 | -4,076371482 | 4,57E-05 | 0,000591141 |
| ENSG00000164109.14 | MAD2L1 | protein_coding | 5663,399014 | -0,568658105 | 0,149126576 | -3,813257973 | 0,000137147 | 0,00158017 |
| ENSG00000182253.15 | SYNM | protein_coding | 1682,200622 | -0,569493632 | 0,152192549 | -3,741928466 | 0,000182614 | 0,002041118 |
| ENSG00000198785.7 | GRIN3A | protein_coding | 524,0519263 | -0,569909886 | 0,203115644 | -2,805839441 | 0,005018569 | 0,033028528 |
| ENSG00000140263.15 | SORD | protein_coding | 8728,702403 | -0,57008872 | 0,131548143 | -4,333688843 | 1,47E-05 | 0,000213769 |
| ENSG00000178104.19 | PDE4DIP | protein_coding | 28097,51037 | -0,570205027 | 0,13027885 | -4,376804272 | 1,20E-05 | 0,000180089 |
| ENSG00000178162.8 | FAR2P2 | transcribed_unproc | 2063,373552 | -0,570450104 | 0,13439188 | -4,244676866 | 2,19E-05 | 0,000304914 |
| ENSG00000112394.17 | SLC16A10 | protein_coding | 3758,064071 | -0,57060085 | 0,120910039 | -4,719218136 | 2,37E-06 | 4,05E-05 |
| ENSG00000063660.9 | GPC1 | protein_coding | 3921,685088 | -0,570691828 | 0,136043196 | -4,194931044 | 2,73E-05 | 0,00037324 |
| ENSG00000272106.1 | AL691432.2 | lncRNA | 767,0813123 | -0,570979379 | 0,167425336 | -3,410352305 | 0,00064879 | 0,006006355 |
| ENSG00000285865.1 | AC010285.3 | lncRNA | 606,3233728 | -0,571388551 | 0,216846598 | -2,634989699 | 0,008413984 | 0,049885075 |
| ENSG00000146802.13 | TMEM168 | protein_coding | 1969,596305 | -0,571797788 | 0,133991525 | -4,267417569 | 1,98E-05 | 0,000279532 |
| ENSG00000116991.11 | SIPA1L2 | protein_coding | 2066,395599 | -0,572123497 | 0,153656054 | -3,723403544 | 0,000196555 | 0,002173734 |
| ENSG00000175548.9 | ALG10B | protein_coding | 1702,722457 | -0,57241477 | 0,164787825 | -3,473647224 | 0,000513436 | 0,004951071 |
| ENSG00000158457.6 | TSPAN33 | protein_coding | 6871,753655 | -0,572424306 | 0,119785067 | -4,778761813 | 1,76E-06 | 3,11E-05 |
| ENSG00000109956.13 | B3GAT1 | protein_coding | 954,4759672 | -0,572640137 | 0,197209062 | -2,903721213 | 0,003687564 | 0,025871027 |
| ENSG00000181751.10 | MACIR | protein_coding | 4202,045841 | -0,574134713 | 0,130314945 | -4,405747252 | 1,05E-05 | 0,000159809 |
| ENSG00000142661.19 | MYOM3 | protein_coding | 1708,954507 | -0,574247161 | 0,161109796 | -3,564321822 | 0,000364798 | 0,003692181 |
| ENSG00000143476.18 | DTL | protein_coding | 4155,380414 | -0,575223626 | 0,14904636 | -3,859360445 | 0,000113684 | 0,001340078 |
| ENSG00000165802.23 | NSMF | protein_coding | 4427,174609 | -0,57544966 | 0,130837951 | -4,398186129 | 1,09E-05 | 0,000164762 |
| ENSG00000274828.1 | AC068473.5 | lncRNA | 728,6824966 | -0,576009015 | 0,17406108 | -3,309234987 | 0,000935513 | 0,008204843 |
| ENSG00000187801.15 | ZFP69B | protein_coding | 447,6726086 | -0,577664205 | 0,196068013 | -2,946243984 | 0,003216586 | 0,023053482 |
| ENSG00000010278.15 | CD9 | protein_coding | 876,7406835 | -0,578081737 | 0,188607786 | -3,064994017 | 0,002176744 | 0,016709548 |
| ENSG00000115526.11 | CHST10 | protein_coding | 2120,612465 | -0,579132472 | 0,131126331 | -4,416599364 | 1,00E-05 | 0,000152548 |
| ENSG00000173894.11 | CBX2 | protein_coding | 3072,988631 | -0,580201674 | 0,128070907 | -4,530315945 | 5,89E-06 | 9,35E-05 |
| ENSG00000158747.15 | NBL1 | protein_coding | 1131,745204 | -0,580472502 | 0,155335555 | -3,736893995 | 0,000186307 | 0,002076861 |
| ENSG00000176349.11 | AC104129.1 | lncRNA | 540,5956701 | -0,580772054 | 0,195263099 | -2,974305214 | 0,002936528 | 0,021390473 |
| ENSG00000282164.3 | PEG13 | lncRNA | 469,8437122 | -0,581332037 | 0,201713874 | -2,881963575 | 0,003952054 | 0,027405185 |
| ENSG00000169783.13 | LINGO1 | protein_coding | 1924,146574 | -0,58211005 | 0,175197006 | -3,322602736 | 0,000891818 | 0,007879106 |
| ENSG00000183763.9 | TRAIP | protein_coding | 636,0486654 | -0,582149504 | 0,217633861 | -2,674903163 | 0,007475084 | 0,045438699 |
| ENSG00000110245.12 | APOC3 | protein_coding | 1547,022739 | -0,583045256 | 0,160818529 | -3,625485559 | 0,000288419 | 0,003019026 |
| ENSG00000001084.13 | GCLC | protein_coding | 3283,033459 | -0,583164366 | 0,124364969 | -4,6891369 | 2,74E-06 | 4,63E-05 |
| ENSG00000185559.16 | DLK1 | protein_coding | 781,3245593 | -0,583793773 | 0,20050722 | -2,911584793 | 0,003596003 | 0,025302819 |
| ENSG00000099904.16 | ZDHHC8 | protein_coding | 904,781158 | -0,584015626 | 0,219363079 | -2,662324161 | 0,00776031 | 0,046792079 |

| | | | | | | | | |
|---|---|---|---|---|---|---|---|---|
| ENSG00000170745.12 | KCNS3 | protein_coding | 414,5330641 | -0,584579263 | 0,214557569 | -2,724580003 | 0,006438333 | 0,040403627 |
| ENSG00000006634.8 | DBF4 | protein_coding | 2026,240529 | -0,585067228 | 0,138887405 | -4,212529061 | 2,53E-05 | 0,000348035 |
| ENSG00000189057.11 | FAM111B | protein_coding | 5283,853891 | -0,585403893 | 0,168818268 | -3,467657266 | 0,000525016 | 0,005033571 |
| ENSG00000152208.13 | GRID2 | protein_coding | 1593,30668 | -0,586092939 | 0,143330913 | -4,089089557 | 4,33E-05 | 0,000564176 |
| ENSG00000198113.3 | TOR4A | protein_coding | 1959,092001 | -0,586220022 | 0,163479529 | -3,58589252 | 0,000335927 | 0,003435583 |
| ENSG00000157870.17 | PRXL2B | protein_coding | 4863,981471 | -0,58747128 | 0,122011728 | -4,814875506 | 1,47E-06 | 2,63E-05 |
| ENSG00000165244.7 | ZNF367 | protein_coding | 3355,477147 | -0,587523897 | 0,149600204 | -3,92729343 | 8,59E-05 | 0,00104562 |
| ENSG00000132405.19 | TBC1D14 | protein_coding | 7417,826308 | -0,58781127 | 0,118011392 | -4,980970582 | 6,33E-07 | 1,20E-05 |
| ENSG00000166224.17 | SGPL1 | protein_coding | 8678,808506 | -0,588185417 | 0,125084333 | -4,702310868 | 2,57E-06 | 4,36E-05 |
| ENSG00000179314.14 | WSCD1 | protein_coding | 5590,447079 | -0,58915446 | 0,125933223 | -4,678308428 | 2,89E-06 | 4,87E-05 |
| ENSG00000180354.16 | MTURN | protein_coding | 1611,82212 | -0,589955927 | 0,139554634 | -4,227419111 | 2,36E-05 | 0,000327299 |
| ENSG00000093009.11 | CDC45 | protein_coding | 1866,008798 | -0,589969968 | 0,188777776 | -3,125208799 | 0,001776789 | 0,014137999 |
| ENSG00000143819.12 | EPHX1 | protein_coding | 27485,72146 | -0,590063855 | 0,136527179 | -4,32195157 | 1,55E-05 | 0,000223444 |
| ENSG00000165730.16 | STOX1 | protein_coding | 1047,711062 | -0,590778001 | 0,162138036 | -3,643673102 | 0,000268775 | 0,00284757 |
| ENSG00000215190.9 | LINC00680 | transcribed_unproc | 676,6200827 | -0,591002374 | 0,173088498 | -3,41445204 | 0,000639104 | 0,005941825 |
| ENSG00000183963.19 | SMTN | protein_coding | 1579,766186 | -0,591110527 | 0,19692716 | -3,001670902 | 0,002685023 | 0,019827496 |
| ENSG00000106617.15 | PRKAG2 | protein_coding | 2900,946696 | -0,591551896 | 0,143483532 | -4,122785989 | 3,74E-05 | 0,000493783 |
| ENSG00000173992.9 | CCS | protein_coding | 2676,212548 | -0,591572785 | 0,137227715 | -4,310884189 | 1,63E-05 | 0,000233519 |
| ENSG00000143473.13 | KCNH1 | protein_coding | 381,720433 | -0,591672533 | 0,208740184 | -2,834492729 | 0,00458985 | 0,030839607 |
| ENSG00000156970.13 | BUB1B | protein_coding | 3336,439454 | -0,593379317 | 0,160649745 | -3,693621276 | 0,000221083 | 0,002420697 |
| ENSG00000112742.10 | TTK | protein_coding | 2359,272887 | -0,593536788 | 0,17453377 | -3,400698839 | 0,000672138 | 0,00618515 |
| ENSG00000177225.17 | GATD1 | protein_coding | 1213,97159 | -0,594896736 | 0,220478149 | -2,69821177 | 0,006971308 | 0,04302604 |
| ENSG00000134184.13 | GSTM1 | protein_coding | 897,3695382 | -0,59598065 | 0,166158766 | -3,586814367 | 0,000334742 | 0,003425455 |
| ENSG00000075131.10 | TIPIN | protein_coding | 1031,872985 | -0,596311553 | 0,1505087 | -3,961973979 | 7,43E-05 | 0,000917003 |
| ENSG00000114251.15 | WNT5A | protein_coding | 2115,476223 | -0,597234316 | 0,146618412 | -4,073392343 | 4,63E-05 | 0,000597651 |
| ENSG00000124145.6 | SDC4 | protein_coding | 2064,625821 | -0,597297686 | 0,144700752 | -4,127813289 | 3,66E-05 | 0,000485254 |
| ENSG00000172167.8 | MTBP | protein_coding | 378,1683071 | -0,597435983 | 0,222044993 | -2,690607767 | 0,0071322 | 0,043761019 |
| ENSG00000040275.17 | SPDL1 | protein_coding | 2580,048837 | -0,597969749 | 0,129690719 | -4,610736631 | 4,01E-06 | 6,61E-05 |
| ENSG00000135324.6 | MRAP2 | protein_coding | 1240,744265 | -0,598236276 | 0,146738406 | -4,076889564 | 4,56E-05 | 0,000590191 |
| ENSG00000075303.13 | SLC25A40 | protein_coding | 2988,333684 | -0,598337048 | 0,126681959 | -4,723143308 | 2,32E-06 | 3,98E-05 |
| ENSG00000101417.12 | PXMP4 | protein_coding | 1433,376841 | -0,599079498 | 0,171767282 | -3,487739283 | 0,000487123 | 0,004741012 |
| ENSG00000134291.12 | TMEM106C | protein_coding | 20843,17957 | -0,60104995 | 0,155505998 | -3,865123905 | 0,000111033 | 0,00131252 |
| ENSG00000144320.14 | LNPK | protein_coding | 9474,071454 | -0,601938666 | 0,121144551 | -4,968763875 | 6,74E-07 | 1,27E-05 |
| ENSG00000138604.10 | GLCE | protein_coding | 2591,228822 | -0,602286036 | 0,13159417 | -4,576844357 | 4,72E-06 | 7,66E-05 |
| ENSG00000085276.19 | MECOM | protein_coding | 3481,245051 | -0,602537663 | 0,131272749 | -4,58996758 | 4,43E-06 | 7,26E-05 |
| ENSG00000154639.19 | CXADR | protein_coding | 3858,928385 | -0,603166534 | 0,122027923 | -4,942856672 | 7,70E-07 | 1,43E-05 |

| | | | | | | | | |
|---|---|---|---|---|---|---|---|---|
| ENSG00000137804.14 | NUSAP1 | protein_coding | 8334,062911 | -0,6058236 | 0,173842769 | -3,48489386 | 0,000492332 | 0,00478282 |
| ENSG00000168071.22 | CCDC88B | protein_coding | 392,0257787 | -0,606003421 | 0,210604758 | -2,877444113 | 0,004009109 | 0,027736517 |
| ENSG00000123179.14 | EBPL | protein_coding | 1947,893214 | -0,606012092 | 0,156891956 | -3,862607784 | 0,000112183 | 0,00132462 |
| ENSG00000198917.13 | SPOUT1 | protein_coding | 1511,426777 | -0,606393734 | 0,22659827 | -2,676073984 | 0,00744902 | 0,045306576 |
| ENSG00000154839.10 | SKA1 | protein_coding | 1494,857257 | -0,606448892 | 0,196522869 | -3,085894769 | 0,002029406 | 0,015762621 |
| ENSG00000157399.16 | ARSL | protein_coding | 3124,700101 | -0,606494149 | 0,150693712 | -4,024681208 | 5,71E-05 | 0,000721253 |
| ENSG00000132938.21 | MTUS2 | protein_coding | 2406,384786 | -0,606608821 | 0,164161171 | -3,695202806 | 0,000219711 | 0,002411986 |
| ENSG00000187244.12 | BCAM | protein_coding | 12750,09943 | -0,606802719 | 0,177359879 | -3,42130769 | 0,000623208 | 0,005827091 |
| ENSG00000204856.12 | FAM216A | protein_coding | 326,1691566 | -0,607469655 | 0,226312698 | -2,684204906 | 0,007270253 | 0,044412825 |
| ENSG00000162804.14 | SNED1 | protein_coding | 1400,773487 | -0,607624923 | 0,21837432 | -2,782492571 | 0,00539431 | 0,035072823 |
| ENSG00000166821.9 | PEX11A | protein_coding | 740,6027025 | -0,60830839 | 0,178346916 | -3,410815305 | 0,000647689 | 0,005999725 |
| ENSG00000119471.15 | HSDL2 | protein_coding | 6211,311899 | -0,608885677 | 0,122882127 | -4,955038542 | 7,23E-07 | 1,36E-05 |
| ENSG00000091651.9 | ORC6 | protein_coding | 1432,306669 | -0,609121493 | 0,14341538 | -4,247253627 | 2,16E-05 | 0,000302435 |
| ENSG00000102048.16 | ASB9 | protein_coding | 2335,569021 | -0,609292997 | 0,16878563 | -3,609862973 | 0,000306359 | 0,003178207 |
| ENSG00000164687.11 | FABP5 | protein_coding | 1721,935132 | -0,609307537 | 0,200938007 | -3,032316019 | 0,00242685 | 0,018262438 |
| ENSG00000100479.13 | POLE2 | protein_coding | 971,3250964 | -0,609869589 | 0,172611091 | -3,533200482 | 0,000410561 | 0,004098758 |
| ENSG00000198093.11 | ZNF649 | protein_coding | 770,2473594 | -0,610274214 | 0,173836875 | -3,510614275 | 0,000447073 | 0,004416925 |
| ENSG00000057019.16 | DCBLD2 | protein_coding | 998,1492443 | -0,610350849 | 0,16980874 | -3,594342948 | 0,000325211 | 0,003335765 |
| ENSG00000164989.17 | CCDC171 | protein_coding | 426,6464717 | -0,610497479 | 0,20092303 | -3,038464433 | 0,002377872 | 0,017971388 |
| ENSG00000100505.13 | TRIM9 | protein_coding | 653,7168721 | -0,610879115 | 0,181311116 | -3,369231457 | 0,000753781 | 0,006825435 |
| ENSG00000169607.13 | CKAP2L | protein_coding | 2182,954163 | -0,611424096 | 0,13334993 | -4,585110005 | 4,54E-06 | 7,39E-05 |
| ENSG00000118600.12 | RXYLT1 | protein_coding | 3808,815434 | -0,611983622 | 0,136005368 | -4,499701978 | 6,80E-06 | 0,000106952 |
| ENSG00000140443.15 | IGF1R | protein_coding | 3956,463961 | -0,612315843 | 0,203660342 | -3,006554131 | 0,00264227 | 0,019560286 |
| ENSG00000184319.16 | RPL23AP82 | transcribed_unproc | 1363,711236 | -0,612450231 | 0,170751614 | -3,586790288 | 0,000334773 | 0,003425455 |
| ENSG00000109265.14 | CRACD | protein_coding | 572,8848275 | -0,612784291 | 0,229464242 | -2,670500137 | 0,007573834 | 0,045892392 |
| ENSG00000175061.18 | SNHG29 | lncRNA | 18840,3611 | -0,613352213 | 0,147653451 | -4,153998497 | 3,27E-05 | 0,000438167 |
| ENSG00000075651.16 | PLD1 | protein_coding | 1456,870013 | -0,613686245 | 0,146233753 | -4,196611476 | 2,71E-05 | 0,000370967 |
| ENSG00000100079.7 | LGALS2 | protein_coding | 565,302324 | -0,613814408 | 0,187652415 | -3,271017904 | 0,001071611 | 0,009201879 |
| ENSG00000269190.6 | FBXO17 | protein_coding | 5038,511864 | -0,613818007 | 0,125049493 | -4,908600533 | 9,17E-07 | 1,69E-05 |
| ENSG00000157388.19 | CACNA1D | protein_coding | 2549,603804 | -0,613835194 | 0,127631077 | -4,809449284 | 1,51E-06 | 2,70E-05 |
| ENSG00000228794.10 | LINC01128 | lncRNA | 2978,226518 | -0,615379003 | 0,137118644 | -4,487930928 | 7,19E-06 | 0,000112695 |
| ENSG00000142528.16 | ZNF473 | protein_coding | 2334,934148 | -0,615775377 | 0,138263501 | -4,453636529 | 8,44E-06 | 0,000130251 |
| ENSG00000175497.17 | DPP10 | protein_coding | 6226,446318 | -0,616480178 | 0,123950844 | -4,973585959 | 6,57E-07 | 1,24E-05 |
| ENSG00000181754.7 | AMIGO1 | protein_coding | 2942,44581 | -0,61669094 | 0,142500033 | -4,327654718 | 1,51E-05 | 0,000219251 |
| ENSG00000280002.1 | AC078925.4 | TEC | 706,8507226 | -0,616905427 | 0,201206904 | -3,066025141 | 0,002169251 | 0,016659305 |
| ENSG00000060656.20 | PTPRU | protein_coding | 12196,92385 | -0,617461139 | 0,154362238 | -4,000078952 | 6,33E-05 | 0,000792842 |

| | | | | | | | | |
|---|---|---|---|---|---|---|---|---|
| ENSG00000164535.15 | DAGLB | protein_coding | 1385,166664 | -0,617633718 | 0,157735223 | -3,915636012 | 9,02E-05 | 0,001093011 |
| ENSG00000171848.16 | RRM2 | protein_coding | 41246,13027 | -0,617917114 | 0,152242986 | -4,058755888 | 4,93E-05 | 0,000633635 |
| ENSG00000157833.13 | GAREM2 | protein_coding | 1371,173641 | -0,618473337 | 0,155957645 | -3,965649365 | 7,32E-05 | 0,000904585 |
| ENSG00000100304.13 | TTLL12 | protein_coding | 5328,048431 | -0,618740662 | 0,116452816 | -5,313230579 | 1,08E-07 | 2,29E-06 |
| ENSG00000112033.14 | PPARD | protein_coding | 997,0995696 | -0,618821515 | 0,176741641 | -3,501277406 | 0,000463034 | 0,004529724 |
| ENSG00000100206.10 | DMC1 | protein_coding | 1400,462673 | -0,618866628 | 0,149361883 | -4,143404026 | 3,42E-05 | 0,000455707 |
| ENSG00000151715.8 | TMEM45B | protein_coding | 422,3055798 | -0,619357393 | 0,224557095 | -2,758128812 | 0,005813329 | 0,037162929 |
| ENSG00000149328.15 | GLB1L2 | protein_coding | 3399,522334 | -0,620369389 | 0,124525118 | -4,981881551 | 6,30E-07 | 1,20E-05 |
| ENSG00000261713.6 | SSTR5-AS1 | lncRNA | 1573,762896 | -0,620875683 | 0,16482916 | -3,766783039 | 0,000165365 | 0,001868272 |
| ENSG00000180879.14 | SSR4 | protein_coding | 16017,38694 | -0,621147582 | 0,125525734 | -4,948368434 | 7,48E-07 | 1,40E-05 |
| ENSG00000198046.12 | ZNF667 | protein_coding | 577,9386214 | -0,622828871 | 0,182587132 | -3,411132346 | 0,000646937 | 0,005997438 |
| ENSG00000012779.11 | ALOX5 | protein_coding | 2128,224247 | -0,623949647 | 0,139057242 | -4,486998571 | 7,22E-06 | 0,000113105 |
| ENSG00000161133.16 | USP41 | protein_coding | 342,4273278 | -0,624754564 | 0,224495107 | -2,782931764 | 0,005387013 | 0,035056614 |
| ENSG00000148985.20 | PGAP2 | protein_coding | 924,5187324 | -0,624927822 | 0,156590221 | -3,990848333 | 6,58E-05 | 0,00082042 |
| ENSG00000135776.5 | ABCB10 | protein_coding | 4914,180282 | -0,625479895 | 0,125133932 | -4,99848349 | 5,78E-07 | 1,10E-05 |
| ENSG00000134508.12 | CABLES1 | protein_coding | 2545,879036 | -0,626921184 | 0,129230806 | -4,851174456 | 1,23E-06 | 2,22E-05 |
| ENSG00000259642.2 | ST20-AS1 | lncRNA | 673,164714 | -0,627473729 | 0,190078293 | -3,301133017 | 0,000962952 | 0,008410264 |
| ENSG00000186193.9 | SAPCD2 | protein_coding | 3254,408929 | -0,62765829 | 0,14425727 | -4,350964693 | 1,36E-05 | 0,000199545 |
| ENSG00000107833.10 | NPM3 | protein_coding | 979,9350417 | -0,628145724 | 0,171751779 | -3,657288014 | 0,000254898 | 0,002719821 |
| ENSG00000125454.12 | SLC25A19 | protein_coding | 912,7494619 | -0,628236626 | 0,158308803 | -3,968425091 | 7,23E-05 | 0,000894643 |
| ENSG00000008513.16 | ST3GAL1 | protein_coding | 1505,559644 | -0,629517634 | 0,150418464 | -4,185108762 | 2,85E-05 | 0,000388691 |
| ENSG00000171885.18 | AQP4 | protein_coding | 1174,264232 | -0,629778622 | 0,157500134 | -3,998591025 | 6,37E-05 | 0,000797242 |
| ENSG00000133119.13 | RFC3 | protein_coding | 5955,381706 | -0,629788253 | 0,140534288 | -4,481385017 | 7,42E-06 | 0,000115861 |
| ENSG00000198729.5 | PPP1R14C | protein_coding | 2242,079761 | -0,629813422 | 0,155577228 | -4,048236551 | 5,16E-05 | 0,000659554 |
| ENSG00000126218.12 | F10 | protein_coding | 9838,712065 | -0,630857534 | 0,139015085 | -4,538050929 | 5,68E-06 | 9,07E-05 |
| ENSG00000198825.15 | INPP5F | protein_coding | 719,2974244 | -0,63158579 | 0,17370929 | -3,635878024 | 0,000277035 | 0,002912976 |
| ENSG00000145642.12 | SHISAL2B | protein_coding | 785,2228149 | -0,631725892 | 0,183778238 | -3,437435789 | 0,00058725 | 0,005534701 |
| ENSG00000176974.20 | SHMT1 | protein_coding | 3155,275249 | -0,632775037 | 0,139338335 | -4,541284605 | 5,59E-06 | 8,94E-05 |
| ENSG00000118271.12 | TTR | protein_coding | 107290,6158 | -0,632931353 | 0,147608946 | -4,287892923 | 1,80E-05 | 0,000257057 |
| ENSG00000169903.7 | TM4SF4 | protein_coding | 3542,432459 | -0,63587004 | 0,150960518 | -4,212161218 | 2,53E-05 | 0,000348144 |
| ENSG00000164663.14 | USP49 | protein_coding | 801,5710767 | -0,635972416 | 0,214915574 | -2,959173246 | 0,003084656 | 0,022248348 |
| ENSG00000116039.13 | ATP6V1B1 | protein_coding | 1635,147519 | -0,636378485 | 0,144261597 | -4,411281306 | 1,03E-05 | 0,000156004 |
| ENSG00000133863.9 | TEX15 | protein_coding | 465,9515555 | -0,637232419 | 0,2162982 | -2,946082854 | 0,003218262 | 0,023057601 |
| ENSG00000186871.7 | ERCC6L | protein_coding | 1349,645391 | -0,640556081 | 0,166008994 | -3,85856251 | 0,000114056 | 0,001342503 |
| ENSG00000166126.11 | AMN | protein_coding | 7350,195431 | -0,640650554 | 0,156226371 | -4,100783684 | 4,12E-05 | 0,00053741 |
| ENSG00000129480.13 | DTD2 | protein_coding | 1313,611571 | -0,640882444 | 0,171005111 | -3,747738535 | 0,000178436 | 0,001999764 |

| | | | | | | | | |
|---|---|---|---|---|---|---|---|---|
| ENSG00000183010.17 | PYCR1 | protein_coding | 5154,417748 | -0,6409065 | 0,157958314 | -4,05744074 | 4,96E-05 | 0,000636822 |
| ENSG00000163006.12 | CCDC138 | protein_coding | 852,7789878 | -0,640937699 | 0,17856885 | -3,589302952 | 0,000331563 | 0,003397591 |
| ENSG00000171320.15 | ESCO2 | protein_coding | 2214,078951 | -0,642151195 | 0,182933278 | -3,510302788 | 0,000447597 | 0,004420017 |
| ENSG00000115419.13 | GLS | protein_coding | 29518,67265 | -0,642304616 | 0,121165913 | -5,301033935 | 1,15E-07 | 2,43E-06 |
| ENSG00000232442.1 | MHENCR | lncRNA | 429,9125755 | -0,645309685 | 0,227909273 | -2,831432331 | 0,004634004 | 0,031063999 |
| ENSG00000143942.5 | CHAC2 | protein_coding | 1061,289025 | -0,645527197 | 0,192215176 | -3,358357078 | 0,000784073 | 0,007054184 |
| ENSG00000090013.11 | BLVRB | protein_coding | 535,6057111 | -0,64575585 | 0,18563081 | -3,478710517 | 0,000503833 | 0,004881742 |
| ENSG00000089094.19 | KDM2B | protein_coding | 1898,238044 | -0,646881922 | 0,154133565 | -4,196891981 | 2,71E-05 | 0,000370751 |
| ENSG00000144749.14 | LRIG1 | protein_coding | 1496,624779 | -0,6478439 | 0,184730869 | -3,50696072 | 0,000453256 | 0,004460994 |
| ENSG00000224078.15 | SNHG14 | lncRNA | 6826,695839 | -0,651616208 | 0,147979976 | -4,40340799 | 1,07E-05 | 0,000161309 |
| ENSG00000006042.12 | TMEM98 | protein_coding | 4249,303979 | -0,652527446 | 0,142136051 | -4,590865165 | 4,41E-06 | 7,23E-05 |
| ENSG00000104369.5 | JPH1 | protein_coding | 519,4930221 | -0,653114738 | 0,189686843 | -3,44312092 | 0,000575042 | 0,005446805 |
| ENSG00000154080.14 | CHST9 | protein_coding | 18337,55206 | -0,653182966 | 0,115817411 | -5,639764893 | 1,70E-08 | 4,03E-07 |
| ENSG00000113649.11 | TCERG1 | protein_coding | 2664,688771 | -0,6539989 | 0,195832493 | -3,339583178 | 0,000839042 | 0,007474616 |
| ENSG00000138835.22 | RGS3 | protein_coding | 1269,676523 | -0,654671832 | 0,190510746 | -3,436403694 | 0,000589492 | 0,005551513 |
| ENSG00000197299.12 | BLM | protein_coding | 928,4008333 | -0,654844175 | 0,230799747 | -2,837282895 | 0,004549928 | 0,0306476 |
| ENSG00000155850.8 | SLC26A2 | protein_coding | 1773,111312 | -0,655535222 | 0,159067914 | -4,121102767 | 3,77E-05 | 0,000496467 |
| ENSG00000205978.6 | NYNRIN | protein_coding | 455,3680731 | -0,656035182 | 0,220672909 | -2,972885002 | 0,00295015 | 0,021452373 |
| ENSG00000147883.12 | CDKN2B | protein_coding | 5191,919339 | -0,656458061 | 0,127109806 | -5,16449583 | 2,41E-07 | 4,88E-06 |
| ENSG00000075240.17 | GRAMD4 | protein_coding | 486,7168334 | -0,656789569 | 0,210308205 | -3,122985954 | 0,001790263 | 0,014207412 |
| ENSG00000103154.9 | NECAB2 | protein_coding | 393,1850163 | -0,657040404 | 0,236755153 | -2,77518945 | 0,005516956 | 0,035680406 |
| ENSG00000063015.20 | SEZ6 | protein_coding | 5185,792364 | -0,657115226 | 0,166243694 | -3,952722721 | 7,73E-05 | 0,000949844 |
| ENSG00000071539.14 | TRIP13 | protein_coding | 1405,119224 | -0,657138945 | 0,159240328 | -4,126711816 | 3,68E-05 | 0,000486966 |
| ENSG00000008256.16 | CYTH3 | protein_coding | 399,4554608 | -0,657275394 | 0,232069666 | -2,832233123 | 0,004622413 | 0,031016098 |
| ENSG00000162949.16 | CAPN13 | protein_coding | 836,2105823 | -0,657768538 | 0,193079532 | -3,406723277 | 0,000657477 | 0,006071589 |
| ENSG00000174371.17 | EXO1 | protein_coding | 2098,665101 | -0,658493628 | 0,158464796 | -4,155456883 | 3,25E-05 | 0,00043594 |
| ENSG00000148950.11 | IMMP1L | protein_coding | 2619,694513 | -0,658745689 | 0,143615349 | -4,586875232 | 4,50E-06 | 7,35E-05 |
| ENSG00000138207.14 | RBP4 | protein_coding | 1187,382542 | -0,65890561 | 0,192781739 | -3,417883938 | 0,0006311 | 0,005882494 |
| ENSG00000162062.15 | TEDC2 | protein_coding | 1322,041162 | -0,66091053 | 0,153291429 | -4,31146435 | 1,62E-05 | 0,000233185 |
| ENSG00000012048.23 | BRCA1 | protein_coding | 2567,543221 | -0,661470203 | 0,178903321 | -3,697361239 | 0,000217852 | 0,002394087 |
| ENSG00000260947.1 | AL356489.2 | lncRNA | 362,543111 | -0,661811342 | 0,223650869 | -2,959127082 | 0,003085118 | 0,022248348 |
| ENSG00000142949.17 | PTPRF | protein_coding | 8672,429159 | -0,662074196 | 0,153032101 | -4,32637461 | 1,52E-05 | 0,000220069 |
| ENSG00000183508.5 | TENT5C | protein_coding | 6510,82763 | -0,662777546 | 0,137633648 | -4,815519711 | 1,47E-06 | 2,62E-05 |
| ENSG00000186188.11 | FFAR4 | protein_coding | 385,3346043 | -0,66397011 | 0,214779857 | -3,091398423 | 0,001992161 | 0,01553871 |
| ENSG00000088992.18 | TESC | protein_coding | 2276,301706 | -0,664092277 | 0,195416412 | -3,398344433 | 0,00067795 | 0,006233151 |
| ENSG00000131944.10 | FAAP24 | protein_coding | 366,9636595 | -0,664535655 | 0,251900343 | -2,638089516 | 0,008337457 | 0,049586552 |

| | | | | | | | | |
|---|---|---|---|---|---|---|---|---|
| ENSG00000196090.12 | PTPRT | protein_coding | 11768,04578 | -0,664621933 | 0,180127498 | -3,689730547 | 0,000224492 | 0,002445231 |
| ENSG00000007402.12 | CACNA2D2 | protein_coding | 4440,593113 | -0,665849031 | 0,144946946 | -4,593743098 | 4,35E-06 | 7,14E-05 |
| ENSG00000124406.16 | ATP8A1 | protein_coding | 19357,70399 | -0,666848214 | 0,141490929 | -4,713010354 | 2,44E-06 | 4,16E-05 |
| ENSG00000167771.6 | RCOR2 | protein_coding | 2434,50377 | -0,66700238 | 0,152462452 | -4,374863252 | 1,22E-05 | 0,000181569 |
| ENSG00000100814.18 | CCNB1IP1 | protein_coding | 2962,501425 | -0,66794648 | 0,138095643 | -4,836839625 | 1,32E-06 | 2,38E-05 |
| ENSG00000118655.7 | DCLRE1B | protein_coding | 1568,78034 | -0,668641897 | 0,151746766 | -4,406300802 | 1,05E-05 | 0,000159517 |
| ENSG00000134871.19 | COL4A2 | protein_coding | 2696,835413 | -0,669294572 | 0,216912245 | -3,085554586 | 0,002031729 | 0,015770738 |
| ENSG00000139044.12 | B4GALNT3 | protein_coding | 1231,261409 | -0,670605883 | 0,170745594 | -3,927515003 | 8,58E-05 | 0,001045265 |
| ENSG00000109576.14 | AADAT | protein_coding | 396,0958505 | -0,671363759 | 0,202133589 | -3,321386435 | 0,000895714 | 0,00790214 |
| ENSG00000216775.3 | AL109918.1 | transcribed_unproc | 1631,584322 | -0,671700018 | 0,139781716 | -4,805349619 | 1,54E-06 | 2,75E-05 |
| ENSG00000132122.12 | SPATA6 | protein_coding | 500,4240593 | -0,672026159 | 0,227735227 | -2,950910009 | 0,003168392 | 0,022778259 |
| ENSG00000110080.20 | ST3GAL4 | protein_coding | 1679,587788 | -0,672883647 | 0,147754044 | -4,554079395 | 5,26E-06 | 8,45E-05 |
| ENSG00000142733.17 | MAP3K6 | protein_coding | 310,7110404 | -0,672916936 | 0,226519792 | -2,970676119 | 0,00297145 | 0,021577281 |
| ENSG00000172554.12 | SNTG2 | protein_coding | 570,5344155 | -0,673359914 | 0,195525427 | -3,443848316 | 0,000573497 | 0,005440039 |
| ENSG00000160072.20 | ATAD3B | protein_coding | 1011,123058 | -0,67569755 | 0,170039354 | -3,97377157 | 7,07E-05 | 0,000876339 |
| ENSG00000100033.16 | PRODH | protein_coding | 4194,714915 | -0,675880263 | 0,154738015 | -4,367900574 | 1,25E-05 | 0,000186257 |
| ENSG00000151338.18 | MIPOL1 | protein_coding | 242,4480412 | -0,676950892 | 0,245754714 | -2,754579477 | 0,005876761 | 0,037497715 |
| ENSG00000174482.10 | LINGO2 | protein_coding | 719,660317 | -0,677554939 | 0,167215348 | -4,051990116 | 5,08E-05 | 0,00065025 |
| ENSG00000163697.17 | APBB2 | protein_coding | 2008,591101 | -0,677603079 | 0,141042849 | -4,804235604 | 1,55E-06 | 2,76E-05 |
| ENSG00000155542.12 | SETD9 | protein_coding | 526,3671263 | -0,67810552 | 0,221850406 | -3,056589048 | 0,00223871 | 0,017086044 |
| ENSG00000165490.13 | DDIAS | protein_coding | 581,9111938 | -0,678364197 | 0,186268192 | -3,641868155 | 0,000270667 | 0,002863268 |
| ENSG00000268516.2 | AC020915.2 | lncRNA | 265,9733122 | -0,678446409 | 0,235643821 | -2,879118182 | 0,003987889 | 0,027607952 |
| ENSG00000164199.18 | ADGRV1 | protein_coding | 1528,172108 | -0,67849452 | 0,168314407 | -4,031113746 | 5,55E-05 | 0,000705627 |
| ENSG00000197016.12 | ZNF470 | protein_coding | 469,3606402 | -0,678519089 | 0,210050357 | -3,230268679 | 0,001236739 | 0,010469526 |
| ENSG00000166402.9 | TUB | protein_coding | 632,5464197 | -0,679131741 | 0,188865648 | -3,595845773 | 0,000323339 | 0,003319818 |
| ENSG00000104774.13 | MAN2B1 | protein_coding | 7886,859751 | -0,680074131 | 0,155869833 | -4,363090148 | 1,28E-05 | 0,000189863 |
| ENSG00000145604.17 | SKP2 | protein_coding | 3536,220127 | -0,681108998 | 0,13772893 | -4,945286338 | 7,60E-07 | 1,42E-05 |
| ENSG00000213853.10 | EMP2 | protein_coding | 8463,184633 | -0,681172747 | 0,165672528 | -4,111561262 | 3,93E-05 | 0,000516466 |
| ENSG00000034533.12 | ASTE1 | protein_coding | 512,1825816 | -0,682093179 | 0,203594252 | -3,350257551 | 0,000807365 | 0,007241721 |
| ENSG00000075043.19 | KCNQ2 | protein_coding | 4828,388104 | -0,682784157 | 0,13209074 | -5,169053897 | 2,35E-07 | 4,77E-06 |
| ENSG00000115616.3 | SLC9A2 | protein_coding | 2525,802066 | -0,682799658 | 0,13415081 | -5,089791542 | 3,58E-07 | 7,09E-06 |
| ENSG00000204347.4 | BTBD17 | protein_coding | 8264,221375 | -0,683984515 | 0,128763872 | -5,31192874 | 1,08E-07 | 2,30E-06 |
| ENSG00000170439.7 | METTL7B | protein_coding | 297,8342138 | -0,684191361 | 0,233481547 | -2,930387311 | 0,003385398 | 0,024032994 |
| ENSG00000163808.17 | KIF15 | protein_coding | 1189,223477 | -0,6842275 | 0,164353795 | -4,163137817 | 3,14E-05 | 0,000423154 |
| ENSG00000011426.11 | ANLN | protein_coding | 6415,984804 | -0,684835172 | 0,141437144 | -4,841975397 | 1,29E-06 | 2,32E-05 |
| ENSG00000144283.22 | PKP4 | protein_coding | 1894,244831 | -0,686033859 | 0,149579344 | -4,586421083 | 4,51E-06 | 7,35E-05 |

| | | | | | | | | |
|---|---|---|---|---|---|---|---|---|
| ENSG00000113205.5 | PCDHB3 | protein_coding | 381,808829 | -0,686621151 | 0,207108578 | -3,315271418 | 0,000915542 | 0,008053306 |
| ENSG00000131791.8 | PRKAB2 | protein_coding | 2051,129776 | -0,686787212 | 0,148756351 | -4,616859772 | 3,90E-06 | 6,44E-05 |
| ENSG00000147003.7 | CLTRN | protein_coding | 423,9137243 | -0,686952742 | 0,226334584 | -3,035120526 | 0,002404396 | 0,018132277 |
| ENSG00000083457.12 | ITGAE | protein_coding | 1530,802238 | -0,687767996 | 0,186856044 | -3,680737214 | 0,000232561 | 0,002521313 |
| ENSG00000111670.16 | GNPTAB | protein_coding | 1664,911689 | -0,687993627 | 0,147456353 | -4,665744229 | 3,08E-06 | 5,17E-05 |
| ENSG00000084710.14 | EFR3B | protein_coding | 718,6244293 | -0,689884782 | 0,183827043 | -3,752901474 | 0,0001748 | 0,001966378 |
| ENSG00000183098.11 | GPC6 | protein_coding | 4547,420901 | -0,69057787 | 0,123446075 | -5,594166265 | 2,22E-08 | 5,17E-07 |
| ENSG00000187266.14 | EPOR | protein_coding | 2139,800696 | -0,691114087 | 0,151060223 | -4,575089819 | 4,76E-06 | 7,72E-05 |
| ENSG00000091704.10 | CPA1 | protein_coding | 1136,630947 | -0,691598257 | 0,162895562 | -4,245654374 | 2,18E-05 | 0,000303992 |
| ENSG00000099953.10 | MMP11 | protein_coding | 391,10446 | -0,691766609 | 0,217905842 | -3,17461249 | 0,001500367 | 0,012303246 |
| ENSG00000155111.15 | CDK19 | protein_coding | 2681,543514 | -0,692209981 | 0,137278063 | -5,042393291 | 4,60E-07 | 8,92E-06 |
| ENSG00000156510.13 | HKDC1 | protein_coding | 753,4970007 | -0,693249135 | 0,205528217 | -3,373011956 | 0,000743507 | 0,006744075 |
| ENSG00000198885.10 | ITPRIPL1 | protein_coding | 896,5428809 | -0,694000561 | 0,158469072 | -4,379406977 | 1,19E-05 | 0,000178334 |
| ENSG00000157851.17 | DPYSL5 | protein_coding | 5616,60118 | -0,695450153 | 0,143066183 | -4,861037997 | 1,17E-06 | 2,12E-05 |
| ENSG00000138193.16 | PLCE1 | protein_coding | 497,3571725 | -0,6972929 | 0,228909035 | -3,046157179 | 0,002317866 | 0,017581352 |
| ENSG00000137868.19 | STRA6 | protein_coding | 10560,05643 | -0,697881997 | 0,128085608 | -5,448559039 | 5,08E-08 | 1,13E-06 |
| ENSG00000197256.11 | KANK2 | protein_coding | 1839,909825 | -0,699376839 | 0,163995793 | -4,264602308 | 2,00E-05 | 0,000282546 |
| ENSG00000215883.10 | CYB5RL | protein_coding | 2130,875423 | -0,700045182 | 0,15654542 | -4,471834318 | 7,76E-06 | 0,00012053 |
| ENSG00000123612.16 | ACVR1C | protein_coding | 67599,54309 | -0,70074055 | 0,121102792 | -5,786328609 | 7,19E-09 | 1,78E-07 |
| ENSG00000188211.9 | NCR3LG1 | protein_coding | 1172,354705 | -0,701100791 | 0,19162891 | -3,658637894 | 0,000253559 | 0,002708298 |
| ENSG00000158555.15 | GDPD5 | protein_coding | 255,4137948 | -0,704992019 | 0,249385758 | -2,826913712 | 0,004699899 | 0,031355126 |
| ENSG00000178761.15 | FAM219B | protein_coding | 3358,986189 | -0,705662909 | 0,138433692 | -5,097479511 | 3,44E-07 | 6,82E-06 |
| ENSG00000214706.10 | IFRD2 | protein_coding | 2935,261361 | -0,705767288 | 0,125209593 | -5,636686999 | 1,73E-08 | 4,10E-07 |
| ENSG00000132561.14 | MATN2 | protein_coding | 8904,151042 | -0,707229585 | 0,17146795 | -4,124558473 | 3,71E-05 | 0,000490305 |
| ENSG00000126953.8 | TIMM8A | protein_coding | 301,2220119 | -0,707313639 | 0,235174341 | -3,007613992 | 0,002633074 | 0,019519742 |
| ENSG00000247828.8 | TMEM161B-AS1 | lncRNA | 372,3323493 | -0,707956639 | 0,242048794 | -2,924850929 | 0,003446213 | 0,024415046 |
| ENSG00000121905.10 | HPCA | protein_coding | 8177,940756 | -0,710023628 | 0,155553112 | -4,564509301 | 5,01E-06 | 8,08E-05 |
| ENSG00000049167.15 | ERCC8 | protein_coding | 1080,698282 | -0,710937658 | 0,157666191 | -4,509131953 | 6,51E-06 | 0,000102693 |
| ENSG00000018236.15 | CNTN1 | protein_coding | 40719,98735 | -0,711756632 | 0,111644077 | -6,37522968 | 1,83E-10 | 5,39E-09 |
| ENSG00000234420.7 | ZNF37BP | transcribed_process | 472,9001419 | -0,711986064 | 0,249227772 | -2,856768562 | 0,004279778 | 0,02921228 |
| ENSG00000171757.17 | LRRC34 | protein_coding | 285,1845905 | -0,713417898 | 0,242048733 | -2,947414307 | 0,003204436 | 0,023005781 |
| ENSG00000121207.12 | LRAT | protein_coding | 408,9828666 | -0,715099331 | 0,202465572 | -3,531955195 | 0,000412499 | 0,004116145 |
| ENSG00000164076.17 | CAMKV | protein_coding | 2162,033496 | -0,715399574 | 0,159729672 | -4,478814525 | 7,51E-06 | 0,00011714 |
| ENSG00000230006.7 | ANKRD36BP2 | transcribed_unproc | 259,3577288 | -0,715802322 | 0,261477675 | -2,737527481 | 0,006190295 | 0,039161954 |
| ENSG00000088386.17 | SLC15A1 | protein_coding | 514,2495219 | -0,715824289 | 0,19169792 | -3,734126529 | 0,000188368 | 0,002094254 |
| ENSG00000203485.14 | INF2 | protein_coding | 557,0138898 | -0,716057008 | 0,214452941 | -3,338993648 | 0,000840825 | 0,007484128 |

| | | | | | | | | |
|---|---|---|---|---|---|---|---|---|
| ENSG00000138346.15 | DNA2 | protein_coding | 1682,155929 | -0,716220664 | 0,160325529 | -4,467290194 | 7,92E-06 | 0,000122572 |
| ENSG00000204860.5 | FAM201A | lncRNA | 302,5812098 | -0,717314761 | 0,230598677 | -3,110662953 | 0,001866679 | 0,014713448 |
| ENSG00000266916.7 | ZNF793-AS1 | lncRNA | 337,4198207 | -0,717402226 | 0,219242275 | -3,272189302 | 0,001067181 | 0,009167597 |
| ENSG00000162512.16 | SDC3 | protein_coding | 1293,521429 | -0,718759516 | 0,189714147 | -3,788644788 | 0,000151471 | 0,001724335 |
| ENSG00000148219.17 | ASTN2 | protein_coding | 313,695571 | -0,71900078 | 0,227462682 | -3,160961495 | 0,001572493 | 0,012814378 |
| ENSG00000248643.5 | RBM14-RBM4 | protein_coding | 445,544997 | -0,719092764 | 0,259320292 | -2,772990726 | 0,00555437 | 0,035866945 |
| ENSG00000144741.18 | SLC25A26 | protein_coding | 801,8730902 | -0,719563081 | 0,244349652 | -2,944809104 | 0,00323154 | 0,023144812 |
| ENSG00000133627.18 | ACTR3B | protein_coding | 2152,400825 | -0,72029241 | 0,132086959 | -5,45316823 | 4,95E-08 | 1,10E-06 |
| ENSG00000287915.1 | AC037433.1 | lncRNA | 489,4153646 | -0,72090671 | 0,20921182 | -3,445822091 | 0,000569325 | 0,005403461 |
| ENSG00000285967.1 | NIPBL-DT | lncRNA | 2698,163972 | -0,721439528 | 0,136887006 | -5,270328769 | 1,36E-07 | 2,84E-06 |
| ENSG00000169962.5 | TAS1R3 | protein_coding | 1158,991348 | -0,722848151 | 0,183315184 | -3,943198468 | 8,04E-05 | 0,000983762 |
| ENSG00000175040.6 | CHST2 | protein_coding | 4586,822313 | -0,72303212 | 0,142462067 | -5,075260633 | 3,87E-07 | 7,57E-06 |
| ENSG00000004399.13 | PLXND1 | protein_coding | 2036,559033 | -0,723044752 | 0,164469601 | -4,396221241 | 1,10E-05 | 0,000166021 |
| ENSG00000111845.5 | PAK1IP1 | protein_coding | 912,5222842 | -0,723328897 | 0,175306591 | -4,126079293 | 3,69E-05 | 0,000487907 |
| ENSG00000257167.2 | TMPO-AS1 | lncRNA | 1023,570054 | -0,723809012 | 0,194918137 | -3,713400009 | 0,000204493 | 0,002253193 |
| ENSG00000161270.20 | NPHS1 | protein_coding | 1878,72177 | -0,724069088 | 0,147738805 | -4,901008159 | 9,53E-07 | 1,75E-05 |
| ENSG00000174498.14 | IGDCC3 | protein_coding | 1122,375803 | -0,724753555 | 0,179853213 | -4,029694794 | 5,58E-05 | 0,0007089 |
| ENSG00000228782.7 | MRPL45P2 | transcribed_unproc | 242,4460739 | -0,724825434 | 0,24987932 | -2,900701967 | 0,003723278 | 0,026104095 |
| ENSG00000150627.16 | WDR17 | protein_coding | 1313,16279 | -0,725433354 | 0,167237321 | -4,337748008 | 1,44E-05 | 0,0002103 |
| ENSG00000260920.2 | AL031985.3 | lncRNA | 323,8196032 | -0,726400359 | 0,226350297 | -3,20918668 | 0,00133111 | 0,011128912 |
| ENSG00000198088.10 | NUP62CL | protein_coding | 310,226845 | -0,726458148 | 0,254356525 | -2,856062565 | 0,004289306 | 0,029240189 |
| ENSG00000156564.9 | LRFN2 | protein_coding | 1174,726912 | -0,727804123 | 0,157977667 | -4,607006384 | 4,09E-06 | 6,72E-05 |
| ENSG00000196169.15 | KIF19 | protein_coding | 562,615898 | -0,728896583 | 0,211062126 | -3,453469348 | 0,000553425 | 0,005271136 |
| ENSG00000229891.2 | LINC01315 | lncRNA | 526,7954565 | -0,729030554 | 0,194363324 | -3,750864823 | 0,000176226 | 0,00197923 |
| ENSG00000256196.2 | AP003721.1 | lncRNA | 1265,402086 | -0,72937063 | 0,166924994 | -4,36945128 | 1,25E-05 | 0,000185466 |
| ENSG00000137843.12 | PAK6 | protein_coding | 907,85933 | -0,729946852 | 0,165158874 | -4,419664751 | 9,89E-06 | 0,00015051 |
| ENSG00000189212.12 | DPY19L2P1 | transcribed_unproc | 241,2795121 | -0,730137036 | 0,258646191 | -2,822918184 | 0,004758871 | 0,031679042 |
| ENSG00000076864.19 | RAP1GAP | protein_coding | 8701,627371 | -0,732538837 | 0,135227163 | -5,41709833 | 6,06E-08 | 1,34E-06 |
| ENSG00000145526.12 | CDH18 | protein_coding | 321,1784561 | -0,732539968 | 0,230172068 | -3,182575432 | 0,001459715 | 0,012007517 |
| ENSG00000162390.18 | ACOT11 | protein_coding | 578,9909953 | -0,733065384 | 0,195920662 | -3,74164407 | 0,00018282 | 0,002042339 |
| ENSG00000109738.11 | GLRB | protein_coding | 1367,519617 | -0,73376423 | 0,14235922 | -5,154314773 | 2,55E-07 | 5,14E-06 |
| ENSG00000250056.8 | LINC01018 | lncRNA | 904,6017012 | -0,733816058 | 0,186117066 | -3,942766089 | 8,05E-05 | 0,000984962 |
| ENSG00000101220.18 | C20orf27 | protein_coding | 1151,312358 | -0,734762634 | 0,166468276 | -4,413829793 | 1,02E-05 | 0,000154402 |
| ENSG00000109674.4 | NEIL3 | protein_coding | 1146,220495 | -0,735201736 | 0,17622442 | -4,171962858 | 3,02E-05 | 0,000408668 |
| ENSG00000129173.13 | E2F8 | protein_coding | 668,3874494 | -0,735644533 | 0,187922942 | -3,914607374 | 9,06E-05 | 0,001096411 |
| ENSG00000165810.17 | BTNL9 | protein_coding | 222,1686932 | -0,735986707 | 0,267842803 | -2,747830811 | 0,005999096 | 0,038185183 |

| | | | | | | | | |
|---|---|---|---|---|---|---|---|---|
| ENSG00000147262.4 | GPR119 | protein_coding | 1969,092846 | -0,738334335 | 0,148750231 | -4,963584445 | 6,92E-07 | 1,30E-05 |
| ENSG00000267270.6 | PARD6G-AS1 | lncRNA | 176,6743871 | -0,738811915 | 0,279963938 | -2,63895386 | 0,00831623 | 0,04952144 |
| ENSG00000088035.18 | ALG6 | protein_coding | 1947,810811 | -0,739470179 | 0,160877118 | -4,596490713 | 4,30E-06 | 7,05E-05 |
| ENSG00000273032.3 | DGCR5 | lncRNA | 687,4983854 | -0,73972778 | 0,168876655 | -4,380284414 | 1,19E-05 | 0,000177744 |
| ENSG00000062282.15 | DGAT2 | protein_coding | 1907,106937 | -0,74029797 | 0,148129389 | -4,997644106 | 5,80E-07 | 1,11E-05 |
| ENSG00000196584.3 | XRCC2 | protein_coding | 337,4037336 | -0,743799412 | 0,262717488 | -2,831175863 | 0,004637721 | 0,031069023 |
| ENSG00000036530.9 | CYP46A1 | protein_coding | 467,4071095 | -0,744142754 | 0,203430514 | -3,657970175 | 0,000254221 | 0,002713977 |
| ENSG00000188338.15 | SLC38A3 | protein_coding | 4329,859198 | -0,745356093 | 0,121949746 | -6,111993817 | 9,84E-10 | 2,71E-08 |
| ENSG00000134072.11 | CAMK1 | protein_coding | 689,1904125 | -0,74841185 | 0,174298026 | -4,293863024 | 1,76E-05 | 0,00025107 |
| ENSG00000117525.14 | F3 | protein_coding | 883,7895867 | -0,751765095 | 0,22478364 | -3,344394173 | 0,000824625 | 0,007366788 |
| ENSG00000130283.9 | GDF1 | protein_coding | 730,0064957 | -0,753560554 | 0,282130519 | -2,670964339 | 0,007563368 | 0,045868804 |
| ENSG00000144354.14 | CDCA7 | protein_coding | 1268,959551 | -0,754700355 | 0,189519346 | -3,982181094 | 6,83E-05 | 0,000847901 |
| ENSG00000143171.13 | RXRG | protein_coding | 708,8467804 | -0,75650988 | 0,182441099 | -4,1465979 | 3,37E-05 | 0,000449972 |
| ENSG00000213366.13 | GSTM2 | protein_coding | 2116,425101 | -0,75658526 | 0,157623336 | -4,799957167 | 1,59E-06 | 2,82E-05 |
| ENSG00000172367.16 | PDZD3 | protein_coding | 619,9533848 | -0,758715027 | 0,274585612 | -2,763127398 | 0,005725042 | 0,036731151 |
| ENSG00000196950.14 | SLC39A10 | protein_coding | 2653,492497 | -0,759403934 | 0,131807615 | -5,761457215 | 8,34E-09 | 2,05E-07 |
| ENSG00000243335.10 | KCTD7 | protein_coding | 1061,866101 | -0,759913196 | 0,15527946 | -4,893842339 | 9,89E-07 | 1,81E-05 |
| ENSG00000198453.13 | ZNF568 | protein_coding | 865,1290518 | -0,7599578 | 0,167890651 | -4,526504583 | 6,00E-06 | 9,50E-05 |
| ENSG00000204175.5 | GPRIN2 | protein_coding | 336,2582534 | -0,76000913 | 0,249276377 | -3,048861422 | 0,002297104 | 0,017474519 |
| ENSG00000112294.14 | ALDH5A1 | protein_coding | 4371,493041 | -0,76048073 | 0,1359352 | -5,594435675 | 2,21E-08 | 5,17E-07 |
| ENSG00000178234.13 | GALNT11 | protein_coding | 4380,186479 | -0,762074078 | 0,118509353 | -6,430497336 | 1,27E-10 | 3,79E-09 |
| ENSG00000118513.19 | MYB | protein_coding | 371,309616 | -0,762590316 | 0,218538358 | -3,489503269 | 0,000483919 | 0,004714215 |
| ENSG00000140848.17 | CPNE2 | protein_coding | 564,7697038 | -0,762638879 | 0,194058395 | -3,929945302 | 8,50E-05 | 0,001035962 |
| ENSG00000157514.16 | TSC22D3 | protein_coding | 5434,130443 | -0,762837543 | 0,12019321 | -6,346760685 | 2,20E-10 | 6,45E-09 |
| ENSG00000242125.3 | SNHG3 | lncRNA | 1154,365119 | -0,76291819 | 0,161275478 | -4,730528164 | 2,24E-06 | 3,86E-05 |
| ENSG00000157227.13 | MMP14 | protein_coding | 863,7685129 | -0,763428523 | 0,165137391 | -4,622990105 | 3,78E-06 | 6,26E-05 |
| ENSG00000083290.20 | ULK2 | protein_coding | 1660,752722 | -0,763822316 | 0,147509784 | -5,178112909 | 2,24E-07 | 4,56E-06 |
| ENSG00000277161.2 | PIGW | protein_coding | 1922,268929 | -0,764225359 | 0,143965165 | -5,308404719 | 1,11E-07 | 2,34E-06 |
| ENSG00000167106.12 | FAM102A | protein_coding | 6511,390312 | -0,76715966 | 0,122209655 | -6,277406292 | 3,44E-10 | 9,91E-09 |
| ENSG00000076641.4 | PAG1 | protein_coding | 1036,199822 | -0,767638295 | 0,1970651 | -3,895353844 | 9,81E-05 | 0,001173024 |
| ENSG00000165887.12 | ANKRD2 | protein_coding | 190,0131328 | -0,767708247 | 0,281444715 | -2,727740848 | 0,006376969 | 0,040102684 |
| ENSG00000254647.7 | INS | protein_coding | 333161,6638 | -0,768639269 | 0,173432249 | -4,431928167 | 9,34E-06 | 0,000142716 |
| ENSG00000167964.13 | RAB26 | protein_coding | 2868,820411 | -0,76977045 | 0,171872693 | -4,47872456 | 7,51E-06 | 0,00011714 |
| ENSG00000137261.14 | KIAA0319 | protein_coding | 858,1732634 | -0,770071313 | 0,16283898 | -4,729035485 | 2,26E-06 | 3,88E-05 |
| ENSG00000127252.7 | PLAAT1 | protein_coding | 294,6584584 | -0,771152353 | 0,257429373 | -2,995588042 | 0,002739162 | 0,020170369 |
| ENSG00000151632.17 | AKR1C2 | protein_coding | 760,2557318 | -0,771819417 | 0,170199072 | -4,534803907 | 5,77E-06 | 9,19E-05 |

| | | | | | | | | |
|---|---|---|---|---|---|---|---|---|
| ENSG00000240204.3 | SMKR1 | protein_coding | 229,5433927 | -0,772132439 | 0,267777925 | -2,883480555 | 0,003933069 | 0,027331415 |
| ENSG00000141753.7 | IGFBP4 | protein_coding | 1561,450021 | -0,772846562 | 0,173045408 | -4,466148919 | 7,96E-06 | 0,000123136 |
| ENSG00000173588.15 | CEP83 | protein_coding | 597,5813393 | -0,77409493 | 0,179611621 | -4,309826527 | 1,63E-05 | 0,000234436 |
| ENSG00000256151.1 | ADGRD1-AS1 | lncRNA | 506,2207424 | -0,774409251 | 0,200318282 | -3,865894021 | 0,000110683 | 0,001309123 |
| ENSG00000157978.12 | LDLRAP1 | protein_coding | 2601,907167 | -0,774724135 | 0,161512464 | -4,796683274 | 1,61E-06 | 2,86E-05 |
| ENSG00000185046.18 | ANKS1B | protein_coding | 544,5198587 | -0,774831194 | 0,202683993 | -3,822853407 | 0,000131916 | 0,001524098 |
| ENSG00000237767.2 | LINC01370 | lncRNA | 324,5487495 | -0,774998167 | 0,265601943 | -2,917893442 | 0,003524048 | 0,024857121 |
| ENSG00000053438.11 | NNAT | protein_coding | 3512,672059 | -0,777504077 | 0,126300085 | -6,156005958 | 7,46E-10 | 2,08E-08 |
| ENSG00000279069.1 | AC015813.5 | TEC | 176,9971018 | -0,777507926 | 0,29150178 | -2,667249323 | 0,007647492 | 0,046245013 |
| ENSG00000065320.9 | NTN1 | protein_coding | 5117,084485 | -0,778442125 | 0,162932179 | -4,777706457 | 1,77E-06 | 3,12E-05 |
| ENSG00000106266.11 | SNX8 | protein_coding | 859,9040187 | -0,779053104 | 0,185260964 | -4,205165989 | 2,61E-05 | 0,000358382 |
| ENSG00000276043.5 | UHRF1 | protein_coding | 2010,674497 | -0,779206908 | 0,180613407 | -4,314225188 | 1,60E-05 | 0,000230607 |
| ENSG00000260804.3 | LINC01963 | lncRNA | 1683,036114 | -0,779603173 | 0,152396457 | -5,115625307 | 3,13E-07 | 6,25E-06 |
| ENSG00000285693.1 | AP002381.2 | lncRNA | 196,5921123 | -0,779644645 | 0,277664037 | -2,807870456 | 0,004987028 | 0,032872619 |
| ENSG00000183077.15 | AFMID | protein_coding | 250,3688027 | -0,781246635 | 0,274574229 | -2,845302121 | 0,004436932 | 0,029992627 |
| ENSG00000173145.11 | NOC3L | protein_coding | 834,8174418 | -0,781751294 | 0,169183623 | -4,620726765 | 3,82E-06 | 6,33E-05 |
| ENSG00000106348.18 | IMPDH1 | protein_coding | 2877,046068 | -0,782086236 | 0,127550655 | -6,131573666 | 8,70E-10 | 2,42E-08 |
| ENSG00000247092.7 | SNHG10 | lncRNA | 513,557194 | -0,78249044 | 0,275111027 | -2,844271453 | 0,004451311 | 0,030056632 |
| ENSG00000185483.12 | ROR1 | protein_coding | 1702,871774 | -0,783628592 | 0,142600294 | -5,495280359 | 3,90E-08 | 8,83E-07 |
| ENSG00000172817.4 | CYP7B1 | protein_coding | 2139,167942 | -0,783749755 | 0,141185553 | -5,551203621 | 2,84E-08 | 6,53E-07 |
| ENSG00000146263.11 | MMS22L | protein_coding | 2106,775785 | -0,784963783 | 0,147403718 | -5,325264478 | 1,01E-07 | 2,15E-06 |
| ENSG00000196511.14 | TPK1 | protein_coding | 856,9635245 | -0,785386491 | 0,177043296 | -4,436126679 | 9,16E-06 | 0,000140372 |
| ENSG00000090447.12 | TFAP4 | protein_coding | 368,7078908 | -0,787595598 | 0,210834973 | -3,735602238 | 0,000187266 | 0,002085332 |
| ENSG00000223802.8 | CERS1 | protein_coding | 792,2566414 | -0,788169704 | 0,214859974 | -3,668294706 | 0,000244174 | 0,002626811 |
| ENSG00000105088.8 | OLFM2 | protein_coding | 3617,793074 | -0,788252184 | 0,149070834 | -5,287769311 | 1,24E-07 | 2,60E-06 |
| ENSG00000163293.12 | NIPAL1 | protein_coding | 924,2034811 | -0,788942798 | 0,165205255 | -4,775530884 | 1,79E-06 | 3,15E-05 |
| ENSG00000086300.16 | SNX10 | protein_coding | 1169,700742 | -0,789143412 | 0,152733364 | -5,166804372 | 2,38E-07 | 4,82E-06 |
| ENSG00000182218.9 | HHIPL1 | protein_coding | 178,3943626 | -0,78927086 | 0,287821778 | -2,74222078 | 0,006102531 | 0,038725762 |
| ENSG00000111186.13 | WNT5B | protein_coding | 678,8247409 | -0,790060353 | 0,188939772 | -4,181546025 | 2,90E-05 | 0,000393852 |
| ENSG00000170100.14 | ZNF778 | protein_coding | 684,6871349 | -0,790425093 | 0,19488356 | -4,0558839 | 4,99E-05 | 0,000639902 |
| ENSG00000196154.12 | S100A4 | protein_coding | 358,6549613 | -0,790770559 | 0,214080615 | -3,69379805 | 0,000220929 | 0,002420697 |
| ENSG00000198300.14 | PEG3 | protein_coding | 1595,402665 | -0,791024417 | 0,165697625 | -4,77390316 | 1,81E-06 | 3,17E-05 |
| ENSG00000162341.18 | TPCN2 | protein_coding | 399,0030284 | -0,791609572 | 0,207047244 | -3,823328216 | 0,000131662 | 0,00152314 |
| ENSG00000261659.2 | Z92544.1 | lncRNA | 332,5521379 | -0,792431074 | 0,243542072 | -3,253774874 | 0,001138825 | 0,009709488 |
| ENSG00000148541.13 | FAM13C | protein_coding | 680,9574221 | -0,792965184 | 0,186782185 | -4,245400505 | 2,18E-05 | 0,000304134 |
| ENSG00000260708.1 | AL118516.1 | lncRNA | 429,6971739 | -0,793556214 | 0,217832783 | -3,64296045 | 0,00026952 | 0,002854025 |

| | | | | | | | | |
|---|---|---|---|---|---|---|---|---|
| ENSG00000197296.6 | FITM2 | protein_coding | 3080,098705 | -0,793736199 | 0,168490199 | -4,710874593 | 2,47E-06 | 4,20E-05 |
| ENSG00000182670.13 | TTC3 | protein_coding | 23016,56326 | -0,794364538 | 0,119117519 | -6,668746502 | 2,58E-11 | 8,32E-10 |
| ENSG00000205002.4 | AARD | protein_coding | 888,5883238 | -0,794831438 | 0,166731926 | -4,767122033 | 1,87E-06 | 3,28E-05 |
| ENSG00000178828.7 | RNF186 | protein_coding | 173,7040082 | -0,79515038 | 0,293485455 | -2,709334879 | 0,006741825 | 0,041893771 |
| ENSG00000107736.21 | CDH23 | protein_coding | 691,845914 | -0,796058223 | 0,173557101 | -4,586722274 | 4,50E-06 | 7,35E-05 |
| ENSG00000136982.6 | DSCC1 | protein_coding | 2382,341077 | -0,796191856 | 0,171325927 | -4,647235082 | 3,36E-06 | 5,61E-05 |
| ENSG00000155465.20 | SLC7A7 | protein_coding | 1950,771756 | -0,797140849 | 0,13227658 | -6,026318865 | 1,68E-09 | 4,51E-08 |
| ENSG00000249042.6 | AC008771.1 | lncRNA | 359,4665797 | -0,797642963 | 0,231875443 | -3,439963086 | 0,000581794 | 0,005499217 |
| ENSG00000164099.3 | PRSS12 | protein_coding | 2187,854137 | -0,798519765 | 0,138077815 | -5,783114125 | 7,33E-09 | 1,82E-07 |
| ENSG00000188677.15 | PARVB | protein_coding | 4394,680472 | -0,799999678 | 0,135277218 | -5,9137798 | 3,34E-09 | 8,71E-08 |
| ENSG00000134323.12 | MYCN | protein_coding | 2218,832448 | -0,800257281 | 0,132034464 | -6,06097271 | 1,35E-09 | 3,68E-08 |
| ENSG00000166920.13 | C15orf48 | protein_coding | 555,0273898 | -0,800327656 | 0,189261312 | -4,228691258 | 2,35E-05 | 0,00032567 |
| ENSG00000064225.12 | ST3GAL6 | protein_coding | 1212,570863 | -0,802373648 | 0,174944128 | -4,5864566 | 4,51E-06 | 7,35E-05 |
| ENSG00000170854.18 | RIOX2 | protein_coding | 519,7422168 | -0,808757034 | 0,251004863 | -3,222077148 | 0,001272649 | 0,010722219 |
| ENSG00000054967.13 | RELT | protein_coding | 2326,968532 | -0,809088927 | 0,136978338 | -5,906692519 | 3,49E-09 | 9,03E-08 |
| ENSG00000115252.18 | PDE1A | protein_coding | 968,4382951 | -0,810401561 | 0,224543947 | -3,609100004 | 0,000307261 | 0,003182836 |
| ENSG00000129965.15 | INS-IGF2 | protein_coding | 285,1720978 | -0,811010853 | 0,235940928 | -3,437347045 | 0,000587442 | 0,005534701 |
| ENSG00000100027.17 | YPEL1 | protein_coding | 414,2029836 | -0,811489802 | 0,286446938 | -2,832949821 | 0,004612063 | 0,030966494 |
| ENSG00000170959.14 | DCDC1 | protein_coding | 1435,981454 | -0,812037487 | 0,15109051 | -5,374510183 | 7,68E-08 | 1,67E-06 |
| ENSG00000253982.2 | AC100810.1 | lncRNA | 417,4189687 | -0,81716286 | 0,2247968 | -3,635117855 | 0,000277854 | 0,002917184 |
| ENSG00000185008.17 | ROBO2 | protein_coding | 6823,985615 | -0,818947533 | 0,122948843 | -6,660880369 | 2,72E-11 | 8,71E-10 |
| ENSG00000256162.2 | SMLR1 | protein_coding | 189,8119613 | -0,819568562 | 0,273405801 | -2,997626825 | 0,002720906 | 0,020057103 |
| ENSG00000174469.23 | CNTNAP2 | protein_coding | 1527,6982 | -0,820525864 | 0,174179577 | -4,710804091 | 2,47E-06 | 4,20E-05 |
| ENSG00000228716.7 | DHFR | protein_coding | 9746,727077 | -0,821646211 | 0,138626549 | -5,927047992 | 3,08E-09 | 8,08E-08 |
| ENSG00000121211.8 | MND1 | protein_coding | 357,8555789 | -0,823989849 | 0,261965514 | -3,145413446 | 0,001658523 | 0,013411036 |
| ENSG00000227627.3 | AL080276.2 | lncRNA | 158,6864754 | -0,82637299 | 0,297549185 | -2,777265174 | 0,005481844 | 0,035519156 |
| ENSG00000137642.13 | SORL1 | protein_coding | 7717,948161 | -0,827112838 | 0,149568391 | -5,529997553 | 3,20E-08 | 7,33E-07 |
| ENSG00000137699.17 | TRIM29 | protein_coding | 1376,63913 | -0,827491474 | 0,148760841 | -5,562562471 | 2,66E-08 | 6,14E-07 |
| ENSG00000117877.11 | POLR1G | protein_coding | 281,5593909 | -0,827899783 | 0,233736866 | -3,542016265 | 0,000397081 | 0,003986998 |
| ENSG00000165521.15 | EML5 | protein_coding | 2066,700555 | -0,8285795 | 0,145766504 | -5,68429289 | 1,31E-08 | 3,16E-07 |
| ENSG00000104691.16 | UBXN8 | protein_coding | 336,2711609 | -0,830109039 | 0,236993434 | -3,502666828 | 0,000460625 | 0,004516163 |
| ENSG00000071282.12 | LMCD1 | protein_coding | 607,0725256 | -0,830346755 | 0,17485401 | -4,748800174 | 2,05E-06 | 3,56E-05 |
| ENSG00000164039.15 | BDH2 | protein_coding | 643,7069317 | -0,830474926 | 0,185892801 | -4,467493752 | 7,91E-06 | 0,000122546 |
| ENSG00000165138.18 | ANKS6 | protein_coding | 664,5030491 | -0,831085282 | 0,19025058 | -4,36837187 | 1,25E-05 | 0,000186114 |
| ENSG00000131781.13 | FMO5 | protein_coding | 263,8453573 | -0,833740565 | 0,242772866 | -3,434241148 | 0,000594215 | 0,005585944 |
| ENSG00000168348.4 | INSM2 | protein_coding | 185,2510934 | -0,833750976 | 0,294735403 | -2,828811762 | 0,004672117 | 0,031219524 |

| | | | | | | | | |
|---|---|---|---|---|---|---|---|---|
| ENSG00000141526.17 | SLC16A3 | protein_coding | 2675,032845 | -0,83627197 | 0,167490544 | -4,992950359 | 5,95E-07 | 1,13E-05 |
| ENSG00000146904.9 | EPHA1 | protein_coding | 164,9269105 | -0,841251833 | 0,315088976 | -2,669886597 | 0,007587687 | 0,045936444 |
| ENSG00000115255.12 | REEP6 | protein_coding | 295,5063316 | -0,841349742 | 0,234311119 | -3,590737595 | 0,000329743 | 0,003380597 |
| ENSG00000167588.13 | GPD1 | protein_coding | 7352,758517 | -0,841494979 | 0,146813349 | -5,731733433 | 9,94E-09 | 2,42E-07 |
| ENSG00000116353.16 | MECR | protein_coding | 585,7124096 | -0,84203336 | 0,189240925 | -4,449530978 | 8,61E-06 | 0,00013257 |
| ENSG00000155380.12 | SLC16A1 | protein_coding | 4461,844351 | -0,842867856 | 0,118071833 | -7,138602288 | 9,43E-13 | 3,42E-11 |
| ENSG00000241697.5 | TMEFF1 | protein_coding | 675,8983155 | -0,844546006 | 0,19695312 | -4,288055994 | 1,80E-05 | 0,000257044 |
| ENSG00000136235.17 | GPNMB | protein_coding | 137,1100404 | -0,852952109 | 0,315926271 | -2,69984546 | 0,006937169 | 0,04289121 |
| ENSG00000175305.17 | CCNE2 | protein_coding | 3527,840242 | -0,854641103 | 0,151206793 | -5,652134319 | 1,58E-08 | 3,77E-07 |
| ENSG00000080200.10 | CRYBG3 | protein_coding | 969,6817076 | -0,854977064 | 0,160601619 | -5,323589308 | 1,02E-07 | 2,17E-06 |
| ENSG00000168243.11 | GNG4 | protein_coding | 14226,46168 | -0,857952366 | 0,129864314 | -6,606529063 | 3,93E-11 | 1,24E-09 |
| ENSG00000185100.11 | ADSS1 | protein_coding | 187,2293725 | -0,859048353 | 0,302943014 | -2,835676397 | 0,004572875 | 0,030752697 |
| ENSG00000170500.13 | LONRF2 | protein_coding | 1371,888206 | -0,85925485 | 0,143850732 | -5,973239348 | 2,33E-09 | 6,19E-08 |
| ENSG00000048342.17 | CC2D2A | protein_coding | 582,7121914 | -0,860311916 | 0,183741968 | -4,682174276 | 2,84E-06 | 4,79E-05 |
| ENSG00000006283.18 | CACNA1G | protein_coding | 381,2545863 | -0,862809234 | 0,251404394 | -3,431957656 | 0,000599241 | 0,00562309 |
| ENSG00000039560.14 | RAI14 | protein_coding | 600,8787063 | -0,863455799 | 0,176616433 | -4,888875764 | 1,01E-06 | 1,86E-05 |
| ENSG00000164683.18 | HEY1 | protein_coding | 409,3471688 | -0,865394407 | 0,231645687 | -3,735853736 | 0,000187079 | 0,002084357 |
| ENSG00000145284.12 | SCD5 | protein_coding | 804,753791 | -0,866023733 | 0,186859696 | -4,634620268 | 3,58E-06 | 5,95E-05 |
| ENSG00000251247.11 | ZNF345 | protein_coding | 301,7946401 | -0,8674375 | 0,249826585 | -3,472158493 | 0,000516291 | 0,004974027 |
| ENSG00000189337.17 | KAZN | protein_coding | 166,2796809 | -0,868588421 | 0,304797432 | -2,849723558 | 0,004375724 | 0,029663794 |
| ENSG00000127946.17 | HIP1 | protein_coding | 450,3966592 | -0,86976611 | 0,223424657 | -3,892883273 | 9,91E-05 | 0,001183686 |
| ENSG00000180914.10 | OXTR | protein_coding | 165,8148365 | -0,870192261 | 0,288502165 | -3,016241699 | 0,002559292 | 0,019094359 |
| ENSG00000236155.6 | ZPLD2P | transcribed_unproc | 271,2744553 | -0,871622004 | 0,277540881 | -3,140517538 | 0,001686496 | 0,013589991 |
| ENSG00000167767.14 | KRT80 | protein_coding | 1547,694775 | -0,872251213 | 0,156950517 | -5,557491799 | 2,74E-08 | 6,31E-07 |
| ENSG00000137474.22 | MYO7A | protein_coding | 649,3978111 | -0,872346067 | 0,206648062 | -4,221409374 | 2,43E-05 | 0,000335042 |
| ENSG00000156113.24 | KCNMA1 | protein_coding | 5912,859182 | -0,872522766 | 0,132413531 | -6,589377695 | 4,42E-11 | 1,39E-09 |
| ENSG00000246465.1 | AC138904.1 | lncRNA | 163,840015 | -0,874334623 | 0,306429898 | -2,853294107 | 0,004326856 | 0,029429085 |
| ENSG00000179978.11 | NAIPP2 | unprocessed_pseud | 210,8500411 | -0,874756013 | 0,265177142 | -3,298761 | 0,000971126 | 0,008467519 |
| ENSG00000093217.11 | XYLB | protein_coding | 313,7222489 | -0,875662561 | 0,230374422 | -3,801040731 | 0,00014409 | 0,001651076 |
| ENSG00000049323.16 | LTBP1 | protein_coding | 298,8073537 | -0,877580096 | 0,229147155 | -3,829766494 | 0,000128265 | 0,001491792 |
| ENSG00000069188.17 | SDK2 | protein_coding | 268,3596633 | -0,877784472 | 0,308127438 | -2,848770877 | 0,004388847 | 0,029715563 |
| ENSG00000049130.16 | KITLG | protein_coding | 695,1416097 | -0,878520998 | 0,179148485 | -4,903870665 | 9,40E-07 | 1,73E-05 |
| ENSG00000173638.19 | SLC19A1 | protein_coding | 818,5373606 | -0,879962514 | 0,196967663 | -4,467548139 | 7,91E-06 | 0,000122546 |
| ENSG00000164684.13 | ZNF704 | protein_coding | 801,3006693 | -0,881737566 | 0,201430838 | -4,377371277 | 1,20E-05 | 0,000179878 |
| ENSG00000196542.8 | SPTSSB | protein_coding | 781,2242157 | -0,883430431 | 0,182369824 | -4,844170003 | 1,27E-06 | 2,30E-05 |
| ENSG00000261526.2 | AC012615.1 | lncRNA | 318,0740535 | -0,884678024 | 0,22956395 | -3,853732367 | 0,000116331 | 0,001364361 |

| | | | | | | | | |
|---|---|---|---|---|---|---|---|---|
| ENSG00000180440.4 | SERTM1 | protein_coding | 246,0594796 | -0,884990446 | 0,25081684 | -3,528433117 | 0,000418028 | 0,004161392 |
| ENSG00000157445.15 | CACNA2D3 | protein_coding | 414,4869586 | -0,887281775 | 0,280050422 | -3,168292944 | 0,001533369 | 0,01253459 |
| ENSG00000205572.10 | SERF1B | protein_coding | 1484,539476 | -0,887645005 | 0,253493987 | -3,501641258 | 0,000462402 | 0,004527774 |
| ENSG00000169891.18 | REPS2 | protein_coding | 945,7985987 | -0,887858603 | 0,171823561 | -5,167269238 | 2,38E-07 | 4,81E-06 |
| ENSG00000123689.6 | G0S2 | protein_coding | 323,7263991 | -0,887920228 | 0,237523648 | -3,738239272 | 0,000185314 | 0,002067985 |
| ENSG00000152578.13 | GRIA4 | protein_coding | 6417,9043 | -0,893236747 | 0,125117871 | -7,139161957 | 9,39E-13 | 3,41E-11 |
| ENSG00000166949.16 | SMAD3 | protein_coding | 1320,838336 | -0,897283286 | 0,184454881 | -4,864513643 | 1,15E-06 | 2,09E-05 |
| ENSG00000171444.18 | MCC | protein_coding | 722,1525215 | -0,898004227 | 0,221243143 | -4,058901959 | 4,93E-05 | 0,000633628 |
| ENSG00000113231.14 | PDE8B | protein_coding | 564,11408 | -0,899591103 | 0,197239545 | -4,560906387 | 5,09E-06 | 8,20E-05 |
| ENSG00000175928.6 | LRRN1 | protein_coding | 175,3573174 | -0,900484039 | 0,290834033 | -3,096212738 | 0,001960097 | 0,015322863 |
| ENSG00000153012.12 | LGI2 | protein_coding | 346,3218155 | -0,901657675 | 0,218064826 | -4,134814816 | 3,55E-05 | 0,000472189 |
| ENSG00000147724.12 | FAM135B | protein_coding | 1581,47362 | -0,902436764 | 0,155370701 | -5,808281462 | 6,31E-09 | 1,58E-07 |
| ENSG00000123836.15 | PFKFB2 | protein_coding | 13202,52205 | -0,912099292 | 0,117635232 | -7,753623464 | 8,93E-15 | 3,75E-13 |
| ENSG00000279348.1 | AC012513.3 | TEC | 716,9850443 | -0,912130825 | 0,21243314 | -4,293731306 | 1,76E-05 | 0,00025107 |
| ENSG00000132840.10 | BHMT2 | protein_coding | 158,4670512 | -0,912321077 | 0,334956324 | -2,723701607 | 0,00645548 | 0,040441456 |
| ENSG00000105173.14 | CCNE1 | protein_coding | 820,7919578 | -0,913139415 | 0,171482084 | -5,324984351 | 1,01E-07 | 2,15E-06 |
| ENSG00000287281.1 | AC012101.2 | lncRNA | 1561,814488 | -0,913465124 | 0,142408347 | -6,414407166 | 1,41E-10 | 4,20E-09 |
| ENSG00000236618.2 | PITPNA-AS1 | lncRNA | 567,2388979 | -0,916420961 | 0,202495912 | -4,52562697 | 6,02E-06 | 9,54E-05 |
| ENSG00000245293.3 | CYP2U1-AS1 | lncRNA | 765,4930814 | -0,917582059 | 0,179832396 | -5,102429147 | 3,35E-07 | 6,65E-06 |
| ENSG00000133216.17 | EPHB2 | protein_coding | 1043,572778 | -0,91779133 | 0,196706323 | -4,665794756 | 3,07E-06 | 5,17E-05 |
| ENSG00000255857.6 | PXN-AS1 | lncRNA | 291,126372 | -0,919342142 | 0,246177982 | -3,734461283 | 0,000188117 | 0,002092581 |
| ENSG00000119408.17 | NEK6 | protein_coding | 467,8130383 | -0,919358568 | 0,202716017 | -4,535204385 | 5,75E-06 | 9,18E-05 |
| ENSG00000099617.4 | EFNA2 | protein_coding | 210,6191444 | -0,92164746 | 0,28361328 | -3,24966257 | 0,00115542 | 0,009820837 |
| ENSG00000163040.15 | CCDC74A | protein_coding | 551,5929671 | -0,921815628 | 0,185178596 | -4,977981505 | 6,43E-07 | 1,22E-05 |
| ENSG00000152102.18 | FAM168B | protein_coding | 10425,4565 | -0,921830489 | 0,114439445 | -8,05518141 | 7,94E-16 | 3,56E-14 |
| ENSG00000174599.5 | TRAM1L1 | protein_coding | 568,0034923 | -0,922413015 | 0,181457139 | -5,083365812 | 3,71E-07 | 7,30E-06 |
| ENSG00000110665.12 | C11orf21 | protein_coding | 181,993684 | -0,924054038 | 0,32232171 | -2,86686875 | 0,004145549 | 0,028533003 |
| ENSG00000188153.14 | COL4A5 | protein_coding | 227,4976771 | -0,925632064 | 0,334480755 | -2,767370172 | 0,005651055 | 0,03635674 |
| ENSG00000100290.3 | BIK | protein_coding | 468,2949307 | -0,925910034 | 0,198657962 | -4,66082519 | 3,15E-06 | 5,28E-05 |
| ENSG00000167191.12 | GPRC5B | protein_coding | 264,0734826 | -0,927217194 | 0,28286972 | -3,277894839 | 0,001045844 | 0,009025035 |
| ENSG00000163873.10 | GRIK3 | protein_coding | 3677,292438 | -0,92726851 | 0,131216978 | -7,066680879 | 1,59E-12 | 5,70E-11 |
| ENSG00000102967.12 | DHODH | protein_coding | 354,9201054 | -0,931753219 | 0,213295825 | -4,36836127 | 1,25E-05 | 0,000186114 |
| ENSG00000108352.13 | RAPGEFL1 | protein_coding | 211,0335434 | -0,933652481 | 0,30496469 | -3,061510108 | 0,002202236 | 0,016863131 |
| ENSG00000270175.1 | AC023509.3 | lncRNA | 228,6981327 | -0,936406997 | 0,286334434 | -3,270326185 | 0,001074235 | 0,00922063 |
| ENSG00000162009.8 | SSTR5 | protein_coding | 405,137752 | -0,937021979 | 0,204908034 | -4,57289038 | 4,81E-06 | 7,79E-05 |
| ENSG00000164849.10 | GPR146 | protein_coding | 192,1416233 | -0,93716242 | 0,278114057 | -3,369705324 | 0,000752486 | 0,006819609 |

| | | | | | | | | |
|---|---|---|---|---|---|---|---|---|
| ENSG00000135835.12 | KIAA1614 | protein_coding | 1098,775381 | -0,937537443 | 0,184670782 | -5,076804422 | 3,84E-07 | 7,52E-06 |
| ENSG00000168913.7 | ENHO | protein_coding | 3084,11927 | -0,938584606 | 0,154948563 | -6,05739471 | 1,38E-09 | 3,75E-08 |
| ENSG00000245750.10 | DRAIC | lncRNA | 437,9915145 | -0,939032726 | 0,220562346 | -4,257448025 | 2,07E-05 | 0,00029111 |
| ENSG00000117069.15 | ST6GALNAC5 | protein_coding | 4758,291062 | -0,939236844 | 0,132331747 | -7,097592703 | 1,27E-12 | 4,57E-11 |
| ENSG00000166073.11 | GPR176 | protein_coding | 1025,337866 | -0,941815843 | 0,167542982 | -5,621338678 | 1,89E-08 | 4,45E-07 |
| ENSG00000125266.8 | EFNB2 | protein_coding | 189,6804284 | -0,943701746 | 0,301529857 | -3,12971245 | 0,001749775 | 0,013992184 |
| ENSG00000237424.1 | FOXD2-AS1 | lncRNA | 144,2140169 | -0,945677346 | 0,337479559 | -2,802176664 | 0,005075907 | 0,033301194 |
| ENSG00000109072.14 | VTN | protein_coding | 5029,24407 | -0,946580417 | 0,173499193 | -5,455820284 | 4,87E-08 | 1,09E-06 |
| ENSG00000163050.18 | COQ8A | protein_coding | 2524,596772 | -0,947151724 | 0,166205681 | -5,698672365 | 1,21E-08 | 2,93E-07 |
| ENSG00000137727.12 | ARHGAP20 | protein_coding | 518,8836661 | -0,949020031 | 0,191532862 | -4,954867911 | 7,24E-07 | 1,36E-05 |
| ENSG00000285662.2 | FAM245B | lncRNA | 284,0557435 | -0,949066238 | 0,259096515 | -3,662983419 | 0,000249295 | 0,002670923 |
| ENSG00000223969.6 | AC002456.1 | lncRNA | 332,0509709 | -0,949190803 | 0,218392007 | -4,346270807 | 1,38E-05 | 0,000203147 |
| ENSG00000278238.1 | AL359513.1 | lncRNA | 153,3988284 | -0,950960222 | 0,298489485 | -3,185908618 | 0,001443002 | 0,011902774 |
| ENSG00000077942.19 | FBLN1 | protein_coding | 170,9954005 | -0,951385575 | 0,355215761 | -2,67833154 | 0,007398994 | 0,045067774 |
| ENSG00000157470.12 | FAM81A | protein_coding | 566,6744927 | -0,951682665 | 0,210499349 | -4,521071767 | 6,15E-06 | 9,73E-05 |
| ENSG00000137962.13 | ARHGAP29 | protein_coding | 927,9479773 | -0,952939965 | 0,158266401 | -6,021113508 | 1,73E-09 | 4,66E-08 |
| ENSG00000007062.12 | PROM1 | protein_coding | 189,6701653 | -0,953660323 | 0,332827719 | -2,865327218 | 0,004165785 | 0,02862183 |
| ENSG00000110324.11 | IL10RA | protein_coding | 599,8988305 | -0,953840129 | 0,20114346 | -4,742088704 | 2,12E-06 | 3,67E-05 |
| ENSG00000124194.17 | GDAP1L1 | protein_coding | 674,6232685 | -0,954079432 | 0,189769366 | -5,027573472 | 4,97E-07 | 9,59E-06 |
| ENSG00000231806.3 | PCAT7 | lncRNA | 343,1135122 | -0,954413424 | 0,241062184 | -3,959200099 | 7,52E-05 | 0,000926081 |
| ENSG00000198363.18 | ASPH | protein_coding | 19067,5424 | -0,955969963 | 0,115733324 | -8,260109831 | 1,46E-16 | 6,82E-15 |
| ENSG00000119927.14 | GPAM | protein_coding | 1131,043264 | -0,957230626 | 0,153357567 | -6,241821945 | 4,33E-10 | 1,24E-08 |
| ENSG00000185829.18 | ARL17A | protein_coding | 294,1743857 | -0,957893934 | 0,232380733 | -4,122088439 | 3,75E-05 | 0,000494658 |
| ENSG00000120594.17 | PLXDC2 | protein_coding | 3104,120992 | -0,959663515 | 0,141782151 | -6,768577758 | 1,30E-11 | 4,31E-10 |
| ENSG00000150893.11 | FREM2 | protein_coding | 766,4941022 | -0,960248977 | 0,198561758 | -4,836021742 | 1,32E-06 | 2,38E-05 |
| ENSG00000072274.13 | TFRC | protein_coding | 22045,40712 | -0,961686069 | 0,108861593 | -8,834025307 | 1,01E-18 | 5,21E-17 |
| ENSG00000169760.17 | NLGN1 | protein_coding | 267,1788557 | -0,962970599 | 0,241532143 | -3,98692525 | 6,69E-05 | 0,000833603 |
| ENSG00000160233.8 | LRRC3 | protein_coding | 1099,638597 | -0,963469462 | 0,172281253 | -5,592421964 | 2,24E-08 | 5,22E-07 |
| ENSG00000230453.9 | ANKRD18B | protein_coding | 286,7719676 | -0,964021008 | 0,247154789 | -3,900474726 | 9,60E-05 | 0,001155086 |
| ENSG00000165449.11 | SLC16A9 | protein_coding | 5059,44254 | -0,964948715 | 0,127695613 | -7,556631703 | 4,14E-14 | 1,67E-12 |
| ENSG00000158258.16 | CLSTN2 | protein_coding | 2916,343758 | -0,965647331 | 0,174514733 | -5,53332841 | 3,14E-08 | 7,20E-07 |
| ENSG00000105664.11 | COMP | protein_coding | 161,0894577 | -0,966043658 | 0,333840149 | -2,893731206 | 0,003806939 | 0,026583812 |
| ENSG00000143416.21 | SELENBP1 | protein_coding | 224,2982307 | -0,966449365 | 0,25857268 | -3,737631392 | 0,000185762 | 0,002071885 |
| ENSG00000288670.1 | AL592295.6 | lncRNA | 308,3982992 | -0,967940784 | 0,246998636 | -3,918810237 | 8,90E-05 | 0,001079752 |
| ENSG00000151835.16 | SACS | protein_coding | 2522,93702 | -0,973891578 | 0,148183397 | -6,572204425 | 4,96E-11 | 1,54E-09 |
| ENSG00000146360.8 | GPR6 | protein_coding | 249,1527569 | -0,975273323 | 0,252759311 | -3,858506019 | 0,000114082 | 0,001342503 |

| | | | | | | | | |
|---|---|---|---|---|---|---|---|---|
| ENSG00000144504.15 | ANKMY1 | protein_coding | 240,2150426 | -0,975649824 | 0,295774898 | -3,2986228 | 0,000971604 | 0,008468162 |
| ENSG00000130558.20 | OLFM1 | protein_coding | 3349,915449 | -0,975658129 | 0,135782127 | -7,185468028 | 6,70E-13 | 2,46E-11 |
| ENSG00000255135.4 | AP002360.1 | lncRNA | 488,9499437 | -0,976432344 | 0,220269152 | -4,432905534 | 9,30E-06 | 0,000142174 |
| ENSG00000105948.13 | TTC26 | protein_coding | 1264,352086 | -0,976715236 | 0,199587032 | -4,893680845 | 9,90E-07 | 1,81E-05 |
| ENSG00000240086.7 | AC092969.1 | lncRNA | 523,3344253 | -0,977228338 | 0,215577238 | -4,533077559 | 5,81E-06 | 9,25E-05 |
| ENSG00000083857.14 | FAT1 | protein_coding | 1232,937892 | -0,977477573 | 0,158639446 | -6,161630009 | 7,20E-10 | 2,01E-08 |
| ENSG00000105647.19 | PIK3R2 | protein_coding | 9917,07414 | -0,977862208 | 0,132026832 | -7,406541485 | 1,30E-13 | 5,04E-12 |
| ENSG00000133466.14 | C1QTNF6 | protein_coding | 2522,898335 | -0,978025336 | 0,132630471 | -7,374062123 | 1,66E-13 | 6,34E-12 |
| ENSG00000185742.7 | C11orf87 | protein_coding | 293,2068942 | -0,979125223 | 0,233353847 | -4,195882078 | 2,72E-05 | 0,00037192 |
| ENSG00000183798.5 | EMILIN3 | protein_coding | 2390,798927 | -0,980166246 | 0,146777194 | -6,677919217 | 2,42E-11 | 7,88E-10 |
| ENSG00000064270.13 | ATP2C2 | protein_coding | 774,6518871 | -0,980430531 | 0,169133952 | -5,796769487 | 6,76E-09 | 1,69E-07 |
| ENSG00000103546.18 | SLC6A2 | protein_coding | 114,1127957 | -0,98424817 | 0,354414021 | -2,777114086 | 0,005484392 | 0,035524677 |
| ENSG00000020181.18 | ADGRA2 | protein_coding | 165,0151648 | -0,987946542 | 0,28959148 | -3,411517984 | 0,000646022 | 0,005994893 |
| ENSG00000004809.14 | SLC22A16 | protein_coding | 631,8684827 | -0,988048369 | 0,218241236 | -4,5273221 | 5,97E-06 | 9,47E-05 |
| ENSG00000115165.10 | CYTIP | protein_coding | 101,5457757 | -0,988569243 | 0,360372476 | -2,74318742 | 0,006084595 | 0,038647145 |
| ENSG00000113196.3 | HAND1 | protein_coding | 233,1336983 | -0,989194989 | 0,249215256 | -3,969239309 | 7,21E-05 | 0,000892646 |
| ENSG00000154358.22 | OBSCN | protein_coding | 210,1214676 | -0,990385692 | 0,330351106 | -2,997979043 | 0,002717764 | 0,020048055 |
| ENSG00000128573.26 | FOXP2 | protein_coding | 1079,063212 | -0,990437675 | 0,22654942 | -4,371839366 | 1,23E-05 | 0,000183709 |
| ENSG00000170962.13 | PDGFD | protein_coding | 4892,648241 | -0,992875208 | 0,129074352 | -7,692273414 | 1,45E-14 | 6,02E-13 |
| ENSG00000169860.7 | P2RY1 | protein_coding | 237,0557487 | -0,998298252 | 0,270373621 | -3,692291613 | 0,000222242 | 0,00243212 |
| ENSG00000100767.17 | PAPLN | protein_coding | 354,2941175 | -0,999601304 | 0,253074741 | -3,949826449 | 7,82E-05 | 0,00095972 |
| ENSG00000074317.11 | SNCB | protein_coding | 328,6543842 | -1,009029614 | 0,218705016 | -4,613655555 | 3,96E-06 | 6,53E-05 |
| ENSG00000188107.15 | EYS | protein_coding | 179,188528 | -1,009035146 | 0,331919346 | -3,04000101 | 0,002365774 | 0,017892872 |
| ENSG00000204219.11 | TCEA3 | protein_coding | 127,3859989 | -1,009910022 | 0,337132216 | -2,995590382 | 0,002739141 | 0,020170369 |
| ENSG00000120820.12 | GLT8D2 | protein_coding | 541,6054245 | -1,0116626 | 0,199285482 | -5,076449082 | 3,85E-07 | 7,53E-06 |
| ENSG00000236810.6 | ELOA-AS1 | lncRNA | 149,5605389 | -1,012222792 | 0,303958278 | -3,330137273 | 0,000868032 | 0,007687075 |
| ENSG00000167619.13 | TMEM145 | protein_coding | 349,4626065 | -1,01291383 | 0,274652532 | -3,687982861 | 0,000226039 | 0,00245697 |
| ENSG00000110148.10 | CCKBR | protein_coding | 139,8794895 | -1,01301395 | 0,310565424 | -3,261837509 | 0,001106926 | 0,00946243 |
| ENSG00000285542.1 | AC013717.1 | protein_coding | 123,3810243 | -1,013433521 | 0,350742283 | -2,889396488 | 0,003859821 | 0,02691717 |
| ENSG00000137124.8 | ALDH1B1 | protein_coding | 2025,787661 | -1,013640332 | 0,152022209 | -6,66771218 | 2,60E-11 | 8,37E-10 |
| ENSG00000136546.16 | SCN7A | protein_coding | 137,9961735 | -1,017744064 | 0,386060563 | -2,636229029 | 0,008383313 | 0,049766367 |
| ENSG00000157087.20 | ATP2B2 | protein_coding | 123,1592071 | -1,019843089 | 0,327452049 | -3,114480706 | 0,00184269 | 0,01455176 |
| ENSG00000287038.1 | AL162388.2 | lncRNA | 525,3863293 | -1,020682076 | 0,193367289 | -5,278462979 | 1,30E-07 | 2,73E-06 |
| ENSG00000182747.5 | SLC35D3 | protein_coding | 92,02715889 | -1,021010164 | 0,37652006 | -2,711701907 | 0,006693876 | 0,041645258 |
| ENSG00000184916.9 | JAG2 | protein_coding | 381,8543398 | -1,028192508 | 0,224805002 | -4,573708324 | 4,79E-06 | 7,76E-05 |
| ENSG00000188859.7 | FAM78B | protein_coding | 222,1639479 | -1,030715791 | 0,262498723 | -3,926555457 | 8,62E-05 | 0,001047614 |

| | | | | | | | | |
|---|---|---|---|---|---|---|---|---|
| ENSG00000152954.12 | NRSN1 | protein_coding | 606,2761868 | -1,035504271 | 0,204907502 | -5,053520536 | 4,34E-07 | 8,44E-06 |
| ENSG00000174611.12 | KY | protein_coding | 373,7340336 | -1,038147794 | 0,228875586 | -4,535860776 | 5,74E-06 | 9,16E-05 |
| ENSG00000242419.5 | PCDHGC4 | protein_coding | 196,7674137 | -1,038582852 | 0,34938114 | -2,972635708 | 0,002952547 | 0,021460801 |
| ENSG00000152990.14 | ADGRA3 | protein_coding | 1616,134559 | -1,041514715 | 0,140223689 | -7,42752328 | 1,11E-13 | 4,33E-12 |
| ENSG00000281128.2 | PTENP1-AS | lncRNA | 168,8265312 | -1,042633988 | 0,340523097 | -3,061859822 | 0,002199664 | 0,016849606 |
| ENSG00000161714.12 | PLCD3 | protein_coding | 260,3990631 | -1,043578137 | 0,298986968 | -3,490380012 | 0,000482334 | 0,004700962 |
| ENSG00000147394.18 | ZNF185 | protein_coding | 294,3241986 | -1,045762229 | 0,267938451 | -3,902994227 | 9,50E-05 | 0,001144438 |
| ENSG00000259881.1 | AC092384.2 | lncRNA | 215,4956551 | -1,04761917 | 0,315305391 | -3,322553944 | 0,000891974 | 0,007879106 |
| ENSG00000188613.7 | NANOS1 | protein_coding | 1547,419662 | -1,051889349 | 0,142164588 | -7,399095412 | 1,37E-13 | 5,31E-12 |
| ENSG00000231764.10 | DLX6-AS1 | lncRNA | 932,6673334 | -1,054063893 | 0,182013384 | -5,791133986 | 6,99E-09 | 1,74E-07 |
| ENSG00000264404.3 | LINC02675 | lncRNA | 109,5689601 | -1,054305294 | 0,372238489 | -2,832338208 | 0,004620894 | 0,031015846 |
| ENSG00000262655.4 | SPON1 | protein_coding | 18509,18123 | -1,054485593 | 0,160439712 | -6,572472481 | 4,95E-11 | 1,54E-09 |
| ENSG00000106003.13 | LFNG | protein_coding | 1299,12986 | -1,055704085 | 0,163411915 | -6,46038624 | 1,04E-10 | 3,14E-09 |
| ENSG00000152315.5 | KCNK13 | protein_coding | 121,8184219 | -1,059248382 | 0,365344381 | -2,899314826 | 0,003739792 | 0,026202323 |
| ENSG00000122783.17 | CYREN | protein_coding | 3242,994894 | -1,059350658 | 0,129451342 | -8,183388787 | 2,76E-16 | 1,26E-14 |
| ENSG00000100351.17 | GRAP2 | protein_coding | 128,2351095 | -1,061994587 | 0,331567453 | -3,20295185 | 0,001360267 | 0,011336463 |
| ENSG00000132837.15 | DMGDH | protein_coding | 178,8251041 | -1,065459573 | 0,290848327 | -3,663282451 | 0,000249004 | 0,002669172 |
| ENSG00000116106.12 | EPHA4 | protein_coding | 1634,5831 | -1,06605966 | 0,212514712 | -5,016404037 | 5,26E-07 | 1,01E-05 |
| ENSG00000260260.1 | SNHG19 | lncRNA | 530,2803568 | -1,066915106 | 0,213022777 | -5,00845554 | 5,49E-07 | 1,05E-05 |
| ENSG00000267260.2 | AC020928.1 | lncRNA | 139,1402827 | -1,068504712 | 0,318523934 | -3,354550783 | 0,00079494 | 0,007142515 |
| ENSG00000204323.5 | SMIM5 | protein_coding | 295,4330015 | -1,070446369 | 0,277497356 | -3,857501149 | 0,000114552 | 0,001345264 |
| ENSG00000144681.10 | STAC | protein_coding | 196,7682474 | -1,071034231 | 0,350886769 | -3,052364256 | 0,002270464 | 0,017284423 |
| ENSG00000286340.1 | AL356776.2 | lncRNA | 163,759471 | -1,07160727 | 0,291478791 | -3,676450231 | 0,000236502 | 0,002554783 |
| ENSG00000140600.17 | SH3GL3 | protein_coding | 432,1862801 | -1,071680366 | 0,22659818 | -4,729430593 | 2,25E-06 | 3,88E-05 |
| ENSG00000102452.18 | NALCN | protein_coding | 312,9566489 | -1,074114552 | 0,237685955 | -4,519049312 | 6,21E-06 | 9,81E-05 |
| ENSG00000196932.12 | TMEM26 | protein_coding | 414,8624591 | -1,0778364 | 0,223917529 | -4,813541867 | 1,48E-06 | 2,64E-05 |
| ENSG00000227877.7 | MRLN | protein_coding | 1603,271553 | -1,078576263 | 0,174740321 | -6,172452119 | 6,72E-10 | 1,89E-08 |
| ENSG00000275163.1 | AC117457.1 | protein_coding | 102,7062699 | -1,078946117 | 0,393257182 | -2,743614525 | 0,006076685 | 0,038608618 |
| ENSG00000078053.17 | AMPH | protein_coding | 161,440602 | -1,081063951 | 0,31162173 | -3,469154582 | 0,000522099 | 0,005018497 |
| ENSG00000150394.14 | CDH8 | protein_coding | 634,1628861 | -1,082143627 | 0,189478726 | -5,711161616 | 1,12E-08 | 2,73E-07 |
| ENSG00000068615.19 | REEP1 | protein_coding | 6126,240752 | -1,084287044 | 0,116888456 | -9,276254317 | 1,76E-20 | 9,85E-19 |
| ENSG00000187240.16 | DYNC2H1 | protein_coding | 695,4518137 | -1,093309644 | 0,257134205 | -4,251902799 | 2,12E-05 | 0,000297016 |
| ENSG00000142798.20 | HSPG2 | protein_coding | 245,8187545 | -1,099710816 | 0,365965961 | -3,004953831 | 0,002656212 | 0,019640624 |
| ENSG00000134250.20 | NOTCH2 | protein_coding | 425,440341 | -1,100483862 | 0,25358625 | -4,339682693 | 1,43E-05 | 0,000208602 |
| ENSG00000176771.17 | NCKAP5 | protein_coding | 187,1476479 | -1,106070531 | 0,323513828 | -3,418928136 | 0,000628683 | 0,005864942 |
| ENSG00000223508.5 | RPL23AP53 | transcribed_process | 106,0447726 | -1,108085514 | 0,348791095 | -3,176931779 | 0,00148842 | 0,01221485 |

| | | | | | | | | |
|---|---|---|---|---|---|---|---|---|
| ENSG00000118515.11 | SGK1 | protein_coding | 173,9727774 | -1,109785419 | 0,293715629 | -3,778435017 | 0,000157817 | 0,001786857 |
| ENSG00000160145.15 | KALRN | protein_coding | 726,0679546 | -1,110054018 | 0,218522327 | -5,079819686 | 3,78E-07 | 7,42E-06 |
| ENSG00000143195.13 | ILDR2 | protein_coding | 2419,237045 | -1,111529086 | 0,217223872 | -5,116974825 | 3,10E-07 | 6,21E-06 |
| ENSG00000183762.13 | KREMEN1 | protein_coding | 788,7870246 | -1,112861879 | 0,204974998 | -5,429256681 | 5,66E-08 | 1,25E-06 |
| ENSG00000138640.15 | FAM13A | protein_coding | 412,0819314 | -1,117060869 | 0,217367519 | -5,139042278 | 2,76E-07 | 5,56E-06 |
| ENSG00000116852.15 | KIF21B | protein_coding | 179,2664336 | -1,118462427 | 0,300706118 | -3,719453512 | 0,000199654 | 0,002203354 |
| ENSG00000187994.14 | RINL | protein_coding | 925,2528364 | -1,119228769 | 0,15655386 | -7,149161103 | 8,73E-13 | 3,18E-11 |
| ENSG00000130829.18 | DUSP9 | protein_coding | 2362,514258 | -1,124863695 | 0,168051354 | -6,693571155 | 2,18E-11 | 7,11E-10 |
| ENSG00000179241.13 | LDLRAD3 | protein_coding | 764,2178869 | -1,130704941 | 0,171702133 | -6,585270197 | 4,54E-11 | 1,43E-09 |
| ENSG00000106236.4 | NPTX2 | protein_coding | 13227,67477 | -1,131845344 | 0,114168772 | -9,913790989 | 3,63E-23 | 2,41E-21 |
| ENSG00000189007.16 | ADAT2 | protein_coding | 187,2278652 | -1,136059455 | 0,275343119 | -4,125977293 | 3,69E-05 | 0,000487907 |
| ENSG00000260063.1 | AL512408.1 | lncRNA | 85,83796012 | -1,140823809 | 0,398458714 | -2,863091631 | 0,004195292 | 0,028786772 |
| ENSG00000197872.11 | CYRIA | protein_coding | 877,7446115 | -1,142560392 | 0,164087604 | -6,963112182 | 3,33E-12 | 1,18E-10 |
| ENSG00000158055.16 | GRHL3 | protein_coding | 481,4774944 | -1,142876681 | 0,252279447 | -4,530201308 | 5,89E-06 | 9,35E-05 |
| ENSG00000286237.1 | ARMCX5-GPRAS | protein_coding | 201,5632519 | -1,143318332 | 0,307241777 | -3,721233306 | 0,000198252 | 0,002189093 |
| ENSG00000106688.12 | SLC1A1 | protein_coding | 169,3971613 | -1,147890924 | 0,295580657 | -3,883511645 | 0,000102959 | 0,001223985 |
| ENSG00000237984.3 | PTENP1 | transcribed_process | 242,6494431 | -1,161783254 | 0,256294111 | -4,533007999 | 5,81E-06 | 9,25E-05 |
| ENSG00000117971.12 | CHRNB4 | protein_coding | 549,1612845 | -1,163026394 | 0,195929046 | -5,935957 | 2,92E-09 | 7,67E-08 |
| ENSG00000183018.9 | SPNS2 | protein_coding | 671,5554636 | -1,164504322 | 0,17846916 | -6,524961082 | 6,80E-11 | 2,08E-09 |
| ENSG00000184949.16 | FAM227A | protein_coding | 156,3411845 | -1,172380431 | 0,303689416 | -3,860458638 | 0,000113174 | 0,001335573 |
| ENSG00000124126.14 | PREX1 | protein_coding | 91,09955478 | -1,174851189 | 0,38377453 | -3,061305786 | 0,002203739 | 0,016868474 |
| ENSG00000130037.5 | KCNA5 | protein_coding | 313,6723705 | -1,178372287 | 0,318979786 | -3,694191098 | 0,000220588 | 0,002419071 |
| ENSG00000224043.8 | CCNT2-AS1 | lncRNA | 609,6035975 | -1,179034797 | 0,22210077 | -5,308557911 | 1,10E-07 | 2,34E-06 |
| ENSG00000163673.7 | DCLK3 | protein_coding | 129,163747 | -1,1800145 | 0,326916817 | -3,609525231 | 0,000306758 | 0,003179966 |
| ENSG00000154330.13 | PGM5 | protein_coding | 2610,524931 | -1,182345779 | 0,190126769 | -6,218723346 | 5,01E-10 | 1,42E-08 |
| ENSG00000119681.12 | LTBP2 | protein_coding | 260,6655707 | -1,185304203 | 0,251144312 | -4,719613964 | 2,36E-06 | 4,04E-05 |
| ENSG00000100234.12 | TIMP3 | protein_coding | 1311,403196 | -1,188312926 | 0,151585833 | -7,839208316 | 4,53E-15 | 1,94E-13 |
| ENSG00000065621.15 | GSTO2 | protein_coding | 1091,41228 | -1,192733703 | 0,183996294 | -6,48237894 | 9,03E-11 | 2,73E-09 |
| ENSG00000104998.4 | IL27RA | protein_coding | 369,5040514 | -1,194791165 | 0,229457883 | -5,207017297 | 1,92E-07 | 3,93E-06 |
| ENSG00000180044.5 | C3orf80 | protein_coding | 1228,254859 | -1,199036721 | 0,157984372 | -7,589590721 | 3,21E-14 | 1,31E-12 |
| ENSG00000138735.16 | PDE5A | protein_coding | 7352,054491 | -1,199859589 | 0,126819061 | -9,4611928 | 3,04E-21 | 1,80E-19 |
| ENSG00000126878.13 | AIF1L | protein_coding | 228,9334773 | -1,211212014 | 0,290145921 | -4,174492642 | 2,99E-05 | 0,000404415 |
| ENSG00000170624.13 | SGCD | protein_coding | 4677,595402 | -1,216842856 | 0,127000831 | -9,58137709 | 9,58E-22 | 5,73E-20 |
| ENSG00000178882.15 | RFLNA | protein_coding | 379,2468963 | -1,223333722 | 0,226677458 | -5,396803596 | 6,78E-08 | 1,49E-06 |
| ENSG00000204588.5 | LINC01123 | lncRNA | 133,2328007 | -1,22368314 | 0,332531103 | -3,679905814 | 0,00023332 | 0,002525624 |
| ENSG00000205795.4 | CYS1 | protein_coding | 1339,589181 | -1,224036898 | 0,186998315 | -6,54571084 | 5,92E-11 | 1,82E-09 |

| | | | | | | | | |
|---|---|---|---|---|---|---|---|---|
| ENSG00000164741.15 | DLC1 | protein_coding | 131,533087 | -1,225641221 | 0,337065104 | -3,636215101 | 0,000276673 | 0,002910846 |
| ENSG00000143434.16 | SEMA6C | protein_coding | 194,6905867 | -1,226901113 | 0,307952263 | -3,984062665 | 6,77E-05 | 0,000842397 |
| ENSG00000161551.14 | ZNF577 | protein_coding | 105,6385078 | -1,233860927 | 0,392533751 | -3,14332442 | 0,001670406 | 0,013501911 |
| ENSG00000176273.15 | SLC35G1 | protein_coding | 1212,843607 | -1,236841805 | 0,148967974 | -8,302736313 | 1,02E-16 | 4,80E-15 |
| ENSG00000184792.16 | OSBP2 | protein_coding | 3335,659963 | -1,239487905 | 0,136482061 | -9,081690981 | 1,07E-19 | 5,80E-18 |
| ENSG00000180730.5 | SHISA2 | protein_coding | 221,2745541 | -1,243606122 | 0,260225732 | -4,778951391 | 1,76E-06 | 3,11E-05 |
| ENSG00000113319.13 | RASGRF2 | protein_coding | 248,1848269 | -1,244906945 | 0,329120966 | -3,782520936 | 0,000155248 | 0,001759963 |
| ENSG00000172575.12 | RASGRP1 | protein_coding | 590,214892 | -1,247762008 | 0,19226958 | -6,489648586 | 8,60E-11 | 2,61E-09 |
| ENSG00000219438.9 | TAFA5 | protein_coding | 2064,519488 | -1,250921538 | 0,133102472 | -9,398184083 | 5,55E-21 | 3,18E-19 |
| ENSG00000267296.2 | CEBPA-DT | lncRNA | 90,8391986 | -1,251564369 | 0,43603659 | -2,870319595 | 0,004100571 | 0,028266532 |
| ENSG00000198835.4 | GJC2 | protein_coding | 1330,634803 | -1,255396183 | 0,227713807 | -5,513043771 | 3,53E-08 | 8,03E-07 |
| ENSG00000229086.5 | LINC01548 | lncRNA | 255,9549412 | -1,256612114 | 0,337687617 | -3,721226511 | 0,000198258 | 0,002189093 |
| ENSG00000165617.15 | DACT1 | protein_coding | 522,070273 | -1,261733241 | 0,189881869 | -6,644832627 | 3,04E-11 | 9,67E-10 |
| ENSG00000120913.23 | PDLIM2 | protein_coding | 221,020299 | -1,262977782 | 0,273720825 | -4,614109211 | 3,95E-06 | 6,52E-05 |
| ENSG00000168032.10 | ENTPD3 | protein_coding | 586,4264435 | -1,267991677 | 0,196030613 | -6,468335005 | 9,91E-11 | 2,99E-09 |
| ENSG00000141449.15 | GREB1L | protein_coding | 257,0209021 | -1,280311183 | 0,272439771 | -4,699428339 | 2,61E-06 | 4,42E-05 |
| ENSG00000138722.10 | MMRN1 | protein_coding | 2097,10939 | -1,281356109 | 0,143369984 | -8,937408483 | 3,98E-19 | 2,10E-17 |
| ENSG00000186897.5 | C1QL4 | protein_coding | 67,80991501 | -1,282984148 | 0,465518234 | -2,756034143 | 0,005850689 | 0,037355862 |
| ENSG00000255198.4 | SNHG9 | lncRNA | 74,27061434 | -1,284443594 | 0,421628429 | -3,046387544 | 0,002316091 | 0,017580625 |
| ENSG00000184489.12 | PTP4A3 | protein_coding | 84,66824003 | -1,289303354 | 0,396810351 | -3,249167648 | 0,001157432 | 0,009833948 |
| ENSG00000184156.17 | KCNQ3 | protein_coding | 180,3025542 | -1,29542598 | 0,303183156 | -4,272750494 | 1,93E-05 | 0,000273112 |
| ENSG00000072182.13 | ASIC4 | protein_coding | 302,0241433 | -1,295460278 | 0,235416746 | -5,502838263 | 3,74E-08 | 8,50E-07 |
| ENSG00000128602.11 | SMO | protein_coding | 182,0026156 | -1,297505532 | 0,302369432 | -4,29112667 | 1,78E-05 | 0,00025386 |
| ENSG00000170091.11 | NSG2 | protein_coding | 211,2924966 | -1,303404456 | 0,300186443 | -4,341983083 | 1,41E-05 | 0,000206718 |
| ENSG00000136541.15 | ERMN | protein_coding | 795,9913159 | -1,305336001 | 0,207221093 | -6,299242911 | 2,99E-10 | 8,68E-09 |
| ENSG00000173406.16 | DAB1 | protein_coding | 357,2186114 | -1,307447884 | 0,373294781 | -3,502454228 | 0,000460993 | 0,004516163 |
| ENSG00000111907.21 | TPD52L1 | protein_coding | 128,9424975 | -1,324836451 | 0,363829606 | -3,641365158 | 0,000271196 | 0,002864751 |
| ENSG00000183723.13 | CMTM4 | protein_coding | 10208,68191 | -1,334550569 | 0,112366673 | -11,87674715 | 1,56E-32 | 1,40E-30 |
| ENSG00000143858.12 | SYT2 | protein_coding | 133,279712 | -1,334976721 | 0,321123466 | -4,157206999 | 3,22E-05 | 0,00043317 |
| ENSG00000169562.13 | GJB1 | protein_coding | 150,9203109 | -1,338178914 | 0,322669866 | -4,14720758 | 3,37E-05 | 0,000449062 |
| ENSG00000173926.6 | MARCHF3 | protein_coding | 440,8907289 | -1,338913698 | 0,314425789 | -4,25828207 | 2,06E-05 | 0,000290222 |
| ENSG00000171724.3 | VAT1L | protein_coding | 112,3521477 | -1,344214979 | 0,360472042 | -3,729040876 | 0,00019221 | 0,002132441 |
| ENSG00000123119.12 | NECAB1 | protein_coding | 138,4321731 | -1,347451783 | 0,348830335 | -3,862771228 | 0,000112108 | 0,001324481 |
| ENSG00000169758.13 | TMEM266 | protein_coding | 75,00716706 | -1,356372125 | 0,447150149 | -3,033370617 | 0,002418384 | 0,018211823 |
| ENSG00000084636.18 | COL16A1 | protein_coding | 137,6682756 | -1,359284007 | 0,354850697 | -3,830580066 | 0,000127842 | 0,001487694 |
| ENSG00000147041.12 | SYTL5 | protein_coding | 207,8263946 | -1,359394515 | 0,282225304 | -4,816699626 | 1,46E-06 | 2,61E-05 |

| | | | | | | | | |
|---|---|---|---|---|---|---|---|---|
| ENSG00000119227.8 | PIGZ | protein_coding | 102,7968127 | -1,384282552 | 0,374414552 | -3,697192173 | 0,000217997 | 0,002394425 |
| ENSG00000153789.13 | CIBAR2 | protein_coding | 932,185961 | -1,384814907 | 0,220567873 | -6,27840712 | 3,42E-10 | 9,86E-09 |
| ENSG00000235026.6 | DPP10-AS1 | lncRNA | 805,7730863 | -1,385650183 | 0,183854883 | -7,536651536 | 4,82E-14 | 1,94E-12 |
| ENSG00000125384.7 | PTGER2 | protein_coding | 894,6951861 | -1,39247608 | 0,176461578 | -7,891100672 | 3,00E-15 | 1,29E-13 |
| ENSG00000132334.17 | PTPRE | protein_coding | 744,2591761 | -1,392786728 | 0,185263386 | -7,517873663 | 5,57E-14 | 2,22E-12 |
| ENSG00000113594.10 | LIFR | protein_coding | 231,4394567 | -1,392944203 | 0,305887355 | -4,553781577 | 5,27E-06 | 8,46E-05 |
| ENSG00000004838.14 | ZMYND10 | protein_coding | 131,0137822 | -1,39458706 | 0,371724384 | -3,751669572 | 0,000175661 | 0,001975006 |
| ENSG00000117600.12 | PLPPR4 | protein_coding | 437,3310704 | -1,396114095 | 0,20391401 | -6,846582502 | 7,56E-12 | 2,58E-10 |
| ENSG00000100433.16 | KCNK10 | protein_coding | 97,89098715 | -1,401719283 | 0,366640422 | -3,823144421 | 0,000131761 | 0,00152314 |
| ENSG00000224165.6 | DNAJC27-AS1 | lncRNA | 62,72412605 | -1,404752959 | 0,496200127 | -2,831020957 | 0,004639968 | 0,031069099 |
| ENSG00000158813.18 | EDA | protein_coding | 1059,01192 | -1,406598483 | 0,160009353 | -8,790726648 | 1,49E-18 | 7,61E-17 |
| ENSG00000225096.3 | AL445250.1 | lncRNA | 118,7018984 | -1,412377935 | 0,346238424 | -4,079206228 | 4,52E-05 | 0,000585425 |
| ENSG00000077348.9 | EXOSC5 | protein_coding | 737,401063 | -1,412883005 | 0,191543486 | -7,376304113 | 1,63E-13 | 6,25E-12 |
| ENSG00000175697.11 | GPR156 | protein_coding | 93,84568869 | -1,418884862 | 0,378935439 | -3,744397372 | 0,000180827 | 0,002022232 |
| ENSG00000167216.17 | KATNAL2 | protein_coding | 74,67711993 | -1,421697213 | 0,454318152 | -3,129298724 | 0,001752241 | 0,014001206 |
| ENSG00000166407.14 | LMO1 | protein_coding | 283,6631412 | -1,423376391 | 0,267466193 | -5,321705807 | 1,03E-07 | 2,19E-06 |
| ENSG00000236502.1 | SIX3-AS1 | lncRNA | 233,5827162 | -1,428988393 | 0,255353957 | -5,596108289 | 2,19E-08 | 5,13E-07 |
| ENSG00000110900.16 | TSPAN11 | protein_coding | 111,557171 | -1,431568666 | 0,356178892 | -4,019240604 | 5,84E-05 | 0,000735446 |
| ENSG00000144339.12 | TMEFF2 | protein_coding | 463,6794739 | -1,443998493 | 0,231623389 | -6,23425163 | 4,54E-10 | 1,30E-08 |
| ENSG00000048540.15 | LMO3 | protein_coding | 116,8893125 | -1,449025264 | 0,365714605 | -3,962174993 | 7,43E-05 | 0,000917003 |
| ENSG00000058335.16 | RASGRF1 | protein_coding | 394,1044302 | -1,450771795 | 0,294454082 | -4,926988224 | 8,35E-07 | 1,55E-05 |
| ENSG00000171812.13 | COL8A2 | protein_coding | 83,94394142 | -1,451491972 | 0,450422537 | -3,222511872 | 0,001270719 | 0,010718172 |
| ENSG00000103647.13 | CORO2B | protein_coding | 67,64703553 | -1,452076323 | 0,453326914 | -3,203154895 | 0,001359308 | 0,011332984 |
| ENSG00000175093.5 | SPSB4 | protein_coding | 377,7152936 | -1,453382656 | 0,227153645 | -6,398236113 | 1,57E-10 | 4,65E-09 |
| ENSG00000270344.3 | POC1B-AS1 | lncRNA | 114,8434648 | -1,457009623 | 0,340145968 | -4,28348345 | 1,84E-05 | 0,000261495 |
| ENSG00000176438.12 | SYNE3 | protein_coding | 321,9354626 | -1,462354333 | 0,310711281 | -4,706473245 | 2,52E-06 | 4,28E-05 |
| ENSG00000112293.15 | GPLD1 | protein_coding | 167,6474409 | -1,462784942 | 0,307212787 | -4,761471542 | 1,92E-06 | 3,36E-05 |
| ENSG00000184709.7 | LRRC26 | protein_coding | 88,00877666 | -1,465903425 | 0,519982707 | -2,819138802 | 0,004815269 | 0,031961845 |
| ENSG00000135625.8 | EGR4 | protein_coding | 73,29763778 | -1,47109305 | 0,432547877 | -3,400994728 | 0,000671411 | 0,00618515 |
| ENSG00000141540.11 | TTYH2 | protein_coding | 251,5677137 | -1,47398824 | 0,259494062 | -5,680238809 | 1,35E-08 | 3,24E-07 |
| ENSG00000178921.14 | PFAS | protein_coding | 1561,347408 | -1,475014587 | 0,158960074 | -9,279151367 | 1,71E-20 | 9,64E-19 |
| ENSG00000168743.13 | NPNT | protein_coding | 1289,537286 | -1,477836976 | 0,157028385 | -9,411272822 | 4,90E-21 | 2,83E-19 |
| ENSG00000111344.11 | RASAL1 | protein_coding | 199,7408708 | -1,486545504 | 0,300633667 | -4,944707348 | 7,63E-07 | 1,42E-05 |
| ENSG00000186603.6 | HPDL | protein_coding | 71,16977539 | -1,48716933 | 0,480206087 | -3,096939769 | 0,001955296 | 0,015291043 |
| ENSG00000244694.7 | PTCHD4 | protein_coding | 98,24235278 | -1,490152378 | 0,393074999 | -3,791012869 | 0,000150034 | 0,001709596 |
| ENSG00000171759.10 | PAH | protein_coding | 138,5961692 | -1,493711442 | 0,314146613 | -4,754822694 | 1,99E-06 | 3,47E-05 |

| | | | | | | | | |
|---|---|---|---|---|---|---|---|---|
| ENSG00000205022.9 | PABPN1L | protein_coding | 84,22703288 | -1,494375337 | 0,503845074 | -2,965942137 | 0,003017573 | 0,02181384 |
| ENSG00000215861.6 | AC245297.1 | transcribed_unproc | 1604,307649 | -1,49797565 | 0,210664122 | -7,110729814 | 1,15E-12 | 4,17E-11 |
| ENSG00000121454.6 | LHX4 | protein_coding | 242,2730113 | -1,499747598 | 0,297446782 | -5,042070342 | 4,61E-07 | 8,93E-06 |
| ENSG00000138079.14 | SLC3A1 | protein_coding | 180,8854935 | -1,506030831 | 0,285647436 | -5,2723415 | 1,35E-07 | 2,81E-06 |
| ENSG00000165566.12 | AMER2 | protein_coding | 105,9811342 | -1,506848004 | 0,421588706 | -3,574213407 | 0,000351282 | 0,003580378 |
| ENSG00000110881.12 | ASIC1 | protein_coding | 430,773145 | -1,507324051 | 0,2471437 | -6,098978252 | 1,07E-09 | 2,93E-08 |
| ENSG00000101695.9 | RNF125 | protein_coding | 44,47625734 | -1,509584735 | 0,538801594 | -2,801745115 | 0,005082701 | 0,033327303 |
| ENSG00000275740.1 | AC091959.1 | protein_coding | 162,4261455 | -1,512895527 | 0,360079218 | -4,201563017 | 2,65E-05 | 0,000363896 |
| ENSG00000198435.4 | NRARP | protein_coding | 138,5330012 | -1,513672428 | 0,318609989 | -4,750863068 | 2,03E-06 | 3,53E-05 |
| ENSG00000169302.16 | STK32A | protein_coding | 226,7385053 | -1,527566441 | 0,304790768 | -5,011852727 | 5,39E-07 | 1,03E-05 |
| ENSG00000169717.7 | ACTRT2 | protein_coding | 154,0992782 | -1,531160138 | 0,342264066 | -4,473622242 | 7,69E-06 | 0,000119704 |
| ENSG00000164626.9 | KCNK5 | protein_coding | 126,1724567 | -1,534773765 | 0,331479938 | -4,630065314 | 3,66E-06 | 6,07E-05 |
| ENSG00000164756.12 | SLC30A8 | protein_coding | 19001,73323 | -1,542603056 | 0,149414553 | -10,32431598 | 5,47E-25 | 3,84E-23 |
| ENSG00000075429.9 | CACNG5 | protein_coding | 888,9413834 | -1,54608162 | 0,178277649 | -8,67232448 | 4,23E-18 | 2,11E-16 |
| ENSG00000285982.1 | AC012213.5 | protein_coding | 93,15008861 | -1,547748274 | 0,426924243 | -3,625346413 | 0,000288574 | 0,003019143 |
| ENSG00000136002.20 | ARHGEF4 | protein_coding | 53,57801531 | -1,548736847 | 0,587512528 | -2,636091613 | 0,008386709 | 0,049766367 |
| ENSG00000149527.18 | PLCH2 | protein_coding | 127,8192487 | -1,549301825 | 0,451411509 | -3,432127433 | 0,000598866 | 0,00562209 |
| ENSG00000086967.10 | MYBPC2 | protein_coding | 73,77851762 | -1,554975884 | 0,48196559 | -3,226321374 | 0,001253925 | 0,010597755 |
| ENSG00000162510.6 | MATN1 | protein_coding | 46,48282325 | -1,570817926 | 0,595165338 | -2,639296724 | 0,008307823 | 0,049494671 |
| ENSG00000279294.1 | AC092135.1 | TEC | 76,8171448 | -1,571178667 | 0,443616062 | -3,541753336 | 0,000397477 | 0,003988605 |
| ENSG00000101098.13 | RIMS4 | protein_coding | 69,05645747 | -1,57271443 | 0,440794137 | -3,567911404 | 0,000359838 | 0,003651581 |
| ENSG00000183688.4 | RFLNB | protein_coding | 3695,782258 | -1,573453503 | 0,129149339 | -12,18320988 | 3,82E-34 | 3,55E-32 |
| ENSG00000132259.13 | CNGA4 | protein_coding | 65,85840373 | -1,576109747 | 0,450973154 | -3,494908136 | 0,000474225 | 0,004628391 |
| ENSG00000113763.12 | UNC5A | protein_coding | 1820,593346 | -1,597110001 | 0,181189072 | -8,814604433 | 1,20E-18 | 6,16E-17 |
| ENSG00000140368.13 | PSTPIP1 | protein_coding | 69,99752437 | -1,598984349 | 0,508265193 | -3,145964689 | 0,0016554 | 0,013390957 |
| ENSG00000103742.12 | IGDCC4 | protein_coding | 452,9171295 | -1,613187106 | 0,221568803 | -7,280750186 | 3,32E-13 | 1,25E-11 |
| ENSG00000253457.3 | SMIM18 | protein_coding | 59,25443179 | -1,620393003 | 0,471052326 | -3,439942689 | 0,000581837 | 0,005499217 |
| ENSG00000196562.14 | SULF2 | protein_coding | 1293,910258 | -1,621813188 | 0,150852592 | -10,75097995 | 5,86E-27 | 4,50E-25 |
| ENSG00000164920.9 | OSR2 | protein_coding | 185,7985552 | -1,623592355 | 0,391402145 | -4,148143732 | 3,35E-05 | 0,000448199 |
| ENSG00000121281.13 | ADCY7 | protein_coding | 47,18295383 | -1,625230982 | 0,596655876 | -2,723900071 | 0,006451602 | 0,040441456 |
| ENSG00000118137.10 | APOA1 | protein_coding | 100,5239489 | -1,626144699 | 0,464121442 | -3,503705174 | 0,000458833 | 0,004507588 |
| ENSG00000154736.6 | ADAMTS5 | protein_coding | 560,4599456 | -1,632438946 | 0,191853417 | -8,508782229 | 1,76E-17 | 8,48E-16 |
| ENSG00000118777.12 | ABCG2 | protein_coding | 2731,356067 | -1,634333223 | 0,15405882 | -10,60850147 | 2,72E-26 | 2,03E-24 |
| ENSG00000100490.9 | CDKL1 | protein_coding | 40,54686795 | -1,636531569 | 0,561741388 | -2,913318483 | 0,003576097 | 0,025179746 |
| ENSG00000166265.13 | CYYR1 | protein_coding | 203,1418649 | -1,638884505 | 0,288180631 | -5,687004366 | 1,29E-08 | 3,12E-07 |
| ENSG00000160401.15 | CFAP157 | protein_coding | 63,91457122 | -1,639727103 | 0,458545805 | -3,575928695 | 0,000348987 | 0,003563557 |

| | | | | | | | | |
|---|---|---|---|---|---|---|---|---|
| ENSG00000130707.18 | ASS1 | protein_coding | 505,2884372 | -1,641049569 | 0,241925353 | -6,783288925 | 1,17E-11 | 3,93E-10 |
| ENSG00000167994.13 | RAB3IL1 | protein_coding | 130,6736469 | -1,641669867 | 0,399728592 | -4,106961319 | 4,01E-05 | 0,000525207 |
| ENSG00000169914.6 | OTUD3 | protein_coding | 623,6281859 | -1,6479392 | 0,216295044 | -7,618941098 | 2,56E-14 | 1,05E-12 |
| ENSG00000135678.12 | CPM | protein_coding | 704,4300737 | -1,652816271 | 0,216512308 | -7,633821303 | 2,28E-14 | 9,37E-13 |
| ENSG00000180549.8 | FUT7 | protein_coding | 42,89497749 | -1,662193531 | 0,562629634 | -2,954329866 | 0,003133489 | 0,022566077 |
| ENSG00000284292.1 | AC004922.1 | protein_coding | 69,5164495 | -1,665028317 | 0,510377585 | -3,262346085 | 0,001104942 | 0,009449327 |
| ENSG00000205929.11 | C21orf62 | protein_coding | 238,9510019 | -1,66518084 | 0,289356028 | -5,754781922 | 8,68E-09 | 2,12E-07 |
| ENSG00000100422.14 | CERK | protein_coding | 6185,837081 | -1,674300935 | 0,11594492 | -14,4404855 | 2,88E-47 | 3,72E-45 |
| ENSG00000074211.14 | PPP2R2C | protein_coding | 609,7977074 | -1,684206679 | 0,193654784 | -8,696953653 | 3,41E-18 | 1,70E-16 |
| ENSG00000260855.1 | AL591848.3 | lncRNA | 53,86548437 | -1,688447817 | 0,495444004 | -3,407948833 | 0,000654532 | 0,006052394 |
| ENSG00000080644.16 | CHRNA3 | protein_coding | 364,2853607 | -1,690878929 | 0,352811472 | -4,792584892 | 1,65E-06 | 2,92E-05 |
| ENSG00000120833.14 | SOCS2 | protein_coding | 52,35479255 | -1,702518781 | 0,542459556 | -3,138517447 | 0,001698048 | 0,013667297 |
| ENSG00000170271.11 | FAXDC2 | protein_coding | 234,9480369 | -1,718479929 | 0,314638536 | -5,461759236 | 4,71E-08 | 1,05E-06 |
| ENSG00000258768.2 | AL356019.2 | lncRNA | 44,30806383 | -1,720732042 | 0,567032899 | -3,034624699 | 0,002408352 | 0,018149332 |
| ENSG00000236824.2 | BCYRN1 | scRNA | 2280,67469 | -1,724936277 | 0,183342399 | -9,408278107 | 5,04E-21 | 2,90E-19 |
| ENSG00000106278.12 | PTPRZ1 | protein_coding | 260,7552464 | -1,730679012 | 0,278915176 | -6,205037086 | 5,47E-10 | 1,55E-08 |
| ENSG00000184502.4 | GAST | protein_coding | 447,430715 | -1,734441112 | 0,311231093 | -5,572840096 | 2,51E-08 | 5,80E-07 |
| ENSG00000189144.14 | ZNF573 | protein_coding | 43,034428 | -1,74791067 | 0,663522905 | -2,634288368 | 0,008431385 | 0,049950403 |
| ENSG00000175161.14 | CADM2 | protein_coding | 96,95462778 | -1,750950195 | 0,610086243 | -2,870004388 | 0,004104661 | 0,028285411 |
| ENSG00000224046.2 | AC005076.1 | lncRNA | 188,4726198 | -1,755732341 | 0,36285515 | -4,838659015 | 1,31E-06 | 2,36E-05 |
| ENSG00000134917.10 | ADAMTS8 | protein_coding | 107,0643354 | -1,757308048 | 0,42513443 | -4,133535007 | 3,57E-05 | 0,000474225 |
| ENSG00000198515.16 | CNGA1 | protein_coding | 58,34188465 | -1,757384216 | 0,485828142 | -3,617296044 | 0,000297697 | 0,003099096 |
| ENSG00000168268.11 | NT5DC2 | protein_coding | 17403,24139 | -1,762104139 | 0,13046673 | -13,50615703 | 1,44E-41 | 1,61E-39 |
| ENSG00000267289.1 | AC008752.2 | lncRNA | 29,42409911 | -1,762363777 | 0,648484967 | -2,717663272 | 0,006574471 | 0,041055927 |
| ENSG00000245614.4 | DDX11-AS1 | lncRNA | 142,8697527 | -1,766270112 | 0,373064674 | -4,734487702 | 2,20E-06 | 3,79E-05 |
| ENSG00000169418.10 | NPR1 | protein_coding | 992,671979 | -1,769367434 | 0,169488425 | -10,4394588 | 1,64E-25 | 1,19E-23 |
| ENSG00000180071.20 | ANKRD18A | protein_coding | 88,22651271 | -1,770113682 | 0,431100016 | -4,10603947 | 4,03E-05 | 0,000526976 |
| ENSG00000151468.11 | CCDC3 | protein_coding | 95,66151103 | -1,774063002 | 0,432250932 | -4,104243329 | 4,06E-05 | 0,000530755 |
| ENSG00000239911.2 | PRKAG2-AS1 | lncRNA | 174,7865652 | -1,781774821 | 0,32485405 | -5,484847184 | 4,14E-08 | 9,33E-07 |
| ENSG00000176014.13 | TUBB6 | protein_coding | 770,0270543 | -1,804507699 | 0,219327727 | -8,227449061 | 1,91E-16 | 8,88E-15 |
| ENSG00000107742.13 | SPOCK2 | protein_coding | 822,7273241 | -1,826132405 | 0,21064183 | -8,669372089 | 4,35E-18 | 2,16E-16 |
| ENSG00000136404.16 | TM6SF1 | protein_coding | 58,2075827 | -1,833384713 | 0,532725804 | -3,441516631 | 0,000578463 | 0,005474738 |
| ENSG00000205363.6 | INSYN1 | protein_coding | 537,2741562 | -1,863615724 | 0,26882436 | -6,932465958 | 4,14E-12 | 1,45E-10 |
| ENSG00000145248.7 | SLC10A4 | protein_coding | 27,41189679 | -1,87683307 | 0,677928225 | -2,768483447 | 0,005631785 | 0,036253067 |
| ENSG00000187800.13 | PEAR1 | protein_coding | 53,44426468 | -1,889147589 | 0,529525101 | -3,567626134 | 0,00036023 | 0,003653786 |
| ENSG00000143847.15 | PPFIA4 | protein_coding | 58,46812067 | -1,902603478 | 0,510599813 | -3,726212643 | 0,000194378 | 0,002151937 |

| | | | | | | | | |
|---|---|---|---|---|---|---|---|---|
| ENSG00000168077.14 | SCARA3 | protein_coding | 210,7595878 | -1,905898428 | 0,28249837 | -6,746582042 | 1,51E-11 | 5,00E-10 |
| ENSG00000279968.3 | CCDC28A-AS1 | lncRNA | 34,01951963 | -1,916908455 | 0,621198658 | -3,085821949 | 0,002029903 | 0,015762621 |
| ENSG00000231473.3 | RB1-DT | lncRNA | 34,82936644 | -1,932063792 | 0,73236553 | -2,638114046 | 0,008336854 | 0,049586552 |
| ENSG00000166450.13 | PRTG | protein_coding | 69,1806863 | -1,936347821 | 0,650751747 | -2,975555318 | 0,002924586 | 0,021310897 |
| ENSG00000259030.7 | FPGT-TNNI3K | protein_coding | 41,46875859 | -1,941864304 | 0,681492853 | -2,849427249 | 0,004379802 | 0,029673514 |
| ENSG00000186732.14 | MPPED1 | protein_coding | 65,07771765 | -1,944432489 | 0,486942613 | -3,993145056 | 6,52E-05 | 0,000813796 |
| ENSG00000236617.2 | AC127070.1 | lncRNA | 27,91138147 | -1,94822568 | 0,70341085 | -2,769683862 | 0,005611073 | 0,036155064 |
| ENSG00000198915.12 | RASGEF1A | protein_coding | 183,2891877 | -1,951397857 | 0,30717264 | -6,352772365 | 2,11E-10 | 6,21E-09 |
| ENSG00000185269.12 | NOTUM | protein_coding | 128,6505639 | -1,970453059 | 0,335935183 | -5,865575138 | 4,48E-09 | 1,14E-07 |
| ENSG00000140807.7 | NKD1 | protein_coding | 264,7664686 | -1,973822428 | 0,367073385 | -5,377187535 | 7,57E-08 | 1,65E-06 |
| ENSG00000129167.9 | TPH1 | protein_coding | 88,83435794 | -1,976507763 | 0,417440276 | -4,734827648 | 2,19E-06 | 3,79E-05 |
| ENSG00000265148.6 | TSPOAP1-AS1 | lncRNA | 31,00829192 | -1,998550175 | 0,662770672 | -3,015447513 | 0,002566003 | 0,019137614 |
| ENSG00000196502.12 | SULT1A1 | protein_coding | 63,51569668 | -2,000387088 | 0,500966854 | -3,993052779 | 6,52E-05 | 0,000813796 |
| ENSG00000224086.6 | PPM1F-AS1 | lncRNA | 71,9139242 | -2,013030194 | 0,562693253 | -3,577491259 | 0,000346908 | 0,003544419 |
| ENSG00000243364.8 | EFNA4 | protein_coding | 88,4137615 | -2,024731545 | 0,487266713 | -4,155283932 | 3,25E-05 | 0,00043599 |
| ENSG00000260469.3 | INSYN1-AS1 | lncRNA | 135,2299837 | -2,030407432 | 0,457609391 | -4,436988121 | 9,12E-06 | 0,000140016 |
| ENSG00000115112.8 | TFCP2L1 | protein_coding | 247,4042951 | -2,038289633 | 0,25327383 | -8,047770392 | 8,43E-16 | 3,77E-14 |
| ENSG00000113303.12 | BTNL8 | protein_coding | 36,66964714 | -2,042758292 | 0,606951163 | -3,365605698 | 0,000763758 | 0,006906814 |
| ENSG00000101445.10 | PPP1R16B | protein_coding | 173,794467 | -2,046563516 | 0,302019257 | -6,776268285 | 1,23E-11 | 4,11E-10 |
| ENSG00000196277.16 | GRM7 | protein_coding | 60,26225226 | -2,048724511 | 0,710382048 | -2,883975623 | 0,003926891 | 0,027312117 |
| ENSG00000228055.3 | FAM245A | lncRNA | 86,66038201 | -2,04920204 | 0,50817056 | -4,03250838 | 5,52E-05 | 0,000701878 |
| ENSG00000120332.16 | TNN | protein_coding | 223,6109082 | -2,065918812 | 0,287954001 | -7,174475106 | 7,26E-13 | 2,66E-11 |
| ENSG00000092445.12 | TYRO3 | protein_coding | 276,9301442 | -2,067231792 | 0,243040678 | -8,505702871 | 1,80E-17 | 8,69E-16 |
| ENSG00000226950.8 | DANCR | lncRNA | 1541,270672 | -2,073058676 | 0,152642762 | -13,58111353 | 5,18E-42 | 5,93E-40 |
| ENSG00000167281.19 | RBFOX3 | protein_coding | 73,84663157 | -2,073689903 | 0,542398929 | -3,823182147 | 0,00013174 | 0,00152314 |
| ENSG00000233087.7 | RAB6D | protein_coding | 45,32495291 | -2,084749375 | 0,677343595 | -3,077831384 | 0,002085129 | 0,016107811 |
| ENSG00000111305.19 | GSG1 | protein_coding | 219,0706325 | -2,138113026 | 0,282291364 | -7,574135433 | 3,62E-14 | 1,46E-12 |
| ENSG00000236901.6 | MIR600HG | lncRNA | 365,8274224 | -2,157171174 | 0,276826263 | -7,792509096 | 6,57E-15 | 2,79E-13 |
| ENSG00000066468.23 | FGFR2 | protein_coding | 77,3848715 | -2,158852288 | 0,457402634 | -4,719807295 | 2,36E-06 | 4,04E-05 |
| ENSG00000183454.18 | GRIN2A | protein_coding | 402,1650593 | -2,198193927 | 0,218519923 | -10,05946686 | 8,34E-24 | 5,58E-22 |
| ENSG00000172031.7 | EPHX4 | protein_coding | 26,50920621 | -2,202039167 | 0,700799203 | -3,142182752 | 0,001676933 | 0,013528554 |
| ENSG00000224568.2 | LINC01886 | lncRNA | 27,1156437 | -2,205272793 | 0,736927165 | -2,992524767 | 0,002766802 | 0,020352426 |
| ENSG00000187922.14 | LCN10 | protein_coding | 458,1365886 | -2,250154837 | 0,27570971 | -8,161318799 | 3,31E-16 | 1,51E-14 |
| ENSG00000260296.1 | AC095057.3 | lncRNA | 41,49573728 | -2,256961007 | 0,615895763 | -3,664517834 | 0,000247805 | 0,002657683 |
| ENSG00000091137.14 | SLC26A4 | protein_coding | 157,5436689 | -2,263880397 | 0,379692136 | -5,962410553 | 2,49E-09 | 6,59E-08 |
| ENSG00000150637.9 | CD226 | protein_coding | 55,97898082 | -2,276446913 | 0,665047497 | -3,422983957 | 0,000619377 | 0,005793863 |

| | | | | | | | | |
|---|---|---|---|---|---|---|---|---|
| ENSG00000145692.15 | BHMT | protein_coding | 35,84415017 | -2,282323757 | 0,708595481 | -3,220912097 | 0,001277833 | 0,010747867 |
| ENSG00000152254.11 | G6PC2 | protein_coding | 601,4026233 | -2,28495439 | 0,224008309 | -10,20031086 | 1,98E-24 | 1,36E-22 |
| ENSG00000141668.10 | CBLN2 | protein_coding | 445,2016128 | -2,285332686 | 0,234525823 | -9,744482125 | 1,95E-22 | 1,22E-20 |
| ENSG00000250770.3 | AC005865.1 | transcribed_unproc | 22,67817527 | -2,301084447 | 0,779432551 | -2,952256029 | 0,003154612 | 0,022686984 |
| ENSG00000005513.10 | SOX8 | protein_coding | 41,11557693 | -2,312348968 | 0,591513642 | -3,909206489 | 9,26E-05 | 0,001119271 |
| ENSG00000163630.11 | SYNPR | protein_coding | 38,82655552 | -2,359571461 | 0,824634428 | -2,861354537 | 0,00421835 | 0,028907091 |
| ENSG00000158748.4 | HTR6 | protein_coding | 18,10231291 | -2,385981841 | 0,885970441 | -2,693071609 | 0,007079706 | 0,043541025 |
| ENSG00000188580.15 | NKAIN2 | protein_coding | 397,5187753 | -2,388086329 | 0,227196634 | -10,51109906 | 7,68E-26 | 5,62E-24 |
| ENSG00000214432.9 | VPS33B-DT | lncRNA | 34,26446552 | -2,398673282 | 0,666822791 | -3,597167517 | 0,000321701 | 0,003309492 |
| ENSG00000124772.12 | CPNE5 | protein_coding | 33,28571859 | -2,401372834 | 0,71327022 | -3,366708389 | 0,000760711 | 0,00688223 |
| ENSG00000273598.1 | AC211486.3 | transcribed_unproc | 30,4330479 | -2,430480219 | 0,888701162 | -2,734867831 | 0,006240533 | 0,039410423 |
| ENSG00000124568.12 | SLC17A1 | protein_coding | 45,04758256 | -2,446979254 | 0,608417612 | -4,021874461 | 5,77E-05 | 0,000728614 |
| ENSG00000128045.7 | RASL11B | protein_coding | 60,0367285 | -2,493769144 | 0,497890935 | -5,008665494 | 5,48E-07 | 1,05E-05 |
| ENSG00000187135.7 | VSTM2B | protein_coding | 19,47065927 | -2,599203776 | 0,852976841 | -3,047214942 | 0,002309725 | 0,017538661 |
| ENSG00000241484.10 | ARHGAP8 | protein_coding | 17,93099477 | -2,631583374 | 0,921708636 | -2,855114156 | 0,004302136 | 0,029299031 |
| ENSG00000187783.12 | TMEM72 | protein_coding | 70,96198166 | -2,697448557 | 0,462011048 | -5,838493623 | 5,27E-09 | 1,33E-07 |
| ENSG00000284431.1 | AL022238.3 | protein_coding | 26,39124217 | -2,742599416 | 0,949972457 | -2,887030455 | 0,003888966 | 0,027075324 |
| ENSG00000233327.11 | USP32P2 | transcribed_unproc | 39,15111933 | -2,745417936 | 0,636060562 | -4,316283857 | 1,59E-05 | 0,000228625 |
| ENSG00000197888.2 | UGT2B17 | protein_coding | 46,85607175 | -2,791922255 | 0,660769008 | -4,225262111 | 2,39E-05 | 0,000330014 |
| ENSG00000256001.2 | AC079949.1 | lncRNA | 27,46516854 | -2,842091787 | 0,789717612 | -3,59887097 | 0,000319602 | 0,003291127 |
| ENSG00000204003.8 | AL355987.1 | protein_coding | 45,94945728 | -2,855574594 | 0,786386415 | -3,631261348 | 0,000282039 | 0,002959646 |
| ENSG00000117115.13 | PADI2 | protein_coding | 88,73659484 | -2,90976467 | 0,421441051 | -6,904321888 | 5,04E-12 | 1,74E-10 |
| ENSG00000151388.11 | ADAMTS12 | protein_coding | 21,65910873 | -2,917043921 | 0,951762862 | -3,064885212 | 0,002177536 | 0,016709548 |
| ENSG00000095464.10 | PDE6C | protein_coding | 18,10151931 | -2,926553426 | 0,89583288 | -3,266851987 | 0,001087505 | 0,009315434 |
| ENSG00000263069.6 | RNF213-AS1 | lncRNA | 56,52895182 | -3,002033425 | 0,658712611 | -4,557425158 | 5,18E-06 | 8,33E-05 |
| ENSG00000258441.1 | LINC00641 | lncRNA | 228,794428 | -3,004449175 | 0,282648807 | -10,62961915 | 2,17E-26 | 1,63E-24 |
| ENSG00000107719.9 | PALD1 | protein_coding | 58,6546708 | -3,008574907 | 0,688741077 | -4,368223425 | 1,25E-05 | 0,000186114 |
| ENSG00000197580.13 | BCO2 | protein_coding | 21,95232934 | -3,102598757 | 0,999439649 | -3,104338277 | 0,001907052 | 0,014992161 |
| ENSG00000237560.7 | LINC01497 | lncRNA | 28,22079426 | -3,111454597 | 0,878841949 | -3,540402915 | 0,000399517 | 0,00400569 |
| ENSG00000214491.8 | SEC14L6 | protein_coding | 13,26882042 | -3,131683826 | 1,124490752 | -2,78497962 | 0,005353109 | 0,034868492 |
| ENSG00000264717.5 | NPY4R2 | protein_coding | 34,29905318 | -3,146822257 | 0,899252834 | -3,4993743 | 0,000466351 | 0,004555152 |
| ENSG00000162511.8 | LAPTM5 | protein_coding | 24,73478642 | -3,255897241 | 1,078594429 | -3,018648301 | 0,002539051 | 0,018956859 |
| ENSG00000183117.19 | CSMD1 | protein_coding | 21,40415359 | -3,262270234 | 0,853240754 | -3,823387734 | 0,000131631 | 0,00152314 |
| ENSG00000141052.18 | MYOCD | protein_coding | 28,54640501 | -3,358032612 | 0,834228359 | -4,025315822 | 5,69E-05 | 0,00072018 |
| ENSG00000171962.18 | DRC3 | protein_coding | 22,18602803 | -3,392032809 | 0,975095566 | -3,478667043 | 0,000503914 | 0,004881742 |
| ENSG00000178568.15 | ERBB4 | protein_coding | 23,82352679 | -3,963876144 | 0,906962445 | -4,370496447 | 1,24E-05 | 0,000184711 |

| | | | | | | | | |
|---|---|---|---|---|---|---|---|---|
| ENSG00000267443.2 | AC010641.1 | lncRNA | 12,33568914 | -4,044641283 | 1,279931944 | -3,160044018 | 0,001577453 | 0,012839805 |
| ENSG00000274628.5 | AL669942.1 | transcribed_unproc | 7,311747492 | -4,435071096 | 1,643597455 | -2,69839253 | 0,006967523 | 0,043015363 |
| ENSG00000188916.9 | INSYN2A | protein_coding | 8,855037945 | -4,591251188 | 1,593030327 | -2,882086493 | 0,003950513 | 0,027403573 |
| ENSG00000215032.2 | GNL3LP1 | processed_pseudog | 9,56434676 | -4,732635064 | 1,559343917 | -3,035016851 | 0,002405222 | 0,018132277 |
| ENSG00000164100.9 | NDST3 | protein_coding | 7,091333447 | -5,848431025 | 1,853295553 | -3,155692581 | 0,001601175 | 0,013012657 |
| ENSG00000205236.6 | AC105052.1 | protein_coding | 16,19166245 | -6,645002534 | 1,536981424 | -4,32341109 | 1,54E-05 | 0,000222627 |
| ENSG00000112486.17 | CCR6 | protein_coding | 13,42923038 | -6,65812314 | 1,584791476 | -4,201261328 | 2,65E-05 | 0,000364143 |
| ENSG00000261069.3 | AC124312.3 | lncRNA | 36,66629918 | -8,404585346 | 1,33914567 | -6,27607999 | 3,47E-10 | 9,98E-09 |

**DEG_significative_IL-1β vs NT 24h**

| **Ensembl_gene_ID** | **gene_name** | **gene_type** | **baseMean** | **log2FoldChange** | **lfcSE** | **stat** | **pvalue** | **padj** |
|---|---|---|---|---|---|---|---|---|
| ENSG00000256500.6 | AL139300.1 | protein_coding | 293,8131854 | 8,485181994 | 2,189331368 | 3,875695621 | 0,000106321 | 0,004639937 |
| ENSG00000276077.4 | CU633904.1 | lncRNA | 80,17648461 | 7,799337898 | 2,538507295 | 3,072411064 | 0,002123371 | 0,044647493 |
| ENSG00000277117.5 | FP565260.3 | protein_coding | 115,6928961 | 6,39857683 | 1,43172954 | 4,469123986 | 7,85E-06 | 0,000579263 |
| ENSG00000243649.9 | CFB | protein_coding | 7,888591343 | 6,205792686 | 1,308322736 | 4,743319453 | 2,10E-06 | 0,000183144 |
| ENSG00000166396.13 | SERPINB7 | protein_coding | 5,289295335 | 5,789467079 | 1,454920245 | 3,979233295 | 6,91E-05 | 0,003273642 |
| ENSG00000105246.6 | EBI3 | protein_coding | 54,79756385 | 5,396762835 | 1,083031197 | 4,983016972 | 6,26E-07 | 6,41E-05 |
| ENSG00000248498.4 | ASNSP1 | transcribed_unproc | 8,762297118 | 5,369329723 | 1,284401562 | 4,180413572 | 2,91E-05 | 0,001646601 |
| ENSG00000227507.3 | LTB | protein_coding | 815,8840032 | 5,244833633 | 0,813928562 | 6,44385008 | 1,16E-10 | 3,89E-08 |
| ENSG00000184845.4 | DRD1 | protein_coding | 9,152543178 | 5,173430055 | 1,207785728 | 4,283400553 | 1,84E-05 | 0,00114075 |
| ENSG00000286239.1 | AC093884.1 | protein_coding | 5,843145653 | 5,049487887 | 1,388343778 | 3,637058752 | 0,000275769 | 0,00974707 |
| ENSG00000242220.9 | TCP10L | protein_coding | 4,865771693 | 5,01049192 | 1,598291132 | 3,134905663 | 0,001719094 | 0,038459728 |
| ENSG00000008517.18 | IL32 | protein_coding | 90,67348884 | 4,957962826 | 0,99212902 | 4,997296446 | 5,81E-07 | 6,04E-05 |
| ENSG00000081041.9 | CXCL2 | protein_coding | 193,635298 | 4,610940466 | 0,73091254 | 6,308470873 | 2,82E-10 | 8,55E-08 |
| ENSG00000253616.5 | AC107959.3 | lncRNA | 8,081674917 | 4,442211828 | 1,193701686 | 3,721375182 | 0,000198141 | 0,007584885 |
| ENSG00000161031.13 | PGLYRP2 | protein_coding | 11,80198839 | 4,381654669 | 1,126962227 | 3,888022655 | 0,000101064 | 0,004454475 |
| ENSG00000167434.10 | CA4 | protein_coding | 1367,295778 | 4,311652009 | 0,753733211 | 5,720395421 | 1,06E-08 | 1,95E-06 |
| ENSG00000049249.9 | TNFRSF9 | protein_coding | 32,900097 | 4,2283496 | 0,700133575 | 6,039346989 | 1,55E-09 | 3,67E-07 |
| ENSG00000255595.5 | AC007368.1 | lncRNA | 45,9058022 | 4,218907643 | 0,638577494 | 6,606727739 | 3,93E-11 | 1,55E-08 |
| ENSG00000139269.3 | INHBE | protein_coding | 150,0228191 | 4,069507716 | 0,745013766 | 5,462325533 | 4,70E-08 | 6,62E-06 |
| ENSG00000090339.9 | ICAM1 | protein_coding | 150,6174767 | 3,990399857 | 0,736402749 | 5,418773708 | 6,00E-08 | 8,15E-06 |
| ENSG00000023445.16 | BIRC3 | protein_coding | 789,0628887 | 3,918518447 | 0,723030028 | 5,419579128 | 5,97E-08 | 8,15E-06 |
| ENSG00000186891.14 | TNFRSF18 | protein_coding | 33,36396352 | 3,89932906 | 0,716723022 | 5,440496458 | 5,31E-08 | 7,39E-06 |
| ENSG00000249494.6 | AC008629.1 | lncRNA | 8,41009991 | 3,895904199 | 1,234866036 | 3,154920521 | 0,001605418 | 0,036544739 |
| ENSG00000232810.4 | TNF | protein_coding | 50,6661187 | 3,623510293 | 0,600119698 | 6,037979268 | 1,56E-09 | 3,67E-07 |
| ENSG00000189325.7 | BNIP5 | protein_coding | 10,52289949 | 3,622011687 | 1,134516301 | 3,192560287 | 0,001410175 | 0,033370624 |

| | | | | | | | | |
|---|---|---|---|---|---|---|---|---|
| ENSG00000163734.4 | CXCL3 | protein_coding | 105,60569 | 3,589942979 | 0,820246055 | 4,376665951 | 1,21E-05 | 0,00080555 |
| ENSG00000140044.13 | JDP2 | protein_coding | 35,5492663 | 3,53604724 | 0,490453159 | 7,20975525 | 5,61E-13 | 3,91E-10 |
| ENSG00000253522.6 | MIR3142HG | lncRNA | 20,96268031 | 3,496711547 | 0,852585349 | 4,101303818 | 4,11E-05 | 0,00216945 |
| ENSG00000137801.11 | THBS1 | protein_coding | 19,38899779 | 3,321520182 | 0,711664344 | 4,667256706 | 3,05E-06 | 0,000257783 |
| ENSG00000163121.10 | NEURL3 | protein_coding | 261,246785 | 3,247190797 | 0,566798357 | 5,729005316 | 1,01E-08 | 1,88E-06 |
| ENSG00000170075.9 | GPR37L1 | protein_coding | 18,27289128 | 3,241712565 | 0,772788658 | 4,194824199 | 2,73E-05 | 0,001567343 |
| ENSG00000160223.17 | ICOSLG | protein_coding | 281,4117934 | 3,240807567 | 0,581537565 | 5,572825838 | 2,51E-08 | 3,93E-06 |
| ENSG00000134339.9 | SAA2 | protein_coding | 14,8682705 | 3,200823258 | 0,775576416 | 4,127024997 | 3,67E-05 | 0,001983172 |
| ENSG00000230183.4 | CNOT6LP1 | processed_pseudog | 22,41848049 | 3,188964148 | 0,775104111 | 4,114239756 | 3,88E-05 | 0,002071087 |
| ENSG00000169884.14 | WNT10B | protein_coding | 59,35410793 | 3,078323481 | 0,488646267 | 6,29969712 | 2,98E-10 | 8,92E-08 |
| ENSG00000135925.9 | WNT10A | protein_coding | 195,802194 | 3,014198304 | 0,534724703 | 5,636916126 | 1,73E-08 | 2,86E-06 |
| ENSG00000148344.11 | PTGES | protein_coding | 49,58623637 | 3,010023861 | 0,479459265 | 6,277955362 | 3,43E-10 | 1,01E-07 |
| ENSG00000158014.15 | SLC30A2 | protein_coding | 30,56527793 | 2,983447467 | 0,538102028 | 5,544389932 | 2,95E-08 | 4,50E-06 |
| ENSG00000163739.5 | CXCL1 | protein_coding | 1075,615019 | 2,927901497 | 0,529208112 | 5,532608875 | 3,16E-08 | 4,75E-06 |
| ENSG00000272341.1 | AL137003.1 | lncRNA | 17,85347583 | 2,808266905 | 0,791230787 | 3,54923867 | 0,000386347 | 0,012552581 |
| ENSG00000261618.2 | LINC02605 | lncRNA | 408,8876075 | 2,756221813 | 0,591846602 | 4,656986797 | 3,21E-06 | 0,000266902 |
| ENSG00000112303.14 | VNN2 | protein_coding | 42,14199637 | 2,754473127 | 0,566793155 | 4,859750162 | 1,18E-06 | 0,000110193 |
| ENSG00000162772.17 | ATF3 | protein_coding | 651,5296305 | 2,709774817 | 0,485318212 | 5,583501198 | 2,36E-08 | 3,75E-06 |
| ENSG00000163735.7 | CXCL5 | protein_coding | 102,6842024 | 2,681004929 | 0,388843855 | 6,894811096 | 5,39E-12 | 2,71E-09 |
| ENSG00000198075.10 | SULT1C4 | protein_coding | 29,53130487 | 2,65741855 | 0,509405964 | 5,216700901 | 1,82E-07 | 2,14E-05 |
| ENSG00000285744.1 | AC083837.1 | lncRNA | 1002,357633 | 2,605320307 | 0,491854131 | 5,296936925 | 1,18E-07 | 1,47E-05 |
| ENSG00000129467.14 | ADCY4 | protein_coding | 13,15085085 | 2,580504606 | 0,739842754 | 3,487909548 | 0,000486813 | 0,014939277 |
| ENSG00000174502.19 | SLC26A9 | protein_coding | 113,7503997 | 2,499863061 | 0,699686652 | 3,572832286 | 0,000353141 | 0,011679517 |
| ENSG00000105327.18 | BBC3 | protein_coding | 47,69752786 | 2,495959046 | 0,528994392 | 4,718309083 | 2,38E-06 | 0,000205541 |
| ENSG00000163131.11 | CTSS | protein_coding | 243,7951307 | 2,484610692 | 0,482651213 | 5,147838911 | 2,64E-07 | 2,97E-05 |
| ENSG00000154930.15 | ACSS1 | protein_coding | 18,8030819 | 2,469483604 | 0,661991707 | 3,730384503 | 0,000191188 | 0,007382584 |
| ENSG00000164236.12 | ANKRD33B | protein_coding | 34,23532644 | 2,423337826 | 0,469593845 | 5,160497424 | 2,46E-07 | 2,79E-05 |
| ENSG00000169429.11 | CXCL8 | protein_coding | 83,64815766 | 2,41799249 | 0,438136454 | 5,518811471 | 3,41E-08 | 5,10E-06 |
| ENSG00000093134.15 | VNN3 | protein_coding | 28,74387005 | 2,411246087 | 0,619203676 | 3,894108158 | 9,86E-05 | 0,004352801 |
| ENSG00000255274.9 | SMIM35 | protein_coding | 167,2016898 | 2,402781982 | 0,338008643 | 7,108640649 | 1,17E-12 | 7,41E-10 |
| ENSG00000085514.16 | PILRA | protein_coding | 59,85053157 | 2,39577083 | 0,460453897 | 5,203063428 | 1,96E-07 | 2,27E-05 |
| ENSG00000125398.8 | SOX9 | protein_coding | 22,98400058 | 2,3954527 | 0,656724811 | 3,647574542 | 0,000264728 | 0,009416979 |
| ENSG00000087085.16 | ACHE | protein_coding | 15,19817245 | 2,391812921 | 0,712400654 | 3,357398549 | 0,000786796 | 0,02154536 |
| ENSG00000112299.8 | VNN1 | protein_coding | 92,68929179 | 2,3819466 | 0,407434481 | 5,846207696 | 5,03E-09 | 1,01E-06 |
| ENSG00000124813.23 | RUNX2 | protein_coding | 17,21265726 | 2,359482662 | 0,751908783 | 3,137990559 | 0,001701104 | 0,038250627 |
| ENSG00000163874.11 | ZC3H12A | protein_coding | 298,6281359 | 2,353659118 | 0,435267413 | 5,407386464 | 6,40E-08 | 8,58E-06 |

| | | | | | | | | |
|---|---|---|---|---|---|---|---|---|
| ENSG00000266903.2 | AC243964.3 | lncRNA | 18,09142705 | 2,339492755 | 0,669907009 | 3,492264935 | 0,000478943 | 0,014779765 |
| ENSG00000124875.10 | CXCL6 | protein_coding | 479,5546622 | 2,278183575 | 0,353637548 | 6,442142767 | 1,18E-10 | 3,89E-08 |
| ENSG00000169715.15 | MT1E | protein_coding | 68,81321717 | 2,233050376 | 0,366322755 | 6,095854935 | 1,09E-09 | 2,74E-07 |
| ENSG00000163501.7 | IHH | protein_coding | 24,28587062 | 2,20094354 | 0,556579724 | 3,954408409 | 7,67E-05 | 0,003573897 |
| ENSG00000064489.23 | BORCS8-MEF2B | protein_coding | 99,39819859 | 2,192027894 | 0,365587229 | 5,995909378 | 2,02E-09 | 4,53E-07 |
| ENSG00000188505.5 | NCCRP1 | protein_coding | 17,8396743 | 2,175151993 | 0,627120444 | 3,468475654 | 0,00052342 | 0,015862633 |
| ENSG00000115353.11 | TACR1 | protein_coding | 41,53338871 | 2,120880387 | 0,45524172 | 4,658800574 | 3,18E-06 | 0,000266902 |
| ENSG00000152785.8 | BMP3 | protein_coding | 123,7026178 | 2,097763906 | 0,300742912 | 6,97527298 | 3,05E-12 | 1,65E-09 |
| ENSG00000107968.10 | MAP3K8 | protein_coding | 195,2911985 | 2,087306294 | 0,245757335 | 8,493363143 | 2,01E-17 | 4,04E-14 |
| ENSG00000157570.12 | TSPAN18 | protein_coding | 101,2963459 | 2,03425155 | 0,338539948 | 6,008896627 | 1,87E-09 | 4,31E-07 |
| ENSG00000105499.14 | PLA2G4C | protein_coding | 373,8061533 | 1,980703648 | 0,405805123 | 4,880923227 | 1,06E-06 | 0,000100702 |
| ENSG00000120889.13 | TNFRSF10B | protein_coding | 32,98580414 | 1,977310621 | 0,571742593 | 3,45839307 | 0,000543408 | 0,016247897 |
| ENSG00000128965.13 | CHAC1 | protein_coding | 555,9735969 | 1,973269021 | 0,349875151 | 5,639923317 | 1,70E-08 | 2,83E-06 |
| ENSG00000263528.8 | IKBKE | protein_coding | 879,3750479 | 1,941042567 | 0,390186431 | 4,974654204 | 6,54E-07 | 6,66E-05 |
| ENSG00000104856.14 | RELB | protein_coding | 1169,213489 | 1,936784103 | 0,393397973 | 4,923218313 | 8,51E-07 | 8,41E-05 |
| ENSG00000254768.6 | AC104009.1 | lncRNA | 17,23543224 | 1,891227427 | 0,60616603 | 3,119982534 | 0,001808618 | 0,040022787 |
| ENSG00000100097.12 | LGALS1 | protein_coding | 67,94820907 | 1,866452415 | 0,373163383 | 5,001703004 | 5,68E-07 | 5,93E-05 |
| ENSG00000177301.16 | KCNA2 | protein_coding | 780,5186667 | 1,857303494 | 0,157944444 | 11,75922016 | 6,33E-32 | 7,00E-28 |
| ENSG00000112562.19 | SMOC2 | protein_coding | 19,20222217 | 1,855003101 | 0,59536035 | 3,115765271 | 0,001834682 | 0,040484194 |
| ENSG00000187621.15 | TCL6 | lncRNA | 26,65660385 | 1,850886815 | 0,536488536 | 3,450002543 | 0,000560581 | 0,016604312 |
| ENSG00000111371.16 | SLC38A1 | protein_coding | 41,87512509 | 1,827083564 | 0,434815797 | 4,201971447 | 2,65E-05 | 0,001540671 |
| ENSG00000285735.2 | LINC02717 | lncRNA | 106,5539667 | 1,825568448 | 0,345221438 | 5,288108576 | 1,24E-07 | 1,53E-05 |
| ENSG00000205436.7 | EXOC3L4 | protein_coding | 152,9393886 | 1,767560134 | 0,269603581 | 6,556144879 | 5,52E-11 | 2,07E-08 |
| ENSG00000235568.7 | NFAM1 | protein_coding | 231,749894 | 1,668988923 | 0,258104312 | 6,466334913 | 1,00E-10 | 3,42E-08 |
| ENSG00000119283.15 | TRIM67 | protein_coding | 123,2940947 | 1,660764608 | 0,286127279 | 5,804286173 | 6,46E-09 | 1,25E-06 |
| ENSG00000056487.16 | PHF21B | protein_coding | 46,51295802 | 1,657379499 | 0,482197678 | 3,437137037 | 0,000587898 | 0,017228918 |
| ENSG00000224511.2 | LINC00365 | lncRNA | 34,42543837 | 1,648969359 | 0,457877412 | 3,601333711 | 0,000316589 | 0,010694421 |
| ENSG00000185101.13 | ANO9 | protein_coding | 899,547857 | 1,594745555 | 0,145598175 | 10,95306008 | 6,42E-28 | 4,74E-24 |
| ENSG00000206341.7 | HLA-H | unprocessed_pseud | 57,89961542 | 1,561403972 | 0,360794645 | 4,327680561 | 1,51E-05 | 0,000972046 |
| ENSG00000250166.3 | AC053513.1 | lncRNA | 305,6414239 | 1,555560481 | 0,217286281 | 7,159036795 | 8,12E-13 | 5,45E-10 |
| ENSG00000129596.5 | CDO1 | protein_coding | 175,7843199 | 1,552909554 | 0,295310495 | 5,258565405 | 1,45E-07 | 1,77E-05 |
| ENSG00000151012.13 | SLC7A11 | protein_coding | 336,0258969 | 1,53125211 | 0,235250448 | 6,509029505 | 7,56E-11 | 2,70E-08 |
| ENSG00000152910.19 | CNTNAP4 | protein_coding | 120,1500242 | 1,491077331 | 0,340018689 | 4,385280512 | 1,16E-05 | 0,000783791 |
| ENSG00000115641.19 | FHL2 | protein_coding | 40,72478028 | 1,480421419 | 0,478745769 | 3,09229139 | 0,001986178 | 0,043126761 |
| ENSG00000077150.20 | NFKB2 | protein_coding | 2327,724578 | 1,478928719 | 0,25189855 | 5,871128342 | 4,33E-09 | 8,95E-07 |
| ENSG00000111981.5 | ULBP1 | protein_coding | 71,22090005 | 1,474745212 | 0,391918326 | 3,762889138 | 0,000167962 | 0,006652519 |

| | | | | | | | | | |
|---|---|---|---|---|---|---|---|---|---|
| ENSG00000205038.12 | PKHD1L1 | protein_coding | 74,27620397 | | 1,463597891 | 0,361810803 | 4,045202298 | 5,23E-05 | 0,002665212 |
| ENSG00000124788.19 | ATXN1 | protein_coding | 217,2934278 | | 1,452494379 | 0,255565565 | 5,68345105 | 1,32E-08 | 2,30E-06 |
| ENSG00000198535.5 | C2CD4A | protein_coding | 4192,238827 | | 1,438872749 | 0,220653483 | 6,520960975 | 6,99E-11 | 2,53E-08 |
| ENSG00000287064.1 | AL606500.1 | lncRNA | 53,69379977 | | 1,433205149 | 0,35519062 | 4,03503096 | 5,46E-05 | 0,002745389 |
| ENSG00000287790.1 | AC025756.1 | lncRNA | 62,9436757 | | 1,416308769 | 0,344836074 | 4,107194331 | 4,00E-05 | 0,002125021 |
| ENSG00000228409.6 | CCT6P1 | transcribed_unproc | 112,5855676 | | 1,414905055 | 0,270733999 | 5,226181634 | 1,73E-07 | 2,07E-05 |
| ENSG00000069399.15 | BCL3 | protein_coding | 309,748653 | | 1,409032354 | 0,190536029 | 7,395096677 | 1,41E-13 | 1,25E-10 |
| ENSG00000175874.10 | CREG2 | protein_coding | 58,11784771 | | 1,408319573 | 0,385502466 | 3,65320509 | 0,000258987 | 0,009257433 |
| ENSG00000074416.14 | MGLL | protein_coding | 199,5656467 | | 1,381150706 | 0,237629956 | 5,812191071 | 6,17E-09 | 1,21E-06 |
| ENSG00000144550.13 | CPNE9 | protein_coding | 89,50200475 | | 1,380494631 | 0,387306266 | 3,564348816 | 0,000364761 | 0,012027859 |
| ENSG00000186204.15 | CYP4F12 | protein_coding | 298,0663042 | | 1,350262349 | 0,225063076 | 5,999484106 | 1,98E-09 | 4,52E-07 |
| ENSG00000135604.10 | STX11 | protein_coding | 226,5594019 | | 1,349339473 | 0,225227734 | 5,991000526 | 2,09E-09 | 4,59E-07 |
| ENSG00000119125.17 | GDA | protein_coding | 42,13562963 | | 1,349243961 | 0,393429429 | 3,429443409 | 0,000604821 | 0,017585098 |
| ENSG00000123095.6 | BHLHE41 | protein_coding | 394,6312115 | | 1,331122463 | 0,179591268 | 7,411955363 | 1,24E-13 | 1,15E-10 |
| ENSG00000090104.12 | RGS1 | protein_coding | 101,9210208 | | 1,328371158 | 0,277293909 | 4,790480843 | 1,66E-06 | 0,000148442 |
| ENSG00000134070.5 | IRAK2 | protein_coding | 476,2974673 | | 1,322426281 | 0,182888254 | 7,230788487 | 4,80E-13 | 3,54E-10 |
| ENSG00000134569.10 | LRP4 | protein_coding | 10694,13585 | | 1,31821645 | 0,098966672 | 13,31980175 | 1,78E-40 | 3,93E-36 |
| ENSG00000126368.6 | NR1D1 | protein_coding | 1168,326061 | | 1,309693851 | 0,149048036 | 8,78705878 | 1,54E-18 | 3,77E-15 |
| ENSG00000135373.13 | EHF | protein_coding | 941,0760311 | | 1,308040655 | 0,283330592 | 4,616658738 | 3,90E-06 | 0,000318392 |
| ENSG00000115380.20 | EFEMP1 | protein_coding | 326,8333313 | | 1,306262768 | 0,207067563 | 6,308389156 | 2,82E-10 | 8,55E-08 |
| ENSG00000118503.15 | TNFAIP3 | protein_coding | 1827,680408 | | 1,29943094 | 0,336817852 | 3,857963378 | 0,000114336 | 0,004931371 |
| ENSG00000103257.9 | SLC7A5 | protein_coding | 597,4298505 | | 1,291382732 | 0,163881465 | 7,879980398 | 3,27E-15 | 4,02E-12 |
| ENSG00000131981.16 | LGALS3 | protein_coding | 81,70256613 | | 1,243062762 | 0,328156776 | 3,788014917 | 0,000151856 | 0,006165064 |
| ENSG00000185022.12 | MAFF | protein_coding | 253,5402365 | | 1,233492585 | 0,203042782 | 6,075037852 | 1,24E-09 | 3,01E-07 |
| ENSG00000154263.17 | ABCA10 | protein_coding | 175,648333 | | 1,223602238 | 0,278377706 | 4,395474969 | 1,11E-05 | 0,000755615 |
| ENSG00000234585.6 | CCT6P3 | transcribed_unproc | 238,7365256 | | 1,219290469 | 0,186517172 | 6,537148584 | 6,27E-11 | 2,31E-08 |
| ENSG00000085465.13 | OVGP1 | protein_coding | 86,67277712 | | 1,213136533 | 0,276011122 | 4,395245098 | 1,11E-05 | 0,000755615 |
| ENSG00000006062.17 | MAP3K14 | protein_coding | 196,7966334 | | 1,203850682 | 0,219859955 | 5,475534114 | 4,36E-08 | 6,27E-06 |
| ENSG00000266312.2 | LINC01927 | lncRNA | 64,41485949 | | 1,192868108 | 0,32443239 | 3,676784888 | 0,000236192 | 0,008623738 |
| ENSG00000101096.20 | NFATC2 | protein_coding | 354,1495891 | | 1,178596804 | 0,193270078 | 6,09818559 | 1,07E-09 | 2,73E-07 |
| ENSG00000257743.8 | MGAM2 | protein_coding | 56,77650284 | | 1,161111133 | 0,36098596 | 3,216499422 | 0,001297648 | 0,031447709 |
| ENSG00000120659.15 | TNFSF11 | protein_coding | 76,71219902 | | 1,152762081 | 0,321725513 | 3,583060824 | 0,000339591 | 0,011344115 |
| ENSG00000125148.7 | MT2A | protein_coding | 451,2426254 | | 1,135035479 | 0,14125504 | 8,035362701 | 9,33E-16 | 1,47E-12 |
| ENSG00000149557.14 | FEZ1 | protein_coding | 158,9146624 | | 1,129156327 | 0,269348939 | 4,192169201 | 2,76E-05 | 0,001579692 |
| ENSG00000168209.5 | DDIT4 | protein_coding | 1203,164553 | | 1,127638335 | 0,150175523 | 7,508802451 | 5,97E-14 | 6,00E-11 |
| ENSG00000213599.10 | SLX1A-SULT1A3 | lncRNA | 104,1812677 | | 1,110473469 | 0,279032016 | 3,979734955 | 6,90E-05 | 0,003273642 |

| | | | | | | | | |
|---|---|---|---|---|---|---|---|---|
| ENSG00000204622.11 | HLA-J | transcribed_unproc | 65,67670378 | 1,10430611 | 0,323390303 | 3,414778055 | 0,00063834 | 0,018247948 |
| ENSG00000136842.14 | TMOD1 | protein_coding | 9037,422284 | 1,101541328 | 0,226125366 | 4,871374435 | 1,11E-06 | 0,000104791 |
| ENSG00000182732.18 | RGS6 | protein_coding | 1347,863592 | 1,100824034 | 0,137568171 | 8,00202567 | 1,22E-15 | 1,81E-12 |
| ENSG00000100628.12 | ASB2 | protein_coding | 89,53113008 | 1,099038966 | 0,348398501 | 3,154545625 | 0,001607482 | 0,036554117 |
| ENSG00000162407.9 | PLPP3 | protein_coding | 1051,828152 | 1,098697915 | 0,14940617 | 7,353765339 | 1,93E-13 | 1,58E-10 |
| ENSG00000121753.12 | ADGRB2 | protein_coding | 2873,266178 | 1,096560177 | 0,116999625 | 9,372339278 | 7,09E-21 | 3,92E-17 |
| ENSG00000019582.15 | CD74 | protein_coding | 11559,90185 | 1,088580029 | 0,177726042 | 6,125045121 | 9,07E-10 | 2,36E-07 |
| ENSG00000180914.10 | OXTR | protein_coding | 343,4188612 | 1,086847233 | 0,174727058 | 6,220257157 | 4,96E-10 | 1,41E-07 |
| ENSG00000205502.4 | C2CD4B | protein_coding | 395,0711374 | 1,079633379 | 0,14546736 | 7,421825624 | 1,16E-13 | 1,11E-10 |
| ENSG00000165238.16 | WNK2 | protein_coding | 116,4529016 | 1,076337816 | 0,234376925 | 4,592336962 | 4,38E-06 | 0,000352349 |
| ENSG00000058085.15 | LAMC2 | protein_coding | 1165,962369 | 1,069397427 | 0,147030265 | 7,273314946 | 3,51E-13 | 2,68E-10 |
| ENSG00000180767.10 | CHST13 | protein_coding | 279,4490281 | 1,063448239 | 0,174078092 | 6,109029722 | 1,00E-09 | 2,58E-07 |
| ENSG00000100906.11 | NFKBIA | protein_coding | 4426,475797 | 1,052768514 | 0,22996096 | 4,578031478 | 4,69E-06 | 0,000370905 |
| ENSG00000078295.17 | ADCY2 | protein_coding | 812,1263983 | 1,051400728 | 0,158040397 | 6,652734015 | 2,88E-11 | 1,20E-08 |
| ENSG00000099251.14 | HSD17B7P2 | transcribed_unproc | 87,31005427 | 1,035096341 | 0,274540159 | 3,770291185 | 0,000163057 | 0,006524057 |
| ENSG00000144802.11 | NFKBIZ | protein_coding | 1445,054775 | 1,025477088 | 0,131098885 | 7,822164808 | 5,19E-15 | 5,74E-12 |
| ENSG00000115604.11 | IL18R1 | protein_coding | 1006,520971 | 1,020286566 | 0,145301986 | 7,021834973 | 2,19E-12 | 1,29E-09 |
| ENSG00000172020.13 | GAP43 | protein_coding | 373,3598453 | 1,017724898 | 0,181632179 | 5,603219126 | 2,10E-08 | 3,42E-06 |
| ENSG00000187608.10 | ISG15 | protein_coding | 61,50215169 | 1,014500097 | 0,328371373 | 3,089490072 | 0,002005004 | 0,043365318 |
| ENSG00000125347.14 | IRF1 | protein_coding | 1023,754203 | 0,990281391 | 0,12528418 | 7,904281246 | 2,69E-15 | 3,51E-12 |
| ENSG00000156920.11 | ADGRG4 | protein_coding | 198,8274777 | 0,989419397 | 0,189315303 | 5,22630437 | 1,73E-07 | 2,07E-05 |
| ENSG00000133624.13 | ZNF767P | transcribed_unproc | 248,1982169 | 0,989080438 | 0,184514128 | 5,360459101 | 8,30E-08 | 1,10E-05 |
| ENSG00000288632.1 | AC133555.6 | lncRNA | 109,8398854 | 0,985813878 | 0,275151371 | 3,582805625 | 0,000339924 | 0,011344115 |
| ENSG00000113739.10 | STC2 | protein_coding | 1028,100168 | 0,976638533 | 0,135486317 | 7,208392359 | 5,66E-13 | 3,91E-10 |
| ENSG00000007062.12 | PROM1 | protein_coding | 432,1033199 | 0,971942591 | 0,171851058 | 5,655726547 | 1,55E-08 | 2,66E-06 |
| ENSG00000155265.11 | GOLGA7B | protein_coding | 1023,271807 | 0,953288492 | 0,109487676 | 8,706810947 | 3,13E-18 | 6,92E-15 |
| ENSG00000006210.7 | CX3CL1 | protein_coding | 2332,545256 | 0,927542544 | 0,09924541 | 9,345949017 | 9,11E-21 | 4,03E-17 |
| ENSG00000020577.14 | SAMD4A | protein_coding | 256,9746062 | 0,912293069 | 0,218496656 | 4,175318222 | 2,98E-05 | 0,001679599 |
| ENSG00000112182.15 | BACH2 | protein_coding | 885,6448295 | 0,912082978 | 0,166122393 | 5,490427638 | 4,01E-08 | 5,84E-06 |
| ENSG00000114270.17 | COL7A1 | protein_coding | 782,3311611 | 0,89336329 | 0,154020031 | 5,80030587 | 6,62E-09 | 1,27E-06 |
| ENSG00000254087.8 | LYN | protein_coding | 401,9338526 | 0,893170396 | 0,205652876 | 4,343097039 | 1,40E-05 | 0,000919656 |
| ENSG00000168497.5 | CAVIN2 | protein_coding | 120,4701048 | 0,892723028 | 0,223480393 | 3,99463692 | 6,48E-05 | 0,003124281 |
| ENSG00000154065.17 | ANKRD29 | protein_coding | 369,2161216 | 0,885434879 | 0,170730088 | 5,186167752 | 2,15E-07 | 2,46E-05 |
| ENSG00000100068.13 | LRP5L | protein_coding | 172,0518912 | 0,875217964 | 0,210555366 | 4,156711753 | 3,23E-05 | 0,00179488 |
| ENSG00000272752.6 | STAG3L5P-PVRIG | lncRNA | 297,5764026 | 0,870409195 | 0,244593034 | 3,558601728 | 0,000372834 | 0,012257554 |
| ENSG00000229644.6 | NAMPTP1 | processed_pseudog | 139,9390359 | 0,868831373 | 0,231877435 | 3,746942319 | 0,000179003 | 0,007034859 |

| | | | | | | | | |
|---|---|---|---|---|---|---|---|---|
| ENSG00000172216.6 | CEBPB | protein_coding | 735,147813 | 0,858688784 | 0,128185567 | 6,698794585 | 2,10E-11 | 9,45E-09 |
| ENSG00000267365.1 | KCNJ2-AS1 | lncRNA | 291,1025413 | 0,854198334 | 0,197012671 | 4,335753277 | 1,45E-05 | 0,000948101 |
| ENSG00000168398.7 | BDKRB2 | protein_coding | 107,2693063 | 0,831226987 | 0,234561086 | 3,543754851 | 0,000394472 | 0,012782985 |
| ENSG00000168394.11 | TAP1 | protein_coding | 877,9292292 | 0,824490133 | 0,148071142 | 5,568202713 | 2,57E-08 | 3,98E-06 |
| ENSG00000163462.18 | TRIM46 | protein_coding | 337,3749951 | 0,824016812 | 0,193608148 | 4,256106072 | 2,08E-05 | 0,001260988 |
| ENSG00000169621.10 | APLF | protein_coding | 225,8101588 | 0,823917741 | 0,171961589 | 4,79128941 | 1,66E-06 | 0,000148442 |
| ENSG00000148468.17 | FAM171A1 | protein_coding | 133,204566 | 0,818514806 | 0,213850139 | 3,827515899 | 0,000129443 | 0,005393703 |
| ENSG00000126458.4 | RRAS | protein_coding | 284,3208874 | 0,81795455 | 0,170662742 | 4,792812668 | 1,64E-06 | 0,00014792 |
| ENSG00000198417.7 | MT1F | protein_coding | 788,1790646 | 0,817477996 | 0,112099135 | 7,292455908 | 3,04E-13 | 2,41E-10 |
| ENSG00000139173.10 | TMEM117 | protein_coding | 619,0516351 | 0,803901558 | 0,121722917 | 6,604356683 | 3,99E-11 | 1,55E-08 |
| ENSG00000197989.14 | SNHG12 | lncRNA | 192,4808949 | 0,799863771 | 0,257874608 | 3,101754681 | 0,001923773 | 0,042019155 |
| ENSG00000180730.5 | SHISA2 | protein_coding | 431,1152354 | 0,794264158 | 0,152313226 | 5,214676223 | 1,84E-07 | 2,14E-05 |
| ENSG00000164056.11 | SPRY1 | protein_coding | 190,4844038 | 0,79129921 | 0,218421826 | 3,622802839 | 0,000291428 | 0,010090978 |
| ENSG00000158220.14 | ESYT3 | protein_coding | 110,9919343 | 0,787936484 | 0,257031016 | 3,06553075 | 0,00217284 | 0,045226966 |
| ENSG00000175197.12 | DDIT3 | protein_coding | 584,3793472 | 0,778791846 | 0,12661511 | 6,15086025 | 7,71E-10 | 2,07E-07 |
| ENSG00000136059.14 | VILL | protein_coding | 482,6402738 | 0,770303065 | 0,173356192 | 4,443470146 | 8,85E-06 | 0,000640059 |
| ENSG00000146674.15 | IGFBP3 | protein_coding | 402,8891734 | 0,766813642 | 0,132879659 | 5,770737589 | 7,89E-09 | 1,49E-06 |
| ENSG00000156510.13 | HKDC1 | protein_coding | 1323,592766 | 0,762720806 | 0,123802054 | 6,160809002 | 7,24E-10 | 1,98E-07 |
| ENSG00000148655.15 | LRMDA | protein_coding | 330,1251006 | 0,762103045 | 0,216424422 | 3,521335703 | 0,000429379 | 0,013669686 |
| ENSG00000205746.9 | PKD1P4 | transcribed_unproc | 289,1030865 | 0,760611685 | 0,219723883 | 3,461670509 | 0,000536834 | 0,016073055 |
| ENSG00000234678.2 | ELF3-AS1 | lncRNA | 574,205167 | 0,75996576 | 0,155154417 | 4,898125198 | 9,68E-07 | 9,31E-05 |
| ENSG00000135069.14 | PSAT1 | protein_coding | 3449,139335 | 0,757586789 | 0,113479367 | 6,675987119 | 2,46E-11 | 1,05E-08 |
| ENSG00000183696.14 | UPP1 | protein_coding | 438,6310912 | 0,754026344 | 0,13623623 | 5,534697658 | 3,12E-08 | 4,72E-06 |
| ENSG00000178078.12 | STAP2 | protein_coding | 3567,551514 | 0,749635621 | 0,082462784 | 9,090593185 | 9,85E-20 | 3,63E-16 |
| ENSG00000145901.16 | TNIP1 | protein_coding | 7299,153569 | 0,736377639 | 0,08251674 | 8,923978804 | 4,50E-19 | 1,24E-15 |
| ENSG00000139289.13 | PHLDA1 | protein_coding | 668,3518534 | 0,733272008 | 0,120343922 | 6,093137034 | 1,11E-09 | 2,75E-07 |
| ENSG00000128512.23 | DOCK4 | protein_coding | 420,469799 | 0,730711912 | 0,188886864 | 3,86851631 | 0,0001095 | 0,004750566 |
| ENSG00000188659.9 | SAXO2 | protein_coding | 354,3226131 | 0,72946323 | 0,236208981 | 3,088211235 | 0,002013653 | 0,043382749 |
| ENSG00000250251.6 | PKD1P6 | transcribed_unproc | 270,9677657 | 0,726236823 | 0,211702725 | 3,430455708 | 0,000602568 | 0,017585098 |
| ENSG00000177133.11 | PRDM16-DT | lncRNA | 307,7179631 | 0,725028376 | 0,168503486 | 4,302750006 | 1,69E-05 | 0,001075637 |
| ENSG00000069535.14 | MAOB | protein_coding | 4174,927218 | 0,719855226 | 0,10286706 | 6,99791776 | 2,60E-12 | 1,44E-09 |
| ENSG00000232721.2 | AC239800.2 | lncRNA | 233,0608879 | 0,716168681 | 0,180170195 | 3,97495647 | 7,04E-05 | 0,003313822 |
| ENSG00000149150.9 | SLC43A1 | protein_coding | 295,4490254 | 0,710455371 | 0,201564135 | 3,524711237 | 0,000423945 | 0,013535641 |
| ENSG00000134352.20 | IL6ST | protein_coding | 5010,696528 | 0,707629498 | 0,085943867 | 8,233624096 | 1,82E-16 | 3,09E-13 |
| ENSG00000166912.17 | MTMR10 | protein_coding | 5411,903016 | 0,701086561 | 0,105023003 | 6,67555242 | 2,46E-11 | 1,05E-08 |
| ENSG00000177606.8 | JUN | protein_coding | 3508,378964 | 0,699191248 | 0,108049664 | 6,47101732 | 9,73E-11 | 3,42E-08 |

| | | | | | | | | |
|---|---|---|---|---|---|---|---|---|
| ENSG00000151615.3 | POU4F2 | protein_coding | 185,479056 | 0,698621191 | 0,199561201 | 3,500786659 | 0,000463887 | 0,014415681 |
| ENSG00000134769.21 | DTNA | protein_coding | 1740,773878 | 0,698045075 | 0,123646776 | 5,645477363 | 1,65E-08 | 2,76E-06 |
| ENSG00000152495.11 | CAMK4 | protein_coding | 3252,563702 | 0,690453908 | 0,208137491 | 3,317297153 | 0,000908929 | 0,02394162 |
| ENSG00000234745.12 | HLA-B | protein_coding | 660,8040549 | 0,683977266 | 0,116967258 | 5,847595953 | 4,99E-09 | 1,01E-06 |
| ENSG00000281571.2 | AC241585.2 | TEC | 3851,97514 | 0,679232365 | 0,100065788 | 6,787858037 | 1,14E-11 | 5,36E-09 |
| ENSG00000146232.17 | NFKBIE | protein_coding | 2024,950567 | 0,67018059 | 0,091030954 | 7,36211764 | 1,81E-13 | 1,54E-10 |
| ENSG00000280649.2 | AC245100.7 | TEC | 1146,129453 | 0,667119745 | 0,11137448 | 5,989879765 | 2,10E-09 | 4,59E-07 |
| ENSG00000074590.13 | NUAK1 | protein_coding | 2222,043294 | 0,656827445 | 0,104832926 | 6,26546899 | 3,72E-10 | 1,08E-07 |
| ENSG00000035664.11 | DAPK2 | protein_coding | 502,5008059 | 0,653948015 | 0,139294893 | 4,694702016 | 2,67E-06 | 0,000228091 |
| ENSG00000146409.12 | SLC18B1 | protein_coding | 13413,37016 | 0,652958868 | 0,172675247 | 3,781427157 | 0,000155932 | 0,006295891 |
| ENSG00000225127.3 | LINC00237 | lncRNA | 606,251886 | 0,652453761 | 0,121989337 | 5,348449107 | 8,87E-08 | 1,15E-05 |
| ENSG00000187193.9 | MT1X | protein_coding | 1381,979991 | 0,652133129 | 0,092889147 | 7,020552456 | 2,21E-12 | 1,29E-09 |
| ENSG00000134107.5 | BHLHE40 | protein_coding | 379,9510438 | 0,646606927 | 0,145042087 | 4,458064135 | 8,27E-06 | 0,000605924 |
| ENSG00000166143.10 | PPP1R14D | protein_coding | 503,2736987 | 0,646216354 | 0,122841132 | 5,260586106 | 1,44E-07 | 1,77E-05 |
| ENSG00000011201.12 | ANOS1 | protein_coding | 282,1289704 | 0,645602033 | 0,154269193 | 4,184905759 | 2,85E-05 | 0,001618514 |
| ENSG00000170099.6 | SERPINA6 | protein_coding | 6183,885886 | 0,644544758 | 0,076479343 | 8,427697334 | 3,53E-17 | 6,50E-14 |
| ENSG00000137198.10 | GMPR | protein_coding | 930,1495892 | 0,641264735 | 0,115598583 | 5,547340797 | 2,90E-08 | 4,46E-06 |
| ENSG00000249464.6 | LINC01091 | lncRNA | 463,0302423 | 0,640475791 | 0,166830547 | 3,839079838 | 0,000123496 | 0,005224624 |
| ENSG00000198363.18 | ASPH | protein_coding | 31470,19079 | 0,633757073 | 0,092044818 | 6,885309611 | 5,77E-12 | 2,77E-09 |
| ENSG00000160200.18 | CBS | protein_coding | 334,7150409 | 0,633597676 | 0,157494646 | 4,022979136 | 5,75E-05 | 0,002844532 |
| ENSG00000197261.11 | C6orf141 | protein_coding | 531,3142753 | 0,632855824 | 0,152695344 | 4,144565305 | 3,40E-05 | 0,001875879 |
| ENSG00000100100.13 | PIK3IP1 | protein_coding | 312,5122961 | 0,630617577 | 0,16519765 | 3,817351986 | 0,000134892 | 0,005537315 |
| ENSG00000196756.13 | SNHG17 | lncRNA | 506,7678323 | 0,629395393 | 0,116126093 | 5,419930828 | 5,96E-08 | 8,15E-06 |
| ENSG00000147100.11 | SLC16A2 | protein_coding | 5800,645968 | 0,623314276 | 0,088647819 | 7,031354889 | 2,05E-12 | 1,26E-09 |
| ENSG00000104611.12 | SH2D4A | protein_coding | 253,7745853 | 0,622938522 | 0,202583559 | 3,074970774 | 0,002105232 | 0,044531892 |
| ENSG00000155792.10 | DEPTOR | protein_coding | 962,1742175 | 0,622242683 | 0,119113901 | 5,223929997 | 1,75E-07 | 2,08E-05 |
| ENSG00000175832.13 | ETV4 | protein_coding | 1958,763994 | 0,603901095 | 0,090181569 | 6,696502401 | 2,13E-11 | 9,45E-09 |
| ENSG00000172986.13 | GXYLT2 | protein_coding | 339,0153143 | 0,59409877 | 0,190402378 | 3,120227687 | 0,001807113 | 0,040022787 |
| ENSG00000105855.10 | ITGB8 | protein_coding | 1634,343567 | 0,584865179 | 0,106688185 | 5,482005156 | 4,21E-08 | 6,08E-06 |
| ENSG00000088970.16 | KIZ | protein_coding | 1075,264662 | 0,582779186 | 0,102097531 | 5,708063479 | 1,14E-08 | 2,06E-06 |
| ENSG00000155974.12 | GRIP1 | protein_coding | 614,3165853 | 0,579037844 | 0,141834224 | 4,082497338 | 4,46E-05 | 0,002325021 |
| ENSG00000144681.10 | STAC | protein_coding | 350,096159 | 0,57528462 | 0,180839895 | 3,181182005 | 0,001466754 | 0,034415069 |
| ENSG00000104312.8 | RIPK2 | protein_coding | 428,7942544 | 0,574062 | 0,158486284 | 3,622155718 | 0,000292158 | 0,010100455 |
| ENSG00000112561.19 | TFEB | protein_coding | 399,239858 | 0,571440678 | 0,129777299 | 4,40324065 | 1,07E-05 | 0,0007397 |
| ENSG00000137331.12 | IER3 | protein_coding | 1180,289271 | 0,569614936 | 0,114259128 | 4,985290436 | 6,19E-07 | 6,37E-05 |
| ENSG00000101255.11 | TRIB3 | protein_coding | 7547,150763 | 0,569535378 | 0,082698522 | 6,886887004 | 5,70E-12 | 2,77E-09 |

| | | | | | | | | |
|---|---|---|---|---|---|---|---|---|
| ENSG00000123700.5 | KCNJ2 | protein_coding | 2256,61618 | 0,568719842 | 0,111104937 | 5,11876299 | 3,08E-07 | 3,40E-05 |
| ENSG00000237036.5 | ZEB1-AS1 | lncRNA | 419,8287045 | 0,567597248 | 0,153335074 | 3,701679165 | 0,000214177 | 0,008003052 |
| ENSG00000163435.16 | ELF3 | protein_coding | 5130,947172 | 0,562901691 | 0,07891008 | 7,133457345 | 9,79E-13 | 6,37E-10 |
| ENSG00000139318.8 | DUSP6 | protein_coding | 1318,040317 | 0,557320761 | 0,12166114 | 4,580926672 | 4,63E-06 | 0,000367117 |
| ENSG00000125869.10 | LAMP5 | protein_coding | 398,3180772 | 0,555807216 | 0,165325242 | 3,361901715 | 0,000774077 | 0,021276051 |
| ENSG00000119714.11 | GPR68 | protein_coding | 313,4702881 | 0,554278325 | 0,147925872 | 3,747000568 | 0,000178962 | 0,007034859 |
| ENSG00000106541.12 | AGR2 | protein_coding | 1259,460272 | 0,554180382 | 0,148287135 | 3,737211466 | 0,000186072 | 0,007235568 |
| ENSG00000152117.17 | SMPD4BP | transcribed_process | 648,2768919 | 0,553066786 | 0,108154254 | 5,113685012 | 3,16E-07 | 3,48E-05 |
| ENSG00000161955.16 | TNFSF13 | protein_coding | 448,1065096 | 0,550919464 | 0,135128402 | 4,077007159 | 4,56E-05 | 0,002364236 |
| ENSG00000157557.13 | ETS2 | protein_coding | 6062,394134 | 0,54946795 | 0,081238006 | 6,763680878 | 1,35E-11 | 6,20E-09 |
| ENSG00000229891.2 | LINC01315 | lncRNA | 840,1557694 | 0,54767452 | 0,11021448 | 4,969170287 | 6,72E-07 | 6,79E-05 |
| ENSG00000131370.16 | SH3BP5 | protein_coding | 215,9692476 | 0,547012759 | 0,177451222 | 3,082609138 | 0,002051945 | 0,043781417 |
| ENSG00000139192.12 | TAPBPL | protein_coding | 470,5166423 | 0,546093659 | 0,135711407 | 4,023933384 | 5,72E-05 | 0,002839377 |
| ENSG00000205220.12 | PSMB10 | protein_coding | 351,5743309 | 0,545645227 | 0,136430687 | 3,999431789 | 6,35E-05 | 0,003087659 |
| ENSG00000226067.7 | LINC00623 | lncRNA | 2268,732148 | 0,545231054 | 0,08877509 | 6,141712222 | 8,16E-10 | 2,15E-07 |
| ENSG00000169436.17 | COL22A1 | protein_coding | 7941,955269 | 0,544763956 | 0,085191688 | 6,394566977 | 1,61E-10 | 5,24E-08 |
| ENSG00000067057.18 | PFKP | protein_coding | 2431,698933 | 0,54278594 | 0,077488227 | 7,004753668 | 2,47E-12 | 1,40E-09 |
| ENSG00000007516.14 | BAIAP3 | protein_coding | 2773,493639 | 0,54243117 | 0,07166741 | 7,568728577 | 3,77E-14 | 3,97E-11 |
| ENSG00000104432.14 | IL7 | protein_coding | 685,9113236 | 0,540121348 | 0,120690544 | 4,475258196 | 7,63E-06 | 0,000565664 |
| ENSG00000164761.9 | TNFRSF11B | protein_coding | 268,3674434 | 0,534593786 | 0,156061888 | 3,425524282 | 0,000613614 | 0,01773922 |
| ENSG00000152932.8 | RAB3C | protein_coding | 10164,58323 | 0,526939777 | 0,084176954 | 6,259905477 | 3,85E-10 | 1,11E-07 |
| ENSG00000171617.14 | ENC1 | protein_coding | 1978,343213 | 0,525799075 | 0,088881412 | 5,915737177 | 3,30E-09 | 6,90E-07 |
| ENSG00000059377.18 | TBXAS1 | protein_coding | 512,104553 | 0,524690331 | 0,150373806 | 3,48924022 | 0,000484396 | 0,014885747 |
| ENSG00000124145.6 | SDC4 | protein_coding | 3104,733457 | 0,524301276 | 0,097932569 | 5,353696736 | 8,62E-08 | 1,13E-05 |
| ENSG00000135318.12 | NT5E | protein_coding | 3892,835895 | 0,522410236 | 0,087131534 | 5,995650638 | 2,03E-09 | 4,53E-07 |
| ENSG00000256043.4 | CTSO | protein_coding | 920,5549017 | 0,520866556 | 0,106233433 | 4,903037957 | 9,44E-07 | 9,16E-05 |
| ENSG00000079689.14 | SCGN | protein_coding | 49085,5127 | 0,514919842 | 0,073879646 | 6,96971179 | 3,18E-12 | 1,67E-09 |
| ENSG00000127946.17 | HIP1 | protein_coding | 724,9999423 | 0,509810242 | 0,113483075 | 4,492390106 | 7,04E-06 | 0,000528236 |
| ENSG00000183604.14 | SMG1P5 | transcribed_unproc | 266,8711009 | 0,507495977 | 0,164560671 | 3,083944503 | 0,002042757 | 0,043627453 |
| ENSG00000087074.8 | PPP1R15A | protein_coding | 1224,830172 | 0,50690656 | 0,100404859 | 5,048625759 | 4,45E-07 | 4,76E-05 |
| ENSG00000288596.2 | C8orf44 | lncRNA | 400,0617478 | 0,506247749 | 0,14038473 | 3,606145414 | 0,000310779 | 0,010546467 |
| ENSG00000106069.23 | CHN2 | protein_coding | 1209,6124 | 0,505037357 | 0,107077953 | 4,716539164 | 2,40E-06 | 0,00020653 |
| ENSG00000206503.13 | HLA-A | protein_coding | 62354,54993 | 0,505029303 | 0,063429318 | 7,962079971 | 1,69E-15 | 2,34E-12 |
| ENSG00000155629.15 | PIK3AP1 | protein_coding | 387,0786684 | 0,494915808 | 0,148752547 | 3,327108122 | 0,000877523 | 0,023421083 |
| ENSG00000115267.8 | IFIH1 | protein_coding | 1647,044832 | 0,494307744 | 0,105541044 | 4,683559371 | 2,82E-06 | 0,000239008 |
| ENSG00000230006.7 | ANKRD36BP2 | transcribed_unproc | 397,6973856 | 0,493953868 | 0,128274655 | 3,850751875 | 0,000117756 | 0,005059153 |

| | | | | | | | | |
|---|---|---|---|---|---|---|---|---|
| ENSG00000178685.14 | PARP10 | protein_coding | 927,5915865 | 0,493273937 | 0,101694149 | 4,850563608 | 1,23E-06 | 0,000114452 |
| ENSG00000099250.18 | NRP1 | protein_coding | 980,3398939 | 0,492435002 | 0,107943369 | 4,56197546 | 5,07E-06 | 0,000397598 |
| ENSG00000130222.11 | GADD45G | protein_coding | 1061,467356 | 0,491437553 | 0,091847687 | 5,350570803 | 8,77E-08 | 1,15E-05 |
| ENSG00000173114.13 | LRRN3 | protein_coding | 325,279366 | 0,488913609 | 0,153563591 | 3,183785983 | 0,001453624 | 0,034242148 |
| ENSG00000250303.4 | LINC02762 | lncRNA | 408,3062838 | 0,485983695 | 0,152044896 | 3,196317058 | 0,001391941 | 0,03305194 |
| ENSG00000261135.1 | AL137802.2 | lncRNA | 584,9842545 | 0,48482693 | 0,112828881 | 4,297010888 | 1,73E-05 | 0,001094394 |
| ENSG00000128271.22 | ADORA2A | protein_coding | 585,3804205 | 0,483715064 | 0,116940617 | 4,13641622 | 3,53E-05 | 0,001922521 |
| ENSG00000170385.10 | SLC30A1 | protein_coding | 5810,399015 | 0,483544135 | 0,0944217 | 5,121112365 | 3,04E-07 | 3,38E-05 |
| ENSG00000132330.18 | SCLY | protein_coding | 2618,557292 | 0,481036459 | 0,085136759 | 5,650161781 | 1,60E-08 | 2,73E-06 |
| ENSG00000163497.3 | FEV | protein_coding | 370,6504802 | 0,478662496 | 0,157525755 | 3,038630076 | 0,002376565 | 0,048065698 |
| ENSG00000197329.12 | PELI1 | protein_coding | 5585,794283 | 0,47677239 | 0,095532226 | 4,990696936 | 6,02E-07 | 6,22E-05 |
| ENSG00000100889.12 | PCK2 | protein_coding | 1454,299869 | 0,47669983 | 0,104801914 | 4,548579445 | 5,40E-06 | 0,000419301 |
| ENSG00000185518.12 | SV2B | protein_coding | 1037,801797 | 0,476196975 | 0,094155025 | 5,057584277 | 4,25E-07 | 4,61E-05 |
| ENSG00000126262.5 | FFAR2 | protein_coding | 1733,256226 | 0,472352022 | 0,089819957 | 5,258876076 | 1,45E-07 | 1,77E-05 |
| ENSG00000178695.6 | KCTD12 | protein_coding | 14871,41931 | 0,467775347 | 0,081941806 | 5,708628752 | 1,14E-08 | 2,06E-06 |
| ENSG00000251192.7 | ZNF674 | protein_coding | 411,3705435 | 0,465894207 | 0,149278614 | 3,120970874 | 0,001802559 | 0,040003422 |
| ENSG00000204264.12 | PSMB8 | protein_coding | 902,2367322 | 0,463033187 | 0,112176274 | 4,127728377 | 3,66E-05 | 0,001981951 |
| ENSG00000196542.8 | SPTSSB | protein_coding | 1249,549378 | 0,462116794 | 0,11438008 | 4,040185983 | 5,34E-05 | 0,002704173 |
| ENSG00000142677.4 | IL22RA1 | protein_coding | 612,637798 | 0,461642859 | 0,119232461 | 3,871788388 | 0,00010804 | 0,004696438 |
| ENSG00000176153.12 | GPX2 | protein_coding | 711,213081 | 0,461109517 | 0,139781114 | 3,298796985 | 0,000971001 | 0,025127918 |
| ENSG00000082497.12 | SERTAD4 | protein_coding | 556,7036089 | 0,458864101 | 0,12118211 | 3,786566346 | 0,000152743 | 0,006189741 |
| ENSG00000167074.15 | TEF | protein_coding | 2313,847177 | 0,458523104 | 0,104177844 | 4,401349521 | 1,08E-05 | 0,000743267 |
| ENSG00000139832.5 | RAB20 | protein_coding | 347,6885039 | 0,456382233 | 0,145388943 | 3,139043605 | 0,001695002 | 0,038152207 |
| ENSG00000133619.17 | KRBA1 | protein_coding | 778,5536546 | 0,455932316 | 0,142024144 | 3,210245125 | 0,001326218 | 0,03191128 |
| ENSG00000184164.15 | CRELD2 | protein_coding | 4871,657173 | 0,455587513 | 0,066002616 | 6,902567497 | 5,11E-12 | 2,63E-09 |
| ENSG00000110619.18 | CARS1 | protein_coding | 2568,268039 | 0,454421682 | 0,093839611 | 4,84253587 | 1,28E-06 | 0,00011795 |
| ENSG00000169136.12 | ATF5 | protein_coding | 6026,941287 | 0,450564743 | 0,075872432 | 5,938451324 | 2,88E-09 | 6,06E-07 |
| ENSG00000188522.15 | FAM83G | protein_coding | 2322,741075 | 0,448795004 | 0,076775467 | 5,845552249 | 5,05E-09 | 1,01E-06 |
| ENSG00000122435.10 | TRMT13 | protein_coding | 770,9654296 | 0,446479381 | 0,142519273 | 3,132764927 | 0,001731681 | 0,038663136 |
| ENSG00000164929.17 | BAALC | protein_coding | 1378,472854 | 0,44444583 | 0,11627905 | 3,822234778 | 0,000132248 | 0,005473351 |
| ENSG00000099625.13 | CBARP | protein_coding | 373,7699772 | 0,441359172 | 0,144240873 | 3,059875902 | 0,002214287 | 0,045830981 |
| ENSG00000151952.16 | TMEM132D | protein_coding | 6001,020195 | 0,440586138 | 0,069580019 | 6,332078405 | 2,42E-10 | 7,54E-08 |
| ENSG00000151883.19 | PARP8 | protein_coding | 2334,174168 | 0,439997887 | 0,106530027 | 4,130271067 | 3,62E-05 | 0,001964961 |
| ENSG00000123240.17 | OPTN | protein_coding | 6768,106284 | 0,43666642 | 0,082107959 | 5,318198431 | 1,05E-07 | 1,34E-05 |
| ENSG00000070540.13 | WIPI1 | protein_coding | 687,4490721 | 0,435523665 | 0,118199229 | 3,684657397 | 0,00022901 | 0,008389215 |
| ENSG00000109171.15 | SLAIN2 | protein_coding | 4063,303091 | 0,434936268 | 0,095271778 | 4,565216217 | 4,99E-06 | 0,000392898 |

| | | | | | | | | |
|---|---|---|---|---|---|---|---|---|
| ENSG00000182704.8 | TSKU | protein_coding | 1123,949898 | 0,432019787 | 0,108578433 | 3,978872939 | 6,92E-05 | 0,003273642 |
| ENSG00000112149.10 | CD83 | protein_coding | 2414,225939 | 0,427347076 | 0,085984072 | 4,970072534 | 6,69E-07 | 6,79E-05 |
| ENSG00000026508.20 | CD44 | protein_coding | 10677,55865 | 0,426507367 | 0,076531948 | 5,572932351 | 2,50E-08 | 3,93E-06 |
| ENSG00000277147.6 | LINC00869 | transcribed_unproc | 1154,168702 | 0,42587252 | 0,097473385 | 4,36911593 | 1,25E-05 | 0,000826416 |
| ENSG00000101298.15 | SNPH | protein_coding | 2843,202431 | 0,422572068 | 0,069492899 | 6,080794886 | 1,20E-09 | 2,94E-07 |
| ENSG00000114268.12 | PFKFB4 | protein_coding | 11564,70163 | 0,421519352 | 0,079451416 | 5,305372461 | 1,12E-07 | 1,42E-05 |
| ENSG00000110328.6 | GALNT18 | protein_coding | 4411,129818 | 0,420580254 | 0,06504041 | 6,466445289 | 1,00E-10 | 3,42E-08 |
| ENSG00000274265.5 | AC245297.3 | lncRNA | 995,1750129 | 0,418817498 | 0,105550814 | 3,967922944 | 7,25E-05 | 0,00340589 |
| ENSG00000120875.9 | DUSP4 | protein_coding | 3020,811849 | 0,417825248 | 0,075955069 | 5,500952762 | 3,78E-08 | 5,54E-06 |
| ENSG00000136048.14 | DRAM1 | protein_coding | 389,389546 | 0,416144309 | 0,135900201 | 3,062131671 | 0,002197668 | 0,045615003 |
| ENSG00000197956.10 | S100A6 | protein_coding | 9652,854053 | 0,414266922 | 0,083973488 | 4,933306145 | 8,08E-07 | 8,07E-05 |
| ENSG00000049759.19 | NEDD4L | protein_coding | 6905,80666 | 0,413420856 | 0,082258831 | 5,025853788 | 5,01E-07 | 5,33E-05 |
| ENSG00000232956.9 | SNHG15 | lncRNA | 572,6031015 | 0,413384441 | 0,111260611 | 3,715460832 | 0,000202834 | 0,007724437 |
| ENSG00000158710.15 | TAGLN2 | protein_coding | 54614,80527 | 0,409534548 | 0,066068497 | 6,198635754 | 5,70E-10 | 1,58E-07 |
| ENSG00000155660.11 | PDIA4 | protein_coding | 24119,33508 | 0,405021884 | 0,061392888 | 6,597211784 | 4,19E-11 | 1,60E-08 |
| ENSG00000198208.12 | RPS6KL1 | protein_coding | 1512,925235 | 0,40289191 | 0,111147117 | 3,624852548 | 0,000289126 | 0,010026975 |
| ENSG00000275964.1 | AL355001.2 | lncRNA | 690,9931852 | 0,398816912 | 0,113408623 | 3,51663659 | 0,000437052 | 0,013834344 |
| ENSG00000133027.18 | PEMT | protein_coding | 1128,847591 | 0,397898201 | 0,093010922 | 4,27797286 | 1,89E-05 | 0,001165651 |
| ENSG00000159348.13 | CYB5R1 | protein_coding | 8171,968832 | 0,396561599 | 0,069647385 | 5,693847648 | 1,24E-08 | 2,20E-06 |
| ENSG00000174738.13 | NR1D2 | protein_coding | 7262,245338 | 0,39642055 | 0,073109897 | 5,422255626 | 5,89E-08 | 8,14E-06 |
| ENSG00000163605.15 | PPP4R2 | protein_coding | 17347,98312 | 0,393996193 | 0,07844663 | 5,02247445 | 5,10E-07 | 5,40E-05 |
| ENSG00000128228.5 | SDF2L1 | protein_coding | 7312,076395 | 0,393828276 | 0,061849341 | 6,36754198 | 1,92E-10 | 6,07E-08 |
| ENSG00000011132.12 | APBA3 | protein_coding | 845,6754247 | 0,390166132 | 0,099990517 | 3,902031355 | 9,54E-05 | 0,004246625 |
| ENSG00000100911.16 | PSME2 | protein_coding | 4932,942095 | 0,387864267 | 0,072107587 | 5,378966121 | 7,49E-08 | 9,99E-06 |
| ENSG00000162924.15 | REL | protein_coding | 2283,129933 | 0,386029994 | 0,096752056 | 3,989889317 | 6,61E-05 | 0,003179609 |
| ENSG00000182325.11 | FBXL6 | protein_coding | 917,2047307 | 0,384822495 | 0,09541469 | 4,033157728 | 5,50E-05 | 0,00274913 |
| ENSG00000139083.11 | ETV6 | protein_coding | 1082,785129 | 0,384116327 | 0,091427135 | 4,201338315 | 2,65E-05 | 0,001540932 |
| ENSG00000165801.10 | ARHGEF40 | protein_coding | 2445,824914 | 0,382480555 | 0,092580143 | 4,131345467 | 3,61E-05 | 0,001960603 |
| ENSG00000176597.12 | B3GNT5 | protein_coding | 4106,23353 | 0,379956169 | 0,086603898 | 4,387287132 | 1,15E-05 | 0,000778979 |
| ENSG00000149289.11 | ZC3H12C | protein_coding | 3711,455921 | 0,375594743 | 0,08904986 | 4,217802708 | 2,47E-05 | 0,001454985 |
| ENSG00000138166.6 | DUSP5 | protein_coding | 1922,926999 | 0,375472586 | 0,087437265 | 4,294194064 | 1,75E-05 | 0,001105222 |
| ENSG00000102554.14 | KLF5 | protein_coding | 1738,81699 | 0,374612557 | 0,082738871 | 4,5276489 | 5,96E-06 | 0,000456634 |
| ENSG00000250799.11 | PRODH2 | protein_coding | 3876,798353 | 0,374441336 | 0,071787402 | 5,215975552 | 1,83E-07 | 2,14E-05 |
| ENSG00000160145.15 | KALRN | protein_coding | 1065,346339 | 0,372386746 | 0,093223312 | 3,994566767 | 6,48E-05 | 0,003124281 |
| ENSG00000166342.19 | NETO1 | protein_coding | 2446,246301 | 0,371538967 | 0,08620892 | 4,309750854 | 1,63E-05 | 0,001045157 |
| ENSG00000143627.19 | PKLR | protein_coding | 943,6772825 | 0,365293307 | 0,118965415 | 3,07058406 | 0,002136405 | 0,044763358 |

| | | | | | | | | |
|---|---|---|---|---|---|---|---|---|
| ENSG00000113924.12 | HGD | protein_coding | 2935,26389 | 0,365074236 | 0,081791236 | 4,46348842 | 8,06E-06 | 0,000592741 |
| ENSG00000259024.6 | TVP23C-CDRT4 | protein_coding | 754,9397865 | 0,363697243 | 0,100602185 | 3,615202243 | 0,000300113 | 0,010343155 |
| ENSG00000157353.17 | FCSK | protein_coding | 1362,395106 | 0,363112489 | 0,094474654 | 3,843491081 | 0,000121296 | 0,005167617 |
| ENSG00000109790.16 | KLHL5 | protein_coding | 3973,955143 | 0,362876134 | 0,083753147 | 4,332686544 | 1,47E-05 | 0,00095577 |
| ENSG00000156384.14 | SFR1 | protein_coding | 1212,039532 | 0,361844622 | 0,10939922 | 3,307561271 | 0,000941121 | 0,024535889 |
| ENSG00000166444.19 | DENND2B | protein_coding | 2672,270173 | 0,361545205 | 0,090429153 | 3,99810452 | 6,39E-05 | 0,00309821 |
| ENSG00000241163.8 | LINC00877 | lncRNA | 1824,656683 | 0,36003203 | 0,092891566 | 3,875831206 | 0,000106261 | 0,004639937 |
| ENSG00000141510.18 | TP53 | protein_coding | 2957,287371 | 0,359934894 | 0,073202316 | 4,916987826 | 8,79E-07 | 8,64E-05 |
| ENSG00000168938.6 | PPIC | protein_coding | 1579,730605 | 0,358625416 | 0,094174506 | 3,808094462 | 0,000140042 | 0,005738082 |
| ENSG00000155090.15 | KLF10 | protein_coding | 1826,624382 | 0,35741617 | 0,081602287 | 4,379977343 | 1,19E-05 | 0,00079581 |
| ENSG00000149599.15 | DUSP15 | protein_coding | 883,7698515 | 0,357026159 | 0,112872754 | 3,163085389 | 0,001561065 | 0,035830013 |
| ENSG00000006652.15 | IFRD1 | protein_coding | 1492,467233 | 0,356665753 | 0,098190096 | 3,632400476 | 0,000280797 | 0,009874809 |
| ENSG00000021488.13 | SLC7A9 | protein_coding | 3576,827399 | 0,355893108 | 0,092603701 | 3,843184495 | 0,000121448 | 0,005167617 |
| ENSG00000165246.14 | NLGN4Y | protein_coding | 2591,994532 | 0,354838877 | 0,095082437 | 3,731907702 | 0,000190035 | 0,007350905 |
| ENSG00000104231.11 | ZFAND1 | protein_coding | 2337,905774 | 0,353993199 | 0,105950383 | 3,341122414 | 0,000834404 | 0,022487245 |
| ENSG00000139793.18 | MBNL2 | protein_coding | 2287,71323 | 0,351772556 | 0,104772867 | 3,357477609 | 0,000786571 | 0,02154536 |
| ENSG00000153822.14 | KCNJ16 | protein_coding | 1624,457687 | 0,350598365 | 0,10521886 | 3,33208672 | 0,000861974 | 0,023117618 |
| ENSG00000151491.14 | EPS8 | protein_coding | 4349,176044 | 0,350080163 | 0,080237483 | 4,363050186 | 1,28E-05 | 0,000847139 |
| ENSG00000179630.11 | LACC1 | protein_coding | 984,8170368 | 0,349104006 | 0,115024225 | 3,035047664 | 0,002404977 | 0,048551566 |
| ENSG00000170542.6 | SERPINB9 | protein_coding | 947,9773521 | 0,34785473 | 0,099922799 | 3,481234854 | 0,000499108 | 0,015274213 |
| ENSG00000178607.16 | ERN1 | protein_coding | 1172,283397 | 0,34413114 | 0,100600052 | 3,420784907 | 0,000624407 | 0,017934084 |
| ENSG00000070669.17 | ASNS | protein_coding | 5990,189081 | 0,343609006 | 0,077501394 | 4,433584837 | 9,27E-06 | 0,000661489 |
| ENSG00000182199.11 | SHMT2 | protein_coding | 10922,39049 | 0,343391446 | 0,080090895 | 4,287521643 | 1,81E-05 | 0,001129286 |
| ENSG00000095209.12 | TMEM38B | protein_coding | 8152,304484 | 0,339430772 | 0,074842602 | 4,535261497 | 5,75E-06 | 0,000441999 |
| ENSG00000145050.19 | MANF | protein_coding | 17648,39482 | 0,33844002 | 0,056038156 | 6,03945673 | 1,55E-09 | 3,67E-07 |
| ENSG00000196455.8 | PIK3R4 | protein_coding | 5243,548522 | 0,338422763 | 0,080156724 | 4,222013417 | 2,42E-05 | 0,001440151 |
| ENSG00000065911.13 | MTHFD2 | protein_coding | 5286,135863 | 0,338334953 | 0,079344165 | 4,264144112 | 2,01E-05 | 0,001219786 |
| ENSG00000130768.15 | SMPDL3B | protein_coding | 7083,066946 | 0,338294264 | 0,061370933 | 5,512288108 | 3,54E-08 | 5,26E-06 |
| ENSG00000127952.17 | STYXL1 | protein_coding | 910,1156563 | 0,334268148 | 0,106844843 | 3,128537975 | 0,001756783 | 0,039184055 |
| ENSG00000185947.15 | ZNF267 | protein_coding | 1595,132285 | 0,333982614 | 0,105550108 | 3,164209117 | 0,00155505 | 0,03577443 |
| ENSG00000059378.13 | PARP12 | protein_coding | 1268,82337 | 0,332122327 | 0,086663419 | 3,832324314 | 0,000126938 | 0,005319386 |
| ENSG00000178538.10 | CA8 | protein_coding | 5966,042076 | 0,329407168 | 0,072348224 | 4,553078849 | 5,29E-06 | 0,000411874 |
| ENSG00000118946.12 | PCDH17 | protein_coding | 1868,177692 | 0,328953687 | 0,080437194 | 4,089571867 | 4,32E-05 | 0,002265923 |
| ENSG00000145687.16 | SSBP2 | protein_coding | 2969,890466 | 0,328433714 | 0,080816218 | 4,063957958 | 4,82E-05 | 0,002488402 |
| ENSG00000095380.11 | NANS | protein_coding | 2737,925125 | 0,327480659 | 0,072361799 | 4,525601424 | 6,02E-06 | 0,000459488 |
| ENSG00000140465.14 | CYP1A1 | protein_coding | 2912,91513 | 0,327387583 | 0,069846873 | 4,687218889 | 2,77E-06 | 0,000235678 |

| | | | | | | | | |
|---|---|---|---|---|---|---|---|---|
| ENSG00000142544.7 | CTU1 | protein_coding | 688,4580465 | 0,327226694 | 0,106615645 | 3,069218357 | 0,002146197 | 0,044888475 |
| ENSG00000108176.15 | DNAJC12 | protein_coding | 3695,409297 | 0,3265303 | 0,083142344 | 3,927364597 | 8,59E-05 | 0,003926073 |
| ENSG00000115290.10 | GRB14 | protein_coding | 814,086808 | 0,326228927 | 0,105704099 | 3,086246719 | 0,002027006 | 0,043480147 |
| ENSG00000109113.20 | RAB34 | protein_coding | 1793,294776 | 0,325025293 | 0,078532899 | 4,138715059 | 3,49E-05 | 0,00191753 |
| ENSG00000184227.8 | ACOT1 | protein_coding | 663,9396625 | 0,322564661 | 0,10326286 | 3,123723874 | 0,00178578 | 0,039670842 |
| ENSG00000136026.14 | CKAP4 | protein_coding | 4035,331224 | 0,32201258 | 0,072381376 | 4,448831957 | 8,63E-06 | 0,000626337 |
| ENSG00000164023.14 | SGMS2 | protein_coding | 6130,516939 | 0,319693493 | 0,100771082 | 3,172472547 | 0,001511468 | 0,035044954 |
| ENSG00000198393.8 | ZNF26 | protein_coding | 1162,108599 | 0,319501225 | 0,100965475 | 3,164460187 | 0,001553709 | 0,03577443 |
| ENSG00000153721.19 | CNKSR3 | protein_coding | 966,7752064 | 0,317413117 | 0,089980558 | 3,527574452 | 0,000419386 | 0,013409437 |
| ENSG00000176887.7 | SOX11 | protein_coding | 4798,792662 | 0,316965572 | 0,076014028 | 4,169829969 | 3,05E-05 | 0,001707494 |
| ENSG00000137094.15 | DNAJB5 | protein_coding | 961,1013409 | 0,314922033 | 0,099037045 | 3,179840773 | 0,00147356 | 0,034465103 |
| ENSG00000109103.12 | UNC119 | protein_coding | 2729,731255 | 0,314579493 | 0,071106216 | 4,42407869 | 9,69E-06 | 0,000684667 |
| ENSG00000131979.20 | GCH1 | protein_coding | 21735,90606 | 0,313542417 | 0,072958937 | 4,297518998 | 1,73E-05 | 0,001094394 |
| ENSG00000145780.8 | FEM1C | protein_coding | 3272,986674 | 0,313426963 | 0,08459859 | 3,704872177 | 0,000211497 | 0,007931509 |
| ENSG00000060558.4 | GNA15 | protein_coding | 1591,935093 | 0,311955886 | 0,078949882 | 3,951315404 | 7,77E-05 | 0,003612804 |
| ENSG00000139410.15 | SDSL | protein_coding | 1169,318044 | 0,31107834 | 0,086253031 | 3,606578645 | 0,000310261 | 0,010545056 |
| ENSG00000166016.6 | ABTB2 | protein_coding | 1437,764749 | 0,310721263 | 0,082150481 | 3,782342581 | 0,000155359 | 0,006284243 |
| ENSG00000171345.13 | KRT19 | protein_coding | 8282,360802 | 0,308834186 | 0,057031383 | 5,415162129 | 6,12E-08 | 8,26E-06 |
| ENSG00000139163.16 | ETNK1 | protein_coding | 46146,71378 | 0,30690378 | 0,074041166 | 4,145042518 | 3,40E-05 | 0,001875879 |
| ENSG00000134058.12 | CDK7 | protein_coding | 1328,472369 | 0,305162685 | 0,087729277 | 3,478458919 | 0,000504306 | 0,015411973 |
| ENSG00000161835.11 | TAMALIN | protein_coding | 3178,297244 | 0,304507128 | 0,062742528 | 4,853281112 | 1,21E-06 | 0,00011337 |
| ENSG00000127666.10 | TICAM1 | protein_coding | 1972,763135 | 0,304132103 | 0,074520898 | 4,081165297 | 4,48E-05 | 0,002332886 |
| ENSG00000166794.5 | PPIB | protein_coding | 54642,4713 | 0,303760958 | 0,058594169 | 5,184149992 | 2,17E-07 | 2,47E-05 |
| ENSG00000109320.13 | NFKB1 | protein_coding | 2960,586044 | 0,303594664 | 0,06986859 | 4,345223881 | 1,39E-05 | 0,000913493 |
| ENSG00000137269.15 | LRRC1 | protein_coding | 3295,808008 | 0,302437829 | 0,076578545 | 3,949380742 | 7,84E-05 | 0,003634492 |
| ENSG00000100908.14 | EMC9 | protein_coding | 1525,860244 | 0,30208155 | 0,082679132 | 3,653661363 | 0,000258527 | 0,009255943 |
| ENSG00000112306.8 | RPS12 | protein_coding | 39818,5417 | 0,299619142 | 0,061782685 | 4,849564886 | 1,24E-06 | 0,000114548 |
| ENSG00000092010.15 | PSME1 | protein_coding | 7805,97877 | 0,29865588 | 0,067493979 | 4,42492629 | 9,65E-06 | 0,00068417 |
| ENSG00000145996.11 | CDKAL1 | protein_coding | 2688,756242 | 0,298649948 | 0,065503176 | 4,559320087 | 5,13E-06 | 0,000401235 |
| ENSG00000134013.16 | LOXL2 | protein_coding | 3324,498107 | 0,298567664 | 0,078506983 | 3,803071442 | 0,000142913 | 0,005844909 |
| ENSG00000182648.13 | LINC01006 | lncRNA | 2203,006087 | 0,297538733 | 0,088250616 | 3,371520174 | 0,000747546 | 0,020753067 |
| ENSG00000113532.13 | ST8SIA4 | protein_coding | 3823,932678 | 0,297374913 | 0,088786649 | 3,349320143 | 0,000810101 | 0,021966056 |
| ENSG00000183878.15 | UTY | protein_coding | 1208,114099 | 0,297169532 | 0,098231818 | 3,025186123 | 0,002484802 | 0,049754499 |
| ENSG00000166012.17 | TAF1D | protein_coding | 1494,913579 | 0,29512078 | 0,087303721 | 3,380391766 | 0,000723826 | 0,02017049 |
| ENSG00000149179.14 | C11orf49 | protein_coding | 5470,880107 | 0,294217496 | 0,057167224 | 5,146611521 | 2,65E-07 | 2,98E-05 |
| ENSG00000171223.6 | JUNB | protein_coding | 4014,951566 | 0,29389843 | 0,071031441 | 4,137582245 | 3,51E-05 | 0,001922249 |

| | | | | | | | | |
|---|---|---|---|---|---|---|---|---|
| ENSG00000137038.8 | DMAC1 | protein_coding | 3875,897497 | 0,292744642 | 0,074955263 | 3,905591573 | 9,40E-05 | 0,004201488 |
| ENSG00000107562.16 | CXCL12 | protein_coding | 4750,417027 | 0,292296678 | 0,072467171 | 4,033504713 | 5,50E-05 | 0,00274913 |
| ENSG00000128272.15 | ATF4 | protein_coding | 6827,842388 | 0,290154101 | 0,057433114 | 5,052035017 | 4,37E-07 | 4,70E-05 |
| ENSG00000163082.10 | SGPP2 | protein_coding | 4138,60243 | 0,289370572 | 0,080874302 | 3,578028677 | 0,000346195 | 0,011518678 |
| ENSG00000166133.18 | RPUSD2 | protein_coding | 1162,265175 | 0,288542048 | 0,08373412 | 3,44593157 | 0,000569094 | 0,016766689 |
| ENSG00000149428.19 | HYOU1 | protein_coding | 36890,30766 | 0,288165329 | 0,051449682 | 5,600915683 | 2,13E-08 | 3,44E-06 |
| ENSG00000196975.16 | ANXA4 | protein_coding | 5326,098914 | 0,288145692 | 0,068391002 | 4,213210577 | 2,52E-05 | 0,001473698 |
| ENSG00000136698.10 | CFC1 | protein_coding | 4748,79729 | 0,287533691 | 0,067578776 | 4,254792797 | 2,09E-05 | 0,001264946 |
| ENSG00000269893.8 | SNHG8 | lncRNA | 1735,65286 | 0,287401568 | 0,084855402 | 3,386956645 | 0,000706726 | 0,019793683 |
| ENSG00000166598.15 | HSP90B1 | protein_coding | 202033,4169 | 0,287208254 | 0,066344762 | 4,329026818 | 1,50E-05 | 0,000968949 |
| ENSG00000169964.8 | TMEM42 | protein_coding | 1247,811402 | 0,286325627 | 0,083525707 | 3,427994067 | 0,000608059 | 0,017632907 |
| ENSG00000068079.8 | IFI35 | protein_coding | 1003,836934 | 0,284770647 | 0,090506591 | 3,146407825 | 0,001652894 | 0,037548175 |
| ENSG00000077232.18 | DNAJC10 | protein_coding | 13343,70166 | 0,282635977 | 0,071368788 | 3,960218254 | 7,49E-05 | 0,003502798 |
| ENSG00000133477.17 | FAM83F | protein_coding | 2549,39919 | 0,282268689 | 0,078714528 | 3,585979553 | 0,000335815 | 0,011240921 |
| ENSG00000165655.17 | ZNF503 | protein_coding | 1582,296212 | 0,28193565 | 0,086300663 | 3,266900163 | 0,00108732 | 0,027526369 |
| ENSG00000125089.17 | SH3TC1 | protein_coding | 2349,010462 | 0,281420462 | 0,076548755 | 3,676355855 | 0,000236589 | 0,008624016 |
| ENSG00000150687.12 | PRSS23 | protein_coding | 2981,253374 | 0,280714814 | 0,076066372 | 3,690393096 | 0,000223908 | 0,008243234 |
| ENSG00000108641.19 | B9D1 | protein_coding | 1451,459442 | 0,279304088 | 0,086002673 | 3,247621029 | 0,001163742 | 0,028834206 |
| ENSG00000100033.16 | PRODH | protein_coding | 5900,203378 | 0,279058637 | 0,063157386 | 4,418464023 | 9,94E-06 | 0,000700456 |
| ENSG00000152779.14 | SLC16A12 | protein_coding | 8116,050655 | 0,278994813 | 0,077398718 | 3,604643863 | 0,000312581 | 0,010591383 |
| ENSG00000116584.20 | ARHGEF2 | protein_coding | 1271,34824 | 0,278134218 | 0,083714622 | 3,322409069 | 0,000892438 | 0,023704769 |
| ENSG00000159388.6 | BTG2 | protein_coding | 2961,901824 | 0,27761637 | 0,072215505 | 3,844276523 | 0,000120909 | 0,005164525 |
| ENSG00000164587.13 | RPS14 | protein_coding | 47693,81715 | 0,27718102 | 0,082770897 | 3,348773905 | 0,0008117 | 0,021982467 |
| ENSG00000125995.16 | ROMO1 | protein_coding | 8717,121035 | 0,276958995 | 0,064448196 | 4,297389424 | 1,73E-05 | 0,001094394 |
| ENSG00000196387.9 | ZNF140 | protein_coding | 2047,46476 | 0,275977306 | 0,080644984 | 3,422126113 | 0,000621335 | 0,017923931 |
| ENSG00000166986.15 | MARS1 | protein_coding | 12484,89195 | 0,275887755 | 0,073276743 | 3,765011182 | 0,000166542 | 0,00661562 |
| ENSG00000231925.12 | TAPBP | protein_coding | 2339,671383 | 0,275324419 | 0,077851952 | 3,536512746 | 0,000405447 | 0,013060223 |
| ENSG00000067082.15 | KLF6 | protein_coding | 1588,58809 | 0,275038456 | 0,088891322 | 3,094097935 | 0,001974123 | 0,042949309 |
| ENSG00000105835.12 | NAMPT | protein_coding | 6538,742456 | 0,273443084 | 0,079602973 | 3,435086335 | 0,000592365 | 0,017336865 |
| ENSG00000176840.12 | MIR7-3HG | lncRNA | 2408,101402 | 0,271515784 | 0,075281225 | 3,606686586 | 0,000310132 | 0,010545056 |
| ENSG00000114115.10 | RBP1 | protein_coding | 1521,123056 | 0,270687905 | 0,089389513 | 3,028184145 | 0,002460281 | 0,049487438 |
| ENSG00000068971.14 | PPP2R5B | protein_coding | 2492,634455 | 0,270431954 | 0,078848357 | 3,429772833 | 0,000604087 | 0,017585098 |
| ENSG00000261455.1 | LINC01003 | lncRNA | 2591,938102 | 0,269684786 | 0,07471185 | 3,609665499 | 0,000306592 | 0,010517298 |
| ENSG00000166704.11 | ZNF606 | protein_coding | 1345,163656 | 0,268521348 | 0,082268892 | 3,263947541 | 0,001098715 | 0,027656621 |
| ENSG00000125848.10 | FLRT3 | protein_coding | 1768,351024 | 0,268084236 | 0,085375396 | 3,140064332 | 0,001689107 | 0,038097034 |
| ENSG00000160752.15 | FDPS | protein_coding | 27348,51391 | 0,2669465 | 0,060259116 | 4,429977048 | 9,42E-06 | 0,00067049 |

| | | | | | | | | |
|---|---|---|---|---|---|---|---|---|
| ENSG00000267673.6 | FDX2 | protein_coding | 1820,903804 | 0,266821728 | 0,085758819 | 3,111303665 | 0,001862633 | 0,041007577 |
| ENSG00000179178.11 | TMEM125 | protein_coding | 3733,656106 | 0,265793287 | 0,06158473 | 4,315895957 | 1,59E-05 | 0,001019443 |
| ENSG00000074842.8 | MYDGF | protein_coding | 19841,39805 | 0,263829088 | 0,058437711 | 4,514706048 | 6,34E-06 | 0,000480442 |
| ENSG00000086619.14 | ERO1B | protein_coding | 238593,6279 | 0,262885275 | 0,065692822 | 4,001735142 | 6,29E-05 | 0,003064483 |
| ENSG00000174276.7 | ZNHIT2 | protein_coding | 1129,562716 | 0,262739419 | 0,086783271 | 3,027535321 | 0,002465569 | 0,049548755 |
| ENSG00000196776.17 | CD47 | protein_coding | 5443,039338 | 0,262088896 | 0,064895201 | 4,038648367 | 5,38E-05 | 0,002709557 |
| ENSG00000183722.9 | LHFPL6 | protein_coding | 2135,409284 | 0,26089673 | 0,083004217 | 3,143174417 | 0,001671262 | 0,037866385 |
| ENSG00000132432.14 | SEC61G | protein_coding | 5862,978131 | 0,260529528 | 0,058538154 | 4,450593533 | 8,56E-06 | 0,000625155 |
| ENSG00000158747.15 | NBL1 | protein_coding | 1726,430558 | 0,259902624 | 0,084037844 | 3,092685534 | 0,001983542 | 0,043111837 |
| ENSG00000136383.7 | ALPK3 | protein_coding | 2446,522078 | 0,259376315 | 0,082208048 | 3,15512071 | 0,001604317 | 0,036544739 |
| ENSG00000157315.5 | TMED6 | protein_coding | 4363,32246 | 0,256469771 | 0,077591931 | 3,305366534 | 0,000948523 | 0,024690603 |
| ENSG00000100902.11 | PSMA6 | protein_coding | 13246,82061 | 0,256177849 | 0,066095421 | 3,875878898 | 0,000106241 | 0,004639937 |
| ENSG00000000457.14 | SCYL3 | protein_coding | 1397,664814 | 0,255037085 | 0,077521513 | 3,289887877 | 0,001002273 | 0,02575644 |
| ENSG00000267795.6 | SMIM22 | protein_coding | 4084,577386 | 0,253765249 | 0,065950031 | 3,847841234 | 0,000119163 | 0,005109701 |
| ENSG00000177380.14 | PPFIA3 | protein_coding | 1893,138901 | 0,250111694 | 0,071657269 | 3,490388313 | 0,000482319 | 0,014863223 |
| ENSG00000157349.17 | DDX19B | protein_coding | 2358,512691 | 0,250057224 | 0,066745659 | 3,74641928 | 0,000179377 | 0,007035385 |
| ENSG00000165644.11 | COMTD1 | protein_coding | 1883,845044 | 0,249129943 | 0,080134635 | 3,108892232 | 0,001877902 | 0,041302644 |
| ENSG00000147883.12 | CDKN2B | protein_coding | 7307,069677 | 0,248946096 | 0,066455917 | 3,746033571 | 0,000179653 | 0,007035385 |
| ENSG00000100219.16 | XBP1 | protein_coding | 29129,52729 | 0,248172349 | 0,055000637 | 4,512172274 | 6,42E-06 | 0,00048456 |
| ENSG00000023330.15 | ALAS1 | protein_coding | 6277,490393 | 0,248075621 | 0,058099673 | 4,269828193 | 1,96E-05 | 0,001198994 |
| ENSG00000114767.7 | RRP9 | protein_coding | 1522,341386 | 0,247697322 | 0,080122801 | 3,091471066 | 0,001991674 | 0,043203703 |
| ENSG00000110330.8 | BIRC2 | protein_coding | 4118,892648 | 0,246506721 | 0,072161442 | 3,416044827 | 0,000635378 | 0,018186767 |
| ENSG00000131966.14 | ACTR10 | protein_coding | 4696,913789 | 0,246467129 | 0,067341232 | 3,659973567 | 0,000252241 | 0,009089724 |
| ENSG00000179085.8 | DPM3 | protein_coding | 3646,892792 | 0,245266696 | 0,063953576 | 3,835073974 | 0,000125527 | 0,005280227 |
| ENSG00000044574.8 | HSPA5 | protein_coding | 117558,206 | 0,244778653 | 0,049621319 | 4,932933209 | 8,10E-07 | 8,07E-05 |
| ENSG00000148444.16 | COMMD3 | protein_coding | 2140,257982 | 0,244354908 | 0,079186557 | 3,085813007 | 0,002029964 | 0,043480147 |
| ENSG00000169567.13 | HINT1 | protein_coding | 24568,68348 | 0,243237899 | 0,065270226 | 3,726628748 | 0,000194058 | 0,007459585 |
| ENSG00000204525.17 | HLA-C | protein_coding | 6099,698931 | 0,242117672 | 0,069027794 | 3,507538883 | 0,000452272 | 0,014169001 |
| ENSG00000152219.5 | ARL14EP | protein_coding | 2745,839617 | 0,241887601 | 0,072141376 | 3,352966278 | 0,000799504 | 0,021812375 |
| ENSG00000131143.10 | COX4I1 | protein_coding | 31725,14233 | 0,239797638 | 0,058977506 | 4,065916854 | 4,78E-05 | 0,002473356 |
| ENSG00000105518.14 | TMEM205 | protein_coding | 3947,585152 | 0,239406572 | 0,069245238 | 3,457372345 | 0,000545471 | 0,01626561 |
| ENSG00000142188.17 | TMEM50B | protein_coding | 4896,689514 | 0,235469496 | 0,071066096 | 3,313387235 | 0,000921733 | 0,024221208 |
| ENSG00000134684.11 | YARS1 | protein_coding | 5430,752688 | 0,235067922 | 0,069617598 | 3,376558922 | 0,000733986 | 0,020402243 |
| ENSG00000066583.12 | ISOC1 | protein_coding | 5950,420915 | 0,234573422 | 0,067746236 | 3,46253069 | 0,000535121 | 0,016065244 |
| ENSG00000171735.19 | CAMTA1 | protein_coding | 5483,775281 | 0,234289429 | 0,073445016 | 3,189997676 | 0,001422739 | 0,033596085 |
| ENSG00000173915.16 | ATP5MD | protein_coding | 8302,15689 | 0,233194449 | 0,062613356 | 3,724356341 | 0,000195814 | 0,007508812 |

| | | | | | | | | |
|---|---|---|---|---|---|---|---|---|
| ENSG00000149273.15 | RPS3 | protein_coding | 114533,4232 | 0,232670117 | 0,055333901 | 4,204838475 | 2,61E-05 | 0,001525285 |
| ENSG00000204498.11 | NFKBIL1 | protein_coding | 2900,506116 | 0,231955302 | 0,070272051 | 3,300818729 | 0,000964031 | 0,025006049 |
| ENSG00000130558.20 | OLFM1 | protein_coding | 5413,518861 | 0,231299063 | 0,068232492 | 3,389866869 | 0,000699266 | 0,019634459 |
| ENSG00000107175.12 | CREB3 | protein_coding | 5055,969822 | 0,228977445 | 0,063059327 | 3,631143158 | 0,000282169 | 0,009894235 |
| ENSG00000154928.18 | EPHB1 | protein_coding | 4483,241428 | 0,228690268 | 0,067585252 | 3,383730345 | 0,000715082 | 0,019952413 |
| ENSG00000106245.11 | BUD31 | protein_coding | 8559,486028 | 0,228511423 | 0,058571629 | 3,901401211 | 9,56E-05 | 0,004249147 |
| ENSG00000101391.21 | CDK5RAP1 | protein_coding | 2877,636811 | 0,227744305 | 0,071792582 | 3,172253999 | 0,001512606 | 0,035044954 |
| ENSG00000100814.18 | CCNB1IP1 | protein_coding | 4119,885296 | 0,226690393 | 0,071418042 | 3,174133424 | 0,001502846 | 0,034928538 |
| ENSG00000198856.13 | OSTC | protein_coding | 9384,712674 | 0,224821429 | 0,069244986 | 3,246753914 | 0,001167293 | 0,028889842 |
| ENSG00000185608.8 | MRPL40 | protein_coding | 3062,759482 | 0,224747583 | 0,072827593 | 3,0860224 | 0,002028535 | 0,043480147 |
| ENSG00000169599.13 | NFU1 | protein_coding | 3532,771987 | 0,223885577 | 0,064631137 | 3,464051365 | 0,000532105 | 0,016018167 |
| ENSG00000186469.8 | GNG2 | protein_coding | 19330,4921 | 0,223361476 | 0,0688481 | 3,24426493 | 0,001177542 | 0,029064791 |
| ENSG00000172379.21 | ARNT2 | protein_coding | 18441,29208 | 0,221397061 | 0,063039736 | 3,512023921 | 0,000444708 | 0,013996598 |
| ENSG00000111678.11 | C12orf57 | protein_coding | 6746,550456 | 0,220606809 | 0,062934601 | 3,50533422 | 0,000456034 | 0,014231618 |
| ENSG00000123992.20 | DNPEP | protein_coding | 3660,674991 | 0,220452427 | 0,061114629 | 3,607195715 | 0,000309524 | 0,010545056 |
| ENSG00000108039.18 | XPNPEP1 | protein_coding | 2955,305142 | 0,21967589 | 0,067824873 | 3,238869144 | 0,001200046 | 0,029502468 |
| ENSG00000204592.9 | HLA-E | protein_coding | 24369,74839 | 0,219381553 | 0,063215729 | 3,470363388 | 0,000519755 | 0,015801053 |
| ENSG00000183421.12 | RIPK4 | protein_coding | 3472,842443 | 0,219263238 | 0,068472232 | 3,202221278 | 0,001363722 | 0,032549852 |
| ENSG00000198814.13 | GK | protein_coding | 5762,888826 | 0,218978586 | 0,068296995 | 3,20626972 | 0,001344679 | 0,032234414 |
| ENSG00000166136.16 | NDUFB8 | protein_coding | 15866,07353 | 0,218956828 | 0,066309421 | 3,302047047 | 0,00095982 | 0,024955312 |
| ENSG00000064102.15 | INTS13 | protein_coding | 3042,434707 | 0,218910199 | 0,064656308 | 3,385751592 | 0,000709836 | 0,019855667 |
| ENSG00000233276.7 | GPX1 | protein_coding | 8372,563845 | 0,2180289 | 0,063562787 | 3,430134356 | 0,000603282 | 0,017585098 |
| ENSG00000165175.16 | MID1IP1 | protein_coding | 8122,743102 | 0,217911174 | 0,0627508 | 3,47264378 | 0,000515359 | 0,015706378 |
| ENSG00000160050.15 | CCDC28B | protein_coding | 3343,427147 | 0,217347664 | 0,063764296 | 3,408610711 | 0,000652946 | 0,01859341 |
| ENSG00000173486.14 | FKBP2 | protein_coding | 8060,737938 | 0,215026094 | 0,061751105 | 3,482141627 | 0,000497421 | 0,015243666 |
| ENSG00000140105.18 | WARS1 | protein_coding | 9805,099727 | 0,212894805 | 0,053006162 | 4,016416128 | 5,91E-05 | 0,002918354 |
| ENSG00000128590.5 | DNAJB9 | protein_coding | 9086,304721 | 0,211992747 | 0,069333323 | 3,05758814 | 0,00223126 | 0,046053042 |
| ENSG00000064490.14 | RFXANK | protein_coding | 2440,149364 | 0,211866069 | 0,068267428 | 3,103472269 | 0,001912642 | 0,0418173 |
| ENSG00000133731.10 | IMPA1 | protein_coding | 4517,965251 | 0,211549365 | 0,061451886 | 3,442520306 | 0,000576321 | 0,016933693 |
| ENSG00000113811.11 | SELENOK | protein_coding | 9028,275679 | 0,208247337 | 0,058349378 | 3,568972675 | 0,000358384 | 0,011835224 |
| ENSG00000079950.14 | STX7 | protein_coding | 6256,56921 | 0,20741811 | 0,068543014 | 3,026101365 | 0,002477292 | 0,049694081 |
| ENSG00000106803.10 | SEC61B | protein_coding | 10203,37945 | 0,207404373 | 0,064409407 | 3,220094418 | 0,001281484 | 0,031192642 |
| ENSG00000002586.20 | CD99 | protein_coding | 57412,32011 | 0,206755807 | 0,051977852 | 3,977767428 | 6,96E-05 | 0,003281883 |
| ENSG00000076604.15 | TRAF4 | protein_coding | 9956,274443 | 0,206606597 | 0,054832516 | 3,767957608 | 0,000164589 | 0,006561601 |
| ENSG00000056998.20 | GYG2 | protein_coding | 4358,576081 | 0,205401 | 0,061303239 | 3,350573369 | 0,000806444 | 0,02192658 |
| ENSG00000134686.18 | PHC2 | protein_coding | 19127,56758 | 0,205396484 | 0,051565254 | 3,983234228 | 6,80E-05 | 0,003248032 |

| | | | | | | | | |
|---|---|---|---|---|---|---|---|---|
| ENSG00000146085.8 | MMUT | protein_coding | 12505,8406 | 0,204540824 | 0,063808096 | 3,205562266 | 0,001347989 | 0,032278788 |
| ENSG00000173545.5 | ZNF622 | protein_coding | 3986,315463 | 0,202927595 | 0,063116275 | 3,215138983 | 0,001303814 | 0,031562565 |
| ENSG00000173442.13 | EHBP1L1 | protein_coding | 4670,112163 | 0,202088439 | 0,066227801 | 3,051413989 | 0,002277663 | 0,046705811 |
| ENSG00000126768.12 | TIMM17B | protein_coding | 20701,55909 | 0,202056064 | 0,060998798 | 3,312459751 | 0,000924794 | 0,024272834 |
| ENSG00000112695.11 | COX7A2 | protein_coding | 14437,85544 | 0,201934127 | 0,065759644 | 3,070791083 | 0,002134925 | 0,044763358 |
| ENSG00000114850.7 | SSR3 | protein_coding | 13335,85448 | 0,200600485 | 0,055278497 | 3,628906263 | 0,000284625 | 0,009964562 |
| ENSG00000031698.13 | SARS1 | protein_coding | 12795,89041 | 0,200015623 | 0,055610225 | 3,596741859 | 0,000322228 | 0,010842419 |
| ENSG00000179218.14 | CALR | protein_coding | 139345,6033 | 0,200011951 | 0,05354944 | 3,735089502 | 0,000187648 | 0,007284051 |
| ENSG00000160326.14 | SLC2A6 | protein_coding | 5490,479028 | 0,199819211 | 0,062921262 | 3,175702514 | 0,001494741 | 0,034813316 |
| ENSG00000051108.15 | HERPUD1 | protein_coding | 7412,607451 | 0,199671602 | 0,060990442 | 3,273817907 | 0,00106105 | 0,026953828 |
| ENSG00000117448.14 | AKR1A1 | protein_coding | 7532,71311 | 0,197332943 | 0,062422571 | 3,161243431 | 0,001570971 | 0,035982728 |
| ENSG00000198931.11 | APRT | protein_coding | 5014,63437 | 0,195372761 | 0,059569648 | 3,279736721 | 0,00103904 | 0,026516493 |
| ENSG00000229117.9 | RPL41 | protein_coding | 52569,27996 | 0,193358838 | 0,057429328 | 3,366900594 | 0,000760181 | 0,020998454 |
| ENSG00000161011.20 | SQSTM1 | protein_coding | 18749,48339 | 0,191100982 | 0,062700334 | 3,047846326 | 0,002304878 | 0,047176432 |
| ENSG00000197879.17 | MYO1C | protein_coding | 12258,91096 | 0,190662534 | 0,060959201 | 3,127707244 | 0,001761756 | 0,039255394 |
| ENSG00000010256.11 | UQCRC1 | protein_coding | 23091,81502 | 0,190435899 | 0,057340378 | 3,321148302 | 0,000896479 | 0,02375508 |
| ENSG00000163682.17 | RPL9 | protein_coding | 27800,69422 | 0,189872692 | 0,061498508 | 3,087435728 | 0,002018914 | 0,043453791 |
| ENSG00000189060.5 | H1-0 | protein_coding | 8009,581127 | 0,188302631 | 0,059179756 | 3,181875748 | 0,001463246 | 0,034369189 |
| ENSG00000163479.14 | SSR2 | protein_coding | 25012,08947 | 0,1869375 | 0,055926587 | 3,34255152 | 0,000830119 | 0,022426401 |
| ENSG00000243678.12 | NME2 | protein_coding | 16232,02583 | 0,186215567 | 0,058554177 | 3,180226865 | 0,001471598 | 0,034465103 |
| ENSG00000178982.10 | EIF3K | protein_coding | 9388,930701 | 0,186155571 | 0,058125855 | 3,202629369 | 0,001361791 | 0,032538868 |
| ENSG00000170315.14 | UBB | protein_coding | 122805,985 | 0,181537277 | 0,05394844 | 3,365014425 | 0,000765397 | 0,021063638 |
| ENSG00000159128.15 | IFNGR2 | protein_coding | 4890,028148 | 0,178445267 | 0,058089092 | 3,071923841 | 0,00212684 | 0,044647493 |
| ENSG00000186635.14 | ARAP1 | protein_coding | 8578,169941 | 0,173662258 | 0,055898018 | 3,106769498 | 0,001891438 | 0,041559048 |
| ENSG00000135148.12 | TRAFD1 | protein_coding | 6341,379473 | 0,172178219 | 0,056412225 | 3,052143766 | 0,002272132 | 0,046678926 |
| ENSG00000197756.10 | RPL37A | protein_coding | 45256,88524 | 0,170463633 | 0,0556976 | 3,060520224 | 0,002209528 | 0,045775304 |
| ENSG00000111057.11 | KRT18 | protein_coding | 11736,06651 | 0,169091105 | 0,05391384 | 3,136320944 | 0,001710819 | 0,038333247 |
| ENSG00000118495.20 | PLAGL1 | protein_coding | 52302,63328 | -0,15189634 | 0,050237151 | -3,023585853 | 0,002497981 | 0,049973179 |
| ENSG00000163069.13 | SGCB | protein_coding | 12601,28338 | -0,156488006 | 0,05148186 | -3,039672732 | 0,002368353 | 0,047987353 |
| ENSG00000140263.15 | SORD | protein_coding | 10839,64802 | -0,159233171 | 0,052295739 | -3,044859393 | 0,002327891 | 0,047603437 |
| ENSG00000116237.16 | ICMT | protein_coding | 33910,98533 | -0,166192337 | 0,050923351 | -3,263578152 | 0,001100148 | 0,027661231 |
| ENSG00000099194.6 | SCD | protein_coding | 282657,4926 | -0,169001897 | 0,051761373 | -3,265019592 | 0,001094565 | 0,027583538 |
| ENSG00000149485.19 | FADS1 | protein_coding | 47926,59066 | -0,169700123 | 0,050058028 | -3,390068075 | 0,000698753 | 0,019634459 |
| ENSG00000132405.19 | TBC1D14 | protein_coding | 9752,093771 | -0,171172308 | 0,055416596 | -3,088827529 | 0,002009481 | 0,043381047 |
| ENSG00000175329.13 | ISX | protein_coding | 15118,98762 | -0,171303193 | 0,056479521 | -3,033014266 | 0,002421242 | 0,048790884 |
| ENSG00000100201.23 | DDX17 | protein_coding | 21263,92406 | -0,172588074 | 0,055597894 | -3,10421963 | 0,001907817 | 0,041794404 |

| | | | | | | | | |
|---|---|---|---|---|---|---|---|---|
| ENSG00000018236.15 | CNTN1 | protein_coding | 53151,69406 | -0,173023916 | 0,052114169 | -3,320093569 | 0,000899873 | 0,023788036 |
| ENSG00000107164.16 | FUBP3 | protein_coding | 7898,884055 | -0,173293556 | 0,057164852 | -3,031470363 | 0,002433658 | 0,048996469 |
| ENSG00000088812.18 | ATRN | protein_coding | 13986,50426 | -0,174112218 | 0,056412264 | -3,08642492 | 0,002025791 | 0,043480147 |
| ENSG00000092964.18 | DPYSL2 | protein_coding | 20910,11667 | -0,174126822 | 0,056496129 | -3,082101837 | 0,002055445 | 0,043813851 |
| ENSG00000185896.11 | LAMP1 | protein_coding | 49922,16149 | -0,175451445 | 0,056869461 | -3,085161015 | 0,00203442 | 0,043533437 |
| ENSG00000122952.17 | ZWINT | protein_coding | 9831,277368 | -0,175595712 | 0,057147276 | -3,072687367 | 0,002121406 | 0,044647493 |
| ENSG00000159267.15 | HLCS | protein_coding | 4687,349894 | -0,177270099 | 0,058278296 | -3,041785928 | 0,002351791 | 0,047783034 |
| ENSG00000146918.20 | NCAPG2 | protein_coding | 8878,403553 | -0,180475504 | 0,056798438 | -3,177473039 | 0,001485645 | 0,034711068 |
| ENSG00000139354.11 | GAS2L3 | protein_coding | 7643,359306 | -0,180652365 | 0,059386099 | -3,041997516 | 0,002350138 | 0,047783034 |
| ENSG00000166197.17 | NOLC1 | protein_coding | 8393,561076 | -0,181550781 | 0,059457026 | -3,053479027 | 0,002262045 | 0,046588476 |
| ENSG00000169679.15 | BUB1 | protein_coding | 5626,588917 | -0,181585086 | 0,056531059 | -3,212129564 | 0,001317549 | 0,031790729 |
| ENSG00000111684.11 | LPCAT3 | protein_coding | 7875,384131 | -0,18488964 | 0,056000083 | -3,301595808 | 0,000961365 | 0,024966151 |
| ENSG00000138160.7 | KIF11 | protein_coding | 8115,515704 | -0,185605762 | 0,060734308 | -3,056028249 | 0,002242901 | 0,04625017 |
| ENSG00000169116.11 | PARM1 | protein_coding | 7960,940559 | -0,185792669 | 0,055351616 | -3,356589765 | 0,000789101 | 0,021581766 |
| ENSG00000133657.16 | ATP13A3 | protein_coding | 26405,29703 | -0,186204285 | 0,052918141 | -3,518723075 | 0,000433629 | 0,013745669 |
| ENSG00000163013.12 | FBXO41 | protein_coding | 4622,760935 | -0,188883498 | 0,059598408 | -3,169270864 | 0,001528219 | 0,035369638 |
| ENSG00000031003.10 | FAM13B | protein_coding | 8628,506019 | -0,18915905 | 0,054892318 | -3,446002211 | 0,000568946 | 0,016766689 |
| ENSG00000101868.13 | POLA1 | protein_coding | 14182,69853 | -0,189446657 | 0,052040108 | -3,640397069 | 0,000272218 | 0,009652394 |
| ENSG00000101126.18 | ADNP | protein_coding | 15220,4967 | -0,190998696 | 0,060845121 | -3,139096315 | 0,001694697 | 0,038152207 |
| ENSG00000160710.17 | ADAR | protein_coding | 14529,61948 | -0,191158925 | 0,059201729 | -3,228941612 | 0,001242493 | 0,030444509 |
| ENSG00000021300.14 | PLEKHB1 | protein_coding | 19266,69842 | -0,19135327 | 0,055214004 | -3,465665535 | 0,000528921 | 0,01596576 |
| ENSG00000141232.5 | TOB1 | protein_coding | 3220,237516 | -0,19218526 | 0,062685538 | -3,065862812 | 0,002170429 | 0,045219309 |
| ENSG00000152291.14 | TGOLN2 | protein_coding | 23139,49888 | -0,192561572 | 0,057463797 | -3,351006748 | 0,000805183 | 0,02192658 |
| ENSG00000132561.14 | MATN2 | protein_coding | 11316,30029 | -0,192611854 | 0,054958653 | -3,504668357 | 0,000457176 | 0,014247162 |
| ENSG00000138796.17 | HADH | protein_coding | 97302,08981 | -0,19265886 | 0,04932622 | -3,905810319 | 9,39E-05 | 0,004201488 |
| ENSG00000180182.11 | MED14 | protein_coding | 13025,62146 | -0,194070953 | 0,062269797 | -3,116614502 | 0,001829406 | 0,040437005 |
| ENSG00000140526.18 | ABHD2 | protein_coding | 23994,88041 | -0,194308316 | 0,049423681 | -3,931482118 | 8,44E-05 | 0,003867413 |
| ENSG00000160007.19 | ARHGAP35 | protein_coding | 17713,15329 | -0,196032589 | 0,060886815 | -3,219622975 | 0,001283593 | 0,031209645 |
| ENSG00000103381.12 | CPPED1 | protein_coding | 4124,631139 | -0,196157728 | 0,061318996 | -3,198971602 | 0,001379188 | 0,03284812 |
| ENSG00000139697.14 | SBNO1 | protein_coding | 6735,293394 | -0,196629643 | 0,058979245 | -3,333878623 | 0,00085644 | 0,022997083 |
| ENSG00000073614.13 | KDM5A | protein_coding | 8581,45976 | -0,196940333 | 0,059729066 | -3,297227736 | 0,000976443 | 0,025209774 |
| ENSG00000152952.12 | PLOD2 | protein_coding | 15133,56259 | -0,197938965 | 0,05370609 | -3,685596291 | 0,000228168 | 0,008372207 |
| ENSG00000156802.13 | ATAD2 | protein_coding | 13547,12326 | -0,198308366 | 0,055853997 | -3,550477618 | 0,000384533 | 0,012539112 |
| ENSG00000196123.13 | KIAA0895L | protein_coding | 3483,394274 | -0,198470783 | 0,064478211 | -3,078106234 | 0,002083206 | 0,044277641 |
| ENSG00000134690.11 | CDCA8 | protein_coding | 6716,111178 | -0,199278088 | 0,057752399 | -3,450559493 | 0,000559426 | 0,016592301 |
| ENSG00000173698.18 | ADGRG2 | protein_coding | 4892,378957 | -0,199282989 | 0,059883783 | -3,327828994 | 0,000875256 | 0,023388779 |

| | | | | | | | | |
|---|---|---|---|---|---|---|---|---|
| ENSG00000070018.9 | LRP6 | protein_coding | 4575,746053 | -0,199444012 | 0,059658544 | -3,343092168 | 0,000828504 | 0,022410115 |
| ENSG00000166257.9 | SCN3B | protein_coding | 3081,240815 | -0,199732033 | 0,065716814 | -3,039283582 | 0,002371415 | 0,048005427 |
| ENSG00000154319.16 | FAM167A | protein_coding | 14275,45074 | -0,200433143 | 0,056061695 | -3,575224456 | 0,000349928 | 0,011607942 |
| ENSG00000156011.17 | PSD3 | protein_coding | 13099,07048 | -0,202029531 | 0,054589572 | -3,700881389 | 0,000214852 | 0,008003052 |
| ENSG00000173542.9 | MOB1B | protein_coding | 26219,23826 | -0,202181356 | 0,053480417 | -3,78047457 | 0,00015653 | 0,006308518 |
| ENSG00000166851.15 | PLK1 | protein_coding | 6767,267601 | -0,202535311 | 0,063798568 | -3,174605928 | 0,001500401 | 0,034908387 |
| ENSG00000072364.13 | AFF4 | protein_coding | 13701,10969 | -0,202614482 | 0,054856363 | -3,693545646 | 0,000221149 | 0,00815523 |
| ENSG00000163781.14 | TOPBP1 | protein_coding | 9756,907767 | -0,202876497 | 0,060570583 | -3,349422893 | 0,000809801 | 0,021966056 |
| ENSG00000163346.17 | PBXIP1 | protein_coding | 6540,547311 | -0,203260752 | 0,06470312 | -3,141436657 | 0,001681212 | 0,037957643 |
| ENSG00000176406.23 | RIMS2 | protein_coding | 3553,002636 | -0,203366871 | 0,063071546 | -3,224383791 | 0,001262441 | 0,030864935 |
| ENSG00000160179.19 | ABCG1 | protein_coding | 6153,688854 | -0,203785755 | 0,055160522 | -3,69441307 | 0,000220395 | 0,008148295 |
| ENSG00000124486.13 | USP9X | protein_coding | 41497,1855 | -0,204204997 | 0,056388135 | -3,621417818 | 0,000292993 | 0,010113511 |
| ENSG00000150712.11 | MTMR12 | protein_coding | 3665,095832 | -0,206104317 | 0,061784502 | -3,335857877 | 0,000850366 | 0,02286173 |
| ENSG00000157240.4 | FZD1 | protein_coding | 9950,749271 | -0,207057267 | 0,054076485 | -3,828970498 | 0,00012868 | 0,005372044 |
| ENSG00000100320.23 | RBFOX2 | protein_coding | 12786,27989 | -0,207095614 | 0,052839065 | -3,919365647 | 8,88E-05 | 0,004017174 |
| ENSG00000132466.19 | ANKRD17 | protein_coding | 12405,26328 | -0,207351898 | 0,062870928 | -3,298056923 | 0,000973564 | 0,025164809 |
| ENSG00000115677.17 | HDLBP | protein_coding | 45430,66927 | -0,207648734 | 0,056102308 | -3,701251203 | 0,000214539 | 0,008003052 |
| ENSG00000184445.12 | KNTC1 | protein_coding | 5245,257541 | -0,207889392 | 0,062531955 | -3,324530481 | 0,000885676 | 0,023581777 |
| ENSG00000011426.11 | ANLN | protein_coding | 8486,077351 | -0,208226829 | 0,057127977 | -3,644918633 | 0,000267476 | 0,009499491 |
| ENSG00000137726.17 | FXYD6 | protein_coding | 11405,48542 | -0,208362671 | 0,061577914 | -3,383724099 | 0,000715098 | 0,019952413 |
| ENSG00000109066.14 | TMEM104 | protein_coding | 5267,879847 | -0,209684907 | 0,068969604 | -3,04025099 | 0,002363811 | 0,047939207 |
| ENSG00000127483.19 | HP1BP3 | protein_coding | 11118,13279 | -0,209769517 | 0,054001792 | -3,884491746 | 0,000102544 | 0,004510721 |
| ENSG00000140718.21 | FTO | protein_coding | 4659,57336 | -0,209902755 | 0,062471764 | -3,359962043 | 0,000779532 | 0,021399406 |
| ENSG00000151835.16 | SACS | protein_coding | 3728,810802 | -0,209976544 | 0,066813218 | -3,142739555 | 0,001673747 | 0,037866385 |
| ENSG00000112290.13 | WASF1 | protein_coding | 4402,185519 | -0,21069083 | 0,068211178 | -3,088802097 | 0,002009653 | 0,043381047 |
| ENSG00000186350.12 | RXRA | protein_coding | 20193,00512 | -0,211110821 | 0,063170405 | -3,341926018 | 0,000831992 | 0,022449588 |
| ENSG00000144043.12 | TEX261 | protein_coding | 12439,03335 | -0,211280865 | 0,053053757 | -3,982392156 | 6,82E-05 | 0,003248032 |
| ENSG00000182534.13 | MXRA7 | protein_coding | 32032,56148 | -0,212276366 | 0,052067188 | -4,076970085 | 4,56E-05 | 0,002364236 |
| ENSG00000139116.19 | KIF21A | protein_coding | 9236,287615 | -0,212330305 | 0,059036881 | -3,596570518 | 0,00032244 | 0,010842419 |
| ENSG00000170571.12 | EMB | protein_coding | 12953,15447 | -0,212390349 | 0,058859244 | -3,608445085 | 0,000308038 | 0,010534223 |
| ENSG00000147526.20 | TACC1 | protein_coding | 4373,708701 | -0,21248351 | 0,065801705 | -3,229149018 | 0,001241592 | 0,030444509 |
| ENSG00000080345.18 | RIF1 | protein_coding | 9117,545616 | -0,212649539 | 0,060155799 | -3,534979869 | 0,000407806 | 0,013115001 |
| ENSG00000006747.15 | SCIN | protein_coding | 8098,087689 | -0,212723031 | 0,054786572 | -3,882758576 | 0,000103278 | 0,00453399 |
| ENSG00000175497.17 | DPP10 | protein_coding | 7873,053978 | -0,212751255 | 0,05742023 | -3,70516203 | 0,000211256 | 0,007931509 |
| ENSG00000103994.18 | ZNF106 | protein_coding | 6124,932707 | -0,213359018 | 0,069588508 | -3,06600936 | 0,002169365 | 0,045219309 |
| ENSG00000054965.11 | FAM168A | protein_coding | 7578,558794 | -0,213640356 | 0,068475715 | -3,119943428 | 0,001808858 | 0,040022787 |

| | | | | | | | | |
|---|---|---|---|---|---|---|---|---|
| ENSG00000185352.8 | HS6ST3 | protein_coding | 14265,75654 | -0,213737444 | 0,061119854 | -3,497021494 | 0,000470484 | 0,01457973 |
| ENSG00000040531.16 | CTNS | protein_coding | 3475,444351 | -0,213819395 | 0,065910621 | -3,244081 | 0,001178302 | 0,029064791 |
| ENSG00000116005.12 | PCYOX1 | protein_coding | 29403,0551 | -0,214442241 | 0,050472726 | -4,24867564 | 2,15E-05 | 0,001296439 |
| ENSG00000068078.20 | FGFR3 | protein_coding | 5963,772821 | -0,214470424 | 0,069878618 | -3,069185266 | 0,002146434 | 0,044888475 |
| ENSG00000179295.18 | PTPN11 | protein_coding | 21024,98816 | -0,214625404 | 0,054400009 | -3,945319263 | 7,97E-05 | 0,003681218 |
| ENSG00000010404.18 | IDS | protein_coding | 22783,56242 | -0,214631345 | 0,054904577 | -3,909170371 | 9,26E-05 | 0,004181978 |
| ENSG00000129116.19 | PALLD | protein_coding | 12139,32542 | -0,215038609 | 0,062990209 | -3,413841819 | 0,000640538 | 0,01828714 |
| ENSG00000178202.13 | POGLUT3 | protein_coding | 4374,427391 | -0,215628212 | 0,06264303 | -3,44217404 | 0,000577059 | 0,016933693 |
| ENSG00000082701.17 | GSK3B | protein_coding | 7612,465865 | -0,216140303 | 0,067633834 | -3,195742276 | 0,001394716 | 0,033075556 |
| ENSG00000137216.19 | TMEM63B | protein_coding | 6114,615581 | -0,216220414 | 0,071029946 | -3,04407401 | 0,002333977 | 0,047639832 |
| ENSG00000165272.16 | AQP3 | protein_coding | 3107,257049 | -0,216745087 | 0,070008422 | -3,095985915 | 0,001961597 | 0,04271879 |
| ENSG00000175426.11 | PCSK1 | protein_coding | 63559,06669 | -0,217918622 | 0,048502373 | -4,492947644 | 7,02E-06 | 0,000528236 |
| ENSG00000088325.16 | TPX2 | protein_coding | 15843,15038 | -0,218395838 | 0,062590751 | -3,489266945 | 0,000484347 | 0,014885747 |
| ENSG00000102781.14 | KATNAL1 | protein_coding | 4986,238271 | -0,218850873 | 0,059556841 | -3,674655496 | 0,000238171 | 0,008667371 |
| ENSG00000162636.16 | FAM102B | protein_coding | 2315,033646 | -0,219420875 | 0,071509672 | -3,068408362 | 0,002152023 | 0,044926508 |
| ENSG00000196090.12 | PTPRT | protein_coding | 15773,44452 | -0,219616188 | 0,063440197 | -3,461782868 | 0,00053661 | 0,016073055 |
| ENSG00000011454.18 | RABGAP1 | protein_coding | 6043,920452 | -0,219790592 | 0,06059671 | -3,627104383 | 0,000286617 | 0,009986925 |
| ENSG00000087460.27 | GNAS | protein_coding | 436063,7203 | -0,222250951 | 0,063083952 | -3,523098081 | 0,000426533 | 0,013598674 |
| ENSG00000168724.17 | DNAJC21 | protein_coding | 4447,93467 | -0,222560178 | 0,068263682 | -3,260301428 | 0,001112939 | 0,027761985 |
| ENSG00000129493.15 | HEATR5A | protein_coding | 4093,19531 | -0,22267171 | 0,067551504 | -3,29632497 | 0,000979586 | 0,025261453 |
| ENSG00000166340.17 | TPP1 | protein_coding | 16568,85109 | -0,222857902 | 0,072491661 | -3,07425571 | 0,002110285 | 0,044553587 |
| ENSG00000139645.10 | ANKRD52 | protein_coding | 10396,24898 | -0,222934023 | 0,073235308 | -3,044078453 | 0,002333943 | 0,047639832 |
| ENSG00000165458.14 | INPPL1 | protein_coding | 10319,66547 | -0,223499078 | 0,064683126 | -3,455291872 | 0,000549697 | 0,016347587 |
| ENSG00000165240.20 | ATP7A | protein_coding | 2024,757386 | -0,223646061 | 0,073514942 | -3,042185085 | 0,002348674 | 0,047783034 |
| ENSG00000006607.14 | FARP2 | protein_coding | 8451,292641 | -0,223766423 | 0,071350253 | -3,136168575 | 0,001711708 | 0,038333247 |
| ENSG00000134909.18 | ARHGAP32 | protein_coding | 2724,777459 | -0,224056402 | 0,070385927 | -3,183255694 | 0,001456289 | 0,034242148 |
| ENSG00000186951.16 | PPARA | protein_coding | 5058,414864 | -0,224087531 | 0,069548156 | -3,222048482 | 0,001272776 | 0,031083268 |
| ENSG00000082397.17 | EPB41L3 | protein_coding | 9059,086248 | -0,224991322 | 0,064059745 | -3,512210727 | 0,000444395 | 0,013996598 |
| ENSG00000120256.11 | LRP11 | protein_coding | 25481,94706 | -0,225398092 | 0,056151939 | -4,014074978 | 5,97E-05 | 0,002940904 |
| ENSG00000118762.8 | PKD2 | protein_coding | 7758,489233 | -0,225794776 | 0,060902078 | -3,707505292 | 0,000209311 | 0,007889636 |
| ENSG00000198420.10 | TCAF1 | protein_coding | 18616,22738 | -0,225799479 | 0,051420678 | -4,391219383 | 1,13E-05 | 0,000767376 |
| ENSG00000169174.11 | PCSK9 | protein_coding | 17222,89864 | -0,22622531 | 0,064279974 | -3,519374636 | 0,000432565 | 0,013731625 |
| ENSG00000175155.11 | YPEL2 | protein_coding | 2939,081442 | -0,226635468 | 0,070740717 | -3,203748513 | 0,001356509 | 0,032447698 |
| ENSG00000188677.15 | PARVB | protein_coding | 5884,371502 | -0,226648454 | 0,064005798 | -3,54106129 | 0,000398521 | 0,012891338 |
| ENSG00000147202.18 | DIAPH2 | protein_coding | 2847,475642 | -0,227022674 | 0,071387689 | -3,180137603 | 0,001472051 | 0,034465103 |
| ENSG00000157540.22 | DYRK1A | protein_coding | 3369,547338 | -0,228208134 | 0,070896317 | -3,218899718 | 0,001286835 | 0,031254124 |

| | | | | | | | | |
|---|---|---|---|---|---|---|---|---|
| ENSG00000060749.16 | QSER1 | protein_coding | 3812,893915 | -0,228706241 | 0,071614122 | -3,193591377 | 0,001405149 | 0,033287278 |
| ENSG00000054523.18 | KIF1B | protein_coding | 7710,502089 | -0,229303223 | 0,063932449 | -3,586648498 | 0,000334955 | 0,011229124 |
| ENSG00000183044.12 | ABAT | protein_coding | 15967,4243 | -0,230247101 | 0,055382039 | -4,157432712 | 3,22E-05 | 0,001793733 |
| ENSG00000169903.7 | TM4SF4 | protein_coding | 4233,741119 | -0,230650168 | 0,065908226 | -3,499565709 | 0,000466017 | 0,014461551 |
| ENSG00000083168.11 | KAT6A | protein_coding | 7916,889921 | -0,230781292 | 0,068522974 | -3,367940392 | 0,00075732 | 0,020945569 |
| ENSG00000092068.20 | SLC7A8 | protein_coding | 34359,94325 | -0,231226087 | 0,070424671 | -3,283310839 | 0,001025955 | 0,026283212 |
| ENSG00000168772.11 | CXXC4 | protein_coding | 2528,606591 | -0,231497551 | 0,076058629 | -3,043672432 | 0,002337095 | 0,047659498 |
| ENSG00000175414.7 | ARL10 | protein_coding | 5401,821234 | -0,232100199 | 0,072303029 | -3,210103394 | 0,001326872 | 0,03191128 |
| ENSG00000091136.15 | LAMB1 | protein_coding | 20945,77371 | -0,232353232 | 0,055691082 | -4,17218023 | 3,02E-05 | 0,001698572 |
| ENSG00000143970.17 | ASXL2 | protein_coding | 3976,204622 | -0,233094443 | 0,06652215 | -3,504012486 | 0,000458304 | 0,01426221 |
| ENSG00000133812.16 | SBF2 | protein_coding | 3103,379904 | -0,233120009 | 0,067441313 | -3,456635082 | 0,000546965 | 0,01628822 |
| ENSG00000115904.14 | SOS1 | protein_coding | 5428,468851 | -0,234106433 | 0,059907538 | -3,907795931 | 9,31E-05 | 0,004187611 |
| ENSG00000158856.18 | DMTN | protein_coding | 19818,97936 | -0,234390849 | 0,077450895 | -3,026315581 | 0,002475538 | 0,049694081 |
| ENSG00000067141.17 | NEO1 | protein_coding | 25702,18937 | -0,23439918 | 0,064247531 | -3,648376454 | 0,000263903 | 0,009402757 |
| ENSG00000102048.16 | ASB9 | protein_coding | 2874,536707 | -0,234887446 | 0,064534785 | -3,639702931 | 0,000272953 | 0,009662964 |
| ENSG00000143776.18 | CDC42BPA | protein_coding | 10365,10418 | -0,235675669 | 0,066382275 | -3,550280094 | 0,000384821 | 0,012539112 |
| ENSG00000135272.12 | MDFIC | protein_coding | 4257,732106 | -0,235740441 | 0,073507274 | -3,207035553 | 0,001341104 | 0,032183592 |
| ENSG00000176463.14 | SLCO3A1 | protein_coding | 5433,458728 | -0,236091734 | 0,074622997 | -3,163793251 | 0,001557274 | 0,035780104 |
| ENSG00000105926.16 | MPP6 | protein_coding | 7802,811759 | -0,236182109 | 0,072388282 | -3,262711896 | 0,001103516 | 0,02771442 |
| ENSG00000167258.15 | CDK12 | protein_coding | 6151,385164 | -0,236397352 | 0,068254562 | -3,46346598 | 0,000533264 | 0,01603125 |
| ENSG00000183688.4 | RFLNB | protein_coding | 6179,173205 | -0,237080874 | 0,065254294 | -3,633184234 | 0,000279945 | 0,00987344 |
| ENSG00000124177.15 | CHD6 | protein_coding | 6896,832896 | -0,237129594 | 0,067491288 | -3,513484495 | 0,00044227 | 0,013975561 |
| ENSG00000099204.20 | ABLIM1 | protein_coding | 3228,065168 | -0,237187435 | 0,063889689 | -3,712452507 | 0,000205261 | 0,007786654 |
| ENSG00000074755.15 | ZZEF1 | protein_coding | 4675,326737 | -0,237436053 | 0,069583461 | -3,412248368 | 0,000644294 | 0,018370676 |
| ENSG00000183337.17 | BCOR | protein_coding | 6529,437167 | -0,237449957 | 0,073160841 | -3,245588145 | 0,001172083 | 0,02897598 |
| ENSG00000009954.11 | BAZ1B | protein_coding | 12110,54828 | -0,237485305 | 0,077780295 | -3,053283672 | 0,002263519 | 0,046588476 |
| ENSG00000133958.13 | UNC79 | protein_coding | 3585,568644 | -0,238578853 | 0,070109573 | -3,402942592 | 0,000666643 | 0,018886223 |
| ENSG00000170145.5 | SIK2 | protein_coding | 5382,600062 | -0,238928525 | 0,077301345 | -3,090871518 | 0,0019957 | 0,04324863 |
| ENSG00000074276.10 | CDHR2 | protein_coding | 8269,199233 | -0,239185031 | 0,072926417 | -3,279813274 | 0,001038758 | 0,026516493 |
| ENSG00000086758.16 | HUWE1 | protein_coding | 23467,93697 | -0,239492939 | 0,077299182 | -3,098259665 | 0,001946608 | 0,042434134 |
| ENSG00000140564.12 | FURIN | protein_coding | 19257,95704 | -0,239658032 | 0,074473665 | -3,218023887 | 0,001290771 | 0,031315342 |
| ENSG00000184661.14 | CDCA2 | protein_coding | 2042,836525 | -0,239737325 | 0,070595714 | -3,395918961 | 0,000683986 | 0,019303413 |
| ENSG00000090863.12 | GLG1 | protein_coding | 20568,64381 | -0,239822273 | 0,065636439 | -3,653797728 | 0,00025839 | 0,009255943 |
| ENSG00000163684.12 | RPP14 | protein_coding | 3092,817997 | -0,24013828 | 0,07226898 | -3,322840294 | 0,000891059 | 0,023696604 |
| ENSG00000174197.16 | MGA | protein_coding | 6933,953251 | -0,240215998 | 0,070519375 | -3,406382955 | 0,000658298 | 0,018673711 |
| ENSG00000144406.19 | UNC80 | protein_coding | 5279,972963 | -0,240654886 | 0,071508464 | -3,365404228 | 0,000764316 | 0,021060093 |

| | | | | | | | | |
|---|---|---|---|---|---|---|---|---|
| ENSG00000189057.11 | FAM111B | protein_coding | 7071,076669 | -0,240686682 | 0,058441213 | -4,118440917 | 3,81E-05 | 0,002043544 |
| ENSG00000004660.15 | CAMKK1 | protein_coding | 2057,606275 | -0,240912747 | 0,074141291 | -3,249373536 | 0,001156595 | 0,028721458 |
| ENSG00000107518.18 | ATRNL1 | protein_coding | 4208,150067 | -0,24105432 | 0,062882066 | -3,833435135 | 0,000126366 | 0,005305459 |
| ENSG00000136861.18 | CDK5RAP2 | protein_coding | 4883,46341 | -0,241209067 | 0,072669643 | -3,319254865 | 0,00090258 | 0,023802726 |
| ENSG00000084674.15 | APOB | protein_coding | 9146,445642 | -0,241294143 | 0,070815623 | -3,407357482 | 0,000655952 | 0,018654992 |
| ENSG00000107263.18 | RAPGEF1 | protein_coding | 7300,294405 | -0,241630099 | 0,073894666 | -3,26992614 | 0,001075756 | 0,0272648 |
| ENSG00000165507.9 | DEPP1 | protein_coding | 3322,788691 | -0,242608283 | 0,066315301 | -3,658405799 | 0,000253789 | 0,009130626 |
| ENSG00000146072.6 | TNFRSF21 | protein_coding | 6994,943566 | -0,242940412 | 0,066567728 | -3,649522376 | 0,000262728 | 0,009376012 |
| ENSG00000112742.10 | TTK | protein_coding | 3050,880116 | -0,243057173 | 0,068207897 | -3,563475517 | 0,000365977 | 0,012050005 |
| ENSG00000101076.17 | HNF4A | protein_coding | 7533,063285 | -0,243237577 | 0,073125103 | -3,326321153 | 0,000880005 | 0,023459018 |
| ENSG00000007402.12 | CACNA2D2 | protein_coding | 5906,828262 | -0,243478242 | 0,076487089 | -3,183259376 | 0,001456271 | 0,034242148 |
| ENSG00000147262.4 | GPR119 | protein_coding | 2500,695564 | -0,244065029 | 0,074285297 | -3,285509215 | 0,001017982 | 0,026129782 |
| ENSG00000074356.17 | NCBP3 | protein_coding | 5437,565493 | -0,244522315 | 0,071776106 | -3,406736972 | 0,000657445 | 0,01867345 |
| ENSG00000164985.15 | PSIP1 | protein_coding | 11740,62747 | -0,244679064 | 0,059829453 | -4,089608926 | 4,32E-05 | 0,002265923 |
| ENSG00000134982.17 | APC | protein_coding | 6139,93084 | -0,244950872 | 0,078887448 | -3,105067759 | 0,001902354 | 0,041757434 |
| ENSG00000167123.19 | CERCAM | protein_coding | 3879,599177 | -0,246253235 | 0,080255444 | -3,068367996 | 0,002152314 | 0,044926508 |
| ENSG00000123612.16 | ACVR1C | protein_coding | 85966,01783 | -0,246463942 | 0,051081348 | -4,824930221 | 1,40E-06 | 0,00012649 |
| ENSG00000102572.15 | STK24 | protein_coding | 12625,29859 | -0,246922114 | 0,060958844 | -4,050636452 | 5,11E-05 | 0,002610077 |
| ENSG00000011105.14 | TSPAN9 | protein_coding | 2288,769072 | -0,24721325 | 0,080306775 | -3,078361075 | 0,002081426 | 0,044277641 |
| ENSG00000138180.16 | CEP55 | protein_coding | 4205,202031 | -0,247258948 | 0,059455737 | -4,158706274 | 3,20E-05 | 0,001788268 |
| ENSG00000164535.15 | DAGLB | protein_coding | 1692,694155 | -0,247800183 | 0,076924886 | -3,221326616 | 0,001275986 | 0,031127316 |
| ENSG00000159640.17 | ACE | protein_coding | 4672,448123 | -0,247949862 | 0,081230328 | -3,05242966 | 0,002269969 | 0,046677825 |
| ENSG00000277053.4 | GTF2IP1 | transcribed_unproc | 15160,12107 | -0,248042141 | 0,066959674 | -3,704351113 | 0,000211933 | 0,007934381 |
| ENSG00000135447.17 | PPP1R1A | protein_coding | 14690,67361 | -0,248116493 | 0,052274902 | -4,74637895 | 2,07E-06 | 0,00018111 |
| ENSG00000079112.10 | CDH17 | protein_coding | 50099,58774 | -0,249002635 | 0,05396271 | -4,61434638 | 3,94E-06 | 0,000320774 |
| ENSG00000064601.20 | CTSA | protein_coding | 34628,0647 | -0,249159125 | 0,072300351 | -3,446167573 | 0,000568598 | 0,016766689 |
| ENSG00000134138.20 | MEIS2 | protein_coding | 27912,13962 | -0,249182085 | 0,065722397 | -3,791433294 | 0,00014978 | 0,006103206 |
| ENSG00000129657.16 | SEC14L1 | protein_coding | 12036,20629 | -0,249776615 | 0,062358063 | -4,005522347 | 6,19E-05 | 0,00302245 |
| ENSG00000078674.18 | PCM1 | protein_coding | 8928,623543 | -0,249990626 | 0,062710141 | -3,986446556 | 6,71E-05 | 0,003212112 |
| ENSG00000125851.10 | PCSK2 | protein_coding | 117934,2378 | -0,250187141 | 0,049494017 | -5,054896682 | 4,31E-07 | 4,65E-05 |
| ENSG00000137868.19 | STRA6 | protein_coding | 13372,61727 | -0,250597542 | 0,070071922 | -3,576290414 | 0,000348504 | 0,011578093 |
| ENSG00000010292.13 | NCAPD2 | protein_coding | 11534,28684 | -0,250826082 | 0,067569333 | -3,712129019 | 0,000205523 | 0,007786654 |
| ENSG00000076351.13 | SLC46A1 | protein_coding | 6644,80238 | -0,251108243 | 0,065699792 | -3,822055363 | 0,000132344 | 0,005473351 |
| ENSG00000141639.12 | MAPK4 | protein_coding | 2180,313019 | -0,251486866 | 0,071168972 | -3,533658839 | 0,00040985 | 0,013123495 |
| ENSG00000149654.11 | CDH22 | protein_coding | 8547,507288 | -0,252105287 | 0,080682212 | -3,124669993 | 0,001780046 | 0,039583222 |
| ENSG00000131747.15 | TOP2A | protein_coding | 41882,65009 | -0,25216773 | 0,054921206 | -4,591445618 | 4,40E-06 | 0,000352349 |

| | | | | | | | | |
|---|---|---|---|---|---|---|---|---|
| ENSG00000137804.14 | NUSAP1 | protein_coding | 10684,2791 | -0,25261027 | 0,059684344 | -4,232437725 | 2,31E-05 | 0,001386155 |
| ENSG00000188070.9 | C11orf95 | protein_coding | 8788,973361 | -0,253260816 | 0,081285835 | -3,115682016 | 0,001835201 | 0,040484194 |
| ENSG00000158258.16 | CLSTN2 | protein_coding | 3985,663804 | -0,25341686 | 0,068004846 | -3,726452945 | 0,000194193 | 0,007459585 |
| ENSG00000270647.6 | TAF15 | protein_coding | 10908,74436 | -0,253852389 | 0,066946761 | -3,791854651 | 0,000149526 | 0,006103206 |
| ENSG00000050030.16 | NEXMIF | protein_coding | 1630,508226 | -0,253997082 | 0,078870766 | -3,220421159 | 0,001280024 | 0,031191421 |
| ENSG00000198586.14 | TLK1 | protein_coding | 4865,607461 | -0,254994351 | 0,082979161 | -3,072992628 | 0,002119237 | 0,044647493 |
| ENSG00000106236.4 | NPTX2 | protein_coding | 19430,67737 | -0,255360722 | 0,058387426 | -4,373556743 | 1,22E-05 | 0,000814653 |
| ENSG00000173598.14 | NUDT4 | protein_coding | 34244,33339 | -0,255731353 | 0,059223324 | -4,31808511 | 1,57E-05 | 0,001012321 |
| ENSG00000136828.19 | RALGPS1 | protein_coding | 4580,428542 | -0,256280955 | 0,066623874 | -3,846683492 | 0,000119727 | 0,005123965 |
| ENSG00000091129.21 | NRCAM | protein_coding | 17096,43164 | -0,256697427 | 0,05558599 | -4,618023828 | 3,87E-06 | 0,000318392 |
| ENSG00000176658.17 | MYO1D | protein_coding | 16950,51685 | -0,256707765 | 0,069281547 | -3,705283396 | 0,000211155 | 0,007931509 |
| ENSG00000172954.14 | LCLAT1 | protein_coding | 4566,174679 | -0,257009595 | 0,06381853 | -4,027193919 | 5,64E-05 | 0,002806595 |
| ENSG00000143437.21 | ARNT | protein_coding | 3079,540259 | -0,257550982 | 0,068177205 | -3,777669976 | 0,000158302 | 0,00634529 |
| ENSG00000090889.12 | KIF4A | protein_coding | 4240,812968 | -0,258040401 | 0,071184008 | -3,624977125 | 0,000288987 | 0,010026975 |
| ENSG00000241973.11 | PI4KA | protein_coding | 18343,15736 | -0,25812032 | 0,062196389 | -4,150085299 | 3,32E-05 | 0,001843009 |
| ENSG00000125384.7 | PTGER2 | protein_coding | 1363,301919 | -0,259199604 | 0,08475621 | -3,058178343 | 0,00222687 | 0,046040201 |
| ENSG00000123836.15 | PFKFB2 | protein_coding | 17707,80756 | -0,260232912 | 0,053743796 | -4,842101444 | 1,28E-06 | 0,00011795 |
| ENSG00000140832.10 | MARVELD3 | protein_coding | 3855,004299 | -0,261112734 | 0,061503228 | -4,245512662 | 2,18E-05 | 0,001311291 |
| ENSG00000100647.8 | SUSD6 | protein_coding | 2027,712163 | -0,261289031 | 0,085717746 | -3,048248976 | 0,002301791 | 0,047156884 |
| ENSG00000163125.15 | RPRD2 | protein_coding | 4733,826207 | -0,261455655 | 0,073919405 | -3,537036761 | 0,000404643 | 0,013060223 |
| ENSG00000086015.22 | MAST2 | protein_coding | 4408,25548 | -0,261924925 | 0,075595971 | -3,464800041 | 0,000530626 | 0,015995406 |
| ENSG00000131389.17 | SLC6A6 | protein_coding | 12540,77825 | -0,261951641 | 0,059523079 | -4,400841561 | 1,08E-05 | 0,000743267 |
| ENSG00000174469.23 | CNTNAP2 | protein_coding | 2077,261463 | -0,261981702 | 0,077273226 | -3,390329556 | 0,000698086 | 0,019634459 |
| ENSG00000133466.14 | C1QTNF6 | protein_coding | 3547,592856 | -0,262722372 | 0,072960162 | -3,600901727 | 0,000317115 | 0,010695878 |
| ENSG00000150760.13 | DOCK1 | protein_coding | 3559,774568 | -0,263073178 | 0,069915598 | -3,762725159 | 0,000168072 | 0,006652519 |
| ENSG00000113300.13 | CNOT6 | protein_coding | 7056,338761 | -0,263973776 | 0,063810611 | -4,136831989 | 3,52E-05 | 0,001922521 |
| ENSG00000174839.13 | DENND6A | protein_coding | 4713,220742 | -0,264214253 | 0,070774882 | -3,733164163 | 0,000189089 | 0,007327126 |
| ENSG00000138944.8 | SHISAL1 | protein_coding | 3944,511588 | -0,264291428 | 0,065379728 | -4,042406357 | 5,29E-05 | 0,00268483 |
| ENSG00000141258.13 | SGSM2 | protein_coding | 4131,890436 | -0,264783165 | 0,082529066 | -3,20836257 | 0,001334931 | 0,03207023 |
| ENSG00000131374.14 | TBC1D5 | protein_coding | 7008,370414 | -0,265059307 | 0,056322864 | -4,706069415 | 2,53E-06 | 0,000216576 |
| ENSG00000114861.23 | FOXP1 | protein_coding | 3086,38241 | -0,26606148 | 0,068898187 | -3,86166154 | 0,000112619 | 0,004866791 |
| ENSG00000091262.16 | ABCC6 | protein_coding | 3495,857425 | -0,266303076 | 0,069699584 | -3,820726888 | 0,000133059 | 0,005476152 |
| ENSG00000185565.12 | LSAMP | protein_coding | 3058,826393 | -0,266520419 | 0,086300128 | -3,088296916 | 0,002013072 | 0,043382749 |
| ENSG00000068650.18 | ATP11A | protein_coding | 8055,347913 | -0,266971044 | 0,077941016 | -3,425295936 | 0,00061413 | 0,01773922 |
| ENSG00000204569.10 | PPP1R10 | protein_coding | 6805,710429 | -0,267155786 | 0,086792135 | -3,078110547 | 0,002083176 | 0,044277641 |
| ENSG00000144040.13 | SFXN5 | protein_coding | 2825,608804 | -0,267363803 | 0,073612176 | -3,632059489 | 0,000281168 | 0,009874809 |

| | | | | | | | | |
|---|---|---|---|---|---|---|---|---|
| ENSG00000133961.20 | NUMB | protein_coding | 3982,468993 | -0,269489412 | 0,067218709 | -4,009142922 | 6,09E-05 | 0,002989685 |
| ENSG00000134318.14 | ROCK2 | protein_coding | 5394,466277 | -0,269873681 | 0,078897569 | -3,420557603 | 0,000624929 | 0,017934084 |
| ENSG00000226137.5 | BAIAP2-DT | lncRNA | 2987,121306 | -0,270437975 | 0,07566874 | -3,57397224 | 0,000351606 | 0,011646167 |
| ENSG00000148019.14 | CEP78 | protein_coding | 4170,378772 | -0,270465513 | 0,074967911 | -3,607750408 | 0,000308863 | 0,010545056 |
| ENSG00000092148.13 | HECTD1 | protein_coding | 15031,15878 | -0,271795429 | 0,080439054 | -3,378898887 | 0,000727768 | 0,020254828 |
| ENSG00000137103.20 | TMEM8B | protein_coding | 1647,083883 | -0,271795903 | 0,08300494 | -3,274454537 | 0,001058662 | 0,02692408 |
| ENSG00000128266.9 | GNAZ | protein_coding | 5404,938866 | -0,272266545 | 0,082941195 | -3,282645552 | 0,001028379 | 0,026305098 |
| ENSG00000213853.10 | EMP2 | protein_coding | 10421,92719 | -0,27348927 | 0,08236325 | -3,320525475 | 0,000898482 | 0,023779669 |
| ENSG00000144619.15 | CNTN4 | protein_coding | 2561,198035 | -0,274061854 | 0,08436068 | -3,24869184 | 0,00115937 | 0,028758095 |
| ENSG00000166341.8 | DCHS1 | protein_coding | 4732,809404 | -0,275239665 | 0,087131453 | -3,158901347 | 0,001583651 | 0,036160846 |
| ENSG00000089057.15 | SLC23A2 | protein_coding | 6496,885287 | -0,275524316 | 0,060750184 | -4,535365929 | 5,75E-06 | 0,000441999 |
| ENSG00000198265.12 | HELZ | protein_coding | 3842,890599 | -0,276360269 | 0,069315099 | -3,987013994 | 6,69E-05 | 0,003211394 |
| ENSG00000143819.12 | EPHX1 | protein_coding | 32656,25748 | -0,276481667 | 0,053970575 | -5,122822347 | 3,01E-07 | 3,36E-05 |
| ENSG00000158417.11 | EIF5B | protein_coding | 7483,91321 | -0,27725578 | 0,085009731 | -3,26145932 | 0,001108403 | 0,027737202 |
| ENSG00000169855.20 | ROBO1 | protein_coding | 17991,73481 | -0,277322678 | 0,074763364 | -3,709339203 | 0,000207801 | 0,007846079 |
| ENSG00000164715.6 | LMTK2 | protein_coding | 6585,867615 | -0,277515222 | 0,083900019 | -3,307689634 | 0,00094069 | 0,024535889 |
| ENSG00000180357.9 | ZNF609 | protein_coding | 3401,926345 | -0,278055923 | 0,079286251 | -3,506987905 | 0,00045321 | 0,014169001 |
| ENSG00000197535.15 | MYO5A | protein_coding | 3391,275967 | -0,278455034 | 0,085276953 | -3,265302349 | 0,001093473 | 0,027583538 |
| ENSG00000113269.14 | RNF130 | protein_coding | 9419,105346 | -0,278535127 | 0,066054417 | -4,216752508 | 2,48E-05 | 0,001454985 |
| ENSG00000100014.20 | SPECC1L | protein_coding | 12380,85355 | -0,279051178 | 0,068686445 | -4,062681907 | 4,85E-05 | 0,00249623 |
| ENSG00000112984.12 | KIF20A | protein_coding | 3724,68697 | -0,279381663 | 0,069088789 | -4,043806043 | 5,26E-05 | 0,002674979 |
| ENSG00000107036.12 | RIC1 | protein_coding | 1849,588066 | -0,279632537 | 0,084896417 | -3,293808467 | 0,000988398 | 0,025459021 |
| ENSG00000185008.17 | ROBO2 | protein_coding | 9074,044839 | -0,279678195 | 0,06192256 | -4,516580017 | 6,28E-06 | 0,000477848 |
| ENSG00000106829.20 | TLE4 | protein_coding | 4529,819766 | -0,282307463 | 0,070651749 | -3,995760422 | 6,45E-05 | 0,003122183 |
| ENSG00000142156.15 | COL6A1 | protein_coding | 2364,076732 | -0,282385019 | 0,093028283 | -3,035474901 | 0,002401572 | 0,048527109 |
| ENSG00000213626.13 | LBH | protein_coding | 3343,206019 | -0,282389892 | 0,071534629 | -3,947597088 | 7,89E-05 | 0,003654007 |
| ENSG00000197077.13 | KIAA1671 | protein_coding | 3111,553484 | -0,282562377 | 0,085190311 | -3,316836995 | 0,000910427 | 0,02395257 |
| ENSG00000138641.18 | HERC3 | protein_coding | 1479,712146 | -0,283194835 | 0,081659258 | -3,468006456 | 0,000524335 | 0,015862633 |
| ENSG00000137177.20 | KIF13A | protein_coding | 5107,695111 | -0,284192696 | 0,074036133 | -3,838567558 | 0,000123754 | 0,005225543 |
| ENSG00000170558.10 | CDH2 | protein_coding | 10141,49959 | -0,284607722 | 0,066735502 | -4,264712387 | 2,00E-05 | 0,001219786 |
| ENSG00000052126.14 | PLEKHA5 | protein_coding | 3294,759316 | -0,284726782 | 0,069086022 | -4,121337073 | 3,77E-05 | 0,002027839 |
| ENSG00000151176.8 | PLBD2 | protein_coding | 6049,647811 | -0,285326909 | 0,090129994 | -3,165726481 | 0,001546962 | 0,035728682 |
| ENSG00000110697.13 | PITPNM1 | protein_coding | 12852,32513 | -0,286227793 | 0,066696715 | -4,291482633 | 1,77E-05 | 0,001115629 |
| ENSG00000081913.14 | PHLPP1 | protein_coding | 2537,477902 | -0,287312431 | 0,086829527 | -3,308925444 | 0,000936548 | 0,024523142 |
| ENSG00000107862.6 | GBF1 | protein_coding | 8189,877592 | -0,287465592 | 0,090953114 | -3,160590988 | 0,001574494 | 0,035988904 |
| ENSG00000102384.13 | CENPI | protein_coding | 1154,951171 | -0,287674014 | 0,093518995 | -3,076102482 | 0,002097257 | 0,044486374 |

| | | | | | | | | |
|---|---|---|---|---|---|---|---|---|
| ENSG00000155465.20 | SLC7A7 | protein_coding | 2349,079861 | -0,287829632 | 0,075008308 | -3,837303344 | 0,000124393 | 0,005242504 |
| ENSG00000170776.22 | AKAP13 | protein_coding | 5538,596038 | -0,288341807 | 0,091191316 | -3,161943716 | 0,001567199 | 0,035933507 |
| ENSG00000198300.14 | PEG3 | protein_coding | 2184,552673 | -0,288540666 | 0,074655051 | -3,864985158 | 0,000111096 | 0,00481039 |
| ENSG00000164176.13 | EDIL3 | protein_coding | 12816,4342 | -0,290019884 | 0,051146937 | -5,670327571 | 1,43E-08 | 2,46E-06 |
| ENSG00000196367.14 | TRRAP | protein_coding | 9684,764522 | -0,290100742 | 0,087390365 | -3,319596409 | 0,000901477 | 0,023801997 |
| ENSG00000133056.13 | PIK3C2B | protein_coding | 2098,381071 | -0,29065063 | 0,089124443 | -3,261177509 | 0,001109506 | 0,027737202 |
| ENSG00000141576.16 | RNF157 | protein_coding | 1490,961969 | -0,290867355 | 0,091558585 | -3,17684416 | 0,00148887 | 0,034741537 |
| ENSG00000114251.15 | WNT5A | protein_coding | 2603,337091 | -0,291084316 | 0,072173566 | -4,033115328 | 5,50E-05 | 0,00274913 |
| ENSG00000160460.16 | SPTBN4 | protein_coding | 1713,452987 | -0,292115536 | 0,094090121 | -3,104635567 | 0,001905136 | 0,041777046 |
| ENSG00000167202.11 | TBC1D2B | protein_coding | 3336,414994 | -0,292136441 | 0,068508713 | -4,264223129 | 2,01E-05 | 0,001219786 |
| ENSG00000101493.11 | ZNF516 | protein_coding | 7544,34114 | -0,292622799 | 0,077225561 | -3,78919615 | 0,000151136 | 0,006147105 |
| ENSG00000186866.17 | POFUT2 | protein_coding | 3046,324972 | -0,292999849 | 0,083460769 | -3,510629631 | 0,000447047 | 0,014050223 |
| ENSG00000130477.15 | UNC13A | protein_coding | 8579,293353 | -0,293274735 | 0,089812859 | -3,265398052 | 0,001093103 | 0,027583538 |
| ENSG00000099991.18 | CABIN1 | protein_coding | 5090,822532 | -0,293345002 | 0,086460623 | -3,392816203 | 0,00069178 | 0,019498513 |
| ENSG00000105088.8 | OLFM2 | protein_coding | 4662,156828 | -0,293578014 | 0,066190232 | -4,435367655 | 9,19E-06 | 0,000658161 |
| ENSG00000087258.16 | GNAO1 | protein_coding | 13256,91454 | -0,294195333 | 0,085984512 | -3,421492146 | 0,000622785 | 0,017934084 |
| ENSG00000182670.13 | TTC3 | protein_coding | 30481,11444 | -0,294890043 | 0,056107776 | -5,255778502 | 1,47E-07 | 1,78E-05 |
| ENSG00000218336.9 | TENM3 | protein_coding | 44550,92238 | -0,295138942 | 0,06056514 | -4,87308278 | 1,10E-06 | 0,000104334 |
| ENSG00000182378.14 | PLCXD1 | protein_coding | 6268,615033 | -0,295593142 | 0,058043623 | -5,092603222 | 3,53E-07 | 3,87E-05 |
| ENSG00000175216.15 | CKAP5 | protein_coding | 13926,09879 | -0,295970516 | 0,071415928 | -4,144320809 | 3,41E-05 | 0,001875879 |
| ENSG00000187555.16 | USP7 | protein_coding | 13963,82157 | -0,296120344 | 0,070596696 | -4,194535457 | 2,73E-05 | 0,001567343 |
| ENSG00000116984.14 | MTR | protein_coding | 2539,650432 | -0,296506436 | 0,096483877 | -3,07311901 | 0,00211834 | 0,044647493 |
| ENSG00000124406.16 | ATP8A1 | protein_coding | 23698,87673 | -0,297518987 | 0,04999162 | -5,951377227 | 2,66E-09 | 5,66E-07 |
| ENSG00000124766.7 | SOX4 | protein_coding | 13345,86172 | -0,297657291 | 0,081233188 | -3,664232539 | 0,000248081 | 0,008969032 |
| ENSG00000183856.11 | IQGAP3 | protein_coding | 2986,283161 | -0,298759826 | 0,078122676 | -3,824239514 | 0,000131176 | 0,005445418 |
| ENSG00000120071.15 | KANSL1 | protein_coding | 2125,509488 | -0,299193393 | 0,077221126 | -3,874501809 | 0,000106843 | 0,004653562 |
| ENSG00000144847.13 | IGSF11 | protein_coding | 1956,767002 | -0,299226468 | 0,07975991 | -3,751589828 | 0,000175717 | 0,00693032 |
| ENSG00000126705.15 | AHDC1 | protein_coding | 2650,534215 | -0,299315332 | 0,09380952 | -3,190671164 | 0,001419427 | 0,033553685 |
| ENSG00000182158.15 | CREB3L2 | protein_coding | 6714,457784 | -0,299566107 | 0,063346845 | -4,728982294 | 2,26E-06 | 0,000195792 |
| ENSG00000013293.6 | SLC7A14 | protein_coding | 3428,572496 | -0,299868913 | 0,065316595 | -4,591006492 | 4,41E-06 | 0,000352349 |
| ENSG00000156970.13 | BUB1B | protein_coding | 4458,370853 | -0,299981475 | 0,067593728 | -4,438007565 | 9,08E-06 | 0,000652253 |
| ENSG00000185219.17 | ZNF445 | protein_coding | 3176,748706 | -0,300053425 | 0,092168602 | -3,25548416 | 0,001131992 | 0,028173743 |
| ENSG00000117298.16 | ECE1 | protein_coding | 16828,62515 | -0,300656358 | 0,081856837 | -3,672953531 | 0,000239763 | 0,008708931 |
| ENSG00000166783.22 | MARF1 | protein_coding | 4349,923838 | -0,300705105 | 0,082772392 | -3,632915478 | 0,000280237 | 0,00987344 |
| ENSG00000197177.15 | ADGRA1 | protein_coding | 1655,007437 | -0,302322926 | 0,088369669 | -3,421116446 | 0,000623646 | 0,017934084 |
| ENSG00000154229.12 | PRKCA | protein_coding | 4173,090005 | -0,302519728 | 0,07628974 | -3,965405163 | 7,33E-05 | 0,003434751 |

| | | | | | | | | |
|---|---|---|---|---|---|---|---|---|
| ENSG00000162877.13 | PM20D1 | protein_coding | 838,8453479 | -0,302837768 | 0,096528576 | -3,13728619 | 0,001705196 | 0,038264873 |
| ENSG00000073060.16 | SCARB1 | protein_coding | 9745,782582 | -0,302979112 | 0,077230432 | -3,92305344 | 8,74E-05 | 0,003972401 |
| ENSG00000151461.20 | UPF2 | protein_coding | 1300,673791 | -0,303091277 | 0,094768492 | -3,198228346 | 0,001382747 | 0,032897494 |
| ENSG00000134744.14 | TUT4 | protein_coding | 4889,887217 | -0,303136665 | 0,073639139 | -4,116515596 | 3,85E-05 | 0,002055705 |
| ENSG00000242110.8 | AMACR | protein_coding | 16644,51296 | -0,303451947 | 0,065151647 | -4,657625079 | 3,20E-06 | 0,000266902 |
| ENSG00000169933.16 | FRMPD4 | protein_coding | 3007,590256 | -0,303584641 | 0,077187413 | -3,933084786 | 8,39E-05 | 0,003849678 |
| ENSG00000107099.16 | DOCK8 | protein_coding | 3817,025822 | -0,303788017 | 0,069325411 | -4,382058651 | 1,18E-05 | 0,000790639 |
| ENSG00000234857.2 | HNRNPUL2-BSCL | protein_coding | 5266,39729 | -0,305398606 | 0,099332679 | -3,074502865 | 0,002108537 | 0,044553587 |
| ENSG00000144749.14 | LRIG1 | protein_coding | 1935,600306 | -0,30552642 | 0,080830771 | -3,779828101 | 0,000156937 | 0,00631342 |
| ENSG00000196632.10 | WNK3 | protein_coding | 1293,711598 | -0,306042755 | 0,091342463 | -3,350498155 | 0,000806663 | 0,02192658 |
| ENSG00000233369.7 | GTF2IP4 | transcribed_process | 30207,27522 | -0,306393221 | 0,060243007 | -5,085954985 | 3,66E-07 | 3,99E-05 |
| ENSG00000204842.18 | ATXN2 | protein_coding | 2003,577817 | -0,307251776 | 0,094206576 | -3,26146845 | 0,001108368 | 0,027737202 |
| ENSG00000179981.11 | TSHZ1 | protein_coding | 18150,46381 | -0,307274529 | 0,085300653 | -3,602252945 | 0,000315471 | 0,010672959 |
| ENSG00000111961.18 | SASH1 | protein_coding | 2581,84599 | -0,308954457 | 0,089587839 | -3,448620484 | 0,000563458 | 0,016667209 |
| ENSG00000006530.18 | AGK | protein_coding | 2194,207584 | -0,309689257 | 0,097795489 | -3,166702866 | 0,001541778 | 0,035646157 |
| ENSG00000069702.11 | TGFBR3 | protein_coding | 8438,14503 | -0,309710019 | 0,067073951 | -4,617441092 | 3,89E-06 | 0,000318392 |
| ENSG00000157851.17 | DPYSL5 | protein_coding | 7465,326891 | -0,310012811 | 0,082034467 | -3,779055601 | 0,000157424 | 0,006321542 |
| ENSG00000079385.23 | CEACAM1 | protein_coding | 4882,300765 | -0,310308099 | 0,076825202 | -4,039144579 | 5,36E-05 | 0,002709557 |
| ENSG00000058404.20 | CAMK2B | protein_coding | 2025,728187 | -0,31072619 | 0,080927084 | -3,839582168 | 0,000123244 | 0,005223935 |
| ENSG00000119711.13 | ALDH6A1 | protein_coding | 1142,860784 | -0,311421302 | 0,090892305 | -3,426266969 | 0,000611939 | 0,01772219 |
| ENSG00000111879.20 | FAM184A | protein_coding | 1337,616689 | -0,313603322 | 0,088676937 | -3,536469937 | 0,000405513 | 0,013060223 |
| ENSG00000108557.19 | RAI1 | protein_coding | 1794,67486 | -0,313964455 | 0,099205648 | -3,164784062 | 0,001551981 | 0,03577443 |
| ENSG00000075213.11 | SEMA3A | protein_coding | 4862,188038 | -0,314600866 | 0,071774972 | -4,383155548 | 1,17E-05 | 0,000789065 |
| ENSG00000074370.18 | ATP2A3 | protein_coding | 136333,9884 | -0,31540377 | 0,075185016 | -4,195034949 | 2,73E-05 | 0,001567343 |
| ENSG00000118482.12 | PHF3 | protein_coding | 7893,76581 | -0,317073884 | 0,073988357 | -4,285456482 | 1,82E-05 | 0,001136268 |
| ENSG00000173166.18 | RAPH1 | protein_coding | 1919,370319 | -0,317077042 | 0,102742545 | -3,08613185 | 0,002027789 | 0,043480147 |
| ENSG00000185697.16 | MYBL1 | protein_coding | 1367,522377 | -0,317615113 | 0,094375458 | -3,365441801 | 0,000764212 | 0,021060093 |
| ENSG00000154122.14 | ANKH | protein_coding | 9044,977703 | -0,320542766 | 0,060390378 | -5,307844972 | 1,11E-07 | 1,41E-05 |
| ENSG00000072062.15 | PRKACA | protein_coding | 9545,67831 | -0,320790607 | 0,078026142 | -4,111322151 | 3,93E-05 | 0,002092395 |
| ENSG00000213366.13 | GSTM2 | protein_coding | 2544,084109 | -0,321369353 | 0,090461206 | -3,552565427 | 0,000381494 | 0,012523649 |
| ENSG00000135835.12 | KIAA1614 | protein_coding | 1587,590344 | -0,325169998 | 0,092011423 | -3,534017697 | 0,000409294 | 0,013123495 |
| ENSG00000066279.18 | ASPM | protein_coding | 4287,38936 | -0,32590965 | 0,072711062 | -4,482256789 | 7,39E-06 | 0,000550228 |
| ENSG00000163536.13 | SERPINI1 | protein_coding | 27506,25263 | -0,326558469 | 0,056519465 | -5,777805346 | 7,57E-09 | 1,44E-06 |
| ENSG00000047410.14 | TPR | protein_coding | 5940,979001 | -0,327971565 | 0,105326841 | -3,113846021 | 0,001846658 | 0,040696377 |
| ENSG00000148408.13 | CACNA1B | protein_coding | 1102,604706 | -0,328707412 | 0,087893619 | -3,739832496 | 0,000184143 | 0,007185796 |
| ENSG00000089199.10 | CHGB | protein_coding | 45591,34119 | -0,329467773 | 0,089301975 | -3,689367143 | 0,000224813 | 0,008262796 |

| | | | | | | | | |
|---|---|---|---|---|---|---|---|---|
| ENSG00000164330.17 | EBF1 | protein_coding | 967,2316427 | -0,329508728 | 0,09282176 | -3,549908224 | 0,000385365 | 0,012539112 |
| ENSG00000102189.17 | EEA1 | protein_coding | 3052,491545 | -0,331179262 | 0,071725337 | -4,617325963 | 3,89E-06 | 0,000318392 |
| ENSG00000154529.15 | CNTNAP3B | protein_coding | 1758,221872 | -0,332078565 | 0,082844465 | -4,008458062 | 6,11E-05 | 0,002991733 |
| ENSG00000172493.22 | AFF1 | protein_coding | 3225,384642 | -0,332189028 | 0,103443798 | -3,211299597 | 0,001321361 | 0,031847966 |
| ENSG00000065361.16 | ERBB3 | protein_coding | 7160,942102 | -0,332680641 | 0,066295346 | -5,018159786 | 5,22E-07 | 5,50E-05 |
| ENSG00000088888.18 | MAVS | protein_coding | 6370,684082 | -0,332699554 | 0,094870432 | -3,506883531 | 0,000453388 | 0,014169001 |
| ENSG00000196549.12 | MME | protein_coding | 10220,77042 | -0,333210023 | 0,060508506 | -5,50682946 | 3,65E-08 | 5,39E-06 |
| ENSG00000157005.4 | SST | protein_coding | 4538,67538 | -0,334131593 | 0,075840698 | -4,405703006 | 1,05E-05 | 0,000733646 |
| ENSG00000157933.10 | SKI | protein_coding | 2591,373376 | -0,335010376 | 0,099381314 | -3,370959404 | 0,000749069 | 0,020754363 |
| ENSG00000119321.9 | FKBP15 | protein_coding | 1819,675323 | -0,335760767 | 0,095705353 | -3,508275742 | 0,000451021 | 0,014155032 |
| ENSG00000152954.12 | NRSN1 | protein_coding | 787,1947388 | -0,336798807 | 0,103284808 | -3,260874591 | 0,001110692 | 0,027737202 |
| ENSG00000145388.15 | METTL14 | protein_coding | 5475,84566 | -0,33778212 | 0,08968616 | -3,766268067 | 0,000165706 | 0,006594259 |
| ENSG00000113649.11 | TCERG1 | protein_coding | 3339,142091 | -0,338148226 | 0,099395387 | -3,402051516 | 0,00066882 | 0,01892368 |
| ENSG00000152818.18 | UTRN | protein_coding | 3487,487749 | -0,339259891 | 0,100646323 | -3,37081256 | 0,000749468 | 0,020754363 |
| ENSG00000170004.17 | CHD3 | protein_coding | 3423,291967 | -0,34078399 | 0,110814591 | -3,075262816 | 0,002103171 | 0,04453088 |
| ENSG00000099834.19 | CDHR5 | protein_coding | 8020,913319 | -0,342245916 | 0,094722018 | -3,613161169 | 0,000302487 | 0,010392574 |
| ENSG00000145555.15 | MYO10 | protein_coding | 12231,62945 | -0,342426168 | 0,087635969 | -3,907370112 | 9,33E-05 | 0,004187611 |
| ENSG00000115355.17 | CCDC88A | protein_coding | 4713,652608 | -0,342933551 | 0,08167532 | -4,198741434 | 2,68E-05 | 0,001554627 |
| ENSG00000106144.20 | CASP2 | protein_coding | 5953,582932 | -0,343014467 | 0,064824378 | -5,291442506 | 1,21E-07 | 1,51E-05 |
| ENSG00000100150.20 | DEPDC5 | protein_coding | 1128,162252 | -0,343098895 | 0,09663988 | -3,550282692 | 0,000384818 | 0,012539112 |
| ENSG00000197555.9 | SIPA1L1 | protein_coding | 3683,142682 | -0,34327045 | 0,101327989 | -3,387716017 | 0,000704772 | 0,019763985 |
| ENSG00000111670.16 | GNPTAB | protein_coding | 2013,923744 | -0,343593894 | 0,085614339 | -4,013275111 | 5,99E-05 | 0,002944334 |
| ENSG00000186111.10 | PIP5K1C | protein_coding | 2323,120363 | -0,344544812 | 0,113246966 | -3,042419796 | 0,002346843 | 0,047783034 |
| ENSG00000101049.17 | SGK2 | protein_coding | 1660,112643 | -0,348334239 | 0,088342998 | -3,942975056 | 8,05E-05 | 0,003709656 |
| ENSG00000029534.21 | ANK1 | protein_coding | 4020,389378 | -0,34958609 | 0,110188254 | -3,172625733 | 0,001510671 | 0,035044954 |
| ENSG00000164742.16 | ADCY1 | protein_coding | 14167,31507 | -0,350763649 | 0,06168483 | -5,686384301 | 1,30E-08 | 2,28E-06 |
| ENSG00000105270.15 | CLIP3 | protein_coding | 2057,061119 | -0,351271192 | 0,089618866 | -3,919612115 | 8,87E-05 | 0,004017174 |
| ENSG00000089177.19 | KIF16B | protein_coding | 2749,096496 | -0,352815913 | 0,071825022 | -4,912158766 | 9,01E-07 | 8,82E-05 |
| ENSG00000152822.14 | GRM1 | protein_coding | 675,5551412 | -0,352845167 | 0,110393213 | -3,196257776 | 0,001392227 | 0,03305194 |
| ENSG00000171219.9 | CDC42BPG | protein_coding | 5845,028155 | -0,352924216 | 0,099592759 | -3,543673463 | 0,000394594 | 0,012782985 |
| ENSG00000169607.13 | CKAP2L | protein_coding | 2961,901545 | -0,353964632 | 0,075982934 | -4,658475424 | 3,19E-06 | 0,000266902 |
| ENSG00000182667.14 | NTM | protein_coding | 1573,37591 | -0,354226414 | 0,099749463 | -3,551161099 | 0,000383536 | 0,012539112 |
| ENSG00000257335.8 | MGAM | protein_coding | 7638,557926 | -0,354252162 | 0,081758052 | -4,332933013 | 1,47E-05 | 0,00095577 |
| ENSG00000155506.18 | LARP1 | protein_coding | 11636,16374 | -0,354678137 | 0,098147606 | -3,613721738 | 0,000301833 | 0,010386247 |
| ENSG00000153317.15 | ASAP1 | protein_coding | 4841,243143 | -0,35574498 | 0,100165563 | -3,551569712 | 0,000382941 | 0,012539112 |
| ENSG00000067208.14 | EVI5 | protein_coding | 966,5491603 | -0,357410467 | 0,099029949 | -3,609114939 | 0,000307243 | 0,010523326 |

| | | | | | | | | |
|---|---|---|---|---|---|---|---|---|
| ENSG00000075568.17 | TMEM131 | protein_coding | 5859,006536 | -0,357945435 | 0,095626828 | -3,743148664 | 0,000181729 | 0,007104113 |
| ENSG00000111206.13 | FOXM1 | protein_coding | 7659,864898 | -0,358503686 | 0,088308062 | -4,059693744 | 4,91E-05 | 0,002522524 |
| ENSG00000100626.17 | GALNT16 | protein_coding | 4570,168155 | -0,359402829 | 0,096797878 | -3,712920523 | 0,000204881 | 0,007786654 |
| ENSG00000145734.19 | BDP1 | protein_coding | 3671,99115 | -0,359633722 | 0,103686235 | -3,468480847 | 0,00052341 | 0,015862633 |
| ENSG00000198826.11 | ARHGAP11A | protein_coding | 7544,712772 | -0,359839149 | 0,062586035 | -5,749511822 | 8,95E-09 | 1,68E-06 |
| ENSG00000167106.12 | FAM102A | protein_coding | 7987,749381 | -0,360069381 | 0,088815734 | -4,054117052 | 5,03E-05 | 0,002577475 |
| ENSG00000167711.14 | SERPINF2 | protein_coding | 583,7890005 | -0,360224717 | 0,117043253 | -3,07770595 | 0,002086006 | 0,044294604 |
| ENSG00000072501.19 | SMC1A | protein_coding | 33813,34203 | -0,360502676 | 0,069485506 | -5,188170878 | 2,12E-07 | 2,45E-05 |
| ENSG00000169418.10 | NPR1 | protein_coding | 1595,939191 | -0,362759511 | 0,106118913 | -3,418424673 | 0,000629847 | 0,018051821 |
| ENSG00000117152.14 | RGS4 | protein_coding | 4624,965969 | -0,363259909 | 0,064814635 | -5,604597007 | 2,09E-08 | 3,42E-06 |
| ENSG00000196782.12 | MAML3 | protein_coding | 1711,357742 | -0,364367008 | 0,09343874 | -3,89952826 | 9,64E-05 | 0,004273568 |
| ENSG00000141526.17 | SLC16A3 | protein_coding | 3084,355198 | -0,365684465 | 0,08148057 | -4,487995936 | 7,19E-06 | 0,000537425 |
| ENSG00000105738.11 | SIPA1L3 | protein_coding | 3722,368047 | -0,366088832 | 0,090807309 | -4,031490816 | 5,54E-05 | 0,002761971 |
| ENSG00000287915.1 | AC037433.1 | lncRNA | 596,7135563 | -0,368352925 | 0,121381402 | -3,034673516 | 0,002407962 | 0,048567517 |
| ENSG00000183963.19 | SMTN | protein_coding | 2116,861835 | -0,369666223 | 0,101959054 | -3,625634099 | 0,000288253 | 0,010026975 |
| ENSG00000035862.12 | TIMP2 | protein_coding | 1810,924517 | -0,370883256 | 0,108190072 | -3,428071057 | 0,000607886 | 0,017632907 |
| ENSG00000048052.22 | HDAC9 | protein_coding | 792,7001047 | -0,371577552 | 0,099959779 | -3,717270643 | 0,000201387 | 0,007695372 |
| ENSG00000182575.8 | NXPH3 | protein_coding | 965,5578556 | -0,373820653 | 0,104354004 | -3,582235832 | 0,000340666 | 0,011351772 |
| ENSG00000113916.18 | BCL6 | protein_coding | 1232,675951 | -0,374367916 | 0,091596703 | -4,087133087 | 4,37E-05 | 0,002284453 |
| ENSG00000187017.17 | ESPN | protein_coding | 2660,869053 | -0,375302995 | 0,103440987 | -3,628184581 | 0,000285421 | 0,009976665 |
| ENSG00000150867.14 | PIP4K2A | protein_coding | 1476,297801 | -0,375599146 | 0,116156979 | -3,233547816 | 0,001222629 | 0,030024286 |
| ENSG00000142949.17 | PTPRF | protein_coding | 11203,23908 | -0,37569536 | 0,08767892 | -4,284899512 | 1,83E-05 | 0,001136268 |
| ENSG00000104936.19 | DMPK | protein_coding | 1004,158636 | -0,376839448 | 0,117277877 | -3,213218538 | 0,001312564 | 0,031739657 |
| ENSG00000104419.17 | NDRG1 | protein_coding | 4854,072903 | -0,377365635 | 0,077639399 | -4,860491474 | 1,17E-06 | 0,000110193 |
| ENSG00000272391.6 | POM121C | protein_coding | 4410,384686 | -0,378470077 | 0,102914308 | -3,677526311 | 0,000235507 | 0,008612931 |
| ENSG00000146555.19 | SDK1 | protein_coding | 3838,101608 | -0,378802209 | 0,096917886 | -3,908486064 | 9,29E-05 | 0,004185298 |
| ENSG00000164855.16 | TMEM184A | protein_coding | 2405,519171 | -0,379588235 | 0,121164408 | -3,13283614 | 0,001731261 | 0,038663136 |
| ENSG00000177225.17 | GATD1 | protein_coding | 1674,134158 | -0,379625238 | 0,114763865 | -3,307881256 | 0,000940047 | 0,024535889 |
| ENSG00000162804.14 | SNED1 | protein_coding | 1610,633712 | -0,381247189 | 0,10200082 | -3,737687501 | 0,000185721 | 0,007234601 |
| ENSG00000144674.16 | GOLGA4 | protein_coding | 2826,045716 | -0,384399088 | 0,116221931 | -3,307457432 | 0,00094147 | 0,024535889 |
| ENSG00000101040.19 | ZMYND8 | protein_coding | 1954,238915 | -0,385239756 | 0,102197291 | -3,76956915 | 0,00016353 | 0,006531148 |
| ENSG00000120645.12 | IQSEC3 | protein_coding | 3157,351883 | -0,385849621 | 0,117690067 | -3,278523249 | 0,001043518 | 0,026569473 |
| ENSG00000163808.17 | KIF15 | protein_coding | 1697,170178 | -0,386076829 | 0,100827941 | -3,829065877 | 0,000128631 | 0,005372044 |
| ENSG00000090924.15 | PLEKHG2 | protein_coding | 1360,290273 | -0,387912059 | 0,114145295 | -3,398406031 | 0,000677797 | 0,019153188 |
| ENSG00000166035.11 | LIPC | protein_coding | 1341,8416 | -0,388744903 | 0,107158696 | -3,627749481 | 0,000285902 | 0,009977726 |
| ENSG00000156687.11 | UNC5D | protein_coding | 1818,644404 | -0,390696529 | 0,073249065 | -5,333809124 | 9,62E-08 | 1,24E-05 |

| | | | | | | | | |
|---|---|---|---|---|---|---|---|---|
| ENSG00000164199.18 | ADGRV1 | protein_coding | 2364,921348 | -0,390826488 | 0,085165106 | -4,589044827 | 4,45E-06 | 0,000354397 |
| ENSG00000132694.19 | ARHGEF11 | protein_coding | 1896,422271 | -0,39194704 | 0,128871048 | -3,041389409 | 0,002354891 | 0,047802118 |
| ENSG00000164989.17 | CCDC171 | protein_coding | 519,360014 | -0,393310859 | 0,117847386 | -3,337459334 | 0,000845481 | 0,022758043 |
| ENSG00000168143.9 | FAM83B | protein_coding | 420,0790901 | -0,395002011 | 0,128992247 | -3,062215142 | 0,002197055 | 0,045615003 |
| ENSG00000128833.13 | MYO5C | protein_coding | 1082,157943 | -0,395859051 | 0,106209872 | -3,727139903 | 0,000193665 | 0,007459585 |
| ENSG00000245293.3 | CYP2U1-AS1 | lncRNA | 1006,20423 | -0,397387841 | 0,09681011 | -4,1048176 | 4,05E-05 | 0,00214185 |
| ENSG00000261713.6 | SSTR5-AS1 | lncRNA | 2141,804428 | -0,397777664 | 0,108309699 | -3,672595055 | 0,0002401 | 0,008708931 |
| ENSG00000136824.19 | SMC2 | protein_coding | 25794,92062 | -0,397822214 | 0,06232621 | -6,382903942 | 1,74E-10 | 5,57E-08 |
| ENSG00000115568.16 | ZNF142 | protein_coding | 1660,783951 | -0,397859479 | 0,107557696 | -3,699033104 | 0,000216422 | 0,008048002 |
| ENSG00000015133.19 | CCDC88C | protein_coding | 1187,207239 | -0,400947773 | 0,126945134 | -3,158433582 | 0,001586194 | 0,036181587 |
| ENSG00000147724.12 | FAM135B | protein_coding | 2052,789473 | -0,402731021 | 0,091581447 | -4,397517539 | 1,09E-05 | 0,000752394 |
| ENSG00000182759.4 | MAFA | protein_coding | 11923,10905 | -0,405726758 | 0,096942618 | -4,185225911 | 2,85E-05 | 0,001618514 |
| ENSG00000182938.7 | OTOP3 | protein_coding | 1156,518279 | -0,406044022 | 0,088023759 | -4,612891162 | 3,97E-06 | 0,000321845 |
| ENSG00000150672.17 | DLG2 | protein_coding | 2325,753442 | -0,408025518 | 0,085310714 | -4,782816846 | 1,73E-06 | 0,000153599 |
| ENSG00000176438.12 | SYNE3 | protein_coding | 444,8567104 | -0,410203502 | 0,129641315 | -3,164141788 | 0,00155541 | 0,03577443 |
| ENSG00000187764.11 | SEMA4D | protein_coding | 1677,294028 | -0,41068159 | 0,104620057 | -3,925457539 | 8,66E-05 | 0,003949151 |
| ENSG00000133226.17 | SRRM1 | protein_coding | 1170,045 | -0,41073286 | 0,134603443 | -3,051429082 | 0,002277548 | 0,046705811 |
| ENSG00000075035.10 | WSCD2 | protein_coding | 554,4102816 | -0,411055288 | 0,119080775 | -3,45190304 | 0,000556648 | 0,016532064 |
| ENSG00000139613.12 | SMARCC2 | protein_coding | 7991,285487 | -0,41198431 | 0,098793478 | -4,170156949 | 3,04E-05 | 0,001707494 |
| ENSG00000102057.10 | KCND1 | protein_coding | 3688,288893 | -0,414361211 | 0,087103852 | -4,75709401 | 1,96E-06 | 0,000173129 |
| ENSG00000107249.23 | GLIS3 | protein_coding | 3282,035226 | -0,41589224 | 0,108704841 | -3,825885172 | 0,000130303 | 0,005419332 |
| ENSG00000114857.18 | NKTR | protein_coding | 1632,197083 | -0,415976921 | 0,128419581 | -3,239201664 | 0,001198648 | 0,029500872 |
| ENSG00000143442.22 | POGZ | protein_coding | 2681,764946 | -0,418863225 | 0,114383214 | -3,661929134 | 0,000250323 | 0,009035317 |
| ENSG00000169992.10 | NLGN2 | protein_coding | 1622,703654 | -0,419961214 | 0,133628637 | -3,142748611 | 0,001673695 | 0,037866385 |
| ENSG00000141068.14 | KSR1 | protein_coding | 1885,560546 | -0,420988893 | 0,099632842 | -4,225402847 | 2,39E-05 | 0,001422467 |
| ENSG00000169507.9 | SLC38A11 | protein_coding | 426,5181395 | -0,422055551 | 0,137851529 | -3,061667543 | 0,002201078 | 0,045642967 |
| ENSG00000287038.1 | AL162388.2 | lncRNA | 712,4317409 | -0,425933312 | 0,114806636 | -3,710006034 | 0,000207254 | 0,007838818 |
| ENSG00000213020.10 | ZNF611 | protein_coding | 545,0114551 | -0,427233275 | 0,128605252 | -3,32205153 | 0,000893582 | 0,023706706 |
| ENSG00000082805.20 | ERC1 | protein_coding | 7319,250931 | -0,428633393 | 0,075204864 | -5,699543469 | 1,20E-08 | 2,14E-06 |
| ENSG00000137812.20 | KNL1 | protein_coding | 1975,685723 | -0,4294535 | 0,09596905 | -4,474916635 | 7,64E-06 | 0,000565664 |
| ENSG00000214595.12 | EML6 | protein_coding | 1110,091599 | -0,429856569 | 0,133814438 | -3,212333243 | 0,001316616 | 0,031790729 |
| ENSG00000090615.15 | GOLGA3 | protein_coding | 1999,465065 | -0,430109396 | 0,122161219 | -3,52083419 | 0,000430192 | 0,013675888 |
| ENSG00000120594.17 | PLXDC2 | protein_coding | 4068,39279 | -0,430221674 | 0,065128777 | -6,60570788 | 3,96E-11 | 1,55E-08 |
| ENSG00000267640.6 | AC016582.3 | transcribed_unproc | 778,6274331 | -0,43346984 | 0,137973674 | -3,141685138 | 0,001679786 | 0,037957643 |
| ENSG00000160712.13 | IL6R | protein_coding | 4500,99014 | -0,434137241 | 0,075901593 | -5,719738208 | 1,07E-08 | 1,95E-06 |
| ENSG00000143473.13 | KCNH1 | protein_coding | 458,0868049 | -0,434228823 | 0,137378797 | -3,160813991 | 0,001573289 | 0,035988904 |

| | | | | | | | | |
|---|---|---|---|---|---|---|---|---|
| ENSG00000177144.7 | NUDT4B | protein_coding | 1662,69505 | -0,43496938 | 0,101427168 | -4,288489845 | 1,80E-05 | 0,001127561 |
| ENSG00000126001.16 | CEP250 | protein_coding | 1876,393271 | -0,436086114 | 0,117923861 | -3,698031159 | 0,000217278 | 0,008066272 |
| ENSG00000184564.11 | SLITRK6 | protein_coding | 895,360074 | -0,440036648 | 0,101176898 | -4,34918107 | 1,37E-05 | 0,000899836 |
| ENSG00000119946.11 | CNNM1 | protein_coding | 3276,002399 | -0,440322202 | 0,077971929 | -5,647188763 | 1,63E-08 | 2,75E-06 |
| ENSG00000005187.12 | ACSM3 | protein_coding | 552,8956125 | -0,440886918 | 0,115398796 | -3,820550397 | 0,000133154 | 0,005476152 |
| ENSG00000135837.16 | CEP350 | protein_coding | 3925,561375 | -0,442058453 | 0,071881089 | -6,149857443 | 7,76E-10 | 2,07E-07 |
| ENSG00000185418.16 | TARS3 | protein_coding | 929,8835143 | -0,443055078 | 0,139474324 | -3,176606742 | 0,001490089 | 0,034741537 |
| ENSG00000107282.8 | APBA1 | protein_coding | 459,1282771 | -0,446534757 | 0,139558213 | -3,199630809 | 0,001376037 | 0,032808406 |
| ENSG00000198198.17 | SZT2 | protein_coding | 1351,687303 | -0,449584893 | 0,107307738 | -4,18967824 | 2,79E-05 | 0,001593017 |
| ENSG00000092853.14 | CLSPN | protein_coding | 1611,268571 | -0,453060622 | 0,123513995 | -3,66809139 | 0,000244368 | 0,008849236 |
| ENSG00000198157.11 | HMGN5 | protein_coding | 1283,195596 | -0,45414886 | 0,092233254 | -4,923916714 | 8,48E-07 | 8,41E-05 |
| ENSG00000168071.22 | CCDC88B | protein_coding | 538,5417956 | -0,457164702 | 0,140466484 | -3,254617674 | 0,001135451 | 0,02822808 |
| ENSG00000167107.13 | ACSF2 | protein_coding | 1710,091978 | -0,458476216 | 0,09470718 | -4,840986874 | 1,29E-06 | 0,000118123 |
| ENSG00000187634.12 | SAMD11 | protein_coding | 385,2016722 | -0,460432324 | 0,149883522 | -3,071934235 | 0,002126766 | 0,044647493 |
| ENSG00000173482.17 | PTPRM | protein_coding | 725,2950359 | -0,460807721 | 0,109241403 | -4,218251589 | 2,46E-05 | 0,001454985 |
| ENSG00000231304.2 | SGO1-AS1 | lncRNA | 304,2263503 | -0,463705719 | 0,142431269 | -3,255645505 | 0,001131349 | 0,028173743 |
| ENSG00000176208.9 | ATAD5 | protein_coding | 511,7651184 | -0,464602532 | 0,141955126 | -3,272883101 | 0,001064565 | 0,027012115 |
| ENSG00000184792.16 | OSBP2 | protein_coding | 4474,968856 | -0,465557709 | 0,079498779 | -5,856161728 | 4,74E-09 | 9,70E-07 |
| ENSG00000116039.13 | ATP6V1B1 | protein_coding | 1894,141857 | -0,467091509 | 0,088163526 | -5,298013021 | 1,17E-07 | 1,47E-05 |
| ENSG00000140092.14 | FBLN5 | protein_coding | 434,6424923 | -0,46890164 | 0,136609036 | -3,432435014 | 0,000598187 | 0,017484135 |
| ENSG00000083857.14 | FAT1 | protein_coding | 1932,282421 | -0,476085179 | 0,096286281 | -4,944475743 | 7,63E-07 | 7,68E-05 |
| ENSG00000083067.23 | TRPM3 | protein_coding | 4804,717014 | -0,476473613 | 0,071918994 | -6,625142898 | 3,47E-11 | 1,42E-08 |
| ENSG00000197296.6 | FITM2 | protein_coding | 4073,403011 | -0,482207086 | 0,092342825 | -5,221922616 | 1,77E-07 | 2,10E-05 |
| ENSG00000204138.13 | PHACTR4 | protein_coding | 1864,60916 | -0,482501508 | 0,104728511 | -4,607164797 | 4,08E-06 | 0,000329626 |
| ENSG00000184481.17 | FOXO4 | protein_coding | 504,2873811 | -0,482828276 | 0,13069951 | -3,694185811 | 0,000220592 | 0,008148295 |
| ENSG00000257315.3 | ZBED6 | protein_coding | 1269,508862 | -0,487648822 | 0,110491877 | -4,413435934 | 1,02E-05 | 0,000714655 |
| ENSG00000262655.4 | SPON1 | protein_coding | 22946,64686 | -0,48937725 | 0,062284459 | -7,857132578 | 3,93E-15 | 4,58E-12 |
| ENSG00000198734.12 | F5 | protein_coding | 535,0194502 | -0,490745438 | 0,125878099 | -3,898576815 | 9,68E-05 | 0,004281811 |
| ENSG00000166435.15 | XRRA1 | protein_coding | 485,9734565 | -0,491072052 | 0,152169195 | -3,227144966 | 0,001250321 | 0,030602436 |
| ENSG00000118263.15 | KLF7 | protein_coding | 1677,447721 | -0,493112866 | 0,100647366 | -4,899411526 | 9,61E-07 | 9,29E-05 |
| ENSG00000066427.24 | ATXN3 | protein_coding | 742,4316158 | -0,49323032 | 0,116970901 | -4,216692478 | 2,48E-05 | 0,001454985 |
| ENSG00000149256.16 | TENM4 | protein_coding | 1709,850807 | -0,495468919 | 0,117475476 | -4,217637034 | 2,47E-05 | 0,001454985 |
| ENSG00000213918.10 | DNASE1 | protein_coding | 1135,257122 | -0,496934643 | 0,120643457 | -4,119035178 | 3,80E-05 | 0,002043229 |
| ENSG00000109265.14 | CRACD | protein_coding | 867,1858155 | -0,497070451 | 0,150943989 | -3,293078804 | 0,000990967 | 0,025495506 |
| ENSG00000170271.11 | FAXDC2 | protein_coding | 376,1975036 | -0,497770634 | 0,141686603 | -3,513180661 | 0,000442776 | 0,013975561 |
| ENSG00000042781.14 | USH2A | protein_coding | 732,5038183 | -0,498845321 | 0,134962753 | -3,696170305 | 0,000218876 | 0,008111984 |

| | | | | | | | | |
|---|---|---|---|---|---|---|---|---|
| ENSG00000140443.15 | IGF1R | protein_coding | 5261,185925 | -0,501639078 | 0,083762575 | -5,9888211 | 2,11E-09 | 4,59E-07 |
| ENSG00000079102.16 | RUNX1T1 | protein_coding | 1503,035478 | -0,506884405 | 0,103379341 | -4,903149895 | 9,43E-07 | 9,16E-05 |
| ENSG00000166987.15 | MBD6 | protein_coding | 3138,694562 | -0,510554437 | 0,105816205 | -4,824917306 | 1,40E-06 | 0,00012649 |
| ENSG00000129965.15 | INS-IGF2 | protein_coding | 377,6628302 | -0,512229478 | 0,140126255 | -3,655485375 | 0,000256696 | 0,009220215 |
| ENSG00000060237.18 | WNK1 | protein_coding | 12884,0407 | -0,513248949 | 0,093849187 | -5,468869407 | 4,53E-08 | 6,47E-06 |
| ENSG00000132938.21 | MTUS2 | protein_coding | 2805,284159 | -0,514125438 | 0,088469299 | -5,811342988 | 6,20E-09 | 1,21E-06 |
| ENSG00000004777.18 | ARHGAP33 | protein_coding | 1023,264415 | -0,518677016 | 0,135730446 | -3,821375608 | 0,000132709 | 0,005476152 |
| ENSG00000076555.15 | ACACB | protein_coding | 1275,896151 | -0,520867159 | 0,11725347 | -4,442232374 | 8,90E-06 | 0,000641656 |
| ENSG00000130749.10 | ZC3H4 | protein_coding | 1722,706762 | -0,521920886 | 0,122114062 | -4,274044078 | 1,92E-05 | 0,001183088 |
| ENSG00000099331.13 | MYO9B | protein_coding | 1760,091419 | -0,527202858 | 0,128901422 | -4,089969302 | 4,31E-05 | 0,002265923 |
| ENSG00000171956.7 | FOXB1 | protein_coding | 987,3153153 | -0,5272088 | 0,107829768 | -4,889269521 | 1,01E-06 | 9,69E-05 |
| ENSG00000141753.7 | IGFBP4 | protein_coding | 1860,985904 | -0,530895277 | 0,099364036 | -5,342931874 | 9,15E-08 | 1,18E-05 |
| ENSG00000077616.11 | NAALAD2 | protein_coding | 200,3832217 | -0,53176219 | 0,173894912 | -3,0579514 | 0,002228557 | 0,046040201 |
| ENSG00000184502.4 | GAST | protein_coding | 523,7134778 | -0,535002216 | 0,125293253 | -4,27000022 | 1,95E-05 | 0,001198994 |
| ENSG00000184156.17 | KCNQ3 | protein_coding | 274,974736 | -0,539107587 | 0,171829129 | -3,137463303 | 0,001704166 | 0,038264873 |
| ENSG00000174804.4 | FZD4 | protein_coding | 272,4315317 | -0,549638106 | 0,177875035 | -3,090023877 | 0,002001404 | 0,04332981 |
| ENSG00000103056.12 | SMPD3 | protein_coding | 592,5553405 | -0,552524382 | 0,118954617 | -4,644833416 | 3,40E-06 | 0,000282045 |
| ENSG00000187240.16 | DYNC2H1 | protein_coding | 932,9000412 | -0,560892766 | 0,12362674 | -4,536985827 | 5,71E-06 | 0,000441467 |
| ENSG00000141574.8 | SECTM1 | protein_coding | 909,2324644 | -0,565056436 | 0,141893108 | -3,982268363 | 6,83E-05 | 0,003248032 |
| ENSG00000270332.2 | SMC2-AS1 | lncRNA | 347,313521 | -0,567608539 | 0,145376015 | -3,904416679 | 9,45E-05 | 0,004213438 |
| ENSG00000021645.20 | NRXN3 | protein_coding | 502,7746579 | -0,567637103 | 0,119241772 | -4,760388035 | 1,93E-06 | 0,000171008 |
| ENSG00000101638.14 | ST8SIA5 | protein_coding | 330,1943733 | -0,573172946 | 0,186346408 | -3,0758465 | 0,002099059 | 0,044486374 |
| ENSG00000072182.13 | ASIC4 | protein_coding | 394,3276021 | -0,57406147 | 0,131348296 | -4,370528494 | 1,24E-05 | 0,000823553 |
| ENSG00000090905.19 | TNRC6A | protein_coding | 2786,295006 | -0,576516272 | 0,10545803 | -5,466784032 | 4,58E-08 | 6,50E-06 |
| ENSG00000171798.18 | KNDC1 | protein_coding | 862,1676884 | -0,579139793 | 0,121935159 | -4,749571795 | 2,04E-06 | 0,000178982 |
| ENSG00000090975.13 | PITPNM2 | protein_coding | 342,8302492 | -0,580574475 | 0,16729847 | -3,470291592 | 0,000519894 | 0,015801053 |
| ENSG00000129244.9 | ATP1B2 | protein_coding | 298,3138049 | -0,595415322 | 0,181351785 | -3,28320629 | 0,001026335 | 0,026283212 |
| ENSG00000119397.17 | CNTRL | protein_coding | 557,0580394 | -0,609979526 | 0,186796501 | -3,265476185 | 0,001092802 | 0,027583538 |
| ENSG00000170962.13 | PDGFD | protein_coding | 5936,318783 | -0,619774438 | 0,068899959 | -8,995280233 | 2,36E-19 | 7,45E-16 |
| ENSG00000079308.19 | TNS1 | protein_coding | 1061,635105 | -0,621912246 | 0,148157794 | -4,197634356 | 2,70E-05 | 0,001558165 |
| ENSG00000106100.11 | NOD1 | protein_coding | 227,4222386 | -0,639220172 | 0,211260017 | -3,025750826 | 0,002480166 | 0,04970666 |
| ENSG00000164920.9 | OSR2 | protein_coding | 248,7404712 | -0,645545239 | 0,171744764 | -3,758747723 | 0,000170766 | 0,006747083 |
| ENSG00000138778.13 | CENPE | protein_coding | 574,8861024 | -0,649086799 | 0,183640439 | -3,534552652 | 0,000408466 | 0,013117155 |
| ENSG00000145642.12 | SHISAL2B | protein_coding | 899,3500124 | -0,654659901 | 0,109438326 | -5,981998475 | 2,20E-09 | 4,73E-07 |
| ENSG00000188011.5 | RTP5 | protein_coding | 175,3469669 | -0,655421222 | 0,198641336 | -3,299520817 | 0,0009685 | 0,025092554 |
| ENSG00000137872.17 | SEMA6D | protein_coding | 683,125957 | -0,661473283 | 0,118720409 | -5,571689733 | 2,52E-08 | 3,93E-06 |

| | | | | | | | | |
|---|---|---|---|---|---|---|---|---|
| ENSG00000120693.14 | SMAD9 | protein_coding | 874,4601876 | -0,663856952 | 0,15061865 | -4,407534862 | 1,05E-05 | 0,000731762 |
| ENSG00000286112.1 | AL441992.2 | protein_coding | 187,6867313 | -0,668421844 | 0,209727071 | -3,18710332 | 0,001437054 | 0,033897935 |
| ENSG00000145362.20 | ANK2 | protein_coding | 1943,016207 | -0,678130548 | 0,12128639 | -5,591151235 | 2,26E-08 | 3,62E-06 |
| ENSG00000058335.16 | RASGRF1 | protein_coding | 611,3156163 | -0,683661098 | 0,177320843 | -3,855503314 | 0,000115492 | 0,004971539 |
| ENSG00000198785.7 | GRIN3A | protein_coding | 588,2243203 | -0,700793425 | 0,128565777 | -5,450855125 | 5,01E-08 | 7,02E-06 |
| ENSG00000122966.17 | CIT | protein_coding | 1863,455104 | -0,71024482 | 0,117862193 | -6,02606145 | 1,68E-09 | 3,91E-07 |
| ENSG00000288534.1 | AP001931.2 | protein_coding | 520,8201321 | -0,715777025 | 0,186378552 | -3,840447397 | 0,00012281 | 0,005215548 |
| ENSG00000140807.7 | NKD1 | protein_coding | 371,663883 | -0,718473004 | 0,207799749 | -3,457525849 | 0,00054516 | 0,01626561 |
| ENSG00000259295.6 | CSPG4P12 | transcribed_unproc | 157,0386147 | -0,751242959 | 0,245150183 | -3,064419331 | 0,00218093 | 0,045352682 |
| ENSG00000259881.1 | AC092384.2 | lncRNA | 284,16299 | -0,780649096 | 0,184662802 | -4,227430144 | 2,36E-05 | 0,001413526 |
| ENSG00000196562.14 | SULF2 | protein_coding | 1826,738527 | -0,789065057 | 0,127196227 | -6,203525644 | 5,52E-10 | 1,55E-07 |
| ENSG00000185829.18 | ARL17A | protein_coding | 375,3206023 | -0,819702497 | 0,169576848 | -4,833811381 | 1,34E-06 | 0,00012196 |
| ENSG00000258311.5 | AC009779.3 | protein_coding | 119,8193896 | -0,869959244 | 0,261410824 | -3,327938879 | 0,000874911 | 0,023388779 |
| ENSG00000183091.19 | NEB | protein_coding | 154,2393869 | -0,907069794 | 0,288407131 | -3,145101829 | 0,00166029 | 0,037677521 |
| ENSG00000101204.18 | CHRNA4 | protein_coding | 102,2648379 | -0,993168354 | 0,284154849 | -3,495165958 | 0,000473767 | 0,014660937 |
| ENSG00000196912.12 | ANKRD36B | protein_coding | 96,68925877 | -0,997094389 | 0,289437555 | -3,44493785 | 0,000571191 | 0,016806071 |
| ENSG00000249249.2 | AC010226.1 | lncRNA | 152,0352497 | -1,015789999 | 0,309827272 | -3,278568712 | 0,00104335 | 0,026569473 |
| ENSG00000230445.4 | LRRC37A6P | transcribed_process | 76,27738771 | -1,038351001 | 0,297160804 | -3,494239439 | 0,000475414 | 0,014691368 |
| ENSG00000163406.11 | SLC15A2 | protein_coding | 95,4925305 | -1,170343159 | 0,314875061 | -3,716849334 | 0,000201723 | 0,007695372 |
| ENSG00000182050.13 | MGAT4C | protein_coding | 46,45627058 | -1,45818059 | 0,470316187 | -3,100426118 | 0,001932424 | 0,042166486 |
| ENSG00000080947.15 | CROCCP3 | transcribed_unproc | 46,97554681 | -1,616867047 | 0,466254957 | -3,467774495 | 0,000524787 | 0,015862633 |
| ENSG00000151006.7 | PRSS53 | protein_coding | 51,59170914 | -1,763616585 | 0,448107281 | -3,93570169 | 8,30E-05 | 0,003815882 |
| ENSG00000170160.17 | CCDC144A | protein_coding | 57,46646575 | -1,772421041 | 0,398301983 | -4,449942799 | 8,59E-06 | 0,000625155 |
| ENSG00000281404.2 | LINC01176 | lncRNA | 26,16085501 | -1,816950967 | 0,523016964 | -3,473980943 | 0,000512797 | 0,015649871 |
| ENSG00000233327.11 | USP32P2 | transcribed_unproc | 47,73710099 | -2,006579187 | 0,511482338 | -3,923066427 | 8,74E-05 | 0,003972401 |
| ENSG00000150637.9 | CD226 | protein_coding | 49,3092695 | -2,083930077 | 0,415925455 | -5,010345133 | 5,43E-07 | 5,70E-05 |
| ENSG00000172789.3 | HOXC5 | protein_coding | 14,38631268 | -2,657650373 | 0,856338213 | -3,103505523 | 0,001912427 | 0,0418173 |
| ENSG00000183117.19 | CSMD1 | protein_coding | 22,25989825 | -2,748828839 | 0,623749571 | -4,406943053 | 1,05E-05 | 0,000731762 |
| ENSG00000274628.5 | AL669942.1 | transcribed_unproc | 14,65260072 | -3,101487166 | 1,005557879 | -3,084344751 | 0,002040011 | 0,043610891 |
| ENSG00000215032.2 | GNL3LP1 | processed_pseudog | 15,40623976 | -3,353818101 | 0,999528049 | -3,355401685 | 0,000792498 | 0,02164792 |
| ENSG00000182557.8 | SPNS3 | protein_coding | 8,738744829 | -3,677569132 | 1,176600919 | -3,125587506 | 0,001774503 | 0,039499644 |
| ENSG00000257527.1 | AC126755.1 | lncRNA | 8,095706165 | -4,219289765 | 1,266518281 | -3,331408498 | 0,000864077 | 0,023145966 |
| ENSG00000130592.17 | LSP1 | protein_coding | 4,878476873 | -4,306435774 | 1,328196771 | -3,242317605 | 0,001185618 | 0,02921268 |
| ENSG00000285839.1 | AL445685.3 | protein_coding | 10,96862909 | -4,37426394 | 1,105053284 | -3,958419022 | 7,54E-05 | 0,003521839 |
| ENSG00000188536.13 | HBA2 | protein_coding | 69,52481733 | -4,604525795 | 1,39048774 | -3,311446525 | 0,00092815 | 0,02433204 |
| ENSG00000284986.1 | AL449043.2 | transcribed_unproc | 6,963249077 | -4,817273225 | 1,339764864 | -3,595610957 | 0,000323631 | 0,010865953 |

| | | | | | | | | |
|---|---|---|---|---|---|---|---|---|
| ENSG00000213285.4 | RPL23AP45 | processed_pseudog | 23,22601642 | -9,860625523 | 3,022793741 | -3,262090096 | 0,00110594 | 0,027737202 |

**DEG_significative_TNFα vs NT 24h**

| **Ensembl_gene_ID** | **gene_name** | **gene_type** | **baseMean** | **log2FoldChange** | **lfcSE** | **stat** | **pvalue** | **padj** |
|---|---|---|---|---|---|---|---|---|
| ENSG00000280800.1 | FP671120.6 | lncRNA | 303,5602014 | 2,585390216 | 0,609109132 | 4,244543514 | 2,19E-05 | 0,013746253 |
| ENSG00000161939.19 | RNASEK-C17orf4 | protein_coding | 54,73144215 | 1,97354201 | 0,464420959 | 4,249468016 | 2,14E-05 | 0,013746253 |
| ENSG00000225840.2 | AC010970.1 | processed_pseudog | 42,68374123 | 1,973108721 | 0,430271864 | 4,585725648 | 4,52E-06 | 0,004320538 |
| ENSG00000085514.16 | PILRA | protein_coding | 45,16952164 | 1,873366244 | 0,439788889 | 4,259694344 | 2,05E-05 | 0,013625398 |
| ENSG00000274290.3 | H2BC6 | protein_coding | 67,58744938 | 1,808098164 | 0,406841386 | 4,444233616 | 8,82E-06 | 0,006680823 |
| ENSG00000163121.10 | NEURL3 | protein_coding | 111,1375409 | 1,65862144 | 0,299048711 | 5,546325324 | 2,92E-08 | 7,12E-05 |
| ENSG00000105499.14 | PLA2G4C | protein_coding | 263,5560652 | 1,450336998 | 0,182709694 | 7,937931295 | 2,06E-15 | 4,52E-11 |
| ENSG00000232573.1 | RPL3P4 | processed_pseudog | 54,87819983 | 1,433772071 | 0,358839347 | 3,995582102 | 6,45E-05 | 0,025773127 |
| ENSG00000167434.10 | CA4 | protein_coding | 648,6799733 | 1,311961507 | 0,210888389 | 6,221117789 | 4,94E-10 | 3,52E-06 |
| ENSG00000283293.1 | RN7SK | snRNA | 285,8904336 | 1,074950604 | 0,224550186 | 4,787128532 | 1,69E-06 | 0,002111173 |
| ENSG00000177301.16 | KCNA2 | protein_coding | 512,9016561 | 1,064996914 | 0,172323814 | 6,180207436 | 6,40E-10 | 3,52E-06 |
| ENSG00000143882.12 | ATP6V1C2 | protein_coding | 152,4845975 | 1,007998289 | 0,258508608 | 3,899283259 | 9,65E-05 | 0,035852191 |
| ENSG00000023445.16 | BIRC3 | protein_coding | 350,7729709 | 1,00610077 | 0,210070871 | 4,789339731 | 1,67E-06 | 0,002111173 |
| ENSG00000227507.3 | LTB | protein_coding | 237,4288641 | 0,913223824 | 0,217257821 | 4,203410577 | 2,63E-05 | 0,015501326 |
| ENSG00000235568.7 | NFAM1 | protein_coding | 172,3050734 | 0,909277204 | 0,209627694 | 4,337581483 | 1,44E-05 | 0,009888309 |
| ENSG00000258017.2 | AC011603.2 | lncRNA | 535,2767821 | 0,869762134 | 0,207959201 | 4,182369084 | 2,88E-05 | 0,015710198 |
| ENSG00000180767.10 | CHST13 | protein_coding | 234,4183537 | 0,851135435 | 0,171400441 | 4,965771571 | 6,84E-07 | 0,00120771 |
| ENSG00000104856.14 | RELB | protein_coding | 630,8839434 | 0,830308622 | 0,14198841 | 5,847721109 | 4,98E-09 | 1,82E-05 |
| ENSG00000227617.9 | CERS6-AS1 | lncRNA | 314,6362925 | 0,82777103 | 0,206677514 | 4,005133472 | 6,20E-05 | 0,025211945 |
| ENSG00000078295.17 | ADCY2 | protein_coding | 692,5188775 | 0,813824412 | 0,196118556 | 4,149655327 | 3,33E-05 | 0,016213015 |
| ENSG00000138442.10 | WDR12 | protein_coding | 1757,093801 | 0,769906176 | 0,144650762 | 5,322517242 | 1,02E-07 | 0,000204357 |
| ENSG00000211459.2 | MT-RNR1 | Mt_rRNA | 30205,55142 | 0,744134424 | 0,130411051 | 5,706068752 | 1,16E-08 | 3,63E-05 |
| ENSG00000109686.19 | SH3D19 | protein_coding | 394,6766628 | 0,66813214 | 0,162322285 | 4,116083874 | 3,85E-05 | 0,017634423 |
| ENSG00000154065.17 | ANKRD29 | protein_coding | 326,8344739 | 0,65965166 | 0,172496982 | 3,824134509 | 0,000131232 | 0,043022636 |
| ENSG00000164008.16 | C1orf50 | protein_coding | 903,5533438 | 0,584427264 | 0,125157898 | 4,669519657 | 3,02E-06 | 0,003157781 |
| ENSG00000118503.15 | TNFAIP3 | protein_coding | 1254,76031 | 0,567137833 | 0,136213521 | 4,163594252 | 3,13E-05 | 0,016002607 |
| ENSG00000100906.11 | NFKBIA | protein_coding | 3460,550982 | 0,558299759 | 0,091291446 | 6,115575811 | 9,62E-10 | 4,23E-06 |
| ENSG00000260342.2 | AC138811.2 | protein_coding | 508,4541193 | 0,558262451 | 0,146303854 | 3,815774059 | 0,000135757 | 0,043851406 |
| ENSG00000168394.11 | TAP1 | protein_coding | 720,8991648 | 0,556578947 | 0,112822625 | 4,933221044 | 8,09E-07 | 0,001269021 |
| ENSG00000019582.15 | CD74 | protein_coding | 9083,73953 | 0,539618268 | 0,0952468 | 5,665473968 | 1,47E-08 | 4,03E-05 |
| ENSG00000234745.12 | HLA-B | protein_coding | 566,7859249 | 0,527532155 | 0,129884797 | 4,061538881 | 4,88E-05 | 0,021416005 |
| ENSG00000134769.21 | DTNA | protein_coding | 1497,034161 | 0,515070737 | 0,113594638 | 4,534287417 | 5,78E-06 | 0,004882843 |

| | | | | | | | | |
|---|---|---|---|---|---|---|---|---|
| ENSG00000178078.12 | STAP2 | protein_coding | 3011,773749 | 0,480040176 | 0,101276819 | 4,739882028 | 2,14E-06 | 0,002348527 |
| ENSG00000131174.7 | COX7B | protein_coding | 4655,959792 | 0,465877978 | 0,103903062 | 4,483775236 | 7,33E-06 | 0,005965849 |
| ENSG00000203875.13 | SNHG5 | lncRNA | 1672,444603 | 0,459027442 | 0,100926855 | 4,54811994 | 5,41E-06 | 0,004755623 |
| ENSG00000164587.13 | RPS14 | protein_coding | 47222,99267 | 0,452330784 | 0,094987256 | 4,76201548 | 1,92E-06 | 0,002215794 |
| ENSG00000136842.14 | TMOD1 | protein_coding | 6680,289428 | 0,430872897 | 0,088392508 | 4,874540916 | 1,09E-06 | 0,001597028 |
| ENSG00000255639.3 | AC005833.1 | protein_coding | 1767,213271 | 0,421634926 | 0,102347544 | 4,119638924 | 3,79E-05 | 0,017634423 |
| ENSG00000134569.10 | LRP4 | protein_coding | 7078,985402 | 0,413891873 | 0,088831997 | 4,659265669 | 3,17E-06 | 0,003168346 |
| ENSG00000213741.11 | RPS29 | protein_coding | 11072,82901 | 0,403358204 | 0,105073383 | 3,838823792 | 0,000123625 | 0,041775777 |
| ENSG00000077150.20 | NFKB2 | protein_coding | 1307,928983 | 0,396758337 | 0,101439951 | 3,911263089 | 9,18E-05 | 0,034770857 |
| ENSG00000146232.17 | NFKBIE | protein_coding | 1683,197068 | 0,380705763 | 0,093944469 | 4,052455341 | 5,07E-05 | 0,02182843 |
| ENSG00000143947.14 | RPS27A | protein_coding | 33887,00553 | 0,360684758 | 0,092895126 | 3,882709175 | 0,000103299 | 0,037196139 |
| ENSG00000169436.17 | COL22A1 | protein_coding | 6838,372383 | 0,358816217 | 0,08153824 | 4,400588226 | 1,08E-05 | 0,007904312 |
| ENSG00000104044.16 | OCA2 | protein_coding | 2760,130014 | 0,350083885 | 0,085836631 | 4,078490492 | 4,53E-05 | 0,020319443 |
| ENSG00000118363.12 | SPCS2 | protein_coding | 19305,22715 | 0,345997129 | 0,086327979 | 4,007937313 | 6,13E-05 | 0,025211945 |
| ENSG00000170296.10 | GABARAP | protein_coding | 25315,47072 | 0,343904589 | 0,082964836 | 4,145184927 | 3,40E-05 | 0,016213015 |
| ENSG00000281571.2 | AC241585.2 | TEC | 3125,370705 | 0,3426415 | 0,089124077 | 3,844544732 | 0,000120776 | 0,041775777 |
| ENSG00000164713.10 | BRI3 | protein_coding | 5522,068689 | 0,328803474 | 0,078686574 | 4,178647724 | 2,93E-05 | 0,015710198 |
| ENSG00000072501.19 | SMC1A | protein_coding | 32685,23438 | -0,312982869 | 0,081208532 | -3,854063849 | 0,000116173 | 0,041157194 |
| ENSG00000012983.11 | MAP4K5 | protein_coding | 3610,59243 | -0,329659514 | 0,078510324 | -4,198932006 | 2,68E-05 | 0,015501326 |
| ENSG00000157600.12 | TMEM164 | protein_coding | 8252,221717 | -0,380943038 | 0,076844814 | -4,957303173 | 7,15E-07 | 0,00120771 |
| ENSG00000204569.10 | PPP1R10 | protein_coding | 6279,803233 | -0,384929639 | 0,098161808 | -3,921378866 | 8,80E-05 | 0,033927719 |
| ENSG00000205726.15 | ITSN1 | protein_coding | 2038,52374 | -0,406797002 | 0,104423315 | -3,895653023 | 9,79E-05 | 0,035852191 |
| ENSG00000134982.17 | APC | protein_coding | 5964,29279 | -0,421715436 | 0,094750042 | -4,450820566 | 8,55E-06 | 0,006680823 |
| ENSG00000196562.14 | SULF2 | protein_coding | 1914,242651 | -0,464112905 | 0,106779456 | -4,346462541 | 1,38E-05 | 0,009802814 |
| ENSG00000067208.14 | EVI5 | protein_coding | 923,942544 | -0,468956478 | 0,122372249 | -3,83221264 | 0,000126996 | 0,042264619 |
| ENSG00000187240.16 | DYNC2H1 | protein_coding | 882,5936091 | -0,527502018 | 0,110295148 | -4,782640285 | 1,73E-06 | 0,002111173 |
| ENSG00000171316.12 | CHD7 | protein_coding | 8848,558782 | -0,568133238 | 0,134259344 | -4,231610416 | 2,32E-05 | 0,014156689 |
| ENSG00000152818.18 | UTRN | protein_coding | 3301,100354 | -0,608887904 | 0,14678544 | -4,148149203 | 3,35E-05 | 0,016213015 |
| ENSG00000152580.8 | IGSF10 | protein_coding | 470,7467442 | -0,636492139 | 0,152173164 | -4,182683223 | 2,88E-05 | 0,015710198 |
| ENSG00000182986.14 | ZNF320 | protein_coding | 187,3094663 | -0,805430765 | 0,209781678 | -3,839376123 | 0,000123347 | 0,041775777 |
| ENSG00000183889.12 | AC138969.1 | protein_coding | 820,451635 | -0,917814436 | 0,169802828 | -5,405177563 | 6,47E-08 | 0,000142211 |
| ENSG00000285542.1 | AC013717.1 | protein_coding | 122,8174922 | -0,981370258 | 0,246622687 | -3,979237562 | 6,91E-05 | 0,027117609 |
| ENSG00000258311.5 | AC009779.3 | protein_coding | 114,6461059 | -1,031460879 | 0,247521384 | -4,167158656 | 3,08E-05 | 0,016002607 |
| ENSG00000156218.13 | ADAMTSL3 | protein_coding | 47,77821788 | -1,315811519 | 0,346198196 | -3,800746323 | 0,000144261 | 0,045923063 |
| ENSG00000183111.12 | ARHGEF37 | protein_coding | 55,74181789 | -1,650332662 | 0,361404511 | -4,566441793 | 4,96E-06 | 0,004540099 |
| ENSG00000274049.4 | INO80B-WBP1 | protein_coding | 17,30460655 | -3,513909608 | 0,868634871 | -4,045324136 | 5,23E-05 | 0,02207093 |

| | | | | | | | | |
|---|---|---|---|---|---|---|---|---|
| ENSG00000183117.19 | CSMD1 | protein_coding | 21,95476739 | -3,889377388 | 0,625038101 | -6,22262448 | 4,89E-10 | 3,52E-06 |
